# PARKINSON’S DISEASE-ASSOCIATED *PINK1* LOSS DISRUPTS ENSHEATHING GLIA AND CAUSES DOPAMINERGIC NEURON SYNAPSE LOSS

**DOI:** 10.1101/2024.12.06.627235

**Authors:** Lorenzo Ghezzi, Sabine Kuenen, Ulrike Pech, Nils Schoovaerts, Ayse Kilic, Suresh Poovathingal, Kristofer Davie, Jochen Lamote, Roman Praschberger, Patrik Verstreken

## Abstract

Parkinson’s disease (PD) is commonly associated with the loss of dopaminergic neurons in the *substantia nigra*, but many other cell types are affected even before neuron loss occurs. Recent studies have linked oligodendrocytes to early stages of PD, though their precise role is still unclear. *PINK1* is mutated in familial PD, and through unbiased single-cell sequencing of the entire brain of *Drosophila Pink1* models, we observed significant gene deregulation in ensheathing glia (EG); cells that share functional similarities with oligodendrocytes. We found that the loss of *Pink1* leads to abnormalities in EG, similar to the reactive response of EG seen upon nerve injury. Using cell-type-specific transcriptomics, we identified deregulated genes in EG as potential functional modifiers. Specifically downregulating two trafficking factors in EG, *Vps35* and *Vps13*, also mutated in PD, was sufficient to rescue neuronal function and protect against dopaminergic synapse loss. Our findings demonstrate that *Pink1* loss in neurons triggers an injury-like response in EG, and that *Pink1* loss in EG in turn disrupts neuronal function. Vesicle trafficking components, which may regulate membrane interactions between organelles in EG, seem to play a role in maintaining neuronal health and ultimately preventing dopaminergic synapse loss. Our work highlights the essential role of glial support cells in the pathogenesis of PD and identifies vesicle trafficking within these cells in disease progression.

## INTRODUCTION

Parkinson’s disease (PD) is characterized by motor symptoms such as bradykinesia, rigidity and tremor that are caused by the progressive loss of dopaminergic neurons in the substantia nigra (Lang & Lozano, 1998). However, most of the patients report non-motor symptoms, such as constipation, hyposmia and sleep defects, even before the onset of motor symptoms (Munhoz et al., 2015). This suggests that PD is a progressive neurodegenerative disease that initiates many years before the diagnosis and involves different neuronal systems, multiple anatomical areas and different cell types. While therapies symptomatically improve motor functioning temporarily by restoring dopaminergic tone, they do not effectively prevent neurodegeneration and disease progression. This highlights the necessity to identify cell-types and biological mechanisms that act at the earliest stages of disease.

With the advent of single-cell sequencing applied to PD and control brains in combination with evidence from genome-wide association studies (GWAS), it is now possible to identify which cells and mechanisms are at play in PD models that recapitulate early stages of PD (Kaempf et al., 2026; Pech et al., 2025). Among these, one study reported that genes related to GWAS loci are enriched for oligodendrocyte-specific gene expression (Bryois et al., 2020). Interestingly, differential gene expression in post-mortem brains with Braak scores of 1-2, 3-4, and 5-6 and controls, suggested a role for oligodendrocytes at an early stage of PD prior to the overt loss of dopaminergic neurons (Bryois et al., 2020). Additionally, two independent single-cell sequencing studies, one of the human substantia nigra and one of the midbrain, both associated PD genetic risk with oligodendrocyte-specific expression patterns (Agarwal et al., 2020; Smajic et al., 2022). While these studies started to link dysfunction of oligodendrocytes to the early stages of PD, they did not address their role in the pathophysiology of the disease.

Here, we describe a *Drosophila* model to start assessing how neuron-glia cross-talk contributes to PD. *Drosophila* has been successfully used to investigate cellular and molecular dysfunction preceding the onset of age-related symptoms, including dopaminergic neuron-dependent motor symptoms (Kaempf et al., 2026; Pech et al., 2025; Valadas et al., 2018). Furthermore, *Drosophila* glial cells display several anatomical and functional features that show remarkable similarity to their mammalian counterparts (Freeman & Doherty, 2006; Kremer et al., 2017). While the small size of the *Drosophila* brain does not necessitate elaborate myelination, ensheathing glia (EG) in this species share not only anatomical features with oligodendrocytes, such as the ability to wrap around nerve tracts in the Central Nervous System (CNS) (Kremer et al., 2017; Yildirim et al., 2019), but also important molecular and functional features, such as providing metabolic support and regulating neuronal activity (Delgado et al., 2018; Otto et al., 2018).

In a brain-wide single-cell sequencing experiment, we identified EG as the most deregulated cell type in a young (pre-motor) *Pink1* loss-of-function *Drosophila* model. Using cell-specific labeling, immunohistochemistry, and electrophysiological recordings, we correlated the transcriptional deregulation in EG with defects that appear similar to those seen upon nerve injury. Additionally, we find that healthy EG are fundamental to support dopaminergic synapse integrity in *Pink1* mutants. Finally, using cell-type-specific transcriptomic, high-throughput screening and immunohistochemistry, we identified vesicle trafficking as a genetic modifier and showed that *Vps13* and *Vps35* are EG-expressed regulators that maintain dopaminergic synapse integrity. Our study shows that neuron-ensheathing glia crosstalk is already disrupted at a young age by the loss of *Pink1* in neurons, but also in EG, and that EG are fundamental to support dopaminergic synapse-integrity, suggesting a role for oligodendrocytes in the progression of PD.

## RESULTS

### Ensheathing glia in Pink1 loss-of-function flies show cell non-autonomous defects

We previously created a whole-brain single-cell RNAseq dataset from different *Drosophila* PD knock-in models. This dataset was generated from young 5-day-old flies to capture ‘early’ changes (Pech et al., 2025). We re-analyzed the data from *Pink1*^*P399L*^ knock-in loss-of-function mutant flies and isogenic controls and performed differential gene expression (DEG) analysis for each cell type using DESeq2 (Figure 1A). Glial cell subtypes show the highest degree of deregulation; particularly, ensheathing glia are strongly affected (Figure 1A). While flies do not myelinate neurons, ensheathing glia (EG) serve similar supporting functions as oligodendrocytes (Otto et al., 2018; Yildirim et al., 2019). These results suggest that ensheathing glia are deregulated in young *Pink1*^*P399L*^ mutants (and also *Pink1*^*KO-WS*^, see below) at an age prior to dopaminergic neuron-dependent motor defects (Kaempf et al., 2026; Pech et al., 2025).

**Figure 1.**
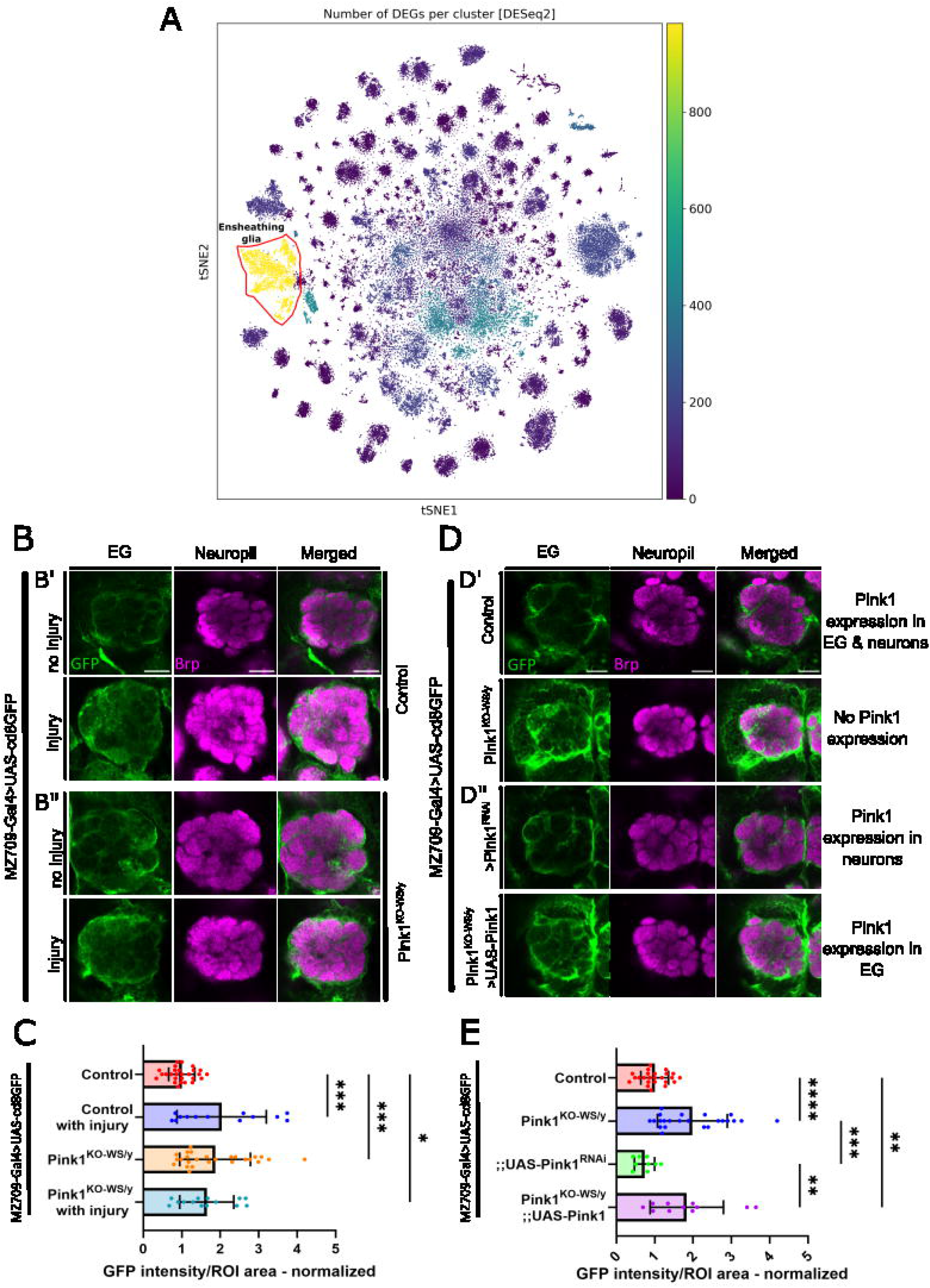
EG are affected non-cell autonomously by *Pink1* loss-of-function (with supplementary fig 1) **(A)** tSNE of the cells of *Pink1*^*P399L*^ knock-in mutants (5-day-old). Cell types are labeled with colors indicating the number of deregulated genes compared to control. EG are encircled and labeled. (**B-B”**) Maximum intensity projections of confocal images of fly brains (5 ± 1-day-old) stained with anti-GFP (Green) and anti-Brp (Magenta), where anti-GFP marks EG and anti-Brp marks presynaptic sites of the antennal lobes in flies where CD8GFP is expressed via the EG driver MZ709-Gal4. Scale bar: 20 µm (**B’**) Maximum intensity projection of confocal images of controls vs. controls 24 hours after ORN-severing (injury). (**B”**) Maximum intensity projection of confocal images of *Pink1*^*KO-WS*^ vs *Pink1*^*KO-WS*^ 24 hours after ORNs severing (injury). (**C**) Quantification of GFP intensity within the glomeruli of the antennal lobe area in 5 ± 1-day-old flies (as in B) relative to controls. ANOVA with Dunnet’s multiple comparison test, * is p<0.05, ** is p<0.01, *** is p<0.001. Effect size: η =0.23. Bars: mean ± SD; points are individual animals N≥13 per genotype, 4 replicates. (**D-D”**) Maximum intensity projection of confocal images of fly brains (5 ± 1-day-old) stained with anti-GFP (Green) and anti-Brp (Magenta), where anti-GFP marks EG and anti-Brp marks presynaptic sites of the antennal lobes in flies where CD8GFP is expressed via the EG driver MZ709-Gal4. Scale bar: 20 µm. (**D’**) Maximum intensity projection of confocal images of control (*w*^*1118*^) and *Pink1*^*KO-WS*^ animals (**D”**) Maximum intensity projection of confocal images of animals with *Pink1* downregulation in EG and *Pink1*^*KO-WS*^ with *Pink1* rescued in EG. (**E**) Quantification of GFP intensity within the glomeruli of the antennal lobe area in 5-day-old flies (as in D) relative to the control. ANOVA with Tukey’s multiple comparison test, * is p<0.05, ** is p<0.01, *** is p<0.001. Effect size: η =0.36 Bars: mean ± SD; points are individual animals N≥10 per genotype, 4 replicates.

When neurons are damaged, activated EG invade the neuropil (Doherty et al., 2009; MacDonald et al., 2006). For example, when fly antennal lobes are removed, severing the Olfactory Receptor Neurons (ORNs), EG invade the antennal lobe neuropil (MacDonald et al., 2006), potentially as a protective response (Figure 1B’-C). To test if EG invaded the neuropil in *Pink1* mutant flies, we expressed a transmembrane fluorescent protein (*UAS-CD8GFP*) under the control of a promoter specific to EG (*MZ709-Gal4*) (Doherty et al., 2009, Supplementary figure 1) in control and *Pink1*^*KO*^ flies. Samples were labeled with anti-GFP and the pre-synaptic protein Bruchpilot (NC82). Similar to ORN-severed controls, GFP signal, lining the membrane of EG, is visible inside the antennal neuropil in *Pink1*^*KO*^, and severing the ORN in *Pink1*^*KO*^ does not further exacerbate this phenotype (Figure 1B”-C). Even though we cannot exclude that in *Pink1* mutants the GFP signal is merely upregulated, *Pink1* loss and neuron-severing that triggers EG activation seem to have a similar phenotype.

In order to determine whether this phenotype is cell-autonomous, we generated cell-type-specific *Pink1* perturbations. We either downregulated *Pink1* using a previously well-validated RNAi line specifically in EG (EG-specific *Pink1* loss-of-function – Figure 1D’), or we re-expressed wild-type *Pink1* specifically in EG in *Pink1*^*KO-WS*^ flies (all cells, but the EG, loss-of-function - Figure 1D”). When *Pink1* is knocked-down in EG (but present in other cells, including neurons), no GFP signal is detected in the neuropil (Figure 1D’’-1E). However, when *Pink1* is not present in neurons and expressed in EG, the GFP is present inside the antennal neuropil (Figure 1D”-E). This suggests that Pink1-defective neurons activate the EG in a cell-nonautonomous fashion to invade, and potentially protect, the neuropil.

### Ensheathing glia function supports synaptic integrity

Our finding that *Pink1*^*KO-WS*^ in neurons elicits a cell non-autonomous response in EG suggested that EG might, in turn, also functionally modulate neuronal integrity in mutant flies. To test this, we made use of a readily accessible model circuit in the fly visual system with an easy electrophysiological read-out, electroretinograms (ERG). Notably, whereas our functional readout here interrogates neuronal activity in the visual system, our analysis of EG morphology was performed in the antennal lobe, a distinct brain region. We therefore make the explicit assumption that EG perform comparable functions across fly brain regions. This assumption is supported by our single-cell sequencing dataset, in which we did not detect region-specific segregation of EG into distinct clusters that could be attributed to different brain areas. The ERG waveforms are susceptible to changes in intracellular signaling, neuronal function, and synaptic transmission, and have been well characterized and amply used to assess neuronal and cellular function (Hardie & Raghu, 2001; Praschberger et al., 2023; Soukup et al., 2016; C. F. Wu & Wong, 1977). We exploited this method to understand the role of *Pink1* in EG on neuronal function and synaptic transmission in young flies.

Comparing control with young *Pink1*^*KO-WS*^ flies does not show a difference in the “depolarization-response”, indicating that -as expected-there is no neurodegeneration occurring in the photoreceptors at this stage (Figure 2A). However, when comparing the ON peak – which represents synaptic transmission from the photoreceptors and within the underlying neuronal circuit, and is a more robust readout of this than the OFF peak (Vilinsky & Johnson, 2012) – we detect a significant reduction in mutant compared to control animals, suggesting that in *Pink1*^*KO-WS*^ flies, synaptic communication in this circuit is partially impaired, already at 5-day-old (Figure 2A-B).

**Figure 2.**
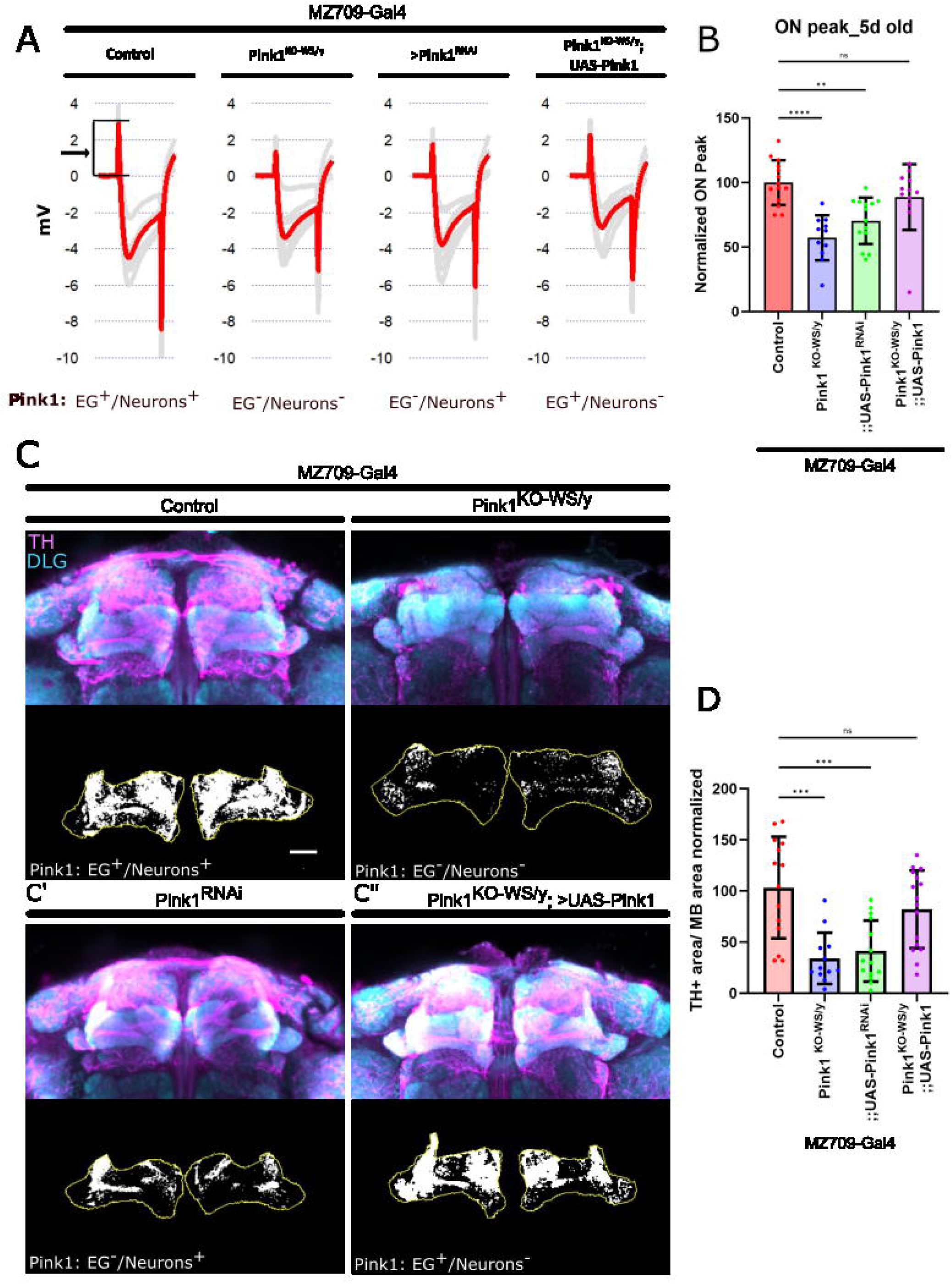
*Pink1* in EG is necessary to support synaptic integrity (with Supplementary figure 2) (**A**) Representative ERG traces of indicated genotypes: ON peak is highlighted by the arrow. (**B**) Normalized ON peak amplitude of flies (5 ± 1-day-old). ANOVA with Tukey’s multiple comparison test, * is p<0.05, ** is p<0.01, *** is p<0.001. Effect size: η =0.55 Bars: mean ± SD; points are individual animals N≥11 per genotype, 3 replicates. (**C-C”**) (**C**) Maximum intensity projection of confocal images of control and *Pink1*^*KO-WS*^ in Mushroom Bodies (MB) of aged flies (22 ± 2-day-old), stained with anti-TH (cyan) and anti-DLG (magenta) antibodies - DLG is used to mark post-synaptic sites of MB. The black and white image is the middle Z-plane within the region of interest of the MB (ROI, yellow), which is used to represent the thresholded TH area (white). Scale bar: 20 µm. (**C’**) Maximum intensity projection of confocal images of *w*^*1118*^ with *Pink1* downregulation in EG. (**C”**) Maximum intensity projection of confocal images of *Pink1*^*KO-WS*^ with *Pink1* rescued in EG. (**D**) Quantification of the dopaminergic synaptic area at MB neuropil in aged flies (22 ± 2-day-old). ANOVA with Tukey’s multiple comparison test, * is p<0.05, ** is p<0.01, *** is p<0.001. Effect size: η = 0.38. Bars: mean ± SD; points are individual animals N≥12 per genotype, 3 replicates.

In order to discern the role of EG in this defective ERG response, we (1) downregulated *Pink1* specifically in EG and (2) we expressed *Pink1* specifically in the EG of *Pink1*^*KO-WS*^. When *Pink1* is knocked down specifically in EG, we observe the same reduction in ON peak response as in the *Pink1*^*KO-WS*^ flies. This indicates that EG integrity is necessary for efficient neurotransmission in this circuit. Conversely, the ON peak defect of *Pink1*^*KO-WS*^ mutants is rescued when *Pink1* is reintroduced in EG only. This suggests that activated EG are able to, at least partially, buffer the detrimental effects of *Pink1* deficiency in neurons (Figure 2A-B).

To also test EG function in relation to PD-relevant dopaminergic (DA) synapses, we evaluated the loss of dopaminergic neuron afferents. In a previous paper from our laboratory, we have shown that the loss of *Pink1* function causes a progressive loss of protocerebral anterior medial dopaminergic neuron (PAM DAN) afferents in the Mushroom Body (MB) neuropil in 25-day-old animals, but not when they are 5-day-old (Kaempf et al., 2026). Indeed, we confirm that 25-day-old animals show a decrease in the dopaminergic synaptic area in MB lobes (Figure 2 C-D). Hence, while ensheathing glia are already activated at a young age in *Pink1*^*KO-WS*^, DA synapses are structurally still intact (at 5 days of age) and only deteriorate in older flies. We then knocked down *Pink1* specifically in EG and found a similar strong decrease in DA synaptic area (Figure 2C’-D). Conversely, when *Pink1* is reintroduced specifically in EG of *Pink1*^*KO-WS*^ animals, DA synapse-loss is significantly rescued (Figure 2C”-D and Supplementary figure 2). Together, these results indicate that *Pink1* in EG is necessary to protect dopaminergic neurons from synapse loss.

### EG cell type-specific transcriptomics reveals modifiers of neuronal dysfunction

To identify pathways in EG that are deregulated by *Pink1* loss-of-function, we optimized a method to isolate EGs from the rest of the brain, enabling us to perform much higher-sensitivity transcriptomics than what we achieved with our 10x droplet-based single-cell sequencing (Pech et al., 2025). We expressed a fluorescently-tagged histone (*UAS-HisTag-eGFP*) in EG (GMR-56-Gal4) and used Fluorescence-activated Cell Sorting (FACS) to isolate EG. We sorted 300 cells per technical replicate and performed Bulk-RNA sequencing using a SMART-seq2 protocol (Figure 3A). The sequencing results of EG-sorted cells and of neuron-sorted cells (*nSyb-Gal4* instead of *GMR-56-Gal4*) show strong enrichment of EG cell markers in the former and strong enrichment of neuronal markers in the latter (Figure 3B). We then compared the transcriptomic profile of EG in *Pink1*^*KO-WS*^ to that of control using differential expression analysis and found 617 deregulated genes in EG (Appendix 1, padj<0.05) (Figure 3C). To understand whether the deregulated genes were enriched for a specific pathway or associated with a cellular component compared to non-significantly deregulated genes, we performed Gene Ontology analysis of the deregulated genes in *Pink1*^*KO-WS*^ flies. Our analysis resulted in generic terms and very diverse pathways that did not allow us to draw further conclusions.

**Figure 3.**
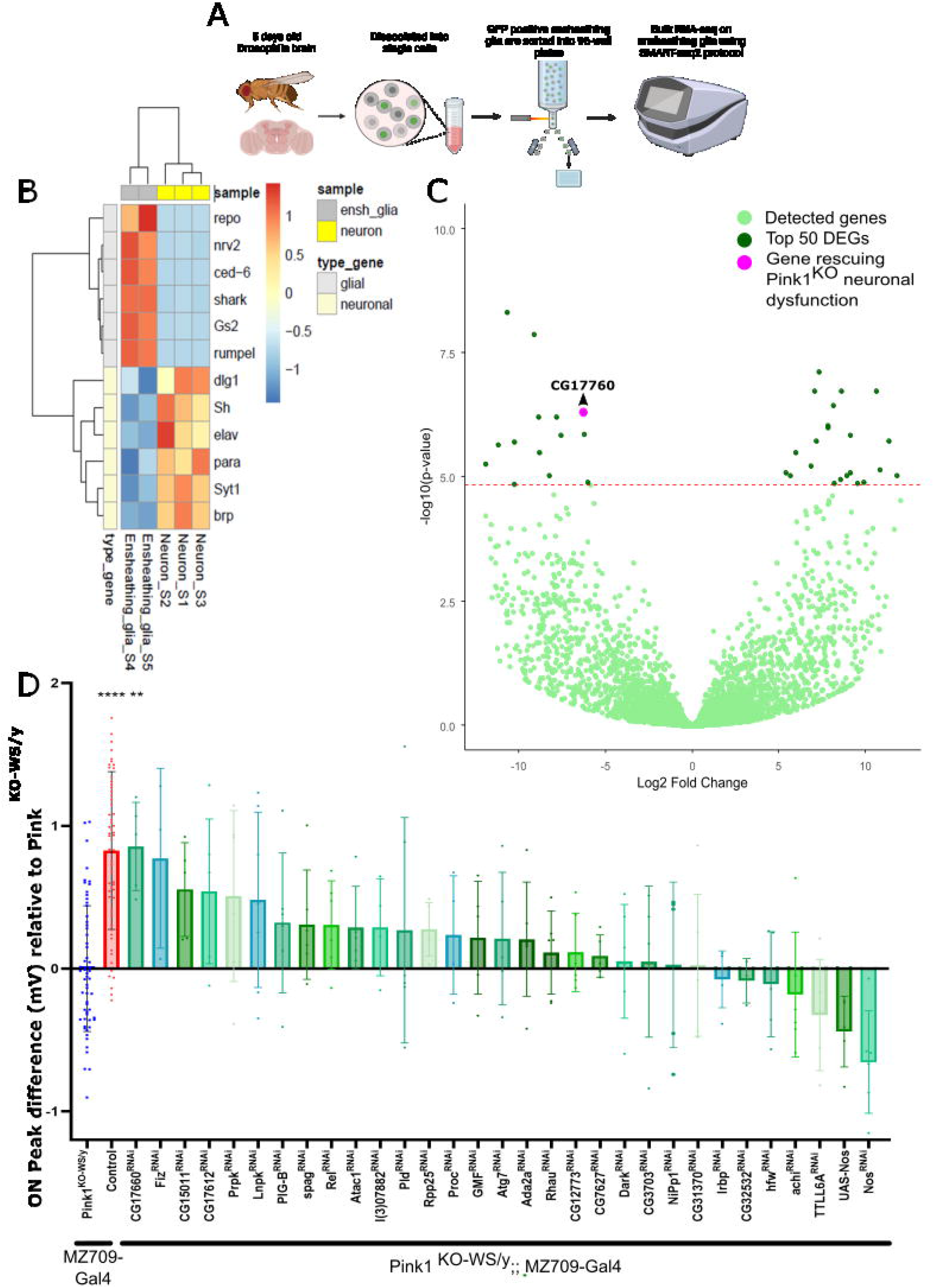
Cell type-specific transcriptomics reveals modifiers of neuronal dysfunction. (**A**)Scheme of cell-type specific transcriptomics (Created in BioRender. Verstreken, P. (2025) https://biorender.com/p56u250). (**B**)Scaled gene expression of representative genes for EG and neurons after sorting EG or neurons using the protocol described in (A) (R Core Team, 2022). N=2, 2 replicates. (**C**)Differentially expressed genes in EG in *Pink1*^*KO-WS*^ compared to control flies, plotted according to their Log2 Fold change and the -Log10 of the adjusted p-value. Intercept in red (-Log10 adjusted p-value = 4.31); light green dots are all detected genes, dark green are the 50 most deregulated genes, and the pink dot is from a gene positive in the genetic screen (D). *Two data points are outside the boundaries of the plot. To determine the transcriptomic profile of each genotype, a N=3 was used, and 3 replicates were performed. (**D**)ERG ON peak value differences to *Pink1*^*KO-WS*^ flies (5 ± 1-day-old) of control (red) and of *Pink1*^*KO-WS*^ flies with DEGs downregulated, or upregulated, specifically in EG (5 ± 1 days old). ON peak values are expressed as the difference to *Pink1*^*KO-WS*^. ANOVA with Dunnett’s test, * is p<0.05, ** is p<0.01, *** is p<0.001. Effect size: η = 0.44. Bars: mean ± SD; points are individual animals N≥3 per genotype. *One data point is outside the boundaries of the plot.

To test if the genes deregulated in EG are modifiers of the neuronal *pink1*^*KO-WS*^ phenotypes, we downregulated and upregulated (when possible) the 50 most deregulated genes (Figure 3C), for which we could find RNAi fly lines, specifically in the EG of *Pink1*^*KO-WS*^ flies and recorded ERGs in 5-day-old animals. Knock down of one of the genes (*CG17660*) resulted in a rescue of the ON transient defects in *Pink1*^*KO-WS*^ (suppressors at p<0.05), and a few exacerbated the defect (Figure 3D). Interestingly, the human orthologs of *CG17660, TMEM87A/B*, have an established role in endosomal sorting (Gaudet et al., 2011; Hirata et al., 2015). Hence, our genetic screening suggests that *Pink1*^*KO-WS*^ phenotypes might be suppressed by downregulating genes involved in vesicle trafficking in EG.

### PD causative vesicle trafficking gene downregulation in EG rescues synaptic dopaminergic neurons afferent loss

Because our ERG-based modifier screen for the *Pink1*^*KO-WS*^ phenotype identified a gene with a known role in vesicle trafficking, we next asked whether *Pink1* in EG also genetically interacts with other PD-causative genes that function in vesicle trafficking, specifically vesicle trafficking components affecting membrane interactions between mitochondria and the endoplasmic reticulum (ER), like *Vps35* and *Vps13* (Brickner & Fuller, 1997; Lesage et al., 2016a; Vilariño-Güell et al., 2011; Zimprich et al., 2011). We find that the knock down of each of these genes in EG also rescues the ON transient defect of *Pink1*^*KO-WS*^ mutants (Figure 4A-B).

**Figure 4.**
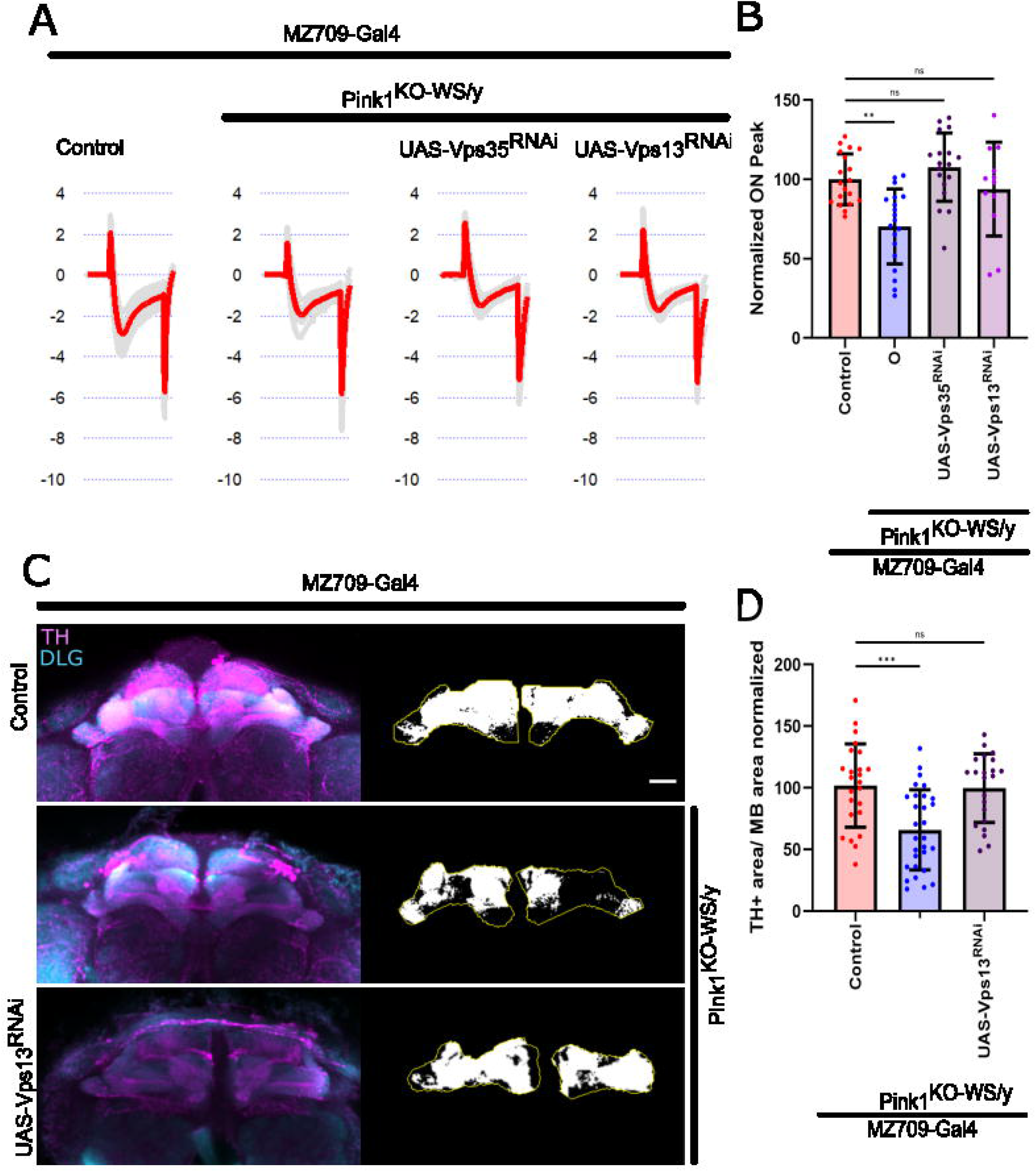
*Vps35* and *Vps13* downregulation in EG rescues synaptic deficits in *Pink1*^*KO-WS*^ flies (with supplementary figure 3) (**A**)Representative ERG traces of control,*Pink1*^*KO-WS*^ flies and *Pink1*^*KO-WS*^ flies with *Vps35* or *Vps13* downregulated in EG. (**B**)Quantification of the normalized ON peak response of flies with the genotypes in (A) (5 ±1-day-old). ANOVA with Dunnett’s multiple comparison test, * is p<0.05, ** is p<0.01, *** is p<0.001. Effect size: η = 0.48. Bars: mean ± SD; points are individual animals N≥10 per genotype, 3 replicates. (**C**)Maximum intensity projection of confocal images of Mushroom Bodies (MB) of aged flies (22 ± 2-day-old) of control, *Pink1*^*KO-WS*^ flies and *Pink1*^*KO-WS*^ flies with *Vps13* downregulated in EG, labeled with anti-TH (cyan) and anti-DLG (magenta), DLG is used to mark the MB neuropil. The black and white image is the middle Z-plane within the region of interest of the MB (ROI, yellow), which is used to represent the thresholded TH area (white). Scale bar: 20 µm. (**D**)Quantification of the dopaminergic synaptic area within MB of aged flies (22 ± 2-day-old). ANOVA with Dunnett’s multiple comparison test, * is p<0.05, ** is p<0.01, *** is p<0.001. Effect size: η =0.23. Bars: mean ± SD; points are individual animals n≥22 per genotype, 5 replicates.

To determine if manipulation of this pathway in EG can also rescue the synaptic innervation defect of PAM DAN onto the mushroom bodies in old (20-25-day-old) *Pink1*^*KO-WS*^ mutants, we expressed RNAi to *Vps13* in EG using *MZ709-Gal4*. Anti-TH labeling (marking DAN synapses) in the MB area (marked by anti-DLG) is significantly reduced in aged *Pink1*^*KO-WS*^, but is rescued to control levels when *Vps13* is knocked down in EG (Figure 4C-D, Supplementary figure 3). These results suggest that manipulation of vesicle trafficking components affecting membrane interactions between mitochondria and the ER in EG regulate glia-neuron crosstalk to maintain synaptic integrity of dopaminergic neuron synapses in the fly brain of *Pink1* mutants.

## DISCUSSION

In this work, we provide evidence for early, non-cell-autonomous activation of EG in a PD-relevant *Drosophila Pink1* model. This occurs already in young *Pink1*-mutant flies when neuronal defects in dopaminergic neurons are not yet measurable. The EG activation and invasion phenotype appears as a protective response, as Pink1-deficient neurons seem to show defects in EG that mimic the response to nerve injury (Doherty et al., 2009; MacDonald et al., 2006). In addition to this, we show that there is also a non-autonomous role of EG to support *Pink1*-deficient neurons. When we manipulate the expression of membrane-lipid trafficking genes, the homologs of the established PD genes VPS13C and VPS35 (Lesage et al., 2016; Vilariño-Güell et al., 2011; Zimprich et al., 2011), specifically in EG, we rescue the dysfunction of *Pink1* mutant neuron defects (ERG and PAM DAN synapse loss). Our work shows the importance of neuron-glia cross-talk in the context of Pink1-deficiency. We suggest (1) there is early EG activation secondary to *Pink1*-loss-induced neuronal impairments; (2) there are specific lipid- and membrane trafficking problems caused by *Pink1* loss in EG that center on mitochondria/ER contacts and (3) there is a convergence of Parkinson-relevant genetic factors that act in EG to maintain neuronal function.

The loss of Pink1 function in the context of Parkinson’s disease has been amply linked to the regulation of mitochondrial health. Pink1 maintains the integrity of the electron transport chain by phosphorylating NDUFA10 (Morais et al., 2009, 2014; Pogson et al., 2014) and when mitochondria are damaged, it phosphorylates Parkin and Ubiquitin to facilitate mitophagy (Kane et al., 2014; Narendra et al., 2010; Narendra & Youle, 2024). The regulation of mitophagy is complex and requires mitochondrial rearrangements controlled by mitochondria-organelle contacts (ER and lysosomes) (Wong et al., 2018, 2019; Yamano et al., 2018). Such contacts mediate inter-organellar lipid exchange as well as facilitate organellar fusion (Kumar et al., 2018; Valadas et al., 2018a). In *Pink1* mutants, these contacts appear to be increased in number, and this causes cellular defects (Grossmann et al., 2023; Valadas et al., 2018).

We found that in EG, *Vps35* functionally interacts with Pink1-induced neuronal dysfunction: its genetic manipulation in EG rescues cell non-autonomous neuronal dysfunction. Loss of *Vps35* itself only in EG is sufficient to rescue neuronal *Pink1* phenotypes. *Pink1* loss-of-function models show increased numbers of mitochondria-ER contact sites (Valadas et al., 2018), affecting mitochondrial calcium levels (Barazzuol et al., 2020; De Brito & Scorrano, 2008; Ham et al., 2023) and dysregulating lipidic ER composition (Valadas et al., 2018) (Figure 5A). Indeed, *Pink1* is necessary for ubiquitination and degradation of Mitofusin (Poole et al., 2010), which is fundamental for the mitochondria-ER tethering (De Brito & Scorrano, 2008). Interestingly, *Vps35* deficiency promotes untethering of mitochondria-ER contact sites by increasing the mitochondrial levels of MUL1, which is necessary for the ubiquitination and degradation of Mitofusin (Puri et al., 2019; Tang et al., 2015; Yun et al., 2014). Hence, conditions that affect mitochondria-ER contact site regulation in EG, by modifying an endosomal trafficking regulator such as *Vps35*, are expected to rescue *Pink1* neuronal dysfunction, possibly by restoring mitochondrial calcium levels and the lipid composition of the ER (Figure 5B); but further work is needed to sort these mechanistic elements in a cell-specific manner.

**Figure 5.**
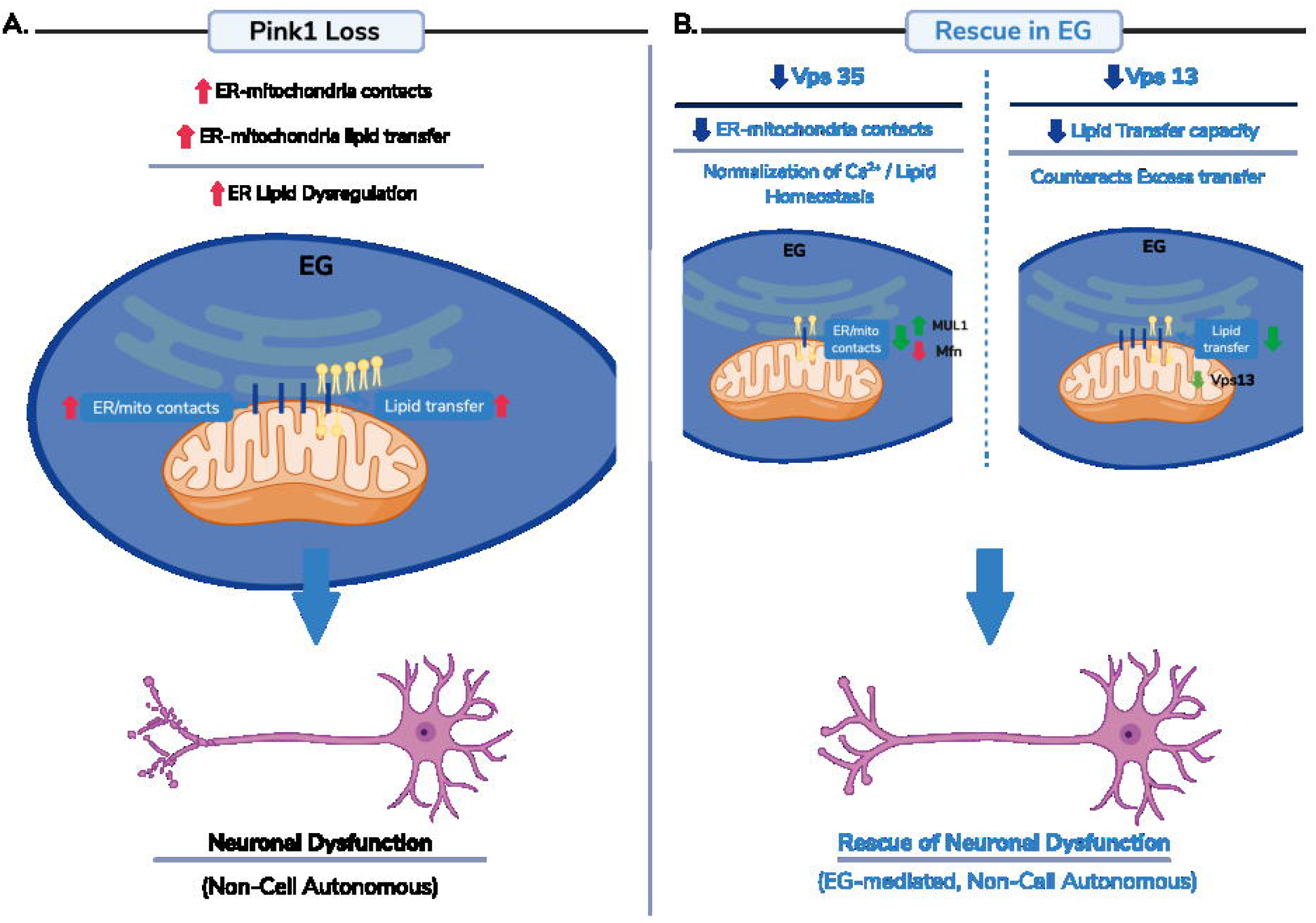
Modulation of ER–mitochondria contact sites and lipid transfer in ensheathing glia (EG) rescues Pink1-dependent neuronal dysfunction. Schematic representation of the suggested model (Created in BioRender. Verstreken, P. (2026) https://BioRender.com/sawj1pw). (A)Loss of *Pink1* leads to an abnormal increase in endoplasmic reticulum (ER)–mitochondria contact sites (represented by blue thick lines), resulting in enhanced ER-to-mitochondria lipid transfer and dysregulation of ER lipid composition (represented by yellow lipids). Increased organelle membrane contacts and lipid flux in EG contribute to neuronal dysfunction in a non–cell autonomous manner. (B)Genetic downregulation of ER–mitochondria contact and lipid transfer regulators in EG rescues Pink1-induced neuronal phenotypes through two convergent mechanisms. Reduction of Vps35 decreases the number of ER–mitochondria contact sites, likely via MUL1-mediated Mitofusin (Mfn) turnover, leading to normalization of calcium and lipid homeostasis. In parallel, downregulation of Vps13, a lipid transfer facilitator at organelle contact sites, limits ER-to-mitochondria lipid transfer capacity, counteracting the excessive lipid flux induced by Pink1 loss. Both interventions restore organelle homeostasis in EG and result in rescue of neuronal dysfunction through a non–cell autonomous mechanism.

Our observation that downregulation of *Vps13* in EG also rescues *Pink1* is in further support of a role for ER-mitochondria contact site regulation. *Vps13* has two mammalian homologs, VPS13A and VPS13C (Velayos-Baeza et al., 2004; Vonk et al., 2017; Vrijsen et al., 2022), which are also mutated in familial Parkinsonism/neurodegenerative disease (Lesage et al., 2016). VPS13A tethers ER to mitochondria and lipid droplets, and VPS13C tethers ER to late endosomes, lysosomes and lipid droplets (Kumar et al., 2018; Muñoz-Braceras et al., 2019; Vrijsen et al., 2022). The function of these proteins is to facilitate lipid transfer between organelles, a function we previously showed to be increased in *Pink1* mutants (Valadas et al., 2018). Hence, the downregulation of *Vps13* would counteract the effects of excessive organelle membrane contact formation and dysregulated lipid homeostasis in *Pink1* mutants (Figure 5B).

Lipid homeostasis in glial cells is crucial for supporting neuronal function by providing energy, protecting against oxidative stress, and regulating synaptic health. In oligodendrocytes, maintaining myelin sheaths is critical (Ettle et al., 2016; Lappe-Siefke et al., 2003). While fly EG do not generate such structures, they do wrap processes around neuropil regions, also requiring lipid membrane production (Kremer et al., 2017; Pogodalla et al., 2021). Through lipid metabolism, glia also supply neurons with essential metabolites, cholesterol, and signaling molecules that aid in membrane integrity, synaptic remodeling, and neuroprotection (Delgado et al., 2018; Otto et al., 2018; Saab et al., 2016; Suárez-Pozos et al., 2020). Further work is now required to define which functions, supported by organelle contact sites in glial cells, drive cell-nonautonomous protective mechanisms in neurons. Our work in flies indicates these glial defects precede dopaminergic synapse loss, and, importantly, that rescuing the glial defects helps to prevent dopaminergic problems later in life.

## METHODS

### Resource availability

#### Lead Contact

Further information and requests for resources and reagents should be directed to the lead contact, Patrik Verstreken.

#### Materials Availability

Data, code, Drosophila models, and reagents are available upon request.

### Fly stocks

The fruit flies were maintained in an incubator at 25°C under a 12h:12h light-dark cycle and provided with a standard diet consisting of corn meal and molasses. For experiments, only male flies were used. Flies were raised in parallel on the same batch of food, which was exchanged every three to four days and kept in only male population of similar density. Flies were aged to 5 ± 1-day-old, and 22 ± 2-day-old, as indicated for immunohistochemistry and ERGs, respectively.

To create a set of *Drosophila* models for Parkinson’s disease, CRISPR/Cas9-based gene editing was utilized, as outlined in (Kaempf et al., 2026; Pech et al., 2025). In brief, each knock-out line contained an attP-flanked w+ reporter cassette that replaced the first shared exon across different isoforms of the targeted gene. For this study, we used knockout flies for *Pink1* from this collection (Kaempf et al., 2026). The “control” strain denotes a semi-isogenized *w*^*1118*^ strain that underwent backcrossing to Canton-S for 10 successive generations, resulting in a strain termed Canton-S-w1118. To generate the fly line *yw; UAS-His2Av::eGFP*.*VK27/TM3Sb*, we created the pUASTattB_His2AV-GFP plasmid, linearizing the pUAST.attB (Bischof et al., 2007) with EcoRI-BamHI. A gBlock, containing the His2AV, GS linker and eGFP sequence, was cloned into this linearized plasmid with Gibson Assembly. The plasmid was inserted into the VK27 landing site by BestGene. The sequence His2AV-GS-eGFP gblock is:

TGAATAGGGAATTGGGcAAacATGGCTGGCGGTAAAGCAGGCAAGGATTCGGGCAAGGCCAAGGCGAAGGCGGTATCGCGTTCCGCGCGCGCGGGTCTTCAGTTCCCCGTGGGTCGCATCCATCGTCATCTCAAGAGCCGCACTACGTCACATGGACGCGTCGGAGCCACTGCAGCCGTGTACTCCGCTGCCATATTGGAATACCTGACCGCCGAGGTCCTGGAGTTGGCAGGCAACGCATCGAAGGACTTGAAAGTGAAACGTATCACTCCTCGCCACTTACAGCTCGCCATTCGCGGAGACGAGGAGCTGGACAGCCTGATCAAGGCAACCATCGCTGGTGGCGGTGTCATTCCGCACATACACAAGTCGCTGATCGGCAAAAAGGAGGAAACGGTGCAGGAcCCGCAGCGGAAGGGCAACGTCATTCTGTCGCAGGCCTACGGTTCAGGCGGAGGTGGCAGCGGCGGTGGCGGATCCATGGTGAGCAAGGGCGAGGAGCTGTTCACCGGGGTGGTGCCCATCCTGGTCGAGCTGGACGGCGACGTAAACGGCCACAAGTTCAGCGTGTCCGGCGAGGGCGAGGGCGATGCCACCTACGGCAAGCTGACCCTGAAGTTCATCTGCACCACCGGCAAGCTGCCCGTGCCCTGGCCCACCCTCGTGACCACCCTGACCTACGGCGTGCAGTGCTTCAGCCGCTACCCCGACCACATGAAGCAGCACGACTTCTTCAAGTCCGCCATGCCCGAAGGCTACGTCCAGGAGCGCACCATCTTCTTCAAGGACGACGGCAACTACAAGACCCGCGCCGAGGTGAAGTTCGAGGGCGACACCCTGGTGAACCGCATCGAGCTGAAGGGCATCGACTTCAAGGAGGACGGCAACATCCTGGGGCACAAGCTGGAGTACAACTACAACAGCCACAACGTCTATATCATGGCCGACAAGCAGAAGAACGGCATCAAGGTGAACTTCAAGATCCGCCACAACATCGAGGACGGCAGCGTGCAGCTCGCCGACCACTACCAGCAGAACACCCCCATCGGCGACGGCCCCGTGCTGCTGCCCGACAACCACTACCTGAGCACCCAGTCCGCCCTGAGCAAAGACCCCAACGAGAAGCGCGATCACATGGTCCTGCTGGAGTTCGTGACCGCCGCCGGGATCACTCTCGGCATGGACGAGCTGTACAAATAAGGGTACCTCTAGAGGATCTT.

Flies were backcrossed for five generations into an inbred Canton-S strain harboring the *w*^*1118*^ mutation. The genotypes used in this study are listed in the key resource table and Supplementary table 1.

### Olfactory Receptor Neurons (ORNs) axotomy

ORN bilateral axotomy was performed on 4±1 day old flies by surgical ablation of the third antennal segment. Briefly, flies were anesthetized with CO_2_, and then both antennae were removed with Dumont #5 forceps and returned to the vial (M. Purice, 2020; M. D. Purice et al., 2017). Flies recovered for 24 h on food at 25°C before dissection.

### Immunohistochemistry and confocal imaging – invasion phenotype

Immunohistochemistry was performed on adult fly brains of 5±1-day-old, with at least two independent experiments. The brains were dissected in ice-cold PBS and fixed for 20 min in freshly prepared 3.7% paraformaldehyde (in 1x PBS, 0.3% Triton X-100 [PBX] [Sigma]) at room temperature (RT), followed by three 15-min washes in PBX at RT on a shaker. Then, the brains were incubated for 1 h in blocking solution (PBX, 10% normal goat serum [NGS]) at RT. Following blocking, the brains were incubated with primary antibodies (rabbit anti-GFP [Thermo Fisher Scientific], 1:1000, mouse anti-brp [DSHB, nc82], 1:100) in blocking solution at 4°C overnight, followed by three 15-min washes in PBX at RT. Secondary antibodies (goat anti-rabbit Alexa488, goat anti-mouse Alexa555, both at 1:1000 [Thermo Fisher Scientific]) in PBX with 10% NGS were applied overnight at 4°C. Afterwards, the brains were washed three times for 15 min in PBT at RT on a shaker and mounted with the anterior facing up in RapiClear 1.47 (SUNJin Lab). Imaging was performed using a Nikon A1R confocal microscope with a 40x (NA 1.15) water immersion lens; Z-stacks of the entire antennal lobes were acquired. The acquisition was carried out using a Galvano scanner, with a zoom factor of 1, scan speed of 0.5, and line averaging set to 2. All images were captured with a pinhole of 0.9 Airy units and a resolution of 1024 × 1024. Z-stacks (with 0,5 μm step intervals) were obtained for data acquisition, and the same imaging settings were applied across all genotypes and sessions. Image analysis was performed using Fiji (Schindelin et al., 2009). Image analysis was performed using Fiji (Schindelin et al., 2009). Each antennal lobe is composed of multiple glomeruli, and EG separate them and invade them when active (Doherty et al., 2009; Pogodalla et al., 2021). To quantify EG invasion in the antennal lobe, we calculated the invasion in every single glomerulus. To do so and be consistent with the section of the antennal lobe analyzed, anti-brp was used to identify the 3 Z-planes where DM6 and DM1 glomeruli were present (B. Wu et al., 2017). Then, to automatically detect and segment each glomerulus of the antennal lobe section, regions of interest (ROIs) were defined in the SUM projection of the 3 Z-planes by using the Fiji plugin Stardist (Weigert et al., 2019).

To quantify the EG invasion in the ROIs, anti-GFP SUM projection of the same 3 Z-planes was used. The GFP intensity was detected in each ROI, and it was normalized to the area of each ROI (GFP intensity/glomeruli area) to calculate the EG invasion in each glomerulus. Then, the normalized GFP intensity of each glomerulus of an antennal lobe was summed and normalized for the number of glomeruli detected in the antennal lobe. For each experiment, the EG antennal lobe invasion of each fly was normalized to the mean of the control. Representative images show the SUM projections of the 3 Z-planes.

### Immunohistochemistry and confocal imaging – TH staining

Immunohistochemistry was performed on adult fly brains of 22±2-day-old, with at least two independent experiments. The brains were dissected in ice-cold PBS and fixed for 20 min in freshly prepared 3.7% paraformaldehyde (in 1x PBS, 0.2% Triton X-100 [PBX]) at room temperature (RT), followed by three 15-min washes in PBX at RT on a shaker. Then, the brains were incubated for 1 h in blocking solution (PBX, 10% normal goat serum [NGS]) at RT. Following blocking, the brains were incubated with primary antibodies (rabbit anti-TH [Sigma], 1:200, mouse anti-DLG [DSHB], 1:100) in blocking solution at 4°C for 1.5 to 2 days, followed by three 15-min washes in PBX at RT. Secondary antibodies (goat anti-rabbit Alexa488, goat anti-mouse Alexa555, both at 1:500 [Thermo Fisher Scientific]) in PBX with 10% NGS were applied overnight at 4°C. Afterwards, the brains were washed three times for 15 min in PBT at RT on a shaker and mounted with the anterior facing up in RapiClear 1.47 (SUNJin Lab). Imaging was performed using a Nikon A1R confocal microscope with a 20x (NA0.95) water immersion lens; Z-stacks of the entire brain were acquired. The acquisition was carried out using a Galvano scanner, with a zoom factor of 1, scan speed of 0.5, and line averaging set to 2. All images were captured with a pinhole of 2.3 Airy units and a resolution of 1024 × 1024. Z-stacks (with 3 μm step intervals) were obtained for data acquisition, and the same imaging settings were applied across all genotypes and sessions. Image analysis was performed using Fiji (Schindelin et al., 2009). To quantify dopaminergic neuron innervation of the mushroom body (MB), anti-DLG was used to identify the five Z-planes containing the synaptic region of the MB lobes. The ROI for the MB was defined in the sum projection of the five z-planes. To exclude background signal comparable to control, the area of anti-TH fluorescence was thresholded (using the default threshold for all z-planes) within the selected z-stacks. Quantification of the thresholded area within the ROI was performed in each z-plane, then summed and normalized to the MB ROI area for each brain individually (TH+ area/MB area). For each experiment, the individual TH+ area/MB area values were normalized to the mean of the control. Representative images show the maximum projection of five z-planes and the thresholded middle z-plane.

### Electroretinograms (ERG)

ERGs were recorded from flies 5±1-day-old as previously described (Heisenberg, 1971; Slabbaert et al., 2016). Flies were immobilized on glass microscope slides using double-sided tape. For recordings, glass electrodes (borosilicate, 1.5 mm outer diameter) filled with 3 M NaCl were placed in the thorax as a reference and on the fly eye for recordings. Each fly was exposed to 5 cycles of 3 s of darkness, followed by a 1-s of light stimuli with LED illumination. Response to the stimuli was recorded using Axoscope 10.7 and analyzed using Clampfit 10.7 software (Molecular Devices). ERG traces were analyzed with Igor Pro 6.37 (Wave Metrics) using a custom-made macro.

### Single-cell dissociation for cell-type specific transcriptomics

To test that EG could be isolated from the rest of the brain, we used two cohorts *GMR56F03> UAS-His2Av::eGFP*.*VK27* and *nSyb-Gal4> UAS-His2Av::eGFP*.*VK27*. To identify differentially expressed genes in EG, two cohorts were used: the control *w*^*1118*^ expressing GMR56F03> UAS-His2Av::eGFP.VK27 and *Pink1*^*KO-WS*^ expressing *GMR56F03> UAS-His2Av::eGFP*.*VK27*. For each genotype, ten brains of flies 5±1-day-old were collected, with most of the laminae removed, from each experimental repeat. The dissections were performed in ice-cold PBS that contained 5 μM Actinomycin D (Sigma) to inhibit changes in gene expression during tissue processing(Y. E. Wu et al., 2017). To prevent any bias caused by different experimental batches, all the genotypes were processed in parallel. To minimize variability across batches, the same reagents and procedures were used throughout the experiment. Dissections were completed within one hour. The dissection order and the assignment of genotypes to dissectors were alternated to reduce any experimenter-related variation.

For the dissociation of brain tissue, a mix of Dispase I (0.6 mg/ml) (Sigma), Collagenase I (30 mg/ml) (Thermo Fisher), Trypsin (0.5x) (Thermo Fisher), and 5 μM Actinomycin D was used. The brains were incubated for 20 min at 25°C with 1000 rpm shaking, performing four triturations at 5-min intervals. After dissociation, the cell suspension was washed with PBS containing 5 μM Actinomycin D and then filtered through a 10 μm strainer (Pluriselect), using 300 μl of PBS EDTA (Sigma) 1μM. DAPI (Sigma) (1:300.000) was added after filtration.

### Fluorescence Activated Cell Sorting (FACS) for cell-type specific transcriptomics

Immediately after the dissociation, cells were sorted using a FACSAria Fusion (BD Biosciences) flow cytometer equipped with 4 lasers (405nm, 488nm, 561nm and 640nm) and a 100 µm nozzle at 20 psi. Live (DAPI-negative), GFP-positive cells were sorted into 96-well PCR plates at 300 cells per well (3 wells per genotype) using a 4-way purity mask with the FACSDiva software v9.0.1 (BD Biosciences).

### Bulk transcriptomics of EG cells

Library preparations and sequencing for the Bulk RNA seq of the EG cells were performed using a modified Smart-seq2 protocol based on a previously published protocol (Picelli et al., 2013). Briefly, 3 μl of cell lysis buffer (0.1% Triton X-100, Sigma Aldrich; 1U/ul of RNase Inhibitor, RNase OUT, Thermo Fisher; 2.5 μM Oligo dT(25), IDT; and 2.5 mM dNTP, Promega) was aliquoted in triplicate to 96-well plates (4titude, Cat. No. 4TI-0960/C). Roughly 300 cells were sorted into the wells with lysis buffer and the plate was spun down at 2000xg for 1 min prior to storage at -80 °C. For the first strand synthesis, the Smart-Seq 2 plates were thawed to RT for a minute and subsequently spun down at 2000 xg for a minute. The RNA denaturation is carried out at 72°C for 10 min and then flash cooled on ice for 5 min. First strand synthesis mix (1X SuperScript II first strand synthesis buffer, Thermo Fisher; 5 mM DTT, Thermo Fisher; 100 U SuperScript II reverse transcriptase enzyme, Thermo Fisher; 10 U RNAse OUT, Thermo Fisher; 1M Betaine, Sigma Aldrich; 6 mM MgCl2, Thermo Fisher; and 1 uM TSO, IDT Technologies) is added to the cell lysis mix to a total volume of 10 μl. The First strand synthesis reaction was carried out with following program: 42°C for 90 min; 10 cycles of {50°C for 2 min; 42°C for 2 min}; 72°C for 15 min; 4°C indefinite hold. PCR amplification of the first strand product was performed by adding 15 μl of the PCR mix to the first strand product (1X KAPA HiFi HotStart Ready Mix, Roche; and 0.1 μM of IS PCR primer, IDT Technologies). 22 cycles of the following PCR program were used to amplify the first strand product: 98°C for 3 min; 22 cycles of {98°C for 20 sec; 67°C for 15 sec; 72°C for 6 min}; 72°C for 5 min; 4C indefinite hold. 20 μl of Ampure XP was added to each well and mixed. Standard Ampure XP purification was carried out as per manufacturer’s recommendation, and the cDNA library was eluted in 17.5 μl of Elution buffer. 17 μl of the eluted library was,transferred to a fresh 96-well plate (4titude, Cat. No. 4TI-0960/C). 5 μl of Tn5 tagmentation mix (1X Tagment DNA buffer, Illumina; ATM mix, Illumina, and cDNA library 1 ng) was prepared for cDNA fragmentation. Tn5 tagmentation was performed with the following program: 55°C for 10 min; 4°C indefinite hold. Tagmentation reaction was stopped by quenching the reaction with 1.25 μl of NT buffer for 5 min (Illumina). 1.25 μl of the i5 and i7 Illumina indexes were added to the plates. Finally 3.75 μl of NPM master mix was added to the plate and mixed well and put for the index PCR amplification: 72°C for 3 min; 95°C for 30 sec; 12 cycles of {95°C for 10 sec; 55°C for 30 sec; 72°C for 30 sec}; 72°C for 5 min, and 4°C indefinite hold.

The indexed PCR products were pooled in a single tube, and 0.8X Ampure XP purification (Beckman Coulter) was carried out as per the manufacturer’s recommendation, and finally, the sequencing library was eluted in 35.5 μl of Elution buffer (Qiagen).

SMART seq 2 libraries were sequenced on the NextSeq 500 (Illumina) sequencing platform. Sequencing was done as per the protocol recommendations: paired-end read of 76 bps (read 1), 76 bps (read 2), 8 bps (index 1) and 8 bps (index 2). For a targeted sequencing depth of ∼30 million reads per sample.

### Analysis of RNAseq data - Neuron vs EG

Raw reads from neuron- vs. glia-specific FACsorting experiments were processed with the nf-core/rnaseq pipeline (Patel et al., 2020) with default parameters and the BDGP6 reference genome. The STAR-aligned and salmon-quantified expression matrix was then subset to genes known to be enriched in neurons or glia and displayed as a per gene scaled heatmap.

### Analysis of RNAseq data - EG-specific differentially expressed genes

After FACsorting of *Pink*^*KO-ws*^ vs. control ensheathing glia, library preparation and sequencing resulting FASTQ files were cleaned with fastp (Chen et al., 2018) with default settings. Aligning and counting were carried out with STAR (Dobin et al., 2013) and the 4th 2020 FlyBase *Drosophila melanogaster* release (r6.35) with the –quantMode flag set as GeneCounts. For differential gene expression testing we used DESeq2 (Love et al., 2014), where we fit a negative binomial model and carried out the Wald-test. We removed the experimental batch as a covariate according to the design formula ∼date+genotype. All genes below a Benjamini-Hochberg corrected p-value of 0.05 were considered deregulated.

### ScRNAseq data loading, filtering, clustering, and differential expression analysis

Raw counts data from (Pech et al., 2025), alongside cell annotations provided by the authors, was loaded into Scanpy (v1.9.1) (Wolf et al., 2018) and subset to only include cells in the young control (<= 6-day-old) or *Pink1*^*KO-WS*^ samples. After subsetting the data, filtering was performed to remove any cell with less than 250 genes expressed or a percentage of counts coming from mitochondrial genes greater than 15%, the data was then normalized to a total of 10.000 counts and log transformed. Highly variable genes were detected, total_counts and percent of mitochondrial reads were regressed out, and finally the data was scaled to unit variance with a zero mean, clipped to a max of 10.

After data pre-processing, a PCA was performed, Harmony (Korsunsky et al., 2019) was used to correct batches using “sample_id” as a batch key, and 122 components were used for dimensionality reduction and clustering analyses. Leiden clustering was performed with a resolution of 8.0 and using the annotations from (Pech et al., 2025), all cells within each cluster were labeled by taking the most common, original annotation per cluster.

Per cluster, a differential analysis was performed. First, pseudobulk counts were generated by summing the counts per cell from all cells per sample. These counts were then used in DESeq2 (v1.44.0) (Love et al., 2014) comparing Control vs *Pink1*^*KO-WS*^ samples. The total number of genes significantly up- or downregulated per cluster (padj <= 0.05 and an abs (log2foldchange) >= 1.5) was counted and plotted.

### Statistical analysis

GraphPad Prism was used for visualization and to determine statistical significance. Datasets were tested for normal distribution using the D’Agostino-Person Omnibus and the Shapiro-Wilk normality test. For a normally distributed dataset ordinary one-way ANOVA was used, followed to correct for multiple comparisons by a post hoc Tukey’s test when comparing all the datasets with each other or Dunnett’s test when comparing all the datasets to a general control. For non-normally distributed datasets Kruskal-Wallis test is used, followed by a post hoc Dunn’s test to correct for multiple comparisons. Significance levels are defined as * is p<0.05, ** is p<0.01, *** is p<0.001, and ns, not significant. Effect size is calculated with eta-squared when one-way ANOVA is used, indicated in the figure legend as η. ‘N’ in the legends is used to indicate how many animals were analyzed. Data are plotted as mean ± SD. Specifics on the statistical test used for each analysis are reported in the figure legends.

## Key resources table

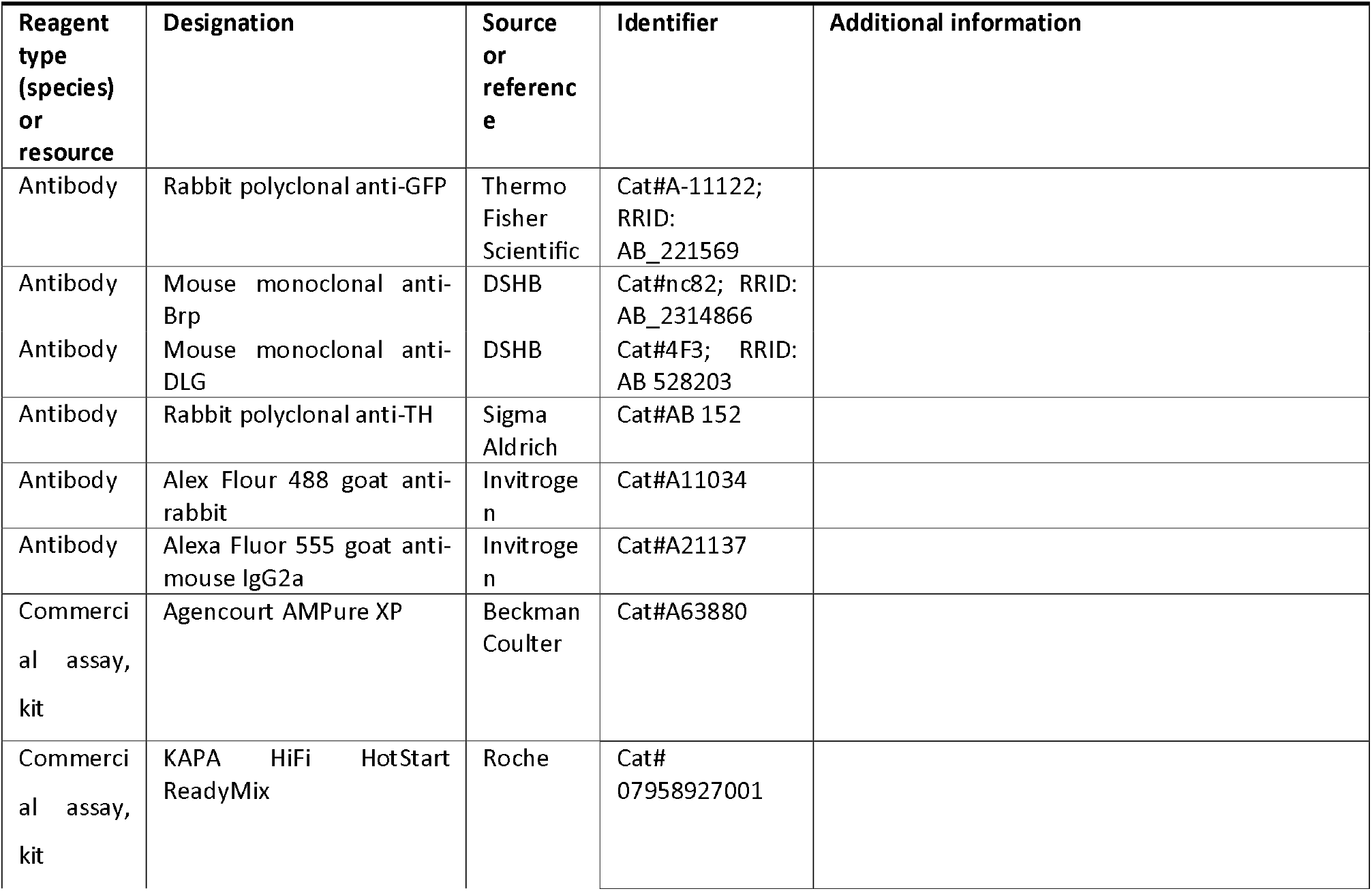
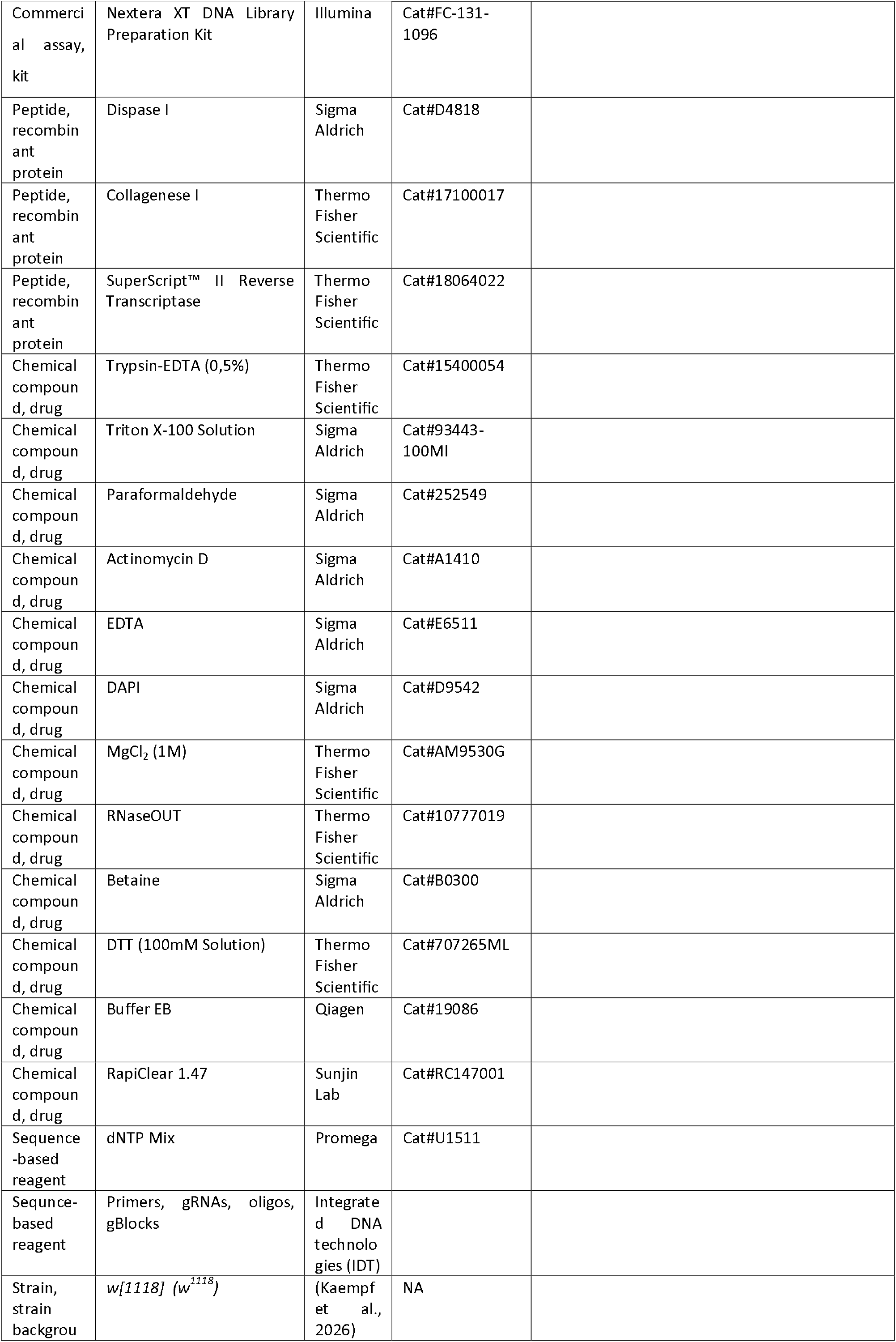
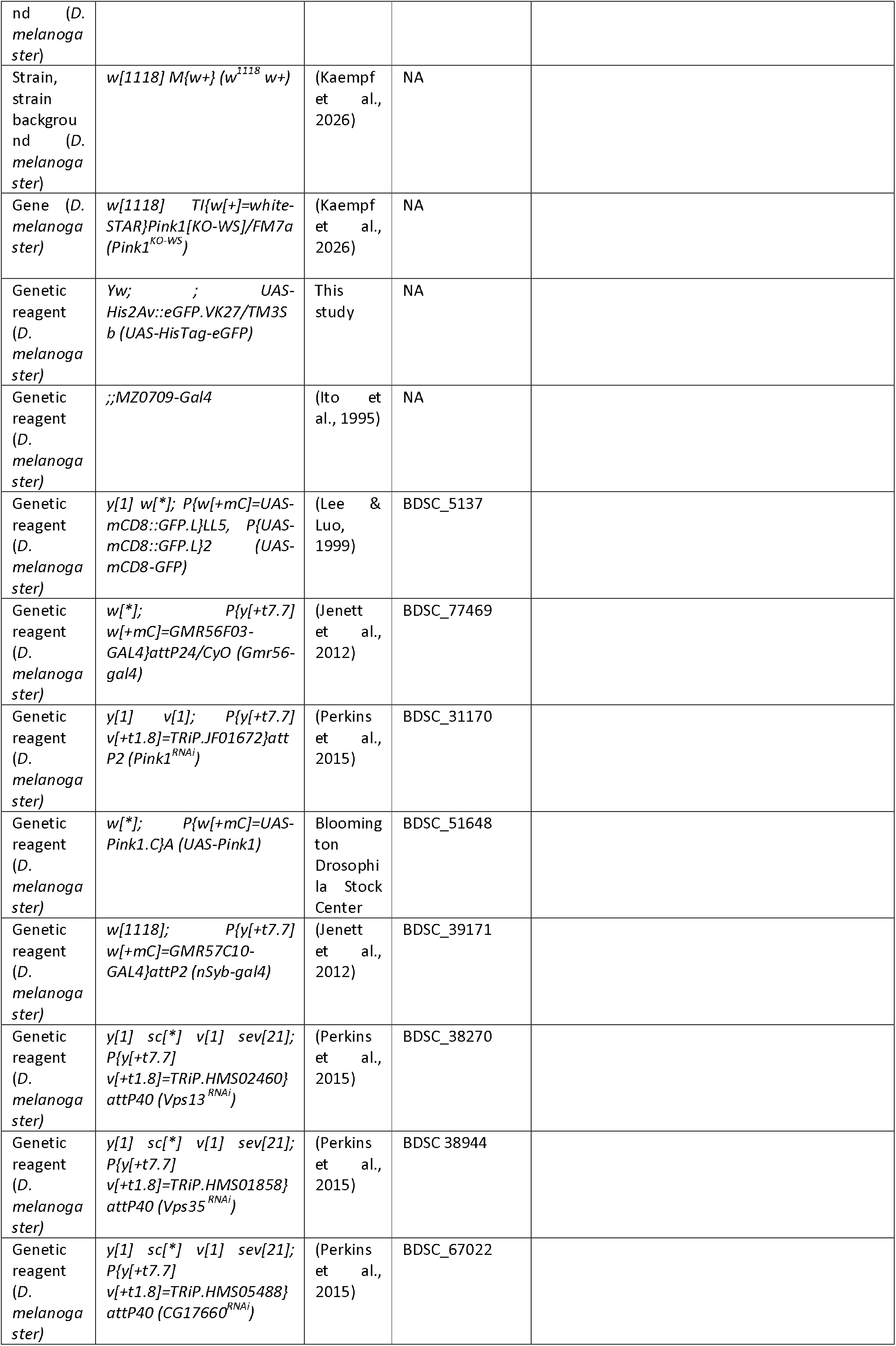
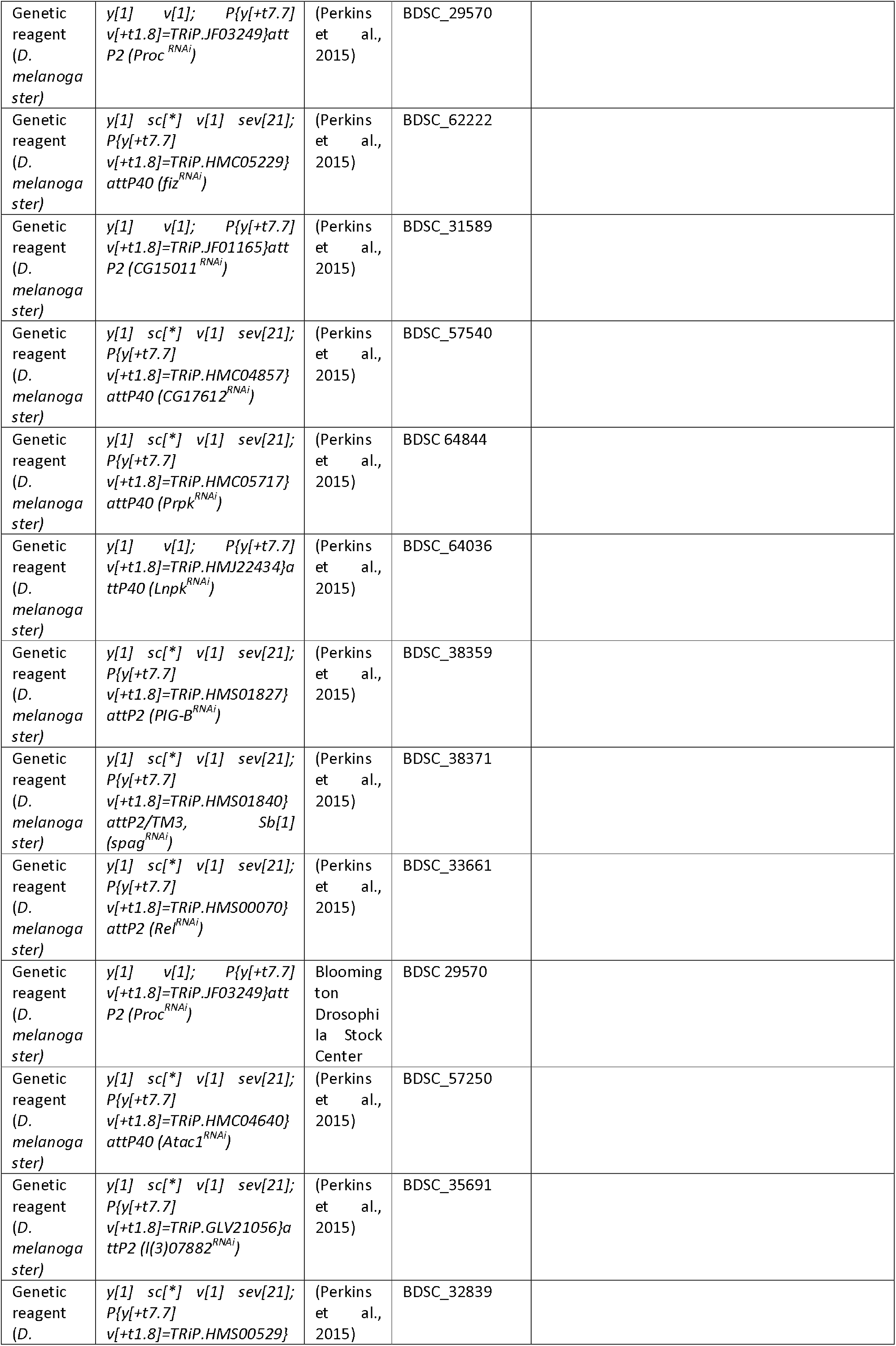
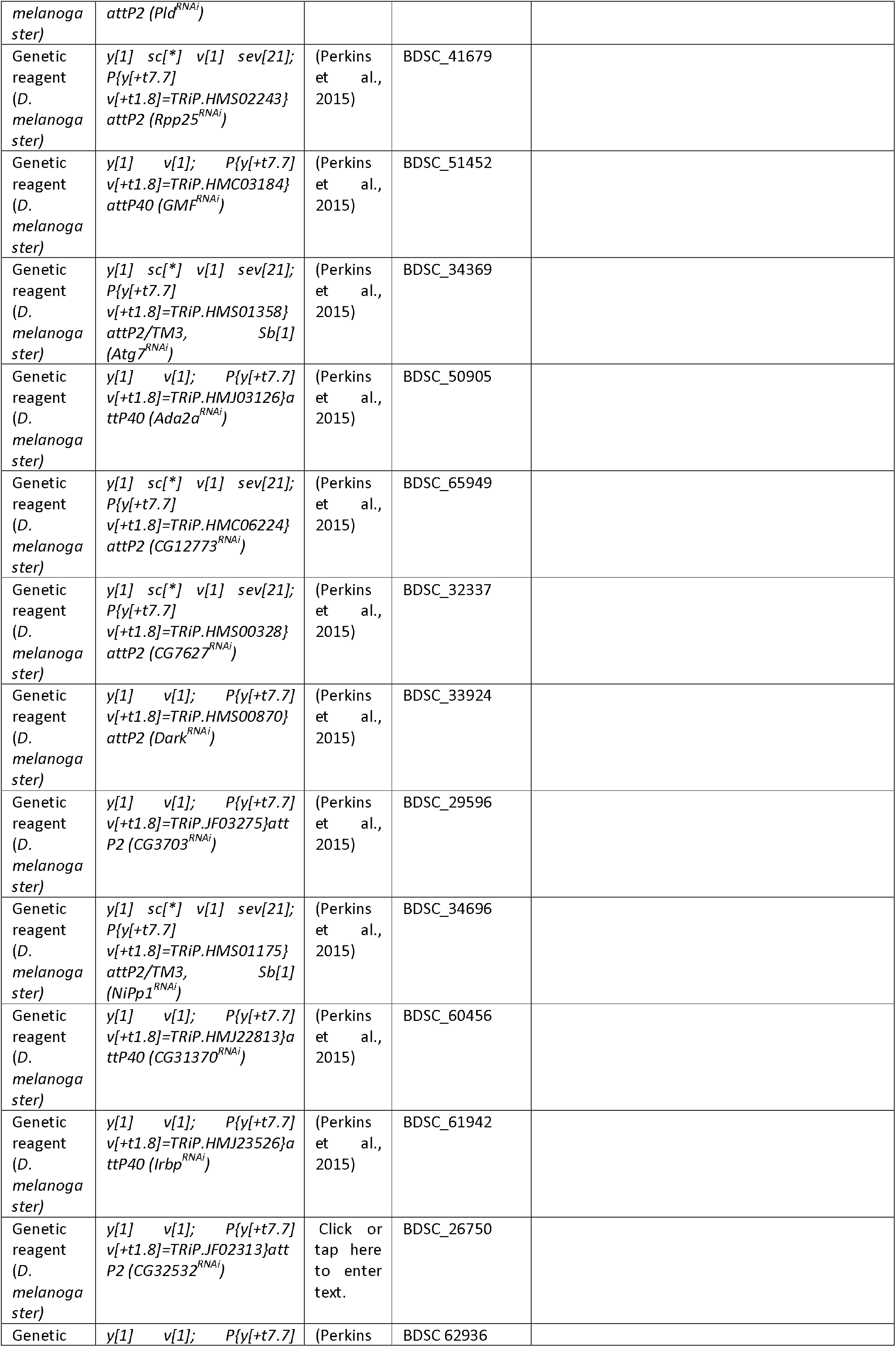
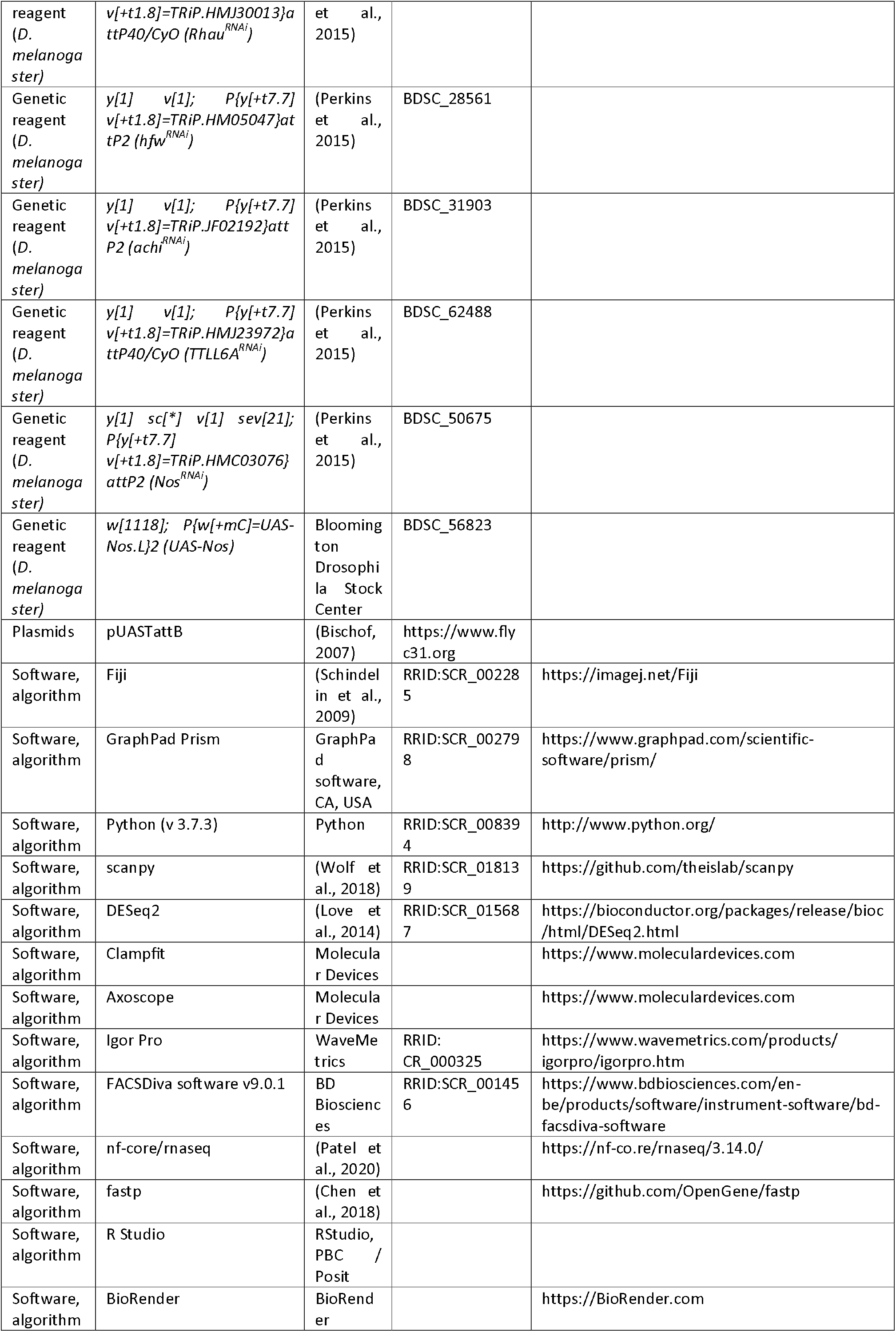
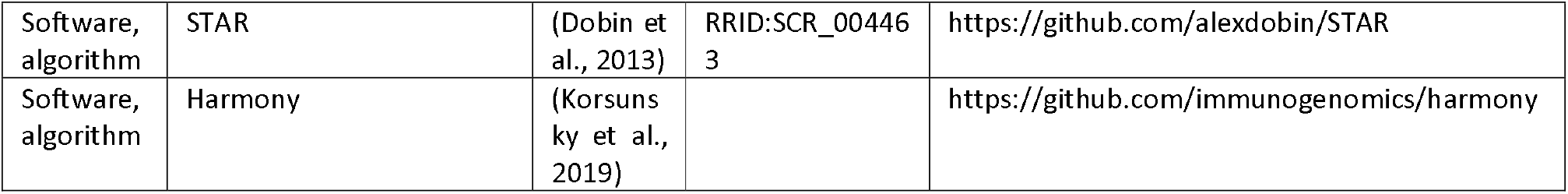

## Data availability statement

RNA sequence data were deposited in GEO (accession number GSE322929). All data generated or analyzed during this study are included in the manuscript and supporting files; source data file has been provided for all figures-SourceData 1.

The following data set was generated

Ghezzi L, Kuenen S, Pech U, Schoovaerts N, Kilic A, Poovathingal S, Davie K, Lamote J, Praschberger R, Verstreken P (2026) **NCBI Gene Expression Omnibus** ID GSE322929. Parkinson’s Disease-Associated *Pink1* Loss Disrupts Ensheathing Glia And Causes Dopaminergic Neuron Synapse Loss.

https://www.ncbi.nlm.nih.gov/geo/query/acc.cgi?acc=GSE322929

The following previously published data sets were used

Janssens JPech UAerts SVerstreken P (2025) NCBI Gene Expression Omnibus ID GSE228843. Rescuing early Parkinson-induced hyposmia prevents dopaminergic system failure [fruit fly].

https://www.ncbi.nlm.nih.gov/geo/query/acc.cgi?acc=GSE228843

## Funding

Leuven University Fund and Opening the Future, ERC, the Chan Zuckerberg Initiative, a Methusalem grant from the Flemish government, KU Leuven BOF, Fund Jacqueline Cigrang administered by the KBS, the KBS, fund Generet and FWO Vlaanderen to PV

EMBO long-term fellowship to RP

DFG fellowship to UP

FWO PhD mandate to AK

## Acknowledgments

We thank the Vlaams Supercomputer Centrum, the VIB Nucleomic and Bioimaging Cores, and the Bloomington *Drosophila* Stock Center (NIH P40OD018537). We are grateful to the members of the Verstreken lab for valuable discussions. Research support was provided by Leuven University Fund and Opening the Future, ERC, the Chan Zuckerberg Initiative, a Methusalem grant from the Flemish government, KU Leuven BOF, Fund Jacqueline Cigrang administered by the KBS, the KBS, fund Generet and FWO Vlaanderen to PV an EMBO long-term fellowship to RP, a DFG fellowship to UP and an FWO PhD mandate to AK. Cartoons/models/illustrations were created in https://BioRender.com. P.V. is an alumnus of the FENS-Kavli Network of Excellence.

## Author contributions

Conceptualization, L.G., R.P., P.V.;

Methodology, L.G., U.P., N.S., J.L., K.D., R.P., P.V., S.K., A.K.;

Software, L.G., K.D., R.P.;

Validation, L.G., K.D., R.P.;

Formal analysis, L.G., K.D., R.P.;

Investigation, L.G., U.P., N.S., J.L., K.D., R.P., S.K., A.K.;

Writing -Original Draft, L.G., R.P., P.V.;

Writing – review and editing, all co-authors read and edited the manuscript.

Visualization, L.G., K.D., R.P., S.K.;

Supervision, R.P., P.V.;

Project administration, L.G.;

Funding acquisition, R.P., P.V.;

---

**Supplementary Figure 1.**
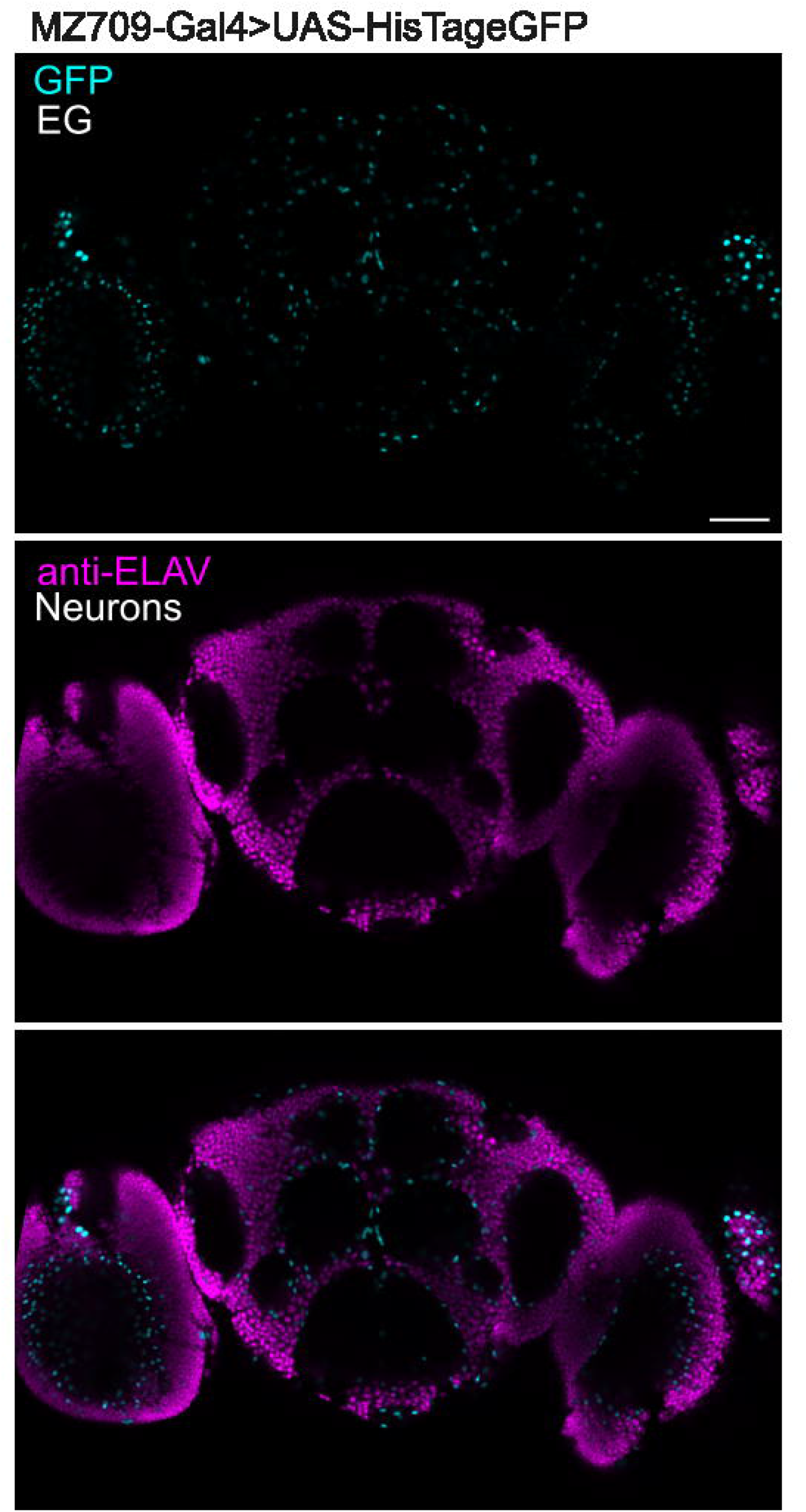
Representative confocal image of a (5 ± 1-day-old) MZ709-Gal4/+;; UAS-HisTageGFP/+ brain stained with anti-GFP (Cyan) and anti-ELAV (Magenta). Scale bar: 50µm; N= 3, 1 replicate.

**Supplementary Figure 2.**
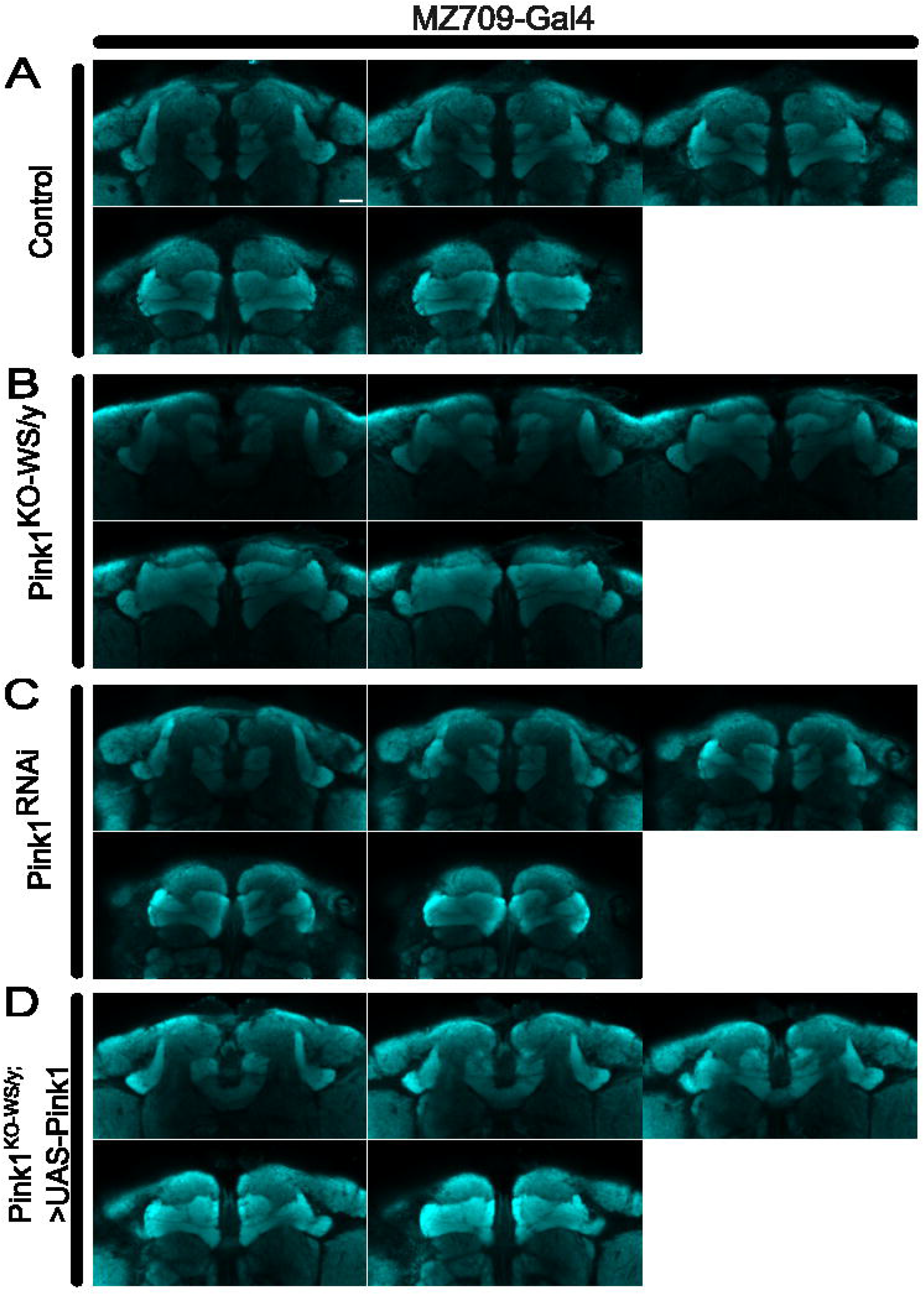
Examples of confocal Z-stacks of anti-DLG labeled fly brains (22 ± 2-day-old) of the indicated genotypes. Such images were used to delineate ROIs to quantify DAN innervation onto the MBs. (A) control (w1118/y;; MZ709-Gal4/+); (B) Pink1KO-WS/Y;; MZ709-Gal4/+; (C) flies where *Pink1* is downregulated in EG (w1118/y;; MZ709-Gal4/ Pink1RNAi); (D) Pink1KO-WS flies with expression of wild type *Pink1* in EG (Pink1KO-WS/Y;; MZ709-Gal4/UAS-Pink1).

**Supplementary Figure 3.**
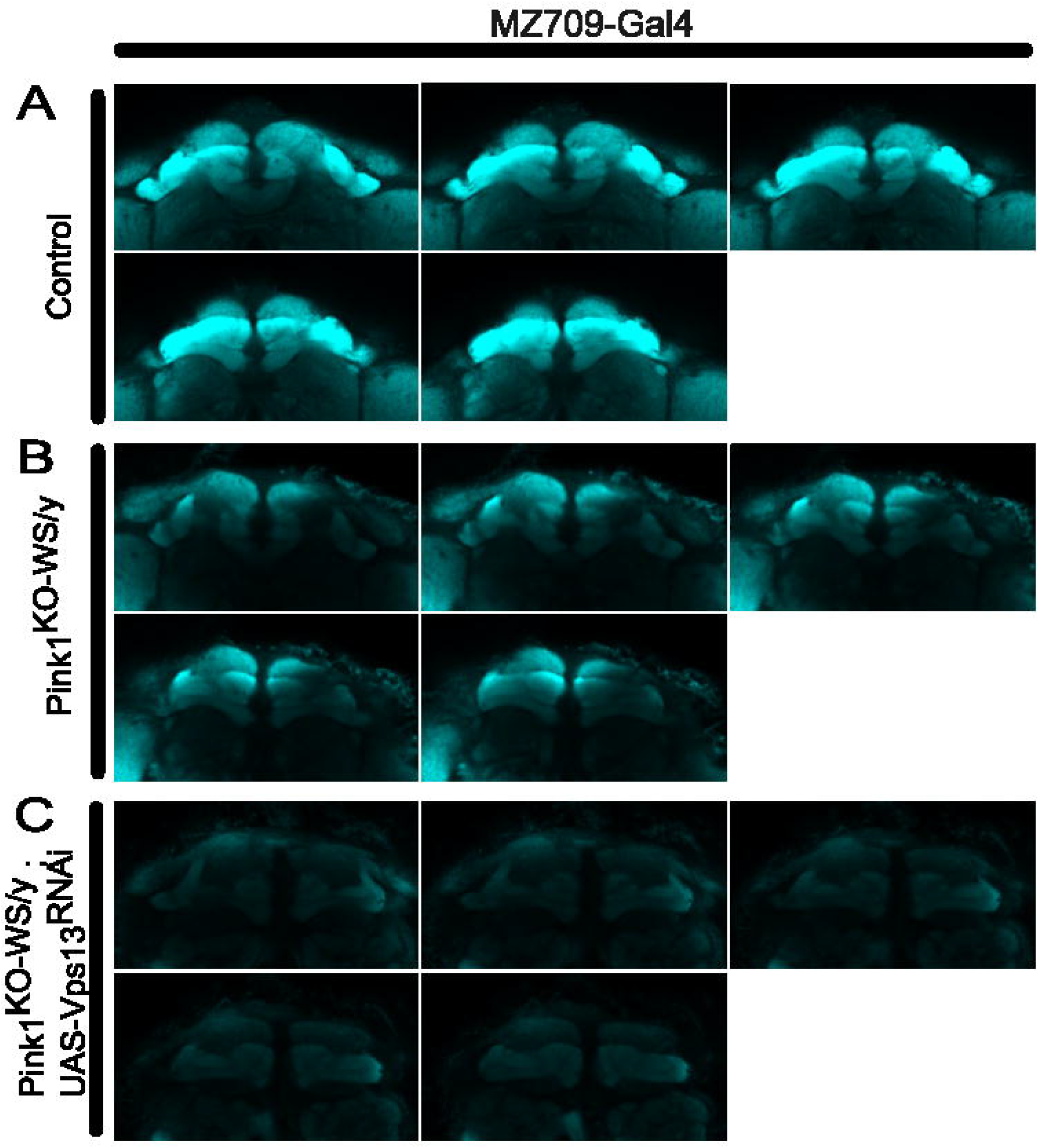
Examples of confocal Z-stacks of anti-DLG labeled fly brains (22 ± 2-day-old) of the indicated genotypes. Such images were used to delineate ROIs to quantify DAN innervation onto the MBs. (A) control (w1118/y;; MZ709-Gal4/+); (B) Pink1KO-WS/Y;; MZ709-Gal4/+; (C) Pink1KO-WS flies with *Vps13* downregulation in EG (Pink1KO-WS/Y;; *Vps13*RNAi/MZ709-Gal4).

## Inclusion and diversity

We support inclusive, diverse, and equitable conduct of research.

