## Appendix 1 for "PARKINSON’S DISEASE-ASSOCIATED *PINK1* LOSS DISRUPTS ENSHEATHING GLIA AND CAUSES DOPAMINERGIC NEURON SYNAPSE LOSS"

| | baseMean | log2FoldCl | lfcSE | stat | pvalue | padj |
|---|---|---|---|---|---|---|
| TTLL6A | 4750.559 | -11.3181 | 1.051882 | -10.7598 | 5.33E-27 | 4.28E-23 |
| l(3)07882 | 1189.898 | -10.1921 | 1.303389 | -7.81967 | 5.30E-15 | 2.13E-11 |
| CG10936 | 791.4433 | -10.6891 | 1.517311 | -7.04474 | 1.86E-12 | 4.97E-09 |
| CG15011 | 937.1086 | -9.15508 | 1.333765 | -6.86408 | 6.69E-12 | 1.34E-08 |
| CG7627 | 1298.357 | 7.298828 | 1.109944 | 6.575853 | 4.84E-11 | 7.77E-08 |
| CG10799 | 2865.974 | 8.638146 | 1.35321 | 6.383447 | 1.73E-10 | 1.86E-07 |
| Nos | 945.041 | 7.084464 | 1.111621 | 6.373091 | 1.85E-10 | 1.86E-07 |
| CG10646 | 546.699 | 10.6305 | 1.665295 | 6.383556 | 1.73E-10 | 1.86E-07 |
| Rhau | 911.6146 | 8.130053 | 1.30148 | 6.246776 | 4.19E-10 | 3.74E-07 |
| CG17660 | 2133.143 | -6.28132 | 1.015593 | -6.18488 | 6.21E-10 | 4.99E-07 |
| Proc | 2353.574 | -7.82446 | 1.278466 | -6.12019 | 9.35E-10 | 6.25E-07 |
| FASN2 | 738.2926 | -8.90389 | 1.451616 | -6.13377 | 8.58E-10 | 6.25E-07 |
| fiz | 953.792 | 7.856676 | 1.299757 | 6.044726 | 1.50E-09 | 9.24E-07 |
| Rpp25 | 822.1356 | 7.853897 | 1.304709 | 6.019656 | 1.75E-09 | 1.00E-06 |
| PIG-B | 2082.336 | -6.23226 | 1.047009 | -5.95244 | 2.64E-09 | 1.41E-06 |
| Lnpk | 1258.999 | 9.121406 | 1.536087 | 5.93808 | 2.88E-09 | 1.45E-06 |
| hfw | 2664.466 | -7.57176 | 1.278148 | -5.92401 | 3.14E-09 | 1.48E-06 |
| CG2126 | 1440.227 | 7.189026 | 1.224281 | 5.87204 | 4.30E-09 | 1.92E-06 |
| lncRNA:CR | 451.2195 | 11.34112 | 1.934365 | 5.862969 | 4.55E-09 | 1.92E-06 |
| spag | 1089.173 | -10.2702 | 1.755652 | -5.8498 | 4.92E-09 | 1.98E-06 |
| Prpk | 976.9641 | -11.1885 | 1.922361 | -5.82017 | 5.88E-09 | 2.25E-06 |
| CG31370 | 978.5736 | -8.8501 | 1.539494 | -5.74871 | 8.99E-09 | 3.28E-06 |
| CG17712 | 2483.84 | 5.98876 | 1.043279 | 5.740324 | 9.45E-09 | 3.30E-06 |
| achi | 416.894 | -11.9462 | 2.117389 | -5.64196 | 1.68E-08 | 5.62E-06 |
| Slc25A46a | 1861.854 | 6.855485 | 1.219843 | 5.619975 | 1.91E-08 | 6.13E-06 |
| CG34232 | 848.2188 | 10.83669 | 1.94161 | 5.581289 | 2.39E-08 | 7.37E-06 |
| NiPp1 | 656.2872 | 9.111064 | 1.640156 | 5.554997 | 2.78E-08 | 8.11E-06 |
| DUBAI | 1722.491 | 5.431373 | 0.978346 | 5.551585 | 2.83E-08 | 8.11E-06 |
| CG12773 | 1275.05 | 8.915078 | 1.614981 | 5.520236 | 3.39E-08 | 9.35E-06 |
| tadr | 1560.365 | 5.683654 | 1.032008 | 5.507375 | 3.64E-08 | 9.35E-06 |
| CG30357 | 397.499 | 11.84152 | 2.151088 | 5.504899 | 3.69E-08 | 9.35E-06 |
| Fic | 1217.805 | -8.27567 | 1.503747 | -5.50336 | 3.73E-08 | 9.35E-06 |
| Ccz1 | 475.0219 | 8.548615 | 1.564171 | 5.46527 | 4.62E-08 | 1.12E-05 |
| Atac1 | 426.3187 | 9.930944 | 1.827597 | 5.43388 | 5.51E-08 | 1.29E-05 |
| CG30424 | 1121.918 | -6.02376 | 1.109398 | -5.42975 | 5.64E-08 | 1.29E-05 |
| CG9004 | 617.8992 | 9.572376 | 1.766005 | 5.420355 | 5.95E-08 | 1.33E-05 |
| Rel | 935.8779 | 8.204279 | 1.516904 | 5.408568 | 6.35E-08 | 1.38E-05 |
| Ada2a | 2019.076 | -10.2619 | 1.901264 | -5.39742 | 6.76E-08 | 1.43E-05 |
| CG32532 | 1547.466 | -5.87735 | 1.091874 | -5.38281 | 7.33E-08 | 1.51E-05 |
| firl | 1009.197 | -8.02402 | 1.512882 | -5.3038 | 1.13E-07 | 2.28E-05 |
| Dark | 573.9756 | 8.110682 | 1.533639 | 5.288523 | 1.23E-07 | 2.41E-05 |
| CG17612 | 1020.991 | 12.03429 | 2.295015 | 5.243666 | 1.57E-07 | 3.01E-05 |
| pwn | 1323.361 | 7.869095 | 1.503215 | 5.234844 | 1.65E-07 | 3.08E-05 |
| Atg7 | 836.3293 | 7.916463 | 1.517504 | 5.216767 | 1.82E-07 | 3.32E-05 |
| Pld | 1231.724 | -5.67404 | 1.089622 | -5.20735 | 1.92E-07 | 3.42E-05 |
| CG6867 | 1366.312 | 7.862875 | 1.513154 | 5.196348 | 2.03E-07 | 3.55E-05 |
| Wdr33 | 1005.735 | 7.185195 | 1.390908 | 5.165829 | 2.39E-07 | 4.09E-05 |
| M6 | 951.0725 | -7.28518 | 1.413304 | -5.15471 | 2.54E-07 | 4.25E-05 |
| GMF | 879.7349 | 7.398752 | 1.445361 | 5.118966 | 3.07E-07 | 4.93E-05 |

| | | | | | | |
|---|---|---|---|---|---|---|
| Camp | 1876.748 | 6.519438 | 1.272894 | 5.121747 | 3.03E-07 | 4.93E-05 |
| Cyfip | 1000.608 | 7.178368 | 1.406078 | 5.105241 | 3.30E-07 | 5.20E-05 |
| Tpc1 | 883.0577 | 9.192462 | 1.802175 | 5.100761 | 3.38E-07 | 5.22E-05 |
| mira | 1037.83 | -7.15616 | 1.407081 | -5.08582 | 3.66E-07 | 5.54E-05 |
| Cyp6w1 | 1435.289 | 6.583727 | 1.297643 | 5.073606 | 3.90E-07 | 5.80E-05 |
| CG34347 | 1347.73 | -7.85718 | 1.552718 | -5.06028 | 4.19E-07 | 6.11E-05 |
| Rh4 | 300.8536 | -11.9419 | 2.363857 | -5.05188 | 4.37E-07 | 6.27E-05 |
| dnt | 855.0223 | 7.602648 | 1.512799 | 5.02555 | 5.02E-07 | 6.95E-05 |
| Usp20-33 | 1164.981 | -7.6412 | 1.520467 | -5.02556 | 5.02E-07 | 6.95E-05 |
| CG6428 | 761.7536 | 6.031842 | 1.208578 | 4.990858 | 6.01E-07 | 8.18E-05 |
| GstD2 | 639.3842 | -11.612 | 2.338095 | -4.96643 | 6.82E-07 | 8.99E-05 |
| lncRNA:CR | 759.4647 | -8.82499 | 1.777071 | -4.96603 | 6.83E-07 | 8.99E-05 |
| CG9547 | 548.1134 | -10.467 | 2.125966 | -4.92339 | 8.51E-07 | 0.00011 |
| pad | 763.5107 | 10.42587 | 2.119736 | 4.918474 | 8.72E-07 | 0.000111 |
| CG4603 | 753.7642 | 6.627124 | 1.348244 | 4.915374 | 8.86E-07 | 0.000111 |
| CG6638 | 473.3985 | 11.66592 | 2.378263 | 4.905225 | 9.33E-07 | 0.000115 |
| CG7601 | 498.7434 | -9.68964 | 1.976787 | -4.90171 | 9.50E-07 | 0.000116 |
| asRNA:CR4 | 363.8035 | 10.10342 | 2.072028 | 4.8761 | 1.08E-06 | 0.00013 |
| CG4676 | 1262.173 | 8.907078 | 1.840739 | 4.838859 | 1.31E-06 | 0.000154 |
| dnk | 1080.378 | -7.87587 | 1.628822 | -4.83531 | 1.33E-06 | 0.000155 |
| mre11 | 532.8426 | -8.54291 | 1.771117 | -4.82346 | 1.41E-06 | 0.000162 |
| ppk12 | 588.873 | -11.6578 | 2.426559 | -4.80426 | 1.55E-06 | 0.000176 |
| alpha-Est8 | 1375.689 | -6.02904 | 1.262605 | -4.77508 | 1.80E-06 | 0.0002 |
| Csl4 | 715.1897 | 7.912328 | 1.661571 | 4.761956 | 1.92E-06 | 0.000211 |
| Mau2 | 534.9165 | 8.877389 | 1.870235 | 4.746671 | 2.07E-06 | 0.000218 |
| Dhap-at | 926.5075 | 6.023835 | 1.26839 | 4.7492 | 2.04E-06 | 0.000218 |
| CG4594 | 305.5028 | -10.6636 | 2.245392 | -4.74908 | 2.04E-06 | 0.000218 |
| CG5704 | 317.6176 | 9.988691 | 2.112231 | 4.728976 | 2.26E-06 | 0.000235 |
| nst | 484.3521 | -11.1984 | 2.374566 | -4.71599 | 2.41E-06 | 0.000248 |
| CG10171 | 748.0433 | 8.011489 | 1.702111 | 4.706797 | 2.52E-06 | 0.000256 |
| CG15436 | 515.9871 | -11.3495 | 2.4133 | -4.70289 | 2.57E-06 | 0.000257 |
| CG4627 | 1152.258 | 7.021276 | 1.505195 | 4.664695 | 3.09E-06 | 0.000306 |
| CG30491 | 1428.071 | 6.5103 | 1.399045 | 4.653388 | 3.27E-06 | 0.00032 |
| adp | 334.5566 | 9.035789 | 1.946097 | 4.643032 | 3.43E-06 | 0.000332 |
| CG13609 | 260.4131 | -9.63685 | 2.082569 | -4.62739 | 3.70E-06 | 0.000342 |
| asRNA:CR4 | 631.3889 | -9.24778 | 1.998025 | -4.62846 | 3.68E-06 | 0.000342 |
| CG6793 | 430.3684 | 10.53007 | 2.272193 | 4.634319 | 3.58E-06 | 0.000342 |
| Blos2 | 438.3986 | 7.294316 | 1.57518 | 4.630783 | 3.64E-06 | 0.000342 |
| Pyroxd1 | 1170.982 | -6.8667 | 1.486109 | -4.62059 | 3.83E-06 | 0.000345 |
| Pdxk | 1438.566 | -6.33072 | 1.369757 | -4.62178 | 3.80E-06 | 0.000345 |
| CG16789 | 607.5918 | 7.164701 | 1.552375 | 4.615317 | 3.92E-06 | 0.00035 |
| CG6830 | 539.0547 | -10.0033 | 2.17103 | -4.60763 | 4.07E-06 | 0.000359 |
| CG9246 | 369.4685 | -8.24787 | 1.792973 | -4.60011 | 4.22E-06 | 0.000368 |
| Est-Q | 469.7577 | -7.83016 | 1.707343 | -4.58617 | 4.51E-06 | 0.00039 |
| CG12448 | 399.8078 | -9.48382 | 2.070497 | -4.58046 | 4.64E-06 | 0.000396 |
| Syx18 | 616.1109 | -8.35214 | 1.827677 | -4.56981 | 4.88E-06 | 0.000412 |
| CG13151 | 432.9034 | -11.6207 | 2.550673 | -4.55594 | 5.22E-06 | 0.000436 |
| Ror | 1022.192 | 6.577376 | 1.444827 | 4.552361 | 5.30E-06 | 0.000439 |
| CG6550 | 585.0607 | -7.65072 | 1.684011 | -4.54315 | 5.54E-06 | 0.000454 |
| pelo | 931.3385 | -6.94618 | 1.531768 | -4.53475 | 5.77E-06 | 0.000468 |

| | | | | | | |
|---|---|---|---|---|---|---|
| PIG-A | 493.3573 | 9.592711 | 2.116743 | 4.531826 | 5.85E-06 | 0.000469 |
| alpha-PheI | 775.3718 | -6.84436 | 1.513692 | -4.52163 | 6.14E-06 | 0.000488 |
| coil | 366.9633 | -10.1972 | 2.256625 | -4.5188 | 6.22E-06 | 0.000489 |
| Cyp28d2 | 695.7862 | 9.641996 | 2.14127 | 4.502934 | 6.70E-06 | 0.000522 |
| phr | 430.8601 | -7.81265 | 1.737268 | -4.49709 | 6.89E-06 | 0.000532 |
| HisCl1 | 1214.398 | -6.87144 | 1.530979 | -4.48826 | 7.18E-06 | 0.000544 |
| CG33978 | 333.9723 | -11.3906 | 2.537149 | -4.48954 | 7.14E-06 | 0.000544 |
| CG8379 | 706.0137 | 8.454017 | 1.88676 | 4.480706 | 7.44E-06 | 0.000554 |
| Dip-B | 354.8804 | 10.39864 | 2.323962 | 4.474529 | 7.66E-06 | 0.000554 |
| CG10321 | 639.2847 | -8.6795 | 1.940645 | -4.47248 | 7.73E-06 | 0.000554 |
| CG7206 | 2811.474 | -3.3786 | 0.754762 | -4.47638 | 7.59E-06 | 0.000554 |
| Rpc53 | 790.2336 | 7.802903 | 1.744342 | 4.473264 | 7.70E-06 | 0.000554 |
| MTF-1 | 687.6926 | -7.04686 | 1.572933 | -4.48008 | 7.46E-06 | 0.000554 |
| CG34357 | 1161.319 | 4.89049 | 1.094028 | 4.470168 | 7.82E-06 | 0.000555 |
| CG15107 | 745.3173 | -6.63378 | 1.485867 | -4.46458 | 8.02E-06 | 0.000565 |
| CG2614 | 641.2481 | -7.44849 | 1.670142 | -4.45979 | 8.20E-06 | 0.000569 |
| CG9008 | 645.4038 | -8.10254 | 1.816992 | -4.45931 | 8.22E-06 | 0.000569 |
| CG6144 | 1510.144 | -5.55041 | 1.246265 | -4.45363 | 8.44E-06 | 0.000579 |
| CG10623 | 751.1332 | 6.476712 | 1.456945 | 4.445406 | 8.77E-06 | 0.000597 |
| CG8298 | 1073.967 | -6.42129 | 1.447333 | -4.43664 | 9.14E-06 | 0.000616 |
| obst-A | 518.7521 | 9.031956 | 2.042299 | 4.422446 | 9.76E-06 | 0.000653 |
| CG14722 | 586.0011 | -7.42685 | 1.681661 | -4.41638 | 1.00E-05 | 0.000666 |
| CG1602 | 604.9766 | 10.05862 | 2.285526 | 4.401008 | 1.08E-05 | 0.000709 |
| Rcd2 | 1008.269 | 6.907473 | 1.576127 | 4.382561 | 1.17E-05 | 0.000765 |
| betaTub60 | 2092.102 | -5.43934 | 1.242521 | -4.37767 | 1.20E-05 | 0.000776 |
| ru | 641.2174 | -9.49179 | 2.1719 | -4.37027 | 1.24E-05 | 0.000797 |
| CG5292 | 580.8667 | -6.52555 | 1.497765 | -4.35686 | 1.32E-05 | 0.00084 |
| dx | 661.8949 | -9.69874 | 2.231913 | -4.34548 | 1.39E-05 | 0.000875 |
| TfIIEalpha | 1301.796 | 6.650406 | 1.530728 | 4.344604 | 1.40E-05 | 0.000875 |
| CG5641 | 2018.42 | -5.22316 | 1.203806 | -4.33887 | 1.43E-05 | 0.000891 |
| RluA-2 | 622.5085 | -7.71479 | 1.780042 | -4.33405 | 1.46E-05 | 0.000904 |
| CG12219 | 674.1425 | -8.00116 | 1.848699 | -4.32799 | 1.50E-05 | 0.000915 |
| cos | 1072.298 | -6.63968 | 1.533869 | -4.32872 | 1.50E-05 | 0.000915 |
| fu | 499.4366 | 7.900491 | 1.829059 | 4.31943 | 1.56E-05 | 0.000944 |
| ymp | 335.7695 | -10.8397 | 2.51436 | -4.3111 | 1.62E-05 | 0.000973 |
| CG5555 | 1051.403 | 5.941167 | 1.381806 | 4.299565 | 1.71E-05 | 0.001017 |
| CG4951 | 606.0654 | -7.95497 | 1.857217 | -4.28328 | 1.84E-05 | 0.001079 |
| rod | 491.4583 | -9.14753 | 2.135462 | -4.28363 | 1.84E-05 | 0.001079 |
| otp | 723.9208 | 6.966963 | 1.629643 | 4.275148 | 1.91E-05 | 0.001111 |
| CG3552 | 357.4748 | 7.997576 | 1.871768 | 4.272738 | 1.93E-05 | 0.001115 |
| CG3407 | 574.2291 | 6.368619 | 1.49309 | 4.265396 | 2.00E-05 | 0.001144 |
| CG13097 | 754.3208 | -7.46558 | 1.757137 | -4.24872 | 2.15E-05 | 0.001224 |
| rig | 369.5297 | 8.56305 | 2.018031 | 4.243271 | 2.20E-05 | 0.001245 |
| CG17802 | 735.7165 | 7.532725 | 1.777363 | 4.238146 | 2.25E-05 | 0.001256 |
| CG5880 | 637.0677 | -7.02653 | 1.657498 | -4.23924 | 2.24E-05 | 0.001256 |
| Camta | 1862.904 | 3.638238 | 0.862456 | 4.218462 | 2.46E-05 | 0.001362 |
| CCHa2 | 523.4574 | -8.118 | 1.925747 | -4.21551 | 2.49E-05 | 0.00137 |
| asRNA:CR4 | 243.6834 | -10.7339 | 2.54745 | -4.21357 | 2.51E-05 | 0.001372 |
| CG3703 | 1143.821 | 6.486977 | 1.540411 | 4.211198 | 2.54E-05 | 0.001378 |
| Smyd4-4 | 1077.187 | 5.559314 | 1.32504 | 4.195581 | 2.72E-05 | 0.001466 |

| | | | | | | |
|---|---|---|---|---|---|---|
| Alg10 | 907.2355 | -7.90167 | 1.888532 | -4.18403 | 2.86E-05 | 0.001532 |
| Alg3 | 343.8811 | -7.80808 | 1.86933 | -4.17694 | 2.95E-05 | 0.00157 |
| Adgf-C | 1431.711 | -6.2167 | 1.48995 | -4.17242 | 3.01E-05 | 0.001571 |
| CG11779 | 520.3206 | 7.336643 | 1.757379 | 4.174764 | 2.98E-05 | 0.001571 |
| CG7208 | 553.4062 | 8.881809 | 2.128733 | 4.172346 | 3.01E-05 | 0.001571 |
| IntS1 | 536.0844 | -7.13695 | 1.711473 | -4.17006 | 3.05E-05 | 0.001577 |
| Rpp30 | 434.4003 | -11.3755 | 2.731802 | -4.16409 | 3.13E-05 | 0.001604 |
| sl | 313.5023 | -7.13075 | 1.713265 | -4.16208 | 3.15E-05 | 0.001604 |
| ssp6 | 1191.295 | -5.51122 | 1.324253 | -4.16176 | 3.16E-05 | 0.001604 |
| CG17026 | 1582.285 | -5.55945 | 1.336467 | -4.15981 | 3.19E-05 | 0.001608 |
| PSR | 450.0398 | -7.96811 | 1.918847 | -4.15255 | 3.29E-05 | 0.001649 |
| Sfmbt | 889.3905 | 6.560389 | 1.582526 | 4.145517 | 3.39E-05 | 0.00168 |
| CG1941 | 358.1842 | 10.41666 | 2.512683 | 4.145632 | 3.39E-05 | 0.00168 |
| Mesh1 | 201.0673 | -11.3017 | 2.729385 | -4.14075 | 3.46E-05 | 0.001696 |
| CG42327 | 2142.227 | -3.69198 | 0.891673 | -4.14051 | 3.47E-05 | 0.001696 |
| Dpit47 | 382.5126 | -10.001 | 2.41638 | -4.13885 | 3.49E-05 | 0.001698 |
| lncRNA:CR | 230.0664 | 9.197144 | 2.228907 | 4.126302 | 3.69E-05 | 0.001782 |
| trbl | 220.8146 | -10.992 | 2.665278 | -4.12415 | 3.72E-05 | 0.001788 |
| Dbp21E2 | 470.2418 | 9.106004 | 2.212632 | 4.115462 | 3.86E-05 | 0.001846 |
| CG32344 | 479.2301 | 8.142051 | 1.984282 | 4.103274 | 4.07E-05 | 0.001935 |
| Cyp12b2 | 3795.63 | -2.76883 | 0.675607 | -4.09829 | 4.16E-05 | 0.001965 |
| CG17726 | 572.5821 | 6.852312 | 1.674336 | 4.092554 | 4.27E-05 | 0.002003 |
| CG4935 | 885.0364 | 7.371235 | 1.808744 | 4.075334 | 4.59E-05 | 0.002132 |
| CG8152 | 438.2612 | -9.28722 | 2.278704 | -4.07566 | 4.59E-05 | 0.002132 |
| CG2794 | 311.3938 | -9.83769 | 2.417395 | -4.06954 | 4.71E-05 | 0.002148 |
| crok | 342.0966 | -9.64099 | 2.368509 | -4.07049 | 4.69E-05 | 0.002148 |
| tex | 325.5767 | 8.753308 | 2.150543 | 4.070278 | 4.70E-05 | 0.002148 |
| CG31457 | 490.7809 | 8.547122 | 2.102339 | 4.065529 | 4.79E-05 | 0.002173 |
| CysRS-m | 346.5391 | 7.146401 | 1.761913 | 4.056047 | 4.99E-05 | 0.00225 |
| CG12288 | 503.5113 | -7.00612 | 1.728305 | -4.05375 | 5.04E-05 | 0.00226 |
| spirit | 234.8125 | 9.373786 | 2.313414 | 4.051928 | 5.08E-05 | 0.002265 |
| TM9SF3 | 2296.193 | 4.931159 | 1.217635 | 4.049784 | 5.13E-05 | 0.002273 |
| ssp | 370.5257 | -8.4051 | 2.07813 | -4.04455 | 5.24E-05 | 0.002312 |
| CG14589 | 231.2725 | -10.685 | 2.650055 | -4.03198 | 5.53E-05 | 0.002412 |
| CG30291 | 928.9416 | -7.99955 | 1.984611 | -4.03079 | 5.56E-05 | 0.002412 |
| CstF64 | 621.4064 | -8.5998 | 2.132895 | -4.03199 | 5.53E-05 | 0.002412 |
| CG10949 | 447.0743 | -6.89439 | 1.711005 | -4.02944 | 5.59E-05 | 0.002413 |
| Rae1 | 1056.038 | -5.60417 | 1.391323 | -4.02794 | 5.63E-05 | 0.002415 |
| jnj | 362.518 | 8.214528 | 2.040043 | 4.026644 | 5.66E-05 | 0.002415 |
| l(2)10685 | 493.5839 | 5.847429 | 1.452879 | 4.024719 | 5.70E-05 | 0.002422 |
| CG13773 | 324.8891 | -7.90106 | 1.968229 | -4.0143 | 5.96E-05 | 0.002505 |
| CG30466 | 227.1361 | 7.940211 | 1.977519 | 4.01524 | 5.94E-05 | 0.002505 |
| lncRNA:CR | 272.0478 | 8.888315 | 2.216865 | 4.009407 | 6.09E-05 | 0.002505 |
| RAF2 | 641.6294 | -8.58163 | 2.139269 | -4.01148 | 6.03E-05 | 0.002505 |
| CG7011 | 190.8441 | -11.2011 | 2.792507 | -4.01113 | 6.04E-05 | 0.002505 |
| CG17593 | 2006.614 | -4.19264 | 1.045596 | -4.00981 | 6.08E-05 | 0.002505 |
| cv-d | 979.4725 | 5.91847 | 1.476721 | 4.007846 | 6.13E-05 | 0.002509 |
| Aos1 | 787.3889 | 6.320943 | 1.578301 | 4.004903 | 6.20E-05 | 0.002528 |
| CG17359 | 281.0305 | 11.20301 | 2.799852 | 4.001285 | 6.30E-05 | 0.002554 |
| CG11334 | 390.1485 | -11.5608 | 2.897887 | -3.98938 | 6.62E-05 | 0.002672 |

| | | | | | | |
|---|---|---|---|---|---|---|
| lncRNA:CR | 207.9927 | 10.75406 | 2.697993 | 3.985948 | 6.72E-05 | 0.002697 |
| Idgf1 | 606.467 | -7.60776 | 1.909333 | -3.98451 | 6.76E-05 | 0.0027 |
| CSN7 | 388.4597 | 6.301425 | 1.582063 | 3.983042 | 6.80E-05 | 0.002703 |
| CG11436 | 942.9732 | 7.477989 | 1.879286 | 3.979166 | 6.92E-05 | 0.002732 |
| Cyp28c1 | 1835.342 | -4.4683 | 1.123189 | -3.97822 | 6.94E-05 | 0.002732 |
| CG12267 | 339.2301 | -8.31067 | 2.092897 | -3.97089 | 7.16E-05 | 0.002776 |
| ProRS-m | 237.4179 | -10.3161 | 2.597517 | -3.97153 | 7.14E-05 | 0.002776 |
| CG9609 | 1001.69 | 6.020454 | 1.515515 | 3.972547 | 7.11E-05 | 0.002776 |
| Rich | 499.5593 | -8.00467 | 2.01887 | -3.96493 | 7.34E-05 | 0.002819 |
| Upf2 | 335.5828 | -8.97987 | 2.264212 | -3.966 | 7.31E-05 | 0.002819 |
| CG6347 | 310.4918 | -8.9772 | 2.268101 | -3.95802 | 7.56E-05 | 0.002888 |
| CG15118 | 1714.726 | 3.908311 | 0.98828 | 3.954661 | 7.66E-05 | 0.002915 |
| Plzf | 625.5818 | 7.397519 | 1.872025 | 3.951613 | 7.76E-05 | 0.002939 |
| Ugt302C1 | 218.0225 | -9.5825 | 2.426706 | -3.94877 | 7.86E-05 | 0.002947 |
| Irbp | 202.0334 | -10.8426 | 2.745879 | -3.94867 | 7.86E-05 | 0.002947 |
| Drep3 | 555.8074 | 7.857601 | 1.991845 | 3.944886 | 7.98E-05 | 0.00298 |
| thoc5 | 208.8412 | -8.97273 | 2.275251 | -3.94362 | 8.03E-05 | 0.002982 |
| CG12814 | 381.5944 | -7.88869 | 2.003464 | -3.93752 | 8.23E-05 | 0.003045 |
| Fam92 | 536.6875 | -5.95175 | 1.513715 | -3.93188 | 8.43E-05 | 0.003073 |
| CG8531 | 1277.986 | -6.6103 | 1.68119 | -3.93192 | 8.43E-05 | 0.003073 |
| U2A | 738.656 | -6.83666 | 1.739676 | -3.92985 | 8.50E-05 | 0.003073 |
| cana | 296.5751 | -7.14343 | 1.816001 | -3.93361 | 8.37E-05 | 0.003073 |
| l(2)37Cg | 473.8694 | 6.68439 | 1.70059 | 3.930629 | 8.47E-05 | 0.003073 |
| Cerk | 2206.886 | -2.90162 | 0.738891 | -3.92699 | 8.60E-05 | 0.003096 |
| TBC1d7 | 303.1996 | -7.81456 | 1.991507 | -3.92394 | 8.71E-05 | 0.003121 |
| CG12224 | 490.9956 | -7.04642 | 1.797329 | -3.9205 | 8.84E-05 | 0.003152 |
| CG18094 | 1190.452 | 5.379339 | 1.372823 | 3.91845 | 8.91E-05 | 0.003165 |
| CG18622 | 353.4835 | -8.32758 | 2.129344 | -3.91087 | 9.20E-05 | 0.003252 |
| Mics1 | 390.3873 | 9.343207 | 2.391251 | 3.907246 | 9.34E-05 | 0.003286 |
| CG15743 | 613.4656 | -7.39825 | 1.895894 | -3.90225 | 9.53E-05 | 0.00334 |
| Hira | 381.2118 | -8.18672 | 2.100676 | -3.89718 | 9.73E-05 | 0.003396 |
| Sin | 385.0068 | -7.8396 | 2.014992 | -3.89064 | 1.00E-04 | 0.003455 |
| DIP-zeta | 437.2753 | -11.0154 | 2.83224 | -3.8893 | 0.000101 | 0.003455 |
| CG10602 | 802.7276 | -6.13685 | 1.578073 | -3.88883 | 0.000101 | 0.003455 |
| CG17027 | 682.8516 | 6.153834 | 1.581975 | 3.88997 | 0.0001 | 0.003455 |
| MED8 | 503.0481 | -7.6591 | 1.973148 | -3.88167 | 0.000104 | 0.003528 |
| LeuRS | 854.3417 | -6.01486 | 1.549283 | -3.88235 | 0.000103 | 0.003528 |
| CG2972 | 342.6367 | 8.260128 | 2.131228 | 3.875759 | 0.000106 | 0.0036 |
| Cyp6a17 | 4346.845 | -2.49194 | 0.644138 | -3.86864 | 0.000109 | 0.003675 |
| CG42393 | 266.6757 | -8.92492 | 2.306571 | -3.86934 | 0.000109 | 0.003675 |
| CG7130 | 403.2081 | 7.993299 | 2.066729 | 3.867608 | 0.00011 | 0.003676 |
| TTLL15 | 596.5043 | -8.45342 | 2.188397 | -3.86284 | 0.000112 | 0.003732 |
| CG15084 | 396.396 | -9.11404 | 2.359995 | -3.86189 | 0.000113 | 0.003732 |
| Mtr4 | 413.491 | 6.240751 | 1.617775 | 3.857615 | 0.000114 | 0.003769 |
| CG8128 | 320.5576 | -8.61998 | 2.234621 | -3.85747 | 0.000115 | 0.003769 |
| SiaT | 450.8944 | 8.684454 | 2.252927 | 3.854742 | 0.000116 | 0.003795 |
| WDR79 | 782.8013 | 7.567826 | 1.964517 | 3.852257 | 0.000117 | 0.003818 |
| Uba4 | 532.9401 | -7.37356 | 1.916329 | -3.84775 | 0.000119 | 0.003873 |
| lncRNA:CR | 243.5297 | -10.4376 | 2.714122 | -3.84567 | 0.00012 | 0.003875 |
| Ing3 | 284.602 | -7.62149 | 1.981818 | -3.8457 | 0.00012 | 0.003875 |

| | | | | | | |
|---|---|---|---|---|---|---|
| Dhod | 707.3627 | -5.69567 | 1.484058 | -3.8379 | 0.000124 | 0.003953 |
| Rpt5 | 1092.509 | -5.91004 | 1.539922 | -3.83788 | 0.000124 | 0.003953 |
| Hyccin | 696.6034 | 6.94532 | 1.809002 | 3.83931 | 0.000123 | 0.003953 |
| NANS | 1057.412 | -4.33274 | 1.129265 | -3.83678 | 0.000125 | 0.003955 |
| chm | 789.445 | -5.46329 | 1.425017 | -3.83384 | 0.000126 | 0.003986 |
| Art3 | 372.1751 | 11.12566 | 2.903438 | 3.831891 | 0.000127 | 0.004002 |
| Vps33B | 479.694 | -9.79747 | 2.562587 | -3.82327 | 0.000132 | 0.004129 |
| CCHa1 | 1732.036 | 6.221105 | 1.627647 | 3.822148 | 0.000132 | 0.004132 |
| CG30010 | 543.8857 | 6.767821 | 1.773099 | 3.816945 | 0.000135 | 0.004203 |
| CG5498 | 948.1067 | 5.525568 | 1.448505 | 3.814671 | 0.000136 | 0.004226 |
| Nrd1 | 483.5141 | -5.34265 | 1.404843 | -3.80302 | 0.000143 | 0.004413 |
| KaiR1D | 1733.27 | 4.229191 | 1.112863 | 3.800281 | 0.000145 | 0.004445 |
| CG13016 | 355.2934 | 7.85528 | 2.06783 | 3.798805 | 0.000145 | 0.004454 |
| Uros1 | 372.3157 | -7.31633 | 1.92866 | -3.79348 | 0.000149 | 0.004533 |
| jumu | 529.7174 | -6.69663 | 1.765959 | -3.79207 | 0.000149 | 0.004542 |
| CG17141 | 352.7418 | 8.885027 | 2.345619 | 3.787924 | 0.000152 | 0.004587 |
| CG7191 | 1381.679 | -5.47969 | 1.446691 | -3.78774 | 0.000152 | 0.004587 |
| sip1 | 666.6298 | 7.233956 | 1.911704 | 3.784036 | 0.000154 | 0.004638 |
| CG32191 | 464.2185 | -5.8942 | 1.558824 | -3.78118 | 0.000156 | 0.004674 |
| Taf7 | 396.902 | 7.411811 | 1.962032 | 3.77762 | 0.000158 | 0.004691 |
| trh | 292.0806 | -8.87308 | 2.348903 | -3.77754 | 0.000158 | 0.004691 |
| tut | 1189.383 | 4.376568 | 1.158074 | 3.779179 | 0.000157 | 0.004691 |
| NC2beta | 740.7553 | 5.594179 | 1.481722 | 3.775458 | 0.00016 | 0.004713 |
| Keap1 | 541.9236 | 7.560549 | 2.005221 | 3.770431 | 0.000163 | 0.004791 |
| CG13408 | 605.9705 | 8.310606 | 2.206864 | 3.765799 | 0.000166 | 0.004863 |
| CG15387 | 230.1346 | -10.7056 | 2.848067 | -3.75892 | 0.000171 | 0.004981 |
| CG9706 | 1015.245 | -4.98589 | 1.328348 | -3.75345 | 0.000174 | 0.005072 |
| CG11168 | 360.3669 | -9.8921 | 2.636829 | -3.75151 | 0.000176 | 0.005079 |
| CG16896 | 399.7606 | -6.29843 | 1.679009 | -3.75128 | 0.000176 | 0.005079 |
| CG42553 | 283.3301 | -9.15774 | 2.4429 | -3.74872 | 0.000178 | 0.005113 |
| UbcE2M | 1278.138 | -3.28822 | 0.877515 | -3.74719 | 0.000179 | 0.005126 |
| CG2681 | 297.3664 | -10.35 | 2.767218 | -3.74023 | 0.000184 | 0.005251 |
| Rev1 | 1074.932 | 7.303795 | 1.953312 | 3.739185 | 0.000185 | 0.005254 |
| CG12129 | 431.8526 | -8.03781 | 2.150822 | -3.73709 | 0.000186 | 0.00528 |
| CG33051 | 382.5251 | -7.11048 | 1.906592 | -3.72942 | 0.000192 | 0.005424 |
| CG8067 | 500.172 | -8.79919 | 2.364193 | -3.72186 | 0.000198 | 0.005561 |
| IleRS-m | 562.8845 | -8.31867 | 2.235385 | -3.72136 | 0.000198 | 0.005561 |
| CG9626 | 173.9439 | -10.7308 | 2.891439 | -3.71123 | 0.000206 | 0.005754 |
| Gcn5 | 1529.812 | 3.272344 | 0.8818 | 3.710982 | 0.000206 | 0.005754 |
| CG18449 | 214.4182 | -9.78152 | 2.637669 | -3.7084 | 0.000209 | 0.005793 |
| ntc | 294.2206 | 10.75939 | 2.908566 | 3.699209 | 0.000216 | 0.005965 |
| Dph1 | 1316.282 | 5.742012 | 1.551933 | 3.69991 | 0.000216 | 0.005965 |
| CG9302 | 1246.95 | -5.41874 | 1.465815 | -3.69674 | 0.000218 | 0.006003 |
| CG6345 | 226.0583 | -10.6879 | 2.896854 | -3.68948 | 0.000225 | 0.006155 |
| CG13876 | 259.6387 | -8.47047 | 2.297426 | -3.68694 | 0.000227 | 0.006196 |
| CG14906 | 271.7683 | 7.37445 | 2.00181 | 3.683891 | 0.00023 | 0.006249 |
| CG43091 | 299.2089 | -8.55033 | 2.324508 | -3.67834 | 0.000235 | 0.006365 |
| CG34183 | 357.9084 | -7.34043 | 1.998076 | -3.67375 | 0.000239 | 0.006459 |
| Psf2 | 226.8873 | 10.37649 | 2.829876 | 3.666765 | 0.000246 | 0.006616 |
| CG31633 | 535.2604 | 7.131974 | 1.946534 | 3.663935 | 0.000248 | 0.006667 |

| | | | | | | |
|---|---|---|---|---|---|---|
| CG6910 | 507.0548 | 8.230874 | 2.248401 | 3.660767 | 0.000251 | 0.006727 |
| CG2875 | 862.5305 | 5.048583 | 1.38635 | 3.641636 | 0.000271 | 0.007224 |
| senju | 390.4874 | -8.65438 | 2.379083 | -3.63769 | 0.000275 | 0.007263 |
| DIP-eta | 372.7961 | 6.311828 | 1.734796 | 3.638369 | 0.000274 | 0.007263 |
| CG8728 | 1945.347 | -3.65005 | 1.003173 | -3.6385 | 0.000274 | 0.007263 |
| ko | 419.3295 | 6.308475 | 1.743579 | 3.618119 | 0.000297 | 0.007777 |
| CG14535 | 1233.562 | 3.410078 | 0.943103 | 3.615805 | 0.000299 | 0.007777 |
| Reg-5 | 383.1895 | -7.10121 | 1.963721 | -3.6162 | 0.000299 | 0.007777 |
| Usp39 | 436.3149 | -8.60387 | 2.378556 | -3.61726 | 0.000298 | 0.007777 |
| skl | 314.2048 | 9.030236 | 2.495126 | 3.61915 | 0.000296 | 0.007777 |
| Strn-Mlck | 763.9584 | -4.93127 | 1.369471 | -3.60086 | 0.000317 | 0.008211 |
| Nup54 | 315.1 | 10.09361 | 2.807216 | 3.595595 | 0.000324 | 0.008352 |
| MFS16 | 359.6643 | 9.821248 | 2.734476 | 3.591638 | 0.000329 | 0.008453 |
| mRRF1 | 264.677 | -10.549 | 2.937842 | -3.59074 | 0.00033 | 0.008455 |
| Akt1 | 947.1016 | -5.10935 | 1.425798 | -3.5835 | 0.000339 | 0.008666 |
| LKRSDH | 192.7737 | -10.6853 | 2.984686 | -3.58003 | 0.000344 | 0.008753 |
| Taf5 | 833.4791 | -6.07965 | 1.700923 | -3.57432 | 0.000351 | 0.008918 |
| CG5114 | 1311.668 | -5.0184 | 1.40511 | -3.57154 | 0.000355 | 0.008985 |
| aph-1 | 623.8694 | 6.354992 | 1.780629 | 3.568958 | 0.000358 | 0.009046 |
| Kr-h2 | 4153.233 | 2.842019 | 0.798998 | 3.556978 | 0.000375 | 0.009439 |
| Nct | 822.04 | 5.820405 | 1.637209 | 3.555077 | 0.000378 | 0.009477 |
| Loxl1 | 205.3145 | 10.15359 | 2.859638 | 3.550656 | 0.000384 | 0.009608 |
| Cp110 | 253.5559 | 10.38475 | 2.928775 | 3.545767 | 0.000391 | 0.009758 |
| r2d2 | 626.9538 | -4.19037 | 1.182723 | -3.54298 | 0.000396 | 0.009831 |
| lncRNA:CR | 244.8413 | -8.59186 | 2.427854 | -3.53887 | 0.000402 | 0.009954 |
| Uhg8 | 257.8805 | -8.85452 | 2.504133 | -3.53596 | 0.000406 | 0.010034 |
| CG31251 | 299.1629 | -8.71856 | 2.467166 | -3.53384 | 0.00041 | 0.010053 |
| CG4882 | 1413.123 | -3.98982 | 1.128984 | -3.53399 | 0.000409 | 0.010053 |
| spn-A | 1079.667 | 3.7897 | 1.072856 | 3.532348 | 0.000412 | 0.010079 |
| CG7131 | 477.8142 | 8.588082 | 2.433788 | 3.528689 | 0.000418 | 0.010152 |
| CG7747 | 282.8831 | 8.147512 | 2.309032 | 3.52854 | 0.000418 | 0.010152 |
| por | 366.7378 | -7.72816 | 2.190506 | -3.52803 | 0.000419 | 0.010152 |
| Pez | 499.477 | -5.4802 | 1.554163 | -3.52615 | 0.000422 | 0.010193 |
| Caf1-55 | 1054.57 | -4.84443 | 1.376359 | -3.51974 | 0.000432 | 0.010411 |
| Ercc1 | 180.7639 | -9.37286 | 2.670403 | -3.5099 | 0.000448 | 0.010772 |
| lncRNA:CR | 385.6089 | 7.219855 | 2.057561 | 3.508939 | 0.00045 | 0.010779 |
| Asciz | 332.4081 | -7.03236 | 2.005208 | -3.50705 | 0.000453 | 0.010823 |
| CG14131 | 300.2395 | 8.52298 | 2.438512 | 3.495156 | 0.000474 | 0.011284 |
| Act79B | 600.4079 | -6.70352 | 1.919909 | -3.49158 | 0.00048 | 0.011308 |
| CG9395 | 2305.633 | 2.634085 | 0.754436 | 3.491464 | 0.00048 | 0.011308 |
| CG8740 | 319.0128 | -8.82065 | 2.52568 | -3.49239 | 0.000479 | 0.011308 |
| CG17030 | 306.4606 | 9.777462 | 2.800415 | 3.491434 | 0.00048 | 0.011308 |
| ste24b | 344.2349 | 9.395568 | 2.692235 | 3.489877 | 0.000483 | 0.011341 |
| esc | 310.8567 | -8.0532 | 2.309069 | -3.48764 | 0.000487 | 0.011403 |
| CG11170 | 243.071 | 8.134664 | 2.333967 | 3.485339 | 0.000492 | 0.011461 |
| hzg | 6959.567 | -2.09789 | 0.60203 | -3.48469 | 0.000493 | 0.011461 |
| Cpr67B | 275.1188 | 8.973047 | 2.575538 | 3.48395 | 0.000494 | 0.011461 |
| lncRNA:CR | 253.743 | 7.890796 | 2.265579 | 3.482905 | 0.000496 | 0.011472 |
| CG14102 | 186.1773 | 10.79347 | 3.100895 | 3.480758 | 0.0005 | 0.011532 |
| CG8326 | 204.4709 | -8.63954 | 2.483313 | -3.47904 | 0.000503 | 0.011573 |

| | | | | | | |
|---|---|---|---|---|---|---|
| slif | 249.2391 | 10.77482 | 3.099956 | 3.475799 | 0.000509 | 0.01168 |
| l(2)k01209 | 896.2943 | 5.946463 | 1.711879 | 3.473647 | 0.000513 | 0.011707 |
| CG14764 | 1841.654 | 3.488649 | 1.004107 | 3.474378 | 0.000512 | 0.011707 |
| ECSIT | 843.2693 | -3.7059 | 1.068319 | -3.46891 | 0.000523 | 0.011721 |
| Zip88E | 168.2501 | 9.323875 | 2.68737 | 3.469517 | 0.000521 | 0.011721 |
| Eip55E | 1125.887 | -4.97588 | 1.434065 | -3.46978 | 0.000521 | 0.011721 |
| CG8314 | 368.3937 | -6.75039 | 1.944553 | -3.47143 | 0.000518 | 0.011721 |
| CR18166 | 1351.337 | -4.74364 | 1.366116 | -3.47235 | 0.000516 | 0.011721 |
| Vps8 | 274.5246 | -6.27679 | 1.809502 | -3.46879 | 0.000523 | 0.011721 |
| CG6752 | 592.638 | -6.3682 | 1.840316 | -3.46038 | 0.000539 | 0.012059 |
| CG4407 | 495.2125 | 5.638813 | 1.630502 | 3.458329 | 0.000544 | 0.012118 |
| Hs3st-B | 672.8181 | 5.845631 | 1.692962 | 3.452901 | 0.000555 | 0.01233 |
| CG14785 | 173.0263 | 10.12 | 2.940355 | 3.441763 | 0.000578 | 0.012811 |
| htl | 2461.282 | -2.60372 | 0.756661 | -3.44106 | 0.000579 | 0.012811 |
| CG15547 | 268.5494 | -7.51151 | 2.18386 | -3.43955 | 0.000583 | 0.012848 |
| CG8478 | 551.4226 | 6.817863 | 1.982794 | 3.438513 | 0.000585 | 0.012862 |
| mt:lrRNA | 406634.3 | 1.225081 | 0.356418 | 3.437201 | 0.000588 | 0.012889 |
| DIP-epsilo | 456.2795 | 5.700916 | 1.662387 | 3.429356 | 0.000605 | 0.013231 |
| Zip48C | 497.4721 | 6.833859 | 1.994191 | 3.426883 | 0.000611 | 0.013315 |
| CG7194 | 323.9607 | -7.26066 | 2.119176 | -3.42617 | 0.000612 | 0.013315 |
| scramb2 | 1511.706 | -4.16989 | 1.217534 | -3.42486 | 0.000615 | 0.013343 |
| RagC-D | 2362.087 | -2.55488 | 0.746699 | -3.42157 | 0.000623 | 0.013469 |
| CG32590 | 541.4213 | 6.765557 | 1.977874 | 3.42062 | 0.000625 | 0.01348 |
| Cdk2 | 904.3587 | 5.045419 | 1.476293 | 3.417627 | 0.000632 | 0.013556 |
| Fdx1 | 402.4264 | 8.166966 | 2.389159 | 3.418344 | 0.00063 | 0.013556 |
| CG3603 | 251.8577 | -8.80068 | 2.57862 | -3.41294 | 0.000643 | 0.013727 |
| CG15784 | 632.6849 | -7.81268 | 2.289246 | -3.41277 | 0.000643 | 0.013727 |
| PolD3 | 625.6669 | 5.490858 | 1.610047 | 3.410371 | 0.000649 | 0.013811 |
| Cyp6d5 | 266.4088 | -8.94168 | 2.63292 | -3.39611 | 0.000684 | 0.014513 |
| Ns4 | 942.7734 | 5.954801 | 1.75442 | 3.394171 | 0.000688 | 0.014577 |
| Uch | 2546.599 | 3.324232 | 0.981297 | 3.387589 | 0.000705 | 0.014892 |
| CG11563 | 331.1363 | -5.28506 | 1.56246 | -3.38253 | 0.000718 | 0.01513 |
| CG14483 | 254.102 | -7.89056 | 2.336186 | -3.37754 | 0.000731 | 0.015367 |
| AhcyL2 | 306.8452 | 9.416037 | 2.794524 | 3.36946 | 0.000753 | 0.015759 |
| Ids | 1309.249 | -3.84726 | 1.141905 | -3.36916 | 0.000754 | 0.015759 |
| PIG-F | 948.425 | -4.30601 | 1.278446 | -3.36816 | 0.000757 | 0.015775 |
| CG18343 | 1006.049 | -3.59776 | 1.068869 | -3.36595 | 0.000763 | 0.015861 |
| eIF2D | 226.6383 | 6.658608 | 1.980654 | 3.361823 | 0.000774 | 0.016058 |
| Mettl14 | 347.0842 | -8.00217 | 2.384406 | -3.35604 | 0.000791 | 0.016355 |
| CG12547 | 4535.787 | -2.17667 | 0.648977 | -3.354 | 0.000797 | 0.016434 |
| CG1231 | 387.4295 | -6.30563 | 1.882746 | -3.34917 | 0.000811 | 0.016681 |
| l(2)03659 | 191.0695 | 9.918963 | 2.964133 | 3.346328 | 0.000819 | 0.016766 |
| Rnmt | 250.6406 | 7.773669 | 2.322942 | 3.346476 | 0.000818 | 0.016766 |
| SecS | 2515.165 | 3.178664 | 0.95113 | 3.341986 | 0.000832 | 0.016952 |
| mute | 152.2992 | -9.53876 | 2.854328 | -3.34186 | 0.000832 | 0.016952 |
| CG6833 | 246.5426 | -8.46486 | 2.533566 | -3.34108 | 0.000835 | 0.016957 |
| CG6227 | 710.4709 | 4.686905 | 1.403516 | 3.339402 | 0.00084 | 0.017017 |
| CG10550 | 1895.179 | 4.201064 | 1.259703 | 3.334964 | 0.000853 | 0.017204 |
| Lst8 | 333.7743 | -7.31565 | 2.193236 | -3.33555 | 0.000851 | 0.017204 |
| CG15525 | 183.9904 | 8.011043 | 2.404609 | 3.331537 | 0.000864 | 0.017304 |

| | | | | | | |
|---|---|---|---|---|---|---|
| CG7857 | 220.6829 | 8.751744 | 2.627155 | 3.331263 | 0.000865 | 0.017304 |
| CG6024 | 4638.892 | 2.072432 | 0.622119 | 3.331248 | 0.000865 | 0.017304 |
| asRNA:CR4 | 189.6959 | 9.893587 | 2.971658 | 3.329315 | 0.000871 | 0.017382 |
| Exo70 | 388.5384 | -6.59708 | 1.982731 | -3.32727 | 0.000877 | 0.017466 |
| pps | 1011.266 | 4.407321 | 1.325195 | 3.325791 | 0.000882 | 0.017516 |
| CG18011 | 371.5926 | -8.05208 | 2.422241 | -3.32423 | 0.000887 | 0.017571 |
| polybromo | 424.3416 | -6.7659 | 2.035927 | -3.32325 | 0.00089 | 0.017589 |
| btz | 914.6689 | -4.55535 | 1.371975 | -3.32029 | 0.000899 | 0.017733 |
| CG10616 | 1051.758 | -3.83995 | 1.156762 | -3.31956 | 0.000902 | 0.017736 |
| CG7497 | 420.8888 | -6.99422 | 2.107758 | -3.31832 | 0.000906 | 0.017771 |
| mTTF | 362.3099 | 7.504594 | 2.262328 | 3.317201 | 0.000909 | 0.017799 |
| CG1129 | 532.725 | -5.66945 | 1.709793 | -3.31587 | 0.000914 | 0.017806 |
| CG10467 | 647.4067 | 6.810735 | 2.054071 | 3.315726 | 0.000914 | 0.017806 |
| twit | 272.4656 | 7.90224 | 2.385011 | 3.313293 | 0.000922 | 0.017875 |
| Cortactin | 1798.371 | -3.11755 | 0.940814 | -3.31367 | 0.000921 | 0.017875 |
| pdm2 | 1182.603 | 4.790153 | 1.447344 | 3.309616 | 0.000934 | 0.017981 |
| Sirt6 | 357.8013 | -6.47529 | 1.956366 | -3.30986 | 0.000933 | 0.017981 |
| Pif1A | 296.7722 | -8.26454 | 2.49707 | -3.30969 | 0.000934 | 0.017981 |
| CG8003 | 362.1163 | -7.14342 | 2.159766 | -3.3075 | 0.000941 | 0.018075 |
| CG13096 | 344.2302 | 6.120375 | 1.85267 | 3.303543 | 0.000955 | 0.018288 |
| BCL7-like | 241.198 | -8.27431 | 2.505907 | -3.30192 | 0.00096 | 0.01835 |
| Naxd | 378.7256 | 7.110286 | 2.155085 | 3.299307 | 0.000969 | 0.018478 |
| mab-21 | 484.5247 | 5.621723 | 1.704856 | 3.297476 | 0.000976 | 0.018555 |
| Psc | 440.7514 | -6.98255 | 2.118231 | -3.29641 | 0.000979 | 0.018581 |
| CG13604 | 1563.386 | 4.385104 | 1.331175 | 3.29416 | 0.000987 | 0.018686 |
| Dlip1 | 219.1201 | 9.404478 | 2.857174 | 3.291532 | 0.000996 | 0.018817 |
| Dph2 | 485.8629 | -4.98967 | 1.517247 | -3.28863 | 0.001007 | 0.018968 |
| Hakai | 259.1976 | -7.70939 | 2.3455 | -3.28689 | 0.001013 | 0.019041 |
| Tip60 | 973.9042 | 3.890049 | 1.184822 | 3.283235 | 0.001026 | 0.019174 |
| Atg14 | 1109.619 | -4.58213 | 1.395266 | -3.28406 | 0.001023 | 0.019174 |
| CG12717 | 1887.21 | 4.238788 | 1.291153 | 3.282949 | 0.001027 | 0.019174 |
| CG8908 | 281.9925 | 8.981193 | 2.742536 | 3.274776 | 0.001057 | 0.019692 |
| CG7218 | 931.4355 | 5.282666 | 1.614116 | 3.272793 | 0.001065 | 0.019785 |
| Ir68a | 1107.767 | -5.08113 | 1.553442 | -3.27089 | 0.001072 | 0.019872 |
| CG12883 | 569.3011 | -7.16612 | 2.193845 | -3.26647 | 0.001089 | 0.020105 |
| ida | 320.7105 | 7.995967 | 2.448026 | 3.266292 | 0.00109 | 0.020105 |
| DIP-lambd | 406.5989 | -6.57254 | 2.012738 | -3.26547 | 0.001093 | 0.020117 |
| Gas41 | 536.9897 | 5.122634 | 1.569148 | 3.264595 | 0.001096 | 0.020133 |
| Pym | 541.4174 | -5.58514 | 1.714182 | -3.2582 | 0.001121 | 0.020545 |
| mRRF2 | 498.7612 | 7.302163 | 2.242386 | 3.256425 | 0.001128 | 0.02058 |
| CG1792 | 1010.659 | 4.07513 | 1.251318 | 3.256671 | 0.001127 | 0.02058 |
| TyrRS | 1220.483 | 3.43126 | 1.054094 | 3.255175 | 0.001133 | 0.020624 |
| CG32944 | 250.7137 | 8.042536 | 2.471367 | 3.254287 | 0.001137 | 0.020642 |
| CG5380 | 159.8673 | -8.22409 | 2.531853 | -3.24825 | 0.001161 | 0.021006 |
| Mrtf | 3643.333 | -2.82294 | 0.869121 | -3.24804 | 0.001162 | 0.021006 |
| Plekhm1 | 201.5792 | 7.661095 | 2.359767 | 3.246547 | 0.001168 | 0.021021 |
| CG33642 | 259.8787 | 8.190736 | 2.522661 | 3.246864 | 0.001167 | 0.021021 |
| CG15365 | 193.9673 | -7.23634 | 2.230426 | -3.24438 | 0.001177 | 0.021135 |
| lncRNA:CR | 503.624 | 6.915868 | 2.133447 | 3.241641 | 0.001188 | 0.021244 |
| nmdyn-D6 | 723.3789 | 6.292849 | 1.940895 | 3.242241 | 0.001186 | 0.021244 |

| | | | | | | |
|---|---|---|---|---|---|---|
| Dgp-1 | 773.7901 | -4.17655 | 1.290005 | -3.23762 | 0.001205 | 0.02148 |
| CR14798 | 218.4587 | 10.29215 | 3.179942 | 3.236586 | 0.00121 | 0.02148 |
| CG5897 | 254.6749 | -8.4828 | 2.620751 | -3.23678 | 0.001209 | 0.02148 |
| CG7453 | 362.3817 | 6.167975 | 1.909654 | 3.229892 | 0.001238 | 0.021941 |
| CG14881 | 428.6932 | 6.218247 | 1.926322 | 3.228041 | 0.001246 | 0.021946 |
| CG1750 | 240.6317 | -8.85493 | 2.742773 | -3.22846 | 0.001245 | 0.021946 |
| SIFa | 8481.706 | 3.696151 | 1.145493 | 3.22669 | 0.001252 | 0.021946 |
| E(var)3-9 | 272.7733 | -8.97347 | 2.780712 | -3.22704 | 0.001251 | 0.021946 |
| Hph | 1983.311 | 3.370618 | 1.044337 | 3.227518 | 0.001249 | 0.021946 |
| brun | 947.2972 | 4.710196 | 1.462455 | 3.220746 | 0.001279 | 0.022357 |
| Ent3 | 1261.506 | 4.156484 | 1.291158 | 3.219191 | 0.001286 | 0.02243 |
| FER | 1894.721 | -3.36506 | 1.046042 | -3.21694 | 0.001296 | 0.022512 |
| trus | 1739.267 | 2.977742 | 0.925658 | 3.216891 | 0.001296 | 0.022512 |
| CG32075 | 454.5363 | -4.91209 | 1.527417 | -3.21595 | 0.0013 | 0.022538 |
| Spn77Ba | 1127.538 | -3.62359 | 1.126974 | -3.21533 | 0.001303 | 0.022538 |
| CG13558 | 944.2302 | -6.68242 | 2.078928 | -3.21436 | 0.001307 | 0.022565 |
| CG10195 | 313.0799 | -6.53526 | 2.035345 | -3.21089 | 0.001323 | 0.022763 |
| Cad96Ca | 205.45 | 6.82565 | 2.125959 | 3.210621 | 0.001324 | 0.022763 |
| Cbp80 | 956.9911 | -4.27861 | 1.332951 | -3.20988 | 0.001328 | 0.022773 |
| CG18467 | 1991.158 | -2.63627 | 0.821798 | -3.20793 | 0.001337 | 0.022879 |
| tho2 | 2408.985 | -2.19941 | 0.687213 | -3.20048 | 0.001372 | 0.023429 |
| CG5180 | 1157.47 | -3.30293 | 1.033279 | -3.19655 | 0.001391 | 0.0237 |
| Rrp6 | 197.3971 | 9.208751 | 2.881931 | 3.195341 | 0.001397 | 0.023749 |
| CG5023 | 249.0937 | 8.14376 | 2.551651 | 3.191565 | 0.001415 | 0.024011 |
| CG10516 | 403.4278 | -6.96909 | 2.191828 | -3.17958 | 0.001475 | 0.024974 |
| Uxt | 185.8906 | -10.7286 | 3.378247 | -3.17578 | 0.001494 | 0.02525 |
| sdk | 1247.649 | -3.63583 | 1.145516 | -3.17396 | 0.001504 | 0.0253 |
| Ent2 | 658.1535 | 4.553409 | 1.434879 | 3.173376 | 0.001507 | 0.0253 |
| CG9143 | 508.7361 | -4.85535 | 1.529518 | -3.17443 | 0.001501 | 0.0253 |
| lncRNA:CR | 521.8903 | 7.691456 | 2.424282 | 3.172674 | 0.00151 | 0.025308 |
| aux | 1179.221 | -4.03266 | 1.271841 | -3.17073 | 0.001521 | 0.025335 |
| CG4572 | 968.3636 | 5.249194 | 1.655081 | 3.171563 | 0.001516 | 0.025335 |
| lncRNA:CR | 181.3571 | 6.531592 | 2.06008 | 3.170552 | 0.001521 | 0.025335 |
| CG7840 | 180.3894 | -9.17453 | 2.8962 | -3.16778 | 0.001536 | 0.025367 |
| lectin-46Cl | 271.0089 | -7.98731 | 2.521147 | -3.16813 | 0.001534 | 0.025367 |
| CG33941 | 826.786 | -4.23065 | 1.335269 | -3.16839 | 0.001533 | 0.025367 |
| Zfrp8 | 144.3175 | -10.1382 | 3.200066 | -3.16812 | 0.001534 | 0.025367 |
| Nf-YC | 575.0382 | -4.22732 | 1.335117 | -3.16625 | 0.001544 | 0.025448 |
| aru | 229.0191 | -6.02552 | 1.903432 | -3.16561 | 0.001548 | 0.025452 |
| asRNA:CR4 | 129.3359 | -9.43195 | 2.980061 | -3.16502 | 0.001551 | 0.025452 |
| SA | 1061.037 | -4.06664 | 1.286691 | -3.16055 | 0.001575 | 0.025794 |
| Dgkepsilor | 1797.896 | -2.59178 | 0.820449 | -3.15898 | 0.001583 | 0.025827 |
| CG32579 | 529.0231 | 7.268353 | 2.300806 | 3.159047 | 0.001583 | 0.025827 |
| CG33958 | 1281.555 | 3.963367 | 1.254939 | 3.158215 | 0.001587 | 0.025842 |
| CG32572 | 256.8676 | -7.56015 | 2.394455 | -3.15736 | 0.001592 | 0.025866 |
| tgo | 1306.939 | 4.154633 | 1.31642 | 3.156009 | 0.001599 | 0.025934 |
| p38b | 610.1778 | 5.326797 | 1.688235 | 3.155247 | 0.001604 | 0.025949 |
| CCT6 | 1177.688 | 3.434527 | 1.089203 | 3.153246 | 0.001615 | 0.026075 |
| Cdc45 | 710.5731 | 5.226303 | 1.660695 | 3.147058 | 0.001649 | 0.026549 |
| CG5087 | 565.8672 | -5.57265 | 1.770888 | -3.14681 | 0.001651 | 0.026549 |

| | | | | | | |
|---|---|---|---|---|---|---|
| CG10361 | 2046.026 | 3.095165 | 0.983797 | 3.146143 | 0.001654 | 0.026556 |
| Pgm1 | 3474.609 | -2.40413 | 0.764433 | -3.14498 | 0.001661 | 0.026608 |
| CG6686 | 2117.859 | -2.7711 | 0.881454 | -3.14379 | 0.001668 | 0.026664 |
| CG11404 | 234.2933 | 7.589843 | 2.418168 | 3.138675 | 0.001697 | 0.02708 |
| Bsg25D | 1912.231 | 2.870322 | 0.917723 | 3.127655 | 0.001762 | 0.02806 |
| E(spl)m3-H | 481.2278 | 5.394114 | 1.725276 | 3.126522 | 0.001769 | 0.028113 |
| CG15725 | 420.5907 | -5.72047 | 1.83056 | -3.12498 | 0.001778 | 0.028204 |
| Npl4 | 858.0111 | -4.78774 | 1.532866 | -3.12339 | 0.001788 | 0.028301 |
| Adck5 | 145.3715 | -11.2562 | 3.60687 | -3.12077 | 0.001804 | 0.028499 |
| gny | 507.4095 | 5.019873 | 1.609154 | 3.119574 | 0.001811 | 0.028558 |
| CG12204 | 326.7437 | -8.37963 | 2.686849 | -3.11876 | 0.001816 | 0.028572 |
| CG4069 | 146.9021 | -10.2128 | 3.275154 | -3.11828 | 0.001819 | 0.028572 |
| CG34220 | 345.882 | -7.14562 | 2.292055 | -3.11756 | 0.001824 | 0.028586 |
| Hmr | 143.8704 | -7.86686 | 2.528288 | -3.11154 | 0.001861 | 0.029092 |
| CG14274 | 2330.026 | 2.308738 | 0.742204 | 3.110651 | 0.001867 | 0.029092 |
| CG34180 | 166.6966 | 9.963093 | 3.202834 | 3.110712 | 0.001866 | 0.029092 |
| eIF2Bdelta | 460.6257 | -5.69431 | 1.831414 | -3.10924 | 0.001876 | 0.029175 |
| Suv3 | 843.8438 | 5.348948 | 1.721428 | 3.107273 | 0.001888 | 0.029313 |
| Vps13B | 475.9299 | -4.4053 | 1.418133 | -3.10641 | 0.001894 | 0.029342 |
| CG1103 | 335.3466 | 7.097025 | 2.286439 | 3.103965 | 0.001909 | 0.029529 |
| CG43340 | 366.5888 | -5.98607 | 1.928897 | -3.10336 | 0.001913 | 0.029532 |
| CG8768 | 1270.57 | 3.796387 | 1.223844 | 3.102019 | 0.001922 | 0.029609 |
| Ntmt | 320.8124 | -7.29746 | 2.353693 | -3.10043 | 0.001932 | 0.029712 |
| Ns2 | 632.969 | 4.901191 | 1.583213 | 3.095724 | 0.001963 | 0.030072 |
| CG17224 | 997.0162 | -4.69924 | 1.517938 | -3.0958 | 0.001963 | 0.030072 |
| CG5828 | 311.7113 | 7.647859 | 2.473747 | 3.091609 | 0.001991 | 0.030434 |
| Nprl2 | 264.0648 | -8.42347 | 2.725847 | -3.09022 | 0.002 | 0.030518 |
| Pex2 | 497.2781 | 6.485832 | 2.106328 | 3.079212 | 0.002075 | 0.031609 |
| lncRNA:CR | 143.9626 | -9.54791 | 3.103335 | -3.07666 | 0.002093 | 0.03182 |
| Hf | 2394.058 | -3.52923 | 1.148151 | -3.07383 | 0.002113 | 0.032063 |
| sxe2 | 748.8704 | -6.00621 | 1.955373 | -3.07165 | 0.002129 | 0.032182 |
| CG14907 | 234.0529 | -8.28743 | 2.698089 | -3.07159 | 0.002129 | 0.032182 |
| Mst87F | 193.7006 | -9.97514 | 3.24947 | -3.06977 | 0.002142 | 0.032289 |
| CG42339 | 1418.206 | -4.39576 | 1.432086 | -3.06948 | 0.002144 | 0.032289 |
| lncRNA:CR | 151.5731 | 10.88626 | 3.549792 | 3.066731 | 0.002164 | 0.032466 |
| cpb | 848.8881 | -4.30298 | 1.403078 | -3.06682 | 0.002164 | 0.032466 |
| Pex3 | 2389.306 | -2.21023 | 0.721061 | -3.06525 | 0.002175 | 0.032567 |
| CG4679 | 1045.302 | 3.632887 | 1.18708 | 3.060355 | 0.002211 | 0.033027 |
| CG9512 | 732.9641 | 3.358927 | 1.097885 | 3.059451 | 0.002217 | 0.033027 |
| smid | 630.3253 | -4.95243 | 1.618771 | -3.05938 | 0.002218 | 0.033027 |
| CG12512 | 843.8631 | -4.96064 | 1.622511 | -3.05739 | 0.002233 | 0.033175 |
| CG14402 | 500.9265 | 6.672982 | 2.183077 | 3.056686 | 0.002238 | 0.033175 |
| CG6040 | 4198.909 | 1.850666 | 0.605725 | 3.05529 | 0.002248 | 0.033175 |
| boly | 163.5486 | -8.40763 | 2.751201 | -3.05599 | 0.002243 | 0.033175 |
| asRNA:CR4 | 187.7005 | 9.518524 | 3.115443 | 3.055271 | 0.002249 | 0.033175 |
| AlkB | 203.6788 | -7.0195 | 2.302971 | -3.04802 | 0.002304 | 0.033924 |
| Exd2 | 291.5768 | 9.393876 | 3.082718 | 3.04727 | 0.002309 | 0.033946 |
| Dph4 | 360.6354 | -5.43862 | 1.788316 | -3.0412 | 0.002356 | 0.034575 |
| CG12698 | 213.9935 | -7.82471 | 2.5755 | -3.03813 | 0.00238 | 0.034865 |
| unc-45 | 1380.353 | 4.545576 | 1.498299 | 3.033825 | 0.002415 | 0.035188 |

| | | | | | | |
|---|---|---|---|---|---|---|
| Gp150 | 740.1867 | -4.61235 | 1.519949 | -3.03454 | 0.002409 | 0.035188 |
| Uba3 | 200.3697 | 7.652632 | 2.52254 | 3.033701 | 0.002416 | 0.035188 |
| CG8461 | 192.838 | 8.20623 | 2.705873 | 3.032748 | 0.002423 | 0.035236 |
| CG13982 | 818.9768 | -4.07116 | 1.344046 | -3.02904 | 0.002453 | 0.035607 |
| noi | 394.9077 | -7.80853 | 2.578809 | -3.02796 | 0.002462 | 0.035669 |
| Cht11 | 550.4085 | 5.511126 | 1.822069 | 3.024653 | 0.002489 | 0.035997 |
| daw | 828.0157 | -4.52335 | 1.496468 | -3.02268 | 0.002505 | 0.036167 |
| VhaSFD | 9605.574 | -1.86681 | 0.618606 | -3.01778 | 0.002546 | 0.036691 |
| CG17279 | 1719.317 | -4.57191 | 1.516746 | -3.01429 | 0.002576 | 0.037049 |
| Task7 | 3641.67 | 3.02253 | 1.003922 | 3.010721 | 0.002606 | 0.037343 |
| Hs6st | 628.9733 | -5.09381 | 1.692147 | -3.01027 | 0.00261 | 0.037343 |
| Arfrp1 | 951.6222 | 4.200139 | 1.395246 | 3.010321 | 0.00261 | 0.037343 |
| Spps | 732.8364 | -4.72143 | 1.572805 | -3.00191 | 0.002683 | 0.038315 |
| Iru | 1738.358 | 3.342589 | 1.113977 | 3.000591 | 0.002695 | 0.038413 |
| CG7381 | 870.7647 | 4.509094 | 1.503387 | 2.999291 | 0.002706 | 0.03848 |
| mahj | 170.3569 | 6.194743 | 2.065615 | 2.998983 | 0.002709 | 0.03848 |
| lncRNA:CR | 823.3874 | 3.348702 | 1.119144 | 2.992199 | 0.00277 | 0.039276 |
| caup | 165.5269 | -7.21652 | 2.413546 | -2.99001 | 0.00279 | 0.039489 |
| Alg13 | 191.867 | -7.4715 | 2.500997 | -2.98741 | 0.002814 | 0.039756 |
| CG31957 | 306.1941 | -5.2922 | 1.772332 | -2.98601 | 0.002826 | 0.039868 |
| CG15296 | 445.8205 | 6.697415 | 2.245151 | 2.983058 | 0.002854 | 0.040184 |
| CG10324 | 151.4422 | -8.43134 | 2.828328 | -2.98103 | 0.002873 | 0.040309 |
| mrt | 2082.552 | 2.211863 | 0.741966 | 2.981086 | 0.002872 | 0.040309 |
| lncRNA:CR | 273.6858 | -5.60465 | 1.881505 | -2.97881 | 0.002894 | 0.040531 |
| CG2006 | 195.9753 | -8.63279 | 2.899004 | -2.97785 | 0.002903 | 0.040589 |
| Smu1 | 1367.777 | -3.75653 | 1.26376 | -2.9725 | 0.002954 | 0.041087 |
| CG43707 | 5219.841 | 1.841053 | 0.619261 | 2.972984 | 0.002949 | 0.041087 |
| Sec3 | 386.4934 | -7.3563 | 2.474207 | -2.97319 | 0.002947 | 0.041087 |
| Rcp | 161.5665 | -10.5392 | 3.546384 | -2.97182 | 0.00296 | 0.041107 |
| CG12003 | 148.1954 | 7.659175 | 2.578375 | 2.970543 | 0.002973 | 0.041208 |
| mRpL51 | 2197.612 | 3.08941 | 1.041075 | 2.967521 | 0.003002 | 0.041543 |
| CG11619 | 576.4201 | 4.562028 | 1.537619 | 2.966944 | 0.003008 | 0.04155 |
| l(2)41Ab | 1348.805 | -2.6731 | 0.901751 | -2.96434 | 0.003033 | 0.04183 |
| CG8301 | 2203.975 | 2.203496 | 0.743542 | 2.963512 | 0.003042 | 0.041872 |
| GluProRS | 853.7945 | -4.41813 | 1.49376 | -2.95772 | 0.003099 | 0.04252 |
| CG15443 | 578.6738 | 4.958264 | 1.676274 | 2.957907 | 0.003097 | 0.04252 |
| kri | 2220.823 | -2.70816 | 0.917408 | -2.95197 | 0.003158 | 0.043247 |
| asRNA:CR4 | 179.5114 | -8.32857 | 2.823215 | -2.95003 | 0.003177 | 0.043445 |
| a6 | 431.2616 | -4.51821 | 1.532178 | -2.94888 | 0.003189 | 0.043532 |
| Slbp | 134.2679 | 9.373119 | 3.179813 | 2.947695 | 0.003202 | 0.043552 |
| CG15747 | 790.2822 | 3.889128 | 1.319245 | 2.947996 | 0.003198 | 0.043552 |
| CG17333 | 162.4258 | -9.51362 | 3.230877 | -2.94459 | 0.003234 | 0.043916 |
| puf | 1049.184 | 3.592071 | 1.221167 | 2.941505 | 0.003266 | 0.044275 |
| mthl10 | 2817.928 | -1.98226 | 0.674121 | -2.94051 | 0.003277 | 0.044275 |
| CG7326 | 529.0427 | 5.029001 | 1.710034 | 2.940878 | 0.003273 | 0.044275 |
| CG8134 | 664.4708 | 4.085597 | 1.390551 | 2.938113 | 0.003302 | 0.044543 |
| Dhfr | 333.323 | -7.17612 | 2.444015 | -2.9362 | 0.003323 | 0.044743 |
| Hen1 | 297.4203 | -5.57907 | 1.902049 | -2.93319 | 0.003355 | 0.045104 |
| CG32095 | 192.163 | -7.26085 | 2.477407 | -2.93083 | 0.003381 | 0.045373 |
| Atg4a | 409.0695 | 6.250511 | 2.133252 | 2.930039 | 0.003389 | 0.045412 |

| | | | | | | |
|---|---|---|---|---|---|---|
| nvy | 209.3376 | 6.883474 | 2.351248 | 2.927583 | 0.003416 | 0.045696 |
| CG6136 | 334.6365 | -6.10499 | 2.086103 | -2.9265 | 0.003428 | 0.045778 |
| cnk | 1040.83 | -3.27176 | 1.119801 | -2.92173 | 0.003481 | 0.046255 |
| CG7650 | 133.6169 | -9.67938 | 3.312632 | -2.92196 | 0.003478 | 0.046255 |
| dock | 2303.497 | 2.464331 | 0.84323 | 2.92249 | 0.003472 | 0.046255 |
| Tango6 | 306.702 | 6.050854 | 2.07325 | 2.918536 | 0.003517 | 0.046602 |
| Mfap1 | 405.6509 | 6.793838 | 2.327958 | 2.918368 | 0.003519 | 0.046602 |
| GluRIID | 166.8356 | -8.9029 | 3.056623 | -2.91266 | 0.003584 | 0.047306 |
| Fancd2 | 556.5936 | -4.96731 | 1.705189 | -2.91306 | 0.003579 | 0.047306 |
| Mocs1 | 319.6345 | -5.25331 | 1.808676 | -2.9045 | 0.003678 | 0.048477 |
| CCDC53 | 566.1721 | -4.16044 | 1.432932 | -2.90345 | 0.003691 | 0.048482 |
| Rrp4 | 189.4188 | 7.628534 | 2.627101 | 2.903784 | 0.003687 | 0.048482 |
| CG6685 | 254.8936 | -6.18806 | 2.133027 | -2.90107 | 0.003719 | 0.048771 |
| Fer3 | 171.4533 | -9.89462 | 3.413201 | -2.89893 | 0.003744 | 0.049012 |
| asRNA:CR4 | 128.511 | -9.31718 | 3.214482 | -2.8985 | 0.00375 | 0.049012 |
| Sf3b3 | 304.9988 | -6.46898 | 2.232349 | -2.89784 | 0.003757 | 0.049037 |
| CG3919 | 222.711 | -7.85099 | 2.711525 | -2.89541 | 0.003787 | 0.049336 |
| CG42806 | 242.4264 | -5.98533 | 2.071086 | -2.88995 | 0.003853 | 0.050121 |
| sr | 1440.362 | 2.463844 | 0.85323 | 2.887668 | 0.003881 | 0.050404 |
| CG12817 | 1005.012 | -3.55558 | 1.23224 | -2.88546 | 0.003908 | 0.050455 |
| Spc25 | 224.3514 | 6.816733 | 2.361626 | 2.886458 | 0.003896 | 0.050455 |
| Ir76a | 760.3641 | 4.564817 | 1.581575 | 2.886247 | 0.003899 | 0.050455 |
| thoc6 | 107.2361 | -11.0941 | 3.845009 | -2.88532 | 0.00391 | 0.050455 |
| CG30060 | 237.3832 | 8.409392 | 2.91634 | 2.883543 | 0.003932 | 0.050659 |
| CG5867 | 1053.993 | 5.170509 | 1.794635 | 2.881092 | 0.003963 | 0.050891 |
| Miga | 1019.141 | 4.04607 | 1.404204 | 2.881397 | 0.003959 | 0.050891 |
| Arc42 | 1029.605 | -3.14927 | 1.09382 | -2.87915 | 0.003988 | 0.051124 |
| BCAS2 | 333.4019 | 4.984891 | 1.732216 | 2.877754 | 0.004005 | 0.051269 |
| CR43086 | 298.2857 | 7.04338 | 2.455724 | 2.868148 | 0.004129 | 0.052767 |
| CG11899 | 341.2765 | 5.986335 | 2.088384 | 2.866491 | 0.00415 | 0.05296 |
| CG13409 | 1237.993 | -3.94998 | 1.378822 | -2.86475 | 0.004173 | 0.053167 |
| Lhr | 143.1335 | 9.774337 | 3.41443 | 2.862656 | 0.004201 | 0.053435 |
| ZIPIC | 315.3225 | 6.781727 | 2.371559 | 2.859608 | 0.004242 | 0.053781 |
| Arpc2 | 1197.437 | -3.94163 | 1.378244 | -2.85989 | 0.004238 | 0.053781 |
| Ak3 | 1070.858 | 3.209396 | 1.123406 | 2.856845 | 0.004279 | 0.053996 |
| CG4282 | 208.6549 | 7.572222 | 2.650075 | 2.857361 | 0.004272 | 0.053996 |
| bonsai | 548.5875 | -4.56433 | 1.597625 | -2.85695 | 0.004277 | 0.053996 |
| CG3386 | 1038.778 | -3.38383 | 1.185445 | -2.85448 | 0.004311 | 0.054314 |
| CG7461 | 755.8778 | -4.5306 | 1.588035 | -2.85296 | 0.004331 | 0.054406 |
| Pkcdelta | 602.1091 | -4.36181 | 1.528879 | -2.85295 | 0.004332 | 0.054406 |
| mt:ND4 | 65940.37 | 1.11424 | 0.390657 | 2.852219 | 0.004342 | 0.054445 |
| Archease | 826.5847 | 4.391931 | 1.540142 | 2.85164 | 0.004349 | 0.05446 |
| CG17652 | 310.1083 | 6.200059 | 2.174917 | 2.85071 | 0.004362 | 0.054534 |
| Gorab | 296.4488 | -5.23129 | 1.836437 | -2.84861 | 0.004391 | 0.05481 |
| CG31921 | 168.7302 | -8.41463 | 2.95582 | -2.8468 | 0.004416 | 0.055037 |
| CG5188 | 145.2386 | -7.87452 | 2.76916 | -2.84365 | 0.00446 | 0.055498 |
| CG43231 | 153.7393 | -7.27298 | 2.561543 | -2.8393 | 0.004521 | 0.056 |
| asRNA:CR4 | 158.9915 | 7.714989 | 2.716436 | 2.840115 | 0.00451 | 0.056 |
| teq | 14286.24 | -1.50949 | 0.531581 | -2.83962 | 0.004517 | 0.056 |
| CG14182 | 1284.321 | 3.559793 | 1.254381 | 2.837888 | 0.004541 | 0.056161 |

| | | | | | | |
|---|---|---|---|---|---|---|
| Nepl16 | 6791.656 | 1.941232 | 0.684467 | 2.836121 | 0.004567 | 0.056386 |
| CG8027 | 359.0962 | 7.438512 | 2.624346 | 2.834424 | 0.004591 | 0.056599 |
| CG3223 | 487.3376 | 4.073228 | 1.438034 | 2.832497 | 0.004619 | 0.056626 |
| geminin | 878.2448 | -2.90306 | 1.024977 | -2.83232 | 0.004621 | 0.056626 |
| CG11710 | 659.1755 | 4.589984 | 1.620019 | 2.833291 | 0.004607 | 0.056626 |
| lncRNA:CR | 173.1715 | 8.423506 | 2.973612 | 2.832752 | 0.004615 | 0.056626 |
| onecut | 10125.62 | 1.31003 | 0.462876 | 2.830198 | 0.004652 | 0.056915 |
| CG3740 | 384.2468 | -5.10767 | 1.805701 | -2.82864 | 0.004675 | 0.057023 |
| MFS9 | 2329.926 | 3.00032 | 1.060914 | 2.828051 | 0.004683 | 0.057023 |
| CG30403 | 381.9495 | 6.12507 | 2.166138 | 2.827645 | 0.004689 | 0.057023 |
| Cdk8 | 294.5117 | -6.53333 | 2.31022 | -2.82801 | 0.004684 | 0.057023 |
| ArfGAP3 | 1767.776 | 2.196935 | 0.777331 | 2.826253 | 0.00471 | 0.057099 |
| lncRNA:CR | 214.6175 | 6.569622 | 2.324426 | 2.826341 | 0.004708 | 0.057099 |
| sls | 563.3432 | 4.179674 | 1.479821 | 2.824447 | 0.004736 | 0.057335 |
| CCKLR-17D | 609.199 | 4.931289 | 1.746316 | 2.823823 | 0.004745 | 0.05736 |
| nerfin-2 | 139.9929 | -9.68463 | 3.431805 | -2.82202 | 0.004772 | 0.057596 |
| Vps13 | 1807.039 | -2.17431 | 0.770622 | -2.82151 | 0.00478 | 0.057603 |
| MED28 | 716.8749 | 3.602928 | 1.277747 | 2.819751 | 0.004806 | 0.057689 |
| Chrac-14 | 344.6763 | 6.823345 | 2.419422 | 2.820237 | 0.004799 | 0.057689 |
| Plod | 1114.121 | -3.72004 | 1.319357 | -2.81958 | 0.004809 | 0.057689 |
| Uck | 3900.445 | 1.795396 | 0.636928 | 2.818836 | 0.00482 | 0.057737 |
| CG43861 | 384.0517 | 5.018219 | 1.781076 | 2.817521 | 0.00484 | 0.057888 |
| HtrA2 | 1058.058 | 3.936277 | 1.397914 | 2.815822 | 0.004865 | 0.058108 |
| lncRNA:CR | 186.2343 | 7.504041 | 2.665606 | 2.815135 | 0.004876 | 0.058146 |
| janA | 305.4322 | -6.43843 | 2.28959 | -2.81204 | 0.004923 | 0.058166 |
| CG8841 | 733.288 | -4.53161 | 1.610552 | -2.8137 | 0.004897 | 0.058166 |
| RYBP | 1370.764 | -4.43925 | 1.578579 | -2.81218 | 0.004921 | 0.058166 |
| Taldo | 2164.103 | -2.45664 | 0.873722 | -2.8117 | 0.004928 | 0.058166 |
| naz | 296.9732 | -5.51785 | 1.961636 | -2.81288 | 0.00491 | 0.058166 |
| CG12716 | 365.2292 | 6.183745 | 2.198519 | 2.812686 | 0.004913 | 0.058166 |
| Naa20A | 1238.03 | -4.05773 | 1.441844 | -2.81427 | 0.004889 | 0.058166 |
| CG14434 | 321.2856 | 5.194838 | 1.849822 | 2.808291 | 0.004981 | 0.058612 |
| lncRNA:CR | 123.3862 | 9.261278 | 3.297781 | 2.808336 | 0.00498 | 0.058612 |
| Fbw5 | 318.6209 | 6.915958 | 2.470034 | 2.799945 | 0.005111 | 0.059972 |
| Kyat | 1521.112 | 3.481576 | 1.24358 | 2.79964 | 0.005116 | 0.059972 |
| CG32687 | 207.9352 | -8.41883 | 3.00777 | -2.79903 | 0.005126 | 0.059972 |
| mri | 2549.194 | -3.16378 | 1.130321 | -2.79901 | 0.005126 | 0.059972 |
| Tace | 4143.692 | 3.025633 | 1.08174 | 2.797005 | 0.005158 | 0.060258 |
| Plap | 421.2174 | 5.296261 | 1.894566 | 2.795501 | 0.005182 | 0.060451 |
| Nhe2 | 4864.39 | 2.090221 | 0.748031 | 2.794299 | 0.005201 | 0.060588 |
| lncRNA:CR | 195.5539 | 7.653577 | 2.743315 | 2.7899 | 0.005272 | 0.061328 |
| Ssu72 | 836.1773 | 5.737415 | 2.057208 | 2.788932 | 0.005288 | 0.061423 |
| Odj | 449.5065 | -4.01 | 1.438726 | -2.78719 | 0.005317 | 0.061574 |
| Regnase-1 | 248.0933 | 8.313861 | 2.983003 | 2.787078 | 0.005319 | 0.061574 |
| lncRNA:CR | 203.786 | 7.750659 | 2.78127 | 2.786734 | 0.005324 | 0.061574 |
| CG33293 | 152.602 | -7.59272 | 2.725868 | -2.78543 | 0.005346 | 0.061699 |
| MICAL-like | 1848.413 | 2.858362 | 1.026461 | 2.784678 | 0.005358 | 0.061699 |
| CG7600 | 745.1858 | -4.35515 | 1.563947 | -2.78472 | 0.005357 | 0.061699 |
| CG4294 | 3745.538 | -1.76303 | 0.633353 | -2.78364 | 0.005375 | 0.061808 |
| wol | 1413.544 | 3.367744 | 1.211283 | 2.780311 | 0.005431 | 0.062356 |

| | | | | | | |
|---|---|---|---|---|---|---|
| CG5022 | 1295.321 | -3.59291 | 1.294346 | -2.77585 | 0.005506 | 0.062948 |
| CG4622 | 567.7456 | -4.15606 | 1.497092 | -2.77609 | 0.005502 | 0.062948 |
| kmr | 12552.25 | -1.3208 | 0.475732 | -2.77635 | 0.005497 | 0.062948 |
| disp | 1190.192 | 3.59178 | 1.295201 | 2.773144 | 0.005552 | 0.063258 |
| CG11836 | 121.2299 | -8.38893 | 3.025871 | -2.7724 | 0.005564 | 0.063258 |
| Marf1 | 1411.1 | 2.717357 | 0.979931 | 2.773007 | 0.005554 | 0.063258 |
| Mip | 3659.284 | 3.150347 | 1.1362 | 2.772705 | 0.005559 | 0.063258 |
| CG4896 | 2435.26 | -1.95025 | 0.703778 | -2.77112 | 0.005586 | 0.063417 |
| nompC | 228.9207 | 6.469343 | 2.335799 | 2.769648 | 0.005612 | 0.063615 |
| ZnT33D | 238.0963 | 7.766818 | 2.805349 | 2.768575 | 0.00563 | 0.063735 |
| Cdk5alpha | 7896.256 | 2.059186 | 0.744014 | 2.767671 | 0.005646 | 0.063763 |
| CCT1 | 566.8566 | -3.7565 | 1.357353 | -2.76752 | 0.005649 | 0.063763 |
| CG11898 | 182.1878 | 6.131679 | 2.216614 | 2.766236 | 0.005671 | 0.063903 |
| flr | 669.0114 | 4.706198 | 1.701518 | 2.765882 | 0.005677 | 0.063903 |
| hb | 137.6619 | 8.401518 | 3.041004 | 2.762745 | 0.005732 | 0.06443 |
| ush | 248.3504 | 6.330136 | 2.294997 | 2.758233 | 0.005811 | 0.065235 |
| FoxP | 9681.467 | 1.464058 | 0.531554 | 2.754297 | 0.005882 | 0.065932 |
| Dcr-2 | 837.441 | -3.75589 | 1.364583 | -2.75241 | 0.005916 | 0.066221 |
| cerv | 377.3708 | -5.74989 | 2.090497 | -2.75049 | 0.005951 | 0.066518 |
| Alr | 369.2748 | -5.81996 | 2.117119 | -2.749 | 0.005978 | 0.066728 |
| SrpRbeta | 5297.872 | -2.05935 | 0.749545 | -2.74746 | 0.006006 | 0.066948 |
| Rnp4F | 504.7707 | -4.69297 | 1.71035 | -2.74386 | 0.006072 | 0.067499 |
| CG7322 | 1264.359 | -2.47462 | 0.901739 | -2.74428 | 0.006064 | 0.067499 |
| CG15317 | 266.545 | -7.7003 | 2.807659 | -2.7426 | 0.006095 | 0.067572 |
| Elp3 | 285.6166 | -8.29952 | 3.025891 | -2.74283 | 0.006091 | 0.067572 |
| CG7728 | 91.67118 | -10.54 | 3.844551 | -2.74155 | 0.006115 | 0.067695 |
| CG18547 | 120.723 | 9.521299 | 3.475221 | 2.739768 | 0.006148 | 0.06797 |
| CG2698 | 4019.654 | 2.183482 | 0.797354 | 2.738409 | 0.006174 | 0.068157 |
| asRNA:CR4 | 190.0945 | 6.047265 | 2.208844 | 2.737751 | 0.006186 | 0.0682 |
| TyrR | 373.1447 | -5.43559 | 1.988818 | -2.73307 | 0.006275 | 0.069081 |
| dnd | 194.9955 | 6.880055 | 2.520515 | 2.729622 | 0.006341 | 0.069522 |
| ClC-b | 1839.227 | 2.561073 | 0.938246 | 2.729639 | 0.00634 | 0.069522 |
| Gasz | 1452.514 | 2.318114 | 0.849186 | 2.729809 | 0.006337 | 0.069522 |
| Ate1 | 1229.926 | -3.09136 | 1.133004 | -2.72846 | 0.006363 | 0.069577 |
| CG41520 | 143.581 | -9.14795 | 3.352404 | -2.72877 | 0.006357 | 0.069577 |
| Oct-TyrR | 886.2351 | 3.085967 | 1.131698 | 2.726847 | 0.006394 | 0.069784 |
| Rep | 1132.465 | -3.17344 | 1.163886 | -2.72659 | 0.006399 | 0.069784 |
| Sas10 | 389.7616 | 5.952047 | 2.183823 | 2.725518 | 0.00642 | 0.069915 |
| CG11686 | 855.127 | -4.58003 | 1.680905 | -2.72474 | 0.006435 | 0.069985 |
| Mfe2 | 846.1458 | 3.431208 | 1.259774 | 2.72367 | 0.006456 | 0.070117 |
| Naa60 | 3148.017 | 1.717837 | 0.630976 | 2.722506 | 0.006479 | 0.07027 |
| CG3842 | 406.8108 | -5.56164 | 2.043258 | -2.72195 | 0.00649 | 0.070294 |
| Fer1 | 913.3424 | -3.69044 | 1.358822 | -2.71591 | 0.006609 | 0.071492 |
| CG12868 | 1099.437 | -3.11565 | 1.147897 | -2.71422 | 0.006643 | 0.07176 |
| CG9485 | 92.76707 | -8.16752 | 3.01052 | -2.71299 | 0.006668 | 0.071833 |
| lawc | 5093.352 | 1.512278 | 0.557404 | 2.713074 | 0.006666 | 0.071833 |
| Eip63F-1 | 2859.852 | 2.441385 | 0.900463 | 2.711254 | 0.006703 | 0.072115 |
| Hmt-1 | 2174.013 | -2.99827 | 1.107654 | -2.70686 | 0.006792 | 0.07277 |
| CG17841 | 8056.388 | -1.67895 | 0.620343 | -2.70648 | 0.0068 | 0.07277 |
| Rhp | 950.2928 | 3.373105 | 1.245815 | 2.70755 | 0.006778 | 0.07277 |

| | | | | | | |
|---|---|---|---|---|---|---|
| Use1 | 120.482 | 9.481931 | 3.502764 | 2.706986 | 0.00679 | 0.07277 |
| CG6903 | 2210.825 | -2.1168 | 0.783523 | -2.70164 | 0.0069 | 0.07358 |
| ATP8A | 5869.129 | 1.451224 | 0.537197 | 2.701476 | 0.006903 | 0.07358 |
| CG5535 | 655.9094 | -4.39312 | 1.626082 | -2.70166 | 0.006899 | 0.07358 |
| ebd2 | 609.2361 | -4.81654 | 1.783324 | -2.70088 | 0.006916 | 0.073614 |
| CG33096 | 169.3463 | -6.99857 | 2.593165 | -2.69885 | 0.006958 | 0.073857 |
| koi | 2400.069 | 2.46713 | 0.914004 | 2.699254 | 0.00695 | 0.073857 |
| CG9951 | 472.1606 | 5.499629 | 2.038062 | 2.698461 | 0.006966 | 0.073857 |
| CG7110 | 208.0684 | 6.511041 | 2.414959 | 2.696129 | 0.007015 | 0.074278 |
| Caf1-105 | 638.3333 | 5.1296 | 1.903025 | 2.695498 | 0.007028 | 0.074321 |
| Lrch | 1916.72 | 2.884784 | 1.070426 | 2.694986 | 0.007039 | 0.074337 |
| nopo | 144.7412 | -9.82702 | 3.647312 | -2.69432 | 0.007053 | 0.074388 |
| Gfat2 | 931.2668 | -3.41709 | 1.268725 | -2.69333 | 0.007074 | 0.074511 |
| CG3645 | 700.2804 | -3.74205 | 1.390523 | -2.69111 | 0.007121 | 0.074714 |
| CG5335 | 542.0544 | -5.87906 | 2.184265 | -2.69155 | 0.007112 | 0.074714 |
| CG2889 | 1588.458 | 2.73283 | 1.015482 | 2.691165 | 0.00712 | 0.074714 |
| lncRNA:CR | 169.9043 | 7.290064 | 2.709598 | 2.69046 | 0.007135 | 0.074763 |
| pkaap | 1399.246 | 2.308616 | 0.860364 | 2.683301 | 0.00729 | 0.076058 |
| htt | 1357.774 | -3.40694 | 1.269489 | -2.68371 | 0.007281 | 0.076058 |
| CG8389 | 1152.404 | -4.74436 | 1.768318 | -2.68298 | 0.007297 | 0.076058 |
| CG15412 | 361.8056 | 6.589387 | 2.455323 | 2.683715 | 0.007281 | 0.076058 |
| NC2alpha | 266.8113 | -6.61556 | 2.467345 | -2.68125 | 0.007335 | 0.076256 |
| lbk | 1977.756 | -2.03935 | 0.760498 | -2.68161 | 0.007327 | 0.076256 |
| CG10431 | 179.9195 | -8.02439 | 2.993999 | -2.68016 | 0.007359 | 0.076405 |
| Nep4 | 5067.539 | -1.57998 | 0.590395 | -2.67615 | 0.007447 | 0.077226 |
| CG15343 | 156.2043 | -8.37741 | 3.131826 | -2.67493 | 0.007474 | 0.077407 |
| dgt2 | 144.4108 | 9.743285 | 3.64486 | 2.673158 | 0.007514 | 0.077617 |
| Nph | 752.0837 | -3.45232 | 1.291419 | -2.67327 | 0.007511 | 0.077617 |
| CG3505 | 229.7952 | 7.117419 | 2.666137 | 2.669563 | 0.007595 | 0.078352 |
| pk | 171.8179 | -7.90012 | 2.962394 | -2.6668 | 0.007658 | 0.078896 |
| SCCRO | 1799.13 | -2.2745 | 0.853371 | -2.66532 | 0.007692 | 0.079144 |
| lncRNA:CR | 300.9669 | -6.14813 | 2.309583 | -2.66201 | 0.007768 | 0.079824 |
| kat80 | 366.7111 | 4.549341 | 1.709405 | 2.661359 | 0.007783 | 0.079876 |
| Atac2 | 356.4966 | 6.934933 | 2.606774 | 2.660351 | 0.007806 | 0.080013 |
| CG9314 | 92.43829 | -9.67628 | 3.639069 | -2.659 | 0.007837 | 0.080233 |
| CG31875 | 457.3288 | 6.529753 | 2.460202 | 2.654153 | 0.007951 | 0.080981 |
| Nse4 | 563.2484 | -4.54551 | 1.711879 | -2.65527 | 0.007924 | 0.080981 |
| CG11103 | 1910.278 | 2.402606 | 0.90508 | 2.654578 | 0.007941 | 0.080981 |
| CG7692 | 589.2173 | 4.364879 | 1.644124 | 2.654836 | 0.007935 | 0.080981 |
| alphaCOP | 1616.733 | -2.26247 | 0.853205 | -2.65173 | 0.008008 | 0.081462 |
| CG11656 | 108.32 | -9.34513 | 3.525179 | -2.65097 | 0.008026 | 0.081542 |
| snoRNA:M | 213.5648 | -7.19975 | 2.71925 | -2.6477 | 0.008104 | 0.08223 |
| CG9286 | 439.9763 | -5.20594 | 1.968417 | -2.64473 | 0.008176 | 0.082849 |
| Mst33A | 127.2006 | 7.724825 | 2.922508 | 2.643217 | 0.008212 | 0.083116 |
| MED27 | 731.8493 | 4.004002 | 1.51511 | 2.642714 | 0.008224 | 0.083135 |
| Klp64D | 2210.577 | 2.037925 | 0.77132 | 2.642126 | 0.008239 | 0.083175 |
| atk | 274.0617 | -7.00388 | 2.652252 | -2.64073 | 0.008273 | 0.083414 |
| CG4813 | 124.4774 | 9.614789 | 3.646828 | 2.63648 | 0.008377 | 0.08436 |
| CG18528 | 187.208 | 7.723485 | 2.930563 | 2.635495 | 0.008401 | 0.084393 |
| CG32335 | 137.9682 | -9.78053 | 3.710522 | -2.63589 | 0.008392 | 0.084393 |

| | | | | | | |
|---|---|---|---|---|---|---|
| asRNA:CR4 | 191.9417 | -7.46051 | 2.831252 | -2.63506 | 0.008412 | 0.084396 |
| CG13126 | 189.8393 | -5.80331 | 2.205042 | -2.63184 | 0.008492 | 0.085094 |
| Wdr24 | 155.6991 | -7.23691 | 2.753955 | -2.62782 | 0.008593 | 0.085997 |
| Dr | 298.6096 | -6.56969 | 2.501428 | -2.62638 | 0.00863 | 0.086257 |
| Kul | 3876.102 | 1.692052 | 0.644665 | 2.624698 | 0.008673 | 0.086575 |
| CG4957 | 942.7696 | 3.934087 | 1.500838 | 2.621261 | 0.008761 | 0.087329 |
| lncRNA:CR | 141446.4 | 1.987161 | 0.7582 | 2.620895 | 0.00877 | 0.087329 |
| CG30467 | 370.5213 | -4.68726 | 1.789729 | -2.61898 | 0.008819 | 0.087604 |
| calypso | 499.9495 | 4.475093 | 1.708643 | 2.619091 | 0.008816 | 0.087604 |
| Prx6005 | 660.7555 | -4.18639 | 1.600616 | -2.61549 | 0.00891 | 0.088396 |
| lncRNA:CR | 114.3273 | -8.04679 | 3.079368 | -2.61313 | 0.008972 | 0.088788 |
| CG43783 | 904.7741 | 3.243639 | 1.241117 | 2.613483 | 0.008962 | 0.088788 |
| CG2861 | 158.8498 | -8.08567 | 3.095161 | -2.61236 | 0.008992 | 0.088879 |
| CG42354 | 154.0031 | -6.42675 | 2.462083 | -2.61029 | 0.009047 | 0.089309 |
| rump | 1125.541 | 2.393972 | 0.918229 | 2.607162 | 0.00913 | 0.090018 |
| CG12877 | 693.336 | 3.912282 | 1.504661 | 2.600108 | 0.009319 | 0.091776 |
| CG4278 | 302.7341 | 4.652477 | 1.790498 | 2.598427 | 0.009365 | 0.092114 |
| didum | 2053.645 | 2.584544 | 0.995793 | 2.595464 | 0.009446 | 0.092798 |
| acj6 | 316.4632 | -4.79472 | 1.848082 | -2.59443 | 0.009475 | 0.092965 |
| CG30148 | 133.7965 | -9.67521 | 3.732203 | -2.59236 | 0.009532 | 0.093152 |
| CG8490 | 391.8206 | -4.34442 | 1.675578 | -2.59279 | 0.00952 | 0.093152 |
| Hip1 | 571.4306 | -4.75902 | 1.835176 | -2.59322 | 0.009508 | 0.093152 |
| CG8549 | 826.0761 | -3.74326 | 1.444128 | -2.59206 | 0.00954 | 0.093152 |
| CG8485 | 1304.715 | -2.72333 | 1.051412 | -2.59017 | 0.009593 | 0.093551 |
| CG11617 | 170.3903 | -6.57674 | 2.543886 | -2.58531 | 0.009729 | 0.094764 |
| CG15211 | 818.8386 | -3.13077 | 1.212832 | -2.58137 | 0.009841 | 0.095736 |
| CG16978 | 768.3719 | 4.942168 | 1.916159 | 2.579205 | 0.009903 | 0.096106 |
| lncRNA:CR | 335.2464 | 6.305685 | 2.444594 | 2.57944 | 0.009896 | 0.096106 |
| CG15522 | 284.9283 | 5.299251 | 2.057541 | 2.575526 | 0.010009 | 0.097017 |
| CG9154 | 488.8327 | -5.61008 | 2.181106 | -2.57212 | 0.010108 | 0.09774 |
| lncRNA:CR | 216.8461 | 7.120063 | 2.768075 | 2.572207 | 0.010105 | 0.09774 |
| sll | 558.3221 | 3.910259 | 1.520714 | 2.571332 | 0.010131 | 0.097846 |
| CG34126 | 2524.589 | -1.84235 | 0.717289 | -2.56848 | 0.010214 | 0.098535 |
| Art7 | 244.0741 | -6.78229 | 2.646764 | -2.56248 | 0.010393 | 0.100134 |
| CG15651 | 1503.816 | -3.40602 | 1.330986 | -2.55902 | 0.010497 | 0.101016 |
| CG32473 | 3123.402 | -1.80668 | 0.706577 | -2.55694 | 0.01056 | 0.101256 |
| CG13605 | 626.9189 | 4.171893 | 1.631359 | 2.557311 | 0.010548 | 0.101256 |
| GXIVsPLA2 | 1344.602 | 3.443096 | 1.346332 | 2.557391 | 0.010546 | 0.101256 |
| CG4936 | 343.1679 | -5.00305 | 1.960115 | -2.55242 | 0.010698 | 0.102457 |
| CR43087 | 192.5759 | 5.829759 | 2.287582 | 2.548437 | 0.010821 | 0.103512 |
| CG5348 | 602.3145 | 4.484237 | 1.761944 | 2.545051 | 0.010926 | 0.104149 |
| Vmat | 46892.74 | -1.12892 | 0.443507 | -2.54544 | 0.010914 | 0.104149 |
| Rint1 | 916.7615 | 4.017065 | 1.578233 | 2.545293 | 0.010919 | 0.104149 |
| Glos | 262.7868 | 6.307239 | 2.480916 | 2.542303 | 0.011012 | 0.104847 |
| CG3149 | 1115.484 | -3.94652 | 1.553025 | -2.54118 | 0.011048 | 0.105059 |
| lace | 2171.316 | 2.022117 | 0.796426 | 2.538988 | 0.011117 | 0.105531 |
| CG5721 | 303.1778 | -5.58584 | 2.200367 | -2.5386 | 0.01113 | 0.105531 |
| lncRNA:CR | 144.2179 | 8.155998 | 3.213081 | 2.538373 | 0.011137 | 0.105531 |
| Obp44a | 102158.4 | -1.1526 | 0.454347 | -2.53683 | 0.011186 | 0.105873 |
| CG12375 | 1815.905 | -3.4134 | 1.3458 | -2.53633 | 0.011202 | 0.105897 |

| | | | | | | |
|---|---|---|---|---|---|---|
| GstE8 | 403.9929 | -6.41212 | 2.530209 | -2.53423 | 0.01127 | 0.10624 |
| Nup98-96 | 323.0065 | 4.55253 | 1.796018 | 2.534791 | 0.011251 | 0.10624 |
| CG42346 | 3096.181 | 2.124174 | 0.838279 | 2.533969 | 0.011278 | 0.10624 |
| Pex13 | 1152.871 | -2.45349 | 0.968647 | -2.53291 | 0.011312 | 0.106437 |
| Cyp6t1 | 1097.167 | 3.50813 | 1.385567 | 2.531909 | 0.011344 | 0.106616 |
| CG10731 | 359.0777 | 5.43658 | 2.147625 | 2.531438 | 0.01136 | 0.106634 |
| Spg7 | 1079.773 | 3.39865 | 1.344133 | 2.528508 | 0.011455 | 0.107403 |
| lncRNA:CR | 153.6518 | -8.29152 | 3.284419 | -2.5245 | 0.011586 | 0.108508 |
| deltaCOP | 1435.02 | 2.263154 | 0.896981 | 2.52308 | 0.011633 | 0.108821 |
| Tak1 | 816.6313 | -3.72735 | 1.477887 | -2.52208 | 0.011666 | 0.108876 |
| Vps29 | 1377.821 | 2.410049 | 0.955557 | 2.52214 | 0.011664 | 0.108876 |
| SrpRalpha | 858.2759 | -3.44519 | 1.366735 | -2.52075 | 0.011711 | 0.109163 |
| CG34354 | 2680.216 | -1.80744 | 0.717144 | -2.52032 | 0.011725 | 0.109167 |
| CG11906 | 775.0646 | 3.378429 | 1.341336 | 2.518704 | 0.011779 | 0.109505 |
| CheB74a | 159.1881 | -7.26945 | 2.886513 | -2.51842 | 0.011788 | 0.109505 |
| CG6834 | 1866.975 | -2.37717 | 0.94432 | -2.51734 | 0.011825 | 0.109716 |
| dgt6 | 441.5481 | 6.967658 | 2.76878 | 2.516508 | 0.011852 | 0.109847 |
| sgll | 2009.447 | 2.523735 | 1.004049 | 2.513557 | 0.011952 | 0.110643 |
| CG2846 | 793.938 | -3.8732 | 1.541716 | -2.51227 | 0.011996 | 0.110874 |
| Vps36 | 450.5631 | -5.56995 | 2.217383 | -2.51195 | 0.012007 | 0.110874 |
| Sec8 | 497.6353 | -3.89612 | 1.551248 | -2.5116 | 0.012018 | 0.110874 |
| ZnT77C | 423.4169 | -4.39462 | 1.750662 | -2.51026 | 0.012064 | 0.111167 |
| CG42797 | 1152.239 | 3.106843 | 1.238169 | 2.509224 | 0.0121 | 0.111367 |
| Ccdc85 | 148.1112 | 8.866447 | 3.539812 | 2.504779 | 0.012253 | 0.112647 |
| CG14712 | 1257.243 | -3.50089 | 1.400067 | -2.50052 | 0.012401 | 0.113363 |
| Nepl17 | 758.5318 | 4.592304 | 1.836455 | 2.500634 | 0.012397 | 0.113363 |
| lncRNA:CR | 75.99087 | -9.6159 | 3.844241 | -2.50138 | 0.012371 | 0.113363 |
| Lkr | 903.8137 | 4.15584 | 1.661877 | 2.50069 | 0.012395 | 0.113363 |
| insv | 83.21657 | 9.614721 | 3.844211 | 2.501091 | 0.012381 | 0.113363 |
| Pa1 | 1738.868 | 3.050449 | 1.22108 | 2.498157 | 0.012484 | 0.11399 |
| CG7168 | 118.135 | -8.79899 | 3.522877 | -2.49767 | 0.012501 | 0.114016 |
| pyr | 123.0127 | -8.60232 | 3.448279 | -2.49467 | 0.012607 | 0.114855 |
| CG4230 | 7765.958 | 2.019711 | 0.810514 | 2.491889 | 0.012707 | 0.115627 |
| MESR4 | 4174.953 | -1.62831 | 0.653571 | -2.49141 | 0.012724 | 0.115641 |
| CG34396 | 146.3505 | 6.395174 | 2.567269 | 2.491042 | 0.012737 | 0.115641 |
| wek | 336.3106 | -3.96231 | 1.593915 | -2.4859 | 0.012922 | 0.11706 |
| CG31323 | 779.6546 | -3.07103 | 1.235182 | -2.4863 | 0.012908 | 0.11706 |
| CG3356 | 1262.201 | 2.747659 | 1.106576 | 2.483027 | 0.013027 | 0.117876 |
| CG31612 | 91.03427 | 9.43331 | 3.800644 | 2.48203 | 0.013064 | 0.118073 |
| CG2662 | 375.5819 | 4.887669 | 1.971976 | 2.478564 | 0.013191 | 0.119092 |
| CG5973 | 677.0084 | -3.65686 | 1.477222 | -2.4755 | 0.013305 | 0.119984 |
| Lectin-gal( | 102.629 | -9.65529 | 3.905256 | -2.47238 | 0.013422 | 0.120899 |
| CG9839 | 875.7462 | 2.520811 | 1.019809 | 2.471846 | 0.013442 | 0.120946 |
| CG17068 | 222.0788 | -5.96187 | 2.413679 | -2.47004 | 0.01351 | 0.121423 |
| Vps25 | 1856.523 | 1.888768 | 0.764805 | 2.469608 | 0.013526 | 0.121432 |
| Atet | 3467.676 | 1.79726 | 0.72799 | 2.468797 | 0.013557 | 0.121572 |
| CG8132 | 152.1612 | 7.778855 | 3.152425 | 2.467578 | 0.013603 | 0.121851 |
| Teh4 | 1293.55 | 3.781414 | 1.533492 | 2.465884 | 0.013668 | 0.122292 |
| hpo | 621.9195 | 3.611933 | 1.465391 | 2.464825 | 0.013708 | 0.122409 |
| CG3831 | 471.7593 | 5.047277 | 2.04779 | 2.464743 | 0.013711 | 0.122409 |

| | | | | | | |
|---|---|---|---|---|---|---|
| CG9664 | 1307.466 | -3.22268 | 1.307988 | -2.46385 | 0.013745 | 0.122579 |
| CG13284 | 236.0507 | -7.92849 | 3.220403 | -2.46196 | 0.013818 | 0.12309 |
| lectin-37Da | 94.2488 | -9.61637 | 3.907623 | -2.46093 | 0.013858 | 0.123308 |
| GABA-B-R1 | 5039.632 | 1.329835 | 0.540497 | 2.460396 | 0.013878 | 0.123353 |
| CG2199 | 459.4961 | 4.119637 | 1.675919 | 2.458136 | 0.013966 | 0.123995 |
| Ptr | 2863.427 | -1.62143 | 0.66013 | -2.45623 | 0.01404 | 0.124518 |
| CG4133 | 406.4583 | 3.880542 | 1.581874 | 2.453129 | 0.014162 | 0.125457 |
| cry | 1694.471 | -2.91271 | 1.188264 | -2.45123 | 0.014237 | 0.125842 |
| msd5 | 604.1853 | 4.554141 | 1.857853 | 2.451292 | 0.014234 | 0.125842 |
| CG13641 | 145.8122 | -8.1817 | 3.343345 | -2.44716 | 0.014399 | 0.127133 |
| RhoGAP15 | 341.7353 | -4.48878 | 1.835045 | -2.44614 | 0.01444 | 0.127213 |
| rept | 180.1645 | 6.481443 | 2.649486 | 2.446302 | 0.014433 | 0.127213 |
| Nep2 | 874.457 | 3.549801 | 1.451942 | 2.444865 | 0.014491 | 0.127479 |
| mld | 2778.85 | -1.62989 | 0.666732 | -2.4446 | 0.014501 | 0.127479 |
| CG1575 | 271.1246 | -6.29201 | 2.575707 | -2.44283 | 0.014573 | 0.127965 |
| Dcr-1 | 157.1839 | 5.916817 | 2.425001 | 2.439923 | 0.01469 | 0.12864 |
| oys | 3934.841 | -1.77069 | 0.725624 | -2.44023 | 0.014678 | 0.12864 |
| Alh | 2939.405 | -1.74438 | 0.7151 | -2.43935 | 0.014714 | 0.12864 |
| CG8209 | 461.013 | -4.17634 | 1.71188 | -2.43962 | 0.014703 | 0.12864 |
| CG32147 | 618.4242 | -3.92536 | 1.609859 | -2.43833 | 0.014755 | 0.128865 |
| Uba1 | 2787.384 | -2.3031 | 0.945605 | -2.43558 | 0.014868 | 0.129707 |
| Pbp95 | 341.825 | 6.611232 | 2.715837 | 2.434326 | 0.01492 | 0.130016 |
| Patsas | 899.3449 | -2.6101 | 1.072657 | -2.4333 | 0.014962 | 0.130242 |
| CG5390 | 584.6631 | -4.20635 | 1.729537 | -2.43207 | 0.015013 | 0.130545 |
| egr | 1191.767 | -2.48694 | 1.025141 | -2.42595 | 0.015269 | 0.132565 |
| Fitm | 383.3514 | -4.41051 | 1.818228 | -2.42572 | 0.015278 | 0.132565 |
| lncRNA:CR | 81.43331 | -9.46333 | 3.907749 | -2.42168 | 0.015449 | 0.133901 |
| TwdlG | 1325.817 | -2.81545 | 1.164374 | -2.418 | 0.015606 | 0.135119 |
| Stam | 699.4173 | 2.802851 | 1.159838 | 2.416589 | 0.015667 | 0.135497 |
| trbd | 1463.228 | -3.30138 | 1.367707 | -2.4138 | 0.015787 | 0.13639 |
| mt:ND4L | 2113.996 | 1.771671 | 0.734235 | 2.412948 | 0.015824 | 0.136513 |
| CG32591 | 68.50143 | -9.42801 | 3.907673 | -2.41269 | 0.015835 | 0.136513 |
| CG12379 | 92.22378 | -8.83953 | 3.664989 | -2.41188 | 0.01587 | 0.136668 |
| lncRNA:CR | 841.2904 | 3.546801 | 1.470985 | 2.411174 | 0.015901 | 0.136788 |
| L2HGDH | 1038.289 | 2.733135 | 1.133954 | 2.410269 | 0.015941 | 0.136981 |
| Fkbp59 | 2166.675 | -2.85251 | 1.183809 | -2.40961 | 0.01597 | 0.137083 |
| CG43347 | 4578.766 | 1.417856 | 0.588641 | 2.408693 | 0.01601 | 0.13728 |
| Es2 | 241.6902 | -5.37239 | 2.234152 | -2.40467 | 0.016187 | 0.138505 |
| Pop2 | 4302.467 | 1.504379 | 0.625558 | 2.404859 | 0.016179 | 0.138505 |
| CG13663 | 1276.561 | -3.45557 | 1.438546 | -2.40213 | 0.0163 | 0.139175 |
| eas | 2981.266 | -1.73189 | 0.720899 | -2.40241 | 0.016287 | 0.139175 |
| mRF1 | 422.1329 | 5.279972 | 2.198602 | 2.401513 | 0.016327 | 0.139221 |
| BORCS6 | 378.519 | 4.74101 | 1.97441 | 2.401228 | 0.01634 | 0.139221 |
| lncRNA:CR | 105.5176 | -7.6655 | 3.194117 | -2.39988 | 0.0164 | 0.139586 |
| Tsp26A | 1808.21 | 2.650644 | 1.105088 | 2.398581 | 0.016459 | 0.139638 |
| ea | 706.8779 | -3.94402 | 1.643803 | -2.39933 | 0.016425 | 0.139638 |
| CG7546 | 1732.132 | -1.89438 | 0.789743 | -2.39873 | 0.016452 | 0.139638 |
| elg1 | 75.70688 | 8.704242 | 3.631228 | 2.397052 | 0.016528 | 0.140074 |
| lncRNA:CR | 309.9374 | 6.503714 | 2.71468 | 2.395757 | 0.016586 | 0.140422 |
| CG5611 | 174.0861 | -6.44903 | 2.695543 | -2.39248 | 0.016735 | 0.141384 |

| | | | | | | |
|---|---|---|---|---|---|---|
| lncRNA:CR | 129.9206 | 8.29753 | 3.46807 | 2.39255 | 0.016732 | 0.141384 |
| Bap55 | 1169.551 | -2.18943 | 0.915421 | -2.39172 | 0.01677 | 0.141528 |
| CG9109 | 176.1035 | 6.686191 | 2.79649 | 2.390922 | 0.016806 | 0.141687 |
| RpI12 | 173.6397 | 5.993842 | 2.510299 | 2.387701 | 0.016954 | 0.142785 |
| Sik2 | 1541.669 | -3.1336 | 1.314035 | -2.38472 | 0.017092 | 0.143798 |
| Cyp6a22 | 641.0827 | -3.35958 | 1.409554 | -2.38344 | 0.017152 | 0.144148 |
| CG11407 | 12694.09 | -1.31438 | 0.553191 | -2.376 | 0.017502 | 0.146521 |
| CG6145 | 1026.99 | 2.275314 | 0.957674 | 2.375876 | 0.017507 | 0.146521 |
| lncRNA:CR | 164.7585 | -8.22063 | 3.459887 | -2.37598 | 0.017502 | 0.146521 |
| CG12992 | 219.2428 | 5.684737 | 2.392524 | 2.376042 | 0.017499 | 0.146521 |
| smo | 117.5145 | 8.081426 | 3.408184 | 2.371182 | 0.017731 | 0.147933 |
| nwk | 7243.603 | 1.212543 | 0.511338 | 2.371312 | 0.017725 | 0.147933 |
| Syx16 | 416.22 | -3.94091 | 1.661689 | -2.37163 | 0.01771 | 0.147933 |
| CG14450 | 986.6928 | -2.95383 | 1.246145 | -2.37037 | 0.01777 | 0.148021 |
| Sgsh | 1638.439 | -3.15921 | 1.332892 | -2.37019 | 0.017779 | 0.148021 |
| CG14561 | 177.7163 | 7.982514 | 3.368934 | 2.369448 | 0.017815 | 0.148166 |
| lncRNA:CR | 95.35342 | 9.250206 | 3.908098 | 2.366933 | 0.017936 | 0.149022 |
| AdSL | 323.6056 | -5.47646 | 2.315089 | -2.36555 | 0.018003 | 0.149426 |
| Sf3a1 | 203.1351 | 4.39143 | 1.856853 | 2.364985 | 0.018031 | 0.149499 |
| CG11755 | 104.0804 | -7.55492 | 3.195825 | -2.364 | 0.018079 | 0.149741 |
| Spn55B | 116.9008 | -8.17721 | 3.459609 | -2.36362 | 0.018097 | 0.149741 |
| CG2921 | 1543.01 | -2.18045 | 0.922737 | -2.36303 | 0.018126 | 0.149828 |
| Pmm2 | 500.6458 | -3.05144 | 1.291547 | -2.36262 | 0.018146 | 0.149836 |
| Cyp6a13 | 3515.743 | -1.50121 | 0.635563 | -2.36201 | 0.018176 | 0.149929 |
| TfIIB | 2254.315 | 2.109657 | 0.893555 | 2.360969 | 0.018227 | 0.150197 |
| Rpb11 | 671.4262 | -3.22433 | 1.366842 | -2.35897 | 0.018326 | 0.150855 |
| CG5916 | 1104.716 | 3.329097 | 1.411734 | 2.358161 | 0.018366 | 0.151028 |
| CG8230 | 3013.541 | 2.274999 | 0.96497 | 2.357586 | 0.018394 | 0.151107 |
| Rgk3 | 2192.08 | -1.8605 | 0.78958 | -2.35632 | 0.018457 | 0.151467 |
| CG1492 | 321.3623 | 5.884543 | 2.498096 | 2.355611 | 0.018492 | 0.151603 |
| CG44085 | 93.3906 | 7.232464 | 3.071906 | 2.354389 | 0.018553 | 0.151947 |
| CG8569 | 610.7515 | 4.306007 | 1.831242 | 2.351414 | 0.018702 | 0.153011 |
| vtd | 7050.009 | 1.238424 | 0.527564 | 2.347438 | 0.018903 | 0.154497 |
| yata | 1402.589 | -2.16019 | 0.920526 | -2.34669 | 0.018941 | 0.154648 |
| Vps37B | 663.3752 | 3.977523 | 1.695936 | 2.345326 | 0.01901 | 0.155059 |
| Scp1 | 242.6894 | -5.90425 | 2.518371 | -2.34447 | 0.019054 | 0.155105 |
| bou | 570.626 | 3.580563 | 1.527246 | 2.344457 | 0.019055 | 0.155105 |
| Ugt35A1 | 789.5732 | -3.68262 | 1.571696 | -2.34308 | 0.019125 | 0.155362 |
| lncRNA:CR | 77.25779 | 9.156098 | 3.907561 | 2.343174 | 0.01912 | 0.155362 |
| Ugt35C1 | 3445.945 | 1.596313 | 0.681514 | 2.342306 | 0.019165 | 0.155529 |
| asRNA:CR4 | 1262.29 | -3.49394 | 1.492283 | -2.34134 | 0.019215 | 0.155774 |
| CG3368 | 963.4852 | -3.40534 | 1.454992 | -2.34046 | 0.01926 | 0.155986 |
| CG1236 | 412.7891 | -4.98633 | 2.132062 | -2.33874 | 0.019349 | 0.156548 |
| HemK1 | 3053.93 | -1.47726 | 0.632041 | -2.33729 | 0.019424 | 0.156996 |
| Adgf-D | 3259.049 | -1.60497 | 0.686966 | -2.33631 | 0.019475 | 0.157249 |
| lncRNA:CR | 251.4618 | -6.08774 | 2.60752 | -2.33469 | 0.01956 | 0.157776 |
| CG31125 | 500.4991 | 3.876234 | 1.661947 | 2.332344 | 0.019683 | 0.158607 |
| Nup44A | 474.221 | -3.99704 | 1.715451 | -2.33002 | 0.019805 | 0.159273 |
| Gbs-70E | 192.4575 | -5.09633 | 2.186924 | -2.33037 | 0.019787 | 0.159273 |
| Hat1 | 830.825 | -3.34964 | 1.439078 | -2.32763 | 0.019932 | 0.159694 |

| | | | | | | |
|---|---|---|---|---|---|---|
| CG3430 | 1082.061 | -3.10055 | 1.331527 | -2.32857 | 0.019882 | 0.159694 |
| ppan | 151.2999 | -6.73352 | 2.892984 | -2.32753 | 0.019937 | 0.159694 |
| CG8237 | 521.6492 | 4.131274 | 1.774671 | 2.32791 | 0.019917 | 0.159694 |
| OdsH | 96.58521 | 9.083209 | 3.903309 | 2.327053 | 0.019962 | 0.159739 |
| DnaJ-H | 4848.167 | 1.735996 | 0.746356 | 2.325964 | 0.02002 | 0.159885 |
| Wnt4 | 614.4947 | 3.872985 | 1.665001 | 2.326116 | 0.020012 | 0.159885 |
| Orc4 | 313.029 | -6.09476 | 2.628698 | -2.31855 | 0.02042 | 0.16291 |
| Task6 | 2805.442 | 1.615557 | 0.698216 | 2.313834 | 0.020677 | 0.164553 |
| U4-U6-60K | 136.2326 | -7.81623 | 3.377703 | -2.31407 | 0.020664 | 0.164553 |
| Agpat3 | 4132.993 | 2.114219 | 0.913803 | 2.313649 | 0.020687 | 0.164553 |
| CG31648 | 1524.647 | 2.948734 | 1.276266 | 2.310438 | 0.020864 | 0.165796 |
| CG13796 | 244.7284 | 6.122127 | 2.650365 | 2.309918 | 0.020893 | 0.16586 |
| Fatp1 | 3355.519 | 1.50382 | 0.651448 | 2.308427 | 0.020975 | 0.166352 |
| Cyp4s3 | 297.2046 | -4.81472 | 2.088024 | -2.30588 | 0.021118 | 0.167314 |
| mei-W68 | 98.58035 | -8.96446 | 3.889801 | -2.30461 | 0.021189 | 0.167712 |
| Atg18b | 2694.032 | -1.57755 | 0.685081 | -2.30273 | 0.021294 | 0.168382 |
| Ctl1 | 1937.828 | 2.540598 | 1.103842 | 2.301597 | 0.021358 | 0.168719 |
| CG7745 | 997.6966 | 2.92938 | 1.273331 | 2.300564 | 0.021416 | 0.169014 |
| Obp57b | 161.8818 | 8.158455 | 3.546959 | 2.300127 | 0.021441 | 0.169043 |
| Sep-01 | 1611.9 | -3.43784 | 1.49542 | -2.29892 | 0.02151 | 0.169418 |
| Gnpnat | 448.705 | 4.052035 | 1.765021 | 2.295743 | 0.021691 | 0.170675 |
| NELF-B | 669.892 | -3.13863 | 1.367593 | -2.295 | 0.021733 | 0.170843 |
| CG15233 | 262.5973 | 7.104894 | 3.09654 | 2.294462 | 0.021764 | 0.170918 |
| Usp30 | 2846.133 | -1.75035 | 0.763463 | -2.29264 | 0.021869 | 0.171404 |
| Prip | 623.0634 | 5.55882 | 2.424518 | 2.292753 | 0.021862 | 0.171404 |
| CG9684 | 191.3764 | -5.42386 | 2.368799 | -2.28971 | 0.022038 | 0.172564 |
| CG11777 | 172.4942 | -7.18477 | 3.138517 | -2.28922 | 0.022066 | 0.172616 |
| Krn | 2152.794 | -3.42936 | 1.498338 | -2.28877 | 0.022092 | 0.172652 |
| bocks | 839.427 | -3.03823 | 1.328785 | -2.28647 | 0.022227 | 0.173532 |
| Tim17b1 | 364.8492 | 4.403774 | 1.927407 | 2.284817 | 0.022324 | 0.173843 |
| ouib | 301.6094 | 5.616679 | 2.458212 | 2.284864 | 0.022321 | 0.173843 |
| dunk | 90.44686 | 8.92958 | 3.908454 | 2.284684 | 0.022331 | 0.173843 |
| CG17597 | 186.4513 | -6.79771 | 2.976413 | -2.28386 | 0.02238 | 0.174051 |
| SCCRO4 | 1886.412 | -1.96453 | 0.860638 | -2.28264 | 0.022451 | 0.174438 |
| CG3756 | 407.1122 | -3.70843 | 1.625547 | -2.28134 | 0.022528 | 0.174697 |
| Jarid2 | 4113.363 | -1.43522 | 0.629044 | -2.28158 | 0.022514 | 0.174697 |
| dpr2 | 1574.85 | 2.186625 | 0.959132 | 2.279797 | 0.02262 | 0.175237 |
| CG30105 | 90.62655 | -8.81139 | 3.866668 | -2.27881 | 0.022679 | 0.175355 |
| lncRNA:CR | 181.9488 | 5.254548 | 2.30575 | 2.278889 | 0.022674 | 0.175355 |
| CG10747 | 349.5569 | 4.214348 | 1.84968 | 2.27842 | 0.022702 | 0.175364 |
| lncRNA:CR | 76.06034 | -8.90418 | 3.909009 | -2.27786 | 0.022735 | 0.175452 |
| CG3061 | 2387.194 | -2.47207 | 1.087027 | -2.27415 | 0.022957 | 0.176655 |
| CG6006 | 3714.915 | -1.68404 | 0.740513 | -2.27416 | 0.022956 | 0.176655 |
| cag | 492.3129 | -4.48653 | 1.972296 | -2.27477 | 0.022919 | 0.176655 |
| CG6959 | 1628.584 | 2.702251 | 1.188949 | 2.272806 | 0.023038 | 0.17677 |
| CG6654 | 722.3553 | 3.552969 | 1.562892 | 2.273331 | 0.023006 | 0.17677 |
| CG3045 | 114.6378 | -6.96571 | 3.064436 | -2.27308 | 0.023021 | 0.17677 |
| CG3335 | 391.5061 | -4.48033 | 1.972115 | -2.27184 | 0.023096 | 0.177048 |
| Arl6IP1 | 3206.035 | -1.69926 | 0.748619 | -2.26986 | 0.023216 | 0.17763 |
| Xpd | 289.5167 | 4.921898 | 2.168366 | 2.269865 | 0.023216 | 0.17763 |

| | | | | | | |
|---|---|---|---|---|---|---|
| goe | 3092.98 | -1.63176 | 0.719216 | -2.26881 | 0.02328 | 0.177779 |
| CG8108 | 3666.338 | -1.39711 | 0.615785 | -2.26882 | 0.023279 | 0.177779 |
| Mys45A | 169.3572 | -6.51632 | 2.875328 | -2.26629 | 0.023434 | 0.178613 |
| CG3163 | 599.8545 | 3.592113 | 1.584883 | 2.266484 | 0.023422 | 0.178613 |
| CG32236 | 125.0404 | 6.052232 | 2.671757 | 2.265263 | 0.023497 | 0.178752 |
| CG6761 | 160.0972 | -8.04656 | 3.551857 | -2.26545 | 0.023485 | 0.178752 |
| CG8520 | 172.6011 | 4.761085 | 2.104732 | 2.262086 | 0.023692 | 0.180069 |
| Ddx1 | 72.83565 | 8.829581 | 3.908548 | 2.259044 | 0.023881 | 0.180988 |
| Tsp42Ek | 556.7387 | 3.171388 | 1.403473 | 2.259672 | 0.023842 | 0.180988 |
| Muc96D | 94.20904 | 8.581291 | 3.798395 | 2.259189 | 0.023872 | 0.180988 |
| muskelin | 260.6175 | 4.066176 | 1.80169 | 2.256867 | 0.024016 | 0.181033 |
| dpr4 | 833.4687 | -2.57718 | 1.1417 | -2.25731 | 0.023988 | 0.181033 |
| su(r) | 641.6428 | -2.83286 | 1.254907 | -2.25742 | 0.023982 | 0.181033 |
| magu | 873.8997 | 2.602194 | 1.153057 | 2.256779 | 0.024022 | 0.181033 |
| CG34377 | 717.9036 | 3.646164 | 1.614645 | 2.258183 | 0.023934 | 0.181033 |
| CG32655 | 95.59628 | -6.79193 | 3.009195 | -2.25706 | 0.024004 | 0.181033 |
| metl | 1479.531 | -2.68994 | 1.192196 | -2.25629 | 0.024053 | 0.181094 |
| bru1 | 970.5266 | -2.33185 | 1.034273 | -2.25458 | 0.02416 | 0.18173 |
| CG4078 | 495.8875 | -4.22007 | 1.872166 | -2.25411 | 0.024189 | 0.181782 |
| CG30108 | 72.31162 | 8.805623 | 3.908578 | 2.252897 | 0.024266 | 0.182185 |
| CG13293 | 1125.94 | 2.411319 | 1.070607 | 2.252291 | 0.024304 | 0.182302 |
| CG9360 | 252.9369 | 3.985997 | 1.770633 | 2.25117 | 0.024375 | 0.182663 |
| mi | 133.5379 | -8.39701 | 3.730968 | -2.25062 | 0.024409 | 0.182751 |
| CycA | 673.3977 | -4.15369 | 1.846572 | -2.2494 | 0.024487 | 0.183161 |
| Not11 | 928.783 | -3.27375 | 1.45639 | -2.24785 | 0.024585 | 0.183727 |
| dve | 362.2043 | -5.98479 | 2.664259 | -2.24632 | 0.024683 | 0.184115 |
| Coq5 | 518.1083 | 4.506086 | 2.005912 | 2.246402 | 0.024678 | 0.184115 |
| IntS14 | 497.4355 | 4.24637 | 1.893583 | 2.242505 | 0.024929 | 0.185774 |
| AnxB11 | 348.5143 | -4.97672 | 2.221542 | -2.24021 | 0.025077 | 0.186706 |
| Cyp6d4 | 424.1692 | 5.88339 | 2.629338 | 2.237593 | 0.025248 | 0.187801 |
| Irp-1A | 669.1596 | 3.327497 | 1.488327 | 2.23573 | 0.025369 | 0.188533 |
| CRAT | 641.6437 | 3.030592 | 1.356936 | 2.233409 | 0.025522 | 0.189491 |
| Ilk | 1125.238 | 2.694077 | 1.206891 | 2.232245 | 0.025599 | 0.189885 |
| Hr38 | 8674.528 | 1.49946 | 0.672027 | 2.231248 | 0.025665 | 0.190199 |
| CG4858 | 2313.593 | -3.21591 | 1.442394 | -2.22957 | 0.025776 | 0.190673 |
| Fmo-2 | 162.7355 | -5.60379 | 2.513235 | -2.22971 | 0.025766 | 0.190673 |
| az2 | 557.2212 | -3.8992 | 1.749401 | -2.22888 | 0.025822 | 0.190836 |
| Rexo5 | 2284.079 | -1.85182 | 0.83127 | -2.2277 | 0.025901 | 0.19124 |
| Arp6 | 129.2275 | 6.461796 | 2.905756 | 2.223792 | 0.026162 | 0.192996 |
| Cog1 | 1003.92 | 3.331914 | 1.499374 | 2.222203 | 0.02627 | 0.193522 |
| pll | 67.57122 | 8.682076 | 3.907289 | 2.222021 | 0.026282 | 0.193522 |
| Hs3st-A | 6238.857 | 1.20447 | 0.542408 | 2.220598 | 0.026378 | 0.194053 |
| CG6724 | 639.4748 | -4.644 | 2.094537 | -2.2172 | 0.02661 | 0.195243 |
| comt | 11570.28 | 0.990628 | 0.446675 | 2.21778 | 0.02657 | 0.195243 |
| comm | 966.2716 | -2.23977 | 1.010205 | -2.21715 | 0.026613 | 0.195243 |
| Nrk | 438.5134 | 4.025555 | 1.816021 | 2.21669 | 0.026644 | 0.195294 |
| Mettl3 | 305.9721 | 6.599848 | 2.978732 | 2.215656 | 0.026715 | 0.195456 |
| CG13531 | 113.806 | 6.776839 | 3.058139 | 2.216001 | 0.026691 | 0.195456 |
| CG7987 | 1589.511 | 2.159252 | 0.975543 | 2.213384 | 0.026871 | 0.196419 |
| TTLL4A | 4575.528 | 1.256983 | 0.568034 | 2.212867 | 0.026907 | 0.196496 |

| | | | | | | |
|---|---|---|---|---|---|---|
| ddbt | 85.5661 | 7.519242 | 3.398495 | 2.212521 | 0.026931 | 0.196496 |
| MFS3 | 177.7059 | 7.547372 | 3.412358 | 2.211776 | 0.026982 | 0.196693 |
| Fs | 428.3627 | 5.392948 | 2.442422 | 2.208032 | 0.027242 | 0.198407 |
| hbs | 134.4268 | -5.4637 | 2.475226 | -2.20736 | 0.027289 | 0.198571 |
| beat-IIIb | 3547.364 | 1.371853 | 0.621891 | 2.205939 | 0.027388 | 0.199111 |
| CG15083 | 1516.16 | -2.12085 | 0.961874 | -2.20491 | 0.02746 | 0.199455 |
| lncRNA:CR | 62.99831 | 8.614641 | 3.908833 | 2.203891 | 0.027532 | 0.199794 |
| ADPS | 1074.743 | 3.90416 | 1.772104 | 2.203121 | 0.027586 | 0.200007 |
| pdgy | 8034.116 | -1.42279 | 0.645933 | -2.20269 | 0.027617 | 0.200047 |
| CG14445 | 160.5348 | -6.8332 | 3.107129 | -2.1992 | 0.027864 | 0.201521 |
| subdued | 2763.808 | 2.159889 | 0.982483 | 2.198399 | 0.027921 | 0.201521 |
| eIF3i | 4522.689 | -1.51623 | 0.689608 | -2.19869 | 0.0279 | 0.201521 |
| CG6175 | 377.5301 | -3.6274 | 1.649795 | -2.1987 | 0.027899 | 0.201521 |
| CG13123 | 92.26905 | 7.471618 | 3.402195 | 2.196117 | 0.028084 | 0.202505 |
| CG2767 | 189.8726 | -5.49712 | 2.503489 | -2.19578 | 0.028107 | 0.202505 |
| Apc | 3846.628 | 1.4109 | 0.64279 | 2.194963 | 0.028166 | 0.202746 |
| Mkk4 | 4046.403 | 1.483089 | 0.676041 | 2.193785 | 0.028251 | 0.203173 |
| Arp5 | 776.4538 | -3.21186 | 1.465594 | -2.19151 | 0.028415 | 0.203443 |
| CG11771 | 789.0679 | -3.78275 | 1.725151 | -2.1927 | 0.028329 | 0.203443 |
| gwl | 363.4277 | -4.66799 | 2.130015 | -2.19153 | 0.028413 | 0.203443 |
| CG42554 | 476.8771 | 3.532079 | 1.61151 | 2.191783 | 0.028395 | 0.203443 |
| CG14229 | 2056.148 | 1.797802 | 0.820211 | 2.191879 | 0.028388 | 0.203443 |
| SMSr | 1285.643 | -2.19867 | 1.003927 | -2.19007 | 0.028519 | 0.204007 |
| CG7791 | 2091.21 | 1.796623 | 0.820752 | 2.188996 | 0.028597 | 0.204382 |
| Nach | 89.08091 | 8.524585 | 3.895767 | 2.188166 | 0.028658 | 0.204631 |
| ebo | 722.832 | -3.27896 | 1.499209 | -2.18713 | 0.028733 | 0.204988 |
| CG11438 | 281.1789 | 6.176511 | 2.82495 | 2.186414 | 0.028785 | 0.205179 |
| CG31694 | 1531.125 | 2.170299 | 0.993347 | 2.184836 | 0.028901 | 0.205819 |
| CG9784 | 1061.758 | -2.4323 | 1.114607 | -2.1822 | 0.029095 | 0.207015 |
| CG31937 | 192.944 | -5.56597 | 2.552066 | -2.18096 | 0.029186 | 0.207482 |
| CG5013 | 75.18197 | -7.81035 | 3.583116 | -2.17976 | 0.029275 | 0.207746 |
| Rpb5 | 621.0788 | 3.40564 | 1.562317 | 2.179865 | 0.029267 | 0.207746 |
| Naam | 63.18637 | 8.391702 | 3.853135 | 2.177889 | 0.029414 | 0.20855 |
| Dl | 125.9047 | -6.54286 | 3.006586 | -2.17618 | 0.029542 | 0.209271 |
| CG44008 | 169.0836 | 5.431971 | 2.498029 | 2.174503 | 0.029667 | 0.209974 |
| CG2701 | 696.7191 | 2.870954 | 1.320746 | 2.173736 | 0.029725 | 0.210166 |
| CadN2 | 1087.193 | 2.893043 | 1.331157 | 2.17333 | 0.029755 | 0.210166 |
| CG9948 | 548.9503 | 3.592292 | 1.653075 | 2.173097 | 0.029773 | 0.210166 |
| Dcp-1 | 253.259 | -5.02236 | 2.31202 | -2.17228 | 0.029834 | 0.210413 |
| CG11902 | 531.6454 | 3.45632 | 1.591844 | 2.171267 | 0.029911 | 0.210419 |
| Cndp2 | 1815.714 | 2.189322 | 1.008113 | 2.171704 | 0.029878 | 0.210419 |
| CG32365 | 120.3512 | -6.3783 | 2.937644 | -2.17123 | 0.029914 | 0.210419 |
| Zif | 1276.869 | -2.81289 | 1.296383 | -2.1698 | 0.030022 | 0.210678 |
| CG6424 | 701.1973 | -2.43649 | 1.122652 | -2.1703 | 0.029984 | 0.210678 |
| CG9760 | 276.5509 | 4.734184 | 2.181951 | 2.169702 | 0.030029 | 0.210678 |
| mRpS33 | 1802.99 | 1.775182 | 0.819338 | 2.166604 | 0.030265 | 0.212146 |
| Lpt | 102.9718 | 5.476798 | 2.529265 | 2.165372 | 0.030359 | 0.212621 |
| Surf6 | 460.5218 | -4.66816 | 2.157707 | -2.16348 | 0.030504 | 0.213449 |
| CG8319 | 555.7655 | -3.70648 | 1.713526 | -2.16307 | 0.030536 | 0.213483 |
| Gclm | 2582.211 | -1.83139 | 0.847192 | -2.16172 | 0.03064 | 0.214027 |

| | | | | | | |
|---|---|---|---|---|---|---|
| CG11454 | 768.7772 | 3.342859 | 1.54697 | 2.160908 | 0.030702 | 0.214064 |
| barr | 109.7205 | 5.970649 | 2.763847 | 2.160267 | 0.030752 | 0.214064 |
| CG16908 | 201.1758 | -6.12852 | 2.836316 | -2.16073 | 0.030716 | 0.214064 |
| GNBP2 | 120.0015 | -6.59264 | 3.051481 | -2.16047 | 0.030736 | 0.214064 |
| PIP82 | 63.74152 | -8.30809 | 3.846828 | -2.15972 | 0.030794 | 0.214171 |
| Pal1 | 1652.901 | -2.48211 | 1.150006 | -2.15835 | 0.030901 | 0.214728 |
| CG18764 | 304.2054 | 4.783735 | 2.218164 | 2.15662 | 0.031035 | 0.215475 |
| CG4716 | 79.04047 | 8.394302 | 3.898249 | 2.153352 | 0.031291 | 0.216875 |
| Bap60 | 1382.455 | -3.06317 | 1.422464 | -2.15343 | 0.031285 | 0.216875 |
| PIG-H | 621.0078 | -3.42469 | 1.590668 | -2.15299 | 0.03132 | 0.216887 |
| Mms19 | 640.0486 | -3.15373 | 1.4659 | -2.15139 | 0.031445 | 0.217568 |
| CG8042 | 919.5113 | 2.915682 | 1.356154 | 2.149964 | 0.031558 | 0.218161 |
| CG4768 | 2212.34 | -1.59395 | 0.742216 | -2.14756 | 0.031749 | 0.219289 |
| hbn | 251.2863 | 3.99483 | 1.860895 | 2.146725 | 0.031815 | 0.219372 |
| ND-49 | 11393 | 1.399654 | 0.651985 | 2.146757 | 0.031813 | 0.219372 |
| X11L | 1344.943 | 2.944026 | 1.37168 | 2.146292 | 0.03185 | 0.219421 |
| CG8435 | 464.5807 | 3.32056 | 1.547561 | 2.145673 | 0.031899 | 0.219573 |
| CG17230 | 2089.011 | -2.22762 | 1.039401 | -2.14317 | 0.032099 | 0.22076 |
| Sec20 | 447.2717 | -4.25055 | 1.983924 | -2.14249 | 0.032154 | 0.220947 |
| CheB53a | 83.55624 | -8.16827 | 3.813624 | -2.14187 | 0.032204 | 0.221105 |
| Oseg5 | 132.0242 | 6.36288 | 2.971199 | 2.141519 | 0.032232 | 0.221107 |
| kcc | 17737.55 | 1.166676 | 0.545762 | 2.137703 | 0.032541 | 0.223034 |
| Lap1 | 1563.849 | 1.84182 | 0.861868 | 2.137008 | 0.032597 | 0.223231 |
| Diap1 | 4600.053 | -1.5179 | 0.71047 | -2.13647 | 0.032641 | 0.223337 |
| FIG4 | 155.7213 | 5.253981 | 2.460516 | 2.135316 | 0.032735 | 0.223793 |
| Tcs3 | 105.0606 | 6.800484 | 3.188357 | 2.132912 | 0.032932 | 0.224946 |
| CG15237 | 778.5703 | 3.129441 | 1.468688 | 2.130774 | 0.033108 | 0.225955 |
| Ggt-1 | 63.99092 | -8.25588 | 3.879545 | -2.12805 | 0.033333 | 0.227297 |
| Rpb8 | 860.6255 | 3.129477 | 1.472109 | 2.125846 | 0.033516 | 0.228353 |
| Urod | 1414.234 | -2.39557 | 1.127625 | -2.12444 | 0.033634 | 0.228528 |
| Gmd | 201.4693 | 4.85936 | 2.288748 | 2.123152 | 0.033741 | 0.228528 |
| jv | 469.8229 | -2.81341 | 1.32499 | -2.12334 | 0.033725 | 0.228528 |
| garz | 1469.966 | 2.66489 | 1.254349 | 2.124521 | 0.033627 | 0.228528 |
| lncRNA:CR | 80.37937 | -8.17469 | 3.849227 | -2.12372 | 0.033693 | 0.228528 |
| CngB | 660.0954 | 4.410523 | 2.076806 | 2.123704 | 0.033695 | 0.228528 |
| ergic53 | 1082.006 | -2.80368 | 1.320497 | -2.1232 | 0.033737 | 0.228528 |
| CG12942 | 122.4057 | -8.12737 | 3.829509 | -2.1223 | 0.033813 | 0.228645 |
| asRNA:CR4 | 194.4818 | -4.03051 | 1.899152 | -2.12227 | 0.033815 | 0.228645 |
| CG3651 | 135.5276 | -5.67296 | 2.674254 | -2.12132 | 0.033895 | 0.228945 |
| lute | 868.7817 | 3.072641 | 1.448635 | 2.12106 | 0.033917 | 0.228945 |
| Rpn2 | 2240.594 | -1.80137 | 0.850142 | -2.1189 | 0.034099 | 0.229596 |
| mesh | 306.9389 | 4.924656 | 2.323947 | 2.119091 | 0.034083 | 0.229596 |
| APP-BP1 | 2111.811 | 2.201601 | 1.039021 | 2.118919 | 0.034097 | 0.229596 |
| Fdh | 720.4438 | -3.18607 | 1.504653 | -2.11748 | 0.034219 | 0.230211 |
| CG14864 | 322.7109 | -3.50263 | 1.655902 | -2.11524 | 0.034409 | 0.230333 |
| CG14502 | 356.2138 | -3.41675 | 1.615086 | -2.11552 | 0.034385 | 0.230333 |
| CG14440 | 839.3655 | 2.517713 | 1.189671 | 2.11631 | 0.034318 | 0.230333 |
| lncRNA:CR | 89.61315 | 8.226942 | 3.887637 | 2.116181 | 0.034329 | 0.230333 |
| CG7509 | 22827.27 | -0.90723 | 0.428727 | -2.1161 | 0.034336 | 0.230333 |
| CG14352 | 213.4933 | -5.7613 | 2.723327 | -2.11554 | 0.034384 | 0.230333 |

| | | | | | | |
|---|---|---|---|---|---|---|
| Xpc | 4124.812 | -2.08365 | 0.985342 | -2.11464 | 0.03446 | 0.230483 |
| asRNA:CR4 | 525.4796 | -3.39319 | 1.606408 | -2.11228 | 0.034662 | 0.231639 |
| CG3309 | 279.2528 | -5.24324 | 2.483492 | -2.11124 | 0.034752 | 0.232045 |
| Rab39 | 1240.972 | 2.484787 | 1.178564 | 2.108318 | 0.035003 | 0.233531 |
| CG5599 | 162.5673 | -6.40979 | 3.043684 | -2.10593 | 0.03521 | 0.234716 |
| lncRNA:CR | 319.4154 | -3.93912 | 1.870968 | -2.10539 | 0.035257 | 0.234832 |
| Myo95E | 688.0318 | 2.797884 | 1.329167 | 2.10499 | 0.035292 | 0.234871 |
| mip130 | 118.6368 | -6.92632 | 3.292182 | -2.10387 | 0.03539 | 0.235326 |
| MetRS | 1074.643 | -2.60502 | 1.238448 | -2.10346 | 0.035426 | 0.235372 |
| RanBP3 | 168.6788 | -6.31335 | 3.004047 | -2.10162 | 0.035587 | 0.236172 |
| snoRNA:gc | 85.6192 | -8.19269 | 3.89867 | -2.10141 | 0.035605 | 0.236172 |
| Ttc30 | 182.5491 | -6.20256 | 2.952857 | -2.10053 | 0.035682 | 0.236209 |
| Srlp | 1201.76 | 2.367055 | 1.126952 | 2.100404 | 0.035693 | 0.236209 |
| Nup58 | 100.5899 | -8.07774 | 3.845923 | -2.10034 | 0.035699 | 0.236209 |
| CG2915 | 955.9955 | -2.27197 | 1.083015 | -2.09782 | 0.035921 | 0.237384 |
| for | 577.1121 | -2.67783 | 1.276585 | -2.09765 | 0.035936 | 0.237384 |
| CG11504 | 845.0313 | 4.255597 | 2.029299 | 2.097078 | 0.035987 | 0.237524 |
| CG33331 | 374.4024 | 3.324712 | 1.585882 | 2.096443 | 0.036043 | 0.2377 |
| CG4752 | 513.9044 | -3.5494 | 1.694393 | -2.09479 | 0.03619 | 0.238471 |
| Spn42Dd | 174.12 | 7.216904 | 3.446476 | 2.093995 | 0.03626 | 0.238742 |
| CG2712 | 194.5193 | 6.15279 | 2.941224 | 2.091915 | 0.036446 | 0.239389 |
| Ppt2 | 815.0611 | -2.80757 | 1.342333 | -2.09156 | 0.036478 | 0.239389 |
| Nprl3 | 606.1558 | -3.40899 | 1.629787 | -2.09168 | 0.036467 | 0.239389 |
| Pus7 | 87.75545 | -8.03684 | 3.841688 | -2.09201 | 0.036438 | 0.239389 |
| RfC3 | 420.2724 | -3.53229 | 1.689831 | -2.09032 | 0.036589 | 0.239921 |
| PolH | 591.1932 | 3.963434 | 1.897521 | 2.088743 | 0.036731 | 0.240067 |
| Ugt302K1 | 516.7517 | -3.0591 | 1.464148 | -2.08934 | 0.036677 | 0.240067 |
| CG10927 | 73.58641 | 8.106286 | 3.880648 | 2.0889 | 0.036717 | 0.240067 |
| mio | 121.5483 | 6.243691 | 2.989018 | 2.088877 | 0.036719 | 0.240067 |
| Cad87A | 1574.861 | -2.09809 | 1.004656 | -2.08837 | 0.036765 | 0.240092 |
| CG31523 | 1724.964 | 1.756317 | 0.841217 | 2.08783 | 0.036813 | 0.240214 |
| CG8272 | 187.0621 | 5.912158 | 2.833257 | 2.086701 | 0.036915 | 0.240684 |
| CG34430 | 436.2869 | 4.649596 | 2.228562 | 2.086367 | 0.036945 | 0.240685 |
| CG42326 | 90.35201 | -8.01929 | 3.845584 | -2.08532 | 0.03704 | 0.241105 |
| CG34376 | 1062.751 | 2.358383 | 1.131695 | 2.083938 | 0.037166 | 0.241728 |
| Top3beta | 446.2045 | -3.44177 | 1.652409 | -2.08288 | 0.037262 | 0.241962 |
| Pp1alpha-9 | 3270.06 | 1.484926 | 0.712869 | 2.083026 | 0.037249 | 0.241962 |
| CG8060 | 166.1092 | 6.381068 | 3.064158 | 2.082486 | 0.037298 | 0.242 |
| CG5554 | 1237.737 | 2.006797 | 0.964233 | 2.081236 | 0.037412 | 0.242545 |
| CG10089 | 1155.306 | 2.680369 | 1.288965 | 2.079473 | 0.037574 | 0.243396 |
| Ir56a | 459.7524 | 4.042581 | 1.944798 | 2.078663 | 0.037648 | 0.243682 |
| Cyp6a15Ps | 1557.116 | -1.99113 | 0.958524 | -2.07729 | 0.037775 | 0.244305 |
| qvr | 34967.11 | 0.814565 | 0.39221 | 2.076857 | 0.037815 | 0.244365 |
| pod1 | 2551.677 | -1.84237 | 0.887812 | -2.07518 | 0.03797 | 0.245169 |
| CG11350 | 136.1122 | -6.78873 | 3.272143 | -2.0747 | 0.038014 | 0.245257 |
| CG10589 | 98.01135 | -7.953 | 3.835568 | -2.07349 | 0.038127 | 0.245591 |
| Ice2 | 93.01024 | -6.96061 | 3.356878 | -2.07354 | 0.038122 | 0.245591 |
| Pex14 | 2982.651 | 1.845371 | 0.890986 | 2.071156 | 0.038344 | 0.246496 |
| CG8086 | 397.2571 | 3.331482 | 1.608323 | 2.071402 | 0.038321 | 0.246496 |
| CG17806 | 269.8614 | -4.59073 | 2.216682 | -2.07099 | 0.03836 | 0.246496 |

| | | | | | | |
|---|---|---|---|---|---|---|
| GCS2beta | 739.3104 | -2.93195 | 1.416597 | -2.06971 | 0.038479 | 0.247069 |
| Alas | 2436.747 | -1.43505 | 0.693843 | -2.06826 | 0.038616 | 0.247746 |
| Nca | 20773.37 | 0.900338 | 0.435672 | 2.066553 | 0.038776 | 0.248379 |
| Est-6 | 213.7106 | 6.067059 | 2.935528 | 2.066769 | 0.038756 | 0.248379 |
| glob1 | 4169.051 | -1.22629 | 0.593877 | -2.06488 | 0.038934 | 0.248993 |
| CG8111 | 130.7741 | 5.625411 | 2.724176 | 2.064995 | 0.038923 | 0.248993 |
| TBC1D23 | 1348.117 | 2.18327 | 1.058586 | 2.06244 | 0.039166 | 0.250275 |
| Hers | 3736.648 | -1.22342 | 0.593433 | -2.06159 | 0.039247 | 0.250591 |
| tomboy40 | 69.7579 | 6.785258 | 3.296241 | 2.058484 | 0.039544 | 0.252288 |
| Axud1 | 2016.971 | 1.697988 | 0.825516 | 2.05688 | 0.039698 | 0.253069 |
| CG17760 | 116.1842 | 7.125761 | 3.465927 | 2.055946 | 0.039788 | 0.253441 |
| CG5026 | 2543.158 | -1.72937 | 0.841553 | -2.05498 | 0.039881 | 0.253835 |
| lncRNA:CR | 540.0747 | -2.68778 | 1.308371 | -2.0543 | 0.039947 | 0.254052 |
| l(2)efl | 80.61701 | -8.00051 | 3.895453 | -2.05381 | 0.039994 | 0.254152 |
| H | 1230.604 | -2.14001 | 1.042203 | -2.05335 | 0.040039 | 0.254233 |
| fat-spondi | 1909.006 | -2.08924 | 1.018153 | -2.05199 | 0.040171 | 0.25487 |
| htk | 886.9545 | -2.19799 | 1.071502 | -2.05132 | 0.040236 | 0.255082 |
| mRpL11 | 376.5976 | 3.636177 | 1.773659 | 2.050098 | 0.040355 | 0.255634 |
| CG5003 | 677.7734 | -2.79238 | 1.363782 | -2.04752 | 0.040607 | 0.257027 |
| CG2200 | 543.8187 | -3.84323 | 1.877705 | -2.04677 | 0.04068 | 0.25729 |
| CG32354 | 368.6616 | -3.46132 | 1.691851 | -2.04588 | 0.040768 | 0.257643 |
| Rh7 | 355.7074 | -4.37476 | 2.13993 | -2.04435 | 0.040919 | 0.258393 |
| mt:ATPase | 216804 | 0.76895 | 0.376255 | 2.043691 | 0.040984 | 0.258599 |
| nAChRalph | 18441.58 | 0.857021 | 0.419418 | 2.043361 | 0.041017 | 0.258602 |
| CDase | 6962.216 | -1.13332 | 0.555208 | -2.04126 | 0.041225 | 0.259104 |
| Clbn | 1129.863 | -2.89929 | 1.420112 | -2.04159 | 0.041192 | 0.259104 |
| lncRNA:CR | 207.0608 | -5.29955 | 2.595274 | -2.042 | 0.041152 | 0.259104 |
| asRNA:CR4 | 257.7987 | 3.50775 | 1.718234 | 2.041485 | 0.041203 | 0.259104 |
| CG1646 | 2775.614 | 1.421185 | 0.696519 | 2.040411 | 0.041309 | 0.259225 |
| CG34310 | 68.31972 | 7.846583 | 3.845004 | 2.040722 | 0.041279 | 0.259225 |
| eIF2Bbeta | 2766.016 | -1.92935 | 0.946255 | -2.03894 | 0.041456 | 0.259945 |
| JMJD5 | 584.9014 | -3.66847 | 1.799983 | -2.03806 | 0.041544 | 0.260293 |
| CG8066 | 449.9857 | -3.53989 | 1.738793 | -2.03583 | 0.041767 | 0.261399 |
| CG13458 | 488.7261 | -3.21436 | 1.579038 | -2.03564 | 0.041786 | 0.261399 |
| CG6907 | 191.8811 | -4.8328 | 2.37486 | -2.03498 | 0.041853 | 0.261407 |
| CG17600 | 636.1221 | 2.92935 | 1.4394 | 2.035118 | 0.041839 | 0.261407 |
| 140up | 321.222 | 4.142577 | 2.037904 | 2.032763 | 0.042076 | 0.262601 |
| brv3 | 89.14538 | 7.866328 | 3.87109 | 2.032071 | 0.042147 | 0.262777 |
| beat-IIb | 2151.074 | 1.784127 | 0.878085 | 2.031839 | 0.04217 | 0.262777 |
| Amacr | 853.4449 | -2.62295 | 1.291728 | -2.03057 | 0.042298 | 0.263167 |
| Arv1 | 268.4205 | 3.892349 | 1.916828 | 2.03062 | 0.042294 | 0.263167 |
| CG33116 | 2234.941 | -1.76144 | 0.867733 | -2.02993 | 0.042364 | 0.263371 |
| CG4953 | 219.1634 | -4.77653 | 2.353471 | -2.02957 | 0.0424 | 0.263393 |
| kar | 222.2859 | 4.883869 | 2.407512 | 2.028596 | 0.042499 | 0.263806 |
| SCCRO3 | 1537.032 | 1.94937 | 0.961461 | 2.027509 | 0.04261 | 0.26429 |
| Hsc70-2 | 144.0694 | -6.35126 | 3.138808 | -2.02346 | 0.043025 | 0.266657 |
| CG31249 | 245.8867 | 4.440477 | 2.198384 | 2.019883 | 0.043396 | 0.268744 |
| CG5757 | 87.48157 | -6.83914 | 3.390036 | -2.01742 | 0.043651 | 0.27012 |
| SPR | 1458.983 | 1.988743 | 0.986027 | 2.016926 | 0.043703 | 0.270233 |
| MTA1-like | 2472.746 | 1.519834 | 0.754049 | 2.015564 | 0.043846 | 0.270629 |

| | | | | | | |
|---|---|---|---|---|---|---|
| CG4459 | 123.1108 | 6.351328 | 3.151484 | 2.015345 | 0.043868 | 0.270629 |
| Pex6 | 547.1948 | -3.18314 | 1.579151 | -2.01573 | 0.043828 | 0.270629 |
| Zn72D | 4576.827 | -1.386 | 0.689089 | -2.01136 | 0.044288 | 0.273006 |
| axed | 589.6423 | 2.414363 | 1.201704 | 2.009115 | 0.044525 | 0.274257 |
| SF1 | 749.1142 | -2.86807 | 1.429411 | -2.00647 | 0.044806 | 0.275355 |
| Non1 | 6265.88 | 1.149376 | 0.572809 | 2.006561 | 0.044796 | 0.275355 |
| mRpL22 | 1673.785 | 1.598837 | 0.796625 | 2.007014 | 0.044748 | 0.275355 |
| mRpL40 | 597.7726 | 2.402406 | 1.198958 | 2.003745 | 0.045097 | 0.276722 |
| fz4 | 117.4064 | 5.739317 | 2.863905 | 2.004018 | 0.045068 | 0.276722 |
| CG11374 | 486.0805 | 3.100183 | 1.549502 | 2.00076 | 0.045418 | 0.278355 |
| CR43170 | 193.4151 | -4.88588 | 2.442177 | -2.00062 | 0.045433 | 0.278355 |
| DIP-delta | 6233.02 | 1.055748 | 0.527931 | 1.999783 | 0.045524 | 0.278519 |
| lncRNA:CR | 133.3537 | 6.044161 | 3.022485 | 1.999732 | 0.045529 | 0.278519 |
| S6kII | 1388.287 | -1.89408 | 0.947476 | -1.99908 | 0.045599 | 0.278523 |
| CG7991 | 871.5946 | 2.486709 | 1.243787 | 1.999304 | 0.045575 | 0.278523 |
| CG7656 | 1285.375 | -2.15362 | 1.077764 | -1.99823 | 0.045691 | 0.278874 |
| CG17646 | 1375.01 | -2.56202 | 1.282452 | -1.99775 | 0.045744 | 0.278982 |
| CG12071 | 2861.088 | -1.38317 | 0.692574 | -1.99714 | 0.04581 | 0.279172 |
| THG | 179.1605 | -6.13375 | 3.075049 | -1.99468 | 0.046077 | 0.280484 |
| mRpS2 | 1145.822 | 2.334855 | 1.170634 | 1.994523 | 0.046095 | 0.280484 |
| Rpt6 | 1025.652 | -2.3663 | 1.186734 | -1.99396 | 0.046156 | 0.280642 |
| CG4497 | 855.1118 | 3.260829 | 1.636114 | 1.993033 | 0.046258 | 0.280876 |
| Acph-1 | 655.6219 | 2.797657 | 1.403761 | 1.992973 | 0.046264 | 0.280876 |
| CG11807 | 2904.932 | 1.419068 | 0.712187 | 1.99255 | 0.046311 | 0.280903 |
| Su(var)2-H | 2245.964 | 1.704816 | 0.855705 | 1.992293 | 0.046339 | 0.280903 |
| mthl8 | 2040.095 | 1.872571 | 0.940275 | 1.991514 | 0.046424 | 0.281209 |
| l(3)72Dp | 373.5511 | 4.820518 | 2.424547 | 1.988214 | 0.046788 | 0.282984 |
| CG15202 | 1535.245 | 2.660906 | 1.338316 | 1.98825 | 0.046784 | 0.282984 |
| CG16868 | 1347.036 | 2.951552 | 1.485187 | 1.987327 | 0.046886 | 0.283152 |
| Snx1 | 2011.478 | 2.090875 | 1.052078 | 1.987375 | 0.046881 | 0.283152 |
| RIOK1 | 299.8654 | 4.25182 | 2.140217 | 1.98663 | 0.046963 | 0.283405 |
| Vps11 | 1406.879 | 2.087446 | 1.05184 | 1.984566 | 0.047193 | 0.284575 |
| CG33310 | 205.0628 | 4.397754 | 2.217272 | 1.983407 | 0.047322 | 0.284926 |
| CG13405 | 107.5731 | 6.239835 | 3.145843 | 1.983517 | 0.04731 | 0.284926 |
| Rubicon | 483.5554 | -2.8002 | 1.412331 | -1.98268 | 0.047403 | 0.285202 |
| GlcAT-S | 4349.922 | -2.10201 | 1.061042 | -1.98108 | 0.047582 | 0.286062 |
| p47 | 809.7203 | -3.24239 | 1.637376 | -1.98024 | 0.047677 | 0.286308 |
| CG8009 | 480.3909 | 2.740225 | 1.384116 | 1.979766 | 0.04773 | 0.286308 |
| CG1486 | 1504.663 | -2.51461 | 1.27009 | -1.97987 | 0.047718 | 0.286308 |
| Vps51 | 704.4347 | -3.87215 | 1.956598 | -1.97902 | 0.047813 | 0.286595 |
| lva | 1151.95 | 2.409867 | 1.218278 | 1.978093 | 0.047918 | 0.287009 |
| cher | 167.4667 | 5.631477 | 2.847842 | 1.977455 | 0.04799 | 0.287226 |
| Cog7 | 1232.668 | 2.831298 | 1.432234 | 1.97684 | 0.04806 | 0.287427 |
| PGRP-LA | 2672.202 | -2.19679 | 1.111666 | -1.97612 | 0.048141 | 0.287698 |
| Nxt1 | 751.911 | -2.36726 | 1.198654 | -1.97493 | 0.048276 | 0.288291 |
| sNPF | 1929.616 | -2.62993 | 1.331962 | -1.97447 | 0.048328 | 0.288386 |
| Hsp68 | 598.0582 | -2.45056 | 1.242363 | -1.9725 | 0.048553 | 0.289512 |
| POLDIP2 | 1184.634 | 2.73736 | 1.388264 | 1.971786 | 0.048634 | 0.289711 |
| CG34015 | 1869.472 | 2.621155 | 1.329636 | 1.971333 | 0.048686 | 0.289711 |
| TORIP | 2034.784 | 1.443678 | 0.732364 | 1.971259 | 0.048694 | 0.289711 |

| | | | | | | |
|---|---|---|---|---|---|---|
| CG46397 | 314.9272 | -3.7676 | 1.912762 | -1.96972 | 0.048871 | 0.290545 |
| CG33640 | 365.6556 | -4.16507 | 2.116711 | -1.96771 | 0.049102 | 0.291696 |
| CG5321 | 1252.076 | -1.91624 | 0.973994 | -1.9674 | 0.049137 | 0.291696 |
| CG4666 | 319.3761 | 4.081799 | 2.075065 | 1.967071 | 0.049175 | 0.291707 |
| pn | 306.3275 | 4.114035 | 2.093134 | 1.965491 | 0.049357 | 0.292572 |
| CG13313 | 95.88356 | -7.38506 | 3.762068 | -1.96303 | 0.049643 | 0.294045 |
| CG32176 | 766.5889 | 2.171783 | 1.107316 | 1.961303 | 0.049844 | 0.295019 |
| CG4045 | 767.3254 | -2.55778 | 1.305294 | -1.95955 | 0.050049 | 0.296015 |
| CG8837 | 66.90387 | 7.580787 | 3.873894 | 1.956891 | 0.05036 | 0.297638 |
| msps | 3941.245 | 1.194725 | 0.610758 | 1.956135 | 0.050449 | 0.297944 |
| tra2 | 3309.114 | 2.815375 | 1.439511 | 1.955785 | 0.05049 | 0.297968 |
| CAH8 | 82.13328 | -7.53683 | 3.856239 | -1.95445 | 0.050648 | 0.298251 |
| Pat1 | 2337.581 | -1.62427 | 0.831172 | -1.9542 | 0.050678 | 0.298251 |
| Rph | 1109.855 | 2.647794 | 1.354981 | 1.95412 | 0.050687 | 0.298251 |
| lncRNA:CR | 177.3416 | -5.00536 | 2.560206 | -1.95506 | 0.050576 | 0.298251 |
| Qtzl | 1189.987 | 2.247248 | 1.150672 | 1.952988 | 0.050821 | 0.29882 |
| Irp-1B | 1079.51 | -2.56837 | 1.31691 | -1.9503 | 0.05114 | 0.299709 |
| Manf | 4386.112 | 1.250203 | 0.640827 | 1.950921 | 0.051066 | 0.299709 |
| mib2 | 1963.713 | 1.589631 | 0.814968 | 1.950544 | 0.051111 | 0.299709 |
| ben | 5201.889 | -1.15161 | 0.590196 | -1.95124 | 0.051029 | 0.299709 |
| Snx21 | 436.0153 | 3.461901 | 1.775202 | 1.950144 | 0.051159 | 0.299709 |
| beat-Ic | 4271.438 | 1.267064 | 0.650121 | 1.948966 | 0.051299 | 0.300069 |
| Mtp | 658.8395 | -2.78131 | 1.426917 | -1.94917 | 0.051275 | 0.300069 |
| CG9521 | 74.61461 | 7.49598 | 3.846679 | 1.948689 | 0.051333 | 0.300069 |
| CG5861 | 3206.156 | 1.252707 | 0.64301 | 1.948193 | 0.051392 | 0.300197 |
| RpLP0-like | 285.5221 | 4.646331 | 2.385755 | 1.94753 | 0.051471 | 0.300442 |
| CG8306 | 68.30741 | -7.5358 | 3.870308 | -1.94708 | 0.051525 | 0.300539 |
| CG30392 | 2626.604 | -2.26045 | 1.16214 | -1.94507 | 0.051766 | 0.301725 |
| Sec31 | 2060.278 | -1.5164 | 0.77997 | -1.94418 | 0.051873 | 0.302131 |
| Cap-D3 | 907.9253 | 3.165024 | 1.628931 | 1.943007 | 0.052015 | 0.302235 |
| Nup75 | 271.7323 | 5.194947 | 2.672925 | 1.943544 | 0.05195 | 0.302235 |
| TBCC | 1276.011 | -1.91918 | 0.987847 | -1.94279 | 0.052042 | 0.302235 |
| Cep97 | 696.958 | 3.175229 | 1.634361 | 1.942795 | 0.052041 | 0.302235 |
| CG4287 | 1690.955 | 1.813781 | 0.934092 | 1.941758 | 0.052166 | 0.302739 |
| CG8910 | 730.1187 | 2.25921 | 1.164033 | 1.940847 | 0.052277 | 0.302802 |
| dpa | 128.5973 | 6.556025 | 3.377789 | 1.940922 | 0.052268 | 0.302802 |
| CG7394 | 1331.925 | 1.818355 | 0.936942 | 1.940734 | 0.052291 | 0.302802 |
| Der-2 | 3347.543 | -2.26961 | 1.169808 | -1.94016 | 0.052361 | 0.302989 |
| CG9646 | 4704.17 | 1.430605 | 0.737487 | 1.939838 | 0.052399 | 0.302995 |
| Nup37 | 148.6359 | 7.017407 | 3.618206 | 1.939471 | 0.052444 | 0.303035 |
| CG11523 | 830.5088 | -2.86157 | 1.475836 | -1.93895 | 0.052508 | 0.303163 |
| Capr | 917.9164 | 2.032056 | 1.048171 | 1.938668 | 0.052542 | 0.303163 |
| Vps13D | 1555.906 | -2.29436 | 1.183728 | -1.93825 | 0.052593 | 0.30324 |
| CG32647 | 3640.303 | -1.22495 | 0.632404 | -1.93697 | 0.052749 | 0.303781 |
| Taf8 | 230.8218 | -4.93558 | 2.548239 | -1.93686 | 0.052762 | 0.303781 |
| Cyp12a4 | 2197.09 | 2.550219 | 1.317172 | 1.936132 | 0.052852 | 0.304076 |
| JMJD7 | 748.7031 | 2.303648 | 1.190108 | 1.935663 | 0.052909 | 0.304189 |
| Atg9 | 747.137 | 4.061287 | 2.09958 | 1.934334 | 0.053072 | 0.30469 |
| CG44098 | 8094.822 | -1.56751 | 0.810235 | -1.93464 | 0.053035 | 0.30469 |
| lost | 7379.066 | -1.00108 | 0.517679 | -1.93378 | 0.053141 | 0.304865 |

| | | | | | | |
|---|---|---|---|---|---|---|
| CG12118 | 92.52845 | -6.81133 | 3.523658 | -1.93303 | 0.053233 | 0.304958 |
| Mul1 | 506.2078 | 4.055104 | 2.097467 | 1.933334 | 0.053195 | 0.304958 |
| CG7943 | 3197.534 | 1.811796 | 0.937801 | 1.931962 | 0.053364 | 0.305493 |
| CG6230 | 2141.307 | -2.57409 | 1.333526 | -1.93029 | 0.053571 | 0.30646 |
| CG31300 | 1315.896 | 2.67464 | 1.386781 | 1.928667 | 0.053772 | 0.30739 |
| CG3847 | 999.425 | -2.24365 | 1.164358 | -1.92694 | 0.053987 | 0.308172 |
| CG2641 | 247.8879 | -4.82642 | 2.504937 | -1.92676 | 0.054009 | 0.308172 |
| Arpc5 | 622.9952 | 2.590756 | 1.3447 | 1.926642 | 0.054024 | 0.308172 |
| TppII | 2101.886 | -2.20548 | 1.146541 | -1.9236 | 0.054405 | 0.310124 |
| CG2016 | 667.0227 | -3.7661 | 1.958613 | -1.92284 | 0.0545 | 0.310445 |
| CG7806 | 1765.415 | -2.46319 | 1.282646 | -1.9204 | 0.054807 | 0.311753 |
| Hsp27 | 169.5863 | -6.07681 | 3.164017 | -1.9206 | 0.054782 | 0.311753 |
| gt | 206.2008 | 4.921831 | 2.563916 | 1.919654 | 0.054902 | 0.311972 |
| Ac13E | 323.0404 | 3.3763 | 1.759811 | 1.918559 | 0.05504 | 0.311972 |
| beat-VII | 8456.244 | 1.135427 | 0.591685 | 1.918973 | 0.054988 | 0.311972 |
| Pdfr | 1553.094 | -1.57166 | 0.819174 | -1.91859 | 0.055037 | 0.311972 |
| Gp210 | 511.7236 | -3.68395 | 1.919765 | -1.91896 | 0.05499 | 0.311972 |
| CG31104 | 68.55416 | 7.393389 | 3.855714 | 1.917515 | 0.055173 | 0.312501 |
| twe | 346.5103 | -4.85365 | 2.536109 | -1.91382 | 0.055644 | 0.314948 |
| Nepl3 | 286.2054 | 5.223376 | 2.729862 | 1.913421 | 0.055694 | 0.315011 |
| lncRNA:CR | 331.1606 | -3.35663 | 1.755361 | -1.91221 | 0.055849 | 0.315663 |
| PCNA2 | 320.7195 | 4.118077 | 2.154854 | 1.91107 | 0.055996 | 0.31627 |
| rnh1 | 993.8837 | -2.99849 | 1.569585 | -1.91037 | 0.056085 | 0.316555 |
| CG15861 | 225.4511 | 4.046489 | 2.119382 | 1.909278 | 0.056226 | 0.317127 |
| CG12105 | 510.8156 | -2.93981 | 1.54142 | -1.90721 | 0.056493 | 0.31841 |
| wuho | 553.6737 | 3.12903 | 1.641122 | 1.90664 | 0.056567 | 0.318602 |
| Prp4k | 662.7402 | 2.261301 | 1.18665 | 1.905617 | 0.0567 | 0.319126 |
| Synj | 7538.994 | -1.22547 | 0.643436 | -1.90457 | 0.056837 | 0.319515 |
| CG1161 | 4370.649 | -1.09172 | 0.57324 | -1.90447 | 0.056849 | 0.319515 |
| CG9896 | 1787.149 | -2.3656 | 1.242661 | -1.90365 | 0.056955 | 0.319889 |
| Sema1b | 3542.466 | 1.668117 | 0.876742 | 1.902631 | 0.057089 | 0.320416 |
| Syx8 | 574.327 | 2.360131 | 1.240973 | 1.901839 | 0.057192 | 0.320772 |
| IKKepsilon | 745.2161 | 3.124227 | 1.644333 | 1.899997 | 0.057434 | 0.3219 |
| CG16854 | 1851.305 | -2.52619 | 1.329898 | -1.89954 | 0.057494 | 0.322013 |
| CG31548 | 791.9065 | -2.03858 | 1.073851 | -1.89838 | 0.057646 | 0.322577 |
| ASPP | 2794.274 | 1.371454 | 0.722517 | 1.898161 | 0.057675 | 0.322577 |
| lncRNA:spl | 379.9648 | 4.00596 | 2.111607 | 1.897114 | 0.057813 | 0.323124 |
| lncRNA:CR | 83.32897 | 7.341261 | 3.870976 | 1.896488 | 0.057895 | 0.323361 |
| CG8516 | 471.8205 | -3.67669 | 1.941006 | -1.89422 | 0.058196 | 0.323913 |
| Rbcn-3A | 5919.533 | 1.20858 | 0.637913 | 1.894586 | 0.058147 | 0.323913 |
| CR40354 | 104.8918 | 5.146533 | 2.716781 | 1.89435 | 0.058179 | 0.323913 |
| CDK2AP1 | 353.7465 | 4.091234 | 2.159753 | 1.894307 | 0.058184 | 0.323913 |
| Cpsf5 | 4632.794 | 1.066185 | 0.562712 | 1.894727 | 0.058129 | 0.323913 |
| CG7328 | 447.8493 | 3.321555 | 1.754687 | 1.892961 | 0.058363 | 0.324616 |
| brp | 26121.68 | 0.878066 | 0.464008 | 1.892351 | 0.058444 | 0.324843 |
| CG17029 | 3150.114 | -1.47769 | 0.781163 | -1.89165 | 0.058537 | 0.325134 |
| mt:ND5 | 77522.58 | 0.700917 | 0.370762 | 1.890474 | 0.058695 | 0.325589 |
| wcy | 3647.274 | 1.177072 | 0.622647 | 1.890433 | 0.0587 | 0.325589 |
| CG17778 | 3370.258 | 1.176813 | 0.622771 | 1.88964 | 0.058806 | 0.32565 |
| CG11425 | 334.593 | -3.60473 | 1.907298 | -1.88997 | 0.058762 | 0.32565 |

| | | | | | | |
|---|---|---|---|---|---|---|
| lncRNA:CR | 420.3659 | 3.048727 | 1.613562 | 1.88944 | 0.058833 | 0.32565 |
| CAH3 | 1469.081 | -1.70347 | 0.902104 | -1.88833 | 0.058982 | 0.326083 |
| CG9691 | 14519.73 | -1.01501 | 0.537541 | -1.88825 | 0.058992 | 0.326083 |
| Apoltp | 232.0738 | -4.41462 | 2.338809 | -1.88755 | 0.059087 | 0.326157 |
| CG8958 | 580.0384 | -3.62025 | 1.917965 | -1.88755 | 0.059087 | 0.326157 |
| Pvr | 714.8702 | -3.12876 | 1.657895 | -1.88719 | 0.059135 | 0.326199 |
| Snp | 254.0938 | -3.87364 | 2.053073 | -1.88675 | 0.059194 | 0.326297 |
| Cyp6a2 | 602.0069 | 2.413353 | 1.280268 | 1.885036 | 0.059425 | 0.327347 |
| CG14894 | 1427.159 | 2.007454 | 1.065759 | 1.883591 | 0.05962 | 0.328198 |
| CG5800 | 205.6131 | 5.397159 | 2.866599 | 1.882774 | 0.059731 | 0.328582 |
| Rcc1 | 365.991 | 2.777751 | 1.476551 | 1.881243 | 0.059939 | 0.3295 |
| Gpdh3 | 209.4834 | -3.8795 | 2.06331 | -1.88023 | 0.060076 | 0.330029 |
| prd1 | 3715.879 | 1.176409 | 0.626687 | 1.877187 | 0.060492 | 0.331408 |
| Pur-alpha | 18041.19 | 0.862725 | 0.459523 | 1.877434 | 0.060459 | 0.331408 |
| CR40190 | 194.8484 | -4.90572 | 2.613008 | -1.87742 | 0.06046 | 0.331408 |
| CG30419 | 600.3822 | 2.923411 | 1.55707 | 1.877508 | 0.060448 | 0.331408 |
| Aef1 | 2269.766 | -1.85114 | 0.987086 | -1.87536 | 0.060744 | 0.331541 |
| CG14459 | 7253.063 | 1.008271 | 0.537559 | 1.875647 | 0.060704 | 0.331541 |
| csul | 751.8566 | -3.21549 | 1.714782 | -1.87516 | 0.060771 | 0.331541 |
| CG3520 | 138.9014 | 5.823264 | 3.103711 | 1.876226 | 0.060624 | 0.331541 |
| CG17032 | 106.4841 | -6.16449 | 3.287541 | -1.87511 | 0.060778 | 0.331541 |
| Adi1 | 1145.647 | 2.779448 | 1.482085 | 1.875363 | 0.060743 | 0.331541 |
| SerRS | 642.2161 | -3.12676 | 1.667692 | -1.8749 | 0.060806 | 0.331541 |
| Tsf2 | 510.7417 | -3.23027 | 1.723284 | -1.87449 | 0.060863 | 0.331628 |
| lncRNA:CR | 67.98101 | 7.253593 | 3.870766 | 1.873942 | 0.060938 | 0.331812 |
| CG42521 | 87.67812 | 7.226693 | 3.864006 | 1.870259 | 0.061448 | 0.33436 |
| Nepl21 | 382.0383 | 3.822447 | 2.04644 | 1.867852 | 0.061783 | 0.335954 |
| lncRNA:CR | 140.4323 | 5.269914 | 2.822133 | 1.867352 | 0.061852 | 0.336106 |
| Gbeta5 | 7447.721 | -0.95771 | 0.513031 | -1.86677 | 0.061933 | 0.336318 |
| lectin-46Ca | 423.5051 | -3.6777 | 1.970741 | -1.86615 | 0.06202 | 0.336561 |
| CG11737 | 1569.039 | -1.8442 | 0.988503 | -1.86565 | 0.06209 | 0.336714 |
| CG18476 | 581.7805 | 2.418526 | 1.297369 | 1.864177 | 0.062297 | 0.337606 |
| HUWE1 | 4044.718 | -1.14951 | 0.616744 | -1.86384 | 0.062345 | 0.337636 |
| CG6013 | 781.5532 | -1.99696 | 1.071715 | -1.86333 | 0.062415 | 0.337671 |
| aft | 326.0835 | 4.041253 | 2.168992 | 1.863194 | 0.062435 | 0.337671 |
| CG32686 | 798.7654 | -3.26087 | 1.751457 | -1.8618 | 0.062631 | 0.337944 |
| lncRNA:CR | 2498.15 | 2.291094 | 1.230392 | 1.862085 | 0.062591 | 0.337944 |
| CG18259 | 975.2053 | 2.003349 | 1.075815 | 1.86217 | 0.062579 | 0.337944 |
| Nufip | 169.5917 | -5.13404 | 2.757803 | -1.86164 | 0.062654 | 0.337944 |
| mRpS10 | 3748.8 | 1.287286 | 0.69215 | 1.859836 | 0.062909 | 0.33909 |
| CG15561 | 134.0692 | -5.38656 | 2.896842 | -1.85946 | 0.062962 | 0.33915 |
| CG43742 | 662.2992 | -3.3781 | 1.817113 | -1.85905 | 0.06302 | 0.339237 |
| gek | 2055.779 | 2.091799 | 1.125966 | 1.857781 | 0.0632 | 0.339976 |
| Gart | 823.3965 | -2.26192 | 1.218199 | -1.85677 | 0.063343 | 0.340175 |
| Mer | 239.5631 | 3.576151 | 1.925869 | 1.856903 | 0.063325 | 0.340175 |
| Rbsn-5 | 810.3907 | -2.19126 | 1.180235 | -1.85663 | 0.063364 | 0.340175 |
| CG5276 | 899.9185 | -1.92384 | 1.036817 | -1.85552 | 0.063522 | 0.340793 |
| Mekk1 | 1904.636 | 1.879422 | 1.01318 | 1.854973 | 0.0636 | 0.340985 |
| Spec2 | 299.9368 | 4.026097 | 2.171083 | 1.854419 | 0.063679 | 0.341035 |
| Arl8 | 4078.682 | -1.62414 | 0.875874 | -1.85431 | 0.063694 | 0.341035 |

| | | | | | | |
|---|---|---|---|---|---|---|
| brm | 1202.508 | -2.48671 | 1.34161 | -1.85353 | 0.063807 | 0.34141 |
| E(z) | 1480.355 | 2.273043 | 1.227327 | 1.852027 | 0.064022 | 0.342332 |
| Shc | 428.4593 | -3.81277 | 2.063213 | -1.84798 | 0.064606 | 0.345224 |
| CG10565 | 1490.537 | 1.914595 | 1.037833 | 1.8448 | 0.065067 | 0.346904 |
| SPoCk | 5272.995 | 1.003393 | 0.543957 | 1.844618 | 0.065093 | 0.346904 |
| pita | 1015.874 | 1.837388 | 0.995794 | 1.845149 | 0.065016 | 0.346904 |
| PDZ-GEF | 3453.046 | 1.400629 | 0.758954 | 1.845472 | 0.064969 | 0.346904 |
| Tailor | 2671.015 | 1.606029 | 0.871196 | 1.843475 | 0.06526 | 0.347561 |
| Alg9 | 2876.802 | 1.509439 | 0.819333 | 1.842279 | 0.065434 | 0.34826 |
| Jupiter | 5764.015 | 1.053958 | 0.572689 | 1.840368 | 0.065714 | 0.349518 |
| Or13a | 134.8253 | 5.128996 | 2.788621 | 1.839259 | 0.065877 | 0.350152 |
| CG44837 | 3610.197 | 1.416407 | 0.770391 | 1.838558 | 0.06598 | 0.350468 |
| CG43085 | 70.91953 | -6.81738 | 3.709617 | -1.83776 | 0.066098 | 0.350861 |
| Parp | 4433.767 | 1.087188 | 0.591745 | 1.837257 | 0.066172 | 0.351022 |
| Eogt | 329.2101 | -4.1788 | 2.275422 | -1.8365 | 0.066284 | 0.351386 |
| Con | 6769.621 | 0.992928 | 0.540769 | 1.836142 | 0.066337 | 0.351431 |
| PAN2 | 1069.256 | 1.979654 | 1.079059 | 1.834612 | 0.066563 | 0.352398 |
| CG2790 | 465.8267 | -2.95234 | 1.60955 | -1.83426 | 0.066615 | 0.35244 |
| CG12241 | 252.3794 | 4.38595 | 2.392474 | 1.833228 | 0.066769 | 0.35266 |
| mip40 | 375.5024 | 2.838931 | 1.548946 | 1.832815 | 0.06683 | 0.35266 |
| ix | 520.4621 | -3.34404 | 1.823916 | -1.83344 | 0.066737 | 0.35266 |
| Sce | 831.0602 | -2.51342 | 1.371353 | -1.8328 | 0.066832 | 0.35266 |
| DNAlig3 | 712.2498 | 2.515635 | 1.373946 | 1.830956 | 0.067107 | 0.353878 |
| mtm | 1184.911 | -2.63853 | 1.443266 | -1.82816 | 0.067525 | 0.355796 |
| fd102C | 402.3852 | -4.39813 | 2.406453 | -1.82764 | 0.067604 | 0.355796 |
| Dref | 2587.334 | -1.64917 | 0.902243 | -1.82786 | 0.067571 | 0.355796 |
| CG15643 | 757.5769 | -2.71133 | 1.484474 | -1.82646 | 0.067781 | 0.356496 |
| CG14805 | 925.7035 | -2.97378 | 1.628942 | -1.82559 | 0.067912 | 0.356527 |
| Nsf2 | 873.4614 | -2.25577 | 1.235672 | -1.82554 | 0.067919 | 0.356527 |
| sti | 146.3123 | -4.90069 | 2.684523 | -1.82553 | 0.06792 | 0.356527 |
| ana1 | 812.7773 | 3.008623 | 1.648579 | 1.824979 | 0.068004 | 0.356703 |
| CG10948 | 13343.22 | 0.953141 | 0.522349 | 1.824722 | 0.068043 | 0.356703 |
| CG12007 | 1158.931 | -2.47655 | 1.357783 | -1.82396 | 0.068157 | 0.35707 |
| atilla | 263.6808 | -5.16806 | 2.837674 | -1.82123 | 0.068571 | 0.359005 |
| osy | 1360.2 | -1.83381 | 1.007576 | -1.82002 | 0.068756 | 0.359737 |
| CG9596 | 334.7507 | 4.170398 | 2.292161 | 1.819418 | 0.068848 | 0.359953 |
| Vps50 | 1161.415 | -2.57356 | 1.415382 | -1.81828 | 0.069022 | 0.359953 |
| HDAC1 | 812.1445 | -3.32251 | 1.827217 | -1.81835 | 0.069011 | 0.359953 |
| CG10353 | 8055.601 | 1.240184 | 0.681974 | 1.818521 | 0.068985 | 0.359953 |
| Twdlalpha | 103.4791 | 5.226577 | 2.873986 | 1.818581 | 0.068975 | 0.359953 |
| RhoGEF2 | 1509.157 | 1.917975 | 1.05554 | 1.817056 | 0.069209 | 0.360693 |
| BtbVII | 3268.098 | -1.16089 | 0.63919 | -1.81619 | 0.069342 | 0.361153 |
| Fsn | 2252.16 | 1.328512 | 0.731886 | 1.815191 | 0.069495 | 0.361645 |
| CG31751 | 8900.564 | 0.930365 | 0.512602 | 1.814984 | 0.069526 | 0.361645 |
| EDTP | 765.3655 | -1.9839 | 1.093495 | -1.81428 | 0.069635 | 0.361974 |
| pasha | 521.1563 | -2.42964 | 1.34066 | -1.81227 | 0.069944 | 0.363111 |
| CG6664 | 1454.547 | -1.69142 | 0.93321 | -1.81247 | 0.069913 | 0.363111 |
| Hsp67Bc | 2670.989 | 1.78097 | 0.982903 | 1.811949 | 0.069994 | 0.363137 |
| lncRNA:CR | 77.80271 | -7.00713 | 3.8692 | -1.811 | 0.07014 | 0.363661 |
| mRpS26 | 1373.438 | 2.321172 | 1.282386 | 1.810042 | 0.070289 | 0.364197 |

| | | | | | | |
|---|---|---|---|---|---|---|
| CG16972 | 530.9608 | 3.166308 | 1.752118 | 1.807132 | 0.070742 | 0.366305 |
| Wdr59 | 1133.936 | -2.33652 | 1.293592 | -1.80623 | 0.070883 | 0.366799 |
| pho | 6194.313 | 0.960421 | 0.53187 | 1.805744 | 0.070958 | 0.366954 |
| fmt | 6312.051 | 1.019006 | 0.564533 | 1.805041 | 0.071068 | 0.367285 |
| CG16717 | 1351.574 | 1.570997 | 0.870606 | 1.804486 | 0.071155 | 0.367497 |
| lncRNA:CR | 145.2738 | 5.862107 | 3.249441 | 1.804035 | 0.071226 | 0.367626 |
| Hexo1 | 1277.902 | -2.12108 | 1.176444 | -1.80296 | 0.071395 | 0.368263 |
| Wdr81 | 2452.952 | 2.226842 | 1.236595 | 1.800785 | 0.071737 | 0.369313 |
| CG16898 | 118.0422 | 6.953107 | 3.860166 | 1.801246 | 0.071664 | 0.369313 |
| CG15027 | 192.3378 | 4.225817 | 2.346595 | 1.80083 | 0.07173 | 0.369313 |
| ana2 | 337.0325 | -3.68783 | 2.049123 | -1.79971 | 0.071906 | 0.369948 |
| CG31785 | 2673.927 | -1.63218 | 0.907063 | -1.79941 | 0.071953 | 0.369954 |
| CG11695 | 207.5974 | -4.05012 | 2.251192 | -1.7991 | 0.072003 | 0.36997 |
| cold | 407.4566 | -3.92242 | 2.182027 | -1.7976 | 0.07224 | 0.370818 |
| Dhx15 | 1806.759 | 1.442428 | 0.802474 | 1.797476 | 0.07226 | 0.370818 |
| ACXD | 1390.762 | -1.73602 | 0.966546 | -1.79611 | 0.072478 | 0.371698 |
| CG1434 | 67.69439 | 6.941592 | 3.866109 | 1.795498 | 0.072574 | 0.371955 |
| CG9044 | 1235.704 | 1.852341 | 1.032513 | 1.794013 | 0.072811 | 0.372692 |
| Ccm3 | 4317.42 | -1.37123 | 0.764266 | -1.79418 | 0.072785 | 0.372692 |
| CG3348 | 191.1902 | 5.046392 | 2.814804 | 1.792804 | 0.073004 | 0.373443 |
| Inos | 29721.36 | -0.84777 | 0.472985 | -1.79239 | 0.073071 | 0.373545 |
| Klp68D | 2554.165 | 1.984026 | 1.107793 | 1.790971 | 0.073298 | 0.374468 |
| CG7523 | 548.9823 | -2.83082 | 1.580893 | -1.79065 | 0.07335 | 0.374495 |
| cyc | 2430.963 | 1.515269 | 0.846637 | 1.789749 | 0.073494 | 0.374755 |
| Sod1 | 36562.62 | -0.76054 | 0.424906 | -1.7899 | 0.07347 | 0.374755 |
| scny | 1536.971 | -1.56433 | 0.874205 | -1.78944 | 0.073544 | 0.374773 |
| Adk2 | 843.2528 | 2.191628 | 1.225277 | 1.78868 | 0.073666 | 0.375157 |
| elB | 2832.568 | 1.578679 | 0.883425 | 1.786998 | 0.073938 | 0.3763 |
| CG8289 | 2628.571 | 1.387424 | 0.776695 | 1.786317 | 0.074048 | 0.376622 |
| CG11399 | 1541.306 | 2.73573 | 1.532577 | 1.785052 | 0.074253 | 0.37725 |
| RNaseZ | 419.721 | -3.15682 | 1.768553 | -1.78497 | 0.074266 | 0.37725 |
| l(1)G0007 | 704.8622 | 2.544029 | 1.425763 | 1.784328 | 0.07437 | 0.377544 |
| CG17637 | 123.9137 | 4.760741 | 2.668629 | 1.783966 | 0.074429 | 0.377604 |
| CG7135 | 2721.666 | 2.321694 | 1.302302 | 1.782762 | 0.074625 | 0.378358 |
| CG40498 | 13134.32 | 0.9355 | 0.52513 | 1.781463 | 0.074837 | 0.379192 |
| Actbeta | 5308.563 | 1.099223 | 0.617229 | 1.7809 | 0.074929 | 0.379419 |
| eIF3a | 2046.975 | 1.408248 | 0.791571 | 1.779055 | 0.075231 | 0.380467 |
| AP-1mu | 2839.664 | -1.48037 | 0.832062 | -1.77916 | 0.075213 | 0.380467 |
| fws | 432.958 | 2.553839 | 1.436357 | 1.777997 | 0.075404 | 0.380852 |
| CG12259 | 281.5784 | -3.45075 | 1.941103 | -1.77772 | 0.075449 | 0.380852 |
| nbs | 673.5599 | 2.566121 | 1.44313 | 1.778163 | 0.075377 | 0.380852 |
| Gmer | 622.5542 | -3.00498 | 1.690663 | -1.7774 | 0.075503 | 0.380882 |
| pds5 | 642.337 | 2.12988 | 1.19879 | 1.776691 | 0.075619 | 0.38123 |
| DCTN3-p2 | 1492.955 | -2.44097 | 1.374623 | -1.77574 | 0.075776 | 0.381781 |
| psidin | 673.2579 | 3.083382 | 1.73676 | 1.775365 | 0.075838 | 0.381794 |
| lncRNA:CR | 131.2057 | -5.3433 | 3.010059 | -1.77515 | 0.075874 | 0.381794 |
| wfs1 | 679.1988 | 3.291455 | 1.856048 | 1.773367 | 0.076168 | 0.382114 |
| Rab10 | 679.2426 | -2.39702 | 1.351933 | -1.77304 | 0.076223 | 0.382114 |
| Haspin | 4871.069 | -0.98596 | 0.555845 | -1.7738 | 0.076097 | 0.382114 |
| Cpsf73 | 346.5571 | 3.373597 | 1.901914 | 1.77379 | 0.076098 | 0.382114 |

| | | | | | | |
|---|---|---|---|---|---|---|
| mRpS6 | 1234.542 | 1.592239 | 0.897921 | 1.773251 | 0.076187 | 0.382114 |
| hay | 493.1412 | 3.826495 | 2.156855 | 1.774108 | 0.076045 | 0.382114 |
| nac | 69.38527 | -6.87461 | 3.878356 | -1.77256 | 0.076302 | 0.382273 |
| RpL40 | 41786.41 | 0.72339 | 0.408321 | 1.77162 | 0.076458 | 0.382813 |
| Ced-12 | 1866.589 | 1.683824 | 0.951613 | 1.769443 | 0.07682 | 0.384148 |
| CRMP | 2998.954 | 1.303455 | 0.736644 | 1.769452 | 0.076818 | 0.384148 |
| CG8229 | 1310.024 | 1.633932 | 0.924475 | 1.767417 | 0.077158 | 0.3856 |
| mRpL41 | 1893.401 | 1.997144 | 1.13078 | 1.766165 | 0.077368 | 0.386344 |
| Rai1 | 1686.577 | -1.86 | 1.053256 | -1.76595 | 0.077404 | 0.386344 |
| CG11403 | 196.6626 | 5.141117 | 2.912226 | 1.765357 | 0.077504 | 0.386604 |
| CG12341 | 1344.274 | -2.4687 | 1.399137 | -1.76445 | 0.077657 | 0.387125 |
| Syx6 | 11566.84 | 0.839666 | 0.475971 | 1.764112 | 0.077713 | 0.387131 |
| l(2)dtl | 86.4685 | 6.43156 | 3.646283 | 1.763868 | 0.077754 | 0.387131 |
| CG8407 | 109.6929 | -5.57263 | 3.160262 | -1.76334 | 0.077843 | 0.387331 |
| ZnT86D | 1549.951 | -2.37837 | 1.349507 | -1.7624 | 0.078002 | 0.387643 |
| ValRS | 759.9496 | -2.67609 | 1.518301 | -1.76256 | 0.077975 | 0.387643 |
| Pal2 | 26280.26 | -0.7353 | 0.417515 | -1.76114 | 0.078215 | 0.388459 |
| CG10904 | 789.5753 | -2.40768 | 1.368373 | -1.75952 | 0.078489 | 0.389581 |
| Tsr1 | 462.3763 | -2.41862 | 1.3769 | -1.75657 | 0.078991 | 0.391651 |
| CG10366 | 1165.204 | 2.519276 | 1.434705 | 1.755953 | 0.079096 | 0.391651 |
| CG6236 | 2084.01 | -1.27221 | 0.7246 | -1.75574 | 0.079133 | 0.391651 |
| MED25 | 125.1943 | -5.94267 | 3.383389 | -1.75643 | 0.079016 | 0.391651 |
| Atf6 | 3582.035 | 1.139116 | 0.648832 | 1.755639 | 0.07915 | 0.391651 |
| Past1 | 1347.871 | -1.92377 | 1.096703 | -1.75414 | 0.079407 | 0.392437 |
| CycK | 1663.283 | 2.129713 | 1.214018 | 1.754268 | 0.079385 | 0.392437 |
| lncRNA:CR | 179.0491 | 5.327767 | 3.039807 | 1.752666 | 0.079659 | 0.393444 |
| gkt | 793.2629 | -2.50866 | 1.432883 | -1.75078 | 0.079984 | 0.394803 |
| Ubc7 | 1294.831 | 1.847837 | 1.055776 | 1.750217 | 0.080081 | 0.395039 |
| Rab9 | 1970.582 | -2.02077 | 1.15508 | -1.74946 | 0.080211 | 0.395439 |
| Idgf4 | 4136.79 | -2.10797 | 1.205205 | -1.74905 | 0.080282 | 0.395459 |
| CG10362 | 1254.57 | -1.80759 | 1.03358 | -1.74887 | 0.080314 | 0.395459 |
| GstE1 | 8408.579 | -0.99195 | 0.567379 | -1.74829 | 0.080413 | 0.395464 |
| Fpps | 796.9782 | -2.09058 | 1.19559 | -1.74858 | 0.080364 | 0.395464 |
| CG45218 | 417.5752 | -2.93086 | 1.678438 | -1.74618 | 0.080779 | 0.397018 |
| CG2182 | 1182.113 | -2.09075 | 1.197715 | -1.74561 | 0.080878 | 0.397029 |
| CG2926 | 3647.431 | -1.23493 | 0.707449 | -1.7456 | 0.08088 | 0.397029 |
| Sccpdh1 | 2141.879 | -2.07632 | 1.189782 | -1.74513 | 0.080963 | 0.397191 |
| Srrm1 | 1105.241 | 1.882 | 1.079295 | 1.743731 | 0.081206 | 0.398143 |
| CG4707 | 237.2776 | -4.14439 | 2.378314 | -1.74257 | 0.081408 | 0.398891 |
| Cdc2rk | 465.9214 | -3.45101 | 1.980786 | -1.74224 | 0.081466 | 0.398929 |
| CG10492 | 1250.73 | -2.79004 | 1.602503 | -1.74105 | 0.081674 | 0.399706 |
| MFS15 | 809.8833 | -2.99296 | 1.719845 | -1.74025 | 0.081815 | 0.400151 |
| Nep1 | 617.1727 | 3.049997 | 1.753022 | 1.739851 | 0.081885 | 0.40025 |
| Daxx | 1485.776 | 1.638056 | 0.942069 | 1.738786 | 0.082072 | 0.400634 |
| Afti | 1507.054 | 2.301241 | 1.323653 | 1.738553 | 0.082113 | 0.400634 |
| Mctp | 1497.537 | 1.598664 | 0.919411 | 1.738791 | 0.082072 | 0.400634 |
| CG12768 | 2203.476 | -1.71448 | 0.986676 | -1.73764 | 0.082275 | 0.401178 |
| Mob4 | 3121.405 | 1.270918 | 0.732093 | 1.736007 | 0.082563 | 0.402336 |
| IntS4 | 148.3766 | -5.63321 | 3.246538 | -1.73514 | 0.082715 | 0.402347 |
| CG8545 | 183.6249 | 4.346481 | 2.504678 | 1.735345 | 0.08268 | 0.402347 |

| | | | | | | |
|---|---|---|---|---|---|---|
| CG9578 | 709.1287 | -2.8851 | 1.662446 | -1.73546 | 0.08266 | 0.402347 |
| CG3156 | 389.5592 | -2.97655 | 1.716814 | -1.73376 | 0.08296 | 0.403048 |
| CG42878 | 376.9589 | 3.719125 | 2.144906 | 1.733933 | 0.08293 | 0.403048 |
| Ssadh | 7984.198 | -0.91602 | 0.52865 | -1.73276 | 0.083139 | 0.403674 |
| Meics | 323.2703 | -3.31094 | 1.911309 | -1.73229 | 0.083222 | 0.403833 |
| CG2993 | 7066.795 | -0.91901 | 0.530708 | -1.73167 | 0.083332 | 0.404123 |
| CG14184 | 3487.115 | 1.699199 | 0.981566 | 1.731111 | 0.083432 | 0.404363 |
| CG6201 | 701.1767 | 2.593316 | 1.498783 | 1.730282 | 0.08358 | 0.404591 |
| CG12213 | 3032.344 | -1.54972 | 0.895611 | -1.73035 | 0.083568 | 0.404591 |
| Ca-alpha1I | 8914.665 | 0.809476 | 0.468116 | 1.729221 | 0.08377 | 0.405265 |
| Snx3 | 3271.862 | 1.142568 | 0.661021 | 1.728489 | 0.083901 | 0.405654 |
| CG7368 | 575.0951 | 3.488049 | 2.018327 | 1.728188 | 0.083955 | 0.405671 |
| CG18596 | 780.5323 | -3.44566 | 1.994665 | -1.72744 | 0.084089 | 0.406076 |
| CG15734 | 68.11227 | 6.666756 | 3.864265 | 1.725233 | 0.084486 | 0.407746 |
| Cypl | 547.5358 | 3.106633 | 1.801831 | 1.724153 | 0.08468 | 0.408439 |
| Vps35 | 1417.326 | 1.943991 | 1.128037 | 1.72334 | 0.084827 | 0.408657 |
| CG12605 | 8934.627 | 0.831352 | 0.482401 | 1.723361 | 0.084823 | 0.408657 |
| Slip1 | 7263.355 | 0.896316 | 0.520242 | 1.722885 | 0.084909 | 0.408808 |
| CG4587 | 5273.268 | 1.046921 | 0.608101 | 1.721624 | 0.085138 | 0.409661 |
| GlnRS | 1439.117 | -1.7834 | 1.036804 | -1.72009 | 0.085416 | 0.410754 |
| RpII15 | 1646.992 | 1.54108 | 0.896577 | 1.718849 | 0.085642 | 0.41096 |
| GstE7 | 1498.172 | 2.147755 | 1.249617 | 1.71873 | 0.085664 | 0.41096 |
| Prp19 | 793.5568 | -2.55942 | 1.488633 | -1.71931 | 0.085559 | 0.41096 |
| Cht6 | 432.4786 | 3.891876 | 2.264138 | 1.718922 | 0.085629 | 0.41096 |
| Rab23 | 243.3842 | -4.07839 | 2.374413 | -1.71764 | 0.085862 | 0.411653 |
| CG8097 | 138.6989 | 5.285984 | 3.078447 | 1.717094 | 0.085962 | 0.411653 |
| Ak1 | 9931.213 | 0.920448 | 0.53598 | 1.717318 | 0.085921 | 0.411653 |
| Kdm4A | 205.4587 | 5.524857 | 3.219177 | 1.716233 | 0.086119 | 0.411726 |
| CG15891 | 1853.25 | 1.298829 | 0.756819 | 1.716169 | 0.086131 | 0.411726 |
| tsr | 4640.557 | 1.063009 | 0.619386 | 1.716229 | 0.08612 | 0.411726 |
| Prat | 158.6672 | 6.081901 | 3.544767 | 1.715741 | 0.086209 | 0.411856 |
| CG14647 | 1546.032 | 1.748627 | 1.020355 | 1.713745 | 0.086576 | 0.412377 |
| CG1105 | 1153.689 | -2.21176 | 1.290368 | -1.71405 | 0.086519 | 0.412377 |
| Pkc53E | 7422.172 | 0.858633 | 0.50096 | 1.713976 | 0.086533 | 0.412377 |
| prel | 2463.407 | 1.557677 | 0.908783 | 1.714025 | 0.086524 | 0.412377 |
| Ptp10D | 3599.312 | 1.271102 | 0.741694 | 1.713781 | 0.086569 | 0.412377 |
| CG7744 | 67.17347 | -6.34244 | 3.701694 | -1.71339 | 0.086641 | 0.412444 |
| CG14186 | 6177.328 | 1.013516 | 0.591646 | 1.713045 | 0.086704 | 0.4125 |
| Dnz1 | 502.0743 | 2.218234 | 1.295725 | 1.711964 | 0.086903 | 0.412987 |
| CG7084 | 14195.49 | -0.91023 | 0.531783 | -1.71165 | 0.086961 | 0.412987 |
| Hr39 | 2420.157 | 1.257323 | 0.73454 | 1.711715 | 0.086949 | 0.412987 |
| CG33521 | 705.6907 | 1.936022 | 1.131479 | 1.711054 | 0.087071 | 0.413266 |
| Sh3beta | 3187.388 | 1.234238 | 0.721563 | 1.710506 | 0.087172 | 0.413501 |
| sage | 728.0742 | 3.134825 | 1.834005 | 1.709278 | 0.0874 | 0.413846 |
| CG31223 | 1044.309 | -2.44407 | 1.429682 | -1.70952 | 0.087354 | 0.413846 |
| c11.1 | 926.3628 | -2.05636 | 1.202932 | -1.70945 | 0.087367 | 0.413846 |
| SelR | 8293.503 | -0.82008 | 0.479916 | -1.70879 | 0.08749 | 0.414031 |
| l(2)37Cd | 367.5942 | 3.566531 | 2.087873 | 1.708212 | 0.087597 | 0.414292 |
| AdoR | 632.4936 | -2.03437 | 1.193419 | -1.70466 | 0.088258 | 0.41626 |
| Gadd45 | 2588.565 | 2.347966 | 1.37728 | 1.704785 | 0.088235 | 0.41626 |

| | | | | | | |
|---|---|---|---|---|---|---|
| mim | 7375.902 | 1.07272 | 0.629316 | 1.704582 | 0.088272 | 0.41626 |
| CG43115 | 265.6082 | -4.44697 | 2.608101 | -1.70506 | 0.088183 | 0.41626 |
| lncRNA:CR | 648.5985 | 2.956856 | 1.733922 | 1.705299 | 0.088139 | 0.41626 |
| smog | 4963.577 | 1.127768 | 0.661726 | 1.704284 | 0.088328 | 0.416278 |
| Nepl20 | 109.0726 | 5.439146 | 3.193466 | 1.703211 | 0.088529 | 0.416568 |
| CG7246 | 469.5489 | 3.339129 | 1.960436 | 1.703259 | 0.08852 | 0.416568 |
| Gale | 665.5568 | -2.62585 | 1.541788 | -1.70312 | 0.088545 | 0.416568 |
| Git | 806.0663 | 2.407655 | 1.414313 | 1.70235 | 0.08869 | 0.417003 |
| Oscillin | 1833.208 | -1.48586 | 0.872987 | -1.70204 | 0.088748 | 0.41703 |
| CG7966 | 768.7711 | -2.22575 | 1.309144 | -1.70016 | 0.089101 | 0.418035 |
| WASp | 1421.6 | -1.51277 | 0.889586 | -1.70053 | 0.089031 | 0.418035 |
| uex | 8204.369 | 0.969429 | 0.570228 | 1.700071 | 0.089118 | 0.418035 |
| tapas | 3252.103 | -1.06317 | 0.625581 | -1.6995 | 0.089225 | 0.418294 |
| SkpA | 9361.337 | 0.937366 | 0.552051 | 1.69797 | 0.089513 | 0.419183 |
| Rac2 | 1673.499 | -1.5573 | 0.917172 | -1.69794 | 0.089519 | 0.419183 |
| CG34242 | 1335.892 | -2.17278 | 1.280805 | -1.69642 | 0.089807 | 0.420041 |
| Tl | 6301.772 | 0.932252 | 0.549515 | 1.696498 | 0.089792 | 0.420041 |
| sty | 8115.958 | -0.86096 | 0.50787 | -1.69524 | 0.09003 | 0.42084 |
| LManI | 978.0983 | -2.223 | 1.31204 | -1.69431 | 0.090206 | 0.421173 |
| CG6791 | 713.9972 | 2.793568 | 1.648561 | 1.694549 | 0.090161 | 0.421173 |
| Cdk12 | 2363.047 | 1.423316 | 0.840333 | 1.693753 | 0.090312 | 0.421422 |
| Appl | 21773.53 | 0.764715 | 0.451573 | 1.69345 | 0.09037 | 0.421446 |
| CG5618 | 1262.18 | -1.49129 | 0.881262 | -1.69222 | 0.090604 | 0.422294 |
| Hsc70-5 | 3293.242 | -1.33333 | 0.788299 | -1.6914 | 0.090761 | 0.422777 |
| CG5946 | 8319.817 | 0.857573 | 0.507295 | 1.69048 | 0.090936 | 0.423349 |
| CG44245 | 1600.986 | 2.285786 | 1.352628 | 1.689885 | 0.09105 | 0.423633 |
| CG31717 | 2297.849 | 1.459818 | 0.864576 | 1.688478 | 0.091319 | 0.423707 |
| lncRNA:CR | 325.2626 | -3.41402 | 2.021766 | -1.68863 | 0.09129 | 0.423707 |
| CG2034 | 363.1803 | -3.80177 | 2.250662 | -1.68918 | 0.091185 | 0.423707 |
| wgn | 1212.215 | -1.51678 | 0.898039 | -1.689 | 0.09122 | 0.423707 |
| CG7668 | 1112.388 | 1.664422 | 0.985945 | 1.688149 | 0.091383 | 0.423707 |
| AIF | 1322.918 | -1.77664 | 1.052347 | -1.68827 | 0.09136 | 0.423707 |
| Ranbp9 | 143.9602 | 4.867092 | 2.885718 | 1.686614 | 0.091678 | 0.424511 |
| Mpc1 | 2395.654 | -1.96745 | 1.166329 | -1.68688 | 0.091627 | 0.424511 |
| lncRNA:CR | 95.51997 | 5.825476 | 3.454343 | 1.686421 | 0.091715 | 0.424511 |
| ph-d | 704.2687 | 2.373731 | 1.408997 | 1.684695 | 0.092047 | 0.425805 |
| CG10877 | 185.9406 | -3.79272 | 2.251932 | -1.68421 | 0.092141 | 0.425994 |
| Acox57D-p | 771.6539 | 3.311368 | 1.966819 | 1.683617 | 0.092256 | 0.426278 |
| CG43778 | 6225.984 | 0.86271 | 0.512643 | 1.682868 | 0.092401 | 0.426701 |
| Gs1 | 8193.521 | 1.027924 | 0.610932 | 1.68255 | 0.092462 | 0.42674 |
| RYa-R | 159.2199 | 5.054892 | 3.006156 | 1.681514 | 0.092663 | 0.427422 |
| CG3568 | 361.5026 | -3.34956 | 1.994819 | -1.67913 | 0.093127 | 0.428518 |
| CG10137 | 766.3777 | 2.472829 | 1.472869 | 1.67892 | 0.093168 | 0.428518 |
| spas | 992.9759 | -2.28556 | 1.361067 | -1.67924 | 0.093105 | 0.428518 |
| ave | 2018.581 | 1.679146 | 0.999578 | 1.679854 | 0.092986 | 0.428518 |
| Kr | 1122.76 | -2.06216 | 1.228103 | -1.67914 | 0.093124 | 0.428518 |
| wake | 1230.07 | -1.62263 | 0.967204 | -1.67765 | 0.093415 | 0.429408 |
| PSMG1 | 857.2849 | 2.576944 | 1.537208 | 1.676379 | 0.093664 | 0.430183 |
| CG4424 | 368.3491 | -3.16477 | 1.888014 | -1.67624 | 0.093691 | 0.430183 |
| t-cup | 290.8421 | -3.46519 | 2.067599 | -1.67595 | 0.093748 | 0.430202 |

| | | | | | | |
|---|---|---|---|---|---|---|
| CG11790 | 6393.447 | 0.879395 | 0.524831 | 1.675577 | 0.093821 | 0.43029 |
| Yif1 | 1732.974 | -2.0605 | 1.23048 | -1.67455 | 0.094022 | 0.430474 |
| CG30373 | 791.3475 | 1.979715 | 1.182195 | 1.67461 | 0.094011 | 0.430474 |
| Urm1 | 532.9823 | 2.918433 | 1.742304 | 1.675043 | 0.093926 | 0.430474 |
| CG15073 | 669.9093 | -2.73597 | 1.634351 | -1.67404 | 0.094122 | 0.430687 |
| CG34266 | 136.3872 | -5.39989 | 3.232269 | -1.67062 | 0.094797 | 0.433438 |
| pHCl-1 | 3429.378 | 1.222012 | 0.731549 | 1.670445 | 0.094831 | 0.433438 |
| Taf1 | 1568.181 | 1.481527 | 0.8873 | 1.669702 | 0.094978 | 0.433862 |
| Edem1 | 683.2102 | -2.45507 | 1.472812 | -1.66693 | 0.095529 | 0.436128 |
| CG8916 | 362.7631 | 3.628966 | 2.178837 | 1.665552 | 0.095803 | 0.437131 |
| RabX4 | 2389.966 | 1.184325 | 0.711403 | 1.664775 | 0.095958 | 0.437589 |
| CG14868 | 2208.504 | 1.71724 | 1.03181 | 1.664299 | 0.096053 | 0.437774 |
| CG6567 | 67.14641 | 6.378528 | 3.836572 | 1.662559 | 0.096401 | 0.437868 |
| CG12662 | 137.0833 | 4.95136 | 2.977935 | 1.662683 | 0.096376 | 0.437868 |
| Prosalpha7 | 1484.364 | -2.03033 | 1.220569 | -1.66343 | 0.096227 | 0.437868 |
| CG14043 | 1208.511 | -1.62761 | 0.978528 | -1.66333 | 0.096247 | 0.437868 |
| FeCH | 2000.702 | -2.12947 | 1.279897 | -1.66378 | 0.096157 | 0.437868 |
| RhoGAP68 | 2436.072 | -1.7547 | 1.055365 | -1.66265 | 0.096384 | 0.437868 |
| Nup62 | 439.7779 | 3.074675 | 1.850677 | 1.661378 | 0.096638 | 0.438695 |
| Pask | 1193.962 | 1.916117 | 1.154127 | 1.66023 | 0.096868 | 0.439494 |
| CG13933 | 1280.758 | -2.07542 | 1.251024 | -1.65898 | 0.09712 | 0.440388 |
| Rnf146 | 566.9266 | -2.69243 | 1.62385 | -1.65805 | 0.097307 | 0.440986 |
| Exn | 1450.239 | 1.484592 | 0.89562 | 1.657613 | 0.097396 | 0.441139 |
| nSMase | 958.2528 | -2.58805 | 1.561674 | -1.65723 | 0.097474 | 0.441243 |
| AP-1sigma | 7638.456 | 0.986213 | 0.5954 | 1.656388 | 0.097643 | 0.441762 |
| CG2540 | 700.645 | 2.317823 | 1.399594 | 1.656069 | 0.097708 | 0.441805 |
| RpII18 | 2540.391 | 1.137924 | 0.68772 | 1.654632 | 0.097999 | 0.442125 |
| CG15270 | 4964.685 | 1.21603 | 0.734701 | 1.655136 | 0.097897 | 0.442125 |
| Mps1 | 65.12655 | -6.29604 | 3.804073 | -1.65508 | 0.097909 | 0.442125 |
| lncRNA:CR | 876.5326 | 1.912534 | 1.155686 | 1.65489 | 0.097947 | 0.442125 |
| Jra | 1270.469 | 1.961908 | 1.186391 | 1.653678 | 0.098193 | 0.442503 |
| Chrac-16 | 172.7549 | 3.537574 | 2.138944 | 1.653889 | 0.09815 | 0.442503 |
| CG4364 | 1688.882 | -1.46115 | 0.883885 | -1.6531 | 0.098311 | 0.442786 |
| CG7718 | 845.2252 | 2.923294 | 1.771335 | 1.650333 | 0.098875 | 0.443505 |
| CG4553 | 151.1856 | 4.563251 | 2.765194 | 1.650246 | 0.098893 | 0.443505 |
| CG9686 | 35266.09 | -0.94104 | 0.569938 | -1.65113 | 0.098713 | 0.443505 |
| Syn | 18924.17 | 0.870227 | 0.527111 | 1.650937 | 0.098752 | 0.443505 |
| GC | 1190.068 | -2.4248 | 1.469213 | -1.65041 | 0.098859 | 0.443505 |
| CG32640 | 139.3121 | 5.051592 | 3.059281 | 1.651235 | 0.098691 | 0.443505 |
| CG1764 | 1764.902 | -1.44805 | 0.877489 | -1.65022 | 0.098897 | 0.443505 |
| CG5290 | 116.5576 | -5.62879 | 3.411082 | -1.65015 | 0.098913 | 0.443505 |
| Cyp4ac3 | 689.6547 | -2.59568 | 1.573722 | -1.64939 | 0.099068 | 0.443952 |
| bves | 3212.948 | 1.375033 | 0.833822 | 1.649073 | 0.099133 | 0.443994 |
| Hip14 | 640.0183 | -2.54893 | 1.546383 | -1.64832 | 0.099288 | 0.44444 |
| La | 1427.991 | 1.614979 | 0.980326 | 1.647389 | 0.099478 | 0.445045 |
| CenG1A | 1130.282 | -2.00698 | 1.218692 | -1.64683 | 0.099593 | 0.445312 |
| Dsp1 | 12166.96 | 0.743836 | 0.452023 | 1.645569 | 0.099852 | 0.446223 |
| Lnk | 3385.038 | -1.53557 | 0.933708 | -1.6446 | 0.100053 | 0.446536 |
| CIA30 | 1406.814 | 2.229892 | 1.356035 | 1.64442 | 0.100089 | 0.446536 |
| Girdin | 785.5866 | -1.90448 | 1.15789 | -1.64479 | 0.100013 | 0.446536 |

| | | | | | | |
|---|---|---|---|---|---|---|
| Dis3 | 275.3901 | -3.49664 | 2.126972 | -1.64395 | 0.100186 | 0.446717 |
| step | 4356.92 | -1.0469 | 0.636959 | -1.6436 | 0.100259 | 0.446797 |
| Tsen54 | 445.149 | -3.36233 | 2.046092 | -1.64329 | 0.100322 | 0.44683 |
| Rab6 | 1701.274 | 1.284035 | 0.781991 | 1.642008 | 0.100588 | 0.446831 |
| Rpn13 | 1502.617 | -1.50966 | 0.919237 | -1.64229 | 0.100529 | 0.446831 |
| Mef2 | 4210.46 | 0.997374 | 0.607322 | 1.642249 | 0.100538 | 0.446831 |
| CG42808 | 945.4676 | -2.2057 | 1.343034 | -1.64232 | 0.100523 | 0.446831 |
| Ice1 | 226.9675 | -4.36888 | 2.66079 | -1.64195 | 0.100601 | 0.446831 |
| GlcT | 761.7291 | -2.37987 | 1.450205 | -1.64106 | 0.100785 | 0.447155 |
| ara | 224.0285 | 4.742649 | 2.889714 | 1.641217 | 0.100752 | 0.447155 |
| Ravus | 658.8213 | 1.921484 | 1.171227 | 1.640573 | 0.100886 | 0.447355 |
| Rpt2 | 1338.952 | 1.530556 | 0.934617 | 1.63763 | 0.101499 | 0.449766 |
| Sytbeta | 1038.658 | -2.1617 | 1.320305 | -1.63727 | 0.101573 | 0.449766 |
| CG14968 | 185.4094 | -3.19071 | 1.948937 | -1.63716 | 0.101598 | 0.449766 |
| Mppe | 1734.948 | -2.32557 | 1.421229 | -1.63631 | 0.101775 | 0.450299 |
| Crag | 928.5932 | -1.98032 | 1.210731 | -1.63564 | 0.101914 | 0.450459 |
| CG16935 | 1371.265 | -1.51387 | 0.925574 | -1.6356 | 0.101923 | 0.450459 |
| kek4 | 2643.414 | 1.088244 | 0.665507 | 1.635211 | 0.102005 | 0.450573 |
| EMC8-9 | 2837.198 | -1.28866 | 0.788411 | -1.6345 | 0.102153 | 0.45098 |
| sofe | 285.2035 | 4.74687 | 2.905685 | 1.633649 | 0.102333 | 0.451523 |
| cm | 1250.54 | 1.721343 | 1.054558 | 1.632289 | 0.102619 | 0.452537 |
| CG6511 | 92.83817 | 5.255734 | 3.221109 | 1.631654 | 0.102752 | 0.452878 |
| Rab8 | 2371.584 | -1.14051 | 0.700389 | -1.62839 | 0.103442 | 0.455669 |
| CG11444 | 1421.768 | 1.788686 | 1.098978 | 1.62759 | 0.103612 | 0.455945 |
| Ptp4E | 2943.217 | -1.12317 | 0.69017 | -1.62739 | 0.103655 | 0.455945 |
| CG42489 | 510.1235 | -2.96992 | 1.825071 | -1.62729 | 0.103675 | 0.455945 |
| Sap130 | 812.6738 | -2.66398 | 1.638195 | -1.62617 | 0.103914 | 0.456745 |
| Srrm234 | 33631.08 | 0.693726 | 0.426816 | 1.625353 | 0.104087 | 0.457057 |
| Meltrin | 1418.118 | 1.653918 | 1.01761 | 1.625297 | 0.104099 | 0.457057 |
| Vps2 | 1436.912 | -1.9981 | 1.229794 | -1.62475 | 0.104216 | 0.457322 |
| MED18 | 560.5644 | -2.50519 | 1.542835 | -1.62375 | 0.104428 | 0.457374 |
| CG17278 | 470.9252 | 2.811907 | 1.731912 | 1.623586 | 0.104464 | 0.457374 |
| beat-IV | 1512.693 | -1.88068 | 1.158516 | -1.62336 | 0.104513 | 0.457374 |
| CG30183 | 241.8088 | -4.2553 | 2.620136 | -1.62408 | 0.10436 | 0.457374 |
| Pdi | 7599.764 | -0.8473 | 0.521637 | -1.62431 | 0.104309 | 0.457374 |
| Eps-15 | 4885.641 | 1.099962 | 0.677944 | 1.622496 | 0.104697 | 0.457929 |
| CG6123 | 1812.059 | 1.242816 | 0.766472 | 1.621478 | 0.104915 | 0.458618 |
| CG14340 | 217.6289 | 4.072199 | 2.5118 | 1.621227 | 0.104969 | 0.458618 |
| Su(fu) | 485.5843 | 2.844322 | 1.756091 | 1.619689 | 0.105299 | 0.458811 |
| CG31357 | 2062.906 | -2.08908 | 1.289601 | -1.61994 | 0.105245 | 0.458811 |
| PIG-X | 1170.741 | -2.38799 | 1.473717 | -1.62039 | 0.105149 | 0.458811 |
| Ptp36E | 1342.21 | -1.44054 | 0.888981 | -1.62044 | 0.105138 | 0.458811 |
| lncRNA:CR | 142.1894 | 4.684732 | 2.891656 | 1.620086 | 0.105214 | 0.458811 |
| rasp | 206.3255 | -4.35765 | 2.691512 | -1.61903 | 0.10544 | 0.459177 |
| Su(H) | 125.9258 | 4.243204 | 2.62233 | 1.618105 | 0.10564 | 0.459798 |
| CG34325 | 311.3529 | 3.687943 | 2.279646 | 1.61777 | 0.105712 | 0.459862 |
| CG5377 | 675.0553 | 3.275541 | 2.026742 | 1.616161 | 0.10606 | 0.461124 |
| lwr | 3930.502 | -1.16572 | 0.722126 | -1.61429 | 0.106464 | 0.46213 |
| CCT2 | 1405.035 | -1.8913 | 1.171566 | -1.61434 | 0.106454 | 0.46213 |
| pck | 475.9226 | 2.618757 | 1.622165 | 1.614359 | 0.10645 | 0.46213 |

| | | | | | | |
|---|---|---|---|---|---|---|
| AstC-R2 | 136.0003 | -4.97071 | 3.079752 | -1.614 | 0.106528 | 0.462158 |
| Nepl10 | 704.528 | -3.24632 | 2.011912 | -1.61355 | 0.106625 | 0.462329 |
| CG43093 | 391.2353 | 2.703421 | 1.676183 | 1.612843 | 0.106779 | 0.462496 |
| dbo | 2274.525 | 1.559854 | 0.967118 | 1.612888 | 0.106769 | 0.462496 |
| lncRNA:CR | 449.0199 | 2.565934 | 1.591399 | 1.612376 | 0.10688 | 0.462686 |
| mAcon1 | 13104.83 | 0.708821 | 0.439704 | 1.612043 | 0.106953 | 0.46275 |
| CG8043 | 487.6947 | -2.82819 | 1.757322 | -1.60938 | 0.107534 | 0.463981 |
| egg | 440.4261 | 2.833191 | 1.760059 | 1.609713 | 0.10746 | 0.463981 |
| Nop60B | 606.0103 | 2.562009 | 1.592153 | 1.609148 | 0.107584 | 0.463981 |
| norpA | 16915.57 | 0.713823 | 0.44356 | 1.609305 | 0.10755 | 0.463981 |
| cN-IIIB | 1256.446 | -1.86408 | 1.157955 | -1.6098 | 0.107441 | 0.463981 |
| Eno | 30156.41 | -0.66826 | 0.415093 | -1.60991 | 0.107417 | 0.463981 |
| MrgBP | 984.4724 | -2.46993 | 1.53532 | -1.60874 | 0.107674 | 0.464119 |
| Ndfip | 2648.77 | 1.112845 | 0.692781 | 1.606344 | 0.108198 | 0.46613 |
| Cap-H2 | 471.2333 | -2.13506 | 1.329618 | -1.60577 | 0.108325 | 0.466355 |
| l(2)SH0834 | 2826.431 | -1.05799 | 0.658944 | -1.60558 | 0.108367 | 0.466355 |
| SMC3 | 989.4937 | 1.567441 | 0.977131 | 1.604125 | 0.108686 | 0.46748 |
| Nnp-1 | 461.5936 | 2.918855 | 1.819934 | 1.603825 | 0.108753 | 0.467514 |
| CG9518 | 78.7764 | -6.19933 | 3.868604 | -1.60247 | 0.109051 | 0.468548 |
| CG3955 | 3615.025 | -0.97183 | 0.60683 | -1.6015 | 0.109267 | 0.469224 |
| kon | 370.4755 | -2.7414 | 1.712473 | -1.60084 | 0.109412 | 0.469594 |
| Art1 | 1265.735 | 2.207107 | 1.379026 | 1.600483 | 0.109492 | 0.469684 |
| KrT95D | 7957.892 | 0.800639 | 0.501024 | 1.598005 | 0.110042 | 0.471212 |
| CG12075 | 95.93914 | 4.673222 | 2.923632 | 1.59843 | 0.109947 | 0.471212 |
| ste24a | 2033.735 | -1.82711 | 1.143294 | -1.59811 | 0.110019 | 0.471212 |
| Cals | 39568.34 | 0.64173 | 0.401628 | 1.597822 | 0.110083 | 0.471212 |
| Pomp | 3232.537 | 1.723893 | 1.07923 | 1.597336 | 0.110191 | 0.471221 |
| Cow | 13741.83 | 0.712183 | 0.44587 | 1.597286 | 0.110202 | 0.471221 |
| CG11317 | 5210.328 | 0.951654 | 0.596851 | 1.594458 | 0.110834 | 0.473204 |
| CG9410 | 1465.753 | 1.754337 | 1.1003 | 1.594417 | 0.110843 | 0.473204 |
| svr | 15276.47 | 0.864467 | 0.542012 | 1.594922 | 0.11073 | 0.473204 |
| CG1307 | 2535.828 | -1.23161 | 0.773199 | -1.59287 | 0.111189 | 0.47443 |
| Syx4 | 2692.018 | 1.2124 | 0.761884 | 1.591318 | 0.111538 | 0.475063 |
| Fas3 | 1065.148 | 1.631031 | 1.024929 | 1.59136 | 0.111529 | 0.475063 |
| tsu | 1095.134 | 1.951777 | 1.226324 | 1.591567 | 0.111482 | 0.475063 |
| CG5877 | 540.0106 | 2.241357 | 1.408829 | 1.590936 | 0.111624 | 0.475063 |
| CG14074 | 358.4236 | 3.123703 | 1.963488 | 1.590895 | 0.111633 | 0.475063 |
| CG31436 | 738.1626 | -2.54347 | 1.59931 | -1.59035 | 0.111755 | 0.475079 |
| mRpL46 | 1391.321 | -1.42649 | 0.896847 | -1.59057 | 0.111707 | 0.475079 |
| mtTFB2 | 484.6938 | 3.594355 | 2.2614 | 1.589438 | 0.111961 | 0.475451 |
| Prosalpha4 | 1310.066 | 1.712333 | 1.077196 | 1.58962 | 0.11192 | 0.475451 |
| Srp68 | 997.0144 | -2.17627 | 1.369505 | -1.58909 | 0.11204 | 0.475532 |
| Ir51a | 159.4092 | 4.80178 | 3.022694 | 1.588576 | 0.112156 | 0.475774 |
| CG2201 | 1964.049 | -1.19333 | 0.751361 | -1.58822 | 0.112237 | 0.475864 |
| nAChRbeta | 4314.761 | -1.24284 | 0.782845 | -1.58759 | 0.112378 | 0.475962 |
| Sil1 | 859.2222 | -2.06902 | 1.303142 | -1.58772 | 0.112351 | 0.475962 |
| LTV1 | 4836.309 | 0.90402 | 0.5696 | 1.587114 | 0.112487 | 0.476171 |
| CG7414 | 3441.374 | 1.00012 | 0.630505 | 1.586222 | 0.112689 | 0.476775 |
| wry | 10105.83 | -0.73938 | 0.466531 | -1.58485 | 0.113001 | 0.477591 |
| CG43056 | 273.6259 | 3.361332 | 2.120585 | 1.585097 | 0.112944 | 0.477591 |

| | | | | | | |
|---|---|---|---|---|---|---|
| Tsp96F | 2990.014 | -1.35491 | 0.855907 | -1.58301 | 0.11342 | 0.478317 |
| elav | 9052.451 | 0.787604 | 0.497607 | 1.582785 | 0.113471 | 0.478317 |
| PIG-S | 290.5485 | -3.47015 | 2.192354 | -1.58284 | 0.113457 | 0.478317 |
| PHGPx | 5849.002 | -0.82176 | 0.518966 | -1.58346 | 0.113316 | 0.478317 |
| fry | 2405.447 | -1.24027 | 0.783165 | -1.58366 | 0.113271 | 0.478317 |
| Lsm11 | 265.0145 | -3.02734 | 1.913184 | -1.58236 | 0.113568 | 0.478478 |
| CG5969 | 1028.368 | 1.948031 | 1.231472 | 1.581871 | 0.113679 | 0.478692 |
| CG17378 | 1980.419 | 1.265575 | 0.800589 | 1.580805 | 0.113923 | 0.478752 |
| lgs | 5240.13 | 0.947898 | 0.599645 | 1.580765 | 0.113932 | 0.478752 |
| stops | 371.4909 | -2.73781 | 1.731594 | -1.58109 | 0.113857 | 0.478752 |
| MKP-4 | 3089.066 | 1.91535 | 1.211464 | 1.581021 | 0.113873 | 0.478752 |
| Tnks | 1122.783 | 2.432126 | 1.539379 | 1.57994 | 0.114121 | 0.479251 |
| Aldh7A1 | 103.1596 | -4.97194 | 3.147864 | -1.57946 | 0.11423 | 0.479251 |
| CG5144 | 180.7383 | -4.55753 | 2.885249 | -1.5796 | 0.114199 | 0.479251 |
| Dop1R1 | 19414.91 | -0.66057 | 0.418449 | -1.57861 | 0.114426 | 0.479572 |
| Tmem63 | 1683.705 | 2.051502 | 1.299512 | 1.57867 | 0.114412 | 0.479572 |
| Cyp49a1 | 699.2814 | 2.298754 | 1.456606 | 1.578157 | 0.114529 | 0.479757 |
| CG7133 | 606.4555 | -2.97343 | 1.884884 | -1.57751 | 0.114678 | 0.480127 |
| CG2906 | 642.9965 | 3.372478 | 2.138287 | 1.577187 | 0.114753 | 0.48019 |
| CG9215 | 320.2362 | -3.59108 | 2.277589 | -1.5767 | 0.114864 | 0.480406 |
| CG32369 | 2282.115 | 1.11731 | 0.709033 | 1.575822 | 0.115067 | 0.481003 |
| viaf | 2158.001 | 1.259268 | 0.799298 | 1.575467 | 0.115149 | 0.481095 |
| GluRIIA | 162.2427 | 4.659235 | 2.959276 | 1.574451 | 0.115383 | 0.481425 |
| a | 2518.998 | 1.068329 | 0.678524 | 1.57449 | 0.115374 | 0.481425 |
| CG13698 | 1063.716 | -2.38349 | 1.513958 | -1.57435 | 0.115408 | 0.481425 |
| scat | 450.4966 | 2.696071 | 1.713363 | 1.573555 | 0.11559 | 0.481937 |
| Ncoa6 | 2649.567 | 1.090133 | 0.693343 | 1.572285 | 0.115884 | 0.482392 |
| agt | 802.4482 | -2.61316 | 1.662268 | -1.57205 | 0.11594 | 0.482392 |
| RagA-B | 1420.986 | -2.15298 | 1.369532 | -1.57206 | 0.115937 | 0.482392 |
| Alg2 | 636.7587 | 2.666717 | 1.695658 | 1.572674 | 0.115794 | 0.482392 |
| Tpr2 | 4849.265 | -1.03771 | 0.660345 | -1.57147 | 0.116074 | 0.482412 |
| CG4293 | 302.1314 | -3.06374 | 1.949581 | -1.57149 | 0.116069 | 0.482412 |
| Ugt317A1 | 385.3954 | -3.55761 | 2.26464 | -1.57094 | 0.116197 | 0.482412 |
| Sac1 | 2116.663 | -1.15272 | 0.733644 | -1.57122 | 0.116131 | 0.482412 |
| eIF5B | 1486.285 | 1.349367 | 0.85907 | 1.570731 | 0.116245 | 0.482412 |
| CG1824 | 956.4565 | -1.98446 | 1.264116 | -1.56984 | 0.116452 | 0.48302 |
| CdGAPr | 2214.09 | 1.220948 | 0.778559 | 1.568214 | 0.116831 | 0.483843 |
| car | 280.3999 | 2.861307 | 1.824528 | 1.568245 | 0.116824 | 0.483843 |
| CG4022 | 470.1736 | -2.47326 | 1.577056 | -1.56828 | 0.116817 | 0.483843 |
| CG16734 | 275.2178 | -3.91211 | 2.49578 | -1.56749 | 0.117001 | 0.484295 |
| S1P | 1039.83 | 2.435525 | 1.554341 | 1.566918 | 0.117134 | 0.484475 |
| ash1 | 1265.69 | -1.39909 | 0.892972 | -1.56678 | 0.117165 | 0.484475 |
| CG11417 | 477.2987 | -2.50164 | 1.597236 | -1.56623 | 0.117294 | 0.484511 |
| GatB | 365.5693 | 3.439107 | 2.195572 | 1.566383 | 0.117259 | 0.484511 |
| EMC3 | 2448.06 | -1.09181 | 0.69725 | -1.56587 | 0.117378 | 0.484607 |
| CG16779 | 8499.432 | 0.768282 | 0.490751 | 1.565522 | 0.11746 | 0.484698 |
| slp2 | 108.7838 | -5.45234 | 3.483751 | -1.56508 | 0.117564 | 0.484878 |
| dao | 1920.911 | -1.25343 | 0.801055 | -1.56472 | 0.117648 | 0.484971 |
| ohgt | 2390.264 | -1.71241 | 1.094761 | -1.56419 | 0.117774 | 0.485242 |
| Pus1 | 760.3401 | 2.426302 | 1.551669 | 1.563672 | 0.117895 | 0.485491 |

| | | | | | | |
|---|---|---|---|---|---|---|
| CG33213 | 100.8003 | 5.78296 | 3.699737 | 1.563073 | 0.118035 | 0.485821 |
| CG11997 | 312.0247 | -3.52563 | 2.259155 | -1.5606 | 0.118619 | 0.485901 |
| CG14721 | 954.4637 | 2.121769 | 1.359069 | 1.561193 | 0.118478 | 0.485901 |
| spn-B | 67.23171 | 6.027094 | 3.862749 | 1.560312 | 0.118686 | 0.485901 |
| CG3631 | 518.8128 | -2.91483 | 1.868586 | -1.55991 | 0.118781 | 0.485901 |
| CG5412 | 355.957 | 2.294481 | 1.469625 | 1.56127 | 0.11846 | 0.485901 |
| PTPMT1 | 322.4637 | -3.19153 | 2.042561 | -1.56251 | 0.118167 | 0.485901 |
| RNaseP:RI | 479.2452 | -2.08043 | 1.332729 | -1.56103 | 0.118517 | 0.485901 |
| Cyp6a20 | 1902.673 | -1.30597 | 0.836392 | -1.56144 | 0.11842 | 0.485901 |
| Prp8 | 3053.256 | 1.486386 | 0.952733 | 1.560129 | 0.11873 | 0.485901 |
| CG14969 | 1418.647 | 1.525325 | 0.977077 | 1.56111 | 0.118498 | 0.485901 |
| Drak | 1964.296 | 1.158061 | 0.741831 | 1.561085 | 0.118504 | 0.485901 |
| CG6891 | 203.7443 | -3.88235 | 2.488816 | -1.55992 | 0.118779 | 0.485901 |
| CG8149 | 1017.116 | -2.24292 | 1.438558 | -1.55914 | 0.118963 | 0.486222 |
| CG43394 | 642.84 | -2.44173 | 1.56615 | -1.55907 | 0.118981 | 0.486222 |
| CG1620 | 1766.278 | -2.01268 | 1.29165 | -1.55822 | 0.119181 | 0.486791 |
| MICU1 | 1376.206 | -1.45934 | 0.937116 | -1.55727 | 0.119406 | 0.486879 |
| CG30122 | 1854.135 | 1.287448 | 0.82682 | 1.557108 | 0.119445 | 0.486879 |
| Ptp69D | 1950.117 | -1.78148 | 1.143979 | -1.55727 | 0.119406 | 0.486879 |
| CG4887 | 3277.813 | -1.1259 | 0.722795 | -1.5577 | 0.119304 | 0.486879 |
| CG32262 | 1970.121 | 1.549924 | 0.99698 | 1.554619 | 0.120037 | 0.489044 |
| mRpL28 | 2107.031 | -1.2132 | 0.780579 | -1.55424 | 0.120128 | 0.489166 |
| CG8032 | 1637.732 | 1.696512 | 1.092356 | 1.553076 | 0.120405 | 0.489893 |
| CAHbeta | 113.8404 | -6.00579 | 3.867991 | -1.55269 | 0.120497 | 0.489893 |
| CG3040 | 1223.173 | 1.644841 | 1.059428 | 1.552575 | 0.120525 | 0.489893 |
| CG10257 | 684.7073 | 2.676815 | 1.724233 | 1.552467 | 0.120551 | 0.489893 |
| unk | 3994.335 | 1.135746 | 0.73253 | 1.550442 | 0.121036 | 0.491413 |
| CG2278 | 425.5499 | -2.73039 | 1.761096 | -1.55039 | 0.121047 | 0.491413 |
| robo3 | 1423.796 | -1.3773 | 0.88853 | -1.55009 | 0.12112 | 0.491459 |
| GAPsec | 70.48179 | 5.441931 | 3.511618 | 1.549693 | 0.121215 | 0.491598 |
| Schip1 | 2782.061 | -1.028 | 0.663826 | -1.54859 | 0.12148 | 0.491937 |
| Ipp | 4784.746 | -1.03361 | 0.667881 | -1.54759 | 0.12172 | 0.491937 |
| Gyf | 8309.189 | -0.83713 | 0.540937 | -1.54756 | 0.121728 | 0.491937 |
| CG9795 | 1633.769 | -1.61313 | 1.041966 | -1.54816 | 0.121583 | 0.491937 |
| CG42404 | 2010.858 | -1.60843 | 1.03905 | -1.54798 | 0.121626 | 0.491937 |
| CG46280 | 2842.81 | -1.01226 | 0.65398 | -1.54785 | 0.121659 | 0.491937 |
| nmo | 11628.4 | 0.753801 | 0.487008 | 1.54782 | 0.121666 | 0.491937 |
| CG10903 | 199.8217 | 3.700759 | 2.39217 | 1.54703 | 0.121856 | 0.492207 |
| MAPk-Ak2 | 1426.195 | 1.391709 | 0.900498 | 1.545488 | 0.122228 | 0.493165 |
| chif | 1222.284 | -1.41958 | 0.918466 | -1.5456 | 0.122202 | 0.493165 |
| sv | 430.9798 | -2.29984 | 1.488297 | -1.54528 | 0.122277 | 0.493165 |
| Tbc1d15-1 | 404.0663 | -2.80472 | 1.816454 | -1.54406 | 0.122574 | 0.493615 |
| SLC22A | 3961.961 | 1.308075 | 0.84703 | 1.544309 | 0.122514 | 0.493615 |
| Ltn1 | 730.5543 | -1.76909 | 1.145736 | -1.54406 | 0.122573 | 0.493615 |
| tkv | 5249.007 | -0.84844 | 0.549805 | -1.54316 | 0.122791 | 0.493994 |
| prod | 1058.426 | -1.65584 | 1.073001 | -1.54318 | 0.122787 | 0.493994 |
| CG9853 | 1394.728 | -2.16407 | 1.403056 | -1.5424 | 0.122977 | 0.494298 |
| CG5027 | 2409.903 | -1.52388 | 0.988026 | -1.54235 | 0.122989 | 0.494298 |
| TfIIS | 622.0018 | -2.04609 | 1.330033 | -1.53837 | 0.123957 | 0.496323 |
| CG10435 | 1726.114 | -1.4558 | 0.94605 | -1.53882 | 0.123849 | 0.496323 |

| | | | | | | |
|---|---|---|---|---|---|---|
| CG15080 | 100.1636 | -5.66775 | 3.684182 | -1.5384 | 0.123951 | 0.496323 |
| Pabp2 | 769.8756 | -1.93991 | 1.260894 | -1.53852 | 0.123922 | 0.496323 |
| CG13404 | 507.358 | 2.062604 | 1.340879 | 1.538248 | 0.123988 | 0.496323 |
| Snap25 | 46482.19 | 0.637707 | 0.414466 | 1.538626 | 0.123896 | 0.496323 |
| pain | 1346.216 | -1.67555 | 1.088197 | -1.53975 | 0.123621 | 0.496323 |
| CG9119 | 325.8469 | -3.32993 | 2.16445 | -1.53846 | 0.123935 | 0.496323 |
| CG4025 | 806.7837 | -2.3506 | 1.529634 | -1.53671 | 0.124364 | 0.497582 |
| CG16865 | 520.1678 | 2.373902 | 1.546699 | 1.534818 | 0.124828 | 0.498941 |
| Orcokinin | 188.1804 | 4.348365 | 2.833142 | 1.534821 | 0.124828 | 0.498941 |
| CG44247 | 11072.77 | 0.889139 | 0.580009 | 1.532975 | 0.125282 | 0.500504 |
| Syt4 | 12650.43 | 0.719383 | 0.469616 | 1.531855 | 0.125558 | 0.501121 |
| c12.2 | 225.5692 | 3.491254 | 2.279119 | 1.531843 | 0.125561 | 0.501121 |
| CG14657 | 1589.229 | -1.2256 | 0.800277 | -1.53146 | 0.125655 | 0.501245 |
| Ciao1 | 813.981 | -1.98672 | 1.297874 | -1.53075 | 0.125831 | 0.501697 |
| CG9663 | 196.7174 | -3.54455 | 2.318025 | -1.52913 | 0.126233 | 0.503053 |
| CG1983 | 2675.205 | 1.244955 | 0.814455 | 1.528574 | 0.12637 | 0.503348 |
| qkr58E-2 | 1717.475 | 1.638822 | 1.072845 | 1.527548 | 0.126625 | 0.503731 |
| CG3402 | 2363.168 | -1.49609 | 0.979238 | -1.52781 | 0.12656 | 0.503731 |
| Taf9 | 333.0177 | -2.65443 | 1.737844 | -1.52743 | 0.126654 | 0.503731 |
| Idh3b | 7045.214 | 0.840546 | 0.550741 | 1.526209 | 0.126958 | 0.504688 |
| CG33298 | 1257.599 | -1.83754 | 1.204449 | -1.52563 | 0.127102 | 0.50501 |
| ver | 220.6708 | -3.28848 | 2.157484 | -1.52422 | 0.127453 | 0.506156 |
| CG17270 | 916.5104 | 1.925985 | 1.264264 | 1.523404 | 0.127658 | 0.506547 |
| CG11586 | 713.6747 | -2.20048 | 1.444526 | -1.52332 | 0.127678 | 0.506547 |
| CG43103 | 247.662 | -3.88211 | 2.548893 | -1.52306 | 0.127745 | 0.50656 |
| CG1662 | 1007.558 | -1.92305 | 1.262862 | -1.52277 | 0.127815 | 0.506591 |
| CG33914 | 149.0754 | 4.04018 | 2.654214 | 1.522176 | 0.127965 | 0.506684 |
| CG12177 | 9507.797 | -0.87551 | 0.575126 | -1.52229 | 0.127936 | 0.506684 |
| CG3652 | 2663.897 | 1.12321 | 0.739526 | 1.518823 | 0.128807 | 0.509766 |
| CG5037 | 4141.105 | -1.60557 | 1.057319 | -1.51853 | 0.128881 | 0.509806 |
| Vta1 | 1551.551 | -1.92475 | 1.268355 | -1.51751 | 0.129137 | 0.51048 |
| CG13455 | 198.5275 | -4.06444 | 2.678643 | -1.51735 | 0.129178 | 0.51048 |
| Coq6 | 1164.421 | -1.80617 | 1.190866 | -1.51669 | 0.129346 | 0.510891 |
| CG43149 | 548.3306 | -2.47263 | 1.63065 | -1.51634 | 0.129432 | 0.510981 |
| CG42233 | 466.6445 | 2.844288 | 1.876336 | 1.515873 | 0.129551 | 0.511199 |
| beat-Vc | 2841.173 | 1.157291 | 0.764052 | 1.514676 | 0.129855 | 0.511892 |
| asRNA:CR4 | 111.2839 | -4.23559 | 2.796356 | -1.51468 | 0.129853 | 0.511892 |
| TrpA1 | 161.342 | 5.293748 | 3.495815 | 1.51431 | 0.129947 | 0.512006 |
| CG7156 | 166.6577 | 3.773275 | 2.493453 | 1.513273 | 0.13021 | 0.512544 |
| CG7834 | 1927.506 | -1.17333 | 0.775358 | -1.51327 | 0.130211 | 0.512544 |
| bdl | 7932.612 | -0.76946 | 0.508909 | -1.51198 | 0.130539 | 0.513582 |
| Sgf29 | 1242.624 | -1.42162 | 0.940736 | -1.51118 | 0.130743 | 0.514132 |
| CG2493 | 2108.074 | 1.16084 | 0.768465 | 1.510596 | 0.130891 | 0.514248 |
| Naa15-16 | 865.0898 | 1.949476 | 1.290566 | 1.510559 | 0.130901 | 0.514248 |
| CG6923 | 1308.458 | 1.939878 | 1.285345 | 1.509228 | 0.131241 | 0.514375 |
| CG33108 | 590.9614 | -3.19894 | 2.119376 | -1.50938 | 0.131202 | 0.514375 |
| Sik3 | 2217.481 | -1.26835 | 0.840352 | -1.5093 | 0.131222 | 0.514375 |
| Pi3K59F | 692.9534 | -2.08491 | 1.381375 | -1.5093 | 0.131222 | 0.514375 |
| sturkopf | 820.8208 | 1.656497 | 1.097777 | 1.508956 | 0.13131 | 0.514375 |
| CG14613 | 463.4021 | 2.49507 | 1.65354 | 1.508926 | 0.131318 | 0.514375 |

| | | | | | | |
|---|---|---|---|---|---|---|
| Sar1 | 4056.974 | -1.02464 | 0.679263 | -1.50846 | 0.131436 | 0.514386 |
| CG2277 | 3599.349 | -1.08065 | 0.716417 | -1.50841 | 0.131449 | 0.514386 |
| H15 | 267.0048 | -3.41275 | 2.264185 | -1.50727 | 0.13174 | 0.514788 |
| Blm | 228.718 | -3.4901 | 2.315545 | -1.50725 | 0.131747 | 0.514788 |
| Gycalpha9 | 9384.103 | -0.74657 | 0.49548 | -1.50676 | 0.131872 | 0.514788 |
| RpL15 | 56965.82 | 0.612429 | 0.406186 | 1.507757 | 0.131617 | 0.514788 |
| CG6843 | 541.8271 | -2.13945 | 1.419673 | -1.507 | 0.13181 | 0.514788 |
| CG3862 | 443.185 | 3.214941 | 2.135395 | 1.505549 | 0.132183 | 0.515 |
| Rsf1 | 4152.281 | -0.97265 | 0.645982 | -1.5057 | 0.132145 | 0.515 |
| Fer2 | 323.1204 | 3.587492 | 2.38207 | 1.50604 | 0.132057 | 0.515 |
| Usp5 | 357.6421 | 2.730318 | 1.812984 | 1.50598 | 0.132072 | 0.515 |
| S6k | 1984.071 | -1.46269 | 0.97178 | -1.50516 | 0.132282 | 0.515135 |
| CG17493 | 6185.989 | 0.867454 | 0.576425 | 1.504887 | 0.132353 | 0.515163 |
| Arp3 | 1097.833 | 1.71834 | 1.14215 | 1.504478 | 0.132458 | 0.515323 |
| rdgB | 10751.56 | -0.73736 | 0.49044 | -1.50347 | 0.132717 | 0.516078 |
| PPO2 | 379.4702 | -2.39525 | 1.594026 | -1.50264 | 0.132931 | 0.51641 |
| sxc | 8421.767 | 0.725258 | 0.482615 | 1.502767 | 0.132899 | 0.51641 |
| Myo31DF | 77.39112 | 5.0096 | 3.334554 | 1.50233 | 0.133012 | 0.516475 |
| Sugb | 295.6368 | -3.10528 | 2.068078 | -1.50153 | 0.133219 | 0.51703 |
| CG31759 | 168.6084 | -4.22117 | 2.814344 | -1.49988 | 0.133647 | 0.518187 |
| CG41378 | 2584.114 | 1.022049 | 0.68133 | 1.500079 | 0.133594 | 0.518187 |
| CG10638 | 346.6404 | 3.093441 | 2.063594 | 1.499055 | 0.133859 | 0.518762 |
| Sk2 | 367.4236 | -3.15587 | 2.105609 | -1.49879 | 0.133928 | 0.518777 |
| CG14562 | 3875.627 | 0.901187 | 0.601747 | 1.497617 | 0.134233 | 0.519456 |
| CG34423 | 63.81692 | 4.548812 | 3.037198 | 1.4977 | 0.134211 | 0.519456 |
| CG14964 | 92.71105 | 4.72995 | 3.162028 | 1.495859 | 0.13469 | 0.520975 |
| CG12299 | 494.2367 | 2.466943 | 1.649917 | 1.495192 | 0.134864 | 0.521029 |
| cu | 1487.269 | -1.7314 | 1.157891 | -1.49531 | 0.134834 | 0.521029 |
| CG43845 | 808.4635 | 1.674335 | 1.119911 | 1.49506 | 0.134899 | 0.521029 |
| sprt | 621.9213 | 1.914588 | 1.280909 | 1.49471 | 0.13499 | 0.52113 |
| CG12948 | 359.0289 | 2.664384 | 1.784445 | 1.493116 | 0.135407 | 0.521382 |
| Blos1 | 2028.545 | 1.392636 | 0.932827 | 1.49292 | 0.135458 | 0.521382 |
| eIF4G2 | 3046.393 | 1.054188 | 0.705713 | 1.493792 | 0.13523 | 0.521382 |
| Gpat4 | 2991.157 | 1.193015 | 0.798854 | 1.493409 | 0.13533 | 0.521382 |
| DIP2 | 4072.507 | -0.87561 | 0.586126 | -1.4939 | 0.135201 | 0.521382 |
| CG7556 | 518.1243 | -2.36016 | 1.581108 | -1.49272 | 0.13551 | 0.521382 |
| CG11658 | 3394.529 | -1.37085 | 0.917888 | -1.49348 | 0.135311 | 0.521382 |
| ics | 1237.579 | 1.3306 | 0.892565 | 1.490761 | 0.136024 | 0.52286 |
| CG1309 | 1007.303 | 2.074193 | 1.39118 | 1.49096 | 0.135972 | 0.52286 |
| stj | 11283.22 | 0.692699 | 0.46487 | 1.490091 | 0.1362 | 0.523286 |
| Nab2 | 885.7175 | 1.721275 | 1.155553 | 1.489568 | 0.136338 | 0.523563 |
| Nop56 | 1387.417 | -1.98567 | 1.333789 | -1.48875 | 0.136554 | 0.523753 |
| Pgant9 | 5272.631 | 0.898687 | 0.603898 | 1.488142 | 0.136713 | 0.523753 |
| Egm | 1172.568 | -1.80612 | 1.213586 | -1.48825 | 0.136685 | 0.523753 |
| CG12301 | 510.0574 | -2.78599 | 1.871482 | -1.48865 | 0.136579 | 0.523753 |
| CG32099 | 229.3789 | -4.03824 | 2.713406 | -1.48826 | 0.136683 | 0.523753 |
| CG10126 | 2401.301 | 1.875192 | 1.260604 | 1.487535 | 0.136874 | 0.52405 |
| E2f1 | 2507.578 | -1.12183 | 0.754619 | -1.48661 | 0.137118 | 0.52405 |
| CG15514 | 1267.743 | -1.50493 | 1.011995 | -1.48709 | 0.136991 | 0.52405 |
| CG18004 | 778.8288 | 2.117208 | 1.424017 | 1.486785 | 0.137072 | 0.52405 |

| | | | | | | |
|---|---|---|---|---|---|---|
| CG34384 | 8904.526 | 0.698692 | 0.469924 | 1.486817 | 0.137063 | 0.52405 |
| RnrL | 90.46017 | -5.7317 | 3.861331 | -1.48438 | 0.137707 | 0.524306 |
| beta4GalT | 174.9152 | 3.885878 | 2.61692 | 1.484905 | 0.137569 | 0.524306 |
| Rbm13 | 433.7091 | 3.390923 | 2.283948 | 1.484677 | 0.13763 | 0.524306 |
| CG8858 | 693.9719 | -2.1452 | 1.444901 | -1.48467 | 0.137632 | 0.524306 |
| CG33639 | 847.7238 | -2.14998 | 1.448229 | -1.48456 | 0.137661 | 0.524306 |
| AGO3 | 291.7059 | -2.62473 | 1.766997 | -1.48542 | 0.137433 | 0.524306 |
| Vlet | 1240.775 | 2.091519 | 1.407759 | 1.485708 | 0.137356 | 0.524306 |
| msl-3 | 1484.693 | 1.519671 | 1.023706 | 1.484479 | 0.137682 | 0.524306 |
| CG30197 | 14331.58 | -0.76572 | 0.516017 | -1.4839 | 0.137836 | 0.524539 |
| CG42671 | 1943.013 | 1.238463 | 0.834735 | 1.48366 | 0.137899 | 0.524539 |
| CG5815 | 479.3659 | -2.58774 | 1.744784 | -1.48313 | 0.13804 | 0.524827 |
| CG2617 | 249.3189 | 3.62788 | 2.448565 | 1.481635 | 0.138437 | 0.526024 |
| CG12945 | 638.6133 | -2.35368 | 1.589026 | -1.48121 | 0.138552 | 0.526024 |
| Bem46 | 263.3584 | 3.455208 | 2.332691 | 1.481211 | 0.13855 | 0.526024 |
| MED17 | 411.1651 | -2.4782 | 1.675357 | -1.4792 | 0.139086 | 0.527803 |
| lncRNA:CR | 319.4573 | -2.68394 | 1.815029 | -1.47873 | 0.139212 | 0.528032 |
| rl | 8425.342 | 0.767698 | 0.519372 | 1.478127 | 0.139374 | 0.528396 |
| eIF3d1 | 2467.067 | -1.52207 | 1.030362 | -1.47722 | 0.139617 | 0.529069 |
| CG5262 | 1612.111 | 1.269273 | 0.860136 | 1.475665 | 0.140034 | 0.529718 |
| CG10621 | 147.4287 | -4.06991 | 2.757304 | -1.47605 | 0.139931 | 0.529718 |
| CG30089 | 1046.761 | -2.01552 | 1.365905 | -1.4756 | 0.140052 | 0.529718 |
| l(1)G0004 | 375.6924 | 2.236484 | 1.515035 | 1.476192 | 0.139892 | 0.529718 |
| CG32815 | 2022.526 | 1.857513 | 1.259552 | 1.474741 | 0.140282 | 0.529833 |
| Corp | 129.5101 | 3.749048 | 2.542589 | 1.4745 | 0.140347 | 0.529833 |
| CG30082 | 89.63715 | 5.653949 | 3.83289 | 1.475114 | 0.140182 | 0.529833 |
| CG42340 | 2688.921 | 1.088605 | 0.738081 | 1.474912 | 0.140236 | 0.529833 |
| CG32809 | 9670.938 | 0.856475 | 0.581073 | 1.473955 | 0.140494 | 0.530138 |
| Idgf6 | 264.168 | 3.647011 | 2.475235 | 1.4734 | 0.140643 | 0.530453 |
| cnn | 2298.075 | -1.04858 | 0.711883 | -1.47297 | 0.140758 | 0.530636 |
| CG13366 | 2365.138 | 1.341181 | 0.910882 | 1.472399 | 0.140913 | 0.530639 |
| eIF3g1 | 3724.851 | 1.324595 | 0.899716 | 1.472236 | 0.140957 | 0.530639 |
| dpr11 | 3107.433 | 1.120096 | 0.760616 | 1.472617 | 0.140854 | 0.530639 |
| CG14757 | 7835.487 | 0.753993 | 0.512358 | 1.471615 | 0.141125 | 0.531021 |
| lid | 3388.855 | -1.09555 | 0.745564 | -1.46943 | 0.141717 | 0.531754 |
| CG11257 | 95.99311 | 5.537329 | 3.768123 | 1.469519 | 0.141692 | 0.531754 |
| l(2)k10201 | 136.4441 | -4.23911 | 2.884029 | -1.46986 | 0.141601 | 0.531754 |
| CG9527 | 295.3469 | -3.10241 | 2.110163 | -1.47022 | 0.141502 | 0.531754 |
| DIP-beta | 997.0238 | 1.966425 | 1.337773 | 1.469924 | 0.141582 | 0.531754 |
| mus302 | 680.6277 | 3.202903 | 2.178081 | 1.470516 | 0.141422 | 0.531754 |
| Tsp3A | 457.5477 | -2.61718 | 1.781857 | -1.46879 | 0.141889 | 0.531809 |
| eco | 313.4622 | 3.58151 | 2.439339 | 1.46823 | 0.142042 | 0.531809 |
| TfAP-2 | 739.5227 | -2.61114 | 1.778357 | -1.46829 | 0.142027 | 0.531809 |
| dind | 602.2724 | 2.419175 | 1.647083 | 1.468763 | 0.141897 | 0.531809 |
| CG7739 | 3948.728 | -0.88267 | 0.601215 | -1.46815 | 0.142063 | 0.531809 |
| CG11414 | 488.5286 | -1.99979 | 1.362475 | -1.46776 | 0.14217 | 0.53196 |
| gukh | 282.6897 | 2.771521 | 1.889414 | 1.466868 | 0.142412 | 0.53237 |
| Secp43 | 1895.907 | -1.977 | 1.347743 | -1.4669 | 0.142403 | 0.53237 |
| CG3726 | 1540.287 | 1.268114 | 0.865776 | 1.464713 | 0.142999 | 0.532437 |
| Wdr82 | 955.8013 | -2.10251 | 1.434909 | -1.46526 | 0.142851 | 0.532437 |

| | | | | | | |
|---|---|---|---|---|---|---|
| CG12811 | 487.9474 | 3.259112 | 2.224278 | 1.465245 | 0.142854 | 0.532437 |
| Sdc | 13131.01 | -0.66106 | 0.450876 | -1.46617 | 0.142602 | 0.532437 |
| Gpo1 | 3291.558 | 1.074898 | 0.733281 | 1.465875 | 0.142682 | 0.532437 |
| del | 622.6705 | 2.337066 | 1.594555 | 1.465653 | 0.142743 | 0.532437 |
| CG43110 | 849.228 | -2.16275 | 1.475019 | -1.46625 | 0.142579 | 0.532437 |
| Vps60 | 2588.738 | 1.088374 | 0.743029 | 1.464779 | 0.142981 | 0.532437 |
| SH3PX1 | 1189.57 | -2.24203 | 1.530798 | -1.46461 | 0.143027 | 0.532437 |
| CG8260 | 176.6705 | 4.368517 | 2.983827 | 1.464065 | 0.143176 | 0.532746 |
| CG5846 | 418.7754 | -3.15695 | 2.159803 | -1.46169 | 0.143827 | 0.534077 |
| CG9098 | 2649.879 | 1.821242 | 1.245406 | 1.462368 | 0.14364 | 0.534077 |
| CG9917 | 6728.721 | -0.83446 | 0.570785 | -1.46195 | 0.143756 | 0.534077 |
| CG32100 | 85.13246 | 5.562411 | 3.80585 | 1.461543 | 0.143867 | 0.534077 |
| Uev1A | 5614.809 | 0.917996 | 0.627908 | 1.461991 | 0.143744 | 0.534077 |
| CG9689 | 16514.27 | -0.7429 | 0.50843 | -1.46116 | 0.143971 | 0.534217 |
| CG13994 | 985.8466 | 2.102227 | 1.439369 | 1.46052 | 0.144147 | 0.534623 |
| ndl | 333.4508 | 3.910843 | 2.678615 | 1.460024 | 0.144283 | 0.534882 |
| Grip91 | 386.4367 | -2.43048 | 1.665483 | -1.45932 | 0.144476 | 0.535349 |
| AsnRS-m | 851.0826 | -2.5743 | 1.765184 | -1.45838 | 0.144737 | 0.536068 |
| Cul1 | 2065.364 | -1.40423 | 0.963092 | -1.45804 | 0.144829 | 0.536161 |
| SLC5A11 | 402.0603 | -2.55712 | 1.754596 | -1.45739 | 0.14501 | 0.536337 |
| crb | 1167.354 | 2.32683 | 1.596366 | 1.45758 | 0.144956 | 0.536337 |
| CG8366 | 469.3196 | 2.399277 | 1.647831 | 1.456021 | 0.145387 | 0.537482 |
| Cyp4p2 | 172.724 | -2.94534 | 2.023283 | -1.45572 | 0.14547 | 0.537542 |
| CG5726 | 516.3498 | 2.857866 | 1.964595 | 1.454684 | 0.145757 | 0.538354 |
| CG17776 | 1747.101 | 1.144914 | 0.787298 | 1.454231 | 0.145882 | 0.538377 |
| CG4393 | 1756.862 | -1.20564 | 0.829088 | -1.45418 | 0.145897 | 0.538377 |
| Spn27A | 476.9826 | 2.430316 | 1.672549 | 1.453061 | 0.146207 | 0.539024 |
| CG14207 | 4177.18 | -0.84892 | 0.584133 | -1.4533 | 0.146141 | 0.539024 |
| larp | 5377.063 | 0.790734 | 0.544494 | 1.452235 | 0.146436 | 0.539622 |
| Eip75B | 19683.61 | 0.651031 | 0.448525 | 1.451492 | 0.146643 | 0.540136 |
| CG2147 | 759.3366 | -2.27565 | 1.568526 | -1.45082 | 0.14683 | 0.540576 |
| mRpL9 | 2271.332 | 1.148933 | 0.792516 | 1.449728 | 0.147134 | 0.540706 |
| Fancm | 632.6024 | 2.847778 | 1.964681 | 1.449486 | 0.147202 | 0.540706 |
| CG33969 | 1401.714 | 1.418472 | 0.978466 | 1.44969 | 0.147145 | 0.540706 |
| lncRNA:CR | 1181.27 | 1.330603 | 0.917727 | 1.449891 | 0.147089 | 0.540706 |
| Klp10A | 3786.374 | -1.10907 | 0.764651 | -1.45043 | 0.146939 | 0.540706 |
| Clamp | 2046.479 | 1.436708 | 0.991718 | 1.448706 | 0.14742 | 0.541258 |
| CG4119 | 996.4364 | 1.403252 | 0.968973 | 1.448185 | 0.147565 | 0.54132 |
| CG9581 | 681.8518 | 2.656938 | 1.835001 | 1.447922 | 0.147639 | 0.54132 |
| CG3281 | 626.9015 | -2.35927 | 1.629366 | -1.44797 | 0.147625 | 0.54132 |
| CG2680 | 2538.968 | -1.10061 | 0.760629 | -1.44697 | 0.147905 | 0.542049 |
| CG10681 | 85.30768 | 4.940443 | 3.416536 | 1.446039 | 0.148166 | 0.542758 |
| CG34409 | 561.4841 | -2.71346 | 1.877209 | -1.44548 | 0.148324 | 0.543089 |
| CG6928 | 341.1599 | -3.59336 | 2.487063 | -1.44482 | 0.148509 | 0.543516 |
| opm | 415.4123 | -2.54614 | 1.763724 | -1.44362 | 0.148846 | 0.544503 |
| net | 1216.872 | -2.13825 | 1.482851 | -1.44198 | 0.149307 | 0.54585 |
| CG4186 | 350.6928 | -2.55927 | 1.775016 | -1.44183 | 0.14935 | 0.54585 |
| abo | 590.9693 | 2.764073 | 1.918943 | 1.440414 | 0.14975 | 0.546378 |
| CG13601 | 1066.387 | -1.57477 | 1.093502 | -1.44011 | 0.149835 | 0.546378 |
| Noa36 | 1222.371 | 2.035302 | 1.412958 | 1.440455 | 0.149739 | 0.546378 |

| | | | | | | |
|---|---|---|---|---|---|---|
| CG34303 | 267.6805 | 3.268337 | 2.269177 | 1.440318 | 0.149778 | 0.546378 |
| Rgk2 | 1128.248 | -1.39155 | 0.965669 | -1.44103 | 0.149577 | 0.546378 |
| gcl | 2004.595 | -1.18827 | 0.825722 | -1.43907 | 0.150131 | 0.546748 |
| CG7839 | 638.3545 | -2.0667 | 1.436034 | -1.43917 | 0.150102 | 0.546748 |
| GABA-B-R3 | 2243.987 | 1.581493 | 1.098996 | 1.439033 | 0.150141 | 0.546748 |
| CG42258 | 2990.836 | -0.97541 | 0.677953 | -1.43876 | 0.15022 | 0.546786 |
| Dhpr | 1189.968 | 1.596967 | 1.110198 | 1.438453 | 0.150306 | 0.546851 |
| Pdss2 | 383.7481 | -2.58997 | 1.801735 | -1.43748 | 0.15058 | 0.546859 |
| CG4839 | 66.4165 | 4.86397 | 3.382187 | 1.438114 | 0.150402 | 0.546859 |
| CG14715 | 1543.92 | -1.17269 | 0.815693 | -1.43766 | 0.15053 | 0.546859 |
| CG17508 | 13780.26 | -0.97429 | 0.677599 | -1.43786 | 0.150474 | 0.546859 |
| CG4074 | 908.7542 | -2.09687 | 1.45984 | -1.43637 | 0.150897 | 0.547019 |
| Ttc19 | 950.0355 | 1.981439 | 1.379269 | 1.436587 | 0.150835 | 0.547019 |
| Ire1 | 755.3975 | -1.63423 | 1.137326 | -1.43691 | 0.150744 | 0.547019 |
| Scamp | 2816.174 | 1.230601 | 0.856605 | 1.436602 | 0.150831 | 0.547019 |
| CG9067 | 720.9948 | 1.579846 | 1.100308 | 1.435821 | 0.151053 | 0.547338 |
| side-VIII | 10155.44 | -0.66366 | 0.462733 | -1.43421 | 0.151512 | 0.548752 |
| CG9662 | 1731.9 | 1.264676 | 0.88231 | 1.43337 | 0.151752 | 0.549374 |
| CG8311 | 699.5235 | 1.80666 | 1.260921 | 1.43281 | 0.151912 | 0.549706 |
| row | 1426.812 | -1.29975 | 0.907403 | -1.43239 | 0.152033 | 0.549896 |
| stan | 587.1955 | -2.10143 | 1.4683 | -1.4312 | 0.152374 | 0.550879 |
| sced | 636.6541 | -2.38636 | 1.668035 | -1.43064 | 0.152533 | 0.551095 |
| CG43759 | 1121.059 | -1.44772 | 1.01203 | -1.43051 | 0.152571 | 0.551095 |
| CG9135 | 939.0272 | -2.15387 | 1.506222 | -1.42998 | 0.152722 | 0.551392 |
| PIG-G | 1310.516 | -1.91269 | 1.338446 | -1.42904 | 0.152993 | 0.552125 |
| neur | 143.3288 | -4.64057 | 3.249462 | -1.4281 | 0.153262 | 0.552596 |
| lncRNA:CR | 75.72149 | 5.476654 | 3.834851 | 1.428127 | 0.153255 | 0.552596 |
| CG12825 | 135.8672 | 4.753607 | 3.32993 | 1.42754 | 0.153424 | 0.55291 |
| CG13484 | 295.0866 | -2.78844 | 1.953615 | -1.42732 | 0.153487 | 0.55291 |
| emb | 4030.954 | 0.866603 | 0.607376 | 1.426798 | 0.153638 | 0.55296 |
| CG11858 | 720.8578 | 2.360572 | 1.654183 | 1.427032 | 0.153571 | 0.55296 |
| Galphao | 14743.61 | 0.691287 | 0.485204 | 1.424734 | 0.154234 | 0.554856 |
| shams | 211.9121 | -3.89967 | 2.738411 | -1.42406 | 0.154428 | 0.554914 |
| casp | 2290.684 | 1.265714 | 0.889092 | 1.423602 | 0.154562 | 0.554914 |
| CG31224 | 5098.122 | -0.88461 | 0.621647 | -1.42301 | 0.154734 | 0.554914 |
| chinmo | 10652.93 | 0.670513 | 0.470912 | 1.423861 | 0.154487 | 0.554914 |
| CG8924 | 1306.821 | -2.01994 | 1.419474 | -1.42302 | 0.15473 | 0.554914 |
| mRpL2 | 1256.706 | -1.69694 | 1.192172 | -1.4234 | 0.15462 | 0.554914 |
| MED24 | 1173.822 | -2.30696 | 1.620456 | -1.42365 | 0.154548 | 0.554914 |
| ssp3 | 526.6432 | -1.85002 | 1.300377 | -1.42268 | 0.154829 | 0.555007 |
| fl(2)d | 5139.727 | 0.824726 | 0.580071 | 1.421768 | 0.155094 | 0.555706 |
| PIG-U | 442.2458 | 3.078142 | 2.16579 | 1.421256 | 0.155242 | 0.55599 |
| Dyb | 5542.19 | 0.807558 | 0.568401 | 1.420754 | 0.155388 | 0.556017 |
| RabX6 | 349.4185 | -2.51083 | 1.767242 | -1.42076 | 0.155386 | 0.556017 |
| poe | 8682.685 | -0.91595 | 0.645357 | -1.4193 | 0.155813 | 0.556498 |
| CG31706 | 20767.01 | -0.64998 | 0.458122 | -1.41879 | 0.15596 | 0.556498 |
| Rfx | 545.0984 | -2.78006 | 1.959311 | -1.4189 | 0.155929 | 0.556498 |
| CG13840 | 1637.847 | 2.231317 | 1.57317 | 1.418357 | 0.156087 | 0.556498 |
| Hrs | 567.8364 | 2.476526 | 1.744571 | 1.419562 | 0.155735 | 0.556498 |
| sau | 1907.855 | -1.38204 | 0.974538 | -1.41815 | 0.156147 | 0.556498 |

| | | | | | | |
|---|---|---|---|---|---|---|
| AhcyL1 | 2669.543 | 0.956273 | 0.674128 | 1.418532 | 0.156035 | 0.556498 |
| CG1640 | 9147.362 | 0.856542 | 0.60347 | 1.419363 | 0.155793 | 0.556498 |
| CG15435 | 886.6881 | 2.90564 | 2.047908 | 1.418833 | 0.155948 | 0.556498 |
| CG33090 | 4320.894 | -0.97237 | 0.686446 | -1.41653 | 0.15662 | 0.557097 |
| EndoU | 455.5697 | 3.535632 | 2.495636 | 1.416726 | 0.156563 | 0.557097 |
| dco | 2173.095 | -1.06734 | 0.753374 | -1.41675 | 0.156556 | 0.557097 |
| CG1910 | 1824.538 | -1.41271 | 0.997327 | -1.4165 | 0.15663 | 0.557097 |
| cora | 7416.278 | -0.89297 | 0.630555 | -1.41617 | 0.156727 | 0.557097 |
| CG12863 | 279.1235 | -3.25809 | 2.300667 | -1.41615 | 0.156731 | 0.557097 |
| mura | 11174.01 | -0.77504 | 0.54757 | -1.41542 | 0.156946 | 0.557119 |
| HP1b | 1178.021 | -1.67862 | 1.185853 | -1.41554 | 0.156911 | 0.557119 |
| Fdx2 | 1702.088 | -1.18309 | 0.835801 | -1.41552 | 0.156916 | 0.557119 |
| CG7706 | 230.5349 | 3.446454 | 2.435828 | 1.4149 | 0.157098 | 0.557132 |
| CR42875 | 665.5368 | -1.67793 | 1.185774 | -1.41505 | 0.157054 | 0.557132 |
| wupA | 499.3521 | 2.167189 | 1.531926 | 1.414682 | 0.157162 | 0.557132 |
| ND-PDSW | 10338.96 | 0.699514 | 0.494545 | 1.41446 | 0.157227 | 0.557132 |
| mRpS9 | 578.0113 | 2.266382 | 1.603825 | 1.41311 | 0.157623 | 0.558044 |
| CG9919 | 3635.577 | 1.45088 | 1.026602 | 1.413283 | 0.157572 | 0.558044 |
| CHOp24 | 3808.348 | 0.897097 | 0.63534 | 1.411994 | 0.157952 | 0.55896 |
| fs(2)ltoPP4 | 1274.371 | -1.83335 | 1.29906 | -1.41129 | 0.15816 | 0.558974 |
| Sp7 | 2108.345 | -1.26463 | 0.895859 | -1.41164 | 0.158056 | 0.558974 |
| CG31007 | 192.5395 | 4.021485 | 2.849548 | 1.411271 | 0.158165 | 0.558974 |
| CG1737 | 515.1454 | -1.81392 | 1.286357 | -1.41012 | 0.158504 | 0.559926 |
| spin | 1476.875 | -1.90905 | 1.355072 | -1.40882 | 0.158888 | 0.561036 |
| Fmr1 | 1734.328 | 1.285964 | 0.914426 | 1.406308 | 0.159633 | 0.56126 |
| CG6912 | 4773.009 | 0.998583 | 0.709117 | 1.408206 | 0.15907 | 0.56126 |
| olf186-M | 204.5306 | -3.21435 | 2.283744 | -1.40749 | 0.159281 | 0.56126 |
| cyp33 | 368.0329 | -3.03065 | 2.154289 | -1.4068 | 0.159487 | 0.56126 |
| Pura | 122.1991 | 3.487473 | 2.478947 | 1.406836 | 0.159476 | 0.56126 |
| crc | 2491.622 | -1.01867 | 0.724338 | -1.40634 | 0.159622 | 0.56126 |
| CG42402 | 20993.25 | 0.581175 | 0.413294 | 1.406201 | 0.159665 | 0.56126 |
| Patronin | 4388.306 | 0.948021 | 0.673405 | 1.407802 | 0.15919 | 0.56126 |
| CG16952 | 3024.201 | 1.570241 | 1.11686 | 1.405943 | 0.159741 | 0.56126 |
| CG32537 | 189.2658 | 3.224629 | 2.293842 | 1.405777 | 0.159791 | 0.56126 |
| CG7990 | 4761.94 | -0.97 | 0.688879 | -1.40808 | 0.159106 | 0.56126 |
| LanA | 65.37706 | 5.003884 | 3.555512 | 1.40736 | 0.159321 | 0.56126 |
| CG4467 | 10884.24 | -0.6412 | 0.456271 | -1.40531 | 0.159928 | 0.561496 |
| CG7685 | 476.9374 | -2.26559 | 1.613458 | -1.40418 | 0.160264 | 0.562432 |
| CG11975 | 2229.322 | 1.82328 | 1.299585 | 1.402971 | 0.160626 | 0.563453 |
| Bdbt | 164.9141 | 3.8195 | 2.723161 | 1.402598 | 0.160737 | 0.563597 |
| Rpb3 | 578.2141 | -2.46879 | 1.760861 | -1.40204 | 0.160905 | 0.563739 |
| asRNA:CR4 | 137.7469 | 4.049491 | 2.888384 | 1.401992 | 0.160918 | 0.563739 |
| Mlp84B | 629.2232 | 2.410036 | 1.720531 | 1.400751 | 0.161289 | 0.564299 |
| CG3907 | 6207.94 | 0.735154 | 0.524816 | 1.400784 | 0.161279 | 0.564299 |
| Fcp1 | 510.4682 | 2.37494 | 1.695061 | 1.401094 | 0.161186 | 0.564299 |
| fog | 3384.284 | -0.95797 | 0.684324 | -1.39988 | 0.161548 | 0.56496 |
| Oga | 254.6163 | -3.6268 | 2.591705 | -1.39939 | 0.161697 | 0.564974 |
| CG17111 | 697.7263 | -2.78166 | 1.988085 | -1.39917 | 0.161763 | 0.564974 |
| CG31098 | 703.8831 | -2.44987 | 1.750782 | -1.3993 | 0.161722 | 0.564974 |
| CaBP1 | 2856.988 | 1.051452 | 0.751863 | 1.398463 | 0.161974 | 0.565465 |

| | | | | | | |
|---|---|---|---|---|---|---|
| mtsh | 144.4536 | -4.67689 | 3.345237 | -1.39807 | 0.162091 | 0.565626 |
| CG16743 | 20626.16 | -0.7323 | 0.524382 | -1.3965 | 0.162563 | 0.567029 |
| Hmt4-20 | 2260.217 | 1.025061 | 0.734512 | 1.395566 | 0.162845 | 0.567676 |
| CG33144 | 4042.335 | 0.809956 | 0.580538 | 1.395181 | 0.162961 | 0.567676 |
| Nep6 | 680.8735 | 2.440836 | 1.749308 | 1.395315 | 0.162921 | 0.567676 |
| CG9992 | 1479.961 | -1.841 | 1.31977 | -1.39494 | 0.163035 | 0.567686 |
| CG6695 | 1416.349 | -1.17742 | 0.844386 | -1.39441 | 0.163195 | 0.567849 |
| Rab19 | 2380.559 | 1.049778 | 0.7529 | 1.394313 | 0.163223 | 0.567849 |
| ph-p | 2078.952 | 1.453625 | 1.043371 | 1.3932 | 0.163559 | 0.568035 |
| CG5062 | 225.5226 | -3.35157 | 2.405403 | -1.39335 | 0.163513 | 0.568035 |
| MED6 | 492.2057 | 2.794352 | 2.005311 | 1.393476 | 0.163476 | 0.568035 |
| Lsm10 | 93.98602 | -5.20251 | 3.733256 | -1.39356 | 0.163451 | 0.568035 |
| Mpp6 | 134.5965 | -3.69067 | 2.650413 | -1.39249 | 0.163774 | 0.568196 |
| Flo1 | 1776.894 | -1.42958 | 1.026657 | -1.39246 | 0.163784 | 0.568196 |
| oxt | 373.7279 | 2.761396 | 1.983272 | 1.392344 | 0.163818 | 0.568196 |
| CG15236 | 2334.04 | 1.449627 | 1.042304 | 1.39079 | 0.164289 | 0.569337 |
| CG8004 | 1332.548 | -1.958 | 1.407604 | -1.39102 | 0.16422 | 0.569337 |
| CG30349 | 896.1089 | -2.41965 | 1.740235 | -1.39041 | 0.164403 | 0.569487 |
| numb | 1447.398 | 1.189196 | 0.856054 | 1.38916 | 0.164784 | 0.570337 |
| Indy | 930.7233 | -1.59785 | 1.150245 | -1.38914 | 0.164791 | 0.570337 |
| Kat60 | 471.3723 | 2.063298 | 1.486599 | 1.387931 | 0.165158 | 0.570606 |
| danr | 385.939 | -2.19373 | 1.580419 | -1.38807 | 0.165117 | 0.570606 |
| CG6051 | 1030.294 | 1.686923 | 1.21568 | 1.387637 | 0.165248 | 0.570606 |
| lncRNA:CR | 429.5792 | 2.812981 | 2.0262 | 1.388303 | 0.165045 | 0.570606 |
| lncRNA:CR | 527.9051 | -1.8386 | 1.325356 | -1.38725 | 0.165366 | 0.570606 |
| CG32022 | 198.6391 | 3.230019 | 2.328263 | 1.387309 | 0.165348 | 0.570606 |
| Snmp2 | 134.0143 | -3.23287 | 2.328774 | -1.38823 | 0.165067 | 0.570606 |
| CG4341 | 1318.881 | 1.392904 | 1.004314 | 1.38692 | 0.165466 | 0.570705 |
| alphaTub8 | 7799.983 | 0.758284 | 0.546862 | 1.386608 | 0.165561 | 0.570788 |
| Cmpk | 8147.516 | 0.693947 | 0.500694 | 1.385972 | 0.165756 | 0.571213 |
| Dif | 1714.218 | -1.12648 | 0.813135 | -1.38536 | 0.165943 | 0.571613 |
| blp | 776.214 | -1.76901 | 1.279374 | -1.38271 | 0.166753 | 0.573348 |
| Smg6 | 1978.47 | 1.586231 | 1.146623 | 1.383393 | 0.166544 | 0.573348 |
| wun | 5166.309 | 0.791715 | 0.572649 | 1.382547 | 0.166804 | 0.573348 |
| sm | 40390.64 | 0.529766 | 0.383134 | 1.382716 | 0.166752 | 0.573348 |
| betaggt-II | 2424.212 | -0.96168 | 0.695464 | -1.38279 | 0.166731 | 0.573348 |
| lncRNA:CR | 150.9703 | -4.41804 | 3.196772 | -1.38203 | 0.166962 | 0.573645 |
| Cdc5 | 532.1473 | -2.79934 | 2.026766 | -1.38118 | 0.167223 | 0.574296 |
| Jheh3 | 1234.144 | -1.75489 | 1.270898 | -1.38083 | 0.167333 | 0.574427 |
| CG4908 | 944.5369 | -1.80033 | 1.305028 | -1.37953 | 0.167731 | 0.57555 |
| Nle | 332.7228 | 3.025846 | 2.194152 | 1.37905 | 0.167879 | 0.575811 |
| CG4291 | 323.7477 | -2.59931 | 1.888236 | -1.37658 | 0.168641 | 0.57608 |
| PolQ | 192.5967 | -4.04653 | 2.938725 | -1.37697 | 0.168522 | 0.57608 |
| CG4562 | 1133.809 | -1.59052 | 1.155103 | -1.37695 | 0.168529 | 0.57608 |
| nAChRalph | 6742.441 | 0.906668 | 0.657725 | 1.378491 | 0.168052 | 0.57608 |
| CG31111 | 211.6566 | 3.244367 | 2.356604 | 1.376713 | 0.168601 | 0.57608 |
| CG12822 | 3477.567 | 1.061411 | 0.770889 | 1.376867 | 0.168553 | 0.57608 |
| HIP | 139.4076 | -3.10271 | 2.254101 | -1.37647 | 0.168675 | 0.57608 |
| Syt12 | 2781.171 | 1.794938 | 1.302852 | 1.377699 | 0.168296 | 0.57608 |
| RabX1 | 1178.277 | 1.406497 | 1.021807 | 1.37648 | 0.168673 | 0.57608 |

| | | | | | | |
|---|---|---|---|---|---|---|
| Syx13 | 1290.677 | -1.67776 | 1.217292 | -1.37827 | 0.168119 | 0.57608 |
| Drep4 | 1416.477 | 1.772632 | 1.289259 | 1.374923 | 0.169155 | 0.577268 |
| CG2258 | 4010.645 | 0.827188 | 0.601743 | 1.374654 | 0.169239 | 0.577268 |
| CG42399 | 626.6748 | -1.96634 | 1.430243 | -1.37483 | 0.169183 | 0.577268 |
| tey | 4916.963 | 0.821306 | 0.597582 | 1.374383 | 0.169323 | 0.577309 |
| CG9727 | 3608.218 | -0.86064 | 0.626854 | -1.37295 | 0.169767 | 0.578088 |
| trsn | 1055.753 | 1.349373 | 0.982499 | 1.373409 | 0.169625 | 0.578088 |
| Fbxl4 | 1077.08 | -2.13466 | 1.554735 | -1.37301 | 0.16975 | 0.578088 |
| Vps26 | 538.6747 | -2.27747 | 1.662151 | -1.3702 | 0.170626 | 0.580765 |
| Trh | 347.3193 | -3.06634 | 2.238285 | -1.36995 | 0.170702 | 0.580777 |
| cac | 22035.36 | 0.620511 | 0.453051 | 1.369628 | 0.170803 | 0.580875 |
| CG1814 | 2844.125 | 0.984259 | 0.718783 | 1.36934 | 0.170893 | 0.580935 |
| Nxf3 | 813.3003 | 1.496197 | 1.093052 | 1.368825 | 0.171054 | 0.581236 |
| CG8414 | 79.93224 | -4.78623 | 3.497368 | -1.36852 | 0.171148 | 0.581309 |
| CG17734 | 8854.647 | 0.667658 | 0.488037 | 1.368048 | 0.171297 | 0.58157 |
| CG14210 | 188.1256 | 3.205454 | 2.345152 | 1.366843 | 0.171674 | 0.582604 |
| CG1847 | 1477.831 | 1.472543 | 1.07754 | 1.366579 | 0.171757 | 0.582639 |
| Mst36Fa | 163.1601 | 3.750496 | 2.745945 | 1.365831 | 0.171992 | 0.582697 |
| Graf | 4790.035 | 0.782808 | 0.572967 | 1.366237 | 0.171865 | 0.582697 |
| CDC50 | 6696.341 | 0.812796 | 0.595056 | 1.365915 | 0.171966 | 0.582697 |
| Aduk | 936.2169 | 1.754612 | 1.285759 | 1.364651 | 0.172363 | 0.582723 |
| GlyT | 1336.543 | 1.855734 | 1.359699 | 1.364813 | 0.172312 | 0.582723 |
| Neto | 331.2866 | -2.28864 | 1.676926 | -1.36478 | 0.172321 | 0.582723 |
| TSG101 | 3934.603 | -0.97221 | 0.7124 | -1.3647 | 0.172347 | 0.582723 |
| CG7504 | 476.1169 | -2.59219 | 1.89908 | -1.36497 | 0.172262 | 0.582723 |
| Mulk | 1018.904 | 1.469204 | 1.077608 | 1.363394 | 0.172758 | 0.583814 |
| CG7985 | 1154.554 | 1.497845 | 1.099636 | 1.362127 | 0.173158 | 0.584562 |
| CG30109 | 680.9177 | 1.88698 | 1.385003 | 1.362437 | 0.17306 | 0.584562 |
| Mvb12 | 905.4134 | 1.717507 | 1.261019 | 1.361999 | 0.173198 | 0.584562 |
| CG5910 | 410.4616 | -3.40019 | 2.497574 | -1.3614 | 0.173388 | 0.584709 |
| E(spl)mbet | 624.3668 | 2.038304 | 1.497165 | 1.361442 | 0.173374 | 0.584709 |
| dtn | 5382.229 | 0.76527 | 0.562428 | 1.360654 | 0.173623 | 0.585258 |
| Tor | 159.3771 | 3.486909 | 2.563978 | 1.359961 | 0.173842 | 0.585751 |
| Arp2 | 2545.703 | -1.32186 | 0.972354 | -1.35944 | 0.174007 | 0.586059 |
| hts | 8684.308 | -0.6596 | 0.485366 | -1.35898 | 0.174153 | 0.586305 |
| MED4 | 808.4909 | -2.17159 | 1.598338 | -1.35865 | 0.174257 | 0.586408 |
| CG6523 | 1707.101 | -1.06243 | 0.782572 | -1.35761 | 0.174587 | 0.586464 |
| Ugt35B1 | 56100.52 | -0.70774 | 0.521306 | -1.35763 | 0.174582 | 0.586464 |
| RpS15Ab | 1303.374 | -1.22323 | 0.900706 | -1.35808 | 0.174438 | 0.586464 |
| CG17765 | 5949.04 | 0.714734 | 0.526511 | 1.357491 | 0.174625 | 0.586464 |
| Sam-S | 12922.61 | 0.786496 | 0.579393 | 1.357449 | 0.174639 | 0.586464 |
| CG5953 | 824.3307 | 1.662448 | 1.225523 | 1.356521 | 0.174933 | 0.587209 |
| HINT1 | 5762.639 | -0.86699 | 0.639329 | -1.35609 | 0.175071 | 0.587425 |
| ear | 728.4163 | 2.05076 | 1.512523 | 1.355854 | 0.175146 | 0.587429 |
| Tgi | 1078.247 | 1.326431 | 0.978618 | 1.355412 | 0.175286 | 0.587655 |
| Ahcy | 335.3047 | -3.06991 | 2.26569 | -1.35496 | 0.175432 | 0.587897 |
| HDAC11 | 703.3393 | -2.14932 | 1.588435 | -1.3531 | 0.176022 | 0.589385 |
| CG17221 | 298.5644 | -2.40258 | 1.775312 | -1.35333 | 0.17595 | 0.589385 |
| Tao | 3207.998 | 0.887727 | 0.656252 | 1.352724 | 0.176144 | 0.589546 |
| CG30054 | 195.9242 | -3.8421 | 2.842682 | -1.35158 | 0.176511 | 0.590529 |

| | | | | | | |
|---|---|---|---|---|---|---|
| dome | 988.8321 | 1.753712 | 1.297864 | 1.351229 | 0.176622 | 0.590654 |
| CG17271 | 1086.277 | -1.65532 | 1.225967 | -1.35022 | 0.176946 | 0.591243 |
| CG7742 | 126.319 | -3.96479 | 2.936004 | -1.3504 | 0.176887 | 0.591243 |
| Ssk | 2686.303 | -0.97905 | 0.725234 | -1.34998 | 0.177023 | 0.591254 |
| Ekar | 396.7464 | -2.68243 | 1.988679 | -1.34885 | 0.177385 | 0.591996 |
| PPP1R15 | 3825.392 | -0.89588 | 0.664192 | -1.34883 | 0.177392 | 0.591996 |
| CG12237 | 3719.235 | 0.817661 | 0.606409 | 1.348366 | 0.177541 | 0.592245 |
| CG10055 | 1407.341 | -1.44494 | 1.072699 | -1.34701 | 0.177977 | 0.592629 |
| bai | 3431.836 | -0.83572 | 0.620496 | -1.34686 | 0.178025 | 0.592629 |
| jus | 4824.041 | -0.76353 | 0.566803 | -1.34709 | 0.177952 | 0.592629 |
| spri | 4384.425 | -0.87249 | 0.647383 | -1.34772 | 0.17775 | 0.592629 |
| Vps15 | 689.0818 | -2.17323 | 1.613081 | -1.34726 | 0.177898 | 0.592629 |
| CG15803 | 791.8939 | 1.63957 | 1.217853 | 1.346278 | 0.178213 | 0.593008 |
| Thd1 | 15964.79 | 0.647561 | 0.481281 | 1.345495 | 0.178465 | 0.593603 |
| CG13255 | 824.714 | -1.40358 | 1.0434 | -1.3452 | 0.17856 | 0.59367 |
| lncRNA:no | 330092.2 | 0.492586 | 0.366544 | 1.343866 | 0.178992 | 0.594633 |
| Taf4 | 3211.838 | -1.0288 | 0.765564 | -1.34385 | 0.178997 | 0.594633 |
| CG12975 | 2659.005 | 1.040721 | 0.774785 | 1.343238 | 0.179195 | 0.595043 |
| Hrd3 | 879.8204 | -1.40405 | 1.045483 | -1.34297 | 0.179283 | 0.595088 |
| CG6041 | 225.8876 | 3.363433 | 2.505113 | 1.342627 | 0.179393 | 0.595207 |
| CG5044 | 416.0274 | -2.39159 | 1.783277 | -1.34112 | 0.179881 | 0.595833 |
| Miro | 1612.178 | 1.549568 | 1.154937 | 1.341691 | 0.179696 | 0.595833 |
| Cul5 | 303.9142 | -3.19433 | 2.382237 | -1.34089 | 0.179955 | 0.595833 |
| CG17002 | 244.9541 | 3.344465 | 2.493805 | 1.341109 | 0.179885 | 0.595833 |
| otk | 185.7193 | 3.693324 | 2.754692 | 1.340739 | 0.180005 | 0.595833 |
| Naa35 | 277.4797 | -2.5511 | 1.903453 | -1.34025 | 0.180164 | 0.595833 |
| gbb | 1525.772 | 1.954152 | 1.458088 | 1.340215 | 0.180175 | 0.595833 |
| e(y)2 | 233.2162 | -3.01689 | 2.250902 | -1.3403 | 0.180148 | 0.595833 |
| nAChRalpł | 11719.16 | 0.614728 | 0.459341 | 1.338282 | 0.180804 | 0.597667 |
| emc | 11060.04 | -0.74562 | 0.557243 | -1.33805 | 0.180879 | 0.597669 |
| mts | 11609.73 | -0.60258 | 0.450834 | -1.33659 | 0.181355 | 0.598255 |
| Ptp61F | 2819.204 | -0.95708 | 0.715933 | -1.33683 | 0.181278 | 0.598255 |
| byn | 138.1799 | -4.0686 | 3.043739 | -1.33671 | 0.181316 | 0.598255 |
| papi | 816.9035 | 1.926013 | 1.440417 | 1.337122 | 0.181183 | 0.598255 |
| Lgr1 | 184.6163 | 3.386089 | 2.53514 | 1.335661 | 0.18166 | 0.598923 |
| Myo81F | 2732.217 | -0.89085 | 0.667041 | -1.33552 | 0.181707 | 0.598923 |
| DAAM | 2460.085 | -0.96267 | 0.721347 | -1.33455 | 0.182023 | 0.599474 |
| nclb | 662.4333 | 2.511167 | 1.881396 | 1.334736 | 0.181963 | 0.599474 |
| CG42663 | 201.2772 | 2.8932 | 2.168522 | 1.334181 | 0.182145 | 0.599628 |
| Nrx-1 | 16165.16 | 0.676062 | 0.506913 | 1.333686 | 0.182307 | 0.599916 |
| CycC | 131.1919 | -3.12235 | 2.343835 | -1.33216 | 0.182809 | 0.601322 |
| rempA | 190.8584 | 3.119382 | 2.3421 | 1.331874 | 0.182902 | 0.601349 |
| mthl9 | 265.7409 | 2.574447 | 1.93324 | 1.331675 | 0.182967 | 0.601349 |
| esn | 690.8467 | -2.1384 | 1.606135 | -1.33139 | 0.183059 | 0.601406 |
| AMPKalph | 1860.454 | -1.2477 | 0.938759 | -1.3291 | 0.183815 | 0.601414 |
| bi | 1418.794 | -1.31219 | 0.98626 | -1.33047 | 0.183365 | 0.601414 |
| SK | 8632.417 | -0.80876 | 0.60862 | -1.32884 | 0.183902 | 0.601414 |
| CG9175 | 829.05 | -1.92216 | 1.446155 | -1.32916 | 0.183797 | 0.601414 |
| CG6180 | 6922.196 | -0.69146 | 0.520597 | -1.3282 | 0.184111 | 0.601414 |
| CG10947 | 1187.334 | -1.50166 | 1.128167 | -1.33106 | 0.183169 | 0.601414 |

| | | | | | | |
|---|---|---|---|---|---|---|
| mAChR-B | 763.5099 | 1.442113 | 1.085665 | 1.328323 | 0.184072 | 0.601414 |
| Osbp | 669.4488 | -1.52506 | 1.147762 | -1.32873 | 0.183938 | 0.601414 |
| eIF3k | 3338.833 | 0.818951 | 0.616382 | 1.328642 | 0.183966 | 0.601414 |
| MESK2 | 12982.12 | 0.627297 | 0.472067 | 1.328831 | 0.183904 | 0.601414 |
| CG14130 | 898.7807 | -2.17307 | 1.635776 | -1.32847 | 0.184024 | 0.601414 |
| CG12038 | 3333.044 | -0.82938 | 0.623226 | -1.33079 | 0.183259 | 0.601414 |
| CG11811 | 1004.605 | 2.005983 | 1.509553 | 1.328859 | 0.183895 | 0.601414 |
| Naprt | 1589.88 | 1.515692 | 1.140172 | 1.329354 | 0.183731 | 0.601414 |
| TBC1D16 | 2141.427 | -1.01471 | 0.764307 | -1.32762 | 0.184304 | 0.601798 |
| lncRNA:let | 505.3404 | -1.73865 | 1.309952 | -1.32726 | 0.184421 | 0.601938 |
| wtrw | 16038.2 | -0.78659 | 0.592758 | -1.32699 | 0.184512 | 0.601988 |
| Tsen34 | 84.58349 | -4.69084 | 3.538978 | -1.32548 | 0.185012 | 0.603376 |
| noc | 1433.717 | -1.19541 | 0.902366 | -1.32475 | 0.185252 | 0.603669 |
| betaTub85 | 106.4195 | -4.67798 | 3.530802 | -1.32491 | 0.185202 | 0.603669 |
| alph | 2676.577 | -0.97232 | 0.734312 | -1.32413 | 0.185461 | 0.604103 |
| PolE4 | 580.2654 | 1.630672 | 1.23222 | 1.323361 | 0.185715 | 0.604653 |
| CG1582 | 1134.817 | -2.35749 | 1.781703 | -1.32316 | 0.185781 | 0.604653 |
| Yippee | 3868.15 | 0.954064 | 0.721173 | 1.322933 | 0.185858 | 0.604659 |
| CG31091 | 94.01192 | -4.61975 | 3.492807 | -1.32265 | 0.185953 | 0.604722 |
| MED22 | 738.2565 | -2.16129 | 1.635021 | -1.32187 | 0.186211 | 0.604827 |
| beta-PheR | 334.5935 | 2.624545 | 1.985347 | 1.321958 | 0.186182 | 0.604827 |
| Fkbp14 | 9792.822 | 0.619121 | 0.46835 | 1.321919 | 0.186195 | 0.604827 |
| Topors | 549.181 | 2.318456 | 1.755036 | 1.32103 | 0.186491 | 0.605494 |
| CG5543 | 871.5534 | -1.64377 | 1.244924 | -1.32038 | 0.186709 | 0.605954 |
| CG2918 | 2452.927 | -1.01681 | 0.770878 | -1.31903 | 0.187158 | 0.606329 |
| CG18661 | 906.8664 | 1.611657 | 1.221591 | 1.31931 | 0.187066 | 0.606329 |
| YL-1 | 365.2716 | 1.969129 | 1.495796 | 1.316443 | 0.188026 | 0.606329 |
| CG11753 | 1860.177 | -1.12705 | 0.856149 | -1.31642 | 0.188033 | 0.606329 |
| Srp72 | 616.6661 | -1.95064 | 1.481666 | -1.31652 | 0.188 | 0.606329 |
| Cog6 | 455.9293 | 2.875003 | 2.183712 | 1.316567 | 0.187984 | 0.606329 |
| blow | 468.5615 | -1.97658 | 1.500374 | -1.31739 | 0.187707 | 0.606329 |
| CG14985 | 575.1443 | -1.91595 | 1.453511 | -1.31815 | 0.187453 | 0.606329 |
| Dek | 3698.758 | -0.81427 | 0.617272 | -1.31915 | 0.187119 | 0.606329 |
| mmy | 2437.445 | 1.681209 | 1.274158 | 1.319467 | 0.187013 | 0.606329 |
| dnr1 | 1025.017 | -1.34297 | 1.01994 | -1.31671 | 0.187935 | 0.606329 |
| eIF2gamm | 1146.708 | -1.71424 | 1.30214 | -1.31648 | 0.188012 | 0.606329 |
| CTPsyn | 979.9537 | -1.55525 | 1.180691 | -1.31724 | 0.187759 | 0.606329 |
| HP5 | 1399.435 | 1.1127 | 0.843678 | 1.318867 | 0.187213 | 0.606329 |
| CG1998 | 7628.41 | -0.75495 | 0.572611 | -1.31843 | 0.187358 | 0.606329 |
| CG12316 | 766.8916 | -2.01693 | 1.529626 | -1.31858 | 0.18731 | 0.606329 |
| CG12926 | 2079.803 | 0.960981 | 0.730795 | 1.31498 | 0.188516 | 0.607644 |
| Rab21 | 3062.92 | -1.09997 | 0.836723 | -1.31461 | 0.188639 | 0.607682 |
| Dll | 1379.283 | -1.13219 | 0.86131 | -1.31449 | 0.18868 | 0.607682 |
| kuk | 3571.993 | -1.05799 | 0.805163 | -1.31401 | 0.188842 | 0.607686 |
| dgt1 | 1022.486 | 1.731558 | 1.317551 | 1.314225 | 0.188771 | 0.607686 |
| sol | 933.2486 | -1.75776 | 1.338133 | -1.31359 | 0.188984 | 0.607686 |
| PNUTS | 7655.751 | -0.67581 | 0.514419 | -1.31374 | 0.188933 | 0.607686 |
| CG34259 | 251.228 | 3.633364 | 2.766788 | 1.313207 | 0.189113 | 0.607859 |
| abs | 347.5255 | 2.857241 | 2.178624 | 1.311489 | 0.189693 | 0.607968 |
| CG8036 | 1505.388 | -1.24247 | 0.947169 | -1.31177 | 0.189597 | 0.607968 |

| | | | | | | |
|---|---|---|---|---|---|---|
| CLS | 1509.487 | -1.10155 | 0.839557 | -1.31206 | 0.1895 | 0.607968 |
| AGBE | 1702.965 | -1.26879 | 0.967408 | -1.31154 | 0.189676 | 0.607968 |
| CG33543 | 4890.402 | 0.850691 | 0.648747 | 1.311284 | 0.189762 | 0.607968 |
| Hsc20 | 989.5524 | -1.40327 | 1.069193 | -1.31246 | 0.189366 | 0.607968 |
| lncRNA:CR | 332.7482 | 2.159982 | 1.646847 | 1.311586 | 0.18966 | 0.607968 |
| CG15019 | 1412.76 | -1.17275 | 0.894489 | -1.31109 | 0.189829 | 0.607968 |
| U2af50 | 1620.861 | 1.888339 | 1.439521 | 1.311783 | 0.189593 | 0.607968 |
| Lac | 282.9318 | 3.160036 | 2.410825 | 1.310769 | 0.189936 | 0.608067 |
| CG31678 | 1311.813 | 1.20991 | 0.924201 | 1.309143 | 0.190486 | 0.608404 |
| CG8199 | 192.0142 | 2.969926 | 2.269384 | 1.308693 | 0.190639 | 0.608404 |
| CG17193 | 240.2711 | 2.549199 | 1.94735 | 1.309061 | 0.190514 | 0.608404 |
| BOD1 | 2278.814 | -0.94751 | 0.723787 | -1.3091 | 0.190499 | 0.608404 |
| betaTub97 | 600.3488 | -1.59387 | 1.217935 | -1.30867 | 0.190648 | 0.608404 |
| Arc1 | 2321.793 | -0.95506 | 0.729074 | -1.30996 | 0.190208 | 0.608404 |
| Eip74EF | 1426.445 | -1.13858 | 0.869554 | -1.30939 | 0.190403 | 0.608404 |
| CG17249 | 662.994 | -2.03592 | 1.555525 | -1.30883 | 0.190592 | 0.608404 |
| RpL5 | 32754.16 | 0.516922 | 0.395467 | 1.307118 | 0.191173 | 0.609838 |
| CG14767 | 18448.42 | 0.541138 | 0.414138 | 1.30666 | 0.191328 | 0.610092 |
| Prosap | 10023.55 | -0.69698 | 0.533839 | -1.30559 | 0.191692 | 0.6109 |
| CG1468 | 339.8211 | 3.243035 | 2.484195 | 1.305467 | 0.191734 | 0.6109 |
| GalT1 | 5537.307 | 1.463063 | 1.122217 | 1.303726 | 0.192327 | 0.612546 |
| CG6565 | 1672.806 | -1.7527 | 1.345572 | -1.30257 | 0.192723 | 0.613564 |
| Arp1 | 2111.76 | -1.45317 | 1.116188 | -1.3019 | 0.19295 | 0.614042 |
| CG8195 | 1575.274 | 1.241394 | 0.954104 | 1.301109 | 0.193221 | 0.614662 |
| Dh31 | 5441.18 | 1.246053 | 0.957873 | 1.300854 | 0.193308 | 0.614696 |
| XRCC1 | 162.5148 | 3.80251 | 2.924666 | 1.300152 | 0.193549 | 0.615217 |
| Optix | 326.9112 | -2.51592 | 1.936059 | -1.29951 | 0.19377 | 0.615676 |
| cal1 | 881.4802 | -1.95638 | 1.508067 | -1.29728 | 0.194536 | 0.616889 |
| flw | 11398.34 | -0.61072 | 0.470621 | -1.29768 | 0.194397 | 0.616889 |
| CG11275 | 1473.553 | 1.561377 | 1.202927 | 1.297982 | 0.194294 | 0.616889 |
| Adf1 | 4652.053 | -0.79663 | 0.613852 | -1.29776 | 0.194369 | 0.616889 |
| r-cup | 244.4275 | 3.619274 | 2.789852 | 1.297299 | 0.194528 | 0.616889 |
| Nckx30C | 44193.02 | 0.553922 | 0.427335 | 1.296224 | 0.194898 | 0.617428 |
| CG10543 | 6082.534 | 0.681331 | 0.525763 | 1.295889 | 0.195014 | 0.617428 |
| CG8777 | 413.5862 | 2.728556 | 2.104683 | 1.296422 | 0.19483 | 0.617428 |
| CG33158 | 1428.149 | -1.32751 | 1.024228 | -1.29611 | 0.194938 | 0.617428 |
| CG6484 | 111.6383 | -3.90257 | 3.013338 | -1.2951 | 0.195286 | 0.618046 |
| lncRNA:CR | 3935.85 | 0.782788 | 0.604694 | 1.29452 | 0.195486 | 0.618245 |
| ed | 1583.768 | -1.37007 | 1.058405 | -1.29447 | 0.195503 | 0.618245 |
| Irk2 | 12786.46 | -0.61592 | 0.476276 | -1.2932 | 0.195943 | 0.618662 |
| xit | 973.3995 | 1.306569 | 1.010267 | 1.293291 | 0.19591 | 0.618662 |
| CG4617 | 296.4546 | 2.416346 | 1.868327 | 1.293321 | 0.1959 | 0.618662 |
| PMCA | 17059.06 | 0.590146 | 0.456329 | 1.293245 | 0.195926 | 0.618662 |
| AP-2alpha | 7072.477 | -0.69958 | 0.541092 | -1.29291 | 0.196044 | 0.618736 |
| CG33170 | 946.3117 | 1.363983 | 1.055462 | 1.292309 | 0.19625 | 0.619145 |
| CG5708 | 6260.548 | 0.697311 | 0.540158 | 1.290939 | 0.196725 | 0.620155 |
| smg | 2676.543 | 0.909361 | 0.704411 | 1.290952 | 0.19672 | 0.620155 |
| Su(var)3-3 | 359.95 | -2.82472 | 2.189274 | -1.29026 | 0.196962 | 0.620657 |
| Col4a1 | 4005.594 | 0.922145 | 0.715102 | 1.289529 | 0.197214 | 0.62071 |
| CG30022 | 1631.41 | -1.33352 | 1.034048 | -1.28961 | 0.197186 | 0.62071 |

| | | | | | | |
|---|---|---|---|---|---|---|
| mtt | 2030.121 | -1.09304 | 0.847768 | -1.28932 | 0.197288 | 0.62071 |
| Nacalpha | 7474.175 | 0.648131 | 0.502463 | 1.289908 | 0.197083 | 0.62071 |
| CG2246 | 3792.122 | -0.86 | 0.667324 | -1.28872 | 0.197494 | 0.620961 |
| CG3558 | 1263.761 | 1.485595 | 1.152837 | 1.288643 | 0.197522 | 0.620961 |
| muc | 5508.812 | -0.70936 | 0.550869 | -1.28771 | 0.197847 | 0.621738 |
| Cat | 2714.974 | -1.00331 | 0.780018 | -1.28626 | 0.198352 | 0.622949 |
| CG4678 | 4836.902 | 0.731364 | 0.56874 | 1.285937 | 0.198465 | 0.622949 |
| CG5989 | 1224.944 | -1.76906 | 1.375587 | -1.28604 | 0.198429 | 0.622949 |
| lncRNA:CR | 328.3448 | 2.076315 | 1.616173 | 1.284711 | 0.198893 | 0.624049 |
| vri | 959.069 | -1.76611 | 1.375313 | -1.28415 | 0.19909 | 0.624421 |
| her | 1039.766 | -1.68471 | 1.313088 | -1.28301 | 0.199489 | 0.625428 |
| CG14435 | 1345.056 | 1.112051 | 0.868002 | 1.281162 | 0.200137 | 0.627216 |
| CG9590 | 779.5023 | -2.05346 | 1.603174 | -1.28087 | 0.200239 | 0.627291 |
| nudE | 2786.142 | 1.246957 | 0.973841 | 1.280453 | 0.200386 | 0.627505 |
| MFS14 | 608.5766 | -2.02894 | 1.584993 | -1.28009 | 0.200513 | 0.627658 |
| Cont | 1576.455 | -1.46342 | 1.144102 | -1.2791 | 0.200861 | 0.628502 |
| CG11700 | 66.89872 | 4.43034 | 3.467875 | 1.277537 | 0.201413 | 0.62997 |
| Tom40 | 3489.958 | -1.02149 | 0.799709 | -1.27733 | 0.201487 | 0.62997 |
| CG17162 | 279.7211 | 2.946366 | 2.308312 | 1.276416 | 0.201809 | 0.63073 |
| Orc5 | 130.8239 | -4.4184 | 3.466305 | -1.27467 | 0.202425 | 0.630937 |
| CG31075 | 311.5856 | -2.59936 | 2.037814 | -1.27556 | 0.202111 | 0.630937 |
| alpha-Man | 2640.025 | 1.189513 | 0.932496 | 1.275623 | 0.202089 | 0.630937 |
| CG10465 | 3644.168 | 0.763364 | 0.598625 | 1.275197 | 0.20224 | 0.630937 |
| tay | 5007.405 | -0.73047 | 0.57297 | -1.27489 | 0.202349 | 0.630937 |
| Sytalpha | 5055.435 | 0.805799 | 0.631984 | 1.275029 | 0.202299 | 0.630937 |
| baz | 267.8752 | 2.430517 | 1.905517 | 1.275516 | 0.202127 | 0.630937 |
| CkIIalpha-i | 2141.313 | 1.018606 | 0.799304 | 1.274365 | 0.202534 | 0.631032 |
| CG6356 | 6201.997 | -0.72883 | 0.572158 | -1.27382 | 0.202727 | 0.631388 |
| cno | 1333.048 | -1.43557 | 1.127213 | -1.27356 | 0.202819 | 0.631431 |
| Ubr3 | 7858.875 | 0.697344 | 0.548492 | 1.271384 | 0.203592 | 0.63359 |
| CG7149 | 5795.769 | -0.86912 | 0.683776 | -1.27107 | 0.203705 | 0.633698 |
| lncRNA:CR | 167.8753 | 2.650836 | 2.085939 | 1.270812 | 0.203796 | 0.633733 |
| CG17829 | 884.664 | -1.96428 | 1.54957 | -1.26763 | 0.204931 | 0.633825 |
| CIAPIN1 | 571.4696 | -1.64192 | 1.294772 | -1.26811 | 0.204757 | 0.633825 |
| Ndg | 81.10155 | -4.90502 | 3.867271 | -1.26834 | 0.204676 | 0.633825 |
| yellow-h | 78.06376 | -4.90433 | 3.867386 | -1.26813 | 0.204753 | 0.633825 |
| CG5789 | 265.2267 | -3.25917 | 2.566222 | -1.27003 | 0.204075 | 0.633825 |
| Tbh | 1845.716 | -1.68928 | 1.330113 | -1.27003 | 0.204075 | 0.633825 |
| mbc | 2899.539 | -0.86272 | 0.679266 | -1.27008 | 0.204057 | 0.633825 |
| betaggt-I | 452.5142 | 2.402832 | 1.894497 | 1.268322 | 0.204683 | 0.633825 |
| nej | 4562.142 | -0.79767 | 0.628906 | -1.26835 | 0.204672 | 0.633825 |
| CG3760 | 1513.529 | 1.112311 | 0.877387 | 1.267754 | 0.204886 | 0.633825 |
| lncRNA:CR | 442.1358 | 2.036288 | 1.606062 | 1.267876 | 0.204842 | 0.633825 |
| sima | 7653.28 | -0.65858 | 0.51907 | -1.26876 | 0.204525 | 0.633825 |
| MsrA | 4159.617 | 0.823374 | 0.649486 | 1.267731 | 0.204894 | 0.633825 |
| D19A | 599.4875 | 2.20738 | 1.739139 | 1.269237 | 0.204356 | 0.633825 |
| CG4462 | 285.7221 | 3.372108 | 2.661835 | 1.266836 | 0.205214 | 0.634456 |
| lig | 2628.84 | -0.92514 | 0.730663 | -1.26617 | 0.205452 | 0.634947 |
| Hsp60C | 131.2162 | 4.178175 | 3.300785 | 1.265813 | 0.20558 | 0.63501 |
| Sply | 3841.673 | 0.844224 | 0.667334 | 1.265069 | 0.205847 | 0.63501 |

| | | | | | | |
|---|---|---|---|---|---|---|
| Rpn6 | 1516.452 | -1.63771 | 1.294622 | -1.26501 | 0.205868 | 0.63501 |
| lncRNA:CR | 260.6201 | 2.692816 | 2.128513 | 1.265116 | 0.20583 | 0.63501 |
| ttv | 5047.587 | -0.68669 | 0.54275 | -1.26521 | 0.205797 | 0.63501 |
| Culd | 205.9061 | -3.41968 | 2.704451 | -1.26446 | 0.206063 | 0.635368 |
| CG10311 | 1594.32 | 1.05607 | 0.835559 | 1.26391 | 0.206262 | 0.635738 |
| ctp | 24644.26 | 0.540212 | 0.427744 | 1.262934 | 0.206613 | 0.636574 |
| CG5937 | 1407.759 | 1.919794 | 1.521304 | 1.26194 | 0.20697 | 0.636945 |
| Sf3b1 | 987.8599 | -1.91577 | 1.519188 | -1.26105 | 0.207291 | 0.636945 |
| Atx2 | 3341.496 | 0.885018 | 0.702183 | 1.260381 | 0.207532 | 0.636945 |
| mRpL55 | 3172.555 | 0.866595 | 0.687462 | 1.260571 | 0.207463 | 0.636945 |
| CG31126 | 1162.398 | 1.890427 | 1.498947 | 1.26117 | 0.207248 | 0.636945 |
| Phb2 | 4157.325 | -0.73523 | 0.58318 | -1.26073 | 0.207408 | 0.636945 |
| GstE3 | 1938.02 | -1.7243 | 1.368194 | -1.26027 | 0.207571 | 0.636945 |
| MED16 | 219.843 | -3.38216 | 2.683878 | -1.26018 | 0.207606 | 0.636945 |
| Cdk5 | 335.8457 | -2.12875 | 1.687786 | -1.26127 | 0.207211 | 0.636945 |
| mxc | 1168.257 | 1.348344 | 1.069128 | 1.261162 | 0.20725 | 0.636945 |
| Pxn | 868.6456 | -2.01016 | 1.594099 | -1.261 | 0.207309 | 0.636945 |
| CG8635 | 1777.355 | 1.11834 | 0.887824 | 1.259641 | 0.207799 | 0.636979 |
| Atg10 | 307.0521 | -2.20727 | 1.752798 | -1.25928 | 0.207929 | 0.636979 |
| Elp2 | 353.5727 | 2.54336 | 2.019719 | 1.259264 | 0.207935 | 0.636979 |
| tea | 1044.278 | 1.215767 | 0.965015 | 1.259843 | 0.207726 | 0.636979 |
| dre4 | 308.8461 | 2.520257 | 2.0024 | 1.258618 | 0.208168 | 0.637451 |
| CG16812 | 243.2139 | -2.65706 | 2.112336 | -1.25788 | 0.208437 | 0.637833 |
| Galk | 1337.483 | -1.35527 | 1.077467 | -1.25783 | 0.208452 | 0.637833 |
| baf | 3490.811 | 0.939566 | 0.748146 | 1.25586 | 0.209167 | 0.638203 |
| CG6583 | 1868.087 | -1.14727 | 0.913383 | -1.25607 | 0.209092 | 0.638203 |
| Sirt1 | 2065.697 | 1.141073 | 0.908014 | 1.256669 | 0.208873 | 0.638203 |
| mRpL4 | 1593.834 | 1.231777 | 0.980915 | 1.255743 | 0.209209 | 0.638203 |
| MED7 | 951.4324 | -1.39443 | 1.110165 | -1.25606 | 0.209094 | 0.638203 |
| CG11391 | 2970.926 | 1.309006 | 1.042136 | 1.25608 | 0.209087 | 0.638203 |
| Acf | 719.1344 | -1.51585 | 1.206215 | -1.2567 | 0.208862 | 0.638203 |
| LeuRS-m | 605.7535 | -1.78134 | 1.418334 | -1.25594 | 0.209139 | 0.638203 |
| Sps1 | 287.4814 | -2.16384 | 1.723749 | -1.25531 | 0.209366 | 0.63844 |
| salr | 193.3387 | 2.939961 | 2.342665 | 1.254964 | 0.209492 | 0.63858 |
| CG8677 | 7353.131 | -0.89382 | 0.713364 | -1.25296 | 0.21022 | 0.640557 |
| PK2-R2 | 134.3573 | 3.560128 | 2.843261 | 1.252129 | 0.210523 | 0.640993 |
| pico | 3945.288 | -0.81443 | 0.650387 | -1.25223 | 0.210487 | 0.640993 |
| Nup154 | 242.9026 | 2.8439 | 2.273016 | 1.251157 | 0.210877 | 0.641828 |
| sno | 3769.053 | -0.89141 | 0.712813 | -1.25056 | 0.211096 | 0.642251 |
| CG4557 | 207.8462 | 3.1145 | 2.492227 | 1.249686 | 0.211414 | 0.642975 |
| Tsp66E | 11477.64 | 0.716969 | 0.573875 | 1.249348 | 0.211538 | 0.643107 |
| beag | 285.9391 | 2.787988 | 2.232222 | 1.248974 | 0.211675 | 0.643279 |
| ctrip | 3923.261 | 0.836345 | 0.670144 | 1.248009 | 0.212028 | 0.644108 |
| CG4751 | 396.2862 | 1.916898 | 1.537876 | 1.246458 | 0.212596 | 0.64556 |
| grp | 1927.927 | -0.96497 | 0.774307 | -1.24624 | 0.212676 | 0.64556 |
| lncRNA:CR | 1129.801 | 1.284615 | 1.030951 | 1.246048 | 0.212747 | 0.64556 |
| CG3191 | 2000.143 | -1.17511 | 0.943487 | -1.24549 | 0.212951 | 0.64569 |
| Ars2 | 4121.408 | -0.74854 | 0.600953 | -1.24558 | 0.212918 | 0.64569 |
| Tm2 | 134.5302 | -3.34653 | 2.688702 | -1.24466 | 0.213255 | 0.646126 |
| RhoGAP18 | 2637.722 | -0.87804 | 0.705432 | -1.24468 | 0.213248 | 0.646126 |

| | | | | | | |
|---|---|---|---|---|---|---|
| lbf2 | 79.74908 | -4.00165 | 3.218618 | -1.24328 | 0.213764 | 0.647423 |
| CG3817 | 130.5576 | -4.02445 | 3.239013 | -1.24249 | 0.214054 | 0.647527 |
| CG30020 | 1146.716 | 1.92419 | 1.548812 | 1.242365 | 0.214102 | 0.647527 |
| Plp | 2755.636 | 0.942799 | 0.758528 | 1.242932 | 0.213893 | 0.647527 |
| CG7872 | 805.4906 | -1.88807 | 1.519805 | -1.24231 | 0.214121 | 0.647527 |
| CG10208 | 1309.959 | -1.27016 | 1.022624 | -1.24206 | 0.214214 | 0.647565 |
| LPCAT | 1371.548 | 1.081023 | 0.871263 | 1.240754 | 0.214697 | 0.648732 |
| CG3732 | 1677.549 | 1.143034 | 0.921403 | 1.240536 | 0.214777 | 0.648732 |
| Cbs | 567.6381 | 1.842847 | 1.485737 | 1.240359 | 0.214843 | 0.648732 |
| Membrin | 592.8414 | 1.724869 | 1.391126 | 1.239908 | 0.215009 | 0.648746 |
| Herc4 | 3008.693 | 0.790433 | 0.637467 | 1.239959 | 0.214991 | 0.648746 |
| phyl | 2308.883 | 0.902306 | 0.728127 | 1.239216 | 0.215266 | 0.649275 |
| CG7382 | 1520.458 | -1.0584 | 0.855019 | -1.23787 | 0.215765 | 0.649292 |
| SCAR | 1377.987 | -1.57662 | 1.273953 | -1.23758 | 0.215871 | 0.649292 |
| GstE2 | 7270.429 | -0.60775 | 0.491192 | -1.2373 | 0.215977 | 0.649292 |
| CG12056 | 771.9236 | -1.42967 | 1.155244 | -1.23755 | 0.215884 | 0.649292 |
| Gga | 670.7408 | 2.111611 | 1.705012 | 1.238473 | 0.215541 | 0.649292 |
| Ir93a | 233.084 | 2.558307 | 2.067178 | 1.237585 | 0.21587 | 0.649292 |
| hoip | 512.045 | -2.03705 | 1.645501 | -1.23795 | 0.215735 | 0.649292 |
| Ncc69 | 6548.738 | -0.9616 | 0.777214 | -1.23724 | 0.215999 | 0.649292 |
| ppk26 | 341.7271 | 3.192921 | 2.580319 | 1.237413 | 0.215934 | 0.649292 |
| xmas | 506.0936 | 2.401945 | 1.942636 | 1.236436 | 0.216297 | 0.649943 |
| prtp | 2358.825 | -0.89514 | 0.724372 | -1.23575 | 0.216553 | 0.650227 |
| Nup205 | 65.88088 | 4.703808 | 3.806042 | 1.235879 | 0.216504 | 0.650227 |
| x16 | 1311.654 | 1.388322 | 1.123792 | 1.235391 | 0.216685 | 0.65038 |
| CG5439 | 465.2087 | -2.2104 | 1.789869 | -1.23495 | 0.216848 | 0.650382 |
| mt:CoI | 777542 | 0.452085 | 0.366028 | 1.235111 | 0.216789 | 0.650382 |
| Eato | 752.0366 | -1.85147 | 1.500071 | -1.23425 | 0.217108 | 0.650676 |
| CG11560 | 1631.962 | 1.760482 | 1.426271 | 1.234325 | 0.217082 | 0.650676 |
| GAPcenA | 2116.868 | -0.99643 | 0.80759 | -1.23383 | 0.217267 | 0.650684 |
| CG8974 | 1977.941 | 1.072374 | 0.869155 | 1.233812 | 0.217273 | 0.650684 |
| TfIIA-L | 988.1075 | -1.34854 | 1.093828 | -1.23286 | 0.217628 | 0.651503 |
| lncRNA:CR | 215.7565 | 2.67458 | 2.169795 | 1.232641 | 0.21771 | 0.651505 |
| CG5028 | 8056.569 | 0.737093 | 0.598818 | 1.230914 | 0.218355 | 0.652497 |
| CG17078 | 1187.418 | 1.735025 | 1.408888 | 1.231485 | 0.218142 | 0.652497 |
| cass | 7337.266 | 0.677884 | 0.550729 | 1.230885 | 0.218366 | 0.652497 |
| RpS5a | 16447.38 | 0.51831 | 0.421087 | 1.230886 | 0.218365 | 0.652497 |
| Dap160 | 8174.955 | 0.637431 | 0.518286 | 1.229883 | 0.218741 | 0.653374 |
| CG7988 | 495.3031 | 2.552677 | 2.076783 | 1.22915 | 0.219016 | 0.653951 |
| su(sable) | 1734.978 | -0.9879 | 0.804833 | -1.22745 | 0.219652 | 0.654875 |
| CG13995 | 2913.147 | -1.43983 | 1.172898 | -1.22758 | 0.219604 | 0.654875 |
| lncRNA:CR | 169.4895 | 3.538555 | 2.88228 | 1.227693 | 0.219562 | 0.654875 |
| mus304 | 116.9229 | -3.69258 | 3.007093 | -1.22796 | 0.219463 | 0.654875 |
| ClC-a | 9087.179 | -0.63516 | 0.517793 | -1.22667 | 0.219946 | 0.655435 |
| CG17839 | 1063.416 | 1.181307 | 0.963136 | 1.226521 | 0.220003 | 0.655435 |
| CG6983 | 3702.264 | 0.864226 | 0.704873 | 1.226073 | 0.220171 | 0.655693 |
| CG13367 | 1005.499 | 1.97166 | 1.611317 | 1.223633 | 0.221091 | 0.656241 |
| CG13248 | 2187.769 | 1.574498 | 1.286311 | 1.224042 | 0.220936 | 0.656241 |
| Mes2 | 2411.409 | 0.871666 | 0.712182 | 1.223938 | 0.220976 | 0.656241 |
| GM130 | 597.0389 | -1.59398 | 1.302621 | -1.22367 | 0.221077 | 0.656241 |

| | | | | | | |
|---|---|---|---|---|---|---|
| CG8388 | 838.5695 | -2.18623 | 1.785422 | -1.22449 | 0.220768 | 0.656241 |
| Tret1-1 | 1582.384 | -1.27493 | 1.041279 | -1.22439 | 0.220807 | 0.656241 |
| mfas | 25780.32 | -0.57955 | 0.47334 | -1.22437 | 0.220811 | 0.656241 |
| CG8173 | 147.2017 | 3.440271 | 2.808678 | 1.224872 | 0.220624 | 0.656241 |
| Ankle2 | 779.7814 | -1.90106 | 1.551526 | -1.22528 | 0.220469 | 0.656241 |
| Pdk | 9913.759 | -0.58582 | 0.478866 | -1.22335 | 0.221197 | 0.656313 |
| syd | 2691.808 | -0.87788 | 0.717727 | -1.22313 | 0.221279 | 0.656315 |
| cyst | 907.8468 | -1.28945 | 1.054532 | -1.22277 | 0.221418 | 0.656482 |
| bwa | 3900.063 | 0.924765 | 0.756927 | 1.221737 | 0.221807 | 0.656667 |
| Smyd5 | 305.8946 | 2.711982 | 2.21938 | 1.221955 | 0.221725 | 0.656667 |
| RpL23A | 9604.552 | 0.5807 | 0.475283 | 1.221799 | 0.221784 | 0.656667 |
| Spn | 4806.605 | -0.68484 | 0.560252 | -1.22237 | 0.221567 | 0.656667 |
| Ppa | 1262.554 | 1.14796 | 0.939843 | 1.221438 | 0.22192 | 0.656759 |
| CG5810 | 175.4711 | 3.134348 | 2.569185 | 1.219978 | 0.222473 | 0.658154 |
| Tsp5D | 14764.38 | -0.57246 | 0.47013 | -1.21767 | 0.223349 | 0.659123 |
| CG6712 | 199.1023 | -2.98005 | 2.446657 | -1.21801 | 0.223221 | 0.659123 |
| CG6937 | 115.8083 | 3.400775 | 2.793015 | 1.2176 | 0.223376 | 0.659123 |
| CG10513 | 18487.94 | -0.67011 | 0.550203 | -1.21793 | 0.223251 | 0.659123 |
| CG1344 | 1911.304 | -1.42437 | 1.169252 | -1.21819 | 0.223151 | 0.659123 |
| Prosalpha2 | 2273.224 | 0.93061 | 0.764168 | 1.217808 | 0.223297 | 0.659123 |
| CG10672 | 944.2735 | -1.50031 | 1.231238 | -1.21854 | 0.22302 | 0.659123 |
| CG3909 | 680.1454 | -1.58361 | 1.301555 | -1.21671 | 0.223716 | 0.659127 |
| Cyp6a23 | 705.8034 | 1.745625 | 1.435147 | 1.216338 | 0.223856 | 0.659127 |
| sff | 5437.269 | 0.739954 | 0.608364 | 1.216301 | 0.22387 | 0.659127 |
| MED14 | 310.5603 | -2.94857 | 2.423194 | -1.21681 | 0.223676 | 0.659127 |
| sNPF-R | 728.8763 | -2.30237 | 1.892569 | -1.21653 | 0.223783 | 0.659127 |
| galla-2 | 536.0149 | -1.61698 | 1.329236 | -1.21648 | 0.223804 | 0.659127 |
| Dmtn | 2464.618 | -0.93197 | 0.766852 | -1.21532 | 0.224245 | 0.659964 |
| CR15345 | 160.9399 | -3.70248 | 3.046993 | -1.21512 | 0.224319 | 0.659964 |
| eIF2Balpha | 1378.201 | 1.116072 | 0.918692 | 1.214849 | 0.224424 | 0.660031 |
| pes | 238.3977 | 2.520642 | 2.076077 | 1.214137 | 0.224695 | 0.660105 |
| Kap3 | 1428.496 | 1.312126 | 1.080395 | 1.214487 | 0.224562 | 0.660105 |
| cactin | 144.5622 | -3.00058 | 2.471075 | -1.21428 | 0.22464 | 0.660105 |
| CG12344 | 1970.836 | 1.421843 | 1.171413 | 1.213785 | 0.22483 | 0.660126 |
| Amun | 839.8067 | -1.49218 | 1.229617 | -1.21354 | 0.224925 | 0.660126 |
| Rgl | 2163.426 | -0.96671 | 0.796645 | -1.21347 | 0.224949 | 0.660126 |
| ifc | 3053.751 | -0.83516 | 0.688999 | -1.21214 | 0.225458 | 0.660894 |
| Ns3 | 403.0116 | -1.93745 | 1.598012 | -1.21241 | 0.225355 | 0.660894 |
| Rh50 | 16298.91 | -0.6396 | 0.527646 | -1.21217 | 0.225447 | 0.660894 |
| unc79 | 14156.73 | 0.788779 | 0.650933 | 1.211768 | 0.225601 | 0.661072 |
| p53 | 240.4001 | 3.750952 | 3.09728 | 1.211047 | 0.225877 | 0.661639 |
| CG3823 | 62.73417 | -4.61156 | 3.809225 | -1.21063 | 0.226037 | 0.661756 |
| CG17292 | 256.6922 | -2.95497 | 2.441602 | -1.21026 | 0.22618 | 0.661756 |
| CG9336 | 605.0463 | 1.560332 | 1.289442 | 1.210083 | 0.226247 | 0.661756 |
| CG33506 | 220.6125 | 2.888851 | 2.387288 | 1.210097 | 0.226241 | 0.661756 |
| Pkc98E | 11494.33 | -0.58382 | 0.482663 | -1.20958 | 0.226441 | 0.661842 |
| CG2064 | 257.0568 | -3.25921 | 2.694032 | -1.20979 | 0.22636 | 0.661842 |
| hrm | 4636.118 | -0.89368 | 0.739013 | -1.20928 | 0.226554 | 0.661929 |
| CG15812 | 156.6184 | 3.449162 | 2.852815 | 1.209038 | 0.226648 | 0.661965 |
| CG9941 | 916.7044 | -1.32939 | 1.100566 | -1.20791 | 0.227081 | 0.662986 |

| | | | | | | |
|---|---|---|---|---|---|---|
| nAChRalpł | 15985.89 | 0.531325 | 0.440269 | 1.206818 | 0.227502 | 0.663976 |
| Lim3 | 220.9549 | 2.815996 | 2.335512 | 1.20573 | 0.227922 | 0.664717 |
| CG6574 | 1193.863 | 1.421905 | 1.179285 | 1.205735 | 0.22792 | 0.664717 |
| chn | 4632.03 | -0.72434 | 0.60099 | -1.20524 | 0.228111 | 0.665028 |
| Mtpalpha | 2406.828 | -1.30071 | 1.081127 | -1.20311 | 0.228934 | 0.666442 |
| CG11920 | 265.8487 | 2.726858 | 2.265785 | 1.203493 | 0.228785 | 0.666442 |
| fkh | 467.756 | 1.89652 | 1.575805 | 1.203525 | 0.228773 | 0.666442 |
| bab1 | 79.9409 | -3.55004 | 2.951213 | -1.20291 | 0.229011 | 0.666442 |
| g | 1520.703 | 0.996734 | 0.828405 | 1.203196 | 0.228901 | 0.666442 |
| CR14033 | 1781.699 | -1.18427 | 0.985323 | -1.20191 | 0.229398 | 0.666599 |
| CG6420 | 1192.745 | -1.2436 | 1.034602 | -1.202 | 0.229362 | 0.666599 |
| chrb | 15125.57 | -0.54924 | 0.456736 | -1.20253 | 0.229158 | 0.666599 |
| Set1 | 7657.342 | 0.615532 | 0.512056 | 1.202079 | 0.229333 | 0.666599 |
| CG30394 | 851.9972 | 1.33879 | 1.114333 | 1.201427 | 0.229585 | 0.666903 |
| CG10265 | 972.724 | 1.709847 | 1.423467 | 1.201185 | 0.229679 | 0.666934 |
| CG14252 | 286.9518 | -2.55366 | 2.126715 | -1.20075 | 0.229848 | 0.667182 |
| AANATL2 | 1476.188 | 1.432324 | 1.193287 | 1.200318 | 0.230016 | 0.667429 |
| CG4658 | 709.9847 | 1.442396 | 1.201935 | 1.200061 | 0.230116 | 0.667437 |
| Tfb1 | 538.654 | 1.805284 | 1.50455 | 1.199883 | 0.230185 | 0.667437 |
| CG32813 | 597.7917 | 1.478238 | 1.237982 | 1.19407 | 0.23245 | 0.667724 |
| N | 5014.033 | -0.65928 | 0.553995 | -1.19005 | 0.234028 | 0.667724 |
| HIPP1 | 2090.968 | -1.72008 | 1.444494 | -1.19078 | 0.233739 | 0.667724 |
| Liprin-alph | 2967.62 | 1.08388 | 0.907229 | 1.194715 | 0.232198 | 0.667724 |
| Rca1 | 1259.624 | 1.571829 | 1.320645 | 1.190198 | 0.233969 | 0.667724 |
| CG1142 | 851.6314 | -1.5781 | 1.325555 | -1.19052 | 0.233843 | 0.667724 |
| Elp1 | 476.5085 | -2.27369 | 1.908133 | -1.19158 | 0.233427 | 0.667724 |
| PGAP3 | 1155.634 | 1.614005 | 1.352381 | 1.193454 | 0.232691 | 0.667724 |
| CG30158 | 2977.399 | 1.391381 | 1.166838 | 1.192437 | 0.23309 | 0.667724 |
| Inx2 | 15020.5 | -0.61011 | 0.510152 | -1.19594 | 0.231721 | 0.667724 |
| CG5065 | 5836.926 | -0.64352 | 0.537664 | -1.19688 | 0.231354 | 0.667724 |
| prim | 2285.633 | -0.94315 | 0.787417 | -1.19778 | 0.231004 | 0.667724 |
| Spred | 5281.025 | 0.810477 | 0.677929 | 1.19552 | 0.231884 | 0.667724 |
| Roe1 | 1799.016 | -1.54049 | 1.287895 | -1.19613 | 0.231645 | 0.667724 |
| pnt | 2119.762 | -0.8635 | 0.723306 | -1.19382 | 0.232549 | 0.667724 |
| Start1 | 1352.743 | -1.14869 | 0.965059 | -1.19028 | 0.233937 | 0.667724 |
| CG3009 | 3651.197 | 0.816236 | 0.685197 | 1.191243 | 0.233558 | 0.667724 |
| spel1 | 1069.664 | -1.90379 | 1.596807 | -1.19225 | 0.233165 | 0.667724 |
| CG14995 | 5389.9 | 0.678489 | 0.569177 | 1.192054 | 0.23324 | 0.667724 |
| CG42232 | 4260.756 | 0.781602 | 0.655443 | 1.192478 | 0.233074 | 0.667724 |
| Pde9 | 7001.444 | 0.701194 | 0.586973 | 1.194593 | 0.232246 | 0.667724 |
| CG42365 | 789.4097 | -2.07861 | 1.73364 | -1.19899 | 0.230532 | 0.667724 |
| wech | 5726.142 | -0.62946 | 0.52822 | -1.19166 | 0.233396 | 0.667724 |
| CG42445 | 249.7165 | -2.91173 | 2.434801 | -1.19588 | 0.231744 | 0.667724 |
| lncRNA:CR | 10905.45 | 0.593106 | 0.495458 | 1.197086 | 0.231273 | 0.667724 |
| Chi | 1229.098 | -1.25774 | 1.055496 | -1.19161 | 0.233414 | 0.667724 |
| CG40486 | 280.2859 | 3.557676 | 2.980019 | 1.193844 | 0.232539 | 0.667724 |
| orb2 | 10745.59 | 0.553779 | 0.463214 | 1.195516 | 0.231885 | 0.667724 |
| CG43813 | 225.282 | 3.437662 | 2.882138 | 1.192747 | 0.232968 | 0.667724 |
| asRNA:CR4 | 596.1106 | 1.800053 | 1.50199 | 1.198446 | 0.230744 | 0.667724 |
| lncRNA:CR | 230.3198 | 3.373812 | 2.813191 | 1.199283 | 0.230418 | 0.667724 |

| | | | | | | |
|---|---|---|---|---|---|---|
| lncRNA:CR | 281.0032 | 2.387525 | 1.99661 | 1.195789 | 0.231779 | 0.667724 |
| gry | 379.8824 | -1.71826 | 1.43556 | -1.19693 | 0.231334 | 0.667724 |
| l(3)80Fg | 30907 | 0.472977 | 0.394683 | 1.198372 | 0.230772 | 0.667724 |
| 18w | 3476.911 | -0.72086 | 0.605373 | -1.19077 | 0.233743 | 0.667724 |
| l(3)02640 | 2253.677 | 0.838077 | 0.703488 | 1.191316 | 0.23353 | 0.667724 |
| dos | 1233.381 | -1.21198 | 1.015494 | -1.19349 | 0.232677 | 0.667724 |
| CG1552 | 29751.57 | -0.54062 | 0.45264 | -1.19437 | 0.232332 | 0.667724 |
| CG1637 | 7310.725 | 0.603353 | 0.505336 | 1.193963 | 0.232492 | 0.667724 |
| Alg14 | 672.5097 | -1.38259 | 1.155777 | -1.19624 | 0.231603 | 0.667724 |
| bnb | 28811.54 | -0.58856 | 0.493031 | -1.19377 | 0.232569 | 0.667724 |
| Prosbeta6 | 3039.106 | -0.76334 | 0.640361 | -1.19205 | 0.233243 | 0.667724 |
| RhoGAP71 | 4999.28 | -0.81447 | 0.683961 | -1.19081 | 0.233727 | 0.667724 |
| Liprin-beta | 819.476 | 1.761068 | 1.473426 | 1.19522 | 0.232001 | 0.667724 |
| Hsp26 | 788.2891 | -1.47271 | 1.235543 | -1.19195 | 0.233279 | 0.667724 |
| Slh | 685.909 | -1.70547 | 1.433511 | -1.18972 | 0.234157 | 0.667856 |
| CG13506 | 563.3044 | 1.808982 | 1.522679 | 1.188026 | 0.234823 | 0.669193 |
| Rap2l | 3148.822 | 0.782164 | 0.658308 | 1.188144 | 0.234777 | 0.669193 |
| Paf-AHalph | 1114.137 | -1.1393 | 0.959097 | -1.18789 | 0.234876 | 0.669193 |
| crn | 81.38455 | 4.502695 | 3.791764 | 1.187493 | 0.235033 | 0.669257 |
| Uba5 | 725.4496 | 2.053387 | 1.729298 | 1.187411 | 0.235065 | 0.669257 |
| CG5021 | 445.2569 | -1.72352 | 1.452332 | -1.18673 | 0.235335 | 0.669788 |
| CG3626 | 970.4633 | -1.3901 | 1.173275 | -1.1848 | 0.236096 | 0.670556 |
| l(2)gl | 1232.111 | -1.59593 | 1.347317 | -1.18453 | 0.236204 | 0.670556 |
| P5CDh1 | 15524.76 | 0.605458 | 0.510894 | 1.185095 | 0.23598 | 0.670556 |
| dpr19 | 9148.842 | 0.623424 | 0.526148 | 1.184884 | 0.236063 | 0.670556 |
| CG9649 | 3874.232 | 0.716514 | 0.605412 | 1.183515 | 0.236605 | 0.670556 |
| CG31469 | 354.1561 | 2.259854 | 1.907994 | 1.184413 | 0.23625 | 0.670556 |
| Not1 | 9373.397 | 0.617469 | 0.52158 | 1.183843 | 0.236475 | 0.670556 |
| CG42458 | 94.19074 | 4.382127 | 3.702273 | 1.183632 | 0.236559 | 0.670556 |
| Brms1 | 943.8747 | -1.66297 | 1.402919 | -1.18537 | 0.235873 | 0.670556 |
| jub | 136.4862 | -3.01597 | 2.544735 | -1.18518 | 0.235946 | 0.670556 |
| RhoGAPp1 | 5726.67 | 0.624522 | 0.527687 | 1.183508 | 0.236608 | 0.670556 |
| lqf | 6614.188 | -0.62442 | 0.527587 | -1.18354 | 0.236594 | 0.670556 |
| Syb | 3137.745 | -0.86247 | 0.729752 | -1.18186 | 0.23726 | 0.672167 |
| Dip-C | 1754.394 | 1.150282 | 0.973983 | 1.181008 | 0.237599 | 0.672833 |
| Atg12 | 1265.998 | -1.08877 | 0.922187 | -1.18064 | 0.237746 | 0.672833 |
| TH1 | 365.9924 | -2.20756 | 1.86963 | -1.18075 | 0.237702 | 0.672833 |
| CYLD | 1957.907 | 0.905991 | 0.767716 | 1.180113 | 0.237955 | 0.673105 |
| CG15706 | 536.9696 | -1.89478 | 1.605783 | -1.17997 | 0.238011 | 0.673105 |
| CG11089 | 562.9503 | 2.169381 | 1.839371 | 1.179415 | 0.238233 | 0.673113 |
| ADD1 | 877.8322 | 1.603022 | 1.359259 | 1.179335 | 0.238265 | 0.673113 |
| mRpS30 | 2111.203 | 0.943685 | 0.8 | 1.179606 | 0.238157 | 0.673113 |
| E23 | 761.7385 | 1.293921 | 1.098015 | 1.178419 | 0.23863 | 0.67367 |
| lncRNA:CR | 308.7644 | 1.865729 | 1.583225 | 1.178436 | 0.238623 | 0.67367 |
| CG6153 | 139.8896 | 3.407445 | 2.893841 | 1.177482 | 0.239003 | 0.674409 |
| CG14694 | 220.1326 | -3.44123 | 2.922887 | -1.17734 | 0.23906 | 0.674409 |
| CG6527 | 189.353 | -2.9274 | 2.487561 | -1.17682 | 0.239269 | 0.674761 |
| Snx6 | 974.867 | 2.117302 | 1.802085 | 1.174918 | 0.240028 | 0.675004 |
| BicD | 3212.223 | 0.76141 | 0.64726 | 1.176359 | 0.239452 | 0.675004 |
| CG10428 | 156.9353 | -3.60782 | 3.068641 | -1.17571 | 0.239712 | 0.675004 |

| | | | | | | |
|---|---|---|---|---|---|---|
| shg | 669.3132 | -1.4514 | 1.234936 | -1.17528 | 0.239881 | 0.675004 |
| Rme-8 | 1268.605 | 1.285134 | 1.093754 | 1.174975 | 0.240005 | 0.675004 |
| CG11127 | 687.1674 | 1.848822 | 1.573267 | 1.175148 | 0.239935 | 0.675004 |
| CG3735 | 385.4759 | 2.229672 | 1.897477 | 1.175072 | 0.239966 | 0.675004 |
| Ptpa | 919.9111 | -1.71045 | 1.455496 | -1.17517 | 0.239927 | 0.675004 |
| CG8671 | 4316.788 | -0.71374 | 0.607986 | -1.17394 | 0.24042 | 0.675736 |
| na | 4939.006 | 0.92643 | 0.789226 | 1.173847 | 0.240456 | 0.675736 |
| PpD3 | 1415.23 | 0.990401 | 0.844272 | 1.173083 | 0.240763 | 0.676092 |
| CG12338 | 714.9725 | -2.19262 | 1.868925 | -1.1732 | 0.240716 | 0.676092 |
| Rim2 | 1561.36 | 1.728588 | 1.473772 | 1.172901 | 0.240836 | 0.676092 |
| ths | 5583.116 | 0.681221 | 0.581472 | 1.171547 | 0.241379 | 0.677187 |
| dpr8 | 3177.965 | 0.7527 | 0.642515 | 1.17149 | 0.241402 | 0.677187 |
| CG7339 | 1041.889 | -1.58688 | 1.354801 | -1.1713 | 0.241479 | 0.677187 |
| zetaCOP | 4920.071 | -0.65385 | 0.558578 | -1.17055 | 0.241778 | 0.677553 |
| Uch-L5 | 1001.046 | -1.31482 | 1.123078 | -1.17073 | 0.241708 | 0.677553 |
| dgrn | 1592.597 | -0.98303 | 0.840126 | -1.1701 | 0.241962 | 0.67781 |
| CG8207 | 158.3078 | -3.15639 | 2.698902 | -1.16951 | 0.242199 | 0.67781 |
| CG43092 | 197.858 | 2.194885 | 1.87713 | 1.169277 | 0.242292 | 0.67781 |
| Hlc | 1303.517 | 1.24532 | 1.064913 | 1.16941 | 0.242238 | 0.67781 |
| CG3008 | 453.9166 | 1.706401 | 1.458606 | 1.169885 | 0.242047 | 0.67781 |
| CG4297 | 3315.401 | -0.74349 | 0.636358 | -1.16836 | 0.242663 | 0.678374 |
| Stacl | 21613.71 | 0.517684 | 0.443081 | 1.168374 | 0.242656 | 0.678374 |
| alrm | 15593.65 | -0.51941 | 0.445023 | -1.16714 | 0.243153 | 0.679508 |
| SmydA-7 | 401.5204 | 2.596013 | 2.225253 | 1.166615 | 0.243366 | 0.679539 |
| Pax | 1755.176 | 0.990246 | 0.848904 | 1.166499 | 0.243413 | 0.679539 |
| asf1 | 1261.061 | -1.80473 | 1.547433 | -1.16628 | 0.243503 | 0.679539 |
| NHP2 | 1894.073 | -1.67013 | 1.431818 | -1.16644 | 0.243436 | 0.679539 |
| CG32221 | 446.5501 | -1.95171 | 1.675116 | -1.16512 | 0.24397 | 0.680606 |
| eIF1A | 5480.751 | -0.73007 | 0.626881 | -1.16461 | 0.244178 | 0.680948 |
| CHKov2 | 2037.539 | 1.610477 | 1.384354 | 1.163341 | 0.244691 | 0.681578 |
| CG1647 | 555.6808 | 1.70795 | 1.468303 | 1.163213 | 0.244743 | 0.681578 |
| CG12112 | 361.2078 | -2.60854 | 2.241761 | -1.16361 | 0.244581 | 0.681578 |
| osp | 1911.165 | -1.07411 | 0.922968 | -1.16376 | 0.244521 | 0.681578 |
| DIP-kappa | 1020.112 | 1.869716 | 1.608134 | 1.162662 | 0.244967 | 0.681964 |
| FRG1 | 230.3828 | -2.3591 | 2.031897 | -1.16104 | 0.245628 | 0.683567 |
| mod(mdg4 | 4023.595 | -0.74529 | 0.642388 | -1.16019 | 0.245973 | 0.683835 |
| Trs23 | 604.7027 | 2.023391 | 1.744177 | 1.160084 | 0.246015 | 0.683835 |
| Ack-like | 903.9007 | 1.205844 | 1.039188 | 1.160372 | 0.245898 | 0.683835 |
| KdelR | 3974.729 | -0.68248 | 0.588368 | -1.15996 | 0.246065 | 0.683835 |
| Rchy1 | 1347.02 | 1.378558 | 1.189629 | 1.158813 | 0.246532 | 0.684128 |
| Uvrag | 464.3265 | -2.01469 | 1.739983 | -1.15788 | 0.246914 | 0.684128 |
| CG10283 | 480.32 | -1.84133 | 1.590111 | -1.15799 | 0.246868 | 0.684128 |
| mfrn | 3146.591 | 0.790423 | 0.682682 | 1.157821 | 0.246937 | 0.684128 |
| CG30389 | 3157.231 | 0.755189 | 0.651833 | 1.158562 | 0.246635 | 0.684128 |
| pix | 5569.846 | 0.653263 | 0.563812 | 1.158654 | 0.246597 | 0.684128 |
| CG3548 | 1704.009 | 1.150613 | 0.992348 | 1.159485 | 0.246258 | 0.684128 |
| mthl1 | 527.6822 | 1.630481 | 1.408024 | 1.157992 | 0.246867 | 0.684128 |
| CG15432 | 418.5515 | 1.961217 | 1.692958 | 1.158455 | 0.246678 | 0.684128 |
| edl | 211.7934 | -2.2704 | 1.963728 | -1.15617 | 0.247613 | 0.685655 |
| CG34367 | 1660.633 | 0.920935 | 0.79662 | 1.156053 | 0.247659 | 0.685655 |

| | | | | | | |
|---|---|---|---|---|---|---|
| RNaseMRF | 198.6895 | -2.34339 | 2.02767 | -1.15571 | 0.247801 | 0.685812 |
| CG6465 | 3215.222 | 0.77196 | 0.668094 | 1.155467 | 0.247899 | 0.685846 |
| CG5273 | 18752.81 | -0.49511 | 0.42943 | -1.15295 | 0.248932 | 0.686853 |
| tub | 1199.08 | 1.241983 | 1.076183 | 1.154062 | 0.248475 | 0.686853 |
| PGAP5 | 770.1189 | -1.62906 | 1.416782 | -1.14983 | 0.250215 | 0.686853 |
| CG9328 | 13281.51 | 0.521009 | 0.451723 | 1.153382 | 0.248754 | 0.686853 |
| alpha-Est9 | 2882.626 | 0.972659 | 0.845963 | 1.149765 | 0.250241 | 0.686853 |
| CG14882 | 289.5724 | 2.936626 | 2.552902 | 1.150309 | 0.250017 | 0.686853 |
| CG14879 | 1491.61 | -1.94845 | 1.694132 | -1.15012 | 0.250095 | 0.686853 |
| Cbp20 | 2118.721 | -1.00139 | 0.870299 | -1.15062 | 0.249887 | 0.686853 |
| CG6142 | 103.0493 | -3.27093 | 2.844378 | -1.14996 | 0.250159 | 0.686853 |
| CG30156 | 75.46855 | 4.083458 | 3.54088 | 1.153233 | 0.248815 | 0.686853 |
| eIF3j | 1345.312 | 1.147193 | 0.997232 | 1.150377 | 0.249989 | 0.686853 |
| Pka-R2 | 8949.556 | -0.53769 | 0.467592 | -1.14992 | 0.250178 | 0.686853 |
| 4E-T | 1567.028 | -1.02712 | 0.893479 | -1.14958 | 0.250317 | 0.686853 |
| spn-F | 393.0129 | -1.98455 | 1.720413 | -1.15353 | 0.248693 | 0.686853 |
| poly | 2065.532 | -1.08821 | 0.943568 | -1.15329 | 0.24879 | 0.686853 |
| CG42336 | 1578.921 | -0.95671 | 0.830621 | -1.1518 | 0.249402 | 0.686853 |
| Nuak | 3448.095 | 0.716608 | 0.622363 | 1.151431 | 0.249555 | 0.686853 |
| CG3065 | 1741.014 | -0.99886 | 0.867325 | -1.15166 | 0.249461 | 0.686853 |
| Vti1b | 1866.032 | -1.1119 | 0.964152 | -1.15324 | 0.24881 | 0.686853 |
| VAChT | 19502.8 | 0.481758 | 0.418278 | 1.151764 | 0.249418 | 0.686853 |
| Mcm3 | 117.7986 | -3.70575 | 3.22285 | -1.14984 | 0.250212 | 0.686853 |
| Elys | 1692.598 | -1.6224 | 1.411227 | -1.14964 | 0.250293 | 0.686853 |
| eIF3e | 2881.713 | -0.83363 | 0.723359 | -1.15245 | 0.249137 | 0.686853 |
| CG1722 | 646.6927 | -1.69268 | 1.469864 | -1.15159 | 0.249491 | 0.686853 |
| CG7785 | 475.1817 | 1.775933 | 1.545153 | 1.149357 | 0.250409 | 0.686869 |
| spir | 16709.17 | 0.602491 | 0.524853 | 1.147923 | 0.251 | 0.688257 |
| Nlp | 1880.364 | 1.070083 | 0.933012 | 1.146913 | 0.251418 | 0.68893 |
| CG14431 | 866.4342 | 1.7052 | 1.486671 | 1.146992 | 0.251385 | 0.68893 |
| Slu7 | 856.2767 | -1.18532 | 1.033906 | -1.14645 | 0.251608 | 0.689216 |
| CtIP | 141.9802 | -3.05201 | 2.665053 | -1.1452 | 0.252127 | 0.689656 |
| CG11147 | 387.1078 | 2.781348 | 2.427848 | 1.145602 | 0.25196 | 0.689656 |
| CG31195 | 239.4712 | -2.70635 | 2.363572 | -1.14503 | 0.252198 | 0.689656 |
| Fs(2)Ket | 8243.913 | 0.580686 | 0.507071 | 1.145176 | 0.252136 | 0.689656 |
| asRNA:CR4 | 671.1373 | -1.30551 | 1.139686 | -1.1455 | 0.252001 | 0.689656 |
| CG7148 | 1191.14 | -1.51606 | 1.325121 | -1.1441 | 0.252584 | 0.690146 |
| mt:CoIII | 646076.6 | 0.418473 | 0.365768 | 1.144093 | 0.252585 | 0.690146 |
| Xrp1 | 11433.55 | 0.607019 | 0.53072 | 1.143765 | 0.252721 | 0.690146 |
| Nup93-1 | 671.7353 | -1.70975 | 1.494626 | -1.14393 | 0.252652 | 0.690146 |
| Tmtc3 | 332.6986 | -2.68818 | 2.350767 | -1.14353 | 0.252817 | 0.690173 |
| CG31549 | 2772.181 | 0.815975 | 0.713794 | 1.143151 | 0.252976 | 0.690372 |
| CG32486 | 12301.36 | 0.549671 | 0.480938 | 1.142916 | 0.253073 | 0.690404 |
| puml | 2783.077 | 0.815048 | 0.714296 | 1.14105 | 0.253849 | 0.690867 |
| CG33523 | 2546.508 | -0.81995 | 0.718269 | -1.14157 | 0.253635 | 0.690867 |
| Syt7 | 15630.06 | 0.486431 | 0.426376 | 1.140851 | 0.253932 | 0.690867 |
| Orct2 | 807.2702 | -1.2147 | 1.063674 | -1.14199 | 0.253458 | 0.690867 |
| lncRNA:CR | 188.1428 | -3.24954 | 2.847357 | -1.14125 | 0.253767 | 0.690867 |
| Sc2 | 4334.589 | 0.688566 | 0.60311 | 1.141692 | 0.253582 | 0.690867 |
| MSBP | 6399.218 | -0.64924 | 0.568691 | -1.14164 | 0.253603 | 0.690867 |

| | | | | | | |
|---|---|---|---|---|---|---|
| arx | 302.4229 | -3.30354 | 2.894264 | -1.14141 | 0.253699 | 0.690867 |
| MtnA | 39144.81 | -0.52259 | 0.458747 | -1.13917 | 0.254633 | 0.691483 |
| imd | 64.6253 | 4.245331 | 3.726343 | 1.139275 | 0.254588 | 0.691483 |
| CG30325 | 193.8318 | -3.80441 | 3.337244 | -1.13998 | 0.254293 | 0.691483 |
| CG6984 | 157.8137 | 3.061553 | 2.687773 | 1.139067 | 0.254675 | 0.691483 |
| lncRNA:CR | 307.6499 | 1.814916 | 1.59278 | 1.139464 | 0.25451 | 0.691483 |
| CG9368 | 4564.955 | -0.78552 | 0.689479 | -1.1393 | 0.254578 | 0.691483 |
| lncRNA:CR | 430.2245 | 2.524186 | 2.216725 | 1.1387 | 0.254828 | 0.691664 |
| CG16989 | 789.0155 | 1.867367 | 1.642302 | 1.137043 | 0.25552 | 0.69284 |
| Sec24CD | 2060.242 | -0.84842 | 0.745928 | -1.1374 | 0.25537 | 0.69284 |
| Flacc | 521.3711 | 2.039837 | 1.793931 | 1.137077 | 0.255506 | 0.69284 |
| mora | 96.80636 | 3.803905 | 3.347276 | 1.136418 | 0.255782 | 0.693314 |
| Sdr | 181.5831 | 2.960967 | 2.607242 | 1.13567 | 0.256095 | 0.69346 |
| Trc8 | 990.1174 | -1.68238 | 1.481248 | -1.13579 | 0.256045 | 0.69346 |
| Vti1a | 638.267 | -1.84703 | 1.626196 | -1.1358 | 0.25604 | 0.69346 |
| sav | 159.4807 | -3.56085 | 3.139077 | -1.13436 | 0.256643 | 0.694009 |
| SelG | 1120.471 | 1.289125 | 1.136406 | 1.134388 | 0.256632 | 0.694009 |
| Prosbeta2 | 1193.36 | -1.50574 | 1.327376 | -1.13438 | 0.256637 | 0.694009 |
| Mpcp2 | 51994.89 | 0.4945 | 0.435782 | 1.13474 | 0.256484 | 0.694009 |
| Irk3 | 4942.495 | -0.76227 | 0.672599 | -1.13332 | 0.257079 | 0.694954 |
| Ak6 | 163.7627 | 3.613072 | 3.190215 | 1.132548 | 0.257404 | 0.695598 |
| CG32756 | 793.2285 | 1.429161 | 1.262543 | 1.13197 | 0.257647 | 0.696019 |
| CG13896 | 83.43297 | -4.38305 | 3.875512 | -1.13096 | 0.258071 | 0.696932 |
| cDIP | 75.37623 | -4.38142 | 3.875742 | -1.13047 | 0.258278 | 0.697019 |
| CG5323 | 760.2928 | 1.54765 | 1.368798 | 1.130664 | 0.258197 | 0.697019 |
| CG32192 | 218.8956 | -2.80574 | 2.483123 | -1.12992 | 0.258509 | 0.697409 |
| CG5445 | 2067.468 | -0.95692 | 0.847049 | -1.12971 | 0.2586 | 0.697421 |
| CG43740 | 970.1144 | 1.128616 | 0.999361 | 1.129337 | 0.258756 | 0.697606 |
| CG3815 | 294.2454 | 2.985739 | 2.645961 | 1.128414 | 0.259145 | 0.698355 |
| Agpat2 | 1182.727 | -1.28378 | 1.137904 | -1.1282 | 0.259237 | 0.698355 |
| ATP8B | 5468.191 | -0.62983 | 0.558626 | -1.12745 | 0.25955 | 0.698355 |
| Set | 1467.85 | -1.33417 | 1.182907 | -1.12787 | 0.259374 | 0.698355 |
| EcR | 7325.367 | -0.58804 | 0.521529 | -1.12754 | 0.259514 | 0.698355 |
| lncRNA:CR | 1329.523 | 0.977835 | 0.867345 | 1.127388 | 0.259578 | 0.698355 |
| CG13001 | 155.8517 | 3.527006 | 3.128897 | 1.127236 | 0.259643 | 0.698355 |
| CG17036 | 104.3377 | -3.86581 | 3.438054 | -1.12442 | 0.260836 | 0.699801 |
| Hs2st | 1326.767 | 0.995149 | 0.885275 | 1.124113 | 0.260965 | 0.699801 |
| Drice | 1124.27 | 2.091549 | 1.860586 | 1.124134 | 0.260956 | 0.699801 |
| sip3 | 4161.995 | 0.725698 | 0.645121 | 1.124903 | 0.26063 | 0.699801 |
| CG4853 | 1085.307 | -1.54922 | 1.377827 | -1.12439 | 0.260848 | 0.699801 |
| CG4593 | 197.7391 | 3.090226 | 2.748175 | 1.124465 | 0.260816 | 0.699801 |
| CG6758 | 513.6772 | -1.97054 | 1.750872 | -1.12546 | 0.260394 | 0.699801 |
| GstE13 | 1454.741 | -1.31851 | 1.17203 | -1.12498 | 0.260596 | 0.699801 |
| mRpL49 | 1633.552 | 0.964927 | 0.857634 | 1.125104 | 0.260545 | 0.699801 |
| CG3803 | 274.5546 | 1.83748 | 1.635864 | 1.123248 | 0.261332 | 0.700552 |
| RpL22 | 5021.203 | -0.60882 | 0.542451 | -1.12235 | 0.261712 | 0.70127 |
| f-cup | 1445.689 | -1.36471 | 1.216095 | -1.12221 | 0.261775 | 0.70127 |
| CG8963 | 1248.168 | 1.058737 | 0.943785 | 1.121799 | 0.261948 | 0.701465 |
| Slob | 7681.319 | 0.613369 | 0.546858 | 1.121624 | 0.262022 | 0.701465 |
| Scgbeta | 3930.308 | 0.729068 | 0.650567 | 1.120665 | 0.262431 | 0.702195 |

| | | | | | | |
|---|---|---|---|---|---|---|
| beat-VI | 1480.323 | -1.50573 | 1.344208 | -1.12016 | 0.262645 | 0.702195 |
| CstF50 | 462.2191 | -1.9585 | 1.748221 | -1.12028 | 0.262595 | 0.702195 |
| CG4598 | 710.8043 | 1.286741 | 1.148495 | 1.120371 | 0.262556 | 0.702195 |
| Cyp4p1 | 281.0208 | 2.648303 | 2.365478 | 1.119564 | 0.2629 | 0.70248 |
| GluRIB | 3448.221 | 0.725745 | 0.648276 | 1.119501 | 0.262927 | 0.70248 |
| hyx | 1171.649 | 1.669129 | 1.492229 | 1.118548 | 0.263333 | 0.703332 |
| PR-Set7 | 198.6371 | 2.433533 | 2.176406 | 1.118143 | 0.263506 | 0.703559 |
| CG31368 | 586.2447 | 1.886614 | 1.688599 | 1.117266 | 0.263881 | 0.703871 |
| Wdr37 | 1194.745 | -1.17169 | 1.048421 | -1.11758 | 0.263747 | 0.703871 |
| mGluR | 2344.979 | -0.81057 | 0.7255 | -1.11725 | 0.263886 | 0.703871 |
| Eb1 | 6655.501 | -0.6165 | 0.552165 | -1.11651 | 0.264204 | 0.704484 |
| msl-2 | 967.5545 | 1.623915 | 1.454826 | 1.116226 | 0.264325 | 0.704575 |
| D12 | 188.4813 | -3.6731 | 3.298392 | -1.1136 | 0.26545 | 0.704701 |
| ClpP | 2268.826 | -0.88827 | 0.797023 | -1.11448 | 0.265072 | 0.704701 |
| ScpX | 1078.024 | -1.27756 | 1.146871 | -1.11396 | 0.265298 | 0.704701 |
| CG1234 | 396.8451 | 2.173032 | 1.950749 | 1.113948 | 0.265302 | 0.704701 |
| RhoGAP93 | 5442.03 | 0.784984 | 0.70369 | 1.115526 | 0.264625 | 0.704701 |
| Bili | 1677.832 | 0.899775 | 0.806969 | 1.115005 | 0.264848 | 0.704701 |
| bigmax | 1198.038 | 1.115931 | 1.002226 | 1.113453 | 0.265514 | 0.704701 |
| eEF1alpha | 17902.9 | 0.556149 | 0.498998 | 1.114533 | 0.265051 | 0.704701 |
| Grip128 | 80.13126 | 3.975933 | 3.564151 | 1.115534 | 0.264621 | 0.704701 |
| mgr | 1384.304 | -0.96397 | 0.865294 | -1.11403 | 0.265264 | 0.704701 |
| AstA-R1 | 1219.926 | -1.42223 | 1.275467 | -1.11507 | 0.264822 | 0.704701 |
| lncRNA:CR | 1521.958 | 1.00277 | 0.899114 | 1.115287 | 0.264728 | 0.704701 |
| CG4962 | 155.474 | -3.6254 | 3.255713 | -1.11355 | 0.265472 | 0.704701 |
| CG32544 | 1656.96 | -0.9135 | 0.820807 | -1.11293 | 0.265738 | 0.705063 |
| Prx3 | 2582.559 | 0.974911 | 0.876683 | 1.112045 | 0.266119 | 0.705839 |
| CG17294 | 1368.364 | -1.07488 | 0.9675 | -1.11098 | 0.266575 | 0.705883 |
| CG10444 | 157.28 | -2.92566 | 2.633136 | -1.11109 | 0.266528 | 0.705883 |
| Hacl | 493.9254 | 2.248623 | 2.023413 | 1.111302 | 0.266438 | 0.705883 |
| NitFhit | 2074.605 | 0.834307 | 0.750589 | 1.111535 | 0.266338 | 0.705883 |
| NO66 | 260.9554 | -2.50538 | 2.25421 | -1.11142 | 0.266386 | 0.705883 |
| mip120 | 386.9731 | -2.36905 | 2.133072 | -1.11063 | 0.266728 | 0.706031 |
| udd | 1072.975 | 1.043806 | 0.939989 | 1.110446 | 0.266807 | 0.706031 |
| lncRNA:CR | 250.1114 | -2.30895 | 2.080439 | -1.10984 | 0.26707 | 0.706494 |
| ATP7 | 561.4912 | -1.4973 | 1.349908 | -1.10919 | 0.26735 | 0.707003 |
| slo | 20086.97 | 0.46791 | 0.422077 | 1.10859 | 0.267607 | 0.707448 |
| CG17544 | 1569.455 | 1.041814 | 0.940093 | 1.108203 | 0.267774 | 0.707449 |
| l(3)05822 | 4657.461 | 0.831718 | 0.750525 | 1.108181 | 0.267784 | 0.707449 |
| CG3630 | 169.4214 | -3.12177 | 2.821197 | -1.10654 | 0.268493 | 0.707924 |
| CG5886 | 1030.859 | -1.67573 | 1.515341 | -1.10585 | 0.268793 | 0.707924 |
| CG15099 | 4837.017 | 0.780555 | 0.705559 | 1.106293 | 0.2686 | 0.707924 |
| Drl-2 | 993.4645 | 1.273117 | 1.151047 | 1.106051 | 0.268704 | 0.707924 |
| Arl4 | 13662.08 | 0.534483 | 0.482978 | 1.106642 | 0.268449 | 0.707924 |
| PRAS40 | 2036.99 | 1.00315 | 0.9063 | 1.106863 | 0.268353 | 0.707924 |
| Pdcd4 | 5113.242 | 0.60161 | 0.543935 | 1.106034 | 0.268712 | 0.707924 |
| Idh3a | 5376.747 | -0.82774 | 0.747963 | -1.10666 | 0.268441 | 0.707924 |
| simj | 2601.226 | 0.74745 | 0.675981 | 1.105726 | 0.268845 | 0.707924 |
| Cdc27 | 762.2309 | 1.663808 | 1.502295 | 1.107511 | 0.268073 | 0.707924 |
| CG5404 | 6370.192 | 0.754131 | 0.682588 | 1.104812 | 0.269241 | 0.708269 |

| | | | | | | |
|---|---|---|---|---|---|---|
| Or43b | 234.8233 | 2.43051 | 2.199728 | 1.104914 | 0.269197 | 0.708269 |
| l(3)72Ab | 1327.119 | -0.95604 | 0.865076 | -1.10515 | 0.269096 | 0.708269 |
| Sf3b2 | 973.5914 | 1.520835 | 1.377201 | 1.104294 | 0.269466 | 0.708628 |
| tzn | 966.8604 | -1.11275 | 1.007896 | -1.10403 | 0.269578 | 0.708691 |
| GlyS | 6100.48 | -0.64631 | 0.585542 | -1.10378 | 0.26969 | 0.708753 |
| CG10628 | 574.2077 | 2.282717 | 2.068535 | 1.103542 | 0.269792 | 0.708788 |
| fray | 5846.944 | 0.623609 | 0.565538 | 1.102682 | 0.270165 | 0.709305 |
| pAbp | 17163.81 | -0.48644 | 0.441092 | -1.10281 | 0.270108 | 0.709305 |
| swm | 1255.28 | 1.226833 | 1.113277 | 1.102001 | 0.270461 | 0.70985 |
| pins | 1432.608 | 1.324216 | 1.20194 | 1.101732 | 0.270578 | 0.709925 |
| Gyg | 1865.013 | -0.84119 | 0.763897 | -1.10118 | 0.270817 | 0.710169 |
| Neurl4 | 685.2491 | -1.63395 | 1.483909 | -1.10111 | 0.270848 | 0.710169 |
| U2af38 | 976.4414 | 1.480858 | 1.346915 | 1.099445 | 0.271574 | 0.711013 |
| Dh44-R2 | 1270.833 | -1.03999 | 0.945626 | -1.09979 | 0.271425 | 0.711013 |
| ACOX1 | 918.2232 | 1.651811 | 1.502118 | 1.099654 | 0.271483 | 0.711013 |
| Torsin | 736.9198 | -1.86683 | 1.69753 | -1.09973 | 0.271449 | 0.711013 |
| sli | 4914.271 | -0.69567 | 0.632802 | -1.09936 | 0.271613 | 0.711013 |
| CG4266 | 1290.257 | -0.99843 | 0.909234 | -1.0981 | 0.27216 | 0.712212 |
| CG33469 | 91.60465 | -4.21538 | 3.8425 | -1.09704 | 0.272624 | 0.713194 |
| CG14478 | 6045.978 | -0.62128 | 0.566658 | -1.09639 | 0.272909 | 0.713371 |
| CG13325 | 233.0095 | -2.77658 | 2.532741 | -1.09628 | 0.272958 | 0.713371 |
| yps | 4925.613 | 0.624602 | 0.569586 | 1.096588 | 0.272822 | 0.713371 |
| CG46339 | 3317.155 | 0.750969 | 0.685193 | 1.095997 | 0.27308 | 0.713457 |
| Nhe1 | 1613.384 | -1.60256 | 1.464662 | -1.09415 | 0.273888 | 0.714333 |
| CG7166 | 5150.843 | 0.591815 | 0.541058 | 1.09381 | 0.274038 | 0.714333 |
| spz | 1812.45 | 1.781292 | 1.627811 | 1.094287 | 0.273829 | 0.714333 |
| Gcn2 | 1088.497 | -1.59036 | 1.453261 | -1.09434 | 0.273807 | 0.714333 |
| Orai | 1502.846 | 0.970544 | 0.887202 | 1.093938 | 0.273982 | 0.714333 |
| lark | 2719.985 | -1.02765 | 0.939479 | -1.09385 | 0.274022 | 0.714333 |
| mRpL34 | 898.1431 | 1.185503 | 1.083098 | 1.094548 | 0.273715 | 0.714333 |
| Usf | 2391.433 | 0.839029 | 0.767353 | 1.093407 | 0.274215 | 0.714562 |
| CG7789 | 861.8082 | 1.344443 | 1.229955 | 1.093083 | 0.274357 | 0.714566 |
| pigs | 4271.437 | -0.71469 | 0.653879 | -1.093 | 0.274395 | 0.714566 |
| Chsy | 1887.547 | 0.91983 | 0.84187 | 1.092603 | 0.274568 | 0.714785 |
| CG12947 | 383.3821 | -1.88953 | 1.73137 | -1.09135 | 0.275118 | 0.715617 |
| Cp190 | 2447.32 | 0.791586 | 0.725262 | 1.091449 | 0.275076 | 0.715617 |
| CG13689 | 375.7243 | 1.725561 | 1.581528 | 1.091072 | 0.275241 | 0.715617 |
| CG13539 | 278.179 | -2.77705 | 2.545263 | -1.09107 | 0.275244 | 0.715617 |
| ssx | 4404.549 | -0.62843 | 0.576615 | -1.08985 | 0.275778 | 0.716299 |
| ihog | 321.9294 | -2.99005 | 2.745873 | -1.08893 | 0.276187 | 0.716299 |
| CG10694 | 208.7764 | -3.22255 | 2.959594 | -1.08885 | 0.276221 | 0.716299 |
| CG10151 | 257.9799 | -2.30785 | 2.117745 | -1.08977 | 0.275815 | 0.716299 |
| SIFaR | 1115.648 | 1.143062 | 1.049599 | 1.089046 | 0.276133 | 0.716299 |
| RpL30 | 6722.976 | 0.659166 | 0.604886 | 1.089736 | 0.275829 | 0.716299 |
| Ca-beta | 4557.56 | -0.61664 | 0.566312 | -1.08887 | 0.276213 | 0.716299 |
| CG9018 | 786.0031 | 1.420765 | 1.304601 | 1.089042 | 0.276136 | 0.716299 |
| Dpm1 | 632.098 | 1.318237 | 1.211083 | 1.088478 | 0.276384 | 0.716366 |
| CG12343 | 1339.14 | -1.56866 | 1.441276 | -1.08839 | 0.276425 | 0.716366 |
| CG17896 | 4854.118 | -0.74275 | 0.682778 | -1.08784 | 0.276667 | 0.716762 |
| RhoGAP92 | 1270.613 | 1.600579 | 1.473851 | 1.085985 | 0.277486 | 0.717957 |

| | | | | | | |
|---|---|---|---|---|---|---|
| Golgin84 | 777.8005 | -1.61804 | 1.489682 | -1.08617 | 0.277405 | 0.717957 |
| ras | 3633.619 | -1.32772 | 1.222033 | -1.08648 | 0.277266 | 0.717957 |
| tfc | 163.8964 | -3.15083 | 2.900335 | -1.08637 | 0.277317 | 0.717957 |
| Bx | 9750.46 | 0.514408 | 0.473788 | 1.085735 | 0.277596 | 0.718011 |
| TBCB | 836.28 | -1.16441 | 1.072922 | -1.08527 | 0.277803 | 0.718313 |
| CR12842 | 802.773 | 1.604807 | 1.479282 | 1.084856 | 0.277986 | 0.718555 |
| CG4686 | 779.3398 | -1.30302 | 1.201366 | -1.08462 | 0.278092 | 0.718598 |
| CG2162 | 1078.4 | 1.142345 | 1.05387 | 1.083953 | 0.278386 | 0.719126 |
| RtGEF | 1817.223 | 0.839517 | 0.775023 | 1.083215 | 0.278713 | 0.719276 |
| Hel89B | 1308.824 | 0.93572 | 0.863717 | 1.083365 | 0.278647 | 0.719276 |
| l(2)k09848 | 632.0946 | 1.719376 | 1.587123 | 1.083329 | 0.278663 | 0.719276 |
| Cbp53E | 14674.93 | 0.523182 | 0.483252 | 1.082628 | 0.278974 | 0.719485 |
| Rac1 | 3144.932 | -0.73511 | 0.679129 | -1.08243 | 0.279063 | 0.719485 |
| Pino | 10863.5 | 0.502745 | 0.464336 | 1.082718 | 0.278933 | 0.719485 |
| CG9377 | 1901.395 | -0.88911 | 0.821776 | -1.08194 | 0.27928 | 0.719814 |
| Smg5 | 2614.35 | -0.7663 | 0.708528 | -1.08154 | 0.279457 | 0.720039 |
| CG31673 | 1434.034 | -1.07833 | 0.997738 | -1.08078 | 0.279795 | 0.720337 |
| snoRNA:2F | 140.0952 | 2.578249 | 2.385777 | 1.080675 | 0.279842 | 0.720337 |
| fln | 1448.676 | -1.06082 | 0.981382 | -1.08094 | 0.279722 | 0.720337 |
| RpS18 | 9912.037 | 0.512745 | 0.474577 | 1.080425 | 0.279953 | 0.720372 |
| Cyp6a14 | 163.9723 | 3.423724 | 3.170297 | 1.079938 | 0.28017 | 0.720372 |
| rho-5 | 2121.433 | -0.89264 | 0.826527 | -1.07999 | 0.280148 | 0.720372 |
| asRNA:CR4 | 2852.355 | 0.954737 | 0.884314 | 1.079636 | 0.280304 | 0.720372 |
| Prps | 4671.652 | -0.60302 | 0.558532 | -1.07966 | 0.280294 | 0.720372 |
| PGRP-LE | 623.4578 | -1.25505 | 1.163388 | -1.07879 | 0.280682 | 0.721113 |
| CG3308 | 1101.102 | -1.52319 | 1.412357 | -1.07847 | 0.280822 | 0.721241 |
| ArgRS-m | 199.0061 | -2.56262 | 2.377373 | -1.07792 | 0.281069 | 0.721415 |
| CG4080 | 2192.001 | 0.774984 | 0.718837 | 1.078109 | 0.280985 | 0.721415 |
| GluClalpha | 23228.57 | 0.474923 | 0.440749 | 1.077538 | 0.28124 | 0.721622 |
| CG12608 | 1275.797 | -1.55577 | 1.444941 | -1.0767 | 0.281615 | 0.721962 |
| mop | 3552.191 | -0.69047 | 0.64132 | -1.07664 | 0.281643 | 0.721962 |
| slgA | 15815.02 | 0.52772 | 0.490037 | 1.076897 | 0.281526 | 0.721962 |
| Atg16 | 605.5786 | -1.93308 | 1.795845 | -1.07642 | 0.281742 | 0.721985 |
| scaRNA:M | 189.3659 | -2.10019 | 1.952812 | -1.07547 | 0.282164 | 0.722837 |
| CG3420 | 2030.224 | 0.846317 | 0.787165 | 1.075146 | 0.282309 | 0.722864 |
| lncRNA:CR | 650.459 | -1.92875 | 1.794112 | -1.07504 | 0.282355 | 0.722864 |
| Act5C | 51633.31 | -0.42437 | 0.39495 | -1.07449 | 0.282605 | 0.723275 |
| Efa6 | 5079.927 | -0.58047 | 0.540358 | -1.07423 | 0.28272 | 0.723338 |
| CG1815 | 666.4409 | 1.376181 | 1.2822 | 1.073296 | 0.283138 | 0.723715 |
| shakB | 16954 | 0.587676 | 0.547416 | 1.073545 | 0.283026 | 0.723715 |
| CG11696 | 569.947 | -1.7441 | 1.624901 | -1.07335 | 0.283112 | 0.723715 |
| CG14312 | 846.7276 | 1.176243 | 1.096696 | 1.072533 | 0.28348 | 0.72411 |
| r | 168.1575 | -3.00665 | 2.803688 | -1.07239 | 0.283544 | 0.72411 |
| CG17598 | 662.8446 | 1.430529 | 1.334014 | 1.072349 | 0.283563 | 0.72411 |
| Ipk2 | 264.1488 | -2.28571 | 2.132366 | -1.07191 | 0.283758 | 0.724148 |
| Arf79F | 10763.87 | 0.483467 | 0.450987 | 1.072021 | 0.283711 | 0.724148 |
| CG1671 | 694.5633 | -1.59261 | 1.486345 | -1.07149 | 0.283947 | 0.724399 |
| GstS1 | 5219.636 | -0.64245 | 0.600094 | -1.07059 | 0.284355 | 0.725026 |
| enok | 1961.959 | -0.79285 | 0.740604 | -1.07055 | 0.284373 | 0.725026 |
| Esyt2 | 3257.601 | 0.740307 | 0.691707 | 1.070261 | 0.284502 | 0.725123 |

| | | | | | | |
|---|---|---|---|---|---|---|
| Nf-YB | 1729.036 | -0.89601 | 0.837904 | -1.06935 | 0.284913 | 0.72571 |
| CG9394 | 59192.99 | -0.54896 | 0.513319 | -1.06944 | 0.284871 | 0.72571 |
| Rcd-1 | 2467.723 | -0.77628 | 0.726072 | -1.06915 | 0.285003 | 0.72571 |
| CG7655 | 284.3944 | 2.060468 | 1.92798 | 1.068719 | 0.285196 | 0.725971 |
| Pi3K21B | 1514.511 | -1.06379 | 0.995937 | -1.06813 | 0.285462 | 0.726418 |
| CoRest | 4771.936 | -0.84471 | 0.791428 | -1.06733 | 0.285824 | 0.727106 |
| CG16890 | 296.3267 | -2.07148 | 1.941675 | -1.06685 | 0.286039 | 0.727423 |
| CG12320 | 160.5985 | 3.18552 | 2.988652 | 1.065872 | 0.286482 | 0.728088 |
| ClC-c | 2045.049 | 1.139768 | 1.069164 | 1.066036 | 0.286407 | 0.728088 |
| sif | 12702.29 | 0.467974 | 0.439221 | 1.065464 | 0.286666 | 0.728096 |
| CG5644 | 1135.773 | -1.39808 | 1.312118 | -1.06551 | 0.286645 | 0.728096 |
| RhoBTB | 3871.33 | 0.942234 | 0.88464 | 1.065105 | 0.286829 | 0.728255 |
| colt | 707.7339 | 1.343711 | 1.26179 | 1.064924 | 0.28691 | 0.728255 |
| ena | 4760.36 | -0.73632 | 0.691577 | -1.0647 | 0.287013 | 0.728286 |
| Hsepi | 600.9777 | -1.93244 | 1.816364 | -1.0639 | 0.287372 | 0.728641 |
| CycJ | 692.1397 | 1.30947 | 1.230758 | 1.063954 | 0.287349 | 0.728641 |
| Nopp140 | 659.4827 | 1.291357 | 1.21402 | 1.063703 | 0.287463 | 0.728641 |
| 312 | 776.2467 | 1.598682 | 1.503105 | 1.063586 | 0.287516 | 0.728641 |
| PlexB | 3773.142 | 0.75327 | 0.708665 | 1.062943 | 0.287808 | 0.729149 |
| bon | 1033.667 | -1.09996 | 1.035352 | -1.0624 | 0.288055 | 0.729545 |
| CG3939 | 751.6175 | -1.66636 | 1.569291 | -1.06186 | 0.288301 | 0.729857 |
| ATPsynG | 25773.17 | 0.447657 | 0.421711 | 1.061526 | 0.288451 | 0.729857 |
| CG1688 | 308.5203 | -1.94853 | 1.835323 | -1.06168 | 0.28838 | 0.729857 |
| CG8245 | 11053.05 | 0.535781 | 0.505238 | 1.060453 | 0.288939 | 0.730649 |
| CG1598 | 1913.308 | -0.99975 | 0.942773 | -1.06044 | 0.288946 | 0.730649 |
| fs(1)K10 | 79.58536 | 3.150344 | 2.975348 | 1.058815 | 0.289684 | 0.731182 |
| CG31908 | 438.1562 | 2.10211 | 1.983986 | 1.059539 | 0.289355 | 0.731182 |
| CG9386 | 199.6294 | -2.46981 | 2.33271 | -1.05877 | 0.289703 | 0.731182 |
| CG4049 | 3841.003 | -0.64087 | 0.605285 | -1.0588 | 0.289692 | 0.731182 |
| eIF6 | 623.3093 | 1.611278 | 1.521313 | 1.059137 | 0.289538 | 0.731182 |
| coro | 990.3169 | 1.454481 | 1.373344 | 1.05908 | 0.289563 | 0.731182 |
| Msr-110 | 48799.58 | -0.44654 | 0.422169 | -1.05773 | 0.290178 | 0.73215 |
| CG5516 | 434.0268 | 1.80535 | 1.707502 | 1.057305 | 0.290373 | 0.73217 |
| Rpn7 | 1892.644 | 0.794975 | 0.752113 | 1.056988 | 0.290517 | 0.73217 |
| SMC1 | 698.4319 | 1.667807 | 1.578295 | 1.056714 | 0.290642 | 0.73217 |
| Tdc2 | 1302.539 | -1.27396 | 1.205198 | -1.05706 | 0.290486 | 0.73217 |
| Smn | 592.3496 | -2.27373 | 2.151481 | -1.05682 | 0.290594 | 0.73217 |
| CG13516 | 269.9987 | -2.45635 | 2.327187 | -1.0555 | 0.291197 | 0.732741 |
| l(3)76BDm | 1433.961 | 1.055825 | 0.999986 | 1.05584 | 0.291041 | 0.732741 |
| lncRNA:CR | 1642.253 | 0.871737 | 0.82596 | 1.055422 | 0.291232 | 0.732741 |
| CG15628 | 5549.737 | -0.56204 | 0.532528 | -1.05542 | 0.291234 | 0.732741 |
| CG11377 | 1119.694 | 1.735627 | 1.644987 | 1.055101 | 0.291379 | 0.732877 |
| P32 | 3498.445 | 0.873194 | 0.828149 | 1.054392 | 0.291703 | 0.733462 |
| Cyp313a1 | 136.0311 | -3.5413 | 3.361303 | -1.05355 | 0.292089 | 0.733973 |
| CG17486 | 2121.32 | -1.33797 | 1.269757 | -1.05372 | 0.29201 | 0.733973 |
| Dscam3 | 2582.033 | 0.717874 | 0.681633 | 1.053168 | 0.292264 | 0.734182 |
| mAChR-A | 7416.16 | 0.533891 | 0.507473 | 1.052059 | 0.292773 | 0.735229 |
| CG9003 | 7000.316 | 0.650099 | 0.618611 | 1.050901 | 0.293304 | 0.735956 |
| sog | 201.4717 | 2.594135 | 2.467843 | 1.051175 | 0.293178 | 0.735956 |
| betaCOP | 1106.389 | -1.41663 | 1.34811 | -1.05083 | 0.293337 | 0.735956 |

| | | | | | | |
|---|---|---|---|---|---|---|
| CG11788 | 272.6797 | -2.80431 | 2.670499 | -1.05011 | 0.293669 | 0.736559 |
| Pex11 | 730.0974 | 1.450944 | 1.382882 | 1.049217 | 0.294078 | 0.737355 |
| CG10425 | 81.69785 | 3.139596 | 2.994154 | 1.048575 | 0.294374 | 0.737634 |
| Tis11 | 937.6043 | 1.717918 | 1.638211 | 1.048655 | 0.294337 | 0.737634 |
| CG31705 | 26501.34 | -0.5234 | 0.499436 | -1.04798 | 0.294646 | 0.737763 |
| mats | 771.1803 | -1.21861 | 1.163544 | -1.04732 | 0.29495 | 0.737763 |
| wde | 3947.246 | -0.72469 | 0.691844 | -1.04748 | 0.294877 | 0.737763 |
| Iml1 | 1488.581 | 1.01647 | 0.970593 | 1.047267 | 0.294977 | 0.737763 |
| CG8993 | 2800.015 | -0.69518 | 0.663618 | -1.04755 | 0.294844 | 0.737763 |
| CG4829 | 10586.97 | -0.60353 | 0.576117 | -1.04758 | 0.29483 | 0.737763 |
| trr | 190.3551 | 2.812927 | 2.687485 | 1.046676 | 0.295249 | 0.737964 |
| CG6700 | 6277.862 | 0.614922 | 0.587493 | 1.046689 | 0.295243 | 0.737964 |
| Atg2 | 560.1758 | -1.57224 | 1.502389 | -1.04649 | 0.295333 | 0.737964 |
| CG3556 | 11642.91 | 0.590554 | 0.564682 | 1.045816 | 0.295646 | 0.738517 |
| drpr | 15961.65 | 0.47681 | 0.456753 | 1.043911 | 0.296526 | 0.740486 |
| Pgm2a | 708.7552 | -1.3843 | 1.327165 | -1.04305 | 0.296926 | 0.741252 |
| CG4615 | 2381.917 | -0.79471 | 0.762816 | -1.04181 | 0.297498 | 0.74245 |
| bchs | 2643.594 | 0.781157 | 0.750401 | 1.040986 | 0.297882 | 0.742839 |
| CG42239 | 698.9471 | 1.641722 | 1.577246 | 1.040879 | 0.297932 | 0.742839 |
| lncRNA:CR | 179.8145 | 2.157371 | 2.072447 | 1.040978 | 0.297886 | 0.742839 |
| E(Pc) | 4402.451 | 0.645046 | 0.620011 | 1.040378 | 0.298164 | 0.743189 |
| vig2 | 1928.827 | -0.80044 | 0.770847 | -1.03839 | 0.29909 | 0.744958 |
| CG11200 | 225.9672 | -2.59554 | 2.499759 | -1.03832 | 0.299122 | 0.744958 |
| zfh2 | 50246 | -0.44839 | 0.43187 | -1.03825 | 0.299153 | 0.744958 |
| CG31637 | 3818.172 | 0.66876 | 0.644535 | 1.037585 | 0.299463 | 0.7455 |
| CG3587 | 271.2875 | -2.77154 | 2.672309 | -1.03713 | 0.299674 | 0.745794 |
| Cse1 | 541.8097 | 1.671657 | 1.612298 | 1.036816 | 0.299821 | 0.745801 |
| ort | 200.8197 | -2.65418 | 2.560151 | -1.03673 | 0.299863 | 0.745801 |
| Ktl | 9573.36 | 0.619362 | 0.597951 | 1.035808 | 0.300292 | 0.746387 |
| Aladin | 307.7817 | -2.14757 | 2.073639 | -1.03565 | 0.300365 | 0.746387 |
| CG11073 | 215.184 | 2.405067 | 2.322337 | 1.035624 | 0.300378 | 0.746387 |
| CG31638 | 2259.446 | -1.03435 | 0.999827 | -1.03453 | 0.300891 | 0.747431 |
| CG31910 | 95.25486 | 4.002026 | 3.875159 | 1.032739 | 0.301726 | 0.747834 |
| tun | 2618.248 | 0.735607 | 0.712531 | 1.032386 | 0.301891 | 0.747834 |
| CG42394 | 2642.43 | 0.742851 | 0.71886 | 1.033374 | 0.301429 | 0.747834 |
| Cap-G | 503.8116 | -2.01067 | 1.945098 | -1.03371 | 0.301272 | 0.747834 |
| cdm | 1581.552 | 1.035663 | 1.003144 | 1.032417 | 0.301877 | 0.747834 |
| CG43788 | 409.7333 | -1.9854 | 1.921866 | -1.03306 | 0.301575 | 0.747834 |
| CG1572 | 748.7853 | -2.16843 | 2.099372 | -1.0329 | 0.301652 | 0.747834 |
| GstT4 | 770.8282 | 1.172374 | 1.134982 | 1.032945 | 0.301629 | 0.747834 |
| CG7889 | 215.6947 | -2.77988 | 2.689855 | -1.03347 | 0.301385 | 0.747834 |
| lncRNA:CR | 4413.864 | -0.5875 | 0.569459 | -1.03168 | 0.302223 | 0.748423 |
| Rabex-5 | 203.3906 | 2.40469 | 2.33683 | 1.029039 | 0.303461 | 0.751258 |
| CG30285 | 4897.681 | 0.652701 | 0.635496 | 1.027073 | 0.304386 | 0.753316 |
| Lrr47 | 819.5136 | -1.36682 | 1.332663 | -1.02563 | 0.305065 | 0.754762 |
| Efr | 2986.325 | -0.92883 | 0.90603 | -1.02516 | 0.305287 | 0.754865 |
| CG7222 | 2388.282 | 0.723258 | 0.705517 | 1.025146 | 0.305294 | 0.754865 |
| jtb | 300.4307 | -2.63185 | 2.568261 | -1.02476 | 0.305476 | 0.755083 |
| Bre1 | 1383.993 | -1.06728 | 1.041959 | -1.0243 | 0.305693 | 0.755154 |
| CG15201 | 8879.216 | 0.601343 | 0.58702 | 1.024399 | 0.305647 | 0.755154 |

| | | | | | | |
|---|---|---|---|---|---|---|
| Gp93 | 2426.25 | 0.751716 | 0.734045 | 1.024074 | 0.3058 | 0.755186 |
| lncRNA:CR | 664.1754 | 1.568512 | 1.534148 | 1.0224 | 0.306592 | 0.756907 |
| kek6 | 1388.883 | 1.168485 | 1.143997 | 1.021405 | 0.307063 | 0.757603 |
| CG2116 | 269.9773 | 2.689626 | 2.632949 | 1.021526 | 0.307005 | 0.757603 |
| CG7702 | 177.9236 | -3.30506 | 3.240079 | -1.02005 | 0.307703 | 0.758555 |
| rho | 8973.304 | -0.55802 | 0.547076 | -1.02001 | 0.307726 | 0.758555 |
| CG7536 | 1302.475 | -1.09514 | 1.073669 | -1.01999 | 0.307732 | 0.758555 |
| CG43867 | 5977.408 | -0.55348 | 0.543559 | -1.01825 | 0.308559 | 0.760126 |
| lncRNA:CR | 1527.315 | -0.86257 | 0.846953 | -1.01844 | 0.308467 | 0.760126 |
| CG17528 | 9104.152 | 0.552521 | 0.542768 | 1.01797 | 0.308692 | 0.760222 |
| TTLL5 | 963.5054 | 1.292797 | 1.270452 | 1.017588 | 0.308874 | 0.760436 |
| CG13192 | 562.1521 | 1.873688 | 1.841961 | 1.017225 | 0.309047 | 0.760628 |
| B4 | 4280.315 | 0.673598 | 0.662385 | 1.016929 | 0.309187 | 0.760741 |
| CG10019 | 222.2548 | -1.97638 | 1.94476 | -1.01626 | 0.309507 | 0.761293 |
| Ufl1 | 3739.857 | -1.13471 | 1.117046 | -1.01582 | 0.309717 | 0.761579 |
| CycD | 632.001 | -2.01469 | 1.985764 | -1.01457 | 0.310311 | 0.762805 |
| CG2685 | 487.5156 | 1.958461 | 1.931269 | 1.01408 | 0.310545 | 0.763145 |
| mRpL43 | 1120.938 | 0.932709 | 0.920224 | 1.013567 | 0.310789 | 0.763512 |
| CG6154 | 795.7913 | -1.33012 | 1.313646 | -1.01254 | 0.311279 | 0.764073 |
| CG1523 | 337.0942 | 1.586737 | 1.567161 | 1.012492 | 0.311303 | 0.764073 |
| CG8323 | 1379.602 | -1.46182 | 1.4435 | -1.01269 | 0.311207 | 0.764073 |
| CG31388 | 1814.003 | -0.9285 | 0.917527 | -1.01195 | 0.31156 | 0.764469 |
| Nepl4 | 132.7151 | -3.03359 | 3.002371 | -1.0104 | 0.312305 | 0.765668 |
| RyR | 2173.731 | 0.856239 | 0.84727 | 1.010586 | 0.312215 | 0.765668 |
| SmB | 1248.628 | -0.88981 | 0.880709 | -1.01034 | 0.312335 | 0.765668 |
| dbr | 1021.124 | 1.144736 | 1.133267 | 1.01012 | 0.312438 | 0.765687 |
| CG8668 | 1418.402 | -1.09251 | 1.081855 | -1.00985 | 0.312567 | 0.76577 |
| CG3436 | 1144.518 | 1.060832 | 1.054212 | 1.00628 | 0.314281 | 0.766131 |
| CG11601 | 196.4442 | 2.536583 | 2.518831 | 1.007048 | 0.313912 | 0.766131 |
| CG13117 | 383.9642 | -2.69357 | 2.672736 | -1.0078 | 0.313552 | 0.766131 |
| CG6443 | 1050.021 | -1.05298 | 1.046264 | -1.00642 | 0.314212 | 0.766131 |
| CG5270 | 899.1843 | 1.101201 | 1.093627 | 1.006926 | 0.31397 | 0.766131 |
| CG14883 | 1055.96 | 1.319954 | 1.311892 | 1.006145 | 0.314346 | 0.766131 |
| mod | 1890.84 | 0.800251 | 0.79487 | 1.00677 | 0.314045 | 0.766131 |
| Pkn | 4866.679 | -0.63729 | 0.632045 | -1.00831 | 0.313308 | 0.766131 |
| CG14664 | 547.1656 | -2.70368 | 2.68474 | -1.00706 | 0.313908 | 0.766131 |
| Nost | 880.7021 | 1.037735 | 1.028709 | 1.008774 | 0.313083 | 0.766131 |
| CG42700 | 198.1141 | 2.957735 | 2.936167 | 1.007346 | 0.313769 | 0.766131 |
| fwd | 8026.003 | -0.6413 | 0.636745 | -1.00715 | 0.313864 | 0.766131 |
| CR43670 | 720.4428 | 1.394704 | 1.38281 | 1.008601 | 0.313166 | 0.766131 |
| EloC | 5192.527 | -0.569 | 0.564346 | -1.00824 | 0.313339 | 0.766131 |
| und | 1397.378 | 1.318375 | 1.310558 | 1.005965 | 0.314432 | 0.766131 |
| bol | 19976.67 | 0.438734 | 0.435944 | 1.006401 | 0.314223 | 0.766131 |
| CG9953 | 4279.662 | -0.79572 | 0.790769 | -1.00626 | 0.314292 | 0.766131 |
| lilli | 9136.666 | 0.488293 | 0.485194 | 1.006387 | 0.314229 | 0.766131 |
| CG15631 | 235.7641 | 2.39362 | 2.380608 | 1.005466 | 0.314672 | 0.766483 |
| Nrg | 4883.509 | 0.661767 | 0.658357 | 1.005179 | 0.31481 | 0.766586 |
| CG12121 | 1365.814 | -0.99846 | 0.994683 | -1.0038 | 0.315474 | 0.76797 |
| mib1 | 967.507 | -1.11005 | 1.106277 | -1.00341 | 0.315665 | 0.768201 |
| Alg1 | 504.9376 | -2.26636 | 2.260839 | -1.00244 | 0.316131 | 0.768791 |

| | | | | | | |
|---|---|---|---|---|---|---|
| CG31997 | 1333.734 | -1.1656 | 1.163021 | -1.00221 | 0.31624 | 0.768791 |
| Rab32 | 297.86 | -2.34538 | 2.340446 | -1.00211 | 0.316291 | 0.768791 |
| CG15601 | 417.7284 | 2.048268 | 2.043474 | 1.002346 | 0.316176 | 0.768791 |
| Sh | 59029.1 | 0.394309 | 0.394073 | 1.000599 | 0.317021 | 0.770131 |
| ArgRS | 444.8042 | 2.051703 | 2.050529 | 1.000572 | 0.317034 | 0.770131 |
| CG2841 | 774.9344 | 1.120787 | 1.120464 | 1.000288 | 0.317171 | 0.770231 |
| CG4660 | 883.5352 | 1.044745 | 1.044674 | 1.000067 | 0.317278 | 0.770258 |
| Jafrac1 | 9950.707 | -0.45825 | 0.458514 | -0.99943 | 0.317585 | 0.770303 |
| Nup35 | 273.4787 | -2.36104 | 2.362169 | -0.99952 | 0.317541 | 0.770303 |
| CG11279 | 1154.59 | 0.98853 | 0.988865 | 0.999661 | 0.317474 | 0.770303 |
| axo | 9942.72 | -0.48714 | 0.487621 | -0.999 | 0.317793 | 0.770553 |
| CG13585 | 4525.553 | 0.557164 | 0.55782 | 0.998825 | 0.31788 | 0.770553 |
| Tctp | 16491.52 | 0.483332 | 0.484035 | 0.998547 | 0.318014 | 0.770646 |
| CG31715 | 1929.817 | 0.75275 | 0.754098 | 0.998211 | 0.318177 | 0.770808 |
| CG6765 | 1403.608 | 1.016528 | 1.018717 | 0.997851 | 0.318352 | 0.770999 |
| CG14695 | 114.124 | 3.325855 | 3.335313 | 0.997164 | 0.318685 | 0.771151 |
| CG30496 | 553.3259 | 1.546773 | 1.551703 | 0.996823 | 0.318851 | 0.771151 |
| jar | 4080.747 | -0.64124 | 0.643321 | -0.99677 | 0.318876 | 0.771151 |
| RpS24 | 16551.02 | 0.482059 | 0.48364 | 0.996731 | 0.318895 | 0.771151 |
| NetB | 3239.737 | -0.65378 | 0.655824 | -0.99688 | 0.318823 | 0.771151 |
| lncRNA:CR | 461.6047 | 2.2927 | 2.300727 | 0.996511 | 0.319002 | 0.771178 |
| Ten-m | 21062.83 | 0.425777 | 0.427377 | 0.996255 | 0.319126 | 0.771246 |
| cmpy | 3339.862 | 0.785183 | 0.78889 | 0.995301 | 0.31959 | 0.77156 |
| CG6770 | 13660.55 | -0.42937 | 0.431524 | -0.995 | 0.319737 | 0.77156 |
| Patj | 1073.839 | -1.19193 | 1.197263 | -0.99554 | 0.319471 | 0.77156 |
| CG7275 | 569.252 | -1.45429 | 1.461369 | -0.99516 | 0.319659 | 0.77156 |
| CG32085 | 4017.657 | -0.75728 | 0.76096 | -0.99517 | 0.319654 | 0.77156 |
| CadN | 11878.78 | 0.442112 | 0.444824 | 0.993903 | 0.32027 | 0.772614 |
| bbx | 126.4531 | 2.499608 | 2.519625 | 0.992055 | 0.321171 | 0.774175 |
| Maf1 | 9214.972 | 0.522287 | 0.526614 | 0.991784 | 0.321303 | 0.774175 |
| CG8814 | 1829.838 | 1.233518 | 1.243132 | 0.992266 | 0.321068 | 0.774175 |
| CG15356 | 2224.198 | -0.74205 | 0.748181 | -0.9918 | 0.321293 | 0.774175 |
| CG42337 | 2619.748 | 0.755072 | 0.761592 | 0.991438 | 0.321472 | 0.774349 |
| HERC2 | 1862.131 | -1.00661 | 1.015513 | -0.99124 | 0.321571 | 0.774356 |
| CG11398 | 420.5757 | 1.991638 | 2.010456 | 0.99064 | 0.321861 | 0.774731 |
| CG7115 | 1030.699 | 1.277678 | 1.289905 | 0.990521 | 0.32192 | 0.774731 |
| CG13384 | 4309.683 | -0.60044 | 0.607349 | -0.98862 | 0.322851 | 0.774784 |
| Acer | 307.0826 | -1.96952 | 1.992918 | -0.98826 | 0.323027 | 0.774784 |
| CG4972 | 3438.071 | -0.658 | 0.666335 | -0.98749 | 0.323404 | 0.774784 |
| Coq3 | 104.519 | -3.10153 | 3.136369 | -0.98889 | 0.322717 | 0.774784 |
| sle | 537.5075 | 1.411299 | 1.427229 | 0.988839 | 0.322742 | 0.774784 |
| spab | 9430.801 | -0.52226 | 0.528614 | -0.98798 | 0.323161 | 0.774784 |
| Gat | 30049.02 | -0.49097 | 0.497474 | -0.98693 | 0.323679 | 0.774784 |
| Etl1 | 830.2061 | -1.51699 | 1.532412 | -0.98994 | 0.322205 | 0.774784 |
| CG40191 | 8118.725 | -0.73742 | 0.747 | -0.98717 | 0.32356 | 0.774784 |
| snoRNA:Ps | 89.91511 | 3.79743 | 3.845043 | 0.987617 | 0.32334 | 0.774784 |
| east | 3667.103 | -0.66903 | 0.676377 | -0.98913 | 0.322597 | 0.774784 |
| twr | 1121.147 | -0.99623 | 1.008437 | -0.9879 | 0.323202 | 0.774784 |
| Snap29 | 866.81 | -1.55587 | 1.576481 | -0.98693 | 0.323679 | 0.774784 |
| Alp2 | 277.8794 | 2.991595 | 3.028312 | 0.987876 | 0.323214 | 0.774784 |

| | | | | | | |
|---|---|---|---|---|---|---|
| hpRNA:CR: | 132.7505 | -2.39816 | 2.422188 | -0.99008 | 0.322135 | 0.774784 |
| Tango1 | 677.6138 | 1.495145 | 1.513957 | 0.987574 | 0.323361 | 0.774784 |
| CG32650 | 140.9033 | -3.68974 | 3.738253 | -0.98702 | 0.323632 | 0.774784 |
| CG12355 | 8809.196 | 0.502938 | 0.508635 | 0.988798 | 0.322762 | 0.774784 |
| l(2)01289 | 128.4773 | -2.44299 | 2.477181 | -0.9862 | 0.324036 | 0.774943 |
| sns | 719.2938 | -1.49641 | 1.517117 | -0.98635 | 0.323961 | 0.774943 |
| CG32264 | 14822.7 | 0.451902 | 0.458082 | 0.98651 | 0.323883 | 0.774943 |
| Sirt7 | 447.6771 | 1.978627 | 2.007124 | 0.985802 | 0.32423 | 0.775178 |
| mago | 923.6279 | 1.379238 | 1.399877 | 0.985257 | 0.324498 | 0.775587 |
| CG11970 | 594.1757 | -1.51167 | 1.536748 | -0.98368 | 0.325272 | 0.776543 |
| aPKC | 5814.562 | -0.56483 | 0.573939 | -0.98413 | 0.32505 | 0.776543 |
| RASSF8 | 2294.576 | 0.937549 | 0.953163 | 0.983618 | 0.325303 | 0.776543 |
| Su(P) | 1462.073 | 0.864892 | 0.878929 | 0.984029 | 0.325101 | 0.776543 |
| Prp40 | 438.121 | -1.5318 | 1.557564 | -0.98346 | 0.325382 | 0.776543 |
| lncRNA:CR | 50381.63 | 0.389583 | 0.396295 | 0.983063 | 0.325577 | 0.776777 |
| barc | 1306.891 | 0.913471 | 0.929734 | 0.982508 | 0.32585 | 0.776812 |
| CG8064 | 213.1209 | 2.56656 | 2.611879 | 0.982649 | 0.32578 | 0.776812 |
| CG8188 | 3924.605 | 0.830794 | 0.846002 | 0.982024 | 0.326088 | 0.776812 |
| AIMP3 | 1405.553 | 0.875547 | 0.891608 | 0.981986 | 0.326107 | 0.776812 |
| Pep | 5845.161 | 0.521669 | 0.53131 | 0.981854 | 0.326172 | 0.776812 |
| AIMP2 | 1528.752 | -1.19286 | 1.214272 | -0.98237 | 0.325919 | 0.776812 |
| Pfas | 144.0031 | -2.83581 | 2.896648 | -0.979 | 0.327581 | 0.778718 |
| Ip259 | 8044.782 | -0.5481 | 0.559945 | -0.97885 | 0.327655 | 0.778718 |
| RpL9 | 11757.99 | 0.464393 | 0.474589 | 0.978517 | 0.327819 | 0.778718 |
| Teh1 | 2927.355 | -0.67985 | 0.694274 | -0.97922 | 0.327471 | 0.778718 |
| Cyp9f2 | 6784.522 | -0.62595 | 0.639284 | -0.97914 | 0.327512 | 0.778718 |
| CG3107 | 6396.059 | 0.556193 | 0.568435 | 0.978463 | 0.327846 | 0.778718 |
| sPLA2 | 1427.616 | 0.936365 | 0.955778 | 0.979688 | 0.32724 | 0.778718 |
| PsGEF | 4857.382 | 0.589826 | 0.602439 | 0.979064 | 0.327548 | 0.778718 |
| CG44153 | 1321.616 | -0.92565 | 0.945948 | -0.97854 | 0.327807 | 0.778718 |
| Rnb | 313.0079 | -1.66568 | 1.703797 | -0.97763 | 0.328257 | 0.779444 |
| Fem-1 | 3358.855 | -0.63142 | 0.645986 | -0.97745 | 0.328346 | 0.779444 |
| per | 285.824 | 2.094103 | 2.157117 | 0.970788 | 0.331654 | 0.779521 |
| brn | 682.9514 | -1.70837 | 1.758883 | -0.97128 | 0.331407 | 0.779521 |
| Tbc1d8-9 | 2387.487 | 0.682799 | 0.70285 | 0.971472 | 0.331313 | 0.779521 |
| Srpk79D | 2955.467 | -0.68507 | 0.705725 | -0.97074 | 0.33168 | 0.779521 |
| CG12170 | 1020.915 | -0.96795 | 0.992433 | -0.97533 | 0.329399 | 0.779521 |
| CG8159 | 278.8926 | -1.87848 | 1.930481 | -0.97306 | 0.330522 | 0.779521 |
| alpha-Man | 2052.427 | 1.219574 | 1.255788 | 0.971163 | 0.331467 | 0.779521 |
| CG8031 | 1145.305 | -1.25786 | 1.29569 | -0.9708 | 0.331647 | 0.779521 |
| Usp8 | 1223.225 | -0.9634 | 0.991071 | -0.97208 | 0.33101 | 0.779521 |
| sba | 1767.138 | 0.861447 | 0.885768 | 0.972543 | 0.33078 | 0.779521 |
| twin | 1755.708 | 0.943708 | 0.965972 | 0.976952 | 0.328593 | 0.779521 |
| CG13436 | 76.3605 | -3.62774 | 3.73614 | -0.97099 | 0.331555 | 0.779521 |
| Lrt | 466.5154 | 1.345004 | 1.382163 | 0.973116 | 0.330496 | 0.779521 |
| Pcl | 2171.092 | 1.089045 | 1.117436 | 0.974593 | 0.329762 | 0.779521 |
| Bap111 | 483.439 | 1.779212 | 1.831456 | 0.971474 | 0.331312 | 0.779521 |
| Sema2a | 2733.451 | -0.73054 | 0.74849 | -0.97602 | 0.329053 | 0.779521 |
| bdg | 485.1129 | 1.741628 | 1.785461 | 0.97545 | 0.329337 | 0.779521 |
| CG8547 | 1994.898 | -1.48378 | 1.52783 | -0.97117 | 0.331466 | 0.779521 |

| | | | | | | |
|---|---|---|---|---|---|---|
| rho-7 | 471.2395 | 2.044224 | 2.104164 | 0.971513 | 0.331293 | 0.779521 |
| VGAT | 1884.953 | -0.98109 | 1.009345 | -0.972 | 0.331049 | 0.779521 |
| CG33643 | 130.1063 | 2.691455 | 2.758003 | 0.975871 | 0.329129 | 0.779521 |
| Mob3 | 2699.574 | -1.20512 | 1.239103 | -0.97258 | 0.330764 | 0.779521 |
| CG30172 | 8278.321 | 0.508866 | 0.523834 | 0.971425 | 0.331336 | 0.779521 |
| Mtch | 3915.145 | 0.738163 | 0.758381 | 0.973341 | 0.330384 | 0.779521 |
| fok | 16275.49 | -0.42723 | 0.438421 | -0.97448 | 0.329816 | 0.779521 |
| Snap24 | 774.2673 | 1.148601 | 1.183053 | 0.970878 | 0.331609 | 0.779521 |
| Gk2 | 316.041 | -2.44307 | 2.510278 | -0.97323 | 0.330441 | 0.779521 |
| ecd | 1090.773 | 1.389808 | 1.423814 | 0.976116 | 0.329007 | 0.779521 |
| CG9911 | 2043.264 | 0.955557 | 0.978092 | 0.97696 | 0.328589 | 0.779521 |
| CG6878 | 2613.079 | 0.662734 | 0.680696 | 0.973613 | 0.330249 | 0.779521 |
| defl | 355.9789 | 2.333383 | 2.401376 | 0.971686 | 0.331207 | 0.779521 |
| Pop4 | 552.7143 | -1.60532 | 1.652748 | -0.9713 | 0.331398 | 0.779521 |
| Mis12 | 673.997 | -1.70365 | 1.747957 | -0.97465 | 0.329732 | 0.779521 |
| CG12576 | 4292.543 | 0.574188 | 0.59091 | 0.971701 | 0.331199 | 0.779521 |
| cact | 2351.531 | 0.67174 | 0.692201 | 0.970441 | 0.331827 | 0.779637 |
| CG33947 | 396.5404 | 2.13115 | 2.197097 | 0.969985 | 0.332054 | 0.779715 |
| Rpn12 | 1524.612 | -0.86898 | 0.895734 | -0.97013 | 0.33198 | 0.779715 |
| chico | 1288.451 | 0.901051 | 0.929705 | 0.969179 | 0.332456 | 0.780199 |
| CG6204 | 249.2087 | 2.815727 | 2.907016 | 0.968597 | 0.332746 | 0.780199 |
| CG5504 | 868.4026 | 1.046122 | 1.079757 | 0.968849 | 0.332621 | 0.780199 |
| CG34457 | 193.3199 | -3.29586 | 3.402046 | -0.96879 | 0.332651 | 0.780199 |
| Gfrl | 8525.677 | -0.5324 | 0.549477 | -0.96892 | 0.332587 | 0.780199 |
| Dredd | 670.8331 | -1.55046 | 1.602612 | -0.96746 | 0.333316 | 0.781128 |
| Pi4KIIIalph | 2177.119 | -0.90288 | 0.933296 | -0.96741 | 0.333337 | 0.781128 |
| Nmdar2 | 3675.405 | 0.60761 | 0.628333 | 0.967019 | 0.333534 | 0.781362 |
| CG30069 | 5374.633 | -0.66554 | 0.688514 | -0.96663 | 0.333727 | 0.781586 |
| Uhg1 | 2473.373 | 0.715965 | 0.741226 | 0.965921 | 0.334084 | 0.781965 |
| Invadolysi | 13578.23 | -0.47094 | 0.487555 | -0.96592 | 0.334084 | 0.781965 |
| rux | 437.9626 | -1.72079 | 1.782866 | -0.96518 | 0.334454 | 0.782375 |
| l(2)09851 | 617.2142 | -1.17628 | 1.218524 | -0.96533 | 0.334379 | 0.782375 |
| CG14231 | 77.65593 | -3.21534 | 3.333794 | -0.96447 | 0.334812 | 0.782983 |
| TER94 | 2683.054 | 0.638256 | 0.661922 | 0.964247 | 0.334922 | 0.783014 |
| Rab18 | 1600.17 | -0.78393 | 0.814582 | -0.96238 | 0.335861 | 0.783295 |
| CG11247 | 3923.313 | -0.67849 | 0.704787 | -0.96269 | 0.335704 | 0.783295 |
| Xe7 | 7186.024 | -0.48453 | 0.504808 | -0.95983 | 0.337141 | 0.783295 |
| Cka | 3496.842 | -0.70343 | 0.729921 | -0.96371 | 0.335192 | 0.783295 |
| CG5676 | 8824.829 | 0.47732 | 0.49676 | 0.960866 | 0.336619 | 0.783295 |
| CG13625 | 152.121 | 2.137926 | 2.223751 | 0.961406 | 0.336348 | 0.783295 |
| CG14543 | 347.6376 | -1.6987 | 1.765405 | -0.96221 | 0.335942 | 0.783295 |
| bys | 648.469 | -1.43807 | 1.495048 | -0.96189 | 0.336106 | 0.783295 |
| c12.1 | 1720.189 | 0.874416 | 0.907657 | 0.963377 | 0.335358 | 0.783295 |
| CG11808 | 194.7298 | 2.524436 | 2.624834 | 0.961751 | 0.336175 | 0.783295 |
| dmpd | 641.866 | -1.17081 | 1.219465 | -0.9601 | 0.337005 | 0.783295 |
| CG1371 | 1939.154 | -0.89591 | 0.932897 | -0.96035 | 0.336879 | 0.783295 |
| LRP1 | 4106.536 | 0.617264 | 0.642875 | 0.960162 | 0.336974 | 0.783295 |
| CG10973 | 1038.759 | 1.270647 | 1.323361 | 0.960167 | 0.336971 | 0.783295 |
| dpr13 | 5405.654 | 0.554464 | 0.576173 | 0.962322 | 0.335888 | 0.783295 |
| sra | 2599.563 | 0.868595 | 0.901455 | 0.963548 | 0.335272 | 0.783295 |

| | | | | | | |
|---|---|---|---|---|---|---|
| CG3663 | 227.7863 | 2.198539 | 2.288417 | 0.960725 | 0.336691 | 0.783295 |
| Nurf-38 | 4470.685 | 0.605314 | 0.629819 | 0.961091 | 0.336507 | 0.783295 |
| Scsalpha1 | 10530.74 | 0.471963 | 0.491765 | 0.959733 | 0.33719 | 0.783295 |
| CG7630 | 28065.69 | 0.399592 | 0.415041 | 0.962778 | 0.335659 | 0.783295 |
| pbl | 1623.153 | 1.186005 | 1.234211 | 0.960941 | 0.336582 | 0.783295 |
| morgue | 921.2938 | -0.97696 | 1.017027 | -0.9606 | 0.336753 | 0.783295 |
| CG8270 | 748.6242 | -1.51481 | 1.580375 | -0.95851 | 0.337804 | 0.784495 |
| Jwa | 2588.54 | 0.712462 | 0.743819 | 0.957843 | 0.338142 | 0.784826 |
| Pvf3 | 7191.96 | 0.576125 | 0.601403 | 0.957968 | 0.338079 | 0.784826 |
| Marc | 1201.302 | 0.868849 | 0.907682 | 0.957218 | 0.338457 | 0.785281 |
| lncRNA:CR | 2859.39 | -0.67998 | 0.710486 | -0.95707 | 0.338534 | 0.785281 |
| CG13954 | 1674.842 | 0.906658 | 0.94774 | 0.956653 | 0.338742 | 0.785538 |
| moody | 9014.071 | 0.531349 | 0.555853 | 0.955916 | 0.339115 | 0.786145 |
| CG16974 | 16866.44 | -0.41001 | 0.429083 | -0.95555 | 0.339298 | 0.786145 |
| CG31473 | 126.5455 | 2.560164 | 2.679151 | 0.955588 | 0.33928 | 0.786145 |
| mof | 111.7639 | -3.15263 | 3.304727 | -0.95398 | 0.340095 | 0.787765 |
| pasi2 | 1461.806 | -1.56408 | 1.641063 | -0.95309 | 0.340546 | 0.78858 |
| pum | 35339.09 | -0.37825 | 0.397551 | -0.95145 | 0.341377 | 0.789163 |
| CG10898 | 447.0038 | 2.023388 | 2.124578 | 0.952371 | 0.340909 | 0.789163 |
| omd | 205.79 | -2.17333 | 2.28399 | -0.95155 | 0.341325 | 0.789163 |
| CG7956 | 2213.184 | 0.90621 | 0.952269 | 0.951633 | 0.341283 | 0.789163 |
| sinu | 743.2687 | 1.256775 | 1.320935 | 0.951428 | 0.341387 | 0.789163 |
| CG13921 | 1688.57 | -0.94782 | 0.995921 | -0.9517 | 0.341249 | 0.789163 |
| CG16721 | 1105.238 | -1.22753 | 1.292103 | -0.95003 | 0.342098 | 0.789198 |
| ZnT41F | 704.7742 | -1.73418 | 1.825545 | -0.94995 | 0.342137 | 0.789198 |
| CG32568 | 371.8308 | 2.250694 | 2.369569 | 0.949833 | 0.342197 | 0.789198 |
| Proc-R | 6965.35 | -0.49534 | 0.521201 | -0.95039 | 0.341913 | 0.789198 |
| inv | 65.53148 | 3.672518 | 3.865588 | 0.950054 | 0.342085 | 0.789198 |
| CG31221 | 27894.87 | 0.425216 | 0.447516 | 0.95017 | 0.342026 | 0.789198 |
| CG34136 | 296.1518 | -2.01781 | 2.123376 | -0.95028 | 0.341969 | 0.789198 |
| Su(Tpl) | 5896.179 | -0.49502 | 0.520467 | -0.95111 | 0.341548 | 0.789198 |
| Srp54k | 2570.91 | 0.915073 | 0.963585 | 0.949655 | 0.342287 | 0.789198 |
| Cyp9h1 | 337.8307 | -2.63063 | 2.772622 | -0.94879 | 0.342729 | 0.789389 |
| wash | 197.9598 | 2.233203 | 2.353815 | 0.948759 | 0.342743 | 0.789389 |
| Sox14 | 367.9939 | -1.94292 | 2.047942 | -0.94872 | 0.342764 | 0.789389 |
| yki | 2188.027 | -1.04446 | 1.100605 | -0.94899 | 0.342626 | 0.789389 |
| Src64B | 2413.692 | -0.71874 | 0.757899 | -0.94833 | 0.342963 | 0.789395 |
| CG14414 | 1177.395 | 0.935253 | 0.986051 | 0.948484 | 0.342883 | 0.789395 |
| Ada2b | 173.6092 | -2.56942 | 2.71041 | -0.94798 | 0.343138 | 0.789572 |
| ppk | 161.4217 | -3.06218 | 3.232133 | -0.94742 | 0.343426 | 0.789742 |
| CG6325 | 962.5465 | -1.13338 | 1.196681 | -0.9471 | 0.343587 | 0.789742 |
| mbo | 263.4695 | -2.38211 | 2.514618 | -0.94731 | 0.343483 | 0.789742 |
| CG17684 | 6017.777 | 0.531651 | 0.561367 | 0.947065 | 0.343606 | 0.789742 |
| Nplp1 | 3233.979 | -0.84326 | 0.890903 | -0.94652 | 0.343884 | 0.789929 |
| CG10984 | 1828.592 | 0.797869 | 0.842885 | 0.946593 | 0.343846 | 0.789929 |
| Nfs1 | 2929.359 | 0.761987 | 0.805671 | 0.94578 | 0.344261 | 0.790262 |
| CG2608 | 802.7585 | -1.40411 | 1.484796 | -0.94566 | 0.344324 | 0.790262 |
| Inx3 | 6084.472 | -0.51892 | 0.548522 | -0.94604 | 0.34413 | 0.790262 |
| Top3alpha | 191.9047 | -2.46362 | 2.605942 | -0.94539 | 0.344462 | 0.790353 |
| CG9471 | 2340.681 | -0.9456 | 1.001043 | -0.94461 | 0.344858 | 0.790808 |

| | | | | | | |
|---|---|---|---|---|---|---|
| side-VI | 4285.273 | 0.680478 | 0.720343 | 0.944658 | 0.344833 | 0.790808 |
| sstn | 569.1393 | 1.790459 | 1.895978 | 0.944346 | 0.344993 | 0.790892 |
| Cog2 | 719.9508 | 1.367396 | 1.448393 | 0.944078 | 0.34513 | 0.79098 |
| 2mit | 2503.635 | 0.73572 | 0.779608 | 0.943705 | 0.34532 | 0.791191 |
| CG10376 | 1450.677 | -0.89844 | 0.952466 | -0.94328 | 0.34554 | 0.791468 |
| RhoGAP1A | 2046.275 | -1.0059 | 1.066959 | -0.94277 | 0.345797 | 0.791605 |
| ALiX | 2080.48 | -0.96582 | 1.02429 | -0.94292 | 0.345724 | 0.791605 |
| bic | 8940.618 | 0.558705 | 0.592975 | 0.942208 | 0.346086 | 0.791621 |
| Drgx | 1534.343 | 0.949953 | 1.008249 | 0.942181 | 0.3461 | 0.791621 |
| CHMP2B | 438.2548 | -1.35289 | 1.43567 | -0.94234 | 0.34602 | 0.791621 |
| RpL18A | 22538.68 | 0.418381 | 0.44429 | 0.941684 | 0.346354 | 0.791752 |
| CG6707 | 3369.884 | -0.63927 | 0.678732 | -0.94186 | 0.346263 | 0.791752 |
| 26-29-p | 4978.001 | 0.658146 | 0.699352 | 0.94108 | 0.346664 | 0.791783 |
| CG12084 | 3991.605 | -0.63519 | 0.674896 | -0.94117 | 0.346616 | 0.791783 |
| sd | 4141.405 | 0.540568 | 0.574385 | 0.941126 | 0.34664 | 0.791783 |
| Khc-73 | 4839.424 | -0.53457 | 0.568389 | -0.94049 | 0.346964 | 0.791792 |
| lncRNA:CR | 108.0571 | 2.723258 | 2.895002 | 0.940676 | 0.346871 | 0.791792 |
| iPLA2-VIA | 1532.416 | 0.868402 | 0.923193 | 0.940651 | 0.346884 | 0.791792 |
| Ldsdh1 | 2547.727 | -0.67269 | 0.715629 | -0.94 | 0.347215 | 0.792141 |
| Dpck | 1756.01 | 0.779651 | 0.829955 | 0.939389 | 0.347531 | 0.792418 |
| Art2 | 122.0676 | -2.83029 | 3.012928 | -0.93938 | 0.347534 | 0.792418 |
| CG1139 | 792.7481 | 1.839664 | 1.958791 | 0.939183 | 0.347637 | 0.792426 |
| CG33143 | 1952.745 | 0.815665 | 0.868734 | 0.938913 | 0.347776 | 0.792518 |
| retm | 5557.421 | -0.51156 | 0.545343 | -0.93805 | 0.348219 | 0.793303 |
| mRpS16 | 1332.72 | 0.884325 | 0.943469 | 0.937312 | 0.348598 | 0.793941 |
| CG12746 | 1103.998 | -1.01074 | 1.079822 | -0.93603 | 0.349259 | 0.794462 |
| Sps2 | 626.296 | -1.5735 | 1.681526 | -0.93576 | 0.349397 | 0.794462 |
| CG5787 | 1857.49 | 1.220095 | 1.303562 | 0.93597 | 0.349289 | 0.794462 |
| CG5285 | 371.6998 | 1.501171 | 1.603003 | 0.936474 | 0.349029 | 0.794462 |
| asrij | 624.3389 | 1.540914 | 1.64673 | 0.935741 | 0.349406 | 0.794462 |
| hiw | 3005.133 | 0.768892 | 0.821718 | 0.935713 | 0.349421 | 0.794462 |
| CG4036 | 391.0741 | 1.456335 | 1.55837 | 0.934524 | 0.350033 | 0.795618 |
| bsk | 12411.32 | -0.43099 | 0.461343 | -0.93421 | 0.350198 | 0.795618 |
| CG5214 | 6675.002 | -0.6662 | 0.71316 | -0.93415 | 0.350226 | 0.795618 |
| CG13392 | 469.3546 | 1.852757 | 1.9861 | 0.932862 | 0.350891 | 0.795675 |
| CG6184 | 1368.178 | 1.095939 | 1.17428 | 0.933286 | 0.350672 | 0.795675 |
| Or65b | 116.7637 | -3.64976 | 3.910797 | -0.93325 | 0.35069 | 0.795675 |
| CG42523 | 115.6105 | -2.62688 | 2.813069 | -0.93381 | 0.3504 | 0.795675 |
| CG43255 | 110.5052 | -3.64897 | 3.910929 | -0.93302 | 0.35081 | 0.795675 |
| CG45071 | 764.3712 | 1.215213 | 1.302819 | 0.932756 | 0.350946 | 0.795675 |
| CG2772 | 177.4731 | -3.26777 | 3.50153 | -0.93324 | 0.350696 | 0.795675 |
| CG32638 | 830.592 | 1.210453 | 1.298227 | 0.932389 | 0.351136 | 0.795881 |
| Spn88Ea | 1822.515 | 0.830752 | 0.891565 | 0.931791 | 0.351445 | 0.796357 |
| CG9331 | 2900.637 | 0.705318 | 0.757735 | 0.930824 | 0.351945 | 0.796769 |
| Gr93a | 240.9474 | 2.945261 | 3.164594 | 0.930691 | 0.352013 | 0.796769 |
| Drep2 | 4726.728 | 0.593423 | 0.637629 | 0.930671 | 0.352024 | 0.796769 |
| ND-SGDH | 9580.064 | 0.46969 | 0.504446 | 0.9311 | 0.351802 | 0.796769 |
| CG8891 | 1481.387 | -0.90386 | 0.971586 | -0.93029 | 0.352218 | 0.796985 |
| Phf5a | 964.3823 | -1.01666 | 1.094206 | -0.92913 | 0.35282 | 0.797359 |
| nop5 | 284.5571 | 1.888537 | 2.032913 | 0.928981 | 0.352899 | 0.797359 |

| | | | | | | |
|---|---|---|---|---|---|---|
| Alg7 | 929.1504 | 1.095338 | 1.179516 | 0.928633 | 0.353079 | 0.797359 |
| nolo | 14314.29 | -0.54593 | 0.587517 | -0.92922 | 0.352773 | 0.797359 |
| HPS1 | 1727.751 | 0.936656 | 1.008444 | 0.928813 | 0.352986 | 0.797359 |
| Shawl | 915.7199 | -0.97345 | 1.047418 | -0.92938 | 0.352693 | 0.797359 |
| Bet1 | 812.514 | -0.97084 | 1.044154 | -0.92978 | 0.352484 | 0.797359 |
| Atxn7 | 725.8517 | -1.51331 | 1.632146 | -0.92719 | 0.353829 | 0.798827 |
| CG5104 | 185.8969 | 2.585644 | 2.791332 | 0.926312 | 0.354284 | 0.798973 |
| lic | 1002.604 | 1.052785 | 1.135959 | 0.926781 | 0.35404 | 0.798973 |
| Ran | 3083.083 | -0.62686 | 0.676506 | -0.92661 | 0.354128 | 0.798973 |
| Tollo | 490.592 | -1.47667 | 1.594164 | -0.9263 | 0.354292 | 0.798973 |
| CG7781 | 31595.28 | 0.367269 | 0.396743 | 0.925709 | 0.354597 | 0.799437 |
| sosie | 230.7544 | 2.649551 | 2.86814 | 0.923787 | 0.355597 | 0.800012 |
| CG5447 | 1711.709 | -0.7996 | 0.865053 | -0.92433 | 0.355312 | 0.800012 |
| IntS12 | 314.3313 | -1.94099 | 2.099304 | -0.92459 | 0.355182 | 0.800012 |
| Ero1L | 593.244 | 1.526319 | 1.651027 | 0.924466 | 0.355244 | 0.800012 |
| fwe | 11123.14 | 0.429077 | 0.464219 | 0.9243 | 0.35533 | 0.800012 |
| asRNA:CR4 | 131.5491 | 2.120164 | 2.295231 | 0.923726 | 0.355629 | 0.800012 |
| rut | 3778.987 | 0.658519 | 0.712903 | 0.923716 | 0.355634 | 0.800012 |
| PVRAP | 1379.392 | -0.783 | 0.847694 | -0.92369 | 0.35565 | 0.800012 |
| Tsc1 | 1122.108 | 0.872812 | 0.945625 | 0.923 | 0.356007 | 0.800593 |
| RhoGAP5A | 91.8586 | -3.05563 | 3.316958 | -0.92122 | 0.356938 | 0.80084 |
| Spx | 324.7356 | -1.6731 | 1.817115 | -0.92075 | 0.357182 | 0.80084 |
| CG7910 | 100.9259 | -3.48229 | 3.783857 | -0.9203 | 0.357415 | 0.80084 |
| hng2 | 198.142 | 2.820941 | 3.062897 | 0.921004 | 0.357048 | 0.80084 |
| JHDM2 | 480.5076 | -2.04507 | 2.222104 | -0.92033 | 0.357399 | 0.80084 |
| Fkbp39 | 658.0338 | -1.28553 | 1.394925 | -0.92157 | 0.356751 | 0.80084 |
| PIG-T | 2016.179 | 0.806418 | 0.875849 | 0.920728 | 0.357193 | 0.80084 |
| Ehbp1 | 5087.117 | -0.61427 | 0.667205 | -0.92065 | 0.357231 | 0.80084 |
| CG12128 | 793.0859 | 1.484475 | 1.610181 | 0.921931 | 0.356565 | 0.80084 |
| Muc14A | 766.3437 | -1.08538 | 1.178652 | -0.92086 | 0.357123 | 0.80084 |
| Spf45 | 1722.791 | -0.86456 | 0.939309 | -0.92042 | 0.357354 | 0.80084 |
| UQCR-11 | 14083.7 | 0.436089 | 0.473785 | 0.920437 | 0.357344 | 0.80084 |
| CG43219 | 293.0469 | 1.590424 | 1.725513 | 0.921711 | 0.356679 | 0.80084 |
| CG32444 | 134.3177 | -3.25011 | 3.532364 | -0.92009 | 0.357523 | 0.80086 |
| CG8641 | 980.1586 | 1.011314 | 1.099545 | 0.919757 | 0.3577 | 0.801032 |
| CG5862 | 605.9803 | -1.40407 | 1.529305 | -0.91811 | 0.358561 | 0.802683 |
| hep | 1738.067 | -0.8295 | 0.903635 | -0.91796 | 0.358637 | 0.802683 |
| SREBP | 4938.479 | 0.573511 | 0.625283 | 0.917202 | 0.359037 | 0.80313 |
| lncRNA:CR | 70.94135 | 3.562236 | 3.883429 | 0.917291 | 0.35899 | 0.80313 |
| sni | 1447.652 | -1.09323 | 1.192485 | -0.91677 | 0.359265 | 0.803193 |
| ND-B12 | 12510.54 | 0.441278 | 0.481334 | 0.91678 | 0.359258 | 0.803193 |
| pie | 378.5146 | 1.83972 | 2.008254 | 0.916079 | 0.359625 | 0.803657 |
| PhKgamma | 6641.777 | 0.55121 | 0.601765 | 0.915988 | 0.359673 | 0.803657 |
| lncRNA:CR | 243.9826 | -1.79177 | 1.956854 | -0.91564 | 0.359857 | 0.803843 |
| CG12253 | 1489.559 | -0.99831 | 1.090788 | -0.91522 | 0.360075 | 0.804101 |
| dod | 621.4571 | -1.20428 | 1.316371 | -0.91485 | 0.360272 | 0.804101 |
| NLaz | 1781.577 | -1.12814 | 1.232933 | -0.915 | 0.36019 | 0.804101 |
| ash2 | 1802.595 | -0.76396 | 0.835353 | -0.91454 | 0.360433 | 0.804235 |
| AANAT1 | 8855.868 | -0.52192 | 0.571122 | -0.91385 | 0.360795 | 0.804817 |
| CG7289 | 1335.63 | -1.08144 | 1.183629 | -0.91366 | 0.360894 | 0.804817 |

| | | | | | | |
|---|---|---|---|---|---|---|
| Nipsnap | 6317.706 | -0.51319 | 0.562263 | -0.91272 | 0.361388 | 0.805023 |
| l(2)k09913 | 12691.94 | -0.45464 | 0.497882 | -0.91315 | 0.361164 | 0.805023 |
| pcm | 692.109 | 1.574428 | 1.724976 | 0.912725 | 0.361387 | 0.805023 |
| Sras | 965.6881 | -1.34637 | 1.47505 | -0.91276 | 0.361368 | 0.805023 |
| HP1c | 496.0432 | 1.425551 | 1.56546 | 0.910627 | 0.362492 | 0.805917 |
| MED26 | 10945.6 | 0.423569 | 0.465026 | 0.910849 | 0.362375 | 0.805917 |
| Pgant2 | 1970.013 | 0.896665 | 0.983705 | 0.911519 | 0.362022 | 0.805917 |
| mol | 582.1971 | -1.25769 | 1.38057 | -0.91099 | 0.362299 | 0.805917 |
| MYPT-75D | 3729.366 | -0.59811 | 0.656789 | -0.91065 | 0.362479 | 0.805917 |
| CG18265 | 1035.932 | -1.23904 | 1.359879 | -0.91114 | 0.362221 | 0.805917 |
| Papst2 | 579.759 | 1.752359 | 1.923084 | 0.911224 | 0.362178 | 0.805917 |
| CG14411 | 1968.767 | -0.67369 | 0.740325 | -0.90999 | 0.362829 | 0.806393 |
| CG6512 | 2551.34 | 0.897889 | 0.986866 | 0.90984 | 0.362907 | 0.806393 |
| jef | 11124.53 | 0.431952 | 0.474897 | 0.90957 | 0.363049 | 0.806423 |
| lncRNA:CR | 144.0951 | -2.39846 | 2.637317 | -0.90943 | 0.363121 | 0.806423 |
| CG2611 | 216.9678 | -2.10001 | 2.311406 | -0.90854 | 0.363593 | 0.807159 |
| mTerf5 | 591.1345 | -1.72189 | 1.895706 | -0.90831 | 0.363715 | 0.807159 |
| CG15890 | 1950.788 | 1.101347 | 1.212624 | 0.908234 | 0.363755 | 0.807159 |
| CG9147 | 584.8346 | -1.35222 | 1.489381 | -0.90791 | 0.363925 | 0.807313 |
| Ranbp16 | 4720.72 | 0.524715 | 0.578057 | 0.907722 | 0.364025 | 0.807313 |
| CG8080 | 544.4963 | -1.71028 | 1.885893 | -0.90688 | 0.364469 | 0.808074 |
| CG15771 | 1573.699 | 0.816151 | 0.900588 | 0.906243 | 0.364807 | 0.808274 |
| DNaseII | 2794.499 | 0.965209 | 1.065186 | 0.906141 | 0.364861 | 0.808274 |
| CG15450 | 130.0104 | -3.53017 | 3.895143 | -0.9063 | 0.364777 | 0.808274 |
| Hsc70-3 | 9062.966 | -0.48126 | 0.531376 | -0.90569 | 0.365102 | 0.808361 |
| Galphai | 2221.447 | 0.67455 | 0.744692 | 0.905811 | 0.365036 | 0.808361 |
| CG31373 | 308.0624 | -2.11949 | 2.340985 | -0.90538 | 0.365262 | 0.808492 |
| CR18275 | 80.74544 | -3.52639 | 3.898595 | -0.90453 | 0.365715 | 0.808604 |
| TM9SF2 | 5331.776 | 0.623482 | 0.689087 | 0.904795 | 0.365574 | 0.808604 |
| snu | 82.06622 | -3.52653 | 3.898572 | -0.90457 | 0.365694 | 0.808604 |
| igl | 8192.356 | -0.53092 | 0.5869 | -0.90462 | 0.365666 | 0.808604 |
| side-VII | 3089.082 | -0.63126 | 0.698067 | -0.9043 | 0.365835 | 0.808645 |
| stwl | 133.9397 | -2.6829 | 2.967482 | -0.9041 | 0.365942 | 0.808659 |
| CG42240 | 1118.946 | -1.04193 | 1.153018 | -0.90366 | 0.366178 | 0.808958 |
| CG32452 | 291.0874 | -2.31278 | 2.561449 | -0.90292 | 0.366569 | 0.809473 |
| PEK | 1573.597 | 0.829743 | 0.919701 | 0.902188 | 0.366957 | 0.809473 |
| rau | 205.3676 | -2.59458 | 2.879022 | -0.9012 | 0.367481 | 0.809473 |
| LManII | 4104.455 | 0.619732 | 0.687644 | 0.90124 | 0.367461 | 0.809473 |
| CG5890 | 6734.473 | 0.691417 | 0.767603 | 0.900748 | 0.367722 | 0.809473 |
| Men-b | 14949.31 | -0.43694 | 0.484497 | -0.90184 | 0.367141 | 0.809473 |
| CG8360 | 1740.369 | 1.205887 | 1.338272 | 0.901077 | 0.367547 | 0.809473 |
| Adar | 10378.13 | -0.41971 | 0.46533 | -0.90196 | 0.367076 | 0.809473 |
| Karybeta3 | 5528.101 | 0.572922 | 0.635313 | 0.901795 | 0.367166 | 0.809473 |
| Opa1 | 4694.819 | 0.569285 | 0.631463 | 0.901533 | 0.367305 | 0.809473 |
| Wnt5 | 876.7172 | 1.077461 | 1.195524 | 0.901246 | 0.367458 | 0.809473 |
| Golgin104 | 940.0645 | -1.1258 | 1.249717 | -0.90084 | 0.367672 | 0.809473 |
| Cbl | 1478.097 | 0.762272 | 0.846193 | 0.900825 | 0.367681 | 0.809473 |
| CG32104 | 66.25976 | 3.281465 | 3.647219 | 0.899717 | 0.368271 | 0.810459 |
| DCP2 | 6279.164 | -0.48571 | 0.540028 | -0.89942 | 0.368427 | 0.810581 |
| CG9743 | 212.2241 | 2.646176 | 2.944639 | 0.898642 | 0.368843 | 0.811051 |

| | | | | | | |
|---|---|---|---|---|---|---|
| Sec6 | 462.2585 | 1.618546 | 1.800859 | 0.898763 | 0.368779 | 0.811051 |
| raw | 1656.245 | -1.22395 | 1.362415 | -0.89837 | 0.368988 | 0.811147 |
| Ctr9 | 1367.366 | 1.026062 | 1.142553 | 0.898044 | 0.369162 | 0.811307 |
| Rga | 839.8854 | 0.955179 | 1.064358 | 0.897423 | 0.369493 | 0.811664 |
| Surf4 | 3054.056 | -0.87277 | 0.973049 | -0.89695 | 0.369747 | 0.811664 |
| CG6231 | 190.1417 | 2.687941 | 2.999168 | 0.896229 | 0.370131 | 0.811664 |
| Hcf | 7812.39 | 0.442362 | 0.49353 | 0.896322 | 0.370081 | 0.811664 |
| AcCoAS | 22526.77 | 0.374225 | 0.417146 | 0.897109 | 0.369661 | 0.811664 |
| CG17807 | 91.98826 | -3.27355 | 3.650533 | -0.89673 | 0.369862 | 0.811664 |
| lncRNA:CR | 192.8984 | -2.11142 | 2.354596 | -0.89672 | 0.369866 | 0.811664 |
| CG3764 | 1513.679 | 1.057086 | 1.17949 | 0.896223 | 0.370134 | 0.811664 |
| E2f2 | 1474.757 | 0.868137 | 0.969425 | 0.895518 | 0.37051 | 0.811955 |
| grsm | 508.771 | -1.58746 | 1.773996 | -0.89485 | 0.370868 | 0.811955 |
| rdgBbeta | 1815.995 | 0.697622 | 0.779606 | 0.894839 | 0.370873 | 0.811955 |
| snz | 1416.385 | -1.15657 | 1.292221 | -0.89502 | 0.370775 | 0.811955 |
| CG4332 | 402.9756 | 2.11223 | 2.359381 | 0.895247 | 0.370655 | 0.811955 |
| Chmp1 | 4388.251 | -0.57016 | 0.636572 | -0.89567 | 0.37043 | 0.811955 |
| Rrp46 | 307.1054 | 2.311834 | 2.584193 | 0.894606 | 0.370998 | 0.812006 |
| CG31869 | 340.8528 | -1.36108 | 1.52412 | -0.89303 | 0.371842 | 0.812303 |
| Ugt49C1 | 137.5463 | -2.82637 | 3.162386 | -0.89375 | 0.371458 | 0.812303 |
| Su(var)2-1( | 1404.282 | 0.826599 | 0.92498 | 0.89364 | 0.371515 | 0.812303 |
| CG5382 | 1210.343 | 0.884047 | 0.989583 | 0.893353 | 0.371668 | 0.812303 |
| CG17691 | 5597.972 | -0.52749 | 0.590506 | -0.89328 | 0.371707 | 0.812303 |
| VepD | 584.5169 | 1.324025 | 1.481801 | 0.893524 | 0.371576 | 0.812303 |
| CG10418 | 597.3618 | -1.07923 | 1.20833 | -0.89316 | 0.371772 | 0.812303 |
| PGRP-LC | 1540.554 | -0.83546 | 0.936405 | -0.8922 | 0.372284 | 0.812882 |
| Su(z)12 | 678.2487 | 1.233604 | 1.382722 | 0.892156 | 0.372309 | 0.812882 |
| Mkrn1 | 1640.732 | -1.09781 | 1.231304 | -0.89158 | 0.372615 | 0.813329 |
| Usp16-45 | 582.1418 | -1.49855 | 1.682108 | -0.89087 | 0.372997 | 0.813555 |
| wdb | 13902.34 | 0.384237 | 0.431327 | 0.890825 | 0.373023 | 0.813555 |
| Toll-7 | 1936.594 | -0.88455 | 0.992744 | -0.89101 | 0.372923 | 0.813555 |
| CG1545 | 14109.92 | -0.49639 | 0.557348 | -0.89063 | 0.373128 | 0.813564 |
| MBD-like | 1165.685 | 0.836596 | 0.93979 | 0.890194 | 0.373362 | 0.813851 |
| CG14591 | 4282.075 | 0.556733 | 0.625832 | 0.889589 | 0.373687 | 0.813891 |
| sqd | 16720.69 | -0.38829 | 0.43657 | -0.88941 | 0.373785 | 0.813891 |
| rdgC | 1480.483 | -1.05653 | 1.187492 | -0.88972 | 0.373619 | 0.813891 |
| Sms | 4317.796 | 0.646678 | 0.726935 | 0.889596 | 0.373683 | 0.813891 |
| CG3226 | 1715.508 | -1.133 | 1.274611 | -0.8889 | 0.374057 | 0.81426 |
| Mtap | 948.3667 | -1.0041 | 1.130558 | -0.88815 | 0.374461 | 0.81492 |
| CG10631 | 3034.963 | -0.56746 | 0.639931 | -0.88676 | 0.375209 | 0.815475 |
| orb | 717.8065 | 1.101728 | 1.242731 | 0.886538 | 0.375328 | 0.815475 |
| CG17883 | 9983.485 | -0.45855 | 0.517343 | -0.88635 | 0.375427 | 0.815475 |
| Mcr | 2400.068 | -1.03351 | 1.165881 | -0.88647 | 0.375366 | 0.815475 |
| CG7970 | 747.2818 | 1.043084 | 1.176036 | 0.886949 | 0.375106 | 0.815475 |
| akirin | 4341.502 | -0.63709 | 0.718237 | -0.88702 | 0.375066 | 0.815475 |
| CG3609 | 2048.48 | -0.7789 | 0.877889 | -0.88724 | 0.374947 | 0.815475 |
| CG17202 | 1534.02 | -0.77727 | 0.877819 | -0.88546 | 0.375909 | 0.815857 |
| Tpi | 15192.49 | -0.40371 | 0.456033 | -0.88527 | 0.376013 | 0.815857 |
| Sbp2 | 236.5719 | -2.08304 | 2.352294 | -0.88554 | 0.375868 | 0.815857 |
| enc | 7431.313 | -0.50237 | 0.567598 | -0.88508 | 0.376112 | 0.815857 |

| | | | | | | |
|---|---|---|---|---|---|---|
| Gem2 | 1389.189 | -0.80906 | 0.913673 | -0.8855 | 0.375885 | 0.815857 |
| EMC10 | 5558.589 | 0.511954 | 0.580839 | 0.881403 | 0.3781 | 0.817862 |
| CG8673 | 93.76508 | -3.1587 | 3.583941 | -0.88135 | 0.378129 | 0.817862 |
| obe | 576.9523 | -1.21512 | 1.378012 | -0.88179 | 0.37789 | 0.817862 |
| Sep-02 | 2209.551 | -0.65555 | 0.743252 | -0.882 | 0.377777 | 0.817862 |
| X11Lbeta | 6347.322 | -0.59374 | 0.672541 | -0.88283 | 0.377329 | 0.817862 |
| mt:Cyt-b | 489384.4 | 0.329395 | 0.373807 | 0.881189 | 0.378215 | 0.817862 |
| CG34205 | 435.8655 | -1.66767 | 1.890967 | -0.88191 | 0.377825 | 0.817862 |
| Trf2 | 4677.061 | -0.64458 | 0.730068 | -0.8829 | 0.377289 | 0.817862 |
| Vps4 | 1958.027 | 0.756365 | 0.85727 | 0.882295 | 0.377617 | 0.817862 |
| CG16758 | 993.4772 | 1.487581 | 1.688304 | 0.88111 | 0.378258 | 0.817862 |
| CG17233 | 1521.2 | 0.788323 | 0.893628 | 0.88216 | 0.37769 | 0.817862 |
| Gs1l | 3601.983 | 0.544821 | 0.618243 | 0.881241 | 0.378188 | 0.817862 |
| CG11723 | 1911.464 | -0.75605 | 0.858714 | -0.88044 | 0.378618 | 0.818419 |
| Mur2B | 1856.643 | 0.691572 | 0.7858 | 0.880086 | 0.378813 | 0.818619 |
| scf | 4254.336 | -0.52692 | 0.599105 | -0.87952 | 0.379122 | 0.819066 |
| CG12935 | 481.4707 | 1.718179 | 1.954707 | 0.878996 | 0.379404 | 0.819234 |
| Trs33 | 393.5066 | 1.249854 | 1.421706 | 0.879123 | 0.379335 | 0.819234 |
| TMEM216 | 1233.529 | -0.93621 | 1.065413 | -0.87873 | 0.379549 | 0.819327 |
| Cyp4d1 | 112.221 | -2.5161 | 2.871154 | -0.87634 | 0.380847 | 0.819908 |
| CG2017 | 1428.289 | 0.883693 | 1.009324 | 0.87553 | 0.381286 | 0.819908 |
| dpr16 | 188.5898 | 2.047909 | 2.340358 | 0.875041 | 0.381552 | 0.819908 |
| IPIP | 705.8317 | 1.033074 | 1.1811 | 0.874671 | 0.381753 | 0.819908 |
| gro | 2214.829 | -0.63119 | 0.719696 | -0.87702 | 0.380476 | 0.819908 |
| CG31030 | 22035.73 | 0.351769 | 0.401972 | 0.875108 | 0.381515 | 0.819908 |
| CG6805 | 3084.894 | -0.58872 | 0.672083 | -0.87597 | 0.381048 | 0.819908 |
| CG30096 | 268.1036 | 2.093636 | 2.393741 | 0.874629 | 0.381776 | 0.819908 |
| lt | 4692.402 | 0.528862 | 0.602742 | 0.877426 | 0.380255 | 0.819908 |
| trio | 8024.333 | 0.454938 | 0.519595 | 0.875562 | 0.381268 | 0.819908 |
| Nna1 | 1487.264 | -0.88123 | 1.00629 | -0.87572 | 0.38118 | 0.819908 |
| CG12024 | 3084.338 | -0.5952 | 0.67831 | -0.87747 | 0.38023 | 0.819908 |
| CG13003 | 66.87175 | 3.418398 | 3.906768 | 0.874994 | 0.381577 | 0.819908 |
| gogo | 1132.991 | -1.11836 | 1.277233 | -0.87561 | 0.381243 | 0.819908 |
| Mbs | 5841.539 | -0.45718 | 0.522437 | -0.87509 | 0.381524 | 0.819908 |
| CG7945 | 2635.81 | 0.7987 | 0.909679 | 0.878002 | 0.379942 | 0.819908 |
| CG5645 | 201.322 | -2.31898 | 2.647968 | -0.87576 | 0.381161 | 0.819908 |
| rt | 269.8823 | -1.96687 | 2.249208 | -0.87447 | 0.381861 | 0.819908 |
| Rbfox1 | 24830.01 | -0.36151 | 0.412691 | -0.87598 | 0.381042 | 0.819908 |
| VGlut | 4118.042 | -1.1249 | 1.281829 | -0.87758 | 0.380173 | 0.819908 |
| Sec61alpha | 7604.989 | -0.43892 | 0.502314 | -0.87379 | 0.382231 | 0.820483 |
| zf30C | 1308.034 | 0.834224 | 0.955034 | 0.873502 | 0.38239 | 0.820604 |
| FucT6 | 2614.846 | 0.583073 | 0.667673 | 0.873291 | 0.382505 | 0.820631 |
| CG14762 | 15801.23 | -0.38211 | 0.43792 | -0.87255 | 0.382908 | 0.821277 |
| FBXO11 | 2716.611 | -0.62222 | 0.715278 | -0.8699 | 0.384356 | 0.822189 |
| CG1951 | 763.0615 | -0.95755 | 1.099075 | -0.87123 | 0.383626 | 0.822189 |
| zda | 3662.031 | 0.976216 | 1.121831 | 0.870199 | 0.384192 | 0.822189 |
| Sptr | 640.1761 | -1.35193 | 1.552955 | -0.87055 | 0.384 | 0.822189 |
| CG12325 | 142.9098 | -3.06272 | 3.520794 | -0.8699 | 0.384357 | 0.822189 |
| Pka-R1 | 8001.332 | -0.49434 | 0.567766 | -0.87067 | 0.383933 | 0.822189 |
| CG12025 | 6932.583 | 0.434444 | 0.499254 | 0.870188 | 0.384198 | 0.822189 |

| | | | | | | |
|---|---|---|---|---|---|---|
| HBS1 | 517.5435 | 1.175183 | 1.350493 | 0.870188 | 0.384198 | 0.822189 |
| CG6479 | 174.42 | 2.646501 | 3.036429 | 0.871583 | 0.383436 | 0.822189 |
| CG16979 | 998.6005 | -1.26378 | 1.451449 | -0.8707 | 0.383916 | 0.822189 |
| CG11378 | 886.7817 | -1.38459 | 1.592085 | -0.86967 | 0.38448 | 0.822232 |
| CG10147 | 543.9991 | 1.69928 | 1.954603 | 0.869374 | 0.384643 | 0.822361 |
| opa | 1594.742 | -0.69744 | 0.802763 | -0.86879 | 0.38496 | 0.822819 |
| fabp | 1459.567 | -0.73845 | 0.850534 | -0.86822 | 0.385273 | 0.823109 |
| CG42268 | 10894.61 | 0.414412 | 0.477339 | 0.868172 | 0.3853 | 0.823109 |
| mt:ND1 | 28179.95 | -0.43765 | 0.504582 | -0.86735 | 0.38575 | 0.82333 |
| koko | 280.0743 | -2.54503 | 2.933596 | -0.86755 | 0.385643 | 0.82333 |
| CG11263 | 268.5058 | -2.53121 | 2.918723 | -0.86723 | 0.385814 | 0.82333 |
| Blimp-1 | 6833.007 | -0.47244 | 0.54475 | -0.86727 | 0.385795 | 0.82333 |
| CG5783 | 153.5847 | 3.147245 | 3.635442 | 0.865712 | 0.386648 | 0.823731 |
| Syx1A | 21475.32 | 0.378579 | 0.437193 | 0.865931 | 0.386528 | 0.823731 |
| CG5098 | 3035.935 | -0.80895 | 0.933534 | -0.86655 | 0.386191 | 0.823731 |
| Vhl | 1324.118 | 0.774195 | 0.893741 | 0.866241 | 0.386358 | 0.823731 |
| Prosbeta5 | 5560.535 | -0.49483 | 0.571138 | -0.86639 | 0.386274 | 0.823731 |
| nSyb | 33665.47 | 0.35484 | 0.409945 | 0.86558 | 0.38672 | 0.823731 |
| Clic | 1543.121 | 0.854854 | 0.987599 | 0.865588 | 0.386716 | 0.823731 |
| px | 2607.364 | -0.58789 | 0.679353 | -0.86537 | 0.386833 | 0.823753 |
| trpl | 191.7333 | 2.31233 | 2.675572 | 0.864237 | 0.387457 | 0.824715 |
| Tim8 | 3333.856 | 0.539604 | 0.624414 | 0.864177 | 0.387491 | 0.824715 |
| CG4793 | 90.83306 | -3.33984 | 3.87256 | -0.86244 | 0.388447 | 0.826176 |
| CG3328 | 2155.19 | 0.661949 | 0.767595 | 0.862366 | 0.388486 | 0.826176 |
| Sxl | 9946.819 | 0.413408 | 0.479335 | 0.862463 | 0.388433 | 0.826176 |
| CG11030 | 2049.369 | 0.680241 | 0.78902 | 0.862134 | 0.388614 | 0.826229 |
| CG3704 | 733.6707 | -1.4601 | 1.696825 | -0.86049 | 0.389521 | 0.826459 |
| GlcAT-I | 432.989 | -1.42581 | 1.656516 | -0.86073 | 0.389386 | 0.826459 |
| Kdm2 | 1665.926 | -0.70977 | 0.824041 | -0.86132 | 0.389059 | 0.826459 |
| cdi | 4551.955 | 0.539838 | 0.627344 | 0.860513 | 0.389506 | 0.826459 |
| CG10254 | 987.1084 | 0.863356 | 1.003388 | 0.860441 | 0.389546 | 0.826459 |
| DMAP1 | 1147.569 | 0.790333 | 0.917871 | 0.86105 | 0.38921 | 0.826459 |
| CG7137 | 256.792 | -2.04134 | 2.369395 | -0.86154 | 0.388939 | 0.826459 |
| Dhc64C | 4947.257 | -0.59351 | 0.689763 | -0.86045 | 0.38954 | 0.826459 |
| ND-13A | 7334.433 | -0.44475 | 0.518198 | -0.85825 | 0.390753 | 0.826498 |
| Bub1 | 1677.8 | 0.82163 | 0.956532 | 0.858967 | 0.390358 | 0.826498 |
| Su(var)205 | 2469.993 | 0.613179 | 0.714184 | 0.858574 | 0.390576 | 0.826498 |
| Akap200 | 3615.878 | 0.620692 | 0.723243 | 0.858207 | 0.390778 | 0.826498 |
| CG31676 | 6195.722 | -0.60663 | 0.706746 | -0.85834 | 0.390702 | 0.826498 |
| CG31248 | 6372.192 | -0.58729 | 0.685417 | -0.85684 | 0.391534 | 0.826498 |
| Alg12 | 834.0444 | 1.312939 | 1.534433 | 0.855651 | 0.392191 | 0.826498 |
| CG11447 | 337.3668 | -1.3971 | 1.632405 | -0.85585 | 0.392078 | 0.826498 |
| His2Av | 4418.701 | -0.50826 | 0.594462 | -0.85499 | 0.392556 | 0.826498 |
| CAH9 | 135.1734 | 3.26516 | 3.817247 | 0.85537 | 0.392346 | 0.826498 |
| superdeatl | 784.1769 | 1.251527 | 1.465036 | 0.854264 | 0.392959 | 0.826498 |
| CG1607 | 5796.429 | -0.76875 | 0.899295 | -0.85483 | 0.392643 | 0.826498 |
| CG15909 | 298.5976 | -1.74081 | 2.03869 | -0.85389 | 0.393169 | 0.826498 |
| icln | 269.6315 | -2.14124 | 2.491954 | -0.85926 | 0.390195 | 0.826498 |
| Pms2 | 65.44615 | 2.849504 | 3.334259 | 0.854614 | 0.392765 | 0.826498 |
| mbt | 3306.17 | 0.635115 | 0.741472 | 0.856561 | 0.391688 | 0.826498 |

| | | | | | | |
|---|---|---|---|---|---|---|
| CG34348 | 788.7679 | -0.91164 | 1.067438 | -0.85404 | 0.393082 | 0.826498 |
| meigo | 1647.159 | -1.14354 | 1.337895 | -0.85473 | 0.392702 | 0.826498 |
| CG42766 | 68.81759 | 3.036111 | 3.540279 | 0.857591 | 0.391118 | 0.826498 |
| Unr | 11723.46 | 0.421817 | 0.492241 | 0.856931 | 0.391483 | 0.826498 |
| mth | 128.9737 | -2.82665 | 3.309618 | -0.85407 | 0.393066 | 0.826498 |
| CG44385 | 66.95445 | -3.34908 | 3.90352 | -0.85796 | 0.390913 | 0.826498 |
| CG32267 | 1751.521 | 0.692944 | 0.810026 | 0.85546 | 0.392297 | 0.826498 |
| Rab40 | 2787.557 | -0.68099 | 0.794294 | -0.85735 | 0.391253 | 0.826498 |
| CG8915 | 405.7039 | -1.91853 | 2.233723 | -0.85889 | 0.390401 | 0.826498 |
| CG6762 | 109.3859 | 2.049118 | 2.395353 | 0.855455 | 0.392299 | 0.826498 |
| CG14082 | 280.7312 | 1.869273 | 2.177166 | 0.85858 | 0.390572 | 0.826498 |
| Nc73EF | 17865.43 | -0.41724 | 0.486861 | -0.85701 | 0.39144 | 0.826498 |
| SsRbeta | 3505.193 | 0.55919 | 0.654561 | 0.854297 | 0.392941 | 0.826498 |
| D | 731.4068 | -1.20207 | 1.400641 | -0.85823 | 0.390766 | 0.826498 |
| vers | 795.7527 | -1.05853 | 1.235691 | -0.85663 | 0.391651 | 0.826498 |
| I-2 | 1523.8 | -1.24308 | 1.454238 | -0.8548 | 0.392661 | 0.826498 |
| loj | 1560.187 | -0.69995 | 0.814565 | -0.85929 | 0.390178 | 0.826498 |
| tank | 932.8746 | -1.3072 | 1.519696 | -0.86017 | 0.389695 | 0.826498 |
| Rrp1 | 686.3456 | 1.393314 | 1.625426 | 0.857199 | 0.391335 | 0.826498 |
| CG5902 | 1388.065 | -1.02026 | 1.195952 | -0.85309 | 0.393609 | 0.826774 |
| Svil | 5702.215 | -0.45566 | 0.534101 | -0.85313 | 0.393589 | 0.826774 |
| Ccn | 4340.777 | 0.51314 | 0.601448 | 0.853174 | 0.393563 | 0.826774 |
| Gpdh1 | 11393.33 | 0.403292 | 0.473159 | 0.852338 | 0.394026 | 0.827218 |
| CG14937 | 125.4011 | -2.82668 | 3.315987 | -0.85244 | 0.39397 | 0.827218 |
| GstE11 | 386.834 | -1.5194 | 1.783713 | -0.85182 | 0.394313 | 0.827604 |
| mEFTu1 | 2039.549 | -0.6264 | 0.735991 | -0.8511 | 0.394716 | 0.828234 |
| GluRIIE | 69.72957 | 2.888601 | 3.395189 | 0.850792 | 0.394885 | 0.828355 |
| asRNA:CR4 | 511.8596 | -1.19832 | 1.40876 | -0.85062 | 0.39498 | 0.828355 |
| Nmdmc | 1238.491 | -0.92749 | 1.091132 | -0.85003 | 0.39531 | 0.828365 |
| gb | 310.5329 | 1.404479 | 1.653988 | 0.849147 | 0.395799 | 0.828365 |
| Mccc2 | 151.0609 | -2.75584 | 3.241666 | -0.85013 | 0.395252 | 0.828365 |
| RpS2 | 21044.33 | 0.35358 | 0.416264 | 0.849414 | 0.395651 | 0.828365 |
| LSm1 | 434.8304 | 1.317961 | 1.552138 | 0.849126 | 0.395811 | 0.828365 |
| HSPC300 | 1084.912 | -0.81502 | 0.959805 | -0.84915 | 0.395799 | 0.828365 |
| Tat | 2759.5 | 0.564487 | 0.663968 | 0.850172 | 0.39523 | 0.828365 |
| CG13305 | 133.0809 | 3.261636 | 3.838855 | 0.849638 | 0.395526 | 0.828365 |
| CG17928 | 1157.336 | -1.18745 | 1.398749 | -0.84893 | 0.395918 | 0.828372 |
| CG3655 | 568.8761 | -1.50614 | 1.776914 | -0.84762 | 0.396652 | 0.829043 |
| CG1890 | 1242.348 | -0.89841 | 1.059708 | -0.84779 | 0.396556 | 0.829043 |
| CG3358 | 282.4558 | 2.029416 | 2.394024 | 0.847701 | 0.396605 | 0.829043 |
| Asator | 15461.91 | 0.369981 | 0.436313 | 0.847973 | 0.396453 | 0.829043 |
| Syn1 | 1753.906 | -0.82045 | 0.969553 | -0.84621 | 0.397434 | 0.829543 |
| TyrRII | 87.32376 | 2.927472 | 3.463854 | 0.845149 | 0.398028 | 0.829543 |
| tmod | 14166.7 | -0.37131 | 0.438833 | -0.84614 | 0.397477 | 0.829543 |
| pen-2 | 1237.931 | -0.76482 | 0.904835 | -0.84526 | 0.397966 | 0.829543 |
| CG42750 | 5380.548 | 0.45007 | 0.532223 | 0.845642 | 0.397752 | 0.829543 |
| hppy | 4430.695 | 0.562401 | 0.665266 | 0.845377 | 0.3979 | 0.829543 |
| CG4554 | 249.1467 | -1.95841 | 2.312674 | -0.84682 | 0.397097 | 0.829543 |
| CG13579 | 2811.589 | 0.65813 | 0.778509 | 0.845373 | 0.397903 | 0.829543 |
| tipE | 1420.113 | -0.72468 | 0.856979 | -0.84562 | 0.397763 | 0.829543 |

| | | | | | | |
|---|---|---|---|---|---|---|
| CG7341 | 381.8788 | 1.663134 | 1.965742 | 0.846059 | 0.39752 | 0.829543 |
| Ldh | 113.3713 | 2.44466 | 2.888472 | 0.846351 | 0.397357 | 0.829543 |
| bib | 950.676 | 1.532457 | 1.814261 | 0.844673 | 0.398294 | 0.829666 |
| CG5727 | 735.6762 | 1.214598 | 1.437755 | 0.844788 | 0.398229 | 0.829666 |
| wb | 110.2687 | -3.22883 | 3.827959 | -0.84349 | 0.398956 | 0.830831 |
| sfl | 1792.123 | 0.754654 | 0.894901 | 0.843282 | 0.399071 | 0.830853 |
| CG13766 | 675.0993 | 1.302434 | 1.54768 | 0.84154 | 0.400046 | 0.831769 |
| Gr32a | 141.3699 | -3.18427 | 3.78154 | -0.84206 | 0.399756 | 0.831769 |
| CG6689 | 794.4127 | 1.197844 | 1.424205 | 0.841061 | 0.400314 | 0.831769 |
| eIF3c | 2036.511 | -0.64635 | 0.767643 | -0.84199 | 0.399793 | 0.831769 |
| Rcd1 | 3876.827 | 0.602241 | 0.715545 | 0.841654 | 0.399982 | 0.831769 |
| Atpalpha | 525640.9 | -0.32186 | 0.382378 | -0.84174 | 0.399936 | 0.831769 |
| Ubi-p63E | 1603.035 | 0.674527 | 0.801759 | 0.84131 | 0.400175 | 0.831769 |
| rad50 | 734.6652 | 1.37192 | 1.631289 | 0.841003 | 0.400346 | 0.831769 |
| Ae2 | 13970.86 | 0.436245 | 0.518827 | 0.84083 | 0.400443 | 0.831769 |
| GAA1 | 946.7898 | -1.33811 | 1.592336 | -0.84034 | 0.400716 | 0.832121 |
| bsf | 1877.443 | 0.672571 | 0.800826 | 0.839847 | 0.400994 | 0.832483 |
| Surf1 | 187.4742 | -2.50865 | 2.990539 | -0.83886 | 0.401546 | 0.833197 |
| D19B | 152.7091 | 1.939138 | 2.311442 | 0.83893 | 0.401509 | 0.833197 |
| M1BP | 1098.152 | -0.85491 | 1.019615 | -0.83846 | 0.401772 | 0.833451 |
| FMRFa | 226.5702 | 1.934468 | 2.309109 | 0.837755 | 0.402168 | 0.833534 |
| Gs2 | 72529.54 | -0.35482 | 0.423587 | -0.83765 | 0.402227 | 0.833534 |
| CG6769 | 338.5005 | 1.624415 | 1.938663 | 0.837905 | 0.402084 | 0.833534 |
| CG32039 | 1292.358 | 0.7693 | 0.917861 | 0.838144 | 0.40195 | 0.833534 |
| Mal-A5 | 5598.572 | -0.55197 | 0.659311 | -0.83719 | 0.402485 | 0.833637 |
| lncRNA:CR | 1733.519 | -0.85921 | 1.02622 | -0.83726 | 0.402448 | 0.833637 |
| Fur2 | 3894.729 | 0.588063 | 0.702674 | 0.836892 | 0.402653 | 0.833771 |
| CG13287 | 364.0618 | -2.52693 | 3.020232 | -0.83667 | 0.402779 | 0.833816 |
| mRpS34 | 540.309 | 1.570827 | 1.878144 | 0.836372 | 0.402946 | 0.833946 |
| CG10804 | 5313.895 | 0.58157 | 0.695952 | 0.835646 | 0.403354 | 0.834576 |
| CG32091 | 251.0335 | 1.416866 | 1.695947 | 0.835443 | 0.403468 | 0.834597 |
| CG12171 | 1400.911 | -1.19309 | 1.42909 | -0.83486 | 0.403795 | 0.835024 |
| CG17904 | 1671.759 | -0.70345 | 0.84313 | -0.83434 | 0.404091 | 0.835024 |
| Psa | 3163.113 | -0.52771 | 0.632328 | -0.83455 | 0.403969 | 0.835024 |
| VhaAC39-1 | 14372.76 | 0.364415 | 0.436764 | 0.834352 | 0.404083 | 0.835024 |
| D1 | 2568.051 | -0.56176 | 0.674142 | -0.8333 | 0.404677 | 0.836021 |
| CG10194 | 870.8975 | 1.532249 | 1.839327 | 0.833049 | 0.404817 | 0.836094 |
| fng | 2682.573 | 0.581344 | 0.698304 | 0.832508 | 0.405122 | 0.836482 |
| CG43373 | 2429.906 | 0.715016 | 0.859037 | 0.832347 | 0.405213 | 0.836482 |
| CG17048 | 64.10158 | -3.24684 | 3.905531 | -0.83134 | 0.405779 | 0.836804 |
| Pdrg1 | 146.1606 | 2.150673 | 2.585832 | 0.831714 | 0.40557 | 0.836804 |
| lncRNA:CR | 3165.193 | 0.774728 | 0.931719 | 0.831504 | 0.405689 | 0.836804 |
| CG6674 | 618.7654 | -1.41292 | 1.699582 | -0.83133 | 0.405786 | 0.836804 |
| CG13843 | 79.30458 | 3.248621 | 3.909106 | 0.831039 | 0.405951 | 0.836812 |
| CG32512 | 2348.45 | -0.57699 | 0.69437 | -0.83096 | 0.405999 | 0.836812 |
| CG12499 | 287.6983 | -1.82194 | 2.194098 | -0.83038 | 0.406322 | 0.836835 |
| Cfp1 | 813.3487 | -1.23246 | 1.483825 | -0.8306 | 0.406202 | 0.836835 |
| lncRNA:CR | 754.1037 | 1.190994 | 1.43423 | 0.830406 | 0.406309 | 0.836835 |
| CG32333 | 1210.624 | 0.955451 | 1.15208 | 0.829326 | 0.40692 | 0.837205 |
| Lztr1 | 1021.382 | 0.976597 | 1.177514 | 0.829372 | 0.406894 | 0.837205 |

| | | | | | | |
|---|---|---|---|---|---|---|
| unc-104 | 59338.19 | 0.367902 | 0.443431 | 0.829672 | 0.406724 | 0.837205 |
| Tnpo | 4307.22 | -0.51406 | 0.619648 | -0.8296 | 0.406764 | 0.837205 |
| Dyrk3 | 12934.05 | -0.40065 | 0.483299 | -0.82898 | 0.407116 | 0.837395 |
| CG10778 | 354.1329 | -1.80716 | 2.183337 | -0.8277 | 0.407838 | 0.837846 |
| ND-75 | 4734.258 | -0.54126 | 0.653538 | -0.8282 | 0.407555 | 0.837846 |
| ND-MWFE | 915.7186 | 0.852612 | 1.03013 | 0.827674 | 0.407855 | 0.837846 |
| Pss | 258.0524 | 1.699074 | 2.052608 | 0.827763 | 0.407804 | 0.837846 |
| Pi3K68D | 1830.327 | -0.62967 | 0.760776 | -0.82767 | 0.407858 | 0.837846 |
| CG14853 | 7140.381 | 0.689877 | 0.83405 | 0.827142 | 0.408157 | 0.838246 |
| CkIalpha | 9054.514 | -0.4035 | 0.487991 | -0.82687 | 0.408311 | 0.838349 |
| Cks85A | 554.2912 | -1.36108 | 1.64737 | -0.82622 | 0.408682 | 0.838466 |
| Nup358 | 1538.342 | 0.773198 | 0.935648 | 0.826378 | 0.40859 | 0.838466 |
| Sgs1 | 162.0292 | -2.98944 | 3.61712 | -0.82647 | 0.408538 | 0.838466 |
| eRF1 | 4991.415 | -0.53353 | 0.646248 | -0.82558 | 0.40904 | 0.838791 |
| beat-IIa | 6378.725 | 0.441641 | 0.535724 | 0.824382 | 0.409722 | 0.838791 |
| Klp98A | 2855.447 | 0.72019 | 0.873582 | 0.82441 | 0.409707 | 0.838791 |
| mys | 3883.083 | 0.576596 | 0.698737 | 0.825196 | 0.40926 | 0.838791 |
| CG3792 | 1706.445 | -0.71212 | 0.8637 | -0.8245 | 0.409657 | 0.838791 |
| Ptpmeg2 | 7848.49 | 0.504764 | 0.612447 | 0.824177 | 0.409839 | 0.838791 |
| CG11000 | 11833.26 | -0.42057 | 0.509629 | -0.82524 | 0.409235 | 0.838791 |
| CG43736 | 2897.51 | 0.552752 | 0.670738 | 0.824096 | 0.409885 | 0.838791 |
| CG11679 | 227.6528 | 2.123927 | 2.576246 | 0.824427 | 0.409697 | 0.838791 |
| mRpL12 | 2147.013 | -0.62892 | 0.762004 | -0.82535 | 0.40917 | 0.838791 |
| CG5245 | 100.8497 | 3.21624 | 3.905071 | 0.823606 | 0.410163 | 0.838933 |
| sigmar | 1817.409 | 0.68369 | 0.830087 | 0.823636 | 0.410146 | 0.838933 |
| Ostgamma | 2153.234 | -0.75033 | 0.912263 | -0.82249 | 0.410799 | 0.839402 |
| CG4901 | 557.0526 | 1.417833 | 1.72378 | 0.822514 | 0.410784 | 0.839402 |
| Tom70 | 1099.94 | -0.76642 | 0.931932 | -0.8224 | 0.410847 | 0.839402 |
| Rlb1 | 67.76455 | 3.211683 | 3.905811 | 0.822283 | 0.410916 | 0.839402 |
| FipoQ | 196.7412 | -1.99624 | 2.42708 | -0.82249 | 0.4108 | 0.839402 |
| Swip-1 | 1423.625 | 0.726846 | 0.884583 | 0.821681 | 0.411258 | 0.839888 |
| Vps39 | 1082.539 | 1.499262 | 1.827164 | 0.820541 | 0.411908 | 0.840003 |
| tnc | 346.3908 | -1.7962 | 2.189337 | -0.82043 | 0.41197 | 0.840003 |
| CG34229 | 168.6121 | 2.043929 | 2.488599 | 0.821317 | 0.411466 | 0.840003 |
| shps | 427.8704 | 2.10232 | 2.562879 | 0.820296 | 0.412047 | 0.840003 |
| asRNA:CR4 | 198.0807 | 1.539201 | 1.875695 | 0.820603 | 0.411872 | 0.840003 |
| ago | 2153.042 | -0.62113 | 0.757156 | -0.82035 | 0.412017 | 0.840003 |
| UBL3 | 3278.867 | -0.53305 | 0.649592 | -0.8206 | 0.411877 | 0.840003 |
| CG30008 | 219.5224 | 2.292363 | 2.797159 | 0.819533 | 0.412483 | 0.840677 |
| vig | 5086.148 | -0.55821 | 0.681915 | -0.81859 | 0.41302 | 0.841346 |
| Cdc42 | 1171.62 | -0.93142 | 1.137604 | -0.81875 | 0.412928 | 0.841346 |
| Ugt49B1 | 695.6854 | -1.30565 | 1.595561 | -0.8183 | 0.413185 | 0.841468 |
| CG42613 | 78696.12 | -0.37105 | 0.453588 | -0.81803 | 0.413342 | 0.841513 |
| Fife | 11430.25 | 0.37145 | 0.454154 | 0.817895 | 0.413417 | 0.841513 |
| bab2 | 741.57 | 0.937419 | 1.1464 | 0.817707 | 0.413525 | 0.841518 |
| tok | 3763.006 | 0.58054 | 0.710871 | 0.816659 | 0.414123 | 0.842253 |
| CG5742 | 513.947 | 1.307374 | 1.601144 | 0.816524 | 0.4142 | 0.842253 |
| CG2818 | 794.132 | -1.18924 | 1.456377 | -0.81657 | 0.414172 | 0.842253 |
| PGAP2 | 706.1011 | 1.29524 | 1.588417 | 0.815428 | 0.414827 | 0.842267 |
| VhaM9.7-b | 4891.771 | 0.487989 | 0.599126 | 0.814502 | 0.415357 | 0.842267 |

| | | | | | | |
|---|---|---|---|---|---|---|
| CCT4 | 1324.896 | -0.78792 | 0.965839 | -0.81579 | 0.414619 | 0.842267 |
| Vha100-2 | 1056.691 | 0.94209 | 1.15639 | 0.814682 | 0.415254 | 0.842267 |
| Idi | 698.4507 | -1.45709 | 1.787144 | -0.81532 | 0.414889 | 0.842267 |
| HmgZ | 9492.146 | -0.38217 | 0.469392 | -0.81419 | 0.415538 | 0.842267 |
| Pgant1 | 650.8556 | 1.08936 | 1.336879 | 0.814853 | 0.415156 | 0.842267 |
| Iswi | 1461.177 | -0.68035 | 0.833903 | -0.81586 | 0.414581 | 0.842267 |
| CG8712 | 299.3107 | 2.086631 | 2.562944 | 0.814154 | 0.415557 | 0.842267 |
| Pde1c | 9760.026 | 0.3841 | 0.470612 | 0.816172 | 0.414402 | 0.842267 |
| ksh | 1396.9 | -0.84402 | 1.035917 | -0.81475 | 0.415214 | 0.842267 |
| CG32137 | 147.2568 | -2.22883 | 2.737689 | -0.81413 | 0.415571 | 0.842267 |
| CG15439 | 652.543 | 0.950117 | 1.166971 | 0.814173 | 0.415546 | 0.842267 |
| eIF2alpha | 1423.381 | -0.7998 | 0.982785 | -0.81381 | 0.415751 | 0.842418 |
| Pde8 | 4121.999 | -0.53424 | 0.657162 | -0.81296 | 0.416244 | 0.843203 |
| CG10874 | 355.5127 | 1.231821 | 1.515728 | 0.812692 | 0.416394 | 0.843296 |
| epsilonCOI | 2096.143 | 0.699945 | 0.862679 | 0.811362 | 0.417158 | 0.843564 |
| Ask1 | 995.5747 | -0.94715 | 1.167904 | -0.81099 | 0.417374 | 0.843564 |
| Cchl | 3162.699 | -0.55453 | 0.684067 | -0.81064 | 0.417573 | 0.843564 |
| CG42322 | 11156.42 | 0.481942 | 0.594486 | 0.810687 | 0.417546 | 0.843564 |
| CG42827 | 153.9038 | -2.11072 | 2.599548 | -0.81195 | 0.416818 | 0.843564 |
| St3 | 4414.085 | 0.532597 | 0.656573 | 0.811177 | 0.417264 | 0.843564 |
| Doa | 7513.797 | 0.571525 | 0.704582 | 0.811155 | 0.417277 | 0.843564 |
| PMP34 | 1611.538 | -1.12543 | 1.386069 | -0.81196 | 0.416817 | 0.843564 |
| CG4661 | 189.3311 | 2.317704 | 2.859136 | 0.810631 | 0.417578 | 0.843564 |
| CG3408 | 309.6275 | 1.673956 | 2.06391 | 0.811061 | 0.417331 | 0.843564 |
| CG12541 | 410.1531 | -1.20517 | 1.487331 | -0.81029 | 0.417773 | 0.843588 |
| angel | 463.7944 | 1.645905 | 2.03137 | 0.810244 | 0.4178 | 0.843588 |
| CG18273 | 389.3723 | -2.07081 | 2.56865 | -0.80619 | 0.420135 | 0.843938 |
| Hem | 1281.177 | 0.807871 | 1.001225 | 0.806883 | 0.419734 | 0.843938 |
| PrBP | 3485.294 | -0.48978 | 0.606851 | -0.80708 | 0.419619 | 0.843938 |
| Samuel | 365.5859 | 1.461647 | 1.805538 | 0.809536 | 0.418207 | 0.843938 |
| sky | 4141.765 | -0.48363 | 0.599848 | -0.80625 | 0.4201 | 0.843938 |
| MAGE | 90.91618 | -3.16127 | 3.916117 | -0.80725 | 0.419524 | 0.843938 |
| CG5500 | 1232.796 | 0.738113 | 0.913112 | 0.808348 | 0.41889 | 0.843938 |
| RpL32 | 6937.9 | 0.472763 | 0.586081 | 0.806651 | 0.419868 | 0.843938 |
| CalpA | 2362.154 | 0.567731 | 0.703578 | 0.80692 | 0.419713 | 0.843938 |
| CG11368 | 160.9572 | 2.258567 | 2.794397 | 0.808248 | 0.418948 | 0.843938 |
| CG2129 | 229.1107 | 2.11789 | 2.616735 | 0.809363 | 0.418306 | 0.843938 |
| LBR | 506.0488 | -1.68539 | 2.08556 | -0.80812 | 0.41902 | 0.843938 |
| CG30495 | 563.3092 | -1.51104 | 1.87355 | -0.80651 | 0.419948 | 0.843938 |
| mtd | 39722.6 | 0.379144 | 0.469032 | 0.808356 | 0.418886 | 0.843938 |
| p | 958.5107 | -1.39753 | 1.733292 | -0.80628 | 0.420079 | 0.843938 |
| inaE | 3742.104 | 0.52002 | 0.643738 | 0.807813 | 0.419199 | 0.843938 |
| CG9861 | 77.54968 | 3.142926 | 3.898573 | 0.806173 | 0.420143 | 0.843938 |
| cpo | 12192.17 | 0.363885 | 0.449802 | 0.808988 | 0.418522 | 0.843938 |
| tn | 75.56184 | 3.142696 | 3.89861 | 0.806107 | 0.420181 | 0.843938 |
| PyK | 22643.69 | -0.33348 | 0.411875 | -0.80965 | 0.418139 | 0.843938 |
| Pcyt1 | 4056.443 | 0.540724 | 0.669267 | 0.807934 | 0.419129 | 0.843938 |
| CG5664 | 647.53 | 1.041635 | 1.293096 | 0.805536 | 0.420511 | 0.844177 |
| CG5522 | 2199.229 | 0.630051 | 0.782109 | 0.80558 | 0.420485 | 0.844177 |
| bru3 | 14445.3 | -0.42058 | 0.522248 | -0.80533 | 0.42063 | 0.844206 |

| | | | | | | |
|---|---|---|---|---|---|---|
| CG7766 | 6101.105 | -0.42228 | 0.52479 | -0.80467 | 0.421009 | 0.844542 |
| CG31687 | 63.05612 | -2.83256 | 3.519758 | -0.80476 | 0.420958 | 0.844542 |
| CG30157 | 130.7416 | 3.098849 | 3.853303 | 0.804206 | 0.421278 | 0.844872 |
| RpLP2 | 20554.14 | 0.373166 | 0.464287 | 0.80374 | 0.421547 | 0.8452 |
| tsh | 235.9984 | 1.857374 | 2.313128 | 0.802971 | 0.421992 | 0.845881 |
| Mob2 | 7699.863 | 0.453128 | 0.564639 | 0.80251 | 0.422258 | 0.846204 |
| csw | 9660.655 | 0.38987 | 0.488401 | 0.798258 | 0.424721 | 0.847107 |
| CG32797 | 70.00773 | -3.08072 | 3.86789 | -0.79649 | 0.425749 | 0.847107 |
| CG11307 | 853.3517 | -1.0027 | 1.257988 | -0.79707 | 0.425413 | 0.847107 |
| Chro | 1428.404 | 0.763891 | 0.956739 | 0.798431 | 0.42462 | 0.847107 |
| corto | 12419.11 | -0.3884 | 0.48758 | -0.79658 | 0.425697 | 0.847107 |
| snRNP-U1- | 1115.636 | 0.810782 | 1.016314 | 0.797767 | 0.425006 | 0.847107 |
| CG5149 | 2408.502 | 1.113234 | 1.397559 | 0.796556 | 0.425709 | 0.847107 |
| Yip1d1 | 1342.426 | 0.762643 | 0.958203 | 0.795909 | 0.426085 | 0.847107 |
| p23 | 3500.221 | -0.49916 | 0.627205 | -0.79584 | 0.426124 | 0.847107 |
| Pnn | 1446.091 | -0.77118 | 0.967492 | -0.79709 | 0.425398 | 0.847107 |
| MRG15 | 626.1295 | 1.147573 | 1.43978 | 0.797048 | 0.425423 | 0.847107 |
| EloB | 1973.273 | 0.89132 | 1.114163 | 0.799991 | 0.423716 | 0.847107 |
| CG13833 | 323.9317 | -2.24911 | 2.825913 | -0.79589 | 0.426098 | 0.847107 |
| CG1571 | 150.1423 | -2.09013 | 2.622705 | -0.79694 | 0.425487 | 0.847107 |
| grau | 998.4872 | -1.01636 | 1.277409 | -0.79564 | 0.426241 | 0.847107 |
| Alk | 12517.09 | 0.36278 | 0.455554 | 0.796349 | 0.425829 | 0.847107 |
| ari-1 | 4378.009 | -0.50322 | 0.632092 | -0.79612 | 0.425961 | 0.847107 |
| OstDelta | 638.194 | 0.988634 | 1.239703 | 0.797477 | 0.425174 | 0.847107 |
| CG17065 | 3572.98 | 0.739353 | 0.923729 | 0.8004 | 0.423479 | 0.847107 |
| Med | 619.2923 | -1.12714 | 1.407217 | -0.80097 | 0.423147 | 0.847107 |
| unc-13 | 50041.25 | 0.369497 | 0.461229 | 0.801112 | 0.423067 | 0.847107 |
| CG33267 | 669.8511 | 0.999314 | 1.255787 | 0.795767 | 0.426167 | 0.847107 |
| sotv | 3618.508 | 0.603631 | 0.753504 | 0.801099 | 0.423075 | 0.847107 |
| CG17568 | 1010.181 | 0.768893 | 0.963792 | 0.797779 | 0.424999 | 0.847107 |
| Frq1 | 11791.2 | 0.37128 | 0.463574 | 0.800908 | 0.423185 | 0.847107 |
| Dbx | 177.0676 | -2.08665 | 2.620109 | -0.7964 | 0.4258 | 0.847107 |
| RpII140 | 1566.42 | -0.8265 | 1.037929 | -0.7963 | 0.425859 | 0.847107 |
| l(3)72Dn | 396.7408 | -1.46711 | 1.844223 | -0.79552 | 0.426313 | 0.847107 |
| asRNA:CR4 | 386.715 | -1.27506 | 1.593098 | -0.80036 | 0.4235 | 0.847107 |
| wg | 84.60345 | 2.921474 | 3.669116 | 0.796234 | 0.425896 | 0.847107 |
| lncRNA:CR | 606.7608 | 1.294944 | 1.626666 | 0.796073 | 0.42599 | 0.847107 |
| CG6179 | 599.7101 | 1.510428 | 1.886194 | 0.800781 | 0.423259 | 0.847107 |
| Mo25 | 5117.076 | 0.561761 | 0.70619 | 0.795482 | 0.426333 | 0.847107 |
| peng | 973.1433 | -0.9622 | 1.205824 | -0.79796 | 0.424894 | 0.847107 |
| CG11927 | 423.3693 | 1.618831 | 2.035341 | 0.795361 | 0.426403 | 0.847107 |
| Rilpl | 3788.693 | -0.47334 | 0.595388 | -0.79502 | 0.426604 | 0.847295 |
| snRNA:7Sk | 5509.63 | -0.49865 | 0.627647 | -0.79448 | 0.426916 | 0.847497 |
| Rab5 | 5198.056 | 0.445922 | 0.561175 | 0.794623 | 0.426833 | 0.847497 |
| Kr-h1 | 6940.633 | 0.50268 | 0.63293 | 0.794212 | 0.427072 | 0.847597 |
| lncRNA:CR | 88.16136 | 2.456208 | 3.093823 | 0.793907 | 0.42725 | 0.847739 |
| Prosbeta4 | 1700.985 | 0.648192 | 0.817128 | 0.793257 | 0.427628 | 0.848281 |
| mus201 | 752.5815 | -1.0517 | 1.327084 | -0.79249 | 0.428076 | 0.84854 |
| CG34401 | 6475.433 | -0.40744 | 0.513993 | -0.79269 | 0.427958 | 0.84854 |
| lncRNA:aca | 120.4714 | 2.712687 | 3.422832 | 0.792527 | 0.428054 | 0.84854 |

| | | | | | | |
|---|---|---|---|---|---|---|
| CG43795 | 232.662 | 1.417579 | 1.790202 | 0.791854 | 0.428446 | 0.848854 |
| tth | 417.2546 | 1.150418 | 1.452624 | 0.791958 | 0.428385 | 0.848854 |
| CG7582 | 9765.512 | -0.40996 | 0.518137 | -0.79121 | 0.428819 | 0.849383 |
| mthl5 | 119.6342 | 2.454427 | 3.103968 | 0.790738 | 0.429097 | 0.849724 |
| SoYb | 829.8415 | -1.31476 | 1.66364 | -0.79029 | 0.429359 | 0.850034 |
| CG8679 | 130.0054 | 2.131587 | 2.697974 | 0.790069 | 0.429487 | 0.850078 |
| RpL11 | 8093.472 | 0.447129 | 0.566172 | 0.789742 | 0.429679 | 0.850247 |
| TBCD | 200.449 | -1.73687 | 2.200785 | -0.7892 | 0.429994 | 0.85066 |
| Paics | 110.7493 | -2.3502 | 2.97916 | -0.78888 | 0.430182 | 0.850823 |
| asRNA:CR4 | 485.3721 | -1.02205 | 1.295918 | -0.78867 | 0.430304 | 0.850856 |
| AstC | 62.03944 | -3.09369 | 3.923819 | -0.78844 | 0.43044 | 0.850915 |
| CG12728 | 153.2474 | 1.689592 | 2.145486 | 0.78751 | 0.430983 | 0.851149 |
| eIF4EHP | 7955.232 | 0.436675 | 0.554492 | 0.787522 | 0.430976 | 0.851149 |
| dysc | 14272.51 | 0.37963 | 0.481972 | 0.787659 | 0.430896 | 0.851149 |
| Fum1 | 3912.761 | -0.5231 | 0.664073 | -0.78771 | 0.430864 | 0.851149 |
| tyf | 4821.624 | 0.431669 | 0.549954 | 0.784917 | 0.432502 | 0.851561 |
| Hpr1 | 557.6595 | -1.04505 | 1.332732 | -0.78414 | 0.432957 | 0.851561 |
| Fgop2 | 2035.653 | 0.648104 | 0.826548 | 0.78411 | 0.432976 | 0.851561 |
| CG33296 | 9680.704 | -0.41188 | 0.524374 | -0.78547 | 0.43218 | 0.851561 |
| CG14401 | 326.5875 | 1.806145 | 2.304062 | 0.783896 | 0.433101 | 0.851561 |
| CG1227 | 183.722 | -2.06656 | 2.633369 | -0.78476 | 0.432596 | 0.851561 |
| dpr15 | 387.417 | 1.279686 | 1.63232 | 0.783968 | 0.433059 | 0.851561 |
| Not10 | 682.911 | -1.03372 | 1.316686 | -0.78509 | 0.432401 | 0.851561 |
| HEATR2 | 488.2222 | 1.131945 | 1.443551 | 0.78414 | 0.432958 | 0.851561 |
| XNP | 2384.784 | -0.68029 | 0.865052 | -0.78641 | 0.431627 | 0.851561 |
| faf | 4366.521 | 0.455024 | 0.579007 | 0.785869 | 0.431944 | 0.851561 |
| AnxB9 | 7788.089 | -0.40322 | 0.513174 | -0.78573 | 0.432025 | 0.851561 |
| NAT1 | 6456.725 | -0.48297 | 0.615207 | -0.78505 | 0.432424 | 0.851561 |
| Arpc4 | 1743.744 | 0.606928 | 0.774148 | 0.783995 | 0.433043 | 0.851561 |
| CG7974 | 508.5267 | 1.188538 | 1.514231 | 0.784912 | 0.432505 | 0.851561 |
| CG32052 | 1943.039 | 0.69776 | 0.88918 | 0.784724 | 0.432616 | 0.851561 |
| CG8005 | 125.8116 | 2.33112 | 2.97137 | 0.784527 | 0.432731 | 0.851561 |
| CG15382 | 247.2584 | -2.18546 | 2.77981 | -0.78619 | 0.431757 | 0.851561 |
| CG7632 | 1836.505 | -0.99313 | 1.268183 | -0.78311 | 0.433562 | 0.852049 |
| smash | 3748.644 | 0.525066 | 0.670485 | 0.783114 | 0.43356 | 0.852049 |
| klar | 2649.794 | 0.534745 | 0.683326 | 0.782563 | 0.433884 | 0.852473 |
| asRNA:CR4 | 97.06374 | -2.09221 | 2.675881 | -0.78188 | 0.434288 | 0.853057 |
| EMC5 | 847.9146 | 0.847375 | 1.0844 | 0.781423 | 0.434554 | 0.853371 |
| Debcl | 962.6124 | -1.06999 | 1.370143 | -0.78094 | 0.43484 | 0.853725 |
| CG14778 | 172.7892 | -2.38544 | 3.055679 | -0.78066 | 0.435003 | 0.853805 |
| RpL35A | 11763.55 | 0.404354 | 0.51931 | 0.778637 | 0.436194 | 0.853805 |
| Vha55 | 19833.55 | 0.382127 | 0.48969 | 0.780345 | 0.435188 | 0.853805 |
| CG31345 | 6255.816 | 0.414285 | 0.531187 | 0.779923 | 0.435436 | 0.853805 |
| CG14321 | 1618.195 | -0.67399 | 0.865599 | -0.77865 | 0.436189 | 0.853805 |
| Rift | 362.0897 | 1.303369 | 1.673629 | 0.778768 | 0.436116 | 0.853805 |
| CG6701 | 1831.218 | -0.71179 | 0.914148 | -0.77864 | 0.436191 | 0.853805 |
| sax | 4589.502 | -0.50587 | 0.648485 | -0.78008 | 0.435342 | 0.853805 |
| CG9902 | 1224.269 | 1.143697 | 1.467272 | 0.779472 | 0.435702 | 0.853805 |
| dac | 574.8334 | -1.28614 | 1.650695 | -0.77915 | 0.435889 | 0.853805 |
| CG34408 | 1607.875 | 0.967543 | 1.242596 | 0.778646 | 0.436188 | 0.853805 |

| | | | | | | |
|---|---|---|---|---|---|---|
| SerT | 252.1073 | -1.97632 | 2.533673 | -0.78002 | 0.435378 | 0.853805 |
| Sec15 | 441.5195 | -1.29519 | 1.66366 | -0.77852 | 0.436264 | 0.853805 |
| CG32795 | 1435.509 | -0.82089 | 1.056445 | -0.77703 | 0.437142 | 0.854243 |
| CG1218 | 123.7079 | -2.21403 | 2.84979 | -0.77691 | 0.437213 | 0.854243 |
| CG9766 | 734.5307 | 0.911088 | 1.172625 | 0.776964 | 0.43718 | 0.854243 |
| CG14505 | 164.3581 | 2.151002 | 2.765237 | 0.777873 | 0.436644 | 0.854243 |
| lncRNA:CR | 158.2017 | 1.83318 | 2.357798 | 0.777497 | 0.436866 | 0.854243 |
| ap | 3494.692 | 0.499922 | 0.642939 | 0.777557 | 0.43683 | 0.854243 |
| CG31690 | 2136.514 | -0.5919 | 0.7619 | -0.77687 | 0.437233 | 0.854243 |
| CG42741 | 332.0562 | 1.486126 | 1.913446 | 0.776675 | 0.437351 | 0.854265 |
| CG17698 | 16127.74 | 0.33071 | 0.425987 | 0.776338 | 0.437549 | 0.854445 |
| CG17272 | 2413.599 | -0.57664 | 0.74299 | -0.77611 | 0.437687 | 0.854506 |
| lncRNA:CR | 80.40356 | 2.92491 | 3.770395 | 0.775757 | 0.437893 | 0.854564 |
| CG13707 | 2475.443 | -0.60306 | 0.777441 | -0.77569 | 0.43793 | 0.854564 |
| hng1 | 378.1758 | -1.39389 | 1.798808 | -0.7749 | 0.4384 | 0.855273 |
| CG11319 | 3892.33 | -0.45787 | 0.591148 | -0.77454 | 0.438613 | 0.855273 |
| PolD1 | 392.5349 | -1.66878 | 2.154259 | -0.77464 | 0.438551 | 0.855273 |
| RpS27A | 16809.3 | 0.344759 | 0.446425 | 0.772267 | 0.439957 | 0.856022 |
| CG10383 | 956.7718 | 0.96116 | 1.244151 | 0.772543 | 0.439793 | 0.856022 |
| Aasdh | 890.6096 | -1.38765 | 1.796193 | -0.77255 | 0.439789 | 0.856022 |
| Epg5 | 387.3322 | 1.653631 | 2.139243 | 0.772998 | 0.439523 | 0.856022 |
| CycE | 149.8854 | 1.74239 | 2.253679 | 0.773132 | 0.439445 | 0.856022 |
| CycT | 3782.41 | 0.46627 | 0.603394 | 0.772745 | 0.439673 | 0.856022 |
| RpS13 | 15575.58 | -0.33692 | 0.435496 | -0.77366 | 0.439134 | 0.856022 |
| CG34150 | 660.2999 | -0.94744 | 1.226415 | -0.77253 | 0.439802 | 0.856022 |
| l(2)gd1 | 353.5257 | -1.39226 | 1.802578 | -0.77237 | 0.439894 | 0.856022 |
| Irc | 1697.069 | 0.740651 | 0.960322 | 0.771254 | 0.440557 | 0.856152 |
| CG13917 | 8957.45 | -0.38983 | 0.505219 | -0.77161 | 0.440346 | 0.856152 |
| CG7879 | 1642.233 | 0.644026 | 0.834919 | 0.771363 | 0.440492 | 0.856152 |
| CG1265 | 4979.58 | 0.477792 | 0.619031 | 0.771838 | 0.440211 | 0.856152 |
| CG32641 | 228.399 | 1.783222 | 2.310496 | 0.771792 | 0.440238 | 0.856152 |
| CG2924 | 1441.688 | -0.73918 | 0.965838 | -0.76533 | 0.444075 | 0.856326 |
| IntS6 | 2103.769 | 0.755103 | 0.986523 | 0.765419 | 0.444022 | 0.856326 |
| DIP-theta | 22119.29 | 0.319922 | 0.41701 | 0.767181 | 0.442974 | 0.856326 |
| CG14005 | 778.4722 | 1.349979 | 1.76456 | 0.765051 | 0.444241 | 0.856326 |
| ND-B22 | 14563.44 | -0.33145 | 0.431053 | -0.76893 | 0.441936 | 0.856326 |
| CG10165 | 851.7026 | -1.23498 | 1.606306 | -0.76883 | 0.441992 | 0.856326 |
| CD98hc | 12783.84 | -0.50429 | 0.657912 | -0.7665 | 0.44338 | 0.856326 |
| CG11986 | 435.8687 | 1.78705 | 2.325882 | 0.768332 | 0.44229 | 0.856326 |
| pont | 145.5152 | -2.23338 | 2.910801 | -0.76727 | 0.44292 | 0.856326 |
| BRWD3 | 724.0637 | -1.20064 | 1.557834 | -0.77071 | 0.440876 | 0.856326 |
| Mccc1 | 284.0727 | -1.40294 | 1.824257 | -0.76905 | 0.441865 | 0.856326 |
| DCTN2-p5( | 1404.762 | -0.72225 | 0.939571 | -0.7687 | 0.442073 | 0.856326 |
| Pgant3 | 226.4621 | -2.00637 | 2.611704 | -0.76822 | 0.442355 | 0.856326 |
| CG11360 | 12789.73 | 0.380783 | 0.496575 | 0.766817 | 0.44319 | 0.856326 |
| Sox102F | 5030.869 | 0.433602 | 0.565194 | 0.767174 | 0.442978 | 0.856326 |
| AnxB10 | 4726.788 | -0.44471 | 0.579148 | -0.76787 | 0.442564 | 0.856326 |
| CG2082 | 10699.26 | 0.383687 | 0.498724 | 0.769338 | 0.441693 | 0.856326 |
| shep | 46925.77 | -0.30457 | 0.397318 | -0.76656 | 0.443346 | 0.856326 |
| mt:srRNA | 7568.816 | -0.39346 | 0.511228 | -0.76964 | 0.441512 | 0.856326 |

| | | | | | | |
|---|---|---|---|---|---|---|
| snoRNA:M | 109.72 | -2.77077 | 3.615794 | -0.7663 | 0.4435 | 0.856326 |
| kra | 13895.17 | 0.3303 | 0.431767 | 0.764997 | 0.444274 | 0.856326 |
| Bet3 | 108.9106 | 1.930166 | 2.509131 | 0.769257 | 0.441741 | 0.856326 |
| tinc | 1966.717 | 0.583824 | 0.762829 | 0.765342 | 0.444068 | 0.856326 |
| CG42788 | 3086.033 | -0.50003 | 0.651886 | -0.76705 | 0.44305 | 0.856326 |
| Myc | 2396.032 | -0.70946 | 0.923075 | -0.76858 | 0.442144 | 0.856326 |
| lncRNA:CR | 86.12334 | 2.694643 | 3.518375 | 0.765877 | 0.443749 | 0.856326 |
| Glo1 | 375.1299 | -1.10228 | 1.436852 | -0.76715 | 0.442991 | 0.856326 |
| mge | 7362.374 | -0.42609 | 0.554158 | -0.7689 | 0.441955 | 0.856326 |
| CG9372 | 73.14625 | 3.003626 | 3.911952 | 0.767808 | 0.442602 | 0.856326 |
| not | 745.4318 | -1.08283 | 1.413047 | -0.76631 | 0.443494 | 0.856326 |
| tant | 1313.444 | 0.948507 | 1.239595 | 0.765174 | 0.444168 | 0.856326 |
| l(3)psg2 | 1387.817 | 0.70423 | 0.919559 | 0.765835 | 0.443775 | 0.856326 |
| Mgstl | 328.6454 | 1.217464 | 1.582276 | 0.769439 | 0.441633 | 0.856326 |
| S | 11480.36 | -0.39179 | 0.509325 | -0.76923 | 0.441757 | 0.856326 |
| IntS3 | 2389.703 | 0.567322 | 0.741912 | 0.764676 | 0.444465 | 0.856488 |
| List | 33205.6 | -0.42266 | 0.553188 | -0.76404 | 0.444846 | 0.856811 |
| CG14100 | 277.1925 | 1.930111 | 2.526159 | 0.76405 | 0.444838 | 0.856811 |
| Wwox | 1058.06 | 0.804654 | 1.055369 | 0.762438 | 0.445798 | 0.856819 |
| GstD9 | 13412.42 | 0.418291 | 0.547616 | 0.763839 | 0.444963 | 0.856819 |
| CG18549 | 2528.518 | -1.04503 | 1.37049 | -0.76252 | 0.445749 | 0.856819 |
| Ctl2 | 3383.558 | -0.47275 | 0.620206 | -0.76224 | 0.445917 | 0.856819 |
| CG1774 | 202.9842 | -1.68415 | 2.207889 | -0.76279 | 0.445589 | 0.856819 |
| ttk | 1935.689 | -0.67934 | 0.891182 | -0.76229 | 0.445887 | 0.856819 |
| CG30344 | 1792.498 | -0.67173 | 0.880699 | -0.76273 | 0.445626 | 0.856819 |
| CG11892 | 1014.426 | -1.28389 | 1.68142 | -0.76358 | 0.44512 | 0.856819 |
| CG11555 | 311.8862 | 1.703038 | 2.233513 | 0.762493 | 0.445766 | 0.856819 |
| Acbp2 | 6262.026 | -0.43386 | 0.56838 | -0.76333 | 0.445265 | 0.856819 |
| His3.3B | 5039.948 | 0.459981 | 0.604016 | 0.761538 | 0.446336 | 0.857021 |
| Utx | 3277.769 | -0.56425 | 0.741084 | -0.76139 | 0.446424 | 0.857021 |
| CG44088 | 321.9156 | 1.608226 | 2.111732 | 0.761567 | 0.446318 | 0.857021 |
| nudC | 198.2298 | -1.83762 | 2.413647 | -0.76135 | 0.44645 | 0.857021 |
| CG32812 | 66.17028 | 2.977038 | 3.918647 | 0.759711 | 0.447428 | 0.857066 |
| nAChRalph | 14940.72 | 0.342667 | 0.450596 | 0.760475 | 0.446971 | 0.857066 |
| CG5674 | 2215.731 | 0.601302 | 0.791812 | 0.7594 | 0.447613 | 0.857066 |
| euc | 5615.372 | 0.404113 | 0.532177 | 0.759358 | 0.447638 | 0.857066 |
| FLASH | 72.55697 | 2.97771 | 3.918498 | 0.759911 | 0.447308 | 0.857066 |
| CG13344 | 1279.868 | -0.80706 | 1.060966 | -0.76069 | 0.446843 | 0.857066 |
| lncRNA:CR | 65.54208 | 2.976964 | 3.918663 | 0.759689 | 0.447441 | 0.857066 |
| CG14971 | 1753.355 | -0.61188 | 0.805797 | -0.75934 | 0.447648 | 0.857066 |
| VhaM9.7-c | 847.5327 | -1.28616 | 1.691139 | -0.76053 | 0.446937 | 0.857066 |
| CG2202 | 602.7867 | -1.03323 | 1.358351 | -0.76065 | 0.446867 | 0.857066 |
| Stt3A | 2900.641 | -0.6529 | 0.858925 | -0.76013 | 0.447174 | 0.857066 |
| Got2 | 6655.511 | -0.38053 | 0.501802 | -0.75834 | 0.448249 | 0.857722 |
| Hsp60A | 8621.746 | -0.3894 | 0.513559 | -0.75823 | 0.448311 | 0.857722 |
| CG15916 | 2481.612 | 0.604742 | 0.797205 | 0.758579 | 0.448105 | 0.857722 |
| SLIRP2 | 184.4511 | 1.807131 | 2.384424 | 0.75789 | 0.448517 | 0.857796 |
| CG9813 | 7422.328 | 0.515453 | 0.680185 | 0.757812 | 0.448563 | 0.857796 |
| Mnn1 | 7329.654 | 0.400904 | 0.529405 | 0.757273 | 0.448886 | 0.858209 |
| NUCB1 | 6686.304 | -0.43784 | 0.578363 | -0.75703 | 0.44903 | 0.858279 |

| | | | | | | |
|---|---|---|---|---|---|---|
| Wnt6 | 283.9176 | 1.235077 | 1.633238 | 0.756213 | 0.449521 | 0.858646 |
| Sgt | 2185.705 | 0.583534 | 0.772118 | 0.755758 | 0.449794 | 0.858646 |
| wge | 4675.344 | -0.48915 | 0.64707 | -0.75594 | 0.449683 | 0.858646 |
| CG11155 | 21241.59 | 0.360421 | 0.476973 | 0.755642 | 0.449864 | 0.858646 |
| PIG-Q | 1475.541 | 1.077593 | 1.425913 | 0.755722 | 0.449816 | 0.858646 |
| CG9231 | 2005.933 | -0.5636 | 0.745028 | -0.75648 | 0.449361 | 0.858646 |
| ApepP | 1383.067 | -0.94122 | 1.24619 | -0.75528 | 0.450081 | 0.858856 |
| RpL24 | 13830.37 | 0.344043 | 0.456104 | 0.754308 | 0.450665 | 0.859595 |
| lili | 1748.725 | 0.658499 | 0.873189 | 0.754131 | 0.450771 | 0.859595 |
| CG10660 | 550.7946 | -1.59767 | 2.118643 | -0.7541 | 0.45079 | 0.859595 |
| CG17724 | 1597.734 | 0.68481 | 0.90846 | 0.753814 | 0.450961 | 0.859688 |
| CG1463 | 693.0889 | 0.85841 | 1.138985 | 0.753662 | 0.451052 | 0.859688 |
| MCTS1 | 2062.717 | -0.63313 | 0.840449 | -0.75332 | 0.451257 | 0.859874 |
| Gel | 5409.385 | -0.53871 | 0.71537 | -0.75305 | 0.451418 | 0.859932 |
| CG8204 | 285.2391 | 1.458799 | 1.937996 | 0.752736 | 0.451609 | 0.859932 |
| kug | 4217.394 | 0.480077 | 0.637666 | 0.752867 | 0.45153 | 0.859932 |
| kn | 471.8443 | 1.271032 | 1.689842 | 0.752161 | 0.451954 | 0.860386 |
| Lrpprc2 | 1018.37 | -0.87788 | 1.167691 | -0.75181 | 0.452167 | 0.860587 |
| MESR6 | 173.7111 | -1.70275 | 2.265451 | -0.75161 | 0.452283 | 0.860603 |
| CCT8 | 1211.117 | 0.803707 | 1.070266 | 0.750942 | 0.452688 | 0.860761 |
| CG11655 | 576.7238 | 1.760331 | 2.344144 | 0.750948 | 0.452684 | 0.860761 |
| aop | 4777.415 | -0.41557 | 0.553283 | -0.7511 | 0.452592 | 0.860761 |
| mt:CoII | 338473.1 | 0.266561 | 0.355175 | 0.750507 | 0.452949 | 0.860851 |
| lncRNA:CR | 66.23663 | 2.575387 | 3.430823 | 0.750661 | 0.452856 | 0.860851 |
| wrapper | 2443.803 | 0.703319 | 0.937901 | 0.749886 | 0.453324 | 0.861358 |
| w | 3525.339 | 0.586216 | 0.785129 | 0.74665 | 0.455275 | 0.861636 |
| Stip1 | 1629.666 | -0.60236 | 0.806396 | -0.74698 | 0.455074 | 0.861636 |
| BNIP3 | 3128.922 | -0.48784 | 0.65366 | -0.74632 | 0.455473 | 0.861636 |
| Lam | 1287.634 | 0.863475 | 1.154233 | 0.748094 | 0.454403 | 0.861636 |
| CG31635 | 6571.184 | -0.44805 | 0.601334 | -0.74509 | 0.456219 | 0.861636 |
| CG7154 | 640.8903 | 0.905384 | 1.213976 | 0.7458 | 0.455788 | 0.861636 |
| Arpc1 | 1064.809 | 0.899395 | 1.207156 | 0.745052 | 0.45624 | 0.861636 |
| Sur-8 | 3551.972 | 0.489065 | 0.654417 | 0.747329 | 0.454865 | 0.861636 |
| CG31475 | 1269.029 | -0.85548 | 1.14512 | -0.74706 | 0.455024 | 0.861636 |
| CR13656 | 1221.684 | 1.115469 | 1.496921 | 0.745176 | 0.456165 | 0.861636 |
| Picot | 10432.54 | -0.37033 | 0.495583 | -0.74726 | 0.454909 | 0.861636 |
| CG7737 | 853.5475 | -0.92856 | 1.246073 | -0.74519 | 0.45616 | 0.861636 |
| CG2269 | 17958.95 | 0.312639 | 0.419588 | 0.745109 | 0.456206 | 0.861636 |
| Tsp42Ef | 1440.7 | 0.771456 | 1.030261 | 0.748797 | 0.45398 | 0.861636 |
| CG32017 | 11486.02 | 0.342428 | 0.459188 | 0.745724 | 0.455834 | 0.861636 |
| CG11594 | 120.9379 | 2.540866 | 3.399084 | 0.747515 | 0.454753 | 0.861636 |
| CG11590 | 577.0832 | -1.02813 | 1.372101 | -0.74931 | 0.453669 | 0.861636 |
| Nelf-A | 2781.051 | 0.522072 | 0.697149 | 0.748867 | 0.453938 | 0.861636 |
| Chpf | 1359.223 | -0.80584 | 1.076761 | -0.74839 | 0.454226 | 0.861636 |
| Ote | 192.9755 | -1.97619 | 2.65255 | -0.74502 | 0.456261 | 0.861636 |
| CG32281 | 1150.381 | 1.081512 | 1.448709 | 0.746535 | 0.455345 | 0.861636 |
| Ulp1 | 2394.4 | -0.58953 | 0.788407 | -0.74775 | 0.45461 | 0.861636 |
| dpr10 | 2449.703 | 0.540259 | 0.722187 | 0.748087 | 0.454408 | 0.861636 |
| Rdl | 40565.36 | 0.288038 | 0.385092 | 0.747972 | 0.454477 | 0.861636 |
| ImpE1 | 1759.076 | 0.673971 | 0.902725 | 0.746597 | 0.455307 | 0.861636 |

| | | | | | | |
|---|---|---|---|---|---|---|
| RpL18 | 11633.61 | -0.39083 | 0.521836 | -0.74896 | 0.453884 | 0.861636 |
| CG13375 | 2018.935 | -0.61575 | 0.828122 | -0.74355 | 0.457151 | 0.861975 |
| Saf6 | 905.7584 | 1.415608 | 1.90455 | 0.743277 | 0.457314 | 0.861975 |
| Ubc6 | 9179.939 | -0.38087 | 0.51396 | -0.74106 | 0.45866 | 0.861975 |
| Mal-B2 | 428.7229 | -1.24004 | 1.675724 | -0.74 | 0.459297 | 0.861975 |
| vls | 397.9759 | 1.487667 | 2.000996 | 0.743463 | 0.457201 | 0.861975 |
| CG10068 | 821.1857 | -0.99931 | 1.346153 | -0.74234 | 0.457879 | 0.861975 |
| CG11722 | 1057.056 | 0.760069 | 1.025077 | 0.741475 | 0.458405 | 0.861975 |
| Leash | 860.1093 | -1.43585 | 1.928679 | -0.74447 | 0.456591 | 0.861975 |
| PK2-R1 | 222.6721 | 1.450718 | 1.960065 | 0.740138 | 0.459216 | 0.861975 |
| CG14322 | 1016.098 | 1.304609 | 1.755286 | 0.743246 | 0.457333 | 0.861975 |
| CG14285 | 72.4917 | 2.559092 | 3.456956 | 0.740273 | 0.459134 | 0.861975 |
| Synd | 3226.087 | -0.51714 | 0.697004 | -0.74195 | 0.458118 | 0.861975 |
| C15 | 90.10606 | 2.814271 | 3.8107 | 0.738518 | 0.4602 | 0.861975 |
| wda | 429.0396 | -1.37843 | 1.864764 | -0.7392 | 0.459788 | 0.861975 |
| CG2218 | 1367.267 | 0.643519 | 0.866791 | 0.742415 | 0.457836 | 0.861975 |
| Tif-IA | 2181.841 | 0.568916 | 0.768322 | 0.740465 | 0.459018 | 0.861975 |
| CG32708 | 568.1927 | 1.262779 | 1.698897 | 0.743293 | 0.457304 | 0.861975 |
| CG13204 | 721.5786 | 1.340348 | 1.812729 | 0.739409 | 0.459659 | 0.861975 |
| Rs1 | 92.31406 | -2.15237 | 2.908839 | -0.73994 | 0.459335 | 0.861975 |
| ChAT | 9468.638 | 0.379369 | 0.512706 | 0.739935 | 0.459339 | 0.861975 |
| Deaf1 | 3346.921 | -0.51122 | 0.691034 | -0.7398 | 0.459423 | 0.861975 |
| Octbeta2R | 4968.37 | 0.459049 | 0.619988 | 0.740416 | 0.459048 | 0.861975 |
| spok | 289.6518 | 1.40344 | 1.890121 | 0.742513 | 0.457776 | 0.861975 |
| CG42524 | 578.2088 | 0.929637 | 1.254409 | 0.741095 | 0.458636 | 0.861975 |
| thoc7 | 1605.448 | 0.633315 | 0.857391 | 0.738653 | 0.460117 | 0.861975 |
| how | 1008.447 | 0.747868 | 1.006747 | 0.742856 | 0.457569 | 0.861975 |
| GckIII | 1947.838 | -0.57542 | 0.778989 | -0.73867 | 0.460108 | 0.861975 |
| Sec10 | 1249.101 | -0.98809 | 1.337279 | -0.73888 | 0.459977 | 0.861975 |
| btsz | 895.4318 | 0.93545 | 1.259068 | 0.74297 | 0.4575 | 0.861975 |
| lncRNA:CR | 195.6963 | -1.64596 | 2.224336 | -0.73998 | 0.459314 | 0.861975 |
| LIMK1 | 1661.424 | -0.76183 | 1.026613 | -0.74208 | 0.458038 | 0.861975 |
| Mpi | 984.4733 | -0.72755 | 0.98157 | -0.74121 | 0.458563 | 0.861975 |
| CG11134 | 579.0939 | 0.928536 | 1.255785 | 0.739407 | 0.45966 | 0.861975 |
| hid | 254.5332 | -1.32064 | 1.77482 | -0.7441 | 0.456816 | 0.861975 |
| Klc | 2617.3 | -0.66836 | 0.904069 | -0.73928 | 0.459739 | 0.861975 |
| CG9515 | 73.17759 | 2.818852 | 3.821699 | 0.737591 | 0.460763 | 0.862828 |
| CG1172 | 951.4316 | -1.13907 | 1.54474 | -0.73739 | 0.460887 | 0.862859 |
| Pzl | 203.3478 | 1.716711 | 2.328731 | 0.737188 | 0.461008 | 0.862885 |
| CG9667 | 167.4295 | 1.596846 | 2.167644 | 0.736674 | 0.461321 | 0.863269 |
| CG4592 | 500.5087 | 1.419294 | 1.927265 | 0.736429 | 0.461469 | 0.863296 |
| zyd | 13107.81 | -0.45057 | 0.611938 | -0.7363 | 0.46155 | 0.863296 |
| Cand1 | 3345.244 | 0.487261 | 0.663374 | 0.734519 | 0.462632 | 0.864201 |
| CG3744 | 753.2376 | 0.908066 | 1.236277 | 0.734517 | 0.462634 | 0.864201 |
| sn | 930.5069 | -0.73527 | 1.000894 | -0.73461 | 0.462575 | 0.864201 |
| ric8a | 364.2918 | -1.4908 | 2.029845 | -0.73444 | 0.46268 | 0.864201 |
| MFS12 | 255.8831 | -1.68345 | 2.291354 | -0.7347 | 0.462523 | 0.864201 |
| asRNA:CR4 | 138.1777 | -2.41974 | 3.293533 | -0.73469 | 0.462526 | 0.864201 |
| CG1358 | 26972.94 | -0.31103 | 0.423783 | -0.73393 | 0.462992 | 0.864583 |
| Rpn5 | 2253.914 | -0.56074 | 0.76454 | -0.73344 | 0.463293 | 0.864943 |

| | | | | | | |
|---|---|---|---|---|---|---|
| PTP-ER | 1568.428 | -0.90259 | 1.231948 | -0.73265 | 0.463769 | 0.86543 |
| RecQ5 | 1096.498 | -0.8065 | 1.100591 | -0.73279 | 0.463688 | 0.86543 |
| CG15772 | 735.4628 | 1.276034 | 1.752237 | 0.728232 | 0.466472 | 0.865818 |
| Ptth | 238.3854 | -2.04866 | 2.806871 | -0.72987 | 0.465467 | 0.865818 |
| isoQC | 535.8507 | -1.05741 | 1.453761 | -0.72736 | 0.467006 | 0.865818 |
| CG11241 | 345.5081 | 1.501731 | 2.054436 | 0.73097 | 0.464798 | 0.865818 |
| Rat1 | 156.1233 | 1.523829 | 2.095178 | 0.727303 | 0.46704 | 0.865818 |
| Megf8 | 829.4812 | -1.17163 | 1.602919 | -0.73093 | 0.464819 | 0.865818 |
| strat | 432.1612 | 1.249166 | 1.709917 | 0.730542 | 0.465059 | 0.865818 |
| AdipoR | 6508.612 | 0.395774 | 0.541665 | 0.730662 | 0.464985 | 0.865818 |
| CG7006 | 251.5007 | 1.666505 | 2.279104 | 0.731211 | 0.464651 | 0.865818 |
| CG31106 | 333.5251 | -2.0464 | 2.807533 | -0.7289 | 0.466064 | 0.865818 |
| Bap170 | 351.1162 | -1.13452 | 1.554696 | -0.72974 | 0.465551 | 0.865818 |
| Cdc7 | 2269.637 | 0.552184 | 0.755051 | 0.73132 | 0.464584 | 0.865818 |
| CG18262 | 207.657 | -1.82754 | 2.51137 | -0.72771 | 0.466793 | 0.865818 |
| CG1440 | 1127.306 | 0.806335 | 1.104527 | 0.730027 | 0.465373 | 0.865818 |
| CG1789 | 936.4295 | 0.7557 | 1.039035 | 0.727309 | 0.467036 | 0.865818 |
| CG30460 | 405.3079 | 1.029863 | 1.413261 | 0.728714 | 0.466176 | 0.865818 |
| beta3GalT | 377.6289 | 1.653008 | 2.25933 | 0.731636 | 0.464391 | 0.865818 |
| Taf3 | 15219.38 | -0.41879 | 0.573559 | -0.73016 | 0.465295 | 0.865818 |
| arm | 10203.15 | -0.37932 | 0.521556 | -0.72728 | 0.467054 | 0.865818 |
| alpha-Cat | 10542.24 | 0.357139 | 0.489571 | 0.729493 | 0.4657 | 0.865818 |
| scramb1 | 4457.623 | 0.459959 | 0.630939 | 0.729008 | 0.465997 | 0.865818 |
| mgl | 19648.73 | -0.34204 | 0.469619 | -0.72833 | 0.466414 | 0.865818 |
| Tasp1 | 447.7153 | 1.375545 | 1.883757 | 0.730214 | 0.46526 | 0.865818 |
| CG44158 | 109.6404 | -2.09733 | 2.880745 | -0.72805 | 0.466581 | 0.865818 |
| p24-1 | 1479.993 | 1.184512 | 1.627991 | 0.727591 | 0.466864 | 0.865818 |
| NFAT | 9670.345 | -0.33933 | 0.466279 | -0.72774 | 0.466771 | 0.865818 |
| Clc | 3763.796 | 0.45176 | 0.619764 | 0.728922 | 0.466049 | 0.865818 |
| roq | 2200.973 | 0.5763 | 0.790112 | 0.72939 | 0.465763 | 0.865818 |
| Mad | 1700.614 | -0.69912 | 0.961398 | -0.7272 | 0.467106 | 0.865818 |
| cype | 21816.36 | 0.331367 | 0.456138 | 0.726461 | 0.467556 | 0.866292 |
| CG10527 | 1713.825 | -0.60569 | 0.834048 | -0.7262 | 0.467713 | 0.866292 |
| gprs | 4586.849 | 0.460343 | 0.633808 | 0.726313 | 0.467647 | 0.866292 |
| CG13912 | 81.16521 | -2.78554 | 3.836434 | -0.72607 | 0.467793 | 0.866292 |
| sds22 | 692.6786 | 0.826424 | 1.139022 | 0.725556 | 0.468111 | 0.86668 |
| CG9016 | 198.5481 | 1.718839 | 2.372388 | 0.724518 | 0.468748 | 0.867059 |
| Kank | 293.1902 | -1.18582 | 1.636466 | -0.72462 | 0.468684 | 0.867059 |
| lncRNA:CR | 309.5646 | 1.671672 | 2.307197 | 0.724547 | 0.46873 | 0.867059 |
| KFase | 551.9931 | -1.20694 | 1.665065 | -0.72486 | 0.468539 | 0.867059 |
| CG31140 | 15416.83 | -0.31071 | 0.429274 | -0.72379 | 0.469192 | 0.867681 |
| CG13654 | 827.4386 | 1.199314 | 1.658326 | 0.723208 | 0.469552 | 0.868147 |
| kek1 | 3516.082 | 0.506856 | 0.701245 | 0.722794 | 0.469807 | 0.868417 |
| Ufd1-like | 1250.311 | -0.76733 | 1.062258 | -0.72236 | 0.470075 | 0.868714 |
| su(f) | 2180.149 | 0.525993 | 0.728528 | 0.721995 | 0.470298 | 0.868925 |
| sta | 13672.84 | 0.310531 | 0.431267 | 0.720042 | 0.471499 | 0.869056 |
| CG12236 | 2467.654 | 0.775894 | 1.076719 | 0.72061 | 0.47115 | 0.869056 |
| Sfxn1-3 | 909.1062 | -1.13673 | 1.579693 | -0.71959 | 0.471777 | 0.869056 |
| srl | 114.1352 | -2.40913 | 3.346028 | -0.72 | 0.471527 | 0.869056 |
| mRpL1 | 1287.344 | 0.693352 | 0.962694 | 0.72022 | 0.47139 | 0.869056 |

| | | | | | | |
|---|---|---|---|---|---|---|
| Coq2 | 324.0969 | 1.133022 | 1.575661 | 0.719077 | 0.472093 | 0.869056 |
| CG9346 | 334.2554 | 1.251308 | 1.739958 | 0.71916 | 0.472042 | 0.869056 |
| CG4744 | 301.6825 | 2.056866 | 2.855447 | 0.720331 | 0.471321 | 0.869056 |
| CG33774 | 502.2453 | 0.930122 | 1.293518 | 0.719064 | 0.472101 | 0.869056 |
| CG44002 | 190.847 | -2.39399 | 3.324807 | -0.72004 | 0.471501 | 0.869056 |
| ghi | 4765.186 | 0.523551 | 0.725691 | 0.721452 | 0.470632 | 0.869056 |
| Nadsyn | 1444.45 | 0.733974 | 1.02058 | 0.719174 | 0.472034 | 0.869056 |
| TrpRS-m | 2422.768 | 0.650042 | 0.903462 | 0.719501 | 0.471832 | 0.869056 |
| CG32032 | 5507.072 | -0.46568 | 0.647296 | -0.71943 | 0.471876 | 0.869056 |
| CG7506 | 1592.777 | -0.63827 | 0.884888 | -0.7213 | 0.470724 | 0.869056 |
| MCU | 5212.421 | 0.389647 | 0.540491 | 0.720914 | 0.470962 | 0.869056 |
| TkR86C | 332.7744 | -1.61189 | 2.24347 | -0.71848 | 0.472461 | 0.869334 |
| CG42709 | 5229.074 | 0.392902 | 0.54686 | 0.718468 | 0.472469 | 0.869334 |
| CG17266 | 196.8683 | 2.076025 | 2.890519 | 0.718219 | 0.472622 | 0.869339 |
| Srp54 | 1482.385 | -0.61734 | 0.859796 | -0.71801 | 0.472751 | 0.869339 |
| CG9164 | 3113.807 | 0.884972 | 1.232661 | 0.717937 | 0.472796 | 0.869339 |
| Eaat1 | 17212.89 | -0.34021 | 0.474152 | -0.71751 | 0.473059 | 0.869424 |
| JIL-1 | 2978.579 | 0.540989 | 0.753813 | 0.717669 | 0.472961 | 0.869424 |
| olf413 | 4741.073 | 0.404275 | 0.564588 | 0.716054 | 0.473958 | 0.870115 |
| l(3)73Ah | 998.7254 | -1.0645 | 1.486412 | -0.71615 | 0.473896 | 0.870115 |
| Snmp1 | 171.1881 | -2.00774 | 2.803188 | -0.71624 | 0.473846 | 0.870115 |
| Ih | 28262.11 | 0.280089 | 0.391108 | 0.716144 | 0.473902 | 0.870115 |
| ZC3H3 | 1124.442 | -0.75462 | 1.053901 | -0.71602 | 0.473977 | 0.870115 |
| CG16711 | 1532.531 | -0.60294 | 0.842306 | -0.71582 | 0.474101 | 0.870142 |
| CG8078 | 456.8956 | 1.201679 | 1.679345 | 0.715564 | 0.474261 | 0.870237 |
| Vinc | 81.8558 | 2.351565 | 3.291413 | 0.714455 | 0.474946 | 0.870303 |
| Dlg5 | 762.336 | -1.07492 | 1.503155 | -0.71511 | 0.47454 | 0.870303 |
| Mondo | 2102.585 | 0.594147 | 0.831863 | 0.714237 | 0.475081 | 0.870303 |
| Bub3 | 491.2479 | -1.00919 | 1.412761 | -0.71434 | 0.475016 | 0.870303 |
| RpL21 | 15608.68 | -0.31698 | 0.443885 | -0.7141 | 0.475164 | 0.870303 |
| rdx | 5867.545 | -0.41109 | 0.575059 | -0.71487 | 0.474689 | 0.870303 |
| lncRNA:CR | 337.3348 | 1.335376 | 1.868534 | 0.714665 | 0.474816 | 0.870303 |
| CG10960 | 13425.29 | -0.33669 | 0.470736 | -0.71525 | 0.474455 | 0.870303 |
| CG2930 | 667.9567 | 0.882595 | 1.243356 | 0.70985 | 0.477797 | 0.870445 |
| CG9586 | 611.4483 | 1.256013 | 1.765431 | 0.711448 | 0.476806 | 0.870445 |
| CG10561 | 1197.625 | 0.679956 | 0.955765 | 0.711426 | 0.47682 | 0.870445 |
| TotF | 62.35554 | 2.776096 | 3.916444 | 0.708831 | 0.47843 | 0.870445 |
| CG1943 | 2618.382 | 0.525783 | 0.738704 | 0.711764 | 0.476611 | 0.870445 |
| ATP6AP2 | 16297.86 | -0.37337 | 0.52424 | -0.71221 | 0.476333 | 0.870445 |
| Best1 | 1540.894 | 0.710753 | 0.997344 | 0.712646 | 0.476065 | 0.870445 |
| Ns1 | 1007.388 | 0.914158 | 1.288004 | 0.709748 | 0.47786 | 0.870445 |
| CG14329 | 110.4822 | 2.778428 | 3.91592 | 0.709521 | 0.478001 | 0.870445 |
| CG14309 | 81.92009 | 2.777373 | 3.916157 | 0.709209 | 0.478195 | 0.870445 |
| CG4000 | 247.1478 | 1.978824 | 2.790699 | 0.709078 | 0.478276 | 0.870445 |
| CG13841 | 363.3178 | -1.13023 | 1.590644 | -0.71055 | 0.477364 | 0.870445 |
| CG13829 | 532.2764 | 1.757596 | 2.469564 | 0.711703 | 0.476649 | 0.870445 |
| mRpS18C | 1132.804 | 0.791393 | 1.111281 | 0.712145 | 0.476375 | 0.870445 |
| CG1544 | 91.78695 | 2.423896 | 3.397824 | 0.713367 | 0.475619 | 0.870445 |
| CG13868 | 12198.44 | 0.38803 | 0.544474 | 0.71267 | 0.47605 | 0.870445 |
| UQCR-6.4 | 7725.56 | 0.351239 | 0.494482 | 0.710317 | 0.477507 | 0.870445 |

| | | | | | | |
|---|---|---|---|---|---|---|
| CG18870 | 926.1136 | -1.31042 | 1.84441 | -0.71048 | 0.477406 | 0.870445 |
| Cyp6t3 | 108.1805 | 2.778364 | 3.915934 | 0.709502 | 0.478013 | 0.870445 |
| kel | 1580.363 | -0.59806 | 0.841033 | -0.7111 | 0.47702 | 0.870445 |
| CG15394 | 69.46992 | 2.776642 | 3.916321 | 0.708992 | 0.478329 | 0.870445 |
| vlc | 2110.33 | 0.679055 | 0.953782 | 0.71196 | 0.476489 | 0.870445 |
| snRNP-U1- | 870.2884 | -1.22059 | 1.717934 | -0.7105 | 0.477397 | 0.870445 |
| RanBPM | 1544.641 | -0.77318 | 1.09094 | -0.70872 | 0.478496 | 0.870445 |
| CG3611 | 95.83498 | 2.777965 | 3.916024 | 0.709384 | 0.478086 | 0.870445 |
| lncRNA:CR | 75.74732 | 2.777041 | 3.916231 | 0.709111 | 0.478256 | 0.870445 |
| lncRNA:CR | 81.71084 | 2.777362 | 3.916159 | 0.709206 | 0.478197 | 0.870445 |
| hop | 129.046 | -2.08353 | 2.927827 | -0.71163 | 0.476694 | 0.870445 |
| mew | 263.7246 | -1.25723 | 1.769808 | -0.71038 | 0.47747 | 0.870445 |
| Atg18a | 2698.119 | -0.57963 | 0.815965 | -0.71036 | 0.477479 | 0.870445 |
| CG10333 | 606.2395 | 0.978401 | 1.381118 | 0.708413 | 0.478689 | 0.8706 |
| Bka | 863.5811 | 0.899143 | 1.271502 | 0.70715 | 0.479473 | 0.870682 |
| CG34155 | 737.996 | 0.89036 | 1.258279 | 0.707602 | 0.479192 | 0.870682 |
| lap | 10568.12 | 0.320379 | 0.452868 | 0.707445 | 0.47929 | 0.870682 |
| lqfR | 2162.961 | -0.60038 | 0.84899 | -0.70717 | 0.47946 | 0.870682 |
| CG13894 | 918.2414 | -1.18778 | 1.679369 | -0.70728 | 0.479395 | 0.870682 |
| lncRNA:CR | 213.0656 | -2.02435 | 2.862156 | -0.70728 | 0.479391 | 0.870682 |
| Ork1 | 7506.966 | -0.37146 | 0.525417 | -0.70699 | 0.479574 | 0.870682 |
| Mgat4a | 3564.169 | -0.4818 | 0.681526 | -0.70694 | 0.479602 | 0.870682 |
| CG12155 | 867.7792 | 0.728017 | 1.030976 | 0.706143 | 0.480099 | 0.871388 |
| CG9123 | 486.3172 | -1.23962 | 1.756416 | -0.70577 | 0.480333 | 0.871615 |
| NP15.6 | 7823.966 | -0.33986 | 0.482184 | -0.70484 | 0.480909 | 0.871986 |
| krz | 2035.097 | 0.593706 | 0.842025 | 0.705093 | 0.480752 | 0.871986 |
| Spt5 | 2549.336 | -0.48409 | 0.6869 | -0.70474 | 0.480972 | 0.871986 |
| RhoGDI | 4226.278 | -0.43624 | 0.619006 | -0.70474 | 0.48097 | 0.871986 |
| santa-mari | 2022.796 | 0.598092 | 0.849148 | 0.704343 | 0.481219 | 0.872113 |
| CCT7 | 1233.138 | -0.92237 | 1.309991 | -0.7041 | 0.481368 | 0.872113 |
| AsnRS | 918.6662 | 0.790754 | 1.12282 | 0.704257 | 0.481272 | 0.872113 |
| CG17075 | 1300.62 | 0.680115 | 0.96699 | 0.703332 | 0.481849 | 0.872787 |
| RpL38 | 25366.78 | 0.291965 | 0.415235 | 0.703132 | 0.481973 | 0.872815 |
| yip2 | 2295.467 | -0.58865 | 0.837778 | -0.70263 | 0.482287 | 0.873167 |
| jing | 4196.093 | 0.450905 | 0.642043 | 0.702297 | 0.482494 | 0.873167 |
| CG13667 | 639.8061 | 1.368221 | 1.947838 | 0.702431 | 0.482411 | 0.873167 |
| ttm50 | 1081.345 | -0.70808 | 1.008838 | -0.70188 | 0.482755 | 0.873442 |
| CG4281 | 1168.981 | 0.758288 | 1.081024 | 0.701453 | 0.48302 | 0.873528 |
| eIF3h | 3359.505 | -0.46476 | 0.662423 | -0.70161 | 0.482923 | 0.873528 |
| mRpL33 | 912.629 | 0.737065 | 1.052829 | 0.70008 | 0.483877 | 0.874094 |
| CanA1 | 90.63127 | 2.742304 | 3.916307 | 0.700227 | 0.483786 | 0.874094 |
| Rcd5 | 1448.192 | -0.7422 | 1.059537 | -0.7005 | 0.483617 | 0.874094 |
| CG9705 | 906.113 | -0.73967 | 1.056192 | -0.70032 | 0.483729 | 0.874094 |
| Tfb4 | 252.3778 | -1.77879 | 2.539437 | -0.70047 | 0.483636 | 0.874094 |
| CG7519 | 1128.192 | 0.661295 | 0.946193 | 0.698901 | 0.484614 | 0.87416 |
| RpL24-like | 8916.794 | 0.358102 | 0.512905 | 0.698184 | 0.485062 | 0.87416 |
| mor | 338.5906 | 1.34884 | 1.931677 | 0.698274 | 0.485006 | 0.87416 |
| CG17786 | 295.0198 | -1.18696 | 1.701348 | -0.69766 | 0.485389 | 0.87416 |
| Tsp42Ed | 905.318 | -1.47303 | 2.110282 | -0.69803 | 0.48516 | 0.87416 |
| CG3184 | 85.98222 | -2.34897 | 3.362316 | -0.69862 | 0.484792 | 0.87416 |

| | | | | | | |
|---|---|---|---|---|---|---|
| ReepB | 3919.519 | 0.415002 | 0.594678 | 0.69786 | 0.485265 | 0.87416 |
| beta4GalN | 1332.037 | -1.2111 | 1.735258 | -0.69794 | 0.485216 | 0.87416 |
| Mat1 | 529.9489 | -0.98152 | 1.403744 | -0.69921 | 0.484419 | 0.87416 |
| Gdh | 9776.842 | 0.351677 | 0.502536 | 0.699805 | 0.484049 | 0.87416 |
| ktub | 1728.018 | -0.69222 | 0.989646 | -0.69947 | 0.48426 | 0.87416 |
| GlyP | 5489.269 | -0.43841 | 0.628468 | -0.69758 | 0.485439 | 0.87416 |
| Nulp1 | 1437.205 | -1.01737 | 1.456462 | -0.69852 | 0.484851 | 0.87416 |
| CG12795 | 191.7762 | -1.71554 | 2.457523 | -0.69808 | 0.485128 | 0.87416 |
| Pfrx | 1569.982 | 0.689502 | 0.989366 | 0.696913 | 0.485857 | 0.874717 |
| 5-HT7 | 2697.434 | 0.745719 | 1.071155 | 0.696182 | 0.486315 | 0.875223 |
| Sec63 | 1928.16 | -0.54408 | 0.7816 | -0.69612 | 0.486356 | 0.875223 |
| CG16790 | 195.5748 | -1.88808 | 2.71652 | -0.69503 | 0.487033 | 0.876128 |
| Galphaq | 4311.102 | 0.493054 | 0.70957 | 0.694863 | 0.487141 | 0.876128 |
| Imp | 5400.698 | 0.378174 | 0.544299 | 0.694791 | 0.487187 | 0.876128 |
| CG3594 | 666.4484 | 1.209138 | 1.740894 | 0.69455 | 0.487337 | 0.876202 |
| Asap | 3072.738 | 0.48361 | 0.696699 | 0.694145 | 0.487592 | 0.876464 |
| Baldspot | 6693.328 | -0.36227 | 0.522343 | -0.69354 | 0.487971 | 0.876598 |
| lncRNA:CR | 1227.98 | -0.70649 | 1.018286 | -0.6938 | 0.487808 | 0.876598 |
| Helz | 3655.987 | -0.48742 | 0.702836 | -0.6935 | 0.487994 | 0.876598 |
| CG14803 | 112.8897 | -2.09504 | 3.022301 | -0.69319 | 0.488188 | 0.876626 |
| CG5746 | 2260.905 | 0.522557 | 0.754098 | 0.692956 | 0.488337 | 0.876626 |
| ND-24 | 6316.163 | 0.400735 | 0.578275 | 0.692983 | 0.48832 | 0.876626 |
| snoRNA:Ps | 122.838 | -2.27792 | 3.289796 | -0.69242 | 0.488674 | 0.877035 |
| Nepl5 | 11707.23 | -0.39734 | 0.574052 | -0.69217 | 0.488829 | 0.877116 |
| Slik | 2823.688 | -0.49784 | 0.719909 | -0.69153 | 0.489232 | 0.877644 |
| sgg | 8859.262 | 0.353384 | 0.511641 | 0.690687 | 0.489762 | 0.87801 |
| CG14671 | 900.2995 | -1.1111 | 1.609705 | -0.69025 | 0.490035 | 0.87801 |
| VhaPPA1-1 | 11852.06 | -0.36064 | 0.522411 | -0.69033 | 0.489986 | 0.87801 |
| PIG-O | 113.9266 | 2.047494 | 2.968183 | 0.689814 | 0.490311 | 0.87801 |
| Nsun5 | 566.2828 | 1.59338 | 2.308007 | 0.69037 | 0.489961 | 0.87801 |
| Usp10 | 2035.399 | -0.51876 | 0.751867 | -0.68996 | 0.490221 | 0.87801 |
| CG8745 | 16053.44 | -0.34726 | 0.502553 | -0.69099 | 0.489573 | 0.87801 |
| unc-13-4A | 3410.095 | -0.48709 | 0.70578 | -0.69014 | 0.490104 | 0.87801 |
| Scox | 1829.719 | -0.69672 | 1.010568 | -0.68943 | 0.490551 | 0.878242 |
| CG33129 | 5024.269 | 0.391359 | 0.567988 | 0.689026 | 0.490807 | 0.878343 |
| CG18659 | 3220.271 | 0.475326 | 0.689882 | 0.688996 | 0.490826 | 0.878343 |
| Ugt35E2 | 298.2688 | 1.330328 | 1.934123 | 0.68782 | 0.491566 | 0.87899 |
| Ctf4 | 136.6344 | 2.139298 | 3.108678 | 0.68817 | 0.491346 | 0.87899 |
| CG42674 | 2977.239 | 0.481909 | 0.700504 | 0.687946 | 0.491487 | 0.87899 |
| Vps16A | 1212.949 | -1.03854 | 1.510105 | -0.68773 | 0.491625 | 0.87899 |
| CG31109 | 2165.404 | -0.69943 | 1.017326 | -0.68752 | 0.491756 | 0.879028 |
| dl | 2486.689 | -0.51277 | 0.746293 | -0.6871 | 0.492023 | 0.879308 |
| MTPAP | 351.473 | -1.24106 | 1.806941 | -0.68683 | 0.492189 | 0.87941 |
| CG11562 | 437.2516 | 1.219612 | 1.776423 | 0.686555 | 0.492363 | 0.879525 |
| stas | 6163.991 | 0.39859 | 0.580824 | 0.686249 | 0.492556 | 0.879674 |
| pr | 1613.879 | -0.99343 | 1.448213 | -0.68597 | 0.492733 | 0.879795 |
| ab | 6457.452 | -0.34908 | 0.509099 | -0.68567 | 0.492919 | 0.879852 |
| CG1532 | 848.6831 | 0.744417 | 1.085836 | 0.68557 | 0.492984 | 0.879852 |
| Nop17l | 3338.866 | -0.43344 | 0.632412 | -0.68537 | 0.49311 | 0.87988 |
| Cyp9f3 | 506.8592 | -1.50663 | 2.200423 | -0.6847 | 0.493534 | 0.880068 |

| | | | | | | |
|---|---|---|---|---|---|---|
| Coq8 | 279.4568 | -1.51125 | 2.20723 | -0.68468 | 0.493545 | 0.880068 |
| CG32069 | 259.724 | -1.18904 | 1.736256 | -0.68483 | 0.493453 | 0.880068 |
| D2hgdh | 792.4445 | -1.37627 | 2.01582 | -0.68274 | 0.494774 | 0.880128 |
| CG4766 | 174.6119 | 1.897577 | 2.782705 | 0.681918 | 0.495291 | 0.880128 |
| siz | 5973.516 | -0.39279 | 0.57604 | -0.68188 | 0.495315 | 0.880128 |
| MICU3 | 2628.226 | -0.47502 | 0.695089 | -0.6834 | 0.494355 | 0.880128 |
| CG5646 | 277.0234 | -1.6314 | 2.388806 | -0.68293 | 0.494648 | 0.880128 |
| Tango5 | 1779.022 | -0.53203 | 0.778482 | -0.68342 | 0.494344 | 0.880128 |
| Magi | 533.0187 | 1.135385 | 1.665149 | 0.681852 | 0.495333 | 0.880128 |
| Tom7 | 3861.672 | 0.412428 | 0.604214 | 0.682586 | 0.494868 | 0.880128 |
| Dhit | 973.9577 | -0.66856 | 0.980751 | -0.68168 | 0.495442 | 0.880128 |
| CG33695 | 455.6287 | 1.330857 | 1.946543 | 0.683703 | 0.494163 | 0.880128 |
| nuf | 7112.881 | -0.40288 | 0.589849 | -0.68302 | 0.494597 | 0.880128 |
| Pak | 3779.02 | -0.45626 | 0.668862 | -0.68215 | 0.495146 | 0.880128 |
| SRPK | 5156.383 | -0.37077 | 0.543431 | -0.68227 | 0.495066 | 0.880128 |
| CG1622 | 1618.364 | 0.921308 | 1.350858 | 0.682017 | 0.495228 | 0.880128 |
| Fim | 6011.146 | 0.380048 | 0.557474 | 0.681732 | 0.495408 | 0.880128 |
| CG14142 | 217.497 | -1.53371 | 2.242154 | -0.68404 | 0.493953 | 0.880128 |
| Fnta | 429.794 | -1.07213 | 1.568391 | -0.68359 | 0.494236 | 0.880128 |
| CG10514 | 4214.79 | -0.51165 | 0.751064 | -0.68124 | 0.495722 | 0.880154 |
| snoRNA:Ps | 65.76418 | 2.259255 | 3.317497 | 0.681012 | 0.495864 | 0.880154 |
| Mdh1 | 12142.72 | -0.31421 | 0.461141 | -0.68137 | 0.495635 | 0.880154 |
| eIF3l | 5723.569 | 0.389908 | 0.572583 | 0.680963 | 0.495895 | 0.880154 |
| PGAP1 | 558.2778 | 1.402079 | 2.062675 | 0.679738 | 0.49667 | 0.880163 |
| chic | 7390.511 | -0.35797 | 0.526032 | -0.68051 | 0.496181 | 0.880163 |
| SppL | 2781.71 | 0.763161 | 1.121839 | 0.680277 | 0.496329 | 0.880163 |
| HLH4C | 140.9175 | -2.28114 | 3.351829 | -0.68057 | 0.496146 | 0.880163 |
| Cadps | 29351.7 | 0.273884 | 0.403027 | 0.679568 | 0.496778 | 0.880163 |
| mIF3 | 327.3981 | 1.639156 | 2.410654 | 0.679963 | 0.496528 | 0.880163 |
| lncRNA:CR | 115.6679 | -2.40641 | 3.540489 | -0.67968 | 0.496705 | 0.880163 |
| FMRFaR | 172.8542 | -1.89801 | 2.791987 | -0.67981 | 0.496627 | 0.880163 |
| CG10584 | 797.0386 | 1.112457 | 1.640445 | 0.678144 | 0.497681 | 0.881419 |
| CG31211 | 2521.042 | -0.48789 | 0.72044 | -0.67721 | 0.498276 | 0.881419 |
| dpr17 | 3099.482 | 0.457324 | 0.676 | 0.676515 | 0.498714 | 0.881419 |
| CG7518 | 1783.738 | -0.53262 | 0.787874 | -0.67602 | 0.499027 | 0.881419 |
| twf | 605.583 | 1.130573 | 1.672659 | 0.675914 | 0.499095 | 0.881419 |
| Hydr1 | 4749.054 | -0.37303 | 0.551456 | -0.67645 | 0.498758 | 0.881419 |
| Clk | 813.9843 | -1.00642 | 1.486648 | -0.67697 | 0.498423 | 0.881419 |
| spen | 53433.15 | -0.26181 | 0.387218 | -0.67614 | 0.498951 | 0.881419 |
| Gapdh1 | 27711.49 | -0.34117 | 0.503858 | -0.67712 | 0.498329 | 0.881419 |
| side-II | 5039.642 | 0.405926 | 0.599621 | 0.676972 | 0.498424 | 0.881419 |
| Vha68-2 | 2230.218 | -0.63003 | 0.929221 | -0.67802 | 0.49776 | 0.881419 |
| slmb | 3599.668 | -0.52695 | 0.779685 | -0.67585 | 0.499134 | 0.881419 |
| trol | 27066.45 | -0.27892 | 0.412201 | -0.67665 | 0.498629 | 0.881419 |
| CG32243 | 1142.397 | 0.673091 | 0.994525 | 0.676797 | 0.498535 | 0.881419 |
| Srp19 | 1328.081 | 0.693135 | 1.023598 | 0.677156 | 0.498307 | 0.881419 |
| 5-HT1A | 1245.088 | -0.6278 | 0.929208 | -0.67563 | 0.499277 | 0.881478 |
| Rassf | 854.733 | -0.76619 | 1.135238 | -0.67491 | 0.499731 | 0.881891 |
| CG43789 | 678.0457 | -0.86629 | 1.283414 | -0.67499 | 0.499685 | 0.881891 |
| RhoGAP16 | 398.6423 | 1.497334 | 2.220428 | 0.674345 | 0.500092 | 0.882335 |

| | | | | | | |
|---|---|---|---|---|---|---|
| CG14814 | 122.5246 | 2.0063 | 2.977131 | 0.673904 | 0.500372 | 0.882513 |
| zip | 2042.099 | 0.556142 | 0.825332 | 0.67384 | 0.500413 | 0.882513 |
| Gasp | 231.7414 | -2.10952 | 3.132572 | -0.67341 | 0.500684 | 0.882602 |
| Mad1 | 233.0724 | 1.19502 | 1.774219 | 0.673547 | 0.500599 | 0.882602 |
| CCHa2-R | 238.1434 | 1.809941 | 2.689769 | 0.672898 | 0.501012 | 0.882987 |
| CG7800 | 824.7562 | 0.837388 | 1.245967 | 0.672079 | 0.501533 | 0.883076 |
| Dlc90F | 3564.354 | 0.485418 | 0.723125 | 0.671278 | 0.502043 | 0.883076 |
| CG6425 | 429.6763 | 1.402932 | 2.089648 | 0.671373 | 0.501983 | 0.883076 |
| Mgat1 | 442.2526 | 1.186069 | 1.766924 | 0.671262 | 0.502054 | 0.883076 |
| CG3032 | 339.0434 | -2.11743 | 3.15304 | -0.67155 | 0.501868 | 0.883076 |
| Rbpn-5 | 90.93689 | 1.765634 | 2.628639 | 0.671691 | 0.50178 | 0.883076 |
| CG8248 | 3271.454 | 0.846802 | 1.260653 | 0.671717 | 0.501764 | 0.883076 |
| mt:ATPase | 3180.874 | 0.438632 | 0.653611 | 0.67109 | 0.502163 | 0.883076 |
| mbf1 | 4987.675 | -0.39784 | 0.592395 | -0.67157 | 0.501856 | 0.883076 |
| CG3530 | 5421.837 | -0.43456 | 0.646543 | -0.67213 | 0.501501 | 0.883076 |
| CG33217 | 1640.318 | 0.544133 | 0.812969 | 0.669317 | 0.503293 | 0.883397 |
| Rtnl1 | 23718.38 | 0.298002 | 0.445075 | 0.669555 | 0.503141 | 0.883397 |
| CG10600 | 3358.068 | 0.542759 | 0.810972 | 0.669269 | 0.503324 | 0.883397 |
| CG5515 | 868.0496 | 0.743905 | 1.110322 | 0.669991 | 0.502864 | 0.883397 |
| CG11893 | 243.1354 | -1.63819 | 2.447635 | -0.6693 | 0.503307 | 0.883397 |
| Cyp6a9 | 467.7191 | -1.43583 | 2.143444 | -0.66987 | 0.502941 | 0.883397 |
| RN-tre | 2358.836 | 0.473611 | 0.707856 | 0.669079 | 0.503445 | 0.883397 |
| Asx | 3924.241 | 0.522813 | 0.781394 | 0.669077 | 0.503446 | 0.883397 |
| Ssrp | 664.5239 | -0.81769 | 1.220659 | -0.66988 | 0.502935 | 0.883397 |
| vib | 11220.62 | 0.362005 | 0.540105 | 0.670248 | 0.502699 | 0.883397 |
| Smyd3 | 217.5945 | 1.651288 | 2.471404 | 0.668158 | 0.504033 | 0.883545 |
| Nhe3 | 6856.702 | 0.489093 | 0.733111 | 0.667148 | 0.504678 | 0.883545 |
| rk | 114.411 | -1.90514 | 2.853403 | -0.66767 | 0.504343 | 0.883545 |
| by | 483.7079 | -1.11658 | 1.675644 | -0.66636 | 0.505181 | 0.883545 |
| CG31278 | 345.3176 | 1.650734 | 2.473714 | 0.66731 | 0.504574 | 0.883545 |
| CG9588 | 364.7077 | 1.19687 | 1.795008 | 0.666777 | 0.504915 | 0.883545 |
| CSN6 | 1092.061 | 1.094836 | 1.638365 | 0.668249 | 0.503975 | 0.883545 |
| side | 3964.271 | -0.39388 | 0.590135 | -0.66744 | 0.504489 | 0.883545 |
| CG1902 | 4202.253 | -0.39633 | 0.594773 | -0.66636 | 0.505182 | 0.883545 |
| Shrm | 131.7915 | 1.566669 | 2.344281 | 0.668294 | 0.503946 | 0.883545 |
| pio | 63.72663 | 2.590077 | 3.882535 | 0.66711 | 0.504702 | 0.883545 |
| Dis3l2 | 1715.283 | -0.70704 | 1.060589 | -0.66665 | 0.504995 | 0.883545 |
| Trs31 | 1144.001 | -0.67889 | 1.016469 | -0.66789 | 0.504203 | 0.883545 |
| CG10483 | 359.9784 | 1.026865 | 1.539927 | 0.666827 | 0.504883 | 0.883545 |
| CG9886 | 690.4774 | 0.934603 | 1.398378 | 0.668348 | 0.503911 | 0.883545 |
| CG3262 | 6082.814 | -0.38532 | 0.578896 | -0.66562 | 0.505654 | 0.883986 |
| CG8818 | 3856.291 | 0.447825 | 0.672732 | 0.665681 | 0.505615 | 0.883986 |
| Or30a | 273.935 | -1.63471 | 2.459614 | -0.66462 | 0.506293 | 0.884871 |
| PIG-C | 98.99526 | -2.28367 | 3.43736 | -0.66437 | 0.506455 | 0.884871 |
| Rtc1 | 1331.301 | 1.146332 | 1.725594 | 0.664311 | 0.506491 | 0.884871 |
| MED1 | 510.4732 | 0.856758 | 1.290281 | 0.664008 | 0.506685 | 0.885017 |
| PKD | 1449.227 | 0.636483 | 0.958818 | 0.663821 | 0.506805 | 0.885022 |
| Uggt | 2973.576 | 0.553512 | 0.83403 | 0.663659 | 0.506908 | 0.885022 |
| alien | 1510.719 | -0.69478 | 1.051921 | -0.66049 | 0.508942 | 0.885874 |
| alphaTub8 | 43825.18 | 0.291277 | 0.440925 | 0.660604 | 0.508866 | 0.885874 |

| | | | | | | |
|---|---|---|---|---|---|---|
| tws | 3680.37 | -0.3995 | 0.604137 | -0.66128 | 0.508434 | 0.885874 |
| ChT | 4152.264 | 0.435412 | 0.658791 | 0.660926 | 0.50866 | 0.885874 |
| CenB1A | 558.5544 | -1.374 | 2.074456 | -0.66234 | 0.507752 | 0.885874 |
| RpS3 | 11486.99 | 0.336706 | 0.508379 | 0.662312 | 0.507771 | 0.885874 |
| APC4 | 233.1734 | 1.545792 | 2.337615 | 0.661269 | 0.50844 | 0.885874 |
| kermit | 655.6254 | -1.20883 | 1.828115 | -0.66125 | 0.508454 | 0.885874 |
| hpRNA:CR | 196.7599 | -1.97848 | 2.993834 | -0.66085 | 0.508707 | 0.885874 |
| Tina-1 | 5675.052 | -0.36873 | 0.556863 | -0.66216 | 0.507867 | 0.885874 |
| hang | 2798.242 | -0.46388 | 0.701734 | -0.66104 | 0.508585 | 0.885874 |
| rudhira | 14132.53 | -0.32276 | 0.488554 | -0.66065 | 0.508835 | 0.885874 |
| Lsm12a | 1695.018 | 0.549803 | 0.831085 | 0.661548 | 0.508261 | 0.885874 |
| CrzR | 160.4071 | 2.113745 | 3.193935 | 0.6618 | 0.5081 | 0.885874 |
| Wdr92 | 589.3077 | 1.34115 | 2.031182 | 0.660281 | 0.509074 | 0.885912 |
| or | 1050.469 | -0.71063 | 1.076816 | -0.65994 | 0.509296 | 0.886106 |
| Dci | 935.275 | 0.654841 | 0.992955 | 0.659487 | 0.509583 | 0.886414 |
| Usp2 | 11145.35 | -0.33722 | 0.511995 | -0.65865 | 0.510123 | 0.887162 |
| brat | 4817.172 | -0.37215 | 0.56535 | -0.65827 | 0.510365 | 0.887389 |
| ppk8 | 234.6271 | -1.58804 | 2.415106 | -0.65755 | 0.51083 | 0.887925 |
| bif | 1031.024 | 0.738548 | 1.123604 | 0.657302 | 0.510986 | 0.887925 |
| Gapdh2 | 46520.3 | -0.27162 | 0.413251 | -0.65727 | 0.511005 | 0.887925 |
| CG43737 | 8824.466 | -0.37424 | 0.569563 | -0.65707 | 0.511134 | 0.887957 |
| PIG-Wa | 216.525 | -2.08538 | 3.176024 | -0.6566 | 0.511438 | 0.888293 |
| Dscam1 | 38509.39 | -0.31336 | 0.477465 | -0.65631 | 0.511626 | 0.888372 |
| Cyp6v1 | 229.1599 | 1.535181 | 2.340657 | 0.655876 | 0.511904 | 0.888372 |
| Mp | 5208.872 | 0.402398 | 0.61356 | 0.655841 | 0.511926 | 0.888372 |
| CG7083 | 2724.6 | -0.43363 | 0.660948 | -0.65607 | 0.511777 | 0.888372 |
| CG6388 | 477.3365 | -1.51131 | 2.305138 | -0.65563 | 0.512065 | 0.888421 |
| CG31191 | 1250.56 | -0.92404 | 1.409771 | -0.65545 | 0.512177 | 0.888423 |
| CG11885 | 510.7569 | 0.864632 | 1.320304 | 0.654874 | 0.512549 | 0.888564 |
| Naus | 3495.989 | -0.45797 | 0.699427 | -0.65478 | 0.512609 | 0.888564 |
| ATPsynF | 30609.92 | 0.281914 | 0.430868 | 0.654293 | 0.512923 | 0.888564 |
| CG44812 | 136.3844 | -1.98554 | 3.034563 | -0.65431 | 0.512913 | 0.888564 |
| shop | 462.7911 | 1.20735 | 1.844998 | 0.654391 | 0.51286 | 0.888564 |
| CG17260 | 135.8309 | -1.45986 | 2.229415 | -0.65482 | 0.512586 | 0.888564 |
| Ilp6 | 3115.925 | 0.435557 | 0.666398 | 0.653599 | 0.51337 | 0.888585 |
| e(r) | 2443.288 | 0.491586 | 0.75191 | 0.653782 | 0.513252 | 0.888585 |
| Zpr1 | 203.8763 | 2.139034 | 3.275163 | 0.653108 | 0.513687 | 0.888585 |
| SmF | 666.5271 | 0.747534 | 1.143892 | 0.653501 | 0.513434 | 0.888585 |
| eIF4H1 | 2293.274 | 0.456827 | 0.698414 | 0.654092 | 0.513053 | 0.888585 |
| Fatp3 | 1499.472 | 0.588694 | 0.901224 | 0.653216 | 0.513617 | 0.888585 |
| Pfdn4 | 1640.999 | -0.56624 | 0.867035 | -0.65307 | 0.51371 | 0.888585 |
| shv | 859.0897 | -1.07019 | 1.640431 | -0.65238 | 0.514154 | 0.8887 |
| CG30423 | 4796.631 | 0.406037 | 0.62265 | 0.65211 | 0.51433 | 0.8887 |
| Spt20 | 629.2001 | 1.11443 | 1.707225 | 0.652772 | 0.513903 | 0.8887 |
| nkd | 810.4714 | -0.69127 | 1.059616 | -0.65237 | 0.51416 | 0.8887 |
| CG13685 | 2118.126 | -0.63879 | 0.979321 | -0.65228 | 0.514222 | 0.8887 |
| CG12081 | 309.5859 | 1.300194 | 1.996072 | 0.651376 | 0.514804 | 0.888753 |
| lin | 261.8796 | 1.534477 | 2.355055 | 0.651567 | 0.51468 | 0.888753 |
| side-V | 1834.511 | -0.49919 | 0.766046 | -0.65164 | 0.514634 | 0.888753 |
| Trpm | 3347.638 | 0.745977 | 1.145156 | 0.65142 | 0.514775 | 0.888753 |

| | | | | | | |
|---|---|---|---|---|---|---|
| red | 4932.723 | 0.373861 | 0.574176 | 0.651126 | 0.514965 | 0.888841 |
| UGP | 2307.477 | -0.61376 | 0.942889 | -0.65094 | 0.515087 | 0.888859 |
| Lis-1 | 10161.17 | 0.354308 | 0.544781 | 0.650368 | 0.515455 | 0.889303 |
| Herp | 3067.867 | 0.444724 | 0.683996 | 0.650186 | 0.515572 | 0.889315 |
| CG18302 | 111.5381 | -2.41411 | 3.716026 | -0.64965 | 0.515919 | 0.88934 |
| Lapsyn | 1935.269 | 0.591635 | 0.910671 | 0.649669 | 0.515906 | 0.88934 |
| CG10737 | 4211.663 | 0.42 | 0.646218 | 0.649936 | 0.515734 | 0.88934 |
| CG13693 | 211.1265 | 1.734953 | 2.679615 | 0.647464 | 0.517332 | 0.889552 |
| CG2091 | 452.9364 | -1.26008 | 1.944934 | -0.64788 | 0.517063 | 0.889552 |
| Aldh | 4072.175 | -0.5315 | 0.820978 | -0.64739 | 0.517377 | 0.889552 |
| ova | 3090.263 | 0.524871 | 0.809419 | 0.648454 | 0.516691 | 0.889552 |
| sqz | 291.7765 | 1.391306 | 2.149233 | 0.64735 | 0.517406 | 0.889552 |
| lncRNA:CR | 90826.26 | 0.248756 | 0.383372 | 0.648864 | 0.516426 | 0.889552 |
| jigr1 | 4342.549 | -0.45755 | 0.706887 | -0.64728 | 0.517452 | 0.889552 |
| CG12065 | 1028.62 | 0.713082 | 1.102035 | 0.647059 | 0.517594 | 0.889552 |
| Atg8a | 40053.62 | 0.252427 | 0.389189 | 0.648598 | 0.516598 | 0.889552 |
| mtDNA-he | 251.2539 | -1.93014 | 2.974197 | -0.64896 | 0.516363 | 0.889552 |
| CG33228 | 443.0396 | 1.309923 | 2.023622 | 0.647316 | 0.517427 | 0.889552 |
| sel | 754.1414 | -0.86843 | 1.341445 | -0.64738 | 0.517384 | 0.889552 |
| Aplip1 | 10321.37 | 0.296504 | 0.457626 | 0.647918 | 0.517038 | 0.889552 |
| IntS2 | 235.5926 | -1.44359 | 2.230643 | -0.64716 | 0.517526 | 0.889552 |
| wcd | 358.3138 | 1.52472 | 2.359949 | 0.646082 | 0.518226 | 0.890285 |
| MsR2 | 167.8904 | 1.902454 | 2.944713 | 0.646058 | 0.518242 | 0.890285 |
| Fcp3C | 132.2965 | 1.887839 | 2.938668 | 0.642413 | 0.520605 | 0.89029 |
| CG3097 | 767.4125 | 0.870599 | 1.353592 | 0.643177 | 0.520109 | 0.89029 |
| CG4788 | 182.9174 | -1.71273 | 2.665164 | -0.64264 | 0.520461 | 0.89029 |
| CG10343 | 474.2308 | -1.28892 | 2.004357 | -0.64306 | 0.520185 | 0.89029 |
| CG3337 | 1950.513 | -0.5131 | 0.796479 | -0.6442 | 0.519443 | 0.89029 |
| fh | 387.9489 | -1.08513 | 1.688804 | -0.64254 | 0.520522 | 0.89029 |
| RpS28b | 14244.88 | 0.301483 | 0.467075 | 0.64547 | 0.518623 | 0.89029 |
| Cpr51A | 115.3776 | 1.547905 | 2.409983 | 0.642289 | 0.520686 | 0.89029 |
| Dad | 1692.974 | 0.64111 | 0.994648 | 0.644559 | 0.519213 | 0.89029 |
| CG34025 | 387.3742 | 0.936321 | 1.456258 | 0.642964 | 0.520248 | 0.89029 |
| Pde11 | 4707.575 | 0.355436 | 0.55174 | 0.644209 | 0.51944 | 0.89029 |
| bma | 2258.817 | 0.567071 | 0.880595 | 0.643963 | 0.519599 | 0.89029 |
| par-1 | 14433.05 | 0.294427 | 0.457484 | 0.643579 | 0.519848 | 0.89029 |
| RpS29 | 23440.04 | 0.274818 | 0.426851 | 0.643826 | 0.519688 | 0.89029 |
| SmydA-3 | 2281.645 | 0.511667 | 0.795552 | 0.643159 | 0.520121 | 0.89029 |
| milt | 5844.845 | 0.340381 | 0.528946 | 0.643507 | 0.519895 | 0.89029 |
| Cpes | 558.8558 | -1.18374 | 1.840452 | -0.64318 | 0.520108 | 0.89029 |
| lncRNA:CR | 153.4766 | 1.82707 | 2.832805 | 0.644968 | 0.518948 | 0.89029 |
| Ric | 8019.51 | 0.35737 | 0.553365 | 0.645813 | 0.518401 | 0.89029 |
| RpL28 | 16371.08 | 0.277441 | 0.430547 | 0.644393 | 0.519321 | 0.89029 |
| gig | 776.4122 | -0.72511 | 1.123071 | -0.64565 | 0.518505 | 0.89029 |
| CG6321 | 542.1382 | 1.112829 | 1.729801 | 0.643327 | 0.520012 | 0.89029 |
| Top1 | 2256.4 | -0.50171 | 0.781678 | -0.64184 | 0.520977 | 0.890599 |
| CG13887 | 3238.517 | 0.500985 | 0.780769 | 0.641656 | 0.521097 | 0.890614 |
| CG32982 | 5537.209 | 0.339508 | 0.53022 | 0.640315 | 0.521968 | 0.891154 |
| holn1 | 648.7415 | -0.80933 | 1.262643 | -0.64098 | 0.521536 | 0.891154 |
| ck | 1328.715 | -0.699 | 1.094528 | -0.63863 | 0.523061 | 0.891154 |

| | | | | | | |
|---|---|---|---|---|---|---|
| Faf2 | 2042.127 | 0.710092 | 1.111734 | 0.638724 | 0.523002 | 0.891154 |
| CG4565 | 399.5587 | 1.574563 | 2.472074 | 0.63694 | 0.524164 | 0.891154 |
| ATPsynCF6 | 14138.89 | 0.289204 | 0.453074 | 0.638314 | 0.523269 | 0.891154 |
| mask | 13134.52 | 0.283256 | 0.44421 | 0.637663 | 0.523693 | 0.891154 |
| Alp4 | 5333.304 | -0.56127 | 0.880517 | -0.63743 | 0.523843 | 0.891154 |
| RpS15 | 17015.54 | 0.270848 | 0.423973 | 0.638832 | 0.522933 | 0.891154 |
| fra | 649.9628 | -0.96014 | 1.502323 | -0.6391 | 0.522757 | 0.891154 |
| CG13185 | 89.49654 | 1.987939 | 3.106859 | 0.639855 | 0.522267 | 0.891154 |
| CG9170 | 2736.51 | -0.46418 | 0.727895 | -0.6377 | 0.523672 | 0.891154 |
| Sirup | 11298.58 | -0.36261 | 0.567793 | -0.63862 | 0.523068 | 0.891154 |
| CG17816 | 12645.92 | 0.288148 | 0.452074 | 0.63739 | 0.523871 | 0.891154 |
| gammaTub | 1152.671 | 0.968492 | 1.511431 | 0.640778 | 0.521667 | 0.891154 |
| osa | 5983.942 | -0.35907 | 0.561744 | -0.6392 | 0.522695 | 0.891154 |
| CG43066 | 12333.65 | -0.31956 | 0.501447 | -0.63727 | 0.523949 | 0.891154 |
| PheRS-m | 1820.385 | 0.826284 | 1.293495 | 0.638799 | 0.522953 | 0.891154 |
| Socs16D | 1813.676 | 0.528469 | 0.826134 | 0.639689 | 0.522375 | 0.891154 |
| CG7332 | 2485.781 | -0.45238 | 0.709986 | -0.63716 | 0.52402 | 0.891154 |
| MED19 | 797.2735 | 1.035888 | 1.624643 | 0.63761 | 0.523728 | 0.891154 |
| mRpS31 | 1235.245 | 0.909312 | 1.426322 | 0.637523 | 0.523784 | 0.891154 |
| CG10077 | 63120.96 | 0.240746 | 0.377958 | 0.636964 | 0.524148 | 0.891154 |
| Txl | 981.9813 | -1.02089 | 1.602904 | -0.6369 | 0.524189 | 0.891154 |
| CG31663 | 14071.79 | -0.3486 | 0.547135 | -0.63714 | 0.524033 | 0.891154 |
| ND-B14.5A | 5627.698 | -0.3467 | 0.549746 | -0.63066 | 0.528262 | 0.891536 |
| Rala | 1848.61 | -0.48804 | 0.770129 | -0.63371 | 0.52627 | 0.891536 |
| Nepl11 | 219.5357 | 1.49785 | 2.375617 | 0.63051 | 0.528361 | 0.891536 |
| SelT | 3492.012 | 0.383911 | 0.606625 | 0.632863 | 0.526823 | 0.891536 |
| Hel25E | 4534.175 | -0.47068 | 0.747367 | -0.62979 | 0.528834 | 0.891536 |
| Pex19 | 4833.979 | -0.45257 | 0.71372 | -0.6341 | 0.526014 | 0.891536 |
| Droj2 | 12246.8 | -0.28414 | 0.448595 | -0.63339 | 0.526479 | 0.891536 |
| CG31287 | 147.5744 | -2.31012 | 3.639166 | -0.63479 | 0.525564 | 0.891536 |
| CG16953 | 6151.41 | -0.40833 | 0.642397 | -0.63564 | 0.525014 | 0.891536 |
| CG10175 | 264.4358 | 1.590733 | 2.516047 | 0.632235 | 0.527233 | 0.891536 |
| GILT3 | 2127.439 | -0.6521 | 1.028309 | -0.63415 | 0.525981 | 0.891536 |
| CG4743 | 455.0622 | 0.899369 | 1.419738 | 0.633476 | 0.526423 | 0.891536 |
| CG31028 | 707.0506 | 0.736569 | 1.157613 | 0.636283 | 0.524592 | 0.891536 |
| CG15097 | 299.1076 | -1.0945 | 1.729263 | -0.63293 | 0.52678 | 0.891536 |
| CCHa1-R | 273.9262 | -1.56022 | 2.459172 | -0.63445 | 0.525787 | 0.891536 |
| Gclc | 4474.919 | -0.4553 | 0.72149 | -0.63105 | 0.528008 | 0.891536 |
| Prx2540-1 | 74.94732 | 2.353638 | 3.731275 | 0.630786 | 0.52818 | 0.891536 |
| Sec24AB | 2471.146 | 0.448919 | 0.709693 | 0.632554 | 0.527025 | 0.891536 |
| CG8243 | 689.0491 | -0.81405 | 1.291538 | -0.6303 | 0.528501 | 0.891536 |
| anne | 27370.16 | 0.249593 | 0.396013 | 0.630266 | 0.528521 | 0.891536 |
| qtc | 2659.401 | 0.55017 | 0.873357 | 0.629948 | 0.528728 | 0.891536 |
| CG12179 | 121.4534 | -1.73236 | 2.749308 | -0.63011 | 0.528624 | 0.891536 |
| CG3662 | 66493.18 | 0.303715 | 0.481903 | 0.630241 | 0.528537 | 0.891536 |
| CG34188 | 87.23839 | -2.07606 | 3.267716 | -0.63533 | 0.525216 | 0.891536 |
| CG43078 | 1731.552 | -0.53763 | 0.850597 | -0.63206 | 0.527345 | 0.891536 |
| wmd | 1206.188 | 0.712795 | 1.126467 | 0.632771 | 0.526883 | 0.891536 |
| Stoml2 | 2061.384 | -0.48779 | 0.771338 | -0.63239 | 0.527132 | 0.891536 |
| lncRNA:CR | 79.22575 | 1.943647 | 3.081574 | 0.630732 | 0.528216 | 0.891536 |

| | | | | | | |
|---|---|---|---|---|---|---|
| asRNA:CR4 | 76.64115 | 1.904662 | 3.016281 | 0.63146 | 0.52774 | 0.891536 |
| Fhos | 4993.53 | -0.44916 | 0.70991 | -0.6327 | 0.526932 | 0.891536 |
| Ttd14 | 2887.71 | 0.448081 | 0.708248 | 0.632661 | 0.526955 | 0.891536 |
| CG8001 | 416.2266 | 0.940502 | 1.494082 | 0.629485 | 0.529032 | 0.891536 |
| PGRP-SA | 378.7193 | -1.93971 | 3.05623 | -0.63467 | 0.525641 | 0.891536 |
| CG15760 | 2318.174 | 0.448189 | 0.706875 | 0.634042 | 0.526053 | 0.891536 |
| sun | 9613.708 | 0.300555 | 0.476001 | 0.631416 | 0.527769 | 0.891536 |
| CG5613 | 138.0225 | -1.94208 | 3.070054 | -0.63259 | 0.527002 | 0.891536 |
| Ranbp21 | 802.1846 | 0.81074 | 1.281851 | 0.632476 | 0.527076 | 0.891536 |
| CG14183 | 140.1748 | -2.04608 | 3.238747 | -0.63175 | 0.52755 | 0.891536 |
| CG5895 | 8815.409 | 0.336792 | 0.53021 | 0.635205 | 0.525295 | 0.891536 |
| CG8833 | 79.46504 | 2.417038 | 3.816191 | 0.633364 | 0.526496 | 0.891536 |
| nkt | 32455.36 | -0.27926 | 0.443682 | -0.62941 | 0.529079 | 0.891536 |
| Dim1 | 856.4791 | -0.99685 | 1.58339 | -0.62956 | 0.528979 | 0.891536 |
| CG12054 | 3752.223 | 0.416573 | 0.662235 | 0.62904 | 0.529323 | 0.891573 |
| CG14669 | 4583.407 | 0.422209 | 0.671044 | 0.629182 | 0.52923 | 0.891573 |
| Vha14-1 | 13091.77 | -0.32352 | 0.514507 | -0.6288 | 0.52948 | 0.891651 |
| Rab1 | 3375.449 | 0.393381 | 0.626817 | 0.627585 | 0.530276 | 0.892803 |
| sei | 2355.386 | -0.76022 | 1.211727 | -0.62739 | 0.530405 | 0.892834 |
| dia | 2019.645 | -0.51927 | 0.828905 | -0.62645 | 0.53102 | 0.89312 |
| TAF1B | 129.7998 | 2.039727 | 3.253702 | 0.626894 | 0.530729 | 0.89312 |
| Hexim | 1571.986 | 0.867751 | 1.384624 | 0.626705 | 0.530852 | 0.89312 |
| scaf6 | 2101.305 | 0.556649 | 0.888387 | 0.626584 | 0.530932 | 0.89312 |
| rst | 500.7574 | -0.92898 | 1.483619 | -0.62616 | 0.53121 | 0.893251 |
| CG6621 | 369.9172 | 0.927056 | 1.481021 | 0.625957 | 0.531343 | 0.893289 |
| CG10132 | 3044.584 | 0.395987 | 0.632852 | 0.625718 | 0.5315 | 0.893365 |
| pch2 | 755.0151 | -0.71531 | 1.143901 | -0.62533 | 0.531755 | 0.893607 |
| CG16791 | 11999.18 | 0.372407 | 0.596621 | 0.624195 | 0.5325 | 0.894109 |
| CG10365 | 2625.523 | 0.430601 | 0.689483 | 0.624527 | 0.532282 | 0.894109 |
| CG34417 | 1355.209 | -0.58433 | 0.935447 | -0.62465 | 0.532199 | 0.894109 |
| Ge-1 | 540.5368 | 0.80804 | 1.29434 | 0.624287 | 0.532439 | 0.894109 |
| CG6310 | 720.2448 | -0.89699 | 1.437502 | -0.62399 | 0.532632 | 0.894144 |
| Nbr | 1617.577 | 0.938132 | 1.504924 | 0.623375 | 0.533038 | 0.894223 |
| CG8701 | 170.3542 | -1.95173 | 3.129411 | -0.62367 | 0.532842 | 0.894223 |
| lncRNA:CR | 478.176 | -0.81914 | 1.314323 | -0.62324 | 0.533124 | 0.894223 |
| Cyp305a1 | 377.2198 | -1.10248 | 1.768653 | -0.62335 | 0.533056 | 0.894223 |
| A16 | 469.6975 | 1.217112 | 1.955008 | 0.622561 | 0.533573 | 0.894788 |
| wls | 1291.475 | -1.0635 | 1.709013 | -0.62229 | 0.533752 | 0.894902 |
| Fibp | 2006.948 | -0.48274 | 0.776072 | -0.62202 | 0.533926 | 0.895006 |
| Su(var)3-7 | 383.24 | -1.21916 | 1.963482 | -0.62092 | 0.534654 | 0.896039 |
| Hnf4 | 3696.465 | 0.378869 | 0.611431 | 0.619644 | 0.535492 | 0.896322 |
| mdy | 1986.605 | -0.59904 | 0.966112 | -0.62005 | 0.535223 | 0.896322 |
| Mdr49 | 103.999 | -2.15043 | 3.468729 | -0.61995 | 0.535292 | 0.896322 |
| cwo | 1642.969 | 0.504064 | 0.812876 | 0.620099 | 0.535192 | 0.896322 |
| Sarm | 6040.64 | -0.38051 | 0.613424 | -0.6203 | 0.535061 | 0.896322 |
| His2Av::GF | 3912.36 | -0.36494 | 0.588842 | -0.61976 | 0.535416 | 0.896322 |
| CG5056 | 1747.243 | 0.532935 | 0.860773 | 0.619136 | 0.535827 | 0.896647 |
| CG42392 | 1894.714 | 0.496521 | 0.802123 | 0.619009 | 0.535911 | 0.896647 |
| CG34125 | 2069.464 | -0.46188 | 0.746608 | -0.61864 | 0.536153 | 0.896858 |
| lncRNA:CR | 1466.087 | 0.622743 | 1.006895 | 0.618479 | 0.53626 | 0.896858 |

| | | | | | | |
|---|---|---|---|---|---|---|
| Oatp26F | 73.58616 | -1.97116 | 3.18968 | -0.61798 | 0.536588 | 0.896972 |
| CG31342 | 2151.856 | -0.55587 | 0.900204 | -0.6175 | 0.536908 | 0.896972 |
| CycG | 19294.45 | 0.252616 | 0.409639 | 0.61668 | 0.537446 | 0.896972 |
| CG33156 | 2088.768 | 0.817181 | 1.323413 | 0.61748 | 0.536918 | 0.896972 |
| GLaz | 4085.414 | -0.45939 | 0.743567 | -0.61781 | 0.536699 | 0.896972 |
| toy | 10655.09 | 0.315275 | 0.510738 | 0.617293 | 0.537041 | 0.896972 |
| Dyrk2 | 7613.795 | 0.309759 | 0.50212 | 0.616902 | 0.537299 | 0.896972 |
| salm | 606.1028 | -0.72999 | 1.182984 | -0.61708 | 0.537183 | 0.896972 |
| CG15210 | 97.13189 | -1.99997 | 3.240383 | -0.6172 | 0.537102 | 0.896972 |
| Ucp4A | 1440.895 | 0.52629 | 0.853306 | 0.616766 | 0.537389 | 0.896972 |
| GstT1 | 1150.841 | 0.61181 | 0.992674 | 0.616325 | 0.53768 | 0.897177 |
| cav | 247.3369 | 1.191902 | 1.93525 | 0.61589 | 0.537967 | 0.897289 |
| CG8405 | 1261.796 | -0.58472 | 0.949626 | -0.61573 | 0.538071 | 0.897289 |
| Cul3 | 3031.229 | -0.40026 | 0.650079 | -0.61572 | 0.538083 | 0.897289 |
| lncRNA:CR | 81.14632 | -1.68957 | 2.74504 | -0.6155 | 0.538226 | 0.897341 |
| CG7920 | 11300.39 | -0.32678 | 0.531078 | -0.61531 | 0.538347 | 0.897357 |
| CG5196 | 2227.147 | -0.59312 | 0.965035 | -0.61461 | 0.538812 | 0.897876 |
| CG5590 | 1443.955 | -0.5947 | 0.967774 | -0.6145 | 0.538884 | 0.897876 |
| asRNA:CR4 | 109.0826 | -1.54698 | 2.518143 | -0.61433 | 0.538994 | 0.897876 |
| cg | 8347.965 | 0.344204 | 0.561515 | 0.612991 | 0.539882 | 0.898297 |
| CG12567 | 1896.2 | -0.52852 | 0.862273 | -0.61294 | 0.539918 | 0.898297 |
| Cdk7 | 620.5896 | -1.00826 | 1.642936 | -0.6137 | 0.539416 | 0.898297 |
| RPA3 | 368.6584 | -1.19479 | 1.947558 | -0.61348 | 0.539559 | 0.898297 |
| CG11164 | 537.0873 | -1.18928 | 1.938914 | -0.61337 | 0.539629 | 0.898297 |
| DCTN1-p15 | 624.3013 | -1.08469 | 1.768876 | -0.61321 | 0.539737 | 0.898297 |
| Hsp60D | 87.51466 | 2.400705 | 3.919125 | 0.612562 | 0.540166 | 0.898523 |
| CG6287 | 5248.991 | -0.3604 | 0.588902 | -0.61199 | 0.540546 | 0.898596 |
| ATPsynC | 72204.06 | 0.238536 | 0.389692 | 0.612115 | 0.540462 | 0.898596 |
| CG42676 | 261.4321 | 1.289891 | 2.107455 | 0.612061 | 0.540498 | 0.898596 |
| mTerf3 | 394.2643 | -1.18111 | 1.931553 | -0.61148 | 0.540881 | 0.898781 |
| Lerp | 1742.221 | 0.802753 | 1.312526 | 0.611609 | 0.540796 | 0.898781 |
| CG15445 | 120.0381 | 1.816726 | 2.974846 | 0.610696 | 0.541401 | 0.899458 |
| Hpd | 255.2598 | 1.535123 | 2.515051 | 0.610374 | 0.541614 | 0.899589 |
| CG6163 | 361.3577 | 0.971995 | 1.59281 | 0.610239 | 0.541703 | 0.899589 |
| Dok | 254.9747 | -1.51004 | 2.476654 | -0.60971 | 0.542055 | 0.899801 |
| JTBR | 851.789 | -0.65928 | 1.081278 | -0.60972 | 0.542048 | 0.899801 |
| CG1213 | 3297.23 | -0.47816 | 0.784541 | -0.60948 | 0.542206 | 0.899865 |
| CG32195 | 2471.763 | 0.503176 | 0.825888 | 0.609255 | 0.542355 | 0.899926 |
| CG12766 | 439.5347 | -1.65368 | 2.717683 | -0.60849 | 0.542864 | 0.900498 |
| CG3529 | 1224.514 | 0.607658 | 0.998786 | 0.608397 | 0.542925 | 0.900498 |
| phtf | 2939.457 | 0.646599 | 1.063784 | 0.607829 | 0.543301 | 0.900565 |
| lncRNA:CR | 12723.08 | 0.305572 | 0.502615 | 0.607964 | 0.543211 | 0.900565 |
| CG7272 | 1297.723 | -0.52637 | 0.865846 | -0.60792 | 0.54324 | 0.900565 |
| Rpn9 | 2407.067 | 0.447701 | 0.736866 | 0.607575 | 0.54347 | 0.900658 |
| Nep3 | 3637.494 | 0.379319 | 0.624553 | 0.607346 | 0.543622 | 0.900724 |
| Usp12-46 | 850.1139 | -0.97398 | 1.605169 | -0.60678 | 0.544 | 0.901165 |
| CG5850 | 817.5763 | 0.775959 | 1.27997 | 0.606232 | 0.544361 | 0.9012 |
| Evi5 | 6897.066 | 0.318615 | 0.525764 | 0.606005 | 0.544512 | 0.9012 |
| CG46387 | 82.17634 | -1.68612 | 2.782848 | -0.6059 | 0.544582 | 0.9012 |
| REG | 4312.574 | -0.53825 | 0.887957 | -0.60617 | 0.544403 | 0.9012 |

| | | | | | | |
|---|---|---|---|---|---|---|
| Mkp3 | 7052.386 | 0.308643 | 0.509339 | 0.605968 | 0.544536 | 0.9012 |
| CG9436 | 3305.809 | -0.37615 | 0.621369 | -0.60536 | 0.544938 | 0.90123 |
| miple2 | 704.0406 | -1.15152 | 1.902157 | -0.60538 | 0.544929 | 0.90123 |
| betaTub56 | 19768.51 | 0.258852 | 0.427412 | 0.605625 | 0.544764 | 0.90123 |
| CG11426 | 2587.409 | 0.446023 | 0.737037 | 0.605157 | 0.545075 | 0.901272 |
| Rpb12 | 1979.112 | -0.45883 | 0.758957 | -0.60456 | 0.545472 | 0.901371 |
| lncRNA:CR | 117.4705 | 2.028295 | 3.354742 | 0.604605 | 0.545441 | 0.901371 |
| CG12096 | 724.5857 | -0.83894 | 1.387241 | -0.60475 | 0.545342 | 0.901371 |
| Actn | 1372.946 | 0.540727 | 0.896746 | 0.602988 | 0.546517 | 0.901797 |
| CG13116 | 131.1612 | -2.05398 | 3.400879 | -0.60396 | 0.545874 | 0.901797 |
| Idgf3 | 6813.566 | -0.39617 | 0.656986 | -0.60301 | 0.546505 | 0.901797 |
| CG18473 | 311.0374 | 1.712611 | 2.838797 | 0.603288 | 0.546317 | 0.901797 |
| GatA | 829.2179 | 0.834565 | 1.383831 | 0.603083 | 0.546453 | 0.901797 |
| mRpL3 | 1614.254 | -0.89244 | 1.479337 | -0.60327 | 0.546329 | 0.901797 |
| Zip71B | 1977.605 | 0.794194 | 1.316561 | 0.603233 | 0.546353 | 0.901797 |
| CG8312 | 1981.554 | 0.584388 | 0.969786 | 0.602595 | 0.546778 | 0.902022 |
| Gip | 128.0248 | -1.57646 | 2.616763 | -0.60245 | 0.546878 | 0.902022 |
| CG31998 | 8035.213 | 0.290586 | 0.482839 | 0.601829 | 0.547288 | 0.902143 |
| conu | 8549.975 | 0.327457 | 0.543993 | 0.60195 | 0.547207 | 0.902143 |
| Uba2 | 423.437 | 1.318232 | 2.189928 | 0.601952 | 0.547206 | 0.902143 |
| Atg4b | 980.6177 | -0.99686 | 1.656943 | -0.60163 | 0.547423 | 0.90218 |
| woc | 3361.35 | 0.37715 | 0.627244 | 0.601281 | 0.547653 | 0.90221 |
| Myd88 | 726.6849 | -0.68964 | 1.14731 | -0.60109 | 0.547778 | 0.90221 |
| fuss | 565.2443 | -0.82504 | 1.372431 | -0.60116 | 0.547736 | 0.90221 |
| Gr39a | 98.24589 | -1.9802 | 3.296699 | -0.60066 | 0.548065 | 0.902497 |
| tna | 7872.569 | -0.31688 | 0.527765 | -0.60042 | 0.548229 | 0.902581 |
| CaMKII | 39790.82 | 0.229378 | 0.38232 | 0.599964 | 0.54853 | 0.902892 |
| CG3164 | 1496.81 | 0.57357 | 0.956302 | 0.599779 | 0.548653 | 0.90291 |
| RpS14b | 2591.636 | 0.459767 | 0.767324 | 0.599183 | 0.549051 | 0.903163 |
| kay | 5672.882 | 0.399717 | 0.667269 | 0.599034 | 0.54915 | 0.903163 |
| AP-2mu | 2060.922 | 0.546469 | 0.912752 | 0.598705 | 0.54937 | 0.903163 |
| lovit | 165.5626 | 1.416533 | 2.363509 | 0.599335 | 0.54895 | 0.903163 |
| Pits | 8470.103 | 0.32245 | 0.538488 | 0.598806 | 0.549302 | 0.903163 |
| Rab27 | 1840.523 | 0.465984 | 0.784167 | 0.59424 | 0.552352 | 0.903273 |
| Rtca | 939.0366 | 0.603054 | 1.019669 | 0.591421 | 0.554238 | 0.903273 |
| CG14812 | 1777.794 | 0.485423 | 0.819773 | 0.592143 | 0.553755 | 0.903273 |
| wds | 717.7125 | 0.859864 | 1.440353 | 0.596982 | 0.55052 | 0.903273 |
| dbe | 344.1825 | 1.347067 | 2.264597 | 0.594837 | 0.551952 | 0.903273 |
| Wdfy2 | 2504.102 | -0.59269 | 1.002964 | -0.59094 | 0.55456 | 0.903273 |
| loh | 63.91622 | 2.098363 | 3.540958 | 0.592598 | 0.55345 | 0.903273 |
| CG9934 | 739.6701 | 0.717063 | 1.204411 | 0.595364 | 0.5516 | 0.903273 |
| grn | 104.7976 | -2.09595 | 3.521843 | -0.59513 | 0.551757 | 0.903273 |
| foxo | 4088.757 | -0.36057 | 0.608532 | -0.59253 | 0.553499 | 0.903273 |
| NK7.1 | 5282.804 | 0.323024 | 0.542758 | 0.595153 | 0.551741 | 0.903273 |
| CG32856 | 363.4132 | 1.018225 | 1.721983 | 0.591309 | 0.554313 | 0.903273 |
| EndoA | 12282.65 | 0.265167 | 0.448001 | 0.591889 | 0.553925 | 0.903273 |
| Calx | 10037.79 | -0.33198 | 0.5614 | -0.59135 | 0.554288 | 0.903273 |
| Rlip | 2996.512 | 0.410635 | 0.694115 | 0.591596 | 0.554121 | 0.903273 |
| glec | 4536.809 | -0.38179 | 0.639345 | -0.59716 | 0.550398 | 0.903273 |
| Sld5 | 90.36049 | -2.29897 | 3.890613 | -0.5909 | 0.554586 | 0.903273 |

| | | | | | | |
|---|---|---|---|---|---|---|
| Fer1HCH | 28902.41 | -0.25083 | 0.421739 | -0.59474 | 0.552016 | 0.903273 |
| CG10081 | 227.0014 | 1.776973 | 2.999041 | 0.592514 | 0.553507 | 0.903273 |
| RhoGAP54 | 534.5396 | 1.130795 | 1.911724 | 0.591505 | 0.554182 | 0.903273 |
| CG8878 | 2738.595 | -0.44533 | 0.753077 | -0.59135 | 0.554286 | 0.903273 |
| Buffy | 176.8453 | -2.29621 | 3.856485 | -0.59541 | 0.551566 | 0.903273 |
| CG1354 | 1166.523 | 0.547936 | 0.927036 | 0.591063 | 0.554478 | 0.903273 |
| Cf2 | 2514.6 | 0.416002 | 0.703753 | 0.59112 | 0.55444 | 0.903273 |
| CG4041 | 314.5195 | -1.21725 | 2.035411 | -0.59804 | 0.549816 | 0.903273 |
| CG10581 | 957.2266 | -0.6606 | 1.107542 | -0.59645 | 0.550873 | 0.903273 |
| Fili | 1263.478 | 0.524038 | 0.879941 | 0.595537 | 0.551484 | 0.903273 |
| glo | 3623.601 | 0.426261 | 0.71815 | 0.593554 | 0.552811 | 0.903273 |
| rdgA | 9228.737 | 0.306661 | 0.519132 | 0.590719 | 0.554709 | 0.903273 |
| mtRNApol | 633.685 | 0.782589 | 1.322737 | 0.591644 | 0.554089 | 0.903273 |
| Nup160 | 394.6581 | -1.39712 | 2.359081 | -0.59223 | 0.553695 | 0.903273 |
| CG4612 | 11423.5 | -0.3226 | 0.539167 | -0.59833 | 0.549617 | 0.903273 |
| uzip | 5973.988 | -0.34026 | 0.571846 | -0.59501 | 0.551835 | 0.903273 |
| Trs20 | 1246.422 | -0.55562 | 0.939563 | -0.59135 | 0.554283 | 0.903273 |
| asRNA:CR4 | 116.6883 | -1.80493 | 3.055619 | -0.59069 | 0.554726 | 0.903273 |
| lncRNA:CR | 319.7124 | -1.01075 | 1.691308 | -0.59762 | 0.550096 | 0.903273 |
| prage | 1161.679 | 0.603529 | 1.01589 | 0.594089 | 0.552452 | 0.903273 |
| sicily | 1536.184 | 0.628712 | 1.060711 | 0.592727 | 0.553364 | 0.903273 |
| AMPdeam | 4028.257 | 0.364276 | 0.6116 | 0.595612 | 0.551435 | 0.903273 |
| CG12672 | 184.8791 | 1.478356 | 2.498262 | 0.591754 | 0.554015 | 0.903273 |
| Rip11 | 1335.854 | -0.56864 | 0.95471 | -0.59562 | 0.551429 | 0.903273 |
| CG14220 | 541.303 | 1.141636 | 1.919497 | 0.594758 | 0.552005 | 0.903273 |
| CG6933 | 217.7172 | -1.87483 | 3.173518 | -0.59077 | 0.554672 | 0.903273 |
| nes | 2842.409 | 0.58917 | 0.989063 | 0.595685 | 0.551386 | 0.903273 |
| Grx1 | 2426.445 | -0.43939 | 0.736282 | -0.59677 | 0.550664 | 0.903273 |
| CG9628 | 445.4332 | -1.02322 | 1.720492 | -0.59472 | 0.552029 | 0.903273 |
| CG4629 | 372.3901 | 0.900569 | 1.511533 | 0.595798 | 0.55131 | 0.903273 |
| Snx27 | 3246.237 | 0.390511 | 0.661524 | 0.590321 | 0.554975 | 0.903496 |
| EMC4 | 4760.178 | -0.39099 | 0.662684 | -0.59001 | 0.555182 | 0.903649 |
| rpr | 195.8241 | -1.72749 | 2.929368 | -0.58971 | 0.555383 | 0.903793 |
| Cog8 | 506.5673 | 1.249788 | 2.120878 | 0.589279 | 0.555674 | 0.904083 |
| eIF3f1 | 8699.77 | -0.29466 | 0.502055 | -0.58691 | 0.557262 | 0.904299 |
| CG3793 | 1505.659 | -0.60991 | 1.03721 | -0.58803 | 0.55651 | 0.904299 |
| rdog | 480.7129 | 0.951705 | 1.616442 | 0.588765 | 0.556019 | 0.904299 |
| botv | 192.3314 | 1.365757 | 2.327998 | 0.586666 | 0.557428 | 0.904299 |
| CG6665 | 884.8015 | 0.648798 | 1.106418 | 0.586395 | 0.55761 | 0.904299 |
| pcs | 5014.924 | -0.3483 | 0.593452 | -0.58691 | 0.557264 | 0.904299 |
| CG14044 | 102.3207 | -1.9477 | 3.317799 | -0.58705 | 0.557173 | 0.904299 |
| cta | 3190.55 | -0.36726 | 0.624213 | -0.58836 | 0.556288 | 0.904299 |
| CG14464 | 2900.312 | 0.431923 | 0.736242 | 0.586659 | 0.557433 | 0.904299 |
| dlt | 391.5648 | -1.17657 | 2.004398 | -0.58699 | 0.557208 | 0.904299 |
| Fie | 16490.39 | -0.24814 | 0.421795 | -0.5883 | 0.556329 | 0.904299 |
| Upf1 | 446.5532 | -0.87709 | 1.495342 | -0.58655 | 0.557507 | 0.904299 |
| Rrp45 | 65.32329 | 2.30112 | 3.923516 | 0.586494 | 0.557543 | 0.904299 |
| CG12991 | 2705.223 | -0.38752 | 0.66079 | -0.58644 | 0.557578 | 0.904299 |
| CG1529 | 272.4934 | 1.451231 | 2.473931 | 0.586609 | 0.557466 | 0.904299 |
| daed | 369.461 | -1.51183 | 2.570434 | -0.58816 | 0.556425 | 0.904299 |

| | | | | | | |
|---|---|---|---|---|---|---|
| Smr | 5582.707 | -0.32627 | 0.556576 | -0.58621 | 0.557734 | 0.904317 |
| mRpS11 | 2677.533 | 0.394354 | 0.673863 | 0.585214 | 0.558404 | 0.904673 |
| Son | 707.5193 | -0.90482 | 1.545997 | -0.58527 | 0.558367 | 0.904673 |
| CG8578 | 958.6806 | 0.646509 | 1.104074 | 0.585567 | 0.558167 | 0.904673 |
| Serinc | 14410.56 | -0.27212 | 0.464855 | -0.58539 | 0.558284 | 0.904673 |
| Or1a | 78.49859 | -2.27746 | 3.952138 | -0.57626 | 0.564438 | 0.904691 |
| Rbf | 599.9219 | 0.842449 | 1.475753 | 0.570861 | 0.568094 | 0.904691 |
| RpL36 | 21958.39 | 0.262773 | 0.457598 | 0.574244 | 0.565803 | 0.904691 |
| CG15912 | 191.8854 | 1.562968 | 2.677774 | 0.583682 | 0.559435 | 0.904691 |
| CG1090 | 5545.897 | -0.38845 | 0.677086 | -0.57371 | 0.566163 | 0.904691 |
| CG7214 | 78.85622 | -2.27749 | 3.952133 | -0.57627 | 0.564434 | 0.904691 |
| Tep2 | 1889.337 | -0.4941 | 0.857571 | -0.57616 | 0.564506 | 0.904691 |
| CG13390 | 328.7372 | 1.378119 | 2.393626 | 0.575745 | 0.564787 | 0.904691 |
| Snx17 | 2123.171 | -0.56517 | 0.983242 | -0.5748 | 0.565426 | 0.904691 |
| w-cup | 82.07484 | -2.27768 | 3.952085 | -0.57632 | 0.564396 | 0.904691 |
| Sec71 | 282.4074 | 1.159924 | 1.991415 | 0.582462 | 0.560255 | 0.904691 |
| BuGZ | 6648.11 | 0.302274 | 0.519041 | 0.58237 | 0.560318 | 0.904691 |
| Ugt301D1 | 4652.88 | -0.39719 | 0.69545 | -0.57113 | 0.567911 | 0.904691 |
| CG8500 | 4069.278 | 0.376402 | 0.659063 | 0.571116 | 0.567921 | 0.904691 |
| RpL3 | 17941.36 | 0.23878 | 0.417897 | 0.571386 | 0.567738 | 0.904691 |
| CG6723 | 7038.124 | 0.366246 | 0.630621 | 0.58077 | 0.561396 | 0.904691 |
| KLHL18 | 2151.245 | -0.43437 | 0.759232 | -0.57212 | 0.567243 | 0.904691 |
| ATPsynO | 24783.12 | 0.257468 | 0.445349 | 0.578126 | 0.563179 | 0.904691 |
| CG5191 | 1258.029 | -0.51509 | 0.884985 | -0.58203 | 0.560546 | 0.904691 |
| AdSS | 4141.445 | -0.33437 | 0.579459 | -0.57704 | 0.563911 | 0.904691 |
| CG5745 | 330.8597 | -0.99925 | 1.747691 | -0.57175 | 0.567489 | 0.904691 |
| Dph5 | 779.5189 | 0.816476 | 1.404438 | 0.581354 | 0.561002 | 0.904691 |
| Elal | 2786.607 | 0.417925 | 0.727835 | 0.574203 | 0.565831 | 0.904691 |
| mino | 1171.358 | -0.70531 | 1.228605 | -0.57407 | 0.565918 | 0.904691 |
| Cisd2 | 7113.038 | -0.35253 | 0.615628 | -0.57263 | 0.566895 | 0.904691 |
| Snx16 | 666.6792 | -0.76812 | 1.337434 | -0.57433 | 0.565748 | 0.904691 |
| CBP | 216.6169 | -1.38673 | 2.411039 | -0.57516 | 0.565184 | 0.904691 |
| RpS14a | 22010.03 | 0.245437 | 0.425453 | 0.576883 | 0.564018 | 0.904691 |
| CG32706 | 312.9273 | -1.25327 | 2.153862 | -0.58187 | 0.560654 | 0.904691 |
| cv-2 | 2825.276 | -0.43236 | 0.754371 | -0.57314 | 0.566548 | 0.904691 |
| CG9304 | 580.9031 | 1.119801 | 1.924154 | 0.581971 | 0.560586 | 0.904691 |
| CG30099 | 72.06135 | -2.27702 | 3.952248 | -0.57613 | 0.564525 | 0.904691 |
| ckn | 6458.757 | -0.31277 | 0.537464 | -0.58193 | 0.560613 | 0.904691 |
| CG8089 | 140.1888 | -2.27965 | 3.951598 | -0.57689 | 0.564011 | 0.904691 |
| bbc | 9275.107 | -0.28833 | 0.496207 | -0.58108 | 0.561189 | 0.904691 |
| Dp | 1654.649 | -0.79833 | 1.39711 | -0.57141 | 0.567719 | 0.904691 |
| CG17019 | 4590.098 | -0.44663 | 0.769007 | -0.58078 | 0.561387 | 0.904691 |
| RpS11 | 22009.93 | 0.238392 | 0.416453 | 0.572434 | 0.567028 | 0.904691 |
| san | 1877.718 | 0.44622 | 0.773724 | 0.576716 | 0.564131 | 0.904691 |
| CG12391 | 1226.876 | -0.52896 | 0.914766 | -0.57824 | 0.563099 | 0.904691 |
| CG7220 | 5490.539 | -0.31504 | 0.543187 | -0.57999 | 0.561921 | 0.904691 |
| babo | 3711.938 | -0.35817 | 0.619818 | -0.57787 | 0.563354 | 0.904691 |
| Corin | 7504.089 | 0.327962 | 0.569208 | 0.576174 | 0.564498 | 0.904691 |
| CG12107 | 1814.961 | -0.48112 | 0.831146 | -0.57886 | 0.562684 | 0.904691 |
| LRR | 3418.901 | 0.475303 | 0.823965 | 0.576849 | 0.564042 | 0.904691 |

| | | | | | | |
|---|---|---|---|---|---|---|
| Tsp42Eh | 2327.234 | -0.49115 | 0.85885 | -0.57187 | 0.567408 | 0.904691 |
| dpr7 | 14812.77 | 0.263617 | 0.461807 | 0.570839 | 0.568109 | 0.904691 |
| Zyx | 12132.91 | 0.270527 | 0.464074 | 0.582939 | 0.559934 | 0.904691 |
| CG18635 | 101.923 | -2.2786 | 3.951856 | -0.57659 | 0.564216 | 0.904691 |
| MRP | 2671.098 | 0.393233 | 0.678065 | 0.579934 | 0.561959 | 0.904691 |
| CROT | 404.6744 | -1.32288 | 2.271043 | -0.5825 | 0.56023 | 0.904691 |
| CG4619 | 918.4712 | -0.70418 | 1.217165 | -0.57854 | 0.562902 | 0.904691 |
| CG5885 | 2160.533 | 0.565783 | 0.972934 | 0.581523 | 0.560888 | 0.904691 |
| fbl | 604.6008 | -0.71861 | 1.251904 | -0.57401 | 0.565959 | 0.904691 |
| mt:ND3 | 3529.506 | 0.352898 | 0.610555 | 0.577995 | 0.563267 | 0.904691 |
| side-III | 2666.279 | 0.385735 | 0.663033 | 0.581773 | 0.56072 | 0.904691 |
| Ufm1 | 1455.777 | -0.50947 | 0.883631 | -0.57657 | 0.56423 | 0.904691 |
| CG42260 | 7438.627 | -0.40326 | 0.702814 | -0.57378 | 0.566117 | 0.904691 |
| CG42673 | 2433.239 | 0.406247 | 0.703756 | 0.577255 | 0.563767 | 0.904691 |
| CG42724 | 3686.109 | -0.35397 | 0.618557 | -0.57226 | 0.567148 | 0.904691 |
| Dscam4 | 3343.253 | 0.374419 | 0.646741 | 0.578932 | 0.562635 | 0.904691 |
| bel | 3280.468 | 0.414111 | 0.72252 | 0.573148 | 0.566544 | 0.904691 |
| CG3500 | 954.0756 | -0.74421 | 1.294132 | -0.57506 | 0.565248 | 0.904691 |
| RpL39 | 9806.521 | 0.28823 | 0.500991 | 0.57532 | 0.565075 | 0.904691 |
| Mtpbeta | 1509.494 | -0.57053 | 0.984681 | -0.57941 | 0.562315 | 0.904691 |
| rno | 2467.033 | 0.462508 | 0.804885 | 0.574627 | 0.565544 | 0.904691 |
| CG43689 | 1401.898 | 0.599763 | 1.04352 | 0.57475 | 0.56546 | 0.904691 |
| CG43693 | 9018.248 | -0.34009 | 0.585995 | -0.58036 | 0.56167 | 0.904691 |
| lncRNA:CR | 233.8483 | 1.309663 | 2.286156 | 0.572867 | 0.566735 | 0.904691 |
| lncRNA:CR | 1083.048 | -0.60913 | 1.054674 | -0.57756 | 0.563563 | 0.904691 |
| asRNA:CR4 | 107.4662 | -2.2788 | 3.951807 | -0.57665 | 0.564177 | 0.904691 |
| lncRNA:CR | 109.2543 | -2.27886 | 3.951793 | -0.57667 | 0.564166 | 0.904691 |
| lncRNA:CR | 165.759 | -2.28052 | 3.948762 | -0.57753 | 0.563583 | 0.904691 |
| Fas1 | 140229.6 | 0.215213 | 0.371432 | 0.579413 | 0.562311 | 0.904691 |
| cv-c | 141.3167 | 1.427063 | 2.488564 | 0.573448 | 0.566341 | 0.904691 |
| lncRNA:CR | 111.5789 | -2.27894 | 3.951774 | -0.57669 | 0.564151 | 0.904691 |
| lncRNA:CR | 122.2139 | 1.674433 | 2.888883 | 0.579613 | 0.562176 | 0.904691 |
| Or63a | 1568.771 | 0.852212 | 1.468058 | 0.580503 | 0.561575 | 0.904691 |
| CG9240 | 1539.023 | 0.635697 | 1.109691 | 0.572859 | 0.56674 | 0.904691 |
| Pgant7 | 719.8031 | 1.071347 | 1.839176 | 0.582515 | 0.56022 | 0.904691 |
| Pex23 | 815.3398 | 0.645437 | 1.128188 | 0.5721 | 0.567254 | 0.904691 |
| Prp3 | 786.0846 | 0.650978 | 1.121211 | 0.580602 | 0.561509 | 0.904691 |
| RasGAP1 | 943.1549 | 0.714809 | 1.229772 | 0.581254 | 0.561069 | 0.904691 |
| CG32409 | 142.1987 | -1.79819 | 3.110548 | -0.57809 | 0.563201 | 0.904691 |
| fipi | 17747.07 | 0.280705 | 0.490952 | 0.571757 | 0.567487 | 0.904691 |
| CG9967 | 97.12068 | 2.067236 | 3.586505 | 0.576393 | 0.56435 | 0.904691 |
| CG13252 | 563.1964 | -0.87339 | 1.530587 | -0.57062 | 0.568257 | 0.904747 |
| Syx5 | 1781.923 | -0.49331 | 0.867017 | -0.56897 | 0.569375 | 0.905138 |
| GstD3 | 6923.459 | -0.32994 | 0.579121 | -0.56973 | 0.568863 | 0.905138 |
| CG6073 | 335.9102 | -1.35197 | 2.378651 | -0.56838 | 0.56978 | 0.905138 |
| NSD | 409.7312 | 0.891925 | 1.570744 | 0.567836 | 0.570146 | 0.905138 |
| Ir47a | 108.2357 | -2.244 | 3.951776 | -0.56785 | 0.57014 | 0.905138 |
| CG34250 | 1875.731 | -0.45155 | 0.793871 | -0.5688 | 0.569493 | 0.905138 |
| RpS25 | 43808.01 | 0.227877 | 0.401246 | 0.567924 | 0.570087 | 0.905138 |
| pea | 333.9283 | 1.327566 | 2.336574 | 0.568167 | 0.569921 | 0.905138 |

| | | | | | | |
|---|---|---|---|---|---|---|
| p120ctn | 10760.93 | 0.285619 | 0.502318 | 0.568602 | 0.569626 | 0.905138 |
| mRpL37 | 1549.96 | -0.49497 | 0.868541 | -0.56989 | 0.568753 | 0.905138 |
| CG3894 | 2174.926 | -0.53188 | 0.936789 | -0.56777 | 0.570194 | 0.905138 |
| kst | 851.5514 | 0.813463 | 1.429745 | 0.568957 | 0.569385 | 0.905138 |
| COX6B | 24257.53 | 0.256898 | 0.451419 | 0.56909 | 0.569295 | 0.905138 |
| caps | 2198.351 | 0.450583 | 0.792427 | 0.568612 | 0.56962 | 0.905138 |
| path | 1078.325 | 0.60866 | 1.071259 | 0.568172 | 0.569918 | 0.905138 |
| CG14015 | 1770.605 | -0.51959 | 0.915928 | -0.56729 | 0.570518 | 0.905474 |
| lncRNA:CR | 96.25529 | -2.23941 | 3.95188 | -0.56667 | 0.570939 | 0.905962 |
| dpr3 | 576.7516 | -0.82618 | 1.458708 | -0.56638 | 0.571137 | 0.906097 |
| CG5282 | 3286.061 | 0.425657 | 0.754786 | 0.563943 | 0.572793 | 0.906105 |
| CG17321 | 2546.801 | -0.3843 | 0.680931 | -0.56438 | 0.572495 | 0.906105 |
| Arfip | 1090.824 | 1.05984 | 1.878495 | 0.564196 | 0.57262 | 0.906105 |
| Pfdn5 | 2017.615 | -0.49726 | 0.878979 | -0.56572 | 0.571583 | 0.906105 |
| CG6330 | 4940.203 | -0.32967 | 0.584345 | -0.56416 | 0.572643 | 0.906105 |
| CG31051 | 978.1189 | -0.9727 | 1.722469 | -0.56471 | 0.57227 | 0.906105 |
| CG13751 | 1000.922 | -0.57377 | 1.013874 | -0.56592 | 0.57145 | 0.906105 |
| nord | 1546.547 | -0.75334 | 1.331203 | -0.56591 | 0.571456 | 0.906105 |
| l(3)72Dr | 118.9076 | 1.396243 | 2.47412 | 0.564339 | 0.572523 | 0.906105 |
| lncRNA:CR | 87.6723 | -2.23538 | 3.951972 | -0.56564 | 0.571641 | 0.906105 |
| Oatp74D | 1499.166 | -0.56676 | 1.005113 | -0.56388 | 0.572835 | 0.906105 |
| Msh6 | 137.5707 | -2.06882 | 3.662575 | -0.56485 | 0.572174 | 0.906105 |
| Pp4-19C | 686.8741 | -0.75003 | 1.328049 | -0.56476 | 0.572236 | 0.906105 |
| fred | 1809.57 | -0.45289 | 0.802209 | -0.56456 | 0.572376 | 0.906105 |
| ND-B14.5E | 6284.807 | 0.309518 | 0.548736 | 0.564057 | 0.572715 | 0.906105 |
| ppk3 | 234.0417 | -2.00115 | 3.550188 | -0.56368 | 0.572975 | 0.906148 |
| RpL23 | 11132.57 | -0.25842 | 0.459402 | -0.56251 | 0.573765 | 0.907218 |
| cin | 397.4153 | 1.096578 | 1.951195 | 0.562003 | 0.574114 | 0.907491 |
| dup | 265.5584 | 1.311405 | 2.334623 | 0.56172 | 0.574307 | 0.907491 |
| PHDP | 188.4748 | 1.667361 | 2.967297 | 0.561912 | 0.574176 | 0.907491 |
| asRNA:CR4 | 96.98808 | -1.71325 | 3.050669 | -0.5616 | 0.57439 | 0.907491 |
| ND-MLRQ | 19126.51 | 0.256878 | 0.458043 | 0.560816 | 0.574923 | 0.907976 |
| PIP5K59B | 11827.71 | 0.26408 | 0.470862 | 0.560843 | 0.574905 | 0.907976 |
| UQCR-Q | 22276.06 | -0.2465 | 0.439673 | -0.56064 | 0.575046 | 0.907991 |
| mRpL44 | 1153.014 | -0.53412 | 0.95434 | -0.55967 | 0.575704 | 0.907994 |
| Oamb | 2611.857 | 0.487807 | 0.872094 | 0.559352 | 0.575922 | 0.907994 |
| RpL29 | 10707.6 | 0.260826 | 0.467286 | 0.558172 | 0.576727 | 0.907994 |
| Pgi | 5703.427 | -0.35245 | 0.631793 | -0.55785 | 0.576947 | 0.907994 |
| Nup153 | 1698.782 | 0.500002 | 0.89553 | 0.558331 | 0.576618 | 0.907994 |
| CG8475 | 5263.98 | -0.38549 | 0.689811 | -0.55883 | 0.57628 | 0.907994 |
| CG34351 | 732.9684 | 0.917965 | 1.644698 | 0.558136 | 0.576751 | 0.907994 |
| cnir | 1299.647 | -0.51025 | 0.914738 | -0.55781 | 0.576971 | 0.907994 |
| CG5004 | 523.2354 | -0.9324 | 1.670516 | -0.55815 | 0.576741 | 0.907994 |
| Syt14 | 853.7427 | 0.760943 | 1.358961 | 0.559945 | 0.575517 | 0.907994 |
| Obp58b | 108.1112 | -2.21116 | 3.951753 | -0.55954 | 0.575794 | 0.907994 |
| CG13594 | 1540.011 | -0.49236 | 0.880077 | -0.55945 | 0.575855 | 0.907994 |
| Cyt-b5 | 4304.404 | -0.31815 | 0.568054 | -0.56008 | 0.575428 | 0.907994 |
| asRNA:CR4 | 72.19286 | 1.564787 | 2.803909 | 0.558073 | 0.576794 | 0.907994 |
| AsnS | 237.9015 | -1.46685 | 2.61832 | -0.56023 | 0.575325 | 0.907994 |
| CG1317 | 3439.364 | -0.3429 | 0.614131 | -0.55835 | 0.576604 | 0.907994 |

| | | | | | | |
|---|---|---|---|---|---|---|
| msk | 1924.372 | 0.6272 | 1.124355 | 0.557831 | 0.57696 | 0.907994 |
| CG14903 | 1501.457 | -0.48748 | 0.874788 | -0.55725 | 0.577357 | 0.908423 |
| mRpL18 | 1035.061 | 0.542622 | 0.975658 | 0.55616 | 0.578101 | 0.909238 |
| tim | 8717.485 | -0.3482 | 0.625988 | -0.55624 | 0.578045 | 0.909238 |
| Mmp2 | 1649.804 | -0.44377 | 0.799142 | -0.55531 | 0.578683 | 0.909512 |
| Jhl-21 | 953.1731 | -0.80837 | 1.456098 | -0.55516 | 0.578783 | 0.909512 |
| E(bx) | 6475.824 | 0.285229 | 0.51347 | 0.555492 | 0.578558 | 0.909512 |
| eIF1 | 10722.28 | 0.25013 | 0.450622 | 0.555077 | 0.578842 | 0.909512 |
| Rok | 1095.829 | -0.53715 | 0.967694 | -0.55508 | 0.57884 | 0.909512 |
| RpL4 | 13085.34 | -0.26282 | 0.473922 | -0.55456 | 0.579194 | 0.909553 |
| RpS3A | 21747.09 | 0.229402 | 0.413788 | 0.554394 | 0.579309 | 0.909553 |
| CG2852 | 14793.33 | -0.25239 | 0.455338 | -0.55429 | 0.57938 | 0.909553 |
| tau | 9056.827 | 0.263575 | 0.475587 | 0.55421 | 0.579435 | 0.909553 |
| CG7101 | 352.1138 | -1.02808 | 1.853286 | -0.55473 | 0.579076 | 0.909553 |
| Raf | 3720.565 | 0.371187 | 0.671337 | 0.552908 | 0.580327 | 0.909719 |
| Nlg2 | 5901.994 | 0.314092 | 0.568193 | 0.552791 | 0.580406 | 0.909719 |
| cni | 2584.783 | 0.383767 | 0.69679 | 0.550764 | 0.581795 | 0.909719 |
| hook | 3835.408 | 0.490784 | 0.893922 | 0.549023 | 0.58299 | 0.909719 |
| Pld3 | 466.8465 | 0.893895 | 1.614864 | 0.553542 | 0.579892 | 0.909719 |
| alpha-Est4 | 281.5022 | -1.83985 | 3.346726 | -0.54975 | 0.582493 | 0.909719 |
| GCC88 | 432.9246 | 0.950168 | 1.716028 | 0.553702 | 0.579783 | 0.909719 |
| Pde6 | 2991.975 | -0.40993 | 0.745185 | -0.55011 | 0.582243 | 0.909719 |
| CG6028 | 807.5304 | -0.94254 | 1.706985 | -0.55217 | 0.580834 | 0.909719 |
| CG6763 | 520.2641 | 1.540615 | 2.7874 | 0.552707 | 0.580464 | 0.909719 |
| RIOK2 | 73.01506 | -1.64373 | 2.981575 | -0.55129 | 0.581432 | 0.909719 |
| neo | 189.1277 | 1.735849 | 3.163466 | 0.548717 | 0.583199 | 0.909719 |
| CG3168 | 2174.649 | -0.40319 | 0.733711 | -0.54952 | 0.582646 | 0.909719 |
| Caf1-180 | 268.5966 | 1.334009 | 2.425586 | 0.549974 | 0.582337 | 0.909719 |
| ZAP3 | 1185.717 | -0.733 | 1.327089 | -0.55234 | 0.580718 | 0.909719 |
| CG33199 | 1077.703 | 0.560722 | 1.021131 | 0.549118 | 0.582924 | 0.909719 |
| CCAP-R | 655.598 | 0.744669 | 1.357403 | 0.548598 | 0.583281 | 0.909719 |
| rumi | 101.7461 | 2.022817 | 3.658837 | 0.552858 | 0.580361 | 0.909719 |
| Desat1 | 25254.21 | -0.22576 | 0.408101 | -0.5532 | 0.580123 | 0.909719 |
| Tudor-SN | 6729.913 | -0.42389 | 0.768203 | -0.55179 | 0.581091 | 0.909719 |
| CG12502 | 1467.978 | -0.49491 | 0.900917 | -0.54934 | 0.582775 | 0.909719 |
| asRNA:CR4 | 313.504 | 1.495169 | 2.715161 | 0.550674 | 0.581857 | 0.909719 |
| Tim23 | 5415.179 | -0.29787 | 0.53825 | -0.55341 | 0.579985 | 0.909719 |
| CG1136 | 201.8391 | -1.83742 | 3.330524 | -0.55169 | 0.581159 | 0.909719 |
| Lsd-2 | 10566.13 | -0.28759 | 0.521822 | -0.55112 | 0.581548 | 0.909719 |
| Obp18a | 98.06753 | 2.042659 | 3.720557 | 0.54902 | 0.582992 | 0.909719 |
| CG32225 | 8480.117 | -0.26639 | 0.484126 | -0.55026 | 0.582143 | 0.909719 |
| CG12713 | 328.7317 | 1.525419 | 2.778807 | 0.548948 | 0.583041 | 0.909719 |
| CG4328 | 172.5487 | 1.409921 | 2.565679 | 0.549531 | 0.582641 | 0.909719 |
| RpL10Ab | 15370.47 | 0.235119 | 0.4264 | 0.551404 | 0.581357 | 0.909719 |
| CG4942 | 1215.611 | 0.977077 | 1.772264 | 0.551316 | 0.581417 | 0.909719 |
| mrva | 4798.098 | 0.334759 | 0.610163 | 0.548638 | 0.583254 | 0.909719 |
| Su(dx) | 5046.235 | -0.33151 | 0.600864 | -0.55172 | 0.581143 | 0.909719 |
| asRNA:CR4 | 220.1415 | -1.31778 | 2.403381 | -0.5483 | 0.583485 | 0.90986 |
| Spase25 | 986.6054 | -0.65139 | 1.18896 | -0.54786 | 0.583785 | 0.910152 |
| RpIII128 | 190.2333 | 1.246778 | 2.280728 | 0.546658 | 0.584614 | 0.910844 |

| | | | | | | |
|---|---|---|---|---|---|---|
| CG33229 | 1128.666 | 0.646251 | 1.182404 | 0.546557 | 0.584683 | 0.910844 |
| CG2186 | 3173.03 | -0.42079 | 0.769485 | -0.54684 | 0.584486 | 0.910844 |
| Bmcp | 494.142 | -0.90134 | 1.649045 | -0.54658 | 0.584665 | 0.910844 |
| Als2 | 1050.736 | 0.738587 | 1.354624 | 0.545234 | 0.585593 | 0.910858 |
| Mst85C | 1471.162 | -0.50253 | 0.922668 | -0.54465 | 0.585992 | 0.910858 |
| CG8176 | 4054.621 | -0.33378 | 0.612478 | -0.54497 | 0.585773 | 0.910858 |
| AOX3 | 504.2757 | -1.0929 | 2.002882 | -0.54566 | 0.585297 | 0.910858 |
| CG11857 | 2955.172 | -0.37365 | 0.685435 | -0.54512 | 0.585669 | 0.910858 |
| CG15098 | 8885.937 | 0.265873 | 0.487248 | 0.545663 | 0.585297 | 0.910858 |
| CG30116 | 1832.459 | 0.445349 | 0.817495 | 0.544773 | 0.58591 | 0.910858 |
| CG3295 | 2765.208 | 0.380088 | 0.69797 | 0.544563 | 0.586054 | 0.910858 |
| mRpL42 | 1429.923 | 0.470008 | 0.862521 | 0.544924 | 0.585806 | 0.910858 |
| pan | 39272.61 | -0.20846 | 0.381611 | -0.54626 | 0.584886 | 0.910858 |
| Glg1 | 1402.096 | 0.681417 | 1.251269 | 0.544581 | 0.586042 | 0.910858 |
| mv | 1419.02 | 0.586517 | 1.076849 | 0.54466 | 0.585987 | 0.910858 |
| DIP-alpha | 245.0775 | 1.23545 | 2.270158 | 0.544213 | 0.586295 | 0.910879 |
| beta-Spec | 6011.778 | -0.32005 | 0.588001 | -0.5443 | 0.586235 | 0.910879 |
| BHD | 360.8405 | 1.136453 | 2.090425 | 0.543647 | 0.586684 | 0.911308 |
| Nsun2 | 241.9519 | -1.07742 | 1.984971 | -0.54279 | 0.587274 | 0.911375 |
| CG9391 | 1712.524 | -0.43359 | 0.798697 | -0.54287 | 0.587222 | 0.911375 |
| TTLL6B | 155.2959 | -1.97222 | 3.634791 | -0.54259 | 0.587409 | 0.911375 |
| Aldh-III | 6007.822 | -0.37417 | 0.68882 | -0.5432 | 0.586991 | 0.911375 |
| CG33111 | 420.8793 | 1.361579 | 2.507626 | 0.542975 | 0.587147 | 0.911375 |
| Hsc70-4 | 20559.47 | -0.22616 | 0.416803 | -0.5426 | 0.587406 | 0.911375 |
| CG8336 | 557.7202 | 1.083342 | 1.997597 | 0.542323 | 0.587596 | 0.91149 |
| Shmt | 413.4624 | 1.308797 | 2.420789 | 0.540649 | 0.58875 | 0.911517 |
| PNKP | 474.9776 | 0.981361 | 1.814141 | 0.540951 | 0.588542 | 0.911517 |
| CG8202 | 1403.347 | 0.946673 | 1.748559 | 0.541402 | 0.588231 | 0.911517 |
| EMC2A | 1401.548 | 0.483311 | 0.893613 | 0.54085 | 0.588611 | 0.911517 |
| CG13603 | 7337.102 | 0.278764 | 0.514985 | 0.541306 | 0.588297 | 0.911517 |
| atl | 1519.318 | 0.469752 | 0.868326 | 0.540986 | 0.588518 | 0.911517 |
| Cul4 | 1369.037 | 0.522989 | 0.965913 | 0.541445 | 0.588201 | 0.911517 |
| mEFTu2 | 453.7255 | -0.74036 | 1.369304 | -0.54069 | 0.588723 | 0.911517 |
| Fer3HCH | 65.70623 | 1.904294 | 3.516672 | 0.541505 | 0.58816 | 0.911517 |
| Sfxn2 | 5340.546 | 0.32137 | 0.594203 | 0.540842 | 0.588616 | 0.911517 |
| kz | 219.5307 | -1.39611 | 2.58389 | -0.54031 | 0.588982 | 0.911526 |
| HDAC6 | 6939.817 | -0.31654 | 0.585712 | -0.54044 | 0.588895 | 0.911526 |
| SPARC | 10148.19 | -0.32134 | 0.594999 | -0.54006 | 0.589155 | 0.911543 |
| robl | 5408.141 | -0.29895 | 0.553643 | -0.53997 | 0.58922 | 0.911543 |
| Cyp1 | 24445.97 | -0.2346 | 0.434852 | -0.53949 | 0.589547 | 0.911872 |
| CG4646 | 1373.639 | 0.478431 | 0.887356 | 0.539165 | 0.589773 | 0.912046 |
| Tsp42Eg | 146.6801 | 1.549415 | 2.877363 | 0.538484 | 0.590243 | 0.91207 |
| ncm | 1240.341 | -0.50699 | 0.941077 | -0.53873 | 0.590073 | 0.91207 |
| CG42238 | 1591.992 | -0.43874 | 0.814091 | -0.53894 | 0.58993 | 0.91207 |
| mRpL39 | 1464.947 | -0.75681 | 1.405351 | -0.53852 | 0.59022 | 0.91207 |
| CG1907 | 2751.198 | -0.42648 | 0.792376 | -0.53823 | 0.59042 | 0.912168 |
| Spt3 | 480.0332 | 1.092098 | 2.029964 | 0.537989 | 0.590585 | 0.912246 |
| G9a | 2358.262 | -0.37994 | 0.714621 | -0.53167 | 0.594958 | 0.912265 |
| Ugalt | 263.6819 | -0.9599 | 1.793892 | -0.53509 | 0.592586 | 0.912265 |
| ICA69 | 2173.356 | -0.67673 | 1.273046 | -0.53159 | 0.595012 | 0.912265 |

| | | | | | | |
|---|---|---|---|---|---|---|
| ND-51 | 11846.07 | -0.25812 | 0.48567 | -0.53148 | 0.595087 | 0.912265 |
| cdc14 | 1120.015 | -0.50259 | 0.937487 | -0.5361 | 0.591888 | 0.912265 |
| mon2 | 1652.183 | -0.42074 | 0.791392 | -0.53165 | 0.594969 | 0.912265 |
| mRpS7 | 1296.464 | -0.48766 | 0.919834 | -0.53016 | 0.595998 | 0.912265 |
| Mal-B1 | 147.74 | -1.95811 | 3.653127 | -0.53601 | 0.591953 | 0.912265 |
| CG4631 | 792.6926 | -0.58273 | 1.091205 | -0.53402 | 0.593328 | 0.912265 |
| Nedd8 | 2405.387 | -0.36663 | 0.6867 | -0.5339 | 0.593413 | 0.912265 |
| TfIIFalpha | 254.8155 | -1.25959 | 2.352212 | -0.53549 | 0.592309 | 0.912265 |
| CG4511 | 838.585 | -0.77618 | 1.461889 | -0.53094 | 0.595459 | 0.912265 |
| CG17187 | 433.8071 | -0.98523 | 1.858476 | -0.53013 | 0.596024 | 0.912265 |
| Hsp70Bb | 76.42786 | -1.49978 | 2.794266 | -0.53673 | 0.591452 | 0.912265 |
| Ace | 29394.31 | 0.227682 | 0.425706 | 0.534834 | 0.592765 | 0.912265 |
| Trap1 | 2827.553 | -0.42254 | 0.78987 | -0.53495 | 0.592685 | 0.912265 |
| CG9945 | 1656.906 | -0.75948 | 1.425033 | -0.53295 | 0.594065 | 0.912265 |
| GstE6 | 5402.919 | -0.37967 | 0.716087 | -0.5302 | 0.595973 | 0.912265 |
| CG5174 | 12014.42 | 0.256428 | 0.477421 | 0.537111 | 0.591191 | 0.912265 |
| HPS4 | 1605.634 | 0.694669 | 1.305694 | 0.532031 | 0.594705 | 0.912265 |
| sca | 359.6923 | 1.074884 | 2.00695 | 0.535581 | 0.592248 | 0.912265 |
| Roc2 | 1731.204 | 0.456844 | 0.854173 | 0.534838 | 0.592762 | 0.912265 |
| CG13743 | 7029.286 | 0.313643 | 0.588407 | 0.533038 | 0.594007 | 0.912265 |
| TBPH | 3250.64 | -0.41839 | 0.788479 | -0.53062 | 0.59568 | 0.912265 |
| Tm1 | 9670.058 | -0.26468 | 0.498644 | -0.53081 | 0.595552 | 0.912265 |
| udt | 342.3257 | -1.25507 | 2.342284 | -0.53583 | 0.592076 | 0.912265 |
| B52 | 10388.76 | -0.3796 | 0.713546 | -0.53199 | 0.594733 | 0.912265 |
| CG41099 | 5452.455 | -0.31737 | 0.594558 | -0.53379 | 0.593487 | 0.912265 |
| CG34132 | 1116.831 | -0.68261 | 1.287363 | -0.53024 | 0.595948 | 0.912265 |
| CG40045 | 8657.394 | 0.260761 | 0.490685 | 0.531423 | 0.595126 | 0.912265 |
| NKAIN | 14650.92 | 0.265992 | 0.498243 | 0.533859 | 0.593439 | 0.912265 |
| cbs | 1462.098 | 0.797695 | 1.505452 | 0.529871 | 0.596201 | 0.912265 |
| CG42235 | 9694.976 | 0.318412 | 0.601052 | 0.529758 | 0.59628 | 0.912265 |
| SdhA | 17981.63 | 0.244339 | 0.455086 | 0.536908 | 0.591331 | 0.912265 |
| Pdk1 | 6008.304 | -0.31154 | 0.58456 | -0.53295 | 0.594066 | 0.912265 |
| CG9205 | 1840.298 | -0.4078 | 0.767261 | -0.5315 | 0.595071 | 0.912265 |
| mRpL45 | 2216.939 | 0.379244 | 0.711832 | 0.532772 | 0.594192 | 0.912265 |
| ocm | 601.6342 | 0.79287 | 1.48855 | 0.532646 | 0.594279 | 0.912265 |
| asRNA:CR4 | 680.4666 | -0.62763 | 1.176682 | -0.53339 | 0.593766 | 0.912265 |
| CG32278 | 5151.602 | 0.292271 | 0.545486 | 0.5358 | 0.592097 | 0.912265 |
| Fit1 | 2266.845 | -0.44873 | 0.839034 | -0.53482 | 0.592773 | 0.912265 |
| Set2 | 1344.442 | -0.46097 | 0.86963 | -0.53007 | 0.596061 | 0.912265 |
| ND-18 | 5139.417 | -0.28681 | 0.541043 | -0.5301 | 0.596041 | 0.912265 |
| CG5589 | 317.8726 | 1.050335 | 1.978506 | 0.530873 | 0.595507 | 0.912265 |
| sina | 3161.442 | -0.33687 | 0.633039 | -0.53215 | 0.594621 | 0.912265 |
| Nrx-IV | 3510.566 | 0.429056 | 0.801368 | 0.535404 | 0.59237 | 0.912265 |
| Taf2 | 287.7418 | 1.436506 | 2.709316 | 0.53021 | 0.595966 | 0.912265 |
| RpS17 | 8054.606 | 0.295456 | 0.55096 | 0.536258 | 0.59178 | 0.912265 |
| Cpsf6 | 1252.458 | -0.51031 | 0.951455 | -0.53634 | 0.591722 | 0.912265 |
| fliI | 176.35 | 1.330642 | 2.503058 | 0.531607 | 0.594999 | 0.912265 |
| RpA-70 | 1830.914 | 0.433364 | 0.81928 | 0.528957 | 0.596836 | 0.912351 |
| dpr14 | 3318.634 | -0.35235 | 0.66617 | -0.52892 | 0.596859 | 0.912351 |
| CG31324 | 4504.729 | -0.29582 | 0.558978 | -0.52922 | 0.59665 | 0.912351 |

| | | | | | | |
|---|---|---|---|---|---|---|
| Mid1 | 5637.93 | -0.29843 | 0.564283 | -0.52886 | 0.596904 | 0.912351 |
| CG3632 | 2417.178 | 0.491153 | 0.928009 | 0.529255 | 0.596629 | 0.912351 |
| CanB2 | 4569.07 | 0.349798 | 0.66205 | 0.528356 | 0.597252 | 0.912474 |
| lncRNA:CR | 731.6116 | -0.62054 | 1.174714 | -0.52825 | 0.597326 | 0.912474 |
| CG32204 | 590.513 | 0.667008 | 1.262489 | 0.528328 | 0.597272 | 0.912474 |
| asRNA:CR4 | 2932.118 | 0.345643 | 0.654537 | 0.528073 | 0.597448 | 0.912487 |
| DOR | 7345.861 | -0.27944 | 0.529653 | -0.52759 | 0.597786 | 0.91283 |
| Sym | 493.6207 | -0.9948 | 1.889666 | -0.52644 | 0.598582 | 0.913761 |
| CG9330 | 2300.206 | 0.402138 | 0.763968 | 0.526381 | 0.598624 | 0.913761 |
| CCT5 | 959.7605 | -0.70022 | 1.331836 | -0.52575 | 0.59906 | 0.914253 |
| Sema1a | 17994.51 | -0.21945 | 0.417903 | -0.52513 | 0.599493 | 0.914739 |
| nrv3 | 84112.69 | 0.210773 | 0.402006 | 0.524303 | 0.600068 | 0.915095 |
| ci | 1514.654 | 0.562129 | 1.071571 | 0.524584 | 0.599872 | 0.915095 |
| CG3700 | 552.6148 | -0.97854 | 1.866126 | -0.52437 | 0.600023 | 0.915095 |
| RpL27A | 18432.33 | 0.234434 | 0.447368 | 0.524029 | 0.600258 | 0.915211 |
| gzl | 4843.716 | -0.28755 | 0.549622 | -0.52317 | 0.600854 | 0.915826 |
| Scm | 911.0006 | 0.845115 | 1.616098 | 0.522935 | 0.601019 | 0.915826 |
| CG6454 | 2138.414 | 0.426704 | 0.815956 | 0.52295 | 0.601009 | 0.915826 |
| Impbeta11 | 573.1419 | 0.905767 | 1.732551 | 0.522794 | 0.601118 | 0.915826 |
| Eip63E | 10597.98 | 0.262892 | 0.503158 | 0.522483 | 0.601334 | 0.915981 |
| tRNA:Thr-/ | 252.8357 | -1.80747 | 3.464942 | -0.52164 | 0.601918 | 0.916201 |
| eIF2Bgamr | 482.3166 | 0.914808 | 1.754334 | 0.521456 | 0.602049 | 0.916201 |
| CG17803 | 223.2137 | 1.5135 | 2.899021 | 0.522073 | 0.60162 | 0.916201 |
| CG6044 | 11884.89 | -0.242 | 0.464051 | -0.5215 | 0.60202 | 0.916201 |
| rho-4 | 156.7972 | 1.181444 | 2.265092 | 0.521588 | 0.601958 | 0.916201 |
| CG5521 | 2157.247 | 0.400411 | 0.768468 | 0.521051 | 0.602331 | 0.916351 |
| Mon1 | 455.491 | 1.196584 | 2.296764 | 0.520987 | 0.602376 | 0.916351 |
| nis | 92.96133 | -1.99631 | 3.836251 | -0.52038 | 0.602798 | 0.916819 |
| beta-Man | 1093.984 | 0.929677 | 1.787099 | 0.520216 | 0.602913 | 0.916821 |
| Rpb10 | 1245.485 | -0.50845 | 0.979117 | -0.51929 | 0.603558 | 0.91728 |
| Psi | 2333.802 | 0.36855 | 0.709476 | 0.519468 | 0.603435 | 0.91728 |
| fidipidine | 632.622 | 0.669145 | 1.288509 | 0.519317 | 0.60354 | 0.91728 |
| SoxN | 515.0276 | 0.861925 | 1.660652 | 0.519028 | 0.603741 | 0.917385 |
| CG14024 | 2106.726 | -0.48735 | 0.940524 | -0.51817 | 0.604342 | 0.918049 |
| CG13999 | 2983.529 | -0.41621 | 0.803659 | -0.51789 | 0.604535 | 0.918049 |
| Usp14 | 1995.183 | 0.403765 | 0.780346 | 0.517418 | 0.604865 | 0.918049 |
| clt | 1242.197 | 0.493889 | 0.954096 | 0.517651 | 0.604702 | 0.918049 |
| CG8765 | 1410.146 | -0.78853 | 1.523594 | -0.51755 | 0.604775 | 0.918049 |
| Pten | 2553.55 | -0.37084 | 0.716513 | -0.51757 | 0.604761 | 0.918049 |
| SP2353 | 3436.481 | 0.479241 | 0.927473 | 0.516717 | 0.605353 | 0.91827 |
| Cyp6g1 | 75.86562 | 1.982047 | 3.834743 | 0.516866 | 0.60525 | 0.91827 |
| CSN1b | 2350.092 | -0.45016 | 0.870936 | -0.51687 | 0.605243 | 0.91827 |
| Cnot4 | 3416.287 | 0.359414 | 0.696968 | 0.515682 | 0.606076 | 0.918472 |
| Ald1 | 23532.39 | -0.21361 | 0.414268 | -0.51562 | 0.60612 | 0.918472 |
| Fer2LCH | 41244.83 | -0.1996 | 0.386823 | -0.51601 | 0.605846 | 0.918472 |
| CG34293 | 640.0554 | 0.612967 | 1.188331 | 0.515821 | 0.605979 | 0.918472 |
| Shab | 15722.82 | 0.25892 | 0.501747 | 0.516037 | 0.605829 | 0.918472 |
| mRpL48 | 1866.727 | -0.42685 | 0.827968 | -0.51554 | 0.606173 | 0.918472 |
| Cyp4p3 | 222.5968 | -1.27753 | 2.484086 | -0.51429 | 0.607052 | 0.918909 |
| Dgk | 1589.243 | -0.44976 | 0.875878 | -0.51349 | 0.607607 | 0.918909 |

| | | | | | | |
|---|---|---|---|---|---|---|
| CG42376 | 2137.544 | 0.378096 | 0.735763 | 0.513883 | 0.607334 | 0.918909 |
| GLS | 1843.482 | -0.65818 | 1.280766 | -0.5139 | 0.607323 | 0.918909 |
| Su(z)2 | 1482.001 | -0.48634 | 0.945439 | -0.51441 | 0.606965 | 0.918909 |
| Gfat1 | 159.4373 | 1.79138 | 3.487103 | 0.513716 | 0.607451 | 0.918909 |
| GlyRS | 1549.614 | -0.41971 | 0.815086 | -0.51493 | 0.6066 | 0.918909 |
| Sox21a | 159.9533 | 1.300568 | 2.532236 | 0.513604 | 0.607529 | 0.918909 |
| CG18178 | 142.3691 | -1.65774 | 3.222965 | -0.51435 | 0.607005 | 0.918909 |
| bip1 | 1581.575 | -0.70626 | 1.374357 | -0.51389 | 0.607331 | 0.918909 |
| bnl | 400.4757 | 0.727465 | 1.417918 | 0.513052 | 0.607915 | 0.918982 |
| Ect3 | 978.3422 | 0.790534 | 1.540546 | 0.513152 | 0.607845 | 0.918982 |
| Tet | 16434.95 | -0.22147 | 0.432041 | -0.51261 | 0.608227 | 0.918982 |
| MAN1 | 891.9826 | -0.62343 | 1.215692 | -0.51282 | 0.608076 | 0.918982 |
| CR43972 | 236.3432 | 1.206871 | 2.353858 | 0.51272 | 0.608147 | 0.918982 |
| Vap33 | 9014.788 | -0.25794 | 0.503425 | -0.51238 | 0.608387 | 0.919051 |
| Arl2 | 367.7509 | 1.128063 | 2.207291 | 0.511062 | 0.609307 | 0.919092 |
| Elk | 1410.185 | 0.790818 | 1.547501 | 0.511029 | 0.609331 | 0.919092 |
| CG6999 | 494.8212 | -0.70617 | 1.378975 | -0.5121 | 0.608582 | 0.919092 |
| Ost48 | 3686.306 | 0.356908 | 0.697427 | 0.511749 | 0.608826 | 0.919092 |
| Mvl | 2634.86 | -0.34031 | 0.665514 | -0.51135 | 0.609105 | 0.919092 |
| h | 1251.816 | -0.56342 | 1.102492 | -0.51104 | 0.609321 | 0.919092 |
| CG14989 | 19622.81 | -0.30276 | 0.591992 | -0.51143 | 0.609053 | 0.919092 |
| CG12194 | 3220.397 | 0.446323 | 0.873267 | 0.511096 | 0.609284 | 0.919092 |
| robo2 | 667.7428 | -0.9029 | 1.76768 | -0.51078 | 0.609504 | 0.91918 |
| Arf102F | 7821.955 | -0.25742 | 0.504293 | -0.51045 | 0.609738 | 0.919361 |
| Pka-C1 | 15836.22 | 0.229521 | 0.450097 | 0.509936 | 0.610096 | 0.919555 |
| CG2811 | 952.2469 | -0.50269 | 0.985597 | -0.51004 | 0.610025 | 0.919555 |
| Rbbp5 | 470.6698 | 0.976184 | 1.922829 | 0.507681 | 0.611677 | 0.919931 |
| Pect | 7121.028 | 0.353741 | 0.697275 | 0.50732 | 0.611931 | 0.919931 |
| beta'COP | 1475.349 | 0.427454 | 0.843162 | 0.506965 | 0.612179 | 0.919931 |
| CG18428 | 813.1736 | 0.600475 | 1.17989 | 0.508924 | 0.610805 | 0.919931 |
| CHKov1 | 667.7485 | -0.59394 | 1.166684 | -0.50909 | 0.610692 | 0.919931 |
| CG4409 | 22676.22 | -0.22752 | 0.448286 | -0.50754 | 0.611778 | 0.919931 |
| mael | 110.1501 | 1.564121 | 3.075386 | 0.508593 | 0.611037 | 0.919931 |
| hoe1 | 10216.2 | 0.245536 | 0.483337 | 0.508001 | 0.611452 | 0.919931 |
| CkIIbeta | 9831.611 | 0.238809 | 0.47104 | 0.506983 | 0.612167 | 0.919931 |
| CG42390 | 1191.73 | -0.4825 | 0.949927 | -0.50793 | 0.6115 | 0.919931 |
| CG42542 | 5298.268 | 0.309758 | 0.610919 | 0.507036 | 0.61213 | 0.919931 |
| SmE | 1458.28 | -0.4607 | 0.906503 | -0.50822 | 0.611301 | 0.919931 |
| COX8 | 28840.4 | 0.212604 | 0.418231 | 0.508342 | 0.611214 | 0.919931 |
| Pfdn2 | 2278.564 | 0.373611 | 0.734828 | 0.508433 | 0.61115 | 0.919931 |
| CG13679 | 73.07235 | 1.995885 | 3.925989 | 0.508378 | 0.611189 | 0.919931 |
| Neos | 421.3694 | 1.116105 | 2.197766 | 0.507836 | 0.611568 | 0.919931 |
| CG42654 | 151.6401 | 1.639093 | 3.235376 | 0.506616 | 0.612424 | 0.919955 |
| ZnT63C | 2581.313 | -0.46305 | 0.913889 | -0.50668 | 0.61238 | 0.919955 |
| Sirt4 | 83.29235 | 1.887941 | 3.729326 | 0.506242 | 0.612687 | 0.920177 |
| Ccs | 634.5602 | -1.01578 | 2.007939 | -0.50588 | 0.612941 | 0.920213 |
| CG42324 | 4671.213 | 0.353219 | 0.698077 | 0.505988 | 0.612865 | 0.920213 |
| CG7231 | 2361.182 | -0.36584 | 0.723461 | -0.50568 | 0.613081 | 0.920252 |
| NPFR | 169.6602 | 1.231996 | 2.437514 | 0.505431 | 0.613256 | 0.920343 |
| CG31219 | 1864.681 | -0.45742 | 0.907378 | -0.50411 | 0.614185 | 0.920653 |

| | | | | | | |
|---|---|---|---|---|---|---|
| CG3618 | 3222.131 | -0.32557 | 0.645628 | -0.50427 | 0.614074 | 0.920653 |
| Ykt6 | 3893.736 | 0.310052 | 0.614759 | 0.504347 | 0.614018 | 0.920653 |
| cnc | 2900.154 | -0.38022 | 0.754284 | -0.50408 | 0.614207 | 0.920653 |
| ppk29 | 188.8585 | 1.296179 | 2.571199 | 0.504114 | 0.614181 | 0.920653 |
| lncRNA:CR | 242.7056 | 1.025797 | 2.035064 | 0.504061 | 0.614218 | 0.920653 |
| CG46338 | 798.7413 | 0.906471 | 1.798575 | 0.503994 | 0.614266 | 0.920653 |
| mthl13 | 574.3111 | 0.901106 | 1.788733 | 0.503768 | 0.614424 | 0.920719 |
| Trim9 | 435.4104 | 0.689373 | 1.369222 | 0.503478 | 0.614628 | 0.920853 |
| Npc2a | 8166.046 | -0.2479 | 0.492638 | -0.5032 | 0.614824 | 0.920973 |
| CG11638 | 642.3112 | 0.701698 | 1.396505 | 0.502467 | 0.615339 | 0.921212 |
| Lip4 | 195.805 | 1.664534 | 3.313432 | 0.50236 | 0.615415 | 0.921212 |
| Oda | 31555.69 | -0.20543 | 0.408633 | -0.50272 | 0.615158 | 0.921212 |
| CNPYb | 1440.876 | 0.429884 | 0.855797 | 0.50232 | 0.615442 | 0.921212 |
| CG7378 | 2989.44 | -0.43444 | 0.86518 | -0.50214 | 0.615567 | 0.921227 |
| CG1513 | 7426.458 | 0.27276 | 0.543554 | 0.501808 | 0.615802 | 0.921236 |
| Tob | 18921.51 | 0.226463 | 0.4512 | 0.501912 | 0.61573 | 0.921236 |
| CG14688 | 4869.55 | -0.30022 | 0.598523 | -0.5016 | 0.615947 | 0.92128 |
| Naa30A | 626.225 | 0.718354 | 1.435386 | 0.50046 | 0.616751 | 0.92134 |
| Orco | 77.33376 | -1.8652 | 3.727386 | -0.5004 | 0.616791 | 0.92134 |
| Sirt2 | 1501.382 | 0.725363 | 1.447988 | 0.500946 | 0.616409 | 0.92134 |
| Chchd2 | 10724.61 | 0.227906 | 0.454745 | 0.501173 | 0.616249 | 0.92134 |
| CG13895 | 1525.277 | 0.454383 | 0.907961 | 0.500443 | 0.616763 | 0.92134 |
| mRpS5 | 3215.154 | 0.315945 | 0.630584 | 0.501036 | 0.616346 | 0.92134 |
| Csas | 7232.866 | -0.30871 | 0.616384 | -0.50084 | 0.616482 | 0.92134 |
| RpI135 | 599.5745 | 0.846736 | 1.696301 | 0.499166 | 0.617663 | 0.921352 |
| gcm2 | 130.3332 | -1.92555 | 3.867608 | -0.49787 | 0.618579 | 0.921352 |
| CG10338 | 1613.65 | 0.438964 | 0.88059 | 0.498488 | 0.61814 | 0.921352 |
| Taf13 | 1437.548 | 0.43251 | 0.865986 | 0.499443 | 0.617468 | 0.921352 |
| m-cup | 1143.903 | 0.682277 | 1.370678 | 0.497766 | 0.618649 | 0.921352 |
| CG6005 | 1578.464 | 0.414607 | 0.832916 | 0.497778 | 0.618641 | 0.921352 |
| CG2224 | 1078.659 | 0.675487 | 1.357429 | 0.497622 | 0.61875 | 0.921352 |
| Pfk | 6729.45 | 0.312116 | 0.625114 | 0.499295 | 0.617572 | 0.921352 |
| CR33317 | 150.5114 | -1.53503 | 3.07595 | -0.49904 | 0.617749 | 0.921352 |
| Gr64f | 206.8007 | -1.52147 | 3.052249 | -0.49847 | 0.61815 | 0.921352 |
| GC1 | 689.2692 | -0.57431 | 1.153308 | -0.49797 | 0.618505 | 0.921352 |
| Rbp1 | 11246.44 | 0.276089 | 0.552756 | 0.499477 | 0.617443 | 0.921352 |
| wdp | 5295.93 | 0.291034 | 0.581998 | 0.50006 | 0.617033 | 0.921352 |
| asRNA:CR4 | 387.3628 | -0.95236 | 1.910451 | -0.4985 | 0.618132 | 0.921352 |
| CG14073 | 2807.646 | -0.35491 | 0.709884 | -0.49996 | 0.617105 | 0.921352 |
| CG6650 | 1098.446 | -0.50902 | 1.021731 | -0.49819 | 0.618351 | 0.921352 |
| Syt1 | 62824.56 | 0.201237 | 0.402712 | 0.499705 | 0.617283 | 0.921352 |
| dpr5 | 1051.062 | -0.58228 | 1.171214 | -0.49716 | 0.619076 | 0.921362 |
| EMRE | 6973.578 | 0.331773 | 0.667944 | 0.496708 | 0.619395 | 0.921362 |
| mRNA-cap | 568.1252 | -0.72986 | 1.467396 | -0.49739 | 0.618917 | 0.921362 |
| CG10417 | 4215.549 | -0.3234 | 0.650998 | -0.49677 | 0.619348 | 0.921362 |
| CG5835 | 2419.747 | 0.370439 | 0.745662 | 0.496792 | 0.619335 | 0.921362 |
| CG14232 | 641.1547 | -0.88329 | 1.778552 | -0.49664 | 0.619445 | 0.921362 |
| 14-3-3epsi | 28066.74 | 0.222594 | 0.44864 | 0.496154 | 0.619786 | 0.921532 |
| CG32645 | 72.69529 | -1.93413 | 3.898283 | -0.49615 | 0.619789 | 0.921532 |
| Non3 | 669.6833 | -0.86249 | 1.740978 | -0.49541 | 0.620314 | 0.921883 |

| | | | | | | |
|---|---|---|---|---|---|---|
| LSm3 | 623.8639 | -0.90607 | 1.829812 | -0.49517 | 0.620478 | 0.921883 |
| OtopLc | 5290.493 | -0.2649 | 0.534978 | -0.49516 | 0.620485 | 0.921883 |
| Lpin | 9237.329 | 0.232489 | 0.46925 | 0.495449 | 0.620283 | 0.921883 |
| Phs | 172.6206 | -1.34788 | 2.725061 | -0.49462 | 0.620866 | 0.922278 |
| Sec16 | 2105.898 | -0.40498 | 0.819668 | -0.49408 | 0.621248 | 0.922676 |
| lncRNA:CR | 459.2512 | 0.782069 | 1.583598 | 0.493856 | 0.621408 | 0.922742 |
| Tk | 1352.538 | -0.63898 | 1.294324 | -0.49367 | 0.621536 | 0.922761 |
| Fas2 | 1751.171 | -0.49134 | 0.996115 | -0.49326 | 0.621832 | 0.922846 |
| mid | 82.93274 | -1.89088 | 3.834633 | -0.49311 | 0.621938 | 0.922846 |
| VhaM9.7-a | 14472.29 | 0.214391 | 0.434627 | 0.493275 | 0.621818 | 0.922846 |
| mRpL16 | 2269.749 | -0.34546 | 0.705021 | -0.49 | 0.624131 | 0.923139 |
| lin-52 | 1630.936 | -0.41763 | 0.857269 | -0.48716 | 0.626145 | 0.923139 |
| Cyp4ac2 | 161.8445 | 1.271577 | 2.588423 | 0.491256 | 0.623246 | 0.923139 |
| CG11070 | 1256.216 | -0.74337 | 1.512682 | -0.49143 | 0.623126 | 0.923139 |
| CG31760 | 1066.002 | -0.52145 | 1.059734 | -0.49206 | 0.622678 | 0.923139 |
| CG9265 | 2294.128 | -0.36614 | 0.747495 | -0.48982 | 0.624259 | 0.923139 |
| CG8526 | 318.4884 | -0.93943 | 1.922434 | -0.48867 | 0.625078 | 0.923139 |
| CG11626 | 77.66168 | -1.92136 | 3.95141 | -0.48625 | 0.626793 | 0.923139 |
| Rh3 | 103.1527 | -1.92179 | 3.951285 | -0.48637 | 0.626703 | 0.923139 |
| e | 5501.324 | -0.31938 | 0.654102 | -0.48826 | 0.625362 | 0.923139 |
| awd | 7909.452 | 0.247145 | 0.50686 | 0.487599 | 0.625834 | 0.923139 |
| CG6675 | 72.11442 | -1.92122 | 3.951448 | -0.48621 | 0.626821 | 0.923139 |
| CG30431 | 131.1059 | -1.73554 | 3.558884 | -0.48766 | 0.625789 | 0.923139 |
| CG4866 | 531.4741 | 0.862406 | 1.75806 | 0.490544 | 0.623749 | 0.923139 |
| l(1)G0320 | 4522.108 | 0.300039 | 0.614654 | 0.488144 | 0.625448 | 0.923139 |
| CARPB | 805.4343 | 0.69056 | 1.404443 | 0.491697 | 0.622934 | 0.923139 |
| Hmg-2 | 830.6368 | 0.801506 | 1.646771 | 0.486714 | 0.626461 | 0.923139 |
| CG5267 | 150.3054 | 1.198684 | 2.460432 | 0.487184 | 0.626128 | 0.923139 |
| Vha36-1 | 10188.81 | -0.23241 | 0.476372 | -0.48788 | 0.625633 | 0.923139 |
| AspRS | 122.0002 | 1.680853 | 3.448944 | 0.487353 | 0.626008 | 0.923139 |
| CG11635 | 143.2164 | 1.050252 | 2.141511 | 0.490426 | 0.623833 | 0.923139 |
| nec | 1618.555 | 0.592037 | 1.205599 | 0.491073 | 0.623375 | 0.923139 |
| eIF5 | 218.0954 | 0.87628 | 1.792907 | 0.488748 | 0.62502 | 0.923139 |
| beat-Va | 1811.016 | -0.40602 | 0.832899 | -0.48748 | 0.625916 | 0.923139 |
| Naa80 | 3942.808 | -0.31858 | 0.655288 | -0.48616 | 0.626851 | 0.923139 |
| CG33777 | 69.07663 | -1.92112 | 3.951472 | -0.48618 | 0.62684 | 0.923139 |
| sev | 1192.991 | -0.61547 | 1.256056 | -0.49 | 0.624135 | 0.923139 |
| CG17163 | 1633.287 | 0.435245 | 0.890143 | 0.488961 | 0.624869 | 0.923139 |
| nub | 148.3085 | -1.45181 | 2.968685 | -0.48904 | 0.624811 | 0.923139 |
| tef | 127.7191 | -1.92205 | 3.951212 | -0.48645 | 0.62665 | 0.923139 |
| l(2)37Cb | 818.6759 | 0.743358 | 1.524556 | 0.48759 | 0.62584 | 0.923139 |
| whip | 561.4844 | -0.61017 | 1.240268 | -0.49197 | 0.622742 | 0.923139 |
| RhoGAP10 | 2971.761 | -0.31783 | 0.64802 | -0.49046 | 0.623809 | 0.923139 |
| robo1 | 3407.561 | -0.29853 | 0.612901 | -0.48708 | 0.626198 | 0.923139 |
| SNF4Agam | 3436.041 | -0.31334 | 0.640493 | -0.48922 | 0.624686 | 0.923139 |
| CG45050 | 11287.69 | -0.21837 | 0.446151 | -0.48946 | 0.624515 | 0.923139 |
| CG7638 | 1776.237 | 0.41351 | 0.84427 | 0.489784 | 0.624287 | 0.923139 |
| pall | 642.935 | -0.66349 | 1.353502 | -0.4902 | 0.62399 | 0.923139 |
| Cyt-c1 | 11645.74 | 0.286663 | 0.58762 | 0.487837 | 0.625665 | 0.923139 |
| CG3165 | 320.9898 | 1.110054 | 2.27687 | 0.487535 | 0.625879 | 0.923139 |

| | | | | | | |
|---|---|---|---|---|---|---|
| Clp | 526.1658 | -1.06454 | 2.161274 | -0.49255 | 0.622329 | 0.923139 |
| CG15914 | 84.28416 | -1.89288 | 3.89715 | -0.48571 | 0.627174 | 0.923445 |
| sqh | 5496.382 | 0.25902 | 0.534457 | 0.484642 | 0.627931 | 0.923542 |
| CG5466 | 2550.215 | -0.37483 | 0.773138 | -0.48481 | 0.627809 | 0.923542 |
| CG18731 | 516.8489 | 0.687024 | 1.41615 | 0.485135 | 0.62758 | 0.923542 |
| mRpS28 | 930.3984 | -0.54231 | 1.118774 | -0.48473 | 0.627866 | 0.923542 |
| nito | 428.7514 | -0.75892 | 1.564459 | -0.4851 | 0.627606 | 0.923542 |
| trx | 9559.306 | 0.256878 | 0.529312 | 0.485305 | 0.62746 | 0.923542 |
| CanB | 23366.9 | 0.212272 | 0.440677 | 0.481694 | 0.630023 | 0.924005 |
| atms | 1556.84 | 0.40299 | 0.839355 | 0.480119 | 0.631143 | 0.924005 |
| Uhg2 | 547.6206 | 0.646485 | 1.345625 | 0.480435 | 0.630918 | 0.924005 |
| ScsbetaA | 12010.11 | -0.23561 | 0.487241 | -0.48355 | 0.628704 | 0.924005 |
| CG7059 | 148.7494 | 1.228164 | 2.553816 | 0.480913 | 0.630578 | 0.924005 |
| Ppox | 193.4762 | 1.055946 | 2.188092 | 0.482587 | 0.629389 | 0.924005 |
| Usp1 | 2173.861 | -0.47873 | 0.99738 | -0.47998 | 0.631239 | 0.924005 |
| CG2059 | 1423.072 | 0.632255 | 1.310816 | 0.482337 | 0.629567 | 0.924005 |
| CG13500 | 104.5781 | -1.90887 | 3.951277 | -0.4831 | 0.629022 | 0.924005 |
| Mhcl | 872.3046 | 0.510279 | 1.057152 | 0.482692 | 0.629314 | 0.924005 |
| TM4SF | 35723.44 | 0.203936 | 0.423077 | 0.482031 | 0.629784 | 0.924005 |
| CG3566 | 2280.927 | -0.34194 | 0.712241 | -0.4801 | 0.631159 | 0.924005 |
| CG34219 | 411.0125 | 0.949667 | 1.967721 | 0.482623 | 0.629364 | 0.924005 |
| Rab26 | 2577.23 | 0.361144 | 0.750192 | 0.481402 | 0.630231 | 0.924005 |
| kuz | 4731.64 | -0.28753 | 0.597591 | -0.48114 | 0.630416 | 0.924005 |
| nsl1 | 2820.483 | -0.33885 | 0.704392 | -0.48105 | 0.630479 | 0.924005 |
| CG30274 | 110.2574 | -1.9096 | 3.951258 | -0.48329 | 0.62889 | 0.924005 |
| spg | 4540.375 | 0.277236 | 0.57731 | 0.480219 | 0.631071 | 0.924005 |
| dop | 4725.037 | -0.35307 | 0.732949 | -0.48172 | 0.630007 | 0.924005 |
| Vha13 | 34086.37 | -0.1957 | 0.405752 | -0.48233 | 0.629575 | 0.924005 |
| CG33233 | 119.7133 | 1.720991 | 3.571616 | 0.481852 | 0.629911 | 0.924005 |
| CG11151 | 3825.763 | -0.28972 | 0.602968 | -0.4805 | 0.630875 | 0.924005 |
| RpL26 | 12016.09 | -0.22718 | 0.47067 | -0.48268 | 0.62932 | 0.924005 |
| CG32112 | 1329.172 | -0.49622 | 1.027123 | -0.48312 | 0.629012 | 0.924005 |
| Grip163 | 1387.669 | 0.455234 | 0.945439 | 0.481505 | 0.630158 | 0.924005 |
| CG5906 | 79.35124 | -1.90442 | 3.951393 | -0.48196 | 0.629834 | 0.924005 |
| Oli | 2256.167 | 0.342281 | 0.713594 | 0.479658 | 0.63147 | 0.924017 |
| Yeti | 8342.639 | 0.291226 | 0.607164 | 0.479649 | 0.631477 | 0.924017 |
| CG8929 | 4680.877 | 0.30326 | 0.633105 | 0.479005 | 0.631935 | 0.924274 |
| CG33017 | 140.4446 | 1.769451 | 3.694698 | 0.478916 | 0.631998 | 0.924274 |
| GramD1B | 4850.862 | 0.316057 | 0.659684 | 0.479103 | 0.631865 | 0.924274 |
| pns | 2379.008 | -0.35186 | 0.735174 | -0.47861 | 0.632214 | 0.924319 |
| Pka-C3 | 325.7479 | 1.001141 | 2.092033 | 0.478549 | 0.632259 | 0.924319 |
| bcn92 | 2221.956 | -0.3556 | 0.744148 | -0.47787 | 0.632745 | 0.924692 |
| fzr | 4653.165 | -0.28665 | 0.599769 | -0.47793 | 0.632697 | 0.924692 |
| CG31650 | 2991.767 | 0.36467 | 0.764444 | 0.47704 | 0.633334 | 0.92483 |
| CG13108 | 1361.396 | 0.433173 | 0.908077 | 0.477022 | 0.633347 | 0.92483 |
| mt:ND2 | 7564.282 | 0.243231 | 0.509999 | 0.476925 | 0.633415 | 0.92483 |
| mr | 627.1974 | 0.718215 | 1.505836 | 0.476954 | 0.633395 | 0.92483 |
| Dlic | 5150.066 | 0.327267 | 0.685626 | 0.477326 | 0.63313 | 0.92483 |
| Pp2C1 | 1099.115 | -0.48364 | 1.015613 | -0.47621 | 0.633925 | 0.924967 |
| Ufc1 | 1492.581 | -0.42813 | 0.898721 | -0.47637 | 0.633808 | 0.924967 |

| | | | | | | |
|---|---|---|---|---|---|---|
| lncRNA:CR | 3216.893 | 0.334914 | 0.703031 | 0.476385 | 0.6338 | 0.924967 |
| ine | 6554.791 | 0.401626 | 0.843492 | 0.476147 | 0.63397 | 0.924967 |
| DCTN4-p6. | 1798.349 | 0.378639 | 0.796429 | 0.47542 | 0.634487 | 0.925385 |
| parvin | 921.1112 | -0.62708 | 1.318906 | -0.47546 | 0.634462 | 0.925385 |
| G6P | 970.5973 | -0.73964 | 1.557025 | -0.47504 | 0.634761 | 0.925617 |
| CrebB | 544.2077 | 0.759697 | 1.599934 | 0.47483 | 0.634908 | 0.925663 |
| NaCP60E | 39181.12 | 0.200337 | 0.422164 | 0.474548 | 0.63511 | 0.925788 |
| Ank | 16090.06 | -0.23888 | 0.503953 | -0.47401 | 0.635493 | 0.926178 |
| SdhB | 1586.664 | -0.39475 | 0.833958 | -0.47335 | 0.635965 | 0.92653 |
| Ubqn | 2779.438 | -0.32164 | 0.679334 | -0.47346 | 0.635882 | 0.92653 |
| CG11050 | 3873.947 | 0.293907 | 0.625818 | 0.469636 | 0.638615 | 0.926691 |
| CG31817 | 331.4724 | 0.891122 | 1.895658 | 0.470086 | 0.638294 | 0.926691 |
| EloA | 474.7691 | 0.611346 | 1.301144 | 0.469853 | 0.63846 | 0.926691 |
| eEF2 | 22201.76 | -0.20573 | 0.436309 | -0.47153 | 0.637262 | 0.926691 |
| Ntan1 | 2765.994 | -0.33562 | 0.710525 | -0.47236 | 0.636669 | 0.926691 |
| spidey | 2665.009 | -0.41937 | 0.888888 | -0.47179 | 0.637077 | 0.926691 |
| Syn2 | 6236.944 | 0.261319 | 0.556244 | 0.469793 | 0.638503 | 0.926691 |
| CG18327 | 107.6761 | -1.62682 | 3.442717 | -0.47254 | 0.636542 | 0.926691 |
| Tapdelta | 2280.004 | 0.342189 | 0.726805 | 0.470813 | 0.637774 | 0.926691 |
| PCB | 1013.042 | -0.55239 | 1.172126 | -0.47127 | 0.637446 | 0.926691 |
| cathD | 14858.61 | -0.20384 | 0.433365 | -0.47036 | 0.638096 | 0.926691 |
| CG13739 | 3455.006 | 0.290047 | 0.616055 | 0.470813 | 0.637774 | 0.926691 |
| alpha-Catr | 634.1409 | 0.706149 | 1.497179 | 0.471653 | 0.637175 | 0.926691 |
| Stlk | 5299.436 | -0.25917 | 0.551468 | -0.46997 | 0.638379 | 0.926691 |
| Tango9 | 4912.437 | -0.30065 | 0.638726 | -0.4707 | 0.637855 | 0.926691 |
| SmD2 | 1417.152 | -0.45649 | 0.967445 | -0.47185 | 0.637035 | 0.926691 |
| CkIIalpha | 3305.731 | -0.34842 | 0.737166 | -0.47265 | 0.636465 | 0.926691 |
| CG45049 | 7782.183 | 0.297855 | 0.633582 | 0.470113 | 0.638274 | 0.926691 |
| CG1927 | 4823.392 | 0.2856 | 0.604355 | 0.47257 | 0.63652 | 0.926691 |
| CG1703 | 412.9353 | 0.702377 | 1.492974 | 0.470455 | 0.63803 | 0.926691 |
| CG14194 | 1099.118 | 0.523783 | 1.114925 | 0.469792 | 0.638504 | 0.926691 |
| CG17361 | 175.728 | -1.27504 | 2.697314 | -0.47271 | 0.636423 | 0.926691 |
| RapGAP1 | 10368.7 | 0.242379 | 0.516507 | 0.469266 | 0.63888 | 0.926907 |
| Trp1 | 5885.767 | -0.28717 | 0.613957 | -0.46774 | 0.639972 | 0.927872 |
| Paip2 | 4117.558 | 0.285635 | 0.610737 | 0.467689 | 0.640007 | 0.927872 |
| Axs | 80.83992 | 1.559901 | 3.332141 | 0.468138 | 0.639686 | 0.927872 |
| CG14515 | 271.0331 | 1.416367 | 3.026699 | 0.467957 | 0.639815 | 0.927872 |
| CG3078 | 3816.066 | 0.284551 | 0.608897 | 0.467322 | 0.64027 | 0.928085 |
| CG3071 | 686.037 | -0.73444 | 1.597081 | -0.45986 | 0.645614 | 0.928162 |
| Ide | 4320.899 | 0.278559 | 0.604645 | 0.460698 | 0.645015 | 0.928162 |
| Hrb27C | 22554.46 | 0.194423 | 0.41958 | 0.463376 | 0.643095 | 0.928162 |
| CG4168 | 681.6987 | 0.547945 | 1.173658 | 0.466869 | 0.640593 | 0.928162 |
| bur | 1001.101 | -0.50296 | 1.084537 | -0.46376 | 0.642822 | 0.928162 |
| Timp | 4031.538 | -0.32331 | 0.702956 | -0.45993 | 0.645566 | 0.928162 |
| Cyp12e1 | 3716.574 | -0.29997 | 0.65235 | -0.45982 | 0.645642 | 0.928162 |
| Hrb87F | 891.0674 | 0.533246 | 1.150348 | 0.463552 | 0.642969 | 0.928162 |
| tara | 1780.466 | 0.408579 | 0.88508 | 0.46163 | 0.644347 | 0.928162 |
| CG17931 | 261.7688 | -0.85355 | 1.85324 | -0.46057 | 0.645105 | 0.928162 |
| aralar1 | 3112.809 | 0.392207 | 0.851999 | 0.460338 | 0.645274 | 0.928162 |
| CG1636 | 1538.467 | 0.388261 | 0.83908 | 0.462722 | 0.643564 | 0.928162 |

| | | | | | | |
|---|---|---|---|---|---|---|
| tud | 5160.209 | -0.26558 | 0.576131 | -0.46098 | 0.644814 | 0.928162 |
| Gbp1 | 323.2196 | -1.11452 | 2.392396 | -0.46586 | 0.641317 | 0.928162 |
| Sec61beta | 3820.087 | -0.27766 | 0.597323 | -0.46483 | 0.642051 | 0.928162 |
| RpS23 | 8640.39 | -0.22799 | 0.489441 | -0.46582 | 0.641346 | 0.928162 |
| CG10375 | 172.295 | -0.96023 | 2.074262 | -0.46292 | 0.643418 | 0.928162 |
| Rbp9 | 13007.11 | 0.205389 | 0.443048 | 0.46358 | 0.642948 | 0.928162 |
| spi | 5226.833 | -0.29144 | 0.626165 | -0.46544 | 0.641616 | 0.928162 |
| AlaRS | 207.2952 | 1.061324 | 2.304292 | 0.460586 | 0.645096 | 0.928162 |
| CG34159 | 2460.09 | 0.377165 | 0.811213 | 0.46494 | 0.641974 | 0.928162 |
| Zdhhc8 | 2558.851 | -0.36974 | 0.793214 | -0.46612 | 0.641127 | 0.928162 |
| 5PtaseI | 3399.765 | -0.2874 | 0.624183 | -0.46045 | 0.645194 | 0.928162 |
| scra | 96.59797 | -1.71366 | 3.675469 | -0.46624 | 0.641042 | 0.928162 |
| nemy | 8503.29 | 0.292725 | 0.634223 | 0.461549 | 0.644405 | 0.928162 |
| YME1L | 908.8513 | -0.53539 | 1.147848 | -0.46643 | 0.640909 | 0.928162 |
| mRpL17 | 2600.612 | 0.367265 | 0.79449 | 0.462266 | 0.643891 | 0.928162 |
| CG43850 | 296.6668 | 0.9389 | 2.012926 | 0.466435 | 0.640904 | 0.928162 |
| Glut1 | 21177.14 | 0.220202 | 0.475244 | 0.463345 | 0.643117 | 0.928162 |
| Rgk1 | 1346.442 | -0.5443 | 1.173876 | -0.46368 | 0.642878 | 0.928162 |
| Spn42Da | 107.6656 | 1.132955 | 2.461792 | 0.460215 | 0.645362 | 0.928162 |
| lncRNA:CR | 2267.489 | -0.3859 | 0.837622 | -0.46071 | 0.645005 | 0.928162 |
| CG45263 | 13539.64 | 0.23391 | 0.505286 | 0.462926 | 0.643418 | 0.928162 |
| lncRNA:CR | 910.6268 | -0.46294 | 0.99846 | -0.46366 | 0.642892 | 0.928162 |
| CG12099 | 2122.497 | 0.383469 | 0.82471 | 0.464974 | 0.64195 | 0.928162 |
| Pgant6 | 494.4196 | 0.659359 | 1.429709 | 0.461184 | 0.644667 | 0.928162 |
| sbr | 4449.043 | -0.28217 | 0.606383 | -0.46534 | 0.64169 | 0.928162 |
| Gapvd1 | 498.5556 | 0.620925 | 1.342092 | 0.462655 | 0.643612 | 0.928162 |
| eag | 7880.206 | 0.249315 | 0.537515 | 0.463829 | 0.64277 | 0.928162 |
| CG6659 | 1443.909 | 0.438647 | 0.943705 | 0.464814 | 0.642065 | 0.928162 |
| Papss | 1058.243 | 0.69596 | 1.508697 | 0.461299 | 0.644584 | 0.928162 |
| rogdi | 4367.726 | 0.269077 | 0.584033 | 0.460723 | 0.644998 | 0.928162 |
| Ubc4 | 1315.133 | 0.405413 | 0.872576 | 0.464616 | 0.642206 | 0.928162 |
| Gug | 6486.696 | -0.24021 | 0.516071 | -0.46547 | 0.641597 | 0.928162 |
| ple | 426.0157 | 1.019522 | 2.21276 | 0.460747 | 0.64498 | 0.928162 |
| CG10031 | 114.2697 | -1.58831 | 3.442413 | -0.46139 | 0.644517 | 0.928162 |
| CG34117 | 1556.783 | -0.37356 | 0.812721 | -0.45964 | 0.645775 | 0.928186 |
| CG14646 | 521.7197 | 0.854468 | 1.861535 | 0.459013 | 0.646225 | 0.9285 |
| CG30428 | 3610.189 | -0.27887 | 0.607403 | -0.45912 | 0.646145 | 0.9285 |
| GstE12 | 11109.89 | 0.230857 | 0.503273 | 0.458711 | 0.646442 | 0.928645 |
| CG6656 | 318.4971 | 0.94641 | 2.065196 | 0.458267 | 0.646761 | 0.928882 |
| gammaCO | 1924.383 | 0.454145 | 0.991239 | 0.45816 | 0.646838 | 0.928882 |
| CG9630 | 325.6449 | 1.110136 | 2.423953 | 0.457986 | 0.646963 | 0.928895 |
| Trx-2 | 5201.594 | -0.27629 | 0.603974 | -0.45746 | 0.64734 | 0.929105 |
| Grd | 122.1479 | 1.754264 | 3.833806 | 0.457578 | 0.647256 | 0.929105 |
| Atf3 | 1758.495 | -0.39079 | 0.878647 | -0.44477 | 0.656488 | 0.929164 |
| Ocrl | 1579.531 | 0.442109 | 0.998121 | 0.442941 | 0.657808 | 0.929164 |
| CG14818 | 2074.595 | -0.32926 | 0.725976 | -0.45355 | 0.650156 | 0.929164 |
| dnc | 15997.57 | -0.18386 | 0.424575 | -0.43305 | 0.664978 | 0.929164 |
| CG15765 | 12486.3 | 0.228454 | 0.511499 | 0.446635 | 0.655139 | 0.929164 |
| drongo | 6342.233 | -0.25165 | 0.575108 | -0.43757 | 0.661696 | 0.929164 |
| CG3625 | 6445.753 | 0.247283 | 0.558996 | 0.44237 | 0.658222 | 0.929164 |

| | | | | | | |
|---|---|---|---|---|---|---|
| kin17 | 907.5863 | 0.713475 | 1.643593 | 0.434095 | 0.66422 | 0.929164 |
| mRpL15 | 962.3753 | -0.51215 | 1.184659 | -0.43232 | 0.66551 | 0.929164 |
| CG33285 | 141.9085 | -1.70748 | 3.951177 | -0.43214 | 0.665637 | 0.929164 |
| CG33288 | 65.34555 | -1.70647 | 3.951495 | -0.43186 | 0.665846 | 0.929164 |
| Wnk | 4966.39 | -0.26098 | 0.580961 | -0.44922 | 0.653271 | 0.929164 |
| Pc | 1189.626 | -0.50978 | 1.148516 | -0.44386 | 0.657147 | 0.929164 |
| CG7458 | 127.947 | -1.70729 | 3.896013 | -0.43821 | 0.661231 | 0.929164 |
| Vha14-2 | 85.43514 | 1.752295 | 3.951516 | 0.443449 | 0.657441 | 0.929164 |
| jagn | 3088.761 | -0.32937 | 0.724975 | -0.45432 | 0.649597 | 0.929164 |
| CG14020 | 102.9148 | -1.7071 | 3.95128 | -0.43204 | 0.665715 | 0.929164 |
| CG11034 | 62.99801 | 1.71957 | 3.951712 | 0.435145 | 0.663457 | 0.929164 |
| CG13784 | 3023.407 | 0.301048 | 0.672185 | 0.447865 | 0.65425 | 0.929164 |
| jp | 353.427 | 0.648236 | 1.493921 | 0.433916 | 0.664349 | 0.929164 |
| Sur | 168.0823 | -1.70761 | 3.951135 | -0.43218 | 0.665609 | 0.929164 |
| CG6729 | 112.7038 | 1.307996 | 2.97228 | 0.440065 | 0.65989 | 0.929164 |
| CG5781 | 116.2053 | 1.643603 | 3.691421 | 0.445249 | 0.656139 | 0.929164 |
| Tap42 | 262.4048 | 1.136417 | 2.556116 | 0.444587 | 0.656618 | 0.929164 |
| Ski6 | 83.4307 | 1.725329 | 3.834725 | 0.449922 | 0.652766 | 0.929164 |
| CG7099 | 425.0317 | -0.71933 | 1.575966 | -0.45644 | 0.648076 | 0.929164 |
| CG16863 | 463.7308 | -0.88455 | 2.028283 | -0.43611 | 0.662757 | 0.929164 |
| Adat1 | 138.4789 | -1.13956 | 2.496948 | -0.45638 | 0.648115 | 0.929164 |
| CG15278 | 66.41387 | -1.7065 | 3.951485 | -0.43186 | 0.66584 | 0.929164 |
| kek3 | 2642.616 | 0.397824 | 0.874704 | 0.454809 | 0.649246 | 0.929164 |
| CG4455 | 104.2811 | 1.752743 | 3.951406 | 0.443575 | 0.65735 | 0.929164 |
| crp | 3543.866 | -0.28134 | 0.632683 | -0.44468 | 0.656551 | 0.929164 |
| yellow-b | 79.57195 | 1.752116 | 3.951561 | 0.443398 | 0.657478 | 0.929164 |
| Rsph9 | 83.44584 | 1.752239 | 3.951531 | 0.443433 | 0.657453 | 0.929164 |
| Acp36DE | 112.0709 | 1.717092 | 3.951359 | 0.434557 | 0.663884 | 0.929164 |
| MESR3 | 3068.762 | 0.301352 | 0.6688 | 0.450586 | 0.652288 | 0.929164 |
| Grip71 | 102.0824 | 1.752699 | 3.951417 | 0.443562 | 0.657359 | 0.929164 |
| CG33120 | 83.68504 | -1.70688 | 3.951366 | -0.43197 | 0.665762 | 0.929164 |
| CG10495 | 992.2899 | 0.633552 | 1.457382 | 0.434719 | 0.663766 | 0.929164 |
| l(2)37Cc | 1645.427 | -0.4019 | 0.901704 | -0.44572 | 0.655802 | 0.929164 |
| pigeon | 1392.257 | -0.44104 | 1.011772 | -0.43591 | 0.662902 | 0.929164 |
| tj | 95.76322 | -1.44857 | 3.273871 | -0.44246 | 0.658154 | 0.929164 |
| COX4 | 37582.9 | 0.182029 | 0.412038 | 0.441777 | 0.658651 | 0.929164 |
| EMC1 | 2267.511 | 0.333108 | 0.732244 | 0.454914 | 0.649171 | 0.929164 |
| CG8369 | 39486.29 | -0.18356 | 0.421912 | -0.43508 | 0.663505 | 0.929164 |
| CG14687 | 785.4553 | 0.771637 | 1.778601 | 0.433845 | 0.664401 | 0.929164 |
| TfIIFbeta | 196.6896 | 1.001615 | 2.225867 | 0.449989 | 0.652718 | 0.929164 |
| CG7488 | 468.087 | 0.831451 | 1.829183 | 0.454548 | 0.649435 | 0.929164 |
| eff | 9272.916 | -0.24186 | 0.560528 | -0.43148 | 0.666121 | 0.929164 |
| SerRS-m | 609.4298 | 0.83673 | 1.853919 | 0.451331 | 0.651751 | 0.929164 |
| CG4546 | 93.39234 | 1.752462 | 3.951464 | 0.443497 | 0.657406 | 0.929164 |
| fru | 13339.04 | -0.19311 | 0.43929 | -0.43961 | 0.660222 | 0.929164 |
| ninaE | 169.5248 | 1.628943 | 3.77684 | 0.431298 | 0.666252 | 0.929164 |
| Trm7-34 | 997.1145 | -0.51113 | 1.168992 | -0.43724 | 0.661938 | 0.929164 |
| CASK | 12314.89 | -0.19276 | 0.438991 | -0.43911 | 0.660584 | 0.929164 |
| CG13850 | 1233.18 | -0.42769 | 0.954465 | -0.44809 | 0.654086 | 0.929164 |
| Ndc1 | 112.3117 | 1.734817 | 3.951364 | 0.439043 | 0.660631 | 0.929164 |

| | | | | | | |
|---|---|---|---|---|---|---|
| Slimp | 476.5417 | 0.987977 | 2.171277 | 0.455021 | 0.649094 | 0.929164 |
| CG10560 | 153.0913 | 1.584577 | 3.654554 | 0.43359 | 0.664586 | 0.929164 |
| wdn | 409.1124 | 0.820948 | 1.800716 | 0.455901 | 0.648461 | 0.929164 |
| eEF1gamm | 2392.952 | -0.35194 | 0.79975 | -0.44006 | 0.659893 | 0.929164 |
| Cnx99A | 3159.406 | -0.29795 | 0.683929 | -0.43564 | 0.663097 | 0.929164 |
| qless | 883.0963 | 0.546442 | 1.243614 | 0.439398 | 0.660373 | 0.929164 |
| Cul2 | 2788.827 | 0.346875 | 0.785834 | 0.44141 | 0.658916 | 0.929164 |
| qsm | 64.91396 | 1.751512 | 3.951709 | 0.443229 | 0.6576 | 0.929164 |
| Hsl | 1690.184 | -0.39199 | 0.873978 | -0.44851 | 0.653784 | 0.929164 |
| CG11041 | 78.52495 | 1.752079 | 3.95157 | 0.443388 | 0.657485 | 0.929164 |
| hrg | 3257.691 | 0.279798 | 0.632297 | 0.442511 | 0.65812 | 0.929164 |
| stau | 6759.425 | 0.241848 | 0.546924 | 0.442196 | 0.658347 | 0.929164 |
| Trf4-1 | 869.3449 | -0.45624 | 1.032224 | -0.442 | 0.658491 | 0.929164 |
| fend | 4591.009 | -0.30049 | 0.681515 | -0.44091 | 0.659277 | 0.929164 |
| CG32683 | 2319.75 | -0.38671 | 0.868448 | -0.44529 | 0.656109 | 0.929164 |
| BTBD9 | 2029.342 | -0.55619 | 1.252702 | -0.44399 | 0.657049 | 0.929164 |
| stx | 2091.233 | -0.34824 | 0.770461 | -0.45199 | 0.651279 | 0.929164 |
| CG10433 | 30636.99 | -0.24077 | 0.547782 | -0.43954 | 0.660273 | 0.929164 |
| Oatp58Dc | 5936.459 | -0.31722 | 0.694196 | -0.45697 | 0.647696 | 0.929164 |
| CG12964 | 70.14895 | 1.751719 | 3.951649 | 0.443288 | 0.657557 | 0.929164 |
| unc-5 | 184.199 | -0.97684 | 2.185317 | -0.447 | 0.654873 | 0.929164 |
| hui | 135.6911 | 1.753227 | 3.951291 | 0.44371 | 0.657252 | 0.929164 |
| drk | 2193.403 | 0.353549 | 0.812447 | 0.435166 | 0.663442 | 0.929164 |
| CG7637 | 140.2225 | -0.97494 | 2.141626 | -0.45523 | 0.648943 | 0.929164 |
| CG30345 | 74.515 | 1.724863 | 3.951591 | 0.436498 | 0.662475 | 0.929164 |
| Shaw | 4007.552 | -0.29597 | 0.686214 | -0.43131 | 0.666243 | 0.929164 |
| ptc | 201.2041 | 1.02995 | 2.374363 | 0.433779 | 0.664449 | 0.929164 |
| CG5033 | 205.1171 | 1.023296 | 2.366028 | 0.432495 | 0.665382 | 0.929164 |
| CR18217 | 223.778 | 1.275031 | 2.893856 | 0.440599 | 0.659503 | 0.929164 |
| REPTOR-BI | 4159.217 | -0.26364 | 0.607161 | -0.43423 | 0.664125 | 0.929164 |
| CG31437 | 65.22805 | 1.751528 | 3.951705 | 0.443233 | 0.657597 | 0.929164 |
| mav | 5907.484 | -0.27981 | 0.628641 | -0.4451 | 0.656249 | 0.929164 |
| IA-2 | 118769.4 | -0.16898 | 0.380801 | -0.44376 | 0.657219 | 0.929164 |
| tral | 4098.652 | 0.271317 | 0.59735 | 0.454201 | 0.649684 | 0.929164 |
| mxt | 2542.345 | -0.32384 | 0.744944 | -0.43472 | 0.663763 | 0.929164 |
| snoRNA:Ps | 555.6862 | -0.55043 | 1.226378 | -0.44883 | 0.653557 | 0.929164 |
| CG34200 | 1669.834 | 0.355978 | 0.815244 | 0.436652 | 0.662364 | 0.929164 |
| CG34288 | 77.80928 | -1.70677 | 3.951401 | -0.43194 | 0.665785 | 0.929164 |
| dpr12 | 6995.61 | -0.24078 | 0.531253 | -0.45323 | 0.650384 | 0.929164 |
| CG40472 | 1753.152 | 0.361209 | 0.824817 | 0.437926 | 0.66144 | 0.929164 |
| stv | 574.1865 | 0.579536 | 1.275557 | 0.45434 | 0.649584 | 0.929164 |
| CG14644 | 153.719 | 1.585716 | 3.64935 | 0.43452 | 0.663911 | 0.929164 |
| Octbeta3R | 2249.158 | 0.339128 | 0.784741 | 0.432153 | 0.66563 | 0.929164 |
| CG42391 | 88.31443 | -1.70694 | 3.951342 | -0.43199 | 0.665748 | 0.929164 |
| Sep-04 | 5395.002 | 0.251766 | 0.578725 | 0.435036 | 0.663537 | 0.929164 |
| Thor | 1289.546 | -0.44109 | 1.0066 | -0.4382 | 0.661242 | 0.929164 |
| LanB1 | 134.2066 | 1.393224 | 3.147052 | 0.442708 | 0.657977 | 0.929164 |
| 5-HT2B | 2036.257 | -0.38627 | 0.890366 | -0.43384 | 0.664407 | 0.929164 |
| Acsl | 11364.53 | 0.21676 | 0.48942 | 0.442892 | 0.657844 | 0.929164 |
| mRpS29 | 1063.834 | 0.490017 | 1.099551 | 0.445652 | 0.655849 | 0.929164 |

| | | | | | | |
|---|---|---|---|---|---|---|
| Liprin-gam | 13551.06 | -0.2124 | 0.482699 | -0.44004 | 0.659911 | 0.929164 |
| CG2765 | 3031.908 | -0.37146 | 0.855404 | -0.43425 | 0.66411 | 0.929164 |
| ttm2 | 93.67721 | -1.70503 | 3.951264 | -0.43151 | 0.666095 | 0.929164 |
| asRNA:CR4 | 85.64363 | -1.70691 | 3.951355 | -0.43198 | 0.665755 | 0.929164 |
| asRNA:CR4 | 772.1605 | -0.90433 | 2.061142 | -0.43875 | 0.66084 | 0.929164 |
| Piezo | 3046.521 | 0.303208 | 0.675489 | 0.448873 | 0.653524 | 0.929164 |
| lncRNA:CR | 92.5877 | -1.70698 | 3.951322 | -0.432 | 0.66574 | 0.929164 |
| CG44325 | 5324.313 | 0.256162 | 0.585062 | 0.437838 | 0.661504 | 0.929164 |
| asRNA:CR4 | 82.04823 | 1.168564 | 2.678744 | 0.436236 | 0.662666 | 0.929164 |
| lncRNA:CR | 689.487 | 0.534473 | 1.209994 | 0.441715 | 0.658695 | 0.929164 |
| Exo84 | 560.0825 | 0.608663 | 1.362983 | 0.446567 | 0.655188 | 0.929164 |
| Ntl | 95.39294 | -1.13323 | 2.56606 | -0.44162 | 0.658762 | 0.929164 |
| lncRNA:CR | 348310.5 | -0.2176 | 0.489562 | -0.44447 | 0.656703 | 0.929164 |
| Der-1 | 3035.608 | 0.348368 | 0.786095 | 0.443163 | 0.657648 | 0.929164 |
| asRNA:CR4 | 125.7056 | -1.70736 | 3.951212 | -0.43211 | 0.66566 | 0.929164 |
| hpRNA:CR4 | 99.36275 | -1.14017 | 2.59304 | -0.4397 | 0.660153 | 0.929164 |
| ND-ACP | 11848.55 | -0.19304 | 0.442854 | -0.43591 | 0.662902 | 0.929164 |
| Pex10 | 501.1399 | 1.07164 | 2.374176 | 0.451373 | 0.651721 | 0.929164 |
| CG12104 | 761.2875 | -0.56992 | 1.27153 | -0.44821 | 0.654 | 0.929164 |
| CG5707 | 440.6934 | 1.074656 | 2.463004 | 0.436319 | 0.662605 | 0.929164 |
| Larp4B | 8275.751 | 0.219924 | 0.5006 | 0.439321 | 0.660429 | 0.929164 |
| promL | 5319.566 | -0.24919 | 0.547371 | -0.45525 | 0.648928 | 0.929164 |
| Ack | 535.2185 | -0.69021 | 1.59464 | -0.43283 | 0.665136 | 0.929164 |
| CG15209 | 3217.599 | -0.29457 | 0.655736 | -0.44921 | 0.653277 | 0.929164 |
| Lint-1 | 565.899 | -0.66834 | 1.508492 | -0.44305 | 0.657728 | 0.929164 |
| Cpr11B | 183.6676 | -1.11242 | 2.503099 | -0.44442 | 0.656742 | 0.929164 |
| Tim9a | 630.5754 | 0.524445 | 1.188195 | 0.441379 | 0.658938 | 0.929164 |
| Rbp1-like | 8190.782 | 0.218441 | 0.487064 | 0.448485 | 0.653803 | 0.929164 |
| Rab3-GEF | 3364.079 | -0.29773 | 0.673152 | -0.4423 | 0.658274 | 0.929164 |
| CG8509 | 90.77484 | 1.75243 | 3.95148 | 0.443487 | 0.657414 | 0.929164 |
| scu | 3088.116 | -0.358 | 0.815701 | -0.43888 | 0.660746 | 0.929164 |
| HisRS | 1052.047 | -0.51019 | 1.124325 | -0.45377 | 0.649993 | 0.929164 |
| CG7058 | 2215.937 | -0.60729 | 1.394426 | -0.43552 | 0.663188 | 0.929164 |
| Sec61gamı | 3987.76 | -0.25484 | 0.589243 | -0.43249 | 0.665384 | 0.929164 |
| CG6951 | 109.0973 | 1.752839 | 3.951384 | 0.443601 | 0.657331 | 0.929164 |
| trc | 1517.876 | -0.43478 | 0.964954 | -0.45057 | 0.652302 | 0.929164 |
| fal | 380.5835 | -0.90911 | 2.078162 | -0.43746 | 0.661779 | 0.929164 |
| CG7484 | 1914.386 | -0.3348 | 0.757314 | -0.44209 | 0.658423 | 0.929164 |
| Pop5 | 385.0351 | 1.008271 | 2.225591 | 0.453035 | 0.650523 | 0.929164 |
| CG32163 | 89.70932 | 1.117758 | 2.556334 | 0.43725 | 0.66193 | 0.929164 |
| Pdh | 84.56619 | 1.728334 | 3.951511 | 0.437386 | 0.661832 | 0.929164 |
| CG7768 | 230.724 | 1.367271 | 3.068457 | 0.445589 | 0.655894 | 0.929164 |
| FoxK | 1698.402 | 0.366539 | 0.84507 | 0.433738 | 0.664479 | 0.929164 |
| Irbp18 | 577.9174 | 0.661769 | 1.460162 | 0.453216 | 0.650393 | 0.929164 |
| CG14174 | 100.244 | -1.7071 | 3.95129 | -0.43204 | 0.665715 | 0.929164 |
| CG3434 | 627.2927 | 0.653929 | 1.434235 | 0.455943 | 0.648431 | 0.929164 |
| foi | 1321.339 | 0.541621 | 1.21585 | 0.445467 | 0.655982 | 0.929164 |
| qm | 1174.538 | -0.41825 | 0.92859 | -0.45042 | 0.65241 | 0.929164 |
| tow | 2835.999 | -0.33088 | 0.746208 | -0.44342 | 0.657464 | 0.929164 |
| mst | 119.0438 | 1.753005 | 3.951344 | 0.443648 | 0.657297 | 0.929164 |

| | | | | | | |
|---|---|---|---|---|---|---|
| Obp19d | 154.7465 | 1.753424 | 3.951243 | 0.443765 | 0.657212 | 0.929164 |
| Ugt36A1 | 128.1984 | -1.70738 | 3.951206 | -0.43212 | 0.665656 | 0.929164 |
| Sbat | 85.80567 | 1.606748 | 3.521026 | 0.456329 | 0.648153 | 0.929164 |
| CG3104 | 1691.831 | 0.453282 | 1.017457 | 0.445504 | 0.655955 | 0.929164 |
| Taf10b | 4420.168 | 0.257255 | 0.562864 | 0.457047 | 0.647638 | 0.929164 |
| CG4238 | 3292.569 | 0.290244 | 0.670734 | 0.432726 | 0.665214 | 0.929164 |
| CkIIalpha-i | 239.3288 | -0.90771 | 2.105781 | -0.43106 | 0.666427 | 0.929246 |
| CG2991 | 5092.015 | 0.280496 | 0.651501 | 0.430538 | 0.666804 | 0.929288 |
| CG32549 | 6834.228 | 0.221996 | 0.515617 | 0.430544 | 0.6668 | 0.929288 |
| Arr2 | 509.3729 | 1.139934 | 2.646463 | 0.430739 | 0.666658 | 0.929288 |
| amon | 7859.766 | 0.220679 | 0.513132 | 0.430063 | 0.66715 | 0.929447 |
| rols | 3380.868 | -0.35697 | 0.829841 | -0.43017 | 0.667072 | 0.929447 |
| Sos | 1187.492 | 0.603275 | 1.403414 | 0.429863 | 0.667296 | 0.929489 |
| Coq7 | 3271.934 | -0.26884 | 0.625669 | -0.42968 | 0.667431 | 0.929516 |
| otu | 92.3363 | -1.6945 | 3.95132 | -0.42884 | 0.668037 | 0.930037 |
| CG15043 | 233.0389 | 1.360157 | 3.171177 | 0.428912 | 0.667987 | 0.930037 |
| Hacd2 | 2140.744 | -0.43171 | 1.008455 | -0.42809 | 0.668583 | 0.930153 |
| CG6908 | 1264.486 | 0.475434 | 1.109423 | 0.428542 | 0.668257 | 0.930153 |
| Dbp80 | 8647.395 | 0.212614 | 0.496483 | 0.42824 | 0.668476 | 0.930153 |
| CG32219 | 129.9427 | -1.34201 | 3.134608 | -0.42813 | 0.668558 | 0.930153 |
| CG13339 | 330.7861 | 0.68255 | 1.59616 | 0.42762 | 0.668928 | 0.930309 |
| Ptip | 940.4413 | -0.70342 | 1.644929 | -0.42763 | 0.668919 | 0.930309 |
| CG10337 | 111.8488 | -1.6865 | 3.951246 | -0.42683 | 0.669504 | 0.930929 |
| Uxs | 379.5396 | -0.94654 | 2.218347 | -0.42669 | 0.669606 | 0.930929 |
| rush | 1053.481 | 0.485343 | 1.138036 | 0.426474 | 0.669762 | 0.930986 |
| hyd | 6850.462 | 0.219603 | 0.51542 | 0.426066 | 0.670059 | 0.931238 |
| Bdp1 | 186.0027 | 0.9676 | 2.276806 | 0.424981 | 0.67085 | 0.93165 |
| p38a | 351.4893 | 0.987893 | 2.325203 | 0.424863 | 0.670937 | 0.93165 |
| mRpL54 | 1562.996 | 0.372042 | 0.875206 | 0.42509 | 0.670771 | 0.93165 |
| CG32055 | 303.4651 | -1.12974 | 2.656758 | -0.42523 | 0.670668 | 0.93165 |
| cert | 1016.453 | 0.423375 | 0.996139 | 0.425016 | 0.670825 | 0.93165 |
| Pli | 6350.466 | 0.229483 | 0.540789 | 0.424349 | 0.671311 | 0.932009 |
| Rif1 | 73.56735 | -1.67547 | 3.951419 | -0.42402 | 0.671553 | 0.932067 |
| dati | 3114.412 | 0.290967 | 0.686286 | 0.423973 | 0.671585 | 0.932067 |
| CG13029 | 71.0746 | -1.67435 | 3.951436 | -0.42373 | 0.67176 | 0.932149 |
| CG11456 | 854.7099 | -0.78379 | 1.851445 | -0.42334 | 0.672048 | 0.932387 |
| dap | 331.7935 | 0.682165 | 1.613439 | 0.422802 | 0.67244 | 0.932551 |
| CG11752 | 5486.186 | 0.245447 | 0.580471 | 0.422842 | 0.672411 | 0.932551 |
| Frq2 | 1846.886 | 0.354549 | 0.838774 | 0.422699 | 0.672515 | 0.932551 |
| Fis1 | 9893.7 | 0.198683 | 0.470507 | 0.422274 | 0.672825 | 0.932659 |
| Zip99C | 399.5928 | 0.642548 | 1.521237 | 0.422385 | 0.672744 | 0.932659 |
| lncRNA:CR | 122.5177 | 1.399076 | 3.316505 | 0.421853 | 0.673133 | 0.932716 |
| cue | 473.8812 | 0.648678 | 1.537064 | 0.422024 | 0.673008 | 0.932716 |
| CG10809 | 1127.801 | 0.395066 | 0.936754 | 0.42174 | 0.673215 | 0.932716 |
| CG13364 | 2122.65 | -0.30306 | 0.719557 | -0.42117 | 0.67363 | 0.932807 |
| CG10341 | 620.9189 | 0.858323 | 2.037761 | 0.421209 | 0.673603 | 0.932807 |
| CG12818 | 281.0869 | -0.96889 | 2.299814 | -0.42129 | 0.673543 | 0.932807 |
| CG6481 | 168.7823 | 1.169811 | 2.779504 | 0.42087 | 0.67385 | 0.932951 |
| CG34353 | 6175.759 | -0.24327 | 0.578732 | -0.42034 | 0.674236 | 0.933268 |
| CG6282 | 7191.796 | -0.21898 | 0.521083 | -0.42024 | 0.674311 | 0.933268 |

| | | | | | | |
|---|---|---|---|---|---|---|
| RpL14 | 31210.57 | 0.16821 | 0.400536 | 0.419961 | 0.674514 | 0.933387 |
| Rpt4 | 1334.861 | -0.40007 | 0.953116 | -0.41975 | 0.674667 | 0.933439 |
| Grip75 | 526.4989 | -0.74889 | 1.786649 | -0.41916 | 0.675099 | 0.933693 |
| CG31183 | 887.3026 | -0.49694 | 1.185365 | -0.41923 | 0.675048 | 0.933693 |
| CG13014 | 504.5531 | 0.757152 | 1.80695 | 0.419022 | 0.6752 | 0.933693 |
| CG1648 | 4350.14 | 0.250655 | 0.598767 | 0.418618 | 0.675495 | 0.933941 |
| ubl | 369.0138 | -0.62859 | 1.503463 | -0.41809 | 0.675879 | 0.934311 |
| CG4042 | 1676.905 | 0.415519 | 0.995295 | 0.417483 | 0.676325 | 0.934449 |
| COX7A | 26580.9 | 0.184846 | 0.442769 | 0.417478 | 0.676329 | 0.934449 |
| S6KL | 1147.359 | 0.410245 | 0.982571 | 0.417522 | 0.676296 | 0.934449 |
| Rox8 | 5160.209 | 0.301491 | 0.723036 | 0.416979 | 0.676694 | 0.934471 |
| CG31038 | 1868.781 | -0.59094 | 1.416733 | -0.41712 | 0.676593 | 0.934471 |
| eEF1alpha | 32671.46 | 0.19597 | 0.469897 | 0.417048 | 0.676643 | 0.934471 |
| Ccdc58 | 351.5019 | 0.866231 | 2.080751 | 0.416307 | 0.677185 | 0.934904 |
| CG7564 | 2752.449 | -0.34183 | 0.821241 | -0.41623 | 0.67724 | 0.934904 |
| Vsp37A | 2568.846 | 0.280928 | 0.675729 | 0.41574 | 0.6776 | 0.93524 |
| CG3987 | 584.8711 | -0.59423 | 1.431012 | -0.41525 | 0.677959 | 0.935554 |
| Cyt-c-p | 45577.01 | 0.16302 | 0.392864 | 0.414952 | 0.678177 | 0.935554 |
| Toll-6 | 2167.774 | -0.38705 | 0.932525 | -0.41506 | 0.678101 | 0.935554 |
| RhoGAP10 | 13026.6 | 0.180396 | 0.435018 | 0.414685 | 0.678372 | 0.935662 |
| CG6293 | 732.6815 | 0.916233 | 2.218637 | 0.412971 | 0.679628 | 0.935913 |
| SdhD | 12263.39 | 0.183557 | 0.443938 | 0.413475 | 0.679259 | 0.935913 |
| vap | 2313.035 | 0.30698 | 0.742037 | 0.413699 | 0.679095 | 0.935913 |
| amn | 398.6986 | -0.5972 | 1.445406 | -0.41317 | 0.679479 | 0.935913 |
| lncRNA:CR | 731.6745 | 0.663891 | 1.606727 | 0.413195 | 0.679464 | 0.935913 |
| lncRNA:CR | 78.36821 | 1.633373 | 3.951512 | 0.413354 | 0.679347 | 0.935913 |
| InR | 5169.912 | -0.26009 | 0.628872 | -0.41358 | 0.679181 | 0.935913 |
| CG18171 | 261.3338 | -0.70998 | 1.719721 | -0.41284 | 0.679721 | 0.935913 |
| IntS10 | 277.6125 | 0.695659 | 1.682789 | 0.413397 | 0.679316 | 0.935913 |
| CG4880 | 193.2392 | 1.566277 | 3.793529 | 0.412881 | 0.679694 | 0.935913 |
| Or45a | 1658.784 | 0.499648 | 1.210798 | 0.41266 | 0.679855 | 0.935938 |
| nAChRalph | 4713.927 | -0.24058 | 0.583718 | -0.41214 | 0.680233 | 0.936298 |
| Alp12 | 760.4206 | -0.45232 | 1.098981 | -0.41158 | 0.680649 | 0.936398 |
| TfIIA-S | 4692.273 | 0.232991 | 0.566107 | 0.411568 | 0.680656 | 0.936398 |
| CG42813 | 4026.488 | -0.24199 | 0.587695 | -0.41176 | 0.680512 | 0.936398 |
| haf | 690.7306 | 0.563781 | 1.370774 | 0.411287 | 0.680862 | 0.936521 |
| Rpn3 | 2312.419 | 0.351826 | 0.857879 | 0.410111 | 0.681725 | 0.937386 |
| Abl | 9586.177 | -0.19505 | 0.475524 | -0.41019 | 0.681669 | 0.937386 |
| UbcE2H | 1581.798 | 0.375596 | 0.917567 | 0.409339 | 0.682291 | 0.938004 |
| ksr | 690.466 | -0.48953 | 1.198393 | -0.40849 | 0.682913 | 0.938538 |
| Drep1 | 398.3133 | 0.758452 | 1.856564 | 0.408525 | 0.682888 | 0.938538 |
| Coprox | 166.8609 | 1.191808 | 2.920397 | 0.408098 | 0.683202 | 0.93865 |
| stumps | 6314.443 | -0.24802 | 0.607809 | -0.40806 | 0.683229 | 0.93865 |
| Charon | 708.3948 | 0.781091 | 1.915448 | 0.407785 | 0.683432 | 0.938768 |
| Cda5 | 1822.57 | 0.469352 | 1.151493 | 0.407603 | 0.683565 | 0.938791 |
| Plc21C | 7707.777 | 0.260326 | 0.640691 | 0.406321 | 0.684507 | 0.938981 |
| Sdhaf3 | 806.3719 | -0.43225 | 1.064249 | -0.40616 | 0.684629 | 0.938981 |
| tou | 4694.162 | 0.251874 | 0.619645 | 0.406482 | 0.684389 | 0.938981 |
| Tpst | 1236.559 | -0.41789 | 1.028414 | -0.40634 | 0.684489 | 0.938981 |
| CG42541 | 16922.28 | 0.189013 | 0.465191 | 0.406313 | 0.684512 | 0.938981 |

| | | | | | | |
|---|---|---|---|---|---|---|
| stai | 7936.671 | 0.195355 | 0.481004 | 0.406141 | 0.684639 | 0.938981 |
| eIF2beta | 2379.383 | -0.31284 | 0.769092 | -0.40677 | 0.684176 | 0.938981 |
| lin-28 | 91.97369 | 1.219555 | 2.996874 | 0.406942 | 0.68405 | 0.938981 |
| Tsp42Ee | 17300.75 | -0.23356 | 0.57559 | -0.40578 | 0.684903 | 0.939182 |
| CNBP | 5850.435 | 0.243832 | 0.601198 | 0.405576 | 0.685054 | 0.939229 |
| Rtf1 | 721.4631 | 0.504843 | 1.245961 | 0.405184 | 0.685343 | 0.939464 |
| Hacd1 | 2276.792 | 0.34047 | 0.841385 | 0.404655 | 0.685731 | 0.939676 |
| CG4970 | 130.2469 | -1.29501 | 3.199582 | -0.40474 | 0.685666 | 0.939676 |
| CG10189 | 270.3544 | 1.222276 | 3.03032 | 0.403349 | 0.686692 | 0.940349 |
| CG7741 | 1021.863 | 0.461053 | 1.141875 | 0.403769 | 0.686383 | 0.940349 |
| jvl | 6972.355 | -0.29665 | 0.735203 | -0.4035 | 0.686583 | 0.940349 |
| CG15203 | 970.736 | -0.43415 | 1.076335 | -0.40335 | 0.686687 | 0.940349 |
| Pex12 | 364.3707 | -0.88432 | 2.194752 | -0.40292 | 0.687005 | 0.940617 |
| SNRPG | 1890.517 | -0.30294 | 0.752955 | -0.40234 | 0.687436 | 0.941047 |
| Obp99c | 89.0742 | -1.54042 | 3.835564 | -0.40161 | 0.687968 | 0.941614 |
| Gld2 | 75.48303 | 1.218434 | 3.035159 | 0.40144 | 0.688096 | 0.94163 |
| AspRS-m | 633.3553 | 0.748173 | 1.865617 | 0.401033 | 0.688396 | 0.94188 |
| CG12730 | 7557.102 | -0.2162 | 0.542043 | -0.39887 | 0.689991 | 0.941908 |
| PLCXD | 5366.895 | -0.34161 | 0.857655 | -0.3983 | 0.690406 | 0.941908 |
| PICK1 | 1645.176 | -0.53789 | 1.349644 | -0.39854 | 0.690232 | 0.941908 |
| CG31808 | 12878.2 | 0.184258 | 0.461665 | 0.399115 | 0.689808 | 0.941908 |
| GCC185 | 486.2588 | -0.71869 | 1.803357 | -0.39853 | 0.69024 | 0.941908 |
| ninaB | 12003.26 | -0.25821 | 0.645908 | -0.39977 | 0.689327 | 0.941908 |
| Rad60 | 569.5295 | -0.73248 | 1.828507 | -0.40059 | 0.688723 | 0.941908 |
| Rab4 | 5249.76 | -0.23382 | 0.585367 | -0.39944 | 0.689572 | 0.941908 |
| Setd3 | 413.0298 | 0.807179 | 2.029011 | 0.397819 | 0.690764 | 0.941908 |
| Echs1 | 1309.645 | -0.48955 | 1.223472 | -0.40014 | 0.689057 | 0.941908 |
| CG32206 | 6769.305 | 0.200089 | 0.502401 | 0.398266 | 0.690434 | 0.941908 |
| Chd1 | 853.6195 | 0.449314 | 1.12577 | 0.399117 | 0.689807 | 0.941908 |
| CG43295 | 72.21207 | 1.557136 | 3.913623 | 0.397876 | 0.690722 | 0.941908 |
| nahoda | 253.7888 | -0.68712 | 1.723308 | -0.39872 | 0.690099 | 0.941908 |
| CG46385 | 21152.99 | -0.17729 | 0.445205 | -0.39823 | 0.690461 | 0.941908 |
| Pcyt2 | 7345.781 | -0.20639 | 0.518479 | -0.39807 | 0.69058 | 0.941908 |
| Tomosyn | 5123.572 | 0.247062 | 0.619952 | 0.398517 | 0.690249 | 0.941908 |
| Grasp65 | 1737.689 | 0.389133 | 0.973455 | 0.399745 | 0.689345 | 0.941908 |
| CG6683 | 328.5606 | 0.624764 | 1.561934 | 0.399994 | 0.689161 | 0.941908 |
| CG4669 | 205.767 | 1.317115 | 3.300194 | 0.399102 | 0.689818 | 0.941908 |
| Oatp33Ea | 1567.141 | 0.393497 | 0.990195 | 0.397394 | 0.691077 | 0.941917 |
| CG8778 | 1418.412 | 0.49044 | 1.234368 | 0.397321 | 0.691131 | 0.941917 |
| GlcAT-P | 20431.41 | -0.18112 | 0.455663 | -0.39749 | 0.691003 | 0.941917 |
| Lrp4 | 7751.45 | -0.20327 | 0.511786 | -0.39717 | 0.69124 | 0.941917 |
| HSPBAP1 | 594.0103 | 0.73472 | 1.850981 | 0.396936 | 0.691415 | 0.941994 |
| mRpS25 | 2146.432 | -0.36029 | 0.90803 | -0.39678 | 0.691531 | 0.941994 |
| CG11964 | 563.0051 | 0.643654 | 1.627092 | 0.395585 | 0.692411 | 0.942873 |
| Pisd | 1404.396 | 0.358176 | 0.905293 | 0.395647 | 0.692365 | 0.942873 |
| Rab11 | 5419.341 | -0.22923 | 0.580122 | -0.39515 | 0.692733 | 0.943103 |
| shrb | 2229.817 | -0.35553 | 0.900003 | -0.39504 | 0.692815 | 0.943103 |
| OtopLa | 10048.16 | 0.212811 | 0.538933 | 0.394875 | 0.692935 | 0.943106 |
| Itpr | 2514.264 | -0.27304 | 0.693181 | -0.39389 | 0.69366 | 0.943932 |
| crm | 1258.487 | 0.524395 | 1.347058 | 0.389289 | 0.697062 | 0.944152 |

| | | | | | | |
|---|---|---|---|---|---|---|
| snf | 3439.647 | -0.25094 | 0.64277 | -0.3904 | 0.696241 | 0.944152 |
| Spat | 131.0929 | 1.538476 | 3.951182 | 0.389371 | 0.697002 | 0.944152 |
| CG11449 | 131.4068 | 1.538478 | 3.951181 | 0.389372 | 0.697001 | 0.944152 |
| CG31643 | 328.5189 | 0.912018 | 2.319673 | 0.393166 | 0.694197 | 0.944152 |
| Wee1 | 3378.898 | 0.286232 | 0.732079 | 0.390985 | 0.695808 | 0.944152 |
| beat-IIIc | 1953.726 | -0.2972 | 0.764528 | -0.38874 | 0.697468 | 0.944152 |
| CG31793 | 1442.846 | 0.436536 | 1.121455 | 0.389258 | 0.697085 | 0.944152 |
| Crz | 71.24844 | 1.537616 | 3.951413 | 0.389131 | 0.69718 | 0.944152 |
| trem | 753.3716 | -0.82849 | 2.130257 | -0.38892 | 0.697338 | 0.944152 |
| tsl | 122.0953 | 1.5384 | 3.951202 | 0.38935 | 0.697017 | 0.944152 |
| Fadd | 602.1836 | -0.66264 | 1.684722 | -0.39332 | 0.69408 | 0.944152 |
| Nepl13 | 137.7888 | 1.538521 | 3.951168 | 0.389384 | 0.696992 | 0.944152 |
| CG13614 | 88.6159 | 1.537977 | 3.951314 | 0.389232 | 0.697105 | 0.944152 |
| Jasper | 932.6639 | 0.619134 | 1.592076 | 0.388885 | 0.697361 | 0.944152 |
| CG16739 | 73.7594 | 1.537661 | 3.951396 | 0.389144 | 0.69717 | 0.944152 |
| CG9294 | 62.66933 | 1.537362 | 3.951482 | 0.389059 | 0.697232 | 0.944152 |
| Ubi-p5E | 6870.265 | 0.199424 | 0.510915 | 0.390327 | 0.696295 | 0.944152 |
| 5-HT2A | 333.47 | -0.85499 | 2.17584 | -0.39295 | 0.694359 | 0.944152 |
| CG42668 | 5364.161 | 0.23248 | 0.596627 | 0.389657 | 0.69679 | 0.944152 |
| SmD1 | 632.9754 | 0.519616 | 1.329817 | 0.390743 | 0.695987 | 0.944152 |
| CG43125 | 75.22412 | 1.537715 | 3.951386 | 0.389158 | 0.697159 | 0.944152 |
| lncRNA:CR | 120.8407 | 0.974811 | 2.479538 | 0.393142 | 0.694214 | 0.944152 |
| CG43245 | 75.1195 | 1.537712 | 3.951387 | 0.389158 | 0.69716 | 0.944152 |
| CG30416 | 110.3775 | 1.538279 | 3.951234 | 0.389316 | 0.697042 | 0.944152 |
| Brca2 | 176.3172 | 0.93732 | 2.390524 | 0.392098 | 0.694986 | 0.944152 |
| lncRNA:CR | 68.10974 | 1.537531 | 3.951436 | 0.389107 | 0.697197 | 0.944152 |
| Cpr64Ab | 81.18765 | 1.537846 | 3.951351 | 0.389195 | 0.697132 | 0.944152 |
| CG12607 | 84.84946 | 1.537916 | 3.951332 | 0.389215 | 0.697117 | 0.944152 |
| ArfGAP1 | 1188.953 | -0.57459 | 1.467468 | -0.39155 | 0.695392 | 0.944152 |
| CG3347 | 84.95409 | 1.537918 | 3.951331 | 0.389215 | 0.697117 | 0.944152 |
| BORCS5 | 1155.874 | 0.440991 | 1.135493 | 0.388369 | 0.697743 | 0.944365 |
| wapl | 2680.651 | -0.25892 | 0.667484 | -0.3879 | 0.698089 | 0.944423 |
| Pdha | 6457.076 | -0.22165 | 0.572231 | -0.38734 | 0.698504 | 0.944423 |
| Pex16 | 622.2818 | -0.60149 | 1.552858 | -0.38734 | 0.698501 | 0.944423 |
| Hex-C | 65.1737 | 1.509168 | 3.897239 | 0.38724 | 0.698578 | 0.944423 |
| Spn43Ab | 13928.57 | -0.17201 | 0.443782 | -0.38759 | 0.698317 | 0.944423 |
| Atg6 | 963.4562 | -0.40874 | 1.055632 | -0.3872 | 0.69861 | 0.944423 |
| CG10576 | 1748.521 | 0.416346 | 1.072692 | 0.388132 | 0.697918 | 0.944423 |
| CG6191 | 140.3805 | 0.842456 | 2.17748 | 0.386895 | 0.698834 | 0.944489 |
| Ca-alpha1T | 15146.71 | 0.201527 | 0.520992 | 0.386814 | 0.698894 | 0.944489 |
| Map205 | 1317.627 | -0.46201 | 1.195182 | -0.38656 | 0.699081 | 0.944496 |
| Hexo2 | 2749.563 | 0.317077 | 0.820402 | 0.38649 | 0.699134 | 0.944496 |
| Rab3 | 37684.72 | 0.158515 | 0.410861 | 0.385812 | 0.699636 | 0.944576 |
| CG2063 | 353.1432 | -0.72288 | 1.874646 | -0.38561 | 0.699786 | 0.944576 |
| lncRNA:CR | 87.59712 | 1.523059 | 3.951314 | 0.385456 | 0.699899 | 0.944576 |
| disco-r | 245.2561 | -0.78569 | 2.037416 | -0.38563 | 0.699768 | 0.944576 |
| lncRNA:CR | 91.9913 | 1.523858 | 3.951295 | 0.38566 | 0.699748 | 0.944576 |
| RhoGAP19 | 5302.179 | -0.22112 | 0.572992 | -0.3859 | 0.699569 | 0.944576 |
| Tbp | 415.3431 | 0.855978 | 2.222257 | 0.385184 | 0.700101 | 0.944689 |
| Rad23 | 5713.38 | 0.216494 | 0.562379 | 0.384962 | 0.700266 | 0.944753 |

| | | | | | | |
|---|---|---|---|---|---|---|
| Rheb | 4839.388 | -0.2224 | 0.578359 | -0.38453 | 0.700583 | 0.944863 |
| snoRNA:M | 239.2719 | 0.802634 | 2.086714 | 0.38464 | 0.700504 | 0.944863 |
| CG12560 | 358.272 | 0.842269 | 2.197787 | 0.383235 | 0.701545 | 0.944891 |
| CG9986 | 841.6395 | -0.58213 | 1.516364 | -0.3839 | 0.701056 | 0.944891 |
| CG14441 | 1702.055 | 0.315546 | 0.823332 | 0.383255 | 0.701531 | 0.944891 |
| Mst77F | 664.0503 | -0.63684 | 1.66102 | -0.3834 | 0.70142 | 0.944891 |
| alphaSnap | 14435.69 | 0.168705 | 0.439315 | 0.384018 | 0.700965 | 0.944891 |
| CG12851 | 2858.006 | -0.30038 | 0.782072 | -0.38408 | 0.700918 | 0.944891 |
| lncRNA:CR | 215.9131 | 0.727163 | 1.896229 | 0.383478 | 0.701365 | 0.944891 |
| Ten-a | 19024.37 | 0.178556 | 0.465844 | 0.383295 | 0.701501 | 0.944891 |
| CG15930 | 149.793 | 0.888766 | 2.323046 | 0.382586 | 0.702026 | 0.944904 |
| mRpL10 | 1039.145 | -0.39717 | 1.037931 | -0.38266 | 0.701973 | 0.944904 |
| Dlip3 | 2090.515 | -0.33846 | 0.884445 | -0.38268 | 0.701959 | 0.944904 |
| rad | 5256.551 | 0.234961 | 0.613777 | 0.382812 | 0.701859 | 0.944904 |
| Lim1 | 1477.308 | -0.3251 | 0.850724 | -0.38215 | 0.702353 | 0.945064 |
| CG6610 | 110.4254 | -1.17995 | 3.087989 | -0.38211 | 0.702381 | 0.945064 |
| CRIF | 1448.77 | 0.318477 | 0.839374 | 0.379422 | 0.704374 | 0.94524 |
| CG11983 | 71.40318 | 1.137857 | 2.992371 | 0.380253 | 0.703758 | 0.94524 |
| wkd | 3211.496 | 0.296516 | 0.779334 | 0.380474 | 0.703594 | 0.94524 |
| Hr96 | 963.9877 | -0.57487 | 1.514472 | -0.37959 | 0.704252 | 0.94524 |
| CG1416 | 2305.115 | 0.295962 | 0.777757 | 0.380533 | 0.70355 | 0.94524 |
| Panx | 471.8407 | 0.73844 | 1.946181 | 0.37943 | 0.704368 | 0.94524 |
| CSN4 | 971.7118 | -0.41262 | 1.085129 | -0.38025 | 0.703759 | 0.94524 |
| Gpb5 | 1213.113 | -0.34991 | 0.920246 | -0.38024 | 0.703769 | 0.94524 |
| AGO2 | 6154.136 | -0.19728 | 0.520029 | -0.37937 | 0.704416 | 0.94524 |
| lncRNA:CR | 85.39159 | -1.49957 | 3.951301 | -0.37951 | 0.704306 | 0.94524 |
| IP3K2 | 1239.091 | 0.346032 | 0.907452 | 0.381322 | 0.702964 | 0.94524 |
| CG4239 | 1901.546 | -0.33155 | 0.874206 | -0.37926 | 0.704492 | 0.94524 |
| par-6 | 2207.338 | -0.27272 | 0.718664 | -0.37948 | 0.704329 | 0.94524 |
| CG3902 | 1521.256 | 0.405598 | 1.069515 | 0.379235 | 0.704513 | 0.94524 |
| CG7430 | 6633.885 | -0.22344 | 0.588146 | -0.37991 | 0.704015 | 0.94524 |
| Notum | 77.3715 | 1.50453 | 3.951362 | 0.380763 | 0.703379 | 0.94524 |
| Prp31 | 427.4255 | 0.693394 | 1.818953 | 0.381205 | 0.703051 | 0.94524 |
| CG4502 | 2831.976 | 0.291801 | 0.775109 | 0.376464 | 0.706572 | 0.945834 |
| Taf11 | 345.4748 | -0.84385 | 2.238792 | -0.37692 | 0.706233 | 0.945834 |
| CG4596 | 342.8043 | -0.82409 | 2.188321 | -0.37658 | 0.706483 | 0.945834 |
| CG6693 | 1800.641 | 0.311371 | 0.826874 | 0.376564 | 0.706498 | 0.945834 |
| Sbf | 2001.508 | -0.32656 | 0.863624 | -0.37812 | 0.705339 | 0.945834 |
| CG5805 | 1368.769 | -0.33922 | 0.90118 | -0.37642 | 0.706606 | 0.945834 |
| CG8320 | 708.6535 | 0.424249 | 1.123511 | 0.37761 | 0.70572 | 0.945834 |
| GstT2 | 227.9505 | 0.663743 | 1.75888 | 0.377367 | 0.705901 | 0.945834 |
| Nfl | 10896.11 | 0.206262 | 0.547782 | 0.376541 | 0.706515 | 0.945834 |
| kek5 | 363.3596 | -0.63327 | 1.68011 | -0.37692 | 0.706232 | 0.945834 |
| CG34404 | 2636.468 | 0.311807 | 0.827227 | 0.37693 | 0.706225 | 0.945834 |
| lncRNA:CR | 81.58778 | -1.40421 | 3.71864 | -0.37761 | 0.705717 | 0.945834 |
| dpr6 | 8856.569 | 0.215368 | 0.569999 | 0.377839 | 0.70555 | 0.945834 |
| Tgt | 220.5575 | 0.981981 | 2.607655 | 0.376576 | 0.706489 | 0.945834 |
| Pdp | 5557.222 | -0.22079 | 0.587312 | -0.37593 | 0.706968 | 0.946042 |
| CG10866 | 1961.503 | -0.30845 | 0.820823 | -0.37579 | 0.707075 | 0.946042 |
| fax | 7310.236 | 0.197256 | 0.525592 | 0.375303 | 0.707435 | 0.946042 |

| | | | | | | |
|---|---|---|---|---|---|---|
| Sting | 633.0212 | 0.513349 | 1.367779 | 0.375316 | 0.707426 | 0.946042 |
| ringer | 5243.462 | 0.216199 | 0.575464 | 0.375695 | 0.707144 | 0.946042 |
| CG11095 | 191.8577 | -1.09346 | 2.913905 | -0.37526 | 0.707469 | 0.946042 |
| Cp1 | 3398.058 | 0.24414 | 0.650958 | 0.375048 | 0.707625 | 0.946093 |
| CG5608 | 905.87 | -0.62505 | 1.667438 | -0.37485 | 0.707769 | 0.946129 |
| Ripalpha | 854.6685 | 0.443484 | 1.187072 | 0.373595 | 0.708706 | 0.946964 |
| mlt | 12984.2 | 0.191056 | 0.511695 | 0.373379 | 0.708866 | 0.946964 |
| CanA-14F | 3935.799 | -0.23347 | 0.625274 | -0.37339 | 0.708858 | 0.946964 |
| OXA1L | 940.0198 | 0.376704 | 1.008069 | 0.373689 | 0.708636 | 0.946964 |
| CG13671 | 522.0322 | -0.56931 | 1.526882 | -0.37286 | 0.709252 | 0.947164 |
| MFS18 | 1050.025 | 0.445871 | 1.195396 | 0.37299 | 0.709156 | 0.947164 |
| CG13083 | 93.35659 | 1.471151 | 3.951275 | 0.372323 | 0.709652 | 0.947541 |
| phol | 895.6517 | -0.57363 | 1.541557 | -0.37211 | 0.709809 | 0.947593 |
| nom | 345.8434 | 0.636873 | 1.717564 | 0.3708 | 0.710786 | 0.947794 |
| CG15536 | 396.2652 | -0.85109 | 2.292729 | -0.37121 | 0.710479 | 0.947794 |
| Sodh-2 | 385.6235 | 0.53115 | 1.429355 | 0.371601 | 0.71019 | 0.947794 |
| Vrp1 | 4142.697 | -0.25754 | 0.693666 | -0.37128 | 0.710432 | 0.947794 |
| Cdep | 14743.6 | -0.15926 | 0.429492 | -0.37081 | 0.710781 | 0.947794 |
| asRNA:CR4 | 142.9108 | 0.951814 | 2.566484 | 0.370863 | 0.71074 | 0.947794 |
| cmb | 246.9786 | 0.905348 | 2.441216 | 0.370859 | 0.710742 | 0.947794 |
| mnd | 2617.995 | 0.310671 | 0.838463 | 0.370524 | 0.710992 | 0.94791 |
| ND-B14.7 | 7324.365 | 0.181904 | 0.491765 | 0.3699 | 0.711457 | 0.948373 |
| CG5903 | 12253.59 | -0.16947 | 0.458534 | -0.3696 | 0.711684 | 0.948518 |
| CG5078 | 247.3124 | -1.23718 | 3.376848 | -0.36637 | 0.714089 | 0.949044 |
| PolZ2 | 499.8365 | -0.79862 | 2.178806 | -0.36654 | 0.713961 | 0.949044 |
| STUB1 | 2862.948 | -0.26177 | 0.713944 | -0.36665 | 0.713881 | 0.949044 |
| CG9467 | 7273.006 | -0.20434 | 0.556638 | -0.3671 | 0.713545 | 0.949044 |
| Mitofilin | 2870.061 | -0.26937 | 0.734054 | -0.36696 | 0.713648 | 0.949044 |
| CG13631 | 7829.689 | -0.22142 | 0.602481 | -0.36751 | 0.713241 | 0.949044 |
| ssh | 620.8564 | -0.45143 | 1.224514 | -0.36866 | 0.712381 | 0.949044 |
| HmgD | 3201.713 | -0.25668 | 0.700478 | -0.36643 | 0.714041 | 0.949044 |
| Hmgs | 1781.183 | -0.46601 | 1.271365 | -0.36654 | 0.71396 | 0.949044 |
| dpr1 | 17025.73 | 0.159 | 0.43387 | 0.36647 | 0.714014 | 0.949044 |
| CNMaR | 660.7139 | 0.65638 | 1.79049 | 0.366592 | 0.713923 | 0.949044 |
| Octbeta1R | 1763.776 | 0.289372 | 0.787416 | 0.367496 | 0.713249 | 0.949044 |
| CG40228 | 5445.466 | -0.20043 | 0.545617 | -0.36735 | 0.713357 | 0.949044 |
| CG42540 | 19522.12 | 0.157636 | 0.428833 | 0.367593 | 0.713177 | 0.949044 |
| Ca-Ma2d | 630.1252 | 0.520985 | 1.421727 | 0.366445 | 0.714033 | 0.949044 |
| smt3 | 8285.176 | -0.17585 | 0.47722 | -0.36848 | 0.712513 | 0.949044 |
| CG14135 | 393.7632 | 0.678889 | 1.843358 | 0.368289 | 0.712658 | 0.949044 |
| CG7903 | 957.0468 | 0.475939 | 1.300772 | 0.36589 | 0.714447 | 0.949206 |
| Aps | 3288.003 | 0.228478 | 0.624319 | 0.365964 | 0.714392 | 0.949206 |
| dpr20 | 1560.778 | -0.30707 | 0.840145 | -0.36549 | 0.714743 | 0.949284 |
| mRpL36 | 1247.282 | -0.3255 | 0.890502 | -0.36552 | 0.71472 | 0.949284 |
| CG7483 | 833.1987 | -0.39051 | 1.072505 | -0.36411 | 0.715776 | 0.949338 |
| RIC-3 | 4434.662 | -0.22372 | 0.615651 | -0.36338 | 0.716321 | 0.949338 |
| Kdm4B | 826.3339 | -0.64696 | 1.780288 | -0.3634 | 0.716306 | 0.949338 |
| CG11076 | 4932.741 | -0.203 | 0.557995 | -0.36381 | 0.715999 | 0.949338 |
| plx | 1854.346 | -0.28028 | 0.768624 | -0.36465 | 0.715374 | 0.949338 |
| RpS6 | 21200.84 | 0.154727 | 0.425243 | 0.363856 | 0.715965 | 0.949338 |

| | | | | | | |
|---|---|---|---|---|---|---|
| pyd | 7260.574 | -0.20397 | 0.560219 | -0.36409 | 0.715794 | 0.949338 |
| CG3511 | 1369.289 | 0.550651 | 1.511198 | 0.364381 | 0.715574 | 0.949338 |
| nod | 101.5602 | 1.442325 | 3.951311 | 0.365024 | 0.715093 | 0.949338 |
| Taf6 | 1313.966 | -0.44337 | 1.215237 | -0.36484 | 0.71523 | 0.949338 |
| MED11 | 607.3525 | 0.444308 | 1.218983 | 0.364491 | 0.715491 | 0.949338 |
| CG17364 | 2761.989 | 0.262489 | 0.722195 | 0.36346 | 0.716261 | 0.949338 |
| ERR | 293.7886 | 0.605989 | 1.659221 | 0.365225 | 0.714943 | 0.949338 |
| CycY | 2147.206 | -0.27105 | 0.748545 | -0.36211 | 0.717271 | 0.949565 |
| Dtd | 260.5691 | 0.659087 | 1.824137 | 0.361314 | 0.717864 | 0.949565 |
| rin | 4875.308 | -0.20254 | 0.559307 | -0.36212 | 0.717262 | 0.949565 |
| His4r | 52266.4 | 0.146224 | 0.404241 | 0.361723 | 0.717559 | 0.949565 |
| CG11110 | 738.7747 | -0.60749 | 1.6831 | -0.36093 | 0.71815 | 0.949565 |
| CG8613 | 3439.179 | -0.2559 | 0.707135 | -0.36189 | 0.717437 | 0.949565 |
| Tmem18 | 558.7611 | -0.45918 | 1.266461 | -0.36257 | 0.716928 | 0.949565 |
| brk | 1515.293 | 0.304489 | 0.841249 | 0.361948 | 0.717391 | 0.949565 |
| slim | 9298.784 | 0.193136 | 0.533471 | 0.362037 | 0.717324 | 0.949565 |
| ebi | 1199.984 | 0.352461 | 0.976755 | 0.360849 | 0.718213 | 0.949565 |
| Tom20 | 6663.989 | 0.187091 | 0.51761 | 0.361451 | 0.717762 | 0.949565 |
| FucTA | 2218.731 | 0.327105 | 0.906669 | 0.360776 | 0.718267 | 0.949565 |
| Tdrd3 | 412.909 | 0.51428 | 1.421904 | 0.361684 | 0.717588 | 0.949565 |
| Srp9 | 1617.52 | 0.301416 | 0.834732 | 0.361094 | 0.718029 | 0.949565 |
| mRpL27 | 1752.405 | -0.30377 | 0.838109 | -0.36245 | 0.717019 | 0.949565 |
| CG5938 | 4004.045 | -0.21811 | 0.604986 | -0.36052 | 0.718458 | 0.949661 |
| lncRNA:Hs | 380337.8 | -0.14106 | 0.391832 | -0.35999 | 0.718853 | 0.950027 |
| mthl14 | 616.625 | 0.567672 | 1.578319 | 0.359669 | 0.719095 | 0.95019 |
| AdamTS-B | 85.83307 | 1.128649 | 3.153413 | 0.357914 | 0.720408 | 0.95047 |
| crq | 3507.32 | 0.220001 | 0.616501 | 0.356854 | 0.721201 | 0.95047 |
| plum | 4619.678 | -0.22124 | 0.619003 | -0.35741 | 0.720785 | 0.95047 |
| CG9344 | 485.4665 | -0.52215 | 1.463041 | -0.35689 | 0.721172 | 0.95047 |
| CG1888 | 1528.566 | 0.39693 | 1.112217 | 0.356881 | 0.721181 | 0.95047 |
| AIMP1 | 661.2927 | 0.702251 | 1.966397 | 0.357126 | 0.720998 | 0.95047 |
| asRNA:CR3 | 114.7376 | 1.400651 | 3.914128 | 0.357845 | 0.720459 | 0.95047 |
| CG17454 | 5802.278 | 0.197754 | 0.5523 | 0.358055 | 0.720302 | 0.95047 |
| snama | 919.8431 | 0.508431 | 1.417575 | 0.358662 | 0.719848 | 0.95047 |
| miple1 | 40312.28 | 0.138079 | 0.386677 | 0.357091 | 0.721024 | 0.95047 |
| Tim17b | 11343.93 | 0.169115 | 0.472266 | 0.358094 | 0.720273 | 0.95047 |
| Sgt1 | 1456.127 | 0.324014 | 0.906489 | 0.357438 | 0.720764 | 0.95047 |
| CR46425 | 381.8444 | 0.613221 | 1.712002 | 0.358189 | 0.720202 | 0.95047 |
| lncRNA:CR | 278.743 | -0.75089 | 2.095149 | -0.35839 | 0.720049 | 0.95047 |
| CG7550 | 271.881 | 0.692484 | 1.931702 | 0.358484 | 0.719981 | 0.95047 |
| slp1 | 486.8744 | 0.802715 | 2.235794 | 0.359029 | 0.719573 | 0.95047 |
| Rpn11 | 2084.183 | 0.265808 | 0.746377 | 0.356131 | 0.721742 | 0.950531 |
| CAH7 | 32752.28 | -0.17683 | 0.496919 | -0.35584 | 0.721958 | 0.950531 |
| e(y)3 | 1176.331 | -0.36063 | 1.012533 | -0.35616 | 0.721718 | 0.950531 |
| pit | 397.3959 | -0.68972 | 1.937651 | -0.35596 | 0.721872 | 0.950531 |
| SF2 | 3064.214 | -0.24601 | 0.690027 | -0.35653 | 0.721445 | 0.950531 |
| RpS21 | 5029.494 | 0.20543 | 0.576943 | 0.356066 | 0.721791 | 0.950531 |
| metro | 1254.64 | -0.45608 | 1.282344 | -0.35566 | 0.722094 | 0.950554 |
| scrt | 27235.48 | 0.154446 | 0.434562 | 0.355406 | 0.722285 | 0.95065 |
| CG3397 | 255.0392 | 0.928484 | 2.615126 | 0.355044 | 0.722557 | 0.950749 |

| | | | | | | |
|---|---|---|---|---|---|---|
| CG11123 | 533.091 | 0.705204 | 1.988321 | 0.354673 | 0.722834 | 0.950749 |
| Sesn | 201.636 | -0.81643 | 2.300099 | -0.35496 | 0.722622 | 0.950749 |
| mRpS17 | 1369.265 | 0.330191 | 0.930717 | 0.35477 | 0.722762 | 0.950749 |
| CG5888 | 4396.653 | -0.23613 | 0.666698 | -0.35418 | 0.723203 | 0.950766 |
| Drat | 10061.78 | 0.168608 | 0.476039 | 0.354188 | 0.723198 | 0.950766 |
| app | 3221.787 | -0.23356 | 0.658901 | -0.35447 | 0.72299 | 0.950766 |
| MED31 | 1921.526 | -0.29321 | 0.829194 | -0.3536 | 0.723636 | 0.951073 |
| grass | 92.57819 | 1.145656 | 3.240397 | 0.353554 | 0.723673 | 0.951073 |
| CG31999 | 3381.486 | 0.243456 | 0.689445 | 0.353119 | 0.723999 | 0.951345 |
| CG11007 | 2009.891 | 0.298103 | 0.846326 | 0.352232 | 0.724665 | 0.951993 |
| Strip | 182.6338 | -0.76661 | 2.176983 | -0.35215 | 0.724729 | 0.951993 |
| lok | 280.5502 | -0.74999 | 2.131017 | -0.35194 | 0.724884 | 0.952041 |
| Lkb1 | 483.9113 | 0.66241 | 1.887089 | 0.351022 | 0.725572 | 0.95266 |
| dare | 585.302 | 0.552559 | 1.574272 | 0.350993 | 0.725593 | 0.95266 |
| CG11839 | 139.1557 | -1.00335 | 2.860765 | -0.35073 | 0.725793 | 0.95272 |
| dom | 3470.887 | -0.23505 | 0.6707 | -0.35046 | 0.725995 | 0.95272 |
| Prosalpha6 | 2596.177 | 0.441806 | 1.260415 | 0.350524 | 0.725945 | 0.95272 |
| T48 | 112.8313 | 0.933716 | 2.666971 | 0.350104 | 0.726261 | 0.952913 |
| Zir | 4009.219 | 0.224068 | 0.641166 | 0.34947 | 0.726736 | 0.953029 |
| CG11882 | 114.8885 | -1.29495 | 3.70282 | -0.34972 | 0.726549 | 0.953029 |
| CG12159 | 1992.359 | -0.26071 | 0.745714 | -0.34961 | 0.726628 | 0.953029 |
| CG9577 | 785.6772 | -0.49865 | 1.427727 | -0.34926 | 0.726892 | 0.953029 |
| CG13928 | 19194.26 | 0.16273 | 0.466021 | 0.349191 | 0.726946 | 0.953029 |
| Tango10 | 1522.239 | 0.32729 | 0.937693 | 0.349037 | 0.727061 | 0.953029 |
| LSm7 | 465.8023 | -0.46502 | 1.33407 | -0.34857 | 0.72741 | 0.953331 |
| Axn | 1019.92 | 0.34635 | 0.994431 | 0.348289 | 0.727623 | 0.953453 |
| Art4 | 551.5491 | -0.57961 | 1.669779 | -0.34712 | 0.728501 | 0.953676 |
| pirk | 464.7927 | 0.693088 | 1.994973 | 0.347417 | 0.728278 | 0.953676 |
| CG42699 | 271.1916 | 0.837726 | 2.408397 | 0.347836 | 0.727964 | 0.953676 |
| nrm | 7044.866 | 0.174327 | 0.502219 | 0.347113 | 0.728506 | 0.953676 |
| CG43367 | 3895.417 | 0.21095 | 0.606855 | 0.347613 | 0.728131 | 0.953676 |
| CG15012 | 5590.322 | 0.18692 | 0.538332 | 0.347221 | 0.728426 | 0.953676 |
| CG8441 | 1611.063 | -0.52314 | 1.510764 | -0.34628 | 0.729135 | 0.953878 |
| SC35 | 2998.917 | 0.225905 | 0.652132 | 0.346409 | 0.729035 | 0.953878 |
| CG13716 | 361.8708 | -0.65468 | 1.890036 | -0.34638 | 0.729054 | 0.953878 |
| CG3808 | 393.024 | -0.52595 | 1.518362 | -0.34639 | 0.729049 | 0.953878 |
| rngo | 345.4813 | 0.706152 | 2.040851 | 0.346009 | 0.729336 | 0.953985 |
| beat-Ia | 1874.236 | -0.29357 | 0.849854 | -0.34543 | 0.729771 | 0.95438 |
| CG11178 | 246.2627 | -0.85123 | 2.466166 | -0.34516 | 0.729971 | 0.95438 |
| CG9723 | 176.1903 | -0.96715 | 2.802263 | -0.34513 | 0.729995 | 0.95438 |
| CG5316 | 551.3561 | -0.49898 | 1.450944 | -0.3439 | 0.730923 | 0.954873 |
| ND-B15 | 4476.666 | 0.195981 | 0.569697 | 0.344009 | 0.73084 | 0.954873 |
| bru2 | 106.6309 | -1.17294 | 3.408447 | -0.34413 | 0.730751 | 0.954873 |
| CG9065 | 6980.891 | 0.234355 | 0.681584 | 0.34384 | 0.730967 | 0.954873 |
| CG6847 | 416.1217 | -0.61827 | 1.796277 | -0.3442 | 0.730698 | 0.954873 |
| RpS20 | 9619.395 | -0.16318 | 0.474894 | -0.34362 | 0.731132 | 0.954911 |
| mRpL52 | 1374.698 | 0.306449 | 0.892354 | 0.343417 | 0.731285 | 0.954911 |
| RNaseX25 | 1737.235 | -0.34042 | 0.991522 | -0.34333 | 0.731353 | 0.954911 |
| CG42271 | 1846.02 | 0.261095 | 0.760927 | 0.343127 | 0.731503 | 0.954952 |
| CG32448 | 756.8622 | 0.416086 | 1.219418 | 0.341217 | 0.732941 | 0.95549 |

| | | | | | | |
|---|---|---|---|---|---|---|
| CG10958 | 290.3258 | 0.955027 | 2.799388 | 0.341156 | 0.732986 | 0.95549 |
| AP-1gamm | 3592.114 | 0.21535 | 0.629617 | 0.342033 | 0.732326 | 0.95549 |
| CG11474 | 2405.73 | 0.351936 | 1.027814 | 0.342413 | 0.73204 | 0.95549 |
| CG8180 | 262.3936 | 0.570339 | 1.669546 | 0.341613 | 0.732642 | 0.95549 |
| hoe2 | 826.4251 | -0.63034 | 1.843029 | -0.34201 | 0.732343 | 0.95549 |
| apolpp | 9364.012 | 0.158352 | 0.463797 | 0.341425 | 0.732784 | 0.95549 |
| CG42748 | 6202.251 | -0.17808 | 0.521722 | -0.34134 | 0.732849 | 0.95549 |
| mbl | 24873.02 | 0.186366 | 0.545675 | 0.341533 | 0.732702 | 0.95549 |
| Hn | 86.67969 | -0.97944 | 2.872693 | -0.34095 | 0.733144 | 0.955539 |
| CG10462 | 525.4799 | -0.57255 | 1.680713 | -0.34066 | 0.733361 | 0.955668 |
| Wbp2 | 9416.726 | 0.15878 | 0.466862 | 0.340102 | 0.73378 | 0.956058 |
| CG15537 | 4850.899 | 0.221861 | 0.653763 | 0.33936 | 0.734339 | 0.956631 |
| Nmd3 | 894.338 | -0.39075 | 1.153427 | -0.33877 | 0.734781 | 0.956746 |
| lncRNA:CR | 116.6638 | 1.299461 | 3.834104 | 0.338922 | 0.734669 | 0.956746 |
| CG10654 | 1963.172 | -0.29779 | 0.879029 | -0.33877 | 0.734785 | 0.956746 |
| RpS27 | 8717.519 | 0.169675 | 0.501175 | 0.338555 | 0.734945 | 0.9568 |
| CG31522 | 3255.197 | 0.213558 | 0.63477 | 0.336434 | 0.736544 | 0.956985 |
| homer | 1728.739 | -0.33508 | 1.001146 | -0.3347 | 0.737851 | 0.956985 |
| CG31897 | 77.56609 | 1.278461 | 3.796358 | 0.33676 | 0.736298 | 0.956985 |
| Ugt37D1 | 909.1081 | 0.574094 | 1.699748 | 0.337752 | 0.73555 | 0.956985 |
| CG12402 | 683.2935 | -0.46096 | 1.376146 | -0.33496 | 0.737653 | 0.956985 |
| CG31156 | 540.0585 | -0.54965 | 1.635157 | -0.33614 | 0.736762 | 0.956985 |
| CG33107 | 1802.7 | 0.324455 | 0.962199 | 0.337201 | 0.735965 | 0.956985 |
| msi | 11326.03 | -0.15422 | 0.459574 | -0.33558 | 0.73719 | 0.956985 |
| galla-1 | 2957.018 | -0.30585 | 0.91432 | -0.33451 | 0.737991 | 0.956985 |
| GIIIspla2 | 975.968 | -0.34048 | 1.007356 | -0.338 | 0.735367 | 0.956985 |
| CG12659 | 343.1878 | 0.748312 | 2.239088 | 0.334204 | 0.738226 | 0.956985 |
| CG15312 | 2124.561 | -0.28082 | 0.83838 | -0.33495 | 0.737661 | 0.956985 |
| CG10082 | 4813.466 | -0.18919 | 0.565309 | -0.33466 | 0.73788 | 0.956985 |
| CG1418 | 705.964 | -0.66281 | 1.968748 | -0.33667 | 0.736369 | 0.956985 |
| ced-6 | 4653.738 | 0.225845 | 0.675862 | 0.334158 | 0.73826 | 0.956985 |
| Coq9 | 618.5577 | 0.509908 | 1.509158 | 0.337876 | 0.735457 | 0.956985 |
| sw | 760.0287 | -0.42936 | 1.273747 | -0.33708 | 0.736056 | 0.956985 |
| sona | 438.1611 | -0.6654 | 1.971464 | -0.33751 | 0.73573 | 0.956985 |
| C3G | 4248.7 | 0.253127 | 0.754029 | 0.335699 | 0.737098 | 0.956985 |
| CG42360 | 1086.59 | 0.331749 | 0.988699 | 0.335541 | 0.737217 | 0.956985 |
| RpL8 | 15388.23 | -0.15627 | 0.467386 | -0.33435 | 0.738118 | 0.956985 |
| TyrRS-m | 159.8571 | 1.012185 | 3.027187 | 0.334365 | 0.738104 | 0.956985 |
| dally | 19831.58 | -0.19125 | 0.570982 | -0.33495 | 0.737665 | 0.956985 |
| Sem1 | 1187.301 | -0.30736 | 0.919981 | -0.3341 | 0.738307 | 0.956985 |
| CG9149 | 472.7486 | -0.62592 | 1.870355 | -0.33465 | 0.737887 | 0.956985 |
| CG15014 | 592.9777 | -0.65948 | 1.960468 | -0.33639 | 0.736576 | 0.956985 |
| Cds | 15423.79 | -0.15214 | 0.453778 | -0.33528 | 0.737417 | 0.956985 |
| CG14621 | 74.13902 | 1.163042 | 3.485961 | 0.333636 | 0.738654 | 0.957281 |
| CG31139 | 209.3177 | 1.228542 | 3.684034 | 0.333477 | 0.738774 | 0.957281 |
| CG6766 | 4923.616 | 0.268522 | 0.806011 | 0.333149 | 0.739022 | 0.957448 |
| inc | 4181.918 | -0.24816 | 0.74608 | -0.33262 | 0.739422 | 0.957812 |
| Pgd | 582.5518 | -0.58773 | 1.78521 | -0.32922 | 0.741986 | 0.958464 |
| CG10979 | 541.4332 | 0.551977 | 1.671006 | 0.330326 | 0.741153 | 0.958464 |
| CG10413 | 1292.82 | 0.446084 | 1.355541 | 0.329082 | 0.742094 | 0.958464 |

| | | | | | | |
|---|---|---|---|---|---|---|
| CG10098 | 1673.352 | -0.43737 | 1.322761 | -0.33065 | 0.740909 | 0.958464 |
| Sap47 | 40196.45 | 0.128257 | 0.389251 | 0.329497 | 0.74178 | 0.958464 |
| ATPsyndel | 9307.079 | -0.16839 | 0.507653 | -0.3317 | 0.740113 | 0.958464 |
| CG12643 | 327.0235 | 0.790165 | 2.396917 | 0.329659 | 0.741658 | 0.958464 |
| CG13148 | 351.3575 | 0.669635 | 2.035684 | 0.328948 | 0.742195 | 0.958464 |
| CG17977 | 1775.828 | -0.28103 | 0.848377 | -0.33126 | 0.740449 | 0.958464 |
| unc80 | 5092.058 | 0.192505 | 0.585098 | 0.329014 | 0.742145 | 0.958464 |
| mRpL24 | 2521.481 | -0.2333 | 0.708998 | -0.32905 | 0.742115 | 0.958464 |
| endos | 5588.047 | -0.19107 | 0.579708 | -0.32959 | 0.74171 | 0.958464 |
| CG3257 | 2618.635 | -0.2398 | 0.727476 | -0.32964 | 0.741673 | 0.958464 |
| alpha-Spec | 8177.289 | 0.168095 | 0.50963 | 0.329838 | 0.741522 | 0.958464 |
| nero | 1408.637 | -0.43128 | 1.305928 | -0.33025 | 0.741212 | 0.958464 |
| CG42751 | 107.457 | -1.16339 | 3.535045 | -0.3291 | 0.742079 | 0.958464 |
| Acn | 4127.881 | 0.194982 | 0.591049 | 0.329892 | 0.741481 | 0.958464 |
| Argk | 53934.54 | -0.12302 | 0.372823 | -0.32996 | 0.741431 | 0.958464 |
| Sf3b6 | 2028.665 | -0.24303 | 0.738439 | -0.32911 | 0.742075 | 0.958464 |
| meng | 1361.341 | 0.305911 | 0.936517 | 0.326648 | 0.743934 | 0.958549 |
| CG13398 | 1545.016 | 0.290838 | 0.89081 | 0.326487 | 0.744056 | 0.958549 |
| Mst89B | 203.2573 | 1.22087 | 3.721649 | 0.328045 | 0.742877 | 0.958549 |
| Vha100-1 | 16898.57 | -0.15907 | 0.484047 | -0.32862 | 0.742443 | 0.958549 |
| Gycbeta10 | 3591.72 | 0.204508 | 0.625687 | 0.326853 | 0.743779 | 0.958549 |
| vimar | 959.8378 | 0.484861 | 1.478729 | 0.32789 | 0.742995 | 0.958549 |
| CG7065 | 855.5398 | -0.41001 | 1.256014 | -0.32643 | 0.744096 | 0.958549 |
| pic | 1336.499 | 0.309202 | 0.9475 | 0.326334 | 0.744171 | 0.958549 |
| tamo | 1692.043 | -0.27745 | 0.849914 | -0.32645 | 0.744086 | 0.958549 |
| CG45002 | 4160.666 | 0.211838 | 0.645686 | 0.328082 | 0.74285 | 0.958549 |
| da | 2255.889 | -0.25045 | 0.764958 | -0.3274 | 0.743365 | 0.958549 |
| Apl | 136.8548 | -0.70811 | 2.16752 | -0.32669 | 0.743901 | 0.958549 |
| Dab | 4265.164 | 0.198347 | 0.605856 | 0.327383 | 0.743378 | 0.958549 |
| CG14153 | 129.5452 | -0.96276 | 2.93949 | -0.32752 | 0.743271 | 0.958549 |
| CG7565 | 1460.777 | -0.36735 | 1.123292 | -0.32703 | 0.743645 | 0.958549 |
| CG5126 | 311.487 | 0.63323 | 1.939778 | 0.326445 | 0.744088 | 0.958549 |
| Fur1 | 3874.548 | -0.19377 | 0.596475 | -0.32486 | 0.745286 | 0.959831 |
| Unc-76 | 10200.46 | 0.147046 | 0.45799 | 0.321069 | 0.748158 | 0.959839 |
| CG16903 | 114.6255 | 0.872991 | 2.713441 | 0.321729 | 0.747658 | 0.959839 |
| CG2931 | 886.1565 | 0.357976 | 1.104237 | 0.324184 | 0.745799 | 0.959839 |
| CG7239 | 466.4155 | -0.4707 | 1.468433 | -0.32055 | 0.748555 | 0.959839 |
| MED20 | 287.5876 | 0.666377 | 2.086615 | 0.319358 | 0.749455 | 0.959839 |
| Gdi | 14600.12 | -0.14288 | 0.445389 | -0.3208 | 0.748361 | 0.959839 |
| Spn31A | 838.5208 | 0.580273 | 1.800311 | 0.322318 | 0.747212 | 0.959839 |
| CG5355 | 1276.237 | -0.31757 | 0.992645 | -0.31992 | 0.749025 | 0.959839 |
| CG18063 | 62.38617 | 1.236756 | 3.857997 | 0.32057 | 0.748537 | 0.959839 |
| Tsp39D | 2588.333 | 0.266776 | 0.824691 | 0.323486 | 0.746327 | 0.959839 |
| CG31301 | 767.6935 | 0.442469 | 1.381652 | 0.320246 | 0.748782 | 0.959839 |
| CG5346 | 160.0005 | 0.938052 | 2.899071 | 0.32357 | 0.746264 | 0.959839 |
| Orct | 7972.286 | 0.190088 | 0.588164 | 0.323189 | 0.746552 | 0.959839 |
| CG6607 | 578.1574 | -0.54815 | 1.714363 | -0.31974 | 0.749167 | 0.959839 |
| CG5913 | 272.9673 | 0.777702 | 2.434526 | 0.319447 | 0.749388 | 0.959839 |
| Sap-r | 10086.89 | 0.162522 | 0.505312 | 0.321628 | 0.747734 | 0.959839 |
| Hsf | 3110.38 | -0.22152 | 0.682775 | -0.32445 | 0.745601 | 0.959839 |

| | | | | | | |
|---|---|---|---|---|---|---|
| CG12895 | 1738.346 | 0.298222 | 0.934369 | 0.319169 | 0.749598 | 0.959839 |
| Csgalnact | 1996.474 | 0.323902 | 1.009839 | 0.320746 | 0.748403 | 0.959839 |
| Eaf | 1471.769 | -0.29613 | 0.917694 | -0.32269 | 0.746931 | 0.959839 |
| zormin | 2782.837 | -0.22415 | 0.694223 | -0.32288 | 0.746785 | 0.959839 |
| CG11883 | 3117.93 | -0.25065 | 0.783195 | -0.32003 | 0.748946 | 0.959839 |
| Ptp99A | 6445.899 | 0.167277 | 0.523857 | 0.319319 | 0.749485 | 0.959839 |
| SteXh:CG4 | 150.3966 | -0.97424 | 3.00851 | -0.32383 | 0.746069 | 0.959839 |
| Hk | 16371.49 | -0.13974 | 0.434707 | -0.32146 | 0.747864 | 0.959839 |
| CG4610 | 767.5352 | -0.63466 | 1.970421 | -0.3221 | 0.74738 | 0.959839 |
| ytr | 5416.028 | 0.197683 | 0.614973 | 0.32145 | 0.74787 | 0.959839 |
| lncRNA:CR | 118.5931 | -0.97167 | 3.025878 | -0.32112 | 0.748118 | 0.959839 |
| dmt | 207.6692 | -0.66351 | 2.053106 | -0.32317 | 0.746564 | 0.959839 |
| CG6023 | 562.9281 | -0.57883 | 1.809562 | -0.31987 | 0.749065 | 0.959839 |
| ftz-f1 | 10531.39 | 0.172655 | 0.532023 | 0.324526 | 0.74554 | 0.959839 |
| CG32187 | 139.8579 | 0.756236 | 2.346348 | 0.322303 | 0.747223 | 0.959839 |
| Arl1 | 630.8581 | -0.47628 | 1.489529 | -0.31976 | 0.749154 | 0.959839 |
| Sse | 175.09 | -0.90137 | 2.784769 | -0.32368 | 0.746181 | 0.959839 |
| ScsbetaG | 435.1072 | -0.72674 | 2.270451 | -0.32009 | 0.748902 | 0.959839 |
| tty | 3207.164 | 0.222254 | 0.688672 | 0.322729 | 0.746901 | 0.959839 |
| CG9396 | 168.9325 | 1.035798 | 3.247133 | 0.318988 | 0.749735 | 0.959862 |
| CG31974 | 3684.34 | -0.22868 | 0.718128 | -0.31843 | 0.750156 | 0.959996 |
| CG10209 | 148.7994 | 0.905768 | 2.844948 | 0.318378 | 0.750199 | 0.959996 |
| GstO2 | 1986.842 | -0.49001 | 1.537956 | -0.31861 | 0.750022 | 0.959996 |
| Sin3A | 9553.707 | -0.15559 | 0.488993 | -0.31818 | 0.75035 | 0.960036 |
| Ccp84Ab | 71.81387 | 1.230281 | 3.870352 | 0.317873 | 0.750581 | 0.960129 |
| Zasp67 | 79.33309 | 1.143229 | 3.597688 | 0.317768 | 0.750661 | 0.960129 |
| CG14427 | 85.10547 | -1.01727 | 3.211277 | -0.31678 | 0.751411 | 0.960934 |
| Top2 | 920.1402 | -0.38299 | 1.209643 | -0.31662 | 0.751535 | 0.96094 |
| Gnf1 | 687.6609 | -0.49967 | 1.579974 | -0.31625 | 0.75181 | 0.961069 |
| 128up | 1104.988 | -0.41693 | 1.320568 | -0.31572 | 0.752214 | 0.961069 |
| Snoo | 1992.938 | -0.28122 | 0.889984 | -0.31598 | 0.752018 | 0.961069 |
| roh | 15429.43 | 0.139926 | 0.443454 | 0.315536 | 0.752355 | 0.961069 |
| kdn | 4785.006 | 0.209558 | 0.663073 | 0.316041 | 0.751971 | 0.961069 |
| RpII215 | 3121.99 | 0.243619 | 0.771698 | 0.315692 | 0.752237 | 0.961069 |
| CG17721 | 794.7781 | -0.5213 | 1.654673 | -0.31505 | 0.752724 | 0.961097 |
| CG14314 | 1891.556 | -0.2986 | 0.948515 | -0.31481 | 0.752904 | 0.961097 |
| REPTOR | 6834.461 | 0.21398 | 0.679373 | 0.314966 | 0.752787 | 0.961097 |
| gammaSna | 2550.753 | 0.225925 | 0.717863 | 0.314719 | 0.752975 | 0.961097 |
| Ugt316A1 | 308.8 | -0.74256 | 2.359041 | -0.31477 | 0.752934 | 0.961097 |
| CG10011 | 2661.143 | -0.23112 | 0.735412 | -0.31427 | 0.753319 | 0.961383 |
| CG31729 | 3923.519 | -0.21988 | 0.704339 | -0.31218 | 0.754905 | 0.96142 |
| CG9356 | 552.1654 | -0.56326 | 1.799149 | -0.31307 | 0.754228 | 0.96142 |
| Abi | 1504.374 | -0.26256 | 0.838489 | -0.31313 | 0.754178 | 0.96142 |
| repo | 3197.007 | 0.247764 | 0.790783 | 0.313315 | 0.754041 | 0.96142 |
| hdc | 6139.781 | 0.179827 | 0.574767 | 0.31287 | 0.75438 | 0.96142 |
| Tbce | 1956.751 | -0.23566 | 0.752915 | -0.313 | 0.75428 | 0.96142 |
| GEFmeso | 2158.767 | -0.23927 | 0.763858 | -0.31323 | 0.754103 | 0.96142 |
| ewg | 2543.663 | 0.212396 | 0.677581 | 0.313463 | 0.753929 | 0.96142 |
| b | 299.3215 | -0.82516 | 2.639934 | -0.31257 | 0.754607 | 0.96142 |
| ps | 89600.41 | 0.120485 | 0.385784 | 0.312312 | 0.754803 | 0.96142 |

| | | | | | | |
|---|---|---|---|---|---|---|
| ATPsynB | 25829.25 | -0.13148 | 0.420877 | -0.3124 | 0.75474 | 0.96142 |
| aay | 6764.415 | 0.183017 | 0.585607 | 0.312525 | 0.754642 | 0.96142 |
| Usp47 | 2483.328 | -0.28893 | 0.920473 | -0.3139 | 0.753601 | 0.96142 |
| Pi4KIIalpha | 715.0948 | -0.46475 | 1.490452 | -0.31182 | 0.75518 | 0.961618 |
| INPP5E | 496.2699 | 0.489185 | 1.569993 | 0.311584 | 0.755357 | 0.96169 |
| Vps16B | 208.8839 | 0.753673 | 2.423896 | 0.310935 | 0.75585 | 0.962166 |
| Taz | 973.9862 | 0.49755 | 1.601336 | 0.310709 | 0.756022 | 0.96221 |
| RpS7 | 14906.94 | 0.136449 | 0.439569 | 0.310416 | 0.756245 | 0.96221 |
| z | 821.7462 | -0.34613 | 1.114806 | -0.31048 | 0.756193 | 0.96221 |
| DppIII | 1206.967 | 0.367449 | 1.184387 | 0.310244 | 0.756375 | 0.962213 |
| Syx17 | 781.7536 | -0.43204 | 1.393948 | -0.30994 | 0.756606 | 0.962213 |
| Cyp4g15 | 5059.36 | 0.190441 | 0.614219 | 0.310054 | 0.75652 | 0.962213 |
| EbpIII | 4792.494 | 0.354238 | 1.143763 | 0.309713 | 0.756779 | 0.96228 |
| Or85f | 240.5136 | -0.8315 | 2.689919 | -0.30912 | 0.757231 | 0.962702 |
| Dg | 3805.672 | -0.22949 | 0.743425 | -0.3087 | 0.757553 | 0.962958 |
| CG6472 | 229.85 | -0.90199 | 2.925751 | -0.30829 | 0.757858 | 0.963067 |
| CG30015 | 2725.062 | -0.22298 | 0.723335 | -0.30827 | 0.757878 | 0.963067 |
| stck | 3742.464 | -0.25866 | 0.839545 | -0.3081 | 0.758009 | 0.96308 |
| RpL34a | 3399.117 | 0.207959 | 0.675478 | 0.30787 | 0.758182 | 0.963147 |
| Idh | 8953.714 | -0.192 | 0.623951 | -0.30771 | 0.758304 | 0.963151 |
| CG8861 | 17099.41 | 0.17483 | 0.568916 | 0.307304 | 0.758612 | 0.963389 |
| nAChRalph | 2550.82 | -0.21858 | 0.713731 | -0.30625 | 0.759416 | 0.964105 |
| EndoB | 5941.038 | -0.16097 | 0.525568 | -0.30627 | 0.7594 | 0.964105 |
| CLIP-190 | 11118.04 | -0.1806 | 0.590677 | -0.30575 | 0.759799 | 0.964317 |
| CG2247 | 1549.45 | 0.291314 | 0.953472 | 0.30553 | 0.759963 | 0.964317 |
| Lon | 925.7232 | 0.31592 | 1.034454 | 0.305398 | 0.760063 | 0.964317 |
| Dronc | 300.5985 | -0.83993 | 2.7486 | -0.30558 | 0.759921 | 0.964317 |
| lncRNA:CR | 129.583 | 0.996061 | 3.263553 | 0.305208 | 0.760208 | 0.964348 |
| PIG-K | 3135.546 | -0.19347 | 0.651984 | -0.29674 | 0.766665 | 0.964358 |
| CG3527 | 481.3181 | -0.53973 | 1.806435 | -0.29878 | 0.765108 | 0.964358 |
| CG12147 | 109.7869 | 1.178901 | 3.951336 | 0.298355 | 0.765432 | 0.964358 |
| eIF4A | 26683.02 | 0.120809 | 0.407859 | 0.296202 | 0.767076 | 0.964358 |
| CG11865 | 108.101 | 1.166099 | 3.95134 | 0.295115 | 0.767906 | 0.964358 |
| CG10466 | 254.5473 | -0.66841 | 2.248186 | -0.29731 | 0.76623 | 0.964358 |
| CG9636 | 5231.857 | 0.180905 | 0.596322 | 0.303368 | 0.761609 | 0.964358 |
| Ras85D | 6313.863 | -0.16601 | 0.555973 | -0.2986 | 0.765247 | 0.964358 |
| CG8483 | 114.4488 | 1.178949 | 3.951319 | 0.298369 | 0.765422 | 0.964358 |
| EMC2B | 930.78 | -0.35261 | 1.166433 | -0.30229 | 0.762428 | 0.964358 |
| wrd | 3626.306 | 0.19341 | 0.646759 | 0.299044 | 0.764906 | 0.964358 |
| CG4733 | 118.2633 | -1.01419 | 3.406451 | -0.29773 | 0.765912 | 0.964358 |
| CG4783 | 76.45458 | 1.178336 | 3.951523 | 0.298198 | 0.765552 | 0.964358 |
| dsd | 3563.431 | 0.214398 | 0.703584 | 0.304723 | 0.760577 | 0.964358 |
| cindr | 8838.798 | 0.16759 | 0.566884 | 0.295633 | 0.76751 | 0.964358 |
| schlank | 3213.494 | 0.228084 | 0.771837 | 0.295508 | 0.767606 | 0.964358 |
| CG14442 | 3691.593 | 0.20568 | 0.696801 | 0.295178 | 0.767858 | 0.964358 |
| CG2124 | 83.97511 | -1.10234 | 3.709248 | -0.29719 | 0.766324 | 0.964358 |
| IntS8 | 77.26701 | -1.17605 | 3.933245 | -0.299 | 0.764938 | 0.964358 |
| CG17385 | 3597.877 | -0.18537 | 0.627409 | -0.29546 | 0.767646 | 0.964358 |
| CG18371 | 103.4934 | 1.178822 | 3.951362 | 0.298333 | 0.765449 | 0.964358 |
| shot | 16806.4 | 0.144762 | 0.483803 | 0.299216 | 0.764775 | 0.964358 |

| | | | | | | |
|---|---|---|---|---|---|---|
| Qsox1 | 2595.335 | 0.226994 | 0.758577 | 0.299237 | 0.764759 | 0.964358 |
| CG13197 | 763.2916 | 0.343564 | 1.155789 | 0.297255 | 0.766272 | 0.964358 |
| CG18858 | 117.2694 | -0.86595 | 2.904708 | -0.29812 | 0.765611 | 0.964358 |
| gus | 8940.389 | 0.160674 | 0.543981 | 0.295368 | 0.767713 | 0.964358 |
| zfh1 | 1603.143 | 0.2889 | 0.9699 | 0.297866 | 0.765805 | 0.964358 |
| ec | 108.0494 | -0.88961 | 2.962561 | -0.30028 | 0.763961 | 0.964358 |
| Glyat | 69.69488 | 1.178163 | 3.951583 | 0.29815 | 0.765589 | 0.964358 |
| snoRNA:Ps | 106.9898 | 1.178867 | 3.951347 | 0.298346 | 0.765439 | 0.964358 |
| vari | 924.7074 | 0.343057 | 1.147623 | 0.298929 | 0.764995 | 0.964358 |
| UQCR-C2 | 7253.932 | 0.159729 | 0.533814 | 0.299223 | 0.76477 | 0.964358 |
| pzg | 1054.478 | 0.389385 | 1.319335 | 0.295137 | 0.767889 | 0.964358 |
| CG42789 | 94.16967 | 1.178687 | 3.951407 | 0.298296 | 0.765478 | 0.964358 |
| CG43229 | 75.98839 | 1.178328 | 3.951527 | 0.298196 | 0.765554 | 0.964358 |
| CG13901 | 356.9434 | 0.57143 | 1.887606 | 0.302727 | 0.762098 | 0.964358 |
| asRNA:CR4 | 834.7282 | -0.3789 | 1.267745 | -0.29888 | 0.765034 | 0.964358 |
| sick | 6517.894 | -0.15116 | 0.507617 | -0.29778 | 0.765874 | 0.964358 |
| lncRNA:CR | 141.7207 | 1.179177 | 3.951239 | 0.298432 | 0.765373 | 0.964358 |
| lncRNA:CR | 77.5994 | 1.007172 | 3.401693 | 0.296079 | 0.767169 | 0.964358 |
| lncRNA:CR | 94.16967 | 1.178687 | 3.951407 | 0.298296 | 0.765478 | 0.964358 |
| lncRNA:CR | 113.7495 | 1.178942 | 3.951321 | 0.298366 | 0.765424 | 0.964358 |
| CG44774 | 2024.629 | 0.222438 | 0.730999 | 0.304293 | 0.760905 | 0.964358 |
| asRNA:CR4 | 76.22149 | 1.178334 | 3.951525 | 0.298197 | 0.765553 | 0.964358 |
| lncRNA:CR | 167.1278 | 1.179344 | 3.951189 | 0.298478 | 0.765338 | 0.964358 |
| lncRNA:CR | 86.94378 | 1.178563 | 3.951449 | 0.298261 | 0.765504 | 0.964358 |
| elgi | 1803.942 | -0.43317 | 1.423414 | -0.30432 | 0.760887 | 0.964358 |
| CR46405 | 82.99096 | -1.16443 | 3.834482 | -0.30367 | 0.761377 | 0.964358 |
| CG46466 | 2468.602 | 0.237698 | 0.804086 | 0.295612 | 0.767526 | 0.964358 |
| CG13920 | 2834.365 | 0.191831 | 0.649361 | 0.295415 | 0.767677 | 0.964358 |
| Jafrac2 | 1106.973 | -0.49437 | 1.630648 | -0.30318 | 0.761756 | 0.964358 |
| CG14965 | 440.0291 | 0.675135 | 2.242771 | 0.301027 | 0.763394 | 0.964358 |
| CG14977 | 1549.504 | 0.271591 | 0.918582 | 0.295663 | 0.767487 | 0.964358 |
| TfIIEbeta | 1076.508 | -0.29181 | 0.978042 | -0.29836 | 0.765425 | 0.964358 |
| nAChRbeta | 61259.03 | 0.126587 | 0.420745 | 0.300864 | 0.763518 | 0.964358 |
| RpS15Aa | 12451.1 | 0.134083 | 0.442783 | 0.302818 | 0.762029 | 0.964358 |
| CtsB | 6794.456 | -0.16462 | 0.549751 | -0.29944 | 0.764606 | 0.964358 |
| CG9915 | 576.9122 | -0.54415 | 1.83151 | -0.2971 | 0.766387 | 0.964358 |
| Edc3 | 2073.286 | -0.26016 | 0.868973 | -0.29939 | 0.764641 | 0.964358 |
| Ilp5 | 134.0286 | 1.179132 | 3.951258 | 0.298419 | 0.765383 | 0.964358 |
| BI-1 | 10893.4 | -0.15029 | 0.505092 | -0.29755 | 0.766047 | 0.964358 |
| frm | 263.0091 | 0.753231 | 2.491262 | 0.302349 | 0.762386 | 0.964358 |
| CG4835 | 130.5322 | 1.179107 | 3.951268 | 0.298412 | 0.765389 | 0.964358 |
| CG15362 | 83.44738 | 1.178494 | 3.951472 | 0.298242 | 0.765519 | 0.964358 |
| CG9641 | 1461.844 | -0.43453 | 1.475574 | -0.29448 | 0.76839 | 0.964815 |
| Mical | 3910.45 | 0.188472 | 0.640515 | 0.294251 | 0.768566 | 0.964849 |
| lncRNA:CR | 77.14827 | 0.947681 | 3.221969 | 0.294131 | 0.768658 | 0.964849 |
| CG5079 | 72.57443 | -0.98707 | 3.365811 | -0.29326 | 0.769321 | 0.964862 |
| CG33641 | 159.7024 | 0.638763 | 2.173263 | 0.293919 | 0.76882 | 0.964862 |
| CG34228 | 844.5113 | 0.385595 | 1.315696 | 0.293073 | 0.769466 | 0.964862 |
| CG43371 | 74.76865 | 1.159906 | 3.95153 | 0.293533 | 0.769114 | 0.964862 |
| CG43901 | 7414.931 | 0.172435 | 0.586945 | 0.293784 | 0.768923 | 0.964862 |

| | | | | | | |
|---|---|---|---|---|---|---|
| vih | 1859.813 | -0.22746 | 0.776265 | -0.29302 | 0.76951 | 0.964862 |
| RpS10b | 5918.257 | 0.153339 | 0.522689 | 0.293366 | 0.769243 | 0.964862 |
| cer | 68.67672 | 0.880328 | 3.012215 | 0.292253 | 0.770093 | 0.965293 |
| CG3589 | 181.5108 | -0.70322 | 2.404998 | -0.2924 | 0.769982 | 0.965293 |
| Flo2 | 3187.239 | -0.1817 | 0.623053 | -0.29162 | 0.770576 | 0.965746 |
| JMJD4 | 411.9368 | -0.86674 | 2.976717 | -0.29117 | 0.770918 | 0.966025 |
| CG32772 | 1813.651 | 0.300486 | 1.033407 | 0.290772 | 0.771225 | 0.966108 |
| eys | 4575.109 | 0.168871 | 0.580624 | 0.290844 | 0.771171 | 0.966108 |
| heix | 118.2783 | -0.78824 | 2.713187 | -0.29052 | 0.771417 | 0.966198 |
| lncRNA:CR | 2197.908 | 0.250682 | 0.86471 | 0.289903 | 0.77189 | 0.966639 |
| PRL-1 | 1466.03 | -0.2413 | 0.833581 | -0.28948 | 0.772217 | 0.966857 |
| CG10237 | 539.1725 | -0.59399 | 2.052753 | -0.28936 | 0.772305 | 0.966857 |
| lncRNA:CR | 90.30182 | -1.14263 | 3.951212 | -0.28919 | 0.772439 | 0.966874 |
| ND-AGGG | 3975.13 | 0.174116 | 0.602476 | 0.289001 | 0.772581 | 0.9669 |
| Eip78C | 4620.303 | 0.216448 | 0.750664 | 0.288342 | 0.773085 | 0.966997 |
| CG33169 | 209.7389 | 0.538484 | 1.872073 | 0.287641 | 0.773622 | 0.966997 |
| nonA-l | 327.9909 | -0.57646 | 1.998402 | -0.28846 | 0.772993 | 0.966997 |
| ETHR | 1083.586 | -0.52543 | 1.826281 | -0.28771 | 0.773571 | 0.966997 |
| tio | 138.1825 | -0.75539 | 2.61839 | -0.28849 | 0.772969 | 0.966997 |
| CG6845 | 1232.878 | -0.37007 | 1.286216 | -0.28772 | 0.77356 | 0.966997 |
| CG2021 | 569.1832 | -0.36522 | 1.269629 | -0.28766 | 0.77361 | 0.966997 |
| CG32109 | 2548.293 | 0.215038 | 0.747431 | 0.287703 | 0.773574 | 0.966997 |
| mt:ND6 | 3578.459 | 0.185618 | 0.646338 | 0.287184 | 0.773971 | 0.967283 |
| ex | 86.46807 | -1.05234 | 3.669513 | -0.28678 | 0.774281 | 0.967351 |
| drosha | 211.8506 | 0.563658 | 1.966446 | 0.286638 | 0.77439 | 0.967351 |
| CG12004 | 2877.734 | -0.18657 | 0.650356 | -0.28687 | 0.774209 | 0.967351 |
| Hydr2 | 3639.365 | -0.17546 | 0.612444 | -0.28648 | 0.774507 | 0.967351 |
| CG7379 | 1784.06 | -0.29119 | 1.017912 | -0.28607 | 0.774824 | 0.967537 |
| CR32745 | 586.6551 | 0.435796 | 1.523896 | 0.285975 | 0.774897 | 0.967537 |
| CG1965 | 1865.595 | -0.23561 | 0.824929 | -0.28561 | 0.775175 | 0.967733 |
| scrib | 5373.386 | 0.152962 | 0.535909 | 0.285426 | 0.775318 | 0.967761 |
| CG5281 | 1887.801 | -0.22821 | 0.802559 | -0.28435 | 0.776139 | 0.967777 |
| Ublcp1 | 363.321 | 0.527992 | 1.857038 | 0.28432 | 0.776165 | 0.967777 |
| Opbp | 470.2406 | -0.64166 | 2.256536 | -0.28435 | 0.776139 | 0.967777 |
| CaMKI | 18345.74 | 0.117169 | 0.412121 | 0.284308 | 0.776175 | 0.967777 |
| AP-1-2beta | 5501.927 | -0.15966 | 0.560904 | -0.28466 | 0.775907 | 0.967777 |
| Or71a | 298.252 | 0.741517 | 2.607813 | 0.284344 | 0.776147 | 0.967777 |
| CG13300 | 532.1499 | -0.40597 | 1.425592 | -0.28477 | 0.775817 | 0.967777 |
| Pex5 | 837.3126 | 0.447097 | 1.581967 | 0.282621 | 0.777468 | 0.967971 |
| CG6379 | 938.16 | 0.301523 | 1.064528 | 0.283246 | 0.776988 | 0.967971 |
| CG7810 | 723.0561 | -0.37659 | 1.335901 | -0.2819 | 0.778019 | 0.967971 |
| RpI1 | 289.5075 | 0.667695 | 2.367088 | 0.282074 | 0.777886 | 0.967971 |
| CG8617 | 974.8407 | -0.43948 | 1.553306 | -0.28293 | 0.777228 | 0.967971 |
| Syngr | 12504.44 | 0.145764 | 0.51702 | 0.281931 | 0.777997 | 0.967971 |
| Cog3 | 209.1248 | -0.77539 | 2.739263 | -0.28307 | 0.777127 | 0.967971 |
| ema | 3811.945 | 0.176535 | 0.623296 | 0.283229 | 0.777002 | 0.967971 |
| RpS16 | 6676.64 | 0.162517 | 0.575419 | 0.282433 | 0.777612 | 0.967971 |
| blw | 42623.09 | -0.11203 | 0.395504 | -0.28326 | 0.776981 | 0.967971 |
| Rpi | 736.3489 | -0.52262 | 1.846912 | -0.28297 | 0.7772 | 0.967971 |
| Dop1R2 | 1038.597 | 0.29313 | 1.036872 | 0.282706 | 0.777402 | 0.967971 |

| | | | | | | |
|---|---|---|---|---|---|---|
| mRpL21 | 1384.549 | -0.2659 | 0.941996 | -0.28227 | 0.777736 | 0.967971 |
| ABCA | 2337.644 | -0.20688 | 0.732585 | -0.2824 | 0.777637 | 0.967971 |
| l(2)k14505 | 1869.013 | 0.336173 | 1.193976 | 0.281558 | 0.778283 | 0.968149 |
| Cyp310a1 | 321.7803 | 0.637377 | 2.267589 | 0.281082 | 0.778648 | 0.968153 |
| Den1 | 1637.3 | 0.2353 | 0.836687 | 0.281228 | 0.778536 | 0.968153 |
| Traf6 | 176.5762 | 0.585652 | 2.082225 | 0.281262 | 0.778509 | 0.968153 |
| MED15 | 238.0603 | 0.499028 | 1.77646 | 0.280912 | 0.778778 | 0.968165 |
| CG7530 | 1809.258 | -0.3537 | 1.26112 | -0.28047 | 0.779121 | 0.968441 |
| Ssb-c31a | 3123.385 | -0.17813 | 0.637431 | -0.27945 | 0.779901 | 0.968477 |
| Klp31E | 311.9918 | -0.51806 | 1.860975 | -0.27838 | 0.780718 | 0.968477 |
| GstZ2 | 497.9078 | 0.586401 | 2.099512 | 0.279303 | 0.780012 | 0.968477 |
| RpL34b | 6989.207 | 0.150115 | 0.536067 | 0.28003 | 0.779455 | 0.968477 |
| CG31220 | 566.3727 | 0.371162 | 1.330056 | 0.279057 | 0.780201 | 0.968477 |
| l(1)G0289 | 6974.961 | 0.1429 | 0.513123 | 0.27849 | 0.780636 | 0.968477 |
| CG17715 | 397.204 | 0.465918 | 1.665506 | 0.279746 | 0.779673 | 0.968477 |
| CG34172 | 181.5799 | 0.839818 | 3.000424 | 0.2799 | 0.779554 | 0.968477 |
| spoon | 2660.749 | -0.21196 | 0.756925 | -0.28002 | 0.77946 | 0.968477 |
| mRpS35 | 1552.402 | 0.246685 | 0.884185 | 0.278998 | 0.780247 | 0.968477 |
| CG11583 | 356.1688 | -0.52687 | 1.88579 | -0.27939 | 0.779945 | 0.968477 |
| CG6340 | 3031.469 | -0.18114 | 0.649997 | -0.27869 | 0.780486 | 0.968477 |
| Nelf-E | 1032.844 | -0.28048 | 1.007319 | -0.27844 | 0.780674 | 0.968477 |
| Ho | 1640.405 | 0.408342 | 1.469619 | 0.277856 | 0.781123 | 0.968829 |
| Lar | 6097.925 | -0.14253 | 0.514272 | -0.27715 | 0.781662 | 0.96893 |
| Arf51F | 5699.026 | 0.187182 | 0.675453 | 0.277121 | 0.781688 | 0.96893 |
| Pih1D1 | 2766.607 | -0.21868 | 0.788242 | -0.27742 | 0.781456 | 0.96893 |
| PCID2 | 262.2778 | 0.755413 | 2.721885 | 0.277533 | 0.781371 | 0.96893 |
| CG43610 | 75.32379 | 1.094215 | 3.951334 | 0.276923 | 0.781839 | 0.968969 |
| Prp18 | 324.8669 | 0.524095 | 1.894871 | 0.276586 | 0.782098 | 0.96914 |
| CG6066 | 234.4436 | 0.540648 | 1.95843 | 0.276062 | 0.782501 | 0.969359 |
| tst | 178.0906 | 0.88203 | 3.197073 | 0.275887 | 0.782635 | 0.969359 |
| rhea | 6722.501 | 0.185335 | 0.671784 | 0.275884 | 0.782637 | 0.969359 |
| CtsF | 7365.542 | -0.14013 | 0.50941 | -0.27509 | 0.783248 | 0.969507 |
| CG5446 | 2938.368 | -0.18365 | 0.66865 | -0.27466 | 0.783579 | 0.969507 |
| TM9SF4 | 2715.049 | 0.21526 | 0.782389 | 0.275132 | 0.783215 | 0.969507 |
| CG15908 | 1812.333 | 0.261153 | 0.950934 | 0.274627 | 0.783602 | 0.969507 |
| boca | 1668.706 | 0.22144 | 0.804646 | 0.275202 | 0.783161 | 0.969507 |
| mnb | 3371.902 | 0.17293 | 0.628898 | 0.274973 | 0.783337 | 0.969507 |
| lsn | 524.7962 | 0.442681 | 1.611259 | 0.274743 | 0.783514 | 0.969507 |
| CG32026 | 754.6312 | -0.35573 | 1.296749 | -0.27432 | 0.783838 | 0.969649 |
| frj | 4064.643 | -0.17314 | 0.632325 | -0.27381 | 0.784231 | 0.969696 |
| CG6357 | 8103.11 | -0.13357 | 0.48782 | -0.2738 | 0.784238 | 0.969696 |
| ana | 12802.76 | -0.13834 | 0.50497 | -0.27395 | 0.784124 | 0.969696 |
| Rack1 | 13510.95 | -0.12242 | 0.447854 | -0.27335 | 0.784587 | 0.969816 |
| CG10286 | 524.0791 | -0.34992 | 1.280814 | -0.2732 | 0.784698 | 0.969816 |
| CG5532 | 3131.773 | 0.189756 | 0.694047 | 0.273405 | 0.784542 | 0.969816 |
| hwt | 6486.63 | -0.14057 | 0.51484 | -0.27304 | 0.784819 | 0.969816 |
| CG32425 | 2526.03 | -0.20728 | 0.759839 | -0.2728 | 0.785009 | 0.969902 |
| Rpn10 | 3496.023 | 0.253807 | 0.932919 | 0.272056 | 0.785579 | 0.969996 |
| CG9253 | 1004.11 | 0.354157 | 1.304577 | 0.271472 | 0.786028 | 0.969996 |
| Kcmf1 | 2929.473 | -0.18009 | 0.660803 | -0.27253 | 0.785218 | 0.969996 |

| | | | | | | |
|---|---|---|---|---|---|---|
| DCTN6-p2 | 353.8972 | -0.44025 | 1.621977 | -0.27143 | 0.78606 | 0.969996 |
| lncRNA:CR | 384.585 | 0.465964 | 1.715688 | 0.27159 | 0.785937 | 0.969996 |
| Patr-1 | 1042.778 | 0.260686 | 0.96094 | 0.271283 | 0.786174 | 0.969996 |
| Hsp83 | 20797.45 | -0.11068 | 0.407877 | -0.27135 | 0.786122 | 0.969996 |
| CG11593 | 1497.826 | -0.24995 | 0.918469 | -0.27214 | 0.785515 | 0.969996 |
| Abp1 | 1332.477 | 0.259874 | 0.958501 | 0.271126 | 0.786294 | 0.969996 |
| bc10 | 1819.802 | -0.22235 | 0.816327 | -0.27238 | 0.785329 | 0.969996 |
| galectin | 6088.387 | -0.15731 | 0.586277 | -0.26832 | 0.788454 | 0.970229 |
| Alg11 | 3615.587 | -0.21772 | 0.809517 | -0.26895 | 0.787972 | 0.970229 |
| CG7369 | 1269.241 | 0.281096 | 1.046834 | 0.26852 | 0.788299 | 0.970229 |
| Snr1 | 1641.372 | -0.37212 | 1.390625 | -0.26759 | 0.789014 | 0.970229 |
| CG9536 | 1919.873 | 0.244909 | 0.920821 | 0.265968 | 0.790263 | 0.970229 |
| CG33303 | 1862.198 | 0.228557 | 0.858308 | 0.266288 | 0.790017 | 0.970229 |
| Trax | 545.1136 | -0.4586 | 1.708805 | -0.26837 | 0.788411 | 0.970229 |
| Srp14 | 580.9267 | -0.35422 | 1.314031 | -0.26957 | 0.787495 | 0.970229 |
| Ythdf | 1911.054 | -0.19882 | 0.747515 | -0.26597 | 0.790263 | 0.970229 |
| DIP-gamm | 1317.309 | 0.260176 | 0.970464 | 0.268094 | 0.788627 | 0.970229 |
| CG8079 | 690.3128 | -0.33912 | 1.26272 | -0.26856 | 0.788268 | 0.970229 |
| Zip102B | 3240.102 | 0.167206 | 0.628938 | 0.265854 | 0.790351 | 0.970229 |
| tacc | 7731.377 | -0.15304 | 0.571716 | -0.26768 | 0.788944 | 0.970229 |
| CG10863 | 2957.909 | -0.19088 | 0.711075 | -0.26844 | 0.788358 | 0.970229 |
| Mtl | 7408.846 | -0.18548 | 0.69688 | -0.26616 | 0.790118 | 0.970229 |
| CG40160 | 717.1885 | 0.555133 | 2.076395 | 0.267354 | 0.789197 | 0.970229 |
| lncRNA:CR | 114.6449 | 0.839315 | 3.138231 | 0.267448 | 0.789124 | 0.970229 |
| mub | 18355.5 | -0.11329 | 0.425804 | -0.26605 | 0.790198 | 0.970229 |
| CG42784 | 2800.913 | 0.17596 | 0.658391 | 0.267257 | 0.789271 | 0.970229 |
| Nup214 | 114.3346 | 0.8187 | 3.028516 | 0.27033 | 0.786906 | 0.970229 |
| Golgin245 | 2187.722 | 0.231393 | 0.860056 | 0.269044 | 0.787896 | 0.970229 |
| ND-19 | 8078.265 | 0.132339 | 0.489094 | 0.27058 | 0.786714 | 0.970229 |
| Pof | 507.2788 | -0.64647 | 2.41129 | -0.2681 | 0.78862 | 0.970229 |
| lncRNA:CR | 1185.354 | 0.251828 | 0.93453 | 0.26947 | 0.787568 | 0.970229 |
| lncRNA:CR | 237.157 | 0.514934 | 1.930202 | 0.266777 | 0.789641 | 0.970229 |
| RpL31 | 13723.7 | 0.121813 | 0.45759 | 0.266205 | 0.790081 | 0.970229 |
| CG1271 | 80.99655 | 1.027366 | 3.858846 | 0.266237 | 0.790057 | 0.970229 |
| CG15814 | 1891.334 | -0.23248 | 0.860259 | -0.27025 | 0.786971 | 0.970229 |
| scyl | 10867.27 | -0.12028 | 0.449716 | -0.26746 | 0.789118 | 0.970229 |
| velo | 1043.606 | 0.288718 | 1.079866 | 0.267365 | 0.789188 | 0.970229 |
| GABPI | 4281.786 | -0.17661 | 0.656045 | -0.2692 | 0.787777 | 0.970229 |
| CG15385 | 2148.051 | 0.285021 | 1.063522 | 0.267997 | 0.788702 | 0.970229 |
| CG5110 | 2119.329 | -0.21334 | 0.804581 | -0.26516 | 0.790888 | 0.970258 |
| CG7029 | 16118.63 | -0.1126 | 0.425608 | -0.26457 | 0.791341 | 0.970258 |
| CG5001 | 682.6446 | 0.428102 | 1.617171 | 0.264723 | 0.791223 | 0.970258 |
| CG8155 | 1372.327 | 0.396747 | 1.498494 | 0.264764 | 0.791191 | 0.970258 |
| jeb | 17864.47 | 0.123685 | 0.466973 | 0.264865 | 0.791113 | 0.970258 |
| Bet5 | 1456.41 | -0.21923 | 0.82807 | -0.26475 | 0.791204 | 0.970258 |
| sov | 4494.083 | -0.1528 | 0.577236 | -0.26472 | 0.791227 | 0.970258 |
| CG32407 | 3631.23 | -0.19712 | 0.743003 | -0.2653 | 0.790782 | 0.970258 |
| CG5254 | 3380.187 | 0.19398 | 0.734092 | 0.264246 | 0.791591 | 0.970346 |
| fne | 35736.66 | 0.101925 | 0.386302 | 0.263848 | 0.791897 | 0.970346 |
| asRNA:CR4 | 136.8358 | 0.580192 | 2.197538 | 0.264019 | 0.791765 | 0.970346 |

| | | | | | | |
|---|---|---|---|---|---|---|
| l(2)k05819 | 1214.877 | 0.402136 | 1.523488 | 0.263957 | 0.791813 | 0.970346 |
| CG4004 | 1536.833 | 0.276768 | 1.049858 | 0.263625 | 0.792069 | 0.970409 |
| CG15877 | 83.70146 | -0.8952 | 3.403857 | -0.263 | 0.792553 | 0.970706 |
| hob | 284.6451 | -0.55652 | 2.115921 | -0.26301 | 0.79254 | 0.970706 |
| Atg13 | 638.3634 | 0.328177 | 1.250963 | 0.262339 | 0.79306 | 0.971178 |
| CG14817 | 2452.494 | -0.18171 | 0.701571 | -0.259 | 0.795632 | 0.971659 |
| pcx | 1052.766 | -0.48469 | 1.865789 | -0.25978 | 0.795036 | 0.971659 |
| CG5381 | 1168.274 | -0.3126 | 1.201797 | -0.26011 | 0.794778 | 0.971659 |
| CG9338 | 306.706 | -0.61273 | 2.349485 | -0.26079 | 0.794252 | 0.971659 |
| Taf12 | 2557.454 | -0.18565 | 0.716616 | -0.25907 | 0.795581 | 0.971659 |
| Smurf | 1699.794 | 0.206281 | 0.790525 | 0.260941 | 0.794138 | 0.971659 |
| APC10 | 257.929 | -0.63857 | 2.462093 | -0.25936 | 0.795358 | 0.971659 |
| ACC | 2693.867 | 0.206452 | 0.792017 | 0.260666 | 0.79435 | 0.971659 |
| CG14982 | 3949.123 | -0.18588 | 0.717025 | -0.25924 | 0.79545 | 0.971659 |
| l(2)05714 | 162.8606 | -0.615 | 2.366974 | -0.25983 | 0.794998 | 0.971659 |
| Dad1 | 1434.915 | 0.220258 | 0.842919 | 0.261304 | 0.793858 | 0.971659 |
| betaGlu | 673.6366 | -0.44381 | 1.706033 | -0.26014 | 0.794756 | 0.971659 |
| msn | 3039.644 | -0.2128 | 0.820972 | -0.2592 | 0.79548 | 0.971659 |
| Chc | 6486.468 | 0.149323 | 0.572962 | 0.260616 | 0.794388 | 0.971659 |
| kto | 114.9294 | -0.6448 | 2.481886 | -0.2598 | 0.795017 | 0.971659 |
| MED10 | 1412.822 | 0.222156 | 0.856342 | 0.259424 | 0.795308 | 0.971659 |
| Nmt | 242.7095 | 0.607357 | 2.322999 | 0.261454 | 0.793742 | 0.971659 |
| CG8620 | 678.7403 | 0.296023 | 1.135595 | 0.260677 | 0.794342 | 0.971659 |
| RtcB | 671.8246 | -0.48747 | 1.885369 | -0.25855 | 0.795981 | 0.971937 |
| Csk | 3628.757 | 0.186167 | 0.72146 | 0.258042 | 0.796375 | 0.97227 |
| wit | 386.1375 | -0.41151 | 1.596829 | -0.2577 | 0.796636 | 0.972441 |
| CG13369 | 660.2418 | 0.292442 | 1.149 | 0.254519 | 0.799095 | 0.972589 |
| usp | 913.5682 | 0.292698 | 1.194073 | 0.245126 | 0.806359 | 0.972589 |
| Spt6 | 1104.262 | 0.352956 | 1.380436 | 0.255685 | 0.798194 | 0.972589 |
| skd | 9983.975 | -0.11855 | 0.478594 | -0.24771 | 0.804358 | 0.972589 |
| Hcs | 685.0453 | -0.31597 | 1.281882 | -0.24649 | 0.805306 | 0.972589 |
| CG2519 | 5471.357 | -0.15807 | 0.617253 | -0.25608 | 0.797886 | 0.972589 |
| Nmdar1 | 2254.898 | -0.29546 | 1.213335 | -0.24351 | 0.807608 | 0.972589 |
| CG9171 | 1223.821 | -0.30713 | 1.246615 | -0.24637 | 0.805395 | 0.972589 |
| Aatf | 229.8468 | 0.517666 | 2.084126 | 0.248385 | 0.803837 | 0.972589 |
| CG8353 | 276.6114 | -0.44217 | 1.742928 | -0.25369 | 0.799732 | 0.972589 |
| CG9555 | 127.6052 | -0.89724 | 3.68007 | -0.24381 | 0.807378 | 0.972589 |
| SamDC | 1220.198 | 0.229776 | 0.928099 | 0.247577 | 0.804462 | 0.972589 |
| Bug22 | 1332.089 | 0.35535 | 1.44699 | 0.245579 | 0.806008 | 0.972589 |
| Nup107 | 350.0994 | 0.426225 | 1.687464 | 0.252583 | 0.800591 | 0.972589 |
| Idgf2 | 1574.545 | -0.37206 | 1.521938 | -0.24446 | 0.806871 | 0.972589 |
| CG31815 | 1661.412 | 0.250622 | 0.979344 | 0.255908 | 0.798022 | 0.972589 |
| CG6115 | 3112.239 | -0.16291 | 0.666409 | -0.24446 | 0.806877 | 0.972589 |
| RPA2 | 268.049 | 0.814149 | 3.350781 | 0.242973 | 0.808026 | 0.972589 |
| mRpL47 | 1384.534 | -0.27867 | 1.104985 | -0.25219 | 0.800894 | 0.972589 |
| CG5038 | 364.7627 | 0.401554 | 1.633746 | 0.245787 | 0.805847 | 0.972589 |
| Arpc3A | 3151.465 | 0.184806 | 0.721369 | 0.256187 | 0.797806 | 0.972589 |
| CG17565 | 1412.078 | -0.37536 | 1.49084 | -0.25178 | 0.801211 | 0.972589 |
| CG7183 | 689.3179 | -0.30638 | 1.192266 | -0.25697 | 0.797201 | 0.972589 |
| CG5793 | 1559.592 | -0.21216 | 0.83956 | -0.2527 | 0.800501 | 0.972589 |

| | | | | | | |
|---|---|---|---|---|---|---|
| mRpL35 | 1688.791 | 0.206268 | 0.842822 | 0.244735 | 0.806662 | 0.972589 |
| Sec13 | 2493.556 | 0.177741 | 0.699741 | 0.25401 | 0.799488 | 0.972589 |
| CG5510 | 909.4875 | 0.436643 | 1.713306 | 0.254854 | 0.798836 | 0.972589 |
| CG31089 | 144.957 | 0.574658 | 2.302787 | 0.249549 | 0.802936 | 0.972589 |
| ro | 317.6605 | -0.56432 | 2.242338 | -0.25167 | 0.801298 | 0.972589 |
| CG31064 | 4049.212 | 0.284648 | 1.136079 | 0.250553 | 0.80216 | 0.972589 |
| ATPsyngar | 24022.12 | 0.099436 | 0.406999 | 0.244314 | 0.806987 | 0.972589 |
| Obp57c | 86.61895 | 0.953155 | 3.85075 | 0.247525 | 0.804502 | 0.972589 |
| Ypel | 842.2218 | 0.406929 | 1.584533 | 0.256813 | 0.797323 | 0.972589 |
| Tango11 | 3799.183 | -0.17419 | 0.685218 | -0.2542 | 0.799338 | 0.972589 |
| Got1 | 17764.09 | 0.163936 | 0.644516 | 0.254355 | 0.799221 | 0.972589 |
| BEAF-32 | 2407.776 | -0.18934 | 0.774752 | -0.24439 | 0.806926 | 0.972589 |
| CG18368 | 130.3121 | 0.794828 | 3.263161 | 0.243576 | 0.807559 | 0.972589 |
| cid | 223.5865 | -0.73687 | 2.920498 | -0.25231 | 0.8008 | 0.972589 |
| ox | 5170.305 | 0.148731 | 0.58327 | 0.254995 | 0.798727 | 0.972589 |
| fdl | 871.065 | -0.41813 | 1.719495 | -0.24317 | 0.807875 | 0.972589 |
| Cam | 43338.03 | 0.104501 | 0.416684 | 0.250792 | 0.801975 | 0.972589 |
| EndoG | 271.0643 | 0.640159 | 2.630206 | 0.243388 | 0.807705 | 0.972589 |
| ERp60 | 8360.769 | -0.14149 | 0.566084 | -0.24995 | 0.802624 | 0.972589 |
| CG13220 | 6493.873 | -0.12676 | 0.520801 | -0.2434 | 0.807694 | 0.972589 |
| CG11825 | 150.4858 | -0.9307 | 3.752033 | -0.24805 | 0.804094 | 0.972589 |
| CG12910 | 166.7843 | 0.735604 | 2.93123 | 0.250954 | 0.80185 | 0.972589 |
| CAP | 10783.76 | 0.124043 | 0.488456 | 0.25395 | 0.799534 | 0.972589 |
| Map60 | 1230.013 | 0.243942 | 0.97514 | 0.250161 | 0.802463 | 0.972589 |
| Prp38 | 718.5229 | -0.36101 | 1.457565 | -0.24768 | 0.804382 | 0.972589 |
| CG10214 | 1356.779 | 0.261358 | 1.038066 | 0.251774 | 0.801216 | 0.972589 |
| mRpL38 | 2319.581 | -0.18518 | 0.73732 | -0.25115 | 0.801697 | 0.972589 |
| CG8939 | 175.8788 | 0.662443 | 2.615755 | 0.253251 | 0.800074 | 0.972589 |
| jim | 40610.95 | -0.09602 | 0.385783 | -0.24891 | 0.803432 | 0.972589 |
| mle | 725.7674 | 0.305876 | 1.248376 | 0.245019 | 0.806442 | 0.972589 |
| nonA | 6831.391 | -0.13133 | 0.537019 | -0.24455 | 0.806804 | 0.972589 |
| His3.3A | 3652.738 | 0.162302 | 0.636385 | 0.255038 | 0.798694 | 0.972589 |
| Tab2 | 455.4144 | -0.41229 | 1.696047 | -0.24309 | 0.807935 | 0.972589 |
| mRpS23 | 316.1498 | -0.55304 | 2.176625 | -0.25408 | 0.799431 | 0.972589 |
| Larp7 | 135.0681 | -0.74879 | 2.96772 | -0.25231 | 0.8008 | 0.972589 |
| NaPi-III | 6623.643 | -0.13322 | 0.545288 | -0.24432 | 0.806984 | 0.972589 |
| CSN8 | 642.1409 | -0.31681 | 1.280409 | -0.24743 | 0.804575 | 0.972589 |
| RNASEK | 23525.37 | 0.108496 | 0.426179 | 0.254579 | 0.799048 | 0.972589 |
| Vha44 | 7836.851 | -0.13406 | 0.544807 | -0.24607 | 0.80563 | 0.972589 |
| CG3253 | 1596.518 | 0.22326 | 0.883317 | 0.252752 | 0.80046 | 0.972589 |
| Usp15-31 | 3556.427 | -0.2014 | 0.827552 | -0.24337 | 0.807721 | 0.972589 |
| lncRNA:CR | 763.563 | -0.42917 | 1.704889 | -0.25173 | 0.801252 | 0.972589 |
| ens | 8895.606 | 0.120626 | 0.47735 | 0.252699 | 0.800501 | 0.972589 |
| lov | 853.0752 | -0.36948 | 1.503401 | -0.24576 | 0.805867 | 0.972589 |
| CG42259 | 195.4076 | 0.708437 | 2.836186 | 0.249785 | 0.802754 | 0.972589 |
| pre-rRNA:( | 282.1062 | 0.44418 | 1.814549 | 0.244788 | 0.80662 | 0.972589 |
| lncRNA:CR | 1336.41 | 0.21518 | 0.865025 | 0.248756 | 0.803549 | 0.972589 |
| CG13807 | 652.4093 | 0.287065 | 1.150376 | 0.24954 | 0.802943 | 0.972589 |
| CG10359 | 353.7353 | -0.491 | 1.953711 | -0.25132 | 0.801569 | 0.972589 |
| Teh3 | 901.149 | 0.337064 | 1.333674 | 0.252733 | 0.800474 | 0.972589 |

| | | | | | | |
|---|---|---|---|---|---|---|
| lncRNA:ro) | 47774.55 | 0.102104 | 0.402839 | 0.253462 | 0.799911 | 0.972589 |
| Kmn1 | 206.6763 | -0.90663 | 3.62529 | -0.25009 | 0.802521 | 0.972589 |
| regucalcin | 1515.114 | -0.24579 | 0.996834 | -0.24657 | 0.805243 | 0.972589 |
| be | 1076.641 | -0.33029 | 1.353104 | -0.2441 | 0.807153 | 0.972589 |
| CG14407 | 2484.995 | 0.186878 | 0.766196 | 0.243903 | 0.807306 | 0.972589 |
| CG8611 | 222.0132 | 0.49441 | 1.932501 | 0.255839 | 0.798075 | 0.972589 |
| CG14234 | 823.874 | -0.31796 | 1.268923 | -0.25058 | 0.802142 | 0.972589 |
| Kap-alpha1 | 4300.977 | -0.16132 | 0.648508 | -0.24876 | 0.803545 | 0.972589 |
| wnd | 1805.383 | -0.19709 | 0.807144 | -0.24419 | 0.807086 | 0.972589 |
| Crtc | 1916.735 | -0.19309 | 0.777288 | -0.24841 | 0.803814 | 0.972589 |
| QIL1 | 11098.11 | -0.11672 | 0.460966 | -0.2532 | 0.800113 | 0.972589 |
| PDCD-5 | 475.9461 | 0.336007 | 1.319147 | 0.254716 | 0.798943 | 0.972589 |
| IntS9 | 1463.311 | 0.270953 | 1.109665 | 0.244176 | 0.807095 | 0.972589 |
| CrebA | 10028.84 | -0.1297 | 0.52584 | -0.24665 | 0.805182 | 0.972589 |
| Sox21b | 67.13454 | -0.89971 | 3.605084 | -0.24957 | 0.802921 | 0.972589 |
| Blos4 | 877.2263 | 0.288188 | 1.145919 | 0.251491 | 0.801435 | 0.972589 |
| vsg | 11880.99 | -0.11967 | 0.484825 | -0.24683 | 0.805036 | 0.972589 |
| Pdp1 | 34037.63 | 0.1033 | 0.420661 | 0.245566 | 0.806018 | 0.972589 |
| tbc | 2835.245 | 0.206313 | 0.840744 | 0.245393 | 0.806152 | 0.972589 |
| kirre | 1316.283 | 0.233383 | 0.961751 | 0.242665 | 0.808265 | 0.972664 |
| CG31550 | 3135.875 | 0.162397 | 0.669462 | 0.242579 | 0.808332 | 0.972664 |
| Fkbp12 | 14725.32 | -0.109 | 0.449698 | -0.24239 | 0.808479 | 0.972696 |
| Syx7 | 2346.615 | -0.17042 | 0.703589 | -0.24221 | 0.808619 | 0.972718 |
| CG31886 | 611.9014 | -0.43438 | 1.796966 | -0.24173 | 0.808991 | 0.972718 |
| Calr | 6707.141 | 0.13526 | 0.559425 | 0.241784 | 0.808947 | 0.972718 |
| Kap-alpha3 | 6817.558 | 0.122208 | 0.506311 | 0.24137 | 0.809269 | 0.972718 |
| CG9399 | 3313.291 | 0.160136 | 0.663116 | 0.24149 | 0.809175 | 0.972718 |
| CG10177 | 482.9596 | -0.49502 | 2.046644 | -0.24187 | 0.808883 | 0.972718 |
| CG13623 | 1200.944 | -0.28329 | 1.174161 | -0.24127 | 0.809346 | 0.972718 |
| Nap1 | 103.1095 | -0.58474 | 2.429105 | -0.24072 | 0.80977 | 0.973082 |
| Las | 822.9315 | -0.35624 | 1.485023 | -0.23989 | 0.810417 | 0.973168 |
| CG4390 | 1616.776 | -0.20352 | 0.847646 | -0.2401 | 0.810252 | 0.973168 |
| NaPi-T | 340.6244 | 0.516145 | 2.148006 | 0.24029 | 0.810105 | 0.973168 |
| ND-20 | 12874.44 | 0.104795 | 0.436906 | 0.239858 | 0.81044 | 0.973168 |
| Upf3 | 248.4179 | -0.74576 | 3.10928 | -0.23985 | 0.810448 | 0.973168 |
| tai | 14190.32 | -0.13288 | 0.55629 | -0.23888 | 0.811201 | 0.973199 |
| beat-Ib | 5145.532 | 0.146421 | 0.612171 | 0.239183 | 0.810964 | 0.973199 |
| Pngl | 561.566 | 0.435056 | 1.818332 | 0.239261 | 0.810903 | 0.973199 |
| amx | 433.5802 | 0.491814 | 2.057806 | 0.238999 | 0.811106 | 0.973199 |
| Ptpmeg | 1425.822 | -0.2396 | 1.002487 | -0.23901 | 0.8111 | 0.973199 |
| Mi-2 | 1660.767 | 0.210509 | 0.879142 | 0.239449 | 0.810758 | 0.973199 |
| RpL13A | 21919.32 | -0.09926 | 0.415935 | -0.23865 | 0.811378 | 0.973265 |
| CG9773 | 1994.331 | -0.18219 | 0.764236 | -0.23839 | 0.811577 | 0.973359 |
| kat-60L1 | 2006.103 | 0.188409 | 0.792117 | 0.237855 | 0.811993 | 0.973713 |
| CG1239 | 495.5667 | 0.334147 | 1.422538 | 0.234895 | 0.81429 | 0.973801 |
| CG9121 | 481.1126 | 0.328693 | 1.383982 | 0.237498 | 0.81227 | 0.973801 |
| Ndf | 2858.888 | -0.1821 | 0.77025 | -0.23642 | 0.813107 | 0.973801 |
| Prosbeta3 | 2376.747 | 0.202094 | 0.862613 | 0.234281 | 0.814767 | 0.973801 |
| beat-Vb | 1534.311 | -0.26758 | 1.135261 | -0.2357 | 0.813667 | 0.973801 |
| SIDL | 513.7019 | -0.32417 | 1.371521 | -0.23636 | 0.813155 | 0.973801 |

| | | | | | | |
|---|---|---|---|---|---|---|
| Sulf1 | 2638.6 | 0.1932 | 0.820869 | 0.23536 | 0.813929 | 0.973801 |
| AP-2sigma | 8980.694 | 0.132687 | 0.560633 | 0.236674 | 0.81291 | 0.973801 |
| Cdc16 | 134.4315 | -0.85419 | 3.614601 | -0.23632 | 0.813187 | 0.973801 |
| Tsp97E | 12808.32 | 0.114082 | 0.486548 | 0.234472 | 0.814619 | 0.973801 |
| sktl | 1035.409 | -0.3686 | 1.572145 | -0.23446 | 0.814628 | 0.973801 |
| Fbl6 | 2858.908 | -0.17169 | 0.726104 | -0.23645 | 0.81308 | 0.973801 |
| CG8046 | 207.0155 | -0.7146 | 3.050022 | -0.23429 | 0.814756 | 0.973801 |
| alt | 2571.919 | -0.16552 | 0.706609 | -0.23425 | 0.814793 | 0.973801 |
| NijA | 1103.777 | 0.233849 | 0.994563 | 0.235127 | 0.81411 | 0.973801 |
| Caper | 5634.888 | 0.136732 | 0.577957 | 0.236579 | 0.812984 | 0.973801 |
| l(1)G0469 | 788.1886 | -0.25328 | 1.077815 | -0.235 | 0.814212 | 0.973801 |
| CG42494 | 71.16537 | 0.905109 | 3.850934 | 0.235036 | 0.814181 | 0.973801 |
| alc | 7543.99 | 0.115881 | 0.491483 | 0.235778 | 0.813605 | 0.973801 |
| lncRNA:CR | 207.9631 | 0.611042 | 2.611203 | 0.234008 | 0.814979 | 0.973801 |
| CG13012 | 1405.527 | -0.22764 | 0.972343 | -0.23411 | 0.814898 | 0.973801 |
| Arp10 | 262.1076 | -0.46204 | 1.952477 | -0.23664 | 0.812935 | 0.973801 |
| Prestin | 466.4171 | 0.448639 | 1.909251 | 0.234982 | 0.814223 | 0.973801 |
| CG7841 | 742.6256 | 0.262214 | 1.119895 | 0.234142 | 0.814875 | 0.973801 |
| CG7139 | 1363.01 | 0.215813 | 0.923862 | 0.233599 | 0.815297 | 0.973853 |
| Hex-A | 13358.92 | 0.105695 | 0.452391 | 0.233638 | 0.815266 | 0.973853 |
| lncRNA:CR | 3176.774 | -0.18934 | 0.810927 | -0.23348 | 0.815387 | 0.973853 |
| mei-P26 | 42998.52 | 0.096245 | 0.413072 | 0.232999 | 0.815762 | 0.97387 |
| CG17985 | 391.4256 | -0.33383 | 1.432406 | -0.23306 | 0.815716 | 0.97387 |
| CG6841 | 96.12012 | -0.63411 | 2.72154 | -0.233 | 0.815764 | 0.97387 |
| CG1077 | 227.1059 | 0.770362 | 3.319784 | 0.232052 | 0.816498 | 0.973876 |
| CG10053 | 100.5875 | 0.727711 | 3.134551 | 0.232158 | 0.816415 | 0.973876 |
| CG11453 | 154.0886 | 0.718587 | 3.092297 | 0.232379 | 0.816243 | 0.973876 |
| lolal | 4716.355 | -0.14971 | 0.643719 | -0.23257 | 0.816091 | 0.973876 |
| CG33710 | 146.2676 | -0.76667 | 3.294372 | -0.23272 | 0.815978 | 0.973876 |
| ebd1 | 456.0908 | -0.32282 | 1.390618 | -0.23214 | 0.81643 | 0.973876 |
| cib | 3031.343 | 0.156159 | 0.676895 | 0.230699 | 0.817549 | 0.974007 |
| Ada1-2 | 294.6762 | 0.417726 | 1.825113 | 0.228877 | 0.818965 | 0.974007 |
| put | 1494.856 | 0.281369 | 1.219358 | 0.230752 | 0.817508 | 0.974007 |
| alpha-Man | 609.0628 | 0.307965 | 1.346617 | 0.228695 | 0.819106 | 0.974007 |
| ClpX | 2225.634 | 0.165312 | 0.719909 | 0.22963 | 0.81838 | 0.974007 |
| Apc2 | 83.94102 | -0.72905 | 3.168358 | -0.2301 | 0.81801 | 0.974007 |
| CG18367 | 155.9686 | 0.752292 | 3.268016 | 0.230198 | 0.817938 | 0.974007 |
| p115 | 80.88604 | 0.758816 | 3.318434 | 0.228667 | 0.819128 | 0.974007 |
| JhI-26 | 96.23324 | -0.58156 | 2.544762 | -0.22853 | 0.819231 | 0.974007 |
| sowah | 1033.527 | 0.255006 | 1.112829 | 0.229151 | 0.818751 | 0.974007 |
| CG13124 | 5914.039 | 0.156005 | 0.677383 | 0.230305 | 0.817854 | 0.974007 |
| Ddr | 1490.246 | -0.19219 | 0.835153 | -0.23013 | 0.817992 | 0.974007 |
| Kif3C | 1273.927 | 0.207897 | 0.900369 | 0.230902 | 0.817391 | 0.974007 |
| Lcch3 | 2082.775 | -0.17226 | 0.754489 | -0.22832 | 0.819399 | 0.974007 |
| lncRNA:CR | 982.1557 | 0.231319 | 1.009604 | 0.229119 | 0.818777 | 0.974007 |
| CG3638 | 3434.385 | -0.14513 | 0.629154 | -0.23067 | 0.817573 | 0.974007 |
| SLO2 | 5126.565 | 0.132412 | 0.573103 | 0.231045 | 0.81728 | 0.974007 |
| nonC | 804.0155 | -0.31356 | 1.371085 | -0.22869 | 0.819109 | 0.974007 |
| CG8944 | 736.3808 | -0.25581 | 1.113514 | -0.22973 | 0.818298 | 0.974007 |
| Max | 931.185 | -0.22777 | 0.997057 | -0.22845 | 0.819299 | 0.974007 |

| | | | | | | |
|---|---|---|---|---|---|---|
| fz2 | 2878.298 | 0.165255 | 0.713824 | 0.231507 | 0.816921 | 0.974007 |
| CG15431 | 606.8031 | 0.292198 | 1.276186 | 0.228962 | 0.818899 | 0.974007 |
| Reph | 4368.86 | -0.14007 | 0.609062 | -0.22997 | 0.818114 | 0.974007 |
| CG6660 | 353.6129 | -0.59474 | 2.612178 | -0.22768 | 0.819896 | 0.974251 |
| CG15602 | 1911.896 | 0.321892 | 1.414373 | 0.227586 | 0.819968 | 0.974251 |
| CG7772 | 1327.046 | -0.19804 | 0.869509 | -0.22776 | 0.819831 | 0.974251 |
| Asph | 1850.193 | -0.19189 | 0.84513 | -0.22705 | 0.820383 | 0.974599 |
| Dp1 | 2493.55 | 0.198535 | 0.87509 | 0.226874 | 0.820522 | 0.97462 |
| CG15111 | 5783.176 | 0.135577 | 0.600188 | 0.22589 | 0.821287 | 0.975252 |
| Amph | 12115.05 | 0.114965 | 0.509021 | 0.225855 | 0.821314 | 0.975252 |
| lncRNA:CR | 153.1808 | 0.618784 | 2.743271 | 0.225564 | 0.82154 | 0.975252 |
| Nf-YA | 117.3714 | -0.63429 | 2.81109 | -0.22564 | 0.821483 | 0.975252 |
| CG11099 | 2420.736 | -0.16378 | 0.728103 | -0.22495 | 0.822022 | 0.97568 |
| Cyp4ac1 | 120.7966 | -0.72635 | 3.251607 | -0.22338 | 0.823239 | 0.97698 |
| CG18540 | 71.92435 | -0.85586 | 3.836663 | -0.22307 | 0.823477 | 0.977118 |
| rb | 1485.589 | -0.24358 | 1.09461 | -0.22252 | 0.823907 | 0.977246 |
| Syp | 29760.11 | 0.095862 | 0.43212 | 0.221842 | 0.824437 | 0.977246 |
| CG30094 | 416.4272 | 0.35077 | 1.574841 | 0.222733 | 0.823743 | 0.977246 |
| 28SrRNA-F | 105.0742 | 0.533571 | 2.402528 | 0.222088 | 0.824246 | 0.977246 |
| CG46448 | 5777.813 | 0.181583 | 0.817027 | 0.222249 | 0.82412 | 0.977246 |
| Nrt | 4114.058 | -0.1494 | 0.673127 | -0.22195 | 0.824357 | 0.977246 |
| CG6418 | 717.9133 | 0.381479 | 1.715633 | 0.222355 | 0.824038 | 0.977246 |
| cbt | 4385.378 | 0.143109 | 0.645932 | 0.221554 | 0.824661 | 0.977367 |
| lncRNA:CR | 134.5743 | 0.765794 | 3.461812 | 0.221212 | 0.824927 | 0.977538 |
| CG5068 | 3095.249 | 0.166912 | 0.755311 | 0.220984 | 0.825105 | 0.977604 |
| mRpL50 | 2010.33 | 0.166438 | 0.756122 | 0.22012 | 0.825778 | 0.978257 |
| MetRS-m | 1032.243 | -0.39508 | 1.802192 | -0.21922 | 0.826475 | 0.978268 |
| veli | 2967.415 | -0.14161 | 0.643871 | -0.21993 | 0.825928 | 0.978268 |
| rdo | 1728.794 | -0.25847 | 1.176828 | -0.21963 | 0.826158 | 0.978268 |
| wap | 1292.461 | 0.2069 | 0.942683 | 0.21948 | 0.826276 | 0.978268 |
| CG18809 | 1810.032 | -0.21285 | 0.968521 | -0.21976 | 0.826055 | 0.978268 |
| CG15715 | 2571.15 | -0.15735 | 0.717936 | -0.21917 | 0.826519 | 0.978268 |
| CG13337 | 69.75943 | -0.83772 | 3.834054 | -0.21849 | 0.827044 | 0.978606 |
| CG8839 | 3056.791 | -0.14597 | 0.668069 | -0.21849 | 0.827047 | 0.978606 |
| CG6329 | 24954.19 | -0.09677 | 0.443569 | -0.21817 | 0.827297 | 0.97862 |
| CG42526 | 71.06923 | -0.84318 | 3.864929 | -0.21816 | 0.827303 | 0.97862 |
| Eph | 10510.73 | -0.09896 | 0.454093 | -0.21792 | 0.827491 | 0.978697 |
| CG13527 | 107.7085 | -0.83443 | 3.833781 | -0.21765 | 0.827701 | 0.978802 |
| PIP4K | 9637.584 | -0.10029 | 0.462058 | -0.21704 | 0.828174 | 0.978951 |
| CG42514 | 393.0896 | -0.38358 | 1.765886 | -0.21722 | 0.828039 | 0.978951 |
| Zcchc7 | 958.6349 | 0.226888 | 1.045471 | 0.21702 | 0.828193 | 0.978951 |
| Ssdp | 5521.937 | 0.135425 | 0.624491 | 0.216857 | 0.82832 | 0.978957 |
| Rbcn-3B | 4280.406 | 0.137882 | 0.636907 | 0.216488 | 0.828608 | 0.979041 |
| l(1)10Bb | 2145.661 | 0.164687 | 0.760846 | 0.216453 | 0.828635 | 0.979041 |
| Roc1a | 3886.038 | 0.127128 | 0.588415 | 0.216052 | 0.828947 | 0.979098 |
| eIF2Bepsil | 142.3833 | -0.61128 | 2.831026 | -0.21592 | 0.829049 | 0.979098 |
| ND-42 | 6334.078 | -0.15132 | 0.700335 | -0.21606 | 0.82894 | 0.979098 |
| CG42764 | 443.6797 | 0.413816 | 1.918914 | 0.215651 | 0.82926 | 0.979202 |
| CG7492 | 833.6684 | 0.36745 | 1.706299 | 0.215349 | 0.829495 | 0.979336 |
| sws | 3095.877 | -0.14018 | 0.651891 | -0.21504 | 0.829735 | 0.979475 |

| | | | | | | |
|---|---|---|---|---|---|---|
| CG4452 | 789.2493 | -0.23348 | 1.086937 | -0.2148 | 0.829921 | 0.979551 |
| Chd64 | 9706.564 | -0.11827 | 0.552158 | -0.21419 | 0.830398 | 0.979969 |
| Ubc2 | 681.4667 | 0.239216 | 1.121677 | 0.213266 | 0.831119 | 0.980007 |
| mRpS22 | 1252.875 | 0.220368 | 1.034032 | 0.213115 | 0.831237 | 0.980007 |
| form3 | 836.1032 | -0.23961 | 1.121069 | -0.21373 | 0.830758 | 0.980007 |
| Ptx1 | 153.3054 | 0.657538 | 3.085466 | 0.213108 | 0.831243 | 0.980007 |
| Ank2 | 22039.78 | 0.086308 | 0.405025 | 0.213094 | 0.831254 | 0.980007 |
| bs | 360.7828 | 0.443521 | 2.080639 | 0.213166 | 0.831198 | 0.980007 |
| ABCB7 | 1237.421 | -0.19322 | 0.906917 | -0.21305 | 0.831284 | 0.980007 |
| lncRNA:CR | 254.9225 | 0.593415 | 2.78837 | 0.212818 | 0.831469 | 0.980081 |
| Cdk4 | 658.2236 | -0.29901 | 1.406599 | -0.21258 | 0.831656 | 0.980111 |
| Ubc10 | 1805.164 | -0.18799 | 0.884765 | -0.21247 | 0.831739 | 0.980111 |
| Prosbeta1 | 2538.019 | -0.14828 | 0.700139 | -0.21179 | 0.832274 | 0.980454 |
| RpL12 | 13642.73 | 0.102157 | 0.482308 | 0.211808 | 0.832257 | 0.980454 |
| CG11878 | 248.9864 | 0.535825 | 2.536462 | 0.211249 | 0.832693 | 0.980659 |
| SCaMC | 6830.622 | -0.13218 | 0.625434 | -0.21134 | 0.83262 | 0.980659 |
| Pis | 8983.544 | -0.10059 | 0.47859 | -0.21019 | 0.83352 | 0.98149 |
| yin | 872.6426 | 0.287367 | 1.369875 | 0.209776 | 0.833843 | 0.981581 |
| AlaRS-m | 1029.876 | -0.41238 | 1.965245 | -0.20984 | 0.833794 | 0.981581 |
| d4 | 2576.886 | -0.1668 | 0.797064 | -0.20927 | 0.834241 | 0.981766 |
| gfzf | 141.9836 | 0.678612 | 3.242897 | 0.209261 | 0.834244 | 0.981766 |
| MED21 | 3305.154 | 0.129032 | 0.61754 | 0.208945 | 0.834491 | 0.981809 |
| UK114 | 771.6636 | -0.22333 | 1.069086 | -0.2089 | 0.834526 | 0.981809 |
| Chchd3 | 3228.695 | -0.1342 | 0.643046 | -0.20869 | 0.83469 | 0.981858 |
| CG33635 | 1323.287 | 0.190242 | 0.913889 | 0.208167 | 0.835099 | 0.982195 |
| RpL37a | 33973.87 | -0.08092 | 0.389058 | -0.208 | 0.835233 | 0.982209 |
| trn | 1271.941 | -0.20202 | 0.97268 | -0.20769 | 0.835471 | 0.982341 |
| Trf5 | 314.8622 | 0.456392 | 2.199071 | 0.207539 | 0.835589 | 0.982341 |
| Socs44A | 2184.061 | 0.191489 | 0.923568 | 0.207337 | 0.835747 | 0.982382 |
| upSET | 11816 | -0.09893 | 0.479386 | -0.20637 | 0.836505 | 0.983129 |
| CG9775 | 4959.758 | -0.11186 | 0.545169 | -0.20518 | 0.837429 | 0.983172 |
| me31B | 5272.768 | -0.18884 | 0.920878 | -0.20506 | 0.837521 | 0.983172 |
| CG9426 | 1444.792 | 0.189263 | 0.918254 | 0.206112 | 0.836703 | 0.983172 |
| MBD-R2 | 940.6868 | 0.212899 | 1.038099 | 0.205086 | 0.837505 | 0.983172 |
| CG14259 | 78.32509 | 0.790693 | 3.844888 | 0.205648 | 0.837066 | 0.983172 |
| dj-1beta | 1547.506 | 0.166635 | 0.811874 | 0.205247 | 0.837379 | 0.983172 |
| wun2 | 5556.433 | 0.130929 | 0.63737 | 0.205422 | 0.837243 | 0.983172 |
| CG46462 | 70.7263 | -0.72856 | 3.551032 | -0.20517 | 0.83744 | 0.983172 |
| Rm62 | 13204.98 | 0.089186 | 0.438098 | 0.203575 | 0.838685 | 0.98324 |
| Rab14 | 8066.48 | -0.11656 | 0.570586 | -0.20428 | 0.838133 | 0.98324 |
| CG18766 | 1803.149 | 0.16258 | 0.801037 | 0.202962 | 0.839165 | 0.98324 |
| Pdhb | 8350.756 | 0.099376 | 0.485196 | 0.204815 | 0.837716 | 0.98324 |
| tra | 739.8706 | 0.227268 | 1.11124 | 0.204517 | 0.837949 | 0.98324 |
| nrv2 | 93494.08 | -0.10114 | 0.495103 | -0.20427 | 0.838139 | 0.98324 |
| Dpm3 | 2095.673 | 0.157069 | 0.771146 | 0.203683 | 0.838601 | 0.98324 |
| brwl | 896.8382 | -0.42973 | 2.116117 | -0.20308 | 0.839075 | 0.98324 |
| l(2)k09022 | 260.2253 | 0.369612 | 1.812021 | 0.203978 | 0.838371 | 0.98324 |
| Xxylt | 3747.875 | 0.13051 | 0.642468 | 0.203139 | 0.839027 | 0.98324 |
| asRNA:CR4 | 68.59508 | 0.78388 | 3.85461 | 0.203362 | 0.838852 | 0.98324 |
| CG44242 | 4714.874 | -0.11668 | 0.573832 | -0.20334 | 0.838869 | 0.98324 |

| | | | | | | |
|---|---|---|---|---|---|---|
| asRNA:CR4 | 117.6064 | -0.77803 | 3.833561 | -0.20295 | 0.839172 | 0.98324 |
| CG4360 | 994.8905 | 0.226008 | 1.11551 | 0.202605 | 0.839443 | 0.983249 |
| CG14257 | 146.6679 | -0.77434 | 3.836307 | -0.20185 | 0.840037 | 0.983249 |
| shf | 336.9299 | -0.45031 | 2.230382 | -0.2019 | 0.839997 | 0.983249 |
| CG42359 | 140.3182 | 0.615985 | 3.04703 | 0.202159 | 0.839792 | 0.983249 |
| CG42672 | 3042.39 | -0.16833 | 0.830265 | -0.20274 | 0.839338 | 0.983249 |
| bin3 | 4574.97 | -0.12733 | 0.629409 | -0.2023 | 0.839685 | 0.983249 |
| Frl | 1344.207 | -0.21201 | 1.050204 | -0.20187 | 0.840016 | 0.983249 |
| Drp1 | 2099.895 | 0.162059 | 0.803773 | 0.201623 | 0.840211 | 0.983309 |
| Pbgs | 529.9634 | -0.3119 | 1.54863 | -0.2014 | 0.840384 | 0.983369 |
| CG31642 | 198.3078 | -0.70988 | 3.535388 | -0.20079 | 0.840861 | 0.983599 |
| ct | 32943.88 | -0.09644 | 0.480938 | -0.20052 | 0.841071 | 0.983599 |
| Ubr1 | 1816.394 | -0.20059 | 0.999905 | -0.20061 | 0.841003 | 0.983599 |
| CG6961 | 251.935 | 0.433239 | 2.157833 | 0.200775 | 0.840874 | 0.983599 |
| Hmu | 2446.811 | 0.152871 | 0.764825 | 0.199876 | 0.841577 | 0.983904 |
| Pmp70 | 1404.371 | -0.17242 | 0.861976 | -0.20003 | 0.841459 | 0.983904 |
| CG11267 | 6622.815 | 0.10787 | 0.540348 | 0.19963 | 0.84177 | 0.983986 |
| AANATL7 | 305.0909 | -0.38783 | 1.95291 | -0.19859 | 0.842583 | 0.984017 |
| CG7694 | 905.5047 | 0.204947 | 1.031229 | 0.19874 | 0.842466 | 0.984017 |
| Dh31-R | 672.6482 | -0.40879 | 2.056895 | -0.19874 | 0.842464 | 0.984017 |
| CG42594 | 737.9709 | -0.22079 | 1.111823 | -0.19859 | 0.842587 | 0.984017 |
| Mmp1 | 3089.289 | 0.147185 | 0.741516 | 0.198492 | 0.84266 | 0.984017 |
| Cdc37 | 1205.161 | 0.186791 | 0.937289 | 0.199289 | 0.842037 | 0.984017 |
| CG32368 | 545.6863 | -0.27812 | 1.398467 | -0.19888 | 0.842358 | 0.984017 |
| CG10075 | 990.037 | 0.205104 | 1.034095 | 0.198342 | 0.842777 | 0.984017 |
| Tlk | 4725.492 | -0.11651 | 0.588861 | -0.19786 | 0.843151 | 0.98431 |
| CG18048 | 198.8484 | 0.47432 | 2.400003 | 0.197633 | 0.843332 | 0.984378 |
| CG15160 | 257.8302 | 0.385179 | 1.951992 | 0.197326 | 0.843572 | 0.984515 |
| Nak | 5087.472 | -0.12264 | 0.624638 | -0.19633 | 0.844351 | 0.984746 |
| mRpS24 | 179.8789 | -0.38231 | 1.945001 | -0.19656 | 0.844173 | 0.984746 |
| RpL13 | 16772.8 | 0.087704 | 0.446811 | 0.19629 | 0.844383 | 0.984746 |
| mt:tRNA:A | 85.40259 | -0.51476 | 2.621427 | -0.19637 | 0.844324 | 0.984746 |
| Cngl | 14108.12 | 0.09303 | 0.47267 | 0.196817 | 0.843971 | 0.984746 |
| egh | 5425.419 | -0.10755 | 0.550399 | -0.1954 | 0.845078 | 0.985093 |
| Atox1 | 4551.807 | 0.137381 | 0.705235 | 0.194802 | 0.845548 | 0.985093 |
| Acbp1 | 1826.273 | -0.15039 | 0.769309 | -0.19549 | 0.845007 | 0.985093 |
| CG31674 | 296.3822 | -0.42233 | 2.168331 | -0.19477 | 0.845573 | 0.985093 |
| Ctr1A | 2268.635 | -0.14322 | 0.73419 | -0.19507 | 0.84534 | 0.985093 |
| PI31 | 2503.971 | -0.16185 | 0.830601 | -0.19486 | 0.845506 | 0.985093 |
| lncRNA:CR | 64.32886 | -0.74842 | 3.834794 | -0.19516 | 0.845264 | 0.985093 |
| SCOT | 103.9344 | -0.5951 | 3.057223 | -0.19466 | 0.845663 | 0.985093 |
| CG14050 | 499.2616 | -0.48025 | 2.491304 | -0.19277 | 0.847138 | 0.985347 |
| msl-1 | 913.117 | -0.20215 | 1.049644 | -0.19259 | 0.84728 | 0.985347 |
| drl | 896.8554 | 0.323211 | 1.665509 | 0.194061 | 0.846128 | 0.985347 |
| Mapmodu | 6307.186 | 0.115337 | 0.598736 | 0.192634 | 0.847246 | 0.985347 |
| ari-2 | 1176.523 | -0.22941 | 1.189941 | -0.19279 | 0.847125 | 0.985347 |
| KCNQ | 656.6538 | -0.23827 | 1.238737 | -0.19235 | 0.847467 | 0.985347 |
| 14-3-3zeta | 36762.8 | -0.07769 | 0.402679 | -0.19293 | 0.847011 | 0.985347 |
| Stat92E | 1936.038 | -0.17749 | 0.915021 | -0.19398 | 0.846194 | 0.985347 |
| PlexA | 19244.32 | -0.08553 | 0.444788 | -0.19229 | 0.847517 | 0.985347 |

| | | | | | | |
|---|---|---|---|---|---|---|
| Sec22 | 1894.591 | 0.193455 | 0.999997 | 0.193455 | 0.846602 | 0.985347 |
| pre-lola-G | 113.9461 | -0.73663 | 3.833863 | -0.19214 | 0.847635 | 0.985347 |
| Pcf11 | 9826.343 | 0.10286 | 0.532436 | 0.193187 | 0.846812 | 0.985347 |
| CG32485 | 3234.237 | 0.120214 | 0.625619 | 0.192153 | 0.847623 | 0.985347 |
| CG4645 | 1300.011 | -0.22469 | 1.166441 | -0.19263 | 0.847246 | 0.985347 |
| GstT3 | 799.4026 | 0.302527 | 1.575459 | 0.192025 | 0.847723 | 0.985347 |
| Sin1 | 583.1408 | 0.409362 | 2.136129 | 0.191637 | 0.848026 | 0.985557 |
| dwg | 87.18679 | -0.63467 | 3.333585 | -0.19039 | 0.849006 | 0.985697 |
| TrxT | 119.6643 | 0.7312 | 3.83368 | 0.190731 | 0.848737 | 0.985697 |
| CARPA | 1088.885 | 0.287525 | 1.505218 | 0.191019 | 0.848511 | 0.985697 |
| CR30279 | 72.00752 | 0.752398 | 3.951211 | 0.190422 | 0.848978 | 0.985697 |
| CG41128 | 6679.887 | 0.096042 | 0.504093 | 0.190524 | 0.848899 | 0.985697 |
| CG10384 | 1486.114 | -0.27057 | 1.420012 | -0.19054 | 0.848886 | 0.985697 |
| CG10347 | 534.3419 | 0.370362 | 1.939151 | 0.190992 | 0.848532 | 0.985697 |
| RhoL | 839.0352 | 0.332737 | 1.759734 | 0.189084 | 0.850027 | 0.985708 |
| CG31441 | 143.723 | 0.662595 | 3.496868 | 0.189482 | 0.849715 | 0.985708 |
| Ppt1 | 291.3919 | 0.402775 | 2.135302 | 0.188627 | 0.850385 | 0.985708 |
| CG4038 | 62.25393 | 0.677945 | 3.565441 | 0.190143 | 0.849197 | 0.985708 |
| CG12384 | 7287.017 | -0.09766 | 0.513893 | -0.19004 | 0.849279 | 0.985708 |
| Vang | 232.1772 | 0.393422 | 2.076874 | 0.18943 | 0.849756 | 0.985708 |
| CG3091 | 2596.033 | -0.21969 | 1.159856 | -0.18941 | 0.849768 | 0.985708 |
| Vps45 | 584.689 | -0.34557 | 1.834832 | -0.18834 | 0.850612 | 0.985708 |
| CG5569 | 2088.8 | -0.13889 | 0.734817 | -0.18901 | 0.850082 | 0.985708 |
| asRNA:CR4 | 93.24062 | -0.67922 | 3.57953 | -0.18975 | 0.849503 | 0.985708 |
| Gbeta13F | 23954.76 | 0.075922 | 0.402567 | 0.188594 | 0.850411 | 0.985708 |
| Rrp47 | 2021.629 | -0.27859 | 1.475426 | -0.18882 | 0.850233 | 0.985708 |
| Hsp67Ba | 89.61623 | -0.6064 | 3.219567 | -0.18835 | 0.850604 | 0.985708 |
| mRpL14 | 1899.609 | -0.15752 | 0.853256 | -0.18461 | 0.853536 | 0.986018 |
| CG18508 | 1449.398 | 0.155955 | 0.830319 | 0.187825 | 0.851014 | 0.986018 |
| Trf | 383.2775 | -0.34617 | 1.881035 | -0.18403 | 0.85399 | 0.986018 |
| CG4968 | 1921.564 | 0.143384 | 0.769334 | 0.186374 | 0.852151 | 0.986018 |
| CG10268 | 297.5112 | -0.3507 | 1.897922 | -0.18478 | 0.853399 | 0.986018 |
| CG10038 | 757.0076 | 0.317572 | 1.734813 | 0.183058 | 0.854752 | 0.986018 |
| mRpS21 | 326.0281 | -0.29335 | 1.580232 | -0.18564 | 0.852731 | 0.986018 |
| smp-30 | 174.0951 | 0.449126 | 2.464326 | 0.182251 | 0.855386 | 0.986018 |
| CG31213 | 64.85324 | 0.635098 | 3.417899 | 0.185815 | 0.85259 | 0.986018 |
| PQBP1 | 982.7398 | 0.184836 | 1.002543 | 0.184367 | 0.853726 | 0.986018 |
| CG10916 | 1306.833 | -0.18267 | 0.994684 | -0.18365 | 0.854291 | 0.986018 |
| Rrp42 | 318.7628 | -0.42555 | 2.280322 | -0.18662 | 0.851959 | 0.986018 |
| Rho1 | 8124.773 | -0.09083 | 0.488398 | -0.18597 | 0.852467 | 0.986018 |
| AstC-R1 | 354.4318 | -0.27557 | 1.509119 | -0.1826 | 0.855109 | 0.986018 |
| Nmda1 | 11458.69 | -0.08828 | 0.483276 | -0.18267 | 0.855054 | 0.986018 |
| Ir41a | 124.6447 | 0.704579 | 3.833785 | 0.183782 | 0.854185 | 0.986018 |
| CG32547 | 131.1415 | 0.590994 | 3.185697 | 0.185515 | 0.852825 | 0.986018 |
| Pink1 | 1678.609 | 0.172911 | 0.935638 | 0.184806 | 0.853381 | 0.986018 |
| CG1024 | 72.70029 | -0.6986 | 3.834242 | -0.1822 | 0.855426 | 0.986018 |
| CG1578 | 2930.471 | -0.13085 | 0.713724 | -0.18334 | 0.854532 | 0.986018 |
| Zw10 | 290.3819 | 0.413433 | 2.22089 | 0.186156 | 0.852322 | 0.986018 |
| CG34195 | 105.3001 | -0.63029 | 3.419464 | -0.18433 | 0.853758 | 0.986018 |
| CG34355 | 751.8575 | 0.232496 | 1.248342 | 0.186244 | 0.852254 | 0.986018 |

| | | | | | | |
|---|---|---|---|---|---|---|
| CG42331 | 269.9834 | -0.34361 | 1.879564 | -0.18281 | 0.854945 | 0.986018 |
| zld | 631.9467 | 0.268391 | 1.450986 | 0.184972 | 0.853251 | 0.986018 |
| Rbp6 | 50090.46 | 0.07479 | 0.403421 | 0.18539 | 0.852923 | 0.986018 |
| CG12848 | 829.8916 | 0.196383 | 1.061016 | 0.185089 | 0.853159 | 0.986018 |
| Hr4 | 327.5325 | -0.34603 | 1.885934 | -0.18348 | 0.854422 | 0.986018 |
| rg | 86.89347 | -0.69899 | 3.835676 | -0.18223 | 0.8554 | 0.986018 |
| otk2 | 64.97449 | -0.7015 | 3.835242 | -0.18291 | 0.85487 | 0.986018 |
| mRpL23 | 2830.584 | 0.121294 | 0.660513 | 0.183636 | 0.854299 | 0.986018 |
| hec | 264.4119 | 0.45856 | 2.503523 | 0.183166 | 0.854668 | 0.986018 |
| sbm | 2858.411 | 0.152159 | 0.832517 | 0.18277 | 0.854979 | 0.986018 |
| CG32196 | 192.1696 | -0.48969 | 2.63564 | -0.1858 | 0.852604 | 0.986018 |
| beg | 364.3977 | -0.42296 | 2.27504 | -0.18591 | 0.852512 | 0.986018 |
| fz | 853.7399 | -0.19074 | 1.038574 | -0.18366 | 0.854283 | 0.986018 |
| CG15456 | 2137.659 | -0.13986 | 0.757345 | -0.18467 | 0.853489 | 0.986018 |
| COX5A | 27630.81 | 0.078376 | 0.434786 | 0.180262 | 0.856947 | 0.986496 |
| CG12321 | 460.6221 | -0.36419 | 2.008236 | -0.18135 | 0.856096 | 0.986496 |
| CG3376 | 2459.295 | 0.159419 | 0.880723 | 0.181009 | 0.85636 | 0.986496 |
| CG40439 | 638.2277 | -0.20867 | 1.157475 | -0.18028 | 0.856931 | 0.986496 |
| ken | 550.9689 | 0.230395 | 1.277011 | 0.180418 | 0.856825 | 0.986496 |
| lncRNA:CR | 200.8819 | 0.333942 | 1.847519 | 0.180752 | 0.856562 | 0.986496 |
| CG2211 | 327.4135 | 0.390888 | 2.16522 | 0.18053 | 0.856736 | 0.986496 |
| CG12093 | 1057.301 | -0.18442 | 1.016844 | -0.18136 | 0.856082 | 0.986496 |
| CG12012 | 2646.748 | 0.135191 | 0.748781 | 0.180548 | 0.856722 | 0.986496 |
| Slc45-1 | 532.4628 | 0.447839 | 2.486746 | 0.18009 | 0.857082 | 0.98651 |
| Sema2b | 1873.442 | 0.151369 | 0.843812 | 0.179387 | 0.857633 | 0.986752 |
| Agpat1 | 831.7529 | -0.28319 | 1.57672 | -0.17961 | 0.857461 | 0.986752 |
| CG14618 | 584.9313 | 0.352051 | 1.962898 | 0.179353 | 0.857661 | 0.986752 |
| SMC5 | 820.0139 | -0.20974 | 1.178457 | -0.17798 | 0.858738 | 0.986796 |
| elm | 5734.819 | 0.09501 | 0.534361 | 0.177801 | 0.858879 | 0.986796 |
| CG14641 | 1225.641 | 0.169976 | 0.961124 | 0.176851 | 0.859625 | 0.986796 |
| CG13397 | 598.6891 | -0.29831 | 1.686299 | -0.1769 | 0.859587 | 0.986796 |
| pxb | 2724.783 | 0.199798 | 1.118056 | 0.178701 | 0.858172 | 0.986796 |
| luna | 4772.698 | 0.123085 | 0.689566 | 0.178497 | 0.858333 | 0.986796 |
| mmd | 8796.535 | -0.08539 | 0.481754 | -0.17724 | 0.85932 | 0.986796 |
| CG42342 | 20763.76 | 0.080364 | 0.45455 | 0.1768 | 0.859666 | 0.986796 |
| dpr21 | 12021.6 | 0.079423 | 0.446581 | 0.177848 | 0.858843 | 0.986796 |
| asRNA:CR4 | 87.94271 | -0.59253 | 3.313849 | -0.17881 | 0.85809 | 0.986796 |
| CG44838 | 9158.091 | 0.093238 | 0.522693 | 0.17838 | 0.858425 | 0.986796 |
| lncRNA:CR | 251.3423 | -0.34256 | 1.93003 | -0.17749 | 0.859123 | 0.986796 |
| 28SrRNA-F | 67.03038 | -0.51345 | 2.886467 | -0.17788 | 0.858816 | 0.986796 |
| lncRNA:CR | 304.7737 | 0.418581 | 2.363528 | 0.1771 | 0.85943 | 0.986796 |
| Gad1 | 11355.09 | 0.105609 | 0.595682 | 0.177291 | 0.85928 | 0.986796 |
| toc | 5540.668 | -0.11907 | 0.6707 | -0.17753 | 0.859092 | 0.986796 |
| CG13694 | 246.3114 | 0.606981 | 3.436286 | 0.176639 | 0.859792 | 0.9868 |
| maf-S | 2141.51 | -0.13203 | 0.748146 | -0.17648 | 0.859919 | 0.986804 |
| Madm | 1212.79 | 0.191359 | 1.088793 | 0.175754 | 0.860487 | 0.986899 |
| CG17991 | 200.4158 | -0.54301 | 3.08975 | -0.17575 | 0.860493 | 0.986899 |
| Df31 | 3403.624 | 0.11721 | 0.666289 | 0.175914 | 0.860361 | 0.986899 |
| RpL19 | 16361.64 | 0.085876 | 0.48734 | 0.176213 | 0.860127 | 0.986899 |
| Rop | 17354.39 | -0.07617 | 0.434767 | -0.1752 | 0.860924 | 0.987252 |

| | | | | | | |
|---|---|---|---|---|---|---|
| CG31100 | 1616.24 | -0.15745 | 0.900648 | -0.17482 | 0.861222 | 0.987449 |
| su(Hw) | 300.0793 | 0.327354 | 1.879037 | 0.174214 | 0.861697 | 0.987449 |
| CG31313 | 172.5414 | -0.50779 | 2.924533 | -0.17363 | 0.862154 | 0.987449 |
| Rpb7 | 362.9142 | -0.32562 | 1.879549 | -0.17324 | 0.862461 | 0.987449 |
| Rab2 | 4551.61 | -0.10509 | 0.604221 | -0.17392 | 0.86193 | 0.987449 |
| CG13428 | 81.219 | -0.46968 | 2.698906 | -0.17403 | 0.861846 | 0.987449 |
| Pu | 1790.224 | 0.136141 | 0.784338 | 0.173574 | 0.8622 | 0.987449 |
| Uhg4 | 812.7021 | 0.18618 | 1.07242 | 0.173608 | 0.862174 | 0.987449 |
| Pgk | 8168.143 | -0.0838 | 0.483768 | -0.17322 | 0.862476 | 0.987449 |
| Uhg3 | 1016.421 | -0.1879 | 1.079007 | -0.17414 | 0.861754 | 0.987449 |
| lncRNA:CR | 64.54478 | -0.62143 | 3.590001 | -0.1731 | 0.862573 | 0.987449 |
| CG5541 | 207.8627 | 0.378325 | 2.183753 | 0.173246 | 0.862458 | 0.987449 |
| CG3918 | 107.2819 | 0.571449 | 3.325515 | 0.171838 | 0.863565 | 0.9875 |
| ORMDL | 1538.426 | 0.146487 | 0.849703 | 0.172398 | 0.863125 | 0.9875 |
| Fip1 | 1266.203 | 0.164909 | 0.957635 | 0.172204 | 0.863277 | 0.9875 |
| mRpS18A | 3055.021 | 0.111735 | 0.649013 | 0.172162 | 0.86331 | 0.9875 |
| Bin1 | 1202.464 | -0.15362 | 0.896972 | -0.17126 | 0.864016 | 0.9875 |
| Shark | 145.8989 | 0.543196 | 3.144318 | 0.172755 | 0.862844 | 0.9875 |
| porin | 26213.55 | -0.07117 | 0.414829 | -0.17157 | 0.863773 | 0.9875 |
| nocte | 6420.909 | -0.0994 | 0.581245 | -0.17101 | 0.864216 | 0.9875 |
| gas | 158.6183 | 0.656985 | 3.83345 | 0.171382 | 0.863923 | 0.9875 |
| pre-rRNA:( | 199.1853 | 0.327731 | 1.911904 | 0.171416 | 0.863897 | 0.9875 |
| CG9132 | 1635.104 | 0.143449 | 0.835197 | 0.171755 | 0.86363 | 0.9875 |
| Usp32 | 1150.674 | -0.24636 | 1.435812 | -0.17158 | 0.863766 | 0.9875 |
| CG1695 | 5775.081 | -0.10018 | 0.58563 | -0.17107 | 0.864171 | 0.9875 |
| LamC | 104.619 | 0.500438 | 2.933108 | 0.170617 | 0.864525 | 0.987571 |
| CG7860 | 129.3092 | -0.6233 | 3.652076 | -0.17067 | 0.864483 | 0.987571 |
| PH4alphaE | 1258.04 | 0.164187 | 0.968721 | 0.169488 | 0.865412 | 0.988411 |
| Rx | 107.0353 | -0.61293 | 3.618922 | -0.16937 | 0.865506 | 0.988411 |
| Mrp4 | 496.5041 | 0.252001 | 1.490649 | 0.169054 | 0.865754 | 0.988553 |
| tos | 81.21191 | 0.582423 | 3.462658 | 0.168201 | 0.866425 | 0.989071 |
| CG43658 | 689.4383 | 0.20669 | 1.229094 | 0.168164 | 0.866454 | 0.989071 |
| ND-B16.6 | 7348.974 | 0.089777 | 0.534467 | 0.167975 | 0.866603 | 0.9891 |
| EndoGI | 528.1745 | -0.21154 | 1.265011 | -0.16722 | 0.867195 | 0.989496 |
| Bruce | 5252.175 | 0.116385 | 0.696649 | 0.167065 | 0.867319 | 0.989496 |
| Etf-QO | 259.1985 | 0.347491 | 2.079889 | 0.167072 | 0.867313 | 0.989496 |
| CG11248 | 644.2836 | 0.205592 | 1.233394 | 0.166688 | 0.867616 | 0.98958 |
| CSN5 | 716.6741 | -0.30636 | 1.838237 | -0.16666 | 0.867639 | 0.98958 |
| Mlc-c | 2299.95 | 0.123435 | 0.741618 | 0.166441 | 0.86781 | 0.989634 |
| CG45676 | 1053.734 | 0.185491 | 1.116409 | 0.16615 | 0.868039 | 0.989755 |
| Scgalpha | 1574.295 | 0.147146 | 0.886614 | 0.165964 | 0.868185 | 0.98978 |
| eIF3b | 3715.867 | -0.0981 | 0.593219 | -0.16537 | 0.868655 | 0.989975 |
| CG10307 | 770.8143 | 0.245535 | 1.486717 | 0.165152 | 0.868824 | 0.989975 |
| DAT | 142.44 | 0.57094 | 3.44977 | 0.165501 | 0.86855 | 0.989975 |
| CG32521 | 13107.09 | -0.10468 | 0.633957 | -0.16512 | 0.868849 | 0.989975 |
| RpL6 | 15094.1 | -0.08329 | 0.505363 | -0.1648 | 0.869099 | 0.99001 |
| COX7C | 17138.91 | -0.06899 | 0.418707 | -0.16477 | 0.869127 | 0.99001 |
| CG15657 | 79.1199 | -0.64094 | 3.898843 | -0.16439 | 0.869421 | 0.990205 |
| SP1029 | 1113.517 | -0.20166 | 1.228851 | -0.1641 | 0.869651 | 0.990327 |
| lncRNA:CR | 167.6861 | 0.462088 | 2.820531 | 0.16383 | 0.869865 | 0.990429 |

| | | | | | | |
|---|---|---|---|---|---|---|
| CG31122 | 2893.669 | 0.132356 | 0.810753 | 0.163251 | 0.870321 | 0.990808 |
| Npc2b | 1473.389 | -0.16491 | 1.011854 | -0.16298 | 0.870537 | 0.990914 |
| if | 3059.008 | -0.12645 | 0.777281 | -0.16268 | 0.870771 | 0.991039 |
| Cyp28a5 | 11392.76 | -0.0907 | 0.559641 | -0.16206 | 0.871257 | 0.991217 |
| CG1360 | 208.6804 | 0.486314 | 3.001395 | 0.162029 | 0.871283 | 0.991217 |
| CG13563 | 174.1058 | 0.427799 | 2.643992 | 0.161801 | 0.871463 | 0.991217 |
| lncRNA:CR | 171.167 | -0.32418 | 2.001232 | -0.16199 | 0.871314 | 0.991217 |
| HP4 | 1252.871 | -0.15827 | 0.97879 | -0.1617 | 0.871545 | 0.991217 |
| CG10702 | 144.4691 | 0.497647 | 3.089269 | 0.161089 | 0.872023 | 0.991476 |
| CG31199 | 68.17882 | 0.617432 | 3.83952 | 0.16081 | 0.872243 | 0.991476 |
| CngA | 140.6707 | 0.345915 | 2.146194 | 0.161176 | 0.871955 | 0.991476 |
| levy | 29446.61 | -0.0649 | 0.404045 | -0.16062 | 0.87239 | 0.991476 |
| CG8675 | 333.4154 | -0.34676 | 2.156882 | -0.16077 | 0.872275 | 0.991476 |
| CG46301 | 498.5398 | -0.20503 | 1.278404 | -0.16038 | 0.872583 | 0.991555 |
| RpL10 | 14946.88 | 0.07064 | 0.442685 | 0.159571 | 0.873219 | 0.991583 |
| CG13365 | 2212.954 | 0.117061 | 0.754891 | 0.155071 | 0.876766 | 0.991583 |
| CG3706 | 307.5044 | 0.253304 | 1.787419 | 0.141715 | 0.887305 | 0.991583 |
| CG13760 | 2102.217 | 0.134467 | 0.861413 | 0.1561 | 0.875954 | 0.991583 |
| CG10802 | 418.3823 | -0.20972 | 1.535039 | -0.13662 | 0.891329 | 0.991583 |
| Parg | 474.1553 | 0.362033 | 2.454414 | 0.147503 | 0.882735 | 0.991583 |
| CG14270 | 1946.841 | 0.143125 | 1.038662 | 0.137797 | 0.890401 | 0.991583 |
| Rpn13R | 194.595 | -0.39868 | 2.739288 | -0.14554 | 0.884283 | 0.991583 |
| RpL35 | 16224.56 | 0.072077 | 0.472692 | 0.152483 | 0.878806 | 0.991583 |
| CG5274 | 450.7229 | 0.311457 | 2.08979 | 0.149038 | 0.881524 | 0.991583 |
| CG10512 | 3141.926 | -0.10041 | 0.705162 | -0.14239 | 0.88677 | 0.991583 |
| SAK | 176.7823 | 0.280398 | 2.010618 | 0.139459 | 0.889088 | 0.991583 |
| RpLP0 | 18841.2 | 0.059881 | 0.434831 | 0.13771 | 0.890469 | 0.991583 |
| Atg17 | 2756.096 | 0.115454 | 0.773928 | 0.149179 | 0.881412 | 0.991583 |
| Pgant5 | 2943.465 | -0.09336 | 0.651643 | -0.14327 | 0.886081 | 0.991583 |
| Vps52 | 608.0301 | -0.19884 | 1.268701 | -0.15673 | 0.875457 | 0.991583 |
| CG9117 | 97.60515 | 0.530377 | 3.579485 | 0.148171 | 0.882208 | 0.991583 |
| Cpr | 1722.884 | -0.15865 | 1.11612 | -0.14215 | 0.886965 | 0.991583 |
| CG9498 | 234.1589 | 0.390979 | 2.861902 | 0.136615 | 0.891335 | 0.991583 |
| COX5B | 18739.53 | -0.06643 | 0.438738 | -0.15142 | 0.879645 | 0.991583 |
| RpL7 | 23287.02 | 0.07673 | 0.506011 | 0.151637 | 0.879473 | 0.991583 |
| Dpy-30L1 | 1786.474 | -0.14711 | 1.048823 | -0.14026 | 0.888454 | 0.991583 |
| CG17124 | 2930.305 | -0.08897 | 0.648875 | -0.13711 | 0.89094 | 0.991583 |
| CG5945 | 450.1044 | 0.360562 | 2.454535 | 0.146896 | 0.883214 | 0.991583 |
| NimC2 | 68.72222 | -0.54075 | 3.951225 | -0.13686 | 0.891144 | 0.991583 |
| Sidpn | 430.7351 | 0.21716 | 1.377149 | 0.157688 | 0.874702 | 0.991583 |
| l(2)37Bb | 754.8768 | -0.23982 | 1.69465 | -0.14152 | 0.887462 | 0.991583 |
| CG31800 | 513.522 | -0.2071 | 1.419721 | -0.14587 | 0.884023 | 0.991583 |
| CG2656 | 220.4888 | 0.418466 | 2.802951 | 0.149295 | 0.881321 | 0.991583 |
| IscU | 29031.46 | -0.06384 | 0.41665 | -0.15323 | 0.878217 | 0.991583 |
| RnpS1 | 371.3262 | -0.28231 | 1.848934 | -0.15269 | 0.878646 | 0.991583 |
| CG9393 | 529.9706 | -0.27383 | 1.924055 | -0.14232 | 0.886829 | 0.991583 |
| CG6254 | 609.144 | 0.202123 | 1.437784 | 0.140579 | 0.888202 | 0.991583 |
| CG12279 | 741.165 | -0.16214 | 1.09537 | -0.14803 | 0.882322 | 0.991583 |
| ND-23 | 12351.66 | -0.06646 | 0.456434 | -0.1456 | 0.884235 | 0.991583 |
| CG3534 | 112.4778 | -0.52258 | 3.361226 | -0.15547 | 0.876449 | 0.991583 |

| | | | | | | |
|---|---|---|---|---|---|---|
| Trm7-32 | 1713.907 | 0.119283 | 0.840231 | 0.141965 | 0.887108 | 0.991583 |
| Mur89F | 98.72409 | 0.531628 | 3.869602 | 0.137386 | 0.890726 | 0.991583 |
| Brf | 240.7105 | -0.29453 | 2.001517 | -0.14715 | 0.883012 | 0.991583 |
| Octalpha2l | 7649.206 | 0.073348 | 0.485481 | 0.151084 | 0.87991 | 0.991583 |
| loco | 1244.325 | 0.135518 | 0.923606 | 0.146727 | 0.883348 | 0.991583 |
| CG31145 | 3907.875 | -0.08905 | 0.649516 | -0.13711 | 0.890944 | 0.991583 |
| CG5986 | 878.1321 | -0.17068 | 1.109992 | -0.15377 | 0.877795 | 0.991583 |
| CG13630 | 342.063 | -0.33774 | 2.191134 | -0.15414 | 0.8775 | 0.991583 |
| CG31510 | 1412.536 | 0.141245 | 0.977933 | 0.144432 | 0.885159 | 0.991583 |
| LpR1 | 3309.923 | -0.10308 | 0.731302 | -0.14096 | 0.887905 | 0.991583 |
| CG4849 | 165.5365 | -0.3649 | 2.409656 | -0.15143 | 0.879635 | 0.991583 |
| eIF4E6 | 306.0611 | -0.29608 | 1.850466 | -0.16 | 0.872881 | 0.991583 |
| CG33203 | 7556.348 | -0.06892 | 0.48822 | -0.14117 | 0.887738 | 0.991583 |
| IntS11 | 618.0799 | 0.231035 | 1.454987 | 0.158788 | 0.873836 | 0.991583 |
| ca | 1056.805 | -0.15692 | 0.9843 | -0.15943 | 0.873332 | 0.991583 |
| TkR99D | 121.6463 | 0.410267 | 2.637051 | 0.155578 | 0.876366 | 0.991583 |
| Sry-delta | 862.4077 | -0.19209 | 1.315491 | -0.14602 | 0.883903 | 0.991583 |
| CG1896 | 366.3091 | -0.34341 | 2.292974 | -0.14977 | 0.880949 | 0.991583 |
| heph | 26052.53 | -0.05883 | 0.405665 | -0.14502 | 0.884698 | 0.991583 |
| CG11159 | 108.0194 | 0.563553 | 3.833835 | 0.146995 | 0.883136 | 0.991583 |
| qkr54B | 3753.904 | 0.089216 | 0.603723 | 0.147777 | 0.882519 | 0.991583 |
| CG3198 | 5548.641 | -0.10376 | 0.742387 | -0.13977 | 0.888841 | 0.991583 |
| CG9034 | 5030.602 | 0.08515 | 0.605513 | 0.140624 | 0.888167 | 0.991583 |
| Khc | 1407.569 | 0.147994 | 1.058049 | 0.139875 | 0.888759 | 0.991583 |
| ATPCL | 6231.981 | 0.080077 | 0.550309 | 0.145513 | 0.884306 | 0.991583 |
| GPHR | 2555.991 | -0.11522 | 0.722269 | -0.15952 | 0.873257 | 0.991583 |
| fand | 312.8577 | 0.307026 | 2.089588 | 0.146931 | 0.883186 | 0.991583 |
| antr | 302.1354 | -0.33632 | 2.435515 | -0.13809 | 0.890171 | 0.991583 |
| Sln | 1143.303 | 0.160331 | 1.029861 | 0.155683 | 0.876283 | 0.991583 |
| CG18336 | 245.4509 | -0.4398 | 2.781385 | -0.15812 | 0.87436 | 0.991583 |
| ND-B14 | 11671.04 | 0.066783 | 0.447617 | 0.149196 | 0.881399 | 0.991583 |
| Prx2540-2 | 110.2676 | 0.471373 | 3.083001 | 0.152894 | 0.878482 | 0.991583 |
| CG1407 | 682.1375 | 0.241297 | 1.745541 | 0.138236 | 0.890054 | 0.991583 |
| CG11191 | 5815.176 | -0.11795 | 0.782387 | -0.15075 | 0.880171 | 0.991583 |
| Tsp42En | 107.638 | -0.55485 | 3.50647 | -0.15824 | 0.874271 | 0.991583 |
| Adk3 | 5663.268 | -0.08438 | 0.553767 | -0.15238 | 0.87889 | 0.991583 |
| Ephrin | 5413.993 | 0.090371 | 0.628232 | 0.14385 | 0.885619 | 0.991583 |
| mRpL13 | 761.731 | -0.16648 | 1.153826 | -0.14429 | 0.885274 | 0.991583 |
| Aprt | 169.4549 | -0.46311 | 3.111949 | -0.14882 | 0.881698 | 0.991583 |
| CG30159 | 444.1092 | -0.28131 | 1.874085 | -0.15011 | 0.880681 | 0.991583 |
| DIP1 | 7562.891 | -0.0791 | 0.572828 | -0.13809 | 0.890173 | 0.991583 |
| GCS2alpha | 12200.17 | -0.07208 | 0.459381 | -0.15691 | 0.875314 | 0.991583 |
| Rab35 | 2863.165 | 0.089399 | 0.650266 | 0.13748 | 0.890651 | 0.991583 |
| CG32576 | 2162.727 | 0.116523 | 0.742094 | 0.157019 | 0.87523 | 0.991583 |
| Gyc88E | 632.1522 | 0.162589 | 1.167208 | 0.139298 | 0.889215 | 0.991583 |
| Mgat2 | 7578.399 | 0.075391 | 0.501719 | 0.150266 | 0.880555 | 0.991583 |
| dlg1 | 13289.21 | 0.0629 | 0.445506 | 0.141187 | 0.887722 | 0.991583 |
| Mtr3 | 362.8816 | 0.393877 | 2.567123 | 0.153431 | 0.878058 | 0.991583 |
| Trl | 445.7769 | 0.278605 | 1.8649 | 0.149394 | 0.881243 | 0.991583 |
| Spt7 | 611.597 | -0.29532 | 1.902974 | -0.15519 | 0.876671 | 0.991583 |

| | | | | | | |
|---|---|---|---|---|---|---|
| CG34120 | 135.8097 | 0.53503 | 3.571903 | 0.149788 | 0.880932 | 0.991583 |
| Nlg4 | 3379.551 | 0.102276 | 0.644782 | 0.158621 | 0.873967 | 0.991583 |
| trv | 6535.813 | 0.090272 | 0.581778 | 0.155166 | 0.876691 | 0.991583 |
| CG34454 | 99.92102 | -0.57468 | 3.951127 | -0.14545 | 0.884358 | 0.991583 |
| Alg-2 | 4206.096 | -0.08702 | 0.600009 | -0.14503 | 0.884685 | 0.991583 |
| CG42361 | 4524.186 | -0.08656 | 0.601886 | -0.14381 | 0.885647 | 0.991583 |
| Dys | 2636.303 | -0.11146 | 0.746176 | -0.14938 | 0.881254 | 0.991583 |
| CG42588 | 2370.2 | 0.137464 | 0.933794 | 0.14721 | 0.882966 | 0.991583 |
| Lmpt | 1284.911 | -0.16168 | 1.170172 | -0.13817 | 0.89011 | 0.991583 |
| lncRNA:CR | 158.3725 | -0.44854 | 3.078842 | -0.14569 | 0.88417 | 0.991583 |
| CG42795 | 12952.33 | 0.082663 | 0.524678 | 0.15755 | 0.874811 | 0.991583 |
| Gprk2 | 2210.768 | -0.12451 | 0.814923 | -0.15279 | 0.878563 | 0.991583 |
| Rbp | 16259.39 | -0.07916 | 0.546992 | -0.14472 | 0.884928 | 0.991583 |
| lncRNA:CR | 464.5007 | 0.293225 | 1.996949 | 0.146836 | 0.883261 | 0.991583 |
| CG43222 | 783.726 | -0.15294 | 1.103037 | -0.13865 | 0.889727 | 0.991583 |
| CG43324 | 11221.87 | 0.06857 | 0.483011 | 0.141963 | 0.887109 | 0.991583 |
| CG13532 | 2266.23 | 0.141062 | 0.937684 | 0.150437 | 0.88042 | 0.991583 |
| jbug | 762.0994 | 0.163195 | 1.181723 | 0.138099 | 0.890162 | 0.991583 |
| vir | 223.7059 | 0.33686 | 2.367131 | 0.142307 | 0.886837 | 0.991583 |
| Galphas | 7877.408 | 0.078073 | 0.519709 | 0.150225 | 0.880587 | 0.991583 |
| CG9380 | 151.3732 | 0.293411 | 2.092184 | 0.140241 | 0.888469 | 0.991583 |
| MED30 | 423.686 | 0.249375 | 1.748 | 0.142663 | 0.886556 | 0.991583 |
| lncRNA:CR | 68.26583 | 0.598868 | 3.834367 | 0.156184 | 0.875888 | 0.991583 |
| gpp | 5484.659 | 0.084375 | 0.603494 | 0.139811 | 0.88881 | 0.991583 |
| CG43902 | 1263.004 | -0.14734 | 0.973365 | -0.15137 | 0.879682 | 0.991583 |
| lncRNA:CR | 245.2738 | 0.390612 | 2.844491 | 0.137322 | 0.890776 | 0.991583 |
| asRNA:CR4 | 65.25641 | 0.484499 | 3.048263 | 0.158943 | 0.873714 | 0.991583 |
| lncRNA:CR | 111.3185 | -0.50183 | 3.496048 | -0.14354 | 0.885861 | 0.991583 |
| PIG-V | 403.4251 | -0.30061 | 2.201486 | -0.13655 | 0.891387 | 0.991583 |
| CG44422 | 1870.815 | 0.127587 | 0.835185 | 0.152765 | 0.878584 | 0.991583 |
| Vamp7 | 3117.256 | 0.114049 | 0.819117 | 0.139234 | 0.889265 | 0.991583 |
| lncRNA:TS | 117.1958 | -0.52589 | 3.71453 | -0.14158 | 0.887416 | 0.991583 |
| pre-rRNA:( | 74.04364 | -0.52684 | 3.362005 | -0.1567 | 0.875478 | 0.991583 |
| Tmc | 89.63613 | 0.595222 | 3.920835 | 0.15181 | 0.879337 | 0.991583 |
| lola | 26827.4 | 0.060266 | 0.431151 | 0.13978 | 0.888834 | 0.991583 |
| caz | 817.8847 | -0.16677 | 1.057984 | -0.15763 | 0.874746 | 0.991583 |
| flz | 177.2091 | -0.48332 | 3.062456 | -0.15782 | 0.874599 | 0.991583 |
| Non2 | 2478.627 | 0.10093 | 0.725234 | 0.139168 | 0.889317 | 0.991583 |
| Rdh | 3102.194 | -0.08625 | 0.629874 | -0.13694 | 0.891081 | 0.991583 |
| CG14997 | 1347.358 | 0.141109 | 0.903411 | 0.156196 | 0.875879 | 0.991583 |
| dsh | 238.4177 | -0.26649 | 1.736676 | -0.15345 | 0.878046 | 0.991583 |
| Sk1 | 2091.656 | 0.119461 | 0.810252 | 0.147436 | 0.882788 | 0.991583 |
| Usp7 | 3595.152 | 0.112687 | 0.724463 | 0.155546 | 0.876391 | 0.991583 |
| Gmap | 430.627 | -0.21114 | 1.537087 | -0.13737 | 0.890741 | 0.991583 |
| Mvd | 314.5302 | -0.2948 | 1.969772 | -0.14966 | 0.881033 | 0.991583 |
| shi | 24198.15 | 0.067296 | 0.461693 | 0.145759 | 0.884111 | 0.991583 |
| Stim | 3418.054 | 0.085811 | 0.627729 | 0.136701 | 0.891267 | 0.991583 |
| SWIP | 123.3909 | -0.54882 | 3.648637 | -0.15042 | 0.880436 | 0.991583 |
| CalpC | 176.6338 | 0.457036 | 2.879136 | 0.158741 | 0.873873 | 0.991583 |
| Sap30 | 1070.135 | -0.17957 | 1.171977 | -0.15322 | 0.878228 | 0.991583 |

| | | | | | | |
|---|---|---|---|---|---|---|
| CG8010 | 339.4257 | 0.280258 | 1.830043 | 0.153143 | 0.878286 | 0.991583 |
| THADA | 222.7057 | -0.32699 | 2.133844 | -0.15324 | 0.878208 | 0.991583 |
| GNBP1 | 217.955 | 0.288233 | 1.993491 | 0.144587 | 0.885037 | 0.991583 |
| Sgf11 | 414.7797 | -0.25831 | 1.885784 | -0.13698 | 0.891049 | 0.991583 |
| Arts | 318.0524 | -0.24153 | 1.647746 | -0.14658 | 0.883464 | 0.991583 |
| Sf3a2 | 710.3976 | 0.181346 | 1.257484 | 0.144214 | 0.885332 | 0.991583 |
| RpS9 | 1703.102 | 0.117913 | 0.820789 | 0.143658 | 0.885771 | 0.991583 |
| Hsp23 | 2334.468 | 0.110439 | 0.7575 | 0.145794 | 0.884084 | 0.991583 |
| eIF4E1 | 15535.72 | -0.07642 | 0.502198 | -0.15216 | 0.879057 | 0.991583 |
| exex | 167.9754 | -0.56083 | 3.668087 | -0.15289 | 0.878482 | 0.991583 |
| vito | 76.9369 | -0.52842 | 3.416013 | -0.15469 | 0.877067 | 0.991583 |
| Dd | 3454.673 | -0.09671 | 0.627183 | -0.1542 | 0.877455 | 0.991583 |
| CG9643 | 296.4149 | 0.357314 | 2.586468 | 0.138147 | 0.890124 | 0.991583 |
| Bacc | 25350.5 | -0.05764 | 0.403551 | -0.14283 | 0.886422 | 0.991583 |
| RFeSP | 21257.31 | 0.068776 | 0.437051 | 0.157365 | 0.874958 | 0.991583 |
| CG31935 | 2955.296 | -0.10315 | 0.654957 | -0.1575 | 0.874853 | 0.991583 |
| Ctr1B | 345.9113 | 0.225077 | 1.653947 | 0.136085 | 0.891754 | 0.991602 |
| lncRNA:CR | 274.993 | 0.314676 | 2.31173 | 0.136121 | 0.891725 | 0.991602 |
| kud | 706.5866 | -0.15047 | 1.107163 | -0.1359 | 0.891898 | 0.991602 |
| CG31689 | 667.1853 | -0.16204 | 1.191126 | -0.13604 | 0.891793 | 0.991602 |
| Npc1a | 2682.145 | 0.110208 | 0.814855 | 0.135249 | 0.892415 | 0.991627 |
| CG31142 | 150.0416 | -0.40656 | 2.999051 | -0.13556 | 0.892167 | 0.991627 |
| CG42495 | 267.5313 | 0.279217 | 2.063139 | 0.135336 | 0.892346 | 0.991627 |
| shtd | 467.2813 | 0.358287 | 2.645367 | 0.135439 | 0.892265 | 0.991627 |
| RpL7A | 14273.58 | -0.06298 | 0.466539 | -0.135 | 0.892615 | 0.991713 |
| CG1116 | 852.4523 | -0.15205 | 1.150584 | -0.13215 | 0.894865 | 0.991751 |
| stc | 911.4923 | -0.15156 | 1.125182 | -0.1347 | 0.892849 | 0.991751 |
| Ibf1 | 495.2381 | 0.235764 | 1.769297 | 0.133253 | 0.893993 | 0.991751 |
| Rbf2 | 119.8487 | -0.43438 | 3.299974 | -0.13163 | 0.895277 | 0.991751 |
| ATPsynD | 25928.63 | -0.05312 | 0.406558 | -0.13065 | 0.896052 | 0.991751 |
| CG11873 | 8048.455 | -0.06405 | 0.49061 | -0.13055 | 0.896134 | 0.991751 |
| kune | 2537.957 | 0.102136 | 0.772716 | 0.132178 | 0.894844 | 0.991751 |
| CG16799 | 2381.956 | -0.09373 | 0.721295 | -0.12995 | 0.896604 | 0.991751 |
| Fak | 3208.594 | 0.08634 | 0.650803 | 0.132668 | 0.894456 | 0.991751 |
| CG17982 | 434.327 | -0.24197 | 1.8557 | -0.13039 | 0.896256 | 0.991751 |
| Bx42 | 1060.269 | -0.12952 | 0.994537 | -0.13023 | 0.896383 | 0.991751 |
| CG7997 | 171.498 | 0.510729 | 3.813195 | 0.133937 | 0.893452 | 0.991751 |
| S2P | 856.1794 | 0.236374 | 1.779225 | 0.132852 | 0.89431 | 0.991751 |
| CNT2 | 133.0804 | -0.45258 | 3.436093 | -0.13171 | 0.895211 | 0.991751 |
| Duba | 1345.484 | -0.134 | 1.030968 | -0.12998 | 0.896585 | 0.991751 |
| CR18228 | 2653.172 | 0.089346 | 0.683266 | 0.130763 | 0.895963 | 0.991751 |
| CG17168 | 1127.708 | -0.14034 | 1.07057 | -0.13109 | 0.895706 | 0.991751 |
| Nlg3 | 5606.743 | -0.07409 | 0.552 | -0.13423 | 0.893223 | 0.991751 |
| CG4218 | 116.8018 | 0.521498 | 3.879701 | 0.134417 | 0.893073 | 0.991751 |
| Ndae1 | 2865.205 | -0.08761 | 0.653316 | -0.1341 | 0.893324 | 0.991751 |
| RpS26 | 26193.5 | 0.055502 | 0.421319 | 0.131734 | 0.895194 | 0.991751 |
| SERCA | 6459.054 | 0.074667 | 0.560681 | 0.133173 | 0.894057 | 0.991751 |
| CG3565 | 66.67185 | -0.51536 | 3.948103 | -0.13053 | 0.896145 | 0.991751 |
| lncRNA:CR | 222.8637 | 0.261871 | 2.009651 | 0.130307 | 0.896324 | 0.991751 |
| CG44195 | 214.04 | 0.2714 | 2.063898 | 0.131499 | 0.895381 | 0.991751 |

| | | | | | | |
|---|---|---|---|---|---|---|
| asRNA:CR4 | 236.5355 | 0.388234 | 2.91617 | 0.133131 | 0.89409 | 0.991751 |
| asRNA:CR4 | 63.23783 | -0.51532 | 3.94812 | -0.13052 | 0.896152 | 0.991751 |
| asRNA:CR4 | 82.22047 | -0.50336 | 3.834344 | -0.13128 | 0.895556 | 0.991751 |
| eEF5 | 7834.083 | -0.06902 | 0.515759 | -0.13382 | 0.893549 | 0.991751 |
| Vav | 3961.032 | -0.0893 | 0.667591 | -0.13377 | 0.893585 | 0.991751 |
| CG6083 | 1455.026 | -0.12654 | 0.955714 | -0.1324 | 0.894668 | 0.991751 |
| CG14341 | 540.7309 | -0.23008 | 1.722851 | -0.13354 | 0.893763 | 0.991751 |
| LysRS | 1005.899 | 0.132044 | 1.024231 | 0.12892 | 0.897421 | 0.991834 |
| eEFSec | 377.3471 | -0.35649 | 2.76425 | -0.12896 | 0.897386 | 0.991834 |
| lncRNA:CR | 98.34661 | 0.366638 | 2.840587 | 0.129071 | 0.897301 | 0.991834 |
| Vha16-1 | 37653.15 | 0.055644 | 0.431396 | 0.128985 | 0.897369 | 0.991834 |
| CG5577 | 283.2672 | 0.248444 | 1.922247 | 0.129247 | 0.897162 | 0.991834 |
| CTCF | 620.1841 | 0.222007 | 1.718606 | 0.129179 | 0.897216 | 0.991834 |
| CG5131 | 1062.278 | -0.19496 | 1.517765 | -0.12845 | 0.89779 | 0.992106 |
| Mybbp1A | 181.338 | 0.283427 | 2.326523 | 0.121824 | 0.903038 | 0.99232 |
| CG3719 | 2540.686 | 0.084612 | 0.714337 | 0.118448 | 0.905713 | 0.99232 |
| CG11596 | 169.0961 | -0.29063 | 2.356496 | -0.12333 | 0.901843 | 0.99232 |
| CG4199 | 1022.258 | 0.1235 | 1.26262 | 0.097812 | 0.922081 | 0.99232 |
| CG2865 | 1622.513 | -0.09031 | 0.854498 | -0.10568 | 0.915835 | 0.99232 |
| boi | 124.0712 | 0.368036 | 3.835604 | 0.095952 | 0.923558 | 0.99232 |
| fs(1)Yb | 117.1318 | -0.33262 | 3.101203 | -0.10725 | 0.914587 | 0.99232 |
| CG32767 | 7023.539 | 0.05867 | 0.534992 | 0.109665 | 0.912675 | 0.99232 |
| Eaat2 | 13594.43 | 0.050186 | 0.55048 | 0.091168 | 0.927359 | 0.99232 |
| Gsc | 130.3772 | 0.400366 | 3.685893 | 0.108621 | 0.913503 | 0.99232 |
| CG5955 | 4485.128 | -0.06015 | 0.620045 | -0.09702 | 0.922714 | 0.99232 |
| ND-39 | 8985.963 | 0.057422 | 0.52642 | 0.10908 | 0.913139 | 0.99232 |
| CG10508 | 7975.705 | -0.0497 | 0.509557 | -0.09754 | 0.922299 | 0.99232 |
| IleRS | 1070.633 | 0.137697 | 1.257251 | 0.109523 | 0.912788 | 0.99232 |
| dsf | 84.44041 | -0.42142 | 3.951159 | -0.10666 | 0.91506 | 0.99232 |
| Tig | 1049.839 | -0.16807 | 1.358139 | -0.12375 | 0.901513 | 0.99232 |
| Hmgcl | 93.47468 | 0.392083 | 3.837599 | 0.102169 | 0.918623 | 0.99232 |
| Pcp | 180.6485 | 0.407384 | 3.361381 | 0.121195 | 0.903536 | 0.99232 |
| Rab30 | 948.6418 | 0.16091 | 1.559776 | 0.103162 | 0.917834 | 0.99232 |
| CG5181 | 998.2097 | -0.13618 | 1.068978 | -0.1274 | 0.898626 | 0.99232 |
| CG8460 | 661.9915 | 0.176894 | 1.40265 | 0.126114 | 0.899642 | 0.99232 |
| PAPLA1 | 1766.763 | -0.09505 | 0.861162 | -0.11038 | 0.912109 | 0.99232 |
| Rcd4 | 427.3817 | -0.17446 | 2.024814 | -0.08616 | 0.931338 | 0.99232 |
| Tsp29Fb | 141.3525 | 0.473734 | 3.833481 | 0.123578 | 0.901649 | 0.99232 |
| eEF1delta | 1416.78 | -0.09167 | 0.851912 | -0.10761 | 0.914305 | 0.99232 |
| nmd | 1314.844 | -0.10866 | 0.92312 | -0.11771 | 0.906301 | 0.99232 |
| trk | 114.4479 | 0.304269 | 2.769564 | 0.109862 | 0.912519 | 0.99232 |
| kek2 | 3264.925 | 0.065915 | 0.68769 | 0.09585 | 0.92364 | 0.99232 |
| crol | 7658.37 | 0.048554 | 0.55212 | 0.087941 | 0.929924 | 0.99232 |
| CG6785 | 77.79686 | 0.477521 | 3.859312 | 0.123732 | 0.901527 | 0.99232 |
| eRF3 | 483.9759 | -0.16502 | 1.790914 | -0.09214 | 0.926583 | 0.99232 |
| ND-B17 | 7472.059 | 0.049142 | 0.492307 | 0.09982 | 0.920487 | 0.99232 |
| Tsen2 | 102.2739 | 0.461176 | 3.833871 | 0.12029 | 0.904254 | 0.99232 |
| mEFTs | 2171.101 | -0.06849 | 0.761071 | -0.08999 | 0.928295 | 0.99232 |
| Arr1 | 107.1821 | 0.392141 | 3.833646 | 0.102289 | 0.918527 | 0.99232 |
| Socs36E | 2185.111 | -0.06858 | 0.797473 | -0.08599 | 0.931472 | 0.99232 |

| | | | | | | |
|---|---|---|---|---|---|---|
| tup | 742.7892 | -0.16393 | 1.937669 | -0.0846 | 0.932577 | 0.99232 |
| Cen | 1897.724 | -0.10778 | 1.088908 | -0.09898 | 0.921155 | 0.99232 |
| CG31677 | 95.86565 | 0.381178 | 3.833807 | 0.099425 | 0.9208 | 0.99232 |
| alpha-Est1 | 101.2451 | 0.34785 | 3.833695 | 0.090735 | 0.927703 | 0.99232 |
| e(y)2b | 78.15769 | 0.36367 | 3.833899 | 0.094856 | 0.924429 | 0.99232 |
| Scr | 416.4562 | 0.158847 | 1.394658 | 0.113897 | 0.909319 | 0.99232 |
| pyd3 | 1348.56 | -0.13 | 1.115845 | -0.1165 | 0.907253 | 0.99232 |
| PolI | 1199.69 | -0.12326 | 1.348273 | -0.09142 | 0.927157 | 0.99232 |
| CG2747 | 3719.463 | -0.06707 | 0.666193 | -0.10068 | 0.919806 | 0.99232 |
| CG11980 | 893.2793 | 0.111853 | 1.168023 | 0.095762 | 0.923709 | 0.99232 |
| TrpRS | 605.9005 | -0.21556 | 2.059815 | -0.10465 | 0.916653 | 0.99232 |
| GstZ1 | 198.2183 | -0.37801 | 3.153453 | -0.11987 | 0.904585 | 0.99232 |
| Unc-115a | 1901.972 | 0.093986 | 0.79543 | 0.118157 | 0.905943 | 0.99232 |
| eca | 1190.39 | 0.095957 | 0.938621 | 0.102232 | 0.918572 | 0.99232 |
| Tsp86D | 1029.776 | 0.163201 | 1.40161 | 0.116438 | 0.907305 | 0.99232 |
| Dph3 | 463.1007 | 0.138638 | 1.376801 | 0.100696 | 0.919792 | 0.99232 |
| SdhC | 24001.05 | 0.050502 | 0.425897 | 0.118578 | 0.90561 | 0.99232 |
| DCAF12 | 127.9526 | -0.35103 | 3.666808 | -0.09573 | 0.923734 | 0.99232 |
| Sccpdh2 | 860.6754 | 0.129985 | 1.242137 | 0.104646 | 0.916657 | 0.99232 |
| Pp1-87B | 8055.138 | -0.04456 | 0.497984 | -0.08948 | 0.928698 | 0.99232 |
| CG5509 | 138.983 | 0.371223 | 3.667879 | 0.101209 | 0.919384 | 0.99232 |
| Dtg | 107.737 | 0.485416 | 3.940717 | 0.12318 | 0.901965 | 0.99232 |
| CG5961 | 1712.996 | -0.09973 | 1.046182 | -0.09533 | 0.924054 | 0.99232 |
| GILT1 | 1343.931 | -0.08981 | 0.945377 | -0.095 | 0.924319 | 0.99232 |
| UQCR-C1 | 17924.72 | 0.046614 | 0.444466 | 0.104877 | 0.916473 | 0.99232 |
| CG6966 | 4195.395 | 0.061683 | 0.704697 | 0.087532 | 0.930249 | 0.99232 |
| CG6171 | 307.9114 | -0.14418 | 1.598905 | -0.09018 | 0.928148 | 0.99232 |
| CG6218 | 5986.178 | -0.05317 | 0.560235 | -0.09491 | 0.924386 | 0.99232 |
| Rh6 | 107.8051 | 0.377588 | 3.834557 | 0.09847 | 0.921559 | 0.99232 |
| asun | 616.1062 | 0.203697 | 1.877415 | 0.108498 | 0.9136 | 0.99232 |
| CG6126 | 344.9233 | -0.22237 | 2.483736 | -0.08953 | 0.928661 | 0.99232 |
| CCT3 | 332.1389 | 0.176954 | 1.522466 | 0.116229 | 0.907471 | 0.99232 |
| CG3995 | 256.6352 | -0.2434 | 2.444074 | -0.09959 | 0.920671 | 0.99232 |
| CG31360 | 1014.806 | -0.10424 | 1.042727 | -0.09996 | 0.920372 | 0.99232 |
| CG5823 | 2299.24 | -0.07844 | 0.822663 | -0.09534 | 0.924041 | 0.99232 |
| CG18600 | 124.4816 | -0.34309 | 3.835414 | -0.08945 | 0.928723 | 0.99232 |
| P5cr | 1414.569 | -0.08018 | 0.900249 | -0.08907 | 0.929027 | 0.99232 |
| CG6195 | 1070.86 | -0.18848 | 1.50804 | -0.12498 | 0.900537 | 0.99232 |
| GABA-B-R2 | 5452.175 | 0.06936 | 0.572846 | 0.121079 | 0.903628 | 0.99232 |
| CG7054 | 169.0087 | 0.372286 | 3.087655 | 0.120572 | 0.90403 | 0.99232 |
| CG31365 | 89.31969 | 0.395281 | 3.29034 | 0.120134 | 0.904377 | 0.99232 |
| GILT2 | 1338.542 | 0.085295 | 0.939658 | 0.090772 | 0.927674 | 0.99232 |
| CG5854 | 162.3446 | 0.337536 | 3.778936 | 0.08932 | 0.928827 | 0.99232 |
| Rab7 | 9935.593 | -0.04217 | 0.482397 | -0.08742 | 0.930341 | 0.99232 |
| Spase22-2 | 1769.385 | -0.08688 | 0.806653 | -0.10771 | 0.914226 | 0.99232 |
| EMC6 | 3345.02 | -0.06862 | 0.617715 | -0.11109 | 0.911544 | 0.99232 |
| dan | 77.16645 | -0.39393 | 3.833886 | -0.10275 | 0.918162 | 0.99232 |
| CG10562 | 768.5116 | 0.160643 | 1.620457 | 0.099135 | 0.921031 | 0.99232 |
| Ets97D | 110.8948 | 0.283075 | 3.247414 | 0.087169 | 0.930537 | 0.99232 |
| CG5934 | 2657.373 | 0.070674 | 0.811516 | 0.087089 | 0.930601 | 0.99232 |

| | | | | | | |
|---|---|---|---|---|---|---|
| dmrt99B | 121.4729 | 0.349086 | 3.727041 | 0.093663 | 0.925377 | 0.99232 |
| Pcd | 1390.611 | 0.073952 | 0.869612 | 0.08504 | 0.93223 | 0.99232 |
| CG1542 | 301.1217 | -0.25866 | 2.466777 | -0.10486 | 0.916489 | 0.99232 |
| CG11539 | 110.9449 | 0.362341 | 3.276139 | 0.1106 | 0.911933 | 0.99232 |
| Ttc7 | 2952.536 | -0.07965 | 0.727746 | -0.10944 | 0.91285 | 0.99232 |
| FAM21 | 613.4874 | 0.113272 | 1.330574 | 0.08513 | 0.932158 | 0.99232 |
| CG13442 | 146.6763 | 0.230696 | 2.698093 | 0.085503 | 0.931861 | 0.99232 |
| CG11180 | 374.8631 | -0.19908 | 2.268965 | -0.08774 | 0.930085 | 0.99232 |
| Nnf1a | 138.7285 | -0.37649 | 3.181461 | -0.11834 | 0.9058 | 0.99232 |
| sano | 785.4091 | -0.14891 | 1.621649 | -0.09182 | 0.926838 | 0.99232 |
| CG4558 | 435.8574 | -0.19454 | 1.758332 | -0.11064 | 0.911904 | 0.99232 |
| CG8300 | 386.266 | -0.17738 | 1.826045 | -0.09714 | 0.922615 | 0.99232 |
| CG9657 | 28004.48 | 0.047216 | 0.516034 | 0.091499 | 0.927096 | 0.99232 |
| Atg5 | 2189.291 | -0.08004 | 0.718201 | -0.11145 | 0.911262 | 0.99232 |
| Smox | 4811.115 | -0.07068 | 0.595852 | -0.11861 | 0.905582 | 0.99232 |
| CG33181 | 4421.835 | -0.05341 | 0.581223 | -0.09189 | 0.926785 | 0.99232 |
| Ogg1 | 64.01919 | 0.34772 | 3.838494 | 0.090588 | 0.92782 | 0.99232 |
| CG11294 | 115.0496 | 0.364267 | 3.836062 | 0.094959 | 0.924348 | 0.99232 |
| CG17754 | 713.735 | -0.14664 | 1.532548 | -0.09569 | 0.92377 | 0.99232 |
| Naxe | 206.4432 | 0.291722 | 2.940828 | 0.099197 | 0.920982 | 0.99232 |
| Treh | 30998.97 | -0.06263 | 0.535409 | -0.11698 | 0.90688 | 0.99232 |
| Xbp1 | 10052.38 | 0.043485 | 0.477518 | 0.091066 | 0.92744 | 0.99232 |
| Acox57D-c | 254.4238 | -0.37832 | 2.977078 | -0.12708 | 0.898879 | 0.99232 |
| CG15673 | 321.597 | -0.34462 | 2.710751 | -0.12713 | 0.898838 | 0.99232 |
| TAF1C-like | 298.9606 | -0.19915 | 1.780801 | -0.11183 | 0.910957 | 0.99232 |
| Sod2 | 8351.821 | -0.04515 | 0.504423 | -0.08952 | 0.928671 | 0.99232 |
| mrj | 3378.478 | -0.05856 | 0.636599 | -0.092 | 0.926702 | 0.99232 |
| CG8187 | 826.266 | -0.11863 | 1.374255 | -0.08632 | 0.93121 | 0.99232 |
| CG10139 | 1214.886 | -0.12578 | 1.157048 | -0.10871 | 0.913435 | 0.99232 |
| SMC2 | 204.7955 | 0.327631 | 3.243694 | 0.101006 | 0.919546 | 0.99232 |
| Dh44-R1 | 419.0895 | -0.24987 | 2.089887 | -0.11956 | 0.904832 | 0.99232 |
| CG13018 | 197.7088 | 0.20951 | 2.17995 | 0.096108 | 0.923435 | 0.99232 |
| CG17739 | 2382.442 | -0.09171 | 1.066689 | -0.08598 | 0.931484 | 0.99232 |
| SmD3 | 120.9734 | 0.455158 | 3.833727 | 0.118725 | 0.905493 | 0.99232 |
| CG9005 | 4756.863 | 0.063992 | 0.576109 | 0.111076 | 0.911556 | 0.99232 |
| Ggamma1 | 89279.56 | 0.032404 | 0.378056 | 0.085712 | 0.931696 | 0.99232 |
| Gle1 | 966.9732 | 0.112249 | 1.194377 | 0.093981 | 0.925124 | 0.99232 |
| Rpt1 | 1850.74 | 0.110926 | 1.01278 | 0.109526 | 0.912785 | 0.99232 |
| CG1603 | 1252.958 | -0.11415 | 1.108517 | -0.10298 | 0.91798 | 0.99232 |
| myo | 9102.865 | -0.05579 | 0.512284 | -0.1089 | 0.913281 | 0.99232 |
| CG9272 | 281.6576 | 0.355092 | 2.951913 | 0.120292 | 0.904252 | 0.99232 |
| CG9451 | 2712.924 | 0.137438 | 1.416686 | 0.097013 | 0.922716 | 0.99232 |
| dpr18 | 8296.601 | -0.06513 | 0.586692 | -0.11102 | 0.9116 | 0.99232 |
| Cyp28d1 | 16267.93 | -0.05465 | 0.544764 | -0.10032 | 0.920088 | 0.99232 |
| ref(2)P | 7252.64 | 0.069547 | 0.797912 | 0.087161 | 0.930543 | 0.99232 |
| 7B2 | 27263.64 | 0.046542 | 0.407674 | 0.114164 | 0.909108 | 0.99232 |
| CG14667 | 1557.452 | -0.08505 | 0.964118 | -0.08821 | 0.929709 | 0.99232 |
| hig | 26720.82 | -0.03947 | 0.397488 | -0.0993 | 0.920902 | 0.99232 |
| cpx | 111504.7 | 0.03825 | 0.413907 | 0.092413 | 0.92637 | 0.99232 |
| CG4630 | 106.8774 | -0.35828 | 3.839484 | -0.09331 | 0.925654 | 0.99232 |

| | | | | | | |
|---|---|---|---|---|---|---|
| HDAC4 | 2831.116 | -0.07116 | 0.685077 | -0.10387 | 0.917274 | 0.99232 |
| gw | 18911.16 | 0.039384 | 0.439956 | 0.089519 | 0.928669 | 0.99232 |
| CG2225 | 2424.07 | -0.06441 | 0.738925 | -0.08716 | 0.930542 | 0.99232 |
| ND-ASHI | 10234.87 | -0.04927 | 0.458283 | -0.10751 | 0.914384 | 0.99232 |
| hth | 31371.35 | -0.04474 | 0.442745 | -0.10104 | 0.919516 | 0.99232 |
| Neb-cGP | 14798.52 | 0.051814 | 0.447191 | 0.115865 | 0.90776 | 0.99232 |
| CG4822 | 432.1501 | 0.229503 | 1.905257 | 0.120458 | 0.904121 | 0.99232 |
| CG32202 | 634.8528 | 0.116661 | 1.161042 | 0.10048 | 0.919963 | 0.99232 |
| CG4313 | 114.4496 | -0.37389 | 3.079672 | -0.12141 | 0.903369 | 0.99232 |
| escl | 274.1542 | -0.20991 | 2.184014 | -0.09611 | 0.923431 | 0.99232 |
| disco | 376.9242 | 0.161913 | 1.852655 | 0.087395 | 0.930358 | 0.99232 |
| su(w[a]) | 7023.789 | -0.04822 | 0.557607 | -0.08648 | 0.931084 | 0.99232 |
| CG9837 | 91.85609 | -0.35183 | 3.834005 | -0.09177 | 0.926884 | 0.99232 |
| Adh | 238.3714 | 0.186494 | 1.781275 | 0.104697 | 0.916616 | 0.99232 |
| CG14408 | 2557.537 | -0.08168 | 0.910965 | -0.08966 | 0.928558 | 0.99232 |
| lncRNA:CR | 76.45461 | 0.390669 | 3.833925 | 0.101898 | 0.918838 | 0.99232 |
| mt:tRNA:C | 93.40189 | -0.27932 | 2.543389 | -0.10982 | 0.912549 | 0.99232 |
| mt:tRNA:G | 132.4312 | 0.28065 | 2.192356 | 0.128013 | 0.898139 | 0.99232 |
| CG7878 | 1437.576 | 0.140484 | 1.211398 | 0.115968 | 0.907678 | 0.99232 |
| CG34134 | 184.9517 | -0.24422 | 2.480959 | -0.09844 | 0.921585 | 0.99232 |
| eIF4B | 1070.595 | 0.09788 | 0.948063 | 0.103242 | 0.917771 | 0.99232 |
| CG34213 | 139.1237 | 0.252696 | 2.772493 | 0.091144 | 0.927378 | 0.99232 |
| smal | 2805.048 | 0.063178 | 0.731192 | 0.086405 | 0.931145 | 0.99232 |
| 28SrRNA-P | 4879.084 | -0.05036 | 0.575446 | -0.08751 | 0.930263 | 0.99232 |
| snoRNA:M | 70.18156 | -0.3287 | 3.834617 | -0.08572 | 0.931689 | 0.99232 |
| Ino80 | 101.567 | 0.364214 | 3.23731 | 0.112505 | 0.910423 | 0.99232 |
| l(3)L1231 | 4601.889 | 0.054296 | 0.592103 | 0.091701 | 0.926936 | 0.99232 |
| Pcmt | 2180.043 | -0.08173 | 0.777857 | -0.10507 | 0.916323 | 0.99232 |
| stmA | 9166.299 | -0.07237 | 0.625324 | -0.11574 | 0.907862 | 0.99232 |
| puc | 2060.784 | -0.08685 | 0.82337 | -0.10548 | 0.915997 | 0.99232 |
| futsch | 33542.28 | 0.047721 | 0.424532 | 0.112408 | 0.9105 | 0.99232 |
| alpha-Man | 4776.622 | 0.061464 | 0.59429 | 0.103424 | 0.917627 | 0.99232 |
| corn | 3898.02 | 0.073533 | 0.630695 | 0.116591 | 0.907184 | 0.99232 |
| CG42450 | 10101.15 | 0.041417 | 0.468015 | 0.088496 | 0.929482 | 0.99232 |
| CG42564 | 254.048 | 0.286034 | 2.529535 | 0.113078 | 0.909969 | 0.99232 |
| Atg1 | 4261.731 | -0.06498 | 0.578129 | -0.11241 | 0.910502 | 0.99232 |
| CG42656 | 170.7304 | -0.3441 | 3.794599 | -0.09068 | 0.927745 | 0.99232 |
| ReepA | 7442.676 | 0.053529 | 0.558227 | 0.095892 | 0.923607 | 0.99232 |
| raskol | 2458.858 | 0.06775 | 0.722799 | 0.093732 | 0.925322 | 0.99232 |
| pdm3 | 4062.208 | 0.06067 | 0.653492 | 0.09284 | 0.926031 | 0.99232 |
| lncRNA:CR | 128.3869 | 0.406129 | 3.724274 | 0.109049 | 0.913164 | 0.99232 |
| sdt | 3726.019 | 0.075191 | 0.603388 | 0.124615 | 0.900828 | 0.99232 |
| l(2)35Be | 1664.143 | 0.108532 | 1.002317 | 0.108282 | 0.913772 | 0.99232 |
| Sec23 | 3168.905 | -0.0771 | 0.800922 | -0.09627 | 0.923307 | 0.99232 |
| VhaAC45 | 15242.14 | 0.050239 | 0.449018 | 0.111886 | 0.910914 | 0.99232 |
| CG43191 | 281.5603 | 0.152294 | 1.620383 | 0.093986 | 0.92512 | 0.99232 |
| CG43322 | 259.6856 | 0.278369 | 3.026791 | 0.091968 | 0.926723 | 0.99232 |
| DnaJ-1 | 5562.403 | 0.067116 | 0.577503 | 0.116217 | 0.90748 | 0.99232 |
| Mitf | 5071.41 | 0.064976 | 0.556932 | 0.116668 | 0.907123 | 0.99232 |
| fd59A | 110.7243 | 0.374897 | 3.764254 | 0.099594 | 0.920667 | 0.99232 |

| | | | | | | |
|---|---|---|---|---|---|---|
| Cdk9 | 390.9311 | 0.161993 | 1.760156 | 0.092033 | 0.926672 | 0.99232 |
| CG9849 | 1272.423 | -0.09148 | 0.891981 | -0.10256 | 0.918316 | 0.99232 |
| CG16787 | 1010.707 | 0.126577 | 1.046263 | 0.12098 | 0.903707 | 0.99232 |
| CG4806 | 309.5938 | 0.210318 | 1.723916 | 0.122 | 0.902899 | 0.99232 |
| CG3776 | 2260.161 | -0.07958 | 0.77553 | -0.10262 | 0.918267 | 0.99232 |
| key | 1404.418 | -0.11224 | 0.880164 | -0.12752 | 0.898529 | 0.99232 |
| wac | 322.3683 | -0.20575 | 1.98208 | -0.10381 | 0.917323 | 0.99232 |
| Hipk | 16369.51 | 0.048682 | 0.430973 | 0.112959 | 0.910063 | 0.99232 |
| Atac3 | 269.1749 | -0.2581 | 3.002368 | -0.08597 | 0.931494 | 0.99232 |
| lncRNA:CR | 96.08929 | -0.34683 | 3.836068 | -0.09041 | 0.927959 | 0.99232 |
| lncRNA:CR | 244.319 | -0.20141 | 1.755089 | -0.11476 | 0.908638 | 0.99232 |
| lncRNA:CR | 88.04109 | -0.3272 | 3.738997 | -0.08751 | 0.930266 | 0.99232 |
| lncRNA:CR | 173.8976 | 0.222141 | 2.594542 | 0.085619 | 0.931769 | 0.99232 |
| asRNA:CR4 | 217.0328 | 0.260812 | 2.877179 | 0.090648 | 0.927772 | 0.99232 |
| lncRNA:CR | 132.4239 | 0.235929 | 2.794423 | 0.084429 | 0.932716 | 0.99232 |
| lncRNA:CR | 83.09523 | 0.367967 | 3.83442 | 0.095964 | 0.923549 | 0.99232 |
| Dscam2 | 16654.81 | 0.039952 | 0.435292 | 0.091783 | 0.92687 | 0.99232 |
| lncRNA:CR | 133.1244 | 0.361301 | 3.127823 | 0.115512 | 0.908039 | 0.99232 |
| lncRNA:CR | 283.032 | 0.301599 | 2.654466 | 0.113619 | 0.90954 | 0.99232 |
| lncRNA:CR | 503.0334 | 0.123941 | 1.428415 | 0.086768 | 0.930856 | 0.99232 |
| lncRNA:CR | 79.71174 | -0.38904 | 3.951298 | -0.09846 | 0.921568 | 0.99232 |
| mamo | 22918.73 | -0.0464 | 0.402505 | -0.11527 | 0.908233 | 0.99232 |
| CG32700 | 3081.849 | 0.053487 | 0.631386 | 0.084713 | 0.932489 | 0.99232 |
| Glut4EF | 23493.39 | 0.046515 | 0.406903 | 0.114314 | 0.908989 | 0.99232 |
| rn | 651.8444 | -0.15688 | 1.842702 | -0.08513 | 0.932155 | 0.99232 |
| LanB2 | 138.0889 | -0.32198 | 3.599507 | -0.08945 | 0.928724 | 0.99232 |
| 28SrRNA-F | 7308.997 | 0.057288 | 0.518629 | 0.110461 | 0.912044 | 0.99232 |
| lncRNA:CR | 72.46495 | 0.380922 | 3.835307 | 0.09932 | 0.920884 | 0.99232 |
| lncRNA:CR | 89.54874 | -0.32473 | 3.834011 | -0.0847 | 0.932503 | 0.99232 |
| lncRNA:CR | 1670.872 | 0.073845 | 0.80421 | 0.091823 | 0.926839 | 0.99232 |
| lncRNA:CR | 194.697 | -0.21851 | 1.946762 | -0.11224 | 0.910629 | 0.99232 |
| Phm | 8635.856 | -0.06314 | 0.501247 | -0.12597 | 0.899758 | 0.99232 |
| Vha26 | 31560.03 | 0.039137 | 0.449384 | 0.08709 | 0.9306 | 0.99232 |
| Letm1 | 2013.657 | 0.074985 | 0.792049 | 0.094672 | 0.924575 | 0.99232 |
| tyn | 870.9136 | -0.14499 | 1.270364 | -0.11413 | 0.909135 | 0.99232 |
| para | 44513.94 | -0.03578 | 0.383263 | -0.09334 | 0.925631 | 0.99232 |
| scro | 17191.85 | -0.05454 | 0.52999 | -0.1029 | 0.918042 | 0.99232 |
| bor | 1267.641 | 0.123874 | 1.147701 | 0.107932 | 0.914049 | 0.99232 |
| PolD2 | 70.41339 | 0.410001 | 3.842933 | 0.10669 | 0.915035 | 0.99232 |
| CG13937 | 4524.692 | -0.05535 | 0.598629 | -0.09246 | 0.926334 | 0.99232 |
| Rap1 | 5313.338 | 0.0687 | 0.554872 | 0.123813 | 0.901463 | 0.99232 |
| CG16986 | 911.3255 | 0.176352 | 1.546307 | 0.114047 | 0.9092 | 0.99232 |
| CG33234 | 499.0198 | 0.186109 | 1.783168 | 0.10437 | 0.916876 | 0.99232 |
| CG32276 | 5876.486 | 0.060405 | 0.536294 | 0.112634 | 0.910321 | 0.99232 |
| CG1316 | 6877.557 | 0.069628 | 0.622177 | 0.111911 | 0.910894 | 0.99232 |
| CG11357 | 1703.719 | 0.095243 | 0.801229 | 0.118871 | 0.905378 | 0.99232 |
| DopEcR | 14036.54 | -0.04392 | 0.437351 | -0.10043 | 0.920002 | 0.99232 |
| sesB | 138454.4 | -0.04935 | 0.390345 | -0.12644 | 0.899386 | 0.99232 |
| Rpt3 | 2833.411 | 0.072437 | 0.689515 | 0.105054 | 0.916333 | 0.99232 |
| Tango4 | 233.8955 | -0.18888 | 1.799063 | -0.10499 | 0.916385 | 0.99232 |

| | | | | | | |
|---|---|---|---|---|---|---|
| CG4318 | 126.0238 | -0.39256 | 3.843428 | -0.10214 | 0.918647 | 0.99232 |
| CG1673 | 172.1847 | -0.24536 | 2.023245 | -0.12127 | 0.903478 | 0.99232 |
| DNAlig4 | 222.4595 | 0.277897 | 2.771438 | 0.100272 | 0.920129 | 0.99232 |
| ND-B18 | 3349.975 | 0.060224 | 0.620153 | 0.097112 | 0.922637 | 0.99232 |
| CG12398 | 76.11111 | 0.37602 | 3.834854 | 0.098053 | 0.92189 | 0.99232 |
| CG9281 | 1312.755 | -0.09278 | 1.011503 | -0.09173 | 0.926915 | 0.99232 |
| MagR | 3405.384 | -0.07345 | 0.610242 | -0.12037 | 0.904191 | 0.99232 |
| CG8206 | 2657.824 | 0.083663 | 0.676957 | 0.123586 | 0.901643 | 0.99232 |
| Rpc160 | 164.2868 | -0.3238 | 2.807667 | -0.11533 | 0.908187 | 0.99232 |
| DENR | 217.6675 | -0.17269 | 1.834076 | -0.09415 | 0.924986 | 0.99232 |
| CG5162 | 194.108 | 0.367994 | 3.259481 | 0.1129 | 0.91011 | 0.99232 |
| Arp8 | 252.0352 | 0.334168 | 3.015674 | 0.11081 | 0.911767 | 0.99232 |
| CG32536 | 150.0401 | -0.25779 | 2.716145 | -0.09491 | 0.924387 | 0.99232 |
| Grip84 | 1246.554 | -0.09123 | 0.951941 | -0.09584 | 0.923647 | 0.99232 |
| mRpS14 | 1318.094 | -0.09646 | 0.865029 | -0.11151 | 0.911209 | 0.99232 |
| Rpn1 | 1240.193 | 0.091217 | 0.951691 | 0.095848 | 0.923642 | 0.99232 |
| CG7646 | 16597.35 | 0.048862 | 0.491821 | 0.099349 | 0.920861 | 0.99232 |
| Oat | 63.8807 | 0.352952 | 3.834216 | 0.092053 | 0.926656 | 0.99232 |
| Gbs-76A | 1670.21 | -0.08612 | 0.85222 | -0.10105 | 0.919508 | 0.99232 |
| CG18135 | 6534.186 | 0.05614 | 0.576406 | 0.097396 | 0.922412 | 0.99232 |
| HipHop | 166.2941 | 0.365537 | 3.468983 | 0.105373 | 0.91608 | 0.99232 |
| Dysb | 174.1023 | 0.396561 | 3.374999 | 0.117499 | 0.906464 | 0.99232 |
| anchor | 4217.266 | 0.055111 | 0.611641 | 0.090104 | 0.928204 | 0.99232 |
| a10 | 140.9046 | -0.31247 | 3.608582 | -0.08659 | 0.930997 | 0.99232 |
| CG9674 | 15242.18 | -0.05042 | 0.428894 | -0.11756 | 0.906417 | 0.99232 |
| Nplp3 | 122.7296 | 0.361947 | 3.377158 | 0.107175 | 0.91465 | 0.99232 |
| CG5151 | 2991.764 | -0.07635 | 0.659713 | -0.11573 | 0.90787 | 0.99232 |
| Cep135 | 75.50308 | 0.414057 | 3.842192 | 0.107766 | 0.914182 | 0.99232 |
| bmm | 1656.309 | 0.099562 | 1.016911 | 0.097906 | 0.922007 | 0.99232 |
| Lmx1a | 66.25414 | -0.35059 | 3.834631 | -0.09143 | 0.927152 | 0.99232 |
| CG11597 | 673.7568 | 0.183689 | 1.598638 | 0.114903 | 0.908522 | 0.99232 |
| Apt1 | 11046.84 | 0.060172 | 0.500964 | 0.120111 | 0.904395 | 0.99232 |
| CG7607 | 12053.11 | -0.04789 | 0.503905 | -0.09504 | 0.924283 | 0.99232 |
| CG7120 | 1333.65 | -0.0942 | 0.857403 | -0.10986 | 0.912518 | 0.99232 |
| CG10274 | 192.1074 | 0.37257 | 3.284909 | 0.113419 | 0.909699 | 0.99232 |
| CG13295 | 387.009 | 0.214333 | 1.98128 | 0.108179 | 0.913854 | 0.99232 |
| Ntf-2 | 4173.075 | 0.065063 | 0.654669 | 0.099384 | 0.920834 | 0.99232 |
| CG1812 | 3860.79 | 0.06537 | 0.737307 | 0.088661 | 0.929351 | 0.99232 |
| capu | 2794.614 | -0.05893 | 0.657178 | -0.08967 | 0.928553 | 0.99232 |
| drm | 138.0714 | 0.329585 | 3.261141 | 0.101064 | 0.919499 | 0.99232 |
| alpha4GT1 | 1713.29 | -0.12467 | 1.262037 | -0.09879 | 0.921308 | 0.99232 |
| Cwc25 | 407.4777 | -0.16274 | 1.473849 | -0.11042 | 0.912076 | 0.99232 |
| Tnpo-SR | 136.6145 | 0.423584 | 3.619822 | 0.117018 | 0.906846 | 0.99232 |
| ND-B17.2 | 12705.13 | 0.037353 | 0.442497 | 0.084415 | 0.932727 | 0.99232 |
| erm | 477.8505 | 0.174924 | 1.390069 | 0.125838 | 0.89986 | 0.99232 |
| Dop2R | 5383.818 | -0.05491 | 0.654981 | -0.08384 | 0.933182 | 0.992674 |
| temp | 2803.372 | -0.06545 | 0.810707 | -0.08074 | 0.93565 | 0.993718 |
| ninaD | 205.2863 | 0.15366 | 1.887573 | 0.081406 | 0.935119 | 0.993718 |
| tefu | 247.3614 | -0.19345 | 2.385832 | -0.08108 | 0.935377 | 0.993718 |
| PIG-L | 1077.349 | -0.14049 | 1.732377 | -0.0811 | 0.935366 | 0.993718 |

| | | | | | | |
|---|---|---|---|---|---|---|
| Trxr-1 | 3677.165 | -0.06671 | 0.816472 | -0.0817 | 0.934884 | 0.993718 |
| eIF4G1 | 23713.27 | 0.034723 | 0.429811 | 0.080787 | 0.935612 | 0.993718 |
| CG32695 | 149.7026 | -0.31231 | 3.833675 | -0.08146 | 0.935073 | 0.993718 |
| CG11406 | 62.47527 | 0.319995 | 3.950509 | 0.081001 | 0.935441 | 0.993718 |
| rtv | 134.6709 | 0.297731 | 3.633664 | 0.081937 | 0.934697 | 0.993718 |
| asRNA:CR4 | 196.1819 | 0.211747 | 2.579156 | 0.082099 | 0.934568 | 0.993718 |
| Hsc70Cb | 4015.734 | 0.050892 | 0.617756 | 0.082382 | 0.934343 | 0.993718 |
| CG3294 | 113.7179 | -0.27666 | 3.387516 | -0.08167 | 0.93491 | 0.993718 |
| CG2652 | 159.5852 | -0.24622 | 3.091257 | -0.07965 | 0.936516 | 0.993807 |
| CG5039 | 68.22504 | -0.30318 | 3.841616 | -0.07892 | 0.937096 | 0.993807 |
| pygo | 1511.863 | -0.07885 | 0.998454 | -0.07897 | 0.937057 | 0.993807 |
| Act42A | 1413.187 | -0.13464 | 1.702769 | -0.07907 | 0.936978 | 0.993807 |
| cpa | 1006.856 | 0.09527 | 1.198934 | 0.079462 | 0.936665 | 0.993807 |
| CG10395 | 823.9088 | 0.093997 | 1.174377 | 0.08004 | 0.936205 | 0.993807 |
| CG40485 | 85.68731 | 0.312499 | 3.950308 | 0.079107 | 0.936947 | 0.993807 |
| CG34370 | 2766.334 | 0.052434 | 0.661825 | 0.079226 | 0.936853 | 0.993807 |
| CG43843 | 99.92819 | 0.307623 | 3.845296 | 0.08 | 0.936237 | 0.993807 |
| CG12531 | 4420.515 | 0.07544 | 0.950845 | 0.07934 | 0.936762 | 0.993807 |
| GluRIA | 3843.615 | 0.04693 | 0.589732 | 0.079578 | 0.936573 | 0.993807 |
| lncRNA:CR | 112.7251 | 0.297895 | 3.787114 | 0.07866 | 0.937303 | 0.993895 |
| Pp2B-14D | 4963.338 | 0.053154 | 0.678315 | 0.078361 | 0.937541 | 0.994016 |
| CG31855 | 1594.732 | 0.06939 | 0.889368 | 0.078022 | 0.937811 | 0.994122 |
| lncRNA:CR | 263.5734 | 0.247422 | 3.175153 | 0.077924 | 0.937888 | 0.994122 |
| CG4404 | 2708.131 | 0.056811 | 0.732393 | 0.077569 | 0.938171 | 0.994159 |
| Shal | 2583.386 | -0.05939 | 0.765339 | -0.0776 | 0.938147 | 0.994159 |
| Uhg5 | 2425.203 | -0.0553 | 0.715269 | -0.07732 | 0.938371 | 0.994207 |
| gol | 5971.232 | 0.044039 | 0.570449 | 0.0772 | 0.938464 | 0.994207 |
| lncRNA:roX | 207945.6 | -0.02782 | 0.368101 | -0.07557 | 0.939763 | 0.994254 |
| RpLP1 | 18897.86 | 0.037241 | 0.486187 | 0.076598 | 0.938943 | 0.994254 |
| mRpS18B | 656.2128 | -0.1457 | 1.922174 | -0.0758 | 0.93958 | 0.994254 |
| P58IPK | 1295.181 | -0.07021 | 0.931134 | -0.0754 | 0.939893 | 0.994254 |
| TFAM | 607.9942 | -0.09367 | 1.231924 | -0.07604 | 0.939391 | 0.994254 |
| CG8321 | 2106.165 | 0.060667 | 0.80788 | 0.075094 | 0.94014 | 0.994254 |
| Crk | 9763.342 | -0.03621 | 0.483027 | -0.07497 | 0.940242 | 0.994254 |
| pros | 35578.59 | -0.03348 | 0.444752 | -0.07527 | 0.940002 | 0.994254 |
| Ac3 | 5136.986 | 0.050145 | 0.6604 | 0.075931 | 0.939474 | 0.994254 |
| Rb97D | 3330.663 | 0.059498 | 0.784357 | 0.075856 | 0.939534 | 0.994254 |
| CG34039 | 248.1529 | 0.178927 | 2.375623 | 0.075318 | 0.939962 | 0.994254 |
| Gprk1 | 11612.81 | 0.036879 | 0.479454 | 0.076919 | 0.938688 | 0.994254 |
| CG9932 | 15726.35 | 0.037448 | 0.496571 | 0.075414 | 0.939885 | 0.994254 |
| Myt1 | 618.543 | -0.12422 | 1.627394 | -0.07633 | 0.939155 | 0.994254 |
| Pfdn1 | 815.3424 | 0.085122 | 1.145694 | 0.074297 | 0.940774 | 0.994385 |
| gce | 4459.786 | 0.047291 | 0.637471 | 0.074186 | 0.940862 | 0.994385 |
| CG4101 | 120.8104 | 0.169222 | 2.280532 | 0.074203 | 0.940849 | 0.994385 |
| Taf10 | 2803.55 | -0.05711 | 0.766354 | -0.07453 | 0.940591 | 0.994385 |
| Mgat3 | 679.6658 | 0.140948 | 1.904153 | 0.074021 | 0.940994 | 0.994393 |
| CR31032 | 610.0655 | -0.16386 | 2.225682 | -0.07362 | 0.941312 | 0.994598 |
| kraken | 615.645 | -0.12987 | 1.768946 | -0.07341 | 0.941476 | 0.994641 |
| Prosalpha3 | 1957.658 | 0.056431 | 0.770462 | 0.073243 | 0.941613 | 0.994654 |
| lbm | 220.4858 | -0.16436 | 2.261838 | -0.07267 | 0.942072 | 0.994698 |

| | | | | | | |
|---|---|---|---|---|---|---|
| qkr58E-3 | 6061.282 | -0.03803 | 0.52107 | -0.07298 | 0.941819 | 0.994698 |
| 28SrRNA-F | 778.9576 | -0.07747 | 1.067536 | -0.07257 | 0.94215 | 0.994698 |
| CG1146 | 1351.451 | -0.06525 | 0.897895 | -0.07266 | 0.942073 | 0.994698 |
| CG17328 | 265.4141 | -0.16287 | 2.280733 | -0.07141 | 0.943069 | 0.994948 |
| CG31626 | 86.44298 | 0.268526 | 3.727266 | 0.072044 | 0.942567 | 0.994948 |
| Cirl | 15424.5 | 0.037595 | 0.523542 | 0.071808 | 0.942755 | 0.994948 |
| pnut | 3319.411 | 0.044673 | 0.623527 | 0.071646 | 0.942884 | 0.994948 |
| Coa7 | 2876.462 | 0.04636 | 0.649885 | 0.071336 | 0.94313 | 0.994948 |
| CG17261 | 938.9018 | -0.09769 | 1.361158 | -0.07177 | 0.942786 | 0.994948 |
| Coa8 | 2119.052 | -0.04717 | 0.732624 | -0.06439 | 0.948664 | 0.994965 |
| tko | 2700.793 | -0.04781 | 0.723064 | -0.06613 | 0.947278 | 0.994965 |
| CG3108 | 94.23146 | 0.19818 | 3.286833 | 0.060295 | 0.95192 | 0.994965 |
| Spp | 1609.45 | -0.07187 | 1.113552 | -0.06454 | 0.948542 | 0.994965 |
| ND-15 | 6857.245 | -0.03026 | 0.527148 | -0.0574 | 0.954225 | 0.994965 |
| Pitslre | 9643.363 | -0.02834 | 0.485695 | -0.05836 | 0.953464 | 0.994965 |
| CG32432 | 6301.457 | 0.031536 | 0.548426 | 0.057503 | 0.954145 | 0.994965 |
| CG33056 | 2094.123 | -0.04964 | 0.752292 | -0.06598 | 0.947395 | 0.994965 |
| CG33054 | 132.8194 | -0.1955 | 2.793297 | -0.06999 | 0.944203 | 0.994965 |
| CG7407 | 3162.369 | -0.04748 | 0.697294 | -0.06809 | 0.945717 | 0.994965 |
| CG14650 | 1708.269 | 0.070489 | 1.166193 | 0.060444 | 0.951802 | 0.994965 |
| cl | 1898.482 | -0.04926 | 0.85957 | -0.0573 | 0.954304 | 0.994965 |
| CG11029 | 171.862 | 0.209773 | 3.224916 | 0.065048 | 0.948136 | 0.994965 |
| TTLL3B | 103.6165 | 0.259606 | 3.841373 | 0.067581 | 0.946119 | 0.994965 |
| MME1 | 104.4091 | 0.269741 | 3.84983 | 0.070066 | 0.944141 | 0.994965 |
| CG15141 | 1573.8 | -0.05756 | 0.851888 | -0.06757 | 0.946126 | 0.994965 |
| CG11035 | 637.0524 | 0.110613 | 1.843351 | 0.060006 | 0.952151 | 0.994965 |
| CG8507 | 2348.088 | -0.07703 | 1.323621 | -0.0582 | 0.953589 | 0.994965 |
| ninaG | 971.7853 | -0.08534 | 1.361766 | -0.06267 | 0.950031 | 0.994965 |
| yrt | 1132.873 | -0.05462 | 0.929964 | -0.05874 | 0.953162 | 0.994965 |
| Nup43 | 418.1251 | -0.1574 | 2.411441 | -0.06527 | 0.947958 | 0.994965 |
| Gos28 | 1239.856 | 0.060313 | 0.923655 | 0.065298 | 0.947937 | 0.994965 |
| mdlc | 129.6168 | -0.19813 | 3.097358 | -0.06397 | 0.948996 | 0.994965 |
| Pi3K92E | 2661.014 | -0.04955 | 0.785497 | -0.06308 | 0.9497 | 0.994965 |
| hdly | 520.2199 | 0.109193 | 1.701946 | 0.064158 | 0.948844 | 0.994965 |
| SKIP | 7419.624 | -0.0325 | 0.538036 | -0.0604 | 0.951836 | 0.994965 |
| CG11791 | 4977.577 | -0.03281 | 0.566946 | -0.05787 | 0.953851 | 0.994965 |
| Stt3B | 1565.332 | 0.103761 | 1.56112 | 0.066466 | 0.947007 | 0.994965 |
| CG12290 | 2037.968 | 0.044808 | 0.736494 | 0.060839 | 0.951487 | 0.994965 |
| Hrb98DE | 6366.078 | 0.036342 | 0.546971 | 0.066443 | 0.947025 | 0.994965 |
| CG15535 | 196.3422 | 0.146533 | 2.418015 | 0.060601 | 0.951677 | 0.994965 |
| Prosalpha5 | 1933.718 | 0.06152 | 0.991383 | 0.062054 | 0.950519 | 0.994965 |
| RpL17 | 10194.53 | 0.035884 | 0.508471 | 0.070572 | 0.943738 | 0.994965 |
| CG9650 | 2545.346 | 0.048502 | 0.716334 | 0.067709 | 0.946018 | 0.994965 |
| mahe | 8020.723 | -0.03239 | 0.535995 | -0.06043 | 0.951814 | 0.994965 |
| CG10555 | 1362.387 | 0.060556 | 1.026117 | 0.059014 | 0.952941 | 0.994965 |
| Dsor1 | 1067.818 | -0.08663 | 1.475794 | -0.0587 | 0.953192 | 0.994965 |
| insc | 72.16912 | 0.258704 | 3.943939 | 0.065595 | 0.9477 | 0.994965 |
| Sara | 3103.344 | -0.0496 | 0.713348 | -0.06954 | 0.944563 | 0.994965 |
| eEF1beta | 5333.051 | -0.03542 | 0.59869 | -0.05916 | 0.952828 | 0.994965 |
| CG7639 | 1274.492 | -0.11433 | 1.616235 | -0.07074 | 0.943604 | 0.994965 |

| | | | | | | |
|---|---|---|---|---|---|---|
| mspo | 1825.469 | 0.056469 | 0.891063 | 0.063373 | 0.94947 | 0.994965 |
| mam | 2668.438 | 0.040975 | 0.706729 | 0.057979 | 0.953766 | 0.994965 |
| CG17574 | 1488.481 | 0.057431 | 0.888997 | 0.064602 | 0.948491 | 0.994965 |
| CG13229 | 111.7774 | 0.252176 | 3.834031 | 0.065773 | 0.947559 | 0.994965 |
| CG1663 | 295.1462 | -0.19674 | 3.287617 | -0.05984 | 0.952281 | 0.994965 |
| CG1909 | 13123.86 | -0.02973 | 0.497151 | -0.05981 | 0.952309 | 0.994965 |
| CG32850 | 4165.967 | 0.036045 | 0.627767 | 0.057417 | 0.954213 | 0.994965 |
| chb | 2600.261 | -0.04376 | 0.681578 | -0.06421 | 0.948804 | 0.994965 |
| mRpL30 | 1484.422 | 0.058425 | 0.879055 | 0.066463 | 0.947009 | 0.994965 |
| exd | 6927.001 | -0.0323 | 0.496681 | -0.06503 | 0.948148 | 0.994965 |
| CG8892 | 662.2281 | -0.10438 | 1.775927 | -0.05878 | 0.95313 | 0.994965 |
| slpr | 3906.951 | 0.049393 | 0.719589 | 0.06864 | 0.945276 | 0.994965 |
| Nipped-A | 9970.498 | -0.03174 | 0.466287 | -0.06807 | 0.945728 | 0.994965 |
| vn | 186.7135 | -0.18147 | 2.836901 | -0.06397 | 0.948995 | 0.994965 |
| HIP-R | 354.6475 | 0.093823 | 1.629982 | 0.05756 | 0.954099 | 0.994965 |
| CtBP | 6483.511 | -0.03556 | 0.529715 | -0.06714 | 0.946472 | 0.994965 |
| Marf | 2566.069 | -0.04028 | 0.687641 | -0.05858 | 0.953288 | 0.994965 |
| CG5758 | 27314.48 | -0.02946 | 0.472577 | -0.06234 | 0.950289 | 0.994965 |
| Pp2A-29B | 5069.449 | 0.045379 | 0.680711 | 0.066664 | 0.946849 | 0.994965 |
| Sod3 | 22009.08 | -0.0306 | 0.478218 | -0.06398 | 0.948988 | 0.994965 |
| Vkor | 203.2995 | 0.172229 | 2.803652 | 0.06143 | 0.951017 | 0.994965 |
| CG34189 | 101.3977 | -0.24378 | 3.567508 | -0.06833 | 0.94552 | 0.994965 |
| bbg | 80.2538 | -0.2067 | 3.48714 | -0.05927 | 0.952733 | 0.994965 |
| ome | 1168.401 | -0.07181 | 1.145529 | -0.06269 | 0.950017 | 0.994965 |
| mtgo | 6637.402 | -0.03024 | 0.506258 | -0.05973 | 0.952368 | 0.994965 |
| Msp300 | 51062.2 | 0.024465 | 0.409902 | 0.059685 | 0.952406 | 0.994965 |
| whd | 2379.213 | -0.06188 | 0.982079 | -0.06301 | 0.94976 | 0.994965 |
| AGO1 | 15203.62 | -0.02827 | 0.432778 | -0.06531 | 0.947924 | 0.994965 |
| CG9890 | 131.1515 | 0.212109 | 3.192522 | 0.066439 | 0.947028 | 0.994965 |
| Rpn8 | 1300.253 | -0.07239 | 1.078192 | -0.06714 | 0.946469 | 0.994965 |
| emp | 407.8485 | -0.16227 | 2.339028 | -0.06937 | 0.944692 | 0.994965 |
| CG1233 | 497.6885 | 0.124215 | 1.856396 | 0.066912 | 0.946652 | 0.994965 |
| Atf-2 | 201.6262 | 0.173736 | 2.500796 | 0.069472 | 0.944614 | 0.994965 |
| FASN1 | 13518.92 | 0.027084 | 0.452554 | 0.059846 | 0.952278 | 0.994965 |
| br | 1649.741 | -0.05128 | 0.85771 | -0.05978 | 0.952328 | 0.994965 |
| Sema5c | 482.7145 | 0.083332 | 1.392006 | 0.059865 | 0.952263 | 0.994965 |
| aqz | 64747.26 | 0.022693 | 0.379793 | 0.059752 | 0.952353 | 0.994965 |
| PAN3 | 10388.47 | 0.030178 | 0.473396 | 0.063748 | 0.949171 | 0.994965 |
| CG14961 | 146.366 | 0.147156 | 2.509648 | 0.058636 | 0.953242 | 0.994965 |
| CG14963 | 105.206 | -0.26847 | 3.849479 | -0.06974 | 0.9444 | 0.994965 |
| Atg3 | 2905.643 | 0.04119 | 0.666238 | 0.061825 | 0.950702 | 0.994965 |
| Strump | 1249.622 | -0.11735 | 1.697527 | -0.06913 | 0.944888 | 0.994965 |
| CG10713 | 907.6211 | -0.0608 | 1.037085 | -0.05863 | 0.953249 | 0.994965 |
| CG7888 | 67446.69 | -0.03352 | 0.51138 | -0.06556 | 0.947732 | 0.994965 |
| pst | 1364.464 | -0.06629 | 0.967449 | -0.06852 | 0.945375 | 0.994965 |
| mal | 82.64799 | -0.26955 | 3.95034 | -0.06824 | 0.945598 | 0.994965 |
| Vps53 | 696.2526 | -0.09824 | 1.382201 | -0.07107 | 0.943339 | 0.994965 |
| CG3702 | 3907.055 | 0.043718 | 0.737431 | 0.059284 | 0.952726 | 0.994965 |
| Wdr62 | 3467.262 | -0.04176 | 0.659116 | -0.06336 | 0.949481 | 0.994965 |
| clu | 1783.205 | 0.04902 | 0.858727 | 0.057084 | 0.954478 | 0.995012 |

| | | | | | | |
|---|---|---|---|---|---|---|
| Vps28 | 2477.271 | 0.039978 | 0.702161 | 0.056936 | 0.954596 | 0.995012 |
| PpV | 5539.259 | 0.032295 | 0.653269 | 0.049436 | 0.960572 | 0.995041 |
| CG32428 | 1176.485 | -0.05017 | 0.984533 | -0.05096 | 0.959359 | 0.995041 |
| Hr78 | 589.0728 | 0.075323 | 1.349529 | 0.055814 | 0.95549 | 0.995041 |
| Csp | 12052.66 | 0.025015 | 0.490204 | 0.051029 | 0.959302 | 0.995041 |
| CG12173 | 1313.784 | -0.04348 | 0.88126 | -0.04934 | 0.960648 | 0.995041 |
| spict | 631.0513 | 0.094679 | 1.78216 | 0.053126 | 0.957632 | 0.995041 |
| c(2)M | 69.86793 | -0.18342 | 3.460429 | -0.053 | 0.957728 | 0.995041 |
| Hasp | 29292.13 | -0.02268 | 0.44584 | -0.05088 | 0.959423 | 0.995041 |
| Dh44 | 82.52315 | 0.133914 | 2.709591 | 0.049422 | 0.960583 | 0.995041 |
| Nup93-2 | 165.9303 | -0.17936 | 3.291182 | -0.0545 | 0.95654 | 0.995041 |
| RpL27 | 9284.887 | 0.029281 | 0.571914 | 0.051198 | 0.959168 | 0.995041 |
| jdp | 19584.29 | -0.02122 | 0.412951 | -0.05138 | 0.959021 | 0.995041 |
| wts | 2858.589 | -0.03405 | 0.651257 | -0.05228 | 0.958304 | 0.995041 |
| mRpL32 | 892.3108 | -0.06983 | 1.335874 | -0.05228 | 0.958309 | 0.995041 |
| SCAP | 2278.888 | 0.035114 | 0.712611 | 0.049276 | 0.9607 | 0.995041 |
| swi2 | 135.8302 | -0.11929 | 2.23712 | -0.05332 | 0.957476 | 0.995041 |
| l(1)G0193 | 3232.436 | -0.03303 | 0.666965 | -0.04953 | 0.9605 | 0.995041 |
| CG10932 | 164.9929 | 0.174805 | 3.232498 | 0.054077 | 0.956874 | 0.995041 |
| CG10795 | 1018.786 | -0.08142 | 1.636056 | -0.04976 | 0.960311 | 0.995041 |
| twz | 820.2648 | -0.07243 | 1.381297 | -0.05243 | 0.958183 | 0.995041 |
| bug | 749.6106 | 0.069156 | 1.375344 | 0.050283 | 0.959897 | 0.995041 |
| fsd | 290.0375 | -0.10903 | 2.038188 | -0.0535 | 0.957337 | 0.995041 |
| Mtor | 992.9823 | -0.07781 | 1.437367 | -0.05413 | 0.95683 | 0.995041 |
| Tsp42Ej | 16001.82 | -0.02364 | 0.475054 | -0.04977 | 0.960308 | 0.995041 |
| Lamp1 | 11269.44 | -0.02494 | 0.490907 | -0.0508 | 0.959485 | 0.995041 |
| CG4537 | 87.23716 | 0.171225 | 3.34165 | 0.05124 | 0.959135 | 0.995041 |
| CHES-1-lik | 2551.679 | 0.036978 | 0.702766 | 0.052617 | 0.958037 | 0.995041 |
| GstD1 | 30023.1 | 0.024384 | 0.459751 | 0.053038 | 0.957702 | 0.995041 |
| side-IV | 1322.358 | 0.059014 | 1.076761 | 0.054807 | 0.956292 | 0.995041 |
| CG34133 | 1492.653 | 0.062235 | 1.237235 | 0.050302 | 0.959882 | 0.995041 |
| Prosbeta7 | 2086.1 | -0.03838 | 0.771147 | -0.04977 | 0.960305 | 0.995041 |
| Ppcs | 3072.613 | 0.035797 | 0.673935 | 0.053116 | 0.957639 | 0.995041 |
| Mdh2 | 19842 | 0.023714 | 0.44395 | 0.053416 | 0.9574 | 0.995041 |
| Skeletor | 271.4584 | -0.1236 | 2.249558 | -0.05494 | 0.956183 | 0.995041 |
| CG13551 | 5531.345 | -0.03071 | 0.58213 | -0.05275 | 0.957931 | 0.995041 |
| CG3860 | 3728.079 | 0.039575 | 0.704309 | 0.056189 | 0.955191 | 0.995041 |
| Vdup1 | 303.9643 | -0.15691 | 2.797709 | -0.05608 | 0.955275 | 0.995041 |
| Eip93F | 8673.085 | -0.02589 | 0.493313 | -0.05249 | 0.958138 | 0.995041 |
| lncRNA:CR | 304.7639 | -0.11719 | 2.259356 | -0.05187 | 0.958633 | 0.995041 |
| asRNA:CR4 | 232.8226 | -0.09602 | 1.753961 | -0.05474 | 0.956344 | 0.995041 |
| Sec5 | 857.8745 | 0.07 | 1.402676 | 0.049905 | 0.960198 | 0.995041 |
| CG12091 | 4048.909 | 0.036729 | 0.671237 | 0.054719 | 0.956363 | 0.995041 |
| CG13008 | 111.9169 | 0.183814 | 3.547283 | 0.051818 | 0.958674 | 0.995041 |
| Ada3 | 1054.25 | -0.05142 | 0.953217 | -0.05394 | 0.956983 | 0.995041 |
| CG18081 | 1519.929 | 0.056472 | 1.030038 | 0.054825 | 0.956278 | 0.995041 |
| ND-13B | 5515.45 | 0.030359 | 0.554328 | 0.054766 | 0.956325 | 0.995041 |
| CalpB | 3003.218 | -0.03584 | 0.666356 | -0.05378 | 0.957106 | 0.995041 |
| Arl5 | 962.9202 | 0.057646 | 1.072164 | 0.053766 | 0.957121 | 0.995041 |
| NTPase | 2944.822 | 0.035678 | 0.670874 | 0.053182 | 0.957587 | 0.995041 |

| | | | | | | |
|---|---|---|---|---|---|---|
| RpL36A | 13167.36 | -0.02465 | 0.502697 | -0.04904 | 0.960885 | 0.995105 |
| NT5E-2 | 1650.781 | 0.040267 | 0.826634 | 0.048712 | 0.961149 | 0.99514 |
| Nipped-B | 20589.67 | -0.02231 | 0.458184 | -0.04869 | 0.961167 | 0.99514 |
| CG5160 | 67.09987 | -0.18587 | 3.834502 | -0.04847 | 0.96134 | 0.995191 |
| Naa40 | 149.0145 | -0.18407 | 3.831723 | -0.04804 | 0.961686 | 0.995292 |
| Eaf6 | 447.9735 | 0.083731 | 1.739559 | 0.048133 | 0.96161 | 0.995292 |
| Oatp30B | 1317.333 | -0.06109 | 1.286697 | -0.04748 | 0.962129 | 0.995298 |
| klg | 6177.742 | -0.02606 | 0.551834 | -0.04722 | 0.962341 | 0.995298 |
| Wsck | 877.8291 | -0.05402 | 1.143351 | -0.04724 | 0.962318 | 0.995298 |
| CG7849 | 921.0052 | -0.08109 | 1.725885 | -0.04698 | 0.962527 | 0.995298 |
| lncRNA:CR | 3580.981 | 0.028569 | 0.613067 | 0.046601 | 0.962831 | 0.995298 |
| dpr9 | 4861.144 | -0.02704 | 0.579502 | -0.04666 | 0.962785 | 0.995298 |
| lncRNA:CR | 113.3845 | -0.157 | 3.378108 | -0.04647 | 0.962932 | 0.995298 |
| Gabat | 28363.75 | -0.01852 | 0.397456 | -0.04659 | 0.962838 | 0.995298 |
| CG8281 | 66.5963 | -0.17899 | 3.835643 | -0.04667 | 0.962779 | 0.995298 |
| oaf | 2199.768 | 0.03693 | 0.778801 | 0.047419 | 0.96218 | 0.995298 |
| Men | 4372.328 | 0.026535 | 0.597079 | 0.044441 | 0.964553 | 0.995731 |
| lncRNA:CR | 2540.535 | 0.038654 | 0.873165 | 0.044269 | 0.96469 | 0.995731 |
| trp | 75.21864 | -0.17222 | 3.833946 | -0.04492 | 0.964172 | 0.995731 |
| SmydA-5 | 546.4073 | -0.09872 | 2.231496 | -0.04424 | 0.964715 | 0.995731 |
| unc-119 | 4259.976 | 0.029905 | 0.668131 | 0.04476 | 0.964299 | 0.995731 |
| shn | 2980.299 | 0.040787 | 0.903609 | 0.045138 | 0.963997 | 0.995731 |
| CG9062 | 283.4173 | 0.076867 | 1.720853 | 0.044668 | 0.964372 | 0.995731 |
| RfC4 | 79.91044 | -0.17029 | 3.792004 | -0.04491 | 0.964181 | 0.995731 |
| 5-HT1B | 1056.006 | 0.046069 | 1.021116 | 0.045116 | 0.964015 | 0.995731 |
| stac | 1287.493 | -0.06619 | 1.494452 | -0.04429 | 0.964671 | 0.995731 |
| mRpL20 | 2185.082 | 0.040524 | 0.907739 | 0.044642 | 0.964392 | 0.995731 |
| Edem2 | 1067.359 | 0.047798 | 1.10023 | 0.043444 | 0.965348 | 0.995744 |
| Tmem131 | 673.953 | 0.05247 | 1.203987 | 0.043581 | 0.965239 | 0.995744 |
| Hr3 | 858.4217 | 0.059027 | 1.345661 | 0.043864 | 0.965012 | 0.995744 |
| CG34231 | 84.43264 | -0.14255 | 3.272748 | -0.04356 | 0.965259 | 0.995744 |
| CG4577 | 33143 | -0.01697 | 0.38793 | -0.04374 | 0.965113 | 0.995744 |
| y | 117.8561 | 0.138852 | 3.253588 | 0.042677 | 0.965959 | 0.995766 |
| fusl | 958.3188 | 0.044521 | 1.047969 | 0.042483 | 0.966114 | 0.995766 |
| GATAd | 2186.15 | -0.03455 | 0.810967 | -0.04261 | 0.966013 | 0.995766 |
| CG33230 | 326.7783 | -0.10692 | 2.508342 | -0.04262 | 0.966001 | 0.995766 |
| Met | 2283.211 | 0.030355 | 0.704954 | 0.043059 | 0.965655 | 0.995766 |
| Prim1 | 89.95923 | 0.141074 | 3.278091 | 0.043035 | 0.965673 | 0.995766 |
| Src42A | 3918.503 | 0.024772 | 0.59744 | 0.041464 | 0.966926 | 0.996475 |
| CG8860 | 66.82152 | 0.118721 | 2.888035 | 0.041108 | 0.96721 | 0.99664 |
| CG6629 | 76.09114 | 0.15715 | 3.87183 | 0.040588 | 0.967624 | 0.996642 |
| Inx7 | 142.327 | 0.156733 | 3.871707 | 0.040482 | 0.967709 | 0.996642 |
| bun | 30435.66 | -0.01618 | 0.398968 | -0.04055 | 0.967653 | 0.996642 |
| Galt | 853.3547 | 0.056762 | 1.397039 | 0.04063 | 0.967591 | 0.996642 |
| CG15653 | 180.9958 | -0.14962 | 3.751538 | -0.03988 | 0.968187 | 0.997006 |
| lncRNA:CR | 72.45577 | -0.13839 | 3.513657 | -0.03939 | 0.968583 | 0.997196 |
| flfl | 3821.326 | -0.02326 | 0.595943 | -0.03903 | 0.968868 | 0.997196 |
| Diap2 | 7738.899 | 0.02079 | 0.529549 | 0.03926 | 0.968683 | 0.997196 |
| kis | 25071.54 | -0.01566 | 0.400602 | -0.03908 | 0.968826 | 0.997196 |
| Ufd4 | 7544.086 | -0.02264 | 0.598347 | -0.03784 | 0.969812 | 0.997855 |

| | | | | | | |
|---|---|---|---|---|---|---|
| CG12929 | 345.0622 | -0.0852 | 2.263859 | -0.03764 | 0.969979 | 0.997855 |
| capt | 1581.252 | -0.03254 | 0.865396 | -0.0376 | 0.970006 | 0.997855 |
| CG43102 | 1808.216 | 0.034798 | 0.920222 | 0.037814 | 0.969836 | 0.997855 |
| CG5339 | 354.6715 | -0.06964 | 1.865691 | -0.03733 | 0.970225 | 0.997953 |
| AstA-R2 | 175.1769 | 0.140743 | 3.79001 | 0.037135 | 0.970377 | 0.997981 |
| PPP4R2r | 1489.031 | 0.038997 | 1.109292 | 0.035155 | 0.971956 | 0.998137 |
| EMC7 | 3485.371 | -0.02181 | 0.622917 | -0.03502 | 0.972067 | 0.998137 |
| arr | 558.0726 | -0.05039 | 1.386178 | -0.03635 | 0.971002 | 0.998137 |
| MsR1 | 826.4999 | -0.05069 | 1.440169 | -0.0352 | 0.971924 | 0.998137 |
| Ir62a | 352.9059 | 0.074286 | 2.083216 | 0.035659 | 0.971554 | 0.998137 |
| mt:tRNA:A | 254.5425 | -0.06146 | 1.714065 | -0.03585 | 0.971398 | 0.998137 |
| rgn | 6123.624 | 0.020053 | 0.55514 | 0.036122 | 0.971185 | 0.998137 |
| lncRNA:CR | 85.61395 | -0.13571 | 3.886599 | -0.03492 | 0.972145 | 0.998137 |
| lncRNA:CR | 82.59849 | 0.108439 | 3.079776 | 0.03521 | 0.971912 | 0.998137 |
| Zasp52 | 470.4454 | 0.048492 | 1.37919 | 0.03516 | 0.971952 | 0.998137 |
| Tsf1 | 19631.83 | 0.021785 | 0.623119 | 0.03496 | 0.972111 | 0.998137 |
| CG32533 | 212.6516 | -0.08952 | 2.5508 | -0.0351 | 0.972004 | 0.998137 |
| CG32066 | 1244.266 | 0.03333 | 0.919992 | 0.036228 | 0.9711 | 0.998137 |
| lncRNA:CR | 1847.072 | 0.030495 | 0.894559 | 0.03409 | 0.972806 | 0.99818 |
| pasi1 | 1254.87 | 0.052734 | 1.521946 | 0.034649 | 0.97236 | 0.99818 |
| CG15922 | 430.0127 | 0.046566 | 1.3573 | 0.034308 | 0.972632 | 0.99818 |
| ATPsynbet | 59151.02 | 0.013116 | 0.387656 | 0.033835 | 0.973009 | 0.99818 |
| HnRNP-K | 4771.175 | 0.019251 | 0.56068 | 0.034336 | 0.972609 | 0.99818 |
| Ac76E | 5485.711 | -0.01894 | 0.558577 | -0.0339 | 0.972953 | 0.99818 |
| mirr | 449.8909 | 0.067081 | 1.986146 | 0.033774 | 0.973057 | 0.99818 |
| CG6094 | 271.38 | -0.0661 | 2.073432 | -0.03188 | 0.974569 | 0.998329 |
| CAH1 | 3487.396 | 0.022604 | 0.70064 | 0.032262 | 0.974263 | 0.998329 |
| Lrrk | 1107.747 | 0.046851 | 1.425351 | 0.03287 | 0.973779 | 0.998329 |
| LpR2 | 8764.929 | -0.01789 | 0.537777 | -0.03326 | 0.973467 | 0.998329 |
| CG17059 | 862.1475 | -0.03454 | 1.067386 | -0.03236 | 0.974186 | 0.998329 |
| DPCoAC | 575.5529 | -0.0449 | 1.380389 | -0.03253 | 0.974051 | 0.998329 |
| mEFG1 | 1067.974 | 0.04138 | 1.297049 | 0.031903 | 0.974549 | 0.998329 |
| Vha68-1 | 23743.6 | -0.0136 | 0.408457 | -0.0333 | 0.973434 | 0.998329 |
| l(3)neo38 | 7411.875 | 0.017556 | 0.533252 | 0.032922 | 0.973737 | 0.998329 |
| CG10352 | 1800.472 | -0.03942 | 1.205563 | -0.0327 | 0.973916 | 0.998329 |
| Pfdn6 | 2907.255 | 0.021001 | 0.658842 | 0.031875 | 0.974571 | 0.998329 |
| Dlip2 | 829.9205 | -0.04738 | 1.494755 | -0.0317 | 0.974711 | 0.998345 |
| TRAM | 951.144 | -0.04027 | 1.361665 | -0.02958 | 0.976404 | 0.998364 |
| CG14777 | 2805.532 | 0.017076 | 0.653624 | 0.026124 | 0.979158 | 0.998364 |
| Xpac | 599.091 | -0.04511 | 1.854962 | -0.02432 | 0.980596 | 0.998364 |
| CSN3 | 1350.799 | 0.029187 | 1.216684 | 0.023989 | 0.980861 | 0.998364 |
| CG7611 | 3438.522 | -0.01436 | 0.641705 | -0.02239 | 0.982141 | 0.998364 |
| CG11999 | 1573.777 | 0.027894 | 0.902636 | 0.030903 | 0.975347 | 0.998364 |
| C1GalTA | 4437.159 | 0.015634 | 0.561649 | 0.027836 | 0.977793 | 0.998364 |
| CG6495 | 2172.241 | 0.022442 | 0.713732 | 0.031443 | 0.974916 | 0.998364 |
| Ing5 | 1332.688 | -0.03102 | 1.027481 | -0.03019 | 0.975914 | 0.998364 |
| Ddc | 8109.749 | -0.01198 | 0.532116 | -0.02251 | 0.982044 | 0.998364 |
| CG11768 | 148.3961 | 0.075229 | 2.596905 | 0.028969 | 0.976889 | 0.998364 |
| CG9801 | 1119.37 | -0.0265 | 0.938931 | -0.02822 | 0.977487 | 0.998364 |
| Sf3b5 | 2040.35 | -0.02017 | 0.756732 | -0.02666 | 0.978731 | 0.998364 |

| | | | | | | |
|---|---|---|---|---|---|---|
| Lk6 | 16243.13 | 0.01191 | 0.456171 | 0.026108 | 0.979171 | 0.998364 |
| Dic1 | 4668.343 | -0.01531 | 0.635616 | -0.02408 | 0.980785 | 0.998364 |
| ATPsynE | 11992.18 | -0.01048 | 0.442533 | -0.02369 | 0.981103 | 0.998364 |
| CG6276 | 1269.627 | 0.02185 | 1.001802 | 0.021811 | 0.982599 | 0.998364 |
| Scp2 | 14390.9 | 0.010034 | 0.443398 | 0.022629 | 0.981946 | 0.998364 |
| RpS30 | 33834.19 | -0.01011 | 0.416758 | -0.02427 | 0.980641 | 0.998364 |
| CG5071 | 957.7975 | -0.04737 | 1.72901 | -0.0274 | 0.978142 | 0.998364 |
| Oseg6 | 515.5986 | -0.04472 | 1.754025 | -0.02549 | 0.979662 | 0.998364 |
| rswl | 693.3795 | -0.0358 | 1.230512 | -0.02909 | 0.976793 | 0.998364 |
| Moe | 4215.516 | -0.01324 | 0.579132 | -0.02285 | 0.981767 | 0.998364 |
| CG2004 | 599.2691 | 0.040339 | 1.374512 | 0.029348 | 0.976587 | 0.998364 |
| eIF3m | 11799.12 | -0.01271 | 0.45179 | -0.02814 | 0.977551 | 0.998364 |
| mars | 112.9574 | -0.07878 | 2.673673 | -0.02946 | 0.976494 | 0.998364 |
| slv | 818.324 | 0.049369 | 1.679271 | 0.029399 | 0.976546 | 0.998364 |
| Gdap2 | 11371.37 | 0.01263 | 0.492257 | 0.025657 | 0.979531 | 0.998364 |
| tutl | 19005.15 | -0.01064 | 0.409771 | -0.02595 | 0.979294 | 0.998364 |
| Tusp | 3305.494 | -0.01462 | 0.665524 | -0.02197 | 0.982469 | 0.998364 |
| CG31712 | 11594.34 | 0.011364 | 0.461137 | 0.024644 | 0.980339 | 0.998364 |
| jet | 897.148 | 0.03588 | 1.594589 | 0.022501 | 0.982048 | 0.998364 |
| Rim | 26210.71 | 0.012768 | 0.414268 | 0.03082 | 0.975413 | 0.998364 |
| ey | 122.728 | 0.097713 | 3.338387 | 0.02927 | 0.97665 | 0.998364 |
| RpS8 | 24716.72 | 0.011739 | 0.434389 | 0.027024 | 0.978441 | 0.998364 |
| RpL41 | 7890.353 | -0.01171 | 0.498763 | -0.02348 | 0.981268 | 0.998364 |
| CG34141 | 247.3106 | 0.055072 | 2.181749 | 0.025242 | 0.979862 | 0.998364 |
| CG40470 | 143.933 | 0.083335 | 3.230059 | 0.0258 | 0.979417 | 0.998364 |
| TrissinR | 2000.713 | 0.019615 | 0.860295 | 0.0228 | 0.98181 | 0.998364 |
| 18SrRNA-F | 343.1505 | 0.037028 | 1.494555 | 0.024775 | 0.980234 | 0.998364 |
| Akr1B | 4452.182 | 0.016597 | 0.560697 | 0.0296 | 0.976386 | 0.998364 |
| vvl | 4924.259 | -0.01869 | 0.751912 | -0.02485 | 0.980173 | 0.998364 |
| fid | 1533.682 | 0.021989 | 0.99588 | 0.02208 | 0.982384 | 0.998364 |
| sgl | 4419.641 | 0.013501 | 0.568037 | 0.023768 | 0.981038 | 0.998364 |
| psq | 9429.077 | 0.012503 | 0.518084 | 0.024133 | 0.980746 | 0.998364 |
| CG31688 | 1563.493 | 0.02333 | 0.862729 | 0.027042 | 0.978426 | 0.998364 |
| qkr58E-1 | 1272.847 | 0.022254 | 0.943675 | 0.023582 | 0.981186 | 0.998364 |
| egl | 1187.646 | 0.027368 | 0.936656 | 0.029219 | 0.97669 | 0.998364 |
| gammaSna | 2129.739 | 0.019975 | 0.718662 | 0.027794 | 0.977826 | 0.998364 |
| DCP1 | 805.2965 | 0.028544 | 1.274172 | 0.022402 | 0.982127 | 0.998364 |
| nerfin-1 | 241.8097 | 0.070228 | 2.276438 | 0.03085 | 0.975389 | 0.998364 |
| CG13907 | 893.2454 | 0.034198 | 1.210156 | 0.028259 | 0.977456 | 0.998364 |
| CG43755 | 260.7312 | -0.06533 | 2.264097 | -0.02885 | 0.976982 | 0.998364 |
| RhoGEF3 | 6571.009 | -0.0159 | 0.565995 | -0.0281 | 0.977583 | 0.998364 |
| hfp | 9463.801 | 0.014722 | 0.496408 | 0.029657 | 0.97634 | 0.998364 |
| Bgb | 4642.623 | 0.017205 | 0.623468 | 0.027596 | 0.977985 | 0.998364 |
| CG11537 | 6153.509 | -0.01855 | 0.648258 | -0.02861 | 0.977177 | 0.998364 |
| Atg101 | 2690.852 | 0.039134 | 1.374033 | 0.028481 | 0.977278 | 0.998364 |
| CG14200 | 1923.615 | -0.02301 | 1.001631 | -0.02298 | 0.981669 | 0.998364 |
| CG7460 | 73.19449 | -0.08701 | 3.83233 | -0.0227 | 0.981887 | 0.998364 |
| RpS4 | 16871.57 | 0.009301 | 0.424089 | 0.021931 | 0.982503 | 0.998364 |
| CG4911 | 17837.65 | -0.00952 | 0.438619 | -0.0217 | 0.982691 | 0.998364 |
| CG4611 | 259.0888 | -0.07396 | 2.66542 | -0.02775 | 0.977862 | 0.998364 |

| | | | | | | |
|---|---|---|---|---|---|---|
| CG32523 | 174.214 | 0.065862 | 2.624397 | 0.025096 | 0.979978 | 0.998364 |
| Nmnat | 1335.052 | -0.02356 | 1.103373 | -0.02136 | 0.982961 | 0.998512 |
| Moca-cyp | 1742.441 | 0.016019 | 0.778345 | 0.020581 | 0.98358 | 0.998627 |
| CG3224 | 998.1129 | 0.020549 | 0.98563 | 0.020849 | 0.983366 | 0.998627 |
| Nf1 | 3504.753 | -0.01275 | 0.618187 | -0.02062 | 0.983546 | 0.998627 |
| Vps20 | 612.157 | -0.02972 | 1.454538 | -0.02044 | 0.983696 | 0.998627 |
| RpS12 | 20454.06 | 0.008909 | 0.42476 | 0.020973 | 0.983267 | 0.998627 |
| Tspo | 374.0332 | 0.02956 | 1.472017 | 0.020081 | 0.983979 | 0.998787 |
| Pgam5 | 769.522 | -0.03751 | 1.888099 | -0.01987 | 0.984151 | 0.998836 |
| CG8372 | 3412.662 | -0.01398 | 0.757661 | -0.01845 | 0.985283 | 0.999003 |
| Reps | 1130.71 | -0.02049 | 1.060069 | -0.01933 | 0.984581 | 0.999003 |
| CG12594 | 2141.679 | 0.013756 | 0.744054 | 0.018488 | 0.985249 | 0.999003 |
| CG5455 | 8381.627 | 0.010853 | 0.576353 | 0.018831 | 0.984976 | 0.999003 |
| spt4 | 635.3579 | -0.02234 | 1.159076 | -0.01927 | 0.984626 | 0.999003 |
| cic | 6713.877 | -0.01 | 0.526857 | -0.01898 | 0.984855 | 0.999003 |
| CG32280 | 1756.095 | 0.017098 | 0.928714 | 0.01841 | 0.985311 | 0.999003 |
| CG4789 | 417.0538 | 0.039153 | 2.061529 | 0.018992 | 0.984848 | 0.999003 |
| ppl | 1654.658 | 0.01389 | 0.902661 | 0.015388 | 0.987723 | 0.999061 |
| CG10470 | 2179.692 | 0.013281 | 0.814351 | 0.016308 | 0.986989 | 0.999061 |
| mRpL19 | 947.1948 | -0.021 | 1.438319 | -0.0146 | 0.988353 | 0.999061 |
| Acyp2 | 912.2747 | 0.014728 | 1.020139 | 0.014438 | 0.988481 | 0.999061 |
| Not3 | 6676.585 | -0.00872 | 0.532872 | -0.01636 | 0.986947 | 0.999061 |
| PCNA | 263.4904 | -0.04513 | 2.846239 | -0.01585 | 0.98735 | 0.999061 |
| CG15611 | 3015.594 | -0.01159 | 0.731446 | -0.01584 | 0.987359 | 0.999061 |
| Mlf | 10008.96 | -0.00823 | 0.552293 | -0.0149 | 0.988115 | 0.999061 |
| CG8726 | 3226.591 | 0.010569 | 0.645374 | 0.016376 | 0.986934 | 0.999061 |
| dpy | 1653.502 | -0.02205 | 1.426239 | -0.01546 | 0.987667 | 0.999061 |
| vir-1 | 23390.44 | 0.007076 | 0.485106 | 0.014586 | 0.988362 | 0.999061 |
| MFS17 | 17120.36 | -0.00823 | 0.47253 | -0.01743 | 0.986096 | 0.999061 |
| gish | 12149.89 | 0.007341 | 0.481527 | 0.015246 | 0.987836 | 0.999061 |
| CG42263 | 198.6739 | -0.04504 | 3.057265 | -0.01473 | 0.988245 | 0.999061 |
| RpL37A | 9475.194 | 0.007631 | 0.468979 | 0.016272 | 0.987017 | 0.999061 |
| Bsg | 20214.18 | -0.00735 | 0.410289 | -0.01792 | 0.985704 | 0.999061 |
| dikar | 5127.009 | 0.008792 | 0.544288 | 0.016152 | 0.987113 | 0.999061 |
| Psn | 1357.248 | -0.01273 | 0.865456 | -0.01471 | 0.988262 | 0.999061 |
| l(3)80Fj | 16666.68 | -0.00659 | 0.430972 | -0.0153 | 0.987792 | 0.999061 |
| CG13919 | 338.6262 | -0.0461 | 2.636032 | -0.01749 | 0.986048 | 0.999061 |
| CG16753 | 405.4688 | 0.031653 | 2.063878 | 0.015337 | 0.987764 | 0.999061 |
| CG17746 | 2568.542 | -0.01326 | 0.900364 | -0.01472 | 0.988252 | 0.999061 |
| Teh2 | 5428.62 | 0.00987 | 0.574745 | 0.017173 | 0.986299 | 0.999061 |
| GCS1 | 423.4571 | -0.02438 | 1.553866 | -0.01569 | 0.987483 | 0.999061 |
| Ist1 | 2345.496 | -0.01577 | 0.917868 | -0.01718 | 0.986293 | 0.999061 |
| CG10340 | 4136.75 | -0.00854 | 0.600023 | -0.01424 | 0.988638 | 0.999094 |
| Mnt | 3937.365 | -0.00844 | 0.603248 | -0.01399 | 0.988836 | 0.999169 |
| CG3021 | 864.1273 | 0.018574 | 1.773335 | 0.010474 | 0.991643 | 0.999203 |
| CG10803 | 1460.413 | 0.010658 | 0.873688 | 0.012199 | 0.990267 | 0.999203 |
| CG33080 | 4088.447 | 0.006942 | 0.739757 | 0.009384 | 0.992513 | 0.999203 |
| CG3781 | 557.7015 | -0.0163 | 1.905659 | -0.00855 | 0.993175 | 0.999203 |
| CG15894 | 2081.362 | 0.009319 | 0.796329 | 0.011703 | 0.990663 | 0.999203 |
| laza | 525.9134 | 0.007274 | 1.436306 | 0.005064 | 0.995959 | 0.999203 |

| | | | | | | |
|---|---|---|---|---|---|---|
| Snm1 | 116.279 | -0.02261 | 3.749471 | -0.00603 | 0.995189 | 0.999203 |
| CG31915 | 1561.981 | 0.009923 | 0.86921 | 0.011416 | 0.990892 | 0.999203 |
| CG31710 | 1545.433 | 0.013636 | 1.397372 | 0.009758 | 0.992214 | 0.999203 |
| ThrRS | 2412.703 | -0.00426 | 0.711293 | -0.00599 | 0.995218 | 0.999203 |
| DCTN5-p2! | 631.2067 | -0.01032 | 1.213492 | -0.0085 | 0.993214 | 0.999203 |
| Ref1 | 3487.729 | 0.008849 | 0.658349 | 0.013442 | 0.989275 | 0.999203 |
| dsx | 2397.043 | 0.008592 | 0.879 | 0.009775 | 0.992201 | 0.999203 |
| CG9922 | 796.2831 | -0.00527 | 1.154906 | -0.00456 | 0.996358 | 0.999203 |
| CG31344 | 6948.334 | 0.002879 | 0.612642 | 0.0047 | 0.99625 | 0.999203 |
| Pak3 | 2840.77 | 0.008495 | 0.661442 | 0.012844 | 0.989753 | 0.999203 |
| CG31229 | 265.1633 | -0.01356 | 1.825456 | -0.00743 | 0.994075 | 0.999203 |
| CG6178 | 134.014 | 0.017515 | 2.259127 | 0.007753 | 0.993814 | 0.999203 |
| fs(1)h | 17161.28 | -0.00559 | 0.528427 | -0.01058 | 0.991557 | 0.999203 |
| CG12123 | 1479.161 | 0.004994 | 0.861989 | 0.005793 | 0.995378 | 0.999203 |
| stum | 283.1163 | 0.015998 | 2.53394 | 0.006314 | 0.994963 | 0.999203 |
| O-fut1 | 460.6669 | -0.01811 | 1.647496 | -0.01099 | 0.991231 | 0.999203 |
| CG17716 | 4028.417 | 0.005521 | 0.939877 | 0.005874 | 0.995313 | 0.999203 |
| ZnT49B | 1696.653 | -0.00417 | 0.791884 | -0.00526 | 0.995802 | 0.999203 |
| Nup188 | 93.71843 | -0.02646 | 3.389874 | -0.00781 | 0.993772 | 0.999203 |
| wal | 1949.834 | 0.004625 | 0.785365 | 0.005889 | 0.995301 | 0.999203 |
| Dbp45A | 1004.47 | -0.01679 | 1.761395 | -0.00953 | 0.992395 | 0.999203 |
| CG12769 | 3603.769 | -0.0034 | 0.637792 | -0.00533 | 0.995744 | 0.999203 |
| CG11141 | 1361.854 | 0.017718 | 1.424818 | 0.012435 | 0.990078 | 0.999203 |
| ABCD | 6845.658 | 0.002697 | 0.568282 | 0.004747 | 0.996213 | 0.999203 |
| CG9776 | 2412.109 | -0.00551 | 0.736314 | -0.00748 | 0.994035 | 0.999203 |
| CG18003 | 416.2089 | -0.03504 | 2.545382 | -0.01377 | 0.989016 | 0.999203 |
| loqs | 1629.473 | 0.008546 | 1.128233 | 0.007574 | 0.993957 | 0.999203 |
| RSG7 | 3810.092 | 0.002943 | 0.644149 | 0.004569 | 0.996354 | 0.999203 |
| Pglym78 | 20283.09 | 0.004732 | 0.419809 | 0.011273 | 0.991006 | 0.999203 |
| Saf-B | 5249.825 | 0.004328 | 0.651754 | 0.00664 | 0.994702 | 0.999203 |
| Epac | 7417.918 | 0.002585 | 0.583178 | 0.004432 | 0.996464 | 0.999203 |
| Ziz | 6646.288 | -0.00269 | 0.549531 | -0.00489 | 0.996098 | 0.999203 |
| CG17683 | 3907.747 | -0.00477 | 0.605554 | -0.00787 | 0.993717 | 0.999203 |
| MtnE | 90.77983 | 0.016519 | 3.334148 | 0.004954 | 0.996047 | 0.999203 |
| p130CAS | 2661.874 | 0.008881 | 0.725225 | 0.012246 | 0.99023 | 0.999203 |
| Gk1 | 2759.957 | 0.0049 | 0.788607 | 0.006213 | 0.995043 | 0.999203 |
| ND-30 | 10386.93 | 0.006817 | 0.500172 | 0.01363 | 0.989125 | 0.999203 |
| Ggamma3( | 53843.4 | -0.00284 | 0.376771 | -0.00754 | 0.993981 | 0.999203 |
| sbb | 10395.52 | -0.00269 | 0.49468 | -0.00543 | 0.995668 | 0.999203 |
| scb | 566.415 | -0.00724 | 1.29371 | -0.0056 | 0.995534 | 0.999203 |
| CG1275 | 4270.676 | 0.005173 | 0.591471 | 0.008746 | 0.993022 | 0.999203 |
| Ythdc1 | 3527.538 | 0.003294 | 0.661895 | 0.004977 | 0.996029 | 0.999203 |
| Ras64B | 2930.519 | -0.00654 | 0.674904 | -0.00969 | 0.992269 | 0.999203 |
| ImpL2 | 413.5248 | -0.00857 | 1.541915 | -0.00556 | 0.995566 | 0.999203 |
| Cip4 | 4343.062 | -0.00422 | 0.587411 | -0.00719 | 0.994264 | 0.999203 |
| CG11122 | 1164.187 | -0.00962 | 1.055253 | -0.00911 | 0.992729 | 0.999203 |
| CG2691 | 399.2411 | 0.011538 | 1.427901 | 0.008081 | 0.993553 | 0.999203 |
| CG6617 | 1228.655 | -0.00843 | 0.920712 | -0.00916 | 0.992692 | 0.999203 |
| rictor | 736.9918 | 0.019361 | 1.510859 | 0.012815 | 0.989776 | 0.999203 |
| CG3961 | 1121.888 | -0.01385 | 1.085179 | -0.01276 | 0.989819 | 0.999203 |

| | | | | | | |
|---|---|---|---|---|---|---|
| Ube3a | 692.1861 | 0.01944 | 1.671134 | 0.011633 | 0.990719 | 0.999203 |
| PGRP-LF | 75.23407 | 0.040167 | 3.930881 | 0.010218 | 0.991847 | 0.999203 |
| CG7182 | 504.8238 | -0.01399 | 1.34855 | -0.01037 | 0.991726 | 0.999203 |
| l(1)G0196 | 5566.315 | -0.00707 | 0.531106 | -0.01331 | 0.98938 | 0.999203 |
| Tango14 | 332.2792 | 0.008939 | 1.819431 | 0.004913 | 0.99608 | 0.999203 |
| Catsup | 1218.332 | 0.003373 | 0.939086 | 0.003592 | 0.997134 | 0.999242 |
| l(3)87Df | 1796.632 | 0.002625 | 0.772122 | 0.003399 | 0.997288 | 0.999242 |
| ogre | 6798.114 | -0.00241 | 0.683194 | -0.00353 | 0.99718 | 0.999242 |
| CG34393 | 450.3883 | -0.0049 | 1.334805 | -0.00367 | 0.997073 | 0.999242 |
| Nedd4 | 4551.866 | -0.00255 | 0.745536 | -0.00342 | 0.997271 | 0.999242 |
| CG9921 | 6665.514 | 0.002082 | 0.55637 | 0.003742 | 0.997015 | 0.999242 |
| RpS19a | 11931.17 | 0.001584 | 0.481312 | 0.00329 | 0.997375 | 0.999242 |
| vkg | 5180.042 | 0.001231 | 0.55241 | 0.002228 | 0.998222 | 0.999343 |
| Slmap | 1771.369 | 0.002007 | 0.815715 | 0.00246 | 0.998037 | 0.999343 |
| CG34289 | 69.06509 | 0.010579 | 3.931524 | 0.002691 | 0.997853 | 0.999343 |
| tweek | 3966.173 | -0.00176 | 0.7634 | -0.00231 | 0.998158 | 0.999343 |
| kibra | 2519.505 | 0.001909 | 0.728806 | 0.00262 | 0.99791 | 0.999343 |
| ITP | 1537.586 | -0.00289 | 1.074146 | -0.00269 | 0.997851 | 0.999343 |
| CG1785 | 101.2624 | -0.00577 | 2.83871 | -0.00203 | 0.998378 | 0.999374 |
| Btk29A | 1862.336 | -0.00125 | 0.875127 | -0.00143 | 0.998861 | 0.999521 |
| l(3)mbt | 577.098 | 0.002484 | 1.485544 | 0.001672 | 0.998666 | 0.999521 |
| CG11961 | 202.2453 | -0.00345 | 2.500199 | -0.00138 | 0.998899 | 0.999521 |
| CG1677 | 3680.574 | 0.000741 | 0.671747 | 0.001103 | 0.99912 | 0.999618 |
| l(2)05287 | 216.8852 | -0.00131 | 1.931712 | -0.00068 | 0.999461 | 0.99971 |
| CG1537 | 5892.174 | 0.000401 | 0.580192 | 0.000692 | 0.999448 | 0.99971 |
| CG46281 | 166.8387 | 0.001964 | 3.825087 | 0.000513 | 0.99959 | 0.999715 |
| CG2938 | 1315.862 | -4.50E-05 | 1.44339 | -3.12E-05 | 0.999975 | 0.999975 |
| tRNA:Pro-( | 0 | NA | NA | NA | NA | NA |
| CG9570 | 0 | NA | NA | NA | NA | NA |
| Or19b | 0 | NA | NA | NA | NA | NA |
| CG15322 | 0.596853 | 0.454699 | 3.988272 | 0.114009 | 0.909231 | NA |
| Or19a | 0 | NA | NA | NA | NA | NA |
| karr | 0 | NA | NA | NA | NA | NA |
| fd19B | 0 | NA | NA | NA | NA | NA |
| CG17003 | 53.64368 | -2.27518 | 3.952705 | -0.5756 | 0.564885 | NA |
| CG12655 | 0 | NA | NA | NA | NA | NA |
| CR32821 | 0 | NA | NA | NA | NA | NA |
| CG3777 | 0 | NA | NA | NA | NA | NA |
| CG12470 | 5.943496 | -1.90071 | 3.957353 | -0.4803 | 0.631016 | NA |
| ac | 0 | NA | NA | NA | NA | NA |
| sc | 0.69928 | 1.019866 | 4.009445 | 0.254366 | 0.799213 | NA |
| l(1)sc | 2.509476 | -1.87226 | 3.965749 | -0.47211 | 0.636851 | NA |
| ase | 0 | NA | NA | NA | NA | NA |
| CG13373 | 0 | NA | NA | NA | NA | NA |
| CG32817 | 2.251365 | 0.241545 | 3.960486 | 0.060989 | 0.951368 | NA |
| CG3176 | 47.98665 | 1.672485 | 3.412548 | 0.490099 | 0.624064 | NA |
| arg | 25.54678 | 1.746595 | 3.952934 | 0.441848 | 0.658599 | NA |
| Cyp4g1 | 0 | NA | NA | NA | NA | NA |
| Pgcl | 0 | NA | NA | NA | NA | NA |
| CG32816 | 12.33805 | -2.25201 | 3.958532 | -0.5689 | 0.569424 | NA |

| | | | | | | |
|---|---|---|---|---|---|---|
| CG3038 | 0.5235 | 1.490714 | 4.02555 | 0.370313 | 0.711149 | NA |
| fz3 | 1.827476 | -1.96736 | 3.880646 | -0.50697 | 0.612179 | NA |
| CG14635 | 0 | NA | NA | NA | NA | NA |
| CG13358 | 0.178053 | -1.16467 | 4.075067 | -0.2858 | 0.775028 | NA |
| CG13359 | 0 | NA | NA | NA | NA | NA |
| CG13360 | 0.1047 | 0.613261 | 4.062799 | 0.150945 | 0.880019 | NA |
| CG13361 | 0.132078 | -1.13211 | 4.069918 | -0.27817 | 0.780885 | NA |
| CG13362 | 16.45073 | -2.25935 | 3.956672 | -0.57102 | 0.567983 | NA |
| CG14634 | 0 | NA | NA | NA | NA | NA |
| CG11664 | 0 | NA | NA | NA | NA | NA |
| CG11663 | 0 | NA | NA | NA | NA | NA |
| mus81 | 29.37713 | 5.548322 | 3.87554 | 1.431625 | 0.152251 | NA |
| CG18823 | 0 | NA | NA | NA | NA | NA |
| CG14631 | 0 | NA | NA | NA | NA | NA |
| CG14629 | 0 | NA | NA | NA | NA | NA |
| CG14628 | 0 | NA | NA | NA | NA | NA |
| TfIIA-S-2 | 0.178812 | -1.08808 | 4.061735 | -0.26789 | 0.788788 | NA |
| CG3690 | 0.666238 | -2.57811 | 4.01049 | -0.64284 | 0.520326 | NA |
| CG3699 | 0 | NA | NA | NA | NA | NA |
| CG3708 | 0 | NA | NA | NA | NA | NA |
| CG3713 | 29.55684 | -1.70427 | 3.952205 | -0.43122 | 0.666308 | NA |
| CG32814 | 14.97075 | -3.48987 | 3.904596 | -0.89379 | 0.371436 | NA |
| CG11379 | 0 | NA | NA | NA | NA | NA |
| CG14627 | 0 | NA | NA | NA | NA | NA |
| CG14626 | 7.000117 | -1.904 | 3.956398 | -0.48124 | 0.630343 | NA |
| CG11380 | 0.233093 | 0.692656 | 4.076246 | 0.169925 | 0.865069 | NA |
| CG14625 | 0 | NA | NA | NA | NA | NA |
| CG11381 | 0 | NA | NA | NA | NA | NA |
| CG14624 | 0 | NA | NA | NA | NA | NA |
| CG11382 | 0 | NA | NA | NA | NA | NA |
| fs(1)N | 15.1631 | 3.24396 | 3.876965 | 0.836727 | 0.402746 | NA |
| Tsp2A | 0.356107 | -1.47099 | 4.037764 | -0.36431 | 0.715628 | NA |
| CG14770 | 7.846744 | 1.52291 | 3.955441 | 0.385017 | 0.700225 | NA |
| png | 14.49113 | 3.125105 | 3.87146 | 0.807216 | 0.419542 | NA |
| CG11409 | 0 | NA | NA | NA | NA | NA |
| CG32811 | 6.52661 | 1.15925 | 3.958017 | 0.292887 | 0.769609 | NA |
| CG11384 | 1.127155 | -1.04467 | 3.985057 | -0.26215 | 0.793209 | NA |
| eIF4E7 | 0 | NA | NA | NA | NA | NA |
| CG14780 | 0 | NA | NA | NA | NA | NA |
| dor | 33.48112 | 6.069143 | 3.829785 | 1.584722 | 0.11303 | NA |
| CG14797 | 14.67598 | 4.458059 | 3.84997 | 1.157946 | 0.246886 | NA |
| b6 | 26.45774 | 1.674484 | 3.952757 | 0.423624 | 0.67184 | NA |
| Scgdelta | 6.972379 | 2.008518 | 3.833463 | 0.523943 | 0.600318 | NA |
| CG3795 | 0 | NA | NA | NA | NA | NA |
| CG14795 | 0 | NA | NA | NA | NA | NA |
| snoRNA:M | 0 | NA | NA | NA | NA | NA |
| CG32808 | 41.08399 | 0.260724 | 3.835126 | 0.067983 | 0.945799 | NA |
| O-fut2 | 24.43814 | -4.11311 | 3.80769 | -1.08021 | 0.280048 | NA |
| CG14787 | 8.033656 | 5.68891 | 3.91627 | 1.452635 | 0.146325 | NA |
| CG32806 | 0 | NA | NA | NA | NA | NA |

| | | | | | | |
|---|---|---|---|---|---|---|
| CG4194 | 11.1405 | -3.47524 | 3.907049 | -0.88948 | 0.373746 | NA |
| CG14054 | 0 | NA | NA | NA | NA | NA |
| CG4325 | 10.46999 | 1.73528 | 3.955796 | 0.438668 | 0.660902 | NA |
| CG3857 | 49.14536 | -5.04624 | 3.804291 | -1.32646 | 0.184687 | NA |
| CG32801 | 0 | NA | NA | NA | NA | NA |
| CG14811 | 0.411147 | -0.29058 | 4.046973 | -0.0718 | 0.94276 | NA |
| Cyp4d2 | 0 | NA | NA | NA | NA | NA |
| Cyp4ae1 | 40.26957 | 1.239645 | 3.835228 | 0.323226 | 0.746524 | NA |
| CG17777 | 0 | NA | NA | NA | NA | NA |
| CG3457 | 0.396233 | -1.6814 | 4.028351 | -0.41739 | 0.676392 | NA |
| Mct1 | 23.21806 | -0.41976 | 3.733453 | -0.11243 | 0.910482 | NA |
| CG14052 | 0.178812 | -1.08808 | 4.061735 | -0.26789 | 0.788788 | NA |
| Or2a | 0 | NA | NA | NA | NA | NA |
| CG2854 | 0.534919 | -2.31746 | 4.015737 | -0.5771 | 0.563875 | NA |
| FarO | 4.947763 | -4.01998 | 3.888438 | -1.03383 | 0.301216 | NA |
| CG14053 | 0.712213 | -1.57119 | 3.998613 | -0.39293 | 0.694369 | NA |
| Cyp4d14 | 0.132078 | -1.13211 | 4.069918 | -0.27817 | 0.780885 | NA |
| Seipin | 51.68751 | 0.221951 | 3.857521 | 0.057537 | 0.954117 | NA |
| Klp3A | 0 | NA | NA | NA | NA | NA |
| Vha36-3 | 54.53617 | 1.513311 | 3.951557 | 0.382966 | 0.701745 | NA |
| CG12496 | 0.53416 | -1.53481 | 4.012395 | -0.38252 | 0.702078 | NA |
| CG2694 | 6.121549 | -3.43093 | 3.914637 | -0.87644 | 0.380793 | NA |
| fs(1)Ya | 4.225249 | -5.62961 | 3.918506 | -1.43667 | 0.150811 | NA |
| CG12498 | 0 | NA | NA | NA | NA | NA |
| dyw | 46.29056 | 0.844852 | 3.951506 | 0.213805 | 0.830699 | NA |
| HLH3B | 5.594238 | 1.155882 | 3.959178 | 0.29195 | 0.770325 | NA |
| ng1 | 0 | NA | NA | NA | NA | NA |
| Sgs4 | 0 | NA | NA | NA | NA | NA |
| CG10801 | 0 | NA | NA | NA | NA | NA |
| CG12206 | 1.007638 | -1.13898 | 3.915925 | -0.29086 | 0.771159 | NA |
| CG16781 | 0 | NA | NA | NA | NA | NA |
| CG16782 | 0 | NA | NA | NA | NA | NA |
| CG14269 | 0 | NA | NA | NA | NA | NA |
| tRNA:Gln-⁻ | 0 | NA | NA | NA | NA | NA |
| CG12535 | 27.93449 | 2.769589 | 3.917916 | 0.706904 | 0.479626 | NA |
| CG10793 | 5.081286 | 4.061506 | 3.872903 | 1.048698 | 0.294317 | NA |
| Pig1 | 0 | NA | NA | NA | NA | NA |
| ng4 | 0 | NA | NA | NA | NA | NA |
| ng3 | 0 | NA | NA | NA | NA | NA |
| ng2 | 2.775422 | -0.26042 | 3.958314 | -0.06579 | 0.947545 | NA |
| CG14265 | 28.31047 | -1.7041 | 3.952262 | -0.43117 | 0.666345 | NA |
| CG3526 | 0.337716 | 1.464149 | 4.04412 | 0.362044 | 0.717319 | NA |
| CG32793 | 12.66808 | 5.031562 | 3.897416 | 1.290999 | 0.196704 | NA |
| CG3592 | 0 | NA | NA | NA | NA | NA |
| CG3598 | 0.178053 | -1.16467 | 4.075067 | -0.2858 | 0.775028 | NA |
| CG14420 | 0.565506 | -0.26224 | 3.986711 | -0.06578 | 0.947554 | NA |
| CG14421 | 0 | NA | NA | NA | NA | NA |
| CG14422 | 0.356866 | -1.92326 | 4.037889 | -0.4763 | 0.633858 | NA |
| CG14423 | 0.282677 | -0.09599 | 4.057934 | -0.02366 | 0.981127 | NA |
| CG17959 | 0 | NA | NA | NA | NA | NA |

| | | | | | | |
|---|---|---|---|---|---|---|
| CG4116 | 0 | NA | NA | NA | NA | NA |
| CG14424 | 0.1047 | 0.613261 | 4.062799 | 0.150945 | 0.880019 | NA |
| CG14419 | 0.178053 | -1.16467 | 4.075067 | -0.2858 | 0.775028 | NA |
| CG14418 | 0 | NA | NA | NA | NA | NA |
| CG14417 | 0.53416 | -1.53481 | 4.012395 | -0.38252 | 0.702078 | NA |
| CG14416 | 0 | NA | NA | NA | NA | NA |
| CG13021 | 0 | NA | NA | NA | NA | NA |
| Ilp7 | 30.57238 | 1.747916 | 3.952603 | 0.442219 | 0.658331 | NA |
| Gas8 | 0 | NA | NA | NA | NA | NA |
| tRNA:Gln-( | 1.120046 | 2.129602 | 3.955197 | 0.538431 | 0.590279 | NA |
| tRNA:Pro-( | 0 | NA | NA | NA | NA | NA |
| CG15239 | 0.178812 | -1.08808 | 4.061735 | -0.26789 | 0.788788 | NA |
| Gem4b | 0.523116 | 1.34877 | 4.007682 | 0.336546 | 0.736459 | NA |
| Gem4c | 0 | NA | NA | NA | NA | NA |
| CG6414 | 0 | NA | NA | NA | NA | NA |
| CG15375 | 0.1047 | 0.613261 | 4.062799 | 0.150945 | 0.880019 | NA |
| dgt4 | 0.132078 | -1.13211 | 4.069918 | -0.27817 | 0.780885 | NA |
| CG15570 | 0.2094 | 1.005034 | 4.045341 | 0.248442 | 0.803792 | NA |
| CG12691 | 0 | NA | NA | NA | NA | NA |
| CG15572 | 0 | NA | NA | NA | NA | NA |
| CG12688 | 0 | NA | NA | NA | NA | NA |
| CG12693 | 0 | NA | NA | NA | NA | NA |
| mei-9 | 15.5429 | -6.22776 | 3.891624 | -1.6003 | 0.109532 | NA |
| CG15579 | 0 | NA | NA | NA | NA | NA |
| CG12692 | 0.628123 | 2.274123 | 4.000539 | 0.568454 | 0.569727 | NA |
| CG15578 | 1.340183 | 1.096454 | 3.924919 | 0.279357 | 0.779971 | NA |
| CG15577 | 0 | NA | NA | NA | NA | NA |
| CG15576 | 0.283512 | -0.2331 | 4.065094 | -0.05734 | 0.954273 | NA |
| Muc4B | 0 | NA | NA | NA | NA | NA |
| Femcoat | 0 | NA | NA | NA | NA | NA |
| CG15571 | 0 | NA | NA | NA | NA | NA |
| CG3546 | 0 | NA | NA | NA | NA | NA |
| lncRNA:CR | 24.15861 | 1.618454 | 3.937837 | 0.411001 | 0.681072 | NA |
| CG7024 | 54.19783 | -3.52218 | 3.899278 | -0.90329 | 0.366371 | NA |
| CG6978 | 0.2094 | 1.005034 | 4.045341 | 0.248442 | 0.803792 | NA |
| CG12682 | 0 | NA | NA | NA | NA | NA |
| boil | 0.282677 | -0.09599 | 4.057934 | -0.02366 | 0.981127 | NA |
| CG6927 | 0.178053 | -1.16467 | 4.075067 | -0.2858 | 0.775028 | NA |
| CG12680 | 0.411147 | -0.29058 | 4.046973 | -0.0718 | 0.94276 | NA |
| CG15468 | 0 | NA | NA | NA | NA | NA |
| CG15471 | 0 | NA | NA | NA | NA | NA |
| CG2871 | 0.178053 | -1.16467 | 4.075067 | -0.2858 | 0.775028 | NA |
| Klf15 | 0 | NA | NA | NA | NA | NA |
| CG15472 | 0 | NA | NA | NA | NA | NA |
| CG15465 | 0 | NA | NA | NA | NA | NA |
| dhd | 0 | NA | NA | NA | NA | NA |
| CG4198 | 0 | NA | NA | NA | NA | NA |
| CG17764 | 10.03708 | -1.02004 | 3.747304 | -0.27221 | 0.785464 | NA |
| CG3323 | 0 | NA | NA | NA | NA | NA |
| CG16756 | 0 | NA | NA | NA | NA | NA |

| | | | | | | |
|---|---|---|---|---|---|---|
| CG4151 | 0 | NA | NA | NA | NA | NA |
| Pop1 | 54.83023 | -0.24009 | 3.230275 | -0.07433 | 0.940751 | NA |
| CG32762 | 0 | NA | NA | NA | NA | NA |
| frma | 0.469871 | 0.864471 | 4.023538 | 0.214854 | 0.829881 | NA |
| NAAT1 | 30.88003 | 0.285581 | 3.838183 | 0.074405 | 0.940688 | NA |
| Gr5a | 0 | NA | NA | NA | NA | NA |
| cv | 6.626178 | 0.834101 | 3.855968 | 0.216314 | 0.828743 | NA |
| CG15773 | 0.178053 | -1.16467 | 4.075067 | -0.2858 | 0.775028 | NA |
| CG15767 | 0 | NA | NA | NA | NA | NA |
| CG12239 | 14.31253 | -0.30075 | 3.845539 | -0.07821 | 0.937664 | NA |
| Cpr5C | 0.356107 | -1.47099 | 4.037764 | -0.36431 | 0.715628 | NA |
| Prosbeta2l | 0 | NA | NA | NA | NA | NA |
| tRNA:Glu-( | 0 | NA | NA | NA | NA | NA |
| CG12729 | 0 | NA | NA | NA | NA | NA |
| CG15764 | 0.356107 | -1.47099 | 4.037764 | -0.36431 | 0.715628 | NA |
| CG4020 | 54.23456 | 1.750878 | 3.951866 | 0.443051 | 0.657729 | NA |
| AgmNAT | 0 | NA | NA | NA | NA | NA |
| Tre1 | 0.178053 | -1.16467 | 4.075067 | -0.2858 | 0.775028 | NA |
| CG32755 | 3.309582 | -0.26274 | 3.956962 | -0.0664 | 0.94706 | NA |
| CG6048 | 0 | NA | NA | NA | NA | NA |
| CG6067 | 0 | NA | NA | NA | NA | NA |
| Grip | 0.566948 | 0.490522 | 3.992077 | 0.122874 | 0.902207 | NA |
| CG5966 | 5.695097 | 5.005505 | 3.924253 | 1.275531 | 0.202121 | NA |
| CG5921 | 23.96085 | -2.26641 | 3.954895 | -0.57307 | 0.566601 | NA |
| Btnd | 0 | NA | NA | NA | NA | NA |
| vanin-like | 0.1047 | 0.613261 | 4.062799 | 0.150945 | 0.880019 | NA |
| CG32751 | 0 | NA | NA | NA | NA | NA |
| CG32750 | 43.45356 | -1.91996 | 3.951805 | -0.48584 | 0.627078 | NA |
| Mipp2 | 6.339729 | -1.90207 | 3.956958 | -0.48069 | 0.630738 | NA |
| raptor | 40.23547 | 2.505067 | 3.563025 | 0.703073 | 0.48201 | NA |
| fs(1)M3 | 0.178053 | -1.16467 | 4.075067 | -0.2858 | 0.775028 | NA |
| mldr | 0.209247 | 1.042804 | 4.047837 | 0.25762 | 0.7967 | NA |
| tRNA:Phe- | 0.2094 | 1.005034 | 4.045341 | 0.248442 | 0.803792 | NA |
| CG3342 | 48.16197 | 1.750376 | 3.951986 | 0.44291 | 0.65783 | NA |
| swa | 0 | NA | NA | NA | NA | NA |
| swaPsi | 0.178053 | -1.16467 | 4.075067 | -0.2858 | 0.775028 | NA |
| CG15880 | 0 | NA | NA | NA | NA | NA |
| Pkg21D | 0 | NA | NA | NA | NA | NA |
| lectin-21Cl | 0 | NA | NA | NA | NA | NA |
| CG33127 | 0 | NA | NA | NA | NA | NA |
| al | 0 | NA | NA | NA | NA | NA |
| CG13692 | 0.535678 | -2.26616 | 4.021604 | -0.5635 | 0.573098 | NA |
| BBS8 | 0.2094 | 1.005034 | 4.045341 | 0.248442 | 0.803792 | NA |
| Pph13 | 0 | NA | NA | NA | NA | NA |
| CG11835 | 0.894061 | -2.00006 | 4.030294 | -0.49626 | 0.619714 | NA |
| Ets21C | 0.63886 | -0.79045 | 3.986802 | -0.19827 | 0.842837 | NA |
| CG13690 | 0.31387 | 1.261782 | 4.037145 | 0.312543 | 0.754628 | NA |
| CG4213 | 0 | NA | NA | NA | NA | NA |
| Ent1 | 10.90318 | -2.34605 | 3.76509 | -0.62311 | 0.533214 | NA |
| CG11911 | 0.1047 | 0.613261 | 4.062799 | 0.150945 | 0.880019 | NA |

| | | | | | | |
|---|---|---|---|---|---|---|
| CG11912 | 3.140998 | 1.694072 | 3.966495 | 0.427096 | 0.66931 | NA |
| CG3345 | 0.178812 | -1.08808 | 4.061735 | -0.26789 | 0.788788 | NA |
| Amnionles | 1.570499 | 1.642769 | 3.980387 | 0.412716 | 0.679815 | NA |
| CG3709 | 34.57163 | 1.654503 | 3.812925 | 0.43392 | 0.664347 | NA |
| CG31975 | 0 | NA | NA | NA | NA | NA |
| ovm | 0 | NA | NA | NA | NA | NA |
| Ir21a | 0.132078 | -1.13211 | 4.069918 | -0.27817 | 0.780885 | NA |
| Spn77Bc | 47.18428 | -5.69436 | 3.830699 | -1.48651 | 0.137145 | NA |
| CG5665 | 27.59826 | -1.70399 | 3.952297 | -0.43114 | 0.666368 | NA |
| in | 0.528311 | -1.72839 | 4.011871 | -0.43082 | 0.6666 | NA |
| CG13247 | 0 | NA | NA | NA | NA | NA |
| zye | 60.78647 | -2.03529 | 3.538433 | -0.5752 | 0.565159 | NA |
| mag | 0 | NA | NA | NA | NA | NA |
| CG18281 | 4.866759 | -1.81691 | 3.846772 | -0.47232 | 0.636697 | NA |
| CG13250 | 0 | NA | NA | NA | NA | NA |
| knrl | 3.426486 | 0.40965 | 3.855283 | 0.106257 | 0.915379 | NA |
| CG5199 | 60.85185 | 0.417134 | 3.834227 | 0.108792 | 0.913367 | NA |
| Toll-9 | 0.178053 | -1.16467 | 4.075067 | -0.2858 | 0.775028 | NA |
| CG13251 | 0.178053 | -1.16467 | 4.075067 | -0.2858 | 0.775028 | NA |
| Spc105R | 0.178053 | -1.16467 | 4.075067 | -0.2858 | 0.775028 | NA |
| CG3288 | 0 | NA | NA | NA | NA | NA |
| CG11458 | 0.132078 | -1.13211 | 4.069918 | -0.27817 | 0.780885 | NA |
| CG3634 | 54.96167 | -3.78299 | 3.801658 | -0.99509 | 0.319693 | NA |
| CG3698 | 0.178053 | -1.16467 | 4.075067 | -0.2858 | 0.775028 | NA |
| Six4 | 0.1047 | 0.613261 | 4.062799 | 0.150945 | 0.880019 | NA |
| CG11396 | 0 | NA | NA | NA | NA | NA |
| Gr77a | 43.45138 | -2.2735 | 3.953123 | -0.57512 | 0.565213 | NA |
| kni | 1.279731 | 0.371782 | 3.896693 | 0.09541 | 0.923989 | NA |
| CG10588 | 0 | NA | NA | NA | NA | NA |
| CR33283 | 0 | NA | NA | NA | NA | NA |
| CG12984 | 22.88797 | -2.26568 | 3.955079 | -0.57285 | 0.566744 | NA |
| CG33284 | 28.27352 | -0.04896 | 3.951524 | -0.01239 | 0.990115 | NA |
| CG10510 | 1.056621 | -1.8126 | 3.984111 | -0.45496 | 0.64914 | NA |
| park | 62.01447 | -0.10618 | 3.097487 | -0.03428 | 0.972655 | NA |
| Ac78C | 52.4844 | -3.22874 | 3.659505 | -0.88229 | 0.377621 | NA |
| CG33286 | 0.461489 | -1.26887 | 4.028229 | -0.31499 | 0.752766 | NA |
| CG33287 | 0.178812 | -1.08808 | 4.061735 | -0.26789 | 0.788788 | NA |
| CG12983 | 49.00082 | -1.92032 | 3.951703 | -0.48595 | 0.627005 | NA |
| CG10587 | 18.62295 | -1.9158 | 3.952995 | -0.48464 | 0.627928 | NA |
| CG11037 | 9.773749 | -1.90931 | 3.95486 | -0.48278 | 0.629255 | NA |
| Sems | 0 | NA | NA | NA | NA | NA |
| Cpr78Cb | 0.53416 | -1.53481 | 4.012395 | -0.38252 | 0.702078 | NA |
| Cpr78Cc | 0 | NA | NA | NA | NA | NA |
| CG32436 | 0.53416 | -1.53481 | 4.012395 | -0.38252 | 0.702078 | NA |
| CG32437 | 0.1047 | 0.613261 | 4.062799 | 0.150945 | 0.880019 | NA |
| CG32440 | 0 | NA | NA | NA | NA | NA |
| ppk5 | 48.05124 | -9.28765 | 3.907707 | -2.37675 | 0.017466 | NA |
| CG33290 | 0 | NA | NA | NA | NA | NA |
| CG7634 | 22.19439 | 6.474651 | 3.90132 | 1.659605 | 0.096994 | NA |
| CG7173 | 0 | NA | NA | NA | NA | NA |

| | | | | | | |
|---|---|---|---|---|---|---|
| CG7202 | 2.824828 | 1.495401 | 3.963153 | 0.377326 | 0.705931 | NA |
| Neu2 | 38.09646 | -0.11829 | 3.951704 | -0.02993 | 0.97612 | NA |
| croc | 4.886874 | -1.89605 | 3.958711 | -0.47896 | 0.631969 | NA |
| sa | 0 | NA | NA | NA | NA | NA |
| CG12971 | 0 | NA | NA | NA | NA | NA |
| CG11309 | 9.06951 | 2.008509 | 3.807331 | 0.527537 | 0.597821 | NA |
| Cpr78Ca | 0.178053 | -1.16467 | 4.075067 | -0.2858 | 0.775028 | NA |
| CG12974 | 0 | NA | NA | NA | NA | NA |
| CG9389 | 47.39566 | -0.42051 | 3.834442 | -0.10967 | 0.912674 | NA |
| CG32445 | 0 | NA | NA | NA | NA | NA |
| CG11249 | 0 | NA | NA | NA | NA | NA |
| CG14573 | 0 | NA | NA | NA | NA | NA |
| CG14572 | 0 | NA | NA | NA | NA | NA |
| CG14564 | 0 | NA | NA | NA | NA | NA |
| CG7370 | 15.9305 | 3.526832 | 3.903133 | 0.90359 | 0.366213 | NA |
| eg | 0 | NA | NA | NA | NA | NA |
| CycH | 3.839014 | -1.92253 | 3.881852 | -0.49526 | 0.620415 | NA |
| CG14563 | 0 | NA | NA | NA | NA | NA |
| CG14565 | 0 | NA | NA | NA | NA | NA |
| CG14566 | 0 | NA | NA | NA | NA | NA |
| CG14567 | 0 | NA | NA | NA | NA | NA |
| CG14568 | 0 | NA | NA | NA | NA | NA |
| CG14569 | 0.1047 | 0.613261 | 4.062799 | 0.150945 | 0.880019 | NA |
| CG14570 | 0 | NA | NA | NA | NA | NA |
| CG32447 | 0 | NA | NA | NA | NA | NA |
| Cpr78E | 0 | NA | NA | NA | NA | NA |
| CR7448 | 0.233093 | 0.692656 | 4.076246 | 0.169925 | 0.865069 | NA |
| Chs2 | 0 | NA | NA | NA | NA | NA |
| P5CS | 0 | NA | NA | NA | NA | NA |
| msopa | 0 | NA | NA | NA | NA | NA |
| CG7140 | 5.831849 | 1.457586 | 3.95798 | 0.368265 | 0.712676 | NA |
| tRNA:Cys-( | 0 | NA | NA | NA | NA | NA |
| CG32450 | 18.95015 | 4.947279 | 3.886956 | 1.27279 | 0.203093 | NA |
| CG11437 | 0 | NA | NA | NA | NA | NA |
| Trxr-2 | 19.89299 | 1.744326 | 3.953505 | 0.44121 | 0.659061 | NA |
| ND-B14.5A | 0 | NA | NA | NA | NA | NA |
| CG14457 | 0.233093 | 0.692656 | 4.076246 | 0.169925 | 0.865069 | NA |
| CG14456 | 0 | NA | NA | NA | NA | NA |
| CG14453 | 0 | NA | NA | NA | NA | NA |
| CG14452 | 0 | NA | NA | NA | NA | NA |
| CG32453 | 0 | NA | NA | NA | NA | NA |
| CG11370 | 0.178053 | -1.16467 | 4.075067 | -0.2858 | 0.775028 | NA |
| CG11367 | 10.02176 | -3.15571 | 3.853013 | -0.81903 | 0.412772 | NA |
| CG32454 | 5.665974 | -0.13909 | 3.910837 | -0.03556 | 0.97163 | NA |
| CG14448 | 0.178812 | -1.08808 | 4.061735 | -0.26789 | 0.788788 | NA |
| l(3)04053 | 57.34553 | -1.70858 | 3.5258 | -0.48459 | 0.627964 | NA |
| CG14451 | 0 | NA | NA | NA | NA | NA |
| CG12546 | 0 | NA | NA | NA | NA | NA |
| CG14454 | 0 | NA | NA | NA | NA | NA |
| CG14455 | 0 | NA | NA | NA | NA | NA |

| | | | | | | |
|---|---|---|---|---|---|---|
| CG11226 | 0.31089 | -1.9003 | 4.033427 | -0.47114 | 0.637542 | NA |
| CG13239 | 0.178053 | -1.16467 | 4.075067 | -0.2858 | 0.775028 | NA |
| CG11109 | 0 | NA | NA | NA | NA | NA |
| Ssl1 | 39.55236 | 3.751206 | 3.819849 | 0.98203 | 0.326085 | NA |
| CG11131 | 0 | NA | NA | NA | NA | NA |
| BoYb | 0 | NA | NA | NA | NA | NA |
| CG32457 | 0.491923 | 0.364479 | 3.983057 | 0.091507 | 0.92709 | NA |
| CG32461 | 0.53416 | -1.53481 | 4.012395 | -0.38252 | 0.702078 | NA |
| snoRNA:M | 1.455566 | 3.686462 | 3.938983 | 0.935892 | 0.349329 | NA |
| CG32459 | 0.178053 | -1.16467 | 4.075067 | -0.2858 | 0.775028 | NA |
| CG32462 | 0 | NA | NA | NA | NA | NA |
| tRNA:Glu-( | 0 | NA | NA | NA | NA | NA |
| CR1075 | 0.282677 | -0.09599 | 4.057934 | -0.02366 | 0.981127 | NA |
| Obp83cd | 0 | NA | NA | NA | NA | NA |
| Osi1 | 0 | NA | NA | NA | NA | NA |
| CR31562 | 0 | NA | NA | NA | NA | NA |
| Osi24 | 18.81464 | 1.644989 | 3.953448 | 0.41609 | 0.677344 | NA |
| Osi2 | 0 | NA | NA | NA | NA | NA |
| Osi3 | 0 | NA | NA | NA | NA | NA |
| Osi5 | 0 | NA | NA | NA | NA | NA |
| Osi6 | 0.178053 | -1.16467 | 4.075067 | -0.2858 | 0.775028 | NA |
| Osi7 | 0.1047 | 0.613261 | 4.062799 | 0.150945 | 0.880019 | NA |
| Osi8 | 0 | NA | NA | NA | NA | NA |
| Osi9 | 0 | NA | NA | NA | NA | NA |
| Osi12 | 0.178053 | -1.16467 | 4.075067 | -0.2858 | 0.775028 | NA |
| CG15594 | 0.1047 | 0.613261 | 4.062799 | 0.150945 | 0.880019 | NA |
| Osi13 | 0 | NA | NA | NA | NA | NA |
| Osi14 | 0 | NA | NA | NA | NA | NA |
| Osi15 | 0.178812 | -1.08808 | 4.061735 | -0.26789 | 0.788788 | NA |
| Osi16 | 0.356107 | -1.47099 | 4.037764 | -0.36431 | 0.715628 | NA |
| CG31560 | 0 | NA | NA | NA | NA | NA |
| Osi17 | 0.356866 | -1.92326 | 4.037889 | -0.4763 | 0.633858 | NA |
| Osi18 | 0 | NA | NA | NA | NA | NA |
| CG15597 | 0 | NA | NA | NA | NA | NA |
| Osi11 | 0.178053 | -1.16467 | 4.075067 | -0.2858 | 0.775028 | NA |
| Osi4 | 6.938443 | 1.866466 | 3.767334 | 0.495434 | 0.620294 | NA |
| CG31559 | 0 | NA | NA | NA | NA | NA |
| Obp83g | 0 | NA | NA | NA | NA | NA |
| Obp83ef | 0.104623 | 0.639075 | 4.067674 | 0.157111 | 0.875158 | NA |
| Or83c | 0.178053 | -1.16467 | 4.075067 | -0.2858 | 0.775028 | NA |
| CG15580 | 0 | NA | NA | NA | NA | NA |
| CG11459 | 0 | NA | NA | NA | NA | NA |
| CG1208 | 59.85759 | -6.72687 | 3.865653 | -1.74016 | 0.08183 | NA |
| CG14676 | 0 | NA | NA | NA | NA | NA |
| CR10991 | 0 | NA | NA | NA | NA | NA |
| CG10280 | 0 | NA | NA | NA | NA | NA |
| Obp83b | 0 | NA | NA | NA | NA | NA |
| Obp83a | 0 | NA | NA | NA | NA | NA |
| CG11373 | 0 | NA | NA | NA | NA | NA |
| Atu | 0 | NA | NA | NA | NA | NA |

| | | | | | | |
|---|---|---|---|---|---|---|
| CG31551 | 0 | NA | NA | NA | NA | NA |
| CG2100 | 31.94744 | 6.509401 | 3.877081 | 1.678944 | 0.093163 | NA |
| cas | 47.91745 | 1.536711 | 3.951659 | 0.388877 | 0.697367 | NA |
| CG2104 | 0.310131 | -1.94616 | 4.045058 | -0.48112 | 0.630431 | NA |
| CG31546 | 22.47914 | 1.661438 | 3.953062 | 0.420291 | 0.674273 | NA |
| NKCC | 24.38506 | 1.336043 | 3.952464 | 0.338028 | 0.735342 | NA |
| HDAC3 | 0 | NA | NA | NA | NA | NA |
| CG12590 | 0 | NA | NA | NA | NA | NA |
| CG1113 | 41.32956 | 6.946828 | 3.877636 | 1.791511 | 0.073211 | NA |
| CG31542 | 0 | NA | NA | NA | NA | NA |
| tRNA:Arg-ˉ | 0.446101 | 0.990384 | 3.987592 | 0.248366 | 0.803851 | NA |
| CG2663 | 0 | NA | NA | NA | NA | NA |
| Or83a | 0 | NA | NA | NA | NA | NA |
| kkv | 59.96358 | -1.38319 | 3.072624 | -0.45017 | 0.652591 | NA |
| CG14668 | 0 | NA | NA | NA | NA | NA |
| CG31538 | 53.86997 | 8.421106 | 3.845727 | 2.189731 | 0.028544 | NA |
| Tim17a2 | 0.132078 | -1.13211 | 4.069918 | -0.27817 | 0.780885 | NA |
| Prosbeta2l | 0 | NA | NA | NA | NA | NA |
| CG1124 | 10.32058 | -3.05598 | 3.917683 | -0.78005 | 0.435363 | NA |
| CG12589 | 0 | NA | NA | NA | NA | NA |
| plh | 0 | NA | NA | NA | NA | NA |
| CG14662 | 0 | NA | NA | NA | NA | NA |
| CG14661 | 17.83634 | -4.83024 | 3.877421 | -1.24574 | 0.212862 | NA |
| snRNA:U1: | 27.01258 | 1.74703 | 3.952825 | 0.44197 | 0.658511 | NA |
| laf | 0.236778 | -0.26946 | 4.071573 | -0.06618 | 0.947234 | NA |
| slx1 | 0.356866 | -1.92326 | 4.037889 | -0.4763 | 0.633858 | NA |
| rpk | 0 | NA | NA | NA | NA | NA |
| spartin | 20.22196 | 2.028623 | 3.773879 | 0.537543 | 0.590893 | NA |
| BBS5 | 0 | NA | NA | NA | NA | NA |
| CG14659 | 0 | NA | NA | NA | NA | NA |
| CG14658 | 0 | NA | NA | NA | NA | NA |
| Hus1-like | 42.38096 | -1.85567 | 3.812901 | -0.48668 | 0.626484 | NA |
| CG17387 | 0.357625 | -1.47707 | 4.044794 | -0.36518 | 0.714979 | NA |
| CG14655 | 43.43676 | 0.361818 | 3.559478 | 0.101649 | 0.919035 | NA |
| CG32945 | 0 | NA | NA | NA | NA | NA |
| MP1 | 0 | NA | NA | NA | NA | NA |
| CG14645 | 0 | NA | NA | NA | NA | NA |
| CG31528 | 3.769197 | 1.70343 | 3.964029 | 0.429722 | 0.667398 | NA |
| CG31525 | 0.178053 | -1.16467 | 4.075067 | -0.2858 | 0.775028 | NA |
| CG1074 | 7.262137 | 0.144289 | 3.84494 | 0.037527 | 0.970065 | NA |
| CG14651 | 0.565506 | -0.26224 | 3.986711 | -0.06578 | 0.947554 | NA |
| lncRNA:CR | 0.104623 | 0.639075 | 4.067674 | 0.157111 | 0.875158 | NA |
| Mur82C | 0 | NA | NA | NA | NA | NA |
| CG9804 | 18.03174 | -1.13858 | 3.800836 | -0.29956 | 0.764511 | NA |
| CG9855 | 25.27218 | 1.148413 | 3.839889 | 0.299074 | 0.764883 | NA |
| Skp2 | 0.233093 | 0.692656 | 4.076246 | 0.169925 | 0.865069 | NA |
| Vps24 | 32.38633 | 2.635777 | 3.757161 | 0.701534 | 0.48297 | NA |
| CG12581 | 0.5892 | -0.44935 | 3.992199 | -0.11256 | 0.910381 | NA |
| TwdlU | 0.365171 | -0.43584 | 4.056359 | -0.10745 | 0.914434 | NA |
| TwdlF | 0.3141 | 1.240997 | 4.03841 | 0.307299 | 0.758616 | NA |

| | | | | | | |
|---|---|---|---|---|---|---|
| TwdlV | 0 | NA | NA | NA | NA | NA |
| Or82a | 0 | NA | NA | NA | NA | NA |
| CG1092 | 49.59256 | 7.483543 | 3.795403 | 1.971739 | 0.048639 | NA |
| CG31516 | 0.283436 | -0.20033 | 4.070792 | -0.04921 | 0.960751 | NA |
| CG14636 | 0 | NA | NA | NA | NA | NA |
| Dsk | 2.377398 | -1.86964 | 3.966533 | -0.47135 | 0.637388 | NA |
| SP555 | 0 | NA | NA | NA | NA | NA |
| CG14036 | 0 | NA | NA | NA | NA | NA |
| CG31917 | 0 | NA | NA | NA | NA | NA |
| tRNA:Asp-( | 0 | NA | NA | NA | NA | NA |
| TotM | 35.38858 | 1.748836 | 3.952374 | 0.442477 | 0.658144 | NA |
| CG14022 | 31.8287 | 2.968918 | 3.920455 | 0.757289 | 0.448877 | NA |
| tRNA:Asp-( | 0 | NA | NA | NA | NA | NA |
| CG18269 | 0 | NA | NA | NA | NA | NA |
| CG14017 | 0 | NA | NA | NA | NA | NA |
| snRNA:U4: | 0 | NA | NA | NA | NA | NA |
| CG18266 | 0.178812 | -1.08808 | 4.061735 | -0.26789 | 0.788788 | NA |
| CG12511 | 0 | NA | NA | NA | NA | NA |
| CG7251 | 0 | NA | NA | NA | NA | NA |
| CG14011 | 0 | NA | NA | NA | NA | NA |
| CG14013 | 0 | NA | NA | NA | NA | NA |
| CG14014 | 0.849019 | -0.87463 | 3.964455 | -0.22062 | 0.825391 | NA |
| tomb | 0 | NA | NA | NA | NA | NA |
| lncRNA:CR | 0.497249 | -0.69954 | 4.001026 | -0.17484 | 0.861205 | NA |
| CG7236 | 0.466186 | 0.959471 | 4.033986 | 0.237847 | 0.812 | NA |
| asRNA:CR3 | 0 | NA | NA | NA | NA | NA |
| CG31913 | 0 | NA | NA | NA | NA | NA |
| CG9029 | 0 | NA | NA | NA | NA | NA |
| CG14000 | 3.036298 | 1.692169 | 3.966999 | 0.426562 | 0.669699 | NA |
| CG9021 | 0 | NA | NA | NA | NA | NA |
| Acp26Aa | 0 | NA | NA | NA | NA | NA |
| Acp26Ab | 0.90868 | 2.395959 | 3.996542 | 0.599508 | 0.548834 | NA |
| CG31644 | 0 | NA | NA | NA | NA | NA |
| obst-E | 1.151699 | 1.609751 | 3.989673 | 0.403479 | 0.686596 | NA |
| CG14006 | 6.409473 | 1.348312 | 3.955806 | 0.340844 | 0.733221 | NA |
| CG11149 | 0 | NA | NA | NA | NA | NA |
| CG14007 | 0 | NA | NA | NA | NA | NA |
| Vm26Ab | 0 | NA | NA | NA | NA | NA |
| Ucp4C | 0 | NA | NA | NA | NA | NA |
| Ucp4B | 0 | NA | NA | NA | NA | NA |
| CG13996 | 0.178053 | -1.16467 | 4.075067 | -0.2858 | 0.775028 | NA |
| CG9222 | 0.178053 | -1.16467 | 4.075067 | -0.2858 | 0.775028 | NA |
| CG13991 | 0 | NA | NA | NA | NA | NA |
| Muc26B | 0 | NA | NA | NA | NA | NA |
| CG13989 | 0.178053 | -1.16467 | 4.075067 | -0.2858 | 0.775028 | NA |
| CG31639 | 0 | NA | NA | NA | NA | NA |
| CG9150 | 8.490945 | -1.76074 | 3.866851 | -0.45534 | 0.648863 | NA |
| slmo | 29.3746 | 0.187224 | 3.735255 | 0.050123 | 0.960024 | NA |
| CG9107 | 0.442493 | 1.796125 | 4.02306 | 0.446457 | 0.655267 | NA |
| H2.0 | 0.536437 | -1.71175 | 4.038042 | -0.42391 | 0.671635 | NA |

| | | | | | | |
|---|---|---|---|---|---|---|
| Gal | 0 | NA | NA | NA | NA | NA |
| CG13992 | 0.178053 | -1.16467 | 4.075067 | -0.2858 | 0.775028 | NA |
| Vm26Aa | 0 | NA | NA | NA | NA | NA |
| CG13998 | 0.3141 | 1.240997 | 4.03841 | 0.307299 | 0.758616 | NA |
| Ugt37B1 | 0 | NA | NA | NA | NA | NA |
| CG13984 | 0.104623 | 0.639075 | 4.067674 | 0.157111 | 0.875158 | NA |
| ppk14 | 0 | NA | NA | NA | NA | NA |
| slam | 0.596546 | 0.421256 | 3.969213 | 0.106131 | 0.915479 | NA |
| CG9500 | 0 | NA | NA | NA | NA | NA |
| CG9497 | 25.90313 | 6.186722 | 3.885785 | 1.592142 | 0.111353 | NA |
| CG13983 | 0 | NA | NA | NA | NA | NA |
| CG9550 | 19.33806 | 2.754453 | 3.923385 | 0.70206 | 0.482642 | NA |
| COX5BL | 0 | NA | NA | NA | NA | NA |
| cup | 0 | NA | NA | NA | NA | NA |
| TTLL3A | 0 | NA | NA | NA | NA | NA |
| AkhR | 0.178053 | -1.16467 | 4.075067 | -0.2858 | 0.775028 | NA |
| Tsp | 0.104623 | 0.639075 | 4.067674 | 0.157111 | 0.875158 | NA |
| CG11327 | 0.209247 | 1.042804 | 4.047837 | 0.25762 | 0.7967 | NA |
| Nse1 | 0 | NA | NA | NA | NA | NA |
| cort | 53.58419 | 0.255901 | 3.951247 | 0.064765 | 0.948361 | NA |
| CG13771 | 0 | NA | NA | NA | NA | NA |
| CG31636 | 0 | NA | NA | NA | NA | NA |
| DIP-iota | 1.602479 | -1.64094 | 3.97352 | -0.41297 | 0.67963 | NA |
| IFT52 | 0 | NA | NA | NA | NA | NA |
| tRNA:Arg-( | 0 | NA | NA | NA | NA | NA |
| CG11236 | 5.697705 | -1.68799 | 3.957546 | -0.42652 | 0.669726 | NA |
| sens-2 | 42.93819 | 8.601066 | 3.908009 | 2.200882 | 0.027744 | NA |
| uif | 0.565506 | -0.26224 | 3.986711 | -0.06578 | 0.947554 | NA |
| sip2 | 1.691239 | 2.13269 | 3.952787 | 0.539541 | 0.589514 | NA |
| CG13775 | 42.02089 | -2.2732 | 3.953197 | -0.57503 | 0.565272 | NA |
| CG18304 | 0.178053 | -1.16467 | 4.075067 | -0.2858 | 0.775028 | NA |
| CG32829 | 16.43789 | 1.742188 | 3.954044 | 0.440609 | 0.659496 | NA |
| Nha1 | 30.21928 | -2.26967 | 3.95408 | -0.57401 | 0.565963 | NA |
| CG17375 | 0 | NA | NA | NA | NA | NA |
| CG17376 | 22.95625 | -0.30077 | 3.912171 | -0.07688 | 0.938719 | NA |
| nrv1 | 55.58639 | 0.250738 | 3.949678 | 0.063483 | 0.949382 | NA |
| LUBEL | 8.957704 | -0.65558 | 3.928837 | -0.16686 | 0.867477 | NA |
| CG11322 | 0 | NA | NA | NA | NA | NA |
| Ugt307A1 | 0 | NA | NA | NA | NA | NA |
| CG31909 | 5.530244 | -0.45535 | 3.868976 | -0.11769 | 0.906311 | NA |
| CG13786 | 0 | NA | NA | NA | NA | NA |
| CG4496 | 14.38375 | 0.953029 | 3.76105 | 0.253394 | 0.799963 | NA |
| Pvf2 | 0.178053 | -1.16467 | 4.075067 | -0.2858 | 0.775028 | NA |
| Nuf2 | 1.856259 | 0.798321 | 3.959062 | 0.201644 | 0.840195 | NA |
| Mst27D | 23.7504 | 5.293715 | 3.876771 | 1.365496 | 0.172097 | NA |
| Wnt10 | 0 | NA | NA | NA | NA | NA |
| ninaC | 17.27331 | 0.338997 | 3.841003 | 0.088257 | 0.929672 | NA |
| gudu | 0.264155 | -1.54099 | 4.048976 | -0.38059 | 0.703509 | NA |
| CG5171 | 0 | NA | NA | NA | NA | NA |
| CG5177 | 0 | NA | NA | NA | NA | NA |

| | | | | | | |
|---|---|---|---|---|---|---|
| CG15818 | 0 | NA | NA | NA | NA | NA |
| CG6055 | 0 | NA | NA | NA | NA | NA |
| CG6441 | 0 | NA | NA | NA | NA | NA |
| Obp28a | 0 | NA | NA | NA | NA | NA |
| Gr28b | 50.58672 | -1.62654 | 3.85182 | -0.42228 | 0.672821 | NA |
| Gr28a | 0.1047 | 0.613261 | 4.062799 | 0.150945 | 0.880019 | NA |
| CG5958 | 0 | NA | NA | NA | NA | NA |
| Spn28B | 0 | NA | NA | NA | NA | NA |
| Cyp4d21 | 0 | NA | NA | NA | NA | NA |
| CG6739 | 0.536437 | -1.71175 | 4.038042 | -0.42391 | 0.671635 | NA |
| Myo28B1 | 9.836325 | 3.136111 | 3.779057 | 0.829866 | 0.406615 | NA |
| Tep3 | 11.85614 | -0.27132 | 3.952006 | -0.06865 | 0.945264 | NA |
| ATPsynGL | 0.104623 | 0.639075 | 4.067674 | 0.157111 | 0.875158 | NA |
| Uro | 0 | NA | NA | NA | NA | NA |
| CG7164 | 0 | NA | NA | NA | NA | NA |
| CG7203 | 0 | NA | NA | NA | NA | NA |
| Acp1 | 0 | NA | NA | NA | NA | NA |
| CG13795 | 0 | NA | NA | NA | NA | NA |
| CG13794 | 0 | NA | NA | NA | NA | NA |
| CG13793 | 0 | NA | NA | NA | NA | NA |
| CG18585 | 0.104623 | 0.639075 | 4.067674 | 0.157111 | 0.875158 | NA |
| CG14537 | 0 | NA | NA | NA | NA | NA |
| tRNA:Gly-( | 0.903015 | -2.03286 | 3.962887 | -0.51297 | 0.607969 | NA |
| Tg | 15.44133 | 0.158208 | 3.8378 | 0.041224 | 0.967118 | NA |
| CG7367 | 0 | NA | NA | NA | NA | NA |
| Spn28Dc | 41.61653 | -3.48473 | 3.923108 | -0.88826 | 0.374402 | NA |
| CG7227 | 0 | NA | NA | NA | NA | NA |
| Spn28Db | 0 | NA | NA | NA | NA | NA |
| Spn28Da | 0.132078 | -1.13211 | 4.069918 | -0.27817 | 0.780885 | NA |
| CG7102 | 0 | NA | NA | NA | NA | NA |
| spz3 | 0.209323 | 1.414133 | 4.032409 | 0.350692 | 0.725819 | NA |
| TwdlE | 0.712213 | -1.57119 | 3.998613 | -0.39293 | 0.694369 | NA |
| CG31606 | 0.209323 | 1.414133 | 4.032409 | 0.350692 | 0.725819 | NA |
| CG8419 | 0.671724 | -0.20025 | 3.964006 | -0.05052 | 0.959711 | NA |
| CG8349 | 0 | NA | NA | NA | NA | NA |
| CG14277 | 0 | NA | NA | NA | NA | NA |
| tRNA:Lys-T | 0 | NA | NA | NA | NA | NA |
| CG14275 | 5.914742 | -1.62872 | 3.882448 | -0.41951 | 0.674844 | NA |
| CG14273 | 0 | NA | NA | NA | NA | NA |
| CG7778 | 0 | NA | NA | NA | NA | NA |
| Mur29B | 0 | NA | NA | NA | NA | NA |
| Spn28F | 0 | NA | NA | NA | NA | NA |
| CG8292 | 0 | NA | NA | NA | NA | NA |
| snRNA:U6a | 0.910416 | 0.505751 | 3.949798 | 0.128045 | 0.898114 | NA |
| CG13385 | 0 | NA | NA | NA | NA | NA |
| CG13386 | 0 | NA | NA | NA | NA | NA |
| CG31898 | 6.728173 | 0.687888 | 3.930528 | 0.175012 | 0.86107 | NA |
| CheA29a | 0 | NA | NA | NA | NA | NA |
| tRNA:Asp-( | 0 | NA | NA | NA | NA | NA |
| CG13088 | 13.74307 | 0.700737 | 3.928104 | 0.178391 | 0.858416 | NA |

| | | | | | | |
|---|---|---|---|---|---|---|
| Sgp | 54.25647 | -1.89598 | 3.951613 | -0.4798 | 0.631371 | NA |
| CG17834 | 61.96254 | -1.70637 | 3.951527 | -0.43183 | 0.665867 | NA |
| tRNA:Asp-( | 0 | NA | NA | NA | NA | NA |
| tRNA:Phe- | 0 | NA | NA | NA | NA | NA |
| tRNA:Asp-( | 13.51941 | 1.169913 | 3.954372 | 0.295853 | 0.767342 | NA |
| tRNA:Asp-( | 0 | NA | NA | NA | NA | NA |
| Bace | 3.869664 | -0.57813 | 3.808855 | -0.15178 | 0.879357 | NA |
| tRNA:Asp-( | 0 | NA | NA | NA | NA | NA |
| grk | 1.015653 | 0.790107 | 3.976718 | 0.198683 | 0.842511 | NA |
| fy | 7.625444 | -0.12463 | 3.874471 | -0.03217 | 0.974338 | NA |
| Peritrophi | 0 | NA | NA | NA | NA | NA |
| Peritrophi | 0 | NA | NA | NA | NA | NA |
| lectin-29Ca | 0 | NA | NA | NA | NA | NA |
| Acp29AB | 0 | NA | NA | NA | NA | NA |
| Glt | 0 | NA | NA | NA | NA | NA |
| tRNA:Asp-( | 0 | NA | NA | NA | NA | NA |
| tRNA:Asp-( | 0 | NA | NA | NA | NA | NA |
| CG12438 | 0 | NA | NA | NA | NA | NA |
| CG32986 | 0 | NA | NA | NA | NA | NA |
| CG32987 | 0 | NA | NA | NA | NA | NA |
| CG32988 | 0 | NA | NA | NA | NA | NA |
| CG32983 | 0 | NA | NA | NA | NA | NA |
| CG9483 | 0 | NA | NA | NA | NA | NA |
| LManVI | 0 | NA | NA | NA | NA | NA |
| LManV | 0 | NA | NA | NA | NA | NA |
| LManIV | 0 | NA | NA | NA | NA | NA |
| LManIII | 0.178053 | -1.16467 | 4.075067 | -0.2858 | 0.775028 | NA |
| CG9289 | 0 | NA | NA | NA | NA | NA |
| tRNA:Asp-( | 0.941609 | 1.421141 | 3.984935 | 0.356628 | 0.72137 | NA |
| CG9287 | 0 | NA | NA | NA | NA | NA |
| Tsp29Fa | 0 | NA | NA | NA | NA | NA |
| CG32985 | 0 | NA | NA | NA | NA | NA |
| CG32984 | 0 | NA | NA | NA | NA | NA |
| CG18088 | 0.69928 | 1.019866 | 4.009445 | 0.254366 | 0.799213 | NA |
| CG18662 | 0.3141 | 1.240997 | 4.03841 | 0.307299 | 0.758616 | NA |
| CG17906 | 0 | NA | NA | NA | NA | NA |
| Try29F | 6.634882 | 3.000912 | 3.871897 | 0.77505 | 0.43831 | NA |
| CG9568 | 23.60178 | -4.34564 | 3.880838 | -1.11977 | 0.262813 | NA |
| CG13102 | 0 | NA | NA | NA | NA | NA |
| CG12439 | 0 | NA | NA | NA | NA | NA |
| Toll-4 | 1.895259 | 0.506147 | 3.963229 | 0.127711 | 0.898378 | NA |
| CG31609 | 0 | NA | NA | NA | NA | NA |
| CG9582 | 0 | NA | NA | NA | NA | NA |
| Rcd-1r | 0 | NA | NA | NA | NA | NA |
| rost | 1.309066 | -0.383 | 3.96689 | -0.09655 | 0.923084 | NA |
| CG13101 | 0 | NA | NA | NA | NA | NA |
| CG9541 | 40.06199 | -1.70532 | 3.951865 | -0.43152 | 0.666088 | NA |
| CG9525 | 0 | NA | NA | NA | NA | NA |
| CG17005 | 0 | NA | NA | NA | NA | NA |
| Cks30A | 0.178812 | -1.08808 | 4.061735 | -0.26789 | 0.788788 | NA |

| | | | | | | |
|---|---|---|---|---|---|---|
| CG3748 | 0.1047 | 0.613261 | 4.062799 | 0.150945 | 0.880019 | NA |
| CG13110 | 0 | NA | NA | NA | NA | NA |
| Mco1 | 0.712213 | -1.57119 | 3.998613 | -0.39293 | 0.694369 | NA |
| CG3769 | 0 | NA | NA | NA | NA | NA |
| CG4438 | 0 | NA | NA | NA | NA | NA |
| borr | 0.104623 | 0.639075 | 4.067674 | 0.157111 | 0.875158 | NA |
| aust | 0.368855 | -0.37404 | 4.015709 | -0.09314 | 0.925789 | NA |
| CG31883 | 25.34651 | 0.665054 | 3.952331 | 0.168269 | 0.866372 | NA |
| CG17855 | 0 | NA | NA | NA | NA | NA |
| Cpr30B | 0 | NA | NA | NA | NA | NA |
| CG13113 | 0.104623 | 0.639075 | 4.067674 | 0.157111 | 0.875158 | NA |
| CG13114 | 0.2094 | 1.005034 | 4.045341 | 0.248442 | 0.803792 | NA |
| CG3841 | 0 | NA | NA | NA | NA | NA |
| CG33299 | 0 | NA | NA | NA | NA | NA |
| CG31882 | 0.132078 | -1.13211 | 4.069918 | -0.27817 | 0.780885 | NA |
| CG4382 | 0.3141 | 1.240997 | 4.03841 | 0.307299 | 0.758616 | NA |
| gcm | 0 | NA | NA | NA | NA | NA |
| CG31709 | 0 | NA | NA | NA | NA | NA |
| CG4017 | 0.178053 | -1.16467 | 4.075067 | -0.2858 | 0.775028 | NA |
| CG17633 | 0.178053 | -1.16467 | 4.075067 | -0.2858 | 0.775028 | NA |
| IP3K1 | 35.75338 | -0.1091 | 3.863007 | -0.02824 | 0.97747 | NA |
| CG4709 | 26.83598 | -7.79137 | 3.909939 | -1.99271 | 0.046293 | NA |
| snoRNA:M | 0.178053 | -1.16467 | 4.075067 | -0.2858 | 0.775028 | NA |
| snoRNA:M | 2.377398 | -1.86964 | 3.966533 | -0.47135 | 0.637388 | NA |
| snoRNA:U: | 0.2094 | 1.005034 | 4.045341 | 0.248442 | 0.803792 | NA |
| snoRNA:U: | 4.249151 | 4.596953 | 3.930552 | 1.169544 | 0.242185 | NA |
| Cyp4e3 | 48.85906 | 1.536765 | 3.951645 | 0.388893 | 0.697356 | NA |
| CG13127 | 0.1047 | 0.613261 | 4.062799 | 0.150945 | 0.880019 | NA |
| obst-B | 1.046999 | 1.598136 | 3.993006 | 0.400234 | 0.688984 | NA |
| Cpr31A | 0 | NA | NA | NA | NA | NA |
| CG13137 | 0 | NA | NA | NA | NA | NA |
| CG5731 | 0 | NA | NA | NA | NA | NA |
| Fum4 | 0 | NA | NA | NA | NA | NA |
| CG13135 | 42.50817 | 1.749815 | 3.952129 | 0.442752 | 0.657945 | NA |
| CG13133 | 0 | NA | NA | NA | NA | NA |
| CG13131 | 0 | NA | NA | NA | NA | NA |
| CG4995 | 0 | NA | NA | NA | NA | NA |
| pim | 0 | NA | NA | NA | NA | NA |
| lft | 0 | NA | NA | NA | NA | NA |
| CG5096 | 0.132078 | -1.13211 | 4.069918 | -0.27817 | 0.780885 | NA |
| Ir31a | 0 | NA | NA | NA | NA | NA |
| CG13144 | 0.64424 | -0.14539 | 3.993669 | -0.0364 | 0.97096 | NA |
| Cog4 | 49.14673 | 5.433623 | 3.768638 | 1.4418 | 0.149359 | NA |
| RluA-1 | 0 | NA | NA | NA | NA | NA |
| Cdk1 | 4.470307 | -2.20441 | 3.970887 | -0.55514 | 0.578797 | NA |
| CG5367 | 43.14144 | 0.51971 | 3.951697 | 0.131516 | 0.895367 | NA |
| CG13138 | 0 | NA | NA | NA | NA | NA |
| Hand | 13.7101 | -1.69969 | 3.953695 | -0.4299 | 0.667268 | NA |
| CG18301 | 0.178053 | -1.16467 | 4.075067 | -0.2858 | 0.775028 | NA |
| CG6138 | 37.36155 | 0.475497 | 3.865059 | 0.123024 | 0.902088 | NA |

| | | | | | | |
|---|---|---|---|---|---|---|
| CG31872 | 0 | NA | NA | NA | NA | NA |
| CG18284 | 0.178053 | -1.16467 | 4.075067 | -0.2858 | 0.775028 | NA |
| CG31871 | 0 | NA | NA | NA | NA | NA |
| CG17104 | 0.178053 | -1.16467 | 4.075067 | -0.2858 | 0.775028 | NA |
| CG17105 | 0 | NA | NA | NA | NA | NA |
| CG17107 | 0 | NA | NA | NA | NA | NA |
| CG6415 | 0 | NA | NA | NA | NA | NA |
| CG6431 | 0 | NA | NA | NA | NA | NA |
| CG17118 | 0 | NA | NA | NA | NA | NA |
| Vha16-5 | 0.282677 | -0.09599 | 4.057934 | -0.02366 | 0.981127 | NA |
| Lip2 | 0 | NA | NA | NA | NA | NA |
| Lip1 | 0.53416 | -1.53481 | 4.012395 | -0.38252 | 0.702078 | NA |
| CG7300 | 0 | NA | NA | NA | NA | NA |
| CG17108 | 2.377398 | -1.86964 | 3.966533 | -0.47135 | 0.637388 | NA |
| CG7294 | 0 | NA | NA | NA | NA | NA |
| CG7296 | 0 | NA | NA | NA | NA | NA |
| CG7299 | 0 | NA | NA | NA | NA | NA |
| CG7329 | 0 | NA | NA | NA | NA | NA |
| CG7384 | 0.466186 | 0.959471 | 4.033986 | 0.237847 | 0.812 | NA |
| CG17127 | 0 | NA | NA | NA | NA | NA |
| CG31870 | 0 | NA | NA | NA | NA | NA |
| CG6508 | 0.178053 | -1.16467 | 4.075067 | -0.2858 | 0.775028 | NA |
| CG17134 | 0 | NA | NA | NA | NA | NA |
| CG12517 | 0 | NA | NA | NA | NA | NA |
| CG14072 | 33.11791 | -1.7047 | 3.952066 | -0.43134 | 0.666218 | NA |
| lectin-33A | 0 | NA | NA | NA | NA | NA |
| Acp32CD | 0 | NA | NA | NA | NA | NA |
| CG14913 | 0 | NA | NA | NA | NA | NA |
| CG18666 | 0.356107 | -1.47099 | 4.037764 | -0.36431 | 0.715628 | NA |
| Art8 | 0 | NA | NA | NA | NA | NA |
| aub | 0.1047 | 0.613261 | 4.062799 | 0.150945 | 0.880019 | NA |
| piwi | 6.032582 | -0.2788 | 3.849484 | -0.07242 | 0.942264 | NA |
| CG14069 | 0 | NA | NA | NA | NA | NA |
| CG7309 | 1.046999 | 1.598136 | 3.993006 | 0.400234 | 0.688984 | NA |
| CG14070 | 0 | NA | NA | NA | NA | NA |
| CG14071 | 0 | NA | NA | NA | NA | NA |
| aurB | 0 | NA | NA | NA | NA | NA |
| Porin2 | 0 | NA | NA | NA | NA | NA |
| CG14915 | 0 | NA | NA | NA | NA | NA |
| hgo | 13.58927 | 0.289593 | 3.848482 | 0.075249 | 0.940017 | NA |
| cmet | 0.356107 | -1.47099 | 4.037764 | -0.36431 | 0.715628 | NA |
| Vm32E | 1.158217 | -3.0773 | 3.989947 | -0.77126 | 0.440551 | NA |
| Gyc32E | 0.954171 | -1.65593 | 3.931926 | -0.42115 | 0.673646 | NA |
| CG31867 | 0 | NA | NA | NA | NA | NA |
| RfC38 | 10.69005 | 1.338687 | 3.954459 | 0.338526 | 0.734967 | NA |
| CG14921 | 2.720205 | 1.493763 | 3.963603 | 0.37687 | 0.70627 | NA |
| AstCC | 0.178053 | -1.16467 | 4.075067 | -0.2858 | 0.775028 | NA |
| sala | 0 | NA | NA | NA | NA | NA |
| spz4 | 0.63886 | -0.79045 | 3.986802 | -0.19827 | 0.842837 | NA |
| CG14930 | 0 | NA | NA | NA | NA | NA |

| | | | | | | |
|---|---|---|---|---|---|---|
| CG14926 | 0 | NA | NA | NA | NA | NA |
| CG4983 | 0 | NA | NA | NA | NA | NA |
| CG4988 | 0.396233 | -1.6814 | 4.028351 | -0.41739 | 0.676392 | NA |
| Vha100-5 | 0.233093 | 0.692656 | 4.076246 | 0.169925 | 0.865069 | NA |
| CG14931 | 0 | NA | NA | NA | NA | NA |
| CG14932 | 0.236778 | -0.26946 | 4.071573 | -0.06618 | 0.947234 | NA |
| Tsp33B | 0.178053 | -1.16467 | 4.075067 | -0.2858 | 0.775028 | NA |
| CG12307 | 0 | NA | NA | NA | NA | NA |
| spag4 | 41.97228 | -6.15664 | 3.864604 | -1.59308 | 0.111141 | NA |
| CG31704 | 8.480694 | 1.730894 | 3.956914 | 0.437435 | 0.661796 | NA |
| CG14933 | 0 | NA | NA | NA | NA | NA |
| Or33a | 0.178053 | -1.16467 | 4.075067 | -0.2858 | 0.775028 | NA |
| Or33b | 0.1047 | 0.613261 | 4.062799 | 0.150945 | 0.880019 | NA |
| Or33c | 0 | NA | NA | NA | NA | NA |
| Crys | 0 | NA | NA | NA | NA | NA |
| CG16964 | 0 | NA | NA | NA | NA | NA |
| CG16965 | 0.712213 | -1.57119 | 3.998613 | -0.39293 | 0.694369 | NA |
| Ada1-1 | 42.6194 | 7.055267 | 3.845109 | 1.834868 | 0.066525 | NA |
| Rh5 | 30.29192 | 4.596794 | 3.802091 | 1.209017 | 0.226656 | NA |
| dmGlut | 0.104623 | 0.639075 | 4.067674 | 0.157111 | 0.875158 | NA |
| CG18788 | 0 | NA | NA | NA | NA | NA |
| RpL7-like | 0 | NA | NA | NA | NA | NA |
| CG14947 | 0 | NA | NA | NA | NA | NA |
| Rab3-GAP | 59.26679 | -2.5013 | 3.768729 | -0.6637 | 0.506883 | NA |
| CG18787 | 22.48819 | -3.74386 | 3.807439 | -0.9833 | 0.325459 | NA |
| CG18789 | 5.122163 | 0.359805 | 3.847332 | 0.093521 | 0.92549 | NA |
| Mt2 | 10.36529 | 1.73509 | 3.955844 | 0.438614 | 0.660941 | NA |
| CG17211 | 31.82862 | 3.548314 | 3.905524 | 0.908537 | 0.363594 | NA |
| Gr33a | 0 | NA | NA | NA | NA | NA |
| CG31861 | 12.05115 | 1.374129 | 3.95414 | 0.347517 | 0.728203 | NA |
| CG17217 | 0 | NA | NA | NA | NA | NA |
| CG31862 | 0 | NA | NA | NA | NA | NA |
| CG17010 | 0 | NA | NA | NA | NA | NA |
| rho-6 | 0.4188 | 1.41068 | 4.034613 | 0.349644 | 0.726606 | NA |
| Pkd2 | 0.3873 | 0.267327 | 4.008308 | 0.066693 | 0.946826 | NA |
| Oatp33Eb | 0.669976 | 0.241146 | 3.936344 | 0.061261 | 0.951151 | NA |
| CR17024 | 0 | NA | NA | NA | NA | NA |
| Ref2 | 25.27357 | -3.15152 | 3.918373 | -0.80429 | 0.421228 | NA |
| CG5418 | 0 | NA | NA | NA | NA | NA |
| CG5421 | 16.20522 | -0.29634 | 3.93457 | -0.07532 | 0.939963 | NA |
| CG5435 | 0 | NA | NA | NA | NA | NA |
| CG15483 | 0.356107 | -1.47099 | 4.037764 | -0.36431 | 0.715628 | NA |
| CG15484 | 0 | NA | NA | NA | NA | NA |
| CG5776 | 30.22693 | 1.683864 | 3.952538 | 0.426021 | 0.670093 | NA |
| IFT43 | 0 | NA | NA | NA | NA | NA |
| Nepl7 | 0.77483 | 0.786875 | 3.969371 | 0.198237 | 0.84286 | NA |
| ACXB | 0.233093 | 0.692656 | 4.076246 | 0.169925 | 0.865069 | NA |
| ACXA | 0 | NA | NA | NA | NA | NA |
| ACXE | 0 | NA | NA | NA | NA | NA |
| CG16800 | 0 | NA | NA | NA | NA | NA |

| | | | | | | |
|---|---|---|---|---|---|---|
| Vha68-3 | 0.314023 | 1.768229 | 4.015745 | 0.440324 | 0.659702 | NA |
| CG5122 | 4.402864 | 0.994202 | 3.785282 | 0.262649 | 0.792821 | NA |
| Pk34A | 0 | NA | NA | NA | NA | NA |
| kmg | 0.3873 | 0.267327 | 4.008308 | 0.066693 | 0.946826 | NA |
| Rsph1 | 2.826131 | 4.274146 | 3.944545 | 1.083558 | 0.278561 | NA |
| snRNA:U2: | 2.601183 | 0.742986 | 3.858991 | 0.192534 | 0.847324 | NA |
| CG16813 | 0 | NA | NA | NA | NA | NA |
| CG16815 | 0 | NA | NA | NA | NA | NA |
| Prosalpha6 | 0 | NA | NA | NA | NA | NA |
| snRNA:U5: | 0 | NA | NA | NA | NA | NA |
| CG16820 | 0.63886 | -0.79045 | 3.986802 | -0.19827 | 0.842837 | NA |
| Naa20B | 0 | NA | NA | NA | NA | NA |
| CG16825 | 0.6282 | 1.523021 | 4.015426 | 0.379293 | 0.704471 | NA |
| CG16826 | 0 | NA | NA | NA | NA | NA |
| CG31848 | 0.178053 | -1.16467 | 4.075067 | -0.2858 | 0.775028 | NA |
| CG31730 | 0 | NA | NA | NA | NA | NA |
| Mabi | 0 | NA | NA | NA | NA | NA |
| CG15480 | 0 | NA | NA | NA | NA | NA |
| snRNA:U2: | 0.547193 | 2.009611 | 4.015384 | 0.500478 | 0.616739 | NA |
| CG15482 | 0 | NA | NA | NA | NA | NA |
| CG9928 | 19.78829 | 1.744272 | 3.953518 | 0.441195 | 0.659072 | NA |
| CG16848 | 0.816913 | -0.91625 | 3.980578 | -0.23018 | 0.817951 | NA |
| Tehao | 4.857655 | -5.24295 | 3.926787 | -1.33518 | 0.181819 | NA |
| CG16852 | 0 | NA | NA | NA | NA | NA |
| CG16853 | 10.6832 | -1.6973 | 3.954478 | -0.42921 | 0.66777 | NA |
| CG9254 | 28.53219 | 0.249276 | 3.745282 | 0.066557 | 0.946934 | NA |
| CG9263 | 0 | NA | NA | NA | NA | NA |
| CG15638 | 1.675199 | 1.648777 | 3.978727 | 0.414398 | 0.678583 | NA |
| CG16956 | 0.104623 | 0.639075 | 4.067674 | 0.157111 | 0.875158 | NA |
| Vm34Ca | 0 | NA | NA | NA | NA | NA |
| CG10859 | 0.31387 | 1.261782 | 4.037145 | 0.312543 | 0.754628 | NA |
| CG16957 | 14.0718 | 1.507016 | 3.953962 | 0.381141 | 0.703099 | NA |
| CG18507 | 5.758419 | 2.904806 | 3.935248 | 0.738151 | 0.460423 | NA |
| Gpo3 | 0.1047 | 0.613261 | 4.062799 | 0.150945 | 0.880019 | NA |
| nur | 28.05906 | -1.66372 | 3.952239 | -0.42096 | 0.673788 | NA |
| asRNA:CR3 | 0 | NA | NA | NA | NA | NA |
| CG7916 | 0.357625 | -1.47707 | 4.044794 | -0.36518 | 0.714979 | NA |
| CG7953 | 0 | NA | NA | NA | NA | NA |
| CG7968 | 0 | NA | NA | NA | NA | NA |
| Ance | 56.27327 | -3.8721 | 3.653932 | -1.05971 | 0.289277 | NA |
| Ance-2 | 0 | NA | NA | NA | NA | NA |
| Acyp | 0 | NA | NA | NA | NA | NA |
| Vajk3 | 0 | NA | NA | NA | NA | NA |
| Vajk1 | 0 | NA | NA | NA | NA | NA |
| CG8997 | 0.64106 | -0.7867 | 4.009822 | -0.19619 | 0.844459 | NA |
| CG33306 | 0 | NA | NA | NA | NA | NA |
| CG33307 | 10.81527 | -3.13106 | 3.912235 | -0.80032 | 0.423523 | NA |
| PolG1 | 0 | NA | NA | NA | NA | NA |
| CG16888 | 0 | NA | NA | NA | NA | NA |
| CG9014 | 0 | NA | NA | NA | NA | NA |

| | | | | | | |
|---|---|---|---|---|---|---|
| NimB1 | 0 | NA | NA | NA | NA | NA |
| NimB5 | 0 | NA | NA | NA | NA | NA |
| bgm | 0.178053 | -1.16467 | 4.075067 | -0.2858 | 0.775028 | NA |
| CG31769 | 0 | NA | NA | NA | NA | NA |
| CG15293 | 0 | NA | NA | NA | NA | NA |
| CG18095 | 0 | NA | NA | NA | NA | NA |
| lncRNA:CR | 0 | NA | NA | NA | NA | NA |
| NimC3 | 0 | NA | NA | NA | NA | NA |
| tRNA:Gln-( | 0.4188 | 1.41068 | 4.034613 | 0.349644 | 0.726606 | NA |
| He | 0 | NA | NA | NA | NA | NA |
| NimB4 | 53.11742 | 0.647127 | 3.920236 | 0.165073 | 0.868886 | NA |
| NimB2 | 0 | NA | NA | NA | NA | NA |
| CG32971 | 0 | NA | NA | NA | NA | NA |
| CG15286 | 0.282677 | -0.09599 | 4.057934 | -0.02366 | 0.981127 | NA |
| Pgant35A | 4.072211 | 1.397994 | 3.844884 | 0.363599 | 0.716158 | NA |
| CG15283 | 0 | NA | NA | NA | NA | NA |
| Cpr35B | 0.466186 | 0.959471 | 4.033986 | 0.237847 | 0.812 | NA |
| tRNA:Pro-/ | 0 | NA | NA | NA | NA | NA |
| tRNA:Pro-/ | 0 | NA | NA | NA | NA | NA |
| tRNA:Pro-/ | 0 | NA | NA | NA | NA | NA |
| CG15282 | 0 | NA | NA | NA | NA | NA |
| tRNA:Pro-/ | 0 | NA | NA | NA | NA | NA |
| CG3473 | 5.025597 | 1.715508 | 3.960878 | 0.433113 | 0.664933 | NA |
| tRNA:Gly-( | 0 | NA | NA | NA | NA | NA |
| tRNA:Gly-( | 0 | NA | NA | NA | NA | NA |
| tRNA:Gly-( | 0 | NA | NA | NA | NA | NA |
| tRNA:Gly-( | 0.2094 | 1.005034 | 4.045341 | 0.248442 | 0.803792 | NA |
| CG31835 | 0 | NA | NA | NA | NA | NA |
| CG4691 | 0 | NA | NA | NA | NA | NA |
| CSN1a | 0.233093 | 0.692656 | 4.076246 | 0.169925 | 0.865069 | NA |
| CG4701 | 0 | NA | NA | NA | NA | NA |
| CG3491 | 0.178053 | -1.16467 | 4.075067 | -0.2858 | 0.775028 | NA |
| ProtB | 0 | NA | NA | NA | NA | NA |
| ProtA | 0 | NA | NA | NA | NA | NA |
| CG4480 | 0 | NA | NA | NA | NA | NA |
| CG15279 | 0.178053 | -1.16467 | 4.075067 | -0.2858 | 0.775028 | NA |
| CG18420 | 0 | NA | NA | NA | NA | NA |
| CG4650 | 0.178053 | -1.16467 | 4.075067 | -0.2858 | 0.775028 | NA |
| CG31775 | 0 | NA | NA | NA | NA | NA |
| yellow-c | 5.647804 | -1.09518 | 3.94024 | -0.27795 | 0.781052 | NA |
| CG15269 | 0 | NA | NA | NA | NA | NA |
| ZnT35C | 0 | NA | NA | NA | NA | NA |
| tRNA:Leu-' | 0 | NA | NA | NA | NA | NA |
| CG31832 | 60.99536 | 1.537303 | 3.951498 | 0.389043 | 0.697244 | NA |
| CG15263 | 0 | NA | NA | NA | NA | NA |
| ms(2)35Ci | 0 | NA | NA | NA | NA | NA |
| CG15262 | 0 | NA | NA | NA | NA | NA |
| CG4161 | 0.946065 | -1.55144 | 3.957279 | -0.39205 | 0.695022 | NA |
| sna | 0 | NA | NA | NA | NA | NA |
| wor | 0 | NA | NA | NA | NA | NA |

| | | | | | | |
|---|---|---|---|---|---|---|
| CG15258 | 0 | NA | NA | NA | NA | NA |
| nht | 0 | NA | NA | NA | NA | NA |
| CG15260 | 17.70242 | -2.26093 | 3.956273 | -0.57148 | 0.567674 | NA |
| CG15256 | 0.178812 | -1.08808 | 4.061735 | -0.26789 | 0.788788 | NA |
| CG7631 | 0.356107 | -1.47099 | 4.037764 | -0.36431 | 0.715628 | NA |
| CG18480 | 0 | NA | NA | NA | NA | NA |
| CG18477 | 0 | NA | NA | NA | NA | NA |
| CG18478 | 0 | NA | NA | NA | NA | NA |
| CG31780 | 0 | NA | NA | NA | NA | NA |
| CG31827 | 0 | NA | NA | NA | NA | NA |
| CG31826 | 0 | NA | NA | NA | NA | NA |
| Ku80 | 37.43781 | -7.99376 | 3.910001 | -2.04444 | 0.04091 | NA |
| Gli | 9.180649 | 5.322493 | 3.909097 | 1.361566 | 0.173335 | NA |
| snRNA:U5: | 0 | NA | NA | NA | NA | NA |
| Or35a | 0 | NA | NA | NA | NA | NA |
| CG15253 | 0.178812 | -1.08808 | 4.061735 | -0.26789 | 0.788788 | NA |
| CG15254 | 0 | NA | NA | NA | NA | NA |
| Semp1 | 0.357625 | -1.47707 | 4.044794 | -0.36518 | 0.714979 | NA |
| CG15255 | 0 | NA | NA | NA | NA | NA |
| CG18109 | 0 | NA | NA | NA | NA | NA |
| CG13243 | 0 | NA | NA | NA | NA | NA |
| Tektin-A | 0 | NA | NA | NA | NA | NA |
| CR31824 | 0 | NA | NA | NA | NA | NA |
| CG31735 | 0.1047 | 0.613261 | 4.062799 | 0.150945 | 0.880019 | NA |
| CG13244 | 0 | NA | NA | NA | NA | NA |
| CG10839 | 0.282677 | -0.09599 | 4.057934 | -0.02366 | 0.981127 | NA |
| CG13245 | 0 | NA | NA | NA | NA | NA |
| BicC | 0 | NA | NA | NA | NA | NA |
| CG31820 | 4.094408 | -1.89106 | 3.960175 | -0.47752 | 0.632993 | NA |
| CG31821 | 0 | NA | NA | NA | NA | NA |
| CG31823 | 0 | NA | NA | NA | NA | NA |
| Tep1 | 0.460806 | -0.60052 | 3.997252 | -0.15023 | 0.880581 | NA |
| CG7653 | 0 | NA | NA | NA | NA | NA |
| CR31822 | 0 | NA | NA | NA | NA | NA |
| CG18518 | 0 | NA | NA | NA | NA | NA |
| CG4891 | 0 | NA | NA | NA | NA | NA |
| CG4892 | 0 | NA | NA | NA | NA | NA |
| CG31819 | 4.139081 | -0.81368 | 3.784958 | -0.21498 | 0.829786 | NA |
| CG17329 | 0.104623 | 0.639075 | 4.067674 | 0.157111 | 0.875158 | NA |
| Cyp303a1 | 0.178053 | -1.16467 | 4.075067 | -0.2858 | 0.775028 | NA |
| fzy | 0 | NA | NA | NA | NA | NA |
| CG4440 | 0 | NA | NA | NA | NA | NA |
| CG13258 | 0.1047 | 0.613261 | 4.062799 | 0.150945 | 0.880019 | NA |
| Nepl8 | 1.238749 | -0.26955 | 3.90623 | -0.069 | 0.944986 | NA |
| CG31816 | 0 | NA | NA | NA | NA | NA |
| CG5968 | 27.36445 | -4.7257 | 3.871879 | -1.22052 | 0.222268 | NA |
| ChLD3 | 0.178053 | -1.16467 | 4.075067 | -0.2858 | 0.775028 | NA |
| Ugt37C2 | 0 | NA | NA | NA | NA | NA |
| Ugt37E1 | 0 | NA | NA | NA | NA | NA |
| CG6012 | 0 | NA | NA | NA | NA | NA |

| | | | | | | |
|---|---|---|---|---|---|---|
| CG31809 | 7.765206 | -0.5101 | 3.950431 | -0.12913 | 0.897258 | NA |
| CG31810 | 0.132078 | -1.13211 | 4.069918 | -0.27817 | 0.780885 | NA |
| Nepl9 | 0 | NA | NA | NA | NA | NA |
| CG13282 | 8.794641 | 3.466039 | 3.919779 | 0.884243 | 0.376565 | NA |
| CG13280 | 11.80165 | 0.084389 | 3.952695 | 0.02135 | 0.982967 | NA |
| ppk17 | 1.179159 | -1.16246 | 3.921297 | -0.29645 | 0.766887 | NA |
| glu | 15.50206 | 4.397129 | 3.808727 | 1.154488 | 0.2483 | NA |
| CG17996 | 0 | NA | NA | NA | NA | NA |
| CG31807 | 7.203293 | -3.22037 | 3.816344 | -0.84384 | 0.39876 | NA |
| Trpgamma | 5.005669 | -0.01925 | 3.950245 | -0.00487 | 0.996112 | NA |
| CG31743 | 0 | NA | NA | NA | NA | NA |
| CG31806 | 21.80443 | 3.09609 | 3.881523 | 0.797648 | 0.425075 | NA |
| CG15136 | 0 | NA | NA | NA | NA | NA |
| CG6304 | 0.233093 | 0.692656 | 4.076246 | 0.169925 | 0.865069 | NA |
| CG31805 | 0.233093 | 0.692656 | 4.076246 | 0.169925 | 0.865069 | NA |
| CG32832 | 42.92525 | -1.91992 | 3.951816 | -0.48583 | 0.627085 | NA |
| Gr36a | 0 | NA | NA | NA | NA | NA |
| Gr36b | 0 | NA | NA | NA | NA | NA |
| Gr36c | 0 | NA | NA | NA | NA | NA |
| CG31750 | 0 | NA | NA | NA | NA | NA |
| CG31804 | 0 | NA | NA | NA | NA | NA |
| CG6380 | 0 | NA | NA | NA | NA | NA |
| CG31784 | 0 | NA | NA | NA | NA | NA |
| CG5043 | 0 | NA | NA | NA | NA | NA |
| CG5050 | 0 | NA | NA | NA | NA | NA |
| Dhc36C | 0.565506 | -0.26224 | 3.986711 | -0.06578 | 0.947554 | NA |
| CG15147 | 0 | NA | NA | NA | NA | NA |
| CG6870 | 0 | NA | NA | NA | NA | NA |
| CG7094 | 0 | NA | NA | NA | NA | NA |
| CG15145 | 36.38597 | 0.021996 | 3.730357 | 0.005897 | 0.995295 | NA |
| CG15144 | 0.178053 | -1.16467 | 4.075067 | -0.2858 | 0.775028 | NA |
| CG15143 | 0.668515 | -2.46469 | 4.009802 | -0.61467 | 0.538775 | NA |
| CG15142 | 0 | NA | NA | NA | NA | NA |
| qua | 0 | NA | NA | NA | NA | NA |
| CG18563 | 0 | NA | NA | NA | NA | NA |
| SPH93 | 0.1047 | 0.613261 | 4.062799 | 0.150945 | 0.880019 | NA |
| CG5681 | 0.2094 | 1.005034 | 4.045341 | 0.248442 | 0.803792 | NA |
| Prosbeta5l | 0 | NA | NA | NA | NA | NA |
| CG5693 | 0 | NA | NA | NA | NA | NA |
| CG31740 | 0 | NA | NA | NA | NA | NA |
| CG31741 | 0 | NA | NA | NA | NA | NA |
| Slc25A46b | 0 | NA | NA | NA | NA | NA |
| CG15152 | 35.70268 | 1.748887 | 3.952361 | 0.442492 | 0.658133 | NA |
| CG15153 | 0.209323 | 1.414133 | 4.032409 | 0.350692 | 0.725819 | NA |
| elfless | 0 | NA | NA | NA | NA | NA |
| btv | 2.12201 | -1.90008 | 3.94883 | -0.48118 | 0.630392 | NA |
| CG15155 | 0 | NA | NA | NA | NA | NA |
| CG5790 | 0.62047 | 1.501254 | 3.987081 | 0.376529 | 0.706523 | NA |
| CG31787 | 0 | NA | NA | NA | NA | NA |
| CG31788 | 14.62664 | 1.617221 | 3.954103 | 0.408998 | 0.682541 | NA |

| | | | | | | |
|---|---|---|---|---|---|---|
| CG31802 | 0 | NA | NA | NA | NA | NA |
| Ntf-2r | 0.233093 | 0.692656 | 4.076246 | 0.169925 | 0.865069 | NA |
| grnd | 0 | NA | NA | NA | NA | NA |
| CG10211 | 0 | NA | NA | NA | NA | NA |
| CG31789 | 0 | NA | NA | NA | NA | NA |
| IFT46 | 0 | NA | NA | NA | NA | NA |
| amos | 0 | NA | NA | NA | NA | NA |
| CG10336 | 0 | NA | NA | NA | NA | NA |
| CG15167 | 0 | NA | NA | NA | NA | NA |
| CG17325 | 0 | NA | NA | NA | NA | NA |
| Ugt36F1 | 0.2094 | 1.005034 | 4.045341 | 0.248442 | 0.803792 | NA |
| Ugt36D1 | 47.69933 | -3.09076 | 3.917316 | -0.789 | 0.430112 | NA |
| Ugt36E1 | 0.282753 | -0.25398 | 4.020204 | -0.06318 | 0.949626 | NA |
| CG31752 | 0.53416 | -1.53481 | 4.012395 | -0.38252 | 0.702078 | NA |
| CG10570 | 0 | NA | NA | NA | NA | NA |
| ham | 26.31472 | 2.744876 | 3.870991 | 0.709089 | 0.478269 | NA |
| CG10348 | 0.411147 | -0.29058 | 4.046973 | -0.0718 | 0.94276 | NA |
| CG15169 | 0 | NA | NA | NA | NA | NA |
| CG31792 | 0 | NA | NA | NA | NA | NA |
| robl37BC | 0 | NA | NA | NA | NA | NA |
| CG15172 | 0 | NA | NA | NA | NA | NA |
| Phlpp | 10.05119 | 1.734496 | 3.955995 | 0.438447 | 0.661062 | NA |
| CG17343 | 0 | NA | NA | NA | NA | NA |
| CG17344 | 0 | NA | NA | NA | NA | NA |
| amd | 36.32136 | -1.91935 | 3.951981 | -0.48567 | 0.627203 | NA |
| CG10700 | 0 | NA | NA | NA | NA | NA |
| CR33315 | 0 | NA | NA | NA | NA | NA |
| CG17572 | 0 | NA | NA | NA | NA | NA |
| CG10650 | 23.8341 | 1.243055 | 3.874244 | 0.320851 | 0.748323 | NA |
| CG15170 | 0 | NA | NA | NA | NA | NA |
| CG10750 | 0.178053 | -1.16467 | 4.075067 | -0.2858 | 0.775028 | NA |
| CG13086 | 0 | NA | NA | NA | NA | NA |
| CG17349 | 0.1047 | 0.613261 | 4.062799 | 0.150945 | 0.880019 | NA |
| CG17350 | 0 | NA | NA | NA | NA | NA |
| CG31798 | 0.892543 | -2.83735 | 3.997814 | -0.70972 | 0.477875 | NA |
| CG13085 | 33.28358 | -1.919 | 3.952078 | -0.48557 | 0.627273 | NA |
| CG17549 | 0 | NA | NA | NA | NA | NA |
| fon | 0.178053 | -1.16467 | 4.075067 | -0.2858 | 0.775028 | NA |
| CG17564 | 0.466186 | 0.959471 | 4.033986 | 0.237847 | 0.812 | NA |
| CG31797 | 18.93681 | 1.532509 | 3.952804 | 0.387702 | 0.698237 | NA |
| gammaTul | 0 | NA | NA | NA | NA | NA |
| RanGAP | 56.74728 | 2.975773 | 3.918927 | 0.759334 | 0.447653 | NA |
| CG10026 | 0.356107 | -1.47099 | 4.037764 | -0.36431 | 0.715628 | NA |
| CG13084 | 0 | NA | NA | NA | NA | NA |
| CG31697 | 0.233093 | 0.692656 | 4.076246 | 0.169925 | 0.865069 | NA |
| CG13082 | 0 | NA | NA | NA | NA | NA |
| CG13081 | 0 | NA | NA | NA | NA | NA |
| Tep4 | 0 | NA | NA | NA | NA | NA |
| CG13078 | 0 | NA | NA | NA | NA | NA |
| CG13077 | 0 | NA | NA | NA | NA | NA |

| | | | | | | |
|---|---|---|---|---|---|---|
| msb1l | 0 | NA | NA | NA | NA | NA |
| scw | 0 | NA | NA | NA | NA | NA |
| CG10463 | 27.97419 | 0.612097 | 3.75119 | 0.163174 | 0.870381 | NA |
| snRNA:U4: | 0.418493 | 1.313669 | 4.019283 | 0.326842 | 0.743788 | NA |
| snRNA:U2: | 0.388212 | 0.094116 | 4.038892 | 0.023303 | 0.981409 | NA |
| snRNA:U5: | 0 | NA | NA | NA | NA | NA |
| CG16772 | 61.53668 | 0.359525 | 3.834959 | 0.093749 | 0.925308 | NA |
| CG10680 | 5.15103 | -1.89739 | 3.95832 | -0.47934 | 0.631695 | NA |
| OS9 | 0 | NA | NA | NA | NA | NA |
| CG13965 | 0 | NA | NA | NA | NA | NA |
| Victoria | 0.732363 | 1.393579 | 3.993417 | 0.348969 | 0.727112 | NA |
| CG10481 | 1.172943 | -0.24009 | 3.970289 | -0.06047 | 0.951781 | NA |
| snRNA:U2: | 4.471821 | 1.090965 | 3.845821 | 0.283675 | 0.776659 | NA |
| CG13962 | 0 | NA | NA | NA | NA | NA |
| snRNA:U5: | 0 | NA | NA | NA | NA | NA |
| fbp | 47.31717 | 1.509425 | 3.951654 | 0.381973 | 0.702481 | NA |
| neb | 20.93999 | 1.744836 | 3.953376 | 0.441353 | 0.658957 | NA |
| nesd | 1.366752 | -3.16859 | 3.965138 | -0.79911 | 0.424225 | NA |
| Kua | 0 | NA | NA | NA | NA | NA |
| CG10730 | 0 | NA | NA | NA | NA | NA |
| AANATL3 | 0 | NA | NA | NA | NA | NA |
| CG12617 | 0 | NA | NA | NA | NA | NA |
| CG16798 | 45.34411 | 0.519731 | 3.879997 | 0.133951 | 0.893441 | NA |
| Ugt37A1 | 0.178053 | -1.16467 | 4.075067 | -0.2858 | 0.775028 | NA |
| CG10651 | 0.178053 | -1.16467 | 4.075067 | -0.2858 | 0.775028 | NA |
| CG15475 | 0.3141 | 1.240997 | 4.03841 | 0.307299 | 0.758616 | NA |
| CG17472 | 0 | NA | NA | NA | NA | NA |
| CG31680 | 0 | NA | NA | NA | NA | NA |
| CG18810 | 0 | NA | NA | NA | NA | NA |
| CarT | 3.94801 | 0.30349 | 3.956229 | 0.076712 | 0.938853 | NA |
| CG9316 | 0 | NA | NA | NA | NA | NA |
| phr6-4 | 0 | NA | NA | NA | NA | NA |
| CG17470 | 0 | NA | NA | NA | NA | NA |
| cad | 3.737851 | 1.338807 | 3.960994 | 0.337998 | 0.735365 | NA |
| CG31675 | 0.209247 | 1.042804 | 4.047837 | 0.25762 | 0.7967 | NA |
| CG9270 | 0.1047 | 0.613261 | 4.062799 | 0.150945 | 0.880019 | NA |
| Spn38F | 0 | NA | NA | NA | NA | NA |
| CheB38a | 0 | NA | NA | NA | NA | NA |
| CG33322 | 0.46073 | -0.17853 | 4.00793 | -0.04454 | 0.964471 | NA |
| Itgbn | 22.86169 | -1.43517 | 3.743857 | -0.38334 | 0.701467 | NA |
| CG9259 | 0 | NA | NA | NA | NA | NA |
| CG9257 | 0.209323 | 1.414133 | 4.032409 | 0.350692 | 0.725819 | NA |
| CG31627 | 0.2094 | 1.005034 | 4.045341 | 0.248442 | 0.803792 | NA |
| Mcm10 | 0.209323 | 1.414133 | 4.032409 | 0.350692 | 0.725819 | NA |
| clumsy | 0.2094 | 1.005034 | 4.045341 | 0.248442 | 0.803792 | NA |
| snRNA:U4: | 1.843226 | -0.62009 | 3.962911 | -0.15647 | 0.87566 | NA |
| CG31988 | 0 | NA | NA | NA | NA | NA |
| CG31624 | 0 | NA | NA | NA | NA | NA |
| Ret | 12.30175 | -3.1116 | 3.915083 | -0.79477 | 0.426746 | NA |
| dimm | 0.1047 | 0.613261 | 4.062799 | 0.150945 | 0.880019 | NA |

| | | | | | | |
|---|---|---|---|---|---|---|
| dtr | 43.86831 | -3.38172 | 3.9033 | -0.86637 | 0.386285 | NA |
| CG8665 | 4.491535 | -2.50294 | 3.842178 | -0.65144 | 0.514764 | NA |
| Gr39b | 0 | NA | NA | NA | NA | NA |
| Cyp6t2Psi | 5.594238 | 1.155882 | 3.959178 | 0.29195 | 0.770325 | NA |
| Ir84a | 0 | NA | NA | NA | NA | NA |
| wa-cup | 0.3141 | 1.240997 | 4.03841 | 0.307299 | 0.758616 | NA |
| CG14598 | 0 | NA | NA | NA | NA | NA |
| tRNA:Val-( | 0 | NA | NA | NA | NA | NA |
| alpha-Est2 | 0.53416 | -1.53481 | 4.012395 | -0.38252 | 0.702078 | NA |
| Nlg1 | 0.178812 | -1.08808 | 4.061735 | -0.26789 | 0.788788 | NA |
| alpha-Est3 | 0 | NA | NA | NA | NA | NA |
| Rtnl2 | 0 | NA | NA | NA | NA | NA |
| alpha-Est6 | 0.356107 | -1.47099 | 4.037764 | -0.36431 | 0.715628 | NA |
| alpha-Est7 | 0 | NA | NA | NA | NA | NA |
| Ubc84D | 0.178053 | -1.16467 | 4.075067 | -0.2858 | 0.775028 | NA |
| alpha-Est1 | 35.72998 | 1.790233 | 3.899643 | 0.459076 | 0.646179 | NA |
| tRNA:Val-( | 0 | NA | NA | NA | NA | NA |
| tRNA:Val-( | 0 | NA | NA | NA | NA | NA |
| tRNA:Gly-1 | 57.31113 | 6.032411 | 3.83801 | 1.571755 | 0.116007 | NA |
| CG31493 | 8.804864 | -5.20569 | 3.890449 | -1.33807 | 0.180874 | NA |
| Sas-4 | 0 | NA | NA | NA | NA | NA |
| sas | 25.00866 | -3.50752 | 3.901674 | -0.89898 | 0.368664 | NA |
| CG14609 | 21.51816 | 1.373109 | 3.872346 | 0.354593 | 0.722894 | NA |
| thw | 0 | NA | NA | NA | NA | NA |
| CG14607 | 0 | NA | NA | NA | NA | NA |
| CG10050 | 6.084524 | -0.3706 | 3.954377 | -0.09372 | 0.925333 | NA |
| CR10032 | 0.178053 | -1.16467 | 4.075067 | -0.2858 | 0.775028 | NA |
| RacGAP84 | 4.124062 | -0.31346 | 3.915857 | -0.08005 | 0.936199 | NA |
| CG31496 | 0.178053 | -1.16467 | 4.075067 | -0.2858 | 0.775028 | NA |
| nxf4 | 0 | NA | NA | NA | NA | NA |
| Mst84Da | 0 | NA | NA | NA | NA | NA |
| Mst84Db | 0 | NA | NA | NA | NA | NA |
| Mst84Dc | 0 | NA | NA | NA | NA | NA |
| Mst84Dd | 0 | NA | NA | NA | NA | NA |
| CG17944 | 0 | NA | NA | NA | NA | NA |
| CG10029 | 0 | NA | NA | NA | NA | NA |
| Gld | 0.46073 | -0.17853 | 4.00793 | -0.04454 | 0.964471 | NA |
| Obp84a | 0 | NA | NA | NA | NA | NA |
| CG14605 | 0 | NA | NA | NA | NA | NA |
| CG14606 | 0 | NA | NA | NA | NA | NA |
| CG14610 | 0 | NA | NA | NA | NA | NA |
| tRNA:Gly-1 | 0.313946 | 1.788224 | 4.01364 | 0.445537 | 0.655932 | NA |
| Ama | 0 | NA | NA | NA | NA | NA |
| Dfd | 39.33256 | -2.64277 | 3.85029 | -0.68638 | 0.492473 | NA |
| tRNA:Lys-1 | 0 | NA | NA | NA | NA | NA |
| tRNA:Lys-1 | 0 | NA | NA | NA | NA | NA |
| 7SLRNA:CF | 0.132078 | -1.13211 | 4.069918 | -0.27817 | 0.780885 | NA |
| ftz | 0.132078 | -1.13211 | 4.069918 | -0.27817 | 0.780885 | NA |
| Sodh-1 | 3.026906 | -1.67104 | 3.963213 | -0.42164 | 0.67329 | NA |
| CG1979 | 0 | NA | NA | NA | NA | NA |

| | | | | | | |
|---|---|---|---|---|---|---|
| djl | 0 | NA | NA | NA | NA | NA |
| Mst84B | 0 | NA | NA | NA | NA | NA |
| tRNA:Lys-T | 0 | NA | NA | NA | NA | NA |
| tRNA:Lys-T | 0 | NA | NA | NA | NA | NA |
| tRNA:Lys-C | 4.502097 | 1.711245 | 3.961986 | 0.431916 | 0.665803 | NA |
| bcd | 0.104623 | 0.639075 | 4.067674 | 0.157111 | 0.875158 | NA |
| zen | 0 | NA | NA | NA | NA | NA |
| zen2 | 0 | NA | NA | NA | NA | NA |
| CG15186 | 40.8306 | -1.89128 | 3.897618 | -0.48524 | 0.627505 | NA |
| CG15185 | 0 | NA | NA | NA | NA | NA |
| Pif2 | 0 | NA | NA | NA | NA | NA |
| CG31482 | 0 | NA | NA | NA | NA | NA |
| sunz | 0.178053 | -1.16467 | 4.075067 | -0.2858 | 0.775028 | NA |
| CG31286 | 0 | NA | NA | NA | NA | NA |
| Ccp84Ag | 0.178053 | -1.16467 | 4.075067 | -0.2858 | 0.775028 | NA |
| Ccp84Ad | 0.178812 | -1.08808 | 4.061735 | -0.26789 | 0.788788 | NA |
| Ccp84Aa | 0.178053 | -1.16467 | 4.075067 | -0.2858 | 0.775028 | NA |
| Ccp84Ac | 0 | NA | NA | NA | NA | NA |
| Ccp84Ae | 0 | NA | NA | NA | NA | NA |
| Ccp84Af | 0 | NA | NA | NA | NA | NA |
| Ass | 0 | NA | NA | NA | NA | NA |
| CG15177 | 0 | NA | NA | NA | NA | NA |
| sowi | 0 | NA | NA | NA | NA | NA |
| CG1138 | 0.178053 | -1.16467 | 4.075067 | -0.2858 | 0.775028 | NA |
| CG1137 | 19.85974 | -0.33589 | 3.76454 | -0.08923 | 0.928902 | NA |
| tRNA:Met- | 3.664497 | 1.702077 | 3.964384 | 0.429342 | 0.667674 | NA |
| Osi19 | 0 | NA | NA | NA | NA | NA |
| Osi20 | 0 | NA | NA | NA | NA | NA |
| CG17917 | 11.2266 | -1.91107 | 3.954354 | -0.48328 | 0.628896 | NA |
| CG10298 | 6.207651 | -1.90163 | 3.957084 | -0.48056 | 0.630826 | NA |
| CG17919 | 9.350061 | -1.7818 | 3.954677 | -0.45055 | 0.652311 | NA |
| Hr83 | 0 | NA | NA | NA | NA | NA |
| Neurochor | 0.356107 | -1.47099 | 4.037764 | -0.36431 | 0.715628 | NA |
| CG2616 | 0 | NA | NA | NA | NA | NA |
| CG10919 | 0.1047 | 0.613261 | 4.062799 | 0.150945 | 0.880019 | NA |
| CG10086 | 0.178053 | -1.16467 | 4.075067 | -0.2858 | 0.775028 | NA |
| CG2678 | 0.104623 | 0.639075 | 4.067674 | 0.157111 | 0.875158 | NA |
| lds | 1.539911 | 0.274724 | 3.966541 | 0.06926 | 0.944782 | NA |
| CG10445 | 0.282753 | -0.25398 | 4.020204 | -0.06318 | 0.949626 | NA |
| CG18268 | 0 | NA | NA | NA | NA | NA |
| CG3014 | 0.209323 | 1.414133 | 4.032409 | 0.350692 | 0.725819 | NA |
| CG18744 | 9.081476 | -2.90706 | 3.924892 | -0.74067 | 0.458892 | NA |
| CG18746 | 0 | NA | NA | NA | NA | NA |
| CG18748 | 0 | NA | NA | NA | NA | NA |
| CG18745 | 0 | NA | NA | NA | NA | NA |
| CG18747 | 0 | NA | NA | NA | NA | NA |
| CG11286 | 0 | NA | NA | NA | NA | NA |
| Os-C | 0 | NA | NA | NA | NA | NA |
| CG7900 | 45.7174 | -3.38716 | 3.903076 | -0.86782 | 0.385494 | NA |
| CG18249 | 10.96245 | -1.91078 | 3.954436 | -0.4832 | 0.628954 | NA |

| | | | | | | |
|---|---|---|---|---|---|---|
| CG15864 | 27.2242 | -4.7603 | 3.866477 | -1.23117 | 0.218258 | NA |
| tRNA:Arg-/ | 0 | NA | NA | NA | NA | NA |
| tRNA:Arg-/ | 0 | NA | NA | NA | NA | NA |
| tRNA:Arg-/ | 0 | NA | NA | NA | NA | NA |
| tRNA:Arg-/ | 0 | NA | NA | NA | NA | NA |
| tRNA:Arg-/ | 0 | NA | NA | NA | NA | NA |
| tRNA:Asn-( | 1.072874 | -2.0339 | 4.019685 | -0.50598 | 0.612867 | NA |
| ImpE3 | 0.660388 | -1.75977 | 4.001266 | -0.4398 | 0.660079 | NA |
| CG11052 | 0.132078 | -1.13211 | 4.069918 | -0.27817 | 0.780885 | NA |
| ELOVL | 3.169864 | -1.88225 | 3.962774 | -0.47498 | 0.634799 | NA |
| Mcm2 | 0 | NA | NA | NA | NA | NA |
| ato | 0.178053 | -1.16467 | 4.075067 | -0.2858 | 0.775028 | NA |
| Nazo | 0 | NA | NA | NA | NA | NA |
| CG31464 | 0 | NA | NA | NA | NA | NA |
| CG31463 | 0 | NA | NA | NA | NA | NA |
| CG11698 | 0 | NA | NA | NA | NA | NA |
| Or85a | 0 | NA | NA | NA | NA | NA |
| CG7443 | 0 | NA | NA | NA | NA | NA |
| COX7AL | 0.357625 | -1.47707 | 4.044794 | -0.36518 | 0.714979 | NA |
| CG7352 | 14.3714 | 3.626327 | 3.843399 | 0.943521 | 0.345415 | NA |
| CG7963 | 0 | NA | NA | NA | NA | NA |
| Obp85a | 0.4188 | 1.41068 | 4.034613 | 0.349644 | 0.726606 | NA |
| Poxm | 0 | NA | NA | NA | NA | NA |
| CG11694 | 0 | NA | NA | NA | NA | NA |
| CG11693 | 0 | NA | NA | NA | NA | NA |
| CG11672 | 0.233093 | 0.692656 | 4.076246 | 0.169925 | 0.865069 | NA |
| CG31454 | 0 | NA | NA | NA | NA | NA |
| CG31259 | 0.178053 | -1.16467 | 4.075067 | -0.2858 | 0.775028 | NA |
| Or85d | 0 | NA | NA | NA | NA | NA |
| CG31460 | 3.623677 | 4.272127 | 3.908648 | 1.092994 | 0.274397 | NA |
| tRNA:Tyr-( | 0 | NA | NA | NA | NA | NA |
| tRNA:Tyr-( | 0 | NA | NA | NA | NA | NA |
| tRNA:Tyr-( | 0 | NA | NA | NA | NA | NA |
| Cyp313b1 | 1.024049 | 0.824862 | 3.952456 | 0.208696 | 0.834686 | NA |
| tRNA:Tyr-( | 0 | NA | NA | NA | NA | NA |
| tRNA:Tyr-( | 0 | NA | NA | NA | NA | NA |
| Or85e | 0 | NA | NA | NA | NA | NA |
| Or85c | 0.356866 | -1.92326 | 4.037889 | -0.4763 | 0.633858 | NA |
| Or85b | 0.178053 | -1.16467 | 4.075067 | -0.2858 | 0.775028 | NA |
| CG8112 | 0 | NA | NA | NA | NA | NA |
| CG11760 | 0.104623 | 0.639075 | 4.067674 | 0.157111 | 0.875158 | NA |
| CG8136 | 0 | NA | NA | NA | NA | NA |
| CG8223 | 33.69399 | 1.570012 | 3.740187 | 0.419768 | 0.674655 | NA |
| CG8236 | 0 | NA | NA | NA | NA | NA |
| Ir85a | 3.347944 | 1.501875 | 3.961309 | 0.379136 | 0.704587 | NA |
| tRNA:Arg-[illegible] | 0 | NA | NA | NA | NA | NA |
| CG33189 | 2.097839 | 1.11873 | 3.972242 | 0.281637 | 0.778222 | NA |
| CG33191 | 0.488184 | -2.32943 | 4.022674 | -0.57908 | 0.562538 | NA |
| CG13318 | 0 | NA | NA | NA | NA | NA |
| ranshi | 20.1734 | -0.0354 | 3.879381 | -0.00912 | 0.99272 | NA |

| | | | | | | |
|---|---|---|---|---|---|---|
| CG33325 | 0 | NA | NA | NA | NA | NA |
| CG11977 | 0 | NA | NA | NA | NA | NA |
| Ada | 0 | NA | NA | NA | NA | NA |
| CG8420 | 11.065 | -2.90387 | 3.761094 | -0.77208 | 0.440066 | NA |
| St2 | 0 | NA | NA | NA | NA | NA |
| CR16735 | 0 | NA | NA | NA | NA | NA |
| CG16736 | 0.6282 | 1.523021 | 4.015426 | 0.379293 | 0.704471 | NA |
| HP1e | 48.96414 | 4.500924 | 3.877655 | 1.160734 | 0.24575 | NA |
| Srr | 0 | NA | NA | NA | NA | NA |
| CG8135 | 0 | NA | NA | NA | NA | NA |
| CG16749 | 23.90606 | -1.9174 | 3.952536 | -0.48511 | 0.6276 | NA |
| CG12951 | 0 | NA | NA | NA | NA | NA |
| SpdS | 39.69083 | -3.50788 | 3.743041 | -0.93717 | 0.34867 | NA |
| Nepl12 | 0.713731 | -2.6372 | 4.003426 | -0.65874 | 0.510065 | NA |
| nmdyn-D7 | 0 | NA | NA | NA | NA | NA |
| CG18542 | 0 | NA | NA | NA | NA | NA |
| CG32939 | 0 | NA | NA | NA | NA | NA |
| alphaTub8 | 0.178053 | -1.16467 | 4.075067 | -0.2858 | 0.775028 | NA |
| p24-2 | 0 | NA | NA | NA | NA | NA |
| Unc-115b | 0 | NA | NA | NA | NA | NA |
| CG9444 | 0.356107 | -1.47099 | 4.037764 | -0.36431 | 0.715628 | NA |
| CG9492 | 0.770937 | -1.65034 | 3.976655 | -0.41501 | 0.678137 | NA |
| Fst | 0.178812 | -1.08808 | 4.061735 | -0.26789 | 0.788788 | NA |
| CG9427 | 10.99349 | 1.736178 | 3.955567 | 0.43892 | 0.660719 | NA |
| CG31415 | 4.502097 | 1.711245 | 3.961986 | 0.431916 | 0.665803 | NA |
| topi | 0 | NA | NA | NA | NA | NA |
| CG8534 | 0 | NA | NA | NA | NA | NA |
| eloF | 0 | NA | NA | NA | NA | NA |
| CG9458 | 0 | NA | NA | NA | NA | NA |
| CG16904 | 0 | NA | NA | NA | NA | NA |
| Whamy | 44.15721 | -1.7056 | 3.951777 | -0.4316 | 0.66603 | NA |
| Rpt3R | 0 | NA | NA | NA | NA | NA |
| Spn85F | 0 | NA | NA | NA | NA | NA |
| Npc2e | 0 | NA | NA | NA | NA | NA |
| Npc2c | 0 | NA | NA | NA | NA | NA |
| ATPsyneps | 0 | NA | NA | NA | NA | NA |
| knk | 0 | NA | NA | NA | NA | NA |
| Alp13 | 2.617498 | 1.683176 | 3.969392 | 0.424039 | 0.671538 | NA |
| Npc2d | 0 | NA | NA | NA | NA | NA |
| Fancl | 0.356107 | -1.47099 | 4.037764 | -0.36431 | 0.715628 | NA |
| CG31407 | 0 | NA | NA | NA | NA | NA |
| CG5359 | 12.49498 | 4.852605 | 3.872106 | 1.253221 | 0.210125 | NA |
| Gr85a | 0 | NA | NA | NA | NA | NA |
| CG12420 | 0 | NA | NA | NA | NA | NA |
| CG3999 | 0.660388 | -1.75977 | 4.001266 | -0.4398 | 0.660079 | NA |
| CG12592 | 13.45584 | -5.37662 | 3.851703 | -1.39591 | 0.162742 | NA |
| CG31406 | 0 | NA | NA | NA | NA | NA |
| tRNA:Ser-( | 0 | NA | NA | NA | NA | NA |
| mthl11 | 12.33805 | -2.25201 | 3.958532 | -0.5689 | 0.569424 | NA |
| CG14684 | 0.1047 | 0.613261 | 4.062799 | 0.150945 | 0.880019 | NA |

| | | | | | | |
|---|---|---|---|---|---|---|
| CG4073 | 0 | NA | NA | NA | NA | NA |
| Desi | 0.2094 | 1.005034 | 4.045341 | 0.248442 | 0.803792 | NA |
| Mcm5 | 25.65141 | 2.965217 | 3.921285 | 0.756185 | 0.449538 | NA |
| CG4570 | 0 | NA | NA | NA | NA | NA |
| CG14693 | 0.283512 | -0.2331 | 4.065094 | -0.05734 | 0.954273 | NA |
| CG31272 | 0 | NA | NA | NA | NA | NA |
| CG14691 | 0 | NA | NA | NA | NA | NA |
| tomboy20 | 0 | NA | NA | NA | NA | NA |
| CG14689 | 6.587971 | -1.69066 | 3.956663 | -0.42729 | 0.669165 | NA |
| CG31467 | 0 | NA | NA | NA | NA | NA |
| CG31391 | 0 | NA | NA | NA | NA | NA |
| Tengl4 | 52.66406 | 1.750763 | 3.951894 | 0.443019 | 0.657752 | NA |
| CG14696 | 22.71952 | 3.432757 | 3.897369 | 0.880788 | 0.378432 | NA |
| mAcon2 | 0.890266 | -1.59482 | 3.989917 | -0.39971 | 0.689368 | NA |
| Ugt302E1 | 0 | NA | NA | NA | NA | NA |
| CG4757 | 0.356107 | -1.47099 | 4.037764 | -0.36431 | 0.715628 | NA |
| Ugt303A1 | 0.411147 | -0.29058 | 4.046973 | -0.0718 | 0.94276 | NA |
| CG18577 | 0 | NA | NA | NA | NA | NA |
| CG4820 | 19.71815 | 4.689282 | 3.878268 | 1.209117 | 0.226618 | NA |
| scpr-C | 18.63659 | 1.743638 | 3.953678 | 0.441017 | 0.659201 | NA |
| scpr-A | 2.931598 | 1.690143 | 3.967537 | 0.425993 | 0.670113 | NA |
| scpr-B | 4.253697 | 2.01957 | 3.943775 | 0.512091 | 0.608588 | NA |
| Ugt304A1 | 0 | NA | NA | NA | NA | NA |
| Ugt35E1 | 0.178053 | -1.16467 | 4.075067 | -0.2858 | 0.775028 | NA |
| PGRP-LB | 1.885233 | -1.25249 | 3.966236 | -0.31579 | 0.752163 | NA |
| sals | 4.012275 | 0.045399 | 3.894374 | 0.011658 | 0.990699 | NA |
| KP78a | 0 | NA | NA | NA | NA | NA |
| KP78b | 0 | NA | NA | NA | NA | NA |
| CG14708 | 0 | NA | NA | NA | NA | NA |
| CG5342 | 0 | NA | NA | NA | NA | NA |
| CG14710 | 37.35103 | -2.6988 | 3.839693 | -0.70287 | 0.482138 | NA |
| CG14711 | 9.437583 | -2.90821 | 3.924596 | -0.74102 | 0.458681 | NA |
| CG14718 | 0 | NA | NA | NA | NA | NA |
| l-t | 0 | NA | NA | NA | NA | NA |
| CG14720 | 0 | NA | NA | NA | NA | NA |
| CG18765 | 0.178053 | -1.16467 | 4.075067 | -0.2858 | 0.775028 | NA |
| CG6813 | 53.92813 | 8.506755 | 3.908876 | 2.176266 | 0.029535 | NA |
| Zaf1 | 23.63529 | -2.82283 | 3.831658 | -0.73671 | 0.461297 | NA |
| CG6790 | 0 | NA | NA | NA | NA | NA |
| sad | 54.27657 | 1.679595 | 3.951813 | 0.425019 | 0.670823 | NA |
| CG14731 | 0 | NA | NA | NA | NA | NA |
| Hsp70Ab | 22.35324 | 0.652205 | 3.746708 | 0.174074 | 0.861807 | NA |
| aurA | 0 | NA | NA | NA | NA | NA |
| GC2 | 0 | NA | NA | NA | NA | NA |
| CG10005 | 10.04383 | 1.526465 | 3.954462 | 0.386011 | 0.699489 | NA |
| Hsp70Aa | 0.670053 | 0.799805 | 3.963306 | 0.201803 | 0.840071 | NA |
| Dnali1 | 0 | NA | NA | NA | NA | NA |
| CG6962 | 38.17913 | -0.50667 | 3.929428 | -0.12894 | 0.897404 | NA |
| ssp5 | 0 | NA | NA | NA | NA | NA |
| tRNA:Lys-T | 0 | NA | NA | NA | NA | NA |

| | | | | | | |
|---|---|---|---|---|---|---|
| Adk1 | 0.178053 | -1.16467 | 4.075067 | -0.2858 | 0.775028 | NA |
| eIF3d2 | 0.2094 | 1.005034 | 4.045341 | 0.248442 | 0.803792 | NA |
| CG4702 | 5.679341 | -1.8997 | 3.957646 | -0.48001 | 0.631222 | NA |
| CG14740 | 0 | NA | NA | NA | NA | NA |
| CG14739 | 0 | NA | NA | NA | NA | NA |
| CG31358 | 0 | NA | NA | NA | NA | NA |
| tRNA:Thr-T | 0 | NA | NA | NA | NA | NA |
| Rsbp15 | 1.883219 | 1.475262 | 3.968901 | 0.371705 | 0.710112 | NA |
| CG3916 | 0 | NA | NA | NA | NA | NA |
| CG3942 | 0 | NA | NA | NA | NA | NA |
| svp | 22.23288 | -1.93366 | 3.736996 | -0.51744 | 0.604851 | NA |
| CG17738 | 0.178053 | -1.16467 | 4.075067 | -0.2858 | 0.775028 | NA |
| CG4066 | 0 | NA | NA | NA | NA | NA |
| CG10013 | 0.178812 | -1.08808 | 4.061735 | -0.26789 | 0.788788 | NA |
| CG4115 | 0 | NA | NA | NA | NA | NA |
| GstD4 | 0 | NA | NA | NA | NA | NA |
| GstD5 | 6.095603 | 1.339992 | 3.956027 | 0.338722 | 0.734819 | NA |
| GstD6 | 0 | NA | NA | NA | NA | NA |
| GstD7 | 0 | NA | NA | NA | NA | NA |
| GstD8 | 0 | NA | NA | NA | NA | NA |
| GstD11 | 0 | NA | NA | NA | NA | NA |
| CG33098 | 0 | NA | NA | NA | NA | NA |
| Desat2 | 0.178053 | -1.16467 | 4.075067 | -0.2858 | 0.775028 | NA |
| CG10035 | 31.82878 | 1.748183 | 3.952537 | 0.442294 | 0.658277 | NA |
| GstD10 | 54.03022 | -0.32344 | 3.836432 | -0.08431 | 0.932813 | NA |
| Tim17a1 | 0.356107 | -1.47099 | 4.037764 | -0.36431 | 0.715628 | NA |
| CG10041 | 0.712213 | -1.57119 | 3.998613 | -0.39293 | 0.694369 | NA |
| tRNA:Thr-T | 0 | NA | NA | NA | NA | NA |
| Cyp313a3 | 0 | NA | NA | NA | NA | NA |
| Cyp313a2 | 0 | NA | NA | NA | NA | NA |
| Cyp313a5 | 0 | NA | NA | NA | NA | NA |
| Pglym87 | 0 | NA | NA | NA | NA | NA |
| CG17404 | 0.178812 | -1.08808 | 4.061735 | -0.26789 | 0.788788 | NA |
| CG4860 | 0 | NA | NA | NA | NA | NA |
| CG4830 | 0.178812 | -1.08808 | 4.061735 | -0.26789 | 0.788788 | NA |
| Hsp70Bbb | 2.615581 | 1.492057 | 3.964088 | 0.376393 | 0.706624 | NA |
| Hsp70Bc | 36.78058 | 0.766546 | 3.835252 | 0.199868 | 0.841583 | NA |
| Hug | 16.19714 | 1.629148 | 3.953821 | 0.412044 | 0.680307 | NA |
| CG18530 | 0 | NA | NA | NA | NA | NA |
| CG11598 | 0.233093 | 0.692656 | 4.076246 | 0.169925 | 0.865069 | NA |
| CG11600 | 0 | NA | NA | NA | NA | NA |
| CG11608 | 0 | NA | NA | NA | NA | NA |
| CG6225 | 0 | NA | NA | NA | NA | NA |
| CG14395 | 9.548171 | 1.40437 | 3.954332 | 0.355147 | 0.722479 | NA |
| Gnmt | 0 | NA | NA | NA | NA | NA |
| NijC | 0.356107 | -1.47099 | 4.037764 | -0.36431 | 0.715628 | NA |
| Ugt37A2 | 0 | NA | NA | NA | NA | NA |
| Ugt37A3 | 0.233093 | 0.692656 | 4.076246 | 0.169925 | 0.865069 | NA |
| CG14391 | 0 | NA | NA | NA | NA | NA |
| CG31347 | 20.06734 | 0.235201 | 3.843993 | 0.061187 | 0.951211 | NA |

| | | | | | | |
|---|---|---|---|---|---|---|
| Cyp313a4 | 0 | NA | NA | NA | NA | NA |
| CG6753 | 0 | NA | NA | NA | NA | NA |
| lncRNA:alp | 14.38293 | 1.72461 | 3.764217 | 0.458159 | 0.646838 | NA |
| Hsp70Ba | 0.774906 | 0.100364 | 3.977538 | 0.025233 | 0.979869 | NA |
| CG17207 | 0 | NA | NA | NA | NA | NA |
| d-cup | 0 | NA | NA | NA | NA | NA |
| CG10909 | 0 | NA | NA | NA | NA | NA |
| Cyp304a1 | 0 | NA | NA | NA | NA | NA |
| CG7091 | 12.45929 | 1.738298 | 3.955028 | 0.439516 | 0.660288 | NA |
| ry | 1.700385 | -0.69042 | 3.913484 | -0.17642 | 0.859963 | NA |
| CG11668 | 1.602479 | -1.64094 | 3.97352 | -0.41297 | 0.67963 | NA |
| CG11670 | 0 | NA | NA | NA | NA | NA |
| CG31157 | 0 | NA | NA | NA | NA | NA |
| sim | 56.00094 | -1.92067 | 3.951604 | -0.48605 | 0.626933 | NA |
| timeout | 35.4428 | -3.51547 | 3.90037 | -0.90132 | 0.367419 | NA |
| snk | 0 | NA | NA | NA | NA | NA |
| CG17327 | 0 | NA | NA | NA | NA | NA |
| yellow-f2 | 0.53416 | -1.53481 | 4.012395 | -0.38252 | 0.702078 | NA |
| yellow-f | 0.178812 | -1.08808 | 4.061735 | -0.26789 | 0.788788 | NA |
| CG14383 | 0.64106 | -0.7867 | 4.009822 | -0.19619 | 0.844459 | NA |
| CG14384 | 0 | NA | NA | NA | NA | NA |
| CG8138 | 0 | NA | NA | NA | NA | NA |
| CG14380 | 0 | NA | NA | NA | NA | NA |
| CG8141 | 0 | NA | NA | NA | NA | NA |
| CG8630 | 0 | NA | NA | NA | NA | NA |
| Osi22 | 0.536437 | -1.71175 | 4.038042 | -0.42391 | 0.671635 | NA |
| CG8773 | 0.356107 | -1.47099 | 4.037764 | -0.36431 | 0.715628 | NA |
| CG8774 | 0.806253 | 0.654822 | 3.981166 | 0.16448 | 0.869353 | NA |
| mthl12 | 0.178053 | -1.16467 | 4.075067 | -0.2858 | 0.775028 | NA |
| CheA87a | 0 | NA | NA | NA | NA | NA |
| CG8870 | 0 | NA | NA | NA | NA | NA |
| CG14377 | 0 | NA | NA | NA | NA | NA |
| yellow-e | 0 | NA | NA | NA | NA | NA |
| yellow-e2 | 0 | NA | NA | NA | NA | NA |
| yellow-e3 | 0 | NA | NA | NA | NA | NA |
| CG14374 | 0 | NA | NA | NA | NA | NA |
| CG9799 | 58.9733 | -0.20209 | 3.836286 | -0.05268 | 0.957989 | NA |
| Lip3 | 0 | NA | NA | NA | NA | NA |
| wntD | 0 | NA | NA | NA | NA | NA |
| apn | 0 | NA | NA | NA | NA | NA |
| TBC1D5 | 0 | NA | NA | NA | NA | NA |
| CG8476 | 0 | NA | NA | NA | NA | NA |
| CG8508 | 0 | NA | NA | NA | NA | NA |
| Ir87a | 0.178812 | -1.08808 | 4.061735 | -0.26789 | 0.788788 | NA |
| Act87E | 0.282753 | -0.25398 | 4.020204 | -0.06318 | 0.949626 | NA |
| CG14370 | 0 | NA | NA | NA | NA | NA |
| CG14369 | 0 | NA | NA | NA | NA | NA |
| CG31337 | 0 | NA | NA | NA | NA | NA |
| CG12538 | 0 | NA | NA | NA | NA | NA |
| asRNA:CR1 | 7.098664 | -3.99432 | 3.837261 | -1.04093 | 0.297908 | NA |

| | | | | | | |
|---|---|---|---|---|---|---|
| CG9269 | 0 | NA | NA | NA | NA | NA |
| CG10841 | 0 | NA | NA | NA | NA | NA |
| CG14367 | 50.74518 | -1.70595 | 3.951664 | -0.4317 | 0.665957 | NA |
| CG9312 | 43.03167 | 1.749869 | 3.952114 | 0.442768 | 0.657934 | NA |
| Cht5 | 42.92689 | 2.972935 | 3.919558 | 0.758488 | 0.448159 | NA |
| CG9297 | 0.461489 | -1.26887 | 4.028229 | -0.31499 | 0.752766 | NA |
| CG9288 | 0 | NA | NA | NA | NA | NA |
| tRNA:Phe- | 0 | NA | NA | NA | NA | NA |
| tRNA:Phe- | 0.1047 | 0.613261 | 4.062799 | 0.150945 | 0.880019 | NA |
| Rbp4 | 0.178053 | -1.16467 | 4.075067 | -0.2858 | 0.775028 | NA |
| CG9757 | 0 | NA | NA | NA | NA | NA |
| CG9759 | 0.178053 | -1.16467 | 4.075067 | -0.2858 | 0.775028 | NA |
| ems | 5.905203 | 1.274247 | 3.957034 | 0.322021 | 0.747437 | NA |
| CG31517 | 0 | NA | NA | NA | NA | NA |
| Orc2 | 0.282753 | -0.25398 | 4.020204 | -0.06318 | 0.949626 | NA |
| CG9925 | 0.74894 | 0.958264 | 3.97722 | 0.240938 | 0.809603 | NA |
| CG9926 | 0.178053 | -1.16467 | 4.075067 | -0.2858 | 0.775028 | NA |
| Art6 | 0.356866 | -1.92326 | 4.037889 | -0.4763 | 0.633858 | NA |
| Art9 | 0.590718 | -0.57492 | 4.003363 | -0.14361 | 0.885809 | NA |
| E5 | 0.2094 | 1.005034 | 4.045341 | 0.248442 | 0.803792 | NA |
| CG14362 | 0 | NA | NA | NA | NA | NA |
| CG31495 | 0.314023 | 1.768229 | 4.015745 | 0.440324 | 0.659702 | NA |
| Or88a | 0 | NA | NA | NA | NA | NA |
| CG3199 | 0 | NA | NA | NA | NA | NA |
| CG9616 | 0 | NA | NA | NA | NA | NA |
| CG9624 | 0 | NA | NA | NA | NA | NA |
| CG9631 | 4.095225 | -1.68037 | 3.96008 | -0.42433 | 0.671327 | NA |
| Sp212 | 0 | NA | NA | NA | NA | NA |
| CG33109 | 0 | NA | NA | NA | NA | NA |
| CG31326 | 0.264155 | -1.54099 | 4.048976 | -0.38059 | 0.703509 | NA |
| CG31327 | 0.411147 | -0.29058 | 4.046973 | -0.0718 | 0.94276 | NA |
| CG31533 | 0.282753 | -0.25398 | 4.020204 | -0.06318 | 0.949626 | NA |
| CG9722 | 0.178053 | -1.16467 | 4.075067 | -0.2858 | 0.775028 | NA |
| CG14355 | 0.132078 | -1.13211 | 4.069918 | -0.27817 | 0.780885 | NA |
| CG14356 | 0 | NA | NA | NA | NA | NA |
| Kif19A | 0.442209 | -2.31142 | 4.024999 | -0.57427 | 0.565788 | NA |
| CG14357 | 0.104623 | 0.639075 | 4.067674 | 0.157111 | 0.875158 | NA |
| CG9920 | 0.178053 | -1.16467 | 4.075067 | -0.2858 | 0.775028 | NA |
| tRNA:Ser-( | 0 | NA | NA | NA | NA | NA |
| CG14839 | 0.178812 | -1.08808 | 4.061735 | -0.26789 | 0.788788 | NA |
| IFT54 | 0 | NA | NA | NA | NA | NA |
| CG31321 | 0 | NA | NA | NA | NA | NA |
| soti | 0 | NA | NA | NA | NA | NA |
| CG14841 | 0 | NA | NA | NA | NA | NA |
| CG14840 | 0 | NA | NA | NA | NA | NA |
| CG14850 | 0 | NA | NA | NA | NA | NA |
| CG14851 | 0.1047 | 0.613261 | 4.062799 | 0.150945 | 0.880019 | NA |
| CG14852 | 0 | NA | NA | NA | NA | NA |
| Cad88C | 0.233093 | 0.692656 | 4.076246 | 0.169925 | 0.865069 | NA |
| BigH1 | 0 | NA | NA | NA | NA | NA |

| | | | | | | |
|---|---|---|---|---|---|---|
| Rad17 | 0 | NA | NA | NA | NA | NA |
| CG7886 | 0 | NA | NA | NA | NA | NA |
| Cys | 0.536437 | -1.71175 | 4.038042 | -0.42391 | 0.671635 | NA |
| CG8087 | 0 | NA | NA | NA | NA | NA |
| CG14854 | 0 | NA | NA | NA | NA | NA |
| CG14855 | 14.55965 | -5.38613 | 3.885812 | -1.3861 | 0.165716 | NA |
| CG14856 | 0 | NA | NA | NA | NA | NA |
| CG14857 | 13.74906 | 6.274049 | 3.852454 | 1.628585 | 0.103401 | NA |
| CG3610 | 2.202539 | 2.272562 | 3.954673 | 0.574652 | 0.565526 | NA |
| CG33332 | 0.357625 | -1.47707 | 4.044794 | -0.36518 | 0.714979 | NA |
| CG14860 | 0.2094 | 1.005034 | 4.045341 | 0.248442 | 0.803792 | NA |
| RpS5b | 0.1047 | 0.613261 | 4.062799 | 0.150945 | 0.880019 | NA |
| CG7362 | 0.460806 | -0.60052 | 3.997252 | -0.15023 | 0.880581 | NA |
| CG14861 | 0 | NA | NA | NA | NA | NA |
| RpL10Aa | 0 | NA | NA | NA | NA | NA |
| CG3984 | 0 | NA | NA | NA | NA | NA |
| Mf | 42.27433 | 5.351597 | 3.865168 | 1.38457 | 0.166184 | NA |
| CG6974 | 0 | NA | NA | NA | NA | NA |
| CG4210 | 57.16616 | 1.751077 | 3.951817 | 0.443107 | 0.657689 | NA |
| CG4338 | 22.83796 | 8.258528 | 3.846797 | 2.146858 | 0.031805 | NA |
| CG14866 | 0 | NA | NA | NA | NA | NA |
| CG6118 | 0.356107 | -1.47099 | 4.037764 | -0.36431 | 0.715628 | NA |
| CG6125 | 24.76767 | -0.60444 | 3.837471 | -0.15751 | 0.874843 | NA |
| h-cup | 11.21736 | -1.69782 | 3.954309 | -0.42936 | 0.667662 | NA |
| l(3)neo43 | 45.17854 | 0.122833 | 3.408704 | 0.036035 | 0.971254 | NA |
| Amt | 0 | NA | NA | NA | NA | NA |
| Spn88Eb | 0.712213 | -1.57119 | 3.998613 | -0.39293 | 0.694369 | NA |
| Act88F | 0.1047 | 0.613261 | 4.062799 | 0.150945 | 0.880019 | NA |
| B9d1 | 0 | NA | NA | NA | NA | NA |
| CG5213 | 0 | NA | NA | NA | NA | NA |
| AOX4 | 0.460806 | -0.60052 | 3.997252 | -0.15023 | 0.880581 | NA |
| AOX2 | 0 | NA | NA | NA | NA | NA |
| CG31296 | 0 | NA | NA | NA | NA | NA |
| Trissin | 6.041173 | 2.443859 | 3.932681 | 0.621423 | 0.534321 | NA |
| CG31446 | 0 | NA | NA | NA | NA | NA |
| CG5399 | 0.178053 | -1.16467 | 4.075067 | -0.2858 | 0.775028 | NA |
| CG4520 | 0 | NA | NA | NA | NA | NA |
| CG12784 | 0 | NA | NA | NA | NA | NA |
| Ttc26 | 0 | NA | NA | NA | NA | NA |
| CG5614 | 0 | NA | NA | NA | NA | NA |
| Uros2 | 0 | NA | NA | NA | NA | NA |
| CG9593 | 1.150856 | 1.439869 | 3.979295 | 0.36184 | 0.717471 | NA |
| CG4576 | 0 | NA | NA | NA | NA | NA |
| spn-E | 0 | NA | NA | NA | NA | NA |
| CG9632 | 0.178812 | -1.08808 | 4.061735 | -0.26789 | 0.788788 | NA |
| Pbp45 | 1.465799 | 1.63603 | 3.98226 | 0.410829 | 0.681198 | NA |
| rec | 0 | NA | NA | NA | NA | NA |
| c(3)G | 16.59954 | 6.419444 | 3.899614 | 1.646174 | 0.099728 | NA |
| CG5623 | 0 | NA | NA | NA | NA | NA |
| CG31294 | 0.1047 | 0.613261 | 4.062799 | 0.150945 | 0.880019 | NA |

| | | | | | | |
|---|---|---|---|---|---|---|
| CR31292 | 0 | NA | NA | NA | NA | NA |
| Fbxl7 | 22.37695 | 1.173937 | 3.953007 | 0.296973 | 0.766487 | NA |
| GATAe | 0 | NA | NA | NA | NA | NA |
| pnr | 0.233093 | 0.692656 | 4.076246 | 0.169925 | 0.865069 | NA |
| CG10264 | 0 | NA | NA | NA | NA | NA |
| CG32855 | 0 | NA | NA | NA | NA | NA |
| CG10185 | 0 | NA | NA | NA | NA | NA |
| IKKbeta | 4.638391 | -1.96453 | 3.907524 | -0.50276 | 0.615136 | NA |
| CG10407 | 24.04478 | -5.41448 | 3.873955 | -1.39766 | 0.162215 | NA |
| CG31088 | 0 | NA | NA | NA | NA | NA |
| CG5478 | 0 | NA | NA | NA | NA | NA |
| CG14877 | 26.00759 | 1.607857 | 3.952694 | 0.406775 | 0.684173 | NA |
| Sb | 0 | NA | NA | NA | NA | NA |
| CG8925 | 0 | NA | NA | NA | NA | NA |
| Zip89B | 0 | NA | NA | NA | NA | NA |
| tRNA:Phe- | 8.391356 | 1.163791 | 3.956459 | 0.29415 | 0.768644 | NA |
| tRNA:Val-/ | 18.64746 | 1.172702 | 3.953425 | 0.296629 | 0.76675 | NA |
| CG6901 | 0 | NA | NA | NA | NA | NA |
| CG17929 | 0 | NA | NA | NA | NA | NA |
| CG17930 | 0.178053 | -1.16467 | 4.075067 | -0.2858 | 0.775028 | NA |
| CG31279 | 1.465799 | 1.63603 | 3.98226 | 0.410829 | 0.681198 | NA |
| tRNA:Thr-/ | 0 | NA | NA | NA | NA | NA |
| CG10405 | 19.58586 | -1.70224 | 3.952863 | -0.43064 | 0.666733 | NA |
| Gyc89Da | 0 | NA | NA | NA | NA | NA |
| Gyc89Db | 13.0246 | 0.398469 | 3.843263 | 0.10368 | 0.917423 | NA |
| CG14891 | 0.356866 | -1.92326 | 4.037889 | -0.4763 | 0.633858 | NA |
| Decay | 4.112682 | -2.19841 | 3.972482 | -0.55341 | 0.579983 | NA |
| Cad89D | 0.132078 | -1.13211 | 4.069918 | -0.27817 | 0.780885 | NA |
| tRNA:Thr-/ | 0 | NA | NA | NA | NA | NA |
| ss | 19.13148 | 0.080845 | 3.833637 | 0.021088 | 0.983175 | NA |
| CG10317 | 0.178812 | -1.08808 | 4.061735 | -0.26789 | 0.788788 | NA |
| CG14880 | 0 | NA | NA | NA | NA | NA |
| CG14892 | 2.745815 | 3.964215 | 3.942889 | 1.005409 | 0.3147 | NA |
| NPF | 29.71301 | 1.535031 | 3.952119 | 0.388407 | 0.697715 | NA |
| CG17562 | 0 | NA | NA | NA | NA | NA |
| CG17560 | 0 | NA | NA | NA | NA | NA |
| CG14893 | 0 | NA | NA | NA | NA | NA |
| Glut3 | 0 | NA | NA | NA | NA | NA |
| lncRNA:bx | 0.3141 | 1.240997 | 4.03841 | 0.307299 | 0.758616 | NA |
| CG31275 | 0 | NA | NA | NA | NA | NA |
| Ubx | 0.178053 | -1.16467 | 4.075067 | -0.2858 | 0.775028 | NA |
| modSP | 49.35567 | 1.672616 | 3.951898 | 0.423244 | 0.672117 | NA |
| CG14905 | 0.178053 | -1.16467 | 4.075067 | -0.2858 | 0.775028 | NA |
| CG10345 | 0 | NA | NA | NA | NA | NA |
| VhaM9.7-c | 0 | NA | NA | NA | NA | NA |
| MESK4 | 11.12828 | -2.88421 | 3.758206 | -0.76744 | 0.442818 | NA |
| CG8907 | 15.71724 | -1.91446 | 3.953377 | -0.48426 | 0.628201 | NA |
| Actn3 | 0 | NA | NA | NA | NA | NA |
| Abd-B | 0.236778 | -0.26946 | 4.071573 | -0.06618 | 0.947234 | NA |
| abd-A | 0.1047 | 0.613261 | 4.062799 | 0.150945 | 0.880019 | NA |

| | | | | | | |
|---|---|---|---|---|---|---|
| CG5225 | 0 | NA | NA | NA | NA | NA |
| CG31268 | 0 | NA | NA | NA | NA | NA |
| CG31269 | 0 | NA | NA | NA | NA | NA |
| CG31265 | 0 | NA | NA | NA | NA | NA |
| CG31267 | 0.6282 | 1.523021 | 4.015426 | 0.379293 | 0.704471 | NA |
| CG5246 | 0 | NA | NA | NA | NA | NA |
| CG5255 | 0 | NA | NA | NA | NA | NA |
| CG5265 | 0 | NA | NA | NA | NA | NA |
| CG31262 | 0 | NA | NA | NA | NA | NA |
| TwdlW | 0.178053 | -1.16467 | 4.075067 | -0.2858 | 0.775028 | NA |
| CG4053 | 0 | NA | NA | NA | NA | NA |
| CG17475 | 0.178053 | -1.16467 | 4.075067 | -0.2858 | 0.775028 | NA |
| CG17477 | 0 | NA | NA | NA | NA | NA |
| CG18213 | 7.741737 | 1.18143 | 3.903954 | 0.302624 | 0.762177 | NA |
| CG4009 | 0 | NA | NA | NA | NA | NA |
| CG31418 | 0 | NA | NA | NA | NA | NA |
| CG31419 | 0 | NA | NA | NA | NA | NA |
| CG17283 | 0 | NA | NA | NA | NA | NA |
| CG5860 | 0 | NA | NA | NA | NA | NA |
| CG5863 | 0 | NA | NA | NA | NA | NA |
| CG5866 | 0 | NA | NA | NA | NA | NA |
| CG33333 | 0 | NA | NA | NA | NA | NA |
| cysu | 1.06832 | -1.61145 | 3.983918 | -0.40449 | 0.685853 | NA |
| CG14330 | 1.398559 | 1.091484 | 3.982137 | 0.274095 | 0.784011 | NA |
| Eh | 0 | NA | NA | NA | NA | NA |
| CG14331 | 0 | NA | NA | NA | NA | NA |
| CG14332 | 0 | NA | NA | NA | NA | NA |
| P5cr-2 | 0 | NA | NA | NA | NA | NA |
| Sgs5bis | 0 | NA | NA | NA | NA | NA |
| Sgs5 | 0 | NA | NA | NA | NA | NA |
| CG14327 | 0 | NA | NA | NA | NA | NA |
| tRNA:Val-( | 0 | NA | NA | NA | NA | NA |
| CG14326 | 0 | NA | NA | NA | NA | NA |
| tRNA:Ala-/ | 0.233093 | 0.692656 | 4.076246 | 0.169925 | 0.865069 | NA |
| tRNA:Pro-⁻ | 0 | NA | NA | NA | NA | NA |
| AttD | 0 | NA | NA | NA | NA | NA |
| CG14325 | 0 | NA | NA | NA | NA | NA |
| tRNA:Pro-⁻ | 0 | NA | NA | NA | NA | NA |
| tRNA:Pro-⁻ | 0 | NA | NA | NA | NA | NA |
| tRNA:Ala-/ | 0 | NA | NA | NA | NA | NA |
| tRNA:Ala-/ | 0 | NA | NA | NA | NA | NA |
| tRNA:Ala-/ | 0 | NA | NA | NA | NA | NA |
| tRNA:Ala-/ | 0 | NA | NA | NA | NA | NA |
| tRNA:Ala-/ | 0 | NA | NA | NA | NA | NA |
| tRNA:Ala-/ | 0 | NA | NA | NA | NA | NA |
| tRNA:Ala-/ | 0 | NA | NA | NA | NA | NA |
| tRNA:Val-/ | 0 | NA | NA | NA | NA | NA |
| tRNA:Thr-/ | 0 | NA | NA | NA | NA | NA |
| tRNA:Thr-/ | 0 | NA | NA | NA | NA | NA |
| CG14323 | 0 | NA | NA | NA | NA | NA |

| | | | | | | |
|---|---|---|---|---|---|---|
| tRNA:Ala-/ | 0 | NA | NA | NA | NA | NA |
| CG14324 | 0 | NA | NA | NA | NA | NA |
| tRNA:Pro-` | 0.178812 | -1.08808 | 4.061735 | -0.26789 | 0.788788 | NA |
| Edg91 | 0 | NA | NA | NA | NA | NA |
| CG12347 | 0.104623 | 0.639075 | 4.067674 | 0.157111 | 0.875158 | NA |
| CG17801 | 0.71449 | -2.47883 | 4.014121 | -0.61753 | 0.536887 | NA |
| Atg8b | 0 | NA | NA | NA | NA | NA |
| CG7794 | 0 | NA | NA | NA | NA | NA |
| CG14317 | 0 | NA | NA | NA | NA | NA |
| tRNA:Gly-1 | 0.702005 | 1.501538 | 3.971695 | 0.37806 | 0.705386 | NA |
| tRNA:Gly-1 | 0 | NA | NA | NA | NA | NA |
| CG14316 | 0 | NA | NA | NA | NA | NA |
| CG14315 | 0 | NA | NA | NA | NA | NA |
| CG18598 | 0 | NA | NA | NA | NA | NA |
| CG18599 | 0.233093 | 0.692656 | 4.076246 | 0.169925 | 0.865069 | NA |
| hmw | 0 | NA | NA | NA | NA | NA |
| lncRNA:CR | 0 | NA | NA | NA | NA | NA |
| CG7675 | 0 | NA | NA | NA | NA | NA |
| WRNexo | 0 | NA | NA | NA | NA | NA |
| CG7126 | 0.178053 | -1.16467 | 4.075067 | -0.2858 | 0.775028 | NA |
| CG7142 | 0.310131 | -1.94616 | 4.045058 | -0.48112 | 0.630431 | NA |
| CG14313 | 0 | NA | NA | NA | NA | NA |
| cona | 0 | NA | NA | NA | NA | NA |
| CG7691 | 0 | NA | NA | NA | NA | NA |
| CG14305 | 0.178053 | -1.16467 | 4.075067 | -0.2858 | 0.775028 | NA |
| CG14304 | 0.732669 | 2.681564 | 3.986192 | 0.672713 | 0.50113 | NA |
| CG10864 | 0.894061 | -2.00006 | 4.030294 | -0.49626 | 0.619714 | NA |
| Vha100-4 | 0.282753 | -0.25398 | 4.020204 | -0.06318 | 0.949626 | NA |
| CG31231 | 0.178053 | -1.16467 | 4.075067 | -0.2858 | 0.775028 | NA |
| CG31230 | 0 | NA | NA | NA | NA | NA |
| CG31226 | 0 | NA | NA | NA | NA | NA |
| CG7720 | 4.416522 | -2.34843 | 3.828177 | -0.61346 | 0.539573 | NA |
| CG7715 | 47.41589 | -1.92023 | 3.95173 | -0.48592 | 0.627024 | NA |
| CG7714 | 0.236701 | -0.24276 | 4.020028 | -0.06039 | 0.951846 | NA |
| Muc91C | 0.638783 | -0.20822 | 3.989517 | -0.05219 | 0.958375 | NA |
| CG14301 | 30.07346 | -6.25491 | 3.869434 | -1.61649 | 0.105988 | NA |
| tRNA:Pro-` | 0.236778 | -0.26946 | 4.071573 | -0.06618 | 0.947234 | NA |
| CG7695 | 0 | NA | NA | NA | NA | NA |
| CG14298 | 0.132078 | -1.13211 | 4.069918 | -0.27817 | 0.780885 | NA |
| CG14297 | 0 | NA | NA | NA | NA | NA |
| Rh2 | 0 | NA | NA | NA | NA | NA |
| CG14286 | 0 | NA | NA | NA | NA | NA |
| CG6026 | 0.488184 | -2.32943 | 4.022674 | -0.57908 | 0.562538 | NA |
| CG14292 | 0.836986 | 1.408818 | 3.9887 | 0.353202 | 0.723937 | NA |
| CG14294 | 0.31387 | 1.261782 | 4.037145 | 0.312543 | 0.754628 | NA |
| Cyp12a5 | 37.02758 | -2.99414 | 3.905974 | -0.76655 | 0.443347 | NA |
| nos | 0.233093 | 0.692656 | 4.076246 | 0.169925 | 0.865069 | NA |
| CG14282 | 0 | NA | NA | NA | NA | NA |
| Gdn1 | 0.86242 | -0.6641 | 3.93138 | -0.16892 | 0.865858 | NA |
| CG14280 | 1.450416 | 2.221915 | 3.921545 | 0.566592 | 0.570992 | NA |

| | | | | | | |
|---|---|---|---|---|---|---|
| CG5217 | 0.178053 | -1.16467 | 4.075067 | -0.2858 | 0.775028 | NA |
| CG5250 | 0.178053 | -1.16467 | 4.075067 | -0.2858 | 0.775028 | NA |
| CG11703 | 11.93579 | 1.737599 | 3.955206 | 0.439319 | 0.66043 | NA |
| CG3734 | 0 | NA | NA | NA | NA | NA |
| CG18493 | 32.61202 | 0.131047 | 3.948353 | 0.03319 | 0.973523 | NA |
| CG3739 | 0 | NA | NA | NA | NA | NA |
| CG15025 | 51.15212 | 2.397978 | 3.919834 | 0.611755 | 0.5407 | NA |
| Scsalpha2 | 0 | NA | NA | NA | NA | NA |
| CG3517 | 49.27478 | -3.16815 | 3.905502 | -0.8112 | 0.417249 | NA |
| CG31244 | 0 | NA | NA | NA | NA | NA |
| CG31245 | 0.178053 | -1.16467 | 4.075067 | -0.2858 | 0.775028 | NA |
| CG31404 | 0 | NA | NA | NA | NA | NA |
| CG3581 | 0.53416 | -1.53481 | 4.012395 | -0.38252 | 0.702078 | NA |
| CG7333 | 0 | NA | NA | NA | NA | NA |
| CG7342 | 0.46073 | -0.17853 | 4.00793 | -0.04454 | 0.964471 | NA |
| CG17752 | 0.356107 | -1.47099 | 4.037764 | -0.36431 | 0.715628 | NA |
| CG16727 | 0.356107 | -1.47099 | 4.037764 | -0.36431 | 0.715628 | NA |
| CG7432 | 3.218621 | -2.17835 | 3.977872 | -0.54762 | 0.583954 | NA |
| tRNA:Val-( | 0 | NA | NA | NA | NA | NA |
| tRNA:Ala-/ | 0 | NA | NA | NA | NA | NA |
| tRNA:Thr-/ | 0 | NA | NA | NA | NA | NA |
| CG11659 | 0.715249 | -1.88064 | 4.034336 | -0.46616 | 0.641102 | NA |
| CG6300 | 0 | NA | NA | NA | NA | NA |
| tRNA:Val-( | 0 | NA | NA | NA | NA | NA |
| CG31459 | 0 | NA | NA | NA | NA | NA |
| CG4770 | 0 | NA | NA | NA | NA | NA |
| CG4854 | 2.81157 | -0.71279 | 3.905405 | -0.18251 | 0.855179 | NA |
| CG17190 | 0 | NA | NA | NA | NA | NA |
| CG4465 | 0.53416 | -1.53481 | 4.012395 | -0.38252 | 0.702078 | NA |
| Xport-A | 0 | NA | NA | NA | NA | NA |
| CG17186 | 1.891372 | -1.25233 | 3.893828 | -0.32162 | 0.747742 | NA |
| CG10887 | 43.71221 | 0.015847 | 3.93167 | 0.004031 | 0.996784 | NA |
| Gr92a | 0 | NA | NA | NA | NA | NA |
| CG17199 | 3.962585 | 1.146346 | 3.962484 | 0.2893 | 0.772352 | NA |
| CG31206 | 0 | NA | NA | NA | NA | NA |
| Ir92a | 0 | NA | NA | NA | NA | NA |
| MtnC | 0 | NA | NA | NA | NA | NA |
| eIF3g2 | 0 | NA | NA | NA | NA | NA |
| Or92a | 0 | NA | NA | NA | NA | NA |
| MtnB | 0 | NA | NA | NA | NA | NA |
| Dic2 | 0 | NA | NA | NA | NA | NA |
| CG4335 | 0 | NA | NA | NA | NA | NA |
| CG4362 | 0 | NA | NA | NA | NA | NA |
| CG4367 | 0 | NA | NA | NA | NA | NA |
| MtnD | 5.594238 | 1.155882 | 3.959178 | 0.29195 | 0.770325 | NA |
| Cpr92F | 0.589959 | -1.15593 | 4.01641 | -0.2878 | 0.773499 | NA |
| CG31205 | 0 | NA | NA | NA | NA | NA |
| CG31200 | 59.5122 | 1.435774 | 3.951593 | 0.36334 | 0.716351 | NA |
| CG17267 | 0 | NA | NA | NA | NA | NA |
| CG15923 | 0.4188 | 1.41068 | 4.034613 | 0.349644 | 0.726606 | NA |

| | | | | | | |
|---|---|---|---|---|---|---|
| tRNA:Leu-[illegible] | 0 | NA | NA | NA | NA | NA |
| CG15695 | 0 | NA | NA | NA | NA | NA |
| CG15696 | 0 | NA | NA | NA | NA | NA |
| TotA | 4.761425 | 1.868764 | 3.77732 | 0.494733 | 0.620789 | NA |
| TotC | 0 | NA | NA | NA | NA | NA |
| TotB | 0 | NA | NA | NA | NA | NA |
| TotZ | 0 | NA | NA | NA | NA | NA |
| tRNA:Thr-[illegible] | 0 | NA | NA | NA | NA | NA |
| tRNA:Thr-[illegible] | 0 | NA | NA | NA | NA | NA |
| tRNA:Thr-[illegible] | 0.2094 | 1.005034 | 4.045341 | 0.248442 | 0.803792 | NA |
| Grik | 3.978597 | 1.705937 | 3.963372 | 0.430426 | 0.666886 | NA |
| TotX | 0 | NA | NA | NA | NA | NA |
| Takl1 | 0 | NA | NA | NA | NA | NA |
| CG5697 | 0 | NA | NA | NA | NA | NA |
| CG12278 | 4.622719 | -1.89457 | 3.959145 | -0.47853 | 0.632273 | NA |
| dmrt93B | 0.178812 | -1.08808 | 4.061735 | -0.26789 | 0.788788 | NA |
| rtet | 7.478238 | -1.69271 | 3.955987 | -0.42789 | 0.668735 | NA |
| Obp93a | 0 | NA | NA | NA | NA | NA |
| CG7044 | 58.97275 | 0.361757 | 3.834322 | 0.094347 | 0.924833 | NA |
| HHEX | 23.3503 | -0.87126 | 3.949683 | -0.22059 | 0.825412 | NA |
| r-l | 50.514 | 0.139459 | 3.306955 | 0.042171 | 0.966362 | NA |
| CG7079 | 13.48854 | 0.426445 | 3.850007 | 0.110765 | 0.911803 | NA |
| CG31207 | 38.3485 | -3.51693 | 3.900132 | -0.90175 | 0.367191 | NA |
| CG31189 | 14.52855 | -1.91377 | 3.953578 | -0.48406 | 0.628344 | NA |
| Dhc93AB | 2.799008 | -3.41842 | 3.863738 | -0.88474 | 0.376295 | NA |
| Alp5 | 11.88699 | -1.91173 | 3.954165 | -0.48347 | 0.628761 | NA |
| CG5892 | 0 | NA | NA | NA | NA | NA |
| CG17298 | 0 | NA | NA | NA | NA | NA |
| tin | 0.209247 | 1.042804 | 4.047837 | 0.25762 | 0.7967 | NA |
| bap | 0.104623 | 0.639075 | 4.067674 | 0.157111 | 0.875158 | NA |
| CG7907 | 0 | NA | NA | NA | NA | NA |
| lbe | 0 | NA | NA | NA | NA | NA |
| lbl | 1.602479 | -1.64094 | 3.97352 | -0.41297 | 0.67963 | NA |
| CG3301 | 0 | NA | NA | NA | NA | NA |
| CG15497 | 0.178053 | -1.16467 | 4.075067 | -0.2858 | 0.775028 | NA |
| CG31176 | 3.739119 | -1.67783 | 3.960931 | -0.4236 | 0.671861 | NA |
| CG15498 | 0 | NA | NA | NA | NA | NA |
| slou | 0.784445 | -1.74882 | 3.934036 | -0.44454 | 0.656656 | NA |
| CG5849 | 0.357625 | -1.47707 | 4.044794 | -0.36518 | 0.714979 | NA |
| CG31343 | 30.49306 | 3.974879 | 3.878962 | 1.024728 | 0.305492 | NA |
| CG31233 | 0.264155 | -1.54099 | 4.048976 | -0.38059 | 0.703509 | NA |
| CG31198 | 8.581471 | -4.22675 | 3.922584 | -1.07754 | 0.281239 | NA |
| Burs | 0 | NA | NA | NA | NA | NA |
| CG6800 | 58.06591 | 1.537193 | 3.951528 | 0.389012 | 0.697267 | NA |
| CG31431 | 17.77187 | -0.90243 | 3.952083 | -0.22834 | 0.81938 | NA |
| CG31178 | 0.528311 | -1.72839 | 4.011871 | -0.43082 | 0.6666 | NA |
| P5CDh2 | 4.711497 | 1.713057 | 3.961514 | 0.432425 | 0.665433 | NA |
| CG31465 | 0 | NA | NA | NA | NA | NA |
| CG6678 | 23.78052 | -4.37543 | 3.898063 | -1.12246 | 0.261665 | NA |
| Qsox3 | 0 | NA | NA | NA | NA | NA |

| | | | | | | |
|---|---|---|---|---|---|---|
| Qsox2 | 0.1047 | 0.613261 | 4.062799 | 0.150945 | 0.880019 | NA |
| CG17819 | 0 | NA | NA | NA | NA | NA |
| fit | 0 | NA | NA | NA | NA | NA |
| CG31174 | 4.843574 | 0.678216 | 3.932375 | 0.17247 | 0.863068 | NA |
| CG6569 | 0 | NA | NA | NA | NA | NA |
| Gr93d | 0 | NA | NA | NA | NA | NA |
| Gr93c | 0 | NA | NA | NA | NA | NA |
| Gr93b | 0 | NA | NA | NA | NA | NA |
| Theg | 0.53416 | -1.53481 | 4.012395 | -0.38252 | 0.702078 | NA |
| BomBc3 | 0.132078 | -1.13211 | 4.069918 | -0.27817 | 0.780885 | NA |
| CG13407 | 0 | NA | NA | NA | NA | NA |
| BomT3 | 14.78675 | 0.385959 | 3.840239 | 0.100504 | 0.919944 | NA |
| CG6015 | 0 | NA | NA | NA | NA | NA |
| Ir94a | 0 | NA | NA | NA | NA | NA |
| CG7080 | 0 | NA | NA | NA | NA | NA |
| CG13862 | 0 | NA | NA | NA | NA | NA |
| rdhB | 40.48828 | 0.319379 | 3.950713 | 0.080841 | 0.935569 | NA |
| CG7071 | 0 | NA | NA | NA | NA | NA |
| CG7069 | 13.49694 | 1.705698 | 3.833984 | 0.444889 | 0.6564 | NA |
| tRNA:Ser-( | 0 | NA | NA | NA | NA | NA |
| tRNA:Ser-( | 0 | NA | NA | NA | NA | NA |
| CG13856 | 1.989299 | 1.663439 | 3.974713 | 0.418505 | 0.675578 | NA |
| CG13855 | 0.1047 | 0.613261 | 4.062799 | 0.150945 | 0.880019 | NA |
| tRNA:Ser-( | 0.95296 | -0.11447 | 3.972323 | -0.02882 | 0.977011 | NA |
| tRNA:Ser-( | 0 | NA | NA | NA | NA | NA |
| CG13857 | 0.357625 | -1.47707 | 4.044794 | -0.36518 | 0.714979 | NA |
| CG5386 | 0 | NA | NA | NA | NA | NA |
| CG5388 | 0 | NA | NA | NA | NA | NA |
| CG5391 | 0.178053 | -1.16467 | 4.075067 | -0.2858 | 0.775028 | NA |
| Ir94c | 0.712213 | -1.57119 | 3.998613 | -0.39293 | 0.694369 | NA |
| Ir94b | 0 | NA | NA | NA | NA | NA |
| Pebp1 | 14.34389 | 1.740403 | 3.954495 | 0.440108 | 0.659859 | NA |
| tHMG1 | 0 | NA | NA | NA | NA | NA |
| tHMG2 | 0 | NA | NA | NA | NA | NA |
| CG33110 | 9.028017 | -3.12802 | 3.9239 | -0.79717 | 0.425352 | NA |
| CG31161 | 56.82071 | 1.715879 | 3.951797 | 0.434202 | 0.664142 | NA |
| btn | 0 | NA | NA | NA | NA | NA |
| sit | 0 | NA | NA | NA | NA | NA |
| CG5326 | 0.936057 | 1.478224 | 3.957115 | 0.373561 | 0.708731 | NA |
| CG33099 | 0 | NA | NA | NA | NA | NA |
| CG5376 | 0 | NA | NA | NA | NA | NA |
| CG13847 | 0.178053 | -1.16467 | 4.075067 | -0.2858 | 0.775028 | NA |
| pinta | 23.76536 | -1.92185 | 3.833443 | -0.50134 | 0.616134 | NA |
| CG17625 | 4.491708 | -2.28312 | 3.879964 | -0.58844 | 0.556238 | NA |
| Nup133 | 33.71021 | -4.52804 | 3.780305 | -1.1978 | 0.230996 | NA |
| CG6972 | 0.178053 | -1.16467 | 4.075067 | -0.2858 | 0.775028 | NA |
| CG6985 | 0.132078 | -1.13211 | 4.069918 | -0.27817 | 0.780885 | NA |
| CG7031 | 0.1047 | 0.613261 | 4.062799 | 0.150945 | 0.880019 | NA |
| CG4704 | 0.388135 | 0.584845 | 4.044798 | 0.144592 | 0.885033 | NA |
| Nepl15 | 6.020741 | -0.43663 | 3.950216 | -0.11053 | 0.911986 | NA |

| | | | | | | |
|---|---|---|---|---|---|---|
| Nepl14 | 0 | NA | NA | NA | NA | NA |
| Takl2 | 0 | NA | NA | NA | NA | NA |
| CG4907 | 0 | NA | NA | NA | NA | NA |
| CG13842 | 0 | NA | NA | NA | NA | NA |
| CCAP | 0 | NA | NA | NA | NA | NA |
| Tpl94D | 4.17304 | -2.3219 | 3.804616 | -0.61028 | 0.541674 | NA |
| Gbp3 | 0.63886 | -0.79045 | 3.986802 | -0.19827 | 0.842837 | NA |
| Or94a | 0 | NA | NA | NA | NA | NA |
| Or94b | 0 | NA | NA | NA | NA | NA |
| CG6688 | 0 | NA | NA | NA | NA | NA |
| CG13838 | 0 | NA | NA | NA | NA | NA |
| CG13837 | 0 | NA | NA | NA | NA | NA |
| Gr94a | 0 | NA | NA | NA | NA | NA |
| CG6726 | 4.170776 | -1.48628 | 3.917111 | -0.37943 | 0.704366 | NA |
| CG17110 | 0 | NA | NA | NA | NA | NA |
| CG17109 | 0.357625 | -1.47707 | 4.044794 | -0.36518 | 0.714979 | NA |
| CG6733 | 3.929571 | 1.013179 | 3.94962 | 0.256526 | 0.797545 | NA |
| CG6738 | 0 | NA | NA | NA | NA | NA |
| CG17119 | 0 | NA | NA | NA | NA | NA |
| CG17121 | 26.25092 | -7.35505 | 3.909707 | -1.88123 | 0.059941 | NA |
| lmd | 0.566189 | -0.21299 | 4.010979 | -0.0531 | 0.957651 | NA |
| CG13827 | 49.12118 | 8.211406 | 3.847206 | 2.134382 | 0.032812 | NA |
| bb8 | 0 | NA | NA | NA | NA | NA |
| wam | 0.178053 | -1.16467 | 4.075067 | -0.2858 | 0.775028 | NA |
| fzo | 46.18135 | 8.178615 | 3.876846 | 2.109605 | 0.034892 | NA |
| VhaAC39-2 | 0.283512 | -0.2331 | 4.065094 | -0.05734 | 0.954273 | NA |
| Ir94f | 0 | NA | NA | NA | NA | NA |
| Ir94g | 0.1047 | 0.613261 | 4.062799 | 0.150945 | 0.880019 | NA |
| Ir94h | 0.337716 | 1.464149 | 4.04412 | 0.362044 | 0.717319 | NA |
| CG10164 | 0.356107 | -1.47099 | 4.037764 | -0.36431 | 0.715628 | NA |
| Ugt303B2 | 0 | NA | NA | NA | NA | NA |
| Ugt303B1 | 0 | NA | NA | NA | NA | NA |
| Ugt303B3 | 0.314023 | 1.768229 | 4.015745 | 0.440324 | 0.659702 | NA |
| CG4374 | 1.090133 | -1.54143 | 3.911316 | -0.3941 | 0.69351 | NA |
| CG17380 | 0 | NA | NA | NA | NA | NA |
| CG31515 | 0 | NA | NA | NA | NA | NA |
| CG4408 | 18.78199 | 0.815491 | 3.826623 | 0.21311 | 0.831241 | NA |
| CG16710 | 0 | NA | NA | NA | NA | NA |
| CG18754 | 1.71701 | -1.85109 | 3.972145 | -0.46602 | 0.641203 | NA |
| SPE | 34.34826 | 0.322011 | 3.844929 | 0.08375 | 0.933256 | NA |
| Gba1a | 0.356866 | -1.92326 | 4.037889 | -0.4763 | 0.633858 | NA |
| Qsox4 | 0 | NA | NA | NA | NA | NA |
| Gba1b | 30.29817 | -0.55558 | 3.887344 | -0.14292 | 0.886354 | NA |
| CG10301 | 0 | NA | NA | NA | NA | NA |
| CG10300 | 0 | NA | NA | NA | NA | NA |
| CG10232 | 0 | NA | NA | NA | NA | NA |
| nau | 0 | NA | NA | NA | NA | NA |
| CG31468 | 0.178053 | -1.16467 | 4.075067 | -0.2858 | 0.775028 | NA |
| prt | 0.310131 | -1.94616 | 4.045058 | -0.48112 | 0.630431 | NA |
| CG10252 | 0 | NA | NA | NA | NA | NA |

| | | | | | | |
|---|---|---|---|---|---|---|
| CG10184 | 0 | NA | NA | NA | NA | NA |
| tRNA:Leu-( | 0 | NA | NA | NA | NA | NA |
| CG10183 | 0 | NA | NA | NA | NA | NA |
| CG16723 | 0.104623 | 0.639075 | 4.067674 | 0.157111 | 0.875158 | NA |
| CG33337 | 0 | NA | NA | NA | NA | NA |
| CG10182 | 0 | NA | NA | NA | NA | NA |
| snRNA:U1: | 0.1047 | 0.613261 | 4.062799 | 0.150945 | 0.880019 | NA |
| snRNA:U1: | 0 | NA | NA | NA | NA | NA |
| CG31141 | 0.356107 | -1.47099 | 4.037764 | -0.36431 | 0.715628 | NA |
| CG13599 | 0.356107 | -1.47099 | 4.037764 | -0.36431 | 0.715628 | NA |
| CG12268 | 0 | NA | NA | NA | NA | NA |
| RpS19b | 0.178053 | -1.16467 | 4.075067 | -0.2858 | 0.775028 | NA |
| tbrd-1 | 0 | NA | NA | NA | NA | NA |
| snRNA:U1: | 0.712213 | -1.57119 | 3.998613 | -0.39293 | 0.694369 | NA |
| CG6000 | 35.60687 | 0.38676 | 3.731527 | 0.103646 | 0.91745 | NA |
| Root | 2.120372 | 1.292103 | 3.861043 | 0.334651 | 0.737888 | NA |
| Npc2f | 0 | NA | NA | NA | NA | NA |
| Kal1 | 13.20616 | 1.757063 | 3.858309 | 0.455397 | 0.648823 | NA |
| CG13606 | 0 | NA | NA | NA | NA | NA |
| Acp95EF | 0 | NA | NA | NA | NA | NA |
| CG13607 | 0 | NA | NA | NA | NA | NA |
| tRNA:Leu-' | 17.06212 | 0.611372 | 3.92928 | 0.155594 | 0.876353 | NA |
| CG6432 | 0 | NA | NA | NA | NA | NA |
| Ms | 6.394311 | -1.60476 | 3.781118 | -0.42441 | 0.671264 | NA |
| CG33339 | 0 | NA | NA | NA | NA | NA |
| CG33340 | 0.178053 | -1.16467 | 4.075067 | -0.2858 | 0.775028 | NA |
| CG13613 | 0 | NA | NA | NA | NA | NA |
| CG33341 | 27.51592 | 1.534667 | 3.952215 | 0.388305 | 0.69779 | NA |
| CG13615 | 0 | NA | NA | NA | NA | NA |
| CG13616 | 0.236701 | -0.24276 | 4.020028 | -0.06039 | 0.951846 | NA |
| CG5768 | 30.572 | 4.601396 | 3.882717 | 1.185097 | 0.235979 | NA |
| CG17780 | 0 | NA | NA | NA | NA | NA |
| CG17781 | 0 | NA | NA | NA | NA | NA |
| CG17782 | 0.283512 | -0.2331 | 4.065094 | -0.05734 | 0.954273 | NA |
| CG17784 | 0 | NA | NA | NA | NA | NA |
| CG5762 | 0 | NA | NA | NA | NA | NA |
| CG5728 | 34.04257 | -0.35453 | 3.86776 | -0.09166 | 0.926966 | NA |
| CG5715 | 4.078721 | -0.2095 | 3.773961 | -0.05551 | 0.95573 | NA |
| CG31128 | 0 | NA | NA | NA | NA | NA |
| DZIP1 | 0 | NA | NA | NA | NA | NA |
| CG13618 | 0.104623 | 0.639075 | 4.067674 | 0.157111 | 0.875158 | NA |
| tRNA:Asp-( | 0 | NA | NA | NA | NA | NA |
| snRNA:U6: | 1.691618 | -4.37744 | 3.934873 | -1.11247 | 0.265935 | NA |
| snRNA:U6: | 0.415513 | -1.071 | 4.002654 | -0.26757 | 0.789028 | NA |
| snRNA:U6: | 0 | NA | NA | NA | NA | NA |
| CG13622 | 0 | NA | NA | NA | NA | NA |
| CG13627 | 0 | NA | NA | NA | NA | NA |
| tld | 5.373551 | -2.10067 | 3.859203 | -0.54433 | 0.586216 | NA |
| asp | 0.178053 | -1.16467 | 4.075067 | -0.2858 | 0.775028 | NA |
| Ets96B | 0 | NA | NA | NA | NA | NA |

| | | | | | | |
|---|---|---|---|---|---|---|
| CG6980 | 0 | NA | NA | NA | NA | NA |
| Esp | 0.1047 | 0.613261 | 4.062799 | 0.150945 | 0.880019 | NA |
| CG7016 | 0.104623 | 0.639075 | 4.067674 | 0.157111 | 0.875158 | NA |
| CG31381 | 60.97474 | -0.60687 | 3.732172 | -0.1626 | 0.87083 | NA |
| CG11069 | 0 | NA | NA | NA | NA | NA |
| CG10845 | 0 | NA | NA | NA | NA | NA |
| tRNA:Asp-( | 0 | NA | NA | NA | NA | NA |
| Mink | 0.233093 | 0.692656 | 4.076246 | 0.169925 | 0.865069 | NA |
| Trf4-2 | 0 | NA | NA | NA | NA | NA |
| tRNA:Asp-( | 0 | NA | NA | NA | NA | NA |
| CG13643 | 0 | NA | NA | NA | NA | NA |
| CG31121 | 0.53416 | -1.53481 | 4.012395 | -0.38252 | 0.702078 | NA |
| CycB3 | 15.60029 | 1.74153 | 3.95421 | 0.440424 | 0.65963 | NA |
| CG13640 | 0 | NA | NA | NA | NA | NA |
| ppk22 | 0.104623 | 0.639075 | 4.067674 | 0.157111 | 0.875158 | NA |
| CAH4 | 0 | NA | NA | NA | NA | NA |
| niki | 0 | NA | NA | NA | NA | NA |
| CG5808 | 29.03263 | 0.186978 | 3.839175 | 0.048703 | 0.961156 | NA |
| Ude | 0.660388 | -1.75977 | 4.001266 | -0.4398 | 0.660079 | NA |
| CG13634 | 0.282677 | -0.09599 | 4.057934 | -0.02366 | 0.981127 | NA |
| AstA | 1.503183 | 2.269606 | 3.970131 | 0.57167 | 0.567545 | NA |
| CG31115 | 2.929451 | 1.496835 | 3.962734 | 0.377728 | 0.705633 | NA |
| CG11786 | 2.564026 | 1.129155 | 3.968527 | 0.284527 | 0.776006 | NA |
| fd96Ca | 0 | NA | NA | NA | NA | NA |
| fd96Cb | 0 | NA | NA | NA | NA | NA |
| Mocs2B | 0 | NA | NA | NA | NA | NA |
| CG13653 | 0 | NA | NA | NA | NA | NA |
| CG31103 | 40.41417 | 1.749563 | 3.952192 | 0.442682 | 0.657996 | NA |
| CG11852 | 0 | NA | NA | NA | NA | NA |
| to | 0 | NA | NA | NA | NA | NA |
| CG11854 | 0 | NA | NA | NA | NA | NA |
| CG13658 | 0 | NA | NA | NA | NA | NA |
| CG13659 | 0.2094 | 1.005034 | 4.045341 | 0.248442 | 0.803792 | NA |
| rha | 0 | NA | NA | NA | NA | NA |
| CG10669 | 22.46979 | 1.48289 | 3.906428 | 0.379603 | 0.704241 | NA |
| CG10553 | 0 | NA | NA | NA | NA | NA |
| CG10559 | 0 | NA | NA | NA | NA | NA |
| CG31087 | 0.1047 | 0.613261 | 4.062799 | 0.150945 | 0.880019 | NA |
| CG31099 | 0 | NA | NA | NA | NA | NA |
| CG31288 | 0.233093 | 0.692656 | 4.076246 | 0.169925 | 0.865069 | NA |
| CG31097 | 2.145747 | -2.13625 | 3.989508 | -0.53547 | 0.592328 | NA |
| CG31102 | 1.072874 | -2.0339 | 4.019685 | -0.50598 | 0.612867 | NA |
| bam | 0 | NA | NA | NA | NA | NA |
| Cad96Cb | 0.104623 | 0.639075 | 4.067674 | 0.157111 | 0.875158 | NA |
| CG5116 | 7.19094 | -0.33858 | 3.795436 | -0.08921 | 0.928917 | NA |
| CG5112 | 0 | NA | NA | NA | NA | NA |
| CG5107 | 51.77445 | -1.92043 | 3.951661 | -0.48598 | 0.626982 | NA |
| CG5111 | 0.104623 | 0.639075 | 4.067674 | 0.157111 | 0.875158 | NA |
| CG4582 | 0 | NA | NA | NA | NA | NA |
| Lgr3 | 8.579107 | 1.524292 | 3.95506 | 0.385403 | 0.699939 | NA |

| | | | | | | |
|---|---|---|---|---|---|---|
| CG5024 | 0 | NA | NA | NA | NA | NA |
| CG17770 | 0 | NA | NA | NA | NA | NA |
| Acam | 0 | NA | NA | NA | NA | NA |
| CG14354 | 0.1047 | 0.613261 | 4.062799 | 0.150945 | 0.880019 | NA |
| CG14545 | 0.178053 | -1.16467 | 4.075067 | -0.2858 | 0.775028 | NA |
| Mst57Da | 0 | NA | NA | NA | NA | NA |
| Mst57Db | 0 | NA | NA | NA | NA | NA |
| Mst57Dc | 0 | NA | NA | NA | NA | NA |
| CG31093 | 0 | NA | NA | NA | NA | NA |
| CG31380 | 0 | NA | NA | NA | NA | NA |
| CG4960 | 0 | NA | NA | NA | NA | NA |
| CG17196 | 0 | NA | NA | NA | NA | NA |
| CG17197 | 0 | NA | NA | NA | NA | NA |
| CG4730 | 0 | NA | NA | NA | NA | NA |
| CG5948 | 51.16004 | -3.52142 | 3.899401 | -0.90307 | 0.36649 | NA |
| Mco3 | 0.132078 | -1.13211 | 4.069918 | -0.27817 | 0.780885 | NA |
| CG14540 | 0 | NA | NA | NA | NA | NA |
| boss | 3.769197 | 1.70343 | 3.964029 | 0.429722 | 0.667398 | NA |
| E(spl)mdel | 0 | NA | NA | NA | NA | NA |
| E(spl)mgar | 0 | NA | NA | NA | NA | NA |
| E(spl)malp | 0 | NA | NA | NA | NA | NA |
| Kaz-m1 | 0 | NA | NA | NA | NA | NA |
| E(spl)m7-H | 0.356107 | -1.47099 | 4.037764 | -0.36431 | 0.715628 | NA |
| E(spl)m8-H | 0.104623 | 0.639075 | 4.067674 | 0.157111 | 0.875158 | NA |
| CG14544 | 2.408098 | 1.677625 | 3.970879 | 0.422482 | 0.672673 | NA |
| dysf | 0 | NA | NA | NA | NA | NA |
| CG14551 | 0 | NA | NA | NA | NA | NA |
| PIG-P | 0 | NA | NA | NA | NA | NA |
| E(spl)m4-B | 0.356107 | -1.47099 | 4.037764 | -0.36431 | 0.715628 | NA |
| E(spl)m2-B | 0 | NA | NA | NA | NA | NA |
| CG14546 | 0.1047 | 0.613261 | 4.062799 | 0.150945 | 0.880019 | NA |
| CG14556 | 0 | NA | NA | NA | NA | NA |
| CG31086 | 0 | NA | NA | NA | NA | NA |
| Ald2 | 0 | NA | NA | NA | NA | NA |
| asRNA:CR3 | 0 | NA | NA | NA | NA | NA |
| Hex-t1 | 0 | NA | NA | NA | NA | NA |
| Hex-t2 | 0 | NA | NA | NA | NA | NA |
| CG14237 | 0 | NA | NA | NA | NA | NA |
| CG14238 | 0 | NA | NA | NA | NA | NA |
| Pdf | 0.178053 | -1.16467 | 4.075067 | -0.2858 | 0.775028 | NA |
| ppk15 | 0 | NA | NA | NA | NA | NA |
| CG6036 | 0 | NA | NA | NA | NA | NA |
| TwdlM | 0 | NA | NA | NA | NA | NA |
| TwdlJ | 0 | NA | NA | NA | NA | NA |
| TwdlN | 0.282753 | -0.25398 | 4.020204 | -0.06318 | 0.949626 | NA |
| TwdlH | 0 | NA | NA | NA | NA | NA |
| TwdlR | 0.466186 | 0.959471 | 4.033986 | 0.237847 | 0.812 | NA |
| TwdlS | 0 | NA | NA | NA | NA | NA |
| TwdlD | 0.1047 | 0.613261 | 4.062799 | 0.150945 | 0.880019 | NA |
| CG14244 | 0 | NA | NA | NA | NA | NA |

| | | | | | | |
|---|---|---|---|---|---|---|
| CG6403 | 0 | NA | NA | NA | NA | NA |
| TwdlQ | 0 | NA | NA | NA | NA | NA |
| TwdlK | 0 | NA | NA | NA | NA | NA |
| TwdlO | 11.93297 | -3.479 | 3.906416 | -0.89059 | 0.373152 | NA |
| TwdlL | 51.08661 | -1.89434 | 3.951656 | -0.47938 | 0.63167 | NA |
| TwdlB | 0 | NA | NA | NA | NA | NA |
| TwdlP | 0.1047 | 0.613261 | 4.062799 | 0.150945 | 0.880019 | NA |
| ms(3)K81 | 0 | NA | NA | NA | NA | NA |
| Spag1 | 1.360102 | 1.453444 | 3.975267 | 0.365622 | 0.714647 | NA |
| CG14253 | 5.547263 | -1.89916 | 3.957803 | -0.47985 | 0.631332 | NA |
| TwdlC | 0 | NA | NA | NA | NA | NA |
| CG31077 | 0 | NA | NA | NA | NA | NA |
| CG6295 | 0 | NA | NA | NA | NA | NA |
| CG6296 | 0 | NA | NA | NA | NA | NA |
| CG31076 | 0 | NA | NA | NA | NA | NA |
| Gr97a | 0 | NA | NA | NA | NA | NA |
| tRNA:Gln-¯ | 0 | NA | NA | NA | NA | NA |
| tRNA:Gln-¯ | 0 | NA | NA | NA | NA | NA |
| ball | 26.98997 | -3.91961 | 3.807038 | -1.02957 | 0.303212 | NA |
| TwdlT | 23.55419 | 6.962447 | 3.913302 | 1.779175 | 0.075211 | NA |
| CG14258 | 22.36002 | -0.85662 | 3.865561 | -0.2216 | 0.824623 | NA |
| CG17189 | 0.365171 | -0.43584 | 4.056359 | -0.10745 | 0.914434 | NA |
| CG5882 | 0 | NA | NA | NA | NA | NA |
| CG5909 | 1.788123 | -2.11324 | 3.996051 | -0.52883 | 0.596922 | NA |
| ppk31 | 0 | NA | NA | NA | NA | NA |
| spn-D | 0.132078 | -1.13211 | 4.069918 | -0.27817 | 0.780885 | NA |
| CG31068 | 20.13944 | -0.66616 | 3.95203 | -0.16856 | 0.866141 | NA |
| PolA2 | 0 | NA | NA | NA | NA | NA |
| Ser | 51.15426 | 2.881849 | 3.865287 | 0.745572 | 0.455926 | NA |
| CG6059 | 0 | NA | NA | NA | NA | NA |
| Cpr97Eb | 0.233093 | 0.692656 | 4.076246 | 0.169925 | 0.865069 | NA |
| Cpr97Ea | 0 | NA | NA | NA | NA | NA |
| Nep5 | 0.356107 | -1.47099 | 4.037764 | -0.36431 | 0.715628 | NA |
| CG6271 | 0 | NA | NA | NA | NA | NA |
| CG6277 | 0 | NA | NA | NA | NA | NA |
| CG6283 | 0 | NA | NA | NA | NA | NA |
| CG17191 | 0.31387 | 1.261782 | 4.037145 | 0.312543 | 0.754628 | NA |
| CG17192 | 0 | NA | NA | NA | NA | NA |
| CG14260 | 0 | NA | NA | NA | NA | NA |
| CG3330 | 0.2094 | 1.005034 | 4.045341 | 0.248442 | 0.803792 | NA |
| CG3339 | 4.862478 | -0.64598 | 3.931505 | -0.16431 | 0.869488 | NA |
| CG14262 | 0 | NA | NA | NA | NA | NA |
| CG5984 | 1.151699 | 1.609751 | 3.989673 | 0.403479 | 0.686596 | NA |
| CG13978 | 0 | NA | NA | NA | NA | NA |
| Cyp6a18 | 0 | NA | NA | NA | NA | NA |
| Gr98a | 0 | NA | NA | NA | NA | NA |
| CG12885 | 0.396233 | -1.6814 | 4.028351 | -0.41739 | 0.676392 | NA |
| Gr98b | 0.178053 | -1.16467 | 4.075067 | -0.2858 | 0.775028 | NA |
| Gr98c | 0.178053 | -1.16467 | 4.075067 | -0.2858 | 0.775028 | NA |
| Gr98d | 0 | NA | NA | NA | NA | NA |

| | | | | | | |
|---|---|---|---|---|---|---|
| CG5639 | 0.466186 | 0.959471 | 4.033986 | 0.237847 | 0.812 | NA |
| CG13972 | 0 | NA | NA | NA | NA | NA |
| CG5402 | 3.664497 | 1.702077 | 3.964384 | 0.429342 | 0.667674 | NA |
| Ets98B | 31.76204 | -6.45629 | 3.870095 | -1.66825 | 0.095266 | NA |
| Or98P | 1.570499 | 1.642769 | 3.980387 | 0.412716 | 0.679815 | NA |
| Or98a | 28.09611 | -0.52971 | 3.87027 | -0.13687 | 0.891137 | NA |
| CG12426 | 0.488184 | -2.32943 | 4.022674 | -0.57908 | 0.562538 | NA |
| Nep7 | 0 | NA | NA | NA | NA | NA |
| CR31054 | 0 | NA | NA | NA | NA | NA |
| CG4815 | 0.178053 | -1.16467 | 4.075067 | -0.2858 | 0.775028 | NA |
| CG31131 | 0 | NA | NA | NA | NA | NA |
| CG31253 | 0 | NA | NA | NA | NA | NA |
| CG4884 | 0 | NA | NA | NA | NA | NA |
| nenya | 0 | NA | NA | NA | NA | NA |
| Mst98Ca | 0.178812 | -1.08808 | 4.061735 | -0.26789 | 0.788788 | NA |
| Mst98Cb | 0 | NA | NA | NA | NA | NA |
| UQCR-14L | 0 | NA | NA | NA | NA | NA |
| CG12516 | 0 | NA | NA | NA | NA | NA |
| CG1894 | 0 | NA | NA | NA | NA | NA |
| Dhc98D | 5.423543 | -2.07187 | 3.783525 | -0.5476 | 0.583965 | NA |
| mIF2 | 7.174845 | -1.8917 | 3.868038 | -0.48906 | 0.624801 | NA |
| Ppn | 44.69223 | -8.65874 | 3.908682 | -2.21526 | 0.026742 | NA |
| CG9988 | 0 | NA | NA | NA | NA | NA |
| CG33346 | 0 | NA | NA | NA | NA | NA |
| Sid | 0 | NA | NA | NA | NA | NA |
| intr | 0 | NA | NA | NA | NA | NA |
| aqrs | 0 | NA | NA | NA | NA | NA |
| CG9997 | 0.178053 | -1.16467 | 4.075067 | -0.2858 | 0.775028 | NA |
| CG10000 | 0 | NA | NA | NA | NA | NA |
| CG31050 | 0 | NA | NA | NA | NA | NA |
| Nepl18 | 3.894907 | 2.998356 | 3.922914 | 0.764318 | 0.444677 | NA |
| CG11828 | 0 | NA | NA | NA | NA | NA |
| Ssl2 | 1.351073 | -1.13435 | 3.970931 | -0.28566 | 0.775136 | NA |
| wat | 0 | NA | NA | NA | NA | NA |
| Nepl19 | 0.411147 | -0.29058 | 4.046973 | -0.0718 | 0.94276 | NA |
| Cpsf100 | 34.97031 | 2.512594 | 3.749483 | 0.670118 | 0.502783 | NA |
| CG11837 | 2.019106 | 1.540265 | 3.919652 | 0.39296 | 0.694349 | NA |
| CG11841 | 59.53394 | 0.420113 | 3.844034 | 0.10929 | 0.912973 | NA |
| CG11842 | 0.178053 | -1.16467 | 4.075067 | -0.2858 | 0.775028 | NA |
| CG11843 | 1.452855 | -1.83946 | 3.975716 | -0.46267 | 0.643599 | NA |
| Brd8 | 10.75107 | -2.54421 | 3.802661 | -0.66906 | 0.503458 | NA |
| stg | 37.0637 | 2.971108 | 3.919966 | 0.757942 | 0.448486 | NA |
| CG31445 | 0.209247 | 1.042804 | 4.047837 | 0.25762 | 0.7967 | NA |
| CR31427 | 0 | NA | NA | NA | NA | NA |
| CG11951 | 15.58734 | 0.407201 | 3.844339 | 0.105922 | 0.915644 | NA |
| CG14507 | 0 | NA | NA | NA | NA | NA |
| Cyt-c1L | 0.178812 | -1.08808 | 4.061735 | -0.26789 | 0.788788 | NA |
| CG14511 | 0 | NA | NA | NA | NA | NA |
| yem | 28.00314 | 3.230829 | 3.850143 | 0.839145 | 0.401388 | NA |
| beta4GalN | 0 | NA | NA | NA | NA | NA |

| | | | | | | |
|---|---|---|---|---|---|---|
| Diedel | 0 | NA | NA | NA | NA | NA |
| CG2310 | 3.112374 | -0.89023 | 3.914142 | -0.22744 | 0.820082 | NA |
| Spase12 | 0 | NA | NA | NA | NA | NA |
| CG2321 | 5.758496 | 1.720238 | 3.959653 | 0.434441 | 0.663968 | NA |
| CG14506 | 0 | NA | NA | NA | NA | NA |
| ND-20L | 0 | NA | NA | NA | NA | NA |
| ppk30 | 6.616054 | -2.22778 | 3.964757 | -0.5619 | 0.574188 | NA |
| Obp99a | 0 | NA | NA | NA | NA | NA |
| ppk19 | 0 | NA | NA | NA | NA | NA |
| Cap-D2 | 0.178053 | -1.16467 | 4.075067 | -0.2858 | 0.775028 | NA |
| Obp99d | 33.99368 | -1.81019 | 3.951967 | -0.45805 | 0.646919 | NA |
| CG15506 | 12.35451 | 2.945353 | 3.925793 | 0.750257 | 0.4531 | NA |
| Obp99b | 0 | NA | NA | NA | NA | NA |
| CG7568 | 0 | NA | NA | NA | NA | NA |
| CG31041 | 0 | NA | NA | NA | NA | NA |
| CG11470 | 0.132078 | -1.13211 | 4.069918 | -0.27817 | 0.780885 | NA |
| CG7567 | 0 | NA | NA | NA | NA | NA |
| Prtl99C | 0 | NA | NA | NA | NA | NA |
| Cad99C | 49.11413 | 1.723013 | 3.637413 | 0.473692 | 0.63572 | NA |
| CG15515 | 0 | NA | NA | NA | NA | NA |
| CG31036 | 8.162549 | 1.532669 | 3.903074 | 0.392682 | 0.694554 | NA |
| Jon99Ciii | 0.932373 | 1.053996 | 3.996193 | 0.26375 | 0.791972 | NA |
| alpha-Man | 0 | NA | NA | NA | NA | NA |
| Jon99Ci | 0 | NA | NA | NA | NA | NA |
| Jon99Cii | 0.1047 | 0.613261 | 4.062799 | 0.150945 | 0.880019 | NA |
| Capa | 0 | NA | NA | NA | NA | NA |
| CG15517 | 0 | NA | NA | NA | NA | NA |
| CG7829 | 0 | NA | NA | NA | NA | NA |
| ncd | 0.310131 | -1.94616 | 4.045058 | -0.48112 | 0.630431 | NA |
| fig | 4.083297 | 1.7071 | 3.963068 | 0.430752 | 0.666649 | NA |
| CG7896 | 0.744924 | -1.68269 | 3.955513 | -0.4254 | 0.670542 | NA |
| Sas-6 | 0 | NA | NA | NA | NA | NA |
| Sry-beta | 0 | NA | NA | NA | NA | NA |
| Sry-alpha | 0 | NA | NA | NA | NA | NA |
| Rpp14b | 0 | NA | NA | NA | NA | NA |
| CG7950 | 0.2094 | 1.005034 | 4.045341 | 0.248442 | 0.803792 | NA |
| CG15529 | 0.837599 | 1.568055 | 4.001801 | 0.391837 | 0.695178 | NA |
| CG15530 | 0 | NA | NA | NA | NA | NA |
| CG15531 | 0 | NA | NA | NA | NA | NA |
| CG9747 | 0 | NA | NA | NA | NA | NA |
| CG11498 | 0 | NA | NA | NA | NA | NA |
| CG31029 | 47.89633 | -1.70452 | 3.951708 | -0.43134 | 0.666223 | NA |
| Ppi1 | 0 | NA | NA | NA | NA | NA |
| CG15528 | 0 | NA | NA | NA | NA | NA |
| janB | 0 | NA | NA | NA | NA | NA |
| ocn | 0 | NA | NA | NA | NA | NA |
| RpS28a | 0 | NA | NA | NA | NA | NA |
| Cec-Psi1 | 0 | NA | NA | NA | NA | NA |
| CecC | 0 | NA | NA | NA | NA | NA |
| CG9682 | 0.178053 | -1.16467 | 4.075067 | -0.2858 | 0.775028 | NA |

| | | | | | | |
|---|---|---|---|---|---|---|
| CG18404 | 0 | NA | NA | NA | NA | NA |
| CG15533 | 0 | NA | NA | NA | NA | NA |
| CG2217 | 0 | NA | NA | NA | NA | NA |
| tRNA:Leu-⁻ | 0.1047 | 0.613261 | 4.062799 | 0.150945 | 0.880019 | NA |
| tRNA:Leu-⁻ | 0 | NA | NA | NA | NA | NA |
| CG9733 | 0 | NA | NA | NA | NA | NA |
| CG9737 | 0 | NA | NA | NA | NA | NA |
| CG15534 | 5.438891 | 2.034849 | 3.783406 | 0.537835 | 0.590691 | NA |
| spdo | 0.178053 | -1.16467 | 4.075067 | -0.2858 | 0.775028 | NA |
| Jon99Fi | 0 | NA | NA | NA | NA | NA |
| PH4alphaS | 0.178053 | -1.16467 | 4.075067 | -0.2858 | 0.775028 | NA |
| CG15539 | 0 | NA | NA | NA | NA | NA |
| CG9698 | 0 | NA | NA | NA | NA | NA |
| CG9702 | 0 | NA | NA | NA | NA | NA |
| CG31019 | 7.428251 | 1.522001 | 3.955692 | 0.384762 | 0.700414 | NA |
| CG31021 | 0 | NA | NA | NA | NA | NA |
| PH4alphaN | 0 | NA | NA | NA | NA | NA |
| PH4alphaP | 0 | NA | NA | NA | NA | NA |
| CG31013 | 0 | NA | NA | NA | NA | NA |
| CG2267 | 0.104623 | 0.639075 | 4.067674 | 0.157111 | 0.875158 | NA |
| CG9717 | 0.104623 | 0.639075 | 4.067674 | 0.157111 | 0.875158 | NA |
| Rpt6R | 0 | NA | NA | NA | NA | NA |
| CG31524 | 0 | NA | NA | NA | NA | NA |
| PH4alphaN | 0.1047 | 0.613261 | 4.062799 | 0.150945 | 0.880019 | NA |
| PH4alphaS | 0 | NA | NA | NA | NA | NA |
| CG31371 | 2.810487 | -2.67968 | 3.85456 | -0.6952 | 0.486932 | NA |
| PH4alphaN | 0 | NA | NA | NA | NA | NA |
| PH4alphaN | 0 | NA | NA | NA | NA | NA |
| Jon99Fii | 0 | NA | NA | NA | NA | NA |
| Osi23 | 0 | NA | NA | NA | NA | NA |
| Spn100A | 0.1047 | 0.613261 | 4.062799 | 0.150945 | 0.880019 | NA |
| eIF4H2 | 0.536437 | -1.71175 | 4.038042 | -0.42391 | 0.671635 | NA |
| CG15543 | 0 | NA | NA | NA | NA | NA |
| CG15544 | 50.30335 | 4.031695 | 3.882542 | 1.038416 | 0.299076 | NA |
| Cpr100A | 0 | NA | NA | NA | NA | NA |
| Cyp4c3 | 0 | NA | NA | NA | NA | NA |
| CG15546 | 0 | NA | NA | NA | NA | NA |
| CG15545 | 0 | NA | NA | NA | NA | NA |
| Npc2h | 0 | NA | NA | NA | NA | NA |
| Npc2g | 0 | NA | NA | NA | NA | NA |
| CG11313 | 0 | NA | NA | NA | NA | NA |
| CG31010 | 0 | NA | NA | NA | NA | NA |
| Pka-C2 | 0 | NA | NA | NA | NA | NA |
| CG12069 | 0 | NA | NA | NA | NA | NA |
| CG15548 | 0 | NA | NA | NA | NA | NA |
| Sox100B | 0 | NA | NA | NA | NA | NA |
| CG15554 | 27.04578 | -0.62809 | 3.927409 | -0.15993 | 0.87294 | NA |
| CG15556 | 1.328986 | 1.486893 | 3.928973 | 0.378443 | 0.705101 | NA |
| CG11318 | 0 | NA | NA | NA | NA | NA |
| CG31008 | 0 | NA | NA | NA | NA | NA |

| | | | | | | |
|---|---|---|---|---|---|---|
| Ctr1C | 0 | NA | NA | NA | NA | NA |
| CG15550 | 0 | NA | NA | NA | NA | NA |
| CG15549 | 0 | NA | NA | NA | NA | NA |
| bnk | 13.76109 | -3.67364 | 3.810035 | -0.9642 | 0.334945 | NA |
| CAH6 | 0.178053 | -1.16467 | 4.075067 | -0.2858 | 0.775028 | NA |
| CAH16 | 0.132078 | -1.13211 | 4.069918 | -0.27817 | 0.780885 | NA |
| CAH5 | 0 | NA | NA | NA | NA | NA |
| ppk24 | 0.178812 | -1.08808 | 4.061735 | -0.26789 | 0.788788 | NA |
| pHCl-2 | 0.178812 | -1.08808 | 4.061735 | -0.26789 | 0.788788 | NA |
| CG15564 | 0 | NA | NA | NA | NA | NA |
| CG15563 | 0 | NA | NA | NA | NA | NA |
| Zwilch | 13.80376 | 4.323275 | 3.857939 | 1.120618 | 0.262451 | NA |
| gskt | 0.132078 | -1.13211 | 4.069918 | -0.27817 | 0.780885 | NA |
| CG11333 | 0.132078 | -1.13211 | 4.069918 | -0.27817 | 0.780885 | NA |
| mey | 0 | NA | NA | NA | NA | NA |
| nyo | 0.209247 | 1.042804 | 4.047837 | 0.25762 | 0.7967 | NA |
| CG1635 | 34.16379 | -3.54834 | 3.767886 | -0.94173 | 0.34633 | NA |
| CG1638 | 0 | NA | NA | NA | NA | NA |
| PNPase | 1.416981 | 1.957611 | 3.905528 | 0.501241 | 0.616201 | NA |
| Ugt35D1 | 0.178053 | -1.16467 | 4.075067 | -0.2858 | 0.775028 | NA |
| CG31204 | 0 | NA | NA | NA | NA | NA |
| Ir100a | 0.1047 | 0.613261 | 4.062799 | 0.150945 | 0.880019 | NA |
| CG1971 | 0 | NA | NA | NA | NA | NA |
| CG2150 | 0 | NA | NA | NA | NA | NA |
| Smvt | 0 | NA | NA | NA | NA | NA |
| CG2187 | 0.209247 | 1.042804 | 4.047837 | 0.25762 | 0.7967 | NA |
| salt | 0.356866 | -1.92326 | 4.037889 | -0.4763 | 0.633858 | NA |
| CG11550 | 0 | NA | NA | NA | NA | NA |
| CG2003 | 4.582433 | -1.5598 | 3.835416 | -0.40668 | 0.68424 | NA |
| CG2053 | 0.132078 | -1.13211 | 4.069918 | -0.27817 | 0.780885 | NA |
| His-Psi:CR3 | 0 | NA | NA | NA | NA | NA |
| His-Psi:CR3 | 0 | NA | NA | NA | NA | NA |
| His-Psi:CR3 | 0 | NA | NA | NA | NA | NA |
| CG11630 | 2.443261 | -3.33948 | 3.872261 | -0.86241 | 0.388461 | NA |
| CG1421 | 0 | NA | NA | NA | NA | NA |
| CG1428 | 0 | NA | NA | NA | NA | NA |
| CG11634 | 0 | NA | NA | NA | NA | NA |
| CR31700 | 0 | NA | NA | NA | NA | NA |
| tplus3a | 0 | NA | NA | NA | NA | NA |
| ttm3 | 0 | NA | NA | NA | NA | NA |
| CG10834 | 0 | NA | NA | NA | NA | NA |
| CG15219 | 0.282753 | -0.25398 | 4.020204 | -0.06318 | 0.949626 | NA |
| tplus3b | 0 | NA | NA | NA | NA | NA |
| CG31601 | 0.178053 | -1.16467 | 4.075067 | -0.2858 | 0.775028 | NA |
| CG2528 | 1.437098 | -2.04622 | 3.922998 | -0.5216 | 0.601952 | NA |
| CG11629 | 0 | NA | NA | NA | NA | NA |
| CG3635 | 1.589402 | -0.20894 | 3.970589 | -0.05262 | 0.958034 | NA |
| CR12628 | 12.77937 | -0.77664 | 3.73913 | -0.20771 | 0.835458 | NA |
| CG7882 | 0 | NA | NA | NA | NA | NA |
| tRNA:Arg-/ | 27.68305 | 0.038243 | 3.931617 | 0.009727 | 0.992239 | NA |

| | | | | | | |
|---|---|---|---|---|---|---|
| tRNA:Arg-/ | 0 | NA | NA | NA | NA | NA |
| tRNA:Asn-( | 0 | NA | NA | NA | NA | NA |
| tRNA:Asn-( | 0.396233 | -1.6814 | 4.028351 | -0.41739 | 0.676392 | NA |
| tRNA:Lys-( | 0 | NA | NA | NA | NA | NA |
| tRNA:Ile-A | 14.30498 | -2.25603 | 3.957513 | -0.57006 | 0.568635 | NA |
| tRNA:Arg-/ | 4.083297 | 1.7071 | 3.963068 | 0.430752 | 0.666649 | NA |
| tRNA:Arg-/ | 0.132078 | -1.13211 | 4.069918 | -0.27817 | 0.780885 | NA |
| CG8343 | 0 | NA | NA | NA | NA | NA |
| tRNA:Asn-( | 0.178053 | -1.16467 | 4.075067 | -0.2858 | 0.775028 | NA |
| eIF3f2 | 0 | NA | NA | NA | NA | NA |
| CG17994 | 0.132078 | -1.13211 | 4.069918 | -0.27817 | 0.780885 | NA |
| CG30432 | 0 | NA | NA | NA | NA | NA |
| tRNA:Asn-( | 0 | NA | NA | NA | NA | NA |
| CG11211 | 43.13637 | 1.749881 | 3.952112 | 0.442771 | 0.657931 | NA |
| tRNA:Arg-/ | 0.519531 | -0.79833 | 3.992004 | -0.19998 | 0.841494 | NA |
| tRNA:Lys-( | 0 | NA | NA | NA | NA | NA |
| tRNA:Lys-( | 0.178812 | -1.08808 | 4.061735 | -0.26789 | 0.788788 | NA |
| tRNA:Lys-( | 0 | NA | NA | NA | NA | NA |
| tRNA:Asn-( | 0 | NA | NA | NA | NA | NA |
| tRNA:Asn-( | 0.233093 | 0.692656 | 4.076246 | 0.169925 | 0.865069 | NA |
| tRNA:Lys-( | 32.29061 | -2.20787 | 3.740898 | -0.5902 | 0.555058 | NA |
| CG7856 | 0 | NA | NA | NA | NA | NA |
| Strica | 0 | NA | NA | NA | NA | NA |
| CG7881 | 0.69928 | 1.019866 | 4.009445 | 0.254366 | 0.799213 | NA |
| Tsp42A | 0.209247 | 1.042804 | 4.047837 | 0.25762 | 0.7967 | NA |
| Or42b | 0.282677 | -0.09599 | 4.057934 | -0.02366 | 0.981127 | NA |
| Or42a | 0.178812 | -1.08808 | 4.061735 | -0.26789 | 0.788788 | NA |
| scaf | 50.36067 | 1.75058 | 3.951939 | 0.442967 | 0.657789 | NA |
| Ugt50B3 | 0 | NA | NA | NA | NA | NA |
| TpnC4 | 0.132078 | -1.13211 | 4.069918 | -0.27817 | 0.780885 | NA |
| CG30440 | 1.788123 | -2.11324 | 3.996051 | -0.52883 | 0.596922 | NA |
| CG3270 | 0.807012 | 0.227825 | 3.984055 | 0.057184 | 0.954399 | NA |
| Zip42C.2 | 0 | NA | NA | NA | NA | NA |
| Zip42C.1 | 0 | NA | NA | NA | NA | NA |
| chk | 0 | NA | NA | NA | NA | NA |
| Spn42Dc | 1.762986 | -3.23443 | 3.951497 | -0.81853 | 0.413053 | NA |
| Spn42De | 0 | NA | NA | NA | NA | NA |
| CheB42a | 0.566265 | -0.7478 | 3.984294 | -0.18769 | 0.851121 | NA |
| ppk25 | 33.27465 | 0.260813 | 3.950091 | 0.066027 | 0.947356 | NA |
| Tsp42Ea | 8.115026 | -2.72109 | 3.809251 | -0.71434 | 0.475018 | NA |
| Tsp42Eb | 0 | NA | NA | NA | NA | NA |
| CG30160 | 0 | NA | NA | NA | NA | NA |
| Tsp42Ec | 0 | NA | NA | NA | NA | NA |
| Cyp6u1 | 45.81478 | 7.075764 | 3.850556 | 1.837596 | 0.066122 | NA |
| tRNA:Lys-( | 6.460049 | 1.480691 | 3.957408 | 0.374157 | 0.708288 | NA |
| tRNA:Lys-( | 5.758496 | 1.720238 | 3.959653 | 0.434441 | 0.663968 | NA |
| tRNA:Lys-( | 3.140998 | 1.694072 | 3.966495 | 0.427096 | 0.66931 | NA |
| tRNA:Lys-( | 0 | NA | NA | NA | NA | NA |
| CheB42c | 24.18874 | -3.28208 | 3.90749 | -0.83994 | 0.400939 | NA |
| CheB42b | 11.51699 | 1.736994 | 3.955359 | 0.43915 | 0.660553 | NA |

| | | | | | | |
|---|---|---|---|---|---|---|
| Spn42Db | 0.264155 | -1.54099 | 4.048976 | -0.38059 | 0.703509 | NA |
| CG9447 | 0 | NA | NA | NA | NA | NA |
| CG9445 | 0.178053 | -1.16467 | 4.075067 | -0.2858 | 0.775028 | NA |
| GNBP-like3 | 0 | NA | NA | NA | NA | NA |
| CG30154 | 0 | NA | NA | NA | NA | NA |
| Bbd | 41.81288 | -0.306 | 3.838729 | -0.07971 | 0.936464 | NA |
| CG13423 | 0.178053 | -1.16467 | 4.075067 | -0.2858 | 0.775028 | NA |
| lms | 0 | NA | NA | NA | NA | NA |
| tRNA:Trp-( | 0 | NA | NA | NA | NA | NA |
| CG13437 | 0 | NA | NA | NA | NA | NA |
| CG13438 | 0 | NA | NA | NA | NA | NA |
| Gr57a | 0 | NA | NA | NA | NA | NA |
| exu | 38.26543 | -7.13272 | 3.84555 | -1.8548 | 0.063625 | NA |
| tRNA:Trp-( | 0 | NA | NA | NA | NA | NA |
| Rcd6 | 5.547263 | -1.89916 | 3.957803 | -0.47985 | 0.631332 | NA |
| Hil | 5.893793 | 2.736316 | 3.846221 | 0.71143 | 0.476818 | NA |
| Cpr57A | 0 | NA | NA | NA | NA | NA |
| Obp57d | 0 | NA | NA | NA | NA | NA |
| Obp57e | 0 | NA | NA | NA | NA | NA |
| CG13427 | 0 | NA | NA | NA | NA | NA |
| CG30151 | 0 | NA | NA | NA | NA | NA |
| CkIIbeta2 | 0 | NA | NA | NA | NA | NA |
| CG13871 | 0 | NA | NA | NA | NA | NA |
| CG13870 | 0 | NA | NA | NA | NA | NA |
| CG16741 | 0 | NA | NA | NA | NA | NA |
| Obp57a | 35.05832 | -1.90778 | 3.831405 | -0.49793 | 0.618532 | NA |
| CG13426 | 0 | NA | NA | NA | NA | NA |
| CG11192 | 0 | NA | NA | NA | NA | NA |
| Cpr56F | 0 | NA | NA | NA | NA | NA |
| Mpcp1 | 0 | NA | NA | NA | NA | NA |
| plu | 0 | NA | NA | NA | NA | NA |
| tRNA:Gly-( | 0 | NA | NA | NA | NA | NA |
| CG11044 | 0 | NA | NA | NA | NA | NA |
| tRNA:Gly-( | 0 | NA | NA | NA | NA | NA |
| Efhc1.2 | 1.712559 | -0.44667 | 3.901753 | -0.11448 | 0.908857 | NA |
| CG9864 | 0.209247 | 1.042804 | 4.047837 | 0.25762 | 0.7967 | NA |
| ppk6 | 0 | NA | NA | NA | NA | NA |
| CG13869 | 0 | NA | NA | NA | NA | NA |
| CG16894 | 0 | NA | NA | NA | NA | NA |
| tRNA:Arg-( | 0 | NA | NA | NA | NA | NA |
| Obp56e | 0 | NA | NA | NA | NA | NA |
| Obp56f | 0.1047 | 0.613261 | 4.062799 | 0.150945 | 0.880019 | NA |
| CG8517 | 0 | NA | NA | NA | NA | NA |
| tRNA:Ile-A | 0 | NA | NA | NA | NA | NA |
| tRNA:iMet | 0 | NA | NA | NA | NA | NA |
| tRNA:Lys-( | 0 | NA | NA | NA | NA | NA |
| Obp56h | 0 | NA | NA | NA | NA | NA |
| Obp56i | 0 | NA | NA | NA | NA | NA |
| CG30447 | 0 | NA | NA | NA | NA | NA |
| CG13872 | 0.178812 | -1.08808 | 4.061735 | -0.26789 | 0.788788 | NA |

| | | | | | | |
|---|---|---|---|---|---|---|
| Obp56g | 0 | NA | NA | NA | NA | NA |
| Or56a | 0 | NA | NA | NA | NA | NA |
| tRNA:Glu-( | 0 | NA | NA | NA | NA | NA |
| tRNA:Glu-⁻ | 0 | NA | NA | NA | NA | NA |
| tRNA:Glu-⁻ | 0 | NA | NA | NA | NA | NA |
| tRNA:Glu-⁻ | 0 | NA | NA | NA | NA | NA |
| tRNA:iMet | 0.357625 | -1.47707 | 4.044794 | -0.36518 | 0.714979 | NA |
| CG15120 | 0.1047 | 0.613261 | 4.062799 | 0.150945 | 0.880019 | NA |
| Ir56b | 0 | NA | NA | NA | NA | NA |
| Ir56c | 0 | NA | NA | NA | NA | NA |
| CG15124 | 0 | NA | NA | NA | NA | NA |
| CG15125 | 52.27531 | -1.89497 | 3.95164 | -0.47954 | 0.631554 | NA |
| CG11018 | 24.69853 | -1.91759 | 3.952484 | -0.48516 | 0.627563 | NA |
| isopeptida | 7.616767 | 3.042683 | 3.856291 | 0.789018 | 0.430101 | NA |
| CG15126 | 0 | NA | NA | NA | NA | NA |
| Obp56a | 4.273279 | -1.68149 | 3.959707 | -0.42465 | 0.671092 | NA |
| Obp56d | 6.075574 | -1.90118 | 3.957216 | -0.48043 | 0.630919 | NA |
| Obp56c | 0 | NA | NA | NA | NA | NA |
| Obp56b | 0 | NA | NA | NA | NA | NA |
| CG15128 | 0 | NA | NA | NA | NA | NA |
| CG15127 | 0 | NA | NA | NA | NA | NA |
| CG15905 | 0.310131 | -1.94616 | 4.045058 | -0.48112 | 0.630431 | NA |
| Ir56d | 25.46162 | -1.70363 | 3.952413 | -0.43104 | 0.666442 | NA |
| tRNA:His-( | 0 | NA | NA | NA | NA | NA |
| CG16926 | 0 | NA | NA | NA | NA | NA |
| wbl | 0 | NA | NA | NA | NA | NA |
| tbrd-2 | 0 | NA | NA | NA | NA | NA |
| rib | 0.550877 | 0.424446 | 3.986474 | 0.106471 | 0.915208 | NA |
| AANATL6 | 0 | NA | NA | NA | NA | NA |
| AANATL4 | 0 | NA | NA | NA | NA | NA |
| fest | 1.098468 | -0.50358 | 3.892767 | -0.12936 | 0.89707 | NA |
| CG10051 | 0.712213 | -1.57119 | 3.998613 | -0.39293 | 0.694369 | NA |
| CG10062 | 0.357625 | -1.47707 | 4.044794 | -0.36518 | 0.714979 | NA |
| CG10073 | 0 | NA | NA | NA | NA | NA |
| tRNA:Ser-( | 0.178812 | -1.08808 | 4.061735 | -0.26789 | 0.788788 | NA |
| tRNA:Val-/ | 2.554805 | 0.923237 | 3.962137 | 0.233015 | 0.81575 | NA |
| tRNA:Val-/ | 45.00211 | -0.51174 | 3.846322 | -0.13305 | 0.894156 | NA |
| Arl6 | 0.132078 | -1.13211 | 4.069918 | -0.27817 | 0.780885 | NA |
| CG9416 | 0.178812 | -1.08808 | 4.061735 | -0.26789 | 0.788788 | NA |
| CG10474 | 0 | NA | NA | NA | NA | NA |
| AANATL5 | 0.357625 | -1.47707 | 4.044794 | -0.36518 | 0.714979 | NA |
| CG33453 | 0.178053 | -1.16467 | 4.075067 | -0.2858 | 0.775028 | NA |
| CG33454 | 0.178053 | -1.16467 | 4.075067 | -0.2858 | 0.775028 | NA |
| CG18605 | 0 | NA | NA | NA | NA | NA |
| CG15115 | 0.178053 | -1.16467 | 4.075067 | -0.2858 | 0.775028 | NA |
| Vha100-3 | 10.98544 | 1.52756 | 3.954161 | 0.386317 | 0.699262 | NA |
| tRNA:Trp-( | 0.4188 | 1.41068 | 4.034613 | 0.349644 | 0.726606 | NA |
| CG15071 | 21.81159 | -0.35488 | 3.840064 | -0.09242 | 0.926368 | NA |
| tRNA:Leu-/ | 31.23449 | 1.175693 | 3.952414 | 0.297462 | 0.766114 | NA |
| CG15082 | 1.884599 | 1.659029 | 3.975915 | 0.41727 | 0.676481 | NA |

| | | | | | | |
|---|---|---|---|---|---|---|
| DptA | 0 | NA | NA | NA | NA | NA |
| DptB | 0 | NA | NA | NA | NA | NA |
| Jheh1 | 0 | NA | NA | NA | NA | NA |
| CG18190 | 0 | NA | NA | NA | NA | NA |
| CG15096 | 55.1811 | -0.51522 | 3.948167 | -0.1305 | 0.896174 | NA |
| CG15093 | 33.5199 | 2.289047 | 3.780817 | 0.605437 | 0.544889 | NA |
| CG17821 | 0 | NA | NA | NA | NA | NA |
| CG18609 | 27.47424 | -0.30682 | 3.951014 | -0.07765 | 0.938103 | NA |
| CG33136 | 0 | NA | NA | NA | NA | NA |
| Gint3 | 0 | NA | NA | NA | NA | NA |
| tRNA:Gly-( | 0 | NA | NA | NA | NA | NA |
| BomS1 | 0 | NA | NA | NA | NA | NA |
| BomS4 | 0 | NA | NA | NA | NA | NA |
| BomS2 | 0 | NA | NA | NA | NA | NA |
| BomS3 | 0.132078 | -1.13211 | 4.069918 | -0.27817 | 0.780885 | NA |
| BomT2 | 0 | NA | NA | NA | NA | NA |
| BomS5 | 20.22782 | 0.357988 | 3.837242 | 0.093293 | 0.925671 | NA |
| Idgf5 | 0 | NA | NA | NA | NA | NA |
| GstE4 | 0 | NA | NA | NA | NA | NA |
| GstE5 | 0 | NA | NA | NA | NA | NA |
| GstE9 | 0 | NA | NA | NA | NA | NA |
| CG5189 | 0 | NA | NA | NA | NA | NA |
| CG30120 | 0 | NA | NA | NA | NA | NA |
| Pepck2 | 0.837599 | 1.568055 | 4.001801 | 0.391837 | 0.695178 | NA |
| Pepck1 | 0 | NA | NA | NA | NA | NA |
| CG5327 | 0 | NA | NA | NA | NA | NA |
| CG5493 | 0 | NA | NA | NA | NA | NA |
| CG17669 | 0.178053 | -1.16467 | 4.075067 | -0.2858 | 0.775028 | NA |
| GstE10 | 0.178053 | -1.16467 | 4.075067 | -0.2858 | 0.775028 | NA |
| BomS6 | 0 | NA | NA | NA | NA | NA |
| BomBc2 | 0 | NA | NA | NA | NA | NA |
| BomBc1 | 0 | NA | NA | NA | NA | NA |
| CG10914 | 11.51333 | 0.103462 | 3.87903 | 0.026672 | 0.978721 | NA |
| CG14500 | 0 | NA | NA | NA | NA | NA |
| CG30114 | 0 | NA | NA | NA | NA | NA |
| fj | 0 | NA | NA | NA | NA | NA |
| CG18536 | 5.232674 | 2.459345 | 3.928366 | 0.626048 | 0.531284 | NA |
| CG18537 | 25.46162 | -1.70363 | 3.952413 | -0.43104 | 0.666442 | NA |
| Tango8 | 0 | NA | NA | NA | NA | NA |
| CG5756 | 2.017234 | -1.62286 | 3.916351 | -0.41438 | 0.678596 | NA |
| CG30323 | 0 | NA | NA | NA | NA | NA |
| CG14490 | 0 | NA | NA | NA | NA | NA |
| CG14491 | 0 | NA | NA | NA | NA | NA |
| CG5084 | 0 | NA | NA | NA | NA | NA |
| CG10912 | 0 | NA | NA | NA | NA | NA |
| CG10911 | 0 | NA | NA | NA | NA | NA |
| Muc55B | 0 | NA | NA | NA | NA | NA |
| CG14495 | 12.92548 | 3.94513 | 3.895408 | 1.012764 | 0.311173 | NA |
| CG5767 | 45.58164 | -1.70567 | 3.95175 | -0.43162 | 0.666015 | NA |
| CG5770 | 0 | NA | NA | NA | NA | NA |

| | | | | | | |
|---|---|---|---|---|---|---|
| CG5773 | 60.31402 | -2.21535 | 3.952429 | -0.5605 | 0.575136 | NA |
| PpY-55A | 0 | NA | NA | NA | NA | NA |
| CG30110 | 18.12623 | 0.036085 | 3.932248 | 0.009177 | 0.992678 | NA |
| CG14480 | 0 | NA | NA | NA | NA | NA |
| POSH | 19.00059 | 0.514523 | 3.799228 | 0.135428 | 0.892273 | NA |
| ND-51L1 | 42.73355 | -2.9315 | 3.918653 | -0.74809 | 0.454407 | NA |
| tRNA:Phe- | 0 | NA | NA | NA | NA | NA |
| snoRNA:M | 0 | NA | NA | NA | NA | NA |
| snoRNA:M | 52.08906 | 0.959997 | 3.30004 | 0.290905 | 0.771124 | NA |
| snoRNA:M | 2.615581 | 1.492057 | 3.964088 | 0.376393 | 0.706624 | NA |
| snoRNA:U: | 1.163877 | -0.74508 | 3.943157 | -0.18896 | 0.850128 | NA |
| snoRNA:U: | 0.743406 | 0.099327 | 3.941247 | 0.025202 | 0.979894 | NA |
| snoRNA:U | 5.831972 | 2.276407 | 3.800113 | 0.599037 | 0.549148 | NA |
| snoRNA:U: | 17.74801 | 0.912382 | 3.752496 | 0.24314 | 0.807897 | NA |
| snoRNA:U: | 1.569349 | 1.463746 | 3.972245 | 0.368493 | 0.712505 | NA |
| snoRNA:M | 3.557267 | 2.696207 | 3.935151 | 0.68516 | 0.493243 | NA |
| snoRNA:sn | 0.732899 | 1.548046 | 4.007786 | 0.38626 | 0.699304 | NA |
| snoRNA:U: | 0 | NA | NA | NA | NA | NA |
| snoRNA:U: | 2.79447 | 1.814247 | 3.907967 | 0.464243 | 0.642473 | NA |
| snoRNA:U: | 0 | NA | NA | NA | NA | NA |
| snoRNA:U: | 1.583959 | 3.294289 | 3.954192 | 0.833113 | 0.404781 | NA |
| snoRNA:sn | 2.092465 | 1.481166 | 3.9672 | 0.373353 | 0.708886 | NA |
| snoRNA:M | 1.046233 | 1.431318 | 3.981858 | 0.35946 | 0.719251 | NA |
| CG10934 | 0 | NA | NA | NA | NA | NA |
| Dlish | 0 | NA | NA | NA | NA | NA |
| CG4975 | 12.77962 | 0.837322 | 3.952701 | 0.211835 | 0.832236 | NA |
| CG4984 | 36.72276 | 1.535867 | 3.951888 | 0.388641 | 0.697541 | NA |
| Ir54a | 0 | NA | NA | NA | NA | NA |
| sub | 0 | NA | NA | NA | NA | NA |
| Oxp | 16.24556 | -1.91474 | 3.953298 | -0.48434 | 0.628144 | NA |
| rhi | 0 | NA | NA | NA | NA | NA |
| tRNA:Phe- | 0 | NA | NA | NA | NA | NA |
| tRNA:Ala-T | 0 | NA | NA | NA | NA | NA |
| CG10939 | 0.460806 | -0.60052 | 3.997252 | -0.15023 | 0.880581 | NA |
| CG10764 | 28.20428 | 1.175216 | 3.952575 | 0.297329 | 0.766215 | NA |
| veil | 0 | NA | NA | NA | NA | NA |
| CG30103 | 0.356107 | -1.47099 | 4.037764 | -0.36431 | 0.715628 | NA |
| CG4847 | 0 | NA | NA | NA | NA | NA |
| insb | 0 | NA | NA | NA | NA | NA |
| Tes | 23.35951 | 5.14908 | 3.837351 | 1.341832 | 0.17965 | NA |
| mthl3 | 0 | NA | NA | NA | NA | NA |
| mthl4 | 0 | NA | NA | NA | NA | NA |
| Vajk4 | 0 | NA | NA | NA | NA | NA |
| tRNA:Ala-T | 0 | NA | NA | NA | NA | NA |
| SmydA-6 | 0 | NA | NA | NA | NA | NA |
| CG17717 | 0 | NA | NA | NA | NA | NA |
| APC7 | 0 | NA | NA | NA | NA | NA |
| CG14443 | 58.92661 | 6.528762 | 3.731039 | 1.749851 | 0.080144 | NA |
| Mcm6 | 50.98199 | -1.92041 | 3.951672 | -0.48598 | 0.626985 | NA |
| CG14439 | 33.71903 | 0.37291 | 3.845652 | 0.096969 | 0.922751 | NA |

| | | | | | | |
|---|---|---|---|---|---|---|
| Atx-1 | 0 | NA | NA | NA | NA | NA |
| mAChR-C | 0 | NA | NA | NA | NA | NA |
| tRNA:Phe- | 0 | NA | NA | NA | NA | NA |
| CR4575 | 0.178053 | -1.16467 | 4.075067 | -0.2858 | 0.775028 | NA |
| CR32733 | 0.574286 | -2.5433 | 4.014753 | -0.63349 | 0.526415 | NA |
| CG4586 | 38.84344 | 4.015504 | 3.891507 | 1.031864 | 0.302136 | NA |
| CG32736 | 0 | NA | NA | NA | NA | NA |
| nullo | 0.2094 | 1.005034 | 4.045341 | 0.248442 | 0.803792 | NA |
| CG4607 | 3.384419 | 1.447151 | 3.896022 | 0.371443 | 0.710307 | NA |
| lncRNA:CR | 0.233093 | 0.692656 | 4.076246 | 0.169925 | 0.865069 | NA |
| CG1958 | 57.55842 | -1.8975 | 3.951574 | -0.48019 | 0.631094 | NA |
| CG18155 | 25.20102 | 0.25905 | 3.767351 | 0.068762 | 0.945179 | NA |
| CG1999 | 0 | NA | NA | NA | NA | NA |
| CG32726 | 0.104623 | 0.639075 | 4.067674 | 0.157111 | 0.875158 | NA |
| CG15036 | 0 | NA | NA | NA | NA | NA |
| CG15035 | 0.132078 | -1.13211 | 4.069918 | -0.27817 | 0.780885 | NA |
| CG15034 | 0.236701 | -0.24276 | 4.020028 | -0.06039 | 0.951846 | NA |
| CG11369 | 0.178053 | -1.16467 | 4.075067 | -0.2858 | 0.775028 | NA |
| Ir7a | 0.357625 | -1.47707 | 4.044794 | -0.36518 | 0.714979 | NA |
| CG15478 | 0 | NA | NA | NA | NA | NA |
| CG12689 | 0 | NA | NA | NA | NA | NA |
| CG32720 | 0 | NA | NA | NA | NA | NA |
| CG32719 | 18.46342 | -1.84699 | 3.95291 | -0.46725 | 0.640322 | NA |
| CG32718 | 0 | NA | NA | NA | NA | NA |
| CG1409 | 0 | NA | NA | NA | NA | NA |
| Ir7c | 0 | NA | NA | NA | NA | NA |
| Ir7g | 0 | NA | NA | NA | NA | NA |
| CG15330 | 0 | NA | NA | NA | NA | NA |
| CG2256 | 24.63898 | 1.467003 | 3.845691 | 0.381467 | 0.702857 | NA |
| CG2233 | 0 | NA | NA | NA | NA | NA |
| CG15333 | 0.446101 | 0.990384 | 3.987592 | 0.248366 | 0.803851 | NA |
| CG15332 | 0.104623 | 0.639075 | 4.067674 | 0.157111 | 0.875158 | NA |
| hdm | 44.09142 | -2.00744 | 3.815864 | -0.52608 | 0.598834 | NA |
| CR33222 | 0 | NA | NA | NA | NA | NA |
| CR11386 | 0 | NA | NA | NA | NA | NA |
| ND-MNLL | 0 | NA | NA | NA | NA | NA |
| Nek2 | 0.53416 | -1.53481 | 4.012395 | -0.38252 | 0.702078 | NA |
| Ir7b | 0.1047 | 0.613261 | 4.062799 | 0.150945 | 0.880019 | NA |
| CG10920 | 0 | NA | NA | NA | NA | NA |
| CG2120 | 0 | NA | NA | NA | NA | NA |
| CG15336 | 0 | NA | NA | NA | NA | NA |
| CG10959 | 0.492912 | 0.154683 | 4.013127 | 0.038544 | 0.969254 | NA |
| CG15337 | 0 | NA | NA | NA | NA | NA |
| CG1632 | 6.812187 | 0.068221 | 3.942824 | 0.017303 | 0.986195 | NA |
| CG1387 | 0.178053 | -1.16467 | 4.075067 | -0.2858 | 0.775028 | NA |
| Or7a | 0 | NA | NA | NA | NA | NA |
| CG10761 | 0.356107 | -1.47099 | 4.037764 | -0.36431 | 0.715628 | NA |
| cyr | 0.178053 | -1.16467 | 4.075067 | -0.2858 | 0.775028 | NA |
| CG2260 | 0.466186 | 0.959471 | 4.033986 | 0.237847 | 0.812 | NA |
| Cp7Fb | 5.448478 | -1.34948 | 3.883391 | -0.3475 | 0.728215 | NA |

| | | | | | | |
|---|---|---|---|---|---|---|
| Cp7Fc | 0.104623 | 0.639075 | 4.067674 | 0.157111 | 0.875158 | NA |
| Cp36 | 0 | NA | NA | NA | NA | NA |
| Cp38 | 2.913353 | -2.39772 | 3.842118 | -0.62406 | 0.532587 | NA |
| CG12111 | 0 | NA | NA | NA | NA | NA |
| CG33223 | 0.1047 | 0.613261 | 4.062799 | 0.150945 | 0.880019 | NA |
| CG12116 | 33.53742 | -5.48574 | 3.818001 | -1.43681 | 0.150772 | NA |
| CG15347 | 0 | NA | NA | NA | NA | NA |
| CG32713 | 0 | NA | NA | NA | NA | NA |
| org-1 | 0.233093 | 0.692656 | 4.076246 | 0.169925 | 0.865069 | NA |
| CG32712 | 0.178053 | -1.16467 | 4.075067 | -0.2858 | 0.775028 | NA |
| CG10962 | 55.62189 | -7.1791 | 3.869574 | -1.85527 | 0.063558 | NA |
| CG12661 | 53.46411 | -3.63361 | 3.913525 | -0.92847 | 0.353162 | NA |
| l(1)G0020 | 39.71398 | -0.44893 | 3.854953 | -0.11645 | 0.907292 | NA |
| CG12057 | 35.00059 | -1.9192 | 3.952021 | -0.48563 | 0.627232 | NA |
| CG12106 | 43.71874 | -3.29433 | 3.773773 | -0.87295 | 0.382688 | NA |
| Obp8a | 2.936409 | 0.416631 | 3.869041 | 0.107683 | 0.914247 | NA |
| CG15368 | 21.97163 | -3.14951 | 3.91884 | -0.80369 | 0.421579 | NA |
| CG15369 | 1.325477 | 0.764512 | 3.893052 | 0.196379 | 0.844314 | NA |
| CG15370 | 0 | NA | NA | NA | NA | NA |
| Gr8a | 4.094408 | -1.89106 | 3.960175 | -0.47752 | 0.632993 | NA |
| lz | 0.178053 | -1.16467 | 4.075067 | -0.2858 | 0.775028 | NA |
| CG17440 | 21.69568 | -1.35849 | 3.783387 | -0.35907 | 0.719545 | NA |
| Erk7 | 0.178053 | -1.16467 | 4.075067 | -0.2858 | 0.775028 | NA |
| CR17438 | 0 | NA | NA | NA | NA | NA |
| Ir8a | 30.63042 | -3.25914 | 3.928093 | -0.8297 | 0.406708 | NA |
| SmydA-9 | 0.178053 | -1.16467 | 4.075067 | -0.2858 | 0.775028 | NA |
| CG12115 | 0 | NA | NA | NA | NA | NA |
| CG10970 | 0.396233 | -1.6814 | 4.028351 | -0.41739 | 0.676392 | NA |
| Cubn | 39.5728 | -0.52138 | 3.772338 | -0.13821 | 0.890074 | NA |
| btd | 0.387453 | 0.098632 | 4.002797 | 0.024641 | 0.980342 | NA |
| Sp1 | 1.886966 | -2.1758 | 3.893319 | -0.55886 | 0.576261 | NA |
| CG3106 | 0 | NA | NA | NA | NA | NA |
| CG1889 | 7.217562 | -4.80075 | 3.84009 | -1.25017 | 0.211239 | NA |
| CG32694 | 34.84252 | 0.118846 | 3.951416 | 0.030077 | 0.976006 | NA |
| CG2990 | 3.455098 | 1.699144 | 3.965156 | 0.428519 | 0.668273 | NA |
| Yp1 | 0 | NA | NA | NA | NA | NA |
| Gr9a | 0 | NA | NA | NA | NA | NA |
| Yp2 | 0.209323 | 1.414133 | 4.032409 | 0.350692 | 0.725819 | NA |
| CG15252 | 0 | NA | NA | NA | NA | NA |
| CG1791 | 0 | NA | NA | NA | NA | NA |
| CG12645 | 0 | NA | NA | NA | NA | NA |
| lncRNA:CR | 1.584932 | -1.84571 | 3.973792 | -0.46447 | 0.642311 | NA |
| CG2909 | 52.7636 | 4.200996 | 3.814257 | 1.101393 | 0.270726 | NA |
| CG2898 | 1.06832 | -1.61145 | 3.983918 | -0.40449 | 0.685853 | NA |
| RabX2 | 0 | NA | NA | NA | NA | NA |
| CG2887 | 0.396233 | -1.6814 | 4.028351 | -0.41739 | 0.676392 | NA |
| Or9a | 0 | NA | NA | NA | NA | NA |
| CG12640 | 0 | NA | NA | NA | NA | NA |
| Psf3 | 17.23342 | -0.5553 | 3.750467 | -0.14806 | 0.882294 | NA |
| CG32681 | 27.46298 | -0.87275 | 3.949352 | -0.22098 | 0.825104 | NA |

| | | | | | | |
|---|---|---|---|---|---|---|
| CG32679 | 0.178812 | -1.08808 | 4.061735 | -0.26789 | 0.788788 | NA |
| CG15306 | 0.233093 | 0.692656 | 4.076246 | 0.169925 | 0.865069 | NA |
| Ipod | 0 | NA | NA | NA | NA | NA |
| CG15308 | 0 | NA | NA | NA | NA | NA |
| CG2962 | 0 | NA | NA | NA | NA | NA |
| Rab9D | 0 | NA | NA | NA | NA | NA |
| CG2111 | 0.283436 | -0.20033 | 4.070792 | -0.04921 | 0.960751 | NA |
| CG12637 | 0 | NA | NA | NA | NA | NA |
| Rab9E | 36.82873 | 1.17637 | 3.952186 | 0.297651 | 0.76597 | NA |
| CG9806 | 0 | NA | NA | NA | NA | NA |
| Rab9Db | 0 | NA | NA | NA | NA | NA |
| CG1628 | 8.07681 | -5.36937 | 3.853869 | -1.39324 | 0.163546 | NA |
| CG15225 | 0 | NA | NA | NA | NA | NA |
| CAH14 | 0 | NA | NA | NA | NA | NA |
| Ipk1 | 31.1851 | 0.646294 | 3.735272 | 0.173025 | 0.862632 | NA |
| Cib2 | 61.87766 | 1.751356 | 3.951748 | 0.443185 | 0.657632 | NA |
| CAH15 | 0 | NA | NA | NA | NA | NA |
| Cht8 | 0.233093 | 0.692656 | 4.076246 | 0.169925 | 0.865069 | NA |
| Cht9 | 0 | NA | NA | NA | NA | NA |
| Cht4 | 0 | NA | NA | NA | NA | NA |
| Cht12 | 0 | NA | NA | NA | NA | NA |
| dgt3 | 0.104623 | 0.639075 | 4.067674 | 0.157111 | 0.875158 | NA |
| CG3216 | 58.08918 | -5.98332 | 3.575188 | -1.67357 | 0.094216 | NA |
| CG15650 | 7.857823 | 2.71295 | 3.907078 | 0.694368 | 0.487451 | NA |
| Dso1 | 34.04859 | -1.8805 | 3.95202 | -0.47583 | 0.634194 | NA |
| CG9993 | 0 | NA | NA | NA | NA | NA |
| CG17999 | 0.178053 | -1.16467 | 4.075067 | -0.2858 | 0.775028 | NA |
| CG9406 | 0 | NA | NA | NA | NA | NA |
| tRNA:Trp-( | 0 | NA | NA | NA | NA | NA |
| tRNA:Gly-( | 0.565353 | 0.142845 | 3.959283 | 0.036079 | 0.97122 | NA |
| BBS9 | 0.52327 | 2.335758 | 3.994212 | 0.584786 | 0.558692 | NA |
| CG17974 | 0 | NA | NA | NA | NA | NA |
| CG9822 | 0 | NA | NA | NA | NA | NA |
| CG9752 | 9.172321 | 3.702066 | 3.8959 | 0.950247 | 0.341987 | NA |
| CG10505 | 4.881497 | 1.238411 | 3.839789 | 0.32252 | 0.747058 | NA |
| CG30393 | 12.28568 | -1.69873 | 3.954016 | -0.42962 | 0.667472 | NA |
| CG30391 | 0.712213 | -1.57119 | 3.998613 | -0.39293 | 0.694369 | NA |
| tRNA:Trp-( | 0 | NA | NA | NA | NA | NA |
| CG4286 | 0 | NA | NA | NA | NA | NA |
| tRNA:Gly-( | 0.337793 | 1.451235 | 4.039844 | 0.35923 | 0.719423 | NA |
| tRNA:Trp-( | 0 | NA | NA | NA | NA | NA |
| Egfr | 0.283512 | -0.2331 | 4.065094 | -0.05734 | 0.954273 | NA |
| CG30222 | 0.282753 | -0.25398 | 4.020204 | -0.06318 | 0.949626 | NA |
| CG33225 | 0 | NA | NA | NA | NA | NA |
| CG15674 | 19.89299 | 1.744326 | 3.953505 | 0.44121 | 0.659061 | NA |
| CG15676 | 0 | NA | NA | NA | NA | NA |
| Ppcdc | 0 | NA | NA | NA | NA | NA |
| Tim10 | 0.356107 | -1.47099 | 4.037764 | -0.36431 | 0.715628 | NA |
| PIG-M | 0 | NA | NA | NA | NA | NA |
| CG30284 | 10.77345 | -5.98626 | 3.923127 | -1.52589 | 0.127037 | NA |

| | | | | | | |
|---|---|---|---|---|---|---|
| CG30283 | 0 | NA | NA | NA | NA | NA |
| CG33226 | 0 | NA | NA | NA | NA | NA |
| CG30287 | 0.178053 | -1.16467 | 4.075067 | -0.2858 | 0.775028 | NA |
| CG30286 | 0 | NA | NA | NA | NA | NA |
| CG30289 | 0 | NA | NA | NA | NA | NA |
| CG30288 | 0 | NA | NA | NA | NA | NA |
| CG10494 | 0.411906 | -0.22397 | 4.020275 | -0.05571 | 0.955572 | NA |
| CG30398 | 0 | NA | NA | NA | NA | NA |
| Grx1t | 0 | NA | NA | NA | NA | NA |
| CG30395 | 0 | NA | NA | NA | NA | NA |
| CG4021 | 0.62774 | 1.374224 | 3.999506 | 0.343599 | 0.731148 | NA |
| lncRNA:CR | 0 | NA | NA | NA | NA | NA |
| CG13488 | 0 | NA | NA | NA | NA | NA |
| Gr58c | 0 | NA | NA | NA | NA | NA |
| Gr58a | 0 | NA | NA | NA | NA | NA |
| dany | 0 | NA | NA | NA | NA | NA |
| Gr58b | 0 | NA | NA | NA | NA | NA |
| CG13494 | 0.356866 | -1.92326 | 4.037889 | -0.4763 | 0.633858 | NA |
| PpN58A | 28.70198 | -2.02706 | 3.814139 | -0.53146 | 0.595101 | NA |
| comr | 0.178053 | -1.16467 | 4.075067 | -0.2858 | 0.775028 | NA |
| tRNA:Gly-T | 0 | NA | NA | NA | NA | NA |
| tRNA:Gly-T | 0.233093 | 0.692656 | 4.076246 | 0.169925 | 0.865069 | NA |
| CG4372 | 0 | NA | NA | NA | NA | NA |
| CG4377 | 0 | NA | NA | NA | NA | NA |
| CG4363 | 0 | NA | NA | NA | NA | NA |
| CG13492 | 0.396233 | -1.6814 | 4.028351 | -0.41739 | 0.676392 | NA |
| tpr | 0 | NA | NA | NA | NA | NA |
| CG18735 | 0 | NA | NA | NA | NA | NA |
| Loxl2 | 0.233093 | 0.692656 | 4.076246 | 0.169925 | 0.865069 | NA |
| CG9308 | 7.478238 | -1.69271 | 3.955987 | -0.42789 | 0.668735 | NA |
| CG30281 | 0 | NA | NA | NA | NA | NA |
| mei-S332 | 0 | NA | NA | NA | NA | NA |
| NtR | 0 | NA | NA | NA | NA | NA |
| CG11269 | 0.932373 | 1.053996 | 3.996193 | 0.26375 | 0.791972 | NA |
| gom | 34.4264 | 0.645301 | 3.731369 | 0.17294 | 0.862699 | NA |
| CG11291 | 0 | NA | NA | NA | NA | NA |
| Oatp58Db | 0 | NA | NA | NA | NA | NA |
| Oatp58Da | 0 | NA | NA | NA | NA | NA |
| CG30278 | 0 | NA | NA | NA | NA | NA |
| Alp8 | 0.209247 | 1.042804 | 4.047837 | 0.25762 | 0.7967 | NA |
| Alp7 | 0 | NA | NA | NA | NA | NA |
| Swim | 0.948342 | -0.9323 | 3.989063 | -0.23371 | 0.815207 | NA |
| CG13502 | 0.132078 | -1.13211 | 4.069918 | -0.27817 | 0.780885 | NA |
| CG11475 | 0 | NA | NA | NA | NA | NA |
| CG13501 | 3.769197 | 1.70343 | 3.964029 | 0.429722 | 0.667398 | NA |
| PpD5 | 0 | NA | NA | NA | NA | NA |
| CG6967 | 5.431375 | 0.663113 | 3.848438 | 0.172307 | 0.863196 | NA |
| Amy-d | 0 | NA | NA | NA | NA | NA |
| Cda9 | 20.20709 | 1.744484 | 3.953465 | 0.441255 | 0.659029 | NA |
| CG11400 | 0.894061 | -2.00006 | 4.030294 | -0.49626 | 0.619714 | NA |

| | | | | | | |
|---|---|---|---|---|---|---|
| Gbp2 | 21.07336 | 0.92445 | 3.737275 | 0.247359 | 0.80463 | NA |
| tRNA:Leu-/ | 0 | NA | NA | NA | NA | NA |
| CG17290 | 0 | NA | NA | NA | NA | NA |
| CG10950 | 0.890266 | -1.59482 | 3.989917 | -0.39971 | 0.689368 | NA |
| CG12699 | 37.23569 | 1.245588 | 3.873662 | 0.321553 | 0.747791 | NA |
| CG18469 | 0 | NA | NA | NA | NA | NA |
| CG10953 | 0 | NA | NA | NA | NA | NA |
| CG30457 | 0 | NA | NA | NA | NA | NA |
| CG30458 | 0 | NA | NA | NA | NA | NA |
| CG17287 | 0 | NA | NA | NA | NA | NA |
| tRNA:Leu-/ | 0 | NA | NA | NA | NA | NA |
| CG15605 | 0 | NA | NA | NA | NA | NA |
| Spn53F | 0 | NA | NA | NA | NA | NA |
| Amy-p | 0 | NA | NA | NA | NA | NA |
| CG8950 | 9.343238 | -5.35141 | 3.891488 | -1.37516 | 0.169082 | NA |
| Arp53D | 0 | NA | NA | NA | NA | NA |
| resilin | 0.178053 | -1.16467 | 4.075067 | -0.2858 | 0.775028 | NA |
| CG15919 | 0 | NA | NA | NA | NA | NA |
| CG5550 | 0 | NA | NA | NA | NA | NA |
| CG6426 | 47.60704 | 2.878717 | 3.919865 | 0.734392 | 0.46271 | NA |
| CG6421 | 0.132078 | -1.13211 | 4.069918 | -0.27817 | 0.780885 | NA |
| CG6429 | 0 | NA | NA | NA | NA | NA |
| inaC | 0.765088 | -0.45272 | 3.976873 | -0.11384 | 0.909366 | NA |
| CG15614 | 1.193706 | 0.462543 | 3.970016 | 0.116509 | 0.907249 | NA |
| tRNA:Gly-( | 0.178053 | -1.16467 | 4.075067 | -0.2858 | 0.775028 | NA |
| CG9010 | 0.178053 | -1.16467 | 4.075067 | -0.2858 | 0.775028 | NA |
| Fen1 | 15.76455 | -2.21017 | 3.871568 | -0.57087 | 0.568087 | NA |
| Ugt37C1 | 0 | NA | NA | NA | NA | NA |
| CG15615 | 0.178812 | -1.08808 | 4.061735 | -0.26789 | 0.788788 | NA |
| Acp53Ea | 0 | NA | NA | NA | NA | NA |
| Acp53C14l | 0 | NA | NA | NA | NA | NA |
| Acp53C14a | 0 | NA | NA | NA | NA | NA |
| CG15617 | 0 | NA | NA | NA | NA | NA |
| CG33458 | 0 | NA | NA | NA | NA | NA |
| S-Lap8 | 0.178053 | -1.16467 | 4.075067 | -0.2858 | 0.775028 | NA |
| CG4945 | 0.71449 | -2.47883 | 4.014121 | -0.61753 | 0.536887 | NA |
| CG5089 | 0.178053 | -1.16467 | 4.075067 | -0.2858 | 0.775028 | NA |
| Amyrel | 0 | NA | NA | NA | NA | NA |
| CG8303 | 0.53416 | -1.53481 | 4.012395 | -0.38252 | 0.702078 | NA |
| Lst | 9.556822 | 1.165775 | 3.955791 | 0.294701 | 0.768222 | NA |
| CG15712 | 0 | NA | NA | NA | NA | NA |
| tRNA:Glu-( | 0.715249 | -1.88064 | 4.034336 | -0.46616 | 0.641102 | NA |
| tRNA:Glu-' | 30.84723 | -3.37581 | 3.869757 | -0.87236 | 0.383013 | NA |
| Tsf3 | 5.749738 | -0.81538 | 3.955041 | -0.20616 | 0.836664 | NA |
| CG15701 | 5.758496 | 1.720238 | 3.959653 | 0.434441 | 0.663968 | NA |
| CG3687 | 0 | NA | NA | NA | NA | NA |
| CG15705 | 0 | NA | NA | NA | NA | NA |
| CG9068 | 0.209247 | 1.042804 | 4.047837 | 0.25762 | 0.7967 | NA |
| CG30100 | 0 | NA | NA | NA | NA | NA |
| Parp16 | 6.86804 | -1.90364 | 3.956501 | -0.48114 | 0.630416 | NA |

| | | | | | | |
|---|---|---|---|---|---|---|
| CG7848 | 0.104623 | 0.639075 | 4.067674 | 0.157111 | 0.875158 | NA |
| CG15711 | 0 | NA | NA | NA | NA | NA |
| CG15710 | 0 | NA | NA | NA | NA | NA |
| CG30324 | 0.178053 | -1.16467 | 4.075067 | -0.2858 | 0.775028 | NA |
| CG7755 | 0.178053 | -1.16467 | 4.075067 | -0.2858 | 0.775028 | NA |
| cato | 0 | NA | NA | NA | NA | NA |
| CG15708 | 0.1047 | 0.613261 | 4.062799 | 0.150945 | 0.880019 | NA |
| krimp | 0 | NA | NA | NA | NA | NA |
| CG7786 | 0 | NA | NA | NA | NA | NA |
| CG7798 | 0 | NA | NA | NA | NA | NA |
| tRNA:Glu-[illegible] | 0 | NA | NA | NA | NA | NA |
| tRNA:Gln-[illegible] | 20.03851 | 0.257894 | 3.836584 | 0.06722 | 0.946407 | NA |
| COX6AL | 0 | NA | NA | NA | NA | NA |
| CG8399 | 2.973451 | 2.154611 | 3.93095 | 0.548115 | 0.583613 | NA |
| CG8401 | 2.272052 | 1.167201 | 3.965566 | 0.294334 | 0.768503 | NA |
| Jhedup | 0 | NA | NA | NA | NA | NA |
| Jhe | 22.74358 | 3.185144 | 3.910177 | 0.814578 | 0.415314 | NA |
| CG30095 | 0.132078 | -1.13211 | 4.069918 | -0.27817 | 0.780885 | NA |
| CysRS | 46.74145 | 7.833318 | 3.907689 | 2.004591 | 0.045007 | NA |
| Ptp52F | 8.188817 | -1.9067 | 3.955613 | -0.48202 | 0.629788 | NA |
| CG10734 | 0.596853 | 0.454699 | 3.988272 | 0.114009 | 0.909231 | NA |
| CG30091 | 0 | NA | NA | NA | NA | NA |
| CG30090 | 0 | NA | NA | NA | NA | NA |
| CG30088 | 0 | NA | NA | NA | NA | NA |
| CG12970 | 0 | NA | NA | NA | NA | NA |
| CG8299 | 0 | NA | NA | NA | NA | NA |
| CG8249 | 0.178053 | -1.16467 | 4.075067 | -0.2858 | 0.775028 | NA |
| Poxn | 0 | NA | NA | NA | NA | NA |
| CG30083 | 0 | NA | NA | NA | NA | NA |
| CG30080 | 0 | NA | NA | NA | NA | NA |
| Ir52a | 0 | NA | NA | NA | NA | NA |
| Ir52b | 0 | NA | NA | NA | NA | NA |
| Ir52c | 0.178053 | -1.16467 | 4.075067 | -0.2858 | 0.775028 | NA |
| Ir52d | 0.132078 | -1.13211 | 4.069918 | -0.27817 | 0.780885 | NA |
| CG30471 | 0 | NA | NA | NA | NA | NA |
| CG12963 | 0.178053 | -1.16467 | 4.075067 | -0.2858 | 0.775028 | NA |
| fus | 11.60363 | -3.07829 | 3.821689 | -0.80548 | 0.420543 | NA |
| CG8192 | 0 | NA | NA | NA | NA | NA |
| CG30472 | 0 | NA | NA | NA | NA | NA |
| Ir51b | 0 | NA | NA | NA | NA | NA |
| ItgaPS4 | 0.356107 | -1.47099 | 4.037764 | -0.36431 | 0.715628 | NA |
| Hr51 | 0 | NA | NA | NA | NA | NA |
| Mtk | 0 | NA | NA | NA | NA | NA |
| CG8160 | 0.104623 | 0.639075 | 4.067674 | 0.157111 | 0.875158 | NA |
| CG8157 | 41.57385 | 7.063577 | 3.888074 | 1.816729 | 0.069259 | NA |
| ND-51L2 | 0 | NA | NA | NA | NA | NA |
| CG8093 | 0 | NA | NA | NA | NA | NA |
| Cyp317a1 | 0.233093 | 0.692656 | 4.076246 | 0.169925 | 0.865069 | NA |
| CG12853 | 0 | NA | NA | NA | NA | NA |
| CG7544 | 36.17436 | -2.1749 | 3.953361 | -0.55014 | 0.582223 | NA |

| | | | | | | |
|---|---|---|---|---|---|---|
| tRNA:Leu-/ | 43.91365 | -0.28895 | 3.835786 | -0.07533 | 0.939952 | NA |
| CG12866 | 0 | NA | NA | NA | NA | NA |
| Dro | 0 | NA | NA | NA | NA | NA |
| AttA | 0 | NA | NA | NA | NA | NA |
| AttB | 0.732899 | 1.548046 | 4.007786 | 0.38626 | 0.699304 | NA |
| CG10205 | 0 | NA | NA | NA | NA | NA |
| Cyp6a19 | 0 | NA | NA | NA | NA | NA |
| Cyp6a8 | 0 | NA | NA | NA | NA | NA |
| Cyp6a21 | 57.01561 | 1.559025 | 3.918779 | 0.397835 | 0.690752 | NA |
| Pgm2b | 27.73631 | -1.91819 | 3.952312 | -0.48533 | 0.62744 | NA |
| CG12857 | 60.43855 | -2.27598 | 3.952504 | -0.57583 | 0.564728 | NA |
| CG30480 | 0.46073 | -0.17853 | 4.00793 | -0.04454 | 0.964471 | NA |
| CG30479 | 0 | NA | NA | NA | NA | NA |
| CG12860 | 0 | NA | NA | NA | NA | NA |
| CG12861 | 0 | NA | NA | NA | NA | NA |
| Adgf-E | 0 | NA | NA | NA | NA | NA |
| CG12862 | 0 | NA | NA | NA | NA | NA |
| CG10104 | 0 | NA | NA | NA | NA | NA |
| Achl | 0.356866 | -1.92326 | 4.037889 | -0.4763 | 0.633858 | NA |
| CG12869 | 0.965993 | 2.477533 | 4.003194 | 0.618889 | 0.53599 | NA |
| Had2 | 0.178053 | -1.16467 | 4.075067 | -0.2858 | 0.775028 | NA |
| CG33468 | 0 | NA | NA | NA | NA | NA |
| CG30076 | 1.015798 | 0.083253 | 3.968257 | 0.02098 | 0.983262 | NA |
| CG18324 | 0.732899 | 1.548046 | 4.007786 | 0.38626 | 0.699304 | NA |
| CG8468 | 16.86233 | 3.834619 | 3.807442 | 1.007138 | 0.313868 | NA |
| SmydA-1 | 0 | NA | NA | NA | NA | NA |
| Arc2 | 4.988115 | 1.214777 | 3.880852 | 0.313018 | 0.754267 | NA |
| Obp50b | 0 | NA | NA | NA | NA | NA |
| Obp50d | 0 | NA | NA | NA | NA | NA |
| CG30075 | 0 | NA | NA | NA | NA | NA |
| Obp50e | 0.104623 | 0.639075 | 4.067674 | 0.157111 | 0.875158 | NA |
| Obp50a | 0 | NA | NA | NA | NA | NA |
| CR10102 | 0 | NA | NA | NA | NA | NA |
| tej | 0.1047 | 0.613261 | 4.062799 | 0.150945 | 0.880019 | NA |
| Sox15 | 1.675199 | 1.648777 | 3.978727 | 0.414398 | 0.678583 | NA |
| CG18568 | 0 | NA | NA | NA | NA | NA |
| St4 | 0 | NA | NA | NA | NA | NA |
| DJ-1alpha | 26.53136 | -2.42901 | 3.746103 | -0.64841 | 0.51672 | NA |
| S-Lap5 | 0 | NA | NA | NA | NA | NA |
| CG6280 | 0.178053 | -1.16467 | 4.075067 | -0.2858 | 0.775028 | NA |
| Cpr50Ca | 0.178053 | -1.16467 | 4.075067 | -0.2858 | 0.775028 | NA |
| Cpr50Cb | 0.178053 | -1.16467 | 4.075067 | -0.2858 | 0.775028 | NA |
| CG30484 | 0.356107 | -1.47099 | 4.037764 | -0.36431 | 0.715628 | NA |
| CG30485 | 0.2094 | 1.005034 | 4.045341 | 0.248442 | 0.803792 | NA |
| CG6553 | 0 | NA | NA | NA | NA | NA |
| CG6337 | 20.7665 | -0.80729 | 3.937839 | -0.20501 | 0.837566 | NA |
| S-Lap7 | 0 | NA | NA | NA | NA | NA |
| CG6220 | 0.1047 | 0.613261 | 4.062799 | 0.150945 | 0.880019 | NA |
| CG6209 | 0.132078 | -1.13211 | 4.069918 | -0.27817 | 0.780885 | NA |
| CG12464 | 0 | NA | NA | NA | NA | NA |

| | | | | | | |
|---|---|---|---|---|---|---|
| AttC | 33.50398 | 1.748507 | 3.952456 | 0.442385 | 0.658211 | NA |
| CG33470 | 0 | NA | NA | NA | NA | NA |
| tRNA:Ile-A | 0 | NA | NA | NA | NA | NA |
| tRNA:Ile-A | 0 | NA | NA | NA | NA | NA |
| tRNA:Ile-A | 0 | NA | NA | NA | NA | NA |
| tRNA:Ile-A | 0 | NA | NA | NA | NA | NA |
| CG30065 | 0.178053 | -1.16467 | 4.075067 | -0.2858 | 0.775028 | NA |
| Ser8 | 0 | NA | NA | NA | NA | NA |
| tRNA:Ile-A | 0 | NA | NA | NA | NA | NA |
| CG13330 | 0 | NA | NA | NA | NA | NA |
| CG13331 | 33.89143 | 1.691092 | 3.95237 | 0.427868 | 0.668747 | NA |
| CG13332 | 52.98589 | -1.39981 | 3.951506 | -0.35425 | 0.723153 | NA |
| link | 19.01268 | 6.056002 | 3.848483 | 1.573608 | 0.115578 | NA |
| CG13334 | 0 | NA | NA | NA | NA | NA |
| tRNA:Lys-C | 0 | NA | NA | NA | NA | NA |
| cbc | 6.033976 | -1.15586 | 3.930565 | -0.29407 | 0.768705 | NA |
| CG30062 | 5.549096 | 1.719007 | 3.959971 | 0.434096 | 0.664219 | NA |
| tRNA:Ile-A | 0 | NA | NA | NA | NA | NA |
| CG30059 | 0.712213 | -1.57119 | 3.998613 | -0.39293 | 0.694369 | NA |
| CG18278 | 9.25877 | -1.69565 | 3.955021 | -0.42873 | 0.668118 | NA |
| CG10814 | 0.3141 | 1.240997 | 4.03841 | 0.307299 | 0.758616 | NA |
| GstE14 | 0 | NA | NA | NA | NA | NA |
| CG17047 | 0 | NA | NA | NA | NA | NA |
| CG4734 | 0 | NA | NA | NA | NA | NA |
| CG4714 | 49.76334 | -3.38268 | 3.921748 | -0.86254 | 0.388388 | NA |
| CG4712 | 16.2704 | -4.33197 | 3.90497 | -1.10935 | 0.26728 | NA |
| CG30058 | 0 | NA | NA | NA | NA | NA |
| Orc3 | 0 | NA | NA | NA | NA | NA |
| CG15870 | 11.21736 | -1.69782 | 3.954309 | -0.42936 | 0.667662 | NA |
| CG13326 | 0 | NA | NA | NA | NA | NA |
| Or49b | 0.104623 | 0.639075 | 4.067674 | 0.157111 | 0.875158 | NA |
| vg | 0.697584 | -1.11707 | 3.977017 | -0.28088 | 0.778801 | NA |
| sug | 0 | NA | NA | NA | NA | NA |
| Sans | 0 | NA | NA | NA | NA | NA |
| CG13321 | 0.178053 | -1.16467 | 4.075067 | -0.2858 | 0.775028 | NA |
| CG13324 | 0 | NA | NA | NA | NA | NA |
| CG13323 | 0 | NA | NA | NA | NA | NA |
| CG30487 | 0 | NA | NA | NA | NA | NA |
| Lfg | 0 | NA | NA | NA | NA | NA |
| Balat | 0 | NA | NA | NA | NA | NA |
| CG30486 | 0 | NA | NA | NA | NA | NA |
| Cyp301a1 | 4.164135 | -1.88038 | 3.815942 | -0.49277 | 0.622176 | NA |
| stil | 0 | NA | NA | NA | NA | NA |
| CG8646 | 55.04438 | -8.22974 | 3.875036 | -2.12379 | 0.033688 | NA |
| wuc | 0 | NA | NA | NA | NA | NA |
| CG12374 | 0.178053 | -1.16467 | 4.075067 | -0.2858 | 0.775028 | NA |
| CG17580 | 0.1047 | 0.613261 | 4.062799 | 0.150945 | 0.880019 | NA |
| Mos | 0.3141 | 1.240997 | 4.03841 | 0.307299 | 0.758616 | NA |
| Obp49a | 0 | NA | NA | NA | NA | NA |
| CG30053 | 0.104623 | 0.639075 | 4.067674 | 0.157111 | 0.875158 | NA |

| | | | | | | |
|---|---|---|---|---|---|---|
| CG8785 | 7.984572 | 0.691827 | 3.929781 | 0.176047 | 0.860257 | NA |
| CG30203 | 33.34724 | 1.501157 | 3.952192 | 0.379829 | 0.704072 | NA |
| CG30046 | 0.178812 | -1.08808 | 4.061735 | -0.26789 | 0.788788 | NA |
| tRNA:His-G | 0 | NA | NA | NA | NA | NA |
| tRNA:His-G | 0 | NA | NA | NA | NA | NA |
| CG30047 | 0.2094 | 1.005034 | 4.045341 | 0.248442 | 0.803792 | NA |
| CG30049 | 0.712972 | -2.55723 | 4.00634 | -0.63829 | 0.523282 | NA |
| CG30043 | 0 | NA | NA | NA | NA | NA |
| CR33013 | 1.099666 | -0.72019 | 3.971513 | -0.18134 | 0.856102 | NA |
| CG33012 | 0.356107 | -1.47099 | 4.037764 | -0.36431 | 0.715628 | NA |
| CG13160 | 0 | NA | NA | NA | NA | NA |
| CG13159 | 0 | NA | NA | NA | NA | NA |
| Cpr49Aa | 1.869955 | -1.27799 | 3.894534 | -0.32815 | 0.742799 | NA |
| Cpr49Ab | 0 | NA | NA | NA | NA | NA |
| CG8501 | 0 | NA | NA | NA | NA | NA |
| Cpr49Ac | 0 | NA | NA | NA | NA | NA |
| Or49a | 0.356107 | -1.47099 | 4.037764 | -0.36431 | 0.715628 | NA |
| Cpr49Ae | 60.83066 | 1.751298 | 3.951762 | 0.443169 | 0.657644 | NA |
| Cpr49Af | 0 | NA | NA | NA | NA | NA |
| Cpr49Ag | 0.2094 | 1.005034 | 4.045341 | 0.248442 | 0.803792 | NA |
| CG30050 | 0.178053 | -1.16467 | 4.075067 | -0.2858 | 0.775028 | NA |
| Cpr49Ah | 0 | NA | NA | NA | NA | NA |
| CG13157 | 0.178053 | -1.16467 | 4.075067 | -0.2858 | 0.775028 | NA |
| Dera | 58.41285 | -3.42598 | 3.843202 | -0.89144 | 0.372694 | NA |
| s-cup | 27.96254 | -0.38576 | 3.940911 | -0.09789 | 0.922023 | NA |
| vis | 47.73975 | -9.05778 | 3.907933 | -2.31779 | 0.020461 | NA |
| dgt5 | 0.209247 | 1.042804 | 4.047837 | 0.25762 | 0.7967 | NA |
| CG30334 | 6.42765 | 1.758232 | 3.853888 | 0.456223 | 0.64823 | NA |
| CG13154 | 0.178053 | -1.16467 | 4.075067 | -0.2858 | 0.775028 | NA |
| CG8834 | 0 | NA | NA | NA | NA | NA |
| Cpr49Ad | 0 | NA | NA | NA | NA | NA |
| CG13155 | 0.178053 | -1.16467 | 4.075067 | -0.2858 | 0.775028 | NA |
| tRNA:His-G | 1.246373 | -1.6238 | 3.979526 | -0.40804 | 0.683245 | NA |
| tRNA:His-G | 0 | NA | NA | NA | NA | NA |
| tRNA:His-G | 0 | NA | NA | NA | NA | NA |
| CG8850 | 0 | NA | NA | NA | NA | NA |
| CG13186 | 0.356107 | -1.47099 | 4.037764 | -0.36431 | 0.715628 | NA |
| CG13171 | 0 | NA | NA | NA | NA | NA |
| CG13170 | 0 | NA | NA | NA | NA | NA |
| CG13168 | 0 | NA | NA | NA | NA | NA |
| Vha36-2 | 0 | NA | NA | NA | NA | NA |
| CG8854 | 0 | NA | NA | NA | NA | NA |
| Sr-CII | 0 | NA | NA | NA | NA | NA |
| Cyp6g2 | 0 | NA | NA | NA | NA | NA |
| OSCP1 | 0 | NA | NA | NA | NA | NA |
| SkpB | 0 | NA | NA | NA | NA | NA |
| CG8888 | 0 | NA | NA | NA | NA | NA |
| CG30039 | 0.1047 | 0.613261 | 4.062799 | 0.150945 | 0.880019 | NA |
| CG13193 | 0 | NA | NA | NA | NA | NA |
| snoRNA:M | 5.78152 | -0.74901 | 3.771122 | -0.19862 | 0.842562 | NA |

| | | | | | | |
|---|---|---|---|---|---|---|
| exp | 9.109486 | -1.8306 | 3.811139 | -0.48033 | 0.630994 | NA |
| RnrS | 0.178053 | -1.16467 | 4.075067 | -0.2858 | 0.775028 | NA |
| CG30037 | 0 | NA | NA | NA | NA | NA |
| CG30036 | 0 | NA | NA | NA | NA | NA |
| reb | 0.178053 | -1.16467 | 4.075067 | -0.2858 | 0.775028 | NA |
| Damm | 4.871543 | -2.71574 | 3.859855 | -0.70359 | 0.48169 | NA |
| Twdlbeta | 0 | NA | NA | NA | NA | NA |
| Sobp | 7.407411 | -1.45501 | 3.891103 | -0.37393 | 0.708454 | NA |
| Ir48c | 15.44553 | -2.96766 | 3.841902 | -0.77244 | 0.439851 | NA |
| tRNA:Met- | 0 | NA | NA | NA | NA | NA |
| CG13198 | 4.842162 | -5.02909 | 3.937067 | -1.27737 | 0.201472 | NA |
| Tret1-2 | 4.163375 | -3.32944 | 3.863315 | -0.86181 | 0.388793 | NA |
| Ir48b | 0 | NA | NA | NA | NA | NA |
| Ir48a | 0 | NA | NA | NA | NA | NA |
| CG13196 | 0 | NA | NA | NA | NA | NA |
| tRNA:Met- | 0.676061 | -1.55209 | 3.991909 | -0.38881 | 0.697417 | NA |
| tRNA:Arg- | 0 | NA | NA | NA | NA | NA |
| CG13228 | 0.356107 | -1.47099 | 4.037764 | -0.36431 | 0.715628 | NA |
| CG13227 | 0 | NA | NA | NA | NA | NA |
| CG13226 | 0 | NA | NA | NA | NA | NA |
| CR30029 | 0 | NA | NA | NA | NA | NA |
| Or47a | 0 | NA | NA | NA | NA | NA |
| Cpr47Eb | 0.233093 | 0.692656 | 4.076246 | 0.169925 | 0.865069 | NA |
| CG13223 | 0 | NA | NA | NA | NA | NA |
| Cpr47Ee | 0.356107 | -1.47099 | 4.037764 | -0.36431 | 0.715628 | NA |
| CG30033 | 53.44971 | -0.65528 | 3.854454 | -0.17001 | 0.865005 | NA |
| sha | 0.282753 | -0.25398 | 4.020204 | -0.06318 | 0.949626 | NA |
| lambdaTry | 0 | NA | NA | NA | NA | NA |
| betaTry | 0.492076 | 0.930335 | 3.992579 | 0.233016 | 0.815749 | NA |
| CG30031 | 0 | NA | NA | NA | NA | NA |
| deltaTry | 0 | NA | NA | NA | NA | NA |
| iotaTry | 0 | NA | NA | NA | NA | NA |
| Gr47b | 0.712213 | -1.57119 | 3.998613 | -0.39293 | 0.694369 | NA |
| tRNA:Thr-1 | 0 | NA | NA | NA | NA | NA |
| tRNA:Thr-1 | 0.178053 | -1.16467 | 4.075067 | -0.2858 | 0.775028 | NA |
| Tsp47F | 0 | NA | NA | NA | NA | NA |
| tRNA:SeC- | 1.631653 | 1.102918 | 3.977952 | 0.277258 | 0.781582 | NA |
| CG13202 | 0 | NA | NA | NA | NA | NA |
| CG30025 | 0 | NA | NA | NA | NA | NA |
| gammaTry | 0 | NA | NA | NA | NA | NA |
| epsilonTry | 0.233093 | 0.692656 | 4.076246 | 0.169925 | 0.865069 | NA |
| alphaTry | 0 | NA | NA | NA | NA | NA |
| thetaTry | 0 | NA | NA | NA | NA | NA |
| etaTry | 0 | NA | NA | NA | NA | NA |
| zetaTry | 0 | NA | NA | NA | NA | NA |
| kappaTry | 0 | NA | NA | NA | NA | NA |
| Or47b | 0 | NA | NA | NA | NA | NA |
| Obp47b | 0 | NA | NA | NA | NA | NA |
| CG18335 | 0 | NA | NA | NA | NA | NA |
| TpnC47D | 0 | NA | NA | NA | NA | NA |

| | | | | | | |
|---|---|---|---|---|---|---|
| Cpr47Ef | 0.732439 | 2.426673 | 3.9948 | 0.607458 | 0.543547 | NA |
| Cpr47Ed | 0.104623 | 0.639075 | 4.067674 | 0.157111 | 0.875158 | NA |
| Cpr47Ec | 0 | NA | NA | NA | NA | NA |
| Cpr47Ea | 0 | NA | NA | NA | NA | NA |
| Listericin | 0 | NA | NA | NA | NA | NA |
| CG13215 | 0 | NA | NA | NA | NA | NA |
| CG13216 | 0.356107 | -1.47099 | 4.037764 | -0.36431 | 0.715628 | NA |
| CG13217 | 0.941609 | 1.421141 | 3.984935 | 0.356628 | 0.72137 | NA |
| CG13218 | 0 | NA | NA | NA | NA | NA |
| CG9084 | 0.178053 | -1.16467 | 4.075067 | -0.2858 | 0.775028 | NA |
| polyph | 0 | NA | NA | NA | NA | NA |
| Obp47a | 0 | NA | NA | NA | NA | NA |
| BBS4 | 7.328995 | 1.727313 | 3.957831 | 0.436429 | 0.662525 | NA |
| CG13231 | 0 | NA | NA | NA | NA | NA |
| CG13230 | 0 | NA | NA | NA | NA | NA |
| Spn47C | 0 | NA | NA | NA | NA | NA |
| CAH13 | 43.29042 | -1.02999 | 3.920407 | -0.26272 | 0.792763 | NA |
| mms4 | 6.024553 | -2.47201 | 3.924044 | -0.62997 | 0.528717 | NA |
| CG30016 | 0 | NA | NA | NA | NA | NA |
| CG33474 | 2.637141 | -1.81621 | 3.890553 | -0.46683 | 0.640625 | NA |
| CG12896 | 23.8804 | -3.71126 | 3.824901 | -0.97029 | 0.331902 | NA |
| CG33475 | 0 | NA | NA | NA | NA | NA |
| CG33477 | 0.178053 | -1.16467 | 4.075067 | -0.2858 | 0.775028 | NA |
| CG12898 | 0 | NA | NA | NA | NA | NA |
| CG12917 | 0 | NA | NA | NA | NA | NA |
| Or46a | 0 | NA | NA | NA | NA | NA |
| Hdc | 44.45506 | -5.83852 | 3.86123 | -1.51209 | 0.130511 | NA |
| CG12914 | 0 | NA | NA | NA | NA | NA |
| CG12911 | 0 | NA | NA | NA | NA | NA |
| CG12909 | 4.711497 | 1.713057 | 3.961514 | 0.432425 | 0.665433 | NA |
| CG12902 | 0 | NA | NA | NA | NA | NA |
| Obp46a | 0 | NA | NA | NA | NA | NA |
| lncRNA:CR | 8.998797 | -1.16537 | 3.822818 | -0.30485 | 0.760484 | NA |
| CG12209 | 0.357625 | -1.47707 | 4.044794 | -0.36518 | 0.714979 | NA |
| CG12912 | 0 | NA | NA | NA | NA | NA |
| Lime | 0.756516 | 2.906385 | 3.966481 | 0.732736 | 0.463719 | NA |
| CG12744 | 42.69616 | -0.47209 | 3.884184 | -0.12154 | 0.903262 | NA |
| CG12923 | 57.31855 | 0.420785 | 3.8345 | 0.109737 | 0.912618 | NA |
| CG12133 | 0 | NA | NA | NA | NA | NA |
| CG12134 | 0 | NA | NA | NA | NA | NA |
| eve | 0 | NA | NA | NA | NA | NA |
| CG12920 | 0.233093 | 0.692656 | 4.076246 | 0.169925 | 0.865069 | NA |
| PIG-N | 58.84852 | -8.91412 | 3.845317 | -2.31818 | 0.02044 | NA |
| Def | 0 | NA | NA | NA | NA | NA |
| CG1441 | 0 | NA | NA | NA | NA | NA |
| tRNA:iMet | 11.93033 | -2.91436 | 3.923013 | -0.74289 | 0.45755 | NA |
| CG30002 | 0 | NA | NA | NA | NA | NA |
| CG1773 | 0 | NA | NA | NA | NA | NA |
| CG10459 | 0.104623 | 0.639075 | 4.067674 | 0.157111 | 0.875158 | NA |
| CG1698 | 6.50284 | 3.679278 | 3.897972 | 0.943896 | 0.345223 | NA |

| | | | | | | |
|---|---|---|---|---|---|---|
| Orc6 | 37.92535 | -1.70516 | 3.951919 | -0.43148 | 0.666123 | NA |
| hebe | 54.7262 | 0.712219 | 3.835088 | 0.185711 | 0.852671 | NA |
| dila | 0 | NA | NA | NA | NA | NA |
| CG30001 | 0.282677 | -0.09599 | 4.057934 | -0.02366 | 0.981127 | NA |
| CG1690 | 0.178053 | -1.16467 | 4.075067 | -0.2858 | 0.775028 | NA |
| Wnt2 | 5.519652 | -1.68736 | 3.957756 | -0.42634 | 0.669859 | NA |
| Or45b | 0 | NA | NA | NA | NA | NA |
| Alp6 | 0 | NA | NA | NA | NA | NA |
| CG1827 | 1.822665 | -0.02704 | 3.958245 | -0.00683 | 0.99455 | NA |
| CG30338 | 56.14706 | -2.2755 | 3.952625 | -0.57569 | 0.564823 | NA |
| CG30339 | 4.576209 | 0.305937 | 3.955371 | 0.077347 | 0.938347 | NA |
| CG30340 | 0 | NA | NA | NA | NA | NA |
| Smyd4-1 | 0.178812 | -1.08808 | 4.061735 | -0.26789 | 0.788788 | NA |
| CG13955 | 0 | NA | NA | NA | NA | NA |
| CG8800 | 0 | NA | NA | NA | NA | NA |
| CNT1 | 0 | NA | NA | NA | NA | NA |
| CG8788 | 0 | NA | NA | NA | NA | NA |
| Rad51D | 42.71757 | 1.749839 | 3.952123 | 0.442759 | 0.65794 | NA |
| unpg | 0 | NA | NA | NA | NA | NA |
| Tsen15 | 0 | NA | NA | NA | NA | NA |
| CG8026 | 0 | NA | NA | NA | NA | NA |
| CG8008 | 3.769197 | 1.70343 | 3.964029 | 0.429722 | 0.667398 | NA |
| Boot | 3.900384 | -5.46701 | 3.920402 | -1.3945 | 0.163166 | NA |
| CG13742 | 0 | NA | NA | NA | NA | NA |
| CG30350 | 0 | NA | NA | NA | NA | NA |
| CG13749 | 7.119595 | 1.726543 | 3.958029 | 0.436213 | 0.662682 | NA |
| CG13748 | 0 | NA | NA | NA | NA | NA |
| CG13747 | 0 | NA | NA | NA | NA | NA |
| CG8197 | 0 | NA | NA | NA | NA | NA |
| Ance-4 | 15.48424 | 1.530975 | 3.953223 | 0.387273 | 0.698554 | NA |
| CG8170 | 0.356107 | -1.47099 | 4.037764 | -0.36431 | 0.715628 | NA |
| CG13744 | 0 | NA | NA | NA | NA | NA |
| CG8172 | 0 | NA | NA | NA | NA | NA |
| FANCI | 0.712213 | -1.57119 | 3.998613 | -0.39293 | 0.694369 | NA |
| Spt | 53.36638 | 0.31979 | 3.950572 | 0.080948 | 0.935484 | NA |
| CG8252 | 0 | NA | NA | NA | NA | NA |
| CG14752 | 0 | NA | NA | NA | NA | NA |
| UQCR-11L | 10.55945 | 0.864524 | 3.954652 | 0.218609 | 0.826954 | NA |
| tRNA:Leu-( | 0 | NA | NA | NA | NA | NA |
| Jon44E | 0 | NA | NA | NA | NA | NA |
| CG8738 | 0.178812 | -1.08808 | 4.061735 | -0.26789 | 0.788788 | NA |
| CG12376 | 0.104623 | 0.639075 | 4.067674 | 0.157111 | 0.875158 | NA |
| PGRP-SC1a | 0 | NA | NA | NA | NA | NA |
| PGRP-SC1b | 0.132078 | -1.13211 | 4.069918 | -0.27817 | 0.780885 | NA |
| PGRP-SC2 | 0.178053 | -1.16467 | 4.075067 | -0.2858 | 0.775028 | NA |
| CG8746 | 0 | NA | NA | NA | NA | NA |
| CG30356 | 0.4188 | 1.41068 | 4.034613 | 0.349644 | 0.726606 | NA |
| CG14743 | 0.466186 | 0.959471 | 4.033986 | 0.237847 | 0.812 | NA |
| CG14744 | 0 | NA | NA | NA | NA | NA |
| CG8584 | 0 | NA | NA | NA | NA | NA |

| | | | | | | |
|---|---|---|---|---|---|---|
| CG8586 | 0 | NA | NA | NA | NA | NA |
| CG30355 | 4.476411 | -4.95604 | 3.938499 | -1.25836 | 0.208262 | NA |
| CG8642 | 0 | NA | NA | NA | NA | NA |
| CG8736 | 3.975684 | 1.507544 | 3.959717 | 0.38072 | 0.703411 | NA |
| CG12780 | 6.053811 | -1.68915 | 3.957162 | -0.42686 | 0.669482 | NA |
| Lcp3 | 0 | NA | NA | NA | NA | NA |
| Lcp4 | 0 | NA | NA | NA | NA | NA |
| Cyp4e2 | 0.4188 | 1.41068 | 4.034613 | 0.349644 | 0.726606 | NA |
| Cyp4e1 | 17.8166 | 6.236091 | 3.906317 | 1.596412 | 0.110397 | NA |
| Mal-A2 | 0 | NA | NA | NA | NA | NA |
| Mal-A4 | 2.144988 | -3.20556 | 3.997331 | -0.80193 | 0.422596 | NA |
| Mal-A6 | 0 | NA | NA | NA | NA | NA |
| Mal-A8 | 0.893302 | -2.63419 | 4.009574 | -0.65698 | 0.511197 | NA |
| Mal-A7 | 0.178812 | -1.08808 | 4.061735 | -0.26789 | 0.788788 | NA |
| Mal-A3 | 0 | NA | NA | NA | NA | NA |
| Mal-A1 | 0.1047 | 0.613261 | 4.062799 | 0.150945 | 0.880019 | NA |
| Cyp4ad1 | 0 | NA | NA | NA | NA | NA |
| Lcp2 | 56.49656 | 1.537131 | 3.951545 | 0.388995 | 0.69728 | NA |
| Lcp1Psi | 0 | NA | NA | NA | NA | NA |
| Lcp1 | 0.466186 | 0.959471 | 4.033986 | 0.237847 | 0.812 | NA |
| CG2121 | 24.91085 | 3.934285 | 3.882925 | 1.013227 | 0.310952 | NA |
| CG2127 | 8.069575 | 5.526343 | 3.91921 | 1.410066 | 0.15852 | NA |
| cola | 0 | NA | NA | NA | NA | NA |
| swif | 0 | NA | NA | NA | NA | NA |
| hubl | 0 | NA | NA | NA | NA | NA |
| spaw | 0 | NA | NA | NA | NA | NA |
| sut1 | 52.40369 | 1.229032 | 3.863023 | 0.318153 | 0.750369 | NA |
| sut2 | 0 | NA | NA | NA | NA | NA |
| sut3 | 44.9711 | -4.00921 | 3.819314 | -1.04972 | 0.293847 | NA |
| sand | 10.55843 | 0.362766 | 3.839992 | 0.094471 | 0.924735 | NA |
| Nup50 | 24.22524 | -1.42522 | 3.906718 | -0.36481 | 0.71525 | NA |
| tRNA:Ile-A | 2.564026 | 1.129155 | 3.968527 | 0.284527 | 0.776006 | NA |
| Mlh1 | 0.722036 | -1.99346 | 3.98794 | -0.49987 | 0.617165 | NA |
| CG14760 | 0 | NA | NA | NA | NA | NA |
| CG30369 | 0 | NA | NA | NA | NA | NA |
| CG14759 | 0 | NA | NA | NA | NA | NA |
| CG30371 | 0.178812 | -1.08808 | 4.061735 | -0.26789 | 0.788788 | NA |
| CG30375 | 0 | NA | NA | NA | NA | NA |
| CG30376 | 0 | NA | NA | NA | NA | NA |
| CG12126 | 0 | NA | NA | NA | NA | NA |
| CG2291 | 0 | NA | NA | NA | NA | NA |
| dpn | 15.70146 | -6.64616 | 3.914944 | -1.69764 | 0.089576 | NA |
| CG14756 | 1.602479 | -1.64094 | 3.97352 | -0.41297 | 0.67963 | NA |
| CG8708 | 0 | NA | NA | NA | NA | NA |
| CR30374 | 0 | NA | NA | NA | NA | NA |
| CG12824 | 0 | NA | NA | NA | NA | NA |
| PolZ1 | 0.356107 | -1.47099 | 4.037764 | -0.36431 | 0.715628 | NA |
| CG30377 | 38.62953 | -5.86643 | 3.829744 | -1.53181 | 0.12557 | NA |
| CG30379 | 0 | NA | NA | NA | NA | NA |
| azot | 0.418493 | 1.313669 | 4.019283 | 0.326842 | 0.743788 | NA |

| | | | | | | |
|---|---|---|---|---|---|---|
| CG30378 | 0 | NA | NA | NA | NA | NA |
| Odc1 | 0.233093 | 0.692656 | 4.076246 | 0.169925 | 0.865069 | NA |
| Odc2 | 0 | NA | NA | NA | NA | NA |
| Hey | 0.178053 | -1.16467 | 4.075067 | -0.2858 | 0.775028 | NA |
| Dic3 | 0 | NA | NA | NA | NA | NA |
| CG14763 | 61.78676 | -3.995 | 3.887632 | -1.02762 | 0.30413 | NA |
| Sep-05 | 20.43145 | -1.6585 | 3.846982 | -0.43112 | 0.666383 | NA |
| CG30380 | 0.1047 | 0.613261 | 4.062799 | 0.150945 | 0.880019 | NA |
| Prosalpha1 | 0 | NA | NA | NA | NA | NA |
| TTLL12 | 57.4356 | -4.32895 | 3.794228 | -1.14093 | 0.253899 | NA |
| Gpo2 | 11.0393 | -1.69765 | 3.954364 | -0.42931 | 0.667697 | NA |
| CG2070 | 0 | NA | NA | NA | NA | NA |
| CG2065 | 0.337793 | 1.451235 | 4.039844 | 0.35923 | 0.719423 | NA |
| Dgat2 | 0 | NA | NA | NA | NA | NA |
| CG1946 | 0 | NA | NA | NA | NA | NA |
| cn | 0.132078 | -1.13211 | 4.069918 | -0.27817 | 0.780885 | NA |
| tor | 0.69928 | 1.019866 | 4.009445 | 0.254366 | 0.799213 | NA |
| CG12826 | 0 | NA | NA | NA | NA | NA |
| CG1339 | 0 | NA | NA | NA | NA | NA |
| Rpe | 13.54535 | 6.212589 | 3.914471 | 1.587083 | 0.112494 | NA |
| CG12164 | 0 | NA | NA | NA | NA | NA |
| CG11112 | 15.66869 | -1.70075 | 3.953349 | -0.43021 | 0.667046 | NA |
| CG11113 | 0 | NA | NA | NA | NA | NA |
| so | 2.8794 | 0.337993 | 3.953187 | 0.085499 | 0.931865 | NA |
| CG11125 | 41.35647 | 1.74968 | 3.952163 | 0.442715 | 0.657972 | NA |
| fa2h | 0 | NA | NA | NA | NA | NA |
| kappaB-Ra | 0 | NA | NA | NA | NA | NA |
| CG30503 | 0 | NA | NA | NA | NA | NA |
| CG11145 | 0 | NA | NA | NA | NA | NA |
| CG1701 | 0 | NA | NA | NA | NA | NA |
| Gr43a | 0.710103 | -0.24484 | 3.975022 | -0.0616 | 0.950885 | NA |
| Tsp42Ei | 0.178053 | -1.16467 | 4.075067 | -0.2858 | 0.775028 | NA |
| Tsp42El | 34.39633 | 0.558152 | 3.883973 | 0.143706 | 0.885732 | NA |
| Tsp42Eo | 0 | NA | NA | NA | NA | NA |
| Tsp42Er | 36.65652 | -2.27186 | 3.953529 | -0.57464 | 0.565534 | NA |
| CG12836 | 0.178053 | -1.16467 | 4.075067 | -0.2858 | 0.775028 | NA |
| Spn43Aa | 0 | NA | NA | NA | NA | NA |
| CG11060 | 0 | NA | NA | NA | NA | NA |
| CG33140 | 0 | NA | NA | NA | NA | NA |
| CG30385 | 0 | NA | NA | NA | NA | NA |
| CG30384 | 0.550724 | 1.292521 | 3.972655 | 0.325355 | 0.744913 | NA |
| Or43a | 0 | NA | NA | NA | NA | NA |
| Spn43Ad | 45.54681 | -0.27118 | 3.835068 | -0.07071 | 0.943628 | NA |
| Cyp9b2 | 26.27968 | 1.746819 | 3.952878 | 0.441911 | 0.658554 | NA |
| Cyp9b1 | 0 | NA | NA | NA | NA | NA |
| CG12831 | 0 | NA | NA | NA | NA | NA |
| tRNA:Thr-/ | 0 | NA | NA | NA | NA | NA |
| tRNA:Thr-/ | 0 | NA | NA | NA | NA | NA |
| Tsp42Eq | 0 | NA | NA | NA | NA | NA |
| Tsp42Ep | 13.9432 | -0.65937 | 3.927737 | -0.16787 | 0.866682 | NA |

| | | | | | | |
|---|---|---|---|---|---|---|
| CG1674 | 29.4972 | -6.15362 | 3.892941 | -1.58071 | 0.113944 | NA |
| CG32006 | 0.715249 | -1.88064 | 4.034336 | -0.46616 | 0.641102 | NA |
| CR32009 | 0 | NA | NA | NA | NA | NA |
| CR32010 | 0 | NA | NA | NA | NA | NA |
| CR32011 | 0.1047 | 0.613261 | 4.062799 | 0.150945 | 0.880019 | NA |
| CG11077 | 0.596546 | 0.421256 | 3.969213 | 0.106131 | 0.915479 | NA |
| CR11023 | 0 | NA | NA | NA | NA | NA |
| Cpsf160 | 9.238209 | -1.83289 | 3.929843 | -0.4664 | 0.640926 | NA |
| betaNACte | 0 | NA | NA | NA | NA | NA |
| pb | 0.341401 | 0.594712 | 4.003898 | 0.148533 | 0.881922 | NA |
| Pxd | 0.178812 | -1.08808 | 4.061735 | -0.26789 | 0.788788 | NA |
| tRNA:Tyr-( | 1.188699 | -1.8233 | 3.980741 | -0.45803 | 0.646931 | NA |
| tRNA:Tyr-( | 0 | NA | NA | NA | NA | NA |
| tRNA:Tyr-( | 0 | NA | NA | NA | NA | NA |
| tRNA:Tyr-( | 0 | NA | NA | NA | NA | NA |
| tRNA:Tyr-( | 0 | NA | NA | NA | NA | NA |
| tRNA:Leu-( | 0.46073 | -0.17853 | 4.00793 | -0.04454 | 0.964471 | NA |
| tRNA:Leu-( | 0.233093 | 0.692656 | 4.076246 | 0.169925 | 0.865069 | NA |
| mia | 17.33667 | -4.8345 | 3.903313 | -1.23856 | 0.215507 | NA |
| CG10725 | 0 | NA | NA | NA | NA | NA |
| Lsp2 | 0 | NA | NA | NA | NA | NA |
| asRNA:CR1 | 0 | NA | NA | NA | NA | NA |
| CG13064 | 0 | NA | NA | NA | NA | NA |
| nompA | 0.356107 | -1.47099 | 4.037764 | -0.36431 | 0.715628 | NA |
| CG13970 | 0 | NA | NA | NA | NA | NA |
| CG15553 | 0 | NA | NA | NA | NA | NA |
| Cyp12d1-d | 0 | NA | NA | NA | NA | NA |
| Cyp12d1-p | 0.492 | 1.04661 | 3.990653 | 0.262265 | 0.793117 | NA |
| CG18853 | 1.251686 | -2.06022 | 4.011654 | -0.51356 | 0.60756 | NA |
| nxf2 | 3.717393 | 0.271825 | 3.852595 | 0.070556 | 0.943751 | NA |
| nan | 50.25597 | 1.750571 | 3.951942 | 0.442965 | 0.657791 | NA |
| CG33489 | 0 | NA | NA | NA | NA | NA |
| CG33490 | 2.439542 | 0.538613 | 3.869507 | 0.139194 | 0.889297 | NA |
| CG10931 | 0 | NA | NA | NA | NA | NA |
| CG5002 | 0 | NA | NA | NA | NA | NA |
| HLH54F | 0 | NA | NA | NA | NA | NA |
| CG6362 | 1.046999 | 1.598136 | 3.993006 | 0.400234 | 0.688984 | NA |
| CG14488 | 0 | NA | NA | NA | NA | NA |
| Sardh | 0.368855 | -0.37404 | 4.015709 | -0.09314 | 0.925789 | NA |
| Zasp66 | 3.674452 | 0.012501 | 3.939617 | 0.003173 | 0.997468 | NA |
| GluRIIB | 0.178053 | -1.16467 | 4.075067 | -0.2858 | 0.775028 | NA |
| CG7730 | 0.178812 | -1.08808 | 4.061735 | -0.26789 | 0.788788 | NA |
| TpnC73F | 0.566189 | -0.21299 | 4.010979 | -0.0531 | 0.957651 | NA |
| CG33493 | 44.05541 | 0.362719 | 3.834619 | 0.094591 | 0.92464 | NA |
| CG33494 | 59.33321 | 4.105825 | 3.886482 | 1.056437 | 0.290768 | NA |
| tRNA:Gly-( | 0 | NA | NA | NA | NA | NA |
| Cec2 | 0 | NA | NA | NA | NA | NA |
| CG10910 | 0 | NA | NA | NA | NA | NA |
| CG12783 | 0.523116 | 1.34877 | 4.007682 | 0.336546 | 0.736459 | NA |
| Obp73a | 1.512901 | 1.047032 | 3.89748 | 0.268643 | 0.788204 | NA |

| | | | | | | |
|---|---|---|---|---|---|---|
| salto | 0 | NA | NA | NA | NA | NA |
| sut4 | 6.850833 | 3.786056 | 3.83481 | 0.987286 | 0.323502 | NA |
| CG14642 | 12.81654 | 7.110847 | 3.909828 | 1.818711 | 0.068956 | NA |
| CG14692 | 0.178053 | -1.16467 | 4.075067 | -0.2858 | 0.775028 | NA |
| CG14736 | 0 | NA | NA | NA | NA | NA |
| Gr89a | 0 | NA | NA | NA | NA | NA |
| Tim17b2 | 0 | NA | NA | NA | NA | NA |
| Mlc2 | 0 | NA | NA | NA | NA | NA |
| CG2336 | 28.02054 | 0.443499 | 3.897798 | 0.113782 | 0.909411 | NA |
| gem | 1.602479 | -1.64094 | 3.97352 | -0.41297 | 0.67963 | NA |
| CG30270 | 0 | NA | NA | NA | NA | NA |
| mthl15 | 19.05693 | -0.75858 | 3.736961 | -0.20299 | 0.83914 | NA |
| bt | 2.366851 | -3.31349 | 3.876975 | -0.85466 | 0.39274 | NA |
| Gss2 | 0 | NA | NA | NA | NA | NA |
| Muc11A | 20.00271 | 3.552593 | 3.864163 | 0.919369 | 0.357902 | NA |
| CG3277 | 1.465799 | 1.63603 | 3.98226 | 0.410829 | 0.681198 | NA |
| CR33318 | 1.569349 | 1.463746 | 3.972245 | 0.368493 | 0.712505 | NA |
| CR33319 | 0 | NA | NA | NA | NA | NA |
| CG33459 | 0 | NA | NA | NA | NA | NA |
| CG33467 | 0.178053 | -1.16467 | 4.075067 | -0.2858 | 0.775028 | NA |
| CREG | 15.51291 | -0.89745 | 3.872436 | -0.23175 | 0.816729 | NA |
| Ugt49B2 | 0 | NA | NA | NA | NA | NA |
| CG7824 | 2.164524 | -2.16108 | 3.916813 | -0.55174 | 0.581124 | NA |
| CG8852 | 0.178053 | -1.16467 | 4.075067 | -0.2858 | 0.775028 | NA |
| lncRNA:CR | 8.094503 | 2.573403 | 3.872154 | 0.664592 | 0.506311 | NA |
| Jheh2 | 13.67116 | 0.36537 | 3.838222 | 0.095193 | 0.924162 | NA |
| att-ORFB | 0 | NA | NA | NA | NA | NA |
| alpha-Est4 | 0 | NA | NA | NA | NA | NA |
| Cyp4aa1 | 0 | NA | NA | NA | NA | NA |
| Lsd-1 | 11.26517 | -2.24925 | 3.959234 | -0.5681 | 0.569965 | NA |
| CG10853 | 2.381275 | 1.651403 | 3.949799 | 0.418098 | 0.675875 | NA |
| FoxL1 | 30.90618 | -1.91869 | 3.952168 | -0.48548 | 0.627338 | NA |
| Gr64a | 0 | NA | NA | NA | NA | NA |
| Gr64b | 0 | NA | NA | NA | NA | NA |
| Gr64c | 0 | NA | NA | NA | NA | NA |
| CG14984 | 0 | NA | NA | NA | NA | NA |
| CG14983 | 0 | NA | NA | NA | NA | NA |
| CG32599 | 0 | NA | NA | NA | NA | NA |
| CG1461 | 0 | NA | NA | NA | NA | NA |
| CG9400 | 0 | NA | NA | NA | NA | NA |
| CG11674 | 0.178053 | -1.16467 | 4.075067 | -0.2858 | 0.775028 | NA |
| peb | 4.354582 | -4.13202 | 3.889311 | -1.0624 | 0.288053 | NA |
| pon | 0 | NA | NA | NA | NA | NA |
| CG15473 | 25.3609 | -7.34659 | 3.91328 | -1.87735 | 0.06047 | NA |
| CG3062 | 0 | NA | NA | NA | NA | NA |
| CG3081 | 0.356866 | -1.92326 | 4.037889 | -0.4763 | 0.633858 | NA |
| CG12684 | 0 | NA | NA | NA | NA | NA |
| Chd3 | 0 | NA | NA | NA | NA | NA |
| CG32214 | 1.450033 | 2.807816 | 3.907765 | 0.718522 | 0.472435 | NA |
| CG12519 | 0.178053 | -1.16467 | 4.075067 | -0.2858 | 0.775028 | NA |

| | | | | | | |
|---|---|---|---|---|---|---|
| CG32213 | 0 | NA | NA | NA | NA | NA |
| CG32212 | 0.209247 | 1.042804 | 4.047837 | 0.25762 | 0.7967 | NA |
| Cpr76Bb | 0.178053 | -1.16467 | 4.075067 | -0.2858 | 0.775028 | NA |
| Cpr76Ba | 0 | NA | NA | NA | NA | NA |
| CG9452 | 11.51692 | 2.942648 | 3.926413 | 0.74945 | 0.453586 | NA |
| CG9449 | 0 | NA | NA | NA | NA | NA |
| brv1 | 0.178053 | -1.16467 | 4.075067 | -0.2858 | 0.775028 | NA |
| hpRNA:CR: | 61.60795 | -3.52692 | 3.901503 | -0.90399 | 0.366001 | NA |
| CG18294 | 0 | NA | NA | NA | NA | NA |
| 825-Oak | 0 | NA | NA | NA | NA | NA |
| CG14096 | 0 | NA | NA | NA | NA | NA |
| CG14095 | 0.178053 | -1.16467 | 4.075067 | -0.2858 | 0.775028 | NA |
| CG33062 | 0 | NA | NA | NA | NA | NA |
| CG1835 | 0 | NA | NA | NA | NA | NA |
| hydra | 0 | NA | NA | NA | NA | NA |
| Ser7 | 0 | NA | NA | NA | NA | NA |
| CG1986 | 0 | NA | NA | NA | NA | NA |
| CG15249 | 0 | NA | NA | NA | NA | NA |
| CG15247 | 24.70918 | 1.746322 | 3.953003 | 0.441771 | 0.658655 | NA |
| CG33502 | 0 | NA | NA | NA | NA | NA |
| CG32819 | 0 | NA | NA | NA | NA | NA |
| CG17450 | 0 | NA | NA | NA | NA | NA |
| CG32820 | 0 | NA | NA | NA | NA | NA |
| RpS10a | 8.103901 | 1.37305 | 3.955703 | 0.347106 | 0.728511 | NA |
| CG12880 | 0.104623 | 0.639075 | 4.067674 | 0.157111 | 0.875158 | NA |
| CG31055 | 1.246373 | -1.6238 | 3.979526 | -0.40804 | 0.683245 | NA |
| Mlc1 | 0.356107 | -1.47099 | 4.037764 | -0.36431 | 0.715628 | NA |
| CG5612 | 1.120353 | 0.8435 | 3.975033 | 0.212199 | 0.831951 | NA |
| Crg-1 | 14.09855 | 6.466143 | 3.912917 | 1.652512 | 0.09843 | NA |
| CG2901 | 0 | NA | NA | NA | NA | NA |
| CR33221 | 0.104623 | 0.639075 | 4.067674 | 0.157111 | 0.875158 | NA |
| CG15625 | 0 | NA | NA | NA | NA | NA |
| CG3036 | 3.70255 | -0.58155 | 3.863881 | -0.15051 | 0.880362 | NA |
| ND-49L | 0 | NA | NA | NA | NA | NA |
| Cdc23 | 6.992797 | 1.160607 | 3.957551 | 0.293264 | 0.769321 | NA |
| CG31683 | 0 | NA | NA | NA | NA | NA |
| CG15130 | 0 | NA | NA | NA | NA | NA |
| TpnC25D | 0.942299 | 1.584447 | 3.996979 | 0.396411 | 0.691802 | NA |
| CG12934 | 0.233093 | 0.692656 | 4.076246 | 0.169925 | 0.865069 | NA |
| CG17601 | 0 | NA | NA | NA | NA | NA |
| Cluap1 | 0 | NA | NA | NA | NA | NA |
| Obp51a | 0.233093 | 0.692656 | 4.076246 | 0.169925 | 0.865069 | NA |
| Smyd4-3 | 0.104623 | 0.639075 | 4.067674 | 0.157111 | 0.875158 | NA |
| CG7763 | 0.2094 | 1.005034 | 4.045341 | 0.248442 | 0.803792 | NA |
| Drip | 0.178812 | -1.08808 | 4.061735 | -0.26789 | 0.788788 | NA |
| CG30026 | 0 | NA | NA | NA | NA | NA |
| COX4L | 0 | NA | NA | NA | NA | NA |
| IFT20 | 0 | NA | NA | NA | NA | NA |
| tRNA:Glu-( | 0 | NA | NA | NA | NA | NA |
| tRNA:Glu-( | 0 | NA | NA | NA | NA | NA |

| | | | | | | |
|---|---|---|---|---|---|---|
| 5SrRNA:CR | 0 | NA | NA | NA | NA | NA |
| 5SrRNA:CR | 0 | NA | NA | NA | NA | NA |
| 5SrRNA:CR | 0 | NA | NA | NA | NA | NA |
| 5SrRNA-Ps | 0 | NA | NA | NA | NA | NA |
| 5SrRNA:CR | 0 | NA | NA | NA | NA | NA |
| 5SrRNA:CR | 0 | NA | NA | NA | NA | NA |
| 5SrRNA:CR | 0 | NA | NA | NA | NA | NA |
| 5SrRNA:CR | 0 | NA | NA | NA | NA | NA |
| 5SrRNA:CR | 0 | NA | NA | NA | NA | NA |
| 5SrRNA:CR | 0 | NA | NA | NA | NA | NA |
| 5SrRNA-Ps | 0 | NA | NA | NA | NA | NA |
| 5SrRNA:CR | 0 | NA | NA | NA | NA | NA |
| 5SrRNA:CR | 0 | NA | NA | NA | NA | NA |
| 5SrRNA:CR | 0 | NA | NA | NA | NA | NA |
| 5SrRNA:CR | 0 | NA | NA | NA | NA | NA |
| 5SrRNA:CR | 0 | NA | NA | NA | NA | NA |
| 5SrRNA:CR | 0 | NA | NA | NA | NA | NA |
| 5SrRNA:CR | 0 | NA | NA | NA | NA | NA |
| 5SrRNA-Ps | 0 | NA | NA | NA | NA | NA |
| 5SrRNA:CR | 0 | NA | NA | NA | NA | NA |
| 5SrRNA:CR | 0 | NA | NA | NA | NA | NA |
| 5SrRNA:CR | 0 | NA | NA | NA | NA | NA |
| 5SrRNA:CR | 0 | NA | NA | NA | NA | NA |
| 5SrRNA:CR | 0 | NA | NA | NA | NA | NA |
| 5SrRNA:CR | 0 | NA | NA | NA | NA | NA |
| 5SrRNA:CR | 0 | NA | NA | NA | NA | NA |
| 5SrRNA:CR | 0 | NA | NA | NA | NA | NA |
| 5SrRNA:CR | 0 | NA | NA | NA | NA | NA |
| 5SrRNA:CR | 0 | NA | NA | NA | NA | NA |
| 5SrRNA:CR | 0 | NA | NA | NA | NA | NA |
| 5SrRNA:CR | 0 | NA | NA | NA | NA | NA |
| 5SrRNA:CR | 0 | NA | NA | NA | NA | NA |
| 5SrRNA:CR | 0 | NA | NA | NA | NA | NA |
| 5SrRNA:CR | 0 | NA | NA | NA | NA | NA |
| 5SrRNA:CR | 0 | NA | NA | NA | NA | NA |
| 5SrRNA:CR | 0 | NA | NA | NA | NA | NA |
| 5SrRNA:CR | 0 | NA | NA | NA | NA | NA |
| 5SrRNA:CR | 0 | NA | NA | NA | NA | NA |
| 5SrRNA:CR | 0 | NA | NA | NA | NA | NA |
| 5SrRNA:CR | 0 | NA | NA | NA | NA | NA |
| 5SrRNA:CR | 0 | NA | NA | NA | NA | NA |
| 5SrRNA:CR | 0 | NA | NA | NA | NA | NA |
| 5SrRNA:CR | 0 | NA | NA | NA | NA | NA |
| 5SrRNA:CR | 0 | NA | NA | NA | NA | NA |
| 5SrRNA:CR | 0 | NA | NA | NA | NA | NA |
| 5SrRNA:CR | 0 | NA | NA | NA | NA | NA |
| 5SrRNA:CR | 0 | NA | NA | NA | NA | NA |
| 5SrRNA:CR | 0 | NA | NA | NA | NA | NA |
| 5SrRNA:CR | 0 | NA | NA | NA | NA | NA |
| 5SrRNA:CR | 0 | NA | NA | NA | NA | NA |

| | | | | | | |
|---|---|---|---|---|---|---|
| 5SrRNA:CF | 0 | NA | NA | NA | NA | NA |
| 5SrRNA:CF | 0 | NA | NA | NA | NA | NA |
| 5SrRNA:CF | 0 | NA | NA | NA | NA | NA |
| 5SrRNA:CF | 0 | NA | NA | NA | NA | NA |
| 5SrRNA:CF | 0 | NA | NA | NA | NA | NA |
| 5SrRNA:CF | 0 | NA | NA | NA | NA | NA |
| 5SrRNA:CF | 0 | NA | NA | NA | NA | NA |
| 5SrRNA:CF | 0 | NA | NA | NA | NA | NA |
| 5SrRNA:CF | 0 | NA | NA | NA | NA | NA |
| 5SrRNA:CF | 0 | NA | NA | NA | NA | NA |
| 5SrRNA:CF | 0 | NA | NA | NA | NA | NA |
| 5SrRNA:CF | 0 | NA | NA | NA | NA | NA |
| 5SrRNA:CF | 0 | NA | NA | NA | NA | NA |
| 5SrRNA-Ps | 0 | NA | NA | NA | NA | NA |
| 5SrRNA:CF | 0 | NA | NA | NA | NA | NA |
| 5SrRNA:CF | 0 | NA | NA | NA | NA | NA |
| 5SrRNA:CF | 0 | NA | NA | NA | NA | NA |
| 5SrRNA:CF | 0 | NA | NA | NA | NA | NA |
| 5SrRNA:CF | 0 | NA | NA | NA | NA | NA |
| 5SrRNA:CF | 0 | NA | NA | NA | NA | NA |
| 5SrRNA:CF | 0 | NA | NA | NA | NA | NA |
| 5SrRNA:CF | 0 | NA | NA | NA | NA | NA |
| 5SrRNA:CF | 0 | NA | NA | NA | NA | NA |
| 5SrRNA:CF | 0 | NA | NA | NA | NA | NA |
| 5SrRNA:CF | 0 | NA | NA | NA | NA | NA |
| 5SrRNA:CF | 0 | NA | NA | NA | NA | NA |
| 5SrRNA:CF | 0 | NA | NA | NA | NA | NA |
| 5SrRNA:CF | 0 | NA | NA | NA | NA | NA |
| 5SrRNA:CF | 0 | NA | NA | NA | NA | NA |
| 5SrRNA:CF | 0 | NA | NA | NA | NA | NA |
| 5SrRNA:CF | 0 | NA | NA | NA | NA | NA |
| 5SrRNA:CF | 0 | NA | NA | NA | NA | NA |
| 5SrRNA:CF | 0 | NA | NA | NA | NA | NA |
| 5SrRNA:CF | 0 | NA | NA | NA | NA | NA |
| 5SrRNA:CF | 0 | NA | NA | NA | NA | NA |
| 5SrRNA:CF | 0 | NA | NA | NA | NA | NA |
| 5SrRNA:CF | 0 | NA | NA | NA | NA | NA |
| 5SrRNA:CF | 0 | NA | NA | NA | NA | NA |
| 5SrRNA:CF | 0 | NA | NA | NA | NA | NA |
| 5SrRNA:CF | 0 | NA | NA | NA | NA | NA |
| 5SrRNA:CF | 0 | NA | NA | NA | NA | NA |
| 5SrRNA:CF | 0 | NA | NA | NA | NA | NA |
| 5SrRNA:CF | 0 | NA | NA | NA | NA | NA |
| 5SrRNA:CF | 0 | NA | NA | NA | NA | NA |
| 5SrRNA:CF | 0 | NA | NA | NA | NA | NA |
| 5SrRNA:CF | 0 | NA | NA | NA | NA | NA |
| 5SrRNA:CF | 0 | NA | NA | NA | NA | NA |
| 5SrRNA:CF | 0 | NA | NA | NA | NA | NA |
| 5SrRNA:CF | 0 | NA | NA | NA | NA | NA |
| 5SrRNA:CF | 0 | NA | NA | NA | NA | NA |

| | | | | | | |
|---|---|---|---|---|---|---|
| TpnC41C | 0 | NA | NA | NA | NA | NA |
| CG12395 | 0 | NA | NA | NA | NA | NA |
| Ste:CG332 | 0 | NA | NA | NA | NA | NA |
| Ste:CG332 | 0 | NA | NA | NA | NA | NA |
| Ste:CG332 | 0 | NA | NA | NA | NA | NA |
| Ste:CG332 | 0 | NA | NA | NA | NA | NA |
| Ste:CG332 | 0 | NA | NA | NA | NA | NA |
| Ste:CG332 | 0 | NA | NA | NA | NA | NA |
| Ste:CG332 | 0 | NA | NA | NA | NA | NA |
| Ste:CG332 | 0 | NA | NA | NA | NA | NA |
| Ste:CG332 | 0 | NA | NA | NA | NA | NA |
| Ste:CG332 | 0 | NA | NA | NA | NA | NA |
| Ste:CG332 | 0 | NA | NA | NA | NA | NA |
| Ste:CG332 | 0 | NA | NA | NA | NA | NA |
| CG5853 | 0 | NA | NA | NA | NA | NA |
| Cpr30F | 0 | NA | NA | NA | NA | NA |
| TbCMF46 | 0 | NA | NA | NA | NA | NA |
| Muc30E | 6.52661 | 1.15925 | 3.958017 | 0.292887 | 0.769609 | NA |
| aspr | 0.178053 | -1.16467 | 4.075067 | -0.2858 | 0.775028 | NA |
| Peritrophi | 0.178053 | -1.16467 | 4.075067 | -0.2858 | 0.775028 | NA |
| CR33498 | 0 | NA | NA | NA | NA | NA |
| Sdic4 | 0 | NA | NA | NA | NA | NA |
| CR33487 | 0 | NA | NA | NA | NA | NA |
| Sdic3 | 0 | NA | NA | NA | NA | NA |
| CR33496 | 0 | NA | NA | NA | NA | NA |
| Sdic2 | 0 | NA | NA | NA | NA | NA |
| CR33491 | 0 | NA | NA | NA | NA | NA |
| Sdic1 | 0 | NA | NA | NA | NA | NA |
| Phf7 | 0 | NA | NA | NA | NA | NA |
| CG9777 | 7.88318 | 3.79196 | 3.882872 | 0.976586 | 0.328774 | NA |
| CG9782 | 0.536437 | -1.71175 | 4.038042 | -0.42391 | 0.671635 | NA |
| CG9903 | 0 | NA | NA | NA | NA | NA |
| CG10822 | 0 | NA | NA | NA | NA | NA |
| CG12158 | 21.13696 | 6.666106 | 3.898853 | 1.709761 | 0.08731 | NA |
| CG12446 | 13.61099 | 1.739652 | 3.954685 | 0.439896 | 0.660012 | NA |
| CG13443 | 0 | NA | NA | NA | NA | NA |
| Br140 | 47.92664 | -5.00969 | 3.781911 | -1.32464 | 0.185289 | NA |
| CG1850 | 0 | NA | NA | NA | NA | NA |
| snsl | 0 | NA | NA | NA | NA | NA |
| Marcal1 | 30.99118 | 1.748008 | 3.952581 | 0.442245 | 0.658312 | NA |
| CG8654 | 0 | NA | NA | NA | NA | NA |
| Jon25Bii | 0 | NA | NA | NA | NA | NA |
| Jon25Biii | 28.00047 | -1.91823 | 3.952298 | -0.48535 | 0.627431 | NA |
| U3-55K | 0 | NA | NA | NA | NA | NA |
| ppk13 | 0 | NA | NA | NA | NA | NA |
| CG33509 | 0.356107 | -1.47099 | 4.037764 | -0.36431 | 0.715628 | NA |
| CG33510 | 0.178053 | -1.16467 | 4.075067 | -0.2858 | 0.775028 | NA |
| CG33511 | 0 | NA | NA | NA | NA | NA |
| CG33514 | 0 | NA | NA | NA | NA | NA |
| Unc-89 | 10.61662 | 2.412086 | 3.927885 | 0.614093 | 0.539154 | NA |

| | | | | | | |
|---|---|---|---|---|---|---|
| Acp53C14( | 0 | NA | NA | NA | NA | NA |
| lectin-37D | 56.69724 | -8.2434 | 3.885137 | -2.12178 | 0.033856 | NA |
| upd3 | 0 | NA | NA | NA | NA | NA |
| tRNA:Arg-˜ | 0 | NA | NA | NA | NA | NA |
| snRNA:U7 | 5.018952 | -1.89674 | 3.958511 | -0.47915 | 0.631829 | NA |
| tRNA:Ala-/ | 0 | NA | NA | NA | NA | NA |
| tRNA:Glu-˜ | 0.53416 | -1.53481 | 4.012395 | -0.38252 | 0.702078 | NA |
| tRNA:Leu-˜ | 0 | NA | NA | NA | NA | NA |
| tRNA:Asn-( | 0 | NA | NA | NA | NA | NA |
| tRNA:Arg-( | 0.1047 | 0.613261 | 4.062799 | 0.150945 | 0.880019 | NA |
| tRNA:Trp-( | 1.361099 | 1.628413 | 3.98439 | 0.408698 | 0.682761 | NA |
| SmydA-8 | 0 | NA | NA | NA | NA | NA |
| CG33552 | 0 | NA | NA | NA | NA | NA |
| CG14300 | 0 | NA | NA | NA | NA | NA |
| CG13376 | 0 | NA | NA | NA | NA | NA |
| CG33557 | 0 | NA | NA | NA | NA | NA |
| eyg | 0.411147 | -0.29058 | 4.046973 | -0.0718 | 0.94276 | NA |
| osk | 0 | NA | NA | NA | NA | NA |
| MstProx | 2.90234 | 0.112359 | 3.880728 | 0.028953 | 0.976902 | NA |
| armi | 2.168104 | -3.94426 | 3.966742 | -0.99433 | 0.320061 | NA |
| mud | 0 | NA | NA | NA | NA | NA |
| oc | 57.59648 | -2.15744 | 3.189781 | -0.67636 | 0.498812 | NA |
| AQP | 22.58384 | 1.661836 | 3.953052 | 0.420393 | 0.674198 | NA |
| polo | 0 | NA | NA | NA | NA | NA |
| l(3)mbn | 0 | NA | NA | NA | NA | NA |
| Gr47a | 0 | NA | NA | NA | NA | NA |
| tll | 0 | NA | NA | NA | NA | NA |
| unc | 0.492153 | 0.308117 | 3.993933 | 0.077146 | 0.938507 | NA |
| lectin-30A | 0 | NA | NA | NA | NA | NA |
| jhamt | 0 | NA | NA | NA | NA | NA |
| bw | 0 | NA | NA | NA | NA | NA |
| dj | 0 | NA | NA | NA | NA | NA |
| Dec-01 | 0 | NA | NA | NA | NA | NA |
| lectin-21C( | 0 | NA | NA | NA | NA | NA |
| CG30048 | 10.65817 | -4.28773 | 3.889354 | -1.10243 | 0.270276 | NA |
| CG31266 | 0 | NA | NA | NA | NA | NA |
| Muc12Ea | 0 | NA | NA | NA | NA | NA |
| gsb | 0 | NA | NA | NA | NA | NA |
| pug | 12.4153 | -1.91223 | 3.954028 | -0.48362 | 0.628659 | NA |
| CG14683 | 0 | NA | NA | NA | NA | NA |
| FucTB | 43.84404 | -1.04519 | 3.779126 | -0.27657 | 0.78211 | NA |
| Mp20 | 0 | NA | NA | NA | NA | NA |
| Doc2 | 0 | NA | NA | NA | NA | NA |
| thr | 0.233093 | 0.692656 | 4.076246 | 0.169925 | 0.865069 | NA |
| CG6614 | 0.2094 | 1.005034 | 4.045341 | 0.248442 | 0.803792 | NA |
| prd | 0.282677 | -0.09599 | 4.057934 | -0.02366 | 0.981127 | NA |
| up | 18.89962 | -7.35467 | 3.911 | -1.88051 | 0.060039 | NA |
| en | 0.282753 | -0.25398 | 4.020204 | -0.06318 | 0.949626 | NA |
| PK1-R | 0.356107 | -1.47099 | 4.037764 | -0.36431 | 0.715628 | NA |
| Gr22b | 0 | NA | NA | NA | NA | NA |

| | | | | | | |
|---|---|---|---|---|---|---|
| CG15109 | 0 | NA | NA | NA | NA | NA |
| CG17681 | 0 | NA | NA | NA | NA | NA |
| ppk7 | 0.357625 | -1.47707 | 4.044794 | -0.36518 | 0.714979 | NA |
| CG6299 | 0 | NA | NA | NA | NA | NA |
| CapaR | 0.1047 | 0.613261 | 4.062799 | 0.150945 | 0.880019 | NA |
| CG15117 | 3.188386 | -2.55369 | 3.944197 | -0.64746 | 0.517337 | NA |
| ppk23 | 0 | NA | NA | NA | NA | NA |
| ppk21 | 0.178053 | -1.16467 | 4.075067 | -0.2858 | 0.775028 | NA |
| ppk20 | 0.178812 | -1.08808 | 4.061735 | -0.26789 | 0.788788 | NA |
| lncRNA:iak | 0.178053 | -1.16467 | 4.075067 | -0.2858 | 0.775028 | NA |
| tRNA:Val-( | 0 | NA | NA | NA | NA | NA |
| Prx5 | 0 | NA | NA | NA | NA | NA |
| CG7215 | 0 | NA | NA | NA | NA | NA |
| CG11889 | 0 | NA | NA | NA | NA | NA |
| CG11891 | 0 | NA | NA | NA | NA | NA |
| CG11714 | 0 | NA | NA | NA | NA | NA |
| Shawn | 0 | NA | NA | NA | NA | NA |
| Tyler | 0 | NA | NA | NA | NA | NA |
| CG33155 | 40.95183 | -0.39539 | 3.583194 | -0.11035 | 0.912134 | NA |
| mRpL53 | 0 | NA | NA | NA | NA | NA |
| CG33095 | 0 | NA | NA | NA | NA | NA |
| CG33626 | 0 | NA | NA | NA | NA | NA |
| CG33627 | 0 | NA | NA | NA | NA | NA |
| snoRNA:U: | 0.4188 | 1.41068 | 4.034613 | 0.349644 | 0.726606 | NA |
| CG33630 | 0 | NA | NA | NA | NA | NA |
| CG33631 | 0 | NA | NA | NA | NA | NA |
| Vajk2 | 0 | NA | NA | NA | NA | NA |
| snoRNA:M | 0 | NA | NA | NA | NA | NA |
| snoRNA:2! | 3.476414 | 2.98224 | 3.925791 | 0.759653 | 0.447462 | NA |
| CheA7a | 0 | NA | NA | NA | NA | NA |
| CG33644 | 46.75667 | -0.65325 | 3.834813 | -0.17035 | 0.864737 | NA |
| CG33645 | 2.670799 | -1.66648 | 3.964795 | -0.42032 | 0.674251 | NA |
| CG33647 | 0 | NA | NA | NA | NA | NA |
| CG33648 | 0 | NA | NA | NA | NA | NA |
| GatC | 0 | NA | NA | NA | NA | NA |
| PolG2 | 0 | NA | NA | NA | NA | NA |
| CG12849 | 0 | NA | NA | NA | NA | NA |
| CG14929 | 0.1047 | 0.613261 | 4.062799 | 0.150945 | 0.880019 | NA |
| CG33654 | 0 | NA | NA | NA | NA | NA |
| snmRNA:2 | 0 | NA | NA | NA | NA | NA |
| Hsp22 | 0 | NA | NA | NA | NA | NA |
| CG4456 | 0 | NA | NA | NA | NA | NA |
| snmRNA:3 | 0 | NA | NA | NA | NA | NA |
| snmRNA:4 | 0 | NA | NA | NA | NA | NA |
| snoRNA:U: | 6.474162 | -0.67346 | 3.787004 | -0.17783 | 0.858854 | NA |
| snoRNA:O | 0 | NA | NA | NA | NA | NA |
| CG33664 | 1.165466 | 1.076041 | 3.987865 | 0.269829 | 0.787292 | NA |
| CG33665 | 0 | NA | NA | NA | NA | NA |
| CG33666 | 1.165466 | 1.076041 | 3.987865 | 0.269829 | 0.787292 | NA |
| CG33667 | 0 | NA | NA | NA | NA | NA |

| | | | | | | |
|---|---|---|---|---|---|---|
| CG33668 | 0.178812 | -1.08808 | 4.061735 | -0.26789 | 0.788788 | NA |
| CG33669 | 0 | NA | NA | NA | NA | NA |
| Phax | 30.16797 | 2.730608 | 3.792957 | 0.719915 | 0.471577 | NA |
| stg1 | 0 | NA | NA | NA | NA | NA |
| CG11566 | 0 | NA | NA | NA | NA | NA |
| CG33671 | 0 | NA | NA | NA | NA | NA |
| CG33672 | 0 | NA | NA | NA | NA | NA |
| CG33673 | 0 | NA | NA | NA | NA | NA |
| snoRNA:M | 4.141397 | 0.891021 | 3.960538 | 0.224975 | 0.821999 | NA |
| CG14246 | 0 | NA | NA | NA | NA | NA |
| CG14245 | 0 | NA | NA | NA | NA | NA |
| LSm-4 | 0 | NA | NA | NA | NA | NA |
| CG17768 | 0 | NA | NA | NA | NA | NA |
| CG15892 | 9.090636 | 1.165026 | 3.956038 | 0.294493 | 0.768381 | NA |
| srp | 0 | NA | NA | NA | NA | NA |
| CG33680 | 0 | NA | NA | NA | NA | NA |
| E(spl)m6-B | 0 | NA | NA | NA | NA | NA |
| lab | 12.87448 | -2.25322 | 3.958224 | -0.56925 | 0.569186 | NA |
| snmRNA:6 | 0 | NA | NA | NA | NA | NA |
| CG18538 | 53.45653 | 0.445875 | 3.851006 | 0.115781 | 0.907826 | NA |
| CG18539 | 11.75377 | 0.698648 | 3.928498 | 0.177841 | 0.858848 | NA |
| CG33483 | 0 | NA | NA | NA | NA | NA |
| snoRNA:22 | 14.74165 | -0.14721 | 3.802205 | -0.03872 | 0.969116 | NA |
| sea | 29.51656 | 0.374199 | 3.83588 | 0.097552 | 0.922288 | NA |
| CheB38c | 0 | NA | NA | NA | NA | NA |
| snmRNA:6 | 0 | NA | NA | NA | NA | NA |
| CG31016 | 0.178053 | -1.16467 | 4.075067 | -0.2858 | 0.775028 | NA |
| CG33687 | 0 | NA | NA | NA | NA | NA |
| CG33688 | 0 | NA | NA | NA | NA | NA |
| CG33689 | 0 | NA | NA | NA | NA | NA |
| CG33690 | 0 | NA | NA | NA | NA | NA |
| snoRNA:Ps | 0.178053 | -1.16467 | 4.075067 | -0.2858 | 0.775028 | NA |
| nompB | 0 | NA | NA | NA | NA | NA |
| snmRNA:4 | 0 | NA | NA | NA | NA | NA |
| CheA75a | 0 | NA | NA | NA | NA | NA |
| CG33700 | 0 | NA | NA | NA | NA | NA |
| CR33701 | 0 | NA | NA | NA | NA | NA |
| CG33702 | 0 | NA | NA | NA | NA | NA |
| CG33703 | 0 | NA | NA | NA | NA | NA |
| CG33704 | 0 | NA | NA | NA | NA | NA |
| snmRNA:2 | 3.177886 | -3.37317 | 3.879402 | -0.86951 | 0.384569 | NA |
| CG10332 | 0 | NA | NA | NA | NA | NA |
| IM18 | 0 | NA | NA | NA | NA | NA |
| snmRNA:1 | 0 | NA | NA | NA | NA | NA |
| alpha4GT2 | 15.07671 | 2.952152 | 3.92424 | 0.752286 | 0.451879 | NA |
| snmRNA:7 | 0 | NA | NA | NA | NA | NA |
| snmRNA:3 | 0 | NA | NA | NA | NA | NA |
| anox | 0 | NA | NA | NA | NA | NA |
| SLIRP1 | 0 | NA | NA | NA | NA | NA |
| scaRNA:M | 10.33588 | 0.019886 | 3.743965 | 0.005311 | 0.995762 | NA |

| | | | | | | |
|---|---|---|---|---|---|---|
| PGRP-LD | 10.67157 | 1.527223 | 3.954255 | 0.386223 | 0.699332 | NA |
| Pmi | 0 | NA | NA | NA | NA | NA |
| Pif1B | 0 | NA | NA | NA | NA | NA |
| CG18749 | 0 | NA | NA | NA | NA | NA |
| CG33722 | 0 | NA | NA | NA | NA | NA |
| CG33723 | 0 | NA | NA | NA | NA | NA |
| CG33725 | 0 | NA | NA | NA | NA | NA |
| scaRNA:M | 0 | NA | NA | NA | NA | NA |
| snmRNA:4 | 0 | NA | NA | NA | NA | NA |
| snmRNA:4 | 0 | NA | NA | NA | NA | NA |
| snmRNA:4 | 0 | NA | NA | NA | NA | NA |
| snmRNA:4 | 0 | NA | NA | NA | NA | NA |
| snmRNA:4 | 0 | NA | NA | NA | NA | NA |
| snmRNA:4 | 0 | NA | NA | NA | NA | NA |
| snmRNA:4 | 0 | NA | NA | NA | NA | NA |
| snmRNA:4 | 0 | NA | NA | NA | NA | NA |
| snmRNA:4 | 0 | NA | NA | NA | NA | NA |
| snmRNA:4 | 0 | NA | NA | NA | NA | NA |
| snmRNA:4 | 0 | NA | NA | NA | NA | NA |
| snmRNA:4 | 0 | NA | NA | NA | NA | NA |
| snmRNA:4 | 0 | NA | NA | NA | NA | NA |
| snmRNA:4 | 0 | NA | NA | NA | NA | NA |
| snmRNA:4 | 0 | NA | NA | NA | NA | NA |
| snmRNA:4 | 0 | NA | NA | NA | NA | NA |
| snmRNA:4 | 0 | NA | NA | NA | NA | NA |
| snmRNA:4 | 0 | NA | NA | NA | NA | NA |
| snmRNA:4 | 0 | NA | NA | NA | NA | NA |
| snmRNA:4 | 0 | NA | NA | NA | NA | NA |
| primo-2 | 0 | NA | NA | NA | NA | NA |
| primo-1 | 0 | NA | NA | NA | NA | NA |
| CG13561 | 0 | NA | NA | NA | NA | NA |
| snmRNA:1 | 0 | NA | NA | NA | NA | NA |
| CG32269 | 0 | NA | NA | NA | NA | NA |
| CG32270 | 0 | NA | NA | NA | NA | NA |
| CG13430 | 0 | NA | NA | NA | NA | NA |
| BORCS7 | 0 | NA | NA | NA | NA | NA |
| CG33752 | 0 | NA | NA | NA | NA | NA |
| CG33137 | 0 | NA | NA | NA | NA | NA |
| CheB93b | 7.656291 | -1.69306 | 3.95587 | -0.42799 | 0.66866 | NA |
| Gr64d | 0 | NA | NA | NA | NA | NA |
| Gr64e | 0 | NA | NA | NA | NA | NA |
| snoRNA:Ps | 2.301712 | 1.486073 | 3.965794 | 0.374723 | 0.707867 | NA |
| Fum2 | 1.337962 | -2.21334 | 3.917153 | -0.56504 | 0.572048 | NA |
| gdl-ORF39 | 0 | NA | NA | NA | NA | NA |
| gdl | 3.559184 | 4.427014 | 3.941475 | 1.123187 | 0.261358 | NA |
| CG33757 | 0.1047 | 0.613261 | 4.062799 | 0.150945 | 0.880019 | NA |
| CG33758 | 0.104623 | 0.639075 | 4.067674 | 0.157111 | 0.875158 | NA |
| CG33764 | 0.356107 | -1.47099 | 4.037764 | -0.36431 | 0.715628 | NA |
| CG33766 | 4.187997 | 1.708209 | 3.962778 | 0.431063 | 0.666422 | NA |
| CG33767 | 0 | NA | NA | NA | NA | NA |

| | | | | | | |
|---|---|---|---|---|---|---|
| CG33768 | 0 | NA | NA | NA | NA | NA |
| CG33769 | 0 | NA | NA | NA | NA | NA |
| CG33770 | 0.104623 | 0.639075 | 4.067674 | 0.157111 | 0.875158 | NA |
| CG33771 | 0 | NA | NA | NA | NA | NA |
| tRNA:CR32 | 0 | NA | NA | NA | NA | NA |
| CG30056 | 0 | NA | NA | NA | NA | NA |
| snoRNA:M | 0.132078 | -1.13211 | 4.069918 | -0.27817 | 0.780885 | NA |
| snoRNA:66 | 0 | NA | NA | NA | NA | NA |
| CG33783 | 0 | NA | NA | NA | NA | NA |
| CG33784 | 0 | NA | NA | NA | NA | NA |
| CG33785 | 0 | NA | NA | NA | NA | NA |
| CG33786 | 0 | NA | NA | NA | NA | NA |
| Or98b | 0 | NA | NA | NA | NA | NA |
| ste24c | 0 | NA | NA | NA | NA | NA |
| CG30461 | 0 | NA | NA | NA | NA | NA |
| CG9130 | 0 | NA | NA | NA | NA | NA |
| CG9133 | 0 | NA | NA | NA | NA | NA |
| snRNA:U2: | 0 | NA | NA | NA | NA | NA |
| CG3588 | 0.236778 | -0.26946 | 4.071573 | -0.06618 | 0.947234 | NA |
| snmRNA:7 | 0 | NA | NA | NA | NA | NA |
| Kaz1-ORFB | 14.44859 | 1.740504 | 3.954469 | 0.440136 | 0.659839 | NA |
| Kaz1-ORFA | 0 | NA | NA | NA | NA | NA |
| CG18170 | 0 | NA | NA | NA | NA | NA |
| CG33791 | 0 | NA | NA | NA | NA | NA |
| CG33792 | 48.96465 | -1.70586 | 3.951691 | -0.43168 | 0.665975 | NA |
| snmRNA:8 | 0 | NA | NA | NA | NA | NA |
| snoRNA:68 | 0 | NA | NA | NA | NA | NA |
| CG13175 | 0 | NA | NA | NA | NA | NA |
| CG33795 | 0 | NA | NA | NA | NA | NA |
| CG33796 | 0 | NA | NA | NA | NA | NA |
| CG14518 | 0 | NA | NA | NA | NA | NA |
| CG33463 | 0 | NA | NA | NA | NA | NA |
| Argl | 3.131606 | -1.39628 | 3.960959 | -0.35251 | 0.724455 | NA |
| CR33797 | 0 | NA | NA | NA | NA | NA |
| Adhr | 0 | NA | NA | NA | NA | NA |
| CG14492 | 0 | NA | NA | NA | NA | NA |
| lncRNA:CR | 18.41437 | 1.172608 | 3.953457 | 0.296603 | 0.766769 | NA |
| His1:CG33 | 2.093079 | 3.901995 | 3.954223 | 0.986792 | 0.323745 | NA |
| His-Psi:CR3 | 0 | NA | NA | NA | NA | NA |
| His3:CG33 | 0 | NA | NA | NA | NA | NA |
| His1:CG31 | 0 | NA | NA | NA | NA | NA |
| His2A:CG3 | 0 | NA | NA | NA | NA | NA |
| His3:CG31 | 0 | NA | NA | NA | NA | NA |
| His1:CG33 | 0 | NA | NA | NA | NA | NA |
| His-Psi:CR3 | 0 | NA | NA | NA | NA | NA |
| His3:CG33 | 0 | NA | NA | NA | NA | NA |
| His1:CG33 | 0 | NA | NA | NA | NA | NA |
| His2A:CG3 | 0.356866 | -1.92326 | 4.037889 | -0.4763 | 0.633858 | NA |
| His3:CG33 | 0 | NA | NA | NA | NA | NA |
| His1:CG33 | 0 | NA | NA | NA | NA | NA |

| | | | | | | |
|---|---|---|---|---|---|---|
| His-Psi:CR3 | 0 | NA | NA | NA | NA | NA |
| His3:CG33 | 0 | NA | NA | NA | NA | NA |
| His1:CG33 | 0 | NA | NA | NA | NA | NA |
| His2A:CG3 | 0 | NA | NA | NA | NA | NA |
| His3:CG33 | 0 | NA | NA | NA | NA | NA |
| His1:CG33 | 0 | NA | NA | NA | NA | NA |
| His2A:CG3 | 0 | NA | NA | NA | NA | NA |
| His3:CG33 | 0 | NA | NA | NA | NA | NA |
| His1:CG33 | 0 | NA | NA | NA | NA | NA |
| His2A:CG3 | 0 | NA | NA | NA | NA | NA |
| His3:CG33 | 0 | NA | NA | NA | NA | NA |
| His1:CG33 | 0 | NA | NA | NA | NA | NA |
| His2A:CG3 | 0 | NA | NA | NA | NA | NA |
| His3:CG33 | 0 | NA | NA | NA | NA | NA |
| His1:CG33 | 0 | NA | NA | NA | NA | NA |
| His2A:CG3 | 0 | NA | NA | NA | NA | NA |
| His3:CG33 | 0 | NA | NA | NA | NA | NA |
| His1:CG33 | 0 | NA | NA | NA | NA | NA |
| His2A:CG3 | 0 | NA | NA | NA | NA | NA |
| His3:CG33 | 0 | NA | NA | NA | NA | NA |
| His1:CG33 | 0 | NA | NA | NA | NA | NA |
| His2A:CG3 | 0 | NA | NA | NA | NA | NA |
| His3:CG33 | 0 | NA | NA | NA | NA | NA |
| His1:CG33 | 0 | NA | NA | NA | NA | NA |
| His2A:CG3 | 0 | NA | NA | NA | NA | NA |
| His3:CG33 | 0 | NA | NA | NA | NA | NA |
| His1:CG33 | 0 | NA | NA | NA | NA | NA |
| His2A:CG3 | 0 | NA | NA | NA | NA | NA |
| His3:CG33 | 0 | NA | NA | NA | NA | NA |
| His1:CG33 | 0 | NA | NA | NA | NA | NA |
| His2A:CG3 | 0 | NA | NA | NA | NA | NA |
| His3:CG33 | 0 | NA | NA | NA | NA | NA |
| His1:CG33 | 0 | NA | NA | NA | NA | NA |
| His2A:CG3 | 0 | NA | NA | NA | NA | NA |
| His3:CG33 | 0 | NA | NA | NA | NA | NA |
| His1:CG33 | 0 | NA | NA | NA | NA | NA |
| His2A:CG3 | 0 | NA | NA | NA | NA | NA |
| His3:CG33 | 0 | NA | NA | NA | NA | NA |
| His1:CG33 | 0 | NA | NA | NA | NA | NA |
| His2A:CG3 | 0 | NA | NA | NA | NA | NA |
| His3:CG33 | 0 | NA | NA | NA | NA | NA |
| His1:CG33 | 0 | NA | NA | NA | NA | NA |
| His2A:CG3 | 0 | NA | NA | NA | NA | NA |
| His3:CG33 | 0 | NA | NA | NA | NA | NA |
| His1:CG33 | 0 | NA | NA | NA | NA | NA |
| His2A:CG3 | 0 | NA | NA | NA | NA | NA |
| His3:CG33 | 0 | NA | NA | NA | NA | NA |
| His1:CG33 | 0 | NA | NA | NA | NA | NA |
| His2A:CG3 | 0 | NA | NA | NA | NA | NA |
| His3:CG33 | 0 | NA | NA | NA | NA | NA |

| | | | | | | |
|---|---|---|---|---|---|---|
| His1:CG33 | 0 | NA | NA | NA | NA | NA |
| His2A:CG3 | 0 | NA | NA | NA | NA | NA |
| His3:CG33 | 0 | NA | NA | NA | NA | NA |
| His1:CG33 | 0 | NA | NA | NA | NA | NA |
| His2A:CG3 | 0 | NA | NA | NA | NA | NA |
| His3:CG33 | 0 | NA | NA | NA | NA | NA |
| His-Psi:CR3 | 0 | NA | NA | NA | NA | NA |
| His-Psi:CR3 | 7.529166 | 1.263047 | 3.766048 | 0.335377 | 0.73734 | NA |
| His2B:CG3 | 7.761965 | -3.16534 | 3.85797 | -0.82047 | 0.411949 | NA |
| His4:CG33 | 0 | NA | NA | NA | NA | NA |
| His2B:CG3 | 0 | NA | NA | NA | NA | NA |
| His4:CG33 | 0 | NA | NA | NA | NA | NA |
| His2B:CG3 | 0 | NA | NA | NA | NA | NA |
| His4:CG33 | 0 | NA | NA | NA | NA | NA |
| His2B:CG3 | 0 | NA | NA | NA | NA | NA |
| His4:CG33 | 0 | NA | NA | NA | NA | NA |
| His2B:CG3 | 0 | NA | NA | NA | NA | NA |
| His4:CG33 | 0 | NA | NA | NA | NA | NA |
| His2B:CG3 | 0 | NA | NA | NA | NA | NA |
| His4:CG33 | 0 | NA | NA | NA | NA | NA |
| His2B:CG3 | 0 | NA | NA | NA | NA | NA |
| His4:CG33 | 0 | NA | NA | NA | NA | NA |
| His2B:CG3 | 0 | NA | NA | NA | NA | NA |
| His4:CG33 | 0 | NA | NA | NA | NA | NA |
| His2B:CG3 | 0 | NA | NA | NA | NA | NA |
| His4:CG33 | 0 | NA | NA | NA | NA | NA |
| His2B:CG3 | 0 | NA | NA | NA | NA | NA |
| His4:CG33 | 0 | NA | NA | NA | NA | NA |
| His2B:CG3 | 0 | NA | NA | NA | NA | NA |
| His4:CG33 | 0 | NA | NA | NA | NA | NA |
| His2B:CG3 | 0 | NA | NA | NA | NA | NA |
| His4:CG33 | 0 | NA | NA | NA | NA | NA |
| His2B:CG3 | 0 | NA | NA | NA | NA | NA |
| His4:CG33 | 0 | NA | NA | NA | NA | NA |
| His2B:CG3 | 0 | NA | NA | NA | NA | NA |
| His4:CG33 | 0 | NA | NA | NA | NA | NA |
| His2B:CG3 | 0 | NA | NA | NA | NA | NA |
| His4:CG33 | 0 | NA | NA | NA | NA | NA |
| His2B:CG3 | 0 | NA | NA | NA | NA | NA |
| His4:CG33 | 0 | NA | NA | NA | NA | NA |
| His2B:CG3 | 0 | NA | NA | NA | NA | NA |
| His4:CG33 | 0 | NA | NA | NA | NA | NA |
| His2B:CG3 | 0 | NA | NA | NA | NA | NA |
| His4:CG33 | 0 | NA | NA | NA | NA | NA |
| His2B:CG3 | 0 | NA | NA | NA | NA | NA |
| His4:CG33 | 0 | NA | NA | NA | NA | NA |
| His2B:CG3 | 0 | NA | NA | NA | NA | NA |
| His4:CG33 | 0 | NA | NA | NA | NA | NA |
| His2B:CG3 | 0 | NA | NA | NA | NA | NA |
| His4:CG31 | 0 | NA | NA | NA | NA | NA |

| | | | | | | |
|---|---|---|---|---|---|---|
| His2B:CG1 | 0 | NA | NA | NA | NA | NA |
| His4:CG33 | 0 | NA | NA | NA | NA | NA |
| His2B:CG3 | 0 | NA | NA | NA | NA | NA |
| seq | 0 | NA | NA | NA | NA | NA |
| CG33912 | 0 | NA | NA | NA | NA | NA |
| snmRNA:1 | 0 | NA | NA | NA | NA | NA |
| snmRNA:1 | 0 | NA | NA | NA | NA | NA |
| snmRNA:4 | 0 | NA | NA | NA | NA | NA |
| CG13203 | 0.2094 | 1.005034 | 4.045341 | 0.248442 | 0.803792 | NA |
| Tim9b | 0 | NA | NA | NA | NA | NA |
| Pstk | 0 | NA | NA | NA | NA | NA |
| waw | 0 | NA | NA | NA | NA | NA |
| Snapin | 0 | NA | NA | NA | NA | NA |
| HemK2 | 0 | NA | NA | NA | NA | NA |
| CG33476 | 0 | NA | NA | NA | NA | NA |
| CG33919 | 0 | NA | NA | NA | NA | NA |
| CG10096 | 0 | NA | NA | NA | NA | NA |
| CG10097 | 0 | NA | NA | NA | NA | NA |
| lncRNA:CR | 0.178812 | -1.08808 | 4.061735 | -0.26789 | 0.788788 | NA |
| CG33920 | 0 | NA | NA | NA | NA | NA |
| snoRNA:U3 | 3.894745 | -3.71361 | 3.877216 | -0.9578 | 0.338161 | NA |
| CG33922 | 0 | NA | NA | NA | NA | NA |
| CG33923 | 0 | NA | NA | NA | NA | NA |
| CG33926 | 0 | NA | NA | NA | NA | NA |
| CG33927 | 33.6958 | 4.558673 | 3.873744 | 1.176813 | 0.23927 | NA |
| CG33928 | 0 | NA | NA | NA | NA | NA |
| CheB38b | 0 | NA | NA | NA | NA | NA |
| CheB93a | 0 | NA | NA | NA | NA | NA |
| CR33929 | 0 | NA | NA | NA | NA | NA |
| snoRNA:18 | 7.158366 | 3.643568 | 3.84445 | 0.947748 | 0.343258 | NA |
| Rpp20 | 0 | NA | NA | NA | NA | NA |
| CG33932 | 0 | NA | NA | NA | NA | NA |
| Indy-2 | 0 | NA | NA | NA | NA | NA |
| CG33934 | 0 | NA | NA | NA | NA | NA |
| mei-218 | 0 | NA | NA | NA | NA | NA |
| mei-217 | 0 | NA | NA | NA | NA | NA |
| lncRNA:CR | 0 | NA | NA | NA | NA | NA |
| CG33939 | 8.557672 | -0.51066 | 3.950186 | -0.12928 | 0.89714 | NA |
| lncRNA:CR | 0 | NA | NA | NA | NA | NA |
| BG642312 | 0 | NA | NA | NA | NA | NA |
| CG33946 | 0 | NA | NA | NA | NA | NA |
| lncRNA:CR | 0 | NA | NA | NA | NA | NA |
| Menl-2 | 0 | NA | NA | NA | NA | NA |
| Menl-1 | 0 | NA | NA | NA | NA | NA |
| CR33629 | 0 | NA | NA | NA | NA | NA |
| CG33632 | 0.660388 | -1.75977 | 4.001266 | -0.4398 | 0.660079 | NA |
| CG33721 | 0 | NA | NA | NA | NA | NA |
| CG6503 | 0 | NA | NA | NA | NA | NA |
| ric8b | 0 | NA | NA | NA | NA | NA |
| Ance-3 | 0.593643 | -1.88241 | 3.976168 | -0.47342 | 0.635911 | NA |

| | | | | | | |
|---|---|---|---|---|---|---|
| CG7179 | 0.178053 | -1.16467 | 4.075067 | -0.2858 | 0.775028 | NA |
| CG14499 | 1.921676 | 3.569804 | 3.94628 | 0.9046 | 0.365677 | NA |
| CG32803 | 44.83741 | 8.292158 | 3.90907 | 2.121261 | 0.0339 | NA |
| CG15888 | 0 | NA | NA | NA | NA | NA |
| CG7025 | 0.460806 | -0.60052 | 3.997252 | -0.15023 | 0.880581 | NA |
| Cpr92A | 0 | NA | NA | NA | NA | NA |
| CG12184 | 4.689767 | -2.12618 | 3.87838 | -0.54821 | 0.583545 | NA |
| twi | 0 | NA | NA | NA | NA | NA |
| CG30280 | 6.651163 | 5.148522 | 3.922546 | 1.312546 | 0.189336 | NA |
| CG12078 | 0.639619 | -1.41861 | 3.980326 | -0.3564 | 0.721538 | NA |
| CG18636 | 0 | NA | NA | NA | NA | NA |
| CR15280 | 0 | NA | NA | NA | NA | NA |
| CecA2 | 0 | NA | NA | NA | NA | NA |
| CG6435 | 28.65416 | 4.501433 | 3.878719 | 1.160546 | 0.245826 | NA |
| E(spl)m5-H | 0 | NA | NA | NA | NA | NA |
| CG13062 | 0 | NA | NA | NA | NA | NA |
| CG13059 | 0 | NA | NA | NA | NA | NA |
| CG13039 | 0 | NA | NA | NA | NA | NA |
| Yp3 | 0.365171 | -0.43584 | 4.056359 | -0.10745 | 0.914434 | NA |
| CG15116 | 0 | NA | NA | NA | NA | NA |
| Cby | 0 | NA | NA | NA | NA | NA |
| Cp7Fa | 26.81579 | 3.055367 | 3.752685 | 0.814182 | 0.415541 | NA |
| lncRNA:CR | 0 | NA | NA | NA | NA | NA |
| CG33964 | 0 | NA | NA | NA | NA | NA |
| CG7255 | 0.178812 | -1.08808 | 4.061735 | -0.26789 | 0.788788 | NA |
| ovo | 9.65601 | 4.262966 | 3.896461 | 1.094061 | 0.273928 | NA |
| CG14400 | 0 | NA | NA | NA | NA | NA |
| CG33965 | 0.178053 | -1.16467 | 4.075067 | -0.2858 | 0.775028 | NA |
| CG33966 | 0 | NA | NA | NA | NA | NA |
| Cyp6a16 | 0 | NA | NA | NA | NA | NA |
| Anp | 31.70405 | -2.16135 | 3.953675 | -0.54667 | 0.584607 | NA |
| Mdr50 | 2.504861 | -0.27875 | 3.903025 | -0.07142 | 0.943065 | NA |
| Cp15 | 0 | NA | NA | NA | NA | NA |
| CG8927 | 0 | NA | NA | NA | NA | NA |
| CG13538 | 0.178053 | -1.16467 | 4.075067 | -0.2858 | 0.775028 | NA |
| glob3 | 0.178812 | -1.08808 | 4.061735 | -0.26789 | 0.788788 | NA |
| CG32857 | 0 | NA | NA | NA | NA | NA |
| bin | 0.209323 | 1.414133 | 4.032409 | 0.350692 | 0.725819 | NA |
| CG9313 | 3.061183 | -4.80925 | 3.930009 | -1.22372 | 0.221056 | NA |
| hh | 0.209323 | 1.414133 | 4.032409 | 0.350692 | 0.725819 | NA |
| CG14302 | 0 | NA | NA | NA | NA | NA |
| bsh | 0.3141 | 1.240997 | 4.03841 | 0.307299 | 0.758616 | NA |
| Obp50c | 0 | NA | NA | NA | NA | NA |
| Elo68alpha | 1.072874 | -2.0339 | 4.019685 | -0.50598 | 0.612867 | NA |
| CG13931 | 0 | NA | NA | NA | NA | NA |
| CG15357 | 0 | NA | NA | NA | NA | NA |
| NetA | 0.907513 | -0.58764 | 3.917565 | -0.15 | 0.880764 | NA |
| CG17195 | 0 | NA | NA | NA | NA | NA |
| CG17198 | 0 | NA | NA | NA | NA | NA |
| Cpr73D | 34.93154 | 1.667959 | 3.898276 | 0.427871 | 0.668745 | NA |

| | | | | | | |
|---|---|---|---|---|---|---|
| CG4956 | 0 | NA | NA | NA | NA | NA |
| CG31904 | 0 | NA | NA | NA | NA | NA |
| CG3355 | 1.584932 | -1.84571 | 3.973792 | -0.46447 | 0.642311 | NA |
| CG13319 | 52.97982 | 9.266427 | 3.844618 | 2.410233 | 0.015942 | NA |
| msd1 | 0 | NA | NA | NA | NA | NA |
| Taf12L | 0 | NA | NA | NA | NA | NA |
| asRNA:Eig( | 34.21758 | -1.64 | 3.951983 | -0.41498 | 0.678156 | NA |
| CG33309 | 0.341478 | 0.247543 | 4.031301 | 0.061405 | 0.951036 | NA |
| tRNA:Asn-( | 0 | NA | NA | NA | NA | NA |
| tRNA:Asn-( | 0.1047 | 0.613261 | 4.062799 | 0.150945 | 0.880019 | NA |
| Cyt-b5-r | 0.233093 | 0.692656 | 4.076246 | 0.169925 | 0.865069 | NA |
| Z600 | 0 | NA | NA | NA | NA | NA |
| CecA1 | 0 | NA | NA | NA | NA | NA |
| Or65c | 46.23827 | -0.87619 | 3.94859 | -0.2219 | 0.824393 | NA |
| mtSSB | 5.800391 | 0.33584 | 3.847088 | 0.087297 | 0.930435 | NA |
| lectin-28C | 0.31089 | -1.9003 | 4.033427 | -0.47114 | 0.637542 | NA |
| gl | 0.132078 | -1.13211 | 4.069918 | -0.27817 | 0.780885 | NA |
| CecB | 0 | NA | NA | NA | NA | NA |
| CG12256 | 0 | NA | NA | NA | NA | NA |
| CG31274 | 0.5892 | -0.44935 | 3.992199 | -0.11256 | 0.910381 | NA |
| CG14247 | 0.934208 | -0.58863 | 3.908901 | -0.15059 | 0.880302 | NA |
| Edg78E | 0.712213 | -1.57119 | 3.998613 | -0.39293 | 0.694369 | NA |
| CG17751 | 0 | NA | NA | NA | NA | NA |
| nAChRbeta | 0 | NA | NA | NA | NA | NA |
| CG13038 | 0 | NA | NA | NA | NA | NA |
| CG17377 | 0.233093 | 0.692656 | 4.076246 | 0.169925 | 0.865069 | NA |
| Fbp2 | 0 | NA | NA | NA | NA | NA |
| CG9459 | 0 | NA | NA | NA | NA | NA |
| CG30087 | 0 | NA | NA | NA | NA | NA |
| CG33460 | 0 | NA | NA | NA | NA | NA |
| CG33461 | 0 | NA | NA | NA | NA | NA |
| CG33462 | 0 | NA | NA | NA | NA | NA |
| Act57B | 0.314023 | 1.768229 | 4.015745 | 0.440324 | 0.659702 | NA |
| CG15403 | 0 | NA | NA | NA | NA | NA |
| CR9337 | 2.406335 | 1.488229 | 3.965178 | 0.375325 | 0.707419 | NA |
| Muc68D | 0.356107 | -1.47099 | 4.037764 | -0.36431 | 0.715628 | NA |
| fab1 | 0 | NA | NA | NA | NA | NA |
| obst-H | 0 | NA | NA | NA | NA | NA |
| CG33985 | 0.2094 | 1.005034 | 4.045341 | 0.248442 | 0.803792 | NA |
| CG33986 | 0.104623 | 0.639075 | 4.067674 | 0.157111 | 0.875158 | NA |
| run | 0 | NA | NA | NA | NA | NA |
| ACXC | 0.178053 | -1.16467 | 4.075067 | -0.2858 | 0.775028 | NA |
| lncRNA:CR | 0 | NA | NA | NA | NA | NA |
| Rpp21 | 57.01968 | 1.537151 | 3.951539 | 0.389001 | 0.697276 | NA |
| sunn | 0 | NA | NA | NA | NA | NA |
| Dso2 | 3.566098 | -1.88656 | 3.961501 | -0.47622 | 0.633916 | NA |
| CG18545 | 0 | NA | NA | NA | NA | NA |
| CG4174 | 2.198698 | 1.671135 | 3.972627 | 0.420662 | 0.674002 | NA |
| CG13380 | 0 | NA | NA | NA | NA | NA |
| Edg84A | 0 | NA | NA | NA | NA | NA |

| | | | | | | |
|---|---|---|---|---|---|---|
| phu | 3.661814 | 1.504942 | 3.960447 | 0.379993 | 0.703951 | NA |
| CG32023 | 0 | NA | NA | NA | NA | NA |
| Andorra | 0 | NA | NA | NA | NA | NA |
| CG32625 | 42.42716 | 3.203495 | 3.907146 | 0.819907 | 0.412269 | NA |
| CG31693 | 0 | NA | NA | NA | NA | NA |
| CG13578 | 9.632393 | 1.733649 | 3.956211 | 0.438209 | 0.661235 | NA |
| tgy | 0 | NA | NA | NA | NA | NA |
| CG7912 | 4.288459 | -4.46903 | 3.945299 | -1.13275 | 0.25732 | NA |
| Jon25Bi | 0 | NA | NA | NA | NA | NA |
| CG12620 | 0 | NA | NA | NA | NA | NA |
| CG14810 | 0 | NA | NA | NA | NA | NA |
| CG30383 | 0.760277 | 0.501824 | 3.968793 | 0.126442 | 0.899382 | NA |
| Vago | 0 | NA | NA | NA | NA | NA |
| eya | 0 | NA | NA | NA | NA | NA |
| CG5928 | 50.2355 | -3.52118 | 3.899441 | -0.903 | 0.366529 | NA |
| CG13044 | 0 | NA | NA | NA | NA | NA |
| yuri | 32.42012 | -6.23816 | 3.871193 | -1.61143 | 0.107086 | NA |
| CG33093 | 0.178053 | -1.16467 | 4.075067 | -0.2858 | 0.775028 | NA |
| CG33993 | 2.951685 | -3.33776 | 3.931437 | -0.84899 | 0.395886 | NA |
| CG15649 | 0 | NA | NA | NA | NA | NA |
| CG6262 | 0.357625 | -1.47707 | 4.044794 | -0.36518 | 0.714979 | NA |
| CG33301 | 0 | NA | NA | NA | NA | NA |
| CG12481 | 0 | NA | NA | NA | NA | NA |
| CG14630 | 0 | NA | NA | NA | NA | NA |
| CG33995 | 0 | NA | NA | NA | NA | NA |
| CG30098 | 0 | NA | NA | NA | NA | NA |
| CG4927 | 0 | NA | NA | NA | NA | NA |
| CG14062 | 0.264155 | -1.54099 | 4.048976 | -0.38059 | 0.703509 | NA |
| CG33679 | 2.097839 | 1.11873 | 3.972242 | 0.281637 | 0.778222 | NA |
| CG33258 | 0 | NA | NA | NA | NA | NA |
| CG33998 | 8.568481 | 0.201195 | 3.845387 | 0.052321 | 0.958273 | NA |
| CG34001 | 0 | NA | NA | NA | NA | NA |
| CG34002 | 18.61218 | 2.713012 | 3.90385 | 0.694958 | 0.487082 | NA |
| NimB3 | 18.0999 | 2.763288 | 3.919351 | 0.705037 | 0.480787 | NA |
| CG34005 | 0 | NA | NA | NA | NA | NA |
| CG33772 | 0.546832 | -1.46767 | 3.995602 | -0.36732 | 0.713379 | NA |
| CR33294 | 0.104623 | 0.639075 | 4.067674 | 0.157111 | 0.875158 | NA |
| CG33773 | 0 | NA | NA | NA | NA | NA |
| CG33775 | 0 | NA | NA | NA | NA | NA |
| CG33658 | 0 | NA | NA | NA | NA | NA |
| lncRNA:CR | 0 | NA | NA | NA | NA | NA |
| CG34007 | 0 | NA | NA | NA | NA | NA |
| CG34008 | 0.264155 | -1.54099 | 4.048976 | -0.38059 | 0.703509 | NA |
| CG34011 | 0.80738 | -2.35989 | 3.971695 | -0.59418 | 0.552393 | NA |
| CG34012 | 0.356107 | -1.47099 | 4.037764 | -0.36431 | 0.715628 | NA |
| CG34021 | 0 | NA | NA | NA | NA | NA |
| lncRNA:CR | 0 | NA | NA | NA | NA | NA |
| CG34026 | 18.85817 | -0.01081 | 3.905228 | -0.00277 | 0.997792 | NA |
| CG34027 | 0 | NA | NA | NA | NA | NA |
| CG34028 | 0 | NA | NA | NA | NA | NA |

| | | | | | | |
|---|---|---|---|---|---|---|
| CG34029 | 0 | NA | NA | NA | NA | NA |
| CG34030 | 0.132078 | -1.13211 | 4.069918 | -0.27817 | 0.780885 | NA |
| CG34031 | 0 | NA | NA | NA | NA | NA |
| CG34033 | 0 | NA | NA | NA | NA | NA |
| CG34034 | 0 | NA | NA | NA | NA | NA |
| CG33465 | 0 | NA | NA | NA | NA | NA |
| CG33798 | 0 | NA | NA | NA | NA | NA |
| CG34038 | 0 | NA | NA | NA | NA | NA |
| CG34040 | 0 | NA | NA | NA | NA | NA |
| CG34041 | 0 | NA | NA | NA | NA | NA |
| ppk10 | 30.21707 | -7.38542 | 3.849408 | -1.91859 | 0.055037 | NA |
| CG34043 | 0 | NA | NA | NA | NA | NA |
| CG34045 | 7.971396 | -3.1233 | 3.925024 | -0.79574 | 0.426183 | NA |
| lncRNA:CR | 0 | NA | NA | NA | NA | NA |
| lncRNA:CR | 0 | NA | NA | NA | NA | NA |
| CG34049 | 0 | NA | NA | NA | NA | NA |
| CG34051 | 2.082181 | -0.96846 | 3.93372 | -0.2462 | 0.805531 | NA |
| lncRNA:CR | 3.769197 | 1.70343 | 3.964029 | 0.429722 | 0.667398 | NA |
| CG34054 | 0 | NA | NA | NA | NA | NA |
| ppk16 | 0 | NA | NA | NA | NA | NA |
| ppk11 | 0 | NA | NA | NA | NA | NA |
| CG34056 | 13.88816 | -1.6998 | 3.95366 | -0.42993 | 0.667246 | NA |
| CG34057 | 0.132078 | -1.13211 | 4.069918 | -0.27817 | 0.780885 | NA |
| CG17140 | 0 | NA | NA | NA | NA | NA |
| CG17139 | 0 | NA | NA | NA | NA | NA |
| mt:tRNA:Il | 56.75501 | -0.46766 | 3.106641 | -0.15053 | 0.880343 | NA |
| mt:tRNA:G | 15.67504 | -0.47805 | 3.736175 | -0.12795 | 0.898188 | NA |
| mt:tRNA:M | 0.442416 | 2.217191 | 3.994664 | 0.555038 | 0.578869 | NA |
| mt:tRNA:T | 3.437871 | 1.622276 | 3.861099 | 0.420159 | 0.674369 | NA |
| mt:tRNA:T | 26.01033 | 0.003978 | 3.731625 | 0.001066 | 0.999149 | NA |
| mt:tRNA:L | 2.372315 | 0.281774 | 3.890111 | 0.072433 | 0.942257 | NA |
| mt:tRNA:L | 29.04862 | -0.02538 | 3.732135 | -0.0068 | 0.994574 | NA |
| mt:tRNA:A | 0.550724 | 1.292521 | 3.972655 | 0.325355 | 0.744913 | NA |
| mt:tRNA:A | 0 | NA | NA | NA | NA | NA |
| mt:tRNA:S | 11.31347 | -0.90891 | 3.748552 | -0.24247 | 0.808417 | NA |
| mt:tRNA:G | 0.341401 | 0.594712 | 4.003898 | 0.148533 | 0.881922 | NA |
| mt:tRNA:P | 2.666879 | -0.602 | 3.892713 | -0.15465 | 0.8771 | NA |
| mt:tRNA:H | 13.90779 | 0.399512 | 3.739452 | 0.106837 | 0.914918 | NA |
| mt:tRNA:T | 1.830721 | -1.23284 | 3.854615 | -0.31984 | 0.749093 | NA |
| mt:tRNA:P | 0.104623 | 0.639075 | 4.067674 | 0.157111 | 0.875158 | NA |
| mt:tRNA:S | 1.844744 | -1.64151 | 3.932701 | -0.4174 | 0.676386 | NA |
| mt:tRNA:L | 1.531116 | 2.649163 | 3.91345 | 0.676938 | 0.498445 | NA |
| mt:tRNA:V | 2.714169 | -1.89778 | 3.845588 | -0.4935 | 0.621663 | NA |
| CG17490 | 0 | NA | NA | NA | NA | NA |
| klhl10 | 0 | NA | NA | NA | NA | NA |
| CG40006 | 51.96363 | 1.257247 | 3.951476 | 0.318171 | 0.750355 | NA |
| CR40005 | 15.49635 | -0.43713 | 3.840737 | -0.11381 | 0.909385 | NA |
| CG17636 | 0 | NA | NA | NA | NA | NA |
| CG17707 | 51.43307 | -3.12903 | 3.765235 | -0.83103 | 0.405956 | NA |
| CG17169 | 0 | NA | NA | NA | NA | NA |

| | | | | | | |
|---|---|---|---|---|---|---|
| CG17167 | 45.7948 | -1.12672 | 3.86437 | -0.29157 | 0.770618 | NA |
| CG12061 | 0 | NA | NA | NA | NA | NA |
| CR12460 | 0 | NA | NA | NA | NA | NA |
| CG40198 | 0 | NA | NA | NA | NA | NA |
| CG40298 | 0.1047 | 0.613261 | 4.062799 | 0.150945 | 0.880019 | NA |
| Acp54A1 | 0 | NA | NA | NA | NA | NA |
| Mkp | 0.357625 | -1.47707 | 4.044794 | -0.36518 | 0.714979 | NA |
| CG14042 | 0 | NA | NA | NA | NA | NA |
| BG642163 | 0 | NA | NA | NA | NA | NA |
| RhoU | 13.2814 | -2.73934 | 3.923108 | -0.69826 | 0.485016 | NA |
| CG34105 | 0 | NA | NA | NA | NA | NA |
| CG34106 | 0 | NA | NA | NA | NA | NA |
| CG34107 | 0.466186 | 0.959471 | 4.033986 | 0.237847 | 0.812 | NA |
| CG34109 | 0.178053 | -1.16467 | 4.075067 | -0.2858 | 0.775028 | NA |
| lobo | 38.37439 | -2.10141 | 3.953086 | -0.53159 | 0.595012 | NA |
| CG34115 | 7.852495 | 1.729066 | 3.957382 | 0.436922 | 0.662168 | NA |
| CG34116 | 0 | NA | NA | NA | NA | NA |
| CG34124 | 1.889787 | -2.72002 | 3.946482 | -0.68923 | 0.490682 | NA |
| CG34129 | 0 | NA | NA | NA | NA | NA |
| CG34130 | 0.209247 | 1.042804 | 4.047837 | 0.25762 | 0.7967 | NA |
| Muted | 0 | NA | NA | NA | NA | NA |
| CG34135 | 0 | NA | NA | NA | NA | NA |
| CG34138 | 17.27549 | 1.742783 | 3.953893 | 0.440776 | 0.659375 | NA |
| CG34140 | 0 | NA | NA | NA | NA | NA |
| CG17672 | 2.241339 | -1.30622 | 3.964211 | -0.3295 | 0.741775 | NA |
| Ir10a | 0 | NA | NA | NA | NA | NA |
| RunxA | 2.529472 | -0.14319 | 3.897549 | -0.03674 | 0.970693 | NA |
| CG34148 | 9.949812 | 4.362198 | 3.829695 | 1.139046 | 0.254684 | NA |
| CG34149 | 37.19161 | -0.81193 | 3.849616 | -0.21091 | 0.832956 | NA |
| snRNA:U1: | 13.6686 | -4.28157 | 3.806563 | -1.12479 | 0.26068 | NA |
| snoRNA:M | 22.10774 | -3.75907 | 3.824501 | -0.98289 | 0.325661 | NA |
| snoRNA:M | 3.245698 | 1.695863 | 3.966021 | 0.427598 | 0.668944 | NA |
| stnA | 0 | NA | NA | NA | NA | NA |
| stnB | 0 | NA | NA | NA | NA | NA |
| CR40282 | 0 | NA | NA | NA | NA | NA |
| CG13177 | 0 | NA | NA | NA | NA | NA |
| asRNA:CR3 | 0 | NA | NA | NA | NA | NA |
| CG41434 | 0 | NA | NA | NA | NA | NA |
| CR41454 | 0 | NA | NA | NA | NA | NA |
| lncRNA:CR | 0 | NA | NA | NA | NA | NA |
| FucTC | 23.03865 | -3.06613 | 3.829362 | -0.80069 | 0.423311 | NA |
| lncRNA:CR | 0 | NA | NA | NA | NA | NA |
| CR41440 | 33.32061 | -2.84761 | 3.745428 | -0.76029 | 0.447081 | NA |
| lncRNA:CR | 0.574571 | 1.337403 | 3.994282 | 0.334829 | 0.737754 | NA |
| CG41106 | 0 | NA | NA | NA | NA | NA |
| snoRNA:Ps | 1.200397 | -2.8489 | 3.970745 | -0.71747 | 0.473083 | NA |
| CR12798 | 0 | NA | NA | NA | NA | NA |
| snoRNA:Ps | 0 | NA | NA | NA | NA | NA |
| snoRNA:Ps | 3.475545 | -4.83221 | 3.887653 | -1.24296 | 0.213881 | NA |
| Uhg7 | 0 | NA | NA | NA | NA | NA |

| | | | | | | |
|---|---|---|---|---|---|---|
| snoRNA:Or | 0 | NA | NA | NA | NA | NA |
| snoRNA:Ps | 25.17407 | 1.174623 | 3.952775 | 0.297164 | 0.766341 | NA |
| snoRNA:Ps | 0.715249 | -1.88064 | 4.034336 | -0.46616 | 0.641102 | NA |
| snoRNA:Ps | 0 | NA | NA | NA | NA | NA |
| ATbp | 11.85165 | 0.437701 | 3.763074 | 0.116315 | 0.907403 | NA |
| snoRNA:Ps | 0 | NA | NA | NA | NA | NA |
| snoRNA:M | 2.225846 | 2.347881 | 3.921221 | 0.598763 | 0.549331 | NA |
| CG13051 | 0 | NA | NA | NA | NA | NA |
| snoRNA:M | 0 | NA | NA | NA | NA | NA |
| snoRNA:Ps | 0 | NA | NA | NA | NA | NA |
| snoRNA:Ps | 0 | NA | NA | NA | NA | NA |
| snoRNA:Ps | 0 | NA | NA | NA | NA | NA |
| snoRNA:Ps | 22.10084 | 2.963994 | 3.7823 | 0.783649 | 0.433246 | NA |
| snoRNA:M | 0 | NA | NA | NA | NA | NA |
| snoRNA:Ps | 5.529851 | 1.269576 | 3.872372 | 0.327855 | 0.743021 | NA |
| snoRNA:Ps | 0 | NA | NA | NA | NA | NA |
| snoRNA:Ps | 0 | NA | NA | NA | NA | NA |
| CR40182 | 0 | NA | NA | NA | NA | NA |
| snoRNA:M | 0 | NA | NA | NA | NA | NA |
| snoRNA:Ps | 14.98477 | -1.01657 | 3.776312 | -0.2692 | 0.787779 | NA |
| snoRNA:Ps | 0 | NA | NA | NA | NA | NA |
| snoRNA:Ps | 0 | NA | NA | NA | NA | NA |
| snoRNA:Ps | 1.818025 | -0.95856 | 3.935933 | -0.24354 | 0.807587 | NA |
| snoRNA:Ps | 3.045501 | 1.793666 | 3.872031 | 0.463236 | 0.643195 | NA |
| snoRNA:Ps | 0 | NA | NA | NA | NA | NA |
| snoRNA:M | 2.63545 | -3.25466 | 3.981159 | -0.81752 | 0.413633 | NA |
| kl-2 | 5.039264 | 1.402987 | 3.944899 | 0.355646 | 0.722106 | NA |
| snoRNA:Ps | 1.884599 | 1.659029 | 3.975915 | 0.41727 | 0.676481 | NA |
| CG14742 | 0 | NA | NA | NA | NA | NA |
| snoRNA:Ps | 0.894061 | -2.00006 | 4.030294 | -0.49626 | 0.619714 | NA |
| snoRNA:Ps | 9.682531 | -0.19292 | 3.844715 | -0.05018 | 0.95998 | NA |
| snoRNA:Ps | 0 | NA | NA | NA | NA | NA |
| snoRNA:M | 4.81267 | 1.512878 | 3.958225 | 0.382211 | 0.702305 | NA |
| snoRNA:M | 0.715249 | -1.88064 | 4.034336 | -0.46616 | 0.641102 | NA |
| snoRNA:Ps | 1.674279 | 3.479487 | 3.968676 | 0.876737 | 0.380629 | NA |
| snoRNA:M | 5.667132 | -4.69551 | 3.945678 | -1.19004 | 0.234031 | NA |
| snoRNA:M | 0.2094 | 1.005034 | 4.045341 | 0.248442 | 0.803792 | NA |
| ORY | 0 | NA | NA | NA | NA | NA |
| snoRNA:Ps | 0.53416 | -1.53481 | 4.012395 | -0.38252 | 0.702078 | NA |
| asRNA:CR3 | 22.25839 | 0.406871 | 3.837539 | 0.106024 | 0.915563 | NA |
| snoRNA:Ps | 0 | NA | NA | NA | NA | NA |
| CR40461 | 8.525377 | -0.95146 | 3.747525 | -0.25389 | 0.79958 | NA |
| snoRNA:Or | 16.6351 | 1.531556 | 3.953064 | 0.387435 | 0.698434 | NA |
| snoRNA:Ps | 0 | NA | NA | NA | NA | NA |
| snoRNA:Ps | 0.178053 | -1.16467 | 4.075067 | -0.2858 | 0.775028 | NA |
| snoRNA:M | 0.535678 | -2.26616 | 4.021604 | -0.5635 | 0.573098 | NA |
| snoRNA:Ps | 0.356107 | -1.47099 | 4.037764 | -0.36431 | 0.715628 | NA |
| snoRNA:Ps | 0.53416 | -1.53481 | 4.012395 | -0.38252 | 0.702078 | NA |
| CR40068 | 0 | NA | NA | NA | NA | NA |
| CG17159 | 0 | NA | NA | NA | NA | NA |

| | | | | | | |
|---|---|---|---|---|---|---|
| snoRNA:M | 0 | NA | NA | NA | NA | NA |
| snoRNA:Ps | 0 | NA | NA | NA | NA | NA |
| snoRNA:Ps | 0 | NA | NA | NA | NA | NA |
| snoRNA:M | 3.062653 | -4.16238 | 3.889479 | -1.07016 | 0.284545 | NA |
| snoRNA:Ps | 0.2094 | 1.005034 | 4.045341 | 0.248442 | 0.803792 | NA |
| snoRNA:Ps | 0 | NA | NA | NA | NA | NA |
| snoRNA:M | 1.646575 | -1.83931 | 3.938458 | -0.46701 | 0.640491 | NA |
| snoRNA:Ps | 0 | NA | NA | NA | NA | NA |
| ARY | 0 | NA | NA | NA | NA | NA |
| CG14397 | 22.25356 | 0.643842 | 3.920878 | 0.164209 | 0.869567 | NA |
| snoRNA:Ps | 5.875758 | -1.68859 | 3.957348 | -0.4267 | 0.6696 | NA |
| snoRNA:Ps | 0 | NA | NA | NA | NA | NA |
| snoRNA:Ps | 0 | NA | NA | NA | NA | NA |
| snoRNA:Ps | 0 | NA | NA | NA | NA | NA |
| snoRNA:Ps | 0 | NA | NA | NA | NA | NA |
| snoRNA:Ps | 13.00586 | 3.176553 | 3.803685 | 0.835125 | 0.403647 | NA |
| snoRNA:Ps | 0 | NA | NA | NA | NA | NA |
| snoRNA:Ps | 7.510116 | -2.23383 | 3.963189 | -0.56365 | 0.572996 | NA |
| snoRNA:Oı | 6.877103 | -1.01854 | 3.952131 | -0.25772 | 0.796624 | NA |
| snoRNA:Ps | 0 | NA | NA | NA | NA | NA |
| CR14578 | 0 | NA | NA | NA | NA | NA |
| snoRNA:Ps | 5.184038 | -4.10985 | 3.943772 | -1.04211 | 0.29736 | NA |
| snoRNA:Ps | 0 | NA | NA | NA | NA | NA |
| snoRNA:Ps | 43.30106 | 1.145615 | 3.951975 | 0.289884 | 0.771905 | NA |
| snoRNA:M | 2.976774 | -2.12883 | 3.8972 | -0.54625 | 0.584897 | NA |
| snoRNA:Ps | 38.22798 | -0.70451 | 3.835459 | -0.18368 | 0.854261 | NA |
| snoRNA:M | 2.082464 | -1.43897 | 3.885873 | -0.37031 | 0.711153 | NA |
| snoRNA:M | 0.357625 | -1.47707 | 4.044794 | -0.36518 | 0.714979 | NA |
| snoRNA:Ps | 0 | NA | NA | NA | NA | NA |
| snoRNA:M | 0.1047 | 0.613261 | 4.062799 | 0.150945 | 0.880019 | NA |
| snoRNA:Ps | 0 | NA | NA | NA | NA | NA |
| snoRNA:Ps | 11.08636 | -2.24874 | 3.959363 | -0.56796 | 0.570065 | NA |
| CG11741 | 0 | NA | NA | NA | NA | NA |
| snoRNA:Ps | 1.984084 | -2.49864 | 3.931494 | -0.63554 | 0.525074 | NA |
| snoRNA:Ps | 0 | NA | NA | NA | NA | NA |
| snoRNA:Ps | 0 | NA | NA | NA | NA | NA |
| snoRNA:M | 0.233093 | 0.692656 | 4.076246 | 0.169925 | 0.865069 | NA |
| snoRNA:Ps | 0 | NA | NA | NA | NA | NA |
| snoRNA:Oı | 0 | NA | NA | NA | NA | NA |
| snoRNA:Ps | 0 | NA | NA | NA | NA | NA |
| snoRNA:Ps | 17.6203 | -0.11337 | 3.735348 | -0.03035 | 0.975788 | NA |
| snoRNA:M | 8.605654 | -1.26164 | 3.766497 | -0.33496 | 0.737653 | NA |
| snoRNA:Ps | 0.357625 | -1.47707 | 4.044794 | -0.36518 | 0.714979 | NA |
| Pp1-Y2 | 0 | NA | NA | NA | NA | NA |
| CG7606 | 0.1047 | 0.613261 | 4.062799 | 0.150945 | 0.880019 | NA |
| asRNA:CR3 | 0 | NA | NA | NA | NA | NA |
| CR40441 | 0 | NA | NA | NA | NA | NA |
| CG16824 | 0 | NA | NA | NA | NA | NA |
| snoRNA:Ps | 0.178053 | -1.16467 | 4.075067 | -0.2858 | 0.775028 | NA |
| snoRNA:Ps | 0 | NA | NA | NA | NA | NA |

| | | | | | | |
|---|---|---|---|---|---|---|
| snoRNA:Ps | 0.178812 | -1.08808 | 4.061735 | -0.26789 | 0.788788 | NA |
| snoRNA:Ps | 0 | NA | NA | NA | NA | NA |
| snoRNA:M | 0 | NA | NA | NA | NA | NA |
| snoRNA:Ps | 0 | NA | NA | NA | NA | NA |
| CG12828 | 0 | NA | NA | NA | NA | NA |
| snoRNA:M | 0 | NA | NA | NA | NA | NA |
| snoRNA:Ps | 0.1047 | 0.613261 | 4.062799 | 0.150945 | 0.880019 | NA |
| snoRNA:M | 0.3141 | 1.240997 | 4.03841 | 0.307299 | 0.758616 | NA |
| snoRNA:Ps | 0 | NA | NA | NA | NA | NA |
| snoRNA:Ps | 0 | NA | NA | NA | NA | NA |
| snoRNA:Ps | 0 | NA | NA | NA | NA | NA |
| snoRNA:Ps | 0.535678 | -2.26616 | 4.021604 | -0.5635 | 0.573098 | NA |
| snoRNA:Ps | 0 | NA | NA | NA | NA | NA |
| CG15140 | 0 | NA | NA | NA | NA | NA |
| Ppr-Y | 0 | NA | NA | NA | NA | NA |
| snoRNA:M | 1.165466 | 1.076041 | 3.987865 | 0.269829 | 0.787292 | NA |
| snoRNA:M | 4.649119 | -2.2071 | 3.970175 | -0.55592 | 0.578266 | NA |
| snoRNA:M | 2.106559 | -2.51934 | 3.952378 | -0.63742 | 0.523849 | NA |
| snoRNA:Or | 1.251686 | -2.06022 | 4.011654 | -0.51356 | 0.60756 | NA |
| snoRNA:M | 1.570499 | 1.642769 | 3.980387 | 0.412716 | 0.679815 | NA |
| snoRNA:Ps | 0 | NA | NA | NA | NA | NA |
| snoRNA:Ps | 0.715249 | -1.88064 | 4.034336 | -0.46616 | 0.641102 | NA |
| lncRNA:Yu | 0 | NA | NA | NA | NA | NA |
| snoRNA:Ps | 0 | NA | NA | NA | NA | NA |
| snoRNA:M | 0 | NA | NA | NA | NA | NA |
| snoRNA:Ps | 0 | NA | NA | NA | NA | NA |
| snoRNA:Ps | 0 | NA | NA | NA | NA | NA |
| snoRNA:Ps | 0 | NA | NA | NA | NA | NA |
| snoRNA:Ps | 0.178053 | -1.16467 | 4.075067 | -0.2858 | 0.775028 | NA |
| snoRNA:Ps | 0.132078 | -1.13211 | 4.069918 | -0.27817 | 0.780885 | NA |
| snoRNA:Ps | 0.396233 | -1.6814 | 4.028351 | -0.41739 | 0.676392 | NA |
| snoRNA:Ps | 0.178053 | -1.16467 | 4.075067 | -0.2858 | 0.775028 | NA |
| snoRNA:Ps | 7.848331 | 2.115625 | 3.936653 | 0.537417 | 0.59098 | NA |
| CR40100 | 7.339494 | -6.11453 | 3.91646 | -1.56124 | 0.118468 | NA |
| snoRNA:Ps | 3.218621 | -2.17835 | 3.977872 | -0.54762 | 0.583954 | NA |
| snoRNA:Ps | 0 | NA | NA | NA | NA | NA |
| snoRNA:Ps | 13.34042 | 0.040251 | 3.904549 | 0.010309 | 0.991775 | NA |
| snoRNA:Ps | 0.2094 | 1.005034 | 4.045341 | 0.248442 | 0.803792 | NA |
| CG34161 | 37.39119 | -1.70511 | 3.951934 | -0.43146 | 0.666132 | NA |
| CG34162 | 2.198698 | 1.671135 | 3.972627 | 0.420662 | 0.674002 | NA |
| CG34163 | 11.51699 | 1.736994 | 3.955359 | 0.43915 | 0.660553 | NA |
| CG34164 | 0 | NA | NA | NA | NA | NA |
| CG34165 | 0 | NA | NA | NA | NA | NA |
| CG34166 | 0 | NA | NA | NA | NA | NA |
| CG34167 | 21.05015 | 5.179901 | 3.853643 | 1.344157 | 0.178898 | NA |
| CG34168 | 0 | NA | NA | NA | NA | NA |
| CG34169 | 0.132078 | -1.13211 | 4.069918 | -0.27817 | 0.780885 | NA |
| CG34170 | 0 | NA | NA | NA | NA | NA |
| CG34171 | 0 | NA | NA | NA | NA | NA |
| CG34173 | 0 | NA | NA | NA | NA | NA |

| | | | | | | |
|---|---|---|---|---|---|---|
| CG34174 | 0 | NA | NA | NA | NA | NA |
| CG34175 | 0 | NA | NA | NA | NA | NA |
| CG34176 | 0 | NA | NA | NA | NA | NA |
| CG34177 | 0 | NA | NA | NA | NA | NA |
| CG34178 | 0 | NA | NA | NA | NA | NA |
| CG34179 | 0 | NA | NA | NA | NA | NA |
| CG34181 | 0 | NA | NA | NA | NA | NA |
| RpS28-like | 0 | NA | NA | NA | NA | NA |
| CG34184 | 0 | NA | NA | NA | NA | NA |
| CG34185 | 0 | NA | NA | NA | NA | NA |
| CG34190 | 0 | NA | NA | NA | NA | NA |
| CG34193 | 0 | NA | NA | NA | NA | NA |
| CG34194 | 0 | NA | NA | NA | NA | NA |
| CG34196 | 0 | NA | NA | NA | NA | NA |
| CG34198 | 0 | NA | NA | NA | NA | NA |
| CG34199 | 0 | NA | NA | NA | NA | NA |
| CG34201 | 16.29153 | -3.49339 | 3.904011 | -0.89482 | 0.370883 | NA |
| CG34202 | 0.565506 | -0.26224 | 3.986711 | -0.06578 | 0.947554 | NA |
| CG34203 | 0.53416 | -1.53481 | 4.012395 | -0.38252 | 0.702078 | NA |
| CG34204 | 0 | NA | NA | NA | NA | NA |
| CG34206 | 26.07028 | 1.746756 | 3.952894 | 0.441893 | 0.658567 | NA |
| CG34207 | 0 | NA | NA | NA | NA | NA |
| CG34208 | 7.132195 | -1.90434 | 3.956298 | -0.48134 | 0.630272 | NA |
| CG34210 | 10.1923 | -2.24595 | 3.960077 | -0.56715 | 0.570614 | NA |
| CG34211 | 0 | NA | NA | NA | NA | NA |
| CG34212 | 0 | NA | NA | NA | NA | NA |
| CG34214 | 0 | NA | NA | NA | NA | NA |
| CG34215 | 33.47263 | 1.690341 | 3.952388 | 0.427676 | 0.668887 | NA |
| CG34216 | 0 | NA | NA | NA | NA | NA |
| CG34217 | 0 | NA | NA | NA | NA | NA |
| CG34221 | 0.1047 | 0.613261 | 4.062799 | 0.150945 | 0.880019 | NA |
| CG34223 | 0.104623 | 0.639075 | 4.067674 | 0.157111 | 0.875158 | NA |
| CG34224 | 0 | NA | NA | NA | NA | NA |
| CG34225 | 0 | NA | NA | NA | NA | NA |
| CG34227 | 5.653796 | 1.719633 | 3.959809 | 0.434272 | 0.664091 | NA |
| CG34230 | 0 | NA | NA | NA | NA | NA |
| CG34234 | 0 | NA | NA | NA | NA | NA |
| CG34235 | 0 | NA | NA | NA | NA | NA |
| CG34236 | 0 | NA | NA | NA | NA | NA |
| CG34238 | 0 | NA | NA | NA | NA | NA |
| CG34244 | 0 | NA | NA | NA | NA | NA |
| CG34245 | 0 | NA | NA | NA | NA | NA |
| CG34247 | 0 | NA | NA | NA | NA | NA |
| CG34248 | 0.178053 | -1.16467 | 4.075067 | -0.2858 | 0.775028 | NA |
| CG34251 | 18.06871 | 1.610685 | 3.900631 | 0.412929 | 0.679658 | NA |
| CG34253 | 0 | NA | NA | NA | NA | NA |
| CR34254 | 0 | NA | NA | NA | NA | NA |
| Blos3 | 57.43077 | -1.06601 | 3.121798 | -0.34147 | 0.732746 | NA |
| CG34256 | 46.84882 | -2.27414 | 3.952963 | -0.5753 | 0.565088 | NA |
| CG34258 | 1.37921 | -3.74935 | 3.951609 | -0.94882 | 0.342714 | NA |

| | | | | | | |
|---|---|---|---|---|---|---|
| CG34260 | 32.95633 | 1.535458 | 3.952 | 0.388527 | 0.697626 | NA |
| CG34261 | 30.66345 | 0.42979 | 3.835644 | 0.112052 | 0.910783 | NA |
| CR34262 | 0 | NA | NA | NA | NA | NA |
| CG34263 | 0 | NA | NA | NA | NA | NA |
| CG34265 | 0 | NA | NA | NA | NA | NA |
| CG34267 | 0 | NA | NA | NA | NA | NA |
| CG34268 | 0 | NA | NA | NA | NA | NA |
| CG34269 | 0 | NA | NA | NA | NA | NA |
| Cpr65Ay | 0 | NA | NA | NA | NA | NA |
| CG34273 | 0 | NA | NA | NA | NA | NA |
| CG34274 | 0.104623 | 0.639075 | 4.067674 | 0.157111 | 0.875158 | NA |
| CG34276 | 0 | NA | NA | NA | NA | NA |
| CG34277 | 0 | NA | NA | NA | NA | NA |
| CG34278 | 0 | NA | NA | NA | NA | NA |
| CG34279 | 5.679341 | -1.8997 | 3.957646 | -0.48001 | 0.631222 | NA |
| CG34280 | 0 | NA | NA | NA | NA | NA |
| CG34281 | 0 | NA | NA | NA | NA | NA |
| CG34282 | 0.178053 | -1.16467 | 4.075067 | -0.2858 | 0.775028 | NA |
| CG34283 | 0 | NA | NA | NA | NA | NA |
| CG34284 | 0 | NA | NA | NA | NA | NA |
| CR34285 | 0 | NA | NA | NA | NA | NA |
| CG34286 | 0 | NA | NA | NA | NA | NA |
| CG34287 | 0 | NA | NA | NA | NA | NA |
| CG34290 | 0.178053 | -1.16467 | 4.075067 | -0.2858 | 0.775028 | NA |
| CG34291 | 0 | NA | NA | NA | NA | NA |
| CG34292 | 0.282753 | -0.25398 | 4.020204 | -0.06318 | 0.949626 | NA |
| CG34295 | 0 | NA | NA | NA | NA | NA |
| CG34296 | 0 | NA | NA | NA | NA | NA |
| CG34297 | 1.546449 | -0.90035 | 3.878965 | -0.23211 | 0.816451 | NA |
| CG34298 | 0 | NA | NA | NA | NA | NA |
| CG34299 | 1.424426 | -1.63334 | 3.97617 | -0.41078 | 0.681232 | NA |
| CG34300 | 26.81957 | -4.85751 | 3.890047 | -1.2487 | 0.211774 | NA |
| CG34301 | 0 | NA | NA | NA | NA | NA |
| CG34302 | 0 | NA | NA | NA | NA | NA |
| CG34305 | 0 | NA | NA | NA | NA | NA |
| CG34307 | 0 | NA | NA | NA | NA | NA |
| CG34308 | 0 | NA | NA | NA | NA | NA |
| CR34311 | 0 | NA | NA | NA | NA | NA |
| CG34312 | 0 | NA | NA | NA | NA | NA |
| CG34313 | 0 | NA | NA | NA | NA | NA |
| Sws1 | 0 | NA | NA | NA | NA | NA |
| CG34315 | 0 | NA | NA | NA | NA | NA |
| CG34316 | 0 | NA | NA | NA | NA | NA |
| Rpp14a | 0 | NA | NA | NA | NA | NA |
| CR34320 | 0 | NA | NA | NA | NA | NA |
| inaF-C | 0 | NA | NA | NA | NA | NA |
| inaF-A | 0.178053 | -1.16467 | 4.075067 | -0.2858 | 0.775028 | NA |
| CG34323 | 0 | NA | NA | NA | NA | NA |
| CG34324 | 4.869825 | 0.168899 | 3.875984 | 0.043576 | 0.965242 | NA |
| CG34327 | 0 | NA | NA | NA | NA | NA |

| | | | | | | |
|---|---|---|---|---|---|---|
| CG34328 | 0 | NA | NA | NA | NA | NA |
| Diedel3 | 0 | NA | NA | NA | NA | NA |
| CG34330 | 0 | NA | NA | NA | NA | NA |
| CG34331 | 0.356866 | -1.92326 | 4.037889 | -0.4763 | 0.633858 | NA |
| CG34332 | 0 | NA | NA | NA | NA | NA |
| Vml | 0 | NA | NA | NA | NA | NA |
| CG34337 | 32.07801 | -0.78824 | 3.836045 | -0.20548 | 0.837196 | NA |
| CG34338 | 0 | NA | NA | NA | NA | NA |
| CG34342 | 0.233093 | 0.692656 | 4.076246 | 0.169925 | 0.865069 | NA |
| CG34345 | 0.178053 | -1.16467 | 4.075067 | -0.2858 | 0.775028 | NA |
| ppk9 | 0 | NA | NA | NA | NA | NA |
| CG34375 | 0 | NA | NA | NA | NA | NA |
| CG34382 | 0.1047 | 0.613261 | 4.062799 | 0.150945 | 0.880019 | NA |
| CG34386 | 0 | NA | NA | NA | NA | NA |
| natalisin | 0 | NA | NA | NA | NA | NA |
| CG34398 | 6.650767 | -2.25569 | 3.816227 | -0.59108 | 0.554468 | NA |
| Nox | 17.90369 | 1.743194 | 3.95379 | 0.440892 | 0.659291 | NA |
| CG34402 | 6.15927 | -2.57152 | 3.928673 | -0.65455 | 0.512757 | NA |
| Lgr4 | 38.88874 | 1.61589 | 3.899551 | 0.414378 | 0.678597 | NA |
| Mthfs | 0 | NA | NA | NA | NA | NA |
| CG34425 | 0 | NA | NA | NA | NA | NA |
| CG34426 | 0.178053 | -1.16467 | 4.075067 | -0.2858 | 0.775028 | NA |
| CG34427 | 0 | NA | NA | NA | NA | NA |
| CG34428 | 0 | NA | NA | NA | NA | NA |
| CG34429 | 0 | NA | NA | NA | NA | NA |
| CG34431 | 4.137111 | 0.483953 | 3.852062 | 0.125635 | 0.900021 | NA |
| CG34432 | 0 | NA | NA | NA | NA | NA |
| CG34433 | 0 | NA | NA | NA | NA | NA |
| CG34435 | 0 | NA | NA | NA | NA | NA |
| CG34436 | 0.712213 | -1.57119 | 3.998613 | -0.39293 | 0.694369 | NA |
| CG34437 | 2.303398 | 1.674512 | 3.971716 | 0.421609 | 0.67331 | NA |
| ImgB | 0 | NA | NA | NA | NA | NA |
| CG34442 | 0 | NA | NA | NA | NA | NA |
| CG34443 | 0 | NA | NA | NA | NA | NA |
| CG34444 | 0 | NA | NA | NA | NA | NA |
| CG34445 | 0 | NA | NA | NA | NA | NA |
| CG34446 | 0 | NA | NA | NA | NA | NA |
| CG34447 | 0 | NA | NA | NA | NA | NA |
| CG34448 | 0 | NA | NA | NA | NA | NA |
| CG34450 | 0 | NA | NA | NA | NA | NA |
| CG34451 | 0 | NA | NA | NA | NA | NA |
| CG34452 | 0 | NA | NA | NA | NA | NA |
| CG34453 | 54.05472 | 0.127405 | 3.9513 | 0.032244 | 0.974278 | NA |
| CG34456 | 0 | NA | NA | NA | NA | NA |
| CG34458 | 0 | NA | NA | NA | NA | NA |
| CG34459 | 0.356107 | -1.47099 | 4.037764 | -0.36431 | 0.715628 | NA |
| CG34460 | 0 | NA | NA | NA | NA | NA |
| CG34462 | 0.178053 | -1.16467 | 4.075067 | -0.2858 | 0.775028 | NA |
| Mst77Y-16 | 6.207651 | -1.90163 | 3.957084 | -0.48056 | 0.630826 | NA |
| CR40601 | 0 | NA | NA | NA | NA | NA |

| | | | | | | |
|---|---|---|---|---|---|---|
| lncRNA:CR | 0 | NA | NA | NA | NA | NA |
| CG40635 | 0.5235 | 1.490714 | 4.02555 | 0.370313 | 0.711149 | NA |
| lncRNA:CR | 0 | NA | NA | NA | NA | NA |
| RYa | 13.40159 | 1.739423 | 3.954743 | 0.439832 | 0.660059 | NA |
| CR40801 | 0.792466 | -1.7823 | 3.993845 | -0.44626 | 0.655408 | NA |
| CG40813 | 3.871061 | 1.506839 | 3.959948 | 0.38052 | 0.70356 | NA |
| Su(Ste):CR | 0.132078 | -1.13211 | 4.069918 | -0.27817 | 0.780885 | NA |
| lncRNA:CR | 0 | NA | NA | NA | NA | NA |
| CR41243 | 0.31387 | 1.261782 | 4.037145 | 0.312543 | 0.754628 | NA |
| CR41501 | 49.29806 | 2.057472 | 3.81542 | 0.539252 | 0.589713 | NA |
| CG41242 | 0 | NA | NA | NA | NA | NA |
| lncRNA:CR | 0 | NA | NA | NA | NA | NA |
| CR41519 | 0.104623 | 0.639075 | 4.067674 | 0.157111 | 0.875158 | NA |
| CR41597 | 0 | NA | NA | NA | NA | NA |
| CR41320 | 20.01437 | 1.693361 | 3.737582 | 0.453063 | 0.650503 | NA |
| CG41284 | 0 | NA | NA | NA | NA | NA |
| lncRNA:CR | 0 | NA | NA | NA | NA | NA |
| CR41423 | 0.178812 | -1.08808 | 4.061735 | -0.26789 | 0.788788 | NA |
| 28SrRNA-F | 41.52785 | -0.11744 | 3.506814 | -0.03349 | 0.973284 | NA |
| CG41562 | 0.732363 | 1.393579 | 3.993417 | 0.348969 | 0.727112 | NA |
| 18SrRNA:C | 0 | NA | NA | NA | NA | NA |
| Su(Ste):CR | 0.2094 | 1.005034 | 4.045341 | 0.248442 | 0.803792 | NA |
| CR41571 | 0 | NA | NA | NA | NA | NA |
| CG41561 | 0 | NA | NA | NA | NA | NA |
| CR40450 | 44.02617 | -4.9226 | 3.860887 | -1.27499 | 0.202313 | NA |
| CR41379 | 1.827286 | 3.319894 | 3.963865 | 0.83754 | 0.402289 | NA |
| 2SrRNA-Ps | 0.1047 | 0.613261 | 4.062799 | 0.150945 | 0.880019 | NA |
| CR41506 | 3.574231 | 0.322748 | 3.853103 | 0.083763 | 0.933245 | NA |
| CR41504 | 0 | NA | NA | NA | NA | NA |
| CR41509 | 0 | NA | NA | NA | NA | NA |
| CR41508 | 0 | NA | NA | NA | NA | NA |
| CR41507 | 1.234675 | -3.13913 | 3.971472 | -0.79042 | 0.429282 | NA |
| CR42201 | 0.313946 | 1.788224 | 4.01364 | 0.445537 | 0.655932 | NA |
| snoRNA:M | 0 | NA | NA | NA | NA | NA |
| snoRNA:M | 0 | NA | NA | NA | NA | NA |
| snoRNA:M | 1.164483 | 1.950196 | 3.904832 | 0.499432 | 0.617475 | NA |
| snoRNA:M | 1.464879 | 3.015441 | 3.967965 | 0.759947 | 0.447287 | NA |
| snoRNA:M | 0 | NA | NA | NA | NA | NA |
| snoRNA:M | 0 | NA | NA | NA | NA | NA |
| snoRNA:Or | 2.825288 | 4.097154 | 3.94606 | 1.03829 | 0.299135 | NA |
| snoRNA:M | 0 | NA | NA | NA | NA | NA |
| snoRNA:M | 0 | NA | NA | NA | NA | NA |
| snoRNA:M | 0 | NA | NA | NA | NA | NA |
| snoRNA:M | 0 | NA | NA | NA | NA | NA |
| snoRNA:M | 0 | NA | NA | NA | NA | NA |
| snoRNA:M | 0 | NA | NA | NA | NA | NA |
| snoRNA:M | 0 | NA | NA | NA | NA | NA |
| snoRNA:M | 0 | NA | NA | NA | NA | NA |
| snoRNA:M | 0.1047 | 0.613261 | 4.062799 | 0.150945 | 0.880019 | NA |
| snoRNA:M | 0 | NA | NA | NA | NA | NA |

| | | | | | | |
|---|---|---|---|---|---|---|
| snoRNA:M | 0.283512 | -0.2331 | 4.065094 | -0.05734 | 0.954273 | NA |
| snoRNA:M | 0 | NA | NA | NA | NA | NA |
| snoRNA:M | 0 | NA | NA | NA | NA | NA |
| snoRNA:M | 0 | NA | NA | NA | NA | NA |
| snoRNA:M | 0 | NA | NA | NA | NA | NA |
| snoRNA:M | 0 | NA | NA | NA | NA | NA |
| snoRNA:M | 0 | NA | NA | NA | NA | NA |
| snoRNA:M | 4.927818 | -5.39244 | 3.922516 | -1.37474 | 0.169212 | NA |
| snoRNA:M | 0 | NA | NA | NA | NA | NA |
| snoRNA:M | 0 | NA | NA | NA | NA | NA |
| snoRNA:M | 1.094576 | -2.45718 | 3.919279 | -0.62695 | 0.530693 | NA |
| snoRNA:M | 0 | NA | NA | NA | NA | NA |
| snoRNA:M | 0 | NA | NA | NA | NA | NA |
| snoRNA:M | 0 | NA | NA | NA | NA | NA |
| snoRNA:M | 0.1047 | 0.613261 | 4.062799 | 0.150945 | 0.880019 | NA |
| snoRNA:M | 0 | NA | NA | NA | NA | NA |
| snoRNA:M | 0 | NA | NA | NA | NA | NA |
| snoRNA:M | 0.984682 | 0.033526 | 3.923473 | 0.008545 | 0.993182 | NA |
| snoRNA:M | 0 | NA | NA | NA | NA | NA |
| snoRNA:M | 0 | NA | NA | NA | NA | NA |
| snoRNA:O | 0 | NA | NA | NA | NA | NA |
| snoRNA:M | 6.288439 | 0.458237 | 3.86805 | 0.118467 | 0.905697 | NA |
| snoRNA:M | 0.178812 | -1.08808 | 4.061735 | -0.26789 | 0.788788 | NA |
| snoRNA:M | 0 | NA | NA | NA | NA | NA |
| snoRNA:M | 0 | NA | NA | NA | NA | NA |
| snoRNA:M | 0 | NA | NA | NA | NA | NA |
| snoRNA:M | 0.670206 | -0.05147 | 3.98115 | -0.01293 | 0.989686 | NA |
| snoRNA:O | 9.810748 | -1.02007 | 3.863705 | -0.26401 | 0.79177 | NA |
| snoRNA:M | 0 | NA | NA | NA | NA | NA |
| snoRNA:O | 8.239336 | -2.29073 | 3.842116 | -0.59622 | 0.551031 | NA |
| snoRNA:O | 0 | NA | NA | NA | NA | NA |
| snoRNA:O | 0 | NA | NA | NA | NA | NA |
| snoRNA:M | 0 | NA | NA | NA | NA | NA |
| snoRNA:M | 7.239783 | 0.346644 | 3.953483 | 0.087681 | 0.930131 | NA |
| snoRNA:M | 0 | NA | NA | NA | NA | NA |
| snoRNA:M | 1.251686 | -2.06022 | 4.011654 | -0.51356 | 0.60756 | NA |
| snoRNA:M | 0 | NA | NA | NA | NA | NA |
| snoRNA:M | 0.461565 | -1.07936 | 4.002266 | -0.26969 | 0.7874 | NA |
| snoRNA:M | 0 | NA | NA | NA | NA | NA |
| snoRNA:M | 0.178053 | -1.16467 | 4.075067 | -0.2858 | 0.775028 | NA |
| snoRNA:O | 0 | NA | NA | NA | NA | NA |
| CR42490 | 0.178053 | -1.16467 | 4.075067 | -0.2858 | 0.775028 | NA |
| psd | 0 | NA | NA | NA | NA | NA |
| Vm26Ac | 0 | NA | NA | NA | NA | NA |
| se | 0 | NA | NA | NA | NA | NA |
| tum | 0 | NA | NA | NA | NA | NA |
| t | 0.442209 | -2.31142 | 4.024999 | -0.57427 | 0.565788 | NA |
| tw | 0 | NA | NA | NA | NA | NA |
| stl | 1.194388 | 0.722286 | 3.956144 | 0.182573 | 0.855133 | NA |
| Cpr47Eg | 0 | NA | NA | NA | NA | NA |

| | | | | | | |
|---|---|---|---|---|---|---|
| Spt-I | 0 | NA | NA | NA | NA | NA |
| snoRNA:M | 0.264155 | -1.54099 | 4.048976 | -0.38059 | 0.703509 | NA |
| snoRNA:Ps | 0 | NA | NA | NA | NA | NA |
| snoRNA:Ps | 0 | NA | NA | NA | NA | NA |
| snoRNA:Ps | 35.25683 | -3.98371 | 3.889472 | -1.02423 | 0.305727 | NA |
| snoRNA:Or | 0 | NA | NA | NA | NA | NA |
| CG34112 | 23.98521 | -6.43086 | 3.846532 | -1.67186 | 0.094552 | NA |
| CG33342 | 0 | NA | NA | NA | NA | NA |
| Lcp65Ag3 | 0 | NA | NA | NA | NA | NA |
| snoRNA:Ps | 0.1047 | 0.613261 | 4.062799 | 0.150945 | 0.880019 | NA |
| snoRNA:Ps | 0 | NA | NA | NA | NA | NA |
| snoRNA:Ps | 0 | NA | NA | NA | NA | NA |
| snoRNA:Ps | 21.38994 | 0.049084 | 3.857496 | 0.012724 | 0.989848 | NA |
| snoRNA:Ps | 7.688928 | -2.23489 | 3.962917 | -0.56395 | 0.572788 | NA |
| snoRNA:Ps | 27.5026 | -7.47493 | 3.912742 | -1.91041 | 0.056081 | NA |
| snoRNA:Ps | 6.464319 | -0.30451 | 3.847808 | -0.07914 | 0.936923 | NA |
| snoRNA:Ps | 6.766025 | -1.69111 | 3.956514 | -0.42742 | 0.66907 | NA |
| snoRNA:Ps | 28.75449 | 0.361046 | 3.731775 | 0.096749 | 0.922926 | NA |
| snoRNA:Ps | 4.807438 | -1.68437 | 3.95875 | -0.42548 | 0.670487 | NA |
| snoRNA:Ps | 0 | NA | NA | NA | NA | NA |
| snoRNA:Ps | 17.44021 | -0.04625 | 3.740045 | -0.01237 | 0.990133 | NA |
| snoRNA:Ps | 0.565506 | -0.26224 | 3.986711 | -0.06578 | 0.947554 | NA |
| snoRNA:Or | 10.2345 | 0.309871 | 3.763927 | 0.082327 | 0.934387 | NA |
| CG13272 | 10.71288 | -1.30008 | 3.811294 | -0.34111 | 0.733018 | NA |
| iav | 0 | NA | NA | NA | NA | NA |
| hd | 7.852418 | 2.924533 | 3.930606 | 0.744041 | 0.456852 | NA |
| frtz | 38.13939 | -1.02958 | 3.920492 | -0.26261 | 0.792847 | NA |
| dgo | 0.178053 | -1.16467 | 4.075067 | -0.2858 | 0.775028 | NA |
| Cpr65Ax1 | 0 | NA | NA | NA | NA | NA |
| Cyt-c-d | 31.27061 | 0.344623 | 3.839481 | 0.089758 | 0.92848 | NA |
| mbm | 42.01534 | -2.0615 | 3.932109 | -0.52427 | 0.600088 | NA |
| snky | 0 | NA | NA | NA | NA | NA |
| rtp | 1.256399 | 1.619735 | 3.986836 | 0.406271 | 0.684544 | NA |
| alphaTub6 | 23.09031 | 3.408532 | 3.747986 | 0.90943 | 0.363123 | NA |
| Hyls1 | 0 | NA | NA | NA | NA | NA |
| eater | 0 | NA | NA | NA | NA | NA |
| Tb | 0 | NA | NA | NA | NA | NA |
| 5.8SrRNA:( | 0 | NA | NA | NA | NA | NA |
| Jon65Aiv | 0 | NA | NA | NA | NA | NA |
| lncRNA:CR | 0 | NA | NA | NA | NA | NA |
| CG34222 | 3.43402 | -1.88523 | 3.961894 | -0.47584 | 0.634189 | NA |
| CG33655 | 0.942299 | 1.584447 | 3.996979 | 0.396411 | 0.691802 | NA |
| CG34241 | 0 | NA | NA | NA | NA | NA |
| CG34160 | 53.41598 | -1.70607 | 3.951626 | -0.43174 | 0.665932 | NA |
| BG642167 | 0 | NA | NA | NA | NA | NA |
| Dup99B | 0 | NA | NA | NA | NA | NA |
| CG34461 | 0.233093 | 0.692656 | 4.076246 | 0.169925 | 0.865069 | NA |
| CG33308 | 0 | NA | NA | NA | NA | NA |
| Acbp3 | 0.178053 | -1.16467 | 4.075067 | -0.2858 | 0.775028 | NA |
| dUTPase | 0 | NA | NA | NA | NA | NA |

| | | | | | | |
|---|---|---|---|---|---|---|
| CG17242 | 0.178053 | -1.16467 | 4.075067 | -0.2858 | 0.775028 | NA |
| CG17575 | 0.233093 | 0.692656 | 4.076246 | 0.169925 | 0.865069 | NA |
| CG1288 | 0.178053 | -1.16467 | 4.075067 | -0.2858 | 0.775028 | NA |
| glob2 | 0.132078 | -1.13211 | 4.069918 | -0.27817 | 0.780885 | NA |
| CG14034 | 4.807438 | -1.68437 | 3.95875 | -0.42548 | 0.670487 | NA |
| CG32388 | 0 | NA | NA | NA | NA | NA |
| CG33981 | 0 | NA | NA | NA | NA | NA |
| CG42237 | 3.978521 | 2.873751 | 3.942721 | 0.728875 | 0.466078 | NA |
| pot | 0.51577 | 1.033715 | 4.015187 | 0.257451 | 0.79683 | NA |
| lmgA | 0 | NA | NA | NA | NA | NA |
| CG34434 | 0.462248 | -0.67622 | 4.039367 | -0.16741 | 0.867049 | NA |
| Cht10 | 17.03506 | 1.591822 | 3.842586 | 0.414258 | 0.678685 | NA |
| CG42245 | 0 | NA | NA | NA | NA | NA |
| CG42246 | 3.830253 | -1.88895 | 3.960794 | -0.47691 | 0.633424 | NA |
| DCX-EMAF | 60.6503 | 0.647423 | 3.920179 | 0.165152 | 0.868825 | NA |
| CG42249 | 6.822581 | -2.42613 | 3.785184 | -0.64095 | 0.521552 | NA |
| PolA1 | 0.837599 | 1.568055 | 4.001801 | 0.391837 | 0.695178 | NA |
| CG42255 | 34.3461 | 1.157374 | 3.873966 | 0.298757 | 0.765125 | NA |
| CG42264 | 0 | NA | NA | NA | NA | NA |
| CG42266 | 2.272811 | 0.29017 | 3.960962 | 0.073257 | 0.941601 | NA |
| RunxB | 0 | NA | NA | NA | NA | NA |
| CG42269 | 0.466186 | 0.959471 | 4.033986 | 0.237847 | 0.812 | NA |
| CG42272 | 8.899494 | 1.731975 | 3.956638 | 0.437739 | 0.661575 | NA |
| CG42284 | 0 | NA | NA | NA | NA | NA |
| CG42287 | 2.012992 | 2.885432 | 3.965929 | 0.727555 | 0.466886 | NA |
| CG42288 | 28.88843 | -3.09444 | 3.800109 | -0.8143 | 0.415471 | NA |
| Ir60b | 0 | NA | NA | NA | NA | NA |
| Ir60c | 0 | NA | NA | NA | NA | NA |
| Ir60d | 0 | NA | NA | NA | NA | NA |
| Ir7f | 0 | NA | NA | NA | NA | NA |
| Ir7e | 0 | NA | NA | NA | NA | NA |
| Ir7d | 0 | NA | NA | NA | NA | NA |
| CG42296 | 0 | NA | NA | NA | NA | NA |
| Ir94d | 0 | NA | NA | NA | NA | NA |
| Ir94e | 0 | NA | NA | NA | NA | NA |
| CG42299 | 0 | NA | NA | NA | NA | NA |
| CG42300 | 0 | NA | NA | NA | NA | NA |
| CG42302 | 0 | NA | NA | NA | NA | NA |
| Snup | 0 | NA | NA | NA | NA | NA |
| CG42304 | 0 | NA | NA | NA | NA | NA |
| CG42305 | 0 | NA | NA | NA | NA | NA |
| CG42306 | 0 | NA | NA | NA | NA | NA |
| CG42307 | 0.356107 | -1.47099 | 4.037764 | -0.36431 | 0.715628 | NA |
| CG42308 | 0 | NA | NA | NA | NA | NA |
| Mlp60A | 0 | NA | NA | NA | NA | NA |
| prom | 25.67408 | 4.169242 | 3.806226 | 1.095374 | 0.273353 | NA |
| grh | 17.05361 | -2.42819 | 3.860112 | -0.62905 | 0.529319 | NA |
| CG42319 | 0.5235 | 1.490714 | 4.02555 | 0.370313 | 0.711149 | NA |
| CG42323 | 0 | NA | NA | NA | NA | NA |
| CG42329 | 0 | NA | NA | NA | NA | NA |

| | | | | | | |
|---|---|---|---|---|---|---|
| lectin-22C | 0 | NA | NA | NA | NA | NA |
| CCKLR-17D | 49.48404 | 0.296766 | 3.834589 | 0.077392 | 0.938312 | NA |
| comm3 | 0 | NA | NA | NA | NA | NA |
| CG42335 | 1.360248 | -0.69345 | 3.975274 | -0.17444 | 0.861519 | NA |
| Prim2 | 1.780533 | -1.64713 | 3.971374 | -0.41475 | 0.678325 | NA |
| sqa | 1.128562 | -2.72595 | 3.934459 | -0.69284 | 0.488411 | NA |
| Jabba | 2.565465 | 0.351309 | 3.959571 | 0.088724 | 0.929301 | NA |
| Ir40a | 0 | NA | NA | NA | NA | NA |
| CG42353 | 0 | NA | NA | NA | NA | NA |
| CG42355 | 0 | NA | NA | NA | NA | NA |
| CG42356 | 0 | NA | NA | NA | NA | NA |
| CG42357 | 0 | NA | NA | NA | NA | NA |
| CG42362 | 0 | NA | NA | NA | NA | NA |
| CG42363 | 0 | NA | NA | NA | NA | NA |
| CG42364 | 0 | NA | NA | NA | NA | NA |
| CG42366 | 0 | NA | NA | NA | NA | NA |
| CG42367 | 0.1047 | 0.613261 | 4.062799 | 0.150945 | 0.880019 | NA |
| CG42369 | 0 | NA | NA | NA | NA | NA |
| Nepl6 | 42.55742 | -0.32424 | 3.835188 | -0.08454 | 0.932624 | NA |
| CG42371 | 0 | NA | NA | NA | NA | NA |
| CG42372 | 0 | NA | NA | NA | NA | NA |
| CG42374 | 0 | NA | NA | NA | NA | NA |
| CG42375 | 0 | NA | NA | NA | NA | NA |
| CG42379 | 0 | NA | NA | NA | NA | NA |
| CG42380 | 0 | NA | NA | NA | NA | NA |
| CG42381 | 0 | NA | NA | NA | NA | NA |
| CG42382 | 0.178053 | -1.16467 | 4.075067 | -0.2858 | 0.775028 | NA |
| CG42383 | 0.2094 | 1.005034 | 4.045341 | 0.248442 | 0.803792 | NA |
| tal-1A | 0 | NA | NA | NA | NA | NA |
| tal-2A | 0 | NA | NA | NA | NA | NA |
| tal-3A | 0 | NA | NA | NA | NA | NA |
| tal-AA | 0 | NA | NA | NA | NA | NA |
| CG42395 | 0 | NA | NA | NA | NA | NA |
| CG42397 | 0 | NA | NA | NA | NA | NA |
| bora | 0 | NA | NA | NA | NA | NA |
| sinah | 0 | NA | NA | NA | NA | NA |
| loopin-1 | 0 | NA | NA | NA | NA | NA |
| CG34309 | 0 | NA | NA | NA | NA | NA |
| out | 16.97128 | -3.24688 | 3.868596 | -0.83929 | 0.401306 | NA |
| Su(Ste):CR | 0.264155 | -1.54099 | 4.048976 | -0.38059 | 0.703509 | NA |
| Su(Ste):CR | 8.415129 | -1.05988 | 3.953582 | -0.26808 | 0.788637 | NA |
| Su(Ste):CR | 1.662101 | -1.09108 | 3.964539 | -0.27521 | 0.783155 | NA |
| Su(Ste):CR | 0.178053 | -1.16467 | 4.075067 | -0.2858 | 0.775028 | NA |
| Su(Ste):CR | 0.776424 | -0.14961 | 3.950397 | -0.03787 | 0.969789 | NA |
| Su(Ste):CR | 0 | NA | NA | NA | NA | NA |
| Su(Ste):CR | 8.14869 | -4.2577 | 3.893911 | -1.09343 | 0.274207 | NA |
| Su(Ste):CR | 0.132078 | -1.13211 | 4.069918 | -0.27817 | 0.780885 | NA |
| Su(Ste):CR | 1.972325 | 3.162291 | 3.909253 | 0.808925 | 0.418558 | NA |
| Su(Ste):CR | 0.178812 | -1.08808 | 4.061735 | -0.26789 | 0.788788 | NA |
| Su(Ste):CR | 0 | NA | NA | NA | NA | NA |

| | | | | | | |
|---|---|---|---|---|---|---|
| Su(Ste):CR | 1.642779 | 0.571925 | 3.935522 | 0.145324 | 0.884455 | NA |
| Su(Ste):CR | 0 | NA | NA | NA | NA | NA |
| Su(Ste):CR | 0 | NA | NA | NA | NA | NA |
| Su(Ste):CR | 0.462324 | -0.59023 | 4.044233 | -0.14594 | 0.883966 | NA |
| Su(Ste):CR | 2.290454 | 0.258093 | 3.867943 | 0.066726 | 0.9468 | NA |
| Su(Ste):CR | 0 | NA | NA | NA | NA | NA |
| Su(Ste):CR | 1.560473 | -0.69219 | 3.96537 | -0.17456 | 0.861426 | NA |
| Su(Ste):CR | 1.267059 | 0.18295 | 3.967592 | 0.046111 | 0.963222 | NA |
| Su(Ste):CR | 0 | NA | NA | NA | NA | NA |
| Su(Ste):CR | 0 | NA | NA | NA | NA | NA |
| Su(Ste):CR | 0 | NA | NA | NA | NA | NA |
| Su(Ste):CR | 12.31828 | -1.26897 | 3.885309 | -0.32661 | 0.743965 | NA |
| Su(Ste):CR | 0 | NA | NA | NA | NA | NA |
| Su(Ste):CR | 0.46073 | -0.17853 | 4.00793 | -0.04454 | 0.964471 | NA |
| Su(Ste):CR | 19.16026 | -0.53338 | 3.759553 | -0.14187 | 0.887179 | NA |
| Su(Ste):CR | 0.1047 | 0.613261 | 4.062799 | 0.150945 | 0.880019 | NA |
| Su(Ste):CR | 7.964415 | 0.86224 | 3.864498 | 0.223118 | 0.823444 | NA |
| Su(Ste):CR | 0.283512 | -0.2331 | 4.065094 | -0.05734 | 0.954273 | NA |
| NimC1 | 8.724611 | -1.69489 | 3.95527 | -0.42851 | 0.668277 | NA |
| CG42446 | 0.132078 | -1.13211 | 4.069918 | -0.27817 | 0.780885 | NA |
| inaF-B | 0.178053 | -1.16467 | 4.075067 | -0.2858 | 0.775028 | NA |
| Skadu | 0 | NA | NA | NA | NA | NA |
| CG42449 | 0.1047 | 0.613261 | 4.062799 | 0.150945 | 0.880019 | NA |
| CG42455 | 0 | NA | NA | NA | NA | NA |
| Dpm2 | 0 | NA | NA | NA | NA | NA |
| Sfp23F | 0 | NA | NA | NA | NA | NA |
| CG42460 | 0 | NA | NA | NA | NA | NA |
| Sfp24Ba | 15.28619 | 1.741265 | 3.954277 | 0.44035 | 0.659684 | NA |
| Sfp24Bb | 11.44399 | -2.24975 | 3.959108 | -0.56825 | 0.569868 | NA |
| Sfp24Bd | 8.061894 | 1.729706 | 3.957218 | 0.437101 | 0.662038 | NA |
| CG42464 | 0 | NA | NA | NA | NA | NA |
| CG42465 | 0 | NA | NA | NA | NA | NA |
| Sfp24C1 | 0 | NA | NA | NA | NA | NA |
| CG42467 | 0.178053 | -1.16467 | 4.075067 | -0.2858 | 0.775028 | NA |
| Sfp24F | 0 | NA | NA | NA | NA | NA |
| Sfp26Ac | 0 | NA | NA | NA | NA | NA |
| CG42470 | 0.31387 | 1.261782 | 4.037145 | 0.312543 | 0.754628 | NA |
| CG42471 | 0 | NA | NA | NA | NA | NA |
| Sfp33A1 | 0 | NA | NA | NA | NA | NA |
| Sfp33A2 | 4.711497 | 1.713057 | 3.961514 | 0.432425 | 0.665433 | NA |
| Sfp33A3 | 0 | NA | NA | NA | NA | NA |
| Sfp35C | 0 | NA | NA | NA | NA | NA |
| Sfp51E | 0 | NA | NA | NA | NA | NA |
| Sfp53D | 0 | NA | NA | NA | NA | NA |
| Sfp60F | 0.104623 | 0.639075 | 4.067674 | 0.157111 | 0.875158 | NA |
| Sfp65A | 0 | NA | NA | NA | NA | NA |
| Sfp70A4 | 0 | NA | NA | NA | NA | NA |
| CG42481 | 0 | NA | NA | NA | NA | NA |
| Sfp77F | 0 | NA | NA | NA | NA | NA |
| Sfp79B | 0.178053 | -1.16467 | 4.075067 | -0.2858 | 0.775028 | NA |

| | | | | | | |
|---|---|---|---|---|---|---|
| Sfp84E | 0 | NA | NA | NA | NA | NA |
| Sfp87B | 0 | NA | NA | NA | NA | NA |
| Tdc1 | 27.30667 | 1.53463 | 3.952225 | 0.388295 | 0.697798 | NA |
| nab | 0 | NA | NA | NA | NA | NA |
| CG42486 | 0 | NA | NA | NA | NA | NA |
| CG42487 | 0 | NA | NA | NA | NA | NA |
| CG42488 | 0 | NA | NA | NA | NA | NA |
| lncRNA:CR | 0 | NA | NA | NA | NA | NA |
| CG17571 | 1.175108 | -0.4198 | 3.916646 | -0.10718 | 0.914644 | NA |
| CG17570 | 0 | NA | NA | NA | NA | NA |
| snoRNA:M | 6.16458 | 0.254405 | 3.784263 | 0.067227 | 0.946401 | NA |
| drd | 0 | NA | NA | NA | NA | NA |
| CG42493 | 38.86737 | 3.201514 | 3.90747 | 0.819332 | 0.412597 | NA |
| NimC4 | 0 | NA | NA | NA | NA | NA |
| CG42496 | 0 | NA | NA | NA | NA | NA |
| CG42497 | 0 | NA | NA | NA | NA | NA |
| CG42498 | 0 | NA | NA | NA | NA | NA |
| CR42499 | 0 | NA | NA | NA | NA | NA |
| CG42500 | 0 | NA | NA | NA | NA | NA |
| CG42502 | 0 | NA | NA | NA | NA | NA |
| Mocs2A | 0 | NA | NA | NA | NA | NA |
| CG42504 | 0 | NA | NA | NA | NA | NA |
| CG42505 | 0 | NA | NA | NA | NA | NA |
| CG42507 | 0 | NA | NA | NA | NA | NA |
| Xport-B | 0 | NA | NA | NA | NA | NA |
| BBIP1 | 0 | NA | NA | NA | NA | NA |
| CG42512 | 0 | NA | NA | NA | NA | NA |
| mtg | 2.097839 | 1.11873 | 3.972242 | 0.281637 | 0.778222 | NA |
| CG42516 | 0 | NA | NA | NA | NA | NA |
| CG42518 | 0 | NA | NA | NA | NA | NA |
| CG17147 | 0.1047 | 0.613261 | 4.062799 | 0.150945 | 0.880019 | NA |
| Pbp49 | 0 | NA | NA | NA | NA | NA |
| MED9 | 0 | NA | NA | NA | NA | NA |
| CG42525 | 0 | NA | NA | NA | NA | NA |
| CG42529 | 0 | NA | NA | NA | NA | NA |
| CR42530 | 0 | NA | NA | NA | NA | NA |
| CG42531 | 0 | NA | NA | NA | NA | NA |
| CR42532 | 0.890266 | -1.59482 | 3.989917 | -0.39971 | 0.689368 | NA |
| CG42534 | 0.282677 | -0.09599 | 4.057934 | -0.02366 | 0.981127 | NA |
| Antp | 4.734085 | -1.48459 | 3.957933 | -0.37509 | 0.707592 | NA |
| CG42535 | 0 | NA | NA | NA | NA | NA |
| CG42536 | 0 | NA | NA | NA | NA | NA |
| CG42537 | 0 | NA | NA | NA | NA | NA |
| CG42538 | 4.856646 | 0.384027 | 3.957599 | 0.097035 | 0.922698 | NA |
| Rrp40 | 1.884522 | 2.777951 | 3.96711 | 0.700245 | 0.483774 | NA |
| serp | 2.314693 | -1.66033 | 3.966843 | -0.41855 | 0.675543 | NA |
| CG42544 | 5.943496 | -1.90071 | 3.957353 | -0.4803 | 0.631016 | NA |
| CR42545 | 0.178053 | -1.16467 | 4.075067 | -0.2858 | 0.775028 | NA |
| CR42546 | 0 | NA | NA | NA | NA | NA |
| CR42548 | 0 | NA | NA | NA | NA | NA |

| | | | | | | |
|---|---|---|---|---|---|---|
| CG42556 | 8.085511 | 4.238261 | 3.901018 | 1.08645 | 0.27728 | NA |
| CG42557 | 0 | NA | NA | NA | NA | NA |
| CG42558 | 0 | NA | NA | NA | NA | NA |
| CG42559 | 0 | NA | NA | NA | NA | NA |
| CG42560 | 0 | NA | NA | NA | NA | NA |
| CG42561 | 5.963526 | 1.517848 | 3.956843 | 0.383601 | 0.701274 | NA |
| CG42562 | 4.019638 | 2.623574 | 3.884633 | 0.675372 | 0.499439 | NA |
| CG42565 | 0 | NA | NA | NA | NA | NA |
| CG42566 | 0 | NA | NA | NA | NA | NA |
| CG42568 | 0 | NA | NA | NA | NA | NA |
| DnaJ-60 | 0 | NA | NA | NA | NA | NA |
| CG42570 | 0 | NA | NA | NA | NA | NA |
| CG42571 | 0 | NA | NA | NA | NA | NA |
| wisp | 7.180792 | -3.22342 | 3.855483 | -0.83606 | 0.40312 | NA |
| inaF-D | 0 | NA | NA | NA | NA | NA |
| CG42577 | 0 | NA | NA | NA | NA | NA |
| CG42578 | 0 | NA | NA | NA | NA | NA |
| CG42579 | 0 | NA | NA | NA | NA | NA |
| CG42580 | 0 | NA | NA | NA | NA | NA |
| CG42581 | 0 | NA | NA | NA | NA | NA |
| CG42583 | 0 | NA | NA | NA | NA | NA |
| cuff | 8.553988 | -1.01778 | 3.922987 | -0.25944 | 0.795295 | NA |
| bond | 0 | NA | NA | NA | NA | NA |
| CG42585 | 0 | NA | NA | NA | NA | NA |
| CG42586 | 0 | NA | NA | NA | NA | NA |
| CG42587 | 0 | NA | NA | NA | NA | NA |
| MCPH1 | 40.86847 | -2.95255 | 3.877731 | -0.76141 | 0.446411 | NA |
| CG42590 | 1.151699 | 1.609751 | 3.989673 | 0.403479 | 0.686596 | NA |
| CG42591 | 0 | NA | NA | NA | NA | NA |
| CG42592 | 0 | NA | NA | NA | NA | NA |
| mei-38 | 0 | NA | NA | NA | NA | NA |
| Incenp | 0.178812 | -1.08808 | 4.061735 | -0.26789 | 0.788788 | NA |
| CG42597 | 0 | NA | NA | NA | NA | NA |
| CG42598 | 0.178053 | -1.16467 | 4.075067 | -0.2858 | 0.775028 | NA |
| asl | 4.397397 | 1.710278 | 3.962238 | 0.431644 | 0.666 | NA |
| snoRNA:83 | 1.000346 | -0.36632 | 3.924715 | -0.09334 | 0.925636 | NA |
| clos | 3.807412 | -3.47991 | 3.840876 | -0.90602 | 0.364925 | NA |
| moi | 0 | NA | NA | NA | NA | NA |
| Cad86C | 12.17836 | -5.29011 | 3.829176 | -1.38153 | 0.167117 | NA |
| Sfp24Bc | 10.30387 | 0.277343 | 3.953034 | 0.070159 | 0.944067 | NA |
| Sfp26Ad | 0 | NA | NA | NA | NA | NA |
| Sfp36F | 0 | NA | NA | NA | NA | NA |
| Sfp38D | 0 | NA | NA | NA | NA | NA |
| Sfp78E | 0.597459 | 1.53156 | 3.979367 | 0.384875 | 0.70033 | NA |
| Sfp93F | 0 | NA | NA | NA | NA | NA |
| Sfp96F | 0 | NA | NA | NA | NA | NA |
| mrn | 29.23132 | -4.59508 | 3.781312 | -1.21521 | 0.224287 | NA |
| sing | 4.471231 | 1.30049 | 3.925251 | 0.331314 | 0.740408 | NA |
| zuc | 0 | NA | NA | NA | NA | NA |
| CG42615 | 0 | NA | NA | NA | NA | NA |

| | | | | | | |
|---|---|---|---|---|---|---|
| conv | 0.104623 | 0.639075 | 4.067674 | 0.157111 | 0.875158 | NA |
| Mat89Ba | 35.75245 | 4.870061 | 3.82557 | 1.273029 | 0.203008 | NA |
| CheB98a | 0 | NA | NA | NA | NA | NA |
| CheA84a | 0 | NA | NA | NA | NA | NA |
| CheA86a | 0 | NA | NA | NA | NA | NA |
| CheB53b | 35.61066 | -1.70495 | 3.951985 | -0.43142 | 0.666165 | NA |
| verm | 0.356107 | -1.47099 | 4.037764 | -0.36431 | 0.715628 | NA |
| Mst36Fb | 0.830552 | 2.118611 | 3.94541 | 0.536981 | 0.591281 | NA |
| Ccdc56 | 0 | NA | NA | NA | NA | NA |
| CG42633 | 0 | NA | NA | NA | NA | NA |
| CG42634 | 0 | NA | NA | NA | NA | NA |
| CG42635 | 0 | NA | NA | NA | NA | NA |
| CG42637 | 0 | NA | NA | NA | NA | NA |
| CG42638 | 0.829716 | 2.152816 | 3.960988 | 0.543505 | 0.586782 | NA |
| PPO3 | 0 | NA | NA | NA | NA | NA |
| mtTFB1 | 0 | NA | NA | NA | NA | NA |
| Gpa2 | 0 | NA | NA | NA | NA | NA |
| alpha-Est5 | 1.235713 | -0.14929 | 3.9675 | -0.03763 | 0.969984 | NA |
| Prosalpha3 | 0 | NA | NA | NA | NA | NA |
| Pp1-Y1 | 0.178053 | -1.16467 | 4.075067 | -0.2858 | 0.775028 | NA |
| Ir75c | 0 | NA | NA | NA | NA | NA |
| Ir75b | 0 | NA | NA | NA | NA | NA |
| lncRNA:CR | 40.20534 | -5.86724 | 3.82114 | -1.53547 | 0.124668 | NA |
| hkb | 1.351755 | -2.03446 | 3.928066 | -0.51793 | 0.604508 | NA |
| DhpD | 14.96136 | 3.644878 | 3.902642 | 0.933951 | 0.350329 | NA |
| CG13377 | 0.53416 | -1.53481 | 4.012395 | -0.38252 | 0.702078 | NA |
| scaRNA:M | 22.66263 | -0.13623 | 3.850084 | -0.03538 | 0.971774 | NA |
| scaRNA:M | 0.132078 | -1.13211 | 4.069918 | -0.27817 | 0.780885 | NA |
| scaRNA:M | 43.67276 | -0.05665 | 3.443182 | -0.01645 | 0.986872 | NA |
| scaRNA:M | 0.707123 | -2.68537 | 4.000063 | -0.67133 | 0.502009 | NA |
| CG42649 | 0 | NA | NA | NA | NA | NA |
| CG42650 | 0 | NA | NA | NA | NA | NA |
| lncRNA:CR | 14.60281 | 0.425506 | 3.752548 | 0.113391 | 0.90972 | NA |
| 7SLRNA:CF | 2.113346 | -2.98132 | 3.916163 | -0.76129 | 0.446486 | NA |
| CR42653 | 4.292697 | 1.709267 | 3.962502 | 0.431361 | 0.666206 | NA |
| CG42655 | 38.51182 | -0.35109 | 3.835396 | -0.09154 | 0.927064 | NA |
| NimA | 37.40553 | 0.004635 | 3.951466 | 0.001173 | 0.999064 | NA |
| lncRNA:CR | 0 | NA | NA | NA | NA | NA |
| CG42658 | 0 | NA | NA | NA | NA | NA |
| NT1 | 0 | NA | NA | NA | NA | NA |
| ms(2)34Fe | 0.356107 | -1.47099 | 4.037764 | -0.36431 | 0.715628 | NA |
| CG42659 | 0 | NA | NA | NA | NA | NA |
| l(2)34Fc | 0 | NA | NA | NA | NA | NA |
| l(2)34Fd | 4.273279 | -1.68149 | 3.959707 | -0.42465 | 0.671092 | NA |
| CG42660 | 0 | NA | NA | NA | NA | NA |
| CG42661 | 0 | NA | NA | NA | NA | NA |
| CG42662 | 0 | NA | NA | NA | NA | NA |
| CG42675 | 0 | NA | NA | NA | NA | NA |
| CG42680 | 0 | NA | NA | NA | NA | NA |
| CG42681 | 0 | NA | NA | NA | NA | NA |

| | | | | | | |
|---|---|---|---|---|---|---|
| CG42682 | 0 | NA | NA | NA | NA | NA |
| CG42685 | 3.552619 | -1.52136 | 3.905383 | -0.38956 | 0.696865 | NA |
| tobi | 0 | NA | NA | NA | NA | NA |
| CG42690 | 0 | NA | NA | NA | NA | NA |
| CG42691 | 0 | NA | NA | NA | NA | NA |
| CG42692 | 0 | NA | NA | NA | NA | NA |
| CG42693 | 0.178812 | -1.08808 | 4.061735 | -0.26789 | 0.788788 | NA |
| CG42694 | 0.178812 | -1.08808 | 4.061735 | -0.26789 | 0.788788 | NA |
| lncRNA:CR | 0 | NA | NA | NA | NA | NA |
| CG42697 | 0 | NA | NA | NA | NA | NA |
| CG42703 | 0 | NA | NA | NA | NA | NA |
| CG42704 | 0 | NA | NA | NA | NA | NA |
| CG42705 | 0 | NA | NA | NA | NA | NA |
| I-3 | 20.38222 | 6.791841 | 3.911947 | 1.736179 | 0.082532 | NA |
| CG42710 | 0.712213 | -1.57119 | 3.998613 | -0.39293 | 0.694369 | NA |
| CG42711 | 0 | NA | NA | NA | NA | NA |
| CG42713 | 0 | NA | NA | NA | NA | NA |
| CG42714 | 0 | NA | NA | NA | NA | NA |
| lncRNA:CR | 0 | NA | NA | NA | NA | NA |
| CG42716 | 0 | NA | NA | NA | NA | NA |
| CG42717 | 0 | NA | NA | NA | NA | NA |
| CG42718 | 0 | NA | NA | NA | NA | NA |
| lncRNA:CR | 0.209247 | 1.042804 | 4.047837 | 0.25762 | 0.7967 | NA |
| CR42722 | 0 | NA | NA | NA | NA | NA |
| CR42723 | 0 | NA | NA | NA | NA | NA |
| Npc1b | 1.479466 | -0.59427 | 3.946397 | -0.15059 | 0.880302 | NA |
| CG42726 | 0 | NA | NA | NA | NA | NA |
| CG42727 | 0 | NA | NA | NA | NA | NA |
| CG42728 | 0 | NA | NA | NA | NA | NA |
| CG42729 | 0 | NA | NA | NA | NA | NA |
| B9d2 | 0 | NA | NA | NA | NA | NA |
| CG42733 | 7.966422 | -3.11497 | 3.915222 | -0.79561 | 0.426261 | NA |
| tctn | 0 | NA | NA | NA | NA | NA |
| lncRNA:CR | 0 | NA | NA | NA | NA | NA |
| asRNA:CR4 | 0 | NA | NA | NA | NA | NA |
| CG42740 | 0 | NA | NA | NA | NA | NA |
| asRNA:CR4 | 0 | NA | NA | NA | NA | NA |
| asRNA:CR4 | 59.53063 | 1.53725 | 3.951513 | 0.389028 | 0.697255 | NA |
| lncRNA:CR | 36.31316 | -7.82544 | 3.911473 | -2.00064 | 0.045431 | NA |
| Cpn | 0 | NA | NA | NA | NA | NA |
| dsx-c73A | 9.76145 | -0.30999 | 3.87651 | -0.07997 | 0.936264 | NA |
| CG42747 | 39.04694 | 6.372122 | 3.869921 | 1.646577 | 0.099645 | NA |
| CG42749 | 0.932373 | 1.053996 | 3.996193 | 0.26375 | 0.791972 | NA |
| CG42752 | 0 | NA | NA | NA | NA | NA |
| CG42753 | 0.2094 | 1.005034 | 4.045341 | 0.248442 | 0.803792 | NA |
| lncRNA:CR | 0 | NA | NA | NA | NA | NA |
| lncRNA:CR | 0 | NA | NA | NA | NA | NA |
| CG42758 | 0 | NA | NA | NA | NA | NA |
| CG42759 | 44.7539 | 1.177044 | 3.95196 | 0.297838 | 0.765827 | NA |
| CG42762 | 0 | NA | NA | NA | NA | NA |

| | | | | | | |
|---|---|---|---|---|---|---|
| CG42763 | 0 | NA | NA | NA | NA | NA |
| lncRNA:CR | 3.452567 | 1.502955 | 3.961005 | 0.379438 | 0.704363 | NA |
| pre-mod(n | 0 | NA | NA | NA | NA | NA |
| pre-mod(n | 0 | NA | NA | NA | NA | NA |
| pre-mod(n | 0 | NA | NA | NA | NA | NA |
| pre-mod(n | 0 | NA | NA | NA | NA | NA |
| pre-mod(n | 0 | NA | NA | NA | NA | NA |
| pre-mod(n | 0 | NA | NA | NA | NA | NA |
| pre-mod(n | 0 | NA | NA | NA | NA | NA |
| pre-mod(n | 0 | NA | NA | NA | NA | NA |
| pre-mod(n | 0 | NA | NA | NA | NA | NA |
| CG42779 | 0 | NA | NA | NA | NA | NA |
| CG42780 | 0 | NA | NA | NA | NA | NA |
| CG42781 | 0 | NA | NA | NA | NA | NA |
| CG42782 | 0 | NA | NA | NA | NA | NA |
| CR42785 | 0 | NA | NA | NA | NA | NA |
| CR42786 | 0 | NA | NA | NA | NA | NA |
| Mzt1 | 0 | NA | NA | NA | NA | NA |
| lncRNA:CR | 0 | NA | NA | NA | NA | NA |
| lncRNA:CR | 0 | NA | NA | NA | NA | NA |
| CG42792 | 0 | NA | NA | NA | NA | NA |
| lncRNA:CR | 0 | NA | NA | NA | NA | NA |
| vnd | 0.945306 | -0.54642 | 3.960875 | -0.13795 | 0.890276 | NA |
| CG42798 | 0 | NA | NA | NA | NA | NA |
| srw | 0 | NA | NA | NA | NA | NA |
| snoRNA:Or | 0 | NA | NA | NA | NA | NA |
| snoRNA:Or | 0 | NA | NA | NA | NA | NA |
| Pxt | 0 | NA | NA | NA | NA | NA |
| CG42807 | 1.072115 | -2.75701 | 4.00642 | -0.68815 | 0.491359 | NA |
| CG42809 | 8.158263 | 1.163335 | 3.956616 | 0.294023 | 0.768741 | NA |
| CG42810 | 23.51641 | 2.298851 | 3.841718 | 0.598391 | 0.549579 | NA |
| CG42811 | 0 | NA | NA | NA | NA | NA |
| CG42812 | 0 | NA | NA | NA | NA | NA |
| CG42814 | 0 | NA | NA | NA | NA | NA |
| CG42815 | 0 | NA | NA | NA | NA | NA |
| CG42816 | 0.337716 | 1.464149 | 4.04412 | 0.362044 | 0.717319 | NA |
| CG42818 | 0.236778 | -0.26946 | 4.071573 | -0.06618 | 0.947234 | NA |
| CG42819 | 0 | NA | NA | NA | NA | NA |
| CG42820 | 0 | NA | NA | NA | NA | NA |
| CG42821 | 0.1047 | 0.613261 | 4.062799 | 0.150945 | 0.880019 | NA |
| CG42822 | 0 | NA | NA | NA | NA | NA |
| CG42823 | 0.178812 | -1.08808 | 4.061735 | -0.26789 | 0.788788 | NA |
| CG42824 | 0 | NA | NA | NA | NA | NA |
| CG42825 | 57.9821 | -1.92075 | 3.95158 | -0.48607 | 0.626916 | NA |
| CG42826 | 0 | NA | NA | NA | NA | NA |
| CG42828 | 35.97601 | -8.3268 | 3.908845 | -2.13025 | 0.033151 | NA |
| CG42830 | 2.773631 | -1.87679 | 3.964396 | -0.47341 | 0.63592 | NA |
| CG42831 | 0.64424 | -0.14539 | 3.993669 | -0.0364 | 0.97096 | NA |
| CG42832 | 0 | NA | NA | NA | NA | NA |
| CG42833 | 0 | NA | NA | NA | NA | NA |

| | | | | | | |
|---|---|---|---|---|---|---|
| CG42834 | 0 | NA | NA | NA | NA | NA |
| CG42835 | 0 | NA | NA | NA | NA | NA |
| lncRNA:CR | 0.178053 | -1.16467 | 4.075067 | -0.2858 | 0.775028 | NA |
| CG42837 | 0 | NA | NA | NA | NA | NA |
| CR42838 | 0 | NA | NA | NA | NA | NA |
| lncRNA:CR | 0.356107 | -1.47099 | 4.037764 | -0.36431 | 0.715628 | NA |
| d | 0 | NA | NA | NA | NA | NA |
| CG42841 | 1.109413 | -0.57335 | 3.886322 | -0.14753 | 0.882713 | NA |
| CR42842 | 0 | NA | NA | NA | NA | NA |
| lncRNA:CR | 0.132078 | -1.13211 | 4.069918 | -0.27817 | 0.780885 | NA |
| CG42846 | 35.64305 | 4.699317 | 3.829871 | 1.227017 | 0.219816 | NA |
| CG42847 | 0 | NA | NA | NA | NA | NA |
| lncRNA:CR | 0 | NA | NA | NA | NA | NA |
| CG42851 | 0 | NA | NA | NA | NA | NA |
| CG42852 | 2.314693 | -1.66033 | 3.966843 | -0.41855 | 0.675543 | NA |
| CG42853 | 0.178053 | -1.16467 | 4.075067 | -0.2858 | 0.775028 | NA |
| CG42854 | 0 | NA | NA | NA | NA | NA |
| CG42855 | 1.71701 | -1.85109 | 3.972145 | -0.46602 | 0.641203 | NA |
| CG42857 | 0 | NA | NA | NA | NA | NA |
| lncRNA:CR | 0 | NA | NA | NA | NA | NA |
| lncRNA:CR | 0.132078 | -1.13211 | 4.069918 | -0.27817 | 0.780885 | NA |
| asRNA:CR4 | 0.104623 | 0.639075 | 4.067674 | 0.157111 | 0.875158 | NA |
| f | 0 | NA | NA | NA | NA | NA |
| sro | 0.765012 | -1.03648 | 3.979317 | -0.26047 | 0.794504 | NA |
| CG42866 | 0.528311 | -1.72839 | 4.011871 | -0.43082 | 0.6666 | NA |
| CG42867 | 14.83604 | 1.618941 | 3.954063 | 0.409437 | 0.682219 | NA |
| lncRNA:CR | 0 | NA | NA | NA | NA | NA |
| CG42869 | 0 | NA | NA | NA | NA | NA |
| CG42870 | 0 | NA | NA | NA | NA | NA |
| lncRNA:CR | 15.00928 | 0.029243 | 3.737531 | 0.007824 | 0.993757 | NA |
| CG42876 | 0 | NA | NA | NA | NA | NA |
| betaNACte | 0 | NA | NA | NA | NA | NA |
| CG43049 | 0 | NA | NA | NA | NA | NA |
| CG43050 | 0.356107 | -1.47099 | 4.037764 | -0.36431 | 0.715628 | NA |
| CG43051 | 11.67348 | -7.01189 | 3.911387 | -1.79269 | 0.073023 | NA |
| CG43052 | 0 | NA | NA | NA | NA | NA |
| lncRNA:CR | 0 | NA | NA | NA | NA | NA |
| CG43054 | 0 | NA | NA | NA | NA | NA |
| CG43055 | 0 | NA | NA | NA | NA | NA |
| CG43057 | 0 | NA | NA | NA | NA | NA |
| CG43058 | 0 | NA | NA | NA | NA | NA |
| CG43059 | 0 | NA | NA | NA | NA | NA |
| CG43060 | 0 | NA | NA | NA | NA | NA |
| CG43061 | 0 | NA | NA | NA | NA | NA |
| CG43062 | 0 | NA | NA | NA | NA | NA |
| CG43063 | 0 | NA | NA | NA | NA | NA |
| CG43064 | 0 | NA | NA | NA | NA | NA |
| CG43068 | 28.3967 | -1.9183 | 3.952279 | -0.48537 | 0.627417 | NA |
| CG43069 | 0.178053 | -1.16467 | 4.075067 | -0.2858 | 0.775028 | NA |
| CG43070 | 0.1047 | 0.613261 | 4.062799 | 0.150945 | 0.880019 | NA |

| | | | | | | |
|---|---|---|---|---|---|---|
| CG43071 | 0 | NA | NA | NA | NA | NA |
| CG43072 | 0 | NA | NA | NA | NA | NA |
| CG43074 | 57.45379 | -1.92073 | 3.951586 | -0.48607 | 0.626921 | NA |
| CG43075 | 0 | NA | NA | NA | NA | NA |
| CG43076 | 0 | NA | NA | NA | NA | NA |
| CG43077 | 0 | NA | NA | NA | NA | NA |
| CR43080 | 0 | NA | NA | NA | NA | NA |
| Vha16-4 | 0 | NA | NA | NA | NA | NA |
| VhaPPA1-2 | 0 | NA | NA | NA | NA | NA |
| Trpml | 0 | NA | NA | NA | NA | NA |
| CG43082 | 0 | NA | NA | NA | NA | NA |
| CG43083 | 0 | NA | NA | NA | NA | NA |
| CG43084 | 0.104623 | 0.639075 | 4.067674 | 0.157111 | 0.875158 | NA |
| CG43088 | 0 | NA | NA | NA | NA | NA |
| CG43089 | 4.289554 | 1.509777 | 3.959091 | 0.381344 | 0.702948 | NA |
| CG43090 | 0 | NA | NA | NA | NA | NA |
| CG43094 | 37.60846 | 0.343026 | 3.839417 | 0.089343 | 0.928809 | NA |
| CG43095 | 55.28149 | 2.975539 | 3.918979 | 0.759264 | 0.447695 | NA |
| CR43096 | 0 | NA | NA | NA | NA | NA |
| CG43098 | 30.78178 | 1.747963 | 3.952592 | 0.442232 | 0.658321 | NA |
| CR43100 | 0 | NA | NA | NA | NA | NA |
| CG43101 | 0 | NA | NA | NA | NA | NA |
| lncRNA:CR | 0 | NA | NA | NA | NA | NA |
| CR43105 | 0.31387 | 1.261782 | 4.037145 | 0.312543 | 0.754628 | NA |
| CG43106 | 0 | NA | NA | NA | NA | NA |
| CG43107 | 41.31225 | -2.64231 | 3.737997 | -0.70688 | 0.479642 | NA |
| CG43108 | 2.092465 | 1.481166 | 3.9672 | 0.373353 | 0.708886 | NA |
| CG43109 | 0 | NA | NA | NA | NA | NA |
| CG43111 | 0.942299 | 1.584447 | 3.996979 | 0.396411 | 0.691802 | NA |
| CG43112 | 0 | NA | NA | NA | NA | NA |
| CG43114 | 0 | NA | NA | NA | NA | NA |
| CG43116 | 0 | NA | NA | NA | NA | NA |
| CG43117 | 10.96245 | -1.91078 | 3.954436 | -0.4832 | 0.628954 | NA |
| CG43120 | 0 | NA | NA | NA | NA | NA |
| CG43121 | 0 | NA | NA | NA | NA | NA |
| CG43123 | 0 | NA | NA | NA | NA | NA |
| CG43124 | 0 | NA | NA | NA | NA | NA |
| CG43127 | 0.1047 | 0.613261 | 4.062799 | 0.150945 | 0.880019 | NA |
| CheA46a | 0 | NA | NA | NA | NA | NA |
| CheA56a | 0 | NA | NA | NA | NA | NA |
| CG43131 | 0 | NA | NA | NA | NA | NA |
| lncRNA:CR | 0 | NA | NA | NA | NA | NA |
| CG43133 | 0 | NA | NA | NA | NA | NA |
| CG43134 | 0 | NA | NA | NA | NA | NA |
| CG43135 | 0 | NA | NA | NA | NA | NA |
| CG43136 | 3.921726 | -4.48744 | 3.92807 | -1.1424 | 0.253286 | NA |
| CG43137 | 0 | NA | NA | NA | NA | NA |
| CG43138 | 0 | NA | NA | NA | NA | NA |
| CG43139 | 0 | NA | NA | NA | NA | NA |
| CR43142 | 0 | NA | NA | NA | NA | NA |

| | | | | | | |
|---|---|---|---|---|---|---|
| DNAlig1 | 13.02091 | -1.82671 | 3.846843 | -0.47486 | 0.634886 | NA |
| lncRNA:CR | 0 | NA | NA | NA | NA | NA |
| CG43145 | 0 | NA | NA | NA | NA | NA |
| lncRNA:CR | 0.233093 | 0.692656 | 4.076246 | 0.169925 | 0.865069 | NA |
| CG43147 | 0.178053 | -1.16467 | 4.075067 | -0.2858 | 0.775028 | NA |
| Tmhs | 0 | NA | NA | NA | NA | NA |
| lncRNA:CR | 0 | NA | NA | NA | NA | NA |
| CG43153 | 0 | NA | NA | NA | NA | NA |
| CG43155 | 17.80533 | -1.70165 | 3.953057 | -0.43046 | 0.666859 | NA |
| CG43156 | 0 | NA | NA | NA | NA | NA |
| lncRNA:CR | 0.104623 | 0.639075 | 4.067674 | 0.157111 | 0.875158 | NA |
| CR43158 | 0 | NA | NA | NA | NA | NA |
| asRNA:CR4 | 5.025597 | 1.715508 | 3.960878 | 0.433113 | 0.664933 | NA |
| asRNA:CR4 | 0 | NA | NA | NA | NA | NA |
| CG43163 | 7.631301 | -1.99716 | 3.828612 | -0.52164 | 0.601921 | NA |
| CG43165 | 0 | NA | NA | NA | NA | NA |
| lncRNA:CR | 0 | NA | NA | NA | NA | NA |
| CG43167 | 0 | NA | NA | NA | NA | NA |
| CG43168 | 0 | NA | NA | NA | NA | NA |
| CG43169 | 0 | NA | NA | NA | NA | NA |
| CG43171 | 0 | NA | NA | NA | NA | NA |
| CG43172 | 0 | NA | NA | NA | NA | NA |
| CG43175 | 0.209247 | 1.042804 | 4.047837 | 0.25762 | 0.7967 | NA |
| CR43176 | 0 | NA | NA | NA | NA | NA |
| CR43177 | 24.56645 | -1.91756 | 3.952492 | -0.48515 | 0.627569 | NA |
| CG43178 | 0 | NA | NA | NA | NA | NA |
| CG43179 | 52.24526 | 1.750729 | 3.951902 | 0.443009 | 0.657759 | NA |
| CG43181 | 0 | NA | NA | NA | NA | NA |
| CG43182 | 0 | NA | NA | NA | NA | NA |
| CG43183 | 0.357625 | -1.47707 | 4.044794 | -0.36518 | 0.714979 | NA |
| CG43184 | 1.849088 | -1.85577 | 3.970721 | -0.46736 | 0.640241 | NA |
| CG43185 | 0 | NA | NA | NA | NA | NA |
| CR43186 | 12.33581 | 0.034503 | 3.952607 | 0.008729 | 0.993035 | NA |
| CG43187 | 0.178053 | -1.16467 | 4.075067 | -0.2858 | 0.775028 | NA |
| CG43188 | 0 | NA | NA | NA | NA | NA |
| CG43190 | 0 | NA | NA | NA | NA | NA |
| CG43192 | 9.789916 | 1.166086 | 3.955676 | 0.294788 | 0.768156 | NA |
| lncRNA:CR | 0.356107 | -1.47099 | 4.037764 | -0.36431 | 0.715628 | NA |
| hemo | 0 | NA | NA | NA | NA | NA |
| CG43195 | 0 | NA | NA | NA | NA | NA |
| CG43198 | 0 | NA | NA | NA | NA | NA |
| CG43200 | 0 | NA | NA | NA | NA | NA |
| CG43201 | 0 | NA | NA | NA | NA | NA |
| BomT1 | 0 | NA | NA | NA | NA | NA |
| CG43203 | 1.056573 | -1.55533 | 3.944973 | -0.39426 | 0.693391 | NA |
| CG43204 | 7.22589 | 1.161221 | 3.95734 | 0.293435 | 0.76919 | NA |
| CG43206 | 0 | NA | NA | NA | NA | NA |
| CG43207 | 0 | NA | NA | NA | NA | NA |
| CG43208 | 0 | NA | NA | NA | NA | NA |
| CG43209 | 0.104623 | 0.639075 | 4.067674 | 0.157111 | 0.875158 | NA |

| | | | | | | |
|---|---|---|---|---|---|---|
| CG43210 | 0 | NA | NA | NA | NA | NA |
| CR43211 | 0 | NA | NA | NA | NA | NA |
| CR43212 | 6.793086 | -0.22491 | 3.755522 | -0.05989 | 0.952246 | NA |
| CR43213 | 0 | NA | NA | NA | NA | NA |
| CR43214 | 0 | NA | NA | NA | NA | NA |
| CR43215 | 0 | NA | NA | NA | NA | NA |
| CR43216 | 0 | NA | NA | NA | NA | NA |
| CR43217 | 0 | NA | NA | NA | NA | NA |
| CG43218 | 0.3141 | 1.240997 | 4.03841 | 0.307299 | 0.758616 | NA |
| CG43220 | 0 | NA | NA | NA | NA | NA |
| CG43221 | 0 | NA | NA | NA | NA | NA |
| Diedel2 | 0 | NA | NA | NA | NA | NA |
| CG43230 | 0 | NA | NA | NA | NA | NA |
| CG43232 | 1.633826 | -0.93012 | 3.966011 | -0.23452 | 0.81458 | NA |
| CG43233 | 0.523193 | 2.185419 | 4.000542 | 0.546281 | 0.584873 | NA |
| lncRNA:CR | 0 | NA | NA | NA | NA | NA |
| CG43235 | 0.104623 | 0.639075 | 4.067674 | 0.157111 | 0.875158 | NA |
| Mtkl | 0 | NA | NA | NA | NA | NA |
| CG43237 | 0.5235 | 1.490714 | 4.02555 | 0.370313 | 0.711149 | NA |
| lncRNA:CR | 0 | NA | NA | NA | NA | NA |
| CR43240 | 0 | NA | NA | NA | NA | NA |
| asRNA:CR4 | 0.356107 | -1.47099 | 4.037764 | -0.36431 | 0.715628 | NA |
| CG43244 | 0 | NA | NA | NA | NA | NA |
| CG43246 | 0 | NA | NA | NA | NA | NA |
| lncRNA:CR | 0 | NA | NA | NA | NA | NA |
| CG43248 | 0.31387 | 1.261782 | 4.037145 | 0.312543 | 0.754628 | NA |
| CG43249 | 0 | NA | NA | NA | NA | NA |
| lncRNA:CR | 0 | NA | NA | NA | NA | NA |
| CG43251 | 0 | NA | NA | NA | NA | NA |
| lncRNA:CR | 0 | NA | NA | NA | NA | NA |
| lncRNA:CR | 0 | NA | NA | NA | NA | NA |
| CG43254 | 0 | NA | NA | NA | NA | NA |
| asRNA:CR4 | 0 | NA | NA | NA | NA | NA |
| lncRNA:CR | 0.104623 | 0.639075 | 4.067674 | 0.157111 | 0.875158 | NA |
| lncRNA:CR | 0 | NA | NA | NA | NA | NA |
| lncRNA:CR | 0 | NA | NA | NA | NA | NA |
| lncRNA:CR | 0 | NA | NA | NA | NA | NA |
| lncRNA:CR | 33.28143 | -1.76241 | 3.952832 | -0.44586 | 0.655698 | NA |
| CG43267 | 0 | NA | NA | NA | NA | NA |
| lncRNA:CR | 0 | NA | NA | NA | NA | NA |
| CG43271 | 0 | NA | NA | NA | NA | NA |
| CG43272 | 0.178812 | -1.08808 | 4.061735 | -0.26789 | 0.788788 | NA |
| CG43273 | 22.18905 | -1.91696 | 3.952661 | -0.48498 | 0.62769 | NA |
| lncRNA:CR | 0 | NA | NA | NA | NA | NA |
| CR43275 | 19.45993 | 1.532694 | 3.952753 | 0.387754 | 0.698198 | NA |
| CR43276 | 22.532 | 2.734609 | 3.820367 | 0.715797 | 0.474116 | NA |
| CG43277 | 0 | NA | NA | NA | NA | NA |
| lncRNA:CR | 0.132078 | -1.13211 | 4.069918 | -0.27817 | 0.780885 | NA |
| lncRNA:CR | 1.71701 | -1.85109 | 3.972145 | -0.46602 | 0.641203 | NA |
| lncRNA:CR | 0 | NA | NA | NA | NA | NA |

| | | | | | | |
|---|---|---|---|---|---|---|
| CR43281 | 0 | NA | NA | NA | NA | NA |
| lncRNA:CR | 0 | NA | NA | NA | NA | NA |
| lncRNA:ch | 28.60885 | -2.67685 | 3.849013 | -0.69546 | 0.486764 | NA |
| lncRNA:CR | 0 | NA | NA | NA | NA | NA |
| lncRNA:CR | 26.85773 | 4.170803 | 3.803627 | 1.096533 | 0.272846 | NA |
| CG43288 | 0 | NA | NA | NA | NA | NA |
| lncRNA:CR | 0.660388 | -1.75977 | 4.001266 | -0.4398 | 0.660079 | NA |
| CG43290 | 0 | NA | NA | NA | NA | NA |
| CG43291 | 0 | NA | NA | NA | NA | NA |
| CG43292 | 0 | NA | NA | NA | NA | NA |
| CG43293 | 14.65799 | 1.740703 | 3.954419 | 0.440192 | 0.659798 | NA |
| CG43294 | 0 | NA | NA | NA | NA | NA |
| CG43296 | 0 | NA | NA | NA | NA | NA |
| lncRNA:CR | 0 | NA | NA | NA | NA | NA |
| lncRNA:CR | 0 | NA | NA | NA | NA | NA |
| asRNA:CR4 | 0.178053 | -1.16467 | 4.075067 | -0.2858 | 0.775028 | NA |
| lncRNA:CR | 0 | NA | NA | NA | NA | NA |
| lncRNA:CR | 0 | NA | NA | NA | NA | NA |
| lncRNA:CR | 4.603653 | 3.529285 | 3.923082 | 0.899621 | 0.368322 | NA |
| lncRNA:CR | 0 | NA | NA | NA | NA | NA |
| CG43307 | 0 | NA | NA | NA | NA | NA |
| CG43308 | 0 | NA | NA | NA | NA | NA |
| CR43309 | 10.43414 | -1.91017 | 3.954613 | -0.48302 | 0.629079 | NA |
| CR43310 | 3.017037 | 1.618643 | 3.847755 | 0.420672 | 0.673995 | NA |
| CG43312 | 9.712619 | 2.649761 | 3.874158 | 0.683958 | 0.494002 | NA |
| snRNA:U4a | 0.132078 | -1.13211 | 4.069918 | -0.27817 | 0.780885 | NA |
| lncRNA:CR | 4.473346 | -0.84823 | 3.837462 | -0.22104 | 0.825062 | NA |
| CG43315 | 0 | NA | NA | NA | NA | NA |
| CG43316 | 0 | NA | NA | NA | NA | NA |
| CG43317 | 26.0193 | -1.91787 | 3.952404 | -0.48524 | 0.627506 | NA |
| CG43318 | 0 | NA | NA | NA | NA | NA |
| CG43319 | 0 | NA | NA | NA | NA | NA |
| CG43321 | 0 | NA | NA | NA | NA | NA |
| CG43325 | 0 | NA | NA | NA | NA | NA |
| CG43326 | 0 | NA | NA | NA | NA | NA |
| CG43327 | 0 | NA | NA | NA | NA | NA |
| CG43328 | 0 | NA | NA | NA | NA | NA |
| CG43329 | 27.62154 | 1.898482 | 3.856334 | 0.492302 | 0.622506 | NA |
| CG43330 | 0 | NA | NA | NA | NA | NA |
| CG43331 | 0 | NA | NA | NA | NA | NA |
| CG43332 | 0.178812 | -1.08808 | 4.061735 | -0.26789 | 0.788788 | NA |
| CG43333 | 0.178053 | -1.16467 | 4.075067 | -0.2858 | 0.775028 | NA |
| CG43335 | 0 | NA | NA | NA | NA | NA |
| CG43336 | 0 | NA | NA | NA | NA | NA |
| CG43337 | 7.665106 | 2.834977 | 3.918606 | 0.723466 | 0.469394 | NA |
| CG43341 | 0 | NA | NA | NA | NA | NA |
| CG43342 | 55.35544 | 8.230467 | 3.901873 | 2.109363 | 0.034913 | NA |
| CG43345 | 0 | NA | NA | NA | NA | NA |
| CG43346 | 0 | NA | NA | NA | NA | NA |
| Klp54D | 2.346923 | -0.0014 | 3.913938 | -0.00036 | 0.999715 | NA |

| | | | | | | |
|---|---|---|---|---|---|---|
| CG43339 | 0 | NA | NA | NA | NA | NA |
| CG43338 | 0.536437 | -1.71175 | 4.038042 | -0.42391 | 0.671635 | NA |
| CG43348 | 0 | NA | NA | NA | NA | NA |
| CG43349 | 0 | NA | NA | NA | NA | NA |
| CG43350 | 0 | NA | NA | NA | NA | NA |
| CG43351 | 0 | NA | NA | NA | NA | NA |
| HP1Lcsd | 0 | NA | NA | NA | NA | NA |
| lncRNA:CR | 0 | NA | NA | NA | NA | NA |
| CG43354 | 0 | NA | NA | NA | NA | NA |
| CG43355 | 0 | NA | NA | NA | NA | NA |
| lncRNA:CR | 0 | NA | NA | NA | NA | NA |
| lncRNA:CR | 0 | NA | NA | NA | NA | NA |
| lncRNA:CR | 0 | NA | NA | NA | NA | NA |
| CR43359 | 0.462248 | -0.67622 | 4.039367 | -0.16741 | 0.867049 | NA |
| CR43360 | 0 | NA | NA | NA | NA | NA |
| CR43361 | 2.735658 | 4.81469 | 3.922071 | 1.227589 | 0.219601 | NA |
| CG43362 | 0 | NA | NA | NA | NA | NA |
| CR43364 | 0 | NA | NA | NA | NA | NA |
| CG43366 | 1.835442 | 2.985712 | 3.902254 | 0.765125 | 0.444197 | NA |
| Cc2d2a | 7.154728 | 4.211453 | 3.881569 | 1.084987 | 0.277927 | NA |
| tx | 0 | NA | NA | NA | NA | NA |
| Prosalpha1 | 0 | NA | NA | NA | NA | NA |
| lncRNA:CR | 0.178053 | -1.16467 | 4.075067 | -0.2858 | 0.775028 | NA |
| vret | 0.53416 | -1.53481 | 4.012395 | -0.38252 | 0.702078 | NA |
| scaRNA:M | 0 | NA | NA | NA | NA | NA |
| CG43376 | 0 | NA | NA | NA | NA | NA |
| CR43377 | 0 | NA | NA | NA | NA | NA |
| CR43378 | 0 | NA | NA | NA | NA | NA |
| CR43379 | 0 | NA | NA | NA | NA | NA |
| CR43380 | 0 | NA | NA | NA | NA | NA |
| CR43381 | 0 | NA | NA | NA | NA | NA |
| CR43382 | 0 | NA | NA | NA | NA | NA |
| CR43383 | 0 | NA | NA | NA | NA | NA |
| CR43384 | 0 | NA | NA | NA | NA | NA |
| CR43385 | 0 | NA | NA | NA | NA | NA |
| CG43386 | 36.2325 | 0.205367 | 3.72082 | 0.055194 | 0.955984 | NA |
| lncRNA:CR | 0 | NA | NA | NA | NA | NA |
| robls54B | 26.38438 | 1.74685 | 3.952871 | 0.441919 | 0.658548 | NA |
| Phae1 | 0.1047 | 0.613261 | 4.062799 | 0.150945 | 0.880019 | NA |
| Phae2 | 0 | NA | NA | NA | NA | NA |
| dar1 | 10.01121 | -4.6891 | 3.916556 | -1.19725 | 0.231209 | NA |
| Coop | 0.1047 | 0.613261 | 4.062799 | 0.150945 | 0.880019 | NA |
| CG43389 | 0 | NA | NA | NA | NA | NA |
| CG43390 | 0 | NA | NA | NA | NA | NA |
| CG43391 | 0 | NA | NA | NA | NA | NA |
| CG43392 | 0 | NA | NA | NA | NA | NA |
| CG43393 | 0 | NA | NA | NA | NA | NA |
| vnc | 0 | NA | NA | NA | NA | NA |
| chas | 27.06311 | -1.08588 | 3.823342 | -0.28401 | 0.776399 | NA |
| CG43396 | 49.10427 | 1.75047 | 3.951966 | 0.442936 | 0.657812 | NA |

| | | | | | | |
|---|---|---|---|---|---|---|
| CG43397 | 0 | NA | NA | NA | NA | NA |
| lncRNA:CR | 34.13531 | 1.47028 | 3.813895 | 0.385506 | 0.699862 | NA |
| CG43400 | 0 | NA | NA | NA | NA | NA |
| CG43401 | 0 | NA | NA | NA | NA | NA |
| CG43402 | 0 | NA | NA | NA | NA | NA |
| CG43403 | 0 | NA | NA | NA | NA | NA |
| lncRNA:CR | 0 | NA | NA | NA | NA | NA |
| CG43405 | 0.178053 | -1.16467 | 4.075067 | -0.2858 | 0.775028 | NA |
| CG43407 | 0 | NA | NA | NA | NA | NA |
| lncRNA:CR | 0 | NA | NA | NA | NA | NA |
| lncRNA:CR | 0 | NA | NA | NA | NA | NA |
| lncRNA:CR | 3.455098 | 1.699144 | 3.965156 | 0.428519 | 0.668273 | NA |
| lncRNA:CR | 0 | NA | NA | NA | NA | NA |
| asRNA:CR4 | 56.58737 | 1.153545 | 3.951728 | 0.291909 | 0.770356 | NA |
| lncRNA:CR | 0 | NA | NA | NA | NA | NA |
| lncRNA:CR | 0 | NA | NA | NA | NA | NA |
| lncRNA:CR | 0 | NA | NA | NA | NA | NA |
| lncRNA:CR | 0 | NA | NA | NA | NA | NA |
| lncRNA:CR | 0 | NA | NA | NA | NA | NA |
| lncRNA:CR | 1.675199 | 1.648777 | 3.978727 | 0.414398 | 0.678583 | NA |
| lncRNA:CR | 0 | NA | NA | NA | NA | NA |
| lncRNA:CR | 0 | NA | NA | NA | NA | NA |
| lncRNA:CR | 0 | NA | NA | NA | NA | NA |
| lncRNA:CR | 0 | NA | NA | NA | NA | NA |
| lncRNA:CR | 0 | NA | NA | NA | NA | NA |
| asRNA:CR4 | 0 | NA | NA | NA | NA | NA |
| asRNA:CR4 | 52.51587 | 0.185741 | 3.83494 | 0.048434 | 0.961371 | NA |
| CG43428 | 2.411839 | -0.34285 | 3.871222 | -0.08856 | 0.929428 | NA |
| asRNA:CR4 | 0 | NA | NA | NA | NA | NA |
| asRNA:CR4 | 0 | NA | NA | NA | NA | NA |
| lncRNA:CR | 19.36949 | 1.74405 | 3.953574 | 0.441132 | 0.659117 | NA |
| lncRNA:CR | 0 | NA | NA | NA | NA | NA |
| asRNA:CR4 | 4.719178 | 3.907158 | 3.916764 | 0.997547 | 0.318499 | NA |
| lncRNA:CR | 0 | NA | NA | NA | NA | NA |
| lncRNA:CR | 0 | NA | NA | NA | NA | NA |
| lncRNA:CR | 0 | NA | NA | NA | NA | NA |
| lncRNA:CR | 0 | NA | NA | NA | NA | NA |
| CG43438 | 0 | NA | NA | NA | NA | NA |
| CG43439 | 0 | NA | NA | NA | NA | NA |
| lncRNA:CR | 0.356107 | -1.47099 | 4.037764 | -0.36431 | 0.715628 | NA |
| CR43441 | 0 | NA | NA | NA | NA | NA |
| Nha2 | 0.104623 | 0.639075 | 4.067674 | 0.157111 | 0.875158 | NA |
| CG43445 | 29.05709 | -1.91841 | 3.952248 | -0.4854 | 0.627395 | NA |
| CG43446 | 34.10726 | 2.77171 | 3.917435 | 0.707532 | 0.479236 | NA |
| CG43447 | 0 | NA | NA | NA | NA | NA |
| CG43448 | 0.2094 | 1.005034 | 4.045341 | 0.248442 | 0.803792 | NA |
| CG43449 | 0 | NA | NA | NA | NA | NA |
| CG43450 | 0 | NA | NA | NA | NA | NA |
| lncRNA:CR | 0.178053 | -1.16467 | 4.075067 | -0.2858 | 0.775028 | NA |
| lncRNA:CR | 39.12912 | 1.190072 | 3.740117 | 0.318191 | 0.75034 | NA |

| | | | | | | |
|---|---|---|---|---|---|---|
| lncRNA:CR | 0.1047 | 0.613261 | 4.062799 | 0.150945 | 0.880019 | NA |
| asRNA:CR4 | 0 | NA | NA | NA | NA | NA |
| lncRNA:CR | 0 | NA | NA | NA | NA | NA |
| lncRNA:CR | 0 | NA | NA | NA | NA | NA |
| lncRNA:CR | 0 | NA | NA | NA | NA | NA |
| lncRNA:CR | 0 | NA | NA | NA | NA | NA |
| lncRNA:CR | 58.22255 | -2.15672 | 3.95241 | -0.54567 | 0.585291 | NA |
| asRNA:CR4 | 0 | NA | NA | NA | NA | NA |
| asRNA:CR4 | 0 | NA | NA | NA | NA | NA |
| CG5819 | 0 | NA | NA | NA | NA | NA |
| pgc | 14.7066 | -3.4891 | 3.904725 | -0.89356 | 0.371559 | NA |
| CG11298 | 0.356866 | -1.92326 | 4.037889 | -0.4763 | 0.633858 | NA |
| CG11362 | 0.178053 | -1.16467 | 4.075067 | -0.2858 | 0.775028 | NA |
| CG10344 | 0 | NA | NA | NA | NA | NA |
| babos | 0 | NA | NA | NA | NA | NA |
| T3dh | 32.75527 | -1.91894 | 3.952097 | -0.48555 | 0.627287 | NA |
| CG3927 | 0 | NA | NA | NA | NA | NA |
| CG4269 | 0.1047 | 0.613261 | 4.062799 | 0.150945 | 0.880019 | NA |
| nsr | 0.283512 | -0.2331 | 4.065094 | -0.05734 | 0.954273 | NA |
| CG13510 | 0 | NA | NA | NA | NA | NA |
| CG13511 | 0 | NA | NA | NA | NA | NA |
| CG4250 | 0.31089 | -1.9003 | 4.033427 | -0.47114 | 0.637542 | NA |
| CG30273 | 0 | NA | NA | NA | NA | NA |
| CG30269 | 0 | NA | NA | NA | NA | NA |
| Obp59a | 0 | NA | NA | NA | NA | NA |
| Obp58d | 0 | NA | NA | NA | NA | NA |
| CG30275 | 0.104623 | 0.639075 | 4.067674 | 0.157111 | 0.875158 | NA |
| CG30268 | 0.236701 | -0.24276 | 4.020028 | -0.06039 | 0.951846 | NA |
| CG13526 | 2.198698 | 1.671135 | 3.972627 | 0.420662 | 0.674002 | NA |
| asRNA:CR3 | 1.264786 | 0.992839 | 3.920274 | 0.253257 | 0.800069 | NA |
| CG30271 | 32.09096 | -1.33351 | 3.951859 | -0.33744 | 0.735787 | NA |
| MFS1 | 0 | NA | NA | NA | NA | NA |
| CG30265 | 0 | NA | NA | NA | NA | NA |
| CG12490 | 0 | NA | NA | NA | NA | NA |
| CG9825 | 0 | NA | NA | NA | NA | NA |
| CG9826 | 0 | NA | NA | NA | NA | NA |
| CG3649 | 0 | NA | NA | NA | NA | NA |
| MED23 | 4.910335 | 4.391786 | 3.877589 | 1.132608 | 0.257379 | NA |
| inaD | 1.412728 | -3.67881 | 3.95877 | -0.92928 | 0.352743 | NA |
| CycB | 0 | NA | NA | NA | NA | NA |
| Obp58c | 0 | NA | NA | NA | NA | NA |
| CG30259 | 0.53416 | -1.53481 | 4.012395 | -0.38252 | 0.702078 | NA |
| CG30196 | 0 | NA | NA | NA | NA | NA |
| Cyp6d2 | 0 | NA | NA | NA | NA | NA |
| CG3746 | 0 | NA | NA | NA | NA | NA |
| CG30195 | 0 | NA | NA | NA | NA | NA |
| CG4329 | 0 | NA | NA | NA | NA | NA |
| tRNA:Ser-T | 0 | NA | NA | NA | NA | NA |
| CG3085 | 4.816197 | 1.713907 | 3.961293 | 0.432664 | 0.665259 | NA |
| Gr59a | 0 | NA | NA | NA | NA | NA |

| | | | | | | |
|---|---|---|---|---|---|---|
| Gr59b | 0.712213 | -1.57119 | 3.998613 | -0.39293 | 0.694369 | NA |
| CG30192 | 0 | NA | NA | NA | NA | NA |
| Klp59D | 0 | NA | NA | NA | NA | NA |
| CG9899 | 0 | NA | NA | NA | NA | NA |
| CG13545 | 0 | NA | NA | NA | NA | NA |
| CG13544 | 0.178812 | -1.08808 | 4.061735 | -0.26789 | 0.788788 | NA |
| Gpdh2 | 3.258635 | -0.08803 | 3.89017 | -0.02263 | 0.981947 | NA |
| Klp59C | 0 | NA | NA | NA | NA | NA |
| Gr59d | 0.233093 | 0.692656 | 4.076246 | 0.169925 | 0.865069 | NA |
| Gr59c | 0 | NA | NA | NA | NA | NA |
| RpL37b | 8.849205 | -1.9079 | 3.955267 | -0.48237 | 0.629543 | NA |
| CG9876 | 0 | NA | NA | NA | NA | NA |
| CG9877 | 0.178053 | -1.16467 | 4.075067 | -0.2858 | 0.775028 | NA |
| Fib | 58.69368 | -1.6645 | 3.477177 | -0.47869 | 0.632158 | NA |
| tRNA:Ser-T | 0 | NA | NA | NA | NA | NA |
| CG9897 | 0 | NA | NA | NA | NA | NA |
| CG32833 | 0.178053 | -1.16467 | 4.075067 | -0.2858 | 0.775028 | NA |
| CG32834 | 0 | NA | NA | NA | NA | NA |
| lncRNA:CR | 0 | NA | NA | NA | NA | NA |
| CG3788 | 56.8878 | -1.20427 | 3.951673 | -0.30475 | 0.760558 | NA |
| CG30187 | 0.651586 | 2.224386 | 4.007948 | 0.554994 | 0.578899 | NA |
| CG3092 | 0 | NA | NA | NA | NA | NA |
| CG13540 | 0 | NA | NA | NA | NA | NA |
| CG13541 | 0.178053 | -1.16467 | 4.075067 | -0.2858 | 0.775028 | NA |
| CG9863 | 0 | NA | NA | NA | NA | NA |
| CR30409 | 0 | NA | NA | NA | NA | NA |
| CG9875 | 0 | NA | NA | NA | NA | NA |
| yellow-d | 0 | NA | NA | NA | NA | NA |
| yellow-d2 | 0.178053 | -1.16467 | 4.075067 | -0.2858 | 0.775028 | NA |
| CG30411 | 0 | NA | NA | NA | NA | NA |
| CG13557 | 0 | NA | NA | NA | NA | NA |
| Or59c | 0 | NA | NA | NA | NA | NA |
| Or59b | 0.178053 | -1.16467 | 4.075067 | -0.2858 | 0.775028 | NA |
| SkpF | 0.104623 | 0.639075 | 4.067674 | 0.157111 | 0.875158 | NA |
| Or59a | 0 | NA | NA | NA | NA | NA |
| CG9815 | 0 | NA | NA | NA | NA | NA |
| CG9812 | 0.341324 | 0.112818 | 4.000379 | 0.028202 | 0.977501 | NA |
| CG30414 | 0.178053 | -1.16467 | 4.075067 | -0.2858 | 0.775028 | NA |
| CG3502 | 0.178053 | -1.16467 | 4.075067 | -0.2858 | 0.775028 | NA |
| CG30413 | 0 | NA | NA | NA | NA | NA |
| tbrd-3 | 0 | NA | NA | NA | NA | NA |
| CG30412 | 0.178053 | -1.16467 | 4.075067 | -0.2858 | 0.775028 | NA |
| Prosbeta5l | 0 | NA | NA | NA | NA | NA |
| CG3124 | 25.87227 | -7.09074 | 3.888022 | -1.82374 | 0.068191 | NA |
| ord | 0 | NA | NA | NA | NA | NA |
| CG12782 | 0 | NA | NA | NA | NA | NA |
| RpL22-like | 0 | NA | NA | NA | NA | NA |
| yip3 | 0 | NA | NA | NA | NA | NA |
| LS2 | 17.1701 | -1.91519 | 3.95317 | -0.48447 | 0.628053 | NA |
| CG13559 | 0.178053 | -1.16467 | 4.075067 | -0.2858 | 0.775028 | NA |

| | | | | | | |
|---|---|---|---|---|---|---|
| Gr59f | 0.660388 | -1.75977 | 4.001266 | -0.4398 | 0.660079 | NA |
| ItgaPS5 | 0 | NA | NA | NA | NA | NA |
| CG30184 | 0 | NA | NA | NA | NA | NA |
| apt | 0.596853 | 0.454699 | 3.988272 | 0.114009 | 0.909231 | NA |
| Eglp1 | 0 | NA | NA | NA | NA | NA |
| retn | 0.44234 | 1.822487 | 4.025091 | 0.452781 | 0.650706 | NA |
| St1 | 7.458983 | 1.161798 | 3.957142 | 0.293595 | 0.769067 | NA |
| CG13560 | 0 | NA | NA | NA | NA | NA |
| CG12491 | 0 | NA | NA | NA | NA | NA |
| CG11300 | 0 | NA | NA | NA | NA | NA |
| CG18128 | 0 | NA | NA | NA | NA | NA |
| Eglp4 | 0 | NA | NA | NA | NA | NA |
| Eglp3 | 0 | NA | NA | NA | NA | NA |
| Eglp2 | 0 | NA | NA | NA | NA | NA |
| Gr59e | 0 | NA | NA | NA | NA | NA |
| CG3906 | 0 | NA | NA | NA | NA | NA |
| CG30177 | 0.283512 | -0.2331 | 4.065094 | -0.05734 | 0.954273 | NA |
| CG5539 | 0.1047 | 0.613261 | 4.062799 | 0.150945 | 0.880019 | NA |
| CG5597 | 0.4188 | 1.41068 | 4.034613 | 0.349644 | 0.726606 | NA |
| CG17658 | 0 | NA | NA | NA | NA | NA |
| CG13562 | 0 | NA | NA | NA | NA | NA |
| CG2812 | 0 | NA | NA | NA | NA | NA |
| fzr2 | 0.388212 | 0.094116 | 4.038892 | 0.023303 | 0.981409 | NA |
| Fmo-1 | 0.178053 | -1.16467 | 4.075067 | -0.2858 | 0.775028 | NA |
| CG30178 | 0 | NA | NA | NA | NA | NA |
| Rsph4a | 0 | NA | NA | NA | NA | NA |
| CG4324 | 0.236778 | -0.26946 | 4.071573 | -0.06618 | 0.947234 | NA |
| CG13566 | 0 | NA | NA | NA | NA | NA |
| bgcn | 0.819872 | -0.85327 | 3.993249 | -0.21368 | 0.830798 | NA |
| shu | 0 | NA | NA | NA | NA | NA |
| CG4763 | 0 | NA | NA | NA | NA | NA |
| CG4797 | 0 | NA | NA | NA | NA | NA |
| CG15800 | 0 | NA | NA | NA | NA | NA |
| CG13564 | 0.178053 | -1.16467 | 4.075067 | -0.2858 | 0.775028 | NA |
| CG16786 | 3.320463 | -2.57495 | 3.942954 | -0.65305 | 0.513723 | NA |
| snoRNA:Ps | 0 | NA | NA | NA | NA | NA |
| CG13575 | 5.387731 | -1.69677 | 3.957058 | -0.4288 | 0.668072 | NA |
| slbo | 0.178053 | -1.16467 | 4.075067 | -0.2858 | 0.775028 | NA |
| CG13577 | 0 | NA | NA | NA | NA | NA |
| Ir60a | 0.356107 | -1.47099 | 4.037764 | -0.36431 | 0.715628 | NA |
| CG10339 | 0.356107 | -1.47099 | 4.037764 | -0.36431 | 0.715628 | NA |
| CG3483 | 0 | NA | NA | NA | NA | NA |
| CG3492 | 15.22063 | -3.95725 | 3.887164 | -1.01803 | 0.308663 | NA |
| CG3494 | 0 | NA | NA | NA | NA | NA |
| CG16837 | 0.356107 | -1.47099 | 4.037764 | -0.36431 | 0.715628 | NA |
| Crtp | 0 | NA | NA | NA | NA | NA |
| CG13581 | 0 | NA | NA | NA | NA | NA |
| CG11413 | 0 | NA | NA | NA | NA | NA |
| uri | 42.21116 | 5.442916 | 3.832312 | 1.42027 | 0.155529 | NA |
| CG13587 | 0 | NA | NA | NA | NA | NA |

| | | | | | | |
|---|---|---|---|---|---|---|
| Samtor | 3.947251 | 1.355039 | 3.960575 | 0.342132 | 0.732252 | NA |
| Adck1 | 47.76895 | -0.36862 | 3.83539 | -0.09611 | 0.923433 | NA |
| Cyp9c1 | 0 | NA | NA | NA | NA | NA |
| CG3640 | 0 | NA | NA | NA | NA | NA |
| CG30161 | 0 | NA | NA | NA | NA | NA |
| Cpr60D | 57.3494 | -0.33305 | 3.835187 | -0.08684 | 0.930798 | NA |
| CG4781 | 0.178053 | -1.16467 | 4.075067 | -0.2858 | 0.775028 | NA |
| CG4741 | 4.290956 | -1.83003 | 3.783302 | -0.48371 | 0.628591 | NA |
| CG4681 | 0.792466 | -1.7823 | 3.993845 | -0.44626 | 0.655408 | NA |
| Ir60e | 0 | NA | NA | NA | NA | NA |
| Prosalpha4 | 0 | NA | NA | NA | NA | NA |
| Ssl | 0 | NA | NA | NA | NA | NA |
| CG13590 | 0 | NA | NA | NA | NA | NA |
| CG13589 | 0 | NA | NA | NA | NA | NA |
| CG4563 | 16.75191 | 2.95528 | 3.92353 | 0.75322 | 0.451318 | NA |
| Pgam5-2 | 0 | NA | NA | NA | NA | NA |
| CG15873 | 0.1047 | 0.613261 | 4.062799 | 0.150945 | 0.880019 | NA |
| spz6 | 26.03894 | 1.673283 | 3.952785 | 0.423318 | 0.672064 | NA |
| CG3880 | 0 | NA | NA | NA | NA | NA |
| CG3650 | 0.1047 | 0.613261 | 4.062799 | 0.150945 | 0.880019 | NA |
| CG9083 | 0 | NA | NA | NA | NA | NA |
| CG3770 | 0.3141 | 1.240997 | 4.03841 | 0.307299 | 0.758616 | NA |
| CG3829 | 6.410969 | -4.42549 | 3.886842 | -1.13858 | 0.254878 | NA |
| CG2736 | 0.69928 | 1.019866 | 4.009445 | 0.254366 | 0.799213 | NA |
| Phk-3 | 0 | NA | NA | NA | NA | NA |
| CG30427 | 8.865743 | -2.22132 | 3.907426 | -0.56849 | 0.569704 | NA |
| Tpc2 | 47.08818 | -1.05688 | 3.951468 | -0.26746 | 0.789112 | NA |
| Ance-5 | 0 | NA | NA | NA | NA | NA |
| ETH | 0 | NA | NA | NA | NA | NA |
| Lcp9 | 0 | NA | NA | NA | NA | NA |
| tRNA:Ala-( | 0 | NA | NA | NA | NA | NA |
| tRNA:Ala-( | 0 | NA | NA | NA | NA | NA |
| tRNA:Ala-( | 3.729492 | 1.144349 | 3.963186 | 0.288745 | 0.772777 | NA |
| CG30430 | 0 | NA | NA | NA | NA | NA |
| CG30429 | 0 | NA | NA | NA | NA | NA |
| Ebp | 0 | NA | NA | NA | NA | NA |
| EbpII | 0 | NA | NA | NA | NA | NA |
| gsb-n | 23.87158 | 1.746033 | 3.953075 | 0.44169 | 0.658714 | NA |
| Lsp1gamm | 0.745683 | -0.62945 | 3.95549 | -0.15913 | 0.873564 | NA |
| CG13875 | 7.844848 | 0.362684 | 3.753528 | 0.096625 | 0.923024 | NA |
| CG13877 | 0 | NA | NA | NA | NA | NA |
| CG32845 | 0.104623 | 0.639075 | 4.067674 | 0.157111 | 0.875158 | NA |
| pyx | 13.76551 | -5.07164 | 3.90693 | -1.29811 | 0.194248 | NA |
| CG7049 | 8.506594 | -2.88022 | 3.80753 | -0.75645 | 0.449377 | NA |
| CG12483 | 0 | NA | NA | NA | NA | NA |
| Roc1b | 0 | NA | NA | NA | NA | NA |
| Ppm1 | 0 | NA | NA | NA | NA | NA |
| Reg-2 | 45.15878 | 6.743671 | 3.874108 | 1.740703 | 0.081736 | NA |
| CG13893 | 53.47464 | -3.14867 | 3.791755 | -0.8304 | 0.406313 | NA |
| Kah | 0.104623 | 0.639075 | 4.067674 | 0.157111 | 0.875158 | NA |

| | | | | | | |
|---|---|---|---|---|---|---|
| CG13891 | 3.140998 | 1.694072 | 3.966495 | 0.427096 | 0.66931 | NA |
| CG13884 | 0 | NA | NA | NA | NA | NA |
| tRNA:Ile-T/ | 0 | NA | NA | NA | NA | NA |
| tRNA:Ile-T/ | 0 | NA | NA | NA | NA | NA |
| Gr61a | 0.132078 | -1.13211 | 4.069918 | -0.27817 | 0.780885 | NA |
| cep290 | 0.1047 | 0.613261 | 4.062799 | 0.150945 | 0.880019 | NA |
| mat | 2.824828 | 1.495401 | 3.963153 | 0.377326 | 0.705931 | NA |
| CG13898 | 0 | NA | NA | NA | NA | NA |
| tRNA:iMet | 0 | NA | NA | NA | NA | NA |
| hng3 | 11.64196 | -6.91318 | 3.912066 | -1.76714 | 0.077204 | NA |
| CG32483 | 0 | NA | NA | NA | NA | NA |
| hiro | 0.282753 | -0.25398 | 4.020204 | -0.06318 | 0.949626 | NA |
| CG17129 | 0 | NA | NA | NA | NA | NA |
| LysB | 0 | NA | NA | NA | NA | NA |
| LysC | 0 | NA | NA | NA | NA | NA |
| LysE | 0 | NA | NA | NA | NA | NA |
| LysS | 0 | NA | NA | NA | NA | NA |
| LysP | 0.178053 | -1.16467 | 4.075067 | -0.2858 | 0.775028 | NA |
| LysD | 0 | NA | NA | NA | NA | NA |
| LysX | 0 | NA | NA | NA | NA | NA |
| galene | 0.178053 | -1.16467 | 4.075067 | -0.2858 | 0.775028 | NA |
| CG9129 | 0.53416 | -1.53481 | 4.012395 | -0.38252 | 0.702078 | NA |
| asRNA:CR4 | 0 | NA | NA | NA | NA | NA |
| asRNA:CR4 | 1.908008 | -0.10149 | 3.903479 | -0.026 | 0.979257 | NA |
| asRNA:CR4 | 0 | NA | NA | NA | NA | NA |
| asRNA:CR4 | 0 | NA | NA | NA | NA | NA |
| asRNA:CR4 | 0 | NA | NA | NA | NA | NA |
| asRNA:CR4 | 0 | NA | NA | NA | NA | NA |
| asRNA:CR4 | 0 | NA | NA | NA | NA | NA |
| lncRNA:CR | 0 | NA | NA | NA | NA | NA |
| asRNA:CR4 | 0 | NA | NA | NA | NA | NA |
| asRNA:CR4 | 0 | NA | NA | NA | NA | NA |
| asRNA:CR4 | 0 | NA | NA | NA | NA | NA |
| asRNA:CR4 | 17.58186 | 3.813812 | 3.886509 | 0.981295 | 0.326447 | NA |
| snoRNA:C( | 0 | NA | NA | NA | NA | NA |
| snoRNA:C( | 0 | NA | NA | NA | NA | NA |
| snoRNA:C( | 0 | NA | NA | NA | NA | NA |
| snoRNA:D( | 1.837946 | 1.501715 | 3.941336 | 0.381017 | 0.703191 | NA |
| snoRNA:ht | 0 | NA | NA | NA | NA | NA |
| snoRNA:ki: | 8.542628 | -1.66664 | 3.940348 | -0.42297 | 0.672319 | NA |
| snoRNA:lo | 0.178053 | -1.16467 | 4.075067 | -0.2858 | 0.775028 | NA |
| snoRNA:nc | 0 | NA | NA | NA | NA | NA |
| snoRNA:S( | 0.837599 | 1.568055 | 4.001801 | 0.391837 | 0.695178 | NA |
| snoRNA:S( | 0.466186 | 0.959471 | 4.033986 | 0.237847 | 0.812 | NA |
| snoRNA:Tu | 1.598336 | -0.35371 | 3.908817 | -0.09049 | 0.927898 | NA |
| scaRNA:M( | 0.62774 | 1.374224 | 3.999506 | 0.343599 | 0.731148 | NA |
| snoRNA:2F | 40.69346 | 2.757735 | 3.734206 | 0.738506 | 0.460207 | NA |
| snoRNA:ab | 0 | NA | NA | NA | NA | NA |
| scaRNA:Ps | 7.383689 | -0.99146 | 3.937704 | -0.25179 | 0.801206 | NA |
| snoRNA:C( | 0.565353 | 0.142845 | 3.959283 | 0.036079 | 0.97122 | NA |

| | | | | | | |
|---|---|---|---|---|---|---|
| snoRNA:C( | 12.28568 | -1.69873 | 3.954016 | -0.42962 | 0.667472 | NA |
| scaRNA:Ps | 6.053811 | -1.68915 | 3.957162 | -0.42686 | 0.669482 | NA |
| snoRNA:Ec | 0 | NA | NA | NA | NA | NA |
| scaRNA:Ps | 0.178053 | -1.16467 | 4.075067 | -0.2858 | 0.775028 | NA |
| snoRNA:Irl | 7.704395 | -3.34433 | 3.835324 | -0.87198 | 0.383219 | NA |
| snoRNA:lo | 0.209247 | 1.042804 | 4.047837 | 0.25762 | 0.7967 | NA |
| snoRNA:lo | 0 | NA | NA | NA | NA | NA |
| snoRNA:lo | 0 | NA | NA | NA | NA | NA |
| snoRNA:Pi | 0.178053 | -1.16467 | 4.075067 | -0.2858 | 0.775028 | NA |
| scaRNA:Ps | 12.66832 | 0.37284 | 3.952392 | 0.094333 | 0.924845 | NA |
| scaRNA:Ps | 0 | NA | NA | NA | NA | NA |
| scaRNA:Ps | 8.755114 | 2.560058 | 3.774782 | 0.6782 | 0.497645 | NA |
| snmRNA:C | 27.09742 | 1.534593 | 3.952235 | 0.388285 | 0.697805 | NA |
| lncRNA:CR | 0.209247 | 1.042804 | 4.047837 | 0.25762 | 0.7967 | NA |
| asRNA:CR4 | 37.93315 | 3.597634 | 3.708242 | 0.970172 | 0.331961 | NA |
| CR43482 | 0 | NA | NA | NA | NA | NA |
| CR43483 | 0 | NA | NA | NA | NA | NA |
| lncRNA:CR | 0 | NA | NA | NA | NA | NA |
| lncRNA:CR | 0 | NA | NA | NA | NA | NA |
| lncRNA:CR | 0 | NA | NA | NA | NA | NA |
| lncRNA:CR | 0 | NA | NA | NA | NA | NA |
| lncRNA:CR | 0 | NA | NA | NA | NA | NA |
| lncRNA:CR | 0 | NA | NA | NA | NA | NA |
| lncRNA:CR | 0 | NA | NA | NA | NA | NA |
| lncRNA:CR | 0 | NA | NA | NA | NA | NA |
| lncRNA:CR | 0 | NA | NA | NA | NA | NA |
| lncRNA:CR | 0 | NA | NA | NA | NA | NA |
| Vsx1 | 43.62684 | -0.08014 | 3.736845 | -0.02145 | 0.982889 | NA |
| Vsx2 | 1.675933 | 0.786673 | 3.895325 | 0.201953 | 0.839953 | NA |
| lncRNA:CR | 0 | NA | NA | NA | NA | NA |
| lncRNA:CR | 0 | NA | NA | NA | NA | NA |
| asRNA:CR4 | 0 | NA | NA | NA | NA | NA |
| lncRNA:CR | 3.455098 | 1.699144 | 3.965156 | 0.428519 | 0.668273 | NA |
| lncRNA:CR | 0 | NA | NA | NA | NA | NA |
| asRNA:CR4 | 0 | NA | NA | NA | NA | NA |
| asRNA:CR4 | 0.236778 | -0.26946 | 4.071573 | -0.06618 | 0.947234 | NA |
| lncRNA:CR | 0 | NA | NA | NA | NA | NA |
| CG43618 | 0 | NA | NA | NA | NA | NA |
| Acp98AB | 0 | NA | NA | NA | NA | NA |
| lncRNA:CR | 5.090129 | 2.804563 | 3.839498 | 0.73045 | 0.465115 | NA |
| lncRNA:CR | 0 | NA | NA | NA | NA | NA |
| lncRNA:CR | 0 | NA | NA | NA | NA | NA |
| lncRNA:CR | 20.17773 | -0.27155 | 3.843595 | -0.07065 | 0.943675 | NA |
| lncRNA:CR | 0 | NA | NA | NA | NA | NA |
| lncRNA:CR | 5.334983 | -0.21854 | 3.8608 | -0.05661 | 0.95486 | NA |
| asRNA:AST | 0 | NA | NA | NA | NA | NA |
| CG43630 | 0 | NA | NA | NA | NA | NA |
| lncRNA:CR | 0 | NA | NA | NA | NA | NA |
| lncRNA:CR | 0.233093 | 0.692656 | 4.076246 | 0.169925 | 0.865069 | NA |
| lncRNA:CR | 0 | NA | NA | NA | NA | NA |

| | | | | | | |
|---|---|---|---|---|---|---|
| lncRNA:CR | 0 | NA | NA | NA | NA | NA |
| lncRNA:CR | 53.738 | 4.435927 | 3.793943 | 1.169213 | 0.242318 | NA |
| asRNA:CR4 | 0 | NA | NA | NA | NA | NA |
| asRNA:CR4 | 3.661814 | 1.504942 | 3.960447 | 0.379993 | 0.703951 | NA |
| CG43638 | 53.19492 | 1.199356 | 3.898806 | 0.307621 | 0.758371 | NA |
| lncRNA:CR | 0 | NA | NA | NA | NA | NA |
| lncRNA:CR | 0 | NA | NA | NA | NA | NA |
| lncRNA:CR | 0 | NA | NA | NA | NA | NA |
| lncRNA:CR | 21.34913 | 0.009894 | 3.73445 | 0.00265 | 0.997886 | NA |
| lncRNA:CR | 0.178053 | -1.16467 | 4.075067 | -0.2858 | 0.775028 | NA |
| CG43645 | 0 | NA | NA | NA | NA | NA |
| CG43646 | 0 | NA | NA | NA | NA | NA |
| CG43647 | 0.2094 | 1.005034 | 4.045341 | 0.248442 | 0.803792 | NA |
| lncRNA:CR | 0 | NA | NA | NA | NA | NA |
| lncRNA:CR | 0 | NA | NA | NA | NA | NA |
| lncRNA:CR | 34.083 | 3.459114 | 3.880687 | 0.891366 | 0.372733 | NA |
| lncRNA:CR | 0 | NA | NA | NA | NA | NA |
| lncRNA:CR | 0 | NA | NA | NA | NA | NA |
| lncRNA:CR | 4.387809 | -5.38802 | 3.922418 | -1.37365 | 0.169551 | NA |
| Myo10A | 1.869329 | -1.24083 | 3.858668 | -0.32157 | 0.747779 | NA |
| CG43659 | 0 | NA | NA | NA | NA | NA |
| lncRNA:CR | 3.034074 | 1.498215 | 3.962342 | 0.378114 | 0.705346 | NA |
| CG43666 | 0 | NA | NA | NA | NA | NA |
| CG43667 | 0.534919 | -2.31746 | 4.015737 | -0.5771 | 0.563875 | NA |
| CG43668 | 0.178053 | -1.16467 | 4.075067 | -0.2858 | 0.775028 | NA |
| CR43669 | 0.178053 | -1.16467 | 4.075067 | -0.2858 | 0.775028 | NA |
| CR43671 | 0.388212 | 0.094116 | 4.038892 | 0.023303 | 0.981409 | NA |
| CG43672 | 0 | NA | NA | NA | NA | NA |
| CG43673 | 0 | NA | NA | NA | NA | NA |
| CG43675 | 36.03047 | 0.629982 | 3.951482 | 0.159429 | 0.873331 | NA |
| Su(var)3-9 | 51.79504 | 2.886521 | 3.919648 | 0.736423 | 0.461473 | NA |
| Rpb4 | 0 | NA | NA | NA | NA | NA |
| CG43677 | 0 | NA | NA | NA | NA | NA |
| CG43678 | 0.1047 | 0.613261 | 4.062799 | 0.150945 | 0.880019 | NA |
| CG43679 | 0 | NA | NA | NA | NA | NA |
| CG43680 | 0 | NA | NA | NA | NA | NA |
| lncRNA:CR | 0.178053 | -1.16467 | 4.075067 | -0.2858 | 0.775028 | NA |
| lncRNA:CR | 0 | NA | NA | NA | NA | NA |
| asRNA:CR4 | 0 | NA | NA | NA | NA | NA |
| lncRNA:CR | 0.104623 | 0.639075 | 4.067674 | 0.157111 | 0.875158 | NA |
| lncRNA:CR | 0 | NA | NA | NA | NA | NA |
| CR43687 | 0 | NA | NA | NA | NA | NA |
| CR43688 | 0.178053 | -1.16467 | 4.075067 | -0.2858 | 0.775028 | NA |
| CG43691 | 17.66118 | 0.448765 | 3.858002 | 0.11632 | 0.907399 | NA |
| CG43694 | 0.819113 | -2.02578 | 3.984237 | -0.50845 | 0.611138 | NA |
| lncRNA:CR | 0.998761 | -0.71333 | 3.993823 | -0.17861 | 0.858245 | NA |
| Hmgcr | 17.55278 | -1.28219 | 3.861796 | -0.33202 | 0.739875 | NA |
| lncRNA:CR | 3.263305 | 1.139496 | 3.96488 | 0.287397 | 0.773808 | NA |
| CG34334 | 0 | NA | NA | NA | NA | NA |
| Gen | 0 | NA | NA | NA | NA | NA |

| | | | | | | |
|---|---|---|---|---|---|---|
| CR43696 | 0 | NA | NA | NA | NA | NA |
| CR43697 | 0 | NA | NA | NA | NA | NA |
| lncRNA:CR | 0 | NA | NA | NA | NA | NA |
| snmRNA:C | 0 | NA | NA | NA | NA | NA |
| CG43702 | 0 | NA | NA | NA | NA | NA |
| CG43703 | 0 | NA | NA | NA | NA | NA |
| lncRNA:CR | 0 | NA | NA | NA | NA | NA |
| lncRNA:CR | 0 | NA | NA | NA | NA | NA |
| lncRNA:CR | 0 | NA | NA | NA | NA | NA |
| snRNA:LU | 0 | NA | NA | NA | NA | NA |
| CG43709 | 0.387453 | 0.098632 | 4.002797 | 0.024641 | 0.980342 | NA |
| CG43710 | 0.178812 | -1.08808 | 4.061735 | -0.26789 | 0.788788 | NA |
| CG43711 | 0 | NA | NA | NA | NA | NA |
| snoRNA:CI | 0 | NA | NA | NA | NA | NA |
| BubR1 | 6.231865 | -1.68968 | 3.956986 | -0.42701 | 0.66937 | NA |
| lncRNA:CR | 1.246373 | -1.6238 | 3.979526 | -0.40804 | 0.683245 | NA |
| lncRNA:CR | 0 | NA | NA | NA | NA | NA |
| lncRNA:CR | 0.178053 | -1.16467 | 4.075067 | -0.2858 | 0.775028 | NA |
| lncRNA:CR | 8.899494 | 1.731975 | 3.956638 | 0.437739 | 0.661575 | NA |
| lncRNA:CR | 0 | NA | NA | NA | NA | NA |
| lncRNA:CR | 0 | NA | NA | NA | NA | NA |
| lncRNA:CR | 0 | NA | NA | NA | NA | NA |
| lncRNA:CR | 0 | NA | NA | NA | NA | NA |
| asRNA:CR4 | 0 | NA | NA | NA | NA | NA |
| CG43725 | 0 | NA | NA | NA | NA | NA |
| qin | 3.601651 | 2.267006 | 3.928796 | 0.577023 | 0.563924 | NA |
| CR43727 | 0.357625 | -1.47707 | 4.044794 | -0.36518 | 0.714979 | NA |
| asRNA:CR4 | 0 | NA | NA | NA | NA | NA |
| asRNA:CR4 | 40.69198 | 0.952196 | 3.745205 | 0.254244 | 0.799307 | NA |
| CG43731 | 0 | NA | NA | NA | NA | NA |
| CG43732 | 0 | NA | NA | NA | NA | NA |
| CR43734 | 0 | NA | NA | NA | NA | NA |
| cd | 0.87953 | 0.922254 | 3.965066 | 0.232595 | 0.816076 | NA |
| Dic61B | 1.073687 | 0.81765 | 3.97163 | 0.205873 | 0.83689 | NA |
| asRNA:CR4 | 0 | NA | NA | NA | NA | NA |
| CG43739 | 0 | NA | NA | NA | NA | NA |
| Hmx | 10.73333 | 0.200181 | 3.952912 | 0.050641 | 0.959611 | NA |
| Cnx14D | 23.18491 | 1.631184 | 3.735382 | 0.436685 | 0.66234 | NA |
| CG43750 | 0 | NA | NA | NA | NA | NA |
| lncRNA:CR | 0 | NA | NA | NA | NA | NA |
| lncRNA:CR | 0.4188 | 1.41068 | 4.034613 | 0.349644 | 0.726606 | NA |
| Send2 | 0.1047 | 0.613261 | 4.062799 | 0.150945 | 0.880019 | NA |
| CG43760 | 0 | NA | NA | NA | NA | NA |
| lncRNA:CR | 0 | NA | NA | NA | NA | NA |
| lncRNA:CR | 0 | NA | NA | NA | NA | NA |
| lncRNA:CR | 0 | NA | NA | NA | NA | NA |
| lncRNA:CR | 4.290474 | 4.790691 | 3.933964 | 1.217777 | 0.223309 | NA |
| lncRNA:CR | 0 | NA | NA | NA | NA | NA |
| asRNA:CR4 | 2.848852 | -1.66884 | 3.963956 | -0.421 | 0.673753 | NA |
| lncRNA:CR | 0 | NA | NA | NA | NA | NA |

| | | | | | | |
|---|---|---|---|---|---|---|
| lncRNA:CR | 0 | NA | NA | NA | NA | NA |
| lncRNA:CR | 0.178053 | -1.16467 | 4.075067 | -0.2858 | 0.775028 | NA |
| mwh | 5.069045 | 2.094511 | 3.864031 | 0.542053 | 0.587782 | NA |
| Det | 10.63722 | -3.13029 | 3.912376 | -0.8001 | 0.423653 | NA |
| CG43773 | 0 | NA | NA | NA | NA | NA |
| CG43774 | 0 | NA | NA | NA | NA | NA |
| CG43775 | 0 | NA | NA | NA | NA | NA |
| CG43776 | 0 | NA | NA | NA | NA | NA |
| CG43777 | 0 | NA | NA | NA | NA | NA |
| CG43779 | 0 | NA | NA | NA | NA | NA |
| wrm1 | 0 | NA | NA | NA | NA | NA |
| wrm2 | 0 | NA | NA | NA | NA | NA |
| CG43784 | 0 | NA | NA | NA | NA | NA |
| lncRNA:CR | 0 | NA | NA | NA | NA | NA |
| PolE1 | 5.859281 | 2.931745 | 3.871197 | 0.757323 | 0.448857 | NA |
| asRNA:CR4 | 0 | NA | NA | NA | NA | NA |
| lncRNA:CR | 0 | NA | NA | NA | NA | NA |
| lncRNA:CR | 0 | NA | NA | NA | NA | NA |
| lncRNA:CR | 0 | NA | NA | NA | NA | NA |
| CG43796 | 0 | NA | NA | NA | NA | NA |
| CG43797 | 0 | NA | NA | NA | NA | NA |
| CG43798 | 0 | NA | NA | NA | NA | NA |
| CG43799 | 0.178053 | -1.16467 | 4.075067 | -0.2858 | 0.775028 | NA |
| CG43800 | 35.35792 | -0.53866 | 3.835721 | -0.14043 | 0.888318 | NA |
| lncRNA:CR | 14.76269 | 1.7408 | 3.954394 | 0.440219 | 0.659778 | NA |
| lncRNA:CR | 0 | NA | NA | NA | NA | NA |
| lncRNA:CR | 51.99156 | -1.706 | 3.951646 | -0.43172 | 0.665945 | NA |
| lncRNA:CR | 0 | NA | NA | NA | NA | NA |
| asRNA:CR4 | 18.67304 | 2.998883 | 3.828897 | 0.783224 | 0.433496 | NA |
| lncRNA:CR | 0 | NA | NA | NA | NA | NA |
| lncRNA:CR | 0 | NA | NA | NA | NA | NA |
| asRNA:CR4 | 0.104623 | 0.639075 | 4.067674 | 0.157111 | 0.875158 | NA |
| lncRNA:CR | 0 | NA | NA | NA | NA | NA |
| lncRNA:CR | 0 | NA | NA | NA | NA | NA |
| asRNA:CR4 | 0 | NA | NA | NA | NA | NA |
| CG43814 | 0 | NA | NA | NA | NA | NA |
| CG43815 | 0 | NA | NA | NA | NA | NA |
| CG43816 | 0 | NA | NA | NA | NA | NA |
| lncRNA:CR | 0.466186 | 0.959471 | 4.033986 | 0.237847 | 0.812 | NA |
| lncRNA:CR | 1.283782 | 0.252057 | 3.902549 | 0.064588 | 0.948502 | NA |
| lncRNA:CR | 0 | NA | NA | NA | NA | NA |
| asRNA:CR4 | 0 | NA | NA | NA | NA | NA |
| lncRNA:CR | 0 | NA | NA | NA | NA | NA |
| lncRNA:CR | 0.178053 | -1.16467 | 4.075067 | -0.2858 | 0.775028 | NA |
| asRNA:CR4 | 0 | NA | NA | NA | NA | NA |
| lncRNA:CR | 15.56811 | 6.714909 | 3.911652 | 1.716643 | 0.086044 | NA |
| lncRNA:CR | 0 | NA | NA | NA | NA | NA |
| lncRNA:CR | 0 | NA | NA | NA | NA | NA |
| CG43829 | 0 | NA | NA | NA | NA | NA |
| lncRNA:CR | 0 | NA | NA | NA | NA | NA |

| | | | | | | |
|---|---|---|---|---|---|---|
| lncRNA:CR | 0 | NA | NA | NA | NA | NA |
| lncRNA:CR | 0 | NA | NA | NA | NA | NA |
| asRNA:CR4 | 0 | NA | NA | NA | NA | NA |
| lncRNA:CR | 0 | NA | NA | NA | NA | NA |
| lncRNA:CR | 0 | NA | NA | NA | NA | NA |
| lncRNA:CR | 22.40578 | 1.745474 | 3.953216 | 0.441533 | 0.658827 | NA |
| lncRNA:CR | 0 | NA | NA | NA | NA | NA |
| lncRNA:CR | 0 | NA | NA | NA | NA | NA |
| CG43841 | 0 | NA | NA | NA | NA | NA |
| asRNA:CR4 | 0 | NA | NA | NA | NA | NA |
| CG43844 | 0 | NA | NA | NA | NA | NA |
| lncRNA:CR | 0 | NA | NA | NA | NA | NA |
| lncRNA:CR | 0.2094 | 1.005034 | 4.045341 | 0.248442 | 0.803792 | NA |
| lncRNA:CR | 0 | NA | NA | NA | NA | NA |
| lncRNA:CR | 0.104623 | 0.639075 | 4.067674 | 0.157111 | 0.875158 | NA |
| CG43851 | 0 | NA | NA | NA | NA | NA |
| lncRNA:CR | 0 | NA | NA | NA | NA | NA |
| lncRNA:CR | 0.178053 | -1.16467 | 4.075067 | -0.2858 | 0.775028 | NA |
| lncRNA:CR | 0 | NA | NA | NA | NA | NA |
| lncRNA:CR | 29.81161 | 0.220283 | 3.866794 | 0.056968 | 0.954571 | NA |
| lncRNA:CR | 0 | NA | NA | NA | NA | NA |
| lncRNA:CR | 0.1047 | 0.613261 | 4.062799 | 0.150945 | 0.880019 | NA |
| lncRNA:CR | 0 | NA | NA | NA | NA | NA |
| lncRNA:CR | 0 | NA | NA | NA | NA | NA |
| asRNA:CR4 | 0 | NA | NA | NA | NA | NA |
| asRNA:CR4 | 0 | NA | NA | NA | NA | NA |
| asRNA:CR4 | 0 | NA | NA | NA | NA | NA |
| asRNA:CR4 | 0 | NA | NA | NA | NA | NA |
| asRNA:CR4 | 0 | NA | NA | NA | NA | NA |
| lncRNA:CR | 5.697705 | -1.68799 | 3.957546 | -0.42652 | 0.669726 | NA |
| asRNA:CR4 | 0 | NA | NA | NA | NA | NA |
| asRNA:CR4 | 0 | NA | NA | NA | NA | NA |
| lncRNA:CR | 0 | NA | NA | NA | NA | NA |
| CG43880 | 0 | NA | NA | NA | NA | NA |
| lncRNA:CR | 0 | NA | NA | NA | NA | NA |
| lncRNA:CR | 0 | NA | NA | NA | NA | NA |
| lncRNA:CR | 6.072596 | 1.721931 | 3.959216 | 0.434917 | 0.663622 | NA |
| lncRNA:CR | 0 | NA | NA | NA | NA | NA |
| asRNA:CR4 | 0 | NA | NA | NA | NA | NA |
| CG43886 | 6.587971 | -1.69066 | 3.956663 | -0.42729 | 0.669165 | NA |
| lncRNA:CR | 32.6204 | 0.432143 | 3.83534 | 0.112674 | 0.910289 | NA |
| lncRNA:CR | 0 | NA | NA | NA | NA | NA |
| lncRNA:CR | 0.178812 | -1.08808 | 4.061735 | -0.26789 | 0.788788 | NA |
| asRNA:CR4 | 0 | NA | NA | NA | NA | NA |
| lncRNA:CR | 0 | NA | NA | NA | NA | NA |
| lncRNA:CR | 0 | NA | NA | NA | NA | NA |
| CG43894 | 0.387453 | 0.098632 | 4.002797 | 0.024641 | 0.980342 | NA |
| CG43895 | 0 | NA | NA | NA | NA | NA |
| CG43896 | 0.283436 | -0.20033 | 4.070792 | -0.04921 | 0.960751 | NA |
| CG43897 | 22.64086 | 0.516917 | 3.880485 | 0.133209 | 0.894028 | NA |

| | | | | | | |
|---|---|---|---|---|---|---|
| asRNA:CR4 | 54.18659 | 0.11364 | 3.378239 | 0.033639 | 0.973165 | NA |
| lncRNA:CR | 16.54259 | 1.742266 | 3.954024 | 0.440631 | 0.65948 | NA |
| asRNA:CR4 | 0.178053 | -1.16467 | 4.075067 | -0.2858 | 0.775028 | NA |
| lncRNA:CR | 10.78858 | -2.22416 | 3.890836 | -0.57164 | 0.567566 | NA |
| lncRNA:CR | 0.357625 | -1.47707 | 4.044794 | -0.36518 | 0.714979 | NA |
| asRNA:CR4 | 0.660388 | -1.75977 | 4.001266 | -0.4398 | 0.660079 | NA |
| lncRNA:CR | 0 | NA | NA | NA | NA | NA |
| asRNA:CR4 | 0 | NA | NA | NA | NA | NA |
| lncRNA:CR | 23.30932 | 1.174184 | 3.952924 | 0.297042 | 0.766434 | NA |
| lncRNA:CR | 0.1047 | 0.613261 | 4.062799 | 0.150945 | 0.880019 | NA |
| lncRNA:CR | 0 | NA | NA | NA | NA | NA |
| lncRNA:CR | 0 | NA | NA | NA | NA | NA |
| CG43915 | 1.176056 | -2.16791 | 3.961611 | -0.54723 | 0.584222 | NA |
| asRNA:CR4 | 18.67427 | -1.27098 | 3.952487 | -0.32157 | 0.747782 | NA |
| lncRNA:CR | 0 | NA | NA | NA | NA | NA |
| lncRNA:CR | 0.466186 | 0.959471 | 4.033986 | 0.237847 | 0.812 | NA |
| CG43919 | 50.3431 | -3.16839 | 3.905461 | -0.81127 | 0.41721 | NA |
| CG43920 | 0 | NA | NA | NA | NA | NA |
| CG43922 | 0.104623 | 0.639075 | 4.067674 | 0.157111 | 0.875158 | NA |
| CG43923 | 0 | NA | NA | NA | NA | NA |
| CG43924 | 0 | NA | NA | NA | NA | NA |
| CG43925 | 5.636447 | -2.58102 | 3.841249 | -0.67192 | 0.501634 | NA |
| asRNA:CR4 | 0 | NA | NA | NA | NA | NA |
| lncRNA:CR | 0 | NA | NA | NA | NA | NA |
| lncRNA:CR | 0 | NA | NA | NA | NA | NA |
| lncRNA:CR | 0.894061 | -2.00006 | 4.030294 | -0.49626 | 0.619714 | NA |
| lncRNA:CR | 0.209247 | 1.042804 | 4.047837 | 0.25762 | 0.7967 | NA |
| CG43931 | 0 | NA | NA | NA | NA | NA |
| lncRNA:CR | 0.132078 | -1.13211 | 4.069918 | -0.27817 | 0.780885 | NA |
| asRNA:CR4 | 1.787364 | -3.08936 | 3.999387 | -0.77246 | 0.439844 | NA |
| CG43935 | 0.178053 | -1.16467 | 4.075067 | -0.2858 | 0.775028 | NA |
| asRNA:CR4 | 2.617498 | 1.683176 | 3.969392 | 0.424039 | 0.671538 | NA |
| lncRNA:CR | 0.104623 | 0.639075 | 4.067674 | 0.157111 | 0.875158 | NA |
| CG43938 | 0 | NA | NA | NA | NA | NA |
| lncRNA:CR | 0.787655 | 0.325119 | 3.927268 | 0.082785 | 0.934022 | NA |
| lncRNA:CR | 0.178053 | -1.16467 | 4.075067 | -0.2858 | 0.775028 | NA |
| lncRNA:CR | 0.411147 | -0.29058 | 4.046973 | -0.0718 | 0.94276 | NA |
| lncRNA:CR | 0.1047 | 0.613261 | 4.062799 | 0.150945 | 0.880019 | NA |
| asRNA:CR4 | 0.605633 | 0.047977 | 3.956223 | 0.012127 | 0.990324 | NA |
| lncRNA:CR | 0 | NA | NA | NA | NA | NA |
| asRNA:CR4 | 0 | NA | NA | NA | NA | NA |
| lncRNA:CR | 0 | NA | NA | NA | NA | NA |
| lncRNA:CR | 0 | NA | NA | NA | NA | NA |
| lncRNA:CR | 0 | NA | NA | NA | NA | NA |
| lncRNA:CR | 0 | NA | NA | NA | NA | NA |
| CG43954 | 0 | NA | NA | NA | NA | NA |
| lncRNA:CR | 9.119426 | -2.24191 | 3.961111 | -0.56598 | 0.571408 | NA |
| lncRNA:CR | 0 | NA | NA | NA | NA | NA |
| lncRNA:CR | 52.44935 | -8.45153 | 3.846053 | -2.19745 | 0.027988 | NA |
| asRNA:CR4 | 0 | NA | NA | NA | NA | NA |

| | | | | | | |
|---|---|---|---|---|---|---|
| Mhc | 1.182007 | -0.53267 | 3.905671 | -0.13638 | 0.891517 | NA |
| CG43965 | 0 | NA | NA | NA | NA | NA |
| lncRNA:CR | 25.11472 | 7.247321 | 3.912065 | 1.852556 | 0.063946 | NA |
| lncRNA:CR | 0 | NA | NA | NA | NA | NA |
| CG43968 | 1.185692 | -1.26854 | 3.910876 | -0.32436 | 0.745664 | NA |
| lncRNA:CR | 0 | NA | NA | NA | NA | NA |
| asRNA:CR4 | 0.356107 | -1.47099 | 4.037764 | -0.36431 | 0.715628 | NA |
| CR43975 | 45.79152 | -8.64584 | 3.908436 | -2.2121 | 0.02696 | NA |
| lncRNA:CR | 0 | NA | NA | NA | NA | NA |
| lncRNA:CR | 0.356107 | -1.47099 | 4.037764 | -0.36431 | 0.715628 | NA |
| asRNA:CR4 | 0 | NA | NA | NA | NA | NA |
| CG43980 | 0.666997 | -3.07779 | 3.974176 | -0.77445 | 0.438666 | NA |
| asRNA:CR4 | 0 | NA | NA | NA | NA | NA |
| CG43982 | 0 | NA | NA | NA | NA | NA |
| lncRNA:CR | 0 | NA | NA | NA | NA | NA |
| lncRNA:CR | 0 | NA | NA | NA | NA | NA |
| lncRNA:CR | 0.178812 | -1.08808 | 4.061735 | -0.26789 | 0.788788 | NA |
| lncRNA:CR | 0 | NA | NA | NA | NA | NA |
| lncRNA:CR | 53.60629 | 2.975255 | 3.919042 | 0.759179 | 0.447745 | NA |
| lncRNA:CR | 0 | NA | NA | NA | NA | NA |
| lncRNA:CR | 0 | NA | NA | NA | NA | NA |
| lncRNA:CR | 0 | NA | NA | NA | NA | NA |
| CG43993 | 2.670799 | -1.66648 | 3.964795 | -0.42032 | 0.674251 | NA |
| lncRNA:CR | 17.48199 | 1.17221 | 3.953592 | 0.296492 | 0.766854 | NA |
| CG43996 | 7.957118 | 2.925265 | 3.930435 | 0.74426 | 0.456719 | NA |
| CG43997 | 0 | NA | NA | NA | NA | NA |
| CG43998 | 0 | NA | NA | NA | NA | NA |
| CG43999 | 0 | NA | NA | NA | NA | NA |
| CG44000 | 0 | NA | NA | NA | NA | NA |
| CG44001 | 0 | NA | NA | NA | NA | NA |
| CR44003 | 4.083297 | 1.7071 | 3.963068 | 0.430752 | 0.666649 | NA |
| CG44004 | 0.264155 | -1.54099 | 4.048976 | -0.38059 | 0.703509 | NA |
| CG44005 | 16.42585 | 1.531457 | 3.953091 | 0.387407 | 0.698455 | NA |
| CG44006 | 1.361099 | 1.628413 | 3.98439 | 0.408698 | 0.682761 | NA |
| CG44013 | 0.178812 | -1.08808 | 4.061735 | -0.26789 | 0.788788 | NA |
| CG44014 | 0 | NA | NA | NA | NA | NA |
| lncRNA:CR | 0 | NA | NA | NA | NA | NA |
| asRNA:CR4 | 0 | NA | NA | NA | NA | NA |
| asRNA:CR4 | 0 | NA | NA | NA | NA | NA |
| lncRNA:CR | 12.0336 | 1.713627 | 3.901155 | 0.439261 | 0.660472 | NA |
| Pburs | 0 | NA | NA | NA | NA | NA |
| lncRNA:CR | 0 | NA | NA | NA | NA | NA |
| CR44028 | 0 | NA | NA | NA | NA | NA |
| lncRNA:CR | 0 | NA | NA | NA | NA | NA |
| lncRNA:CR | 5.246801 | 1.5796 | 3.7704 | 0.418948 | 0.675254 | NA |
| lncRNA:CR | 0 | NA | NA | NA | NA | NA |
| asRNA:CR4 | 0.178053 | -1.16467 | 4.075067 | -0.2858 | 0.775028 | NA |
| lncRNA:CR | 0 | NA | NA | NA | NA | NA |
| CG44037 | 0.1047 | 0.613261 | 4.062799 | 0.150945 | 0.880019 | NA |
| CG44038 | 0 | NA | NA | NA | NA | NA |

| | | | | | | |
|---|---|---|---|---|---|---|
| lncRNA:CR | 0 | NA | NA | NA | NA | NA |
| CG44040 | 0.178053 | -1.16467 | 4.075067 | -0.2858 | 0.775028 | NA |
| lncRNA:CR | 0 | NA | NA | NA | NA | NA |
| lncRNA:CR | 0.519377 | -0.84038 | 3.994824 | -0.21037 | 0.833381 | NA |
| CG44044 | 8.391356 | 1.163791 | 3.956459 | 0.29415 | 0.768644 | NA |
| asRNA:CR4 | 0 | NA | NA | NA | NA | NA |
| asRNA:CR4 | 1.539076 | 1.809495 | 3.954298 | 0.457602 | 0.647238 | NA |
| asRNA:CR4 | 12.35459 | 1.738163 | 3.955063 | 0.439478 | 0.660315 | NA |
| lncRNA:CR | 0.1047 | 0.613261 | 4.062799 | 0.150945 | 0.880019 | NA |
| asRNA:CR4 | 0 | NA | NA | NA | NA | NA |
| lncRNA:CR | 4.296714 | 1.161077 | 3.83414 | 0.302826 | 0.762023 | NA |
| lncRNA:CR | 18.64746 | 1.172702 | 3.953425 | 0.296629 | 0.76675 | NA |
| asRNA:CR4 | 0 | NA | NA | NA | NA | NA |
| Tep5Psi | 0 | NA | NA | NA | NA | NA |
| lncRNA:iab | 0.282753 | -0.25398 | 4.020204 | -0.06318 | 0.949626 | NA |
| asRNA:CR4 | 0 | NA | NA | NA | NA | NA |
| asRNA:CR4 | 22.09455 | -6.28991 | 3.881714 | -1.6204 | 0.105147 | NA |
| lncRNA:CR | 0 | NA | NA | NA | NA | NA |
| lncRNA:CR | 0 | NA | NA | NA | NA | NA |
| lncRNA:CR | 0.104623 | 0.639075 | 4.067674 | 0.157111 | 0.875158 | NA |
| lncRNA:CR | 0 | NA | NA | NA | NA | NA |
| lncRNA:CR | 0 | NA | NA | NA | NA | NA |
| lncRNA:CR | 0 | NA | NA | NA | NA | NA |
| asRNA:CR4 | 1.012366 | -0.44098 | 3.915726 | -0.11262 | 0.910333 | NA |
| asRNA:CR4 | 0 | NA | NA | NA | NA | NA |
| asRNA:CR4 | 0 | NA | NA | NA | NA | NA |
| asRNA:CR4 | 49.25079 | -2.0781 | 3.839737 | -0.54121 | 0.588364 | NA |
| lncRNA:CR | 0 | NA | NA | NA | NA | NA |
| asRNA:CR4 | 0.178053 | -1.16467 | 4.075067 | -0.2858 | 0.775028 | NA |
| lncRNA:CR | 0 | NA | NA | NA | NA | NA |
| lncRNA:CR | 0 | NA | NA | NA | NA | NA |
| asRNA:CR4 | 5.601706 | -0.71134 | 3.846202 | -0.18495 | 0.853272 | NA |
| lncRNA:CR | 14.70171 | 4.588677 | 3.790777 | 1.210485 | 0.226093 | NA |
| lncRNA:CR | 30.62342 | 0.645379 | 3.920576 | 0.164613 | 0.869248 | NA |
| lncRNA:CR | 61.46341 | -7.3543 | 3.810193 | -1.93017 | 0.053586 | NA |
| lncRNA:CR | 0.178053 | -1.16467 | 4.075067 | -0.2858 | 0.775028 | NA |
| lncRNA:CR | 5.018952 | -1.89674 | 3.958511 | -0.47915 | 0.631829 | NA |
| lncRNA:CR | 14.8957 | -2.05089 | 3.95628 | -0.51839 | 0.604187 | NA |
| CR44079 | 0 | NA | NA | NA | NA | NA |
| lncRNA:CR | 0.31089 | -1.9003 | 4.033427 | -0.47114 | 0.637542 | NA |
| lncRNA:CR | 0 | NA | NA | NA | NA | NA |
| lncRNA:CR | 0.660388 | -1.75977 | 4.001266 | -0.4398 | 0.660079 | NA |
| asRNA:CR4 | 13.60102 | 1.529869 | 3.953541 | 0.386962 | 0.698785 | NA |
| lncRNA:CR | 0 | NA | NA | NA | NA | NA |
| asRNA:CR4 | 0 | NA | NA | NA | NA | NA |
| lncRNA:CR | 0 | NA | NA | NA | NA | NA |
| CG44090 | 0 | NA | NA | NA | NA | NA |
| lncRNA:CR | 0 | NA | NA | NA | NA | NA |
| asRNA:CR4 | 35.80241 | -0.76211 | 3.835757 | -0.19868 | 0.842509 | NA |
| CG44094 | 0 | NA | NA | NA | NA | NA |

| | | | | | | |
|---|---|---|---|---|---|---|
| asRNA:CR4 | 25.6553 | -0.01617 | 3.838211 | -0.00421 | 0.996639 | NA |
| lncRNA:CR | 0 | NA | NA | NA | NA | NA |
| lncRNA:CR | 0 | NA | NA | NA | NA | NA |
| lncRNA:CR | 0 | NA | NA | NA | NA | NA |
| CG44102 | 27.00065 | -2.26818 | 3.954453 | -0.57358 | 0.566255 | NA |
| lncRNA:CR | 0 | NA | NA | NA | NA | NA |
| lncRNA:CR | 0 | NA | NA | NA | NA | NA |
| asRNA:CR4 | 0 | NA | NA | NA | NA | NA |
| asRNA:CR4 | 0.906754 | -0.02643 | 3.908482 | -0.00676 | 0.994604 | NA |
| asRNA:CR4 | 0 | NA | NA | NA | NA | NA |
| lncRNA:CR | 56.07563 | -1.89572 | 3.833098 | -0.49457 | 0.620907 | NA |
| lncRNA:CR | 0 | NA | NA | NA | NA | NA |
| lncRNA:CR | 16.56183 | 0.395184 | 3.841278 | 0.102878 | 0.91806 | NA |
| lncRNA:CR | 0 | NA | NA | NA | NA | NA |
| lncRNA:CR | 0 | NA | NA | NA | NA | NA |
| asRNA:CR4 | 0 | NA | NA | NA | NA | NA |
| asRNA:CR4 | 0 | NA | NA | NA | NA | NA |
| asRNA:CR4 | 11.09014 | 2.752213 | 3.921894 | 0.701756 | 0.482831 | NA |
| lncRNA:CR | 0 | NA | NA | NA | NA | NA |
| lncRNA:CR | 0 | NA | NA | NA | NA | NA |
| asRNA:CR4 | 10.53854 | -5.91314 | 3.921221 | -1.50799 | 0.131558 | NA |
| lncRNA:CR | 0 | NA | NA | NA | NA | NA |
| lncRNA:CR | 0 | NA | NA | NA | NA | NA |
| fd3F | 0.356107 | -1.47099 | 4.037764 | -0.36431 | 0.715628 | NA |
| asRNA:CR4 | 0 | NA | NA | NA | NA | NA |
| lncRNA:CR | 0 | NA | NA | NA | NA | NA |
| lncRNA:CR | 0 | NA | NA | NA | NA | NA |
| lncRNA:CR | 46.09172 | -3.63267 | 3.81368 | -0.95254 | 0.340824 | NA |
| asRNA:CR4 | 0 | NA | NA | NA | NA | NA |
| CG4267 | 0 | NA | NA | NA | NA | NA |
| lncRNA:CR | 0 | NA | NA | NA | NA | NA |
| lncRNA:CR | 0 | NA | NA | NA | NA | NA |
| asRNA:CR4 | 0 | NA | NA | NA | NA | NA |
| lncRNA:CR | 6.437242 | -2.22638 | 3.96512 | -0.56149 | 0.574463 | NA |
| lncRNA:CR | 0 | NA | NA | NA | NA | NA |
| CG44139 | 0.712213 | -1.57119 | 3.998613 | -0.39293 | 0.694369 | NA |
| CG44140 | 0 | NA | NA | NA | NA | NA |
| CG44141 | 0 | NA | NA | NA | NA | NA |
| CG44142 | 0 | NA | NA | NA | NA | NA |
| lncRNA:CR | 0 | NA | NA | NA | NA | NA |
| asRNA:CR4 | 0 | NA | NA | NA | NA | NA |
| lncRNA:CR | 0.1047 | 0.613261 | 4.062799 | 0.150945 | 0.880019 | NA |
| lncRNA:CR | 0 | NA | NA | NA | NA | NA |
| asRNA:CR4 | 0 | NA | NA | NA | NA | NA |
| lncRNA:CR | 0 | NA | NA | NA | NA | NA |
| lncRNA:CR | 27.208 | -1.91809 | 3.952339 | -0.48531 | 0.62746 | NA |
| lncRNA:CR | 30.33163 | 1.684094 | 3.952533 | 0.42608 | 0.67005 | NA |
| lncRNA:CR | 56.77181 | 1.702996 | 3.810244 | 0.446952 | 0.65491 | NA |
| ppk18 | 0.178812 | -1.08808 | 4.061735 | -0.26789 | 0.788788 | NA |
| Np | 0.178053 | -1.16467 | 4.075067 | -0.2858 | 0.775028 | NA |

| | | | | | | |
|---|---|---|---|---|---|---|
| lncRNA:CR | 0.209247 | 1.042804 | 4.047837 | 0.25762 | 0.7967 | NA |
| asRNA:CR4 | 0 | NA | NA | NA | NA | NA |
| lncRNA:CR | 0 | NA | NA | NA | NA | NA |
| lncRNA:CR | 55.96105 | -0.31099 | 3.834692 | -0.0811 | 0.935363 | NA |
| Irk1 | 0.233093 | 0.692656 | 4.076246 | 0.169925 | 0.865069 | NA |
| asRNA:CR4 | 0 | NA | NA | NA | NA | NA |
| CG44163 | 0 | NA | NA | NA | NA | NA |
| FDY | 50.10731 | -0.07241 | 3.912127 | -0.01851 | 0.985233 | NA |
| lncRNA:CR | 54.85822 | -3.52234 | 3.899253 | -0.90334 | 0.366347 | NA |
| lncRNA:CR | 0 | NA | NA | NA | NA | NA |
| asRNA:CR4 | 0 | NA | NA | NA | NA | NA |
| asRNA:CR4 | 0 | NA | NA | NA | NA | NA |
| lncRNA:CR | 0 | NA | NA | NA | NA | NA |
| lncRNA:CR | 12.35394 | 1.168965 | 3.954696 | 0.295589 | 0.767544 | NA |
| lncRNA:CR | 0 | NA | NA | NA | NA | NA |
| lncRNA:CR | 0 | NA | NA | NA | NA | NA |
| CG44174 | 0 | NA | NA | NA | NA | NA |
| lncRNA:CR | 0 | NA | NA | NA | NA | NA |
| asRNA:CR4 | 0 | NA | NA | NA | NA | NA |
| lncRNA:CR | 3.245698 | 1.695863 | 3.966021 | 0.427598 | 0.668944 | NA |
| asRNA:CR4 | 36.54536 | 0.972774 | 3.836399 | 0.253564 | 0.799832 | NA |
| lncRNA:CR | 0 | NA | NA | NA | NA | NA |
| asRNA:CR4 | 0 | NA | NA | NA | NA | NA |
| asRNA:CR4 | 0 | NA | NA | NA | NA | NA |
| lncRNA:CR | 0 | NA | NA | NA | NA | NA |
| lncRNA:CR | 0 | NA | NA | NA | NA | NA |
| lncRNA:CR | 58.04537 | -1.70624 | 3.951569 | -0.43179 | 0.665895 | NA |
| lncRNA:CR | 0.178053 | -1.16467 | 4.075067 | -0.2858 | 0.775028 | NA |
| lncRNA:CR | 0 | NA | NA | NA | NA | NA |
| lncRNA:CR | 0 | NA | NA | NA | NA | NA |
| lncRNA:CR | 0 | NA | NA | NA | NA | NA |
| asRNA:CR4 | 0.132078 | -1.13211 | 4.069918 | -0.27817 | 0.780885 | NA |
| CG44194 | 0 | NA | NA | NA | NA | NA |
| lncRNA:CR | 0 | NA | NA | NA | NA | NA |
| lncRNA:CR | 0 | NA | NA | NA | NA | NA |
| CG44198 | 0 | NA | NA | NA | NA | NA |
| lncRNA:CR | 0 | NA | NA | NA | NA | NA |
| eIF4E3 | 0.209247 | 1.042804 | 4.047837 | 0.25762 | 0.7967 | NA |
| lncRNA:CR | 0 | NA | NA | NA | NA | NA |
| asRNA:CR4 | 0.715249 | -1.88064 | 4.034336 | -0.46616 | 0.641102 | NA |
| lncRNA:CR | 0.178053 | -1.16467 | 4.075067 | -0.2858 | 0.775028 | NA |
| lncRNA:CR | 1.050006 | 0.081601 | 3.967032 | 0.02057 | 0.983589 | NA |
| lncRNA:CR | 0 | NA | NA | NA | NA | NA |
| Oseg1 | 56.67021 | 1.966039 | 3.772312 | 0.521176 | 0.602244 | NA |
| asRNA:CR4 | 0 | NA | NA | NA | NA | NA |
| asRNA:CR4 | 0.1047 | 0.613261 | 4.062799 | 0.150945 | 0.880019 | NA |
| lncRNA:CR | 0 | NA | NA | NA | NA | NA |
| Gr22c | 0 | NA | NA | NA | NA | NA |
| Gr2a | 0.2094 | 1.005034 | 4.045341 | 0.248442 | 0.803792 | NA |
| Ir47b | 28.08548 | -3.92041 | 3.842805 | -1.0202 | 0.307636 | NA |

| | | | | | | |
|---|---|---|---|---|---|---|
| Ir56e | 0 | NA | NA | NA | NA | NA |
| lncRNA:CR | 0 | NA | NA | NA | NA | NA |
| lncRNA:CR | 0 | NA | NA | NA | NA | NA |
| lncRNA:CR | 0 | NA | NA | NA | NA | NA |
| CG44215 | 0 | NA | NA | NA | NA | NA |
| lncRNA:CR | 0.104623 | 0.639075 | 4.067674 | 0.157111 | 0.875158 | NA |
| lncRNA:CR | 0 | NA | NA | NA | NA | NA |
| lncRNA:CR | 0 | NA | NA | NA | NA | NA |
| lncRNA:CR | 19.58586 | -1.70224 | 3.952863 | -0.43064 | 0.666733 | NA |
| lncRNA:CR | 28.59406 | 0.113972 | 3.951527 | 0.028843 | 0.97699 | NA |
| lncRNA:CR | 6.491396 | 1.723946 | 3.958697 | 0.435483 | 0.663212 | NA |
| Ir60f | 0 | NA | NA | NA | NA | NA |
| asRNA:CR4 | 0 | NA | NA | NA | NA | NA |
| lncRNA:CR | 0 | NA | NA | NA | NA | NA |
| lncRNA:CR | 0 | NA | NA | NA | NA | NA |
| CG44227 | 0 | NA | NA | NA | NA | NA |
| asRNA:CR4 | 0 | NA | NA | NA | NA | NA |
| lncRNA:CR | 0 | NA | NA | NA | NA | NA |
| lncRNA:CR | 0 | NA | NA | NA | NA | NA |
| lncRNA:CR | 0 | NA | NA | NA | NA | NA |
| asRNA:CR4 | 0 | NA | NA | NA | NA | NA |
| lncRNA:CR | 48.07628 | -1.92027 | 3.951719 | -0.48593 | 0.627015 | NA |
| asRNA:CR4 | 0.178053 | -1.16467 | 4.075067 | -0.2858 | 0.775028 | NA |
| asRNA:CR4 | 0 | NA | NA | NA | NA | NA |
| lncRNA:CR | 0 | NA | NA | NA | NA | NA |
| CG44243 | 0.1047 | 0.613261 | 4.062799 | 0.150945 | 0.880019 | NA |
| CG44249 | 0 | NA | NA | NA | NA | NA |
| CG44250 | 0 | NA | NA | NA | NA | NA |
| CG44251 | 0 | NA | NA | NA | NA | NA |
| Fatp2 | 16.75199 | 1.742418 | 3.953985 | 0.440674 | 0.659449 | NA |
| CG44253 | 0 | NA | NA | NA | NA | NA |
| sud1 | 13.45938 | -2.22969 | 3.889841 | -0.57321 | 0.566503 | NA |
| lncRNA:CR | 0 | NA | NA | NA | NA | NA |
| CG44257 | 60.53811 | -1.70633 | 3.951542 | -0.43181 | 0.665877 | NA |
| lncRNA:CR | 0 | NA | NA | NA | NA | NA |
| CG44259 | 0.528311 | -1.72839 | 4.011871 | -0.43082 | 0.6666 | NA |
| lncRNA:CR | 0 | NA | NA | NA | NA | NA |
| CG44261 | 0 | NA | NA | NA | NA | NA |
| lncRNA:CR | 0 | NA | NA | NA | NA | NA |
| lncRNA:CR | 0 | NA | NA | NA | NA | NA |
| lncRNA:CR | 0 | NA | NA | NA | NA | NA |
| lncRNA:CR | 45.63199 | 1.147331 | 3.951921 | 0.290322 | 0.77157 | NA |
| asRNA:CR4 | 0 | NA | NA | NA | NA | NA |
| asRNA:CR4 | 0 | NA | NA | NA | NA | NA |
| lncRNA:CR | 56.96877 | 4.991552 | 3.805778 | 1.311572 | 0.189665 | NA |
| lncRNA:CR | 3.026728 | 2.697288 | 3.902197 | 0.691223 | 0.489426 | NA |
| CG44270 | 0 | NA | NA | NA | NA | NA |
| CG44271 | 0 | NA | NA | NA | NA | NA |
| CG42688 | 0 | NA | NA | NA | NA | NA |
| CG17097 | 0 | NA | NA | NA | NA | NA |

| | | | | | | |
|---|---|---|---|---|---|---|
| CG32727 | 0 | NA | NA | NA | NA | NA |
| CG13639 | 0 | NA | NA | NA | NA | NA |
| CG18258 | 0 | NA | NA | NA | NA | NA |
| CG18234 | 0 | NA | NA | NA | NA | NA |
| CG15425 | 0 | NA | NA | NA | NA | NA |
| lncRNA:CR | 0.178053 | -1.16467 | 4.075067 | -0.2858 | 0.775028 | NA |
| CG14717 | 39.02448 | 1.53608 | 3.95183 | 0.388701 | 0.697497 | NA |
| asRNA:CR4 | 0 | NA | NA | NA | NA | NA |
| lncRNA:CR | 0 | NA | NA | NA | NA | NA |
| lncRNA:CR | 0 | NA | NA | NA | NA | NA |
| lncRNA:CR | 0 | NA | NA | NA | NA | NA |
| asRNA:CR4 | 0 | NA | NA | NA | NA | NA |
| lncRNA:CR | 0 | NA | NA | NA | NA | NA |
| lncRNA:CR | 0 | NA | NA | NA | NA | NA |
| lncRNA:CR | 0 | NA | NA | NA | NA | NA |
| lncRNA:CR | 0 | NA | NA | NA | NA | NA |
| lncRNA:CR | 0.178053 | -1.16467 | 4.075067 | -0.2858 | 0.775028 | NA |
| lncRNA:CR | 0 | NA | NA | NA | NA | NA |
| lncRNA:CR | 0 | NA | NA | NA | NA | NA |
| lncRNA:CR | 0 | NA | NA | NA | NA | NA |
| lncRNA:CR | 0.5235 | 1.490714 | 4.02555 | 0.370313 | 0.711149 | NA |
| lncRNA:CR | 0 | NA | NA | NA | NA | NA |
| CG44286 | 0 | NA | NA | NA | NA | NA |
| asRNA:CR4 | 0 | NA | NA | NA | NA | NA |
| lncRNA:CR | 0 | NA | NA | NA | NA | NA |
| lncRNA:CR | 0 | NA | NA | NA | NA | NA |
| CG44290 | 0 | NA | NA | NA | NA | NA |
| asRNA:CR4 | 50.06972 | 9.06589 | 3.907623 | 2.320052 | 0.020338 | NA |
| lncRNA:CR | 0 | NA | NA | NA | NA | NA |
| asRNA:CR4 | 0 | NA | NA | NA | NA | NA |
| CG44296 | 0.132078 | -1.13211 | 4.069918 | -0.27817 | 0.780885 | NA |
| asRNA:CR4 | 0 | NA | NA | NA | NA | NA |
| lncRNA:CR | 0.53416 | -1.53481 | 4.012395 | -0.38252 | 0.702078 | NA |
| lncRNA:CR | 0.178812 | -1.08808 | 4.061735 | -0.26789 | 0.788788 | NA |
| lncRNA:CR | 0.1047 | 0.613261 | 4.062799 | 0.150945 | 0.880019 | NA |
| lncRNA:CR | 26.73231 | 0.108549 | 3.837492 | 0.028287 | 0.977434 | NA |
| Sfp33A4 | 0.178053 | -1.16467 | 4.075067 | -0.2858 | 0.775028 | NA |
| CG44303 | 0 | NA | NA | NA | NA | NA |
| asRNA:CR4 | 0 | NA | NA | NA | NA | NA |
| lncRNA:CR | 0 | NA | NA | NA | NA | NA |
| lncRNA:CR | 0.233093 | 0.692656 | 4.076246 | 0.169925 | 0.865069 | NA |
| lncRNA:CR | 0 | NA | NA | NA | NA | NA |
| lncRNA:CR | 0 | NA | NA | NA | NA | NA |
| lncRNA:CR | 0 | NA | NA | NA | NA | NA |
| lncRNA:CR | 0 | NA | NA | NA | NA | NA |
| lncRNA:CR | 0 | NA | NA | NA | NA | NA |
| lncRNA:CR | 0 | NA | NA | NA | NA | NA |
| CG44314 | 0.5235 | 1.490714 | 4.02555 | 0.370313 | 0.711149 | NA |
| lncRNA:CR | 0 | NA | NA | NA | NA | NA |
| lncRNA:CR | 0 | NA | NA | NA | NA | NA |

| | | | | | | |
|---|---|---|---|---|---|---|
| lncRNA:CR | 5.273793 | -1.01041 | 3.929562 | -0.25713 | 0.797079 | NA |
| lncRNA:CR | 0 | NA | NA | NA | NA | NA |
| lncRNA:CR | 0 | NA | NA | NA | NA | NA |
| lncRNA:CR | 0 | NA | NA | NA | NA | NA |
| lncRNA:CR | 0 | NA | NA | NA | NA | NA |
| lncRNA:CR | 0 | NA | NA | NA | NA | NA |
| lncRNA:CR | 0.132078 | -1.13211 | 4.069918 | -0.27817 | 0.780885 | NA |
| CG44329 | 0 | NA | NA | NA | NA | NA |
| lncRNA:CR | 0 | NA | NA | NA | NA | NA |
| lncRNA:CR | 36.88016 | 3.915449 | 3.797741 | 1.030994 | 0.302543 | NA |
| lncRNA:CR | 0 | NA | NA | NA | NA | NA |
| lncRNA:CR | 0 | NA | NA | NA | NA | NA |
| CG44335 | 0.356107 | -1.47099 | 4.037764 | -0.36431 | 0.715628 | NA |
| lncRNA:TS: | 0 | NA | NA | NA | NA | NA |
| lncRNA:CR | 0 | NA | NA | NA | NA | NA |
| lncRNA:CR | 0 | NA | NA | NA | NA | NA |
| lncRNA:CR | 0 | NA | NA | NA | NA | NA |
| asRNA:CR4 | 51.7467 | 1.177464 | 3.951818 | 0.297955 | 0.765738 | NA |
| CG44341 | 0.5235 | 1.490714 | 4.02555 | 0.370313 | 0.711149 | NA |
| lncRNA:CR | 0 | NA | NA | NA | NA | NA |
| lncRNA:CR | 14.13231 | -1.91351 | 3.953652 | -0.48398 | 0.628397 | NA |
| lncRNA:CR | 0.178053 | -1.16467 | 4.075067 | -0.2858 | 0.775028 | NA |
| lncRNA:CR | 0 | NA | NA | NA | NA | NA |
| lncRNA:CR | 39.24263 | -7.94987 | 3.910145 | -2.03314 | 0.042038 | NA |
| lncRNA:CR | 0 | NA | NA | NA | NA | NA |
| lncRNA:CR | 0 | NA | NA | NA | NA | NA |
| asRNA:CR4 | 0 | NA | NA | NA | NA | NA |
| lncRNA:CR | 0 | NA | NA | NA | NA | NA |
| lncRNA:CR | 0 | NA | NA | NA | NA | NA |
| lncRNA:CR | 0 | NA | NA | NA | NA | NA |
| CG44355 | 0.536437 | -1.71175 | 4.038042 | -0.42391 | 0.671635 | NA |
| lncRNA:CR | 0.466186 | 0.959471 | 4.033986 | 0.237847 | 0.812 | NA |
| asRNA:CR4 | 7.328995 | 1.727313 | 3.957831 | 0.436429 | 0.662525 | NA |
| lncRNA:CR | 2.509476 | -1.87226 | 3.965749 | -0.47211 | 0.636851 | NA |
| lncRNA:CR | 0 | NA | NA | NA | NA | NA |
| CG44362 | 0 | NA | NA | NA | NA | NA |
| lncRNA:CR | 0.178053 | -1.16467 | 4.075067 | -0.2858 | 0.775028 | NA |
| lncRNA:CR | 0.807012 | 0.227825 | 3.984055 | 0.057184 | 0.954399 | NA |
| lncRNA:CR | 50.98879 | 2.974775 | 3.919148 | 0.759036 | 0.447831 | NA |
| asRNA:CR4 | 0 | NA | NA | NA | NA | NA |
| asRNA:CR4 | 57.45379 | -1.92073 | 3.951586 | -0.48607 | 0.626921 | NA |
| lncRNA:CR | 0 | NA | NA | NA | NA | NA |
| asRNA:CR4 | 15.35954 | 1.623059 | 3.953965 | 0.410489 | 0.681447 | NA |
| lncRNA:CR | 0 | NA | NA | NA | NA | NA |
| asRNA:CR4 | 0 | NA | NA | NA | NA | NA |
| lncRNA:CR | 0 | NA | NA | NA | NA | NA |
| asRNA:CR4 | 0.69928 | 1.019866 | 4.009445 | 0.254366 | 0.799213 | NA |
| asRNA:CR4 | 0.178812 | -1.08808 | 4.061735 | -0.26789 | 0.788788 | NA |
| lncRNA:CR | 0 | NA | NA | NA | NA | NA |
| lncRNA:CR | 0 | NA | NA | NA | NA | NA |

| | | | | | | |
|---|---|---|---|---|---|---|
| lncRNA:CR | 0.233093 | 0.692656 | 4.076246 | 0.169925 | 0.865069 | NA |
| lncRNA:CR | 0.178053 | -1.16467 | 4.075067 | -0.2858 | 0.775028 | NA |
| asRNA:CR4 | 0 | NA | NA | NA | NA | NA |
| asRNA:CR4 | 0 | NA | NA | NA | NA | NA |
| CR44383 | 2.043558 | 0.685373 | 3.968489 | 0.172704 | 0.862884 | NA |
| asRNA:CR4 | 1.251686 | -2.06022 | 4.011654 | -0.51356 | 0.60756 | NA |
| lncRNA:CR | 0 | NA | NA | NA | NA | NA |
| CG44388 | 0 | NA | NA | NA | NA | NA |
| lncRNA:CR | 0 | NA | NA | NA | NA | NA |
| asRNA:CR4 | 36.40609 | 5.614564 | 3.869983 | 1.450798 | 0.146836 | NA |
| CR44391 | 0 | NA | NA | NA | NA | NA |
| lncRNA:CR | 0 | NA | NA | NA | NA | NA |
| lncRNA:CR | 0 | NA | NA | NA | NA | NA |
| CR44394 | 0 | NA | NA | NA | NA | NA |
| lncRNA:CR | 2.165895 | 0.59106 | 3.93154 | 0.150338 | 0.880498 | NA |
| lncRNA:CR | 0 | NA | NA | NA | NA | NA |
| lncRNA:CR | 0 | NA | NA | NA | NA | NA |
| lncRNA:CR | 0 | NA | NA | NA | NA | NA |
| lncRNA:CR | 0 | NA | NA | NA | NA | NA |
| lncRNA:CR | 0 | NA | NA | NA | NA | NA |
| CG44403 | 0.209323 | 1.414133 | 4.032409 | 0.350692 | 0.725819 | NA |
| IBIN | 0 | NA | NA | NA | NA | NA |
| CG44405 | 0 | NA | NA | NA | NA | NA |
| CG44406 | 0.178812 | -1.08808 | 4.061735 | -0.26789 | 0.788788 | NA |
| lncRNA:CR | 0 | NA | NA | NA | NA | NA |
| lncRNA:CR | 0 | NA | NA | NA | NA | NA |
| lncRNA:CR | 0 | NA | NA | NA | NA | NA |
| lncRNA:CR | 0 | NA | NA | NA | NA | NA |
| lncRNA:CR | 0 | NA | NA | NA | NA | NA |
| lncRNA:CR | 0 | NA | NA | NA | NA | NA |
| lncRNA:CR | 0 | NA | NA | NA | NA | NA |
| lncRNA:CR | 0 | NA | NA | NA | NA | NA |
| asRNA:CR4 | 0 | NA | NA | NA | NA | NA |
| lncRNA:CR | 0 | NA | NA | NA | NA | NA |
| lncRNA:CR | 0 | NA | NA | NA | NA | NA |
| lncRNA:CR | 0 | NA | NA | NA | NA | NA |
| lncRNA:CR | 0.178053 | -1.16467 | 4.075067 | -0.2858 | 0.775028 | NA |
| Prosalpha4 | 0 | NA | NA | NA | NA | NA |
| beat-IIIa | 0 | NA | NA | NA | NA | NA |
| CG44426 | 0 | NA | NA | NA | NA | NA |
| asRNA:CR4 | 0 | NA | NA | NA | NA | NA |
| CG44428 | 0 | NA | NA | NA | NA | NA |
| lncRNA:CR | 0 | NA | NA | NA | NA | NA |
| asRNA:CR4 | 6.800512 | 1.520433 | 3.956126 | 0.384324 | 0.700738 | NA |
| lncRNA:CR | 0 | NA | NA | NA | NA | NA |
| CG44433 | 0.356107 | -1.47099 | 4.037764 | -0.36431 | 0.715628 | NA |
| CG44434 | 0 | NA | NA | NA | NA | NA |
| CG44435 | 0 | NA | NA | NA | NA | NA |
| CG44437 | 57.97741 | -0.51855 | 3.729977 | -0.13902 | 0.889433 | NA |
| CG44438 | 0 | NA | NA | NA | NA | NA |

| | | | | | | |
|---|---|---|---|---|---|---|
| CG44439 | 0 | NA | NA | NA | NA | NA |
| lncRNA:CR | 36.53199 | -2.17586 | 3.953338 | -0.55038 | 0.582056 | NA |
| lncRNA:CR | 0.2094 | 1.005034 | 4.045341 | 0.248442 | 0.803792 | NA |
| lncRNA:CR | 0 | NA | NA | NA | NA | NA |
| lncRNA:CR | 0.356107 | -1.47099 | 4.037764 | -0.36431 | 0.715628 | NA |
| lncRNA:CR | 18.76824 | -3.57913 | 3.799552 | -0.94199 | 0.346199 | NA |
| lncRNA:CR | 0 | NA | NA | NA | NA | NA |
| lncRNA:CR | 16.75199 | 1.742418 | 3.953985 | 0.440674 | 0.659449 | NA |
| lncRNA:CR | 0.2094 | 1.005034 | 4.045341 | 0.248442 | 0.803792 | NA |
| lncRNA:TS | 19.80125 | 1.470231 | 3.952641 | 0.371962 | 0.709921 | NA |
| lncRNA:CR | 53.04399 | 1.536977 | 3.951587 | 0.388952 | 0.697312 | NA |
| lncRNA:CR | 0 | NA | NA | NA | NA | NA |
| lncRNA:CR | 0 | NA | NA | NA | NA | NA |
| lncRNA:CR | 0 | NA | NA | NA | NA | NA |
| lncRNA:CR | 0 | NA | NA | NA | NA | NA |
| lncRNA:CR | 0 | NA | NA | NA | NA | NA |
| lncRNA:CR | 0 | NA | NA | NA | NA | NA |
| lncRNA:CR | 0 | NA | NA | NA | NA | NA |
| lncRNA:CR | 0.178053 | -1.16467 | 4.075067 | -0.2858 | 0.775028 | NA |
| lncRNA:CR | 0 | NA | NA | NA | NA | NA |
| lncRNA:CR | 0 | NA | NA | NA | NA | NA |
| lncRNA:CR | 0 | NA | NA | NA | NA | NA |
| lncRNA:CR | 0 | NA | NA | NA | NA | NA |
| lncRNA:CR | 0.178812 | -1.08808 | 4.061735 | -0.26789 | 0.788788 | NA |
| lncRNA:CR | 3.04686 | 1.344349 | 3.854011 | 0.348818 | 0.727226 | NA |
| lncRNA:CR | 0 | NA | NA | NA | NA | NA |
| lncRNA:CR | 0 | NA | NA | NA | NA | NA |
| lncRNA:CR | 0 | NA | NA | NA | NA | NA |
| lncRNA:CR | 0 | NA | NA | NA | NA | NA |
| lncRNA:CR | 0 | NA | NA | NA | NA | NA |
| lncRNA:CR | 0 | NA | NA | NA | NA | NA |
| lncRNA:CR | 0.104623 | 0.639075 | 4.067674 | 0.157111 | 0.875158 | NA |
| lncRNA:CR | 4.880176 | -1.84563 | 3.862392 | -0.47785 | 0.63276 | NA |
| CG44475 | 0 | NA | NA | NA | NA | NA |
| CG44476 | 2.503372 | -2.15366 | 3.984641 | -0.54049 | 0.588858 | NA |
| CG44477 | 0 | NA | NA | NA | NA | NA |
| lncRNA:CR | 0 | NA | NA | NA | NA | NA |
| lncRNA:CR | 0 | NA | NA | NA | NA | NA |
| lncRNA:CR | 0 | NA | NA | NA | NA | NA |
| lncRNA:CR | 0 | NA | NA | NA | NA | NA |
| lncRNA:CR | 0.178053 | -1.16467 | 4.075067 | -0.2858 | 0.775028 | NA |
| lncRNA:CR | 0 | NA | NA | NA | NA | NA |
| lncRNA:CR | 0 | NA | NA | NA | NA | NA |
| lncRNA:CR | 0 | NA | NA | NA | NA | NA |
| lncRNA:CR | 0.282753 | -0.25398 | 4.020204 | -0.06318 | 0.949626 | NA |
| lncRNA:CR | 0 | NA | NA | NA | NA | NA |
| lncRNA:CR | 0 | NA | NA | NA | NA | NA |
| lncRNA:CR | 0 | NA | NA | NA | NA | NA |
| lncRNA:CR | 0 | NA | NA | NA | NA | NA |
| lncRNA:CR | 0 | NA | NA | NA | NA | NA |

| | | | | | | |
|---|---|---|---|---|---|---|
| lncRNA:CR | 13.76214 | -1.20321 | 3.952973 | -0.30438 | 0.760838 | NA |
| lncRNA:CR | 3.782135 | -0.08243 | 3.888969 | -0.0212 | 0.98309 | NA |
| lncRNA:CR | 0 | NA | NA | NA | NA | NA |
| lncRNA:CR | 0.178053 | -1.16467 | 4.075067 | -0.2858 | 0.775028 | NA |
| lncRNA:CR | 32.77108 | 1.748369 | 3.95249 | 0.442346 | 0.658239 | NA |
| lncRNA:CR | 0 | NA | NA | NA | NA | NA |
| lncRNA:CR | 41.3135 | 2.474548 | 3.884939 | 0.636959 | 0.524151 | NA |
| lncRNA:CR | 0 | NA | NA | NA | NA | NA |
| lncRNA:CR | 5.989472 | -3.42885 | 3.914999 | -0.87582 | 0.381125 | NA |
| lncRNA:CR | 0.418493 | 1.313669 | 4.019283 | 0.326842 | 0.743788 | NA |
| lncRNA:CR | 0 | NA | NA | NA | NA | NA |
| lncRNA:CR | 2.809039 | -0.20446 | 3.817847 | -0.05355 | 0.95729 | NA |
| lncRNA:CR | 0 | NA | NA | NA | NA | NA |
| lncRNA:CR | 0.536437 | -1.71175 | 4.038042 | -0.42391 | 0.671635 | NA |
| lncRNA:CR | 36.73885 | -0.44728 | 3.950028 | -0.11323 | 0.909844 | NA |
| lncRNA:CR | 0 | NA | NA | NA | NA | NA |
| lncRNA:CR | 0 | NA | NA | NA | NA | NA |
| asRNA:CR4 | 0 | NA | NA | NA | NA | NA |
| lncRNA:CR | 0 | NA | NA | NA | NA | NA |
| lncRNA:CR | 22.85558 | 7.152658 | 3.910546 | 1.829069 | 0.067389 | NA |
| asRNA:CR4 | 11.98042 | -2.25114 | 3.958753 | -0.56865 | 0.569594 | NA |
| lncRNA:CR | 0 | NA | NA | NA | NA | NA |
| lncRNA:CR | 0 | NA | NA | NA | NA | NA |
| lncRNA:CR | 0.178053 | -1.16467 | 4.075067 | -0.2858 | 0.775028 | NA |
| lncRNA:CR | 0.310131 | -1.94616 | 4.045058 | -0.48112 | 0.630431 | NA |
| lncRNA:CR | 0.178053 | -1.16467 | 4.075067 | -0.2858 | 0.775028 | NA |
| lncRNA:CR | 0.104623 | 0.639075 | 4.067674 | 0.157111 | 0.875158 | NA |
| lncRNA:CR | 0.837599 | 1.568055 | 4.001801 | 0.391837 | 0.695178 | NA |
| CG44521 | 0 | NA | NA | NA | NA | NA |
| lncRNA:CR | 0 | NA | NA | NA | NA | NA |
| lncRNA:CR | 0 | NA | NA | NA | NA | NA |
| lncRNA:CR | 0 | NA | NA | NA | NA | NA |
| lncRNA:CR | 0 | NA | NA | NA | NA | NA |
| lncRNA:CR | 0.356107 | -1.47099 | 4.037764 | -0.36431 | 0.715628 | NA |
| lncRNA:CR | 0 | NA | NA | NA | NA | NA |
| lncRNA:CR | 0 | NA | NA | NA | NA | NA |
| lncRNA:CR | 0 | NA | NA | NA | NA | NA |
| lncRNA:CR | 0 | NA | NA | NA | NA | NA |
| CG44532 | 41.08433 | -3.16599 | 3.905889 | -0.81057 | 0.417614 | NA |
| lncRNA:CR | 0 | NA | NA | NA | NA | NA |
| lncRNA:CR | 0.104623 | 0.639075 | 4.067674 | 0.157111 | 0.875158 | NA |
| lncRNA:CR | 0.356107 | -1.47099 | 4.037764 | -0.36431 | 0.715628 | NA |
| lncRNA:CR | 0 | NA | NA | NA | NA | NA |
| lncRNA:CR | 0 | NA | NA | NA | NA | NA |
| lncRNA:CR | 0 | NA | NA | NA | NA | NA |
| lncRNA:CR | 0 | NA | NA | NA | NA | NA |
| lncRNA:CR | 0 | NA | NA | NA | NA | NA |
| lncRNA:CR | 0 | NA | NA | NA | NA | NA |
| lncRNA:CR | 0 | NA | NA | NA | NA | NA |
| lncRNA:CR | 56.83034 | -1.66616 | 3.951565 | -0.42165 | 0.673284 | NA |

| | | | | | | |
|---|---|---|---|---|---|---|
| lncRNA:CR | 0 | NA | NA | NA | NA | NA |
| lncRNA:CR | 0 | NA | NA | NA | NA | NA |
| lncRNA:CR | 0.178053 | -1.16467 | 4.075067 | -0.2858 | 0.775028 | NA |
| lncRNA:CR | 0 | NA | NA | NA | NA | NA |
| lncRNA:CR | 0 | NA | NA | NA | NA | NA |
| lncRNA:CR | 0 | NA | NA | NA | NA | NA |
| lncRNA:CR | 0 | NA | NA | NA | NA | NA |
| lncRNA:CR | 0.732516 | 2.619177 | 3.987823 | 0.656794 | 0.511314 | NA |
| lncRNA:CR | 0 | NA | NA | NA | NA | NA |
| lncRNA:CR | 17.30218 | -1.91525 | 3.953153 | -0.48449 | 0.628041 | NA |
| lncRNA:CR | 0 | NA | NA | NA | NA | NA |
| lncRNA:CR | 0 | NA | NA | NA | NA | NA |
| lncRNA:CR | 21.67767 | 1.173743 | 3.953075 | 0.296919 | 0.766528 | NA |
| lncRNA:CR | 0 | NA | NA | NA | NA | NA |
| lncRNA:CR | 0.712213 | -1.57119 | 3.998613 | -0.39293 | 0.694369 | NA |
| lncRNA:CR | 0 | NA | NA | NA | NA | NA |
| lncRNA:CR | 0 | NA | NA | NA | NA | NA |
| lncRNA:CR | 0 | NA | NA | NA | NA | NA |
| lncRNA:CR | 0 | NA | NA | NA | NA | NA |
| lncRNA:CR | 0 | NA | NA | NA | NA | NA |
| lncRNA:CR | 0 | NA | NA | NA | NA | NA |
| lncRNA:CR | 1.452855 | -1.83946 | 3.975716 | -0.46267 | 0.643599 | NA |
| lncRNA:CR | 35.62268 | -0.27627 | 3.83623 | -0.07202 | 0.94259 | NA |
| lncRNA:CR | 0.178053 | -1.16467 | 4.075067 | -0.2858 | 0.775028 | NA |
| CG44569 | 0 | NA | NA | NA | NA | NA |
| lncRNA:CR | 1.42283 | -0.01324 | 3.949901 | -0.00335 | 0.997325 | NA |
| lncRNA:CR | 0 | NA | NA | NA | NA | NA |
| lncRNA:CR | 0.178053 | -1.16467 | 4.075067 | -0.2858 | 0.775028 | NA |
| lncRNA:CR | 0 | NA | NA | NA | NA | NA |
| CG44574 | 0 | NA | NA | NA | NA | NA |
| lncRNA:CR | 0 | NA | NA | NA | NA | NA |
| lncRNA:CR | 0 | NA | NA | NA | NA | NA |
| lncRNA:CR | 0 | NA | NA | NA | NA | NA |
| lncRNA:CR | 0 | NA | NA | NA | NA | NA |
| lncRNA:CR | 0 | NA | NA | NA | NA | NA |
| lncRNA:CR | 0 | NA | NA | NA | NA | NA |
| lncRNA:CR | 0.178812 | -1.08808 | 4.061735 | -0.26789 | 0.788788 | NA |
| lncRNA:CR | 0 | NA | NA | NA | NA | NA |
| lncRNA:CR | 0 | NA | NA | NA | NA | NA |
| lncRNA:CR | 0 | NA | NA | NA | NA | NA |
| lncRNA:CR | 0 | NA | NA | NA | NA | NA |
| lncRNA:CR | 0 | NA | NA | NA | NA | NA |
| lncRNA:CR | 0 | NA | NA | NA | NA | NA |
| lncRNA:CR | 0 | NA | NA | NA | NA | NA |
| lncRNA:CR | 0 | NA | NA | NA | NA | NA |
| asRNA:CR4 | 0 | NA | NA | NA | NA | NA |
| lncRNA:CR | 5.299592 | -1.24712 | 3.956386 | -0.31522 | 0.752597 | NA |
| lncRNA:CR | 0.282753 | -0.25398 | 4.020204 | -0.06318 | 0.949626 | NA |
| lncRNA:CR | 28.0941 | 0.324571 | 3.837217 | 0.084585 | 0.932591 | NA |
| lncRNA:CR | 34.73333 | -3.05451 | 3.802354 | -0.80332 | 0.421789 | NA |

| | | | | | | |
|---|---|---|---|---|---|---|
| lncRNA:CR | 0.837599 | 1.568055 | 4.001801 | 0.391837 | 0.695178 | NA |
| lncRNA:CR | 1.434452 | 0.968283 | 3.97124 | 0.243824 | 0.807367 | NA |
| asRNA:CR4 | 0 | NA | NA | NA | NA | NA |
| asRNA:CR4 | 0 | NA | NA | NA | NA | NA |
| lncRNA:CR | 0 | NA | NA | NA | NA | NA |
| lncRNA:CR | 0.310131 | -1.94616 | 4.045058 | -0.48112 | 0.630431 | NA |
| lncRNA:CR | 0 | NA | NA | NA | NA | NA |
| lncRNA:CR | 0 | NA | NA | NA | NA | NA |
| lncRNA:CR | 0 | NA | NA | NA | NA | NA |
| lncRNA:CR | 0.492153 | 0.308117 | 3.993933 | 0.077146 | 0.938507 | NA |
| lncRNA:CR | 0 | NA | NA | NA | NA | NA |
| lncRNA:CR | 0 | NA | NA | NA | NA | NA |
| lncRNA:CR | 0 | NA | NA | NA | NA | NA |
| lncRNA:CR | 0 | NA | NA | NA | NA | NA |
| lncRNA:CR | 0 | NA | NA | NA | NA | NA |
| lncRNA:CR | 0.104623 | 0.639075 | 4.067674 | 0.157111 | 0.875158 | NA |
| lncRNA:CR | 0 | NA | NA | NA | NA | NA |
| lncRNA:CR | 0 | NA | NA | NA | NA | NA |
| lncRNA:CR | 0 | NA | NA | NA | NA | NA |
| lncRNA:CR | 0 | NA | NA | NA | NA | NA |
| lncRNA:CR | 0 | NA | NA | NA | NA | NA |
| lncRNA:CR | 0 | NA | NA | NA | NA | NA |
| lncRNA:CR | 60.90505 | 6.471408 | 3.858031 | 1.677386 | 0.093467 | NA |
| lncRNA:CR | 0.104623 | 0.639075 | 4.067674 | 0.157111 | 0.875158 | NA |
| CG44622 | 0 | NA | NA | NA | NA | NA |
| CG44623 | 0 | NA | NA | NA | NA | NA |
| CG44624 | 0 | NA | NA | NA | NA | NA |
| CG44625 | 0 | NA | NA | NA | NA | NA |
| lncRNA:CR | 0 | NA | NA | NA | NA | NA |
| lncRNA:CR | 0 | NA | NA | NA | NA | NA |
| lncRNA:CR | 0 | NA | NA | NA | NA | NA |
| lncRNA:CR | 0 | NA | NA | NA | NA | NA |
| lncRNA:CR | 0.104623 | 0.639075 | 4.067674 | 0.157111 | 0.875158 | NA |
| lncRNA:CR | 4.920897 | 1.714724 | 3.961081 | 0.432893 | 0.665093 | NA |
| lncRNA:CR | 0 | NA | NA | NA | NA | NA |
| lncRNA:CR | 25.63925 | -5.92924 | 3.896742 | -1.52159 | 0.128112 | NA |
| lncRNA:CR | 0 | NA | NA | NA | NA | NA |
| lncRNA:CR | 0 | NA | NA | NA | NA | NA |
| lncRNA:CR | 0.132078 | -1.13211 | 4.069918 | -0.27817 | 0.780885 | NA |
| lncRNA:CR | 0 | NA | NA | NA | NA | NA |
| lncRNA:CR | 11.0393 | -1.69765 | 3.954364 | -0.42931 | 0.667697 | NA |
| lncRNA:CR | 2.814602 | 0.57184 | 3.871466 | 0.147706 | 0.882575 | NA |
| lncRNA:CR | 12.14519 | 1.737886 | 3.955133 | 0.4394 | 0.660372 | NA |
| lncRNA:CR | 17.69429 | 1.74306 | 3.953824 | 0.440854 | 0.659319 | NA |
| lncRNA:CR | 0 | NA | NA | NA | NA | NA |
| lncRNA:CR | 0 | NA | NA | NA | NA | NA |
| lncRNA:CR | 0 | NA | NA | NA | NA | NA |
| lncRNA:CR | 0 | NA | NA | NA | NA | NA |
| lncRNA:CR | 0 | NA | NA | NA | NA | NA |
| lncRNA:CR | 0 | NA | NA | NA | NA | NA |

| | | | | | | |
|---|---|---|---|---|---|---|
| lncRNA:CR | 0 | NA | NA | NA | NA | NA |
| asRNA:CR4 | 0 | NA | NA | NA | NA | NA |
| lncRNA:CR | 0.1047 | 0.613261 | 4.062799 | 0.150945 | 0.880019 | NA |
| lncRNA:CR | 1.351073 | -1.13435 | 3.970931 | -0.28566 | 0.775136 | NA |
| lncRNA:CR | 0.356866 | -1.92326 | 4.037889 | -0.4763 | 0.633858 | NA |
| lncRNA:CR | 0 | NA | NA | NA | NA | NA |
| CR44659 | 0 | NA | NA | NA | NA | NA |
| lncRNA:CR | 0 | NA | NA | NA | NA | NA |
| lncRNA:CR | 0 | NA | NA | NA | NA | NA |
| lncRNA:CR | 0.104623 | 0.639075 | 4.067674 | 0.157111 | 0.875158 | NA |
| lncRNA:CR | 0 | NA | NA | NA | NA | NA |
| CG44666 | 0.178053 | -1.16467 | 4.075067 | -0.2858 | 0.775028 | NA |
| lncRNA:CR | 0 | NA | NA | NA | NA | NA |
| lncRNA:CR | 0 | NA | NA | NA | NA | NA |
| lncRNA:CR | 0 | NA | NA | NA | NA | NA |
| lncRNA:CR | 0 | NA | NA | NA | NA | NA |
| lncRNA:CR | 0 | NA | NA | NA | NA | NA |
| lncRNA:CR | 0 | NA | NA | NA | NA | NA |
| lncRNA:CR | 0 | NA | NA | NA | NA | NA |
| lncRNA:CR | 0.1047 | 0.613261 | 4.062799 | 0.150945 | 0.880019 | NA |
| lncRNA:CR | 0 | NA | NA | NA | NA | NA |
| lncRNA:CR | 0 | NA | NA | NA | NA | NA |
| lncRNA:CR | 0 | NA | NA | NA | NA | NA |
| lncRNA:CR | 0 | NA | NA | NA | NA | NA |
| lncRNA:CR | 0.132078 | -1.13211 | 4.069918 | -0.27817 | 0.780885 | NA |
| lncRNA:CR | 0 | NA | NA | NA | NA | NA |
| lncRNA:CR | 0 | NA | NA | NA | NA | NA |
| lncRNA:CR | 0 | NA | NA | NA | NA | NA |
| lncRNA:CR | 0 | NA | NA | NA | NA | NA |
| lncRNA:CR | 0 | NA | NA | NA | NA | NA |
| lncRNA:CR | 0 | NA | NA | NA | NA | NA |
| lncRNA:CR | 0 | NA | NA | NA | NA | NA |
| lncRNA:CR | 0 | NA | NA | NA | NA | NA |
| lncRNA:CR | 0 | NA | NA | NA | NA | NA |
| lncRNA:CR | 0 | NA | NA | NA | NA | NA |
| asRNA:CR4 | 3.030212 | 1.13655 | 3.965915 | 0.28658 | 0.774434 | NA |
| lncRNA:CR | 0 | NA | NA | NA | NA | NA |
| lncRNA:CR | 3.204959 | -1.67301 | 3.96255 | -0.42221 | 0.672875 | NA |
| lncRNA:CR | 0 | NA | NA | NA | NA | NA |
| lncRNA:CR | 0 | NA | NA | NA | NA | NA |
| lncRNA:CR | 0 | NA | NA | NA | NA | NA |
| lncRNA:CR | 0 | NA | NA | NA | NA | NA |
| lncRNA:CR | 0 | NA | NA | NA | NA | NA |
| lncRNA:CR | 0 | NA | NA | NA | NA | NA |
| lncRNA:CR | 0 | NA | NA | NA | NA | NA |
| lncRNA:CR | 0 | NA | NA | NA | NA | NA |
| lncRNA:CR | 0 | NA | NA | NA | NA | NA |
| lncRNA:CR | 24.18053 | 0.369972 | 3.836136 | 0.096444 | 0.923168 | NA |
| asRNA:CR4 | 9.520716 | 1.525762 | 3.954655 | 0.385814 | 0.699634 | NA |
| asRNA:CR4 | 12.65941 | 1.529109 | 3.953735 | 0.386751 | 0.698941 | NA |

| | | | | | | |
|---|---|---|---|---|---|---|
| lncRNA:CR | 0 | NA | NA | NA | NA | NA |
| CR44706 | 0 | NA | NA | NA | NA | NA |
| lncRNA:CR | 0 | NA | NA | NA | NA | NA |
| lncRNA:CR | 0 | NA | NA | NA | NA | NA |
| lncRNA:CR | 0 | NA | NA | NA | NA | NA |
| lncRNA:CR | 0 | NA | NA | NA | NA | NA |
| lncRNA:CR | 0 | NA | NA | NA | NA | NA |
| lncRNA:CR | 0 | NA | NA | NA | NA | NA |
| lncRNA:CR | 0 | NA | NA | NA | NA | NA |
| lncRNA:CR | 9.004194 | 1.73223 | 3.956573 | 0.437811 | 0.661523 | NA |
| lncRNA:CR | 13.52991 | 4.302045 | 3.889439 | 1.106084 | 0.26869 | NA |
| lncRNA:CR | 0 | NA | NA | NA | NA | NA |
| lncRNA:CR | 8.132723 | -5.4006 | 3.911847 | -1.38058 | 0.167409 | NA |
| lncRNA:CR | 0 | NA | NA | NA | NA | NA |
| lncRNA:CR | 0 | NA | NA | NA | NA | NA |
| lncRNA:CR | 0 | NA | NA | NA | NA | NA |
| lncRNA:CR | 0 | NA | NA | NA | NA | NA |
| lncRNA:CR | 0.752547 | 0.662441 | 3.960528 | 0.167261 | 0.867165 | NA |
| asRNA:CR4 | 32.72265 | -2.27063 | 3.95384 | -0.57428 | 0.565776 | NA |
| lncRNA:CR | 0 | NA | NA | NA | NA | NA |
| lncRNA:CR | 0 | NA | NA | NA | NA | NA |
| lncRNA:CR | 0.488184 | -2.32943 | 4.022674 | -0.57908 | 0.562538 | NA |
| lncRNA:CR | 0 | NA | NA | NA | NA | NA |
| lncRNA:CR | 0 | NA | NA | NA | NA | NA |
| lncRNA:CR | 0 | NA | NA | NA | NA | NA |
| lncRNA:CR | 0 | NA | NA | NA | NA | NA |
| lncRNA:CR | 0.233093 | 0.692656 | 4.076246 | 0.169925 | 0.865069 | NA |
| lncRNA:CR | 0 | NA | NA | NA | NA | NA |
| lncRNA:CR | 0 | NA | NA | NA | NA | NA |
| lncRNA:CR | 0 | NA | NA | NA | NA | NA |
| lncRNA:CR | 0 | NA | NA | NA | NA | NA |
| lncRNA:CR | 21.36913 | -1.85672 | 3.95265 | -0.46974 | 0.63854 | NA |
| lncRNA:CR | 0.357625 | -1.47707 | 4.044794 | -0.36518 | 0.714979 | NA |
| lncRNA:CR | 0 | NA | NA | NA | NA | NA |
| lncRNA:CR | 0 | NA | NA | NA | NA | NA |
| lncRNA:CR | 0 | NA | NA | NA | NA | NA |
| lncRNA:CR | 0.178053 | -1.16467 | 4.075067 | -0.2858 | 0.775028 | NA |
| lncRNA:CR | 28.74697 | -1.04237 | 3.91275 | -0.2664 | 0.789928 | NA |
| lncRNA:CR | 0 | NA | NA | NA | NA | NA |
| lncRNA:CR | 0 | NA | NA | NA | NA | NA |
| lncRNA:CR | 0 | NA | NA | NA | NA | NA |
| lncRNA:CR | 0 | NA | NA | NA | NA | NA |
| lncRNA:CR | 0 | NA | NA | NA | NA | NA |
| lncRNA:CR | 0 | NA | NA | NA | NA | NA |
| lncRNA:CR | 0.132078 | -1.13211 | 4.069918 | -0.27817 | 0.780885 | NA |
| lncRNA:CR | 1.259804 | -3.02464 | 3.93518 | -0.76862 | 0.442122 | NA |
| Tfb5 | 0 | NA | NA | NA | NA | NA |
| lncRNA:CR | 0 | NA | NA | NA | NA | NA |
| lncRNA:CR | 0.233093 | 0.692656 | 4.076246 | 0.169925 | 0.865069 | NA |
| lncRNA:CR | 9.416093 | 1.525613 | 3.954696 | 0.385773 | 0.699665 | NA |

| | | | | | | |
|---|---|---|---|---|---|---|
| lncRNA:CR | 51.4574 | -1.70598 | 3.951653 | -0.43171 | 0.66595 | NA |
| lncRNA:CR | 0 | NA | NA | NA | NA | NA |
| lncRNA:CR | 0 | NA | NA | NA | NA | NA |
| lncRNA:CR | 0 | NA | NA | NA | NA | NA |
| lncRNA:CR | 0 | NA | NA | NA | NA | NA |
| lncRNA:CR | 0 | NA | NA | NA | NA | NA |
| lncRNA:CR | 0 | NA | NA | NA | NA | NA |
| lncRNA:CR | 0 | NA | NA | NA | NA | NA |
| lncRNA:CR | 0 | NA | NA | NA | NA | NA |
| lncRNA:CR | 0 | NA | NA | NA | NA | NA |
| lncRNA:CR | 0.1047 | 0.613261 | 4.062799 | 0.150945 | 0.880019 | NA |
| lncRNA:CR | 0.337793 | 1.451235 | 4.039844 | 0.35923 | 0.719423 | NA |
| lncRNA:CR | 0 | NA | NA | NA | NA | NA |
| lncRNA:CR | 0 | NA | NA | NA | NA | NA |
| lncRNA:CR | 0 | NA | NA | NA | NA | NA |
| lncRNA:CR | 0.104623 | 0.639075 | 4.067674 | 0.157111 | 0.875158 | NA |
| lncRNA:CR | 49.68049 | -3.09354 | 3.917238 | -0.78972 | 0.429689 | NA |
| lncRNA:CR | 0 | NA | NA | NA | NA | NA |
| lncRNA:CR | 0.132078 | -1.13211 | 4.069918 | -0.27817 | 0.780885 | NA |
| lncRNA:CR | 0 | NA | NA | NA | NA | NA |
| lncRNA:CR | 0 | NA | NA | NA | NA | NA |
| lncRNA:CR | 0 | NA | NA | NA | NA | NA |
| lncRNA:CR | 0.1047 | 0.613261 | 4.062799 | 0.150945 | 0.880019 | NA |
| lncRNA:CR | 0 | NA | NA | NA | NA | NA |
| lncRNA:CR | 0 | NA | NA | NA | NA | NA |
| lncRNA:CR | 28.3967 | -1.9183 | 3.952279 | -0.48537 | 0.627417 | NA |
| lncRNA:CR | 0 | NA | NA | NA | NA | NA |
| lncRNA:CR | 0 | NA | NA | NA | NA | NA |
| asRNA:TS1 | 0 | NA | NA | NA | NA | NA |
| asRNA:CR4 | 0 | NA | NA | NA | NA | NA |
| lncRNA:CR | 0.233093 | 0.692656 | 4.076246 | 0.169925 | 0.865069 | NA |
| lncRNA:CR | 0 | NA | NA | NA | NA | NA |
| lncRNA:CR | 0 | NA | NA | NA | NA | NA |
| lncRNA:CR | 0 | NA | NA | NA | NA | NA |
| lncRNA:CR | 0 | NA | NA | NA | NA | NA |
| lncRNA:CR | 0 | NA | NA | NA | NA | NA |
| lncRNA:CR | 26.32944 | -3.50887 | 3.901453 | -0.89938 | 0.368453 | NA |
| lncRNA:CR | 0 | NA | NA | NA | NA | NA |
| lncRNA:CR | 0.178812 | -1.08808 | 4.061735 | -0.26789 | 0.788788 | NA |
| lncRNA:CR | 0.356107 | -1.47099 | 4.037764 | -0.36431 | 0.715628 | NA |
| lncRNA:CR | 0 | NA | NA | NA | NA | NA |
| asRNA:CR4 | 0 | NA | NA | NA | NA | NA |
| lncRNA:CR | 0 | NA | NA | NA | NA | NA |
| lncRNA:CR | 37.06377 | 1.749099 | 3.952308 | 0.442551 | 0.65809 | NA |
| lncRNA:CR | 0 | NA | NA | NA | NA | NA |
| lncRNA:CR | 0 | NA | NA | NA | NA | NA |
| lncRNA:CR | 0.536437 | -1.71175 | 4.038042 | -0.42391 | 0.671635 | NA |
| CG44815 | 48.71464 | -6.40365 | 3.803354 | -1.68369 | 0.092243 | NA |
| lncRNA:CR | 0 | NA | NA | NA | NA | NA |
| lncRNA:CR | 33.01942 | -1.91897 | 3.952088 | -0.48556 | 0.62728 | NA |

| | | | | | | |
|---|---|---|---|---|---|---|
| asRNA:CR4 | 0 | NA | NA | NA | NA | NA |
| lncRNA:CR | 0 | NA | NA | NA | NA | NA |
| lncRNA:CR | 0.178053 | -1.16467 | 4.075067 | -0.2858 | 0.775028 | NA |
| lncRNA:CR | 0.1047 | 0.613261 | 4.062799 | 0.150945 | 0.880019 | NA |
| lncRNA:CR | 0.178053 | -1.16467 | 4.075067 | -0.2858 | 0.775028 | NA |
| lncRNA:CR | 0 | NA | NA | NA | NA | NA |
| lncRNA:CR | 0 | NA | NA | NA | NA | NA |
| lncRNA:CR | 0 | NA | NA | NA | NA | NA |
| lncRNA:CR | 33.36378 | -0.87424 | 3.949021 | -0.22138 | 0.824796 | NA |
| lncRNA:CR | 0 | NA | NA | NA | NA | NA |
| lncRNA:CR | 26.25761 | 0.295888 | 3.835781 | 0.077139 | 0.938513 | NA |
| lncRNA:CR | 0.178053 | -1.16467 | 4.075067 | -0.2858 | 0.775028 | NA |
| lncRNA:CR | 6.386696 | 1.723466 | 3.95882 | 0.435348 | 0.66331 | NA |
| lncRNA:CR | 45.16509 | 1.606671 | 3.397908 | 0.472841 | 0.636327 | NA |
| lncRNA:CR | 23.42968 | -1.65532 | 3.952493 | -0.41881 | 0.675359 | NA |
| lncRNA:CR | 0.233093 | 0.692656 | 4.076246 | 0.169925 | 0.865069 | NA |
| CG44836 | 0.675302 | -2.05376 | 3.984422 | -0.51545 | 0.606241 | NA |
| CR44839 | 0 | NA | NA | NA | NA | NA |
| ana3 | 2.703587 | -2.11665 | 3.896693 | -0.54319 | 0.586998 | NA |
| asRNA:CR4 | 0 | NA | NA | NA | NA | NA |
| lncRNA:CR | 0 | NA | NA | NA | NA | NA |
| asRNA:CR4 | 3.86362 | -1.36225 | 3.963244 | -0.34372 | 0.731057 | NA |
| Gyc76C | 10.67704 | -1.33672 | 3.747935 | -0.35666 | 0.72135 | NA |
| lncRNA:CR | 29.95418 | -1.87492 | 3.952167 | -0.4744 | 0.635212 | NA |
| lncRNA:CR | 0.1047 | 0.613261 | 4.062799 | 0.150945 | 0.880019 | NA |
| lncRNA:CR | 0.1047 | 0.613261 | 4.062799 | 0.150945 | 0.880019 | NA |
| asRNA:CR4 | 0 | NA | NA | NA | NA | NA |
| lncRNA:CR | 1.849088 | -1.85577 | 3.970721 | -0.46736 | 0.640241 | NA |
| asRNA:CR4 | 0 | NA | NA | NA | NA | NA |
| asRNA:CR4 | 0 | NA | NA | NA | NA | NA |
| lncRNA:CR | 40.77652 | -0.3465 | 3.951463 | -0.08769 | 0.930124 | NA |
| lncRNA:CR | 0 | NA | NA | NA | NA | NA |
| lncRNA:CR | 0 | NA | NA | NA | NA | NA |
| lncRNA:CR | 0 | NA | NA | NA | NA | NA |
| lncRNA:CR | 0 | NA | NA | NA | NA | NA |
| lncRNA:CR | 0 | NA | NA | NA | NA | NA |
| lncRNA:CR | 0 | NA | NA | NA | NA | NA |
| lncRNA:CR | 0 | NA | NA | NA | NA | NA |
| lncRNA:CR | 0 | NA | NA | NA | NA | NA |
| lncRNA:CR | 0 | NA | NA | NA | NA | NA |
| lncRNA:CR | 0 | NA | NA | NA | NA | NA |
| lncRNA:CR | 0.178812 | -1.08808 | 4.061735 | -0.26789 | 0.788788 | NA |
| lncRNA:CR | 0 | NA | NA | NA | NA | NA |
| asRNA:CR4 | 0 | NA | NA | NA | NA | NA |
| lncRNA:CR | 0 | NA | NA | NA | NA | NA |
| lncRNA:CR | 0.732899 | 1.548046 | 4.007786 | 0.38626 | 0.699304 | NA |
| CG44869 | 0 | NA | NA | NA | NA | NA |
| lncRNA:CR | 0 | NA | NA | NA | NA | NA |
| lncRNA:CR | 0.712213 | -1.57119 | 3.998613 | -0.39293 | 0.694369 | NA |
| asRNA:CR4 | 0 | NA | NA | NA | NA | NA |

| | | | | | | |
|---|---|---|---|---|---|---|
| asRNA:CR4 | 13.47192 | -1.91304 | 3.953785 | -0.48385 | 0.628491 | NA |
| lncRNA:CR | 58.58902 | 1.537214 | 3.951522 | 0.389018 | 0.697263 | NA |
| lncRNA:CR | 0 | NA | NA | NA | NA | NA |
| pre-mod(n | 0 | NA | NA | NA | NA | NA |
| pre-mod(n | 0 | NA | NA | NA | NA | NA |
| pre-mod(n | 0 | NA | NA | NA | NA | NA |
| pre-mod(n | 0 | NA | NA | NA | NA | NA |
| pre-mod(n | 0 | NA | NA | NA | NA | NA |
| pre-mod(n | 0 | NA | NA | NA | NA | NA |
| pre-mod(n | 0 | NA | NA | NA | NA | NA |
| pre-mod(n | 0 | NA | NA | NA | NA | NA |
| pre-mod(n | 0 | NA | NA | NA | NA | NA |
| lncRNA:CR | 0 | NA | NA | NA | NA | NA |
| lncRNA:CR | 14.34389 | 1.740403 | 3.954495 | 0.440108 | 0.659859 | NA |
| lncRNA:CR | 0 | NA | NA | NA | NA | NA |
| lncRNA:CR | 0 | NA | NA | NA | NA | NA |
| lncRNA:CR | 0 | NA | NA | NA | NA | NA |
| Tgs1 | 0 | NA | NA | NA | NA | NA |
| asRNA:CR4 | 0 | NA | NA | NA | NA | NA |
| lncRNA:CR | 0 | NA | NA | NA | NA | NA |
| asRNA:CR4 | 2.128938 | -1.0769 | 3.887431 | -0.27702 | 0.781765 | NA |
| lncRNA:CR | 0.357625 | -1.47707 | 4.044794 | -0.36518 | 0.714979 | NA |
| asRNA:CR4 | 0 | NA | NA | NA | NA | NA |
| asRNA:CR4 | 0 | NA | NA | NA | NA | NA |
| lncRNA:CR | 0 | NA | NA | NA | NA | NA |
| lncRNA:CR | 0 | NA | NA | NA | NA | NA |
| lncRNA:CR | 14.91797 | 1.17086 | 3.95405 | 0.296117 | 0.767141 | NA |
| lncRNA:CR | 0 | NA | NA | NA | NA | NA |
| lncRNA:CR | 0 | NA | NA | NA | NA | NA |
| lncRNA:CR | 0 | NA | NA | NA | NA | NA |
| lncRNA:CR | 0 | NA | NA | NA | NA | NA |
| CR44906 | 0 | NA | NA | NA | NA | NA |
| asRNA:CR4 | 0 | NA | NA | NA | NA | NA |
| lncRNA:CR | 0 | NA | NA | NA | NA | NA |
| asRNA:CR4 | 0 | NA | NA | NA | NA | NA |
| asRNA:CR4 | 46.52949 | 2.372784 | 3.844405 | 0.617204 | 0.5371 | NA |
| lncRNA:CR | 0 | NA | NA | NA | NA | NA |
| lncRNA:CR | 0 | NA | NA | NA | NA | NA |
| lncRNA:CR | 19.96634 | 1.650754 | 3.953312 | 0.417562 | 0.676267 | NA |
| lncRNA:CR | 0 | NA | NA | NA | NA | NA |
| lncRNA:CR | 0 | NA | NA | NA | NA | NA |
| lncRNA:CR | 0 | NA | NA | NA | NA | NA |
| lncRNA:CR | 0.387376 | 0.625691 | 4.009146 | 0.156066 | 0.875981 | NA |
| lncRNA:CR | 0 | NA | NA | NA | NA | NA |
| lncRNA:CR | 0 | NA | NA | NA | NA | NA |
| lncRNA:CR | 0.233093 | 0.692656 | 4.076246 | 0.169925 | 0.865069 | NA |
| lncRNA:CR | 0 | NA | NA | NA | NA | NA |
| lncRNA:CR | 0 | NA | NA | NA | NA | NA |
| lncRNA:CR | 0 | NA | NA | NA | NA | NA |
| lncRNA:CR | 0 | NA | NA | NA | NA | NA |

| | | | | | | |
|---|---|---|---|---|---|---|
| lncRNA:CR | 0.104623 | 0.639075 | 4.067674 | 0.157111 | 0.875158 | NA |
| lncRNA:CR | 0.178053 | -1.16467 | 4.075067 | -0.2858 | 0.775028 | NA |
| lncRNA:CR | 0 | NA | NA | NA | NA | NA |
| lncRNA:CR | 0 | NA | NA | NA | NA | NA |
| lncRNA:CR | 0 | NA | NA | NA | NA | NA |
| lncRNA:CR | 0 | NA | NA | NA | NA | NA |
| lncRNA:CR | 0 | NA | NA | NA | NA | NA |
| lncRNA:CR | 0 | NA | NA | NA | NA | NA |
| lncRNA:CR | 0 | NA | NA | NA | NA | NA |
| lncRNA:CR | 0 | NA | NA | NA | NA | NA |
| lncRNA:CR | 6.247973 | 0.287943 | 3.851599 | 0.074759 | 0.940406 | NA |
| lncRNA:CR | 0 | NA | NA | NA | NA | NA |
| lncRNA:CR | 0 | NA | NA | NA | NA | NA |
| lncRNA:CR | 0 | NA | NA | NA | NA | NA |
| lncRNA:CR | 0 | NA | NA | NA | NA | NA |
| lncRNA:CR | 0.282677 | -0.09599 | 4.057934 | -0.02366 | 0.981127 | NA |
| lncRNA:CR | 0 | NA | NA | NA | NA | NA |
| lncRNA:CR | 0 | NA | NA | NA | NA | NA |
| lncRNA:CR | 0.178053 | -1.16467 | 4.075067 | -0.2858 | 0.775028 | NA |
| lncRNA:CR | 0.466186 | 0.959471 | 4.033986 | 0.237847 | 0.812 | NA |
| lncRNA:CR | 0 | NA | NA | NA | NA | NA |
| lncRNA:CR | 0 | NA | NA | NA | NA | NA |
| lncRNA:CR | 0 | NA | NA | NA | NA | NA |
| lncRNA:CR | 0.233093 | 0.692656 | 4.076246 | 0.169925 | 0.865069 | NA |
| lncRNA:CR | 0 | NA | NA | NA | NA | NA |
| lncRNA:CR | 0.132078 | -1.13211 | 4.069918 | -0.27817 | 0.780885 | NA |
| CG44954 | 0 | NA | NA | NA | NA | NA |
| lncRNA:CR | 0.337793 | 1.451235 | 4.039844 | 0.35923 | 0.719423 | NA |
| CG44956 | 0 | NA | NA | NA | NA | NA |
| lncRNA:CR | 0.2094 | 1.005034 | 4.045341 | 0.248442 | 0.803792 | NA |
| asRNA:CR4 | 0 | NA | NA | NA | NA | NA |
| asRNA:CR4 | 0 | NA | NA | NA | NA | NA |
| asRNA:CR4 | 1.070597 | -3.05742 | 3.990949 | -0.76609 | 0.443624 | NA |
| asRNA:CR4 | 0.806253 | 0.654822 | 3.981166 | 0.16448 | 0.869353 | NA |
| asRNA:CR4 | 2.860997 | -2.16734 | 3.980875 | -0.54444 | 0.586141 | NA |
| lncRNA:CR | 0 | NA | NA | NA | NA | NA |
| lncRNA:CR | 0 | NA | NA | NA | NA | NA |
| lncRNA:CR | 21.43243 | -3.03925 | 3.86974 | -0.78539 | 0.432226 | NA |
| lncRNA:CR | 0 | NA | NA | NA | NA | NA |
| lncRNA:CR | 0 | NA | NA | NA | NA | NA |
| asRNA:CR4 | 0 | NA | NA | NA | NA | NA |
| lncRNA:CR | 0 | NA | NA | NA | NA | NA |
| lncRNA:CR | 0 | NA | NA | NA | NA | NA |
| asRNA:CR4 | 0 | NA | NA | NA | NA | NA |
| lncRNA:CR | 0.5235 | 1.490714 | 4.02555 | 0.370313 | 0.711149 | NA |
| lncRNA:CR | 0 | NA | NA | NA | NA | NA |
| asRNA:CR4 | 10.67157 | 1.527223 | 3.954255 | 0.386223 | 0.699332 | NA |
| asRNA:CR4 | 0 | NA | NA | NA | NA | NA |
| lncRNA:CR | 24.70517 | 0.356184 | 3.838128 | 0.092801 | 0.926061 | NA |
| lncRNA:CR | 0 | NA | NA | NA | NA | NA |

| | | | | | | |
|---|---|---|---|---|---|---|
| asRNA:CR4 | 0 | NA | NA | NA | NA | NA |
| lncRNA:CR | 0 | NA | NA | NA | NA | NA |
| asRNA:CR4 | 0 | NA | NA | NA | NA | NA |
| lncRNA:CR | 0.1047 | 0.613261 | 4.062799 | 0.150945 | 0.880019 | NA |
| lncRNA:CR | 0 | NA | NA | NA | NA | NA |
| lncRNA:CR | 0 | NA | NA | NA | NA | NA |
| asRNA:CR4 | 0 | NA | NA | NA | NA | NA |
| asRNA:CR4 | 0 | NA | NA | NA | NA | NA |
| lncRNA:CR | 18.94521 | -0.23186 | 3.861384 | -0.06005 | 0.952119 | NA |
| lncRNA:CR | 33.64127 | -0.93046 | 3.896581 | -0.23879 | 0.811269 | NA |
| lncRNA:CR | 0.178053 | -1.16467 | 4.075067 | -0.2858 | 0.775028 | NA |
| asRNA:CR4 | 0 | NA | NA | NA | NA | NA |
| asRNA:CR4 | 1.06832 | -1.61145 | 3.983918 | -0.40449 | 0.685853 | NA |
| asRNA:CR4 | 6.568725 | -2.05969 | 3.83095 | -0.53764 | 0.590823 | NA |
| lncRNA:CR | 0 | NA | NA | NA | NA | NA |
| asRNA:CR4 | 4.816197 | 1.713907 | 3.961293 | 0.432664 | 0.665259 | NA |
| asRNA:CR4 | 0 | NA | NA | NA | NA | NA |
| lncRNA:CR | 0 | NA | NA | NA | NA | NA |
| lncRNA:CR | 24.91851 | 2.96466 | 3.92141 | 0.756019 | 0.449638 | NA |
| lncRNA:CR | 0 | NA | NA | NA | NA | NA |
| lncRNA:CR | 0 | NA | NA | NA | NA | NA |
| asRNA:CR4 | 0 | NA | NA | NA | NA | NA |
| asRNA:CR4 | 0 | NA | NA | NA | NA | NA |
| lncRNA:CR | 0 | NA | NA | NA | NA | NA |
| CG45011 | 0.178053 | -1.16467 | 4.075067 | -0.2858 | 0.775028 | NA |
| CG45012 | 0 | NA | NA | NA | NA | NA |
| lncRNA:CR | 0 | NA | NA | NA | NA | NA |
| lncRNA:CR | 0 | NA | NA | NA | NA | NA |
| asRNA:CR4 | 0 | NA | NA | NA | NA | NA |
| asRNA:CR4 | 7.642194 | -3.40016 | 3.943094 | -0.86231 | 0.388519 | NA |
| lncRNA:CR | 0 | NA | NA | NA | NA | NA |
| lncRNA:CR | 0 | NA | NA | NA | NA | NA |
| lncRNA:CR | 0.1047 | 0.613261 | 4.062799 | 0.150945 | 0.880019 | NA |
| lncRNA:CR | 0 | NA | NA | NA | NA | NA |
| lncRNA:CR | 3.978597 | 1.705937 | 3.963372 | 0.430426 | 0.666886 | NA |
| asRNA:CR4 | 0 | NA | NA | NA | NA | NA |
| asRNA:CR4 | 0 | NA | NA | NA | NA | NA |
| lncRNA:CR | 0 | NA | NA | NA | NA | NA |
| lncRNA:CR | 0 | NA | NA | NA | NA | NA |
| lncRNA:CR | 0 | NA | NA | NA | NA | NA |
| CR45033 | 0 | NA | NA | NA | NA | NA |
| lncRNA:CR | 0 | NA | NA | NA | NA | NA |
| lncRNA:CR | 21.51768 | -0.07372 | 3.880056 | -0.019 | 0.984841 | NA |
| lncRNA:CR | 18.52259 | 5.861174 | 3.857864 | 1.519279 | 0.128692 | NA |
| lncRNA:CR | 0 | NA | NA | NA | NA | NA |
| lncRNA:CR | 0 | NA | NA | NA | NA | NA |
| lncRNA:CR | 0 | NA | NA | NA | NA | NA |
| lncRNA:CR | 0 | NA | NA | NA | NA | NA |
| asRNA:CR4 | 0 | NA | NA | NA | NA | NA |
| asRNA:CR4 | 0 | NA | NA | NA | NA | NA |

| | | | | | | |
|---|---|---|---|---|---|---|
| lncRNA:CR | 0 | NA | NA | NA | NA | NA |
| asRNA:CR4 | 3.729492 | 1.144349 | 3.963186 | 0.288745 | 0.772777 | NA |
| CG45045 | 0 | NA | NA | NA | NA | NA |
| asRNA:CR4 | 0 | NA | NA | NA | NA | NA |
| lncRNA:CR | 15.06118 | -1.62798 | 3.95333 | -0.4118 | 0.680487 | NA |
| lncRNA:CR | 54.756 | -9.28132 | 3.844412 | -2.41424 | 0.015768 | NA |
| lncRNA:CR | 25.81773 | -1.70371 | 3.952393 | -0.43106 | 0.666426 | NA |
| asRNA:CR4 | 6.972663 | -1.74081 | 3.955814 | -0.44006 | 0.65989 | NA |
| lncRNA:CR | 0 | NA | NA | NA | NA | NA |
| asRNA:CR4 | 0.712213 | -1.57119 | 3.998613 | -0.39293 | 0.694369 | NA |
| lncRNA:CR | 0 | NA | NA | NA | NA | NA |
| asRNA:CR4 | 0 | NA | NA | NA | NA | NA |
| CG45060 | 0 | NA | NA | NA | NA | NA |
| CG45061 | 4.397397 | 1.710278 | 3.962238 | 0.431644 | 0.666 | NA |
| CG45062 | 3.830253 | -1.88895 | 3.960794 | -0.47691 | 0.633424 | NA |
| CG45063 | 0.178053 | -1.16467 | 4.075067 | -0.2858 | 0.775028 | NA |
| CG45064 | 0 | NA | NA | NA | NA | NA |
| CG45065 | 0 | NA | NA | NA | NA | NA |
| CG45066 | 0 | NA | NA | NA | NA | NA |
| CG45067 | 0 | NA | NA | NA | NA | NA |
| PIG-Z | 0 | NA | NA | NA | NA | NA |
| CG45069 | 0 | NA | NA | NA | NA | NA |
| CG45072 | 0.132078 | -1.13211 | 4.069918 | -0.27817 | 0.780885 | NA |
| CG45073 | 0 | NA | NA | NA | NA | NA |
| CG45075 | 0 | NA | NA | NA | NA | NA |
| CG45076 | 0.233093 | 0.692656 | 4.076246 | 0.169925 | 0.865069 | NA |
| CG45078 | 0 | NA | NA | NA | NA | NA |
| fau | 0 | NA | NA | NA | NA | NA |
| CG45079 | 0 | NA | NA | NA | NA | NA |
| CG45080 | 0 | NA | NA | NA | NA | NA |
| CG45081 | 0 | NA | NA | NA | NA | NA |
| lncRNA:CR | 26.78423 | -0.33788 | 3.8373 | -0.08805 | 0.929836 | NA |
| CG45084 | 0 | NA | NA | NA | NA | NA |
| CG45085 | 0 | NA | NA | NA | NA | NA |
| CG45086 | 0 | NA | NA | NA | NA | NA |
| CG45087 | 0 | NA | NA | NA | NA | NA |
| CG45088 | 0 | NA | NA | NA | NA | NA |
| CG45089 | 0 | NA | NA | NA | NA | NA |
| SclA | 0 | NA | NA | NA | NA | NA |
| SclB | 0 | NA | NA | NA | NA | NA |
| CG45092 | 0 | NA | NA | NA | NA | NA |
| CG45095 | 0 | NA | NA | NA | NA | NA |
| lncRNA:CR | 0 | NA | NA | NA | NA | NA |
| lncRNA:CR | 0 | NA | NA | NA | NA | NA |
| lncRNA:CR | 0 | NA | NA | NA | NA | NA |
| CG45099 | 0 | NA | NA | NA | NA | NA |
| CG45100 | 0 | NA | NA | NA | NA | NA |
| CG45101 | 0 | NA | NA | NA | NA | NA |
| lncRNA:CR | 0 | NA | NA | NA | NA | NA |
| asRNA:CR4 | 1.762219 | -0.18545 | 3.949645 | -0.04695 | 0.962551 | NA |

| | | | | | | |
|---|---|---|---|---|---|---|
| CG45105 | 0.178053 | -1.16467 | 4.075067 | -0.2858 | 0.775028 | NA |
| lncRNA:yaı | 0.104623 | 0.639075 | 4.067674 | 0.157111 | 0.875158 | NA |
| tRNA:Thr-T | 0 | NA | NA | NA | NA | NA |
| tRNA:Thr-T | 0 | NA | NA | NA | NA | NA |
| asRNA:CR4 | 0 | NA | NA | NA | NA | NA |
| lncRNA:CR | 0 | NA | NA | NA | NA | NA |
| tRNA:Leu-( | 0.357625 | -1.47707 | 4.044794 | -0.36518 | 0.714979 | NA |
| lncRNA:CR | 0 | NA | NA | NA | NA | NA |
| lncRNA:CR | 7.264273 | -1.90467 | 3.956202 | -0.48144 | 0.630204 | NA |
| lncRNA:CR | 0 | NA | NA | NA | NA | NA |
| lncRNA:CR | 0 | NA | NA | NA | NA | NA |
| lncRNA:CR | 34.10083 | 1.69146 | 3.952362 | 0.427962 | 0.668679 | NA |
| CG45116 | 0 | NA | NA | NA | NA | NA |
| lncRNA:CR | 0 | NA | NA | NA | NA | NA |
| lncRNA:CR | 0.132078 | -1.13211 | 4.069918 | -0.27817 | 0.780885 | NA |
| asRNA:CR4 | 8.475891 | 5.42735 | 3.881918 | 1.39811 | 0.16208 | NA |
| Kmn2 | 0.984306 | 0.3138 | 3.972938 | 0.078984 | 0.937045 | NA |
| CG45122 | 0 | NA | NA | NA | NA | NA |
| lncRNA:CR | 0 | NA | NA | NA | NA | NA |
| asRNA:CR4 | 13.07791 | 1.529441 | 3.953645 | 0.386843 | 0.698872 | NA |
| lncRNA:CR | 0 | NA | NA | NA | NA | NA |
| asRNA:CR4 | 0.715249 | -1.88064 | 4.034336 | -0.46616 | 0.641102 | NA |
| asRNA:CR4 | 27.02402 | -0.38229 | 3.951361 | -0.09675 | 0.922926 | NA |
| lncRNA:CR | 0.6282 | 1.523021 | 4.015426 | 0.379293 | 0.704471 | NA |
| asRNA:CR4 | 0 | NA | NA | NA | NA | NA |
| asRNA:CR4 | 30.40563 | -3.24507 | 3.925567 | -0.82665 | 0.408435 | NA |
| asRNA:CR4 | 0 | NA | NA | NA | NA | NA |
| asRNA:CR4 | 1.788123 | -2.11324 | 3.996051 | -0.52883 | 0.596922 | NA |
| asRNA:CR4 | 0 | NA | NA | NA | NA | NA |
| asRNA:CR4 | 0 | NA | NA | NA | NA | NA |
| asRNA:CR4 | 6.502747 | 3.328933 | 3.858011 | 0.862862 | 0.388213 | NA |
| asRNA:CR4 | 13.47192 | -1.91304 | 3.953785 | -0.48385 | 0.628491 | NA |
| lncRNA:CR | 0.178053 | -1.16467 | 4.075067 | -0.2858 | 0.775028 | NA |
| lncRNA:CR | 0.528311 | -1.72839 | 4.011871 | -0.43082 | 0.6666 | NA |
| asRNA:CR4 | 0.209247 | 1.042804 | 4.047837 | 0.25762 | 0.7967 | NA |
| lncRNA:CR | 0.132078 | -1.13211 | 4.069918 | -0.27817 | 0.780885 | NA |
| asRNA:CR4 | 0 | NA | NA | NA | NA | NA |
| asRNA:CR4 | 0 | NA | NA | NA | NA | NA |
| lncRNA:CR | 0 | NA | NA | NA | NA | NA |
| CG45154 | 0 | NA | NA | NA | NA | NA |
| lncRNA:CR | 58.58902 | 1.537214 | 3.951522 | 0.389018 | 0.697263 | NA |
| lncRNA:CR | 0 | NA | NA | NA | NA | NA |
| lncRNA:CR | 0 | NA | NA | NA | NA | NA |
| lncRNA:CR | 22.55783 | -1.86002 | 3.952562 | -0.47059 | 0.637936 | NA |
| asRNA:CR4 | 0 | NA | NA | NA | NA | NA |
| asRNA:CR4 | 0 | NA | NA | NA | NA | NA |
| asRNA:CR4 | 0 | NA | NA | NA | NA | NA |
| asRNA:CR4 | 5.444396 | 1.718358 | 3.960139 | 0.433914 | 0.664351 | NA |
| lncRNA:CR | 0 | NA | NA | NA | NA | NA |
| lncRNA:CR | 0.466186 | 0.959471 | 4.033986 | 0.237847 | 0.812 | NA |

| | | | | | | |
|---|---|---|---|---|---|---|
| asRNA:CR4 | 1.001866 | -0.01426 | 3.93462 | -0.00362 | 0.997108 | NA |
| lncRNA:CR | 0 | NA | NA | NA | NA | NA |
| lncRNA:CR | 0 | NA | NA | NA | NA | NA |
| asRNA:CR4 | 0 | NA | NA | NA | NA | NA |
| lncRNA:CR | 0 | NA | NA | NA | NA | NA |
| lncRNA:CR | 0 | NA | NA | NA | NA | NA |
| lncRNA:CR | 0 | NA | NA | NA | NA | NA |
| asRNA:CR4 | 5.842825 | -1.3066 | 3.855864 | -0.33886 | 0.734715 | NA |
| lncRNA:CR | 0.368779 | -0.59727 | 3.999361 | -0.14934 | 0.881285 | NA |
| lncRNA:CR | 0 | NA | NA | NA | NA | NA |
| lncRNA:CR | 0.178053 | -1.16467 | 4.075067 | -0.2858 | 0.775028 | NA |
| lncRNA:CR | 1.574509 | -1.80121 | 3.864174 | -0.46613 | 0.641122 | NA |
| lncRNA:CR | 0 | NA | NA | NA | NA | NA |
| asRNA:CR4 | 24.1766 | -0.41914 | 3.836896 | -0.10924 | 0.913013 | NA |
| lncRNA:CR | 0 | NA | NA | NA | NA | NA |
| lncRNA:CR | 0 | NA | NA | NA | NA | NA |
| lncRNA:CR | 0.536437 | -1.71175 | 4.038042 | -0.42391 | 0.671635 | NA |
| asRNA:CR4 | 36.14557 | -2.93027 | 3.918963 | -0.74771 | 0.454632 | NA |
| asRNA:CR4 | 15.8231 | 1.086724 | 3.773657 | 0.287977 | 0.773365 | NA |
| lncRNA:CR | 0 | NA | NA | NA | NA | NA |
| asRNA:CR4 | 0 | NA | NA | NA | NA | NA |
| lncRNA:CR | 0 | NA | NA | NA | NA | NA |
| asRNA:CR4 | 10.00445 | 2.535938 | 3.852337 | 0.658286 | 0.510355 | NA |
| lncRNA:CR | 43.16435 | -0.24515 | 3.81209 | -0.06431 | 0.948725 | NA |
| lncRNA:CR | 0 | NA | NA | NA | NA | NA |
| Cep89 | 0 | NA | NA | NA | NA | NA |
| Zmynd10 | 0 | NA | NA | NA | NA | NA |
| hog | 0.942299 | 1.584447 | 3.996979 | 0.396411 | 0.691802 | NA |
| asRNA:CR4 | 0 | NA | NA | NA | NA | NA |
| lncRNA:CR | 0 | NA | NA | NA | NA | NA |
| lncRNA:CR | 0 | NA | NA | NA | NA | NA |
| asRNA:CR4 | 0 | NA | NA | NA | NA | NA |
| lncRNA:CR | 55.95161 | 3.2621 | 3.911881 | 0.833895 | 0.40434 | NA |
| lncRNA:CR | 0 | NA | NA | NA | NA | NA |
| asRNA:CR4 | 0 | NA | NA | NA | NA | NA |
| asRNA:CR4 | 0.660388 | -1.75977 | 4.001266 | -0.4398 | 0.660079 | NA |
| CG45207 | 0 | NA | NA | NA | NA | NA |
| asRNA:CR4 | 0 | NA | NA | NA | NA | NA |
| asRNA:CR4 | 0 | NA | NA | NA | NA | NA |
| asRNA:CR4 | 0 | NA | NA | NA | NA | NA |
| lncRNA:CR | 51.87386 | -3.06763 | 3.877788 | -0.79108 | 0.428899 | NA |
| lncRNA:CR | 0 | NA | NA | NA | NA | NA |
| asRNA:CR4 | 0 | NA | NA | NA | NA | NA |
| lncRNA:CR | 0 | NA | NA | NA | NA | NA |
| lncRNA:CR | 0.178812 | -1.08808 | 4.061735 | -0.26789 | 0.788788 | NA |
| lncRNA:CR | 4.292697 | 1.709267 | 3.962502 | 0.431361 | 0.666206 | NA |
| lncRNA:CR | 2.485045 | -1.42724 | 3.888448 | -0.36705 | 0.713585 | NA |
| lncRNA:CR | 34.2073 | -0.339 | 3.836652 | -0.08836 | 0.929592 | NA |
| lncRNA:CR | 0 | NA | NA | NA | NA | NA |
| mfr | 0 | NA | NA | NA | NA | NA |

| | | | | | | |
|---|---|---|---|---|---|---|
| CR45227 | 0.546963 | 2.042449 | 4.016608 | 0.508501 | 0.611102 | NA |
| lncRNA:CR | 0 | NA | NA | NA | NA | NA |
| lncRNA:CR | 0 | NA | NA | NA | NA | NA |
| lncRNA:CR | 0.178053 | -1.16467 | 4.075067 | -0.2858 | 0.775028 | NA |
| lncRNA:CR | 1.046999 | 1.598136 | 3.993006 | 0.400234 | 0.688984 | NA |
| lncRNA:CR | 0 | NA | NA | NA | NA | NA |
| lncRNA:CR | 0 | NA | NA | NA | NA | NA |
| lncRNA:CR | 59.71745 | 0.86843 | 3.668413 | 0.236732 | 0.812865 | NA |
| lncRNA:CR | 0.566112 | -1.1989 | 3.981989 | -0.30108 | 0.763353 | NA |
| lncRNA:CR | 0 | NA | NA | NA | NA | NA |
| lncRNA:CR | 0.132078 | -1.13211 | 4.069918 | -0.27817 | 0.780885 | NA |
| lncRNA:CR | 0 | NA | NA | NA | NA | NA |
| lncRNA:CR | 52.30276 | -1.92048 | 3.951653 | -0.48599 | 0.626971 | NA |
| lncRNA:CR | 53.39712 | -0.42981 | 3.835303 | -0.11207 | 0.91077 | NA |
| lncRNA:CR | 0 | NA | NA | NA | NA | NA |
| lncRNA:CR | 0 | NA | NA | NA | NA | NA |
| lncRNA:CR | 3.034074 | 1.498215 | 3.962342 | 0.378114 | 0.705346 | NA |
| lncRNA:CR | 0 | NA | NA | NA | NA | NA |
| lncRNA:CR | 0 | NA | NA | NA | NA | NA |
| lncRNA:CR | 0 | NA | NA | NA | NA | NA |
| lncRNA:CR | 0.356107 | -1.47099 | 4.037764 | -0.36431 | 0.715628 | NA |
| lncRNA:CR | 0 | NA | NA | NA | NA | NA |
| lncRNA:CR | 0 | NA | NA | NA | NA | NA |
| lncRNA:CR | 0.104623 | 0.639075 | 4.067674 | 0.157111 | 0.875158 | NA |
| lncRNA:CR | 0.178053 | -1.16467 | 4.075067 | -0.2858 | 0.775028 | NA |
| asRNA:CR4 | 0 | NA | NA | NA | NA | NA |
| lncRNA:CR | 0.233093 | 0.692656 | 4.076246 | 0.169925 | 0.865069 | NA |
| lncRNA:CR | 0 | NA | NA | NA | NA | NA |
| asRNA:CR4 | 0 | NA | NA | NA | NA | NA |
| lncRNA:CR | 0 | NA | NA | NA | NA | NA |
| lncRNA:CR | 0.660388 | -1.75977 | 4.001266 | -0.4398 | 0.660079 | NA |
| asRNA:CR4 | 14.91797 | 1.17086 | 3.95405 | 0.296117 | 0.767141 | NA |
| lncRNA:CR | 0 | NA | NA | NA | NA | NA |
| lncRNA:CR | 0 | NA | NA | NA | NA | NA |
| lncRNA:CR | 17.79899 | 1.743127 | 3.953807 | 0.440873 | 0.659305 | NA |
| asRNA:CR4 | 0 | NA | NA | NA | NA | NA |
| lncRNA:CR | 0 | NA | NA | NA | NA | NA |
| asRNA:CR4 | 0 | NA | NA | NA | NA | NA |
| asRNA:CR4 | 0.178053 | -1.16467 | 4.075067 | -0.2858 | 0.775028 | NA |
| CG45273 | 0 | NA | NA | NA | NA | NA |
| lncRNA:CR | 59.64785 | -1.7063 | 3.951551 | -0.4318 | 0.665883 | NA |
| asRNA:CR4 | 0 | NA | NA | NA | NA | NA |
| lncRNA:CR | 0 | NA | NA | NA | NA | NA |
| lncRNA:CR | 1.089612 | 1.039859 | 3.940679 | 0.263878 | 0.791874 | NA |
| lncRNA:CR | 0 | NA | NA | NA | NA | NA |
| lncRNA:CR | 1.188699 | -1.8233 | 3.980741 | -0.45803 | 0.646931 | NA |
| lncRNA:CR | 0 | NA | NA | NA | NA | NA |
| lncRNA:CR | 0 | NA | NA | NA | NA | NA |
| lncRNA:CR | 0.233093 | 0.692656 | 4.076246 | 0.169925 | 0.865069 | NA |
| lncRNA:CR | 11.70063 | -4.87591 | 3.85987 | -1.26323 | 0.206506 | NA |

| | | | | | | |
|---|---|---|---|---|---|---|
| lncRNA:CR | 0 | NA | NA | NA | NA | NA |
| lncRNA:CR | 0.356107 | -1.47099 | 4.037764 | -0.36431 | 0.715628 | NA |
| lncRNA:CR | 0 | NA | NA | NA | NA | NA |
| lncRNA:CR | 9.79293 | -1.69634 | 3.954799 | -0.42893 | 0.667972 | NA |
| lncRNA:CR | 0 | NA | NA | NA | NA | NA |
| lncRNA:CR | 0 | NA | NA | NA | NA | NA |
| lncRNA:CR | 0 | NA | NA | NA | NA | NA |
| lncRNA:CR | 0 | NA | NA | NA | NA | NA |
| lncRNA:CR | 0 | NA | NA | NA | NA | NA |
| lncRNA:CR | 0 | NA | NA | NA | NA | NA |
| lncRNA:CR | 0 | NA | NA | NA | NA | NA |
| lncRNA:CR | 0 | NA | NA | NA | NA | NA |
| lncRNA:CR | 0.282753 | -0.25398 | 4.020204 | -0.06318 | 0.949626 | NA |
| lncRNA:CR | 0 | NA | NA | NA | NA | NA |
| asRNA:CR4 | 0 | NA | NA | NA | NA | NA |
| lncRNA:CR | 0 | NA | NA | NA | NA | NA |
| lncRNA:CR | 0 | NA | NA | NA | NA | NA |
| lncRNA:CR | 0.282753 | -0.25398 | 4.020204 | -0.06318 | 0.949626 | NA |
| lncRNA:CR | 0 | NA | NA | NA | NA | NA |
| asRNA:CR4 | 0 | NA | NA | NA | NA | NA |
| lncRNA:CR | 0 | NA | NA | NA | NA | NA |
| CG45307 | 0 | NA | NA | NA | NA | NA |
| lncRNA:CR | 0.104623 | 0.639075 | 4.067674 | 0.157111 | 0.875158 | NA |
| lncRNA:CR | 0 | NA | NA | NA | NA | NA |
| lncRNA:CR | 0.356107 | -1.47099 | 4.037764 | -0.36431 | 0.715628 | NA |
| lncRNA:CR | 0 | NA | NA | NA | NA | NA |
| lncRNA:CR | 0 | NA | NA | NA | NA | NA |
| CR45315 | 0.178053 | -1.16467 | 4.075067 | -0.2858 | 0.775028 | NA |
| lncRNA:CR | 0 | NA | NA | NA | NA | NA |
| lncRNA:CR | 0 | NA | NA | NA | NA | NA |
| lncRNA:CR | 0 | NA | NA | NA | NA | NA |
| lncRNA:CR | 0 | NA | NA | NA | NA | NA |
| lncRNA:CR | 0 | NA | NA | NA | NA | NA |
| lncRNA:CR | 0 | NA | NA | NA | NA | NA |
| lncRNA:CR | 0 | NA | NA | NA | NA | NA |
| lncRNA:CR | 1.879809 | 1.619289 | 3.898416 | 0.415371 | 0.67787 | NA |
| lncRNA:CR | 3.51509 | -3.04388 | 3.928899 | -0.77474 | 0.438493 | NA |
| asRNA:CR4 | 0.69928 | 1.019866 | 4.009445 | 0.254366 | 0.799213 | NA |
| lncRNA:CR | 0 | NA | NA | NA | NA | NA |
| lncRNA:CR | 0 | NA | NA | NA | NA | NA |
| asRNA:CR4 | 0 | NA | NA | NA | NA | NA |
| lncRNA:CR | 0 | NA | NA | NA | NA | NA |
| lncRNA:CR | 0 | NA | NA | NA | NA | NA |
| lncRNA:CR | 0 | NA | NA | NA | NA | NA |
| lncRNA:CR | 0 | NA | NA | NA | NA | NA |
| lncRNA:CR | 0 | NA | NA | NA | NA | NA |
| lncRNA:CR | 0 | NA | NA | NA | NA | NA |
| lncRNA:CR | 34.25344 | 8.910206 | 3.845848 | 2.316838 | 0.020513 | NA |
| asRNA:CR4 | 0 | NA | NA | NA | NA | NA |
| asRNA:CR4 | 0 | NA | NA | NA | NA | NA |

| | | | | | | |
|---|---|---|---|---|---|---|
| asRNA:CR4 | 0 | NA | NA | NA | NA | NA |
| lncRNA:CR | 0 | NA | NA | NA | NA | NA |
| lncRNA:CR | 0 | NA | NA | NA | NA | NA |
| lncRNA:CR | 0 | NA | NA | NA | NA | NA |
| lncRNA:CR | 0.807511 | 1.368689 | 3.976452 | 0.344199 | 0.730697 | NA |
| lncRNA:CR | 0 | NA | NA | NA | NA | NA |
| asRNA:CR4 | 17.43425 | -1.91531 | 3.953136 | -0.4845 | 0.628029 | NA |
| lncRNA:CR | 22.55783 | -1.86002 | 3.952562 | -0.47059 | 0.637936 | NA |
| lncRNA:CR | 0 | NA | NA | NA | NA | NA |
| lncRNA:CR | 0 | NA | NA | NA | NA | NA |
| lncRNA:CR | 0.132078 | -1.13211 | 4.069918 | -0.27817 | 0.780885 | NA |
| lncRNA:CR | 0 | NA | NA | NA | NA | NA |
| lncRNA:CR | 0.132078 | -1.13211 | 4.069918 | -0.27817 | 0.780885 | NA |
| lncRNA:CR | 0 | NA | NA | NA | NA | NA |
| lncRNA:CR | 0 | NA | NA | NA | NA | NA |
| lncRNA:CR | 0 | NA | NA | NA | NA | NA |
| lncRNA:CR | 0.236701 | -0.24276 | 4.020028 | -0.06039 | 0.951846 | NA |
| lncRNA:CR | 2.053389 | -0.89329 | 3.886943 | -0.22982 | 0.818233 | NA |
| lncRNA:CR | 0 | NA | NA | NA | NA | NA |
| lncRNA:CR | 0 | NA | NA | NA | NA | NA |
| lncRNA:CR | 12.02044 | -0.34591 | 3.842436 | -0.09002 | 0.928269 | NA |
| asRNA:CR4 | 0 | NA | NA | NA | NA | NA |
| asRNA:CR4 | 0 | NA | NA | NA | NA | NA |
| lncRNA:CR | 0 | NA | NA | NA | NA | NA |
| lncRNA:CR | 0 | NA | NA | NA | NA | NA |
| asRNA:CR4 | 0 | NA | NA | NA | NA | NA |
| lncRNA:CR | 0 | NA | NA | NA | NA | NA |
| lncRNA:CR | 0 | NA | NA | NA | NA | NA |
| lncRNA:CR | 0 | NA | NA | NA | NA | NA |
| lncRNA:CR | 26.28346 | -1.91792 | 3.952389 | -0.48526 | 0.627494 | NA |
| Cenp-C | 0.178053 | -1.16467 | 4.075067 | -0.2858 | 0.775028 | NA |
| asRNA:CR4 | 0 | NA | NA | NA | NA | NA |
| lncRNA:CR | 0 | NA | NA | NA | NA | NA |
| lncRNA:CR | 1.311343 | -1.38229 | 3.935643 | -0.35122 | 0.72542 | NA |
| lncRNA:CR | 0 | NA | NA | NA | NA | NA |
| lncRNA:CR | 12.66861 | 2.946279 | 3.925581 | 0.750533 | 0.452933 | NA |
| lncRNA:CR | 0 | NA | NA | NA | NA | NA |
| lncRNA:CR | 0.264155 | -1.54099 | 4.048976 | -0.38059 | 0.703509 | NA |
| asRNA:CR4 | 0 | NA | NA | NA | NA | NA |
| CR45385 | 0.178053 | -1.16467 | 4.075067 | -0.2858 | 0.775028 | NA |
| lncRNA:CR | 17.06609 | 1.74264 | 3.95393 | 0.440736 | 0.659404 | NA |
| lncRNA:CR | 0 | NA | NA | NA | NA | NA |
| lncRNA:CR | 0 | NA | NA | NA | NA | NA |
| lncRNA:CR | 0 | NA | NA | NA | NA | NA |
| lncRNA:CR | 0 | NA | NA | NA | NA | NA |
| CR45391 | 0 | NA | NA | NA | NA | NA |
| lncRNA:CR | 0.1047 | 0.613261 | 4.062799 | 0.150945 | 0.880019 | NA |
| lncRNA:CR | 0 | NA | NA | NA | NA | NA |
| lncRNA:CR | 0 | NA | NA | NA | NA | NA |
| lncRNA:CR | 0 | NA | NA | NA | NA | NA |

| | | | | | | |
|---|---|---|---|---|---|---|
| asRNA:CR4 | 0.418493 | 1.313669 | 4.019283 | 0.326842 | 0.743788 | NA |
| asRNA:CR4 | 0 | NA | NA | NA | NA | NA |
| lncRNA:CR | 0 | NA | NA | NA | NA | NA |
| lncRNA:CR | 0.104623 | 0.639075 | 4.067674 | 0.157111 | 0.875158 | NA |
| lncRNA:CR | 0 | NA | NA | NA | NA | NA |
| lncRNA:CR | 0 | NA | NA | NA | NA | NA |
| lncRNA:CR | 0 | NA | NA | NA | NA | NA |
| lncRNA:CR | 31.03826 | -1.91871 | 3.952163 | -0.48548 | 0.627334 | NA |
| lncRNA:CR | 4.470307 | -2.20441 | 3.970887 | -0.55514 | 0.578797 | NA |
| lncRNA:CR | 0 | NA | NA | NA | NA | NA |
| lncRNA:CR | 0.1047 | 0.613261 | 4.062799 | 0.150945 | 0.880019 | NA |
| lncRNA:CR | 0 | NA | NA | NA | NA | NA |
| lncRNA:CR | 0.178053 | -1.16467 | 4.075067 | -0.2858 | 0.775028 | NA |
| lncRNA:CR | 0 | NA | NA | NA | NA | NA |
| lncRNA:CR | 20.91648 | -5.61529 | 3.897424 | -1.44077 | 0.14965 | NA |
| lncRNA:CR | 0.178053 | -1.16467 | 4.075067 | -0.2858 | 0.775028 | NA |
| CG45413 | 0 | NA | NA | NA | NA | NA |
| lncRNA:CR | 5.811418 | -1.90022 | 3.957496 | -0.48016 | 0.631116 | NA |
| lncRNA:CR | 0 | NA | NA | NA | NA | NA |
| lncRNA:CR | 0 | NA | NA | NA | NA | NA |
| lncRNA:CR | 31.54074 | -1.71799 | 3.812837 | -0.45058 | 0.652293 | NA |
| lncRNA:CR | 19.78865 | -8.32814 | 3.846501 | -2.16512 | 0.030378 | NA |
| lncRNA:CR | 0 | NA | NA | NA | NA | NA |
| lncRNA:CR | 0 | NA | NA | NA | NA | NA |
| lncRNA:CR | 0 | NA | NA | NA | NA | NA |
| lncRNA:CR | 0 | NA | NA | NA | NA | NA |
| lncRNA:CR | 0 | NA | NA | NA | NA | NA |
| lncRNA:CR | 0 | NA | NA | NA | NA | NA |
| lncRNA:CR | 0 | NA | NA | NA | NA | NA |
| lncRNA:CR | 0 | NA | NA | NA | NA | NA |
| CG45428 | 0 | NA | NA | NA | NA | NA |
| lncRNA:CR | 0 | NA | NA | NA | NA | NA |
| lncRNA:CR | 1.981645 | 1.940655 | 3.957663 | 0.490354 | 0.623884 | NA |
| lncRNA:CR | 0.341324 | 0.112818 | 4.000379 | 0.028202 | 0.977501 | NA |
| lncRNA:CR | 12.24092 | 1.528761 | 3.953831 | 0.386653 | 0.699013 | NA |
| lncRNA:CR | 0 | NA | NA | NA | NA | NA |
| lncRNA:CR | 0 | NA | NA | NA | NA | NA |
| lncRNA:CR | 27.4313 | 2.96645 | 3.921008 | 0.756553 | 0.449318 | NA |
| lncRNA:CR | 0 | NA | NA | NA | NA | NA |
| lncRNA:CR | 0 | NA | NA | NA | NA | NA |
| lncRNA:CR | 0 | NA | NA | NA | NA | NA |
| lncRNA:CR | 0 | NA | NA | NA | NA | NA |
| lncRNA:CR | 0 | NA | NA | NA | NA | NA |
| lncRNA:CR | 0 | NA | NA | NA | NA | NA |
| lncRNA:CR | 0 | NA | NA | NA | NA | NA |
| lncRNA:CR | 2.097839 | 1.11873 | 3.972242 | 0.281637 | 0.778222 | NA |
| lncRNA:CR | 0 | NA | NA | NA | NA | NA |
| asRNA:CR4 | 0 | NA | NA | NA | NA | NA |
| lncRNA:CR | 0 | NA | NA | NA | NA | NA |
| lncRNA:CR | 0 | NA | NA | NA | NA | NA |

| | | | | | | |
|---|---|---|---|---|---|---|
| lncRNA:CR | 0 | NA | NA | NA | NA | NA |
| lncRNA:CR | 0 | NA | NA | NA | NA | NA |
| lncRNA:CR | 0.178053 | -1.16467 | 4.075067 | -0.2858 | 0.775028 | NA |
| lncRNA:CR | 0 | NA | NA | NA | NA | NA |
| lncRNA:CR | 0.178053 | -1.16467 | 4.075067 | -0.2858 | 0.775028 | NA |
| lncRNA:CR | 0 | NA | NA | NA | NA | NA |
| lncRNA:CR | 0 | NA | NA | NA | NA | NA |
| lncRNA:CR | 0 | NA | NA | NA | NA | NA |
| lncRNA:CR | 0 | NA | NA | NA | NA | NA |
| lncRNA:CR | 0 | NA | NA | NA | NA | NA |
| CR45458 | 4.467006 | 0.772124 | 3.77768 | 0.204391 | 0.838048 | NA |
| lncRNA:CR | 43.67991 | 0.357798 | 3.834646 | 0.093307 | 0.92566 | NA |
| lncRNA:CR | 0 | NA | NA | NA | NA | NA |
| lncRNA:CR | 20.20579 | -2.26351 | 3.955621 | -0.57223 | 0.567168 | NA |
| lncRNA:CR | 25.61656 | 3.064656 | 3.899904 | 0.785829 | 0.431968 | NA |
| lncRNA:CR | 0 | NA | NA | NA | NA | NA |
| asRNA:CR4 | 0 | NA | NA | NA | NA | NA |
| asRNA:CR4 | 5.758496 | 1.720238 | 3.959653 | 0.434441 | 0.663968 | NA |
| asRNA:CR4 | 0 | NA | NA | NA | NA | NA |
| asRNA:CR4 | 15.87189 | 0.102102 | 3.878599 | 0.026324 | 0.978999 | NA |
| CR45470 | 4.173233 | -1.2228 | 3.859453 | -0.31683 | 0.75137 | NA |
| lncRNA:CR | 4.117185 | -4.39498 | 3.860397 | -1.13848 | 0.254921 | NA |
| lncRNA:CR | 4.027089 | 2.292019 | 3.823753 | 0.599416 | 0.548895 | NA |
| lncRNA:CR | 0 | NA | NA | NA | NA | NA |
| lncRNA:CR | 0 | NA | NA | NA | NA | NA |
| lncRNA:CR | 0 | NA | NA | NA | NA | NA |
| asRNA:CR4 | 18.42939 | 1.302761 | 3.855833 | 0.337867 | 0.735463 | NA |
| lncRNA:CR | 4.905149 | -4.66714 | 3.94586 | -1.18279 | 0.23689 | NA |
| asRNA:CR4 | 0 | NA | NA | NA | NA | NA |
| lncRNA:CR | 0.178053 | -1.16467 | 4.075067 | -0.2858 | 0.775028 | NA |
| lncRNA:CR | 0 | NA | NA | NA | NA | NA |
| lncRNA:CR | 0 | NA | NA | NA | NA | NA |
| asRNA:CR4 | 10.7016 | -4.57918 | 3.854492 | -1.18801 | 0.234829 | NA |
| CG45486 | 0 | NA | NA | NA | NA | NA |
| CG45487 | 0 | NA | NA | NA | NA | NA |
| CG45488 | 0 | NA | NA | NA | NA | NA |
| CG45489 | 0 | NA | NA | NA | NA | NA |
| CG45490 | 0 | NA | NA | NA | NA | NA |
| CG45491 | 0 | NA | NA | NA | NA | NA |
| CG45492 | 0 | NA | NA | NA | NA | NA |
| CG45493 | 0 | NA | NA | NA | NA | NA |
| CG45494 | 0 | NA | NA | NA | NA | NA |
| CR45495 | 0 | NA | NA | NA | NA | NA |
| CR45496 | 0 | NA | NA | NA | NA | NA |
| CR45497 | 0 | NA | NA | NA | NA | NA |
| CR45498 | 0 | NA | NA | NA | NA | NA |
| CR45499 | 0 | NA | NA | NA | NA | NA |
| lncRNA:CR | 0 | NA | NA | NA | NA | NA |
| asRNA:CR4 | 0 | NA | NA | NA | NA | NA |
| lncRNA:CR | 1.165466 | 1.076041 | 3.987865 | 0.269829 | 0.787292 | NA |

| | | | | | | |
|---|---|---|---|---|---|---|
| lncRNA:CR | 0 | NA | NA | NA | NA | NA |
| lncRNA:CR | 0 | NA | NA | NA | NA | NA |
| lncRNA:CR | 0 | NA | NA | NA | NA | NA |
| lncRNA:CR | 0 | NA | NA | NA | NA | NA |
| lncRNA:CR | 0.178053 | -1.16467 | 4.075067 | -0.2858 | 0.775028 | NA |
| lncRNA:CR | 0 | NA | NA | NA | NA | NA |
| lncRNA:CR | 0 | NA | NA | NA | NA | NA |
| lncRNA:CR | 0 | NA | NA | NA | NA | NA |
| lncRNA:CR | 0 | NA | NA | NA | NA | NA |
| lncRNA:CR | 0 | NA | NA | NA | NA | NA |
| lncRNA:CR | 0 | NA | NA | NA | NA | NA |
| lncRNA:CR | 0 | NA | NA | NA | NA | NA |
| lncRNA:CR | 14.1377 | 0.321112 | 3.845108 | 0.083512 | 0.933445 | NA |
| lncRNA:CR | 0 | NA | NA | NA | NA | NA |
| asRNA:CR4 | 0.104623 | 0.639075 | 4.067674 | 0.157111 | 0.875158 | NA |
| lncRNA:CR | 0 | NA | NA | NA | NA | NA |
| asRNA:CR4 | 22.7084 | -3.5958 | 3.920039 | -0.91729 | 0.358993 | NA |
| lncRNA:CR | 0 | NA | NA | NA | NA | NA |
| lncRNA:CR | 0 | NA | NA | NA | NA | NA |
| lncRNA:CR | 0 | NA | NA | NA | NA | NA |
| lncRNA:CR | 0 | NA | NA | NA | NA | NA |
| asRNA:CR4 | 0 | NA | NA | NA | NA | NA |
| lncRNA:CR | 0.178812 | -1.08808 | 4.061735 | -0.26789 | 0.788788 | NA |
| asRNA:CR4 | 0.466186 | 0.959471 | 4.033986 | 0.237847 | 0.812 | NA |
| lncRNA:CR | 0 | NA | NA | NA | NA | NA |
| lncRNA:CR | 0 | NA | NA | NA | NA | NA |
| lncRNA:CR | 0.178053 | -1.16467 | 4.075067 | -0.2858 | 0.775028 | NA |
| lncRNA:CR | 0 | NA | NA | NA | NA | NA |
| lncRNA:CR | 0 | NA | NA | NA | NA | NA |
| lncRNA:CR | 0.357625 | -1.47707 | 4.044794 | -0.36518 | 0.714979 | NA |
| asRNA:CR4 | 0 | NA | NA | NA | NA | NA |
| lncRNA:CR | 0.356107 | -1.47099 | 4.037764 | -0.36431 | 0.715628 | NA |
| lncRNA:CR | 0 | NA | NA | NA | NA | NA |
| lncRNA:CR | 0 | NA | NA | NA | NA | NA |
| lncRNA:CR | 0 | NA | NA | NA | NA | NA |
| lncRNA:CR | 0 | NA | NA | NA | NA | NA |
| CG45544 | 0 | NA | NA | NA | NA | NA |
| CG45545 | 0 | NA | NA | NA | NA | NA |
| CG45546 | 9.773749 | -1.90931 | 3.95486 | -0.48278 | 0.629255 | NA |
| lncRNA:CR | 20.86827 | -1.91658 | 3.952771 | -0.48487 | 0.627769 | NA |
| CR45548 | 0 | NA | NA | NA | NA | NA |
| asRNA:CR4 | 21.27488 | 0.66979 | 3.951899 | 0.169486 | 0.865415 | NA |
| lncRNA:CR | 0 | NA | NA | NA | NA | NA |
| lncRNA:CR | 3.871196 | -3.05496 | 3.926716 | -0.77799 | 0.436573 | NA |
| lncRNA:CR | 0.1047 | 0.613261 | 4.062799 | 0.150945 | 0.880019 | NA |
| lncRNA:CR | 0 | NA | NA | NA | NA | NA |
| lncRNA:CR | 0 | NA | NA | NA | NA | NA |
| lncRNA:CR | 0 | NA | NA | NA | NA | NA |
| lncRNA:CR | 8.674033 | 2.146231 | 3.887461 | 0.552091 | 0.580886 | NA |
| lncRNA:CR | 0 | NA | NA | NA | NA | NA |

| | | | | | | |
|---|---|---|---|---|---|---|
| lncRNA:CR | 13.2445 | -6.27204 | 3.895023 | -1.61027 | 0.107339 | NA |
| lncRNA:CR | 0 | NA | NA | NA | NA | NA |
| lncRNA:CR | 0 | NA | NA | NA | NA | NA |
| lncRNA:CR | 0 | NA | NA | NA | NA | NA |
| lncRNA:CR | 0 | NA | NA | NA | NA | NA |
| lncRNA:CR | 0 | NA | NA | NA | NA | NA |
| lncRNA:CR | 0 | NA | NA | NA | NA | NA |
| lncRNA:CR | 17.75203 | -0.87337 | 3.952279 | -0.22098 | 0.825109 | NA |
| lncRNA:CR | 0 | NA | NA | NA | NA | NA |
| lncRNA:CR | 0 | NA | NA | NA | NA | NA |
| lncRNA:CR | 0 | NA | NA | NA | NA | NA |
| lncRNA:CR | 0 | NA | NA | NA | NA | NA |
| lncRNA:CR | 0 | NA | NA | NA | NA | NA |
| lncRNA:CR | 0 | NA | NA | NA | NA | NA |
| lncRNA:CR | 0.53416 | -1.53481 | 4.012395 | -0.38252 | 0.702078 | NA |
| lncRNA:CR | 0 | NA | NA | NA | NA | NA |
| lncRNA:CR | 0 | NA | NA | NA | NA | NA |
| lncRNA:CR | 0.282677 | -0.09599 | 4.057934 | -0.02366 | 0.981127 | NA |
| lncRNA:CR | 0 | NA | NA | NA | NA | NA |
| lncRNA:CR | 0.1047 | 0.613261 | 4.062799 | 0.150945 | 0.880019 | NA |
| lncRNA:CR | 26.46968 | 1.534479 | 3.952266 | 0.388253 | 0.697829 | NA |
| lncRNA:CR | 0.178053 | -1.16467 | 4.075067 | -0.2858 | 0.775028 | NA |
| lncRNA:CR | 0 | NA | NA | NA | NA | NA |
| lncRNA:CR | 0 | NA | NA | NA | NA | NA |
| lncRNA:CR | 0 | NA | NA | NA | NA | NA |
| lncRNA:CR | 0 | NA | NA | NA | NA | NA |
| lncRNA:CR | 0.178053 | -1.16467 | 4.075067 | -0.2858 | 0.775028 | NA |
| lncRNA:CR | 0 | NA | NA | NA | NA | NA |
| lncRNA:CR | 0 | NA | NA | NA | NA | NA |
| lncRNA:CR | 0 | NA | NA | NA | NA | NA |
| lncRNA:CR | 1.072874 | -2.0339 | 4.019685 | -0.50598 | 0.612867 | NA |
| lncRNA:CR | 0 | NA | NA | NA | NA | NA |
| lncRNA:CR | 0 | NA | NA | NA | NA | NA |
| lncRNA:CR | 0 | NA | NA | NA | NA | NA |
| lncRNA:CR | 0 | NA | NA | NA | NA | NA |
| lncRNA:CR | 0 | NA | NA | NA | NA | NA |
| lncRNA:CR | 43.37157 | -0.27373 | 3.950641 | -0.06929 | 0.94476 | NA |
| lncRNA:CR | 0 | NA | NA | NA | NA | NA |
| lncRNA:CR | 0 | NA | NA | NA | NA | NA |
| lncRNA:CR | 0 | NA | NA | NA | NA | NA |
| asRNA:CR4 | 0 | NA | NA | NA | NA | NA |
| lncRNA:CR | 5.831849 | 1.457586 | 3.95798 | 0.368265 | 0.712676 | NA |
| CG45603 | 3.350398 | 1.697551 | 3.965576 | 0.428072 | 0.668599 | NA |
| asRNA:CR4 | 0.337793 | 1.451235 | 4.039844 | 0.35923 | 0.719423 | NA |
| lncRNA:CR | 0 | NA | NA | NA | NA | NA |
| lncRNA:CR | 0 | NA | NA | NA | NA | NA |
| lncRNA:CR | 0 | NA | NA | NA | NA | NA |
| lncRNA:CR | 0 | NA | NA | NA | NA | NA |
| lncRNA:CR | 0 | NA | NA | NA | NA | NA |
| lncRNA:CR | 0 | NA | NA | NA | NA | NA |

| | | | | | | |
|---|---|---|---|---|---|---|
| asRNA:CR4 | 16.01909 | 1.741867 | 3.954124 | 0.440519 | 0.659561 | NA |
| asRNA:CR4 | 6.281996 | 1.722971 | 3.958948 | 0.435209 | 0.66341 | NA |
| lncRNA:CR | 0 | NA | NA | NA | NA | NA |
| lncRNA:CR | 9.323729 | 1.165404 | 3.955911 | 0.294598 | 0.768301 | NA |
| lncRNA:CR | 2.113243 | -1.8635 | 3.968379 | -0.46959 | 0.63865 | NA |
| lncRNA:CR | 3.993031 | 0.347252 | 3.851261 | 0.090166 | 0.928155 | NA |
| lncRNA:CR | 8.452972 | -1.9072 | 3.955468 | -0.48217 | 0.629686 | NA |
| lncRNA:CR | 0 | NA | NA | NA | NA | NA |
| lncRNA:CR | 0 | NA | NA | NA | NA | NA |
| asRNA:CR4 | 0 | NA | NA | NA | NA | NA |
| lncRNA:CR | 0 | NA | NA | NA | NA | NA |
| asRNA:CR4 | 0 | NA | NA | NA | NA | NA |
| lncRNA:CR | 0 | NA | NA | NA | NA | NA |
| lncRNA:CR | 10.25653 | -2.77849 | 3.836392 | -0.72425 | 0.468914 | NA |
| lncRNA:CR | 2.512798 | 1.684724 | 3.970151 | 0.424348 | 0.671312 | NA |
| lncRNA:CR | 0 | NA | NA | NA | NA | NA |
| lncRNA:CR | 4.18493 | 1.509067 | 3.95929 | 0.381146 | 0.703095 | NA |
| asRNA:CR4 | 0 | NA | NA | NA | NA | NA |
| lncRNA:CR | 0 | NA | NA | NA | NA | NA |
| lncRNA:CR | 0 | NA | NA | NA | NA | NA |
| lncRNA:CR | 0 | NA | NA | NA | NA | NA |
| lncRNA:CR | 0.3141 | 1.240997 | 4.03841 | 0.307299 | 0.758616 | NA |
| lncRNA:CR | 0 | NA | NA | NA | NA | NA |
| lncRNA:CR | 0.62774 | 1.374224 | 3.999506 | 0.343599 | 0.731148 | NA |
| lncRNA:CR | 0 | NA | NA | NA | NA | NA |
| lncRNA:CR | 0 | NA | NA | NA | NA | NA |
| lncRNA:CR | 0 | NA | NA | NA | NA | NA |
| lncRNA:CR | 0 | NA | NA | NA | NA | NA |
| lncRNA:CR | 0 | NA | NA | NA | NA | NA |
| lncRNA:CR | 0 | NA | NA | NA | NA | NA |
| lncRNA:CR | 0 | NA | NA | NA | NA | NA |
| lncRNA:CR | 4.4988 | 1.511101 | 3.958721 | 0.381714 | 0.702673 | NA |
| lncRNA:CR | 0 | NA | NA | NA | NA | NA |
| lncRNA:CR | 0.178053 | -1.16467 | 4.075067 | -0.2858 | 0.775028 | NA |
| lncRNA:CR | 0.365171 | -0.43584 | 4.056359 | -0.10745 | 0.914434 | NA |
| lncRNA:CR | 0 | NA | NA | NA | NA | NA |
| asRNA:CR4 | 0 | NA | NA | NA | NA | NA |
| lncRNA:CR | 0 | NA | NA | NA | NA | NA |
| lncRNA:CR | 43.6302 | -2.27354 | 3.953114 | -0.57513 | 0.565206 | NA |
| lncRNA:CR | 0 | NA | NA | NA | NA | NA |
| lncRNA:CR | 0 | NA | NA | NA | NA | NA |
| lncRNA:CR | 0 | NA | NA | NA | NA | NA |
| lncRNA:CR | 0 | NA | NA | NA | NA | NA |
| lncRNA:CR | 0 | NA | NA | NA | NA | NA |
| lncRNA:CR | 0 | NA | NA | NA | NA | NA |
| lncRNA:CR | 0 | NA | NA | NA | NA | NA |
| lncRNA:CR | 0 | NA | NA | NA | NA | NA |
| lncRNA:CR | 37.04619 | 6.605198 | 3.827551 | 1.725698 | 0.084402 | NA |
| lncRNA:CR | 0 | NA | NA | NA | NA | NA |
| lncRNA:CR | 17.58959 | 1.742992 | 3.953841 | 0.440835 | 0.659332 | NA |

| | | | | | | |
|---|---|---|---|---|---|---|
| asRNA:CR4 | 0 | NA | NA | NA | NA | NA |
| lncRNA:CR | 0 | NA | NA | NA | NA | NA |
| lncRNA:CR | 0 | NA | NA | NA | NA | NA |
| lncRNA:CR | 0 | NA | NA | NA | NA | NA |
| lncRNA:CR | 0 | NA | NA | NA | NA | NA |
| lncRNA:CR | 1.361099 | 1.628413 | 3.98439 | 0.408698 | 0.682761 | NA |
| lncRNA:CR | 2.451623 | 0.000923 | 3.913399 | 0.000236 | 0.999812 | NA |
| lncRNA:CR | 0 | NA | NA | NA | NA | NA |
| lncRNA:CR | 0 | NA | NA | NA | NA | NA |
| lncRNA:CR | 37.90137 | 1.749223 | 3.952277 | 0.442586 | 0.658065 | NA |
| lncRNA:CR | 3.010332 | -1.56487 | 3.960938 | -0.39507 | 0.692788 | NA |
| lncRNA:CR | 0 | NA | NA | NA | NA | NA |
| lncRNA:CR | 0 | NA | NA | NA | NA | NA |
| lncRNA:CR | 0 | NA | NA | NA | NA | NA |
| lncRNA:CR | 0 | NA | NA | NA | NA | NA |
| lncRNA:CR | 30.60546 | 1.052815 | 3.952339 | 0.266378 | 0.789948 | NA |
| asRNA:CR4 | 0 | NA | NA | NA | NA | NA |
| lncRNA:CR | 3.036298 | 1.692169 | 3.966999 | 0.426562 | 0.669699 | NA |
| lncRNA:CR | 1.602479 | -1.64094 | 3.97352 | -0.41297 | 0.67963 | NA |
| asRNA:CR4 | 0 | NA | NA | NA | NA | NA |
| lncRNA:CR | 0.178053 | -1.16467 | 4.075067 | -0.2858 | 0.775028 | NA |
| lncRNA:CR | 0 | NA | NA | NA | NA | NA |
| lncRNA:CR | 0 | NA | NA | NA | NA | NA |
| CG45690 | 0 | NA | NA | NA | NA | NA |
| CG45691 | 0.1047 | 0.613261 | 4.062799 | 0.150945 | 0.880019 | NA |
| lncRNA:CR | 0 | NA | NA | NA | NA | NA |
| lncRNA:CR | 1.133183 | -2.25669 | 3.937492 | -0.57313 | 0.566557 | NA |
| lncRNA:CR | 0 | NA | NA | NA | NA | NA |
| lncRNA:CR | 0 | NA | NA | NA | NA | NA |
| lncRNA:CR | 0 | NA | NA | NA | NA | NA |
| lncRNA:CR | 0.534919 | -2.31746 | 4.015737 | -0.5771 | 0.563875 | NA |
| lncRNA:CR | 0 | NA | NA | NA | NA | NA |
| lncRNA:CR | 0.178812 | -1.08808 | 4.061735 | -0.26789 | 0.788788 | NA |
| lncRNA:CR | 0 | NA | NA | NA | NA | NA |
| asRNA:CR4 | 0 | NA | NA | NA | NA | NA |
| lncRNA:CR | 0.1047 | 0.613261 | 4.062799 | 0.150945 | 0.880019 | NA |
| lncRNA:CR | 0 | NA | NA | NA | NA | NA |
| lncRNA:CR | 0.356107 | -1.47099 | 4.037764 | -0.36431 | 0.715628 | NA |
| lncRNA:CR | 0 | NA | NA | NA | NA | NA |
| lncRNA:CR | 0.104623 | 0.639075 | 4.067674 | 0.157111 | 0.875158 | NA |
| lncRNA:CR | 13.07791 | 1.529441 | 3.953645 | 0.386843 | 0.698872 | NA |
| lncRNA:CR | 0 | NA | NA | NA | NA | NA |
| lncRNA:CR | 0 | NA | NA | NA | NA | NA |
| asRNA:CR4 | 0.62774 | 1.374224 | 3.999506 | 0.343599 | 0.731148 | NA |
| asRNA:CR4 | 0 | NA | NA | NA | NA | NA |
| asRNA:CR4 | 0 | NA | NA | NA | NA | NA |
| asRNA:CR4 | 0 | NA | NA | NA | NA | NA |
| CR45716 | 0 | NA | NA | NA | NA | NA |
| lncRNA:CR | 0 | NA | NA | NA | NA | NA |
| lncRNA:CR | 0 | NA | NA | NA | NA | NA |

| | | | | | | |
|---|---|---|---|---|---|---|
| lncRNA:CR | 0 | NA | NA | NA | NA | NA |
| lncRNA:CR | 0 | NA | NA | NA | NA | NA |
| lncRNA:CR | 0.178053 | -1.16467 | 4.075067 | -0.2858 | 0.775028 | NA |
| lncRNA:CR | 0 | NA | NA | NA | NA | NA |
| lncRNA:CR | 0 | NA | NA | NA | NA | NA |
| lncRNA:CR | 0.357625 | -1.47707 | 4.044794 | -0.36518 | 0.714979 | NA |
| lncRNA:CR | 0 | NA | NA | NA | NA | NA |
| lncRNA:CR | 0 | NA | NA | NA | NA | NA |
| lncRNA:CR | 0 | NA | NA | NA | NA | NA |
| lncRNA:CR | 0 | NA | NA | NA | NA | NA |
| lncRNA:CR | 0 | NA | NA | NA | NA | NA |
| lncRNA:CR | 0 | NA | NA | NA | NA | NA |
| lncRNA:CR | 0 | NA | NA | NA | NA | NA |
| lncRNA:CR | 6.177296 | 1.72246 | 3.95908 | 0.435066 | 0.663515 | NA |
| lncRNA:CR | 0 | NA | NA | NA | NA | NA |
| lncRNA:CR | 0 | NA | NA | NA | NA | NA |
| lncRNA:CR | 0.178053 | -1.16467 | 4.075067 | -0.2858 | 0.775028 | NA |
| lncRNA:CR | 0 | NA | NA | NA | NA | NA |
| asRNA:CR4 | 0 | NA | NA | NA | NA | NA |
| lncRNA:CR | 0 | NA | NA | NA | NA | NA |
| lncRNA:CR | 0 | NA | NA | NA | NA | NA |
| lncRNA:CR | 0 | NA | NA | NA | NA | NA |
| lncRNA:CR | 0 | NA | NA | NA | NA | NA |
| lncRNA:CR | 0.713731 | -2.6372 | 4.003426 | -0.65874 | 0.510065 | NA |
| lncRNA:CR | 0 | NA | NA | NA | NA | NA |
| lncRNA:CR | 0 | NA | NA | NA | NA | NA |
| lncRNA:CR | 0 | NA | NA | NA | NA | NA |
| lncRNA:CR | 0 | NA | NA | NA | NA | NA |
| lncRNA:CR | 0 | NA | NA | NA | NA | NA |
| lncRNA:CR | 0 | NA | NA | NA | NA | NA |
| lncRNA:CR | 0.5235 | 1.490714 | 4.02555 | 0.370313 | 0.711149 | NA |
| lncRNA:TS | 0.233093 | 0.692656 | 4.076246 | 0.169925 | 0.865069 | NA |
| lncRNA:CR | 0 | NA | NA | NA | NA | NA |
| lncRNA:CR | 0 | NA | NA | NA | NA | NA |
| lncRNA:CR | 0 | NA | NA | NA | NA | NA |
| lncRNA:CR | 0 | NA | NA | NA | NA | NA |
| lncRNA:CR | 0 | NA | NA | NA | NA | NA |
| lncRNA:CR | 0 | NA | NA | NA | NA | NA |
| lncRNA:CR | 0 | NA | NA | NA | NA | NA |
| Acp33A | 45.58164 | -1.70567 | 3.95175 | -0.43162 | 0.666015 | NA |
| p38c | 0 | NA | NA | NA | NA | NA |
| squ | 0.894061 | -2.00006 | 4.030294 | -0.49626 | 0.619714 | NA |
| JYalpha | 0.104623 | 0.639075 | 4.067674 | 0.157111 | 0.875158 | NA |
| mil | 0 | NA | NA | NA | NA | NA |
| Ubc87F | 0 | NA | NA | NA | NA | NA |
| lncRNA:CR | 0 | NA | NA | NA | NA | NA |
| lncRNA:CR | 48.79017 | 1.750442 | 3.951972 | 0.442929 | 0.657817 | NA |
| AOX1 | 26.87653 | 1.675648 | 3.95273 | 0.423922 | 0.671623 | NA |
| CG45764 | 0 | NA | NA | NA | NA | NA |
| CG45765 | 0 | NA | NA | NA | NA | NA |

| | | | | | | |
|---|---|---|---|---|---|---|
| CG45766 | 0 | NA | NA | NA | NA | NA |
| CG45770 | 0 | NA | NA | NA | NA | NA |
| CR45771 | 0 | NA | NA | NA | NA | NA |
| CR45772 | 0 | NA | NA | NA | NA | NA |
| CR45773 | 0 | NA | NA | NA | NA | NA |
| CR45775 | 0 | NA | NA | NA | NA | NA |
| CR45776 | 0 | NA | NA | NA | NA | NA |
| CR45777 | 0 | NA | NA | NA | NA | NA |
| asRNA:CR4 | 0 | NA | NA | NA | NA | NA |
| CR45779 | 0 | NA | NA | NA | NA | NA |
| CR45780 | 0 | NA | NA | NA | NA | NA |
| kl-3 | 0.310131 | -1.94616 | 4.045058 | -0.48112 | 0.630431 | NA |
| kl-5 | 0.209323 | 1.414133 | 4.032409 | 0.350692 | 0.725819 | NA |
| chp | 14.02979 | 1.740091 | 3.954574 | 0.44002 | 0.659923 | NA |
| lncRNA:CR | 0 | NA | NA | NA | NA | NA |
| asRNA:CR4 | 17.69135 | -2.63201 | 3.819493 | -0.6891 | 0.490761 | NA |
| lncRNA:CR | 0 | NA | NA | NA | NA | NA |
| lncRNA:CR | 0 | NA | NA | NA | NA | NA |
| lncRNA:CR | 0 | NA | NA | NA | NA | NA |
| lncRNA:CR | 0 | NA | NA | NA | NA | NA |
| Su(Ste):CR | 8.63859 | 2.129376 | 3.882221 | 0.548494 | 0.583353 | NA |
| Su(Ste):CR | 0.104623 | 0.639075 | 4.067674 | 0.157111 | 0.875158 | NA |
| Su(Ste):CR | 0.209247 | 1.042804 | 4.047837 | 0.25762 | 0.7967 | NA |
| asRNA:CR4 | 0 | NA | NA | NA | NA | NA |
| asRNA:CR4 | 4.751466 | -4.28836 | 3.925119 | -1.09254 | 0.274595 | NA |
| WDY | 0 | NA | NA | NA | NA | NA |
| asRNA:CR4 | 0 | NA | NA | NA | NA | NA |
| asRNA:CR4 | 0.178812 | -1.08808 | 4.061735 | -0.26789 | 0.788788 | NA |
| asRNA:CR4 | 40.83052 | 1.504801 | 3.951769 | 0.380792 | 0.703358 | NA |
| lncRNA:CR | 0 | NA | NA | NA | NA | NA |
| lncRNA:CR | 15.74203 | 2.861944 | 3.86825 | 0.739855 | 0.459388 | NA |
| asRNA:CR4 | 0 | NA | NA | NA | NA | NA |
| asRNA:CR4 | 0.357625 | -1.47707 | 4.044794 | -0.36518 | 0.714979 | NA |
| lncRNA:CR | 12.05347 | 5.69107 | 3.888842 | 1.463436 | 0.143348 | NA |
| lncRNA:CR | 46.03423 | 1.536597 | 3.95169 | 0.388846 | 0.69739 | NA |
| lncRNA:CR | 0 | NA | NA | NA | NA | NA |
| lncRNA:CR | 0 | NA | NA | NA | NA | NA |
| asRNA:CR4 | 0 | NA | NA | NA | NA | NA |
| asRNA:CR4 | 0 | NA | NA | NA | NA | NA |
| asRNA:CR4 | 0.357625 | -1.47707 | 4.044794 | -0.36518 | 0.714979 | NA |
| asRNA:CR4 | 11.75152 | -1.69829 | 3.954156 | -0.42949 | 0.667564 | NA |
| lncRNA:CR | 2.136639 | -1.65662 | 3.968112 | -0.41748 | 0.676326 | NA |
| lncRNA:CR | 0 | NA | NA | NA | NA | NA |
| asRNA:CR4 | 0.675586 | 2.168561 | 4.004381 | 0.541547 | 0.58813 | NA |
| lncRNA:CR | 6.759704 | 1.159951 | 3.957776 | 0.293082 | 0.76946 | NA |
| lncRNA:CR | 0.233093 | 0.692656 | 4.076246 | 0.169925 | 0.865069 | NA |
| lncRNA:CR | 0 | NA | NA | NA | NA | NA |
| asRNA:CR4 | 0 | NA | NA | NA | NA | NA |
| asRNA:CR4 | 0 | NA | NA | NA | NA | NA |
| asRNA:CR4 | 0 | NA | NA | NA | NA | NA |

| | | | | | | |
|---|---|---|---|---|---|---|
| asRNA:CR4 | 0.233093 | 0.692656 | 4.076246 | 0.169925 | 0.865069 | NA |
| asRNA:CR4 | 0 | NA | NA | NA | NA | NA |
| asRNA:CR4 | 26.36506 | 1.534459 | 3.952272 | 0.388247 | 0.697833 | NA |
| lncRNA:CR | 15.07537 | -0.74317 | 3.839833 | -0.19354 | 0.846534 | NA |
| asRNA:CR4 | 24.45125 | -4.6558 | 3.831429 | -1.21516 | 0.224305 | NA |
| PRY | 0 | NA | NA | NA | NA | NA |
| Mst77Y-4 | 0 | NA | NA | NA | NA | NA |
| Mst77Y-13 | 0 | NA | NA | NA | NA | NA |
| Mst77Y-7 | 0 | NA | NA | NA | NA | NA |
| Mst77Y-9 | 0 | NA | NA | NA | NA | NA |
| Mst77Y-8F | 0 | NA | NA | NA | NA | NA |
| asRNA:CR4 | 2.992556 | -1.1917 | 3.874831 | -0.30755 | 0.758426 | NA |
| 2SrRNA:CF | 0 | NA | NA | NA | NA | NA |
| 28SrRNA:C | 0 | NA | NA | NA | NA | NA |
| 18SrRNA:C | 0 | NA | NA | NA | NA | NA |
| 5.8SrRNA:( | 0 | NA | NA | NA | NA | NA |
| 2SrRNA:CF | 0 | NA | NA | NA | NA | NA |
| 18SrRNA:C | 0 | NA | NA | NA | NA | NA |
| 5.8SrRNA:( | 0 | NA | NA | NA | NA | NA |
| 2SrRNA:CF | 0 | NA | NA | NA | NA | NA |
| 28SrRNA:C | 0 | NA | NA | NA | NA | NA |
| pre-rRNA:( | 1.105046 | 0.352707 | 3.9248 | 0.089866 | 0.928394 | NA |
| 5.8SrRNA-I | 0 | NA | NA | NA | NA | NA |
| 2SrRNA-Ps | 0 | NA | NA | NA | NA | NA |
| 28SrRNA-F | 10.77055 | -0.35551 | 3.74939 | -0.09482 | 0.924459 | NA |
| 5.8SrRNA:( | 0 | NA | NA | NA | NA | NA |
| 28SrRNA-F | 33.79481 | -0.61642 | 3.732525 | -0.16515 | 0.868827 | NA |
| 5.8SrRNA-I | 2.765401 | 1.298228 | 3.824442 | 0.339456 | 0.734267 | NA |
| 5.8SrRNA-I | 1.882835 | 1.186085 | 3.870456 | 0.306446 | 0.759265 | NA |
| 2SrRNA-Ps | 0.822293 | -0.3043 | 3.943495 | -0.07717 | 0.938492 | NA |
| 28SrRNA-F | 30.24361 | -1.48273 | 3.741689 | -0.39627 | 0.691904 | NA |
| 18SrRNA-F | 0 | NA | NA | NA | NA | NA |
| 28SrRNA-F | 0 | NA | NA | NA | NA | NA |
| 5.8SrRNA-I | 6.317042 | -0.01047 | 3.760156 | -0.00278 | 0.997779 | NA |
| 2SrRNA:CF | 2.595543 | 0.932825 | 3.870466 | 0.241011 | 0.809547 | NA |
| asRNA:CR4 | 0 | NA | NA | NA | NA | NA |
| lncRNA:CR | 0 | NA | NA | NA | NA | NA |
| asRNA:CR4 | 0 | NA | NA | NA | NA | NA |
| asRNA:CR4 | 28.38777 | -3.16628 | 3.910246 | -0.80974 | 0.41809 | NA |
| asRNA:CR4 | 0 | NA | NA | NA | NA | NA |
| asRNA:CR4 | 0 | NA | NA | NA | NA | NA |
| asRNA:CR4 | 0 | NA | NA | NA | NA | NA |
| asRNA:CR4 | 0 | NA | NA | NA | NA | NA |
| asRNA:CR4 | 0.466186 | 0.959471 | 4.033986 | 0.237847 | 0.812 | NA |
| asRNA:CR4 | 14.77113 | -0.51583 | 3.827631 | -0.13477 | 0.892797 | NA |
| asRNA:CR4 | 0 | NA | NA | NA | NA | NA |
| lncRNA:CR | 0.209247 | 1.042804 | 4.047837 | 0.25762 | 0.7967 | NA |
| lncRNA:CR | 0.546909 | -1.52446 | 3.990381 | -0.38203 | 0.702437 | NA |
| lncRNA:CR | 5.392151 | -0.24879 | 3.939994 | -0.06315 | 0.949651 | NA |
| CR45879 | 0 | NA | NA | NA | NA | NA |

| | | | | | | |
|---|---|---|---|---|---|---|
| lncRNA:CR | 0 | NA | NA | NA | NA | NA |
| lncRNA:CR | 14.39647 | -1.91368 | 3.953602 | -0.48403 | 0.628361 | NA |
| asRNA:CR4 | 0 | NA | NA | NA | NA | NA |
| lncRNA:CR | 0 | NA | NA | NA | NA | NA |
| lncRNA:CR | 0 | NA | NA | NA | NA | NA |
| asRNA:CR4 | 0 | NA | NA | NA | NA | NA |
| asRNA:CR4 | 0 | NA | NA | NA | NA | NA |
| lncRNA:CR | 0 | NA | NA | NA | NA | NA |
| lncRNA:CR | 0 | NA | NA | NA | NA | NA |
| asRNA:CR4 | 0.712213 | -1.57119 | 3.998613 | -0.39293 | 0.694369 | NA |
| asRNA:CR4 | 0.104623 | 0.639075 | 4.067674 | 0.157111 | 0.875158 | NA |
| asRNA:CR4 | 0 | NA | NA | NA | NA | NA |
| asRNA:CR4 | 0 | NA | NA | NA | NA | NA |
| lncRNA:CR | 0 | NA | NA | NA | NA | NA |
| asRNA:CR4 | 0 | NA | NA | NA | NA | NA |
| asRNA:CR4 | 0.236701 | -0.24276 | 4.020028 | -0.06039 | 0.951846 | NA |
| asRNA:CR4 | 0 | NA | NA | NA | NA | NA |
| lncRNA:CR | 0 | NA | NA | NA | NA | NA |
| lncRNA:CR | 0 | NA | NA | NA | NA | NA |
| lncRNA:CR | 0 | NA | NA | NA | NA | NA |
| lncRNA:CR | 0 | NA | NA | NA | NA | NA |
| lncRNA:CR | 0 | NA | NA | NA | NA | NA |
| lncRNA:CR | 4.502097 | 1.711245 | 3.961986 | 0.431916 | 0.665803 | NA |
| lncRNA:CR | 0 | NA | NA | NA | NA | NA |
| lncRNA:CR | 0 | NA | NA | NA | NA | NA |
| lncRNA:CR | 0 | NA | NA | NA | NA | NA |
| lncRNA:CR | 7.896637 | -0.10602 | 3.843338 | -0.02759 | 0.977993 | NA |
| asRNA:CR4 | 23.49128 | -7.65283 | 3.910018 | -1.95724 | 0.05032 | NA |
| asRNA:CR4 | 19.44283 | -0.35691 | 3.940359 | -0.09058 | 0.927827 | NA |
| asRNA:CR4 | 0 | NA | NA | NA | NA | NA |
| asRNA:CR4 | 7.405948 | -4.33579 | 3.832055 | -1.13145 | 0.257865 | NA |
| lncRNA:CR | 0.178053 | -1.16467 | 4.075067 | -0.2858 | 0.775028 | NA |
| lncRNA:CR | 1.251686 | -2.06022 | 4.011654 | -0.51356 | 0.60756 | NA |
| lncRNA:CR | 0 | NA | NA | NA | NA | NA |
| lncRNA:CR | 0 | NA | NA | NA | NA | NA |
| lncRNA:CR | 0 | NA | NA | NA | NA | NA |
| lncRNA:CR | 0.1047 | 0.613261 | 4.062799 | 0.150945 | 0.880019 | NA |
| lncRNA:CR | 0 | NA | NA | NA | NA | NA |
| lncRNA:CR | 0 | NA | NA | NA | NA | NA |
| lncRNA:dn | 23.13868 | 1.745763 | 3.953144 | 0.441614 | 0.658769 | NA |
| asRNA:CR4 | 2.377398 | -1.86964 | 3.966533 | -0.47135 | 0.637388 | NA |
| asRNA:CR4 | 0 | NA | NA | NA | NA | NA |
| asRNA:CR4 | 0 | NA | NA | NA | NA | NA |
| asRNA:CR4 | 18.51832 | 1.532353 | 3.952846 | 0.387658 | 0.698269 | NA |
| asRNA:CR4 | 0 | NA | NA | NA | NA | NA |
| CCY | 0 | NA | NA | NA | NA | NA |
| CR45931 | 0 | NA | NA | NA | NA | NA |
| CR45932 | 0 | NA | NA | NA | NA | NA |
| CR45933 | 0 | NA | NA | NA | NA | NA |
| CR45934 | 0.104623 | 0.639075 | 4.067674 | 0.157111 | 0.875158 | NA |

| | | | | | | |
|---|---|---|---|---|---|---|
| lncRNA:CR | 0.236701 | -0.24276 | 4.020028 | -0.06039 | 0.951846 | NA |
| lncRNA:CR | 19.57889 | 1.744162 | 3.953546 | 0.441164 | 0.659094 | NA |
| lncRNA:CR | 0 | NA | NA | NA | NA | NA |
| lncRNA:CR | 34.21651 | 5.408294 | 3.867203 | 1.398503 | 0.161962 | NA |
| lncRNA:CR | 0 | NA | NA | NA | NA | NA |
| lncRNA:CR | 0.536437 | -1.71175 | 4.038042 | -0.42391 | 0.671635 | NA |
| lncRNA:CR | 60.09046 | 0.108146 | 3.877936 | 0.027888 | 0.977752 | NA |
| lncRNA:CR | 22.58528 | -1.91707 | 3.95263 | -0.48501 | 0.627668 | NA |
| asRNA:CR4 | 0 | NA | NA | NA | NA | NA |
| asRNA:CR4 | 2.089576 | -1.64443 | 3.854892 | -0.42658 | 0.669683 | NA |
| lncRNA:CR | 6.242746 | -5.91587 | 3.917721 | -1.51003 | 0.131037 | NA |
| CR45946 | 0 | NA | NA | NA | NA | NA |
| CR45947 | 0 | NA | NA | NA | NA | NA |
| CR45948 | 0 | NA | NA | NA | NA | NA |
| lncRNA:CR | 1.150933 | 2.556099 | 3.971366 | 0.643632 | 0.519814 | NA |
| lncRNA:CR | 0 | NA | NA | NA | NA | NA |
| asRNA:CR4 | 3.689025 | 2.035225 | 3.827627 | 0.53172 | 0.59492 | NA |
| asRNA:CR4 | 0 | NA | NA | NA | NA | NA |
| lncRNA:CR | 14.55648 | 2.381884 | 3.924069 | 0.606993 | 0.543855 | NA |
| lncRNA:CR | 0 | NA | NA | NA | NA | NA |
| asRNA:CR4 | 0 | NA | NA | NA | NA | NA |
| lncRNA:CR | 0 | NA | NA | NA | NA | NA |
| lncRNA:CR | 0 | NA | NA | NA | NA | NA |
| asRNA:CR4 | 0.1047 | 0.613261 | 4.062799 | 0.150945 | 0.880019 | NA |
| lncRNA:CR | 0.1047 | 0.613261 | 4.062799 | 0.150945 | 0.880019 | NA |
| lncRNA:CR | 0 | NA | NA | NA | NA | NA |
| lncRNA:CR | 22.26746 | 4.480072 | 3.881716 | 1.154147 | 0.24844 | NA |
| lncRNA:CR | 0.1047 | 0.613261 | 4.062799 | 0.150945 | 0.880019 | NA |
| lncRNA:CR | 0 | NA | NA | NA | NA | NA |
| lncRNA:CR | 0 | NA | NA | NA | NA | NA |
| asRNA:CR4 | 0 | NA | NA | NA | NA | NA |
| asRNA:CR4 | 4.273279 | -1.68149 | 3.959707 | -0.42465 | 0.671092 | NA |
| asRNA:CR4 | 0 | NA | NA | NA | NA | NA |
| lncRNA:CR | 3.406565 | -1.59698 | 3.959971 | -0.40328 | 0.686741 | NA |
| lncRNA:CR | 8.375918 | 2.928025 | 3.929792 | 0.745084 | 0.456221 | NA |
| asRNA:CR4 | 0 | NA | NA | NA | NA | NA |
| lncRNA:CR | 0 | NA | NA | NA | NA | NA |
| lncRNA:CR | 0 | NA | NA | NA | NA | NA |
| lncRNA:CR | 0 | NA | NA | NA | NA | NA |
| lncRNA:CR | 0.233093 | 0.692656 | 4.076246 | 0.169925 | 0.865069 | NA |
| lncRNA:CR | 0 | NA | NA | NA | NA | NA |
| lncRNA:CR | 0 | NA | NA | NA | NA | NA |
| lncRNA:CR | 61.3842 | -0.37547 | 3.848515 | -0.09756 | 0.92228 | NA |
| lncRNA:CR | 0 | NA | NA | NA | NA | NA |
| lncRNA:CR | 0 | NA | NA | NA | NA | NA |
| lncRNA:CR | 0 | NA | NA | NA | NA | NA |
| pre-mod(n | 0 | NA | NA | NA | NA | NA |
| pre-mod(n | 0 | NA | NA | NA | NA | NA |
| pre-mod(n | 0 | NA | NA | NA | NA | NA |
| pre-mod(n | 0 | NA | NA | NA | NA | NA |

| | | | | | | |
|---|---|---|---|---|---|---|
| pre-mod(n | 0 | NA | NA | NA | NA | NA |
| lncRNA:CR | 0 | NA | NA | NA | NA | NA |
| asRNA:CR4 | 0 | NA | NA | NA | NA | NA |
| lncRNA:CR | 18.32249 | 1.743452 | 3.953725 | 0.440964 | 0.659239 | NA |
| lncRNA:CR | 0.178053 | -1.16467 | 4.075067 | -0.2858 | 0.775028 | NA |
| lncRNA:CR | 0 | NA | NA | NA | NA | NA |
| asRNA:CR4 | 0 | NA | NA | NA | NA | NA |
| lncRNA:CR | 0 | NA | NA | NA | NA | NA |
| lncRNA:CR | 0 | NA | NA | NA | NA | NA |
| lncRNA:CR | 0 | NA | NA | NA | NA | NA |
| lncRNA:CR | 15.79724 | -2.14689 | 3.83328 | -0.56007 | 0.575435 | NA |
| lncRNA:CR | 24.17597 | 1.976401 | 3.849234 | 0.513453 | 0.607634 | NA |
| asRNA:CR4 | 0.31089 | -1.9003 | 4.033427 | -0.47114 | 0.637542 | NA |
| lncRNA:CR | 7.300184 | -1.69234 | 3.956109 | -0.42778 | 0.668812 | NA |
| lncRNA:CR | 5.027078 | -3.91478 | 3.923105 | -0.99788 | 0.318338 | NA |
| lncRNA:CR | 0 | NA | NA | NA | NA | NA |
| lncRNA:CR | 10.15877 | 1.720737 | 3.854791 | 0.446389 | 0.655316 | NA |
| lncRNA:CR | 0 | NA | NA | NA | NA | NA |
| asRNA:CR4 | 0 | NA | NA | NA | NA | NA |
| lncRNA:CR | 4.397397 | 1.710278 | 3.962238 | 0.431644 | 0.666 | NA |
| lncRNA:CR | 0 | NA | NA | NA | NA | NA |
| lncRNA:CR | 1.788123 | -2.11324 | 3.996051 | -0.52883 | 0.596922 | NA |
| lncRNA:CR | 0 | NA | NA | NA | NA | NA |
| lncRNA:CR | 0 | NA | NA | NA | NA | NA |
| asRNA:CR4 | 0 | NA | NA | NA | NA | NA |
| lncRNA:CR | 0 | NA | NA | NA | NA | NA |
| asRNA:CR4 | 0 | NA | NA | NA | NA | NA |
| lncRNA:CR | 25.17312 | 6.917906 | 3.91192 | 1.768417 | 0.076991 | NA |
| lncRNA:CR | 38.69348 | 1.176553 | 3.952124 | 0.297701 | 0.765931 | NA |
| lncRNA:CR | 25.75618 | 1.74666 | 3.952918 | 0.441866 | 0.658586 | NA |
| lncRNA:CR | 0 | NA | NA | NA | NA | NA |
| CG46025 | 0 | NA | NA | NA | NA | NA |
| CG46026 | 0 | NA | NA | NA | NA | NA |
| CR46027 | 0 | NA | NA | NA | NA | NA |
| lncRNA:CR | 0 | NA | NA | NA | NA | NA |
| asRNA:CR4 | 3.587175 | 0.666222 | 3.934688 | 0.16932 | 0.865545 | NA |
| lncRNA:CR | 0 | NA | NA | NA | NA | NA |
| lncRNA:CR | 0 | NA | NA | NA | NA | NA |
| lncRNA:CR | 0 | NA | NA | NA | NA | NA |
| asRNA:CR4 | 0 | NA | NA | NA | NA | NA |
| asRNA:CR4 | 0 | NA | NA | NA | NA | NA |
| lncRNA:CR | 7.060928 | 0.241064 | 3.929271 | 0.061351 | 0.95108 | NA |
| lncRNA:fla | 37.98432 | -0.00713 | 3.671125 | -0.00194 | 0.998451 | NA |
| asRNA:CR4 | 0 | NA | NA | NA | NA | NA |
| lncRNA:CR | 0 | NA | NA | NA | NA | NA |
| lncRNA:CR | 0 | NA | NA | NA | NA | NA |
| lncRNA:CR | 0 | NA | NA | NA | NA | NA |
| CG46042 | 0 | NA | NA | NA | NA | NA |
| lncRNA:CR | 0 | NA | NA | NA | NA | NA |
| asRNA:CR4 | 6.192502 | -0.21435 | 3.95184 | -0.05424 | 0.956744 | NA |

| | | | | | | |
|---|---|---|---|---|---|---|
| asRNA:CR4 | 0 | NA | NA | NA | NA | NA |
| lncRNA:CR | 0 | NA | NA | NA | NA | NA |
| asRNA:CR4 | 0 | NA | NA | NA | NA | NA |
| asRNA:CR4 | 36.44735 | -2.05857 | 3.837895 | -0.53638 | 0.591696 | NA |
| asRNA:CR4 | 0 | NA | NA | NA | NA | NA |
| asRNA:CR4 | 3.557191 | 1.503975 | 3.960718 | 0.379723 | 0.704151 | NA |
| lncRNA:CR | 0 | NA | NA | NA | NA | NA |
| lncRNA:CR | 0 | NA | NA | NA | NA | NA |
| asRNA:CR4 | 0 | NA | NA | NA | NA | NA |
| asRNA:CR4 | 0 | NA | NA | NA | NA | NA |
| lncRNA:CR | 2.680666 | -3.72746 | 3.969815 | -0.93895 | 0.347756 | NA |
| lncRNA:CR | 0 | NA | NA | NA | NA | NA |
| asRNA:CR4 | 0 | NA | NA | NA | NA | NA |
| CG46059 | 0 | NA | NA | NA | NA | NA |
| lncRNA:CR | 0 | NA | NA | NA | NA | NA |
| lncRNA:CR | 0 | NA | NA | NA | NA | NA |
| lncRNA:CR | 0 | NA | NA | NA | NA | NA |
| asRNA:CR4 | 0 | NA | NA | NA | NA | NA |
| asRNA:CR4 | 10.02672 | -0.48795 | 3.879095 | -0.12579 | 0.899899 | NA |
| asRNA:CR4 | 0 | NA | NA | NA | NA | NA |
| asRNA:CR4 | 5.519652 | -1.68736 | 3.957756 | -0.42634 | 0.669859 | NA |
| asRNA:CR4 | 0 | NA | NA | NA | NA | NA |
| asRNA:CR4 | 0 | NA | NA | NA | NA | NA |
| asRNA:CR4 | 0 | NA | NA | NA | NA | NA |
| CR46072 | 0 | NA | NA | NA | NA | NA |
| lncRNA:CR | 29.83767 | 0.042745 | 3.951496 | 0.010817 | 0.991369 | NA |
| asRNA:CR4 | 0 | NA | NA | NA | NA | NA |
| lncRNA:CR | 0.178812 | -1.08808 | 4.061735 | -0.26789 | 0.788788 | NA |
| lncRNA:CR | 0 | NA | NA | NA | NA | NA |
| lncRNA:CR | 0 | NA | NA | NA | NA | NA |
| lncRNA:CR | 0.4188 | 1.41068 | 4.034613 | 0.349644 | 0.726606 | NA |
| lncRNA:CR | 0 | NA | NA | NA | NA | NA |
| asRNA:CR4 | 0 | NA | NA | NA | NA | NA |
| asRNA:CR4 | 30.44537 | 1.53513 | 3.95209 | 0.388435 | 0.697694 | NA |
| lncRNA:CR | 0 | NA | NA | NA | NA | NA |
| lncRNA:CR | 0 | NA | NA | NA | NA | NA |
| lncRNA:CR | 0 | NA | NA | NA | NA | NA |
| lncRNA:CR | 0 | NA | NA | NA | NA | NA |
| lncRNA:CR | 0 | NA | NA | NA | NA | NA |
| asRNA:CR4 | 0 | NA | NA | NA | NA | NA |
| lncRNA:CR | 1.320777 | -1.83209 | 3.977996 | -0.46056 | 0.645117 | NA |
| asRNA:CR4 | 0 | NA | NA | NA | NA | NA |
| asRNA:CR4 | 0 | NA | NA | NA | NA | NA |
| asRNA:CR4 | 37.16358 | 7.864241 | 3.908785 | 2.01194 | 0.044226 | NA |
| lncRNA:CR | 0.5235 | 1.490714 | 4.02555 | 0.370313 | 0.711149 | NA |
| asRNA:CR4 | 0 | NA | NA | NA | NA | NA |
| asRNA:CR4 | 0 | NA | NA | NA | NA | NA |
| lncRNA:CR | 2.082181 | -0.96846 | 3.93372 | -0.2462 | 0.805531 | NA |
| asRNA:CR4 | 0 | NA | NA | NA | NA | NA |
| lncRNA:CR | 0 | NA | NA | NA | NA | NA |

| | | | | | | |
|---|---|---|---|---|---|---|
| asRNA:CR4 | 0 | NA | NA | NA | NA | NA |
| lncRNA:CR | 0 | NA | NA | NA | NA | NA |
| asRNA:CR4 | 0.3141 | 1.240997 | 4.03841 | 0.307299 | 0.758616 | NA |
| asRNA:CR4 | 0.712213 | -1.57119 | 3.998613 | -0.39293 | 0.694369 | NA |
| lncRNA:CR | 0 | NA | NA | NA | NA | NA |
| asRNA:CR4 | 1.849088 | -1.85577 | 3.970721 | -0.46736 | 0.640241 | NA |
| lncRNA:CR | 1.675199 | 1.648777 | 3.978727 | 0.414398 | 0.678583 | NA |
| asRNA:CR4 | 8.452972 | -1.9072 | 3.955468 | -0.48217 | 0.629686 | NA |
| lncRNA:CR | 0.837599 | 1.568055 | 4.001801 | 0.391837 | 0.695178 | NA |
| lncRNA:CR | 0.178053 | -1.16467 | 4.075067 | -0.2858 | 0.775028 | NA |
| lncRNA:CR | 0 | NA | NA | NA | NA | NA |
| lncRNA:CR | 0 | NA | NA | NA | NA | NA |
| lncRNA:CR | 0 | NA | NA | NA | NA | NA |
| lncRNA:CR | 35.78123 | 2.772161 | 3.917333 | 0.707665 | 0.479153 | NA |
| lncRNA:CR | 50.36067 | 1.75058 | 3.951939 | 0.442967 | 0.657789 | NA |
| lncRNA:CR | 0 | NA | NA | NA | NA | NA |
| lncRNA:CR | 0 | NA | NA | NA | NA | NA |
| lncRNA:CR | 0 | NA | NA | NA | NA | NA |
| rgr | 24.45421 | -0.95665 | 3.870901 | -0.24714 | 0.804801 | NA |
| lncRNA:CR | 0.712972 | -2.55723 | 4.00634 | -0.63829 | 0.523282 | NA |
| lncRNA:CR | 0 | NA | NA | NA | NA | NA |
| lncRNA:CR | 0 | NA | NA | NA | NA | NA |
| lncRNA:CR | 1.111482 | -3.16801 | 3.986608 | -0.79466 | 0.426809 | NA |
| lncRNA:CR | 10.7864 | -0.32907 | 3.842081 | -0.08565 | 0.931746 | NA |
| lncRNA:CR | 17.77561 | -6.78916 | 3.913679 | -1.73473 | 0.082789 | NA |
| lncRNA:CR | 0.233093 | 0.692656 | 4.076246 | 0.169925 | 0.865069 | NA |
| asRNA:CR4 | 11.18848 | 1.167813 | 3.955086 | 0.295269 | 0.767789 | NA |
| asRNA:CR4 | 0 | NA | NA | NA | NA | NA |
| lncRNA:CR | 0 | NA | NA | NA | NA | NA |
| lncRNA:CR | 27.32668 | 1.747117 | 3.952803 | 0.441994 | 0.658493 | NA |
| lncRNA:CR | 0.178812 | -1.08808 | 4.061735 | -0.26789 | 0.788788 | NA |
| asRNA:CR4 | 0 | NA | NA | NA | NA | NA |
| asRNA:CR4 | 0 | NA | NA | NA | NA | NA |
| asRNA:CR4 | 0.388212 | 0.094116 | 4.038892 | 0.023303 | 0.981409 | NA |
| asRNA:CR4 | 0.178812 | -1.08808 | 4.061735 | -0.26789 | 0.788788 | NA |
| asRNA:CR4 | 51.86084 | 1.254897 | 3.729034 | 0.336521 | 0.736478 | NA |
| asRNA:CR4 | 0 | NA | NA | NA | NA | NA |
| asRNA:CR4 | 31.48333 | 1.686518 | 3.952477 | 0.426699 | 0.669598 | NA |
| asRNA:CR4 | 0 | NA | NA | NA | NA | NA |
| lncRNA:CR | 8.606562 | 1.391373 | 3.954671 | 0.35183 | 0.724965 | NA |
| lncRNA:CR | 0.388059 | -0.03373 | 4.021335 | -0.00839 | 0.993308 | NA |
| lncRNA:CR | 0 | NA | NA | NA | NA | NA |
| asRNA:CR4 | 9.614877 | -1.69611 | 3.95487 | -0.42887 | 0.66802 | NA |
| lncRNA:CR | 0 | NA | NA | NA | NA | NA |
| lncRNA:CR | 0 | NA | NA | NA | NA | NA |
| asRNA:CR4 | 0 | NA | NA | NA | NA | NA |
| CR46150 | 0 | NA | NA | NA | NA | NA |
| CR46151 | 0 | NA | NA | NA | NA | NA |
| CR46152 | 0 | NA | NA | NA | NA | NA |
| lncRNA:CR | 0 | NA | NA | NA | NA | NA |

| | | | | | | |
|---|---|---|---|---|---|---|
| CR46154 | 0 | NA | NA | NA | NA | NA |
| CR46155 | 0 | NA | NA | NA | NA | NA |
| lncRNA:CR | 0 | NA | NA | NA | NA | NA |
| CR46157 | 0 | NA | NA | NA | NA | NA |
| CR46158 | 0 | NA | NA | NA | NA | NA |
| CR46159 | 0 | NA | NA | NA | NA | NA |
| CR46160 | 0 | NA | NA | NA | NA | NA |
| CR46161 | 0 | NA | NA | NA | NA | NA |
| CR46162 | 0 | NA | NA | NA | NA | NA |
| CR46163 | 0 | NA | NA | NA | NA | NA |
| CR46164 | 0 | NA | NA | NA | NA | NA |
| CR46165 | 0 | NA | NA | NA | NA | NA |
| CR46166 | 0 | NA | NA | NA | NA | NA |
| CR46167 | 0 | NA | NA | NA | NA | NA |
| CR46168 | 0 | NA | NA | NA | NA | NA |
| CR46169 | 0 | NA | NA | NA | NA | NA |
| CR46170 | 0 | NA | NA | NA | NA | NA |
| CR46171 | 0 | NA | NA | NA | NA | NA |
| CR46172 | 0 | NA | NA | NA | NA | NA |
| CR46173 | 0 | NA | NA | NA | NA | NA |
| CR46174 | 0 | NA | NA | NA | NA | NA |
| CR46175 | 0 | NA | NA | NA | NA | NA |
| CR46176 | 0 | NA | NA | NA | NA | NA |
| CR46177 | 0 | NA | NA | NA | NA | NA |
| CR46178 | 0 | NA | NA | NA | NA | NA |
| CR46179 | 0 | NA | NA | NA | NA | NA |
| CR46180 | 0 | NA | NA | NA | NA | NA |
| CR46181 | 0 | NA | NA | NA | NA | NA |
| CR46182 | 0 | NA | NA | NA | NA | NA |
| CR46183 | 0 | NA | NA | NA | NA | NA |
| CR46184 | 0 | NA | NA | NA | NA | NA |
| CR46185 | 0 | NA | NA | NA | NA | NA |
| CR46186 | 0 | NA | NA | NA | NA | NA |
| CR46187 | 0 | NA | NA | NA | NA | NA |
| CR46188 | 0 | NA | NA | NA | NA | NA |
| CR46189 | 0 | NA | NA | NA | NA | NA |
| CR46190 | 0 | NA | NA | NA | NA | NA |
| CG46191 | 0 | NA | NA | NA | NA | NA |
| CG46192 | 0 | NA | NA | NA | NA | NA |
| CG46193 | 0 | NA | NA | NA | NA | NA |
| asRNA:CR4 | 0 | NA | NA | NA | NA | NA |
| lncRNA:CR | 0 | NA | NA | NA | NA | NA |
| lncRNA:CR | 0 | NA | NA | NA | NA | NA |
| lncRNA:CR | 0 | NA | NA | NA | NA | NA |
| lncRNA:CR | 0 | NA | NA | NA | NA | NA |
| lncRNA:CR | 1.778595 | 1.471835 | 3.969892 | 0.370749 | 0.710824 | NA |
| lncRNA:CR | 0 | NA | NA | NA | NA | NA |
| lncRNA:CR | 0 | NA | NA | NA | NA | NA |
| lncRNA:CR | 8.030548 | 1.525016 | 3.95632 | 0.385463 | 0.699894 | NA |
| lncRNA:CR | 0 | NA | NA | NA | NA | NA |

| | | | | | | |
|---|---|---|---|---|---|---|
| lncRNA:CR | 0 | NA | NA | NA | NA | NA |
| lncRNA:CR | 0 | NA | NA | NA | NA | NA |
| lncRNA:CR | 0 | NA | NA | NA | NA | NA |
| lncRNA:CR | 0 | NA | NA | NA | NA | NA |
| lncRNA:CR | 0 | NA | NA | NA | NA | NA |
| lncRNA:CR | 0 | NA | NA | NA | NA | NA |
| lncRNA:CR | 0 | NA | NA | NA | NA | NA |
| lncRNA:CR | 0 | NA | NA | NA | NA | NA |
| lncRNA:CR | 0 | NA | NA | NA | NA | NA |
| lncRNA:CR | 0.356107 | -1.47099 | 4.037764 | -0.36431 | 0.715628 | NA |
| lncRNA:CR | 0 | NA | NA | NA | NA | NA |
| lncRNA:CR | 0 | NA | NA | NA | NA | NA |
| lncRNA:CR | 0 | NA | NA | NA | NA | NA |
| lncRNA:CR | 0.233093 | 0.692656 | 4.076246 | 0.169925 | 0.865069 | NA |
| lncRNA:CR | 0 | NA | NA | NA | NA | NA |
| lncRNA:CR | 0 | NA | NA | NA | NA | NA |
| lncRNA:CR | 0 | NA | NA | NA | NA | NA |
| lncRNA:CR | 0 | NA | NA | NA | NA | NA |
| lncRNA:CR | 13.78281 | -3.1405 | 3.92095 | -0.80095 | 0.423159 | NA |
| lncRNA:CR | 0.233093 | 0.692656 | 4.076246 | 0.169925 | 0.865069 | NA |
| lncRNA:CR | 0 | NA | NA | NA | NA | NA |
| lncRNA:CR | 0 | NA | NA | NA | NA | NA |
| lncRNA:CR | 0 | NA | NA | NA | NA | NA |
| CR46232 | 0 | NA | NA | NA | NA | NA |
| lncRNA:CR | 0 | NA | NA | NA | NA | NA |
| lncRNA:CR | 0 | NA | NA | NA | NA | NA |
| lncRNA:CR | 4.187997 | 1.708209 | 3.962778 | 0.431063 | 0.666422 | NA |
| lncRNA:CR | 0 | NA | NA | NA | NA | NA |
| lncRNA:CR | 0 | NA | NA | NA | NA | NA |
| lncRNA:CR | 0 | NA | NA | NA | NA | NA |
| lncRNA:CR | 0 | NA | NA | NA | NA | NA |
| lncRNA:CR | 0 | NA | NA | NA | NA | NA |
| lncRNA:CR | 0 | NA | NA | NA | NA | NA |
| lncRNA:CR | 0 | NA | NA | NA | NA | NA |
| lncRNA:CR | 0 | NA | NA | NA | NA | NA |
| CG46244 | 0 | NA | NA | NA | NA | NA |
| lncRNA:CR | 0 | NA | NA | NA | NA | NA |
| asRNA:CR4 | 21.34314 | 1.533287 | 3.952591 | 0.387919 | 0.698076 | NA |
| corolla | 0 | NA | NA | NA | NA | NA |
| lncRNA:CR | 0.365171 | -0.43584 | 4.056359 | -0.10745 | 0.914434 | NA |
| lncRNA:CR | 0 | NA | NA | NA | NA | NA |
| mh | 32.79769 | 0.40497 | 3.835261 | 0.105591 | 0.915907 | NA |
| lncRNA:CR | 0 | NA | NA | NA | NA | NA |
| lncRNA:CR | 1.224286 | 0.5443 | 3.941384 | 0.138099 | 0.890162 | NA |
| asRNA:CR4 | 0 | NA | NA | NA | NA | NA |
| lncRNA:CR | 0 | NA | NA | NA | NA | NA |
| lncRNA:CR | 49.26498 | -1.92033 | 3.951699 | -0.48595 | 0.627001 | NA |
| asRNA:CR4 | 0 | NA | NA | NA | NA | NA |
| asRNA:CR4 | 0 | NA | NA | NA | NA | NA |
| lncRNA:CR | 0.1047 | 0.613261 | 4.062799 | 0.150945 | 0.880019 | NA |

| | | | | | | |
|---|---|---|---|---|---|---|
| lncRNA:CR | 60.80649 | -5.92873 | 3.798939 | -1.56063 | 0.118612 | NA |
| lncRNA:CR | 0 | NA | NA | NA | NA | NA |
| lncRNA:CR | 3.399867 | 0.625277 | 3.948538 | 0.158357 | 0.874176 | NA |
| asRNA:CR4 | 0.6282 | 1.523021 | 4.015426 | 0.379293 | 0.704471 | NA |
| asRNA:CR4 | 7.175252 | 2.865514 | 3.90792 | 0.733258 | 0.463401 | NA |
| ltl | 33.89211 | 2.282995 | 3.888173 | 0.587164 | 0.557094 | NA |
| CG4836 | 0.76953 | -0.78481 | 3.994657 | -0.19646 | 0.844247 | NA |
| asRNA:CR4 | 0 | NA | NA | NA | NA | NA |
| lncRNA:CR | 4.195678 | 1.148144 | 3.961859 | 0.289799 | 0.77197 | NA |
| asRNA:CR4 | 0 | NA | NA | NA | NA | NA |
| lncRNA:CR | 0 | NA | NA | NA | NA | NA |
| asRNA:CR4 | 0 | NA | NA | NA | NA | NA |
| CG46270 | 0 | NA | NA | NA | NA | NA |
| lncRNA:CR | 21.00638 | -4.09826 | 3.910875 | -1.04791 | 0.294679 | NA |
| CR46272 | 0 | NA | NA | NA | NA | NA |
| CR46273 | 0 | NA | NA | NA | NA | NA |
| CR46274 | 0 | NA | NA | NA | NA | NA |
| SdicA | 0 | NA | NA | NA | NA | NA |
| SdicB | 0 | NA | NA | NA | NA | NA |
| SdicC | 0 | NA | NA | NA | NA | NA |
| CR46279 | 0.132078 | -1.13211 | 4.069918 | -0.27817 | 0.780885 | NA |
| PPO1 | 0 | NA | NA | NA | NA | NA |
| solo | 0 | NA | NA | NA | NA | NA |
| vas | 2.826898 | 1.687978 | 3.968112 | 0.425386 | 0.670556 | NA |
| Drs | 0 | NA | NA | NA | NA | NA |
| IMPPP | 0 | NA | NA | NA | NA | NA |
| Dhc16F | 1.775574 | -1.56317 | 3.850786 | -0.40593 | 0.684791 | NA |
| Alp1 | 0 | NA | NA | NA | NA | NA |
| Ak2 | 36.24403 | -0.32411 | 3.715943 | -0.08722 | 0.930495 | NA |
| lncRNA:CR | 0.178812 | -1.08808 | 4.061735 | -0.26789 | 0.788788 | NA |
| nw | 0 | NA | NA | NA | NA | NA |
| Duox | 2.702574 | -1.65369 | 3.877152 | -0.42652 | 0.669727 | NA |
| vilya | 46.89963 | -7.63202 | 3.912315 | -1.95077 | 0.051085 | NA |
| CG46298 | 0 | NA | NA | NA | NA | NA |
| lncRNA:CR | 0 | NA | NA | NA | NA | NA |
| CR46300 | 0 | NA | NA | NA | NA | NA |
| Osi21 | 0 | NA | NA | NA | NA | NA |
| CG31600 | 0 | NA | NA | NA | NA | NA |
| CG46302 | 2.241108 | -0.36611 | 3.822753 | -0.09577 | 0.923702 | NA |
| CG46303 | 5.777958 | -5.10368 | 3.905934 | -1.30665 | 0.191332 | NA |
| CG46304 | 0 | NA | NA | NA | NA | NA |
| CG46305 | 0 | NA | NA | NA | NA | NA |
| CG46306 | 0 | NA | NA | NA | NA | NA |
| CG46307 | 0 | NA | NA | NA | NA | NA |
| CG46308 | 0 | NA | NA | NA | NA | NA |
| CG46309 | 0 | NA | NA | NA | NA | NA |
| CG46310 | 0 | NA | NA | NA | NA | NA |
| CG46311 | 0 | NA | NA | NA | NA | NA |
| CG46312 | 0 | NA | NA | NA | NA | NA |
| CG46313 | 0 | NA | NA | NA | NA | NA |

| | | | | | | |
|---|---|---|---|---|---|---|
| CG46314 | 0 | NA | NA | NA | NA | NA |
| CG46315 | 0 | NA | NA | NA | NA | NA |
| CG46316 | 0 | NA | NA | NA | NA | NA |
| CG46317 | 0 | NA | NA | NA | NA | NA |
| CG46318 | 0 | NA | NA | NA | NA | NA |
| CG46319 | 0 | NA | NA | NA | NA | NA |
| CG46320 | 0 | NA | NA | NA | NA | NA |
| CG46321 | 0 | NA | NA | NA | NA | NA |
| lncRNA:CR | 0.1047 | 0.613261 | 4.062799 | 0.150945 | 0.880019 | NA |
| l(2)k05911 | 0 | NA | NA | NA | NA | NA |
| ds | 2.796185 | -0.43502 | 3.958512 | -0.10989 | 0.912493 | NA |
| Oaz | 55.08358 | 0.295828 | 3.950253 | 0.074888 | 0.940303 | NA |
| lncRNA:TS | 0.104623 | 0.639075 | 4.067674 | 0.157111 | 0.875158 | NA |
| lncRNA:TS | 0 | NA | NA | NA | NA | NA |
| lncRNA:TS | 0 | NA | NA | NA | NA | NA |
| lncRNA:CR | 0 | NA | NA | NA | NA | NA |
| lncRNA:TS | 0.411147 | -0.29058 | 4.046973 | -0.0718 | 0.94276 | NA |
| lncRNA:CR | 0 | NA | NA | NA | NA | NA |
| lncRNA:TS | 0 | NA | NA | NA | NA | NA |
| lncRNA:TS | 0.104623 | 0.639075 | 4.067674 | 0.157111 | 0.875158 | NA |
| lncRNA:TS | 0 | NA | NA | NA | NA | NA |
| lncRNA:TS | 0 | NA | NA | NA | NA | NA |
| lncRNA:TS | 0 | NA | NA | NA | NA | NA |
| lncRNA:TS | 0.283436 | -0.20033 | 4.070792 | -0.04921 | 0.960751 | NA |
| btl | 0 | NA | NA | NA | NA | NA |
| mah | 0 | NA | NA | NA | NA | NA |
| CG46335 | 0.4188 | 1.41068 | 4.034613 | 0.349644 | 0.726606 | NA |
| lncRNA:CR | 0 | NA | NA | NA | NA | NA |
| knon | 0 | NA | NA | NA | NA | NA |
| Fuca | 0 | NA | NA | NA | NA | NA |
| sisRNA:1 | 0 | NA | NA | NA | NA | NA |
| hpRNA:1 | 0 | NA | NA | NA | NA | NA |
| sisRNA:4 | 0.132078 | -1.13211 | 4.069918 | -0.27817 | 0.780885 | NA |
| CG32500 | 2.823422 | -0.23191 | 3.951154 | -0.05869 | 0.953196 | NA |
| prg | 24.01921 | 5.432886 | 3.853673 | 1.409794 | 0.1586 | NA |
| hpRNA:CR | 0 | NA | NA | NA | NA | NA |
| hpRNA:CR | 0 | NA | NA | NA | NA | NA |
| lncRNA:CR | 24.21525 | -1.70339 | 3.952491 | -0.43097 | 0.666493 | NA |
| lncRNA:CR | 0.894061 | -2.00006 | 4.030294 | -0.49626 | 0.619714 | NA |
| lncRNA:CR | 26.05119 | 1.534399 | 3.952288 | 0.388231 | 0.697845 | NA |
| sisRNA:CR | 0.3141 | 1.240997 | 4.03841 | 0.307299 | 0.758616 | NA |
| sisRNA:CR | 0 | NA | NA | NA | NA | NA |
| sisRNA:3 | 41.16015 | 1.065863 | 3.91908 | 0.271968 | 0.785647 | NA |
| sisRNA:CR | 0 | NA | NA | NA | NA | NA |
| sisRNA:CR | 0 | NA | NA | NA | NA | NA |
| sisRNA:2 | 0 | NA | NA | NA | NA | NA |
| sisRNA:CR | 0 | NA | NA | NA | NA | NA |
| sisRNA:CR | 0 | NA | NA | NA | NA | NA |
| AdamTS-A | 0.932373 | 1.053996 | 3.996193 | 0.26375 | 0.791972 | NA |
| stw | 30.98428 | 1.72462 | 3.898599 | 0.442369 | 0.658222 | NA |

| | | | | | | |
|---|---|---|---|---|---|---|
| ich | 46.84806 | -3.6295 | 3.914224 | -0.92726 | 0.353792 | NA |
| vrs | 0 | NA | NA | NA | NA | NA |
| Had1 | 0 | NA | NA | NA | NA | NA |
| hll | 0.132078 | -1.13211 | 4.069918 | -0.27817 | 0.780885 | NA |
| snoRNA:Oı | 0.5235 | 1.490714 | 4.02555 | 0.370313 | 0.711149 | NA |
| snoRNA:Oı | 3.455098 | 1.699144 | 3.965156 | 0.428519 | 0.668273 | NA |
| snoRNA:Oı | 0 | NA | NA | NA | NA | NA |
| snoRNA:Oı | 0 | NA | NA | NA | NA | NA |
| snoRNA:Oı | 0 | NA | NA | NA | NA | NA |
| snoRNA:Oı | 30.4167 | 4.477644 | 3.850131 | 1.162985 | 0.244836 | NA |
| snoRNA:M | 0 | NA | NA | NA | NA | NA |
| snoRNA:M | 0.536437 | -1.71175 | 4.038042 | -0.42391 | 0.671635 | NA |
| scaRNA:M | 0 | NA | NA | NA | NA | NA |
| snoRNA:Oı | 31.51468 | 1.748118 | 3.952553 | 0.442276 | 0.65829 | NA |
| snoRNA:Oı | 0 | NA | NA | NA | NA | NA |
| scaRNA:M | 0 | NA | NA | NA | NA | NA |
| snoRNA:M | 0 | NA | NA | NA | NA | NA |
| snoRNA:Oı | 0 | NA | NA | NA | NA | NA |
| CG46383 | 0 | NA | NA | NA | NA | NA |
| snoRNA:M | 0 | NA | NA | NA | NA | NA |
| Orc1 | 0.31387 | 1.261782 | 4.037145 | 0.312543 | 0.754628 | NA |
| CG46388 | 0 | NA | NA | NA | NA | NA |
| lncRNA:osl | 0 | NA | NA | NA | NA | NA |
| lncRNA:SM | 0 | NA | NA | NA | NA | NA |
| lncRNA:CR | 0 | NA | NA | NA | NA | NA |
| lncRNA:en | 0 | NA | NA | NA | NA | NA |
| lncRNA:CR | 40.77362 | 6.844073 | 3.846532 | 1.779284 | 0.075193 | NA |
| CG46394 | 0.104623 | 0.639075 | 4.067674 | 0.157111 | 0.875158 | NA |
| CG46395 | 0 | NA | NA | NA | NA | NA |
| CG46396 | 0 | NA | NA | NA | NA | NA |
| CG46398 | 0 | NA | NA | NA | NA | NA |
| CG46399 | 0 | NA | NA | NA | NA | NA |
| sbRNA:1 | 0 | NA | NA | NA | NA | NA |
| CR46401 | 0.356107 | -1.47099 | 4.037764 | -0.36431 | 0.715628 | NA |
| CR46402 | 0.924544 | -1.7993 | 3.988349 | -0.45114 | 0.651889 | NA |
| CR46403 | 0 | NA | NA | NA | NA | NA |
| CR46404 | 0.104623 | 0.639075 | 4.067674 | 0.157111 | 0.875158 | NA |
| CR46406 | 0 | NA | NA | NA | NA | NA |
| CR46407 | 0 | NA | NA | NA | NA | NA |
| lncRNA:CR | 0 | NA | NA | NA | NA | NA |
| lncRNA:CR | 0 | NA | NA | NA | NA | NA |
| lncRNA:CR | 0 | NA | NA | NA | NA | NA |
| lncRNA:CR | 0 | NA | NA | NA | NA | NA |
| CG46412 | 0 | NA | NA | NA | NA | NA |
| CR46413 | 0 | NA | NA | NA | NA | NA |
| lncRNA:CR | 0 | NA | NA | NA | NA | NA |
| CR46415 | 0 | NA | NA | NA | NA | NA |
| lncRNA:CR | 0 | NA | NA | NA | NA | NA |
| CR46417 | 0 | NA | NA | NA | NA | NA |
| CR46418 | 0 | NA | NA | NA | NA | NA |

| | | | | | | |
|---|---|---|---|---|---|---|
| CR46419 | 0 | NA | NA | NA | NA | NA |
| CR46420 | 0 | NA | NA | NA | NA | NA |
| CR46421 | 0 | NA | NA | NA | NA | NA |
| CR46422 | 0 | NA | NA | NA | NA | NA |
| CR46423 | 0 | NA | NA | NA | NA | NA |
| CR46424 | 0 | NA | NA | NA | NA | NA |
| CG46426 | 0 | NA | NA | NA | NA | NA |
| CG46427 | 0 | NA | NA | NA | NA | NA |
| CG46428 | 0 | NA | NA | NA | NA | NA |
| CG46429 | 0 | NA | NA | NA | NA | NA |
| CG46430 | 0.178053 | -1.16467 | 4.075067 | -0.2858 | 0.775028 | NA |
| lncRNA:CR | 0 | NA | NA | NA | NA | NA |
| CR46432 | 0 | NA | NA | NA | NA | NA |
| CG46433 | 0 | NA | NA | NA | NA | NA |
| CG46434 | 0 | NA | NA | NA | NA | NA |
| CG46435 | 0 | NA | NA | NA | NA | NA |
| Osi10a | 28.00047 | -1.91823 | 3.952298 | -0.48535 | 0.627431 | NA |
| Osi10b | 4.754797 | -1.89533 | 3.958922 | -0.47875 | 0.632117 | NA |
| CG46439 | 0 | NA | NA | NA | NA | NA |
| Iyd | 0 | NA | NA | NA | NA | NA |
| FASN3 | 0.732899 | 1.548046 | 4.007786 | 0.38626 | 0.699304 | NA |
| nvd | 0.872712 | -0.58069 | 3.940867 | -0.14735 | 0.882854 | NA |
| CG46440 | 0 | NA | NA | NA | NA | NA |
| CG46441 | 0 | NA | NA | NA | NA | NA |
| CG46442 | 0 | NA | NA | NA | NA | NA |
| CG46443 | 0 | NA | NA | NA | NA | NA |
| lncRNA:CR | 0 | NA | NA | NA | NA | NA |
| lncRNA:CR | 0 | NA | NA | NA | NA | NA |
| lncRNA:CR | 0 | NA | NA | NA | NA | NA |
| CR46447 | 0 | NA | NA | NA | NA | NA |
| slf | 3.21584 | -3.35157 | 3.92887 | -0.85306 | 0.393624 | NA |
| Nplp2 | 0.132078 | -1.13211 | 4.069918 | -0.27817 | 0.780885 | NA |
| lncRNA:CR | 0 | NA | NA | NA | NA | NA |
| lncRNA:CR | 0.985641 | 0.934855 | 3.921487 | 0.238393 | 0.811576 | NA |
| asRNA:CR4 | 0 | NA | NA | NA | NA | NA |
| asRNA:CR4 | 0 | NA | NA | NA | NA | NA |
| lncRNA:CR | 5.347781 | -1.55895 | 3.918344 | -0.39786 | 0.690734 | NA |
| lncRNA:CR | 0 | NA | NA | NA | NA | NA |
| CG46456 | 0 | NA | NA | NA | NA | NA |
| CG46457 | 0 | NA | NA | NA | NA | NA |
| CG46458 | 0 | NA | NA | NA | NA | NA |
| CG46459 | 0 | NA | NA | NA | NA | NA |
| CG46460 | 0 | NA | NA | NA | NA | NA |
| CR46461 | 0 | NA | NA | NA | NA | NA |
| CG46463 | 0 | NA | NA | NA | NA | NA |
| CG46464 | 0 | NA | NA | NA | NA | NA |
| CG46465 | 0.178053 | -1.16467 | 4.075067 | -0.2858 | 0.775028 | NA |
| CG46467 | 0 | NA | NA | NA | NA | NA |
| CG46468 | 0 | NA | NA | NA | NA | NA |
| asRNA:CR4 | 0 | NA | NA | NA | NA | NA |

| | | | | | | |
|---|---|---|---|---|---|---|
| asRNA:CR4 | 0 | NA | NA | NA | NA | NA |
| asRNA:CR4 | 0.46073 | -0.17853 | 4.00793 | -0.04454 | 0.964471 | NA |
| lncRNA:CR | 0 | NA | NA | NA | NA | NA |
| asRNA:CR4 | 1.188699 | -1.8233 | 3.980741 | -0.45803 | 0.646931 | NA |
| lncRNA:CR | 0 | NA | NA | NA | NA | NA |
| asRNA:CR4 | 20.14764 | 0.653009 | 3.773981 | 0.173029 | 0.862629 | NA |
| asRNA:CR4 | 6.040806 | -0.34722 | 3.850518 | -0.09017 | 0.928149 | NA |
| lncRNA:CR | 0 | NA | NA | NA | NA | NA |
| dbf | 0.396233 | -1.6814 | 4.028351 | -0.41739 | 0.676392 | NA |
| CG46491 | 0.1047 | 0.613261 | 4.062799 | 0.150945 | 0.880019 | NA |
| CG46492 | 0.53416 | -1.53481 | 4.012395 | -0.38252 | 0.702078 | NA |
| lncRNA:Vir | 0.732899 | 1.548046 | 4.007786 | 0.38626 | 0.699304 | NA |
| ccb | 20.7453 | 1.173452 | 3.953172 | 0.296838 | 0.76659 | NA |
| lncRNA:CR | 0 | NA | NA | NA | NA | NA |
| lncRNA:CR | 0.178053 | -1.16467 | 4.075067 | -0.2858 | 0.775028 | NA |
| lncRNA:CR | 0 | NA | NA | NA | NA | NA |
| Pen | 31.99226 | -0.29486 | 3.849831 | -0.07659 | 0.93895 | NA |
| esg | 0 | NA | NA | NA | NA | NA |
| Datp | 0 | NA | NA | NA | NA | NA |
| Myo61F | 0.178053 | -1.16467 | 4.075067 | -0.2858 | 0.775028 | NA |
| tRNA:Glu-( | 0 | NA | NA | NA | NA | NA |
| tRNA:Glu-( | 0 | NA | NA | NA | NA | NA |
| tRNA:Glu-( | 0 | NA | NA | NA | NA | NA |
| tRNA:Glu-( | 0 | NA | NA | NA | NA | NA |
| tRNA:Glu-( | 0 | NA | NA | NA | NA | NA |
| tRNA:Glu-( | 0 | NA | NA | NA | NA | NA |
| tRNA:Glu-( | 0 | NA | NA | NA | NA | NA |
| CG32320 | 0.104623 | 0.639075 | 4.067674 | 0.157111 | 0.875158 | NA |
| tRNA:Glu-( | 0 | NA | NA | NA | NA | NA |
| tRNA:Glu-( | 0 | NA | NA | NA | NA | NA |
| CG9168 | 0 | NA | NA | NA | NA | NA |
| FucTD | 0 | NA | NA | NA | NA | NA |
| robl62A | 0 | NA | NA | NA | NA | NA |
| stet | 49.14683 | 2.326131 | 3.923627 | 0.592852 | 0.55328 | NA |
| Naa30B | 0 | NA | NA | NA | NA | NA |
| SA-2 | 0 | NA | NA | NA | NA | NA |
| CG9173 | 0 | NA | NA | NA | NA | NA |
| CG9184 | 0 | NA | NA | NA | NA | NA |
| Psf1 | 0 | NA | NA | NA | NA | NA |
| CG32318 | 0 | NA | NA | NA | NA | NA |
| Klp61F | 0 | NA | NA | NA | NA | NA |
| CG9192 | 0 | NA | NA | NA | NA | NA |
| Bro | 0 | NA | NA | NA | NA | NA |
| CG12011 | 0 | NA | NA | NA | NA | NA |
| CG12035 | 0.1047 | 0.613261 | 4.062799 | 0.150945 | 0.880019 | NA |
| CG12020 | 0 | NA | NA | NA | NA | NA |
| GV1 | 0 | NA | NA | NA | NA | NA |
| Cpr62Bb | 0 | NA | NA | NA | NA | NA |
| Cpr62Ba | 0.492153 | 0.308117 | 3.993933 | 0.077146 | 0.938507 | NA |
| Dhc62B | 2.057989 | 0.390629 | 3.876247 | 0.100775 | 0.919729 | NA |

| | | | | | | |
|---|---|---|---|---|---|---|
| Cht2 | 0.178053 | -1.16467 | 4.075067 | -0.2858 | 0.775028 | NA |
| PIG-Wb | 0 | NA | NA | NA | NA | NA |
| Oseg4 | 0.209247 | 1.042804 | 4.047837 | 0.25762 | 0.7967 | NA |
| CG13930 | 0 | NA | NA | NA | NA | NA |
| RabX5 | 0 | NA | NA | NA | NA | NA |
| CG13926 | 7.597237 | -0.04835 | 3.749374 | -0.01289 | 0.989712 | NA |
| CG32313 | 7.94478 | 1.493767 | 3.901153 | 0.382904 | 0.701791 | NA |
| CG13924 | 0.178053 | -1.16467 | 4.075067 | -0.2858 | 0.775028 | NA |
| tRNA:Cys-( | 0 | NA | NA | NA | NA | NA |
| Cnb | 0 | NA | NA | NA | NA | NA |
| CG11815 | 0 | NA | NA | NA | NA | NA |
| mu2 | 1.510528 | -3.57574 | 3.956385 | -0.90379 | 0.366107 | NA |
| CG5687 | 37.58727 | 1.749177 | 3.952288 | 0.442573 | 0.658074 | NA |
| dsb | 0.1047 | 0.613261 | 4.062799 | 0.150945 | 0.880019 | NA |
| CNMa | 0 | NA | NA | NA | NA | NA |
| Cpr62Bc | 0 | NA | NA | NA | NA | NA |
| CG13810 | 0.178053 | -1.16467 | 4.075067 | -0.2858 | 0.775028 | NA |
| CG8960 | 0 | NA | NA | NA | NA | NA |
| Oseg2 | 0 | NA | NA | NA | NA | NA |
| CG32301 | 0.1047 | 0.613261 | 4.062799 | 0.150945 | 0.880019 | NA |
| CG32305 | 0.209247 | 1.042804 | 4.047837 | 0.25762 | 0.7967 | NA |
| CG13806 | 0 | NA | NA | NA | NA | NA |
| obst-I | 0 | NA | NA | NA | NA | NA |
| CG32302 | 0 | NA | NA | NA | NA | NA |
| tRNA:Cys-( | 0.104623 | 0.639075 | 4.067674 | 0.157111 | 0.875158 | NA |
| yellow-g2 | 0 | NA | NA | NA | NA | NA |
| CG13801 | 0.178053 | -1.16467 | 4.075067 | -0.2858 | 0.775028 | NA |
| CG32298 | 0 | NA | NA | NA | NA | NA |
| CG32299 | 0 | NA | NA | NA | NA | NA |
| yellow-g | 0.178053 | -1.16467 | 4.075067 | -0.2858 | 0.775028 | NA |
| CG15878 | 0 | NA | NA | NA | NA | NA |
| CG15820 | 0 | NA | NA | NA | NA | NA |
| CR15821 | 0 | NA | NA | NA | NA | NA |
| CG15879 | 18.41716 | 1.242528 | 3.952837 | 0.314338 | 0.753264 | NA |
| CG15822 | 0.2094 | 1.005034 | 4.045341 | 0.248442 | 0.803792 | NA |
| CG16984 | 0.932373 | 1.053996 | 3.996193 | 0.26375 | 0.791972 | NA |
| CG16985 | 0.178812 | -1.08808 | 4.061735 | -0.26789 | 0.788788 | NA |
| MEP-1 | 18.65959 | -7.94727 | 3.909133 | -2.033 | 0.042052 | NA |
| CG1143 | 0.178053 | -1.16467 | 4.075067 | -0.2858 | 0.775028 | NA |
| CG1246 | 0 | NA | NA | NA | NA | NA |
| Acp62F | 0 | NA | NA | NA | NA | NA |
| CG14949 | 2.720205 | 1.493763 | 3.963603 | 0.37687 | 0.70627 | NA |
| CG12182 | 0.547193 | 2.009611 | 4.015384 | 0.500478 | 0.616739 | NA |
| CG32295 | 0.732899 | 1.548046 | 4.007786 | 0.38626 | 0.699304 | NA |
| Cyp4d20 | 0 | NA | NA | NA | NA | NA |
| CG16762 | 0 | NA | NA | NA | NA | NA |
| CPT2 | 32.15775 | -8.55648 | 3.908352 | -2.18928 | 0.028576 | NA |
| CG2113 | 0 | NA | NA | NA | NA | NA |
| aly | 0 | NA | NA | NA | NA | NA |
| CG32487 | 0 | NA | NA | NA | NA | NA |

| | | | | | | |
|---|---|---|---|---|---|---|
| CG32488 | 0 | NA | NA | NA | NA | NA |
| CG32846 | 15.82342 | 6.300527 | 3.898026 | 1.616338 | 0.106021 | NA |
| CG9970 | 14.75188 | 1.530561 | 3.953337 | 0.387157 | 0.69864 | NA |
| spz5 | 0.1047 | 0.613261 | 4.062799 | 0.150945 | 0.880019 | NA |
| snRNA:U5: | 2.376752 | 1.184528 | 3.965092 | 0.298739 | 0.765139 | NA |
| tRNA:Cys-( | 0 | NA | NA | NA | NA | NA |
| tRNA:Cys-( | 0 | NA | NA | NA | NA | NA |
| tRNA:Cys-( | 0 | NA | NA | NA | NA | NA |
| tRNA:Lys-C | 0 | NA | NA | NA | NA | NA |
| tRNA:Cys-( | 0 | NA | NA | NA | NA | NA |
| CG32284 | 0 | NA | NA | NA | NA | NA |
| CG14957 | 0 | NA | NA | NA | NA | NA |
| CG14958 | 0 | NA | NA | NA | NA | NA |
| ckd | 0.341478 | 0.247543 | 4.031301 | 0.061405 | 0.951036 | NA |
| CG14960 | 0.264155 | -1.54099 | 4.048976 | -0.38059 | 0.703509 | NA |
| CG12017 | 0.1047 | 0.613261 | 4.062799 | 0.150945 | 0.880019 | NA |
| CG12009 | 13.05737 | -3.85746 | 3.884392 | -0.99307 | 0.320678 | NA |
| Drsl2 | 0 | NA | NA | NA | NA | NA |
| Drsl3 | 0 | NA | NA | NA | NA | NA |
| Drsl4 | 0.178812 | -1.08808 | 4.061735 | -0.26789 | 0.788788 | NA |
| Drsl5 | 0 | NA | NA | NA | NA | NA |
| CG12016 | 0 | NA | NA | NA | NA | NA |
| CG12010 | 0 | NA | NA | NA | NA | NA |
| Drsl6 | 0 | NA | NA | NA | NA | NA |
| Drsl1 | 0 | NA | NA | NA | NA | NA |
| CG33160 | 0 | NA | NA | NA | NA | NA |
| CG32277 | 0 | NA | NA | NA | NA | NA |
| CG33159 | 0 | NA | NA | NA | NA | NA |
| CG32271 | 0.178053 | -1.16467 | 4.075067 | -0.2858 | 0.775028 | NA |
| CG14966 | 45.76658 | 0.270517 | 3.850027 | 0.070264 | 0.943984 | NA |
| tRNA:Met- | 0 | NA | NA | NA | NA | NA |
| Cht7 | 0 | NA | NA | NA | NA | NA |
| tRNA:Lys-C | 0 | NA | NA | NA | NA | NA |
| CG10357 | 0 | NA | NA | NA | NA | NA |
| CG10862 | 0 | NA | NA | NA | NA | NA |
| CG14974 | 0 | NA | NA | NA | NA | NA |
| ImpE2 | 0 | NA | NA | NA | NA | NA |
| CG32266 | 21.44256 | -3.50308 | 3.902405 | -0.89767 | 0.36936 | NA |
| Awh | 42.05878 | 0.319444 | 3.950691 | 0.080858 | 0.935555 | NA |
| Gr63a | 0 | NA | NA | NA | NA | NA |
| Acp63F | 0 | NA | NA | NA | NA | NA |
| CG32263 | 0 | NA | NA | NA | NA | NA |
| ppk27 | 1.570499 | 1.642769 | 3.980387 | 0.412716 | 0.679815 | NA |
| Ugt305A1 | 0 | NA | NA | NA | NA | NA |
| dib | 0 | NA | NA | NA | NA | NA |
| CG32259 | 24.70788 | 1.174519 | 3.95281 | 0.297135 | 0.766363 | NA |
| CG14990 | 0 | NA | NA | NA | NA | NA |
| Akh | 5.549096 | 1.719007 | 3.959971 | 0.434096 | 0.664219 | NA |
| CG32260 | 0.178053 | -1.16467 | 4.075067 | -0.2858 | 0.775028 | NA |
| CG1299 | 28.85338 | -2.37636 | 3.824083 | -0.62142 | 0.534324 | NA |

| | | | | | | |
|---|---|---|---|---|---|---|
| CG18675 | 0 | NA | NA | NA | NA | NA |
| Cpr64Ad | 0.2094 | 1.005034 | 4.045341 | 0.248442 | 0.803792 | NA |
| pav | 0 | NA | NA | NA | NA | NA |
| CG1273 | 0 | NA | NA | NA | NA | NA |
| asRNA:CR3 | 0 | NA | NA | NA | NA | NA |
| Cpr64Ac | 0 | NA | NA | NA | NA | NA |
| Cpr64Aa | 0.589959 | -1.15593 | 4.01641 | -0.2878 | 0.773499 | NA |
| CG15005 | 5.15103 | -1.89739 | 3.95832 | -0.47934 | 0.631695 | NA |
| mas | 0.46073 | -0.17853 | 4.00793 | -0.04454 | 0.964471 | NA |
| Faa | 29.13396 | 0.70714 | 3.92688 | 0.180077 | 0.857092 | NA |
| Dpy-30L2 | 0 | NA | NA | NA | NA | NA |
| Claspin | 17.58978 | 4.862249 | 3.865835 | 1.257749 | 0.208483 | NA |
| slow | 22.04207 | 5.969532 | 3.832821 | 1.557478 | 0.119357 | NA |
| CG15020 | 0 | NA | NA | NA | NA | NA |
| CG15021 | 0 | NA | NA | NA | NA | NA |
| CG32249 | 0 | NA | NA | NA | NA | NA |
| CG32248 | 6.766025 | -1.69111 | 3.956514 | -0.42742 | 0.66907 | NA |
| CG11349 | 0 | NA | NA | NA | NA | NA |
| CG7465 | 42.57865 | -6.91802 | 3.832657 | -1.80502 | 0.071072 | NA |
| CG11353 | 0 | NA | NA | NA | NA | NA |
| CG32246 | 0 | NA | NA | NA | NA | NA |
| Tie | 2.202539 | 2.272562 | 3.954673 | 0.574652 | 0.565526 | NA |
| CG13720 | 0 | NA | NA | NA | NA | NA |
| CG13721 | 0 | NA | NA | NA | NA | NA |
| CG13722 | 0.132078 | -1.13211 | 4.069918 | -0.27817 | 0.780885 | NA |
| CG32241 | 0.178053 | -1.16467 | 4.075067 | -0.2858 | 0.775028 | NA |
| CG15024 | 0 | NA | NA | NA | NA | NA |
| CG15023 | 0 | NA | NA | NA | NA | NA |
| CG15022 | 0.1047 | 0.613261 | 4.062799 | 0.150945 | 0.880019 | NA |
| CG11345 | 0.178053 | -1.16467 | 4.075067 | -0.2858 | 0.775028 | NA |
| CG32240 | 0 | NA | NA | NA | NA | NA |
| CG11342 | 27.55977 | 3.191976 | 3.909044 | 0.816562 | 0.414179 | NA |
| dyl | 0 | NA | NA | NA | NA | NA |
| TTLL1B | 43.85347 | 4.462231 | 3.748827 | 1.190301 | 0.233928 | NA |
| RhoGEF64 | 53.19271 | -4.35312 | 3.873036 | -1.12396 | 0.261032 | NA |
| CG13713 | 0 | NA | NA | NA | NA | NA |
| CG13711 | 0 | NA | NA | NA | NA | NA |
| Dnah3 | 3.754299 | -3.35495 | 3.965996 | -0.84593 | 0.397592 | NA |
| CG13705 | 35.5289 | -1.91926 | 3.952005 | -0.48564 | 0.627221 | NA |
| CG13704 | 6.794867 | -2.22911 | 3.964412 | -0.56228 | 0.573926 | NA |
| CG32235 | 0 | NA | NA | NA | NA | NA |
| CG32237 | 0.132078 | -1.13211 | 4.069918 | -0.27817 | 0.780885 | NA |
| CG13708 | 20.45977 | -1.90968 | 3.739211 | -0.51072 | 0.609549 | NA |
| CG18808 | 0 | NA | NA | NA | NA | NA |
| CG12493 | 0 | NA | NA | NA | NA | NA |
| CG13712 | 0 | NA | NA | NA | NA | NA |
| CG15876 | 0 | NA | NA | NA | NA | NA |
| CG18418 | 1.022344 | -2.80547 | 3.981013 | -0.70471 | 0.480989 | NA |
| CG7514 | 0.178053 | -1.16467 | 4.075067 | -0.2858 | 0.775028 | NA |
| ATPsynCF6 | 0 | NA | NA | NA | NA | NA |

| | | | | | | |
|---|---|---|---|---|---|---|
| CG32232 | 24.32907 | 0.933374 | 3.903833 | 0.239092 | 0.811035 | NA |
| CG2157 | 4.273279 | -1.68149 | 3.959707 | -0.42465 | 0.671092 | NA |
| CG15208 | 0.1047 | 0.613261 | 4.062799 | 0.150945 | 0.880019 | NA |
| v | 0.178053 | -1.16467 | 4.075067 | -0.2858 | 0.775028 | NA |
| CG2145 | 50.46788 | -6.82068 | 3.836142 | -1.778 | 0.075403 | NA |
| SmydA-4 | 0.95493 | -0.53313 | 3.924428 | -0.13585 | 0.89194 | NA |
| C901 | 1.072874 | -2.0339 | 4.019685 | -0.50598 | 0.612867 | NA |
| Rab9Fb | 0 | NA | NA | NA | NA | NA |
| Rab9Fa | 0 | NA | NA | NA | NA | NA |
| feo | 0.816913 | -0.91625 | 3.980578 | -0.23018 | 0.817951 | NA |
| CG32669 | 11.0323 | -0.23399 | 3.853354 | -0.06072 | 0.951579 | NA |
| Ant2 | 41.14964 | 5.687613 | 3.840096 | 1.481112 | 0.138577 | NA |
| CG2076 | 0 | NA | NA | NA | NA | NA |
| CG2061 | 0 | NA | NA | NA | NA | NA |
| CG15200 | 0 | NA | NA | NA | NA | NA |
| CG11106 | 0 | NA | NA | NA | NA | NA |
| CG15198 | 0.104623 | 0.639075 | 4.067674 | 0.157111 | 0.875158 | NA |
| CG11756 | 0 | NA | NA | NA | NA | NA |
| CG1738 | 11.20282 | 2.941535 | 3.926668 | 0.749117 | 0.453786 | NA |
| sisA | 0 | NA | NA | NA | NA | NA |
| CG1394 | 0.178053 | -1.16467 | 4.075067 | -0.2858 | 0.775028 | NA |
| antdh | 0 | NA | NA | NA | NA | NA |
| CG32668 | 10.69829 | -1.91048 | 3.954523 | -0.48311 | 0.629015 | NA |
| CG15199 | 0.590718 | -0.57492 | 4.003363 | -0.14361 | 0.885809 | NA |
| ssp7 | 0.356107 | -1.47099 | 4.037764 | -0.36431 | 0.715628 | NA |
| CG15196 | 1.960396 | -3.42446 | 3.91235 | -0.87529 | 0.381414 | NA |
| CG11699 | 0 | NA | NA | NA | NA | NA |
| Reepl1 | 0 | NA | NA | NA | NA | NA |
| CG2444 | 0.132078 | -1.13211 | 4.069918 | -0.27817 | 0.780885 | NA |
| CG2533 | 0.104623 | 0.639075 | 4.067674 | 0.157111 | 0.875158 | NA |
| CG1561 | 9.643054 | 1.306255 | 3.954754 | 0.3303 | 0.741173 | NA |
| CG33235 | 1.015653 | 0.790107 | 3.976718 | 0.198683 | 0.842511 | NA |
| Or10a | 0 | NA | NA | NA | NA | NA |
| Gr10a | 0 | NA | NA | NA | NA | NA |
| Gr10b | 0 | NA | NA | NA | NA | NA |
| CG15221 | 1.989299 | 1.663439 | 3.974713 | 0.418505 | 0.675578 | NA |
| m | 24.44335 | -0.39632 | 3.846162 | -0.10304 | 0.917929 | NA |
| Karl | 20.15605 | 1.648222 | 3.818865 | 0.4316 | 0.666032 | NA |
| CG15739 | 11.80318 | 1.681662 | 3.822762 | 0.439908 | 0.660004 | NA |
| CG1840 | 32.49042 | -2.60768 | 3.813826 | -0.68374 | 0.494137 | NA |
| CG1806 | 6.920855 | 1.189058 | 3.955844 | 0.300583 | 0.763733 | NA |
| fw | 10.94123 | 2.606354 | 3.914804 | 0.665769 | 0.505559 | NA |
| CG18130 | 8.404177 | -2.23868 | 3.961941 | -0.56505 | 0.572043 | NA |
| tsg | 0 | NA | NA | NA | NA | NA |
| gd | 1.027601 | -2.18695 | 3.915354 | -0.55856 | 0.576463 | NA |
| Sclp | 0 | NA | NA | NA | NA | NA |
| lncRNA:CR | 0 | NA | NA | NA | NA | NA |
| lncRNA:CR | 0 | NA | NA | NA | NA | NA |
| CG15740 | 0.178053 | -1.16467 | 4.075067 | -0.2858 | 0.775028 | NA |
| CG15741 | 0 | NA | NA | NA | NA | NA |

| | | | | | | |
|---|---|---|---|---|---|---|
| dy | 3.336187 | -0.97256 | 3.796706 | -0.25616 | 0.797829 | NA |
| Cyp318a1 | 0 | NA | NA | NA | NA | NA |
| lncRNA:CR | 34.66276 | -0.75215 | 3.836304 | -0.19606 | 0.844563 | NA |
| CG2750 | 5.440409 | 1.515841 | 3.9574 | 0.38304 | 0.70169 | NA |
| CG1924 | 24.73324 | 0.508069 | 3.762734 | 0.135027 | 0.892591 | NA |
| CG11356 | 0.732363 | 1.393579 | 3.993417 | 0.348969 | 0.727112 | NA |
| moon | 0 | NA | NA | NA | NA | NA |
| Uch-L5R | 0.178053 | -1.16467 | 4.075067 | -0.2858 | 0.775028 | NA |
| Cyp311a1 | 16.4892 | 0.545269 | 3.862506 | 0.14117 | 0.887736 | NA |
| CG2577 | 0.178812 | -1.08808 | 4.061735 | -0.26789 | 0.788788 | NA |
| CG2574 | 0 | NA | NA | NA | NA | NA |
| CG15927 | 0.528311 | -1.72839 | 4.011871 | -0.43082 | 0.6666 | NA |
| Cpr11A | 0.178812 | -1.08808 | 4.061735 | -0.26789 | 0.788788 | NA |
| Lsp1alpha | 0 | NA | NA | NA | NA | NA |
| CG2556 | 0 | NA | NA | NA | NA | NA |
| Fpgs | 10.26059 | 1.734896 | 3.955893 | 0.43856 | 0.660981 | NA |
| CG11085 | 0 | NA | NA | NA | NA | NA |
| Mks1 | 0 | NA | NA | NA | NA | NA |
| CG15731 | 0 | NA | NA | NA | NA | NA |
| CG32651 | 0 | NA | NA | NA | NA | NA |
| Ir11a | 1.361099 | 1.628413 | 3.98439 | 0.408698 | 0.682761 | NA |
| CG15728 | 0 | NA | NA | NA | NA | NA |
| Aven | 1.613947 | 1.895597 | 3.896457 | 0.486492 | 0.626618 | NA |
| CG32642 | 0 | NA | NA | NA | NA | NA |
| Mur11Da | 0 | NA | NA | NA | NA | NA |
| CG15721 | 7.074029 | 4.68901 | 3.889581 | 1.205531 | 0.227998 | NA |
| CG32643 | 0 | NA | NA | NA | NA | NA |
| CG2209 | 4.816197 | 1.713907 | 3.961293 | 0.432664 | 0.665259 | NA |
| CG15717 | 7.075951 | 0.348699 | 3.843569 | 0.090723 | 0.927713 | NA |
| MFS10 | 10.27892 | -5.98386 | 3.878613 | -1.54278 | 0.122884 | NA |
| CG15742 | 0.178053 | -1.16467 | 4.075067 | -0.2858 | 0.775028 | NA |
| CG32639 | 0 | NA | NA | NA | NA | NA |
| CG12723 | 0 | NA | NA | NA | NA | NA |
| CG12715 | 0 | NA | NA | NA | NA | NA |
| CG15719 | 2.493505 | -2.83145 | 3.945175 | -0.7177 | 0.472943 | NA |
| CG15744 | 27.08613 | -0.40611 | 3.741707 | -0.10854 | 0.91357 | NA |
| lncRNA:CR | 53.93676 | -0.8499 | 3.834632 | -0.22164 | 0.824595 | NA |
| dmrt11E | 0 | NA | NA | NA | NA | NA |
| CG15754 | 0 | NA | NA | NA | NA | NA |
| CG15756 | 0.356107 | -1.47099 | 4.037764 | -0.36431 | 0.715628 | NA |
| Cpr12A | 0.178053 | -1.16467 | 4.075067 | -0.2858 | 0.775028 | NA |
| CG11816 | 0 | NA | NA | NA | NA | NA |
| CG12725 | 0 | NA | NA | NA | NA | NA |
| CG32631 | 0.104623 | 0.639075 | 4.067674 | 0.157111 | 0.875158 | NA |
| CG32633 | 0 | NA | NA | NA | NA | NA |
| BthD | 0 | NA | NA | NA | NA | NA |
| Tango2 | 37.23489 | 0.382619 | 3.78468 | 0.101097 | 0.919474 | NA |
| CG11162 | 0 | NA | NA | NA | NA | NA |
| CG11158 | 6.9072 | -0.95341 | 3.846074 | -0.24789 | 0.804219 | NA |
| mus101 | 4.840354 | 2.903273 | 3.920944 | 0.740452 | 0.459025 | NA |

| | | | | | | |
|---|---|---|---|---|---|---|
| CG10996 | 0 | NA | NA | NA | NA | NA |
| CG10993 | 0.837599 | 1.568055 | 4.001801 | 0.391837 | 0.695178 | NA |
| Ndc80 | 0.178053 | -1.16467 | 4.075067 | -0.2858 | 0.775028 | NA |
| CG11068 | 37.42258 | -7.43411 | 3.913063 | -1.89982 | 0.057457 | NA |
| Ste12DOR | 0.341478 | 0.247543 | 4.031301 | 0.061405 | 0.951036 | NA |
| tRNA:Arg-T | 0 | NA | NA | NA | NA | NA |
| tRNA:Ser-( | 0 | NA | NA | NA | NA | NA |
| tRNA:Ser-( | 0 | NA | NA | NA | NA | NA |
| tRNA:Arg-T | 0 | NA | NA | NA | NA | NA |
| tRNA:Arg-T | 0 | NA | NA | NA | NA | NA |
| tRNA:Arg-T | 0 | NA | NA | NA | NA | NA |
| tRNA:Arg-T | 0 | NA | NA | NA | NA | NA |
| tRNA:Ser-/ | 0 | NA | NA | NA | NA | NA |
| tRNA:Ser-/ | 0 | NA | NA | NA | NA | NA |
| CG12480 | 0 | NA | NA | NA | NA | NA |
| CG12479 | 0 | NA | NA | NA | NA | NA |
| tRNA:Ser-/ | 0 | NA | NA | NA | NA | NA |
| tRNA:Arg-T | 0.209323 | 1.414133 | 4.032409 | 0.350692 | 0.725819 | NA |
| tRNA:Ser-( | 0 | NA | NA | NA | NA | NA |
| tRNA:Ser-/ | 0 | NA | NA | NA | NA | NA |
| tRNA:Ser-/ | 0 | NA | NA | NA | NA | NA |
| CG1368 | 0 | NA | NA | NA | NA | NA |
| CG11581 | 0 | NA | NA | NA | NA | NA |
| CG11584 | 0 | NA | NA | NA | NA | NA |
| CG32603 | 0 | NA | NA | NA | NA | NA |
| CG11585 | 0 | NA | NA | NA | NA | NA |
| yl | 0.356107 | -1.47099 | 4.037764 | -0.36431 | 0.715628 | NA |
| betaNACte | 16.24556 | -1.91474 | 3.953298 | -0.48434 | 0.628144 | NA |
| betaNACte | 0 | NA | NA | NA | NA | NA |
| CG9411 | 0.104623 | 0.639075 | 4.067674 | 0.157111 | 0.875158 | NA |
| betaNACte | 2.197088 | 1.483728 | 3.966465 | 0.374068 | 0.708354 | NA |
| betaNACte | 0 | NA | NA | NA | NA | NA |
| CG13403 | 0 | NA | NA | NA | NA | NA |
| CG12540 | 0 | NA | NA | NA | NA | NA |
| CG5334 | 2.386561 | -0.84224 | 3.87503 | -0.21735 | 0.827935 | NA |
| CG5347 | 0 | NA | NA | NA | NA | NA |
| Ag5r | 1.636199 | 1.416598 | 3.959479 | 0.357774 | 0.720512 | NA |
| Ag5r2 | 0 | NA | NA | NA | NA | NA |
| CG14410 | 0 | NA | NA | NA | NA | NA |
| dob | 50.366 | 3.758038 | 3.841771 | 0.978205 | 0.327973 | NA |
| CG33177 | 0 | NA | NA | NA | NA | NA |
| CG33178 | 0 | NA | NA | NA | NA | NA |
| CG9030 | 0 | NA | NA | NA | NA | NA |
| CG9503 | 22.52959 | -3.5953 | 3.920126 | -0.91714 | 0.35907 | NA |
| Eo | 0 | NA | NA | NA | NA | NA |
| CG14406 | 18.23161 | 2.284207 | 3.922196 | 0.582379 | 0.560311 | NA |
| CG9514 | 46.61855 | -1.92039 | 3.729988 | -0.51485 | 0.606656 | NA |
| CG9519 | 0.178053 | -1.16467 | 4.075067 | -0.2858 | 0.775028 | NA |
| CG9522 | 0 | NA | NA | NA | NA | NA |
| CG12539 | 0 | NA | NA | NA | NA | NA |

| | | | | | | |
|---|---|---|---|---|---|---|
| CG5662 | 0 | NA | NA | NA | NA | NA |
| dah | 46.2728 | -1.89472 | 3.812621 | -0.49696 | 0.619217 | NA |
| Pp1-13C | 3.004951 | 1.268861 | 3.962825 | 0.320191 | 0.748823 | NA |
| CG9114 | 0 | NA | NA | NA | NA | NA |
| CG9106 | 0.233093 | 0.692656 | 4.076246 | 0.169925 | 0.865069 | NA |
| PPYR1 | 0.932373 | 1.053996 | 3.996193 | 0.26375 | 0.791972 | NA |
| CG9101 | 0 | NA | NA | NA | NA | NA |
| CG32588 | 0 | NA | NA | NA | NA | NA |
| CG9095 | 2.255529 | -2.74072 | 3.879247 | -0.70651 | 0.479871 | NA |
| CG9072 | 0.104623 | 0.639075 | 4.067674 | 0.157111 | 0.875158 | NA |
| CG6294 | 0 | NA | NA | NA | NA | NA |
| CG15641 | 0 | NA | NA | NA | NA | NA |
| CG6324 | 0 | NA | NA | NA | NA | NA |
| Alp11 | 11.80161 | -2.25069 | 3.958868 | -0.56852 | 0.569683 | NA |
| CG8117 | 0 | NA | NA | NA | NA | NA |
| CG8119 | 0 | NA | NA | NA | NA | NA |
| CG12708 | 0 | NA | NA | NA | NA | NA |
| CG15599 | 0.178053 | -1.16467 | 4.075067 | -0.2858 | 0.775028 | NA |
| CG15646 | 0.178053 | -1.16467 | 4.075067 | -0.2858 | 0.775028 | NA |
| CG9213 | 0 | NA | NA | NA | NA | NA |
| CG15642 | 59.97658 | 1.515634 | 3.951499 | 0.383559 | 0.701305 | NA |
| CG8191 | 1.728708 | -4.08437 | 3.944501 | -1.03546 | 0.300454 | NA |
| CG33172 | 53.53591 | 0.647144 | 3.920233 | 0.165078 | 0.868883 | NA |
| lncRNA:CR | 2.96562 | -1.05157 | 3.948849 | -0.2663 | 0.790011 | NA |
| CG8565 | 0 | NA | NA | NA | NA | NA |
| Myb | 0 | NA | NA | NA | NA | NA |
| CG32581 | 53.63374 | 0.70988 | 3.926378 | 0.180798 | 0.856526 | NA |
| Efhc1.1 | 33.23222 | 5.611975 | 3.80839 | 1.473582 | 0.140594 | NA |
| CG8952 | 0 | NA | NA | NA | NA | NA |
| snRNA:U2: | 5.611999 | -3.46526 | 3.88618 | -0.89169 | 0.372559 | NA |
| UQCR-14 | 0 | NA | NA | NA | NA | NA |
| CG3679 | 6.996679 | 3.207923 | 3.880624 | 0.826651 | 0.408435 | NA |
| CG12507 | 0 | NA | NA | NA | NA | NA |
| snRNA:U5: | 0 | NA | NA | NA | NA | NA |
| mei-41 | 34.89398 | 2.711137 | 3.829482 | 0.707965 | 0.478967 | NA |
| CG4301 | 7.50187 | -0.36741 | 3.752704 | -0.0979 | 0.922008 | NA |
| Traf-like | 0 | NA | NA | NA | NA | NA |
| CG9981 | 0.209247 | 1.042804 | 4.047837 | 0.25762 | 0.7967 | NA |
| CG32573 | 0 | NA | NA | NA | NA | NA |
| CG4653 | 0 | NA | NA | NA | NA | NA |
| TwdlX | 1.191279 | 2.587225 | 3.944347 | 0.655932 | 0.511868 | NA |
| TwdlY | 1.157666 | 0.845933 | 3.944497 | 0.214459 | 0.830189 | NA |
| TwdlZ | 0.1047 | 0.613261 | 4.062799 | 0.150945 | 0.880019 | NA |
| CG13005 | 8.179791 | -1.22454 | 3.954419 | -0.30966 | 0.756817 | NA |
| ppk28 | 0 | NA | NA | NA | NA | NA |
| CG4872 | 29.67269 | 0.452298 | 3.836636 | 0.117889 | 0.906155 | NA |
| CG13002 | 0 | NA | NA | NA | NA | NA |
| CG4928 | 0.787578 | 1.071505 | 3.94115 | 0.271876 | 0.785717 | NA |
| CG33252 | 0 | NA | NA | NA | NA | NA |
| CG13004 | 14.33141 | -5.1087 | 3.872918 | -1.31908 | 0.187141 | NA |

| | | | | | | |
|---|---|---|---|---|---|---|
| CG9672 | 20.0718 | 0.317621 | 3.951304 | 0.080384 | 0.935932 | NA |
| CG9673 | 4.680909 | 0.306279 | 3.955251 | 0.077436 | 0.938277 | NA |
| sphe | 1.684167 | -1.89559 | 3.906369 | -0.48526 | 0.627495 | NA |
| CG9676 | 20.72232 | -1.00423 | 3.935683 | -0.25516 | 0.798598 | NA |
| CG13010 | 0 | NA | NA | NA | NA | NA |
| CG15865 | 0 | NA | NA | NA | NA | NA |
| CG4949 | 26.35188 | -1.70379 | 3.952363 | -0.43108 | 0.66641 | NA |
| CG4955 | 0 | NA | NA | NA | NA | NA |
| CG16700 | 0 | NA | NA | NA | NA | NA |
| CG4991 | 0.178053 | -1.16467 | 4.075067 | -0.2858 | 0.775028 | NA |
| CG5070 | 0 | NA | NA | NA | NA | NA |
| CG32564 | 0 | NA | NA | NA | NA | NA |
| CG32563 | 0 | NA | NA | NA | NA | NA |
| CG12998 | 0.1047 | 0.613261 | 4.062799 | 0.150945 | 0.880019 | NA |
| CG5172 | 0 | NA | NA | NA | NA | NA |
| CG10597 | 0 | NA | NA | NA | NA | NA |
| CG10598 | 0 | NA | NA | NA | NA | NA |
| CG12995 | 0.178812 | -1.08808 | 4.061735 | -0.26789 | 0.788788 | NA |
| TTLL1A | 0 | NA | NA | NA | NA | NA |
| Arpc3B | 47.66106 | 1.103248 | 3.951587 | 0.279191 | 0.780098 | NA |
| CG8945 | 0.396233 | -1.6814 | 4.028351 | -0.41739 | 0.676392 | NA |
| CG13000 | 0 | NA | NA | NA | NA | NA |
| wus | 0.1047 | 0.613261 | 4.062799 | 0.150945 | 0.880019 | NA |
| B-H2 | 0 | NA | NA | NA | NA | NA |
| B-H1 | 0.282677 | -0.09599 | 4.057934 | -0.02366 | 0.981127 | NA |
| CG8568 | 37.93296 | 1.263721 | 3.750677 | 0.336932 | 0.736169 | NA |
| p-cup | 0 | NA | NA | NA | NA | NA |
| CG12994 | 0 | NA | NA | NA | NA | NA |
| CG8661 | 0 | NA | NA | NA | NA | NA |
| CG8664 | 0 | NA | NA | NA | NA | NA |
| unc-4 | 30.24016 | 1.289554 | 3.867131 | 0.333465 | 0.738783 | NA |
| CG6398 | 1.046999 | 1.598136 | 3.993006 | 0.400234 | 0.688984 | NA |
| Mco4 | 0.356107 | -1.47099 | 4.037764 | -0.36431 | 0.715628 | NA |
| CG32554 | 2.229917 | -0.25931 | 3.961959 | -0.06545 | 0.947816 | NA |
| CG8142 | 41.67252 | 0.371839 | 3.835694 | 0.096942 | 0.922773 | NA |
| CG12986 | 0 | NA | NA | NA | NA | NA |
| CG12990 | 12.99789 | -1.69923 | 3.953847 | -0.42977 | 0.667366 | NA |
| CG12985 | 0 | NA | NA | NA | NA | NA |
| CG6788 | 0 | NA | NA | NA | NA | NA |
| Gss1 | 0.466186 | 0.959471 | 4.033986 | 0.237847 | 0.812 | NA |
| CR32496 | 0 | NA | NA | NA | NA | NA |
| CG15373 | 0 | NA | NA | NA | NA | NA |
| lncRNA:CR | 0.732899 | 1.548046 | 4.007786 | 0.38626 | 0.699304 | NA |
| Him | 0 | NA | NA | NA | NA | NA |
| lncRNA:CR | 0.989172 | 0.122741 | 3.950507 | 0.03107 | 0.975214 | NA |
| CG6106 | 2.544779 | -0.269 | 3.959075 | -0.06795 | 0.945829 | NA |
| CG15056 | 0 | NA | NA | NA | NA | NA |
| CG6290 | 0 | NA | NA | NA | NA | NA |
| CG32551 | 0.178053 | -1.16467 | 4.075067 | -0.2858 | 0.775028 | NA |
| CG32548 | 0 | NA | NA | NA | NA | NA |

| | | | | | | |
|---|---|---|---|---|---|---|
| upd1 | 0.178053 | -1.16467 | 4.075067 | -0.2858 | 0.775028 | NA |
| CG15059 | 0.356107 | -1.47099 | 4.037764 | -0.36431 | 0.715628 | NA |
| upd2 | 3.039809 | -2.17312 | 3.979294 | -0.54611 | 0.584992 | NA |
| Hesr | 0 | NA | NA | NA | NA | NA |
| Xrcc2 | 0 | NA | NA | NA | NA | NA |
| CG15047 | 0.1047 | 0.613261 | 4.062799 | 0.150945 | 0.880019 | NA |
| tRNA:Pro-( | 0 | NA | NA | NA | NA | NA |
| phm | 0 | NA | NA | NA | NA | NA |
| Cyp308a1 | 0.337793 | 1.451235 | 4.039844 | 0.35923 | 0.719423 | NA |
| CG12609 | 0.178053 | -1.16467 | 4.075067 | -0.2858 | 0.775028 | NA |
| CG6873 | 0 | NA | NA | NA | NA | NA |
| Cyp18a1 | 0.282753 | -0.25398 | 4.020204 | -0.06318 | 0.949626 | NA |
| CG6696 | 0.132078 | -1.13211 | 4.069918 | -0.27817 | 0.780885 | NA |
| CG15040 | 31.09191 | 0.655504 | 3.927741 | 0.166891 | 0.867456 | NA |
| CG15042 | 0 | NA | NA | NA | NA | NA |
| CG6470 | 0.314023 | 1.768229 | 4.015745 | 0.440324 | 0.659702 | NA |
| CG15044 | 0 | NA | NA | NA | NA | NA |
| CG15046 | 1.529126 | -1.18057 | 3.969053 | -0.29744 | 0.766127 | NA |
| psh | 0.1047 | 0.613261 | 4.062799 | 0.150945 | 0.880019 | NA |
| Hayan | 0 | NA | NA | NA | NA | NA |
| CR6900 | 47.63831 | 3.559934 | 3.903583 | 0.911966 | 0.361787 | NA |
| Pvf1 | 0.770937 | -1.65034 | 3.976655 | -0.41501 | 0.678137 | NA |
| SdhBL | 0.314023 | 1.768229 | 4.015745 | 0.440324 | 0.659702 | NA |
| CG14190 | 0 | NA | NA | NA | NA | NA |
| CG7406 | 0 | NA | NA | NA | NA | NA |
| CG14191 | 0 | NA | NA | NA | NA | NA |
| CG33253 | 0.282753 | -0.25398 | 4.020204 | -0.06318 | 0.949626 | NA |
| Mec2 | 0 | NA | NA | NA | NA | NA |
| HP1D3csd | 0 | NA | NA | NA | NA | NA |
| Mur18B | 0 | NA | NA | NA | NA | NA |
| Muc18B | 0 | NA | NA | NA | NA | NA |
| CG7884 | 0 | NA | NA | NA | NA | NA |
| CG8051 | 0.178053 | -1.16467 | 4.075067 | -0.2858 | 0.775028 | NA |
| CG8034 | 0.1047 | 0.613261 | 4.062799 | 0.150945 | 0.880019 | NA |
| CG8028 | 0 | NA | NA | NA | NA | NA |
| CG7992 | 0 | NA | NA | NA | NA | NA |
| CG14196 | 4.816197 | 1.713907 | 3.961293 | 0.432664 | 0.665259 | NA |
| CG14195 | 32.4569 | 2.969217 | 3.920388 | 0.757378 | 0.448823 | NA |
| CG7914 | 14.5227 | 0.316307 | 3.951755 | 0.080042 | 0.936204 | NA |
| Inx5 | 0 | NA | NA | NA | NA | NA |
| CG15882 | 0 | NA | NA | NA | NA | NA |
| CG7423 | 0 | NA | NA | NA | NA | NA |
| narya | 8.549286 | 4.495795 | 3.877208 | 1.159544 | 0.246234 | NA |
| CG14204 | 0.178053 | -1.16467 | 4.075067 | -0.2858 | 0.775028 | NA |
| CG14205 | 0.357625 | -1.47707 | 4.044794 | -0.36518 | 0.714979 | NA |
| CG14212 | 0 | NA | NA | NA | NA | NA |
| Zw | 4.167137 | -2.13584 | 3.890062 | -0.54905 | 0.582972 | NA |
| CG14227 | 53.87678 | -2.20788 | 3.952601 | -0.55859 | 0.576442 | NA |
| et | 27.08669 | 0.318515 | 3.951 | 0.080616 | 0.935747 | NA |
| CG14223 | 0.236778 | -0.26946 | 4.071573 | -0.06618 | 0.947234 | NA |

| | | | | | | |
|---|---|---|---|---|---|---|
| CG14221 | 0.368779 | -0.59727 | 3.999361 | -0.14934 | 0.881285 | NA |
| CG14219 | 0.523116 | 1.34877 | 4.007682 | 0.336546 | 0.736459 | NA |
| CG14218 | 0 | NA | NA | NA | NA | NA |
| CG12231 | 0 | NA | NA | NA | NA | NA |
| CG14230 | 61.28999 | 0.451195 | 3.846158 | 0.11731 | 0.906614 | NA |
| SkpD | 0 | NA | NA | NA | NA | NA |
| SkpC | 0 | NA | NA | NA | NA | NA |
| SkpE | 0 | NA | NA | NA | NA | NA |
| CG12702 | 0 | NA | NA | NA | NA | NA |
| meso18E | 0.578178 | 0.709574 | 3.962639 | 0.179066 | 0.857886 | NA |
| CG14185 | 0 | NA | NA | NA | NA | NA |
| CG6996 | 0.233093 | 0.692656 | 4.076246 | 0.169925 | 0.865069 | NA |
| CG17145 | 0 | NA | NA | NA | NA | NA |
| CG13813 | 9.843966 | -1.74148 | 3.755972 | -0.46366 | 0.642895 | NA |
| CG32228 | 3.734244 | -3.93648 | 3.895617 | -1.01049 | 0.31226 | NA |
| Spn77Bb | 0.178053 | -1.16467 | 4.075067 | -0.2858 | 0.775028 | NA |
| CR6434 | 1.296331 | 0.32833 | 3.890767 | 0.084387 | 0.932749 | NA |
| CG17122 | 0 | NA | NA | NA | NA | NA |
| CG7017 | 3.204959 | -1.67301 | 3.96255 | -0.42221 | 0.672875 | NA |
| asRNA:CR3 | 0 | NA | NA | NA | NA | NA |
| CG7290 | 0 | NA | NA | NA | NA | NA |
| CG7298 | 0 | NA | NA | NA | NA | NA |
| obst-F | 0 | NA | NA | NA | NA | NA |
| CG7335 | 0 | NA | NA | NA | NA | NA |
| obst-J | 0 | NA | NA | NA | NA | NA |
| CG7365 | 0 | NA | NA | NA | NA | NA |
| CG14187 | 53.67173 | 1.537006 | 3.951579 | 0.38896 | 0.697306 | NA |
| Ir76b | 0.104623 | 0.639075 | 4.067674 | 0.157111 | 0.875158 | NA |
| CG17732 | 0.620262 | -2.71754 | 3.999923 | -0.6794 | 0.496886 | NA |
| schuy | 44.46488 | 1.536494 | 3.951717 | 0.388817 | 0.697412 | NA |
| hale | 0.414754 | -1.14733 | 4.015515 | -0.28573 | 0.775088 | NA |
| CG14104 | 31.23582 | 0.74645 | 3.836369 | 0.194572 | 0.845728 | NA |
| Cpr76Bc | 0 | NA | NA | NA | NA | NA |
| Cpr76Bd | 0.1047 | 0.613261 | 4.062799 | 0.150945 | 0.880019 | NA |
| Rcd7 | 1.430498 | -2.08135 | 4.005348 | -0.51964 | 0.603313 | NA |
| CG9300 | 0.236778 | -0.26946 | 4.071573 | -0.06618 | 0.947234 | NA |
| ms(3)76Cc | 0 | NA | NA | NA | NA | NA |
| Gbeta76C | 3.893142 | 0.325078 | 3.852087 | 0.08439 | 0.932746 | NA |
| ms(3)76Ca | 0 | NA | NA | NA | NA | NA |
| lush | 0 | NA | NA | NA | NA | NA |
| CG9279 | 0 | NA | NA | NA | NA | NA |
| CG9666 | 0 | NA | NA | NA | NA | NA |
| CG14085 | 0.178053 | -1.16467 | 4.075067 | -0.2858 | 0.775028 | NA |
| CG14086 | 0.847327 | -2.62258 | 4.005321 | -0.65477 | 0.512614 | NA |
| ms(3)76Ba | 1.251686 | -2.06022 | 4.011654 | -0.51356 | 0.60756 | NA |
| CR33255 | 0 | NA | NA | NA | NA | NA |
| hpRNA:CR: | 4.411568 | -1.49887 | 3.862047 | -0.3881 | 0.69794 | NA |
| pip | 1.180677 | -1.30885 | 3.976465 | -0.32915 | 0.742044 | NA |
| CG14089 | 0 | NA | NA | NA | NA | NA |
| CG14088 | 0 | NA | NA | NA | NA | NA |

| | | | | | | |
|---|---|---|---|---|---|---|
| CG6836 | 35.04917 | 4.270749 | 3.883235 | 1.099791 | 0.271423 | NA |
| CG14075 | 0.679117 | 1.734471 | 3.96601 | 0.437334 | 0.661869 | NA |
| CG14079 | 0.178053 | -1.16467 | 4.075067 | -0.2858 | 0.775028 | NA |
| Spn75F | 0 | NA | NA | NA | NA | NA |
| Acp76A | 0.1047 | 0.613261 | 4.062799 | 0.150945 | 0.880019 | NA |
| SmydA-2 | 19.05246 | -2.92317 | 3.920759 | -0.74556 | 0.455931 | NA |
| CG3819 | 27.81959 | 0.318582 | 3.950977 | 0.080634 | 0.935733 | NA |
| CG6893 | 0 | NA | NA | NA | NA | NA |
| CG6885 | 0.414754 | -1.14733 | 4.015515 | -0.28573 | 0.775088 | NA |
| CG6865 | 0 | NA | NA | NA | NA | NA |
| NijB | 0.283512 | -0.2331 | 4.065094 | -0.05734 | 0.954273 | NA |
| CG6839 | 0.178053 | -1.16467 | 4.075067 | -0.2858 | 0.775028 | NA |
| CG18223 | 0 | NA | NA | NA | NA | NA |
| CG14077 | 0 | NA | NA | NA | NA | NA |
| Ir75d | 0.264155 | -1.54099 | 4.048976 | -0.38059 | 0.703509 | NA |
| Rad9 | 42.63364 | -1.88881 | 3.951801 | -0.47796 | 0.632678 | NA |
| CG12477 | 0.178053 | -1.16467 | 4.075067 | -0.2858 | 0.775028 | NA |
| Dic4 | 0 | NA | NA | NA | NA | NA |
| Cyp12c1 | 0 | NA | NA | NA | NA | NA |
| CG32198 | 0.233093 | 0.692656 | 4.076246 | 0.169925 | 0.865069 | NA |
| CG7271 | 0.2094 | 1.005034 | 4.045341 | 0.248442 | 0.803792 | NA |
| CG18233 | 0.1047 | 0.613261 | 4.062799 | 0.150945 | 0.880019 | NA |
| CG18231 | 0.264155 | -1.54099 | 4.048976 | -0.38059 | 0.703509 | NA |
| CG32201 | 0 | NA | NA | NA | NA | NA |
| CG32199 | 0.356107 | -1.47099 | 4.037764 | -0.36431 | 0.715628 | NA |
| tRNA:Pro-( | 0 | NA | NA | NA | NA | NA |
| term | 0.178812 | -1.08808 | 4.061735 | -0.26789 | 0.788788 | NA |
| Met75Ca | 0 | NA | NA | NA | NA | NA |
| Met75Cb | 0 | NA | NA | NA | NA | NA |
| CG4306 | 0 | NA | NA | NA | NA | NA |
| geko | 26.67969 | -1.918 | 3.952367 | -0.48528 | 0.627479 | NA |
| CG7330 | 6.207651 | -1.90163 | 3.957084 | -0.48056 | 0.630826 | NA |
| CG7320 | 0 | NA | NA | NA | NA | NA |
| grim | 6.347558 | -0.81697 | 3.860925 | -0.2116 | 0.832419 | NA |
| CG13700 | 0 | NA | NA | NA | NA | NA |
| CG5103 | 2.453201 | -4.08442 | 3.895709 | -1.04844 | 0.294436 | NA |
| CG13699 | 0.178053 | -1.16467 | 4.075067 | -0.2858 | 0.775028 | NA |
| Cyp312a1 | 0 | NA | NA | NA | NA | NA |
| lncRNA:CR | 0 | NA | NA | NA | NA | NA |
| CG7408 | 47.55102 | 1.177227 | 3.951898 | 0.297889 | 0.765788 | NA |
| CG7402 | 0 | NA | NA | NA | NA | NA |
| Tsp74F | 0 | NA | NA | NA | NA | NA |
| CG16775 | 0 | NA | NA | NA | NA | NA |
| CG5506 | 0 | NA | NA | NA | NA | NA |
| CG5567 | 49.36089 | -7.70823 | 3.843115 | -2.00572 | 0.044886 | NA |
| CG32189 | 0 | NA | NA | NA | NA | NA |
| CG32188 | 0 | NA | NA | NA | NA | NA |
| Ir75a | 0 | NA | NA | NA | NA | NA |
| Cln3 | 0 | NA | NA | NA | NA | NA |
| edin | 0 | NA | NA | NA | NA | NA |

| | | | | | | |
|---|---|---|---|---|---|---|
| CG7542 | 0 | NA | NA | NA | NA | NA |
| CG32182 | 0 | NA | NA | NA | NA | NA |
| CG32181 | 0 | NA | NA | NA | NA | NA |
| Adgf-A | 2.346336 | -0.97604 | 3.932042 | -0.24823 | 0.803959 | NA |
| Adgf-A2 | 1.557478 | -1.35228 | 3.96773 | -0.34082 | 0.733239 | NA |
| Adgf-B | 0.178053 | -1.16467 | 4.075067 | -0.2858 | 0.775028 | NA |
| CG6034 | 0 | NA | NA | NA | NA | NA |
| CG6052 | 14.13449 | 1.740196 | 3.954547 | 0.440049 | 0.659901 | NA |
| Jon74E | 28.17852 | -3.51055 | 3.901177 | -0.89987 | 0.36819 | NA |
| CG7707 | 0 | NA | NA | NA | NA | NA |
| CG13723 | 0 | NA | NA | NA | NA | NA |
| CG13724 | 0.178812 | -1.08808 | 4.061735 | -0.26789 | 0.788788 | NA |
| tap | 6.386696 | 1.723466 | 3.95882 | 0.435348 | 0.66331 | NA |
| SecCl | 0 | NA | NA | NA | NA | NA |
| qjt | 1.609311 | -2.0987 | 4.000255 | -0.52464 | 0.599832 | NA |
| CG6333 | 0.178053 | -1.16467 | 4.075067 | -0.2858 | 0.775028 | NA |
| CG13733 | 0 | NA | NA | NA | NA | NA |
| CG12229 | 1.387557 | 0.928325 | 3.968008 | 0.233952 | 0.815022 | NA |
| Coq4 | 0.418493 | 1.313669 | 4.019283 | 0.326842 | 0.743788 | NA |
| frc | 0 | NA | NA | NA | NA | NA |
| blot | 15.78941 | -2.00358 | 3.760538 | -0.53279 | 0.594178 | NA |
| CG13731 | 0 | NA | NA | NA | NA | NA |
| CG13728 | 0.462324 | -0.59023 | 4.044233 | -0.14594 | 0.883966 | NA |
| Cad74A | 3.428902 | -0.09856 | 3.948141 | -0.02496 | 0.980084 | NA |
| brv2 | 0.387453 | 0.098632 | 4.002797 | 0.024641 | 0.980342 | NA |
| CG13727 | 0.178053 | -1.16467 | 4.075067 | -0.2858 | 0.775028 | NA |
| tRNA:Pro-( | 0 | NA | NA | NA | NA | NA |
| Or74a | 0.1047 | 0.613261 | 4.062799 | 0.150945 | 0.880019 | NA |
| CG13725 | 0.356107 | -1.47099 | 4.037764 | -0.36431 | 0.715628 | NA |
| ND-24L | 0 | NA | NA | NA | NA | NA |
| CG6497 | 2.895532 | 1.407909 | 3.855446 | 0.365174 | 0.714981 | NA |
| CG32170 | 0.469871 | 0.864471 | 4.023538 | 0.214854 | 0.829881 | NA |
| CG13028 | 0 | NA | NA | NA | NA | NA |
| Fit2 | 50.34066 | 2.476387 | 3.842284 | 0.644509 | 0.519245 | NA |
| CG7724 | 4.292697 | 1.709267 | 3.962502 | 0.431361 | 0.666206 | NA |
| Ilp8 | 0 | NA | NA | NA | NA | NA |
| CG6652 | 0.356107 | -1.47099 | 4.037764 | -0.36431 | 0.715628 | NA |
| CG13022 | 0 | NA | NA | NA | NA | NA |
| CG11905 | 0.1047 | 0.613261 | 4.062799 | 0.150945 | 0.880019 | NA |
| CG13023 | 0 | NA | NA | NA | NA | NA |
| snRNA:U1: | 0 | NA | NA | NA | NA | NA |
| CG13032 | 1.945653 | -0.39222 | 3.885108 | -0.10095 | 0.919587 | NA |
| CG9692 | 0 | NA | NA | NA | NA | NA |
| CG13031 | 0 | NA | NA | NA | NA | NA |
| PGRP-SB2 | 0 | NA | NA | NA | NA | NA |
| CG9701 | 0 | NA | NA | NA | NA | NA |
| CG32161 | 0 | NA | NA | NA | NA | NA |
| CG13024 | 0.178053 | -1.16467 | 4.075067 | -0.2858 | 0.775028 | NA |
| Dbp73D | 25.62709 | -7.41624 | 3.907554 | -1.89792 | 0.057706 | NA |
| PGRP-SB1 | 1.958586 | -1.65227 | 3.969602 | -0.41623 | 0.677241 | NA |

| | | | | | | |
|---|---|---|---|---|---|---|
| CG13026 | 0 | NA | NA | NA | NA | NA |
| Lasp | 0 | NA | NA | NA | NA | NA |
| Galphaf | 26.59378 | 1.746911 | 3.952855 | 0.441936 | 0.658535 | NA |
| CG4098 | 50.45739 | 0.169876 | 3.501337 | 0.048517 | 0.961304 | NA |
| spd-2 | 0.411147 | -0.29058 | 4.046973 | -0.0718 | 0.94276 | NA |
| CG13063 | 0 | NA | NA | NA | NA | NA |
| lncRNA:CR | 0 | NA | NA | NA | NA | NA |
| CG13060 | 0 | NA | NA | NA | NA | NA |
| CG13058 | 0 | NA | NA | NA | NA | NA |
| CG33061 | 0 | NA | NA | NA | NA | NA |
| CG33060 | 0 | NA | NA | NA | NA | NA |
| retinin | 43.13629 | 2.972992 | 3.919545 | 0.758504 | 0.448149 | NA |
| CG13056 | 39.75538 | -1.91967 | 3.951888 | -0.48576 | 0.627137 | NA |
| CG4998 | 0 | NA | NA | NA | NA | NA |
| CG33257 | 0 | NA | NA | NA | NA | NA |
| tRNA:Met- | 0 | NA | NA | NA | NA | NA |
| Cpr72Ea | 0 | NA | NA | NA | NA | NA |
| Cpr72Eb | 0 | NA | NA | NA | NA | NA |
| CG4842 | 0 | NA | NA | NA | NA | NA |
| CG18814 | 0 | NA | NA | NA | NA | NA |
| Gagr | 0 | NA | NA | NA | NA | NA |
| st | 50.10738 | -0.01706 | 3.262872 | -0.00523 | 0.995828 | NA |
| CG4229 | 12.99789 | -1.69923 | 3.953847 | -0.42977 | 0.667366 | NA |
| Mipp1 | 0.732899 | 1.548046 | 4.007786 | 0.38626 | 0.699304 | NA |
| CG13033 | 0.132078 | -1.13211 | 4.069918 | -0.27817 | 0.780885 | NA |
| aos | 0.178053 | -1.16467 | 4.075067 | -0.2858 | 0.775028 | NA |
| GluRS-m | 0.1047 | 0.613261 | 4.062799 | 0.150945 | 0.880019 | NA |
| Agpat4 | 0 | NA | NA | NA | NA | NA |
| Cpr72Ec | 0 | NA | NA | NA | NA | NA |
| CG13040 | 0 | NA | NA | NA | NA | NA |
| CG13041 | 0 | NA | NA | NA | NA | NA |
| CG13042 | 0.178053 | -1.16467 | 4.075067 | -0.2858 | 0.775028 | NA |
| CG13043 | 0 | NA | NA | NA | NA | NA |
| CG13075 | 9.715876 | 0.790869 | 3.749971 | 0.2109 | 0.832965 | NA |
| ATPsynbet | 0 | NA | NA | NA | NA | NA |
| tRNA:Met- | 0 | NA | NA | NA | NA | NA |
| CG13071 | 0 | NA | NA | NA | NA | NA |
| CG13070 | 0 | NA | NA | NA | NA | NA |
| CG13069 | 0.178053 | -1.16467 | 4.075067 | -0.2858 | 0.775028 | NA |
| CG13068 | 0.178053 | -1.16467 | 4.075067 | -0.2858 | 0.775028 | NA |
| CG13067 | 0 | NA | NA | NA | NA | NA |
| CG13066 | 0 | NA | NA | NA | NA | NA |
| CG13065 | 0 | NA | NA | NA | NA | NA |
| CG4982 | 0 | NA | NA | NA | NA | NA |
| CG13045 | 0 | NA | NA | NA | NA | NA |
| CG13046 | 0 | NA | NA | NA | NA | NA |
| CG13047 | 0.178812 | -1.08808 | 4.061735 | -0.26789 | 0.788788 | NA |
| CG13048 | 0 | NA | NA | NA | NA | NA |
| CG13049 | 0.178053 | -1.16467 | 4.075067 | -0.2858 | 0.775028 | NA |
| CG13050 | 0 | NA | NA | NA | NA | NA |

| | | | | | | |
|---|---|---|---|---|---|---|
| CG4950 | 1.228742 | -0.20401 | 3.923681 | -0.052 | 0.958532 | NA |
| CG13053 | 37.03508 | -1.70508 | 3.951944 | -0.43145 | 0.666139 | NA |
| CG13054 | 0 | NA | NA | NA | NA | NA |
| CG13055 | 0.597612 | 0.189345 | 3.999077 | 0.047347 | 0.962237 | NA |
| CG13073 | 0 | NA | NA | NA | NA | NA |
| CG32152 | 0 | NA | NA | NA | NA | NA |
| CG5157 | 0 | NA | NA | NA | NA | NA |
| CG13074 | 2.124287 | -3.00082 | 3.903937 | -0.76867 | 0.442092 | NA |
| CG5235 | 12.66869 | 1.738562 | 3.954961 | 0.43959 | 0.660234 | NA |
| CG6244 | 23.32498 | -1.70323 | 3.952551 | -0.43092 | 0.666526 | NA |
| CG13445 | 0 | NA | NA | NA | NA | NA |
| CG17028 | 51.25665 | 0.217308 | 3.834572 | 0.056671 | 0.954807 | NA |
| meru | 0.598264 | -0.35879 | 4.007615 | -0.08953 | 0.928662 | NA |
| CG7276 | 0 | NA | NA | NA | NA | NA |
| yellow-k | 0 | NA | NA | NA | NA | NA |
| Eig71Eb | 0.178053 | -1.16467 | 4.075067 | -0.2858 | 0.775028 | NA |
| Eig71Ed | 0.104623 | 0.639075 | 4.067674 | 0.157111 | 0.875158 | NA |
| Eig71Eg | 0 | NA | NA | NA | NA | NA |
| Eig71Ei | 0 | NA | NA | NA | NA | NA |
| Eig71Ek | 0 | NA | NA | NA | NA | NA |
| CG7304 | 0 | NA | NA | NA | NA | NA |
| Pgant8 | 0 | NA | NA | NA | NA | NA |
| comm2 | 0 | NA | NA | NA | NA | NA |
| CG7579 | 0 | NA | NA | NA | NA | NA |
| Eig71Ej | 0 | NA | NA | NA | NA | NA |
| Eig71Eh | 0 | NA | NA | NA | NA | NA |
| Eig71Ef | 0 | NA | NA | NA | NA | NA |
| Eig71Ee | 0 | NA | NA | NA | NA | NA |
| Eig71Ec | 0.178053 | -1.16467 | 4.075067 | -0.2858 | 0.775028 | NA |
| Eig71Ea | 0 | NA | NA | NA | NA | NA |
| CG13449 | 0.233093 | 0.692656 | 4.076246 | 0.169925 | 0.865069 | NA |
| mex1 | 0 | NA | NA | NA | NA | NA |
| CG13454 | 0 | NA | NA | NA | NA | NA |
| Best4 | 0 | NA | NA | NA | NA | NA |
| CG33259 | 0.233093 | 0.692656 | 4.076246 | 0.169925 | 0.865069 | NA |
| Ran-like | 0 | NA | NA | NA | NA | NA |
| cocoon | 0 | NA | NA | NA | NA | NA |
| Best3 | 0 | NA | NA | NA | NA | NA |
| Tom | 0 | NA | NA | NA | NA | NA |
| Brd | 0 | NA | NA | NA | NA | NA |
| Ocho | 0 | NA | NA | NA | NA | NA |
| CG3349 | 0.461565 | -1.07936 | 4.002266 | -0.26969 | 0.7874 | NA |
| CG32148 | 0 | NA | NA | NA | NA | NA |
| EAChm | 0 | NA | NA | NA | NA | NA |
| CG32847 | 0 | NA | NA | NA | NA | NA |
| CG6888 | 0 | NA | NA | NA | NA | NA |
| CG13457 | 0.71449 | -2.47883 | 4.014121 | -0.61753 | 0.536887 | NA |
| CG16959 | 0 | NA | NA | NA | NA | NA |
| CG13460 | 0 | NA | NA | NA | NA | NA |
| CG18649 | 0 | NA | NA | NA | NA | NA |

| | | | | | | |
|---|---|---|---|---|---|---|
| CG13461 | 0 | NA | NA | NA | NA | NA |
| CG12310 | 0 | NA | NA | NA | NA | NA |
| CG18581 | 0 | NA | NA | NA | NA | NA |
| ind | 0.283512 | -0.2331 | 4.065094 | -0.05734 | 0.954273 | NA |
| Rpn12R | 0 | NA | NA | NA | NA | NA |
| Reck | 14.18074 | -6.2064 | 3.89028 | -1.59536 | 0.110632 | NA |
| BobA | 0 | NA | NA | NA | NA | NA |
| CG13465 | 0 | NA | NA | NA | NA | NA |
| shd | 0 | NA | NA | NA | NA | NA |
| dlp | 52.72043 | 0.619272 | 3.841596 | 0.161202 | 0.871934 | NA |
| tRNA:iMet | 0 | NA | NA | NA | NA | NA |
| CG5048 | 0 | NA | NA | NA | NA | NA |
| goddard | 0 | NA | NA | NA | NA | NA |
| CG13476 | 0 | NA | NA | NA | NA | NA |
| CG13473 | 0 | NA | NA | NA | NA | NA |
| gnu | 0 | NA | NA | NA | NA | NA |
| Magt4b | 0.536437 | -1.71175 | 4.038042 | -0.42391 | 0.671635 | NA |
| CG13471 | 0 | NA | NA | NA | NA | NA |
| CG17177 | 0 | NA | NA | NA | NA | NA |
| CG13474 | 0 | NA | NA | NA | NA | NA |
| CG4914 | 0 | NA | NA | NA | NA | NA |
| tRNA:iMet | 0 | NA | NA | NA | NA | NA |
| CG9592 | 0 | NA | NA | NA | NA | NA |
| HGTX | 29.13665 | 1.175374 | 3.952522 | 0.297373 | 0.766182 | NA |
| Lk | 0 | NA | NA | NA | NA | NA |
| CG4613 | 0 | NA | NA | NA | NA | NA |
| CG3868 | 0 | NA | NA | NA | NA | NA |
| CG13481 | 0 | NA | NA | NA | NA | NA |
| saturn | 0 | NA | NA | NA | NA | NA |
| CG7924 | 0.356107 | -1.47099 | 4.037764 | -0.36431 | 0.715628 | NA |
| CG7906 | 0 | NA | NA | NA | NA | NA |
| CG13482 | 5.413809 | 0.308289 | 3.954543 | 0.077958 | 0.937861 | NA |
| Fbp1 | 1.487119 | -0.47654 | 3.965238 | -0.12018 | 0.904342 | NA |
| Tsp68C | 0 | NA | NA | NA | NA | NA |
| Hml | 34.03076 | -6.55315 | 3.869613 | -1.69349 | 0.090362 | NA |
| CG13737 | 41.57107 | -0.41604 | 3.941747 | -0.10555 | 0.915941 | NA |
| ssp2 | 0 | NA | NA | NA | NA | NA |
| Hsc70-1 | 0.712213 | -1.57119 | 3.998613 | -0.39293 | 0.694369 | NA |
| CG17362 | 0 | NA | NA | NA | NA | NA |
| CG9040 | 0 | NA | NA | NA | NA | NA |
| CG6661 | 0 | NA | NA | NA | NA | NA |
| Pex1 | 48.02639 | -0.87637 | 3.948549 | -0.22195 | 0.824354 | NA |
| CG8100 | 0 | NA | NA | NA | NA | NA |
| CG13738 | 1.154422 | -3.36701 | 3.968232 | -0.84849 | 0.396164 | NA |
| CG8750 | 0 | NA | NA | NA | NA | NA |
| CG8757 | 6.161573 | 0.633982 | 3.820035 | 0.165962 | 0.868186 | NA |
| CG10738 | 37.18266 | -6.09488 | 3.823513 | -1.59405 | 0.110924 | NA |
| CG10710 | 3.853845 | -0.30875 | 3.956334 | -0.07804 | 0.937796 | NA |
| CG10116 | 0.104623 | 0.639075 | 4.067674 | 0.157111 | 0.875158 | NA |
| CG33262 | 0 | NA | NA | NA | NA | NA |

| | | | | | | |
|---|---|---|---|---|---|---|
| tRNA:Pro-/ | 0 | NA | NA | NA | NA | NA |
| tRNA:Pro-/ | 0 | NA | NA | NA | NA | NA |
| tRNA:Pro-/ | 0 | NA | NA | NA | NA | NA |
| tRNA:Leu-( | 0 | NA | NA | NA | NA | NA |
| tRNA:Asp-( | 0 | NA | NA | NA | NA | NA |
| SP | 6.759704 | 1.159951 | 3.957776 | 0.293082 | 0.76946 | NA |
| CG14113 | 0 | NA | NA | NA | NA | NA |
| CG17687 | 0.178053 | -1.16467 | 4.075067 | -0.2858 | 0.775028 | NA |
| SNCF | 0 | NA | NA | NA | NA | NA |
| CG14111 | 0 | NA | NA | NA | NA | NA |
| ImpL1 | 0 | NA | NA | NA | NA | NA |
| CG14110 | 0.233093 | 0.692656 | 4.076246 | 0.169925 | 0.865069 | NA |
| tRNA:Val-/ | 0.178812 | -1.08808 | 4.061735 | -0.26789 | 0.788788 | NA |
| CG14109 | 0 | NA | NA | NA | NA | NA |
| CG10140 | 0.178053 | -1.16467 | 4.075067 | -0.2858 | 0.775028 | NA |
| CG10154 | 0 | NA | NA | NA | NA | NA |
| tRNA:Val-/ | 0 | NA | NA | NA | NA | NA |
| CG14105 | 2.121376 | 0.634749 | 3.940869 | 0.161068 | 0.87204 | NA |
| CG14106 | 0.356107 | -1.47099 | 4.037764 | -0.36431 | 0.715628 | NA |
| CG33263 | 0 | NA | NA | NA | NA | NA |
| CG32121 | 57.86031 | -2.25893 | 3.898804 | -0.57939 | 0.562325 | NA |
| CG10222 | 33.40808 | 1.599386 | 3.815985 | 0.419128 | 0.675123 | NA |
| sens | 0.660388 | -1.75977 | 4.001266 | -0.4398 | 0.660079 | NA |
| Poc1 | 0.178053 | -1.16467 | 4.075067 | -0.2858 | 0.775028 | NA |
| CG14107 | 0 | NA | NA | NA | NA | NA |
| CG32119 | 0 | NA | NA | NA | NA | NA |
| CG14120 | 0.567024 | -0.28781 | 3.989037 | -0.07215 | 0.942482 | NA |
| CG14118 | 0.178053 | -1.16467 | 4.075067 | -0.2858 | 0.775028 | NA |
| CG10749 | 0 | NA | NA | NA | NA | NA |
| CG11262 | 0 | NA | NA | NA | NA | NA |
| CR14117 | 7.611441 | 3.315511 | 3.918634 | 0.846088 | 0.397503 | NA |
| ste14 | 7.884683 | 0.438792 | 3.857335 | 0.113755 | 0.909432 | NA |
| CG11251 | 0 | NA | NA | NA | NA | NA |
| CG17300 | 0 | NA | NA | NA | NA | NA |
| CG14115 | 0.104623 | 0.639075 | 4.067674 | 0.157111 | 0.875158 | NA |
| Cul6 | 0 | NA | NA | NA | NA | NA |
| CG10748 | 0.955237 | -0.24643 | 3.926835 | -0.06275 | 0.949962 | NA |
| Or69a | 0 | NA | NA | NA | NA | NA |
| CG10752 | 0 | NA | NA | NA | NA | NA |
| CG32115 | 12.87809 | 1.738817 | 3.954896 | 0.439662 | 0.660182 | NA |
| CG32117 | 0 | NA | NA | NA | NA | NA |
| CG12520 | 0 | NA | NA | NA | NA | NA |
| lncRNA:CR | 25.07789 | 0.224222 | 3.836219 | 0.058449 | 0.953391 | NA |
| CG17666 | 0.178053 | -1.16467 | 4.075067 | -0.2858 | 0.775028 | NA |
| CG32110 | 0 | NA | NA | NA | NA | NA |
| CG10943 | 0.1047 | 0.613261 | 4.062799 | 0.150945 | 0.880019 | NA |
| CG32107 | 3.306991 | 0.324254 | 3.865979 | 0.083874 | 0.933157 | NA |
| CG10969 | 0 | NA | NA | NA | NA | NA |
| CG32106 | 0 | NA | NA | NA | NA | NA |
| Obp69a | 0.104623 | 0.639075 | 4.067674 | 0.157111 | 0.875158 | NA |

| | | | | | | |
|---|---|---|---|---|---|---|
| CG10657 | 0 | NA | NA | NA | NA | NA |
| CG10663 | 1.256399 | 1.619735 | 3.986836 | 0.406271 | 0.684544 | NA |
| CG32102 | 0 | NA | NA | NA | NA | NA |
| toe | 17.06609 | 1.74264 | 3.95393 | 0.440736 | 0.659404 | NA |
| Smyd4-2 | 0 | NA | NA | NA | NA | NA |
| CG17154 | 0 | NA | NA | NA | NA | NA |
| DEF8 | 44.16891 | 2.265868 | 3.820182 | 0.593131 | 0.553093 | NA |
| CAH2 | 0.282753 | -0.25398 | 4.020204 | -0.06318 | 0.949626 | NA |
| Est-P | 0 | NA | NA | NA | NA | NA |
| CG11529 | 6.86804 | -1.90364 | 3.956501 | -0.48114 | 0.630416 | NA |
| Ir68b | 0.233093 | 0.692656 | 4.076246 | 0.169925 | 0.865069 | NA |
| CG10907 | 61.1854 | -4.04898 | 3.863031 | -1.04813 | 0.294576 | NA |
| CG7264 | 2.560185 | 3.612111 | 3.952886 | 0.913791 | 0.360827 | NA |
| Sprn | 0.565506 | -0.26224 | 3.986711 | -0.06578 | 0.947554 | NA |
| CCDC151 | 0 | NA | NA | NA | NA | NA |
| CG11588 | 0.31089 | -1.9003 | 4.033427 | -0.47114 | 0.637542 | NA |
| Rpt4R | 0 | NA | NA | NA | NA | NA |
| CG7252 | 0.178053 | -1.16467 | 4.075067 | -0.2858 | 0.775028 | NA |
| CG17826 | 0 | NA | NA | NA | NA | NA |
| CG7248 | 0 | NA | NA | NA | NA | NA |
| CG14125 | 0 | NA | NA | NA | NA | NA |
| CG6938 | 0.1047 | 0.613261 | 4.062799 | 0.150945 | 0.880019 | NA |
| CG6931 | 0.104623 | 0.639075 | 4.067674 | 0.157111 | 0.875158 | NA |
| Muc68E | 0.104623 | 0.639075 | 4.067674 | 0.157111 | 0.875158 | NA |
| prc | 1.246373 | -1.6238 | 3.979526 | -0.40804 | 0.683245 | NA |
| CG11570 | 0 | NA | NA | NA | NA | NA |
| obst-G | 0.178053 | -1.16467 | 4.075067 | -0.2858 | 0.775028 | NA |
| CG5883 | 0 | NA | NA | NA | NA | NA |
| SdhAL | 0 | NA | NA | NA | NA | NA |
| CG12521 | 0 | NA | NA | NA | NA | NA |
| tRNA:Gln-ˉ | 0 | NA | NA | NA | NA | NA |
| CG5964 | 0 | NA | NA | NA | NA | NA |
| Tim13 | 0 | NA | NA | NA | NA | NA |
| Vha16-3 | 0 | NA | NA | NA | NA | NA |
| Vha16-2 | 0 | NA | NA | NA | NA | NA |
| CG33500 | 0 | NA | NA | NA | NA | NA |
| Sgs7 | 0 | NA | NA | NA | NA | NA |
| Sgs3 | 0.178053 | -1.16467 | 4.075067 | -0.2858 | 0.775028 | NA |
| CG7512 | 0 | NA | NA | NA | NA | NA |
| CG7377 | 0 | NA | NA | NA | NA | NA |
| CG32087 | 0.357625 | -1.47707 | 4.044794 | -0.36518 | 0.714979 | NA |
| Gr68a | 0 | NA | NA | NA | NA | NA |
| CG14132 | 1.452855 | -1.83946 | 3.975716 | -0.46267 | 0.643599 | NA |
| crim | 5.549096 | 1.719007 | 3.959971 | 0.434096 | 0.664219 | NA |
| Dnai2 | 0 | NA | NA | NA | NA | NA |
| Pldn | 54.52358 | -1.63077 | 3.835842 | -0.42514 | 0.670734 | NA |
| CG6071 | 3.830253 | -1.88895 | 3.960794 | -0.47691 | 0.633424 | NA |
| Muc68Ca | 1.743906 | 0.331025 | 3.909891 | 0.084664 | 0.932529 | NA |
| CG14137 | 0.209247 | 1.042804 | 4.047837 | 0.25762 | 0.7967 | NA |
| CG33268 | 0.104623 | 0.639075 | 4.067674 | 0.157111 | 0.875158 | NA |

| | | | | | | |
|---|---|---|---|---|---|---|
| CG32086 | 0 | NA | NA | NA | NA | NA |
| CG33269 | 2.796185 | -0.43502 | 3.958512 | -0.10989 | 0.912493 | NA |
| CG33270 | 30.43823 | 0.793074 | 3.951793 | 0.200687 | 0.840943 | NA |
| CG33271 | 0.1047 | 0.613261 | 4.062799 | 0.150945 | 0.880019 | NA |
| CG33272 | 0.2094 | 1.005034 | 4.045341 | 0.248442 | 0.803792 | NA |
| Sgs8 | 0 | NA | NA | NA | NA | NA |
| CG7573 | 0.233093 | 0.692656 | 4.076246 | 0.169925 | 0.865069 | NA |
| CG11726 | 0.178053 | -1.16467 | 4.075067 | -0.2858 | 0.775028 | NA |
| CG7560 | 0 | NA | NA | NA | NA | NA |
| CG7557 | 0 | NA | NA | NA | NA | NA |
| CG7551 | 0 | NA | NA | NA | NA | NA |
| Fum3 | 0.283436 | -0.20033 | 4.070792 | -0.04921 | 0.960751 | NA |
| CG12289 | 0 | NA | NA | NA | NA | NA |
| CG6149 | 0 | NA | NA | NA | NA | NA |
| CG6168 | 0 | NA | NA | NA | NA | NA |
| Cpr67Fa1 | 0 | NA | NA | NA | NA | NA |
| Cpr67Fa2 | 0 | NA | NA | NA | NA | NA |
| Cpr67Fb | 0 | NA | NA | NA | NA | NA |
| CG14147 | 0 | NA | NA | NA | NA | NA |
| Fad2 | 0 | NA | NA | NA | NA | NA |
| CG32079 | 0 | NA | NA | NA | NA | NA |
| CG32081 | 0 | NA | NA | NA | NA | NA |
| SuUR | 0 | NA | NA | NA | NA | NA |
| CG12522 | 0 | NA | NA | NA | NA | NA |
| nol | 27.35154 | 0.351828 | 3.838529 | 0.091657 | 0.926971 | NA |
| CG14143 | 0 | NA | NA | NA | NA | NA |
| IRSp53 | 10.45349 | -2.27299 | 3.895083 | -0.58355 | 0.559521 | NA |
| CG6216 | 0 | NA | NA | NA | NA | NA |
| CG32074 | 0 | NA | NA | NA | NA | NA |
| CG32073 | 0 | NA | NA | NA | NA | NA |
| CG32071 | 0 | NA | NA | NA | NA | NA |
| Elo68beta | 12.51686 | -2.25242 | 3.958427 | -0.56902 | 0.569342 | NA |
| klu | 2.315095 | -3.77244 | 3.930617 | -0.95976 | 0.337177 | NA |
| CG18628 | 37.37177 | -2.27194 | 3.953479 | -0.57467 | 0.565515 | NA |
| CG14154 | 0.915764 | -1.24472 | 3.920815 | -0.31746 | 0.750892 | NA |
| CG12523 | 0 | NA | NA | NA | NA | NA |
| CG6409 | 50.3216 | -1.92039 | 3.951682 | -0.48597 | 0.62699 | NA |
| CG14151 | 0 | NA | NA | NA | NA | NA |
| CG12362 | 0 | NA | NA | NA | NA | NA |
| CG32061 | 0 | NA | NA | NA | NA | NA |
| S-Lap3 | 0.282753 | -0.25398 | 4.020204 | -0.06318 | 0.949626 | NA |
| S-Lap4 | 1.17302 | -1.0781 | 3.973292 | -0.27134 | 0.786132 | NA |
| Gem3 | 0.209323 | 1.414133 | 4.032409 | 0.350692 | 0.725819 | NA |
| Ir67c | 0 | NA | NA | NA | NA | NA |
| Ir67b | 0 | NA | NA | NA | NA | NA |
| CG8072 | 39.88746 | -1.91968 | 3.951885 | -0.48576 | 0.627135 | NA |
| CG8065 | 0 | NA | NA | NA | NA | NA |
| Or67d | 0.104623 | 0.639075 | 4.067674 | 0.157111 | 0.875158 | NA |
| can | 20.70922 | -0.66171 | 3.927095 | -0.1685 | 0.866191 | NA |
| Or67c | 0.209247 | 1.042804 | 4.047837 | 0.25762 | 0.7967 | NA |

| | | | | | | |
|---|---|---|---|---|---|---|
| ect | 0 | NA | NA | NA | NA | NA |
| CG6628 | 0.5892 | -0.44935 | 3.992199 | -0.11256 | 0.910381 | NA |
| CG32054 | 0 | NA | NA | NA | NA | NA |
| CG32053 | 0 | NA | NA | NA | NA | NA |
| CG14160 | 0.660388 | -1.75977 | 4.001266 | -0.4398 | 0.660079 | NA |
| Ilp1 | 0 | NA | NA | NA | NA | NA |
| Ilp2 | 1.150856 | 1.439869 | 3.979295 | 0.36184 | 0.717471 | NA |
| Hez | 0 | NA | NA | NA | NA | NA |
| CG6709 | 0 | NA | NA | NA | NA | NA |
| Ilp4 | 3.90674 | 1.303404 | 3.85744 | 0.337893 | 0.735443 | NA |
| Ilp3 | 0.178053 | -1.16467 | 4.075067 | -0.2858 | 0.775028 | NA |
| Or67a | 56.01438 | 2.975657 | 3.918953 | 0.759299 | 0.447674 | NA |
| CG3222 | 0 | NA | NA | NA | NA | NA |
| CG3306 | 0 | NA | NA | NA | NA | NA |
| CG8329 | 0 | NA | NA | NA | NA | NA |
| CG18179 | 0 | NA | NA | NA | NA | NA |
| CG18180 | 0.178053 | -1.16467 | 4.075067 | -0.2858 | 0.775028 | NA |
| CG16719 | 0 | NA | NA | NA | NA | NA |
| CG6749 | 39.33834 | 1.536107 | 3.951823 | 0.388708 | 0.697492 | NA |
| Or67b | 0.209247 | 1.042804 | 4.047837 | 0.25762 | 0.7967 | NA |
| CG3088 | 0 | NA | NA | NA | NA | NA |
| Ir67a | 0.178053 | -1.16467 | 4.075067 | -0.2858 | 0.775028 | NA |
| CG4476 | 9.102223 | 1.525146 | 3.954825 | 0.385642 | 0.699762 | NA |
| CG4477 | 0 | NA | NA | NA | NA | NA |
| CG32040 | 16.62954 | -2.25959 | 3.956611 | -0.57109 | 0.567936 | NA |
| CG4461 | 0 | NA | NA | NA | NA | NA |
| CG32037 | 1.06832 | -1.61145 | 3.983918 | -0.40449 | 0.685853 | NA |
| CG32036 | 0.178812 | -1.08808 | 4.061735 | -0.26789 | 0.788788 | NA |
| CG3437 | 1.188699 | -1.8233 | 3.980741 | -0.45803 | 0.646931 | NA |
| CG3448 | 0 | NA | NA | NA | NA | NA |
| CG3982 | 0.6282 | 1.523021 | 4.015426 | 0.379293 | 0.704471 | NA |
| Klp67A | 39.72894 | 2.852469 | 3.814609 | 0.747775 | 0.454596 | NA |
| CG4483 | 0 | NA | NA | NA | NA | NA |
| tRNA:Thr-/ | 0 | NA | NA | NA | NA | NA |
| GNBP3 | 42.19407 | 1.749778 | 3.952138 | 0.442742 | 0.657952 | NA |
| CG13314 | 0 | NA | NA | NA | NA | NA |
| Doc3 | 0 | NA | NA | NA | NA | NA |
| Doc1 | 0 | NA | NA | NA | NA | NA |
| CG13315 | 54.53546 | 2.089514 | 3.81177 | 0.548174 | 0.583572 | NA |
| CG5194 | 0 | NA | NA | NA | NA | NA |
| CG5280 | 0 | NA | NA | NA | NA | NA |
| CG5653 | 0.178812 | -1.08808 | 4.061735 | -0.26789 | 0.788788 | NA |
| ValRS-m | 20.56194 | -6.87598 | 3.848295 | -1.78676 | 0.073976 | NA |
| Cpr66D | 0 | NA | NA | NA | NA | NA |
| Pex7 | 2.601344 | -1.94058 | 3.843265 | -0.50493 | 0.613608 | NA |
| Cp18 | 47.86426 | -0.73412 | 3.838201 | -0.19127 | 0.848316 | NA |
| Cp19 | 0 | NA | NA | NA | NA | NA |
| Cp16 | 0 | NA | NA | NA | NA | NA |
| CG6576 | 16.94897 | 1.531701 | 3.953025 | 0.387476 | 0.698404 | NA |
| CG32024 | 0 | NA | NA | NA | NA | NA |

| | | | | | | |
|---|---|---|---|---|---|---|
| CG13308 | 0 | NA | NA | NA | NA | NA |
| CG13309 | 0 | NA | NA | NA | NA | NA |
| Mcm7 | 23.63412 | 0.363931 | 3.8372 | 0.094843 | 0.92444 | NA |
| CG13312 | 0.178053 | -1.16467 | 4.075067 | -0.2858 | 0.775028 | NA |
| CG13311 | 0 | NA | NA | NA | NA | NA |
| CG13310 | 0.178053 | -1.16467 | 4.075067 | -0.2858 | 0.775028 | NA |
| Acbp5 | 0.209247 | 1.042804 | 4.047837 | 0.25762 | 0.7967 | NA |
| CG13306 | 0 | NA | NA | NA | NA | NA |
| Prm | 0.71449 | -2.47883 | 4.014121 | -0.61753 | 0.536887 | NA |
| Cpr66Cb | 0 | NA | NA | NA | NA | NA |
| Jon66Ci | 0.178053 | -1.16467 | 4.075067 | -0.2858 | 0.775028 | NA |
| mtrm | 0 | NA | NA | NA | NA | NA |
| GstO3 | 48.21994 | -0.6997 | 3.843113 | -0.18207 | 0.85553 | NA |
| snoRNA:U4 | 0.528311 | -1.72839 | 4.011871 | -0.43082 | 0.6666 | NA |
| snoRNA:U4 | 0 | NA | NA | NA | NA | NA |
| S-Lap1 | 23.50303 | -1.70325 | 3.952538 | -0.43093 | 0.666522 | NA |
| S-Lap2 | 0.178053 | -1.16467 | 4.075067 | -0.2858 | 0.775028 | NA |
| Cdc6 | 0 | NA | NA | NA | NA | NA |
| GstO1 | 0 | NA | NA | NA | NA | NA |
| mkg-p | 0 | NA | NA | NA | NA | NA |
| Jon66Cii | 0.536437 | -1.71175 | 4.038042 | -0.42391 | 0.671635 | NA |
| CG33057 | 0 | NA | NA | NA | NA | NA |
| fan | 0 | NA | NA | NA | NA | NA |
| ldbr | 0.132078 | -1.13211 | 4.069918 | -0.27817 | 0.780885 | NA |
| mus301 | 0.460806 | -0.60052 | 3.997252 | -0.15023 | 0.880581 | NA |
| CG8006 | 0.578255 | 0.214162 | 3.950818 | 0.054207 | 0.95677 | NA |
| CG8012 | 1.250927 | -2.85877 | 4.004056 | -0.71397 | 0.475246 | NA |
| CG13678 | 0 | NA | NA | NA | NA | NA |
| CG13670 | 0 | NA | NA | NA | NA | NA |
| Gr66a | 0 | NA | NA | NA | NA | NA |
| CG7201 | 0 | NA | NA | NA | NA | NA |
| CG7213 | 32.66638 | 1.748349 | 3.952495 | 0.442341 | 0.658243 | NA |
| CG13674 | 0 | NA | NA | NA | NA | NA |
| CG7366 | 0 | NA | NA | NA | NA | NA |
| CG7387 | 0 | NA | NA | NA | NA | NA |
| mthl7 | 0 | NA | NA | NA | NA | NA |
| RecQ4 | 10.31535 | 0.546884 | 3.885881 | 0.140736 | 0.888078 | NA |
| CG13675 | 0.697584 | -1.11707 | 3.977017 | -0.28088 | 0.778801 | NA |
| CG13676 | 0 | NA | NA | NA | NA | NA |
| tRNA:Leu-( | 2.955291 | 2.974173 | 3.948131 | 0.753312 | 0.451263 | NA |
| tRNA:Leu-( | 0 | NA | NA | NA | NA | NA |
| tRNA:Leu-( | 0 | NA | NA | NA | NA | NA |
| tRNA:Leu-( | 0 | NA | NA | NA | NA | NA |
| tRNA:Leu-( | 0 | NA | NA | NA | NA | NA |
| tRNA:Leu-( | 0 | NA | NA | NA | NA | NA |
| eIF4E5 | 0 | NA | NA | NA | NA | NA |
| tRNA:Ala-/ | 0 | NA | NA | NA | NA | NA |
| CG32371 | 51.73315 | -2.1338 | 3.836203 | -0.55623 | 0.578056 | NA |
| mthl6 | 0 | NA | NA | NA | NA | NA |
| Cyp316a1 | 0.178053 | -1.16467 | 4.075067 | -0.2858 | 0.775028 | NA |

| | | | | | | |
|---|---|---|---|---|---|---|
| frac | 0 | NA | NA | NA | NA | NA |
| CG14838 | 0.178812 | -1.08808 | 4.061735 | -0.26789 | 0.788788 | NA |
| CG32373 | 0.932373 | 1.053996 | 3.996193 | 0.26375 | 0.791972 | NA |
| PGRP-SD | 0.64106 | -0.7867 | 4.009822 | -0.19619 | 0.844459 | NA |
| Mcad | 1.603238 | -2.77841 | 3.960305 | -0.70157 | 0.48295 | NA |
| CG7409 | 0 | NA | NA | NA | NA | NA |
| CG8492 | 0.3873 | 0.267327 | 4.008308 | 0.066693 | 0.946826 | NA |
| CG7457 | 6.195187 | -0.9937 | 3.870835 | -0.25671 | 0.797399 | NA |
| CG7716 | 0 | NA | NA | NA | NA | NA |
| CG14837 | 3.978597 | 1.705937 | 3.963372 | 0.430426 | 0.666886 | NA |
| CG32376 | 0 | NA | NA | NA | NA | NA |
| CG32374 | 0 | NA | NA | NA | NA | NA |
| CG16998 | 0 | NA | NA | NA | NA | NA |
| CG7548 | 0 | NA | NA | NA | NA | NA |
| CG8539 | 3.026906 | -1.67104 | 3.963213 | -0.42164 | 0.67329 | NA |
| Cyp4d8 | 0 | NA | NA | NA | NA | NA |
| CG14826 | 0 | NA | NA | NA | NA | NA |
| mei-P22 | 0 | NA | NA | NA | NA | NA |
| CG8607 | 54.06369 | 0.355682 | 3.835105 | 0.092744 | 0.926107 | NA |
| mus312 | 0 | NA | NA | NA | NA | NA |
| CG8563 | 0.836986 | 1.408818 | 3.9887 | 0.353202 | 0.723937 | NA |
| CG32379 | 0.535678 | -2.26616 | 4.021604 | -0.5635 | 0.573098 | NA |
| CG8562 | 0 | NA | NA | NA | NA | NA |
| CG18417 | 0.178053 | -1.16467 | 4.075067 | -0.2858 | 0.775028 | NA |
| CG33278 | 0 | NA | NA | NA | NA | NA |
| Tsp66A | 0 | NA | NA | NA | NA | NA |
| CG8541 | 0 | NA | NA | NA | NA | NA |
| CG8543 | 0 | NA | NA | NA | NA | NA |
| sphinx2 | 0 | NA | NA | NA | NA | NA |
| sphinx1 | 0.132078 | -1.13211 | 4.069918 | -0.27817 | 0.780885 | NA |
| CG14835 | 0 | NA | NA | NA | NA | NA |
| CG8560 | 0.233093 | 0.692656 | 4.076246 | 0.169925 | 0.865069 | NA |
| CG8564 | 0 | NA | NA | NA | NA | NA |
| CG14834 | 0 | NA | NA | NA | NA | NA |
| Cln7 | 2.617498 | 1.683176 | 3.969392 | 0.424039 | 0.671538 | NA |
| RhoGEF4 | 0 | NA | NA | NA | NA | NA |
| CG14830 | 0.4188 | 1.41068 | 4.034613 | 0.349644 | 0.726606 | NA |
| CG14829 | 0 | NA | NA | NA | NA | NA |
| CG14820 | 0 | NA | NA | NA | NA | NA |
| CG14823 | 33.69012 | 0.334761 | 3.835151 | 0.087288 | 0.930443 | NA |
| Cpr65Ec | 0 | NA | NA | NA | NA | NA |
| BBS1 | 1.172943 | -0.24009 | 3.970289 | -0.06047 | 0.951781 | NA |
| Acbp6 | 0 | NA | NA | NA | NA | NA |
| melt | 0.712972 | -2.55723 | 4.00634 | -0.63829 | 0.523282 | NA |
| Acbp4 | 0 | NA | NA | NA | NA | NA |
| Cpr65Eb | 0.178053 | -1.16467 | 4.075067 | -0.2858 | 0.775028 | NA |
| Cpr65Ea | 0 | NA | NA | NA | NA | NA |
| CG32391 | 0 | NA | NA | NA | NA | NA |
| CG17744 | 0.356107 | -1.47099 | 4.037764 | -0.36431 | 0.715628 | NA |
| CG10063 | 0 | NA | NA | NA | NA | NA |

| | | | | | | |
|---|---|---|---|---|---|---|
| CG10064 | 0.233093 | 0.692656 | 4.076246 | 0.169925 | 0.865069 | NA |
| CG8519 | 0.965478 | 0.284617 | 3.909784 | 0.072796 | 0.941968 | NA |
| Prat2 | 0 | NA | NA | NA | NA | NA |
| Rsph3 | 6.534036 | 0.399922 | 3.95438 | 0.101134 | 0.919444 | NA |
| SP1173 | 51.4057 | -0.30865 | 3.840908 | -0.08036 | 0.935952 | NA |
| CG32395 | 0 | NA | NA | NA | NA | NA |
| eIF4E4 | 0.337793 | 1.451235 | 4.039844 | 0.35923 | 0.719423 | NA |
| CG32396 | 5.618481 | 0.256232 | 3.939283 | 0.065045 | 0.948138 | NA |
| zpg | 0 | NA | NA | NA | NA | NA |
| Lcp65Ad | 0 | NA | NA | NA | NA | NA |
| Lcp65Ac | 0 | NA | NA | NA | NA | NA |
| Lcp65Ab2 | 0 | NA | NA | NA | NA | NA |
| Lcp65APsi | 0 | NA | NA | NA | NA | NA |
| Lcp65Ab1 | 0 | NA | NA | NA | NA | NA |
| Lcp65Aa | 0 | NA | NA | NA | NA | NA |
| Cpr65Az | 0 | NA | NA | NA | NA | NA |
| Prdm13 | 0 | NA | NA | NA | NA | NA |
| CG7376 | 0 | NA | NA | NA | NA | NA |
| CG7386 | 0.178812 | -1.08808 | 4.061735 | -0.26789 | 0.788788 | NA |
| CG8219 | 0 | NA | NA | NA | NA | NA |
| Or65a | 0 | NA | NA | NA | NA | NA |
| CG10163 | 0 | NA | NA | NA | NA | NA |
| Best2 | 0 | NA | NA | NA | NA | NA |
| Mdr65 | 9.063163 | -1.59667 | 3.901916 | -0.4092 | 0.682393 | NA |
| CG10226 | 31.2421 | 2.010797 | 3.841139 | 0.52349 | 0.600633 | NA |
| CG13299 | 0 | NA | NA | NA | NA | NA |
| CG13297 | 0 | NA | NA | NA | NA | NA |
| Acp65Aa | 0 | NA | NA | NA | NA | NA |
| Cpr65Ax2 | 0 | NA | NA | NA | NA | NA |
| Cpr65Aw | 0 | NA | NA | NA | NA | NA |
| Cpr65Av | 0 | NA | NA | NA | NA | NA |
| CG5568 | 0.178053 | -1.16467 | 4.075067 | -0.2858 | 0.775028 | NA |
| CG18586 | 0.104623 | 0.639075 | 4.067674 | 0.157111 | 0.875158 | NA |
| CG5592 | 0.715249 | -1.88064 | 4.034336 | -0.46616 | 0.641102 | NA |
| CG13288 | 15.49551 | 2.952995 | 3.924049 | 0.752538 | 0.451728 | NA |
| yip7 | 0.356866 | -1.92326 | 4.037889 | -0.4763 | 0.633858 | NA |
| CG6462 | 0 | NA | NA | NA | NA | NA |
| Jon65Aiii | 0.1047 | 0.613261 | 4.062799 | 0.150945 | 0.880019 | NA |
| Jon65Aii | 1.165466 | 1.076041 | 3.987865 | 0.269829 | 0.787292 | NA |
| CG6592 | 0.356866 | -1.92326 | 4.037889 | -0.4763 | 0.633858 | NA |
| CG6602 | 0 | NA | NA | NA | NA | NA |
| Ets65A | 33.62193 | -1.20414 | 3.737232 | -0.3222 | 0.747301 | NA |
| Lcp65Ae | 0 | NA | NA | NA | NA | NA |
| Lcp65Af | 0 | NA | NA | NA | NA | NA |
| Lcp65Ag1 | 0 | NA | NA | NA | NA | NA |
| Lcp65Ag2 | 0.178053 | -1.16467 | 4.075067 | -0.2858 | 0.775028 | NA |
| Cpr65Au | 0 | NA | NA | NA | NA | NA |
| CG10469 | 0.132078 | -1.13211 | 4.069918 | -0.27817 | 0.780885 | NA |
| CG10472 | 0 | NA | NA | NA | NA | NA |
| Jon65Ai | 0.53416 | -1.53481 | 4.012395 | -0.38252 | 0.702078 | NA |

| | | | | | | |
|---|---|---|---|---|---|---|
| CG10477 | 49.80582 | 1.710623 | 3.951918 | 0.432859 | 0.665117 | NA |
| alphaKap4 | 0.178053 | -1.16467 | 4.075067 | -0.2858 | 0.775028 | NA |
| CG10479 | 35.69879 | -0.80072 | 3.862535 | -0.2073 | 0.835773 | NA |
| CG10486 | 0 | NA | NA | NA | NA | NA |
| CG32413 | 0 | NA | NA | NA | NA | NA |
| QC | 0.178053 | -1.16467 | 4.075067 | -0.2858 | 0.775028 | NA |
| PolE2 | 0 | NA | NA | NA | NA | NA |
| Alp9 | 0 | NA | NA | NA | NA | NA |
| mthl2 | 0.104623 | 0.639075 | 4.067674 | 0.157111 | 0.875158 | NA |
| mad2 | 0.756593 | 2.288499 | 4.007508 | 0.571053 | 0.567964 | NA |
| Tektin-C | 0.104623 | 0.639075 | 4.067674 | 0.157111 | 0.875158 | NA |
| Cralbp | 0 | NA | NA | NA | NA | NA |
| Ppat-Dpck | 44.5857 | 4.998179 | 3.815108 | 1.310102 | 0.190161 | NA |
| CG10591 | 18.16143 | -1.70178 | 3.953016 | -0.4305 | 0.666832 | NA |
| Alp10 | 0.104623 | 0.639075 | 4.067674 | 0.157111 | 0.875158 | NA |
| spo | 2.704526 | -0.12723 | 3.962037 | -0.03211 | 0.974384 | NA |
| CG13285 | 20.41649 | 1.744587 | 3.953439 | 0.441283 | 0.659008 | NA |
| tRNA:Ser-/ | 0 | NA | NA | NA | NA | NA |
| tRNA:Val-T | 0 | NA | NA | NA | NA | NA |
| blanks | 0.104623 | 0.639075 | 4.067674 | 0.157111 | 0.875158 | NA |
| Ir64a | 0 | NA | NA | NA | NA | NA |
| tRNA:Val-T | 0.178053 | -1.16467 | 4.075067 | -0.2858 | 0.775028 | NA |
| lama | 52.53549 | 5.177539 | 3.863175 | 1.340229 | 0.180171 | NA |
| CG4597 | 0 | NA | NA | NA | NA | NA |
| CG15212 | 0.178053 | -1.16467 | 4.075067 | -0.2858 | 0.775028 | NA |
| Gdap1 | 0 | NA | NA | NA | NA | NA |
| CG10674 | 11.07347 | 0.05554 | 3.740588 | 0.014848 | 0.988153 | NA |
| CG15213 | 0 | NA | NA | NA | NA | NA |
| tRNA:Arg-' | 14.37984 | -3.72513 | 3.833179 | -0.97181 | 0.331144 | NA |
| CG1494 | 0.712213 | -1.57119 | 3.998613 | -0.39293 | 0.694369 | NA |
| Ir20a | 0.178053 | -1.16467 | 4.075067 | -0.2858 | 0.775028 | NA |
| CG10918 | 0 | NA | NA | NA | NA | NA |
| whe | 0 | NA | NA | NA | NA | NA |
| lcs | 0 | NA | NA | NA | NA | NA |
| snoRNA:M | 0 | NA | NA | NA | NA | NA |
| CG1801 | 2.166279 | 2.499239 | 3.918069 | 0.637875 | 0.523555 | NA |
| tRNA:Ile-A | 1.631653 | 1.102918 | 3.977952 | 0.277258 | 0.781582 | NA |
| CG11666 | 0 | NA | NA | NA | NA | NA |
| CG1724 | 0 | NA | NA | NA | NA | NA |
| CR32511 | 0 | NA | NA | NA | NA | NA |
| CG1503 | 37.76609 | 1.837408 | 3.89293 | 0.471986 | 0.636937 | NA |
| CG15446 | 26.35188 | -1.70379 | 3.952363 | -0.43108 | 0.66641 | NA |
| CG15449 | 0 | NA | NA | NA | NA | NA |
| CG1304 | 0 | NA | NA | NA | NA | NA |
| Ser6 | 0.1047 | 0.613261 | 4.062799 | 0.150945 | 0.880019 | NA |
| CG11227 | 0 | NA | NA | NA | NA | NA |
| CR11235 | 0 | NA | NA | NA | NA | NA |
| CG1314 | 0.31387 | 1.261782 | 4.037145 | 0.312543 | 0.754628 | NA |
| CG32507 | 0.233093 | 0.692656 | 4.076246 | 0.169925 | 0.865069 | NA |
| CG15452 | 0 | NA | NA | NA | NA | NA |

| | | | | | | |
|---|---|---|---|---|---|---|
| CG1324 | 0.104623 | 0.639075 | 4.067674 | 0.157111 | 0.875158 | NA |
| CG1631 | 0.178053 | -1.16467 | 4.075067 | -0.2858 | 0.775028 | NA |
| Obp19a | 0.357625 | -1.47707 | 4.044794 | -0.36518 | 0.714979 | NA |
| Obp19b | 0 | NA | NA | NA | NA | NA |
| CG15461 | 1.251686 | -2.06022 | 4.011654 | -0.51356 | 0.60756 | NA |
| Obp19c | 0 | NA | NA | NA | NA | NA |
| CG15458 | 0.104623 | 0.639075 | 4.067674 | 0.157111 | 0.875158 | NA |
| CG15459 | 0 | NA | NA | NA | NA | NA |
| jb | 0 | NA | NA | NA | NA | NA |
| CG12679 | 0 | NA | NA | NA | NA | NA |
| CG1504 | 59.88496 | -0.92728 | 3.834286 | -0.24184 | 0.808905 | NA |
| Inx6 | 0 | NA | NA | NA | NA | NA |
| tilB | 0 | NA | NA | NA | NA | NA |
| Cda4 | 0 | NA | NA | NA | NA | NA |
| CG14615 | 0 | NA | NA | NA | NA | NA |
| CG15635 | 0.209247 | 1.042804 | 4.047837 | 0.25762 | 0.7967 | NA |
| CG11929 | 0 | NA | NA | NA | NA | NA |
| Elba3 | 0.53416 | -1.53481 | 4.012395 | -0.38252 | 0.702078 | NA |
| CG15629 | 0 | NA | NA | NA | NA | NA |
| Ir25a | 0.3873 | 0.267327 | 4.008308 | 0.066693 | 0.946826 | NA |
| CG3225 | 38.42487 | 1.749297 | 3.952259 | 0.442607 | 0.65805 | NA |
| CG3251 | 0 | NA | NA | NA | NA | NA |
| Bsg25A | 0 | NA | NA | NA | NA | NA |
| CG15429 | 0 | NA | NA | NA | NA | NA |
| Traf4 | 0 | NA | NA | NA | NA | NA |
| Tps1 | 0 | NA | NA | NA | NA | NA |
| CG33003 | 0 | NA | NA | NA | NA | NA |
| ND-B8 | 0.53416 | -1.53481 | 4.012395 | -0.38252 | 0.702078 | NA |
| HP6 | 0 | NA | NA | NA | NA | NA |
| tRNA:Trp-( | 7.328995 | 1.727313 | 3.957831 | 0.436429 | 0.662525 | NA |
| Sr-CIII | 0 | NA | NA | NA | NA | NA |
| Sr-CI | 0 | NA | NA | NA | NA | NA |
| CG2955 | 0 | NA | NA | NA | NA | NA |
| lectin-24D | 0.178812 | -1.08808 | 4.061735 | -0.26789 | 0.788788 | NA |
| ft | 25.18057 | 2.402066 | 3.924325 | 0.612097 | 0.540474 | NA |
| Or24a | 0 | NA | NA | NA | NA | NA |
| bowl | 0 | NA | NA | NA | NA | NA |
| CG31960 | 0 | NA | NA | NA | NA | NA |
| CR31958 | 0 | NA | NA | NA | NA | NA |
| bark | 41.52989 | -4.2848 | 3.850407 | -1.11282 | 0.265786 | NA |
| TTLL4B | 0.178053 | -1.16467 | 4.075067 | -0.2858 | 0.775028 | NA |
| CG15422 | 0.178053 | -1.16467 | 4.075067 | -0.2858 | 0.775028 | NA |
| CG15423 | 0 | NA | NA | NA | NA | NA |
| CG31773 | 47.01027 | 1.750282 | 3.952013 | 0.442884 | 0.65785 | NA |
| CG31776 | 18.95069 | 1.743818 | 3.953632 | 0.441067 | 0.659164 | NA |
| Pgant4 | 36.49196 | 2.1296 | 3.827282 | 0.556426 | 0.57792 | NA |
| CG8851 | 0 | NA | NA | NA | NA | NA |
| IFT57 | 0.178812 | -1.08808 | 4.061735 | -0.26789 | 0.788788 | NA |
| CG2816 | 0 | NA | NA | NA | NA | NA |
| lectin-24A | 0 | NA | NA | NA | NA | NA |

| | | | | | | |
|---|---|---|---|---|---|---|
| CG31955 | 11.51692 | 2.942648 | 3.926413 | 0.74945 | 0.453586 | NA |
| cutlet | 0.178053 | -1.16467 | 4.075067 | -0.2858 | 0.775028 | NA |
| CG31777 | 0 | NA | NA | NA | NA | NA |
| CG31778 | 0 | NA | NA | NA | NA | NA |
| CG3604 | 0 | NA | NA | NA | NA | NA |
| IM33 | 0 | NA | NA | NA | NA | NA |
| CG16713 | 10.43414 | -1.91017 | 3.954613 | -0.48302 | 0.629079 | NA |
| Acp24A4 | 0 | NA | NA | NA | NA | NA |
| CG3513 | 0 | NA | NA | NA | NA | NA |
| CG16704 | 1.71701 | -1.85109 | 3.972145 | -0.46602 | 0.641203 | NA |
| CG15418 | 0 | NA | NA | NA | NA | NA |
| odd | 0.178812 | -1.08808 | 4.061735 | -0.26789 | 0.788788 | NA |
| sob | 0.104623 | 0.639075 | 4.067674 | 0.157111 | 0.875158 | NA |
| Spindly | 0 | NA | NA | NA | NA | NA |
| Sr-CIV | 0.104623 | 0.639075 | 4.067674 | 0.157111 | 0.875158 | NA |
| CG3213 | 1.032167 | -2.82383 | 3.952034 | -0.71452 | 0.474903 | NA |
| CG31954 | 0 | NA | NA | NA | NA | NA |
| CG15414 | 0.590718 | -0.57492 | 4.003363 | -0.14361 | 0.885809 | NA |
| Pif1 | 10.68396 | -2.91163 | 3.923713 | -0.74206 | 0.458051 | NA |
| CG3246 | 0.178053 | -1.16467 | 4.075067 | -0.2858 | 0.775028 | NA |
| CR31953 | 0 | NA | NA | NA | NA | NA |
| tRNA:Ser-/ | 0 | NA | NA | NA | NA | NA |
| CG15404 | 0 | NA | NA | NA | NA | NA |
| CG15406 | 0 | NA | NA | NA | NA | NA |
| CG33281 | 0 | NA | NA | NA | NA | NA |
| CG33282 | 0 | NA | NA | NA | NA | NA |
| CG3285 | 0.264155 | -1.54099 | 4.048976 | -0.38059 | 0.703509 | NA |
| CG15408 | 0 | NA | NA | NA | NA | NA |
| CG8838 | 0 | NA | NA | NA | NA | NA |
| CG8840 | 0 | NA | NA | NA | NA | NA |
| CG31952 | 17.27549 | 1.742783 | 3.953893 | 0.440776 | 0.659375 | NA |
| Fign | 1.246373 | -1.6238 | 3.979526 | -0.40804 | 0.683245 | NA |
| OtopLb | 0.178053 | -1.16467 | 4.075067 | -0.2858 | 0.775028 | NA |
| CG31698 | 0 | NA | NA | NA | NA | NA |
| tRNA:Ser-/ | 0 | NA | NA | NA | NA | NA |
| Cpr23B | 0 | NA | NA | NA | NA | NA |
| CG2975 | 0.1047 | 0.613261 | 4.062799 | 0.150945 | 0.880019 | NA |
| CG2983 | 0 | NA | NA | NA | NA | NA |
| CG8813 | 30.01746 | 2.825397 | 3.921362 | 0.720514 | 0.471209 | NA |
| CG17264 | 0.178053 | -1.16467 | 4.075067 | -0.2858 | 0.775028 | NA |
| CG17219 | 15.78326 | -4.82319 | 3.879528 | -1.24324 | 0.213779 | NA |
| snRNA:U5: | 0 | NA | NA | NA | NA | NA |
| PpD6 | 0 | NA | NA | NA | NA | NA |
| CG17258 | 0.807012 | 0.227825 | 3.984055 | 0.057184 | 0.954399 | NA |
| okr | 0 | NA | NA | NA | NA | NA |
| Ts | 0 | NA | NA | NA | NA | NA |
| CG3117 | 0 | NA | NA | NA | NA | NA |
| CG18557 | 0 | NA | NA | NA | NA | NA |
| CG18558 | 0 | NA | NA | NA | NA | NA |
| CG3119 | 0 | NA | NA | NA | NA | NA |

| | | | | | | |
|---|---|---|---|---|---|---|
| Elba2 | 0 | NA | NA | NA | NA | NA |
| CG15395 | 0 | NA | NA | NA | NA | NA |
| Prosbeta4I | 0 | NA | NA | NA | NA | NA |
| Or23a | 0.53416 | -1.53481 | 4.012395 | -0.38252 | 0.702078 | NA |
| Prosbeta4I | 0 | NA | NA | NA | NA | NA |
| Trf4 | 0.1047 | 0.613261 | 4.062799 | 0.150945 | 0.880019 | NA |
| Gr23a | 0.178053 | -1.16467 | 4.075067 | -0.2858 | 0.775028 | NA |
| CG2964 | 0.178053 | -1.16467 | 4.075067 | -0.2858 | 0.775028 | NA |
| CG3123 | 0.53416 | -1.53481 | 4.012395 | -0.38252 | 0.702078 | NA |
| CG15399 | 0 | NA | NA | NA | NA | NA |
| CG9961 | 0 | NA | NA | NA | NA | NA |
| CG33125 | 13.17699 | -0.04195 | 3.934483 | -0.01066 | 0.991492 | NA |
| CG9962 | 0 | NA | NA | NA | NA | NA |
| Rad1 | 0 | NA | NA | NA | NA | NA |
| Cyp309a2 | 0.64424 | -0.14539 | 3.993669 | -0.0364 | 0.97096 | NA |
| Cyp309a1 | 0.132078 | -1.13211 | 4.069918 | -0.27817 | 0.780885 | NA |
| CG3515 | 0 | NA | NA | NA | NA | NA |
| CG17234 | 0 | NA | NA | NA | NA | NA |
| CG4270 | 0 | NA | NA | NA | NA | NA |
| CG4271 | 0 | NA | NA | NA | NA | NA |
| CG31949 | 20.92335 | 0.036905 | 3.932004 | 0.009386 | 0.992511 | NA |
| CG16995 | 1.608552 | -3.02178 | 4.000667 | -0.75532 | 0.450057 | NA |
| CG15390 | 12.70381 | 2.601219 | 3.852935 | 0.675126 | 0.499595 | NA |
| dpp | 0.492 | 1.04661 | 3.990653 | 0.262265 | 0.793117 | NA |
| CG3528 | 0 | NA | NA | NA | NA | NA |
| CG31948 | 0 | NA | NA | NA | NA | NA |
| CG18641 | 0 | NA | NA | NA | NA | NA |
| CG3557 | 0.387453 | 0.098632 | 4.002797 | 0.024641 | 0.980342 | NA |
| CG3597 | 0 | NA | NA | NA | NA | NA |
| robl22E | 0.178053 | -1.16467 | 4.075067 | -0.2858 | 0.775028 | NA |
| CG17237 | 0 | NA | NA | NA | NA | NA |
| CG31681 | 0 | NA | NA | NA | NA | NA |
| Send1 | 0 | NA | NA | NA | NA | NA |
| CG17239 | 0 | NA | NA | NA | NA | NA |
| Ser12 | 0 | NA | NA | NA | NA | NA |
| Tengl1 | 0 | NA | NA | NA | NA | NA |
| Tengl2 | 0 | NA | NA | NA | NA | NA |
| Tengl3 | 0 | NA | NA | NA | NA | NA |
| tRNA:Arg-1 | 0 | NA | NA | NA | NA | NA |
| Nplp4 | 0.1047 | 0.613261 | 4.062799 | 0.150945 | 0.880019 | NA |
| tRNA:Gln-( | 0.2094 | 1.005034 | 4.045341 | 0.248442 | 0.803792 | NA |
| tRNA:Gln-( | 0 | NA | NA | NA | NA | NA |
| tRNA:Gly-( | 0.178053 | -1.16467 | 4.075067 | -0.2858 | 0.775028 | NA |
| Kebab | 0 | NA | NA | NA | NA | NA |
| Or22c | 0 | NA | NA | NA | NA | NA |
| CG12674 | 0 | NA | NA | NA | NA | NA |
| tRNA:Arg-1 | 0 | NA | NA | NA | NA | NA |
| CG15386 | 0 | NA | NA | NA | NA | NA |
| CG33124 | 0.71449 | -2.47883 | 4.014121 | -0.61753 | 0.536887 | NA |
| CG31668 | 0.233093 | 0.692656 | 4.076246 | 0.169925 | 0.865069 | NA |

| | | | | | | |
|---|---|---|---|---|---|---|
| CG4259 | 0.233093 | 0.692656 | 4.076246 | 0.169925 | 0.865069 | NA |
| tRNA:Gln-( | 23.99654 | -0.2631 | 3.937905 | -0.06681 | 0.946732 | NA |
| tRNA:Gln-( | 1.981396 | -0.40108 | 3.850773 | -0.10416 | 0.917045 | NA |
| tRNA:Gln-( | 12.82013 | 1.16938 | 3.954559 | 0.295704 | 0.767456 | NA |
| CG15353 | 0.31387 | 1.261782 | 4.037145 | 0.312543 | 0.754628 | NA |
| Obp22a | 0 | NA | NA | NA | NA | NA |
| tRNA:Gly-( | 20.12002 | -1.7024 | 3.952811 | -0.43068 | 0.6667 | NA |
| Or22a | 0.53416 | -1.53481 | 4.012395 | -0.38252 | 0.702078 | NA |
| Or22b | 0.233093 | 0.692656 | 4.076246 | 0.169925 | 0.865069 | NA |
| CG17648 | 0 | NA | NA | NA | NA | NA |
| Gr22e | 0 | NA | NA | NA | NA | NA |
| CG7295 | 0 | NA | NA | NA | NA | NA |
| CG15358 | 1.303633 | 3.749586 | 3.93937 | 0.951824 | 0.341186 | NA |
| c-cup | 0 | NA | NA | NA | NA | NA |
| CG31664 | 0 | NA | NA | NA | NA | NA |
| CG31933 | 0 | NA | NA | NA | NA | NA |
| Gr22a | 0 | NA | NA | NA | NA | NA |
| Gr22d | 0 | NA | NA | NA | NA | NA |
| CG17650 | 0 | NA | NA | NA | NA | NA |
| Gr22f | 0 | NA | NA | NA | NA | NA |
| halo | 0 | NA | NA | NA | NA | NA |
| CG18132 | 0 | NA | NA | NA | NA | NA |
| CG5397 | 0 | NA | NA | NA | NA | NA |
| a5 | 0.660388 | -1.75977 | 4.001266 | -0.4398 | 0.660079 | NA |
| CG5440 | 0 | NA | NA | NA | NA | NA |
| Cdlc2 | 1.754666 | -0.19119 | 3.90635 | -0.04894 | 0.960963 | NA |
| CG14346 | 0.132078 | -1.13211 | 4.069918 | -0.27817 | 0.780885 | NA |
| snRNA:U3 | 0 | NA | NA | NA | NA | NA |
| CG31928 | 0 | NA | NA | NA | NA | NA |
| CG33128 | 0 | NA | NA | NA | NA | NA |
| CG31926 | 0 | NA | NA | NA | NA | NA |
| CG31661 | 0 | NA | NA | NA | NA | NA |
| CG7420 | 37.72406 | -6.09319 | 3.83485 | -1.5889 | 0.112083 | NA |
| CG18131 | 0 | NA | NA | NA | NA | NA |
| CG31659 | 0 | NA | NA | NA | NA | NA |
| CG5565 | 0 | NA | NA | NA | NA | NA |
| CG31924 | 0 | NA | NA | NA | NA | NA |
| CG5561 | 0 | NA | NA | NA | NA | NA |
| CG5556 | 0 | NA | NA | NA | NA | NA |
| CG4415 | 18.53189 | 1.743577 | 3.953693 | 0.441 | 0.659213 | NA |
| ast | 3.261816 | -4.09457 | 3.920422 | -1.04442 | 0.296291 | NA |
| CG14339 | 0 | NA | NA | NA | NA | NA |
| CG31922 | 18.89855 | 0.335597 | 3.734209 | 0.089871 | 0.92839 | NA |
| CG5011 | 0 | NA | NA | NA | NA | NA |
| CG14342 | 0 | NA | NA | NA | NA | NA |
| CG5139 | 0 | NA | NA | NA | NA | NA |
| CG5080 | 0.67891 | -1.66339 | 3.984816 | -0.41743 | 0.676362 | NA |
| Iris | 0 | NA | NA | NA | NA | NA |
| CG4375 | 0 | NA | NA | NA | NA | NA |
| CG2839 | 3.796853 | -0.83753 | 3.822349 | -0.21912 | 0.82656 | NA |

| | | | | | | |
|---|---|---|---|---|---|---|
| Hsp60B | 0 | NA | NA | NA | NA | NA |
| CG12506 | 0 | NA | NA | NA | NA | NA |
| CG13946 | 0 | NA | NA | NA | NA | NA |
| CG13947 | 0 | NA | NA | NA | NA | NA |
| Gr21a | 0.178053 | -1.16467 | 4.075067 | -0.2858 | 0.775028 | NA |
| CG3544 | 0 | NA | NA | NA | NA | NA |
| Nnf1b | 0 | NA | NA | NA | NA | NA |
| CG13949 | 0 | NA | NA | NA | NA | NA |
| CG13950 | 0 | NA | NA | NA | NA | NA |
| ninaA | 0 | NA | NA | NA | NA | NA |
| Lsp1beta | 0.64424 | -0.14539 | 3.993669 | -0.0364 | 0.97096 | NA |
| GluRIIC | 0.104623 | 0.639075 | 4.067674 | 0.157111 | 0.875158 | NA |
| snRNA:U1: | 0 | NA | NA | NA | NA | NA |
| CG15824 | 0 | NA | NA | NA | NA | NA |
